# Multi-Platform Proteomic Analysis of Alzheimer’s Disease Cerebrospinal Fluid and Plasma Reveals Network Biomarkers Associated with Proteostasis and the Matrisome

**DOI:** 10.1101/2022.06.20.494087

**Authors:** Eric B. Dammer, Lingyan Ping, Duc M. Duong, Erica S. Modeste, Nicholas T. Seyfried, James J. Lah, Allan I. Levey, Erik C.B. Johnson

## Abstract

Robust and accessible biomarkers that can capture the heterogeneity of Alzheimer’s disease and its diverse pathological processes are urgently needed. Here, we undertook an investigation of Alzheimer’s disease cerebrospinal fluid (CSF) and plasma from the same subjects using three different proteomic platforms—SomaLogic SomaScan, Olink proximity extension assay, and tandem mass tag-based mass spectrometry—to assess which protein markers in these two biofluids may serve as reliable biomarkers of AD pathophysiology observed from unbiased brain proteomics studies. Median correlation of overlapping protein measurements across platforms in CSF (*r*∼0.7) and plasma (*r*∼0.6) was good, with more variability in plasma. The SomaScan technology provided the most measurements in plasma. Surprisingly, many proteins altered in AD CSF were found to be altered in the opposite direction in plasma, including important members of AD brain co-expression modules. An exception was SMOC1, a key member of the brain matrisome module associated with amyloid-β deposition in AD, which was found to be elevated in both CSF and plasma. Protein co-expression analysis on greater than 7000 protein measurements in CSF and 9500 protein measurements in plasma across all proteomic platforms revealed strong changes in modules related to autophagy, ubiquitination, and sugar metabolism in CSF, and endocytosis and the matrisome in plasma. Cross-platform and cross-biofluid proteomics represents a promising approach for AD biomarker development.

## Introduction

Alzheimer’s disease (AD) is a growing public health problem with no available disease-modifying therapies. Multi-omic analyses of AD brain have illustrated the varied and complex pathophysiology beyond amyloid-β (Aβ) plaques and tau neurofibrillary tangles^1–5^, but how these pathological processes develop over time during the disease course is unclear. Given that AD is a heterogeneous disease, composed of different combinations and degrees of brain pathologies in a given individual, multiple biomarkers beyond Aβ and tau will be required to advance our understanding of the complex disease processes underlying AD. One of the current limitations for advancement of AD research, clinical care, and therapeutic development is the lack of easily-accessible biofluid biomarkers for these varied pathological processes.

Protein biomarkers represent a promising class of biomarkers for AD given the large diversity of potential markers, their direct role in subserving biological processes, and the fact that standard protein affinity-based measurement approaches are already deployed in most clinical laboratories around the world. Three main quantitative protein measurement technologies are currently available to conduct proteomic discovery experiments in biofluids at scale: mass spectrometry, multiplexed nucleic acid aptamers, and multiplexed antibodies. Mass spectrometry (MS) has been used most extensively to date in AD biofluid biomarker discovery research, and provides a direct measurement of protein identity and abundance through the measurement of peptides^3, 6–8^. Through the use of isobaric tandem mass tags (TMT) or data-independent acquisition (DIA) techniques^9^, cohorts of hundreds of subjects can be analyzed at a depth of thousands of proteins in CSF and plasma. Traditionally, depth of analysis by MS is limited by the large dynamic range of protein concentration present in CSF and, especially, in plasma, as well as the problem of missing values that accumulate when analyzing larger cohorts^10–12^. Specificity and accuracy may be affected by ion interference as the complexity of the matrix increases. More recently, two affinity-based proteomic technologies have become available for biofluid measurements that offer different advantages and disadvantages compared to MS for protein measurement in biofluids: the SomaScan® aptamer-based technology from SomaLogic, and proximity extension assay (PEA) technology from Olink®. SomaScan uses modified DNA aptamers (SOMAmers) with slow off-rate binding kinetics to measure relative protein levels in multiplex fashion^13–15^. Multiple SOMAmers can be generated towards a given protein target and included in the microarray-based readout of relative fluorescence intensity. PEA uses a sandwich antibody-based approach where the capture and detection antibodies are conjugated to a complementary oligonucleotide probe pair, the levels of which are ultimately measured by quantitative PCR or next-generation sequencing approaches to provide a relative protein abundance value^16, 17^. Measurement specificity is provided both through the dual epitope antibody binding as well as through specific oligonucleotide probe hybridization. As affinity-based approaches, both SomaScan and PEA (subsequently referred to as “Olink”) theoretically suffer less from dynamic range and missing value challenges compared to MS. However, because both are affinity-based, they are indirect measurements of protein identity and abundance. Specificity and accuracy may be affected by protein modifications or off-target binding, and sensitive and specific reagents must be designed for each protein.

In order to further the development of robust biofluid biomarkers of AD that can reflect multiple pathological processes in brain, we conducted a proteomic analysis on CSF and plasma by applying each proteomic technology described above to the same discovery set of AD and control samples. By analyzing the same samples with each proteomic technology, we were able to conduct an in-depth cross-platform technical analysis to better understand the current strengths and limitations inherent in each platform for AD biofluid biomarker development in CSF and plasma, and increase confidence in proteins that show promise as potential AD biomarkers. Furthermore, because the CSF and plasma samples we analyzed were matched within subject, we were able to explore the relationship between CSF and plasma levels of promising AD biomarkers within subject. We leveraged the combined proteomic datasets to generate an AD protein co-expression network in CSF and plasma, and explored how protein co-expression in these two fluids might be related to each other and to AD brain protein co-expression. We found strong co-expression signals for proteostasis, synaptic biology, sugar metabolism, complement, and TGF-β signaling in AD CSF, and endocytosis, matrisome, and complement in AD plasma. Multi-platform proteomic analysis of AD biofluids holds promise for identification of robust biomarkers for clinical translation.

## Results

### Pre-Processing and Technical Analyses of Proteomic Measurements in CSF and Plasma

For our discovery experiments and cross-platform proteomic analyses, we used CSF and plasma from a cohort of control (*n*=18) and AD (*n*=18) patients in the Emory Goizueta Alzheimer’s Disease Research Center (ADRC) (**Figure 1, Supplementary Table 1**). The CSF and plasma samples were drawn at or near the same time point for each subject. All samples were analyzed by each proteomic platform except for one subject (*n*=35/36), whose CSF and plasma were analyzed only by SomaScan. This subject was excluded from all direct cross-platform comparisons, and therefore, such analyses were restricted to *N*=35 subjects. For both CSF and plasma, mass spectrometry measurements were performed using isobaric tandem mass tags (TMT-MS) with pre-fractionation^7^, including both with and without prior depletion of highly abundant proteins in each fluid. For PEA analyses, we used all thirteen qPCR-based human biomarker panels available through Olink, encompassing 1196 protein assays (1160 unique proteins). For aptamer-based analyses, we used the SomaScan assay (v4.1) from SomaLogic, which provides 7288 SOMAmers targeting 6596 unique proteins.

**Figure 1.**
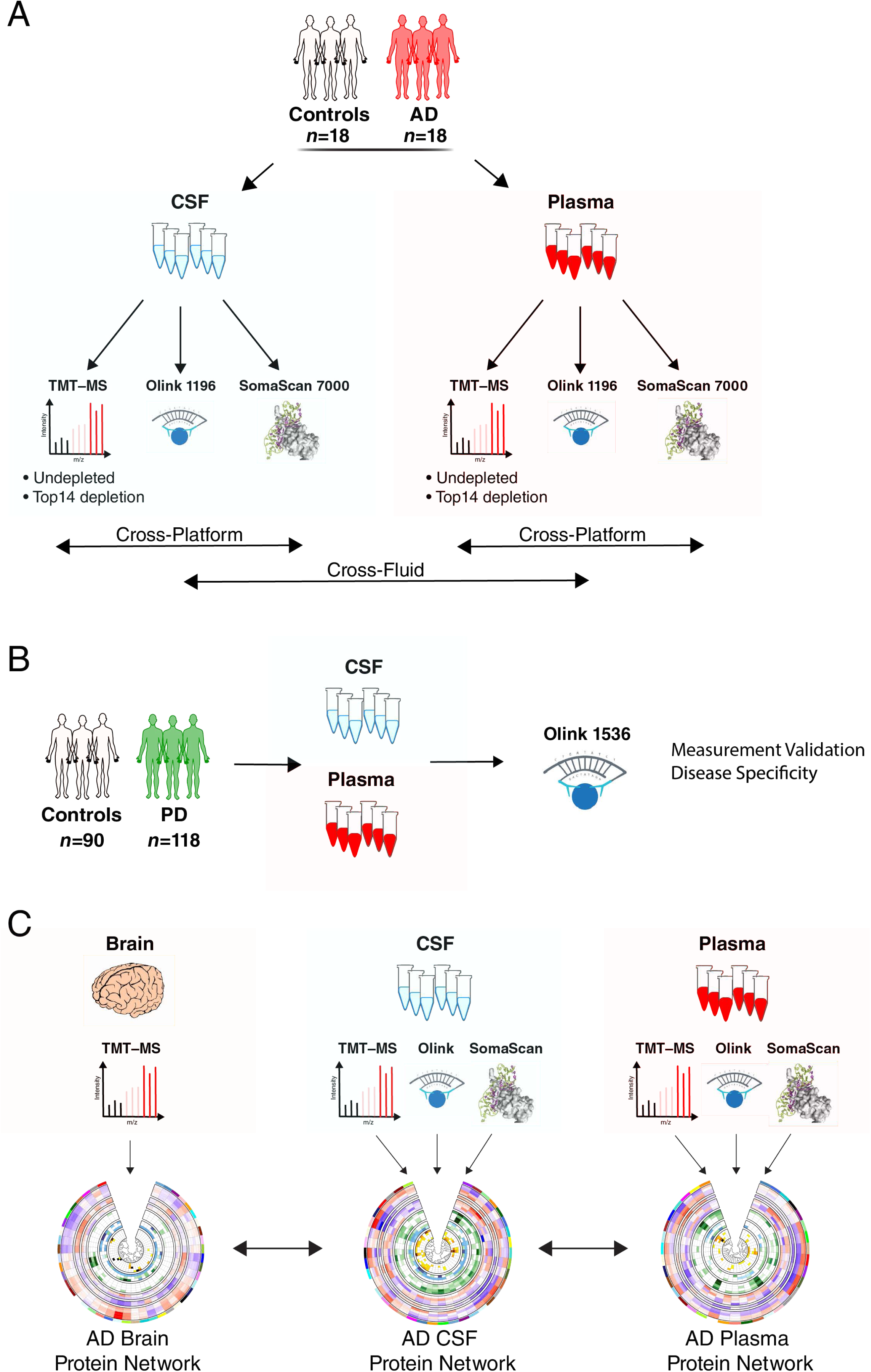
Study Overview. (A-C) Cerebrospinal fluid (CSF) and plasma sampled within subject at or near the same time point from control (*n*=18) and AD (*n*=18) subjects was analyzed by three proteomic platforms: untargeted tandem mass tag mass spectrometry (TMT-MS) on both undepleted fluid and fluid depleted of the top 14 most abundant proteins; proximity extension assay (PEA) targeting 1160 unique proteins (Olink 1196); and a modified aptamer-based assay targeting 6596 unique proteins (SomaLogic 7000). (B) Matched CSF and plasma from 90 control and 118 Parkinson’s disease (PD) subjects were analyzed by PEA targeting 1536 unique proteins (Olink 1536). (C) Protein co-expression networks for CSF and plasma were separately constructed using the union measurements from all three platforms in each biofluid. Networks were compared across biofluids and with an AD brain network previously described in Johnson *et al*.^2^

Analysis of the signal-to-noise (S:N) properties of each platform showed that a large proportion of the SomaScan measurements in CSF were at or near background noise level (**Supplementary Figure 1A**). By contrast, S:N was acceptable for nearly all SOMAmers in plasma (**Supplementary Figure 1B**). To address this limitation, we empirically determined a S:N cutoff for SOMAmers in CSF by correlating measurements in common across all three proteomic platforms at different SOMAmer S:N thresholds, and selected a S:N threshold where the correlations were maximized (**Supplementary Figure 1C**). This S:N threshold was 0.45. Applying this threshold to the SomaScan data reduced the number of quantified SOMAmers in CSF from 6776 to 3624 (**Supplementary Figure 1D, Supplementary Table 2**). This reduced set of SomaScan CSF measurements was used for most subsequent analyses.

We also analyzed missing measurements present in each platform across 36 CSF and plasma samples (**Supplementary Figure 2**). The Olink and SomaScan platforms had a similar increase in missing values across samples, which was greater in CSF than in plasma. TMT-MS suffered more from missing values in both fluids, particularly in plasma. In plasma undepleted of highly abundant plasma proteins, a maximum of approximately 500 quantified proteins was reached within 2 batches of TMT-MS. In plasma depleted of the top fourteen most abundant proteins, this threshold had nearly been reached at the point when all 36 samples had been analyzed. We decided to set a threshold of <75% missing values (or measurement in at least 9 out of 36 samples) for subsequent individual protein analyses, and exclude proteins with higher levels of missing values from consideration. Protein measurements that met this threshold were well balanced across AD and control cases. After applying the S:N and missingness filters, a large proportion (50.6%) of the SomaScan measurements in CSF, and a significant proportion (13.5%) of the TMT-MS measurements in depleted plasma, were removed from consideration for individual protein analyses.

In order to assess whether highly abundant protein depletion significantly affected TMT-MS measurements in CSF and plasma, we correlated protein values before and after depletion of these top fourteen most abundant proteins within subject (**Supplementary Figure 3A**). Median correlation after depletion in CSF was excellent in both CSF (*r*=0.78) and plasma (*r*=0.69), with greater variability introduced by depletion in plasma. Correlation was also good at the group level when proteins that were significantly altered in AD in either depleted or undepleted CSF (**Supplementary Figure 3B**) or plasma (**Supplementary Figure 3C**) were correlated with depletion *versus* no depletion. To avoid setting an arbitrary correlation threshold, we excluded all proteins measured by TMT-MS in CSF and plasma that had correlation values of zero or below (*i.e*., anticorrelated), or that had an opposite direction of change in AD, after highly abundant protein depletion. This totaled 32 proteins in CSF, and 27 proteins in plasma (**Supplementary Table 3**). The final protein abundance matrices used for individual protein analyses and cross-platform comparisons therefore included the <75% missingness filter across all platforms and fluids, the S:N filter for SomaScan CSF, and excluded proteins that were strongly affected by highly abundant protein depletion in CSF and plasma from TMT-MS measurements.

### Cross-Platform Comparisons

Across all three platforms, we were able to measure a total of 4655 unique proteins (as represented by unique gene symbols) in CSF, and 6794 unique proteins in plasma (**Figure 2A, Supplementary Tables 4-7**). The SomaScan platform provided the deepest proteomic coverage in plasma, measuring 4662 proteins not measured by Olink or MS. Most of the proteins that could be measured in plasma could also be measured in CSF by Olink (**Figure 2B, Supplementary Table 8**), whereas due primarily to our S:N filter, only approximately half of the proteins that could be measured in plasma by SomaScan could also be reliably measured in CSF (**Supplementary Table 9**). Over twice as many proteins could be measured in CSF compared to plasma on depleted fluid using TMT-MS due to the larger number of highly abundant proteins in plasma and the protein dynamic range limitations of unbiased discovery MS-based approaches. Only marginal improvement in proteomic coverage was observed in MS with depleted versus undepleted fluid in both CSF and plasma (**Supplementary Figure 4A, Supplementary Tables 10 and 11**), with depth of coverage improvement more apparent in CSF than in plasma (**Supplementary Figure 4B**). Ontologies of proteins uniquely measured by the SomaScan platform in CSF and plasma included nucleic acid metabolism and binding, and nucleus (**Supplementary Figure 4C**), suggesting that the platform was enriched for measurement of nuclear proteins. Ontologies unique to Olink included mitotic cell cycle, immune processes, and plasma membrane, reflecting selection bias of these biological pathways in the Olink platform compared to the other platforms. Ontologies unique to MS included transmembrane transport, complement, and cytoskeleton/structural proteins, representing more highly abundant proteins.

**Figure 2.**
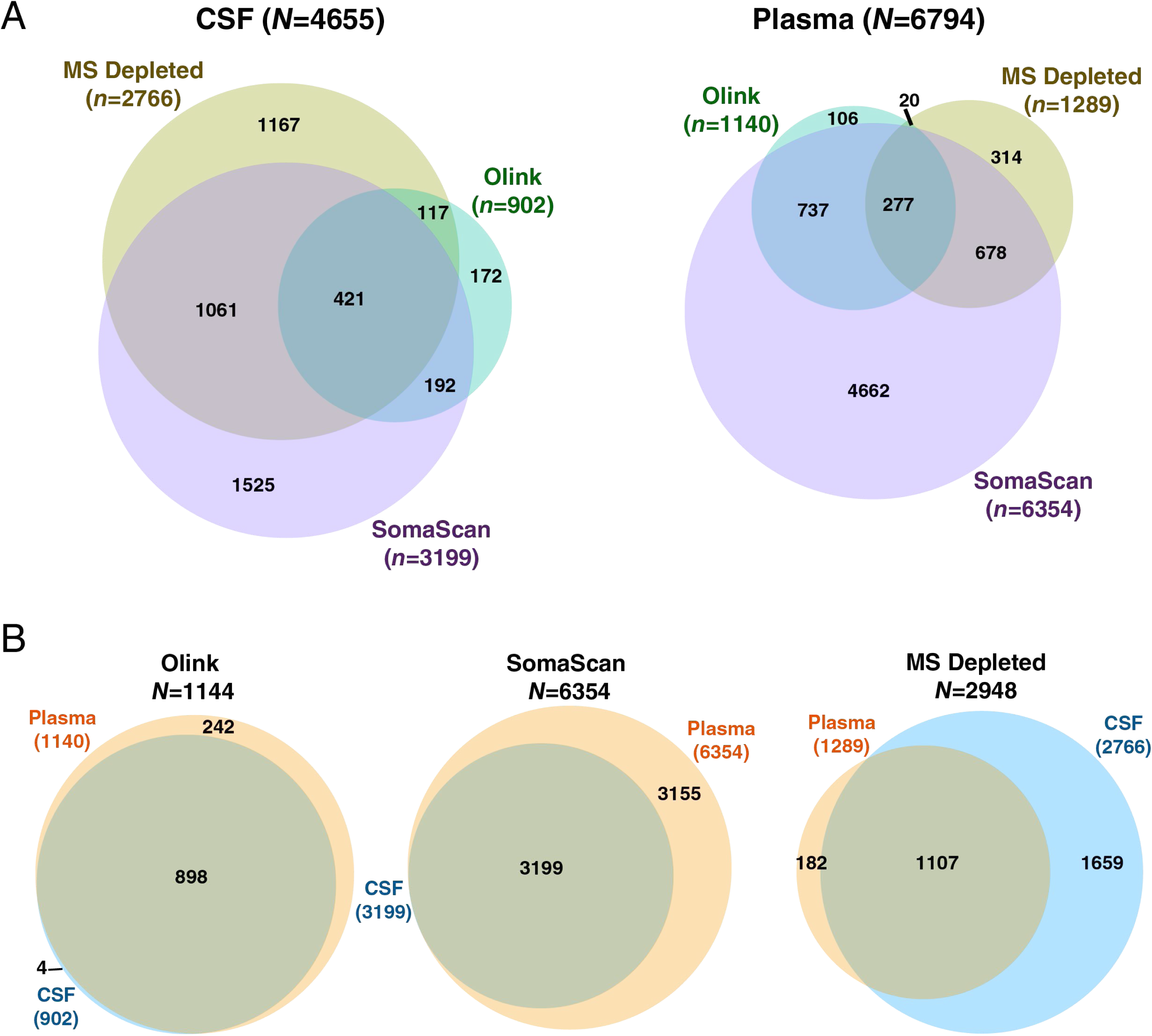
Depth and Overlap of Proteomic Coverage by Platform in CSF and Plasma. (A, B) Number and overlap of proteins measured by TMT-MS in depleted fluid, Olink 1196, and SomaScan 7000 platforms in CSF (left) and plasma (right) from *n*=36 subjects. The threshold for inclusion was measurement in at least 9 subjects (or missing values <75%). CSF measurements on the SomaScan 7000 platform underwent signal-to-noise filtering (**Supplementary Figure 1)** prior to subsequent analyses. (B) Number and overlap of proteins measured in CSF and plasma within each proteomic platform.

We correlated protein measurements across all three platforms in CSF and plasma (**Figure 3, Supplementary Tables 12-17, Extended Data**). Median correlation within subject was approximately 0.7 for CSF, with a similar distribution of correlation values among platforms. Median correlation was slightly lower in plasma at approximately 0.6, with more variability between the MS and affinity-based measurements than between SomaScan and Olink affinity-based measurements. Other than slightly lower overall correlation, these correlation patterns were generally similar when analyzed at the group level rather than within subject (**Supplementary Figure 5**). Improvement in median correlation was generally observed in plasma when only proteins that were significantly altered in AD in a given platform were used for correlation (**Figure 3B**), suggesting that inclusion of proteins with lower S:N led to decreased correlation. Interestingly, in CSF, the improvement in correlation was only observed with MS-based measurements. In summary, median correlation of proteomic measurements between platforms within the same subject was quite good, with better correlation in CSF than in plasma.

**Figure 3.**
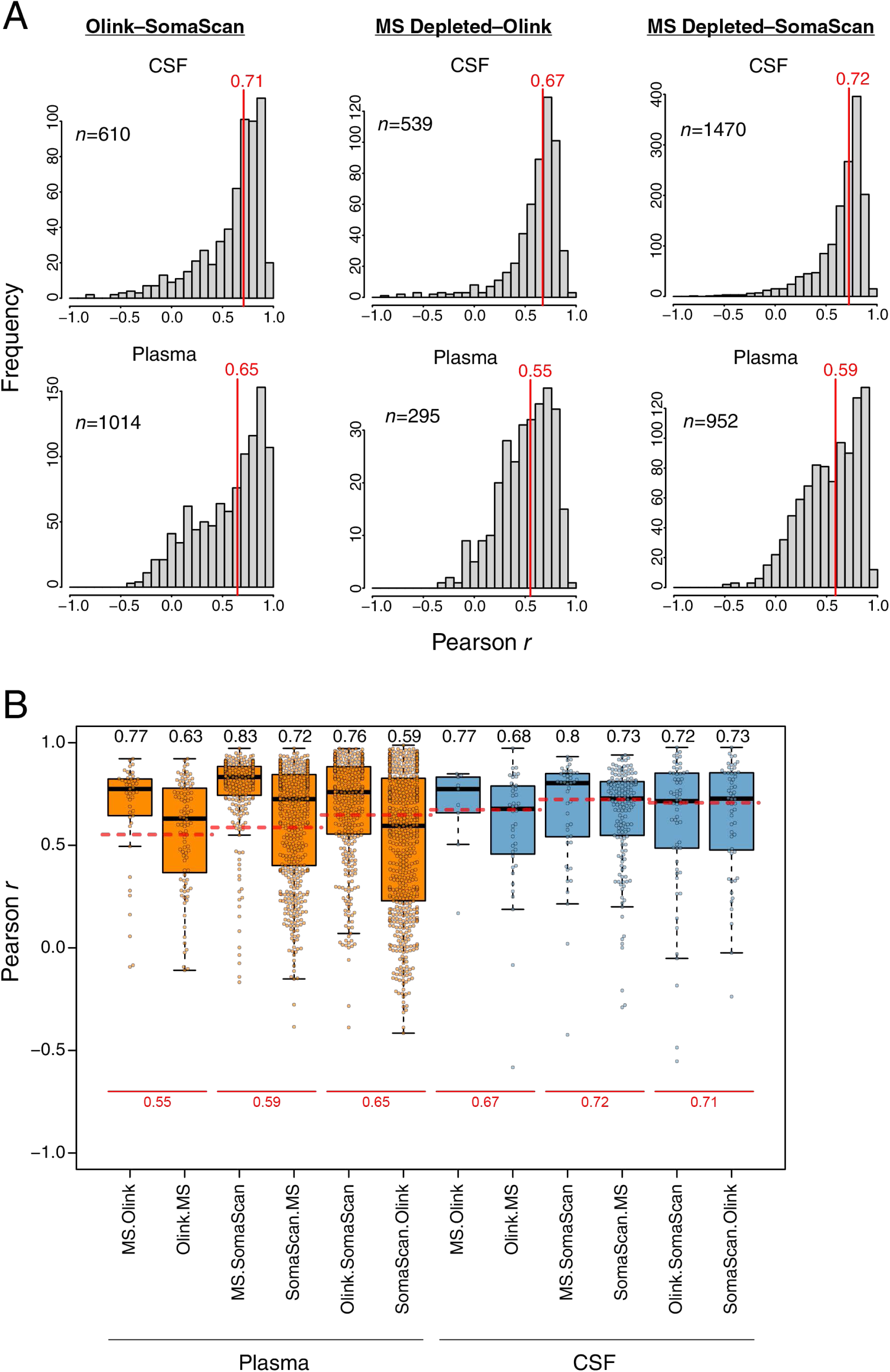
Correlation Between Platforms in CSF and Plasma. (A) Frequency distribution of correlation *r* values within subject between proteins commonly measured in the Olink and SomaScan platforms (left), MS depleted and Olink platforms (center), and MS depleted and SomaScan platforms (right), in CSF (top) and plasma (bottom). The vertical red lines indicate the median correlations. The number of proteins used for each correlation analysis is provided in the respective panel. All correlation values are provided in **Supplementary Tables 12-17,** with individual correlation plots provided in **Extended Data**. (B) Distribution of correlation *r* values for each cross-platform measurement in CSF and plasma considering only proteins that are significantly differentially expressed in a given platform. For each comparison, differentially expressed proteins are restricted to the first platform listed (e.g., MS.Olink indicates proteins differentially expressed by MS that have a corresponding measurement in Olink). The median correlation for each comparison is provided above the respective boxplot. The red dashed lines for each cross-platform measurement correspond to the median correlation considering all proteins as shown in (A), with the correlation value provided below in red. Boxplots represent the median, 25^th^, and 75^th^ percentiles, and box hinges represent the interquartile range of the two middle quartiles within a group. Datapoints up to 1.5 times the interquartile range from box hinge define the extent of whiskers (error bars).

To compare how proteomic measurements in our discovery cohort compared to Olink and SomaScan measurements in other AD cohorts, we performed correlation analyses with plasma Olink data from a Hong Kong-based cohort^18^, CSF and plasma Olink data from the BioFinder cohort^19^, and SomaScan plasma data from the AddNeuroMed cohort^20^ (**Supplementary Figure 6**). Correlations were restricted to proteins significantly altered in AD in each biofluid to maximize S:N. In AD plasma, correlation of Olink measurements in the Hong Kong cohort with our discovery cohort Olink measurements was excellent (*r*=0.82), with lower but strong correlation with SomaScan (*r*=0.57) and MS (*r*=0.63) measurements in our cohort (**Supplementary Figure 6A**). When comparing BioFinder Olink CSF measurements with our CSF measurements, correlation was also excellent across all measurement platforms (*r*∼0.7) (**Supplementary Figure 6B**). However, BioFinder Olink plasma measurements did not correlate with our discovery cohort platform plasma measurements (**Supplementary Figure 6C**). SomaScan plasma measurements in the AddNeuroMed cohort also did not correlate with any of our platform plasma measurements (**Supplementary Figure 6D**). These findings suggest that our cohort was most similar to the Hong Kong cohort, and that pre-analytical factors unique to each cohort likely significantly influenced the plasma measurements in each cohort.

### Proteins of Lower Abundance Are Decreased in AD Plasma

To determine which proteins were significantly altered across platforms in AD CSF and plasma, we performed differential abundance analyses within each fluid for each proteomic platform (**Figure 4, Supplementary Figure 7**). The analyses were performed without median normalization of overall protein abundance levels between AD and control cases, given that biomarker measurements in a clinical setting do not undergo median normalization^21, 22^. While a greater number of proteins were found to be decreased in AD CSF across all platforms, the decrease in plasma proteins in AD was much greater than in CSF and was strikingly apparent across all platforms (**Figure 4A**). This finding was consistent with the strong bias towards lower protein abundance observed in AD plasma in the Hong Kong cohort, in which the data had undergone some degree of median normalization prior to differential abundance analysis^18^. Overlap of differentially abundant proteins was low to modest across platforms (**Figure 4B**), due likely in part to the smaller size of the cohort and less statistical power to observe significant differences. Given the clear abundance differences observed in AD plasma across platforms, we further investigated this phenomenon by exploring which proteins were driving the difference in abundance. We first ranked each platform measurement by its contribution to the overall signal within each fluid, and tested the difference between the top 5% strongest signals in each platform compared with the total signal (**Supplementary Figure 8**). By this approach, we found that both the top 5% and overall protein levels were decreased in AD plasma by SomaScan and Olink, but not MS. However, because SOMAmer relative fluorescence units and Olink normalized protein expression values do not necessarily correlate with absolute protein abundance, we also calibrated measurements in each platform to known absolute protein concentrations in plasma obtained from the Human Protein Atlas^23^. After calibration, we observed that the top 5% most highly abundant proteins were not decreased in AD plasma in any platform, but that the overall decrease in AD plasma proteins was driven by proteins of lower abundance. This was consistently observed across platforms except for undepleted plasma analyzed by MS, in which many fewer proteins of lower abundance were measured. In summary, we observed a decrease in CSF and plasma proteins in AD compared to control, with the striking bias in plasma driven by proteins of lower abundance.

**Figure 4.**
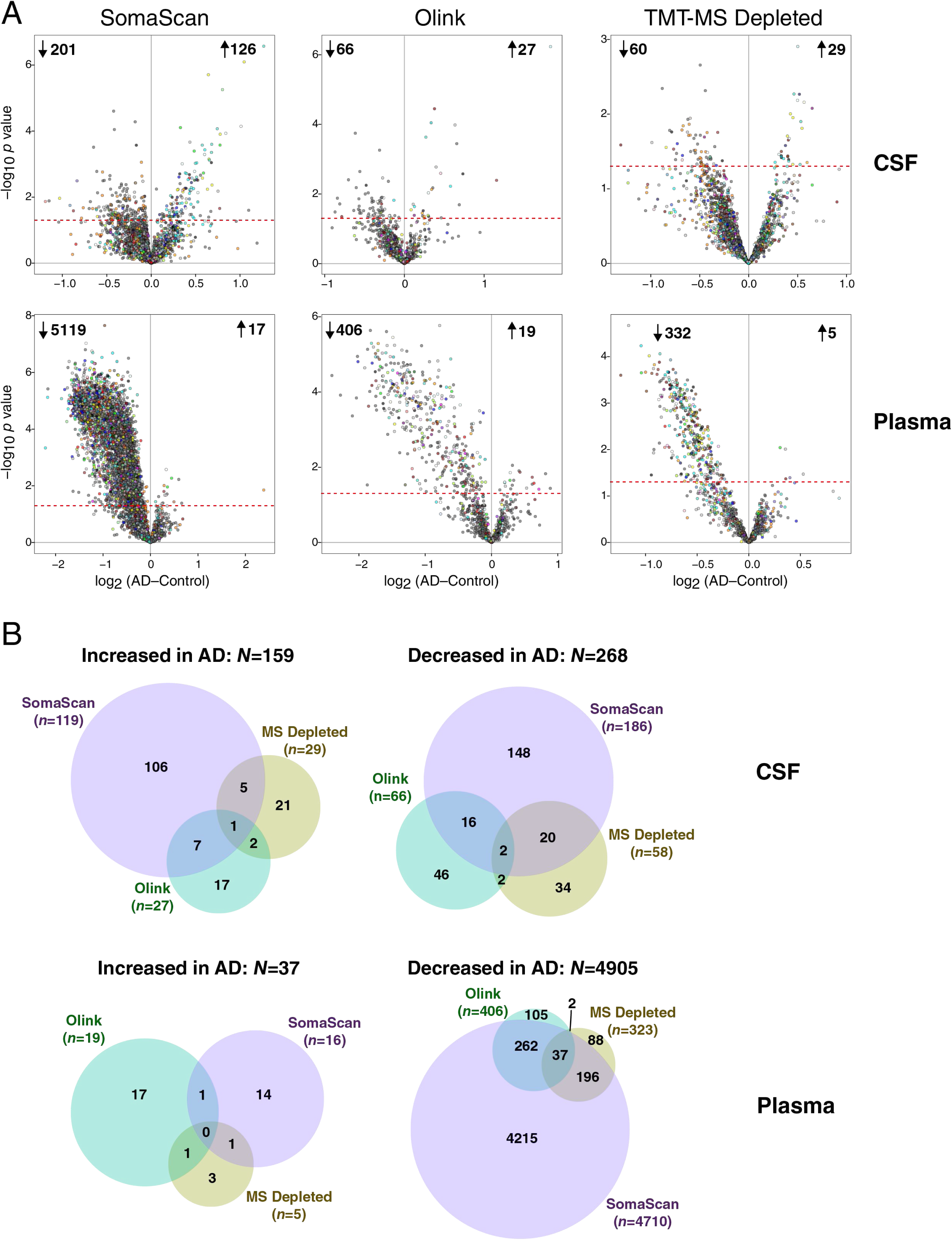
Differential Protein Abundance in AD by Platform in CSF and Plasma. (A) Differential protein abundance at the individual protein level between AD and control cases on the SomaScan (left), Olink (center), and TMT-MS Depleted (right) platforms, in CSF (top) and plasma (bottom). AD-control values less than 0 indicate decreased levels in AD, and values greater than 0 indicate increased levels in AD. Proteins that are above the dashed red line are significantly altered in AD by *t* test at *p*<0.05. Proteins are colored by the brain co-expression module in which they reside, as described in Johnson *et al*.^2^ (B) Overlap of differential protein expression in CSF and plasma across the three platforms. Overlap includes all commonly and uniquely measured proteins in each platform, and is restricted to unique gene symbols (i.e., excluding gene isoforms or redundant aptamers, which are included in (A)).

### Brain Protein Network Module Coverage by Platform in CSF and Plasma

We recently generated a consensus AD brain protein co-expression network from over 500 brain tissues as part of the Accelerating Medicines Partnership for Alzheimer’s Disease (AMP-AD) initiative that revealed many modules strongly correlated to AD neuropathological traits and cognitive decline^2^ (**Figure 5A**). To determine the potential for AD-relevant brain modules to be measured by markers in CSF and plasma, we calculated the percent coverage in CSF and plasma for each of the 44 brain network modules by proteomic platform (**Figure 5B**). All modules had at least some coverage in CSF, with SomaScan and MS providing the most module coverage in CSF compared to the Olink 1196 platform. In plasma, SomaScan provided the most module coverage. Brain module M26 complement/acute phase was particularly well covered by SomaScan and MS in both CSF and plasma, and M42 matrisome was well covered in both fluids by all platforms. Given that M42 matrisome had the strongest correlation to AD neuropathological traits in brain, we more closely examined this module across platforms in both fluids. M42 hub proteins—or proteins that contribute most to the module eigenprotein and are drivers of module co-expression—were generally well measured by all platforms in CSF (**Figure 6A**). In plasma, M42 hub protein coverage was best with SomaScan and Olink, and especially with SomaScan. SMOC1, which was the strongest driver of M42 co-expression, could be measured in CSF and plasma by both Olink and SomaScan, and measurements were well correlated in both fluids between the two platforms (**Supplementary Figure 9A, B**). Levels of SMOC1 were elevated in both AD CSF and plasma despite the decrease in lower abundant proteins in AD plasma (**Figure 6B**). We leveraged Olink CSF and plasma data from control (*n*=90) and Parkinson’s disease (PD, *n*=118) subjects in the Accelerating Medicines Partnership for Parkinson’s Disease (AMP-PD) consortium to test the specificity of SMOC1 for AD (**Figure 1B**). We did not observe an increase in SMOC1 in PD CSF, and observed a weak increase in PD plasma (**Figure 6C**). SMOC1 levels were generally well correlated between CSF and plasma within AD subjects but not controls (**Figure 6D**). CSF and plasma SMOC1 levels also correlated with Aβ/Tau levels in CSF (**Figure 6E**). In plasma, this correlation was driven by group differences, and was not significant within group (**Supplementary Figure 9C**). SMOC1 levels did not correlate strongly with cognitive function in AD (**Supplementary Figure 9D**) or PD (**Supplementary Figure 9E**). Interestingly, SMOC1 levels correlated weakly with age in both CSF and plasma (**Figure 6F**), an association which has been previously described in plasma^24, 25^.

**Figure 5.**
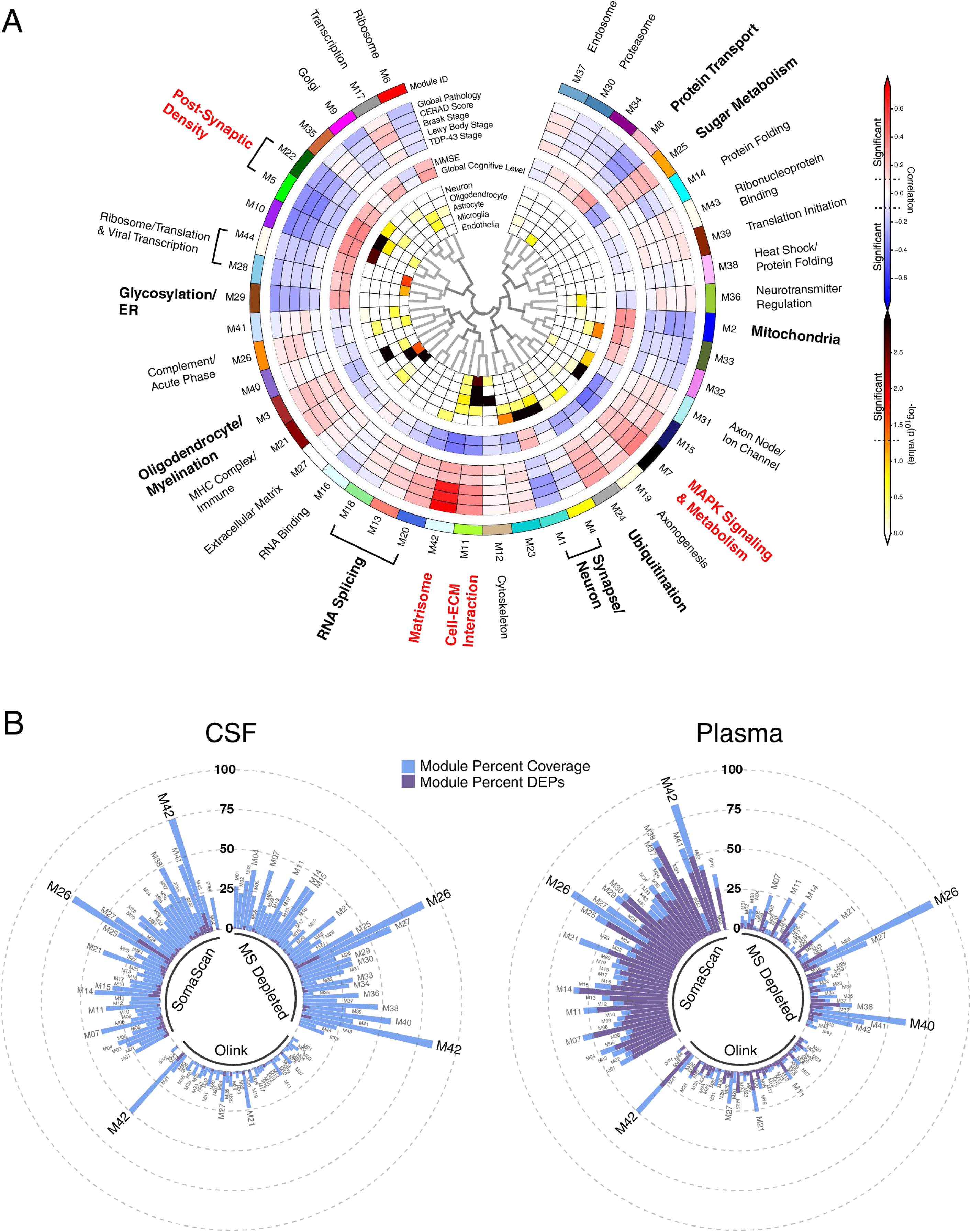
Brain Protein Network Module Coverage in CSF and Plasma by Platform. (A, B) AD brain protein co-expression network constructed from AD, asymptomatic AD, and control brains (total *n*=488) as described in Johnson *et al*.^2^ (A). The network was constructed using the weighted co-expression network algorithm (WGCNA), and consisted of 44 protein co-expression modules. Module relatedness is shown in the central dendrogram. The principal biology for each module was defined by gene ontology analysis. Modules that did not have a clear ontology were not assigned an ontology term. As previously described in Johnson *et al*., brain module eigenproteins were correlated with neuropathological traits in the ROSMAP and Banner cohorts (global pathology, neuritic amyloid plaque burden (CERAD score), tau tangle burden (Braak stage), lewy body burden, and pathological TDP-43 burden); cognitive function (mini-mental state examination (MMSE) score and global cognitive function, higher scores are better); and brain weight and the number of *APOE* ε4 alleles (0, 1, 2). Red indicates positive correlation, whereas blue indicates negative correlation, with statistical significance threshold at *r*=0.1 (dotted lines). Twelve of the 44 modules that were most highly correlated to neuropathological and/or cognitive traits are in bold, with the four most strongly trait-related modules highlighted in red. The cell type nature of each module was assessed by module protein overlap with cell type specific marker lists of neurons, oligodendrocytes, astrocytes, microglia, and endothelia. (B) Percent module coverage (dashed rings) by each analysis platform in CSF (left) and plasma (right). The percent of module proteins that are measured as differentially expressed proteins (DEPs) by each platform is shown in purple.

**Figure 6.**
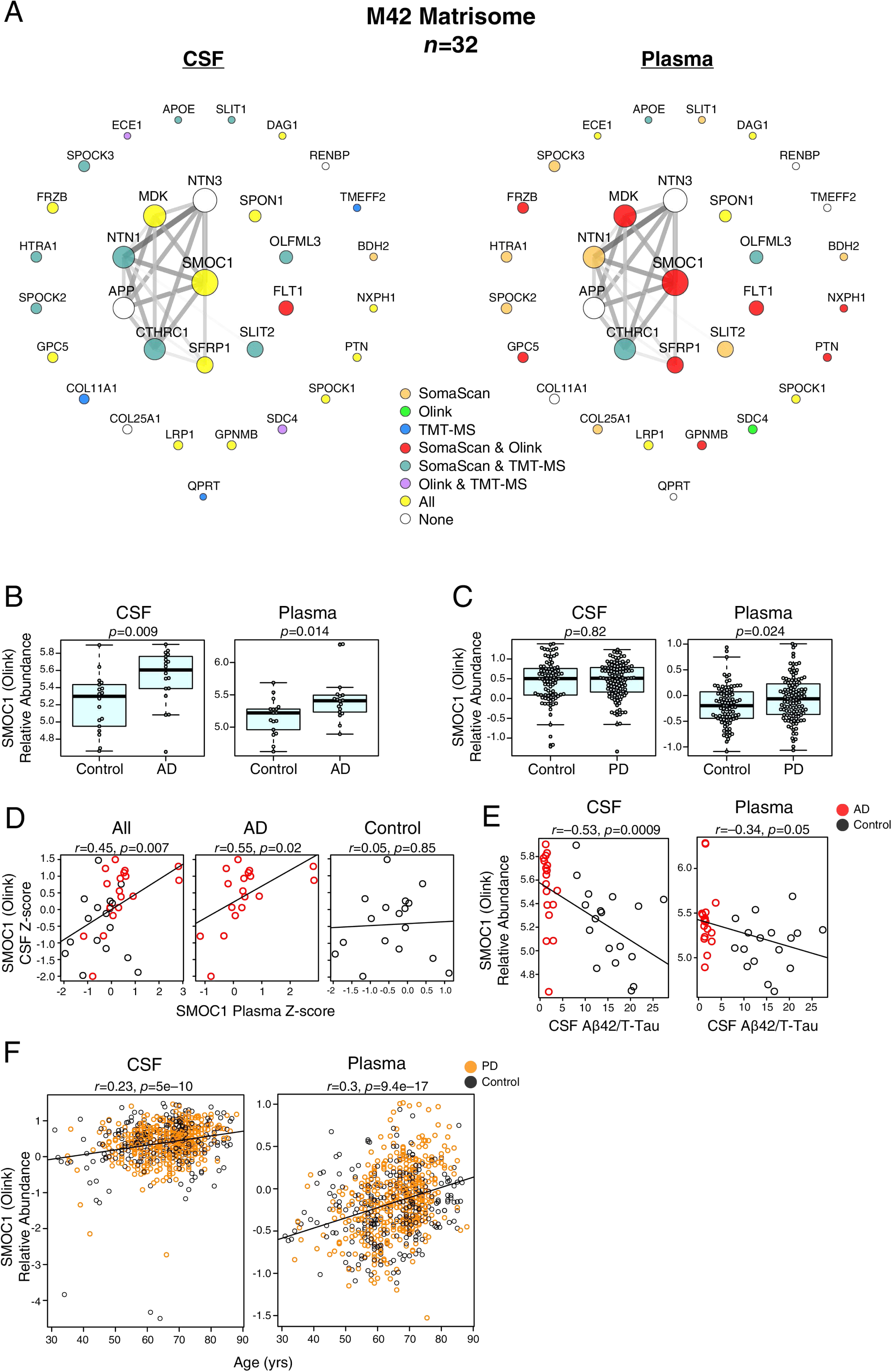
Brain M42 Matrisome Coverage and SMOC1 Levels in CSF and Plasma. (A-F) Coverage of the M42 matrisome module by proteomic platform in CSF and plasma (A). (B) Differences in SMOC1 relative abundance as measured by Olink between control and AD cases in CSF (left) and plasma (right). (C) Differences in SMOC1 relative abundance as measured by Olink between control and PD cases in CSF (left) and plasma (right) in the Accelerating Medicines Partnership – Parkinson’s Disease (AMP-PD) cohort. (D) Correlation of relative SMOC1 levels between CSF and plasma. (E) Correlation of SMOC1 relative abundance with CSF Aβ/T-Tau ratio in CSF (left) and plasma (right). (F) Correlation of SMOC1 relative abundance with age in CSF (left) and plasma (right) in the AMP-PD cohort. Differences between groups were assessed by *t* test. Correlations were performed using Pearson correlation. Aβ, amyloid-β; TMT-MS, tandem mass tag mass spectrometry; SMOC1, SPARC-related modular calcium-binding protein 1; T-Tau, total tau.

Hub proteins of other AD-relevant brain co-expression modules could also be measured in CSF and plasma (**Figure 7**). These included HOMER1 in the M5 Post-Synaptic Density module (**Supplementary Figure 10**), NEFL in the M3 Oligodendrocyte/Myelination module (**Supplementary Figure 11**), CHI3L1—also known as YKL-40—in the M21 MHC Complex/Immune module (**Supplementary Figure 12**), YWHAZ in the M4 Synapse/Neuron module (**Supplementary Figure 13**), ENO1 in the M7 MAPK Signaling/Metabolism Module (**Supplementary Figure 14**), and PEBP1 in the M25 Sugar Metabolism Module (**Supplementary Figure 15**). All of these proteins were increased in AD CSF, yet only NEFL and CHI3L1 were also increased in AD plasma, demonstrating the diversity of potential AD biomarker changes across different fluids. Furthermore, not all proteins within a brain module were found to behave similarly in CSF and plasma. For instance, YWHAZ in the M4 Synapse/Neuron module was observed to be increased in CSF and decreased in plasma, whereas NPTXR, another M4 protein, was found to be decreased in both CSF and plasma (**Supplementary Figure 16**). NPTXR also illustrated a discrepancy in platform measurements, where the Olink measurement was significantly negatively correlated with the MS and SomaScan measurements in CSF, but was more similar to the SomaScan measurement than the MS measurement in plasma (**Supplementary Figure 16A**). Another example of a protein with discrepancy in measurements across fluids was SPP1 in the M21 MHC Complex/Immune module (**Supplementary Figure 17**). SPP1 measurements best correlated between MS and SomaScan in CSF, with the Olink measurement being anticorrelated to the other two platforms. However, in plasma, SomaScan and Olink SPP1 measurements correlated well, whereas MS did not with either affinity-based platform. NEFL was also highly correlated among all platforms in CSF, but was anticorrelated between SomaScan and Olink in plasma. As has been previously demonstrated, NEFL levels were strongly correlated with increasing age^26^ (**Supplementary Figure 11F**). In summary, AD brain co-expression module proteins could be measured by all three platforms in CSF and plasma, but SomaScan had the best coverage in plasma, especially for the M42 matrisome module. SMOC1, a hub of M42, was elevated in both AD CSF and plasma and levels correlated within subject between CSF and plasma. Other AD brain module protein hubs could also be measured in CSF and plasma, but opposite directions of change were often observed between CSF and plasma protein levels of these hubs, and protein measurements did not always positively correlate across proteomic platforms.

**Figure 7.**
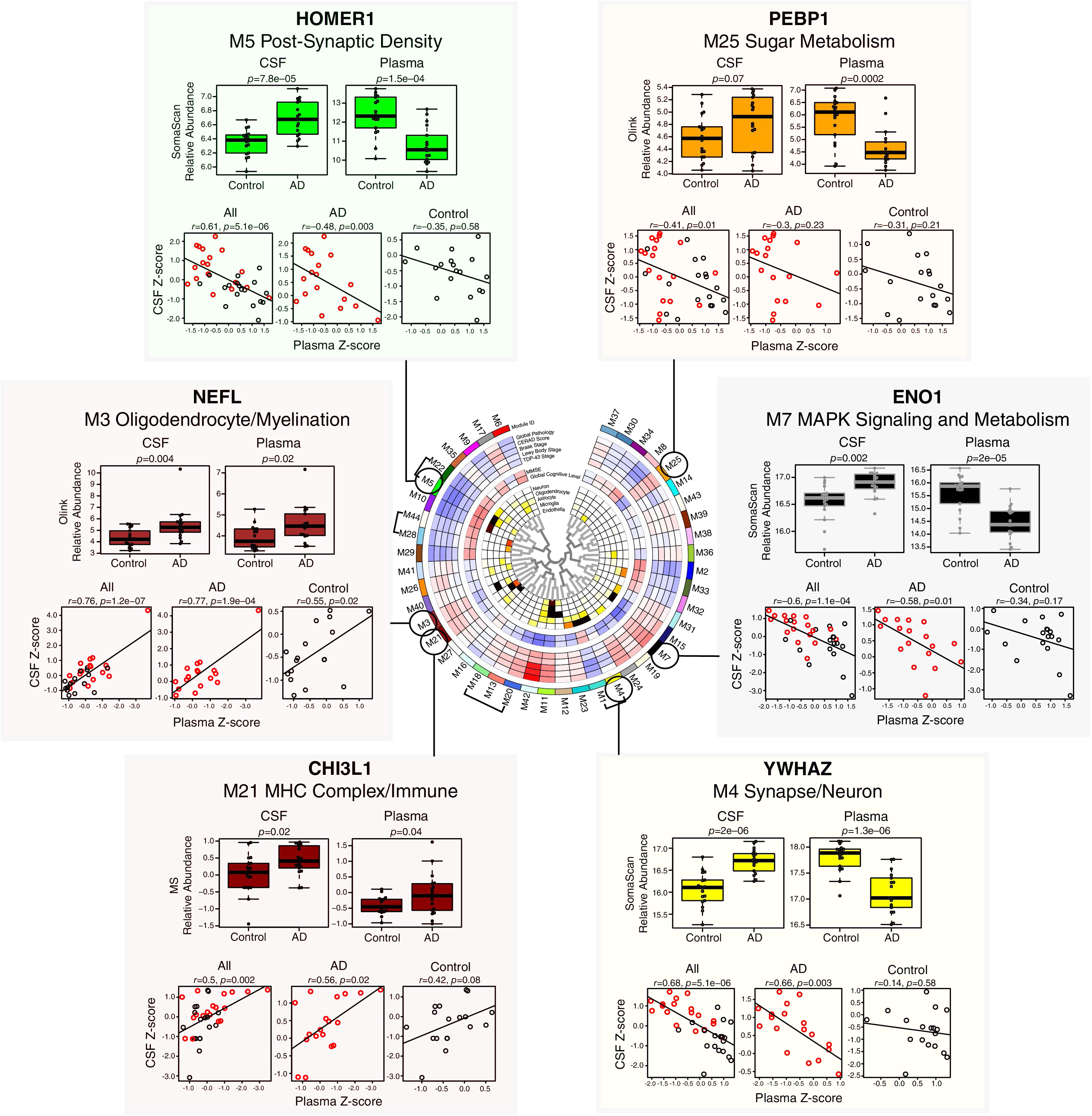
AD Brain Module Proteins in CSF and Plasma. Proteins that are members of AD brain co-expression modules as described in Johnson *et al*.^2^ and can also be measured in CSF and plasma by one or more proteomic platforms are illustrated for selected disease-related modules. Further details of the brain network are provided in Figure 5. Relative abundance differences between AD and control in CSF and plasma for each marker were determined by *t* test, and correlation between relative levels in CSF and plasma within subject was performed by Pearson correlation. CHI3L1, Chitinase-3-like protein 1; ENO1, alpha-enolase; HOMER1, Homer protein homolog 1; NEFL, neurofilament light polypeptide; PEBP1, phosphatidylethanolamine-binding protein 1; YWHAZ, 14-3-3 protein zeta.

### AD CSF Co-Expression Network Reveals Strong Disease-Related Modules Reflecting Proteostasis, Synaptic, Complement, and Sugar Metabolism Pathophysiology

While AD-related protein co-expression modules have been reliably identified in brain, to date it has been unclear whether co-expression modules related to AD are also present in biofluids. To address this question, we leveraged all three proteomic platforms and harmonized their measurements by median normalization into separate protein abundance matrices for CSF and plasma, and used these harmonized abundance matrices to build co-expression networks for each fluid. We were then able to compare these networks to one another, and to the consensus AD brain network (**Figure 1C**). Using this approach, we built a CSF co-expression network from 7158 protein assays targeting 4154 unique gene symbols (**Figure 8A, Supplementary Tables 18 and 19, Extended Data**). The network consisted of 38 modules, with each platform contributing measurements to nearly all modules (**Supplementary Figure 18**). Modules that were most strongly correlated to Aβ and tau pathological measures in CSF and/or cognitive function included M15 post-synaptic membrane, M8 autophagy, M7 SNAP receptor/SNARE complex, M32 synaptic membrane/matrisome, M16 sugar metabolism, M29 sugar metabolism/actin depolymerization, M24 ubiquitination, M26 TGF-β signaling, and M3 complement/protein activation cascade. M8 autophagy and M24 ubiquitination modules were particularly strongly correlated to total tau and p-tau181 levels. The M8 autophagy module contained microtubule associated protein tau (MAPT) and SMOC1 as members, as well as other markers previously associated with AD such as NEFL and PEBP1 (**Figure 8B**). M8 module eigenprotein levels were strongly negatively correlated with cognitive function (*r*= –0.67) and Aβ42/tau ratio (*r*= –0.82), and strongly positively correlated with total tau (*r*=0.86) and p-tau181 (*r*=0.78) (**Figure 8C**), reflecting its close association to AD brain amyloid-β and tau pathology. Among the other modules that correlated strongly with traits were M29 sugar metabolism/actin depolymerization, which was most strongly correlated to *APOE* ε4, and M26 TGF-β signaling, which was strongly positively correlated with age. The M8 autophagy and M15/M32 synaptic modules were enriched in neuronal and oligodendrocyte cell type markers, potentially reflecting the brain cell type origin of these CSF modules. The M24 ubiquitination and M29 sugar metabolism/actin depolymerization modules did not have cell type character, whereas the M3 complement/protein activation cascade module was enriched in endothelial and microglial markers.

**Figure 8.**
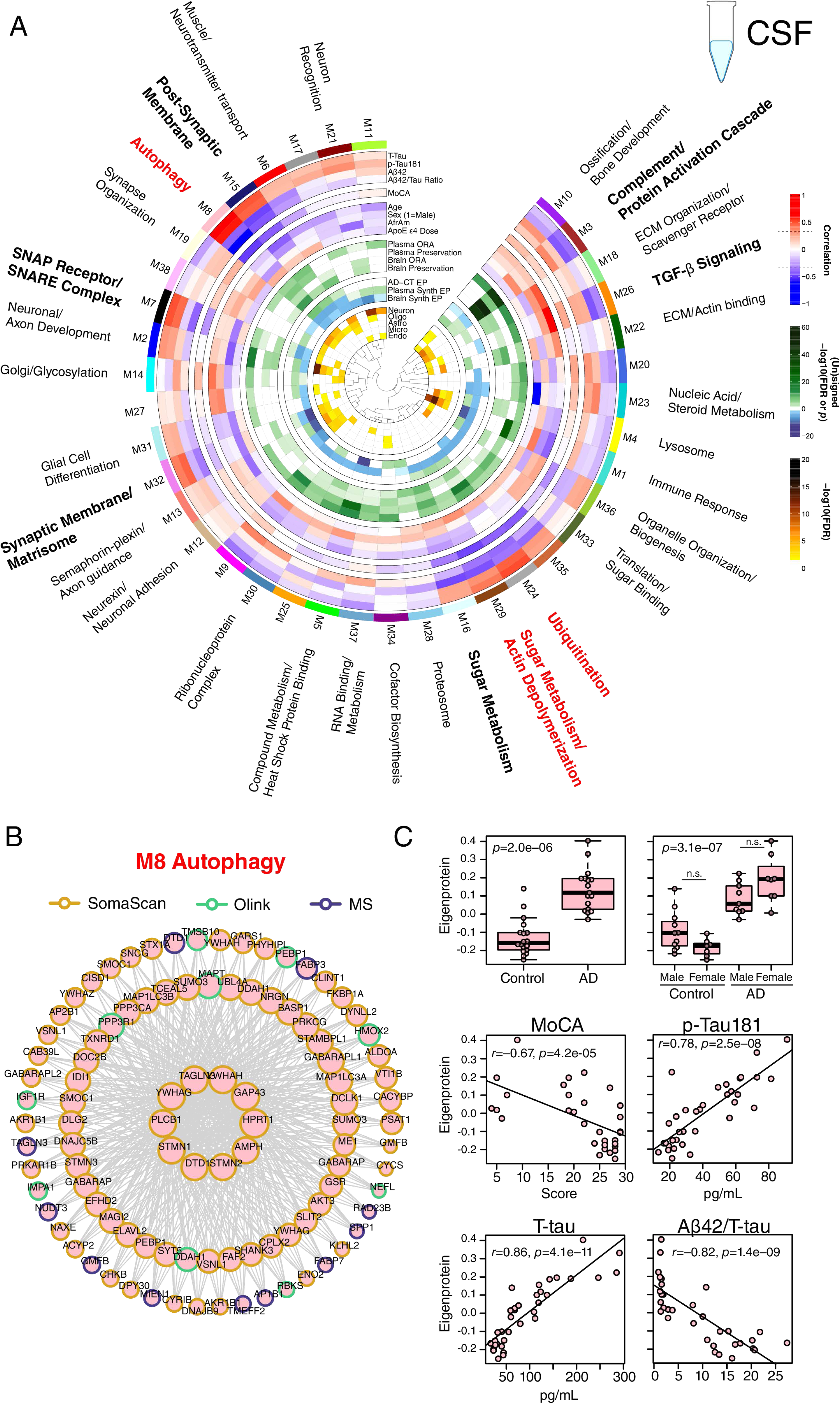
AD CSF Protein Co-Expression Network. (A-C) Control (*n*=18) and AD (*n*=18) CSF samples were analyzed by TMT-MS, Olink, and Somalogic proteomic platforms, and a protein co-expression network was constructed using the union protein measurements harmonized across all three platforms (*n*=3594 SomaScan, 902 Olink, and 2662 TMT-MS depleted protein measurements, total *n*=7158 measurements, *n*=4154 unique gene symbols). The network consisted of 38 protein co-expression modules. Module relatedness is shown in the central dendrogram. The principal biology for each module was defined by gene ontology analysis. Modules that did not have a clear ontology were not assigned an ontology term. CSF network module eigenproteins were correlated with CSF total tau levels (T-Tau), CSF phospho-tau 181 (p-Tau181) levels, CSF amyloid beta 42 (Aβ42) levels, CSF Aβ42/Tau ratio, Montreal Cognitive Assessment Score (MoCA, higher scores represent better cognitive function), age, sex (1=male, 0=female), African American (AfrAm) status, and the number of apolipoprotein E epsilon 4 alleles (0, 1, 2; ApoE ε4 dose). Red indicates positive correlation, whereas blue indicates negative correlation, with statistical significance threshold at *r*=0.35 (dotted lines). Nine of the 38 modules that were most highly correlated to CSF pathological or cognitive traits are in bold, with the three most strongly trait-related modules highlighted in red. CSF modules were tested for their presence in the plasma and brain networks by module protein overrepresentation analysis (ORA) and network preservation (preservation) statistics, as previously described^2^. Only modules that reached statistical significance after FDR correction are colored by degree of significance. ORA *p* values are for the module with the strongest overlap. In addition to module overlap and preservation analyses, the difference in CSF module eigenprotein between control and AD, or CSF synthetic eigenprotein in plasma and brain between control and AD, was determined. A significantly increased eigenprotein in AD is indicated in green, whereas a significantly decreased eigenprotein is indicated in blue. The cell type nature of each module was assessed by module protein overlap with cell type specific marker lists of neurons, oligodendrocytes, astrocytes, microglia, and endothelia. (B) The top 100 proteins by module eigenprotein correlation value (kME) for the M8 Autophagy module (*n*=156 proteins). Larger nodes represent larger kME values, or module hubs. Module proteins are highlighted according to the proteomic platform in which they were measured. (C) M8 Autophagy eigenprotein levels between control and AD (top panels), and eigenprotein correlation with MoCA and p-Tau181 (middle panels) and T-tau and Aβ42/T-tau levels (bottom panels). Eigenprotein differences between sexes in control and AD were not significant (*p*=0.13). Further information on the CSF network is provided in **Supplementary Tables 18 and 19, and Extended Data**. Boxplots represent the median, 25^th^, and 75^th^ percentiles, and box hinges represent the interquartile range of the two middle quartiles within a group. Datapoints up to 1.5 times the interquartile range from box hinge define the extent of whiskers (error bars). Differences between groups were assessed by *t* test or one-way ANOVA with Tukey test. Correlations were performed using Pearson correlation. n.s., not significant.

We tested whether CSF modules were present in brain or plasma by two different approaches: over-representation analysis (ORA), and network preservation statistics. We also tested how the module eigenproteins changed in CSF between AD and control, and whether the cognate module eigenprotein (or “synthetic” eigenprotein) in plasma and brain were altered in AD (**Figure 8A, Extended Data**). The CSF M3 complement/protein activation cascade module was most strongly preserved in plasma and brain, and was decreased in AD CSF. M26 TGF-β signaling was also decreased in AD CSF. Modules that were increased in AD CSF included the M15 and M32 synaptic, and M8 and M24 proteostasis modules, along with M16 sugar metabolism. Most of the synthetic eigenproteins for these modules were decreased in plasma except for the M3 complement/protein activation cascade module, which was increased in both plasma and brain.

In summary, we were able to construct an AD CSF protein co-expression network from >7000 protein assays in CSF which revealed strong disease-associated modules related to proteostasis, synaptic biology, sugar metabolism, and complement. All module eigenproteins were increased in CSF and decreased in plasma except for complement, which was decreased in CSF and increased in plasma.

### AD Plasma Co-Expression Network Reveals Strong Disease-Related Modules Reflecting Endocytosis and Matrisome Pathophysiology

The plasma network included 9589 protein assays targeting 6614 unique gene symbols, and consisted of 35 modules with good platform measurement representation across the network, similar to the CSF network (**Figure 9A, Supplementary Tables 20 and 21, Supplementary Figure 18**). The SomaScan platform contributed approximately 80% of the measurements in the network. A striking feature of the plasma network was the number of modules related to extracellular matrix biology and the matrisome that correlated with AD CSF Aβ and tau biomarkers. One such module was the M33 adhesion/ECM/wound response module whose eigenprotein—along with other matrisome-related module eigenproteins—was elevated in AD plasma despite the decrease in lower abundance plasma proteins in AD (**Figure 9B, C**). M33 module co-expression was driven by tenascin (TNC), an extracellular matrix protein involved in neuronal migration and regeneration, as well as synaptic plasticity. TNC was measured by ten separate assays across the three platforms, eight of which were SOMAmers. Eight of the ten TNC assays fell within M33, including six SOMAmers and one Olink and one MS measurement, suggesting good correlation for most TNC measurements across platforms. SPP1 as measured by Olink and SomaScan were also members of M33. Another module strongly related to AD was the M24 endocytosis module, which was the module most strongly correlated to total tau levels in CSF. Interestingly, this module was not strongly preserved in CSF or brain, potentially reflecting a more systemic process associated with AD. Plasma modules were generally less well preserved in CSF and brain than CSF modules were preserved in plasma and brain. Brain protein co-expression was generally not strongly preserved in CSF or plasma except for the complement module, which was highly preserved across all tissues (**Supplementary Figure 19 and 20, Supplementary Table 22**).

**Figure 9.**
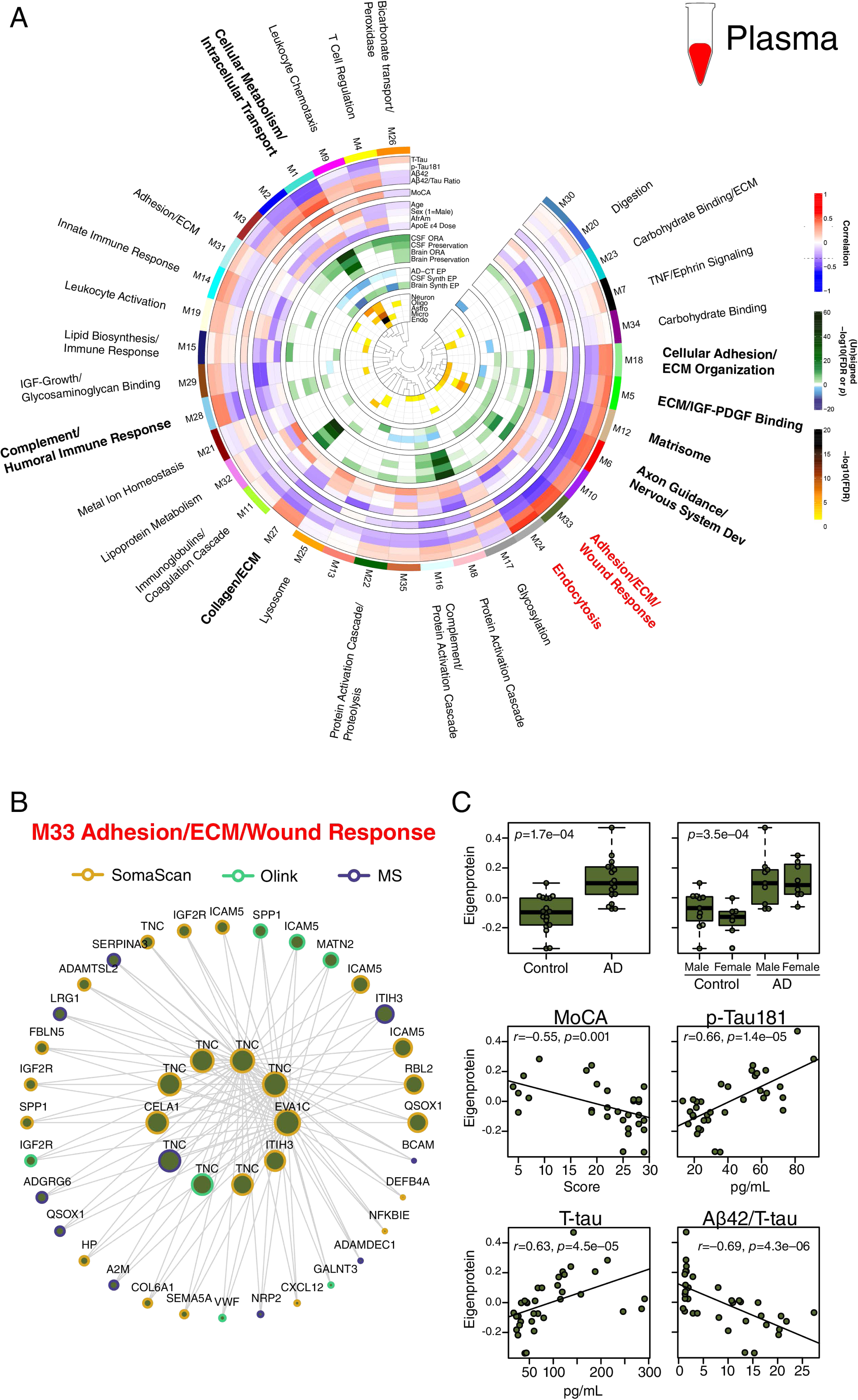
AD Plasma Protein Co-Expression Network. (A-C) Control (*n*=18) and AD (*n*=18) plasma samples from *n*=35/36 of the same subjects from which the CSF network was generated were similarly analyzed by TMT-MS, Olink, and Somalogic proteomic platforms, and a protein co-expression network was constructed using the union protein measurements harmonized across all three platforms (*n*=7284 SomaScan, 1140 Olink, and 1165 TMT-MS depleted protein measurements, total *n*=9589 measurements, *n*=6614 unique gene symbols). The network consisted of 35 protein co-expression modules. Nine of the 35 modules that were most highly correlated to CSF pathological or cognitive traits are in bold, with the two most strongly trait-related modules highlighted in red. Module eigenprotein correlations with traits, preservation analyses, eigenprotein differences between AD and control, and cell type overlap tests were performed in the same fashion as the CSF network as described in Figure 8. Correlation was performed with CSF levels of T-Tau, p-Tau181, Aβ42, and Aβ42/T-Tau ratio. (B) The 42 proteins comprising the M33 Adhesions/ECM/Wound Response module. Larger nodes represent larger kME values, or module hubs. Module proteins are highlighted according to the proteomic platform in which they were measured. (C) M33 Adhesion/ECM/Wound Response levels between control and AD (top panels), and eigenprotein correlation with MoCA and CSF p-Tau181 (middle panels) and T-tau and Aβ42/T-tau levels (bottom panels). Further information on the plasma network is provided in **Supplementary Tables 20 and 21, and Extended Data**. Boxplots represent the median, 25^th^, and 75^th^ percentiles, and box hinges represent the interquartile range of the two middle quartiles within a group. Datapoints up to 1.5 times the interquartile range from box hinge define the extent of whiskers (error bars). Differences between groups were assessed by *t* test or one-way ANOVA with Tukey test. Correlations were performed using Pearson correlation.

We compared our 3-platform plasma network using module ORA to a serum network built from approximately 5000 SOMAmers previously reported by Emilsson *et al*.^27^ (**Supplementary Figure 21**). The serum network had fewer modules than the plasma network (27 *versus* 38). Over half of the serum modules had significant overlap in plasma by ORA. One of these modules was serum module 11—a lipid module with many module protein levels affected by variation in the *APOE* locus. Serum module 11 overlapped with plasma modules M15 Lipid Biosynthesis/Immune Response and M32 Lipoprotein Metabolism. M15 co-expression was driven by ApoE and levels of this module decreased most strongly with increasing number of *APOE* ε4 alleles, whereas M32 co-expression was strongly driven by ApoB and module levels increased most strongly with the number of *APOE* ε4 alleles (**Figure 9A**). Therefore, the 3-platform plasma network provided sufficient resolution to identify lipoprotein-related protein co-expression modules divergent in their relationship to *APOE* ε4 genotype.

Given our findings with the discrepancy in levels of individual AD-related proteins between CSF and plasma, we tested whether CSF and plasma co-expression modules also showed discrepancy in the levels of their eigenproteins and synthetic eigenproteins in the paired fluid (**Figure 10**). Like many individual proteins, we also observed an inverse relationship of module levels in plasma compared to the levels in CSF in AD. One notable exception was the M3 complement/protein activation cascade module, where the within-subject module eigenprotein was increased in AD plasma compared to AD CSF. In plasma, the within-subject eigenprotein relationship to CSF was noisier, and did not show a strong discrepancy between fluids for most modules. An exception again was the plasma M8 protein activation cascade module, which was increased in AD plasma compared to CSF for most subjects despite the general decrease in protein abundances in AD plasma.

**Figure 10.**
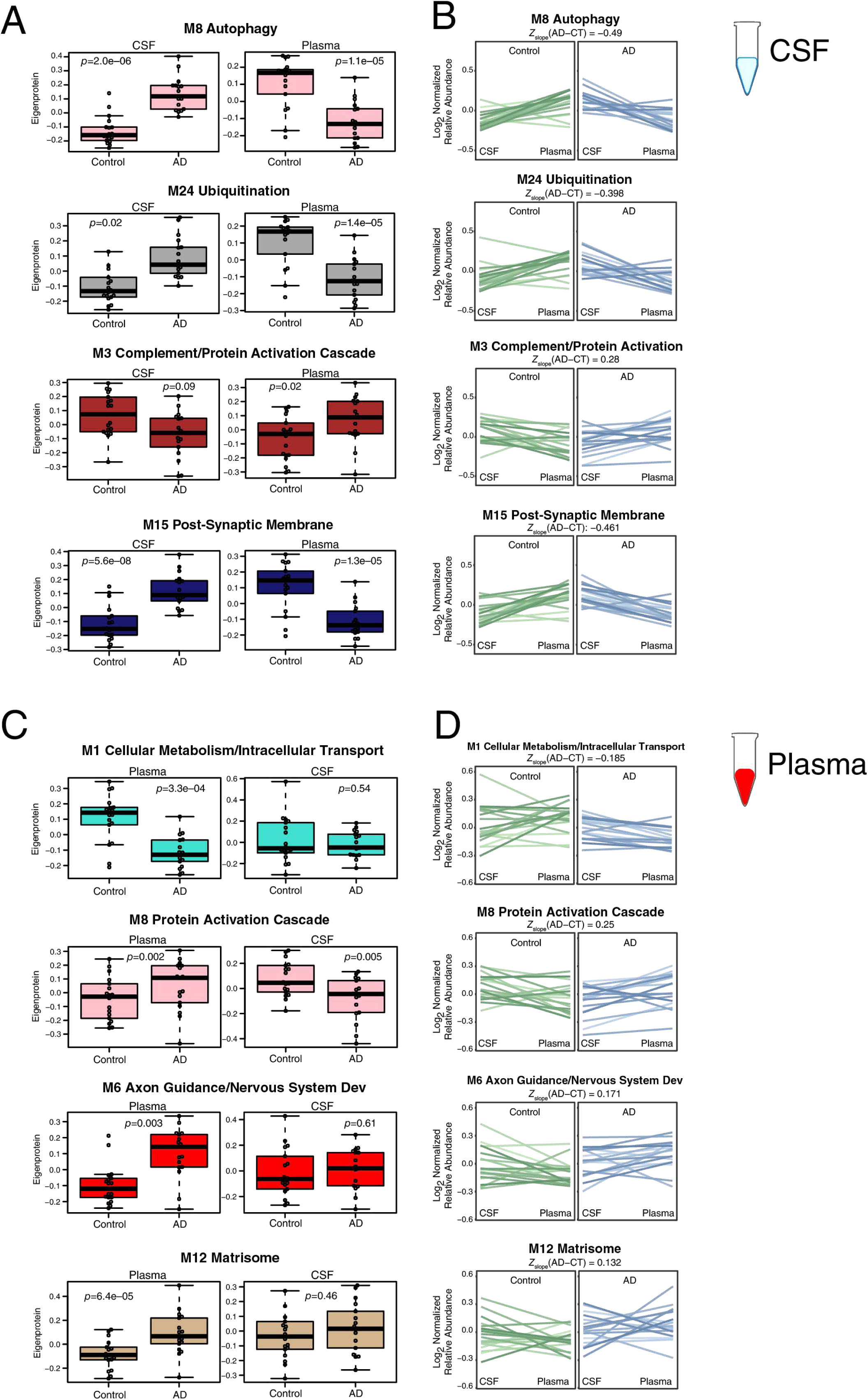
Cross-Fluid Eigenproteins. (A-D) CSF protein network module eigenproteins and their synthetic eigenproteins in plasma were compared between control and AD groups (A). (B) Relative eigenprotein levels were compared within subject across fluids in control and AD cases. The difference in average slope (*Z*_slope_) between AD and control was calculated for each module. (C) Plasma protein network module eigenproteins and their synthetic eigenproteins in CSF were compared between control and AD groups. (D) Relative eigenprotein levels were compared within subject across fluids in control and AD cases. *n*=18 control, 17 AD for all analyses. Selected modules shown include those that had significant AD trait relationships and were also preserved in CSF by at least one preservation method. Boxplots, within subject cross-fluid eigenproteins, and *Z*_slope_ calculations for all CSF and plasma modules are provided in **Extended Data.** Differences between groups were assessed by *t* test.

In summary, we were able to build AD CSF and plasma protein co-expression networks using measurements from all three proteomic platforms, providing excellent proteomic depth in each fluid. Synaptic, proteostasis, sugar metabolism, and complement modules were strongly altered in AD CSF, while matrisome, endocytosis, and complement modules were altered in AD plasma. The complement modules were the modules best preserved across brain, CSF, and plasma, and were observed to be increased in brain and plasma, but decreased in CSF. AD-related CSF module eigenproteins were generally increased in AD but decreased in plasma, likely due in part to the overall decrease of low abundance plasma proteins in AD. The relationship of plasma modules to CSF was more variable, reflecting the contribution of many tissues other than brain to plasma protein co-expression.

## Discussion

In this study, we used three different proteomic technologies to interrogate matched CSF and plasma from a discovery cohort of AD subjects in order to identify and assess promising protein biomarkers for AD. We found that overall correlation among the proteomic platforms was good, with weaker correlation in plasma. We observed a general decreased expression level of lower abundance proteins in AD CSF and plasma, most notably in plasma. However, despite this general decrease, proteins that are important drivers of AD brain co-expression modules such as SMOC1 were increased in AD plasma, and show promise as accessible biomarkers of AD brain pathology. Co-expression analysis showed strong changes in AD CSF related to proteostasis, sugar metabolism, synaptic biology, and complement pathways. Analysis of plasma showed matrisome, complement, and endocytosis modules strongly correlated with AD. These modules may themselves represent promising AD biofluid biomarkers potentially more robust to analytical and natural biological variation than individual protein markers.

SOMAmer signals were much lower in CSF compared to plasma, an important consideration when interpreting SomaScan assay data in this fluid. We used an empirically-derived threshold to remove assays that did not meet our criteria for acceptable S:N, which removed approximately half of the assays from consideration for most of our analyses. For co-expression analyses, we decided to keep all SomaScan CSF assays due to the robust ability of network co-expression to handle noise. Interestingly, most of the SOMAmers that were removed by our S:N threshold for individual protein analyses fell within a network co-expression module, suggesting that the low signal for many SOMAmers in CSF may still contain biological meaning. The approach to handle S:N for SomaScan in CSF will depend on the context and needs of the desired analysis. There were no issues with S:N for SomaScan in plasma, reflecting optimization of the platform for this matrix.

Correlation of protein measurements across platforms was good, with a median *r* of approximately 0.7 in CSF and 0.6 in plasma. The median correlation in plasma was generally higher than what has previously been observed comparing Olink and SomaScan assays^28, 29^, perhaps because of the larger number of assays compared between these two platforms in this study and the lack of normalization of the SomaScan data^29^. The correlations improved when considering only proteins that are altered in AD, suggesting that measurement variability within a given assay reduced the correlation when S:N for a given assay was low. The source of this noise is likely biological, particularly in plasma where protein levels are influenced by multiple tissues and organ systems and other factors unrelated to AD. An important consideration when interpreting cross-platform correlation is also protein isoform, or “proteoform,” complexity in plasma versus CSF, including splicing variation and post-translational modifications (PTMs), as well as protein complex formation. Such proteoform complexity is very likely to be a large driver of variation across proteomic measurements within and across platforms, where a given platform is targeting a particular epitope or peptide for measurement of protein levels that could be obscured by splice variation, PTMs, or complex formation^29, 30^. This is likely the source of variation in SPP1 measurement noted above, which has greater than 40 known phosphorylation sites and 5 glycosylation sites. Another source of variation is also off-target binding, or ion co-isolation and interference in the case of MS, which is a larger issue in more complex matrices such as plasma. In this context, one current advantage for the SomaScan and MS platforms is the use of multiple aptamers or peptides for the measurement of some proteins, which could allow for consideration of such biological and technical variation. An example of this is the SomaScan measurement of TNC described above, where six of the eight SOMAmers correlated with one another. Multiple assays for other AD-related proteins, such as NEFL, CHI3L1, NPTXR, and SPP1 would be a welcome addition to proteomic platforms. In-depth characterization of the actual protein species being measured by an assay in a given platform will significantly advance our understanding of proteoforms related to disease.

A key observation that arose from our analyses was the strong bias for proteins of medium to low abundance to be decreased in AD plasma. Our AD plasma data were similar to those in a recently-described Hong Kong cohort where this strong bias was also observed^18^. The basis of this observation is not currently clear, but could represent general protein translation reduction^31^ in a systemic fashion in AD that somewhat spares proteins of higher abundance such as albumin, which are the primary drivers of gross plasma protein level measurements in clinical assays. Alteration of the blood-brain barrier in AD may also contribute to the observed discrepancy between the CSF and plasma levels of some proteins^8, 32^. We chose not to perform median normalization prior to analysis given that we are most interested in actual measured levels for clinical biomarker discovery and translation. Therefore, proteins that remain elevated in plasma without median normalization, such as SMOC1, represent highly promising markers for AD. SMOC1 may be an excellent marker for the M42 brain matrisome module in CSF and plasma given the fact that it is a key driver of M42 co-expression in brain. M42 is strongly related to amyloid deposition^2, 33^, and the fact that CSF and plasma SMOC1 did not correlate with cognitive function is consistent with this association given that amyloid also does not strongly correlate with cognitive function^34^ and M42 levels are not an independent driver of cognitive decline after adjustment for AD neuropathology^2^. Other brain module hub proteins as described above may also represent promising biofluid biomarkers for brain processes. The relationship in the levels of many of these markers between CSF and plasma appears more complicated than for SMOC1, with some showing opposite directions of change in AD plasma compared to CSF. While some of this variation may be due to peripheral sources of the protein, it is also possible that exchange of certain proteins from CSF to the plasma compartment is regulated. Further investigation into such possible regulatory mechanisms is warranted.

One potential way to deal with variation in any one biomarker is to construct panels of markers that reflect a particular biological process, where the composite level of the panel becomes the measurement of the biological process of interest^7^. In this study, we constructed protein co-expression networks in CSF and plasma to illustrate this potential approach to AD biomarker development. We were able to incorporate all three proteomic platform measurements into these networks, which were based on >7100 protein measurements in CSF, and >9500 protein measurements in plasma. To our knowledge, these are the deepest proteomic analyses of these two fluids to date. The CSF network revealed autophagy and ubiquitination modules that were strongly correlated to current AD CSF biomarkers, indicating disruption of proteostasis is a strong disease-related signal in AD CSF. Other strong disease-related signals included alterations in synaptic biology, sugar metabolism, and complement, all of which have been previously described in AD brain^2, 3^. In plasma, these modules were not as well defined. Whether the CSF eigenproteins for these modules will translate into potential plasma biomarkers is currently unknown, and will require further study in larger cohorts. One module that will likely translate across tissues is the complement module, which was highly preserved across brain, CSF, and plasma. Interestingly, while complement module levels were increased in brain and plasma, they were decreased in CSF, suggesting that complement deposition in brain leads to discordance in brain-CSF levels similar to that seen with brain Aβ deposition^35^. However, in contrast to Aβ, the source of complement in the brain may be derived largely from peripheral sources given the observed elevation in plasma. This hypothesis would be consistent with recent findings on peripheral factors that influence brain pathophysiology in AD^36^.

In summary, multi-platform proteomic analysis of AD CSF and plasma is a promising approach to further development of biomarkers that can reflect the complex and multifaceted processes that comprise AD, and that can enable patient stratification, diagnostics and disease monitoring, and therapeutic development.

## Supporting information

Dammer et al 2022 Extended Data

Dammer et al 2022 Supplementary Tables

## Acknowledgements

We are grateful to those who agreed to donate their CSF and blood for this study. The authors would like to thank SomaLogic and Olink research customer support teams for their consultations on data analysis. This study was supported by the following National Institutes of Health funding mechanisms: K08AG068604, P30AG066511, and U54AG065187.

## Data and Code Availability

Raw data, case traits, and analyses related to this manuscript are available at https://www.synapse.org/3platformEmory. Code available in the research compendium for the current study is available from https://www.synapse.org/3platformEmory. The algorithm used for batch correction is fully documented and available as an R function, which can be downloaded from https://github.com/edammer/TAMPOR. Data used in the preparation of this article were obtained from the Accelerating Medicine Partnership® (AMP®) Parkinson’s Disease (AMP PD) Knowledge Platform. For up-to-date information on the study, visit https://www.amp-pd.org. The AMP® PD program is a public-private partnership managed by the Foundation for the National Institutes of Health and funded by the National Institute of Neurological Disorders and Stroke (NINDS) in partnership with the Aligning Science Across Parkinson’s (ASAP) initiative; Celgene Corporation, a subsidiary of Bristol-Myers Squibb Company; GlaxoSmithKline plc (GSK); The Michael J. Fox Foundation for Parkinson’s Research; Pfizer Inc.; Sanofi US Services Inc.; and Verily Life Sciences. ACCELERATING MEDICINES PARTNERSHIP and AMP are registered service marks of the U.S. Department of Health and Human Services. Absolute quantitative plasma protein data was obtained from the Human Protein Atlas at proteinatlas.org.

## Author Contributions

ECBJ, EBD, LP, DMD, and JJL designed the experiments; LP and DMD carried out experiments; EBD and ECBJ analyzed data; LP, DMD, ESM, NTS, JJL, and AIL provided advice on the interpretation of data; ECBJ wrote the manuscript with input from coauthors. All authors approved the final manuscript.

## Methods

### CSF and Plasma Samples and Case Classification

All CSF and plasma samples used in this study were collected under the auspices of the Emory Goizueta Alzheimer’s Disease Research Center (ADRC). The cohort consisted of 18 healthy controls and 18 patients with AD. Basic demographic data were obtained from the Goizueta ADRC. Controls and patients with AD received standardized cognitive assessments in the Emory Cognitive Neurology Clinic or Goizueta ADRC. CSF and plasma were collected at or near the same time point in each individual and banked according to the 2014 National Institute on Aging best practice guidelines for Alzheimer’s Disease Centers (https://alz.washington.edu/BiospecimenTaskForce.html). CSF samples were subjected to ELISA Aβ_1–42_, total tau, and p-tau181 analysis by the INNO-BIA AlzBio3 Luminex Assay^37^.

ELISA values were used to support diagnostic classification based on established AD biomarker cutoff criteria^38, 39^. *APOE* genotype was determined by extracting DNA from the plasma buffy using the GenePure kit (Qiagen) following the manufacturer’s recommended protocol, then determining the rs7412 and rs429358 genotypes using either an Affymetrix Precision Medicine Array (Affymetrix) or TaqMan assays (ThermoFisher Scientific C_904973_10 and C_3084793_20). All samples were analyzed by each proteomic platform except for one subject, whose CSF and plasma were analyzed only by SomaScan. All Emory research participants provided informed consent under protocols approved by the Institutional Review Board at Emory University. Summarized case metadata is provided in **Supplementary Table 1**.

### Quantification of Proteins by Olink Proximity Extension Assay (PEA)

Proteins were quantified by PEA as previously described^17^. Aliquots of CSF and plasma from each subject were sent to Olink (Olink Proteomics, Uppsala, Sweden) for analysis on 13 human Olink Target 96 panels (cardiometabolic, cardiovascular II, cardiovascular III, cell regulation, development, immune response, inflammation, metabolism, neuro exploratory, neurology, oncology II, oncology III, and organ damage). All samples passed quality control measures and were randomized by Olink prior to analysis on single plates. Results were reported as Normalized Protein eXpression (NPX) values in log2 scale for relative quantification of protein abundance.

### Quantification of Proteins by SomaLogic SomaScan Modified Aptamers

Proteins were quantified by SomaScan as previously described^13, 14^. Aliquots of CSF and plasma from each subject were sent to SomaLogic (SomaLogic, Boulder, CO) for analysis using the modified aptamer SomaScan assay (v4.1). All samples passed quality control measures and were randomized by SomaLogic prior to analysis on single plates. Results were reported as relative fluorescence units (RFUs) for relative quantification of protein abundance.

### CSF Protein Preparation and Digestion for Tandem Mass Tag Mass Spectrometry (TMT-MS) Analysis

#### CSF Undepleted of Highly Abundant Plasma Proteins

Equal volumes (50 μl of each sample) of CSF were digested with lysyl endopeptidase (LysC, Wako 125-05061) and trypsin (ThermoFisher Scientific 90058). Briefly, each sample was reduced and alkylated with 1 μl of 0.5 M tris-2(-carboxyethyl)-phosphine (TCEP) and 5 μl of 0.4 M chloroacetamide (CAA) at 90°C for 10 min, followed by water bath sonication for 15 min. The same volume of 8 M urea buffer [56 μl, 8 M urea in 10 mM Tris, 100 mM NaH_2_PO_4_ (pH 8.5)] was added to each sample after cooling the samples to room temperature, along with LysC (2.5 μg). After overnight digestion, 336 μl of 50 mM ammonium bicarbonate (ABC) was added to each sample to dilute the urea concentration to 1 M, along with trypsin (5 μg). After 12 hours, the trypsin digestion was stopped by adding final concentration of 1% formic acid (FA) and 0.1% trifluoroacetic acid (TFA).

#### CSF Depleted of Highly Abundant Plasma Proteins

To increase the depth of proteome coverage, immunodepletion of highly abundant proteins was performed as previously described^7^. For CSF samples, 130 μl was incubated with equal volume (130 μl) of High Select Top14 Abundant Protein Depletion Resin (ThermoFisher Scientific, A36372) at room temperature in centrifuge columns (ThermoFisher Scientific, A89868). After 15 min of mixing with gentle rotation, the samples were centrifuged at 1000 x *g* for 2 min. Sample flow-through was concentrated with a 3K Ultra Centrifugal Filter Device (Millipore, UFC500396) by centrifugation at 14,000 x *g* for 30 min, and then the immunodepleted samples were diluted to equal volumes of 75 μl with phosphate-buffered saline. Immunodepleted CSF (60 μl) was then digested with LysC and trypsin. Briefly, the samples were reduced and alkylated with 1.2 μl of 0.5 M TCEP and 3 μl of 0.8 M CAA at 90°C for 10 min, followed by water bath sonication for 15 min. Samples were diluted with 193 μl of 8 M urea buffer [8 M urea in 10 mM Tris, 100 mM NaH_2_PO_4_ (pH 8.5)] to a final concentration of 6 M urea. LysC (4.5 μg) was used for overnight digestion at room temperature. Samples were then diluted to 1 M urea with 50 mM ABC. Trypsin (4.5 μg) was then added, and the samples were subsequently incubated for 12 hours. The digestion was then stopped by adding final concentration of 1% FA and 0.1% TFA.

### Plasma Protein Preparation and Digestion for TMT-MS Analysis

#### Plasma Undepleted of Highly Abundant Plasma Proteins

Equal volumes (2 μl of each sample) of plasma were digested with LysC and trypsin. Briefly, each sample was diluted 10-fold with 50 mM ABC, following by reduction and alkylation with 0.4 μl of 0.5 M of TCEP and 2 μl of 0.4 M CAA with heating at 90°C for 10 min. The samples were sonicated for 15 min with water bath sonication to help sample solubilization. 8 M urea buffer [22.4 μl, 8 M urea, 10 mM Tris, 100 mM NaH_2_PO_4_ (pH 8.5)] was added to each sample after cooling to room temperature, along with LysC (10 μg). After overnight digestion, 134.4 μl of 50 mM ABC was added to each sample to dilute the urea concentration to 1 M, along with trypsin (20 μg). After 12 hours, the trypsin digestion was stopped by adding final concentration of 1% FA and 0.1% TFA.

#### Plasma Depleted of Highly Abundant Plasma Proteins

The High Select Top14 Abundant Protein Depletion Resin was also utilized for plasma samples prior to digestion. Following mixing, 500 μl of resin was aliquoted into each spin column. After the resin settled to the bottom of the spin column, 8 μL of each sample was added and depletion was performed by gentle rotation for 15 min at room temperature, followed by centrifugation at 1000 x *g* for 2 min. Sample flow-through was concentrated with a 3K Ultra Centrifugal Filter Device by centrifugation at 14,000 x *g* for 30 min. Immunodepleted samples were diluted to equal volumes of 75 μl with phosphate-buffered saline. Immunodepleted plasma (60 μl) was then digested with LysC and trypsin using the same protocol used for CSF depleted samples.

#### Isobaric TMT Peptide Labeling

Before TMT labeling, the digested peptides were desalted using 50 mg of Sep-Pak C18 columns (Waters). Briefly, the columns were activated with 1 mL of methanol, then equilibrated with 2 × 1 mL 0.1% TFA. The acidified samples were loaded following by washing with 2 × 1 mL 0.1% TFA. Elution was performed with 1 mL 50% acetonitrile. To normalize protein quantification across batches, global internal standard (GIS) samples were generated for each sample set by combining 100 μl aliquots from each sample elution. All individual samples and GIS pooled standards were dried by speed vacuum (Labconco).

Both depleted CSF and depleted plasma samples were divided into five TMT batches, labeled using an 11-plex TMT kit (ThermoFisher Scientific, A34808, lot number for TMT 10-plex: SI258088, 131C channel SJ258847), and derivatized as previously described^7^. For the sample and channel distribution, please see **Supplementary Table 1**. 5 mg of each channel reagent was dissolved in 256 μL anhydrous acetonitrile. Each peptide sample was resuspended in 50 μl of 100 mM triethylammonium bicarbonate (TEAB) buffer, and 20.5 μl of TMT reagent solution was subsequently added. After 1 hour, the reaction was quenched with 4 μl of 5% hydroxylamine (ThermoFisher Scientific, 90115) for 15 min. The peptide solutions were then combined according to the batch arrangement. Each TMT sample was desalted with 100 mg of Sep-Pak C18 columns and dried by speed vacuum. Notably, there were 9 TMT channels used for depleted CSF samples with one GIS sample on channel 127N, whereas 10 channels were used for depleted plasma samples with two GIS samples included on both 127C and 131C channels. Channel 126 was left empty on both sample sets.

For undepleted CSF and plasma samples, the TMT 16-plex kit (ThermoFisher Scientific, A44520, lot number VH311511) was used for labeling, which divided both CSF and plasma sample sets into 3 TMT batches with 12 samples plus 1 GIS in each batch. The sample and channel distribution were the same for CSF and plasma samples (**Supplementary Table 1**). 5 mg of each channel reagent was dissolved in 200 μL anhydrous acetonitrile. Each CSF peptide sample was resuspended in 50 μl of 100 mM TEAB buffer, and 10 μl of TMT reagent solution was subsequently added. For plasma samples, each peptide sample was resuspended in 150 μl of 100 mM TEAB buffer, and 30 μl of TMT reagent solution was subsequently added. The labeling was stopped after 1 hr with 4 μl of 5% hydroxylamine for CSF and 12 μl of 5% hydroxylamine for plasma, and the peptide solutions were then combined according to the batch arrangement. The combined TMT samples were desalted with 100 mg of Sep-Pak C18 columns except for each undepleted plasma TMT sample, which was split and desalted using 2 x 100 mg of Sep-Pak C18 columns. The elutions were dried under speed vacuum.

### High-pH Off-line Fractionation

#### CSF and Plasma Undepleted of Highly Abundant Plasma Proteins

Dried samples were re-suspended in high pH loading buffer (0.07% vol/vol NH_4_OH, 0.045% vol/vol FA, 2% vol/vol ACN) and loaded onto a Water’s BEH column (2.1 mm x 150 mm with 1.7 µm particles). A Vanquish UPLC system (ThermoFisher Scientific) was used to carry out the fractionation. Solvent A consisted of 0.0175% (vol/vol) NH_4_OH, 0.01125% (vol/vol) FA, and 2% (vol/vol) ACN; solvent B consisted of 0.0175% (vol/vol) NH_4_OH, 0.01125% (vol/vol) FA, and 90% (vol/vol) ACN. The sample elution was performed over a 25 min gradient with a flow rate of 0.6 mL/min with a gradient from 0 to 50% solvent B. A total of 192 individual equal volume fractions were collected across the gradient. Fractions were concatenated to either 48 or 96 fractions and dried to completeness using vacuum centrifugation.

#### CSF and Plasma Depleted of Highly Abundant Plasma Proteins

Dried samples were re-suspended in high pH loading buffer (0.07% vol/vol NH_4_OH, 0.045% vol/vol FA, 2% vol/vol ACN) and loaded onto an Agilent ZORBAX 300 Extend-C18 column (2.1 mm × 150 mm with 3.5 µm beads). An Agilent 1100 HPLC system was used to carry out the fractionation. Solvent A consisted of 0.0175% (vol/vol) NH_4_OH, 0.01125% (vol/vol) FA and 2% (vol/vol) ACN; solvent B consisted of 0.0175% (vol/vol) NH_4_OH, 0.01125% (vol/vol) FA and 90% (vol/vol) ACN. The sample elution was performed over a 60 min gradient with a flow rate of 0.4 mL/min with a gradient from 0 to 60% solvent B. A total of 96 individual equal volume fractions were collected across the gradient and subsequently pooled by concatenation into 30 fractions and dried to completeness under vacuum centrifugation.

### TMT Mass Spectrometry

#### CSF Undepleted of Highly Abundant Plasma Proteins

For batch 1, all samples (∼1ug) were loaded and eluted using a Dionex Ultimate 3000 RSLCnano (ThermoFisher Scientific) on an in-house packed 25 cm, 100 μm internal diameter (i.d.) capillary column with 1.9 μm Reprosil-Pur C18 beads (Dr. Maisch, Ammerbuch, Germany) over a 60 min gradient. Mass spectrometry was performed with a high-field asymmetric waveform ion mobility spectrometry (FAIMS) Pro-equipped Orbitrap Eclipse (ThermoFisher Scientific) in positive ion mode using data-dependent acquisition with 2 second top speed cycles. Each cycle consisted of one full MS scan followed by as many MS/MS events that could fit within the given 2 second cycle time limit. MS scans were collected at a resolution of 120,000 (410-1600 m/z range, 4×10^5 AGC, 50 ms maximum ion injection time, FAIMS compensation voltage of −50 and −70). All higher energy collision-induced dissociation (HCD) MS/MS spectra were acquired at a resolution of 30,000 (0.7 m/z isolation width, 35% collision energy, 1.25×10^5 AGC target, 54 ms maximum ion time, TurboTMT on). Dynamic exclusion was set to exclude previously sequenced peaks for 30 seconds within a 10-ppm isolation window. For batches 2 and 3, samples were eluted over a 21 min gradient. Mass spectrometry was performed the same as batch 1 except with a FAIMS compensation voltage of −45, and dynamic exclusion set to exclude previously sequenced peaks for 6 seconds within a 10-ppm isolation window.

#### Plasma Undepleted of Highly Abundant Plasma Proteins

Mass spectrometry was performed the same as for CSF undepleted batches 2 and 3 except FAIMS compensation voltage was set at −40 and −60, and dynamic exclusion was set to exclude previously sequenced peaks for 20 seconds within a 10-ppm isolation window.

#### CSF and Plasma Depleted of Highly Abundant Plasma Proteins

All fractions (∼1ug) were loaded and eluted using an Easy-nLC 1200 (ThermoFisher Scientific) on an in-house packed 30 cm, 750 μm i.d. capillary column with 1.9 μm Reprosil-Pur C18 beads over a 120 min gradient. Mass spectrometry was performed with a Q-Exactive HFX (ThermoFisher Scientific) in positive ion mode using data-dependent acquisition with a top 10 method. Each cycle consisted of one full MS scan followed by 10 MS/MS events. MS scans were collected at a resolution of 120,000 (400-1600 m/z range, 3×10^6 AGC, 100 ms maximum ion injection time). All higher energy collision-induced dissociation (HCD) MS/MS spectra were acquired at a resolution of 45,000 (1.6 m/z isolation width, 35% collision energy, 1×10^5 AGC target, 86 ms maximum ion time). Dynamic exclusion was set to exclude previously sequenced peaks for 20 seconds within a 10-ppm isolation window.

#### Database Searches and Protein Quantification

All raw files were searched using Proteome Discoverer (version 2.4.1.15, ThermoFisher Scientific) with Sequest HT. The spectra were searched against a human UniProt database downloaded April 2015 (90300 target sequences). Search parameters included 20 ppm precursor mass window, 0.05 Da product mass window, dynamic modifications methionine (+15.995 Da), deamidated asparagine and glutamine (+0.984 Da), phosphorylated serine, threonine and tyrosine (+79.966 Da), and static modifications for carbamidomethyl cysteines (+57.021 Da) and N-terminal and lysine-tagged TMT (+229.163 or +304.207 Da depending on the dataset). Percolator was used to filter peptide spectral matches (PSMs) to 1% FDR. Peptides were grouped using strict parsimony and only razor and unique peptides were used for protein level quantitation. Reporter ions were quantified from MS2 scans using an integration tolerance of 20 ppm with the most confident centroid setting. Only unique and razor (i.e., parsimonious) peptides were considered for quantification.

### Protein Abundance Data Processing

#### Tandem Mass Tag Mass Spectrometry (TMT-MS)

Only proteins that were identified and summarized as high confidence (<1% FDR) by Proteome Discoverer (PD) were used for analysis. The 3730 UniProt protein identifier accessions provided by PD were further annotated with Hugo Gene Nomenclature Committee (HGNC) official gene symbols. TMT-MS data were processed identically for both CSF and plasma, including fluid depleted of highly abundant proteins. TMT reporter intensities (abundances) that had not undergone normalization by Proteome Discoverer (PD) were used for analysis to preserve inherent protein abundance differences between control and AD subjects. Four separate datasets were used for analysis: CSF undepleted, CSF depleted, plasma undepleted, and plasma depleted of highly abundant proteins. For each dataset, batch correction was performed by dividing abundances for each protein within each batch by the global internal standard (GIS). GIS measurements were then removed, and proteins with more than 75 percent (*n*=27/36) missing values were excluded from consideration. The number of remaining protein isoforms after missing value control was 1128 in undepleted plasma, 2229 in undepleted CSF, 1385 in plasma depleted of highly abundant proteins, and 2944 in CSF depleted of highly abundant proteins.

#### Olink Proximity Extension Assay (PEA) and SomaLogic SomaScan Assay

Olink NPX values were analyzed using the OlinkAnalyze R package v1.2.1. NPX values that were flagged with quality control (QC) warnings were removed from further consideration. SomaScan RFU data and assay metadata for CSF and plasma were analyzed using the SomaDataIO R package v3.1.0. Olink NPX and SomaLogic SomaScan data included blank buffer replicate measurements (noise). Buffer measurements were used to calculate signal-to-noise (S:N) ratios for both unnormalized NPX or RFU abundance data. S:N ratios were calculated by subtracting the within-assay median buffer signal from the unlogged assay signal (2^NPX^ or RFU), then dividing by the median buffer signal. Protein assay-specific limit of detection (LOD) was defined as median log_2_ buffer signal plus 3 standard deviations (SD) of the assay’s buffer measurements. NPX background signal SD for PEA was defined from historically recorded background variance of the assays and included as a component of predetermined LOD, whereas SomaScan background SD was calculated from available buffer replicate data for the assays performed. Sample measurements in CSF or plasma that were below LOD were retained but considered as missing values. Olink repeated measurements of the same UniProt protein assayed in different panels (*N*=36 duplicated assays) were reduced to their representative single best replicate of the assay in one of the panels based on criteria including the highest pairwise correlation to other replicate assays, highest signal, and largest dynamic range.

Both platform assays underwent a first-pass filter allowing up to 75 percent (*n*=27/36) missing values. Because SomaScan CSF data had low signal, a second-pass filter step was applied to remove assays that did not meet an empirically-derived S:N threshold. This threshold was determined by correlating SomaScan assays with Olink and TMT-MS undepleted assays at varying S:N cutoff values (0, 0.15, 0.25, 0.3, 0.35, 0.4, 0.45, 0.50, 0.625, 0.75, 1, 2, 4, and 8), and selecting the S:N value (≥0.45) that maximized median Pearson correlation with the other platforms. After application of first- and second-pass filters and removal of control aptamers, 3594 CSF and 7284 plasma human SomaScan assays were kept for subsequent analyses. For Olink assays, after applying the first-pass filter, 902 CSF and 1140 plasma proteins were kept for subsequent analyses.

#### Proteome Coverage Overlap, Ontology Enrichment, and Missing Data Analysis

Unique gene symbols measured in each platform were counted, and overlap was visualized using the venneuler R package (v1.1-0) venneuler function. Enrichment of gene ontologies (GO) in different Venn categories was calculated as a Fisher’s exact test *p* value transformed to *z* score using GO-Elite (v1.2.5) and visualized using a custom in-house R script. The same procedure was used to determine ontology enrichment for network modules. Missing data (**Supplementary Figure 2**) in Olink and SomaScan included assays flagged by QC warnings, below LOD, and truly missing measurements. Missing data in TMT-MS was considered at the level of batch, as all measurements within a batch result from the same MS/MS fragmentation.

#### Censoring of Proteins Affected by Depletion of Highly Abundant Proteins

Proteins considered for analysis were those measured by TMT-MS before and after depletion of highly abundant proteins that had the same UniProt accession, at least 9 paired abundance measurements, and at least 3 measurements per case status group (AD or control; *n*=1932 CSF proteins, *n*=852 plasma proteins). Pairwise measurements were correlated using Pearson correlation on the difference in abundance between AD versus control subjects. Proteins that were discordant in their differential abundance, as well as proteins with negative Pearson rho across depleted and undepleted matched protein measurements across the 36 case samples, were considered significantly affected by depletion. In total, 32 proteins in CSF and 27 in plasma were censored from the TMT-MS depleted data due to effects of depletion on their abundance levels (**Supplementary Table 3**).

#### Protein Abundance Correlation Analysis

Proteins measured in common across two platforms within the same biofluid were correlated across all samples using the corAndPvalue function in the WGCNA R package (v1.69) (**Supplementary Tables 12-17, Extended Data**). In the case of multiple SomaScan assays for the same protein, the assay with the identical UniProt protein accession, or secondarily, a SOMAmer measuring an identical gene product, was selected. When multiple cross-platform UniProt accession or gene symbol matches occurred, the SOMAmer with the highest correlation was selected. We constructed a population histogram of all Pearson correlations for distinct gene products or UniProt accessions (representing distinct protein isoforms), and identified the median rho for each population of paired measurements between two platforms (**Figure 3**).

### Cumulative Signal and Total Protein Abundance Comparison

Mean NPX, RFU, or ion intensity signal without prior normalization or log transformation for each protein across the 36 samples was ranked for each platform and biofluid from highest to lowest abundance. Curves of incremental median cumulative abundance were constructed for AD and control groups (**Supplementary Figure 8**, left *uncalibrated* panels). Absolute protein abundance differences in plasma were also assessed by calibrating relative platform signals to absolute plasma protein concentrations (*n*=4226) as provided in the Human Protein Atlas (HPA) on March 5, 2022, https://www.proteinatlas.org/search. For the calibrated abundance calculations, unlogged abundance (signal minus buffer median, including positive values below LOD) was calibrated so that the geometric mean of control group measurements was set to the known absolute blood concentration with all individual measurements varying relative to this value. Missing values were considered as one-half the minimum assayed nonzero signal for geometric mean calculations. When multiple assays for a gene product were available within a platform, the assay with maximum mean signal was selected for calibration. HPA-calibrated values were plotted for all 4226 proteins regardless of presence in the platform as a ranked absolute abundance curve (non-cumulative, black trace in **Supplementary Figure 8B, D, F**, and **H**). Then, the cumulative log_10_-transformed abundance of all lesser abundant ranked proteins up to each represented rank of any protein measured within the platform was plotted as the median such value for AD or control (**Supplementary Figure 8**). In contrast to the uncalibrated cumulative abundance curves, the value at each point in these left-truncated cumulative sum curves represents the sum (cumulative) abundance of only lesser abundant proteins up to that rank, and not those ranked with higher abundance.

### Differential Expression Analysis

Differences between AD and control were assessed on the log_2_(abundance) measurements over all proteins after data processing as described above, which included signal cleanup, filtering on missingness, removal of proteins affected by top-14 highly abundant protein depletion, and, in the case of SomaScan CSF data, control of excessively low S:N assays. Volcano plots were made using a custom in house script via the plotly (v4.9.2.1) R package function ggplotly. Individual volcano points were colored by membership in the 44 brain network modules described in Johnson *et al*^2^.

### Comparison to External Datasets

External Olink datasets used for correlation included plasma AD effect sizes from a Hong Kong-based cohort provided in Jiang *et al.*^18^ Appendix Table 1, from the BioFinder cohort described in Whelan *et al.*^19^ Table S19, and from the Accelerating Medicines Partnership for Parkinson’s Disease (AMP-PD) 2021 v2-5 (May 10) release. The external SomaScan dataset was obtained from the ANMerge version of the AddNeuroMed study data as described in Birkenbihl *et al.*^20^ Only sample data collected at the last visit in the AMP-PD and ANMerge datasets were used for correlations.

Data provided in Jiang *et al*. and Whelan *et al*. was used directly for correlation without additional processing. AMP-PD Olink raw data from 212 study participants and all four 384-assay panels available (Cardiometabolic, Inflammation, Neurology, and Oncology) was loaded and processed in the same fashion as Emory Olink data. Four participants with a diagnosis of multiple systems atrophy were excluded. Final number of subjects analyzed was 118 PD and 90 control. As described above, values below LOD were censored as missing but otherwise retained for correlations, and proteins duplicated in the different panels were reduced to one representative assay. Final assay numbers included 1054 CSF and 1398 plasma assays. Log_2_ fold change values that remained significant after FDR correction were correlated with log_2_ fold change values in the current study using Pearson correlation and Student’s *p* values as implemented in the WGCNA package verboseScatterplot function.

### Harmonization of Platform Protein Abundance Prior to Network Analysis

TMT-MS ion counts in fluid depleted of highly abundant proteins, SomaScan RFUs, and Olink unlogged NPX values, totaling 9,589 assays in plasma and 7,158 assays in CSF, were assembled for the 35 case samples commonly measured on all three platforms. Only truly missing values were considered as unavailable; values below LOD or those subject to S:N threshold-based filtering were retained. Data were transposed prior to removal of platform-specific effects as a batch effect using the TAMPOR algorithm. Proteins were considered as samples (columns) and samples as rows for the two-way table median polish of ratio using TAMPOR. Common proteins measured across all three platforms were used as the GIS (*n*=101 in plasma and *n*=201 in CSF) to calculate the central tendency of data within and across platforms used for the denominators in the TAMPOR algorithm, as previously described^2^. Normalized data used in subsequent network analyses was of the form log_2_(abundance/central tendency) of the common proteins in all platforms. No protein assay had more than fifty percent missing values.

### Protein Co-Expression Network Analysis

Networks for CSF and plasma were constructed using the harmonized protein abundances for each biofluid. The Weighted Correlation Network Analysis (WGCNA) algorithm (v1.69) was used for network generation. No outliers were detected using the WGCNA sample network connectivity outlier algorithm. The WGCNA blockwiseModules function was run on the CSF and plasma harmonized abundances with the following parameters: power=6.5 (CSF) or 11 (plasma), deepSplit=4, minModuleSize=10, mergeCutHeight=0.07, TOMdenom=”mean”, bicor correlation, signed network type, PAM staging and PAM respects dendro as TRUE, and a maxBlockSize larger than the total number of protein assays. Module memberships were then iteratively reassigned to enforce kME table consistency, as previously described^3^. The resulting network assignments were visualized as modules using the iGraph(v1.2.5) package. Module eigenprotein correlations and significance were visualized in circular heatmaps using the circlize(v0.4.10), dendextend(v1.13.4), and dendsort(0.3.3) R packages. Synthetic eigenproteins for each network (CSF, plasma, and brain^2^) were calculated as previously described^3^. For synthetic eigenproteins translated either from or to brain, the existing data for 8,619 proteins underlying the brain network were mapped to labels in the biofluid network using a mapping rubric to cross-reference protein labels. Specifically, (1) an exact Uniprot ID match to that in labels of the form Symbol|UniprotID|platform|biofluid took precedence for labels with MS as the platform, followed by (2) symbol matches with MS as the platform. This was followed by (3) an exact Uniprot ID match to an Olink row in a biofluid dataset, and then (4) an exact Uniprot ID to a SomaScan row, followed by (5) a symbol match with Olink as the platform, and finally, (6) a symbol match with SomaScan as the platform. In this way, unmatched proteins across pairs of networks were minimized. The same 6-point rubric was used for matching (relabeling) brain network member labels before performing module preservation (below).

### Network Module Overlap

Overrepresentation analysis (ORA) of module gene symbols between networks was determined using two-tailed Fisher’s exact test, followed by correction of *p* values for multiple testing using the Benjamini-Hochberg method. The plasma network was compared to a SomaScan human serum network constructed from 4,137 proteins as described in Emilsson *et al.*^27^ Serum network assignments in 27 modules plus grey were curated from Table S7 in Emilsson *et al*^27^. Overlap of module gene symbols between the two networks was determined as described above. Overlap was visualized using a custom in-house script.

### Network Preservation

Pairwise, directional preservation between CSF and plasma, plasma and CSF, and brain to each of the biofluid networks and vice versa was performed using the WGCNA (v1.69) modulePreservation function with 500 permutations after harmonizing protein assay labels as described above. *Z*_summary_ composite *z* score for 8 underlying network parameters was calculated and visualized by circular heatmap as significance (minus log_10_(Benjamini-Hochberg adjusted *p* values), corresponding to the *Z*_summary_ scores obtained.

### Cell Type Marker Enrichment Analyses

Cell type-specific enriched marker gene symbol lists were used as previously published to perform a Fisher’s exact one-tailed test for enrichment^2^. Benjamini-Hochberg correction was applied to all resulting *p* values.

### Other Statistics

All statistical analyses were performed in R (v4.0.2). Boxplots represent the median, 25^th^, and 75^th^ percentile extremes; thus, hinges of a box represent the interquartile range of the two middle quartiles of data within a group. The farthest data points up to 1.5 times the interquartile range away from box hinges define the extent of whiskers (error bars). Correlations were performed using the biweight midcorrelation function as implemented in the WGCNA R package or Pearson correlation. Comparisons between two groups were performed by two-sided *t* test. Comparisons among three or more groups were performed with Kruskal-Wallis nonparametric ANOVA or standard ANOVA with Tukey or *post hoc* pairwise comparison of significance. *P* values were adjusted for multiple comparisons by false discovery rate (FDR) correction according to the Benjamini-Hochberg method where indicated. *Z* score conversion of normalized protein data and normalized protein eigenproteins or synthetic eigenproteins were calculated as fold of standard deviation from the mean.

## Supplementary Figures

**Supplementary Figure 1.**
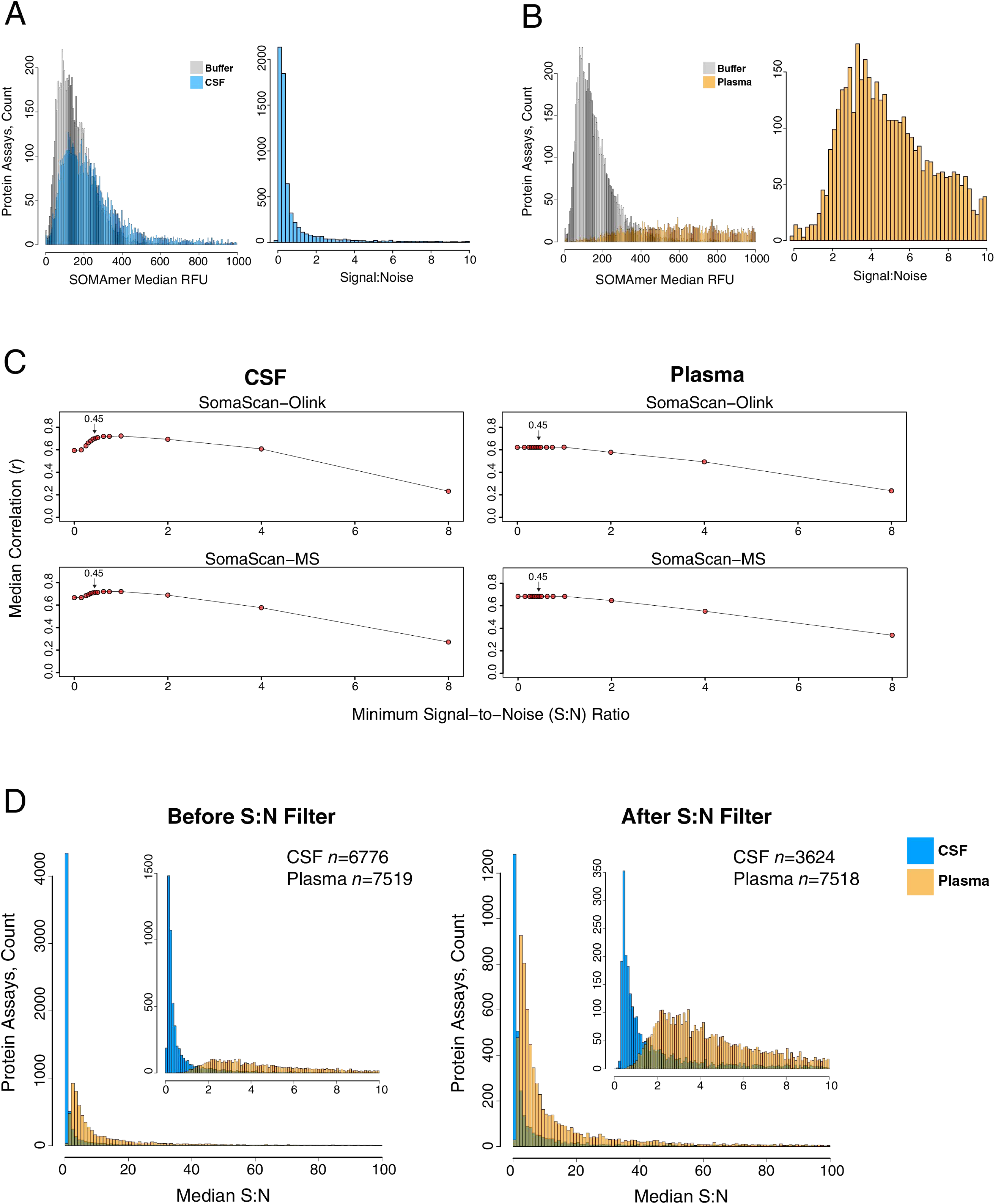
Signal-to-Noise Analysis in SomaScan. (A-D) Signal-to-noise was analyzed in the SomaScan 7000 data in CSF and plasma. (A) The frequency of median relative fluorescence units (RFU) for each protein assay in CSF across *n*=36 subjects and the frequency of median RFU for the buffer control of each aptamer reagent (left), with frequency of calculated signal:noise ratio for each aptamer reagent (right). (B) The same analysis as shown in (A), except in plasma. (C) The median correlation across all proteins commonly measured in the SomaScan 7000 platform with those in the Olink 1196 and TMT-MS platforms at a given minimum SomaScan signal:noise threshold, in CSF (left) and plasma (right). A signal:noise (S:N) ratio of 0.45 is indicated on the curves (noise is subtracted from signal prior to ratio calculation, and therefore S:N is <1). (D) Frequency of aptamer median S:N in CSF and plasma prior to (left) and after (right) applying a S:N filter of 0.45. Total aptamer numbers in each fluid before and after applying the filter are provided. Numbers include both human and non-human assays but no control probes, with a <75% missingness threshold by limit of detection applied prior to S:N filter. (Insets) S:N from 0-10.

**Supplementary Figure 2.**
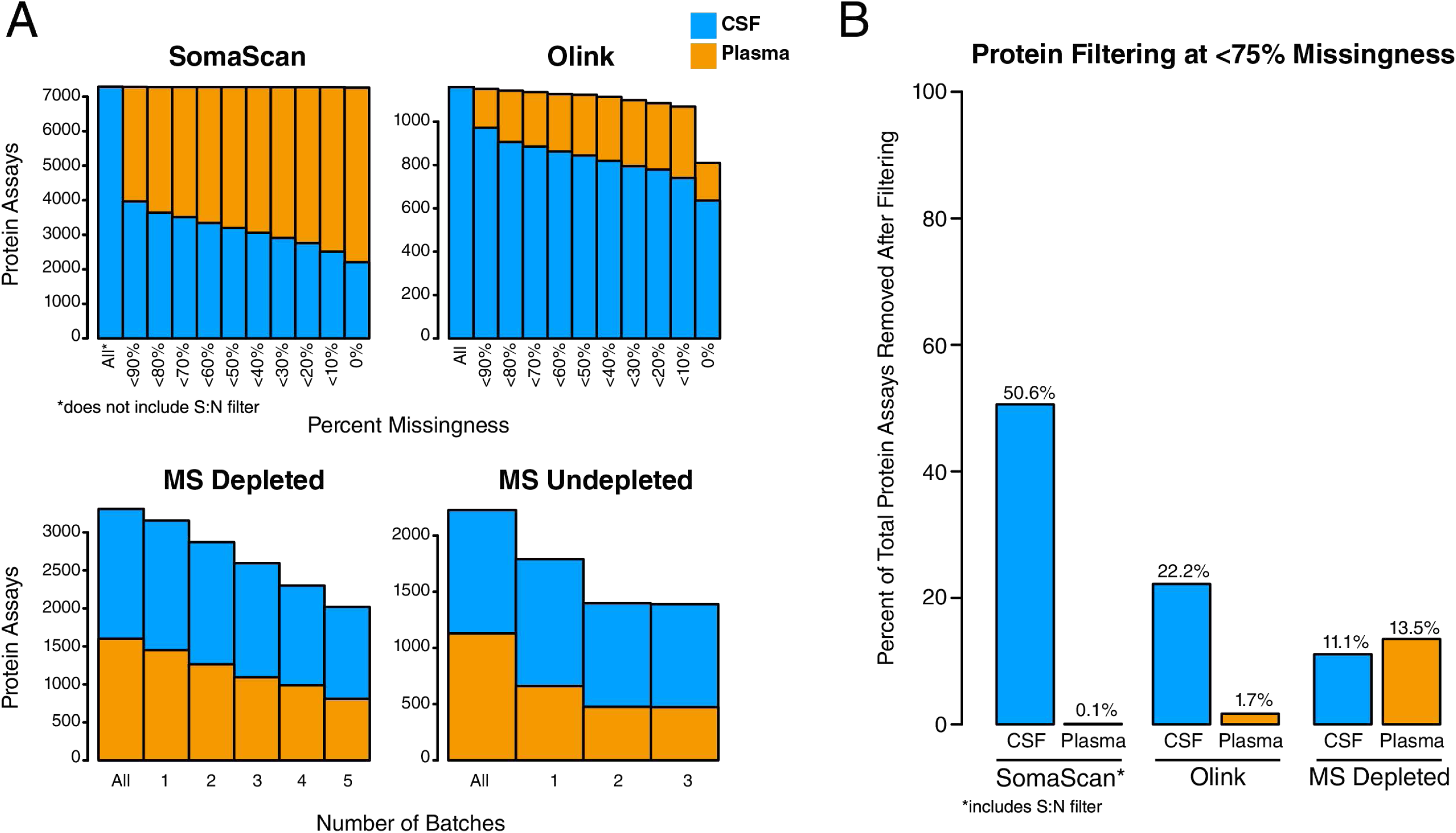
Missing Values. (A, B) The number of quantified protein assays by percent missingness across *n*=36 samples in CSF and plasma for the targeted SomaScan and Olink platforms, and the number of quantified proteins across TMT batches in the TMT-MS depleted and undepleted analyses (A). The SomaScan analysis includes application of the signal-to-noise (S:N) filter in CSF. “All” in each panel indicates the total number of human measurements for each platform and analysis, including assays that fall below S:N threshold in SomaScan, or do not pass QC criteria in Olink, or proteins that are identified but not quantified by reporter ions in MS (*n*=7288 total SomaScan human assays; *n*=1160 total Olink assays; *n*=1602 plasma, *n*=3310 CSF total proteins identified in MS depleted fluid; *n*=1129 plasma, *n*=2229 CSF total proteins identified in MS undepleted fluid). (B) The number of proteins removed after applying a <75% missingness filter. The SomaScan analysis also includes application of the S:N filter in CSF.

**Supplementary Figure 3.**
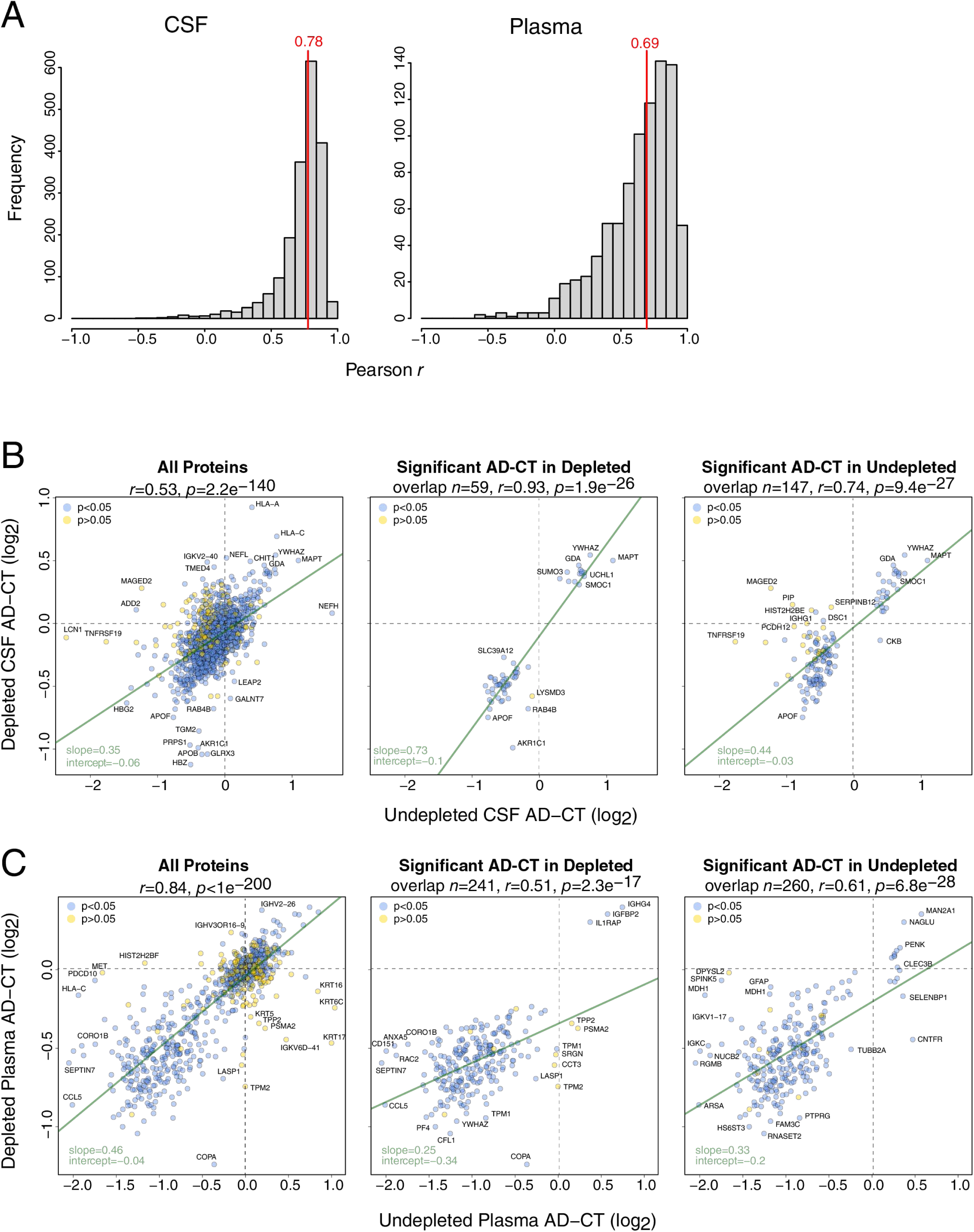
Effects of Highly Abundant Protein Depletion. (A-C) Frequency distribution of the correlation of TMT-MS measurements before and after depletion of the top 14 most highly abundant plasma proteins, in CSF (left) and plasma (right) (A). The vertical red line indicates the median correlation across all measurements after depletion. (B) Correlation of proteins between depleted and undepleted CSF, considering all proteins (left), proteins that are significantly different between AD and control in the depleted analysis and that have a corresponding measurement in the undepleted analysis (*n*=59, center), and proteins that are significantly different between AD and control in the undepleted analysis and that have a corresponding measurement in the depleted analysis (*n*=147, right). The individual proteins are colored according to whether they are correlated across paired measurements of the same subjects in depleted and undepleted measurements. (C) Same as in (B), except in plasma. Correlations were performed by Pearson test.

**Supplementary Figure 4.**
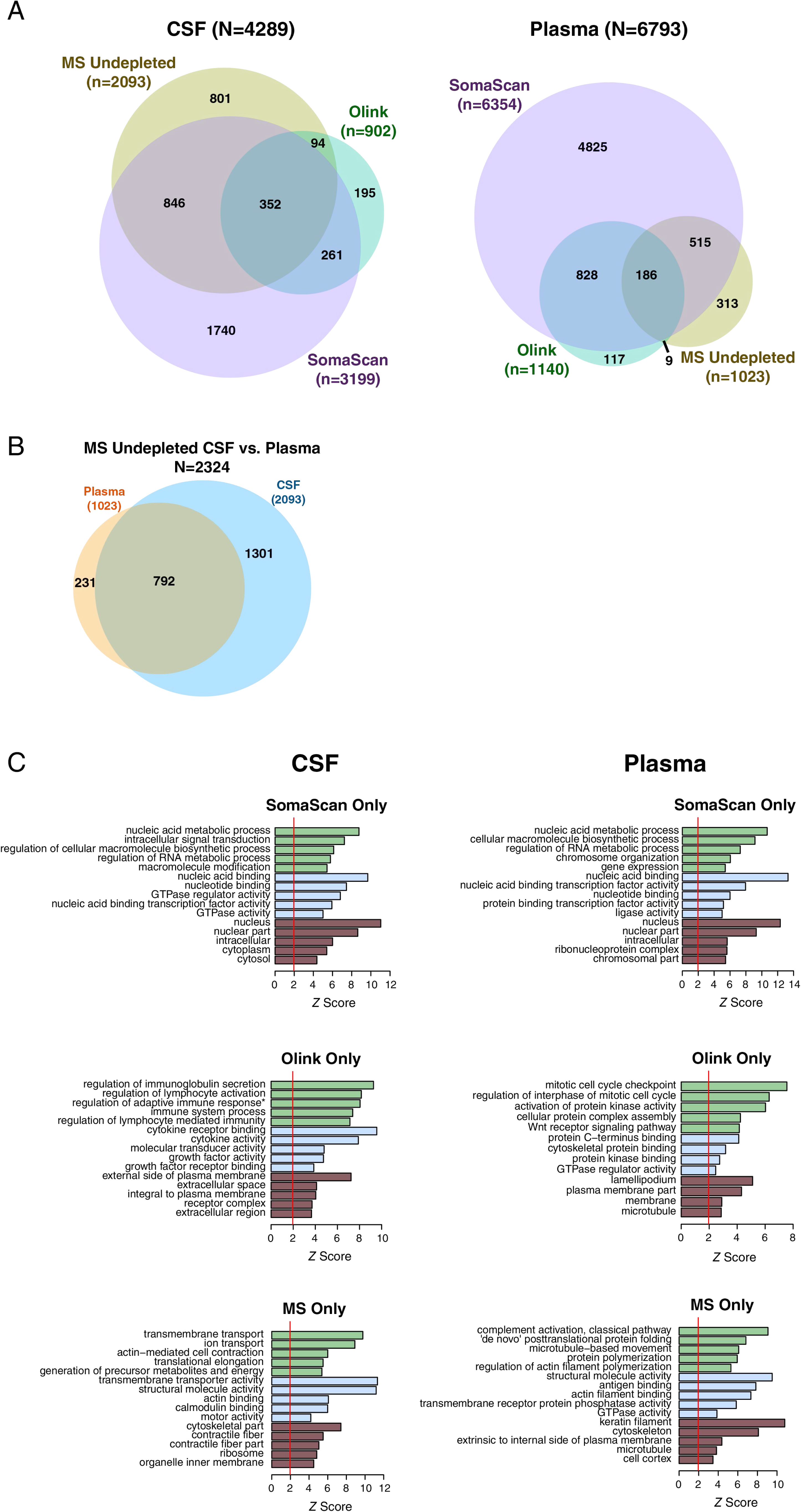
Proteomic Coverage with Undepleted Fluid and Unique Ontology Coverage. (A-C) Number and overlap of proteins measured by TMT-MS in undepleted fluid, Olink 1196, and SomaScan 7000 platforms in CSF (left) and plasma (right) from *n*=36 subjects. The threshold for inclusion was measurement in at least 9 subjects (or missing values <75%). CSF measurements on the SomaScan 7000 platform underwent signal-to-noise filtering (**Supplementary** Figure 1) prior to subsequent analyses. (B) Number and overlap of proteins measured in CSF and plasma by TMT-MS in undepleted fluid. (C) Gene ontology of proteins uniquely measured in each platform in CSF and plasma as shown in Figure 2. The vertical red line indicates significance at a *z* score of 1.96.

**Supplementary Figure 5.**
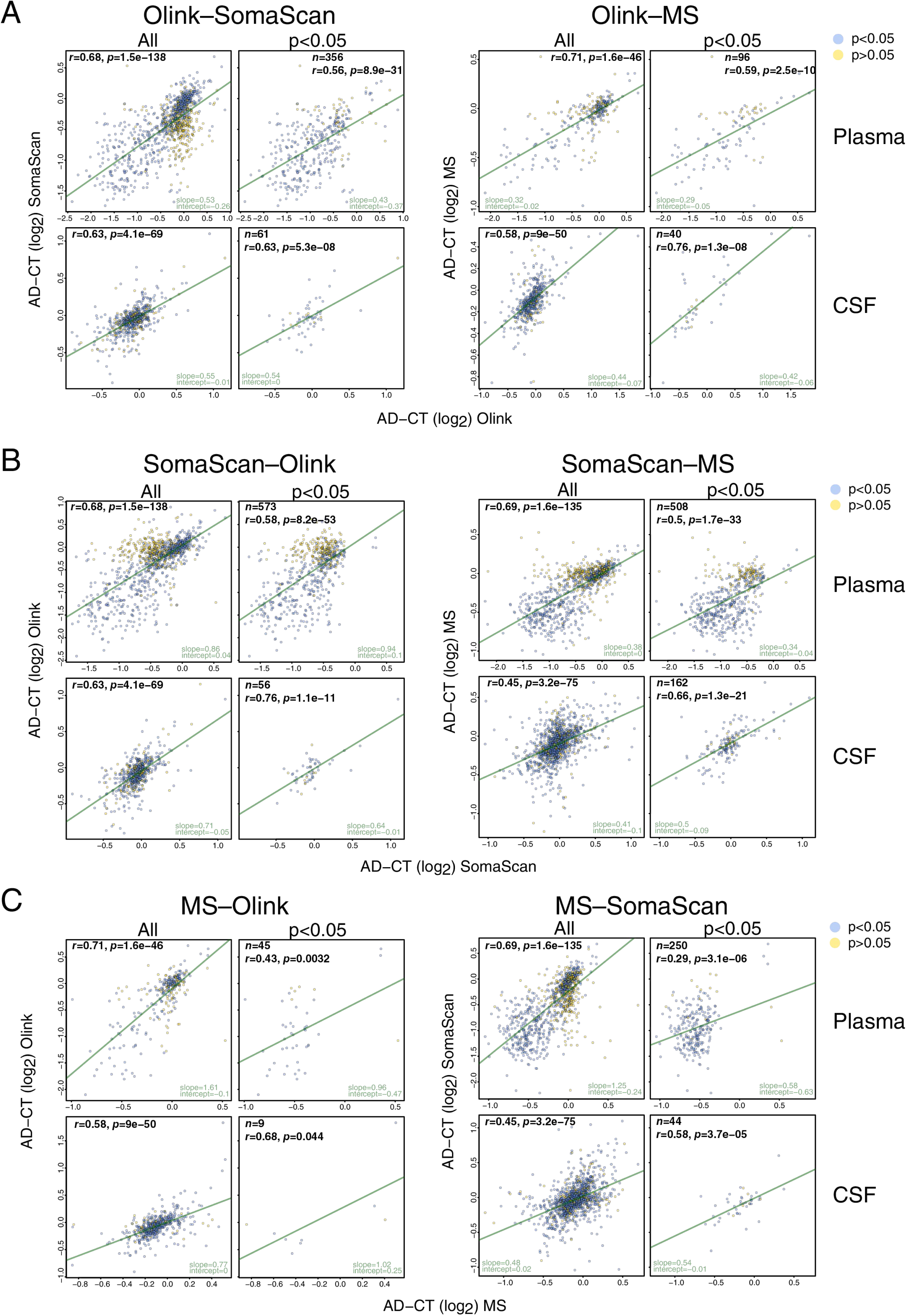
Correlation of AD Minus Control Differences Between Platforms Across Subjects. (A) Correlation between Olink and SomaScan (left) and Olink and MS (right) AD minus control (AD–CT) values, in plasma (top) and CSF (bottom). Correlations are provided for all AD–CT values (“All”), and AD–CT values in Olink that are significant at a level of *p*<0.05 (“*p*<0.05”). The individual proteins are colored according to whether they are significantly correlated within subject between platforms as shown in Figure 3. (B) Same as in (A), except with correlation between SomaScan and Olink (left) and SomaScan and MS (right) AD–CT values. (C) Same as in (A) and (B), except with correlation between MS and Olink (left) and MS and SomaScan (right). The number of significantly differentially expressed proteins used for correlation is provided in the respective panel. Correlations were performed with Pearson test. The green lines indicate lines of best fit for each dataset.

**Supplementary Figure 6.**
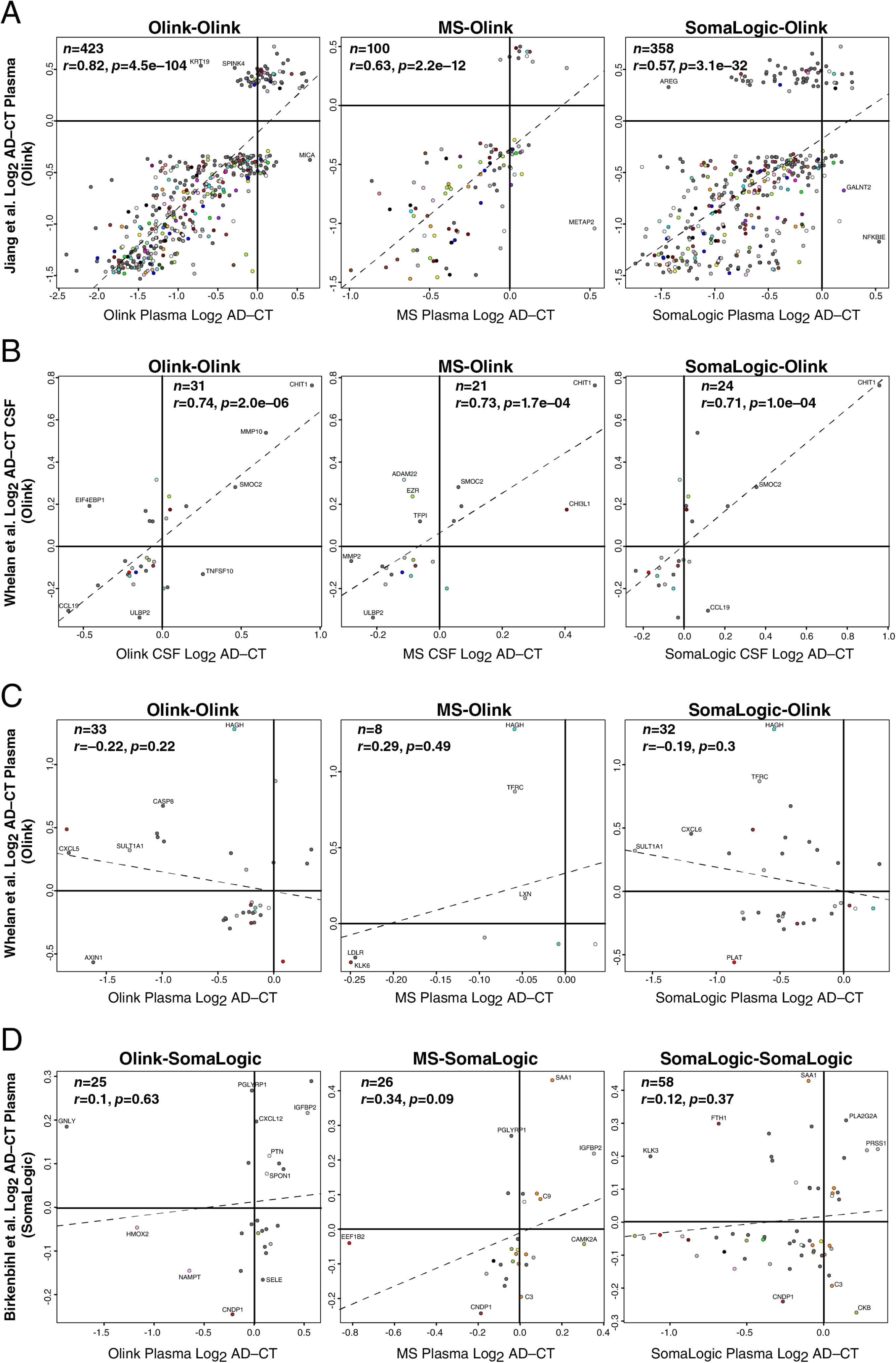
Correlation of CSF and Plasma Measurements with Other Cohorts. (A-D) Correlation of Emory cohort SomaLogic, Olink, and MS data with SomaLogic and Olink data from other cohorts. Only proteins considered significantly different between AD and control cases at 5% FDR in each cohort were considered for correlation. (A) Emory data from each platform was correlated with Olink plasma data from Jiang *et al*.^18^ (B) Correlation with CSF Olink data from the BioFinder cohort as described in Whelan *et al*.^19^ (C) Correlation with plasma Olink data from Whelan *et al*. (D) Correlation with plasma SomaLogic data from the ANMerge version of the AddNeuroMed dataset as described in Birkenbihl *et al*.^20^ Proteins are colored by the brain co-expression module in which they reside, as described in Johnson *et al*.^2^

**Supplementary Figure 7.**
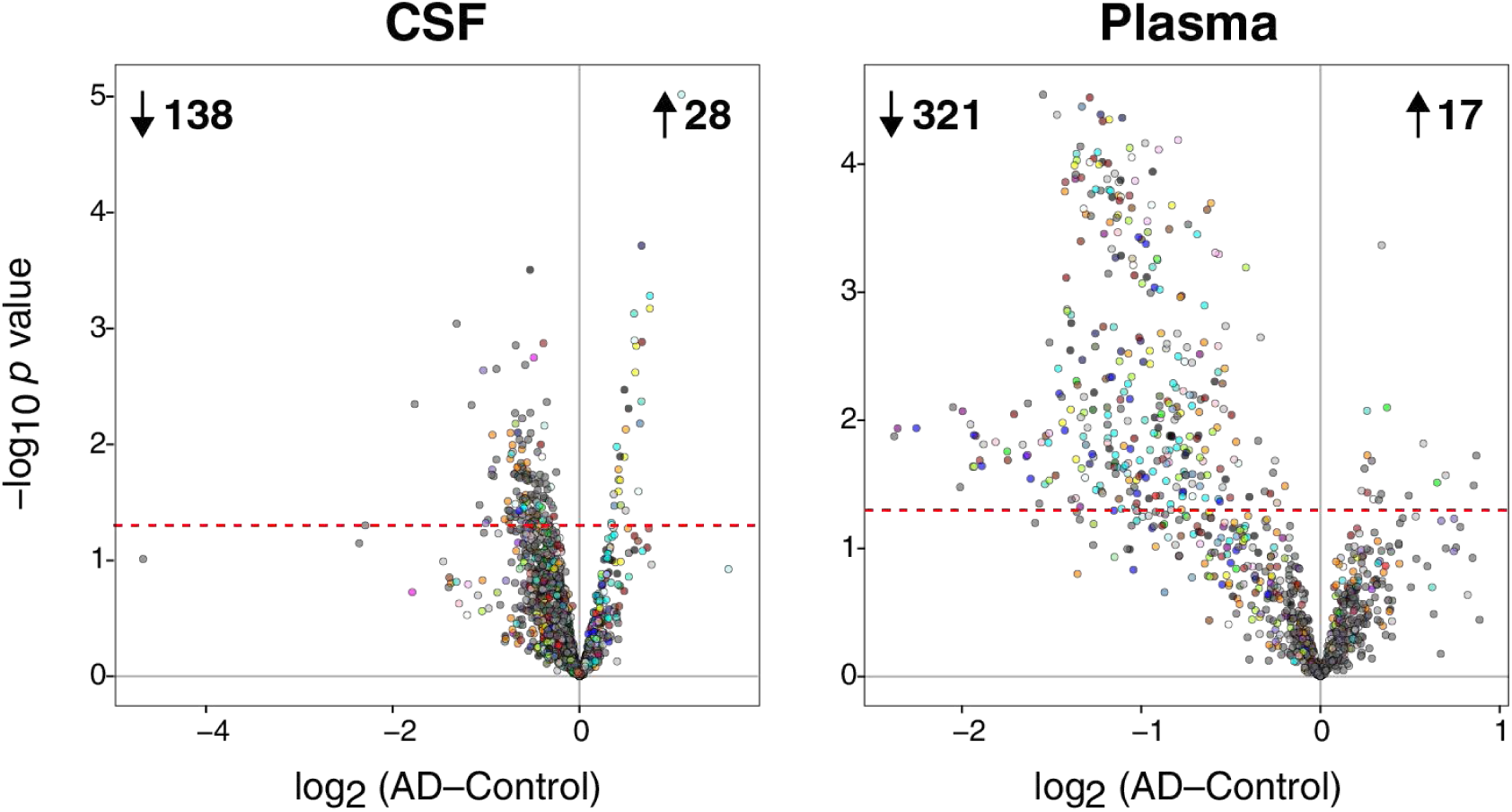
Differential Protein Abundance in AD by Undepleted TMT-MS. Differential protein abundance between AD and control cases in CSF (left) and plasma (right) on the TMT-MS undepleted platform. Proteins that are above the dashed red line are significantly altered in AD by *t* test at *p*<0.05. Proteins are colored by the brain co-expression module in which they reside, as described in Johnson *et al*.^2^

**Supplementary Figure 8.**
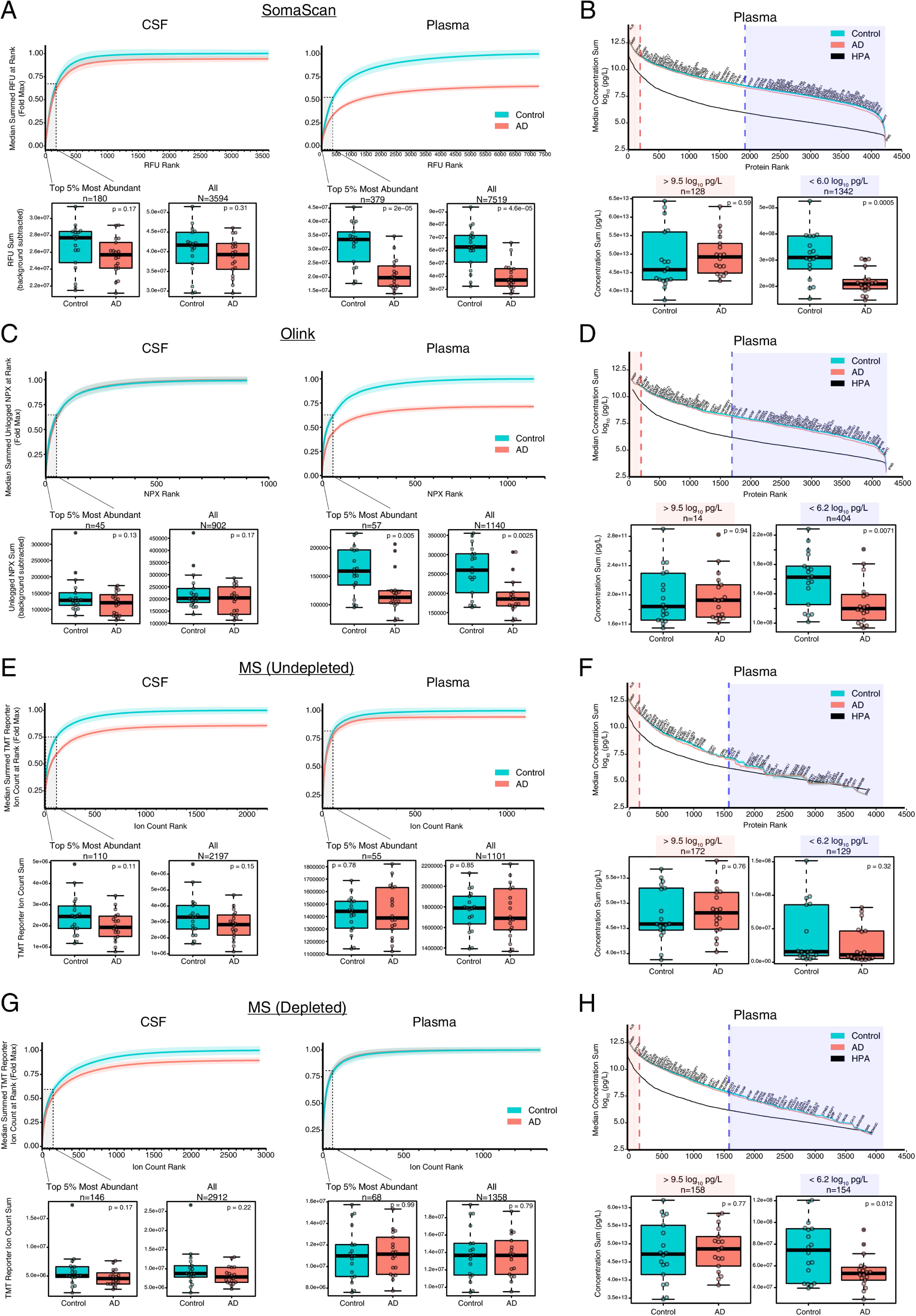
Protein Abundance Analysis By Platform in CSF and Plasma. (A-H) Individual and total protein signal levels as measures of relative abundance in CSF and plasma were analyzed on the SomaScan (A, B), Olink (C, D), undepleted tandem mass tag mass spectrometry (TMT-MS) (E, F), and depleted TMT-MS platforms (G, H) in control (*n*=18) and AD (*n*=18) subjects. (A) Aptamer relative fluorescence units (RFUs) were ranked and analyzed by the contribution of each aptamer RFU to the cumulative RFU in CSF (left) or plasma (right). The difference in RFUs between AD and control cases for the top 5% protein RFUs (shaded box) and all protein RFUs for each fluid are shown in the boxplots below. Background signal was subtracted for each RFU. RFUs below limit of detection (LOD) were not considered. (B) SomaScan aptamer RFUs were calibrated to proteins of known absolute plasma concentration from the Human Protein Atlas^23^ (*n*=2685 out of 4226 proteins in the HPA), and proteins ranked by concentration. The black line indicates the absolute concentration values from the HPA and is not summed, whereas the turquoise and red lines indicate the summed total protein concentration for all proteins below rank in control and AD cases, respectively. Boxplots represent summed concentration in control and AD cases for the top 5% (>9.5 log_10_ pg/L) and bottom 50% (<6.0 log_10_ pg/L) of proteins measured that overlap with HPA proteins. (C) Olink unlogged normalized protein expression (NPX) values were ranked and analyzed in CSF (left) and plasma (right) in similar fashion as shown in (A). Background signal was subtracted from unlogged NPX values. NPX values below LOD were not considered. (D) Olink NPX values were calibrated to proteins of known absolute plasma concentration and ranked, as described in (B) (*n*=808 out of 4226 proteins in the HPA). Boxplots represent summed concentration in control and AD cases for the top 5% (>9.5 log_10_ pg/L) and bottom 50% (<6.2 log_10_ pg/L) of proteins measured that overlap with HPA proteins. (E) Undepleted TMT mass spectrometry summed reporter ion counts for each protein were ranked and analyzed in CSF (left) and plasma (right) in similar fashion as shown in (A) and (C). (F) Undepleted TMT-MS ion counts were calibrated to proteins of known absolute plasma concentration and ranked, as described in (B) and (D) (*n*=925 out of 4226 proteins in the HPA). The red shaded area represents the top 10% most abundant proteins, whereas the blue shaded area represents the bottom 30% of abundance at a threshold of <6.2 pg/L in the undepleted MS data. (G) Depleted TMT-MS ion counts were analyzed as described in (E). (H) Depleted TMT-MS ion counts were analyzed as described in (F) (*n*=1163 out of 4226 proteins in the HPA). The blue shaded area represents the bottom 30% of abundance at a threshold of <6.2 pg/L in the depleted MS data. Shaded areas represent +/-SEM. Differences between control and AD were determined by *t* test.

**Supplementary Figure 9.**
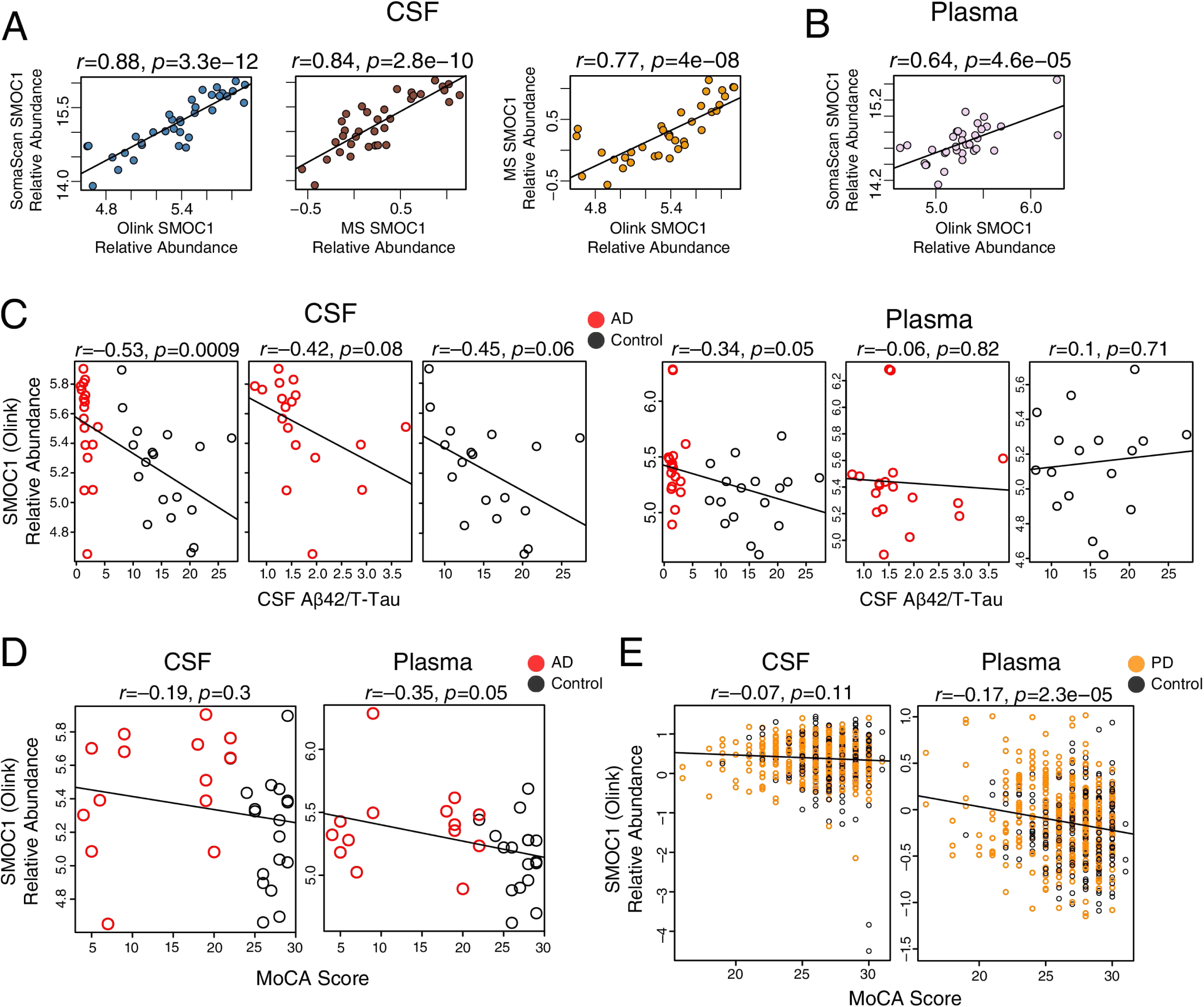
SMOC1 Analyses. (A-E) Correlation of SMOC1 relative abundance in CSF between proteomic platforms (A). (B) Correlation of SMOC1 relative abundance in plasma between Olink and SomaScan platforms. SMOC1 was not measured by MS in plasma. Correlations in (A) and (B) were performed with the SMOC1 5694.57 SOMAmer. (C) Correlation of SMOC1 relative abundance as measured by Olink with CSF Aβ/T-Tau ratio in CSF (left) and plasma (right). (D) Correlation of SMOC1 relative abundance with MoCA score, a measure of cognitive performance (higher scores reflect better cognitive performance), in CSF (left) and plasma (right). (E) Correlation of SMOC1 relative abundance with MoCA score in CSF (left) and plasma (right) in the Accelerating Medicines Partnership – Parkinson’s Disease (AMP-PD) cohort. Correlations were performed using Pearson correlation. Aβ, amyloid-β; MoCA, Montreal Cognitive Assessment; MS, mass spectrometry; SMOC1, SPARC-related modular calcium-binding protein 1; T-Tau, total tau.

**Supplementary Figure 10.**
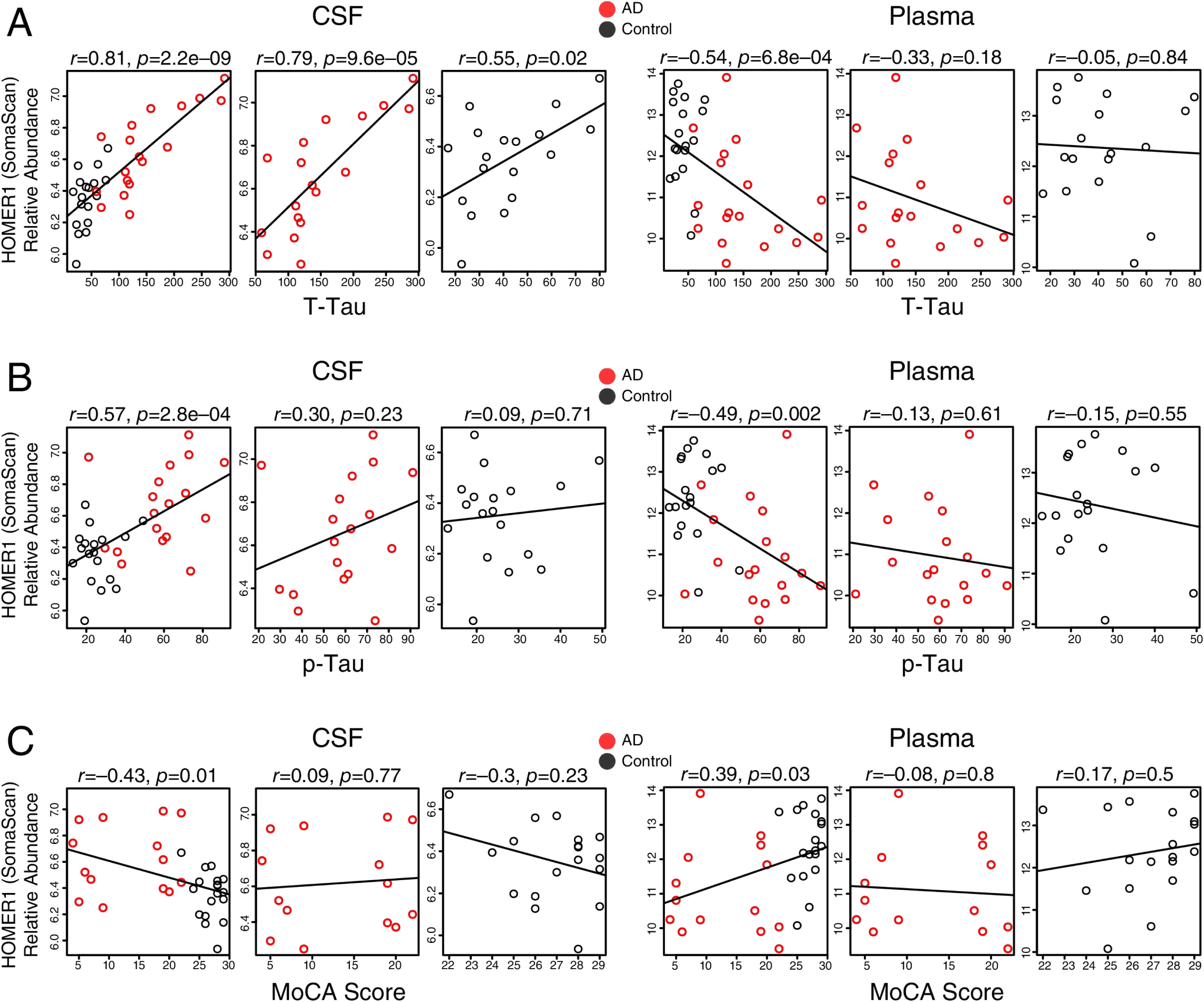
HOMER1 Analyses. (A-C) Correlation of HOMER1 relative abundance as measured by SomaScan in CSF (left) and plasma (right) with CSF total tau (T-Tau) levels (A). (B) Correlation of HOMER1 relative abundance in CSF (left) and plasma (right) with CSF phosphorylated tau181 (p-Tau) levels. (C) Correlation of HOMER1 relative abundance in CSF (left) and plasma (right) with Montreal Cognitive Assessment score (MoCA, higher scores reflect better cognitive performance). HOMER1 was measured in CSF and plasma only by SomaScan. Correlations were performed using Pearson correlation. HOMER1, Homer protein homolog 1.

**Supplementary Figure 11.**
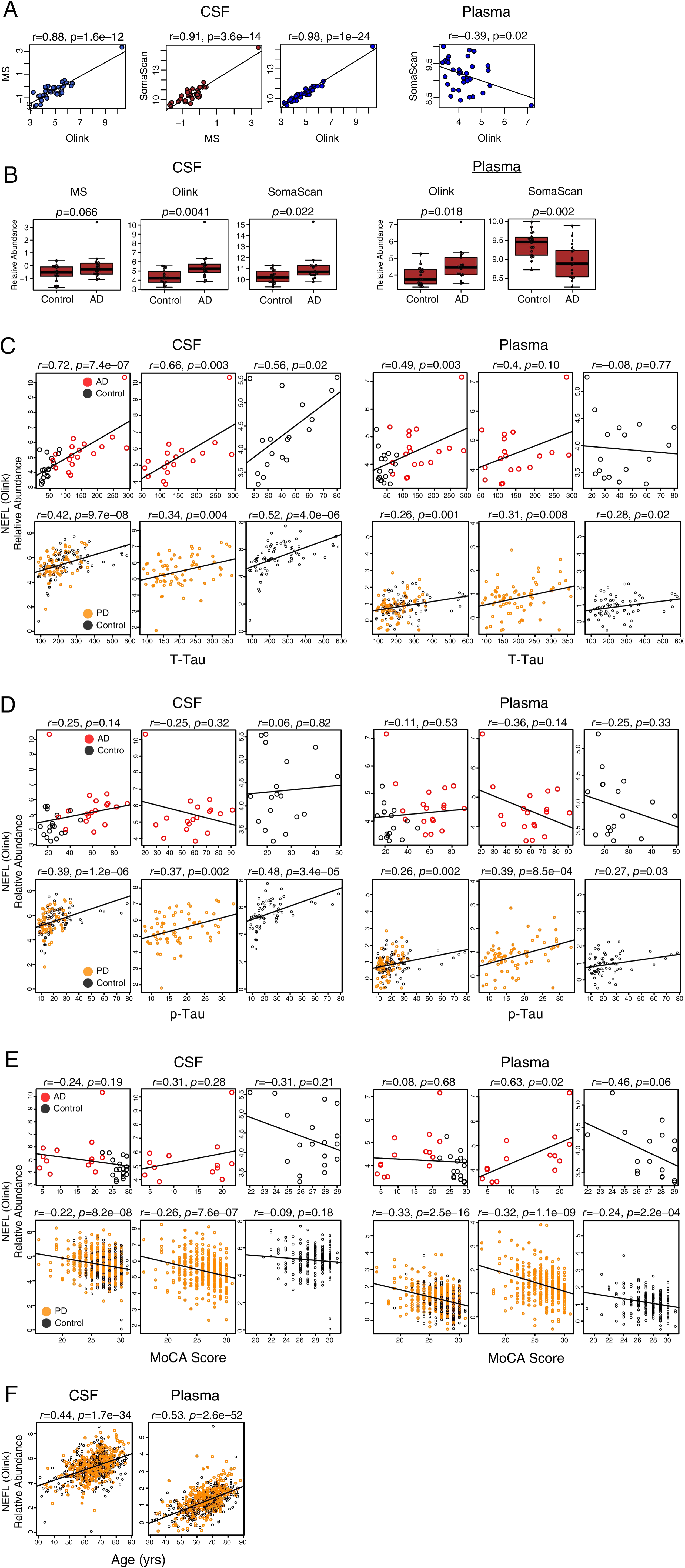
NEFL Analyses. (A-F) Correlation of NEFL relative abundance in CSF (left) and plasma (right) between proteomic platforms (A). NEFL was not measured in plasma by MS. (B) Differences in NEFL relative abundance between control and AD cases by platform in CSF (left) and plasma (right). (C) Correlation of NEFL relative abundance as measured by Olink in CSF (left) and plasma (right) with CSF T-Tau levels in the AD discovery cohort (top) and AMP-PD cohort (bottom). (D) Correlation of NEFL relative abundance in CSF (left) and plasma (right) with CSF p-Tau levels in the AD discovery cohort (top) and AMP-PD cohort (bottom). (E) Correlation of NEFL relative abundance in CSF (left) and plasma (right) with MoCA score in the AD discovery cohort (top) and AMP-PD cohort (bottom). (F) Correlation of NEFL relative abundance in CSF (left) and plasma (right) with age in the AMP-PD cohort. Correlations were performed using Pearson correlation. Group differences were assessed using *t* test. Boxplots represent the median, 25^th^, and 75^th^ percentiles, and box hinges represent the interquartile range of the two middle quartiles within a group. Datapoints up to 1.5 times the interquartile range from box hinge define the extent of whiskers (error bars). AMP-PD, Accelerating Medicines Partnership – Parkinson’s Disease; MoCA, Montreal Cognitive Assessment (higher scores reflect better cognitive performance); MS, mass spectrometry; NEFL, neurofilament light polypeptide; p-Tau, phosphorylated tau181; T-Tau, total tau.

**Supplementary Figure 12.**
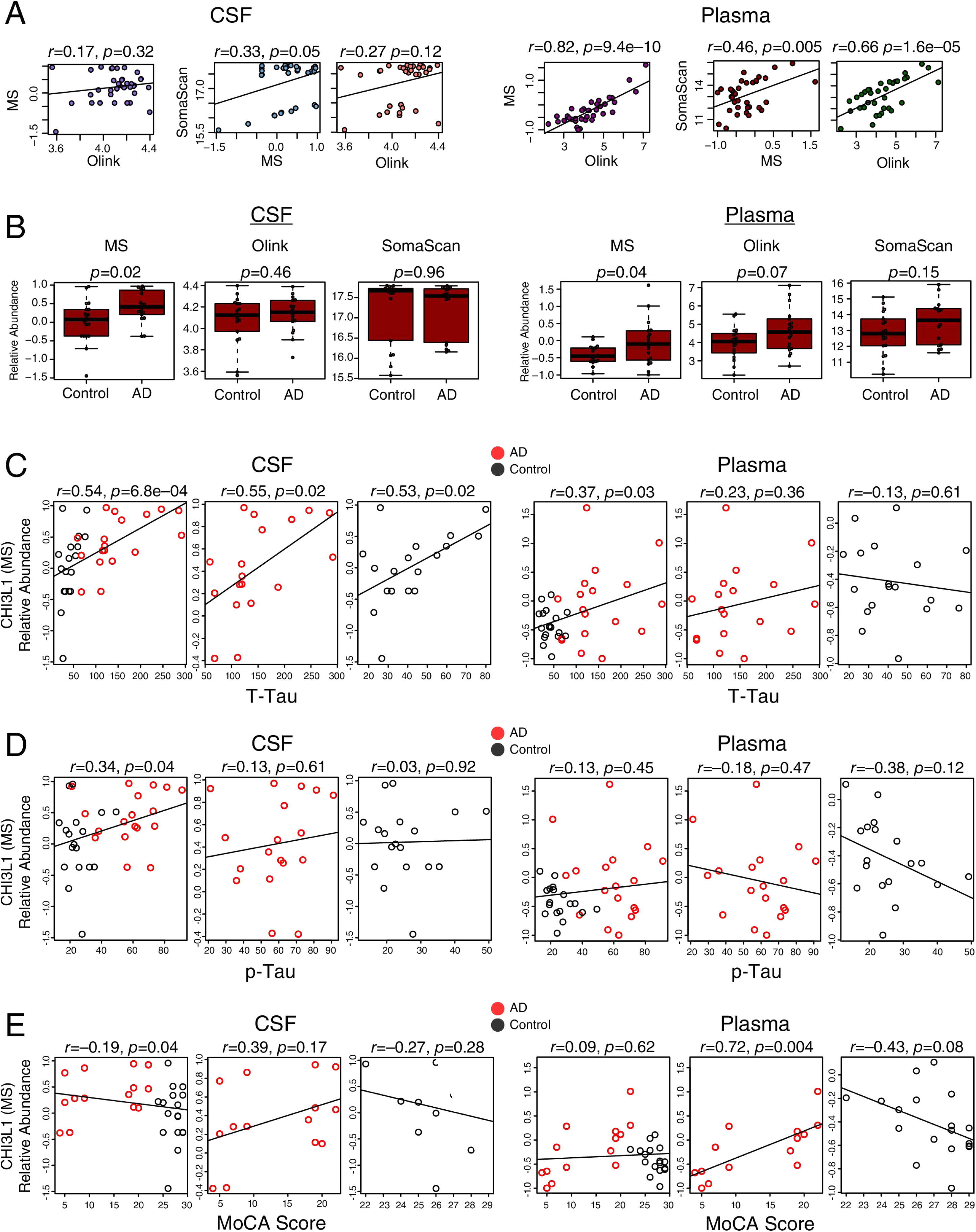
CHI3L1 Analyses. (A-E) Correlation of CHI3L1 relative abundance in CSF (left) and plasma (right) between proteomic platforms (A). (B) Differences in CHI3L1 relative abundance between control and AD cases by platform in CSF (left) and plasma (right). (C) Correlation of CHI3L1 relative abundance as measured by MS in CSF (left) and plasma (right) with CSF T-Tau levels. (D) Correlation of CHI3L1 relative abundance in CSF (left) and plasma (right) with CSF p-Tau levels. (E) Correlation of CHI3L1 relative abundance in CSF (left) and plasma (right) with MoCA score. Correlations were performed using Pearson correlation. Group differences were assessed using *t* test. Boxplots represent the median, 25^th^, and 75^th^ percentiles, and box hinges represent the interquartile range of the two middle quartiles within a group. Datapoints up to 1.5 times the interquartile range from box hinge define the extent of whiskers (error bars). CHI3L1, Chitinase-3-like protein 1; MoCA, Montreal Cognitive Assessment (higher scores reflect better cognitive performance); MS, mass spectrometry; p-Tau, phosphorylated tau181; T-Tau, total tau.

**Supplementary Figure 13.**
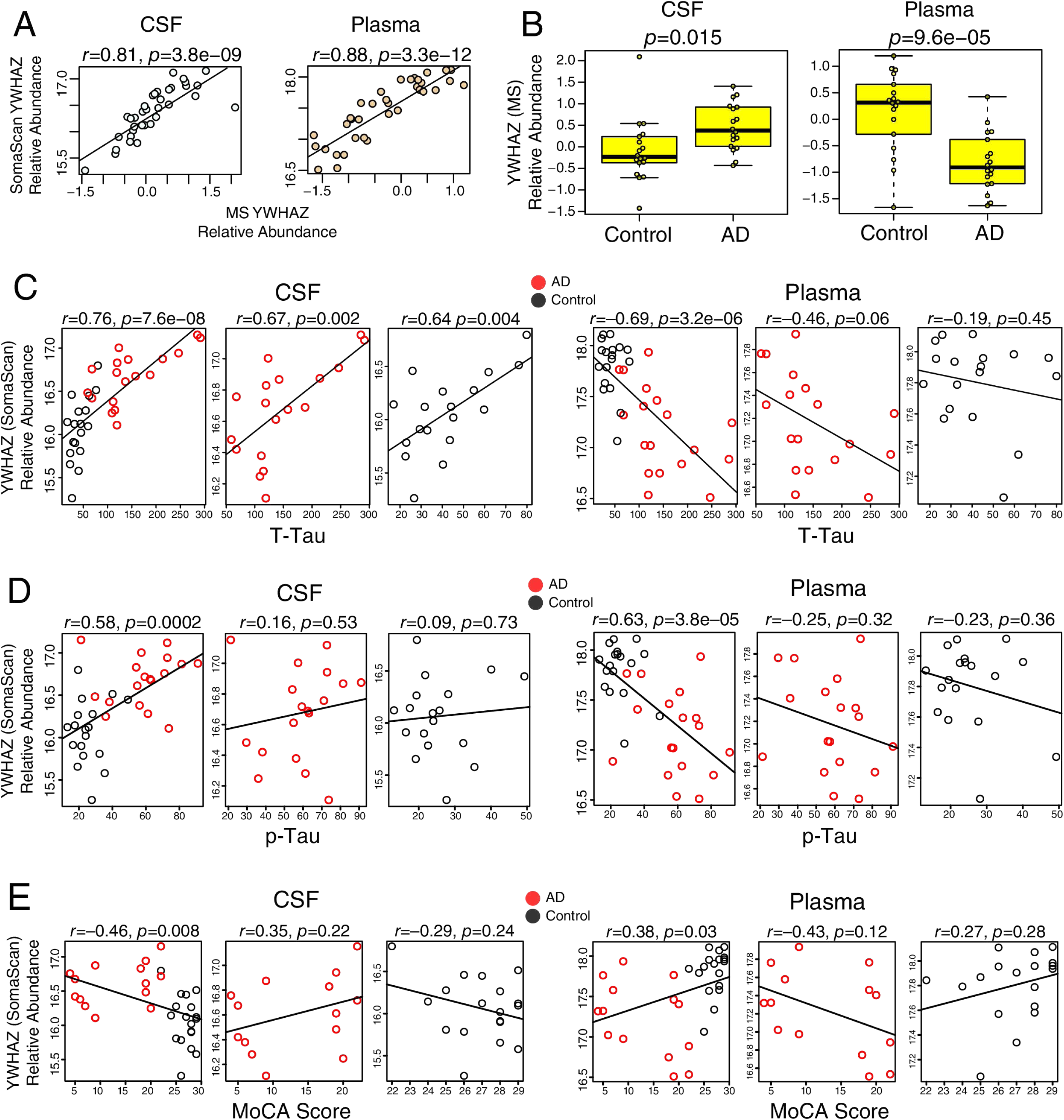
YWHAZ Analyses. (A-E) Correlation of YWHAZ relative abundance in CSF (left) and plasma (right) between SomaScan and MS platforms (A). YWHAZ was not measured by Olink. (B) Differences in YWHAZ relative abundance as measured by MS between control and AD cases in CSF (left) and plasma (right). (C) Correlation of YWHAZ relative abundance as measured by MS in CSF (left) and plasma (right) with CSF T-Tau levels. (D) Correlation of YWHAZ relative abundance in CSF (left) and plasma (right) with CSF p-Tau levels. (E) Correlation of YWHAZ relative abundance in CSF (left) and plasma (right) with MoCA score. Correlations were performed using Pearson correlation. Group differences were assessed using *t* test. Boxplots represent the median, 25^th^, and 75^th^ percentiles, and box hinges represent the interquartile range of the two middle quartiles within a group. Datapoints up to 1.5 times the interquartile range from box hinge define the extent of whiskers (error bars). MoCA, Montreal Cognitive Assessment (higher scores reflect better cognitive performance); MS, mass spectrometry; p-Tau, phosphorylated tau181; T-Tau, total tau; YWHAZ, 14-3-3 protein zeta.

**Supplementary Figure 14.**
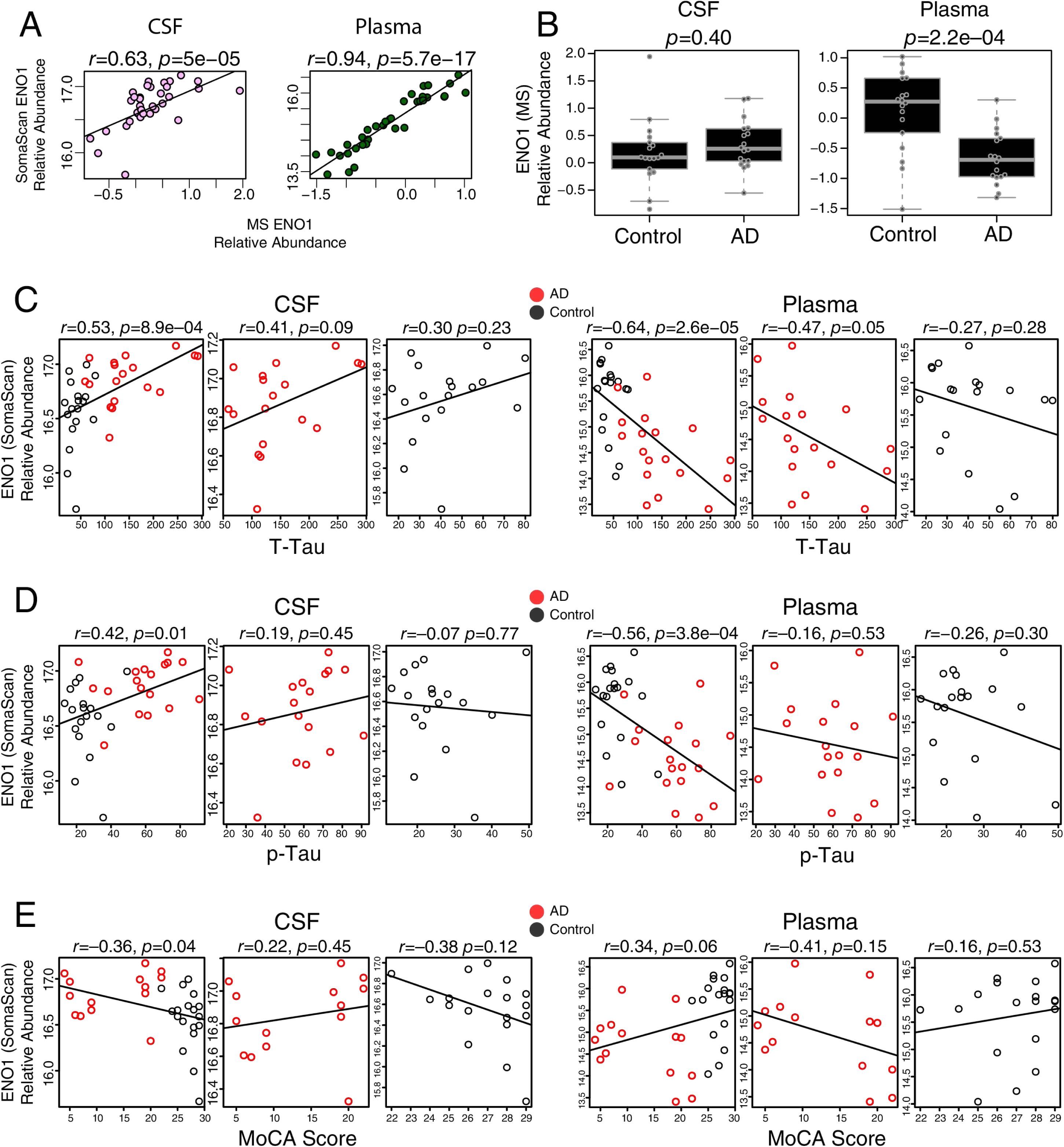
ENO1 Analyses. (A-E) Correlation of ENO1 relative abundance in CSF (left) and plasma (right) between SomaScan and MS platforms (A). ENO1 was not measured by Olink. (B) Differences in ENO1 relative abundance as measured by MS between control and AD cases in CSF (left) and plasma (right). (C) Correlation of ENO1 relative abundance as measured by SomaScan in CSF (left) and plasma (right) with CSF T-Tau levels. (D) Correlation of ENO1 relative abundance in CSF (left) and plasma (right) with CSF p-Tau levels. (E) Correlation of ENO1 relative abundance in CSF (left) and plasma (right) with MoCA score. Correlations were performed using Pearson correlation. Group differences were assessed using *t* test. Boxplots represent the median, 25^th^, and 75^th^ percentiles, and box hinges represent the interquartile range of the two middle quartiles within a group. Datapoints up to 1.5 times the interquartile range from box hinge define the extent of whiskers (error bars). ENO1, alpha-enolase; MoCA, Montreal Cognitive Assessment (higher scores reflect better cognitive performance); MS, mass spectrometry; p-Tau, phosphorylated tau181; T-Tau, total tau.

**Supplementary Figure 15.**
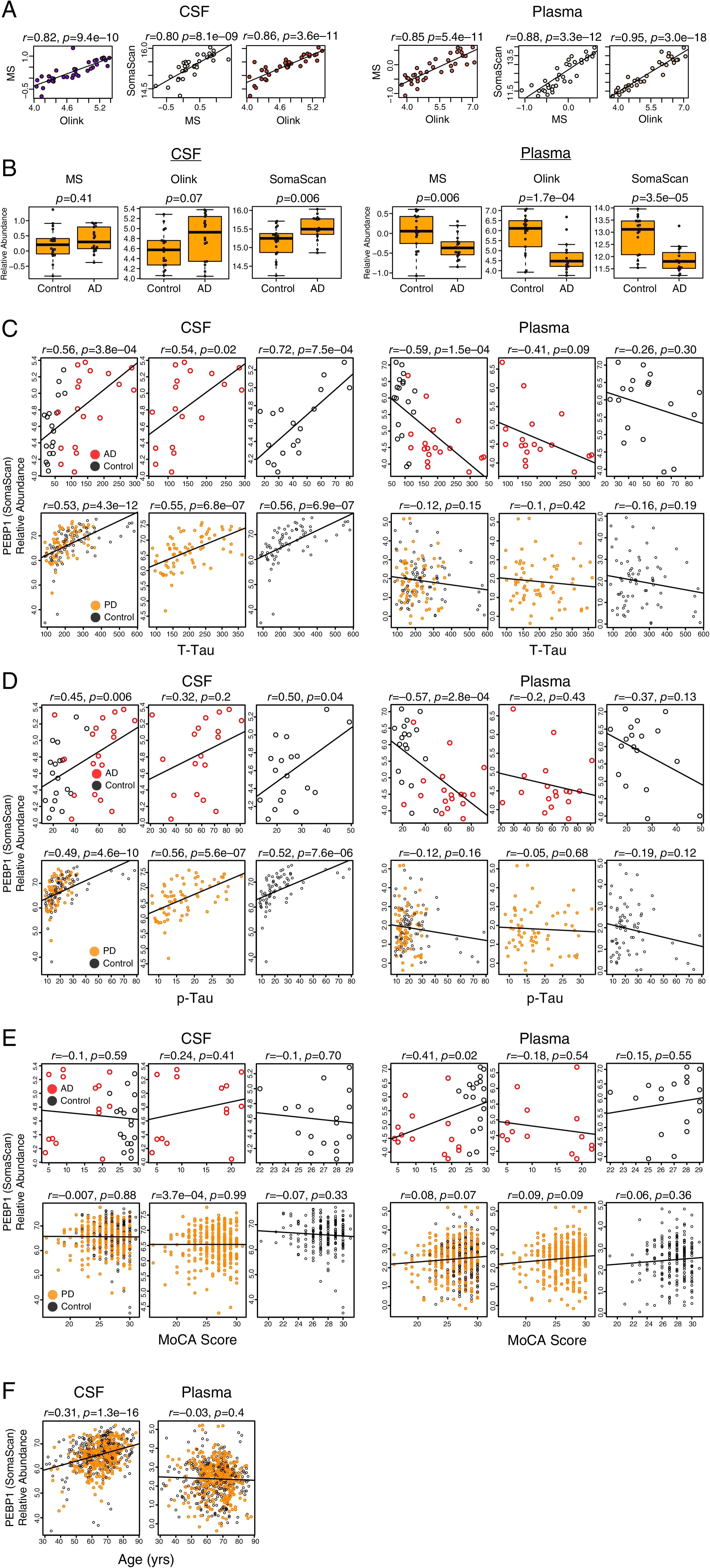
PEBP1 Analyses. (A-F) Correlation of PEBP1 relative abundance in CSF (left) and plasma (right) between proteomic platforms (A). (B) Differences in PEBP1 relative abundance between control and AD cases by platform in CSF (left) and plasma (right). (C) Correlation of PEBP1 relative abundance as measured by SomaScan in CSF (left) and plasma (right) with CSF T-Tau levels in the AD discovery cohort (top) and AMP-PD cohort (bottom). (D) Correlation of PEBP1 relative abundance in CSF (left) and plasma (right) with CSF p-Tau levels in the AD discovery cohort (top) and AMP-PD cohort (bottom). (E) Correlation of PEBP1 relative abundance in CSF (left) and plasma (right) with MoCA score in the AD discovery cohort (top) and AMP-PD cohort (bottom). (F) Correlation of PEBP1 relative abundance in CSF (left) and plasma (right) with age in the AMP-PD cohort. Correlations were performed using Pearson correlation. Group differences were assessed using *t* test. Boxplots represent the median, 25^th^, and 75^th^ percentiles, and box hinges represent the interquartile range of the two middle quartiles within a group. Datapoints up to 1.5 times the interquartile range from box hinge define the extent of whiskers (error bars). AMP-PD, Accelerating Medicines Partnership – Parkinson’s Disease; MoCA, Montreal Cognitive Assessment (higher scores reflect better cognitive performance); MS, mass spectrometry; PEBP1, phosphatidylethanolamine-binding protein 1; p-Tau, phosphorylated tau181; T-Tau, total tau.

**Supplementary Figure 16.**
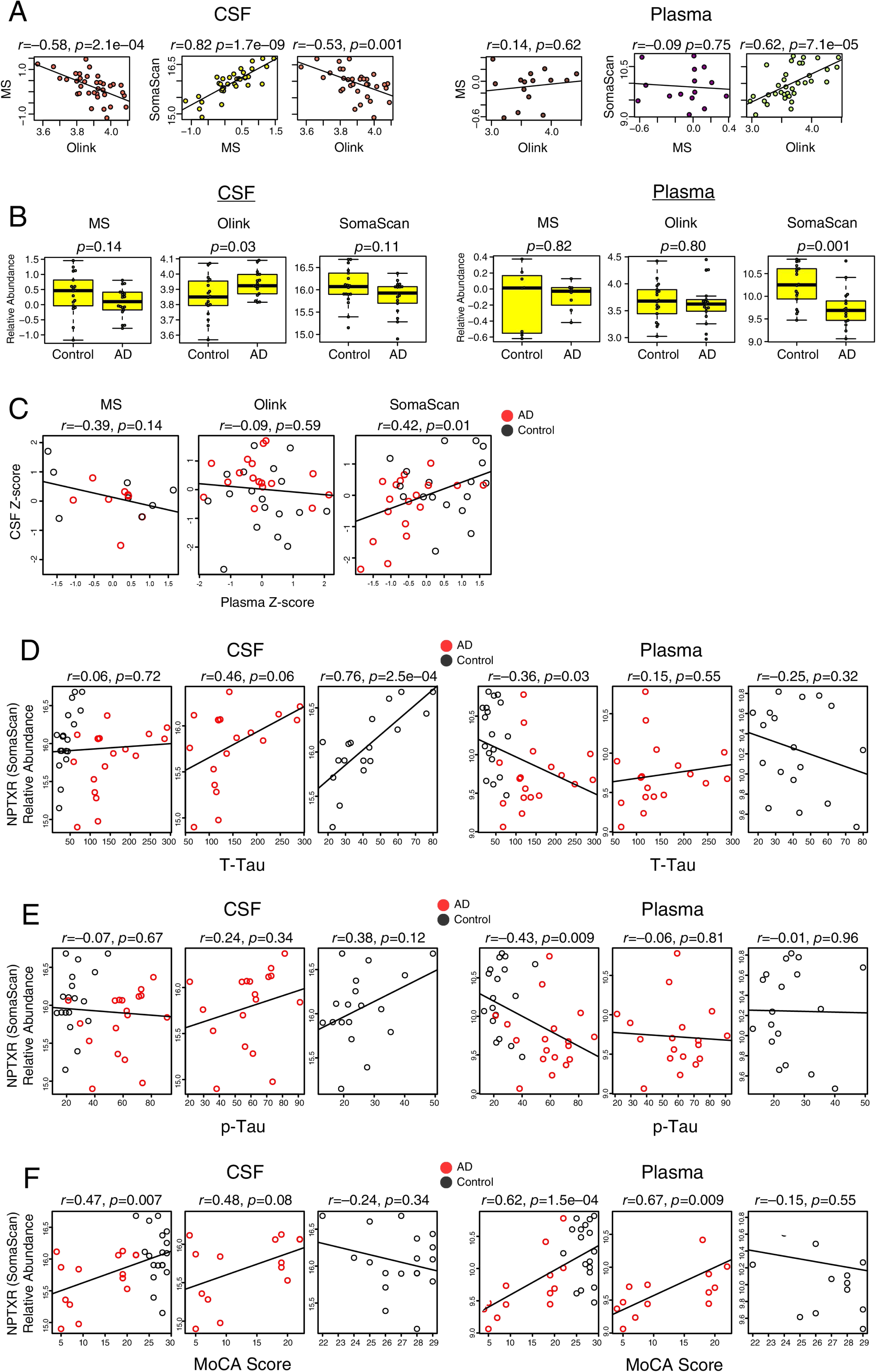
NPTXR Analyses. (A-F) Correlation of NPTXR relative abundance in CSF (left) and plasma (right) between proteomic platforms (A). (B) Differences in NPTXR relative abundance between control and AD cases by platform in CSF (left) and plasma (right). (C) Correlation of relative NPTXR levels measured in each platform between CSF and plasma. (D) Correlation of NPTXR relative abundance as measured by SomaScan in CSF (left) and plasma (right) with CSF T-Tau levels. (E) Correlation of NPTXR relative abundance in CSF (left) and plasma (right) with CSF p-Tau levels. (F) Correlation of PEBP1 relative abundance in CSF (left) and plasma (right) with MoCA score. Correlations were performed using Pearson correlation. Group differences were assessed using *t* test. Boxplots represent the median, 25^th^, and 75^th^ percentiles, and box hinges represent the interquartile range of the two middle quartiles within a group. Datapoints up to 1.5 times the interquartile range from box hinge define the extent of whiskers (error bars). MoCA, Montreal Cognitive Assessment (higher scores reflect better cognitive performance); MS, mass spectrometry; NPTXR, neuronal pentraxin receptor; p-Tau, phosphorylated tau181; T-Tau, total tau.

**Supplementary Figure 17.**
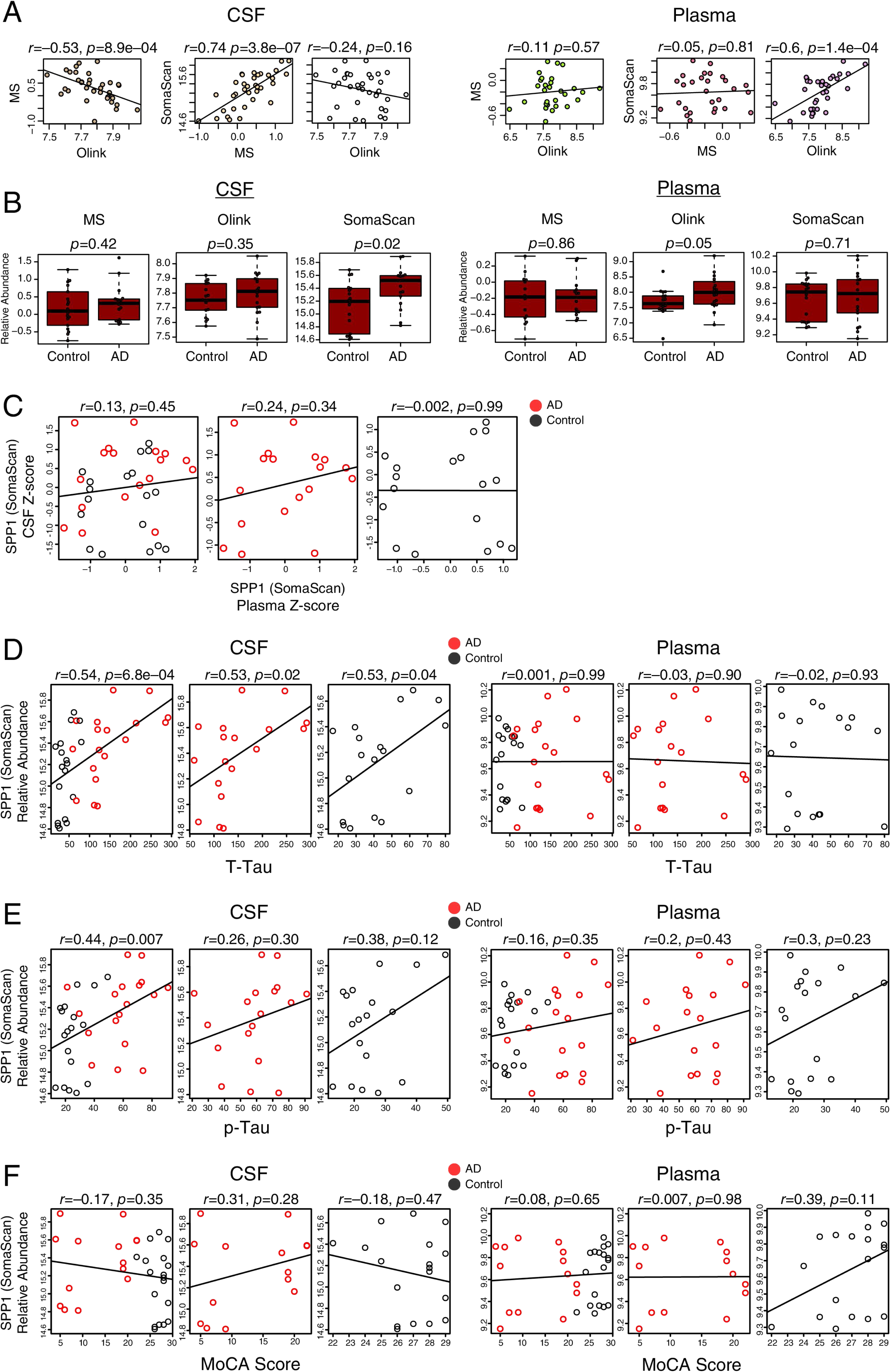
SPP1 Analyses. (A-F) Correlation of SPP1 relative abundance in CSF (left) and plasma (right) between proteomic platforms (A). (B) Differences in SPP1 relative abundance between control and AD cases by platform in CSF (left) and plasma (right). (C) Correlation of relative SPP1 levels measured by SomaScan between CSF and plasma. (D) Correlation of SPP1 relative abundance in CSF (left) and plasma (right) with CSF T-Tau levels. (E) Correlation of SPP1 relative abundance in CSF (left) and plasma (right) with CSF p-Tau levels. (F) Correlation of SPP1 relative abundance in CSF (left) and plasma (right) with MoCA score. Correlations were performed using Pearson correlation. Group differences were assessed using *t* test. Boxplots represent the median, 25^th^, and 75^th^ percentiles, and box hinges represent the interquartile range of the two middle quartiles within a group. Datapoints up to 1.5 times the interquartile range from box hinge define the extent of whiskers (error bars). MoCA, Montreal Cognitive Assessment (higher scores reflect better cognitive performance); MS, mass spectrometry; p-Tau, phosphorylated tau181; SPP1, osteopontin; T-Tau, total tau.

**Supplementary Figure 18.**
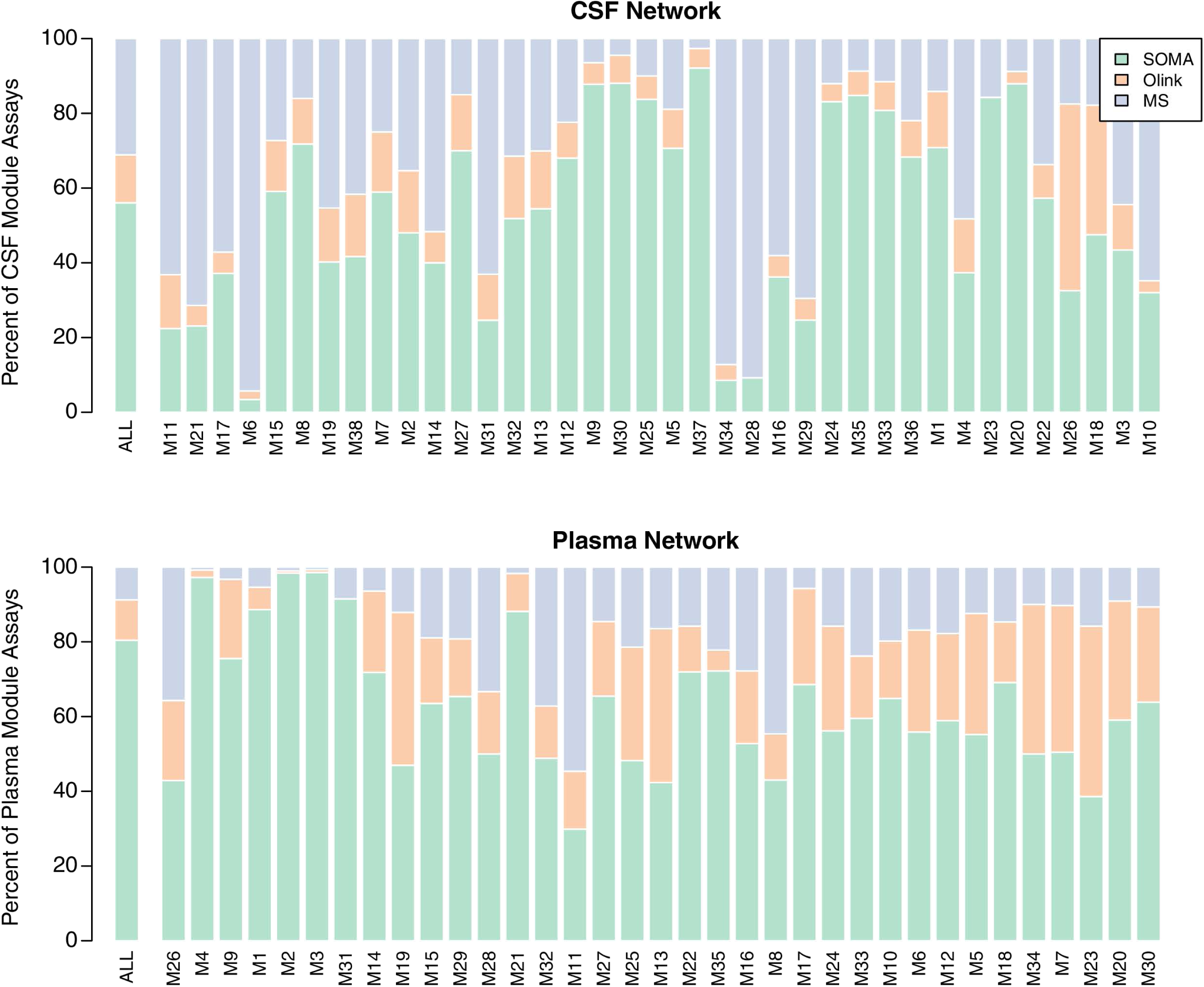
CSF and Plasma Network Platform Representation. Percent coverage of CSF (top) and plasma (bottom) network modules by proteomic platform. Modules are listed in relationship order. “All” indicates percent coverage across the entire network.

**Supplementary Figure 19.**
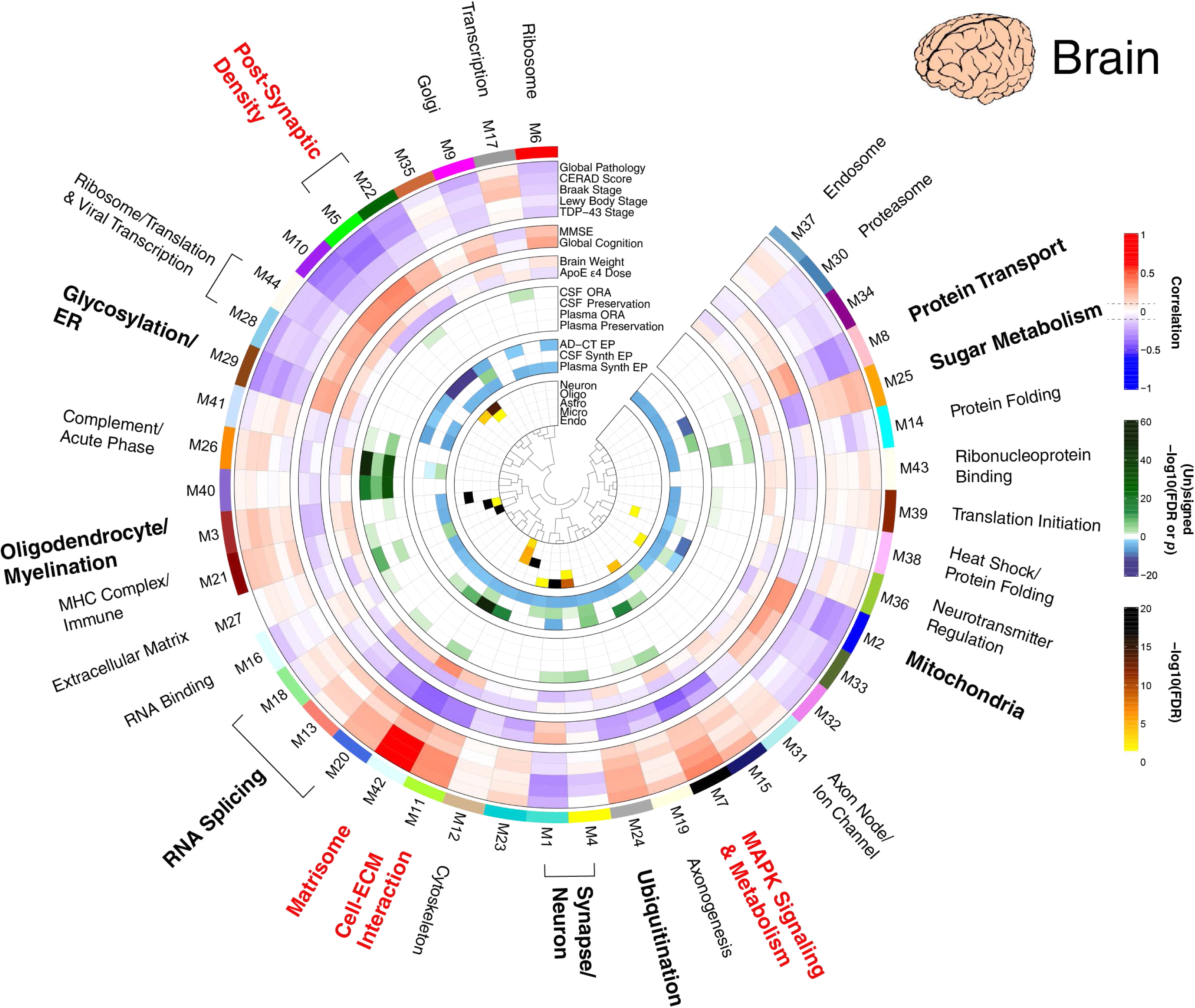
AD Brain Protein Co-Expression Network. AD brain consensus protein correlation network as shown in Figure 5, including integration with CSF and plasma data. Descriptions of module eigenprotein (EP) correlations with traits and cell type overlap testing are provided in Figure 5. Brain modules were tested for their presence in the CSF and plasma networks by brain module protein overrepresentation analysis (ORA) and network preservation (preservation) statistics, as previously described^2^. ORA *p* values are for the module with the strongest overlap. Only modules that reached statistical significance after FDR correction are colored by degree of significance. In addition to module overlap and preservation analyses, the difference in brain module eigenprotein between control and AD, or brain synthetic eigenprotein in CSF or plasma between control and AD, was determined. A significantly increased eigenprotein in AD is indicated in green, whereas a significantly decreased eigenprotein is indicated in blue.

**Supplementary Figure 20.**
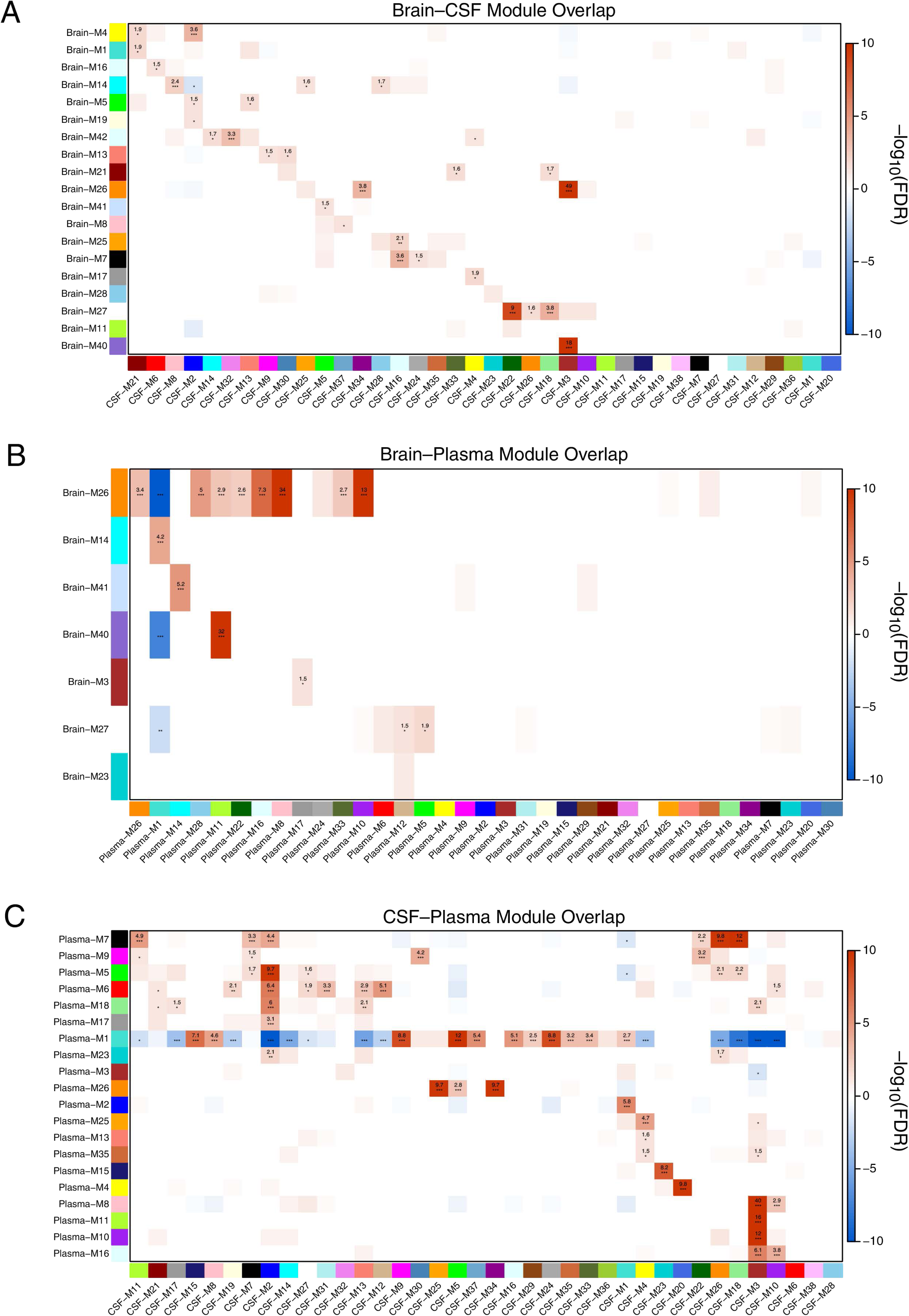
Module Over-Representation Analysis Across Brain and Biofluid Networks. (A-C) Module member overrepresentation analysis (ORA) of the brain and CSF networks (A), brain and plasma networks (B), and CSF and plasma networks (C). The numbers in each box represent the –log_10_(FDR) value for the overlap after Benjamini-Hochberg correction. Modules on the y-axis (rows) without an overlap FDR value of – log_10_ (FDR) > 1 were not included in the heatmaps.

**Supplementary Figure 21.**
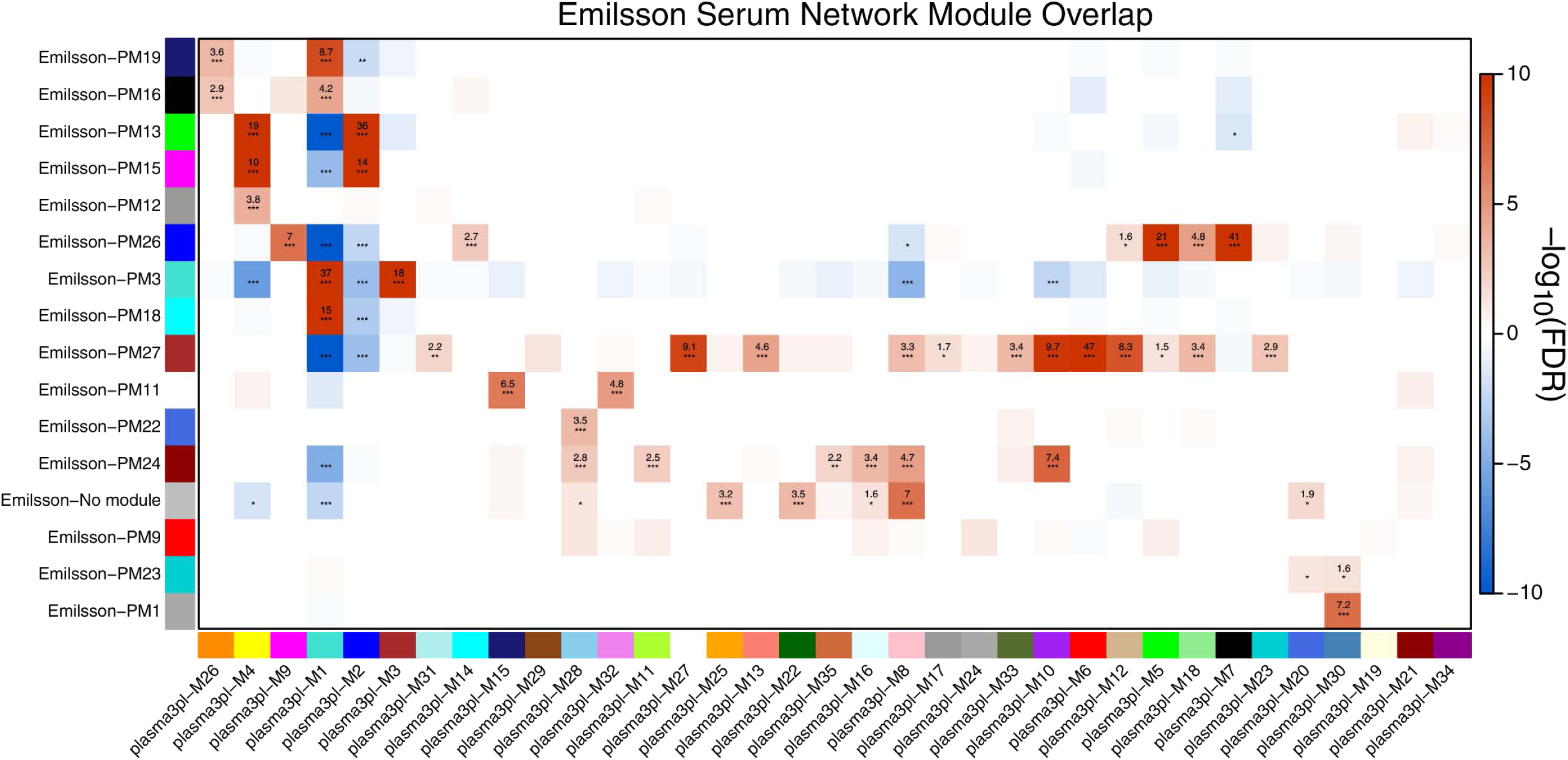
Plasma Network Module Over-Representation Analysis with a SomaScan Serum Network. Module member overrepresentation analysis (ORA) of a serum protein co-expression network (Emilsson-PM) obtained using the SomaScan platform, as described in Emilsson *et al.*^27^, with the plasma 3-platform (plasma-3pl) network. Ontologies for each plasma network module are provided in Figure 9. The numbers in each box represent the –log_10_(FDR) value for the overlap after Benjamini-Hochberg correction. Modules on the y-axis (rows) without an overlap FDR value of – log_10_ (FDR) > 1 were not included in the heatmaps.

## Extended Data

**<u>Extended Data 1.</u> Correlation of Proteins Commonly Measured in CSF by MS-TMT and Olink.** *n*=36 unless otherwise indicated. Measurements were from TMT-MS on CSF depleted of highly abundant plasma proteins. Olink NPX values are shown on the x-axis, and TMT-MS log2 relative values are shown on the y-axis. Measurements that were below LOD in Olink are not outlined. Correlations include those proteins that were matched by gene symbol only. Correlations were performed using Pearson test.

**<u>Extended Data 2.</u> Correlation of Proteins Commonly Measured in CSF by MS-TMT and SomaScan.** *n*=35 unless otherwise indicated. Measurements were from TMT-MS on CSF depleted of highly abundant plasma proteins. TMT-MS log_2_ relative values are shown on the x-axis, and SomaScan log_2_(RFU) values are shown on the y-axis. Measurements that were below LOD in SomaScan are not outlined. Correlations were performed for all SOMAmers including multiple SOMAmers targeting the same protein, and include those proteins that were matched by gene symbol only. Correlations were performed using Pearson test.

**<u>Extended Data 3.</u> Correlation of Proteins Commonly Measured in CSF by Olink and SomaScan.** *n*=35 unless otherwise indicated. Olink NPX values are shown on the x-axis, and SomaScan log2(RFU) values are shown on the y-axis. Measurements that were below LOD in either platform are not outlined. Correlations were performed for all SOMAmers including multiple SOMAmers targeting the same protein, and include those proteins that were matched by gene symbol only. Correlations were performed using Pearson test.

**<u>Extended Data 4.</u> GO Analysis on AD CSF Network Modules.** Gene ontology (GO) analysis was performed to gain insight into the biological meaning of each AD CSF protein network module. Enrichment for a given ontology is shown by *z* score.

**<u>Extended Data 5.</u> AD CSF Network Module Protein Graphs.** The size of each circle indicates the relative eigenprotein correlation value (kME) in each network module. Those proteins with the largest kME are considered “hub” proteins within the module, and explain the largest variance in module expression. Proteins outlined in gold are from the SomaScan platform. Proteins outlined in green are from the Olink platform. Proteins outlined in purple are from the TMT-MS platform. Only the top 100 proteins by kME for each module are shown.

**<u>Extended Data 6.</u> AD CSF Network Module Eigenprotein Levels and Correlations.** *n*=18 control, 17 AD. Differences between case groups were assessed by *t* test or one-way ANOVA. Correlations were performed by bicor or Pearson test (cor). Significance at p<0.05 is highlighted in red. Boxplots represent the median, 25th, and 75th percentiles, and box hinges represent the interquartile range of the two middle quartiles within a group. Datapoints up to 1.5 times the interquartile range from box hinge define the extent of whiskers (error bars).

**<u>Extended Data 7.</u> AD CSF Network Module Synthetic Eigenprotein Levels and Correlations in Brain**. *n*=18 control, 17 AD. Differences between case groups were assessed by *t* test or one-way ANOVA. Correlations were performed by bicor or Pearson test (cor). Significance at p<0.05 is highlighted in red. Boxplots represent the median, 25th, and 75th percentiles, and box hinges represent the interquartile range of the two middle quartiles within a group. Datapoints up to 1.5 times the interquartile range from box hinge define the extent of whiskers (error bars).

**<u>Extended Data 8.</u> AD CSF Network Module Synthetic Eigenprotein Levels and Correlations in Plasma**. *n*=18 control, 17 AD. Differences between case groups were assessed by *t* test or one-way ANOVA. Correlations were performed by bicor or Pearson test (cor). Significance at p<0.05 is highlighted in red. Boxplots represent the median, 25th, and 75th percentiles, and box hinges represent the interquartile range of the two middle quartiles within a group. Datapoints up to 1.5 times the interquartile range from box hinge define the extent of whiskers (error bars).

**<u>Extended Data 9.</u> Within-Subject CSF Network Module Eigenprotein Levels in Plasma and CSF**. *n*=18 control, 17 AD. Relative CSF protein network module eigenprotein levels and their synthetic eigenprotein levels in plasma were compared within subject across fluids in control and AD cases. The difference in average slope (*Z*_slope_) between AD and control was calculated for each module.

**<u>Extended Data 10.</u> Correlation of Proteins Commonly Measured in Plasma by MS-TMT and Olink.** *n*=36 unless otherwise indicated. Measurements were from TMT-MS on plasma depleted of highly abundant plasma proteins. Olink NPX values are shown on the x-axis, and TMT-MS log_2_ relative values are shown on the y-axis. Measurements that were below LOD in Olink are not outlined. Correlations include those proteins that were matched by gene symbol only. Correlations were performed using Pearson test.

**<u>Extended Data 11.</u> Correlation of Proteins Commonly Measured in Plasma by MS-TMT and SomaScan.** *n*=35 unless otherwise indicated. Measurements were from TMT-MS on plasma depleted of highly abundant plasma proteins. TMT-MS log_2_ relative values are shown on the x-axis, and SomaScan log_2_(RFU) values are shown on the y-axis. Measurements that were below LOD in SomaScan are not outlined. Correlations were performed for all SOMAmers including multiple SOMAmers targeting the same protein, and include those proteins that were matched by gene symbol only. Correlations were performed using Pearson test.

**<u>Extended Data 12.</u> Correlation of Proteins Commonly Measured in Plasma by Olink and SomaScan.** *n*=35 unless otherwise indicated. Olink NPX values are shown on the x-axis, and SomaScan log_2_(RFU) values are shown on the y-axis. Measurements that were below LOD in either platform are not outlined. Correlations were performed for all SOMAmers including multiple SOMAmers targeting the same protein, and include those proteins that were matched by gene symbol only. Correlations were performed using Pearson test.

**<u>Extended Data 13.</u> GO Analysis on AD Plasma Network Modules.** Gene ontology (GO) analysis was performed to gain insight into the biological meaning of each AD plasma protein network module. Enrichment for a given ontology is shown by *z* score.

**<u>Extended Data 14.</u> AD Plasma Network Module Protein Graphs.** The size of each circle indicates the relative eigenprotein correlation value (kME) in each network module. Those proteins with the largest kME are considered “hub” proteins within the module, and explain the largest variance in module expression. Proteins outlined in gold are from the SomaScan platform. Proteins outlined in green are from the Olink platform. Proteins outlined in purple are from the TMT-MS platform. Only the top 100 proteins by kME for each module are shown.

**<u>Extended Data 15.</u> AD Plasma Network Module Eigenprotein Levels and Correlations.** *n*=18 control, 17 AD. Differences between case groups were assessed by *t* test or one-way ANOVA. Correlations were performed by bicor or Pearson test (cor). Significance at p<0.05 is highlighted in red. Boxplots represent the median, 25th, and 75th percentiles, and box hinges represent the interquartile range of the two middle quartiles within a group. Datapoints up to 1.5 times the interquartile range from box hinge define the extent of whiskers (error bars).

**<u>Extended Data 16.</u> AD Plasma Network Module Synthetic Eigenprotein Levels and Correlations in Brain**. *n*=18 control, 17 AD. Differences between case groups were assessed by *t* test or one-way ANOVA. Correlations were performed by bicor or Pearson test (cor). Significance at p<0.05 is highlighted in red. Boxplots represent the median, 25th, and 75th percentiles, and box hinges represent the interquartile range of the two middle quartiles within a group. Datapoints up to 1.5 times the interquartile range from box hinge define the extent of whiskers (error bars).

**<u>Extended Data 17.</u> AD Plasma Network Module Synthetic Eigenprotein Levels and Correlations in CSF**. *n*=18 control, 17 AD. Differences between case groups were assessed by *t* test or one-way ANOVA. Correlations were performed by bicor or Pearson test (cor). Significance at p<0.05 is highlighted in red. Boxplots represent the median, 25th, and 75th percentiles, and box hinges represent the interquartile range of the two middle quartiles within a group. Datapoints up to 1.5 times the interquartile range from box hinge define the extent of whiskers (error bars).

**<u>Extended Data 18.</u> Within-Subject Plasma Network Module Eigenprotein Levels in CSF and Plasma**. *n*=18 control, 17 AD. Relative plasma protein network module eigenprotein levels and their synthetic eigenprotein levels in CSF were compared within subject across fluids in control and AD cases. The difference in average slope (*Z*_slope_) between AD and control was calculated for each module.

**<u>Extended Data 19.</u> Brain TMT-MS AD Network Module Synthetic Eigenprotein Levels and Correlations in CSF**. *n*=18 control, 17 AD. Differences between case groups were assessed by *t* test or one-way ANOVA. Correlations were performed by bicor or Pearson test (cor). Significance at p<0.05 is highlighted in red. Boxplots represent the median, 25th, and 75th percentiles, and box hinges represent the interquartile range of the two middle quartiles within a group. Datapoints up to 1.5 times the interquartile range from box hinge define the extent of whiskers (error bars).

**<u>Extended Data 20.</u> Brain TMT-MS AD Network Module Synthetic Eigenprotein Levels and Correlations in Plasma**. *n*=18 control, 17 AD. Differences between case groups were assessed by *t* test or one-way ANOVA. Correlations were performed by bicor or Pearson test (cor). Significance at p<0.05 is highlighted in red. Boxplots represent the median, 25th, and 75th percentiles, and box hinges represent the interquartile range of the two middle quartiles within a group. Datapoints up to 1.5 times the interquartile range from box hinge define the extent of whiskers (error bars).

