## Supplementary material for "Multi-Platform Proteomic Analysis of Alzheimer’s Disease Cerebrospinal Fluid and Plasma Reveals Network Biomarkers Associated with Proteostasis and the Matrisome": Dammer et al 2022 Extended Data

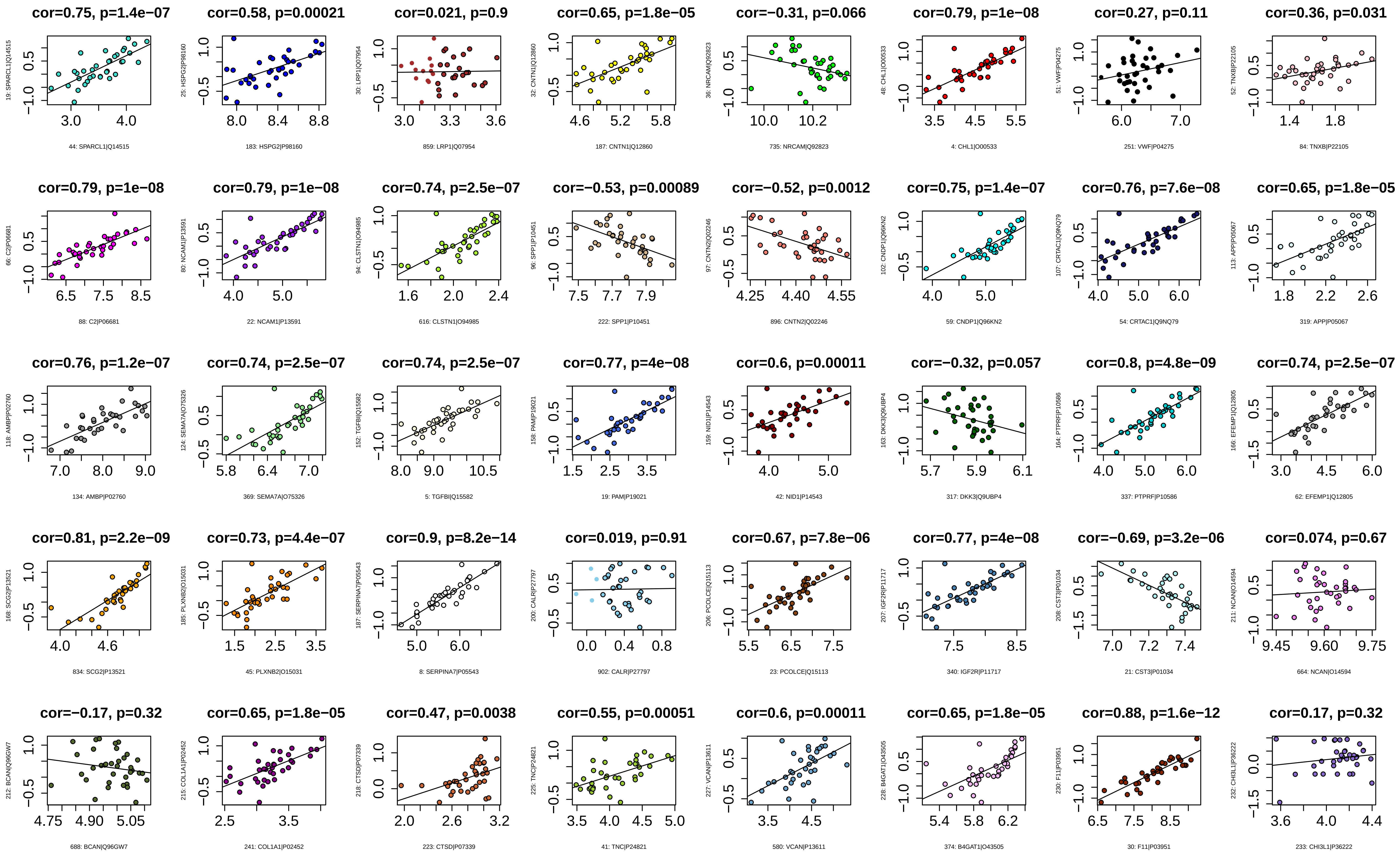
cor=0.75, p=1.4e−07
19: SPARCL1|Q14515
44: SPARCL1|Q14515
cor=0.58, p=0.00021
25: HSPG2|P98160
183: HSPG2|P98160
cor=0.021, p=0.9
30: LRP1|Q07954
859: LRP1|Q07954
cor=0.65, p=1.8e−05
32: CNTN1|Q12860
187: CNTN1|Q12860
cor=−0.31, p=0.066
36: NRCAM|Q92823
735: NRCAM|Q92823
cor=0.79, p=1e−08
48: CHL1|O00533
4: CHL1|O00533
cor=0.27, p=0.11
51: VWF|P04275
251: VWF|P04275
cor=0.36, p=0.031
52: TNXB|P22105
84: TNXB|P22105
cor=0.79, p=1e−08
66: C2|P06681
88: C2|P06681
cor=0.79, p=1e−08
80: NCAM1|P13591
22: NCAM1|P13591
cor=0.74, p=2.5e−07
94: CLSTN1|O94985
616: CLSTN1|O94985
cor=−0.53, p=0.00089
96: SPP1|P10451
222: SPP1|P10451
cor=−0.52, p=0.0012
97: CNTN2|Q02246
896: CNTN2|Q02246
cor=0.75, p=1.4e−07
102: CNDP1|Q96KN2
59: CNDP1|Q96KN2
cor=0.76, p=7.6e−08
107: CRTAC1|Q9NQ79
54: CRTAC1|Q9NQ79
cor=0.65, p=1.8e−05
113: APP|P05067
319: APP|P05067
cor=0.76, p=1.2e−07
118: AMBP|P02760
134: AMBP|P02760
cor=0.74, p=2.5e−07
124: SEMA7A|O75326
369: SEMA7A|O75326
cor=0.74, p=2.5e−07
152: TGFBI|Q15582
5: TGFBI|Q15582
cor=0.77, p=4e−08
158: PAM|P19021
19: PAM|P19021
cor=0.6, p=0.00011
159: NID1|P14543
42: NID1|P14543
cor=−0.32, p=0.057
163: DKK3|Q9UBP4
317: DKK3|Q9UBP4
cor=0.8, p=4.8e−09
164: PTPRF|P10586
337: PTPRF|P10586
cor=0.74, p=2.5e−07
166: EFEMP1|Q12805
62: EFEMP1|Q12805
cor=0.81, p=2.2e−09
168: SCG2|P13521
834: SCG2|P13521
cor=0.73, p=4.4e−07
185: PLXNB2|O15031
45: PLXNB2|O15031
cor=0.9, p=8.2e−14
187: SERPINA7|P05543
8: SERPINA7|P05543
cor=0.019, p=0.91
200: CALR|P27797
902: CALR|P27797
cor=0.67, p=7.8e−06
206: PCOLCE|Q15113
23: PCOLCE|Q15113
cor=0.77, p=4e−08
207: IGF2R|P11717
340: IGF2R|P11717
cor=−0.69, p=3.2e−06
208: CST3|P01034
21: CST3|P01034
cor=0.074, p=0.67
211: NCAN|O14594
664: NCAN|O14594
cor=−0.17, p=0.32
212: BCAN|Q96GW7
688: BCAN|Q96GW7
cor=0.65, p=1.8e−05
215: COL1A1|P02452
241: COL1A1|P02452
cor=0.47, p=0.0038
218: CTSD|P07339
223: CTSD|P07339
cor=0.55, p=0.00051
225: TNC|P24821
41: TNC|P24821
cor=0.6, p=0.00011
227: VCAN|P13611
580: VCAN|P13611
cor=0.65, p=1.8e−05
228: B4GAT1|O43505
374: B4GAT1|O43505
cor=0.88, p=1.6e−12
230: F11|P03951
30: F11|P03951
cor=0.17, p=0.32
232: CHI3L1|P36222
233: CHI3L1|P36222

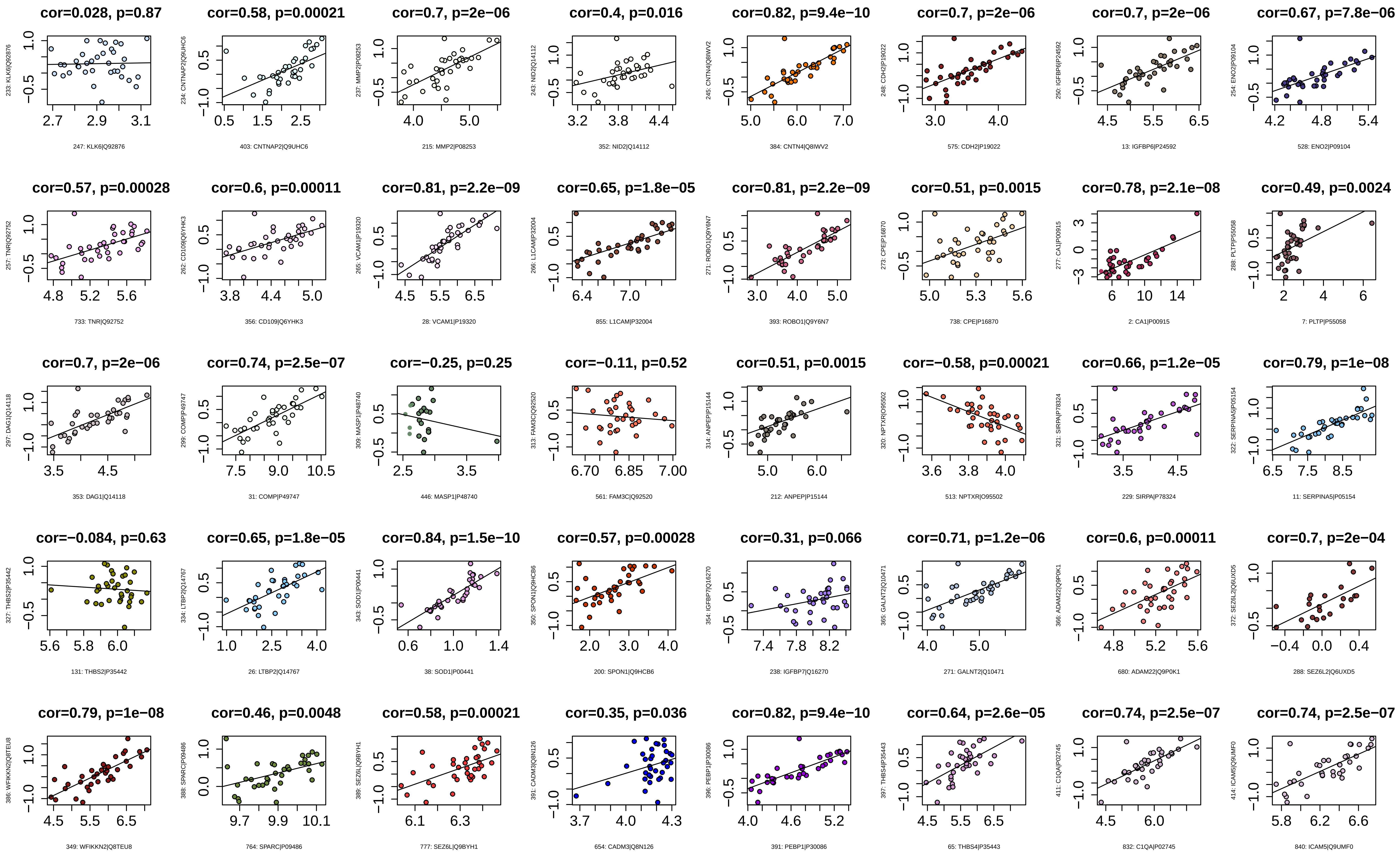
cor=0.028, p=0.87
233: KLK6|Q92876
247: KLK6|Q92876
cor=0.58, p=0.00021
234: CNTNAP2|Q9UHC6
403: CNTNAP2|Q9UHC6
cor=0.7, p=2e−06
237: MMP2|P08253
215: MMP2|P08253
cor=0.4, p=0.016
243: NID2|Q14112
352: NID2|Q14112
cor=0.82, p=9.4e−10
245: CNTN4|Q8IWV2
384: CNTN4|Q8IWV2
cor=0.7, p=2e−06
248: CDH2|P19022
575: CDH2|P19022
cor=0.7, p=2e−06
250: IGFBP6|P24592
13: IGFBP6|P24592
cor=0.67, p=7.8e−06
254: ENO2|P09104
528: ENO2|P09104
cor=0.57, p=0.00028
257: TNR|Q92752
733: TNR|Q92752
cor=0.6, p=0.00011
262: CD109|Q6YHK3
356: CD109|Q6YHK3
cor=0.81, p=2.2e−09
265: VCAM1|P19320
28: VCAM1|P19320
cor=0.65, p=1.8e−05
266: L1CAM|P32004
855: L1CAM|P32004
cor=0.81, p=2.2e−09
271: ROBO1|Q9Y6N7
393: ROBO1|Q9Y6N7
cor=0.51, p=0.0015
273: CPE|P16870
738: CPE|P16870
cor=0.78, p=2.1e−08
277: CA1|P00915
2: CA1|P00915
cor=0.49, p=0.0024
288: PLTP|P55058
7: PLTP|P55058
cor=0.7, p=2e−06
297: DAG1|Q14118
353: DAG1|Q14118
cor=0.74, p=2.5e−07
299: COMP|P49747
31: COMP|P49747
cor=−0.25, p=0.25
309: MASP1|P48740
446: MASP1|P48740
cor=−0.11, p=0.52
313: FAM3C|Q92520
561: FAM3C|Q92520
cor=0.51, p=0.0015
314: ANPEP|P15144
212: ANPEP|P15144
cor=−0.58, p=0.00021
320: NPTXR|O95502
513: NPTXR|O95502
cor=0.66, p=1.2e−05
321: SIRPA|P78324
229: SIRPA|P78324
cor=0.79, p=1e−08
322: SERPINA5|P05154
11: SERPINA5|P05154
cor=−0.084, p=0.63
327: THBS2|P35442
131: THBS2|P35442
cor=0.65, p=1.8e−05
334: LTBP2|Q14767
26: LTBP2|Q14767
cor=0.84, p=1.5e−10
343: SOD1|P00441
38: SOD1|P00441
cor=0.57, p=0.00028
350: SPON1|Q9HCB6
200: SPON1|Q9HCB6
cor=0.31, p=0.066
354: IGFBP7|Q16270
238: IGFBP7|Q16270
cor=0.71, p=1.2e−06
365: GALNT2|Q10471
271: GALNT2|Q10471
cor=0.6, p=0.00011
366: ADAM22|Q9P0K1
680: ADAM22|Q9P0K1
cor=0.7, p=2e−04
372: SEZ6L2|Q6UXD5
288: SEZ6L2|Q6UXD5
cor=0.79, p=1e−08
386: WFIKKN2|Q8TEU8
349: WFIKKN2|Q8TEU8
cor=0.46, p=0.0048
388: SPARC|P09486
764: SPARC|P09486
cor=0.58, p=0.00021
389: SEZ6L|Q9BYH1
777: SEZ6L|Q9BYH1
cor=0.35, p=0.036
391: CADM3|Q8N126
654: CADM3|Q8N126
cor=0.82, p=9.4e−10
396: PEBP1|P30086
391: PEBP1|P30086
cor=0.64, p=2.6e−05
397: THBS4|P35443
65: THBS4|P35443
cor=0.74, p=2.5e−07
411: C1QA|P02745
832: C1QA|P02745
cor=0.74, p=2.5e−07
414: ICAM5|Q9UMF0
840: ICAM5|Q9UMF0

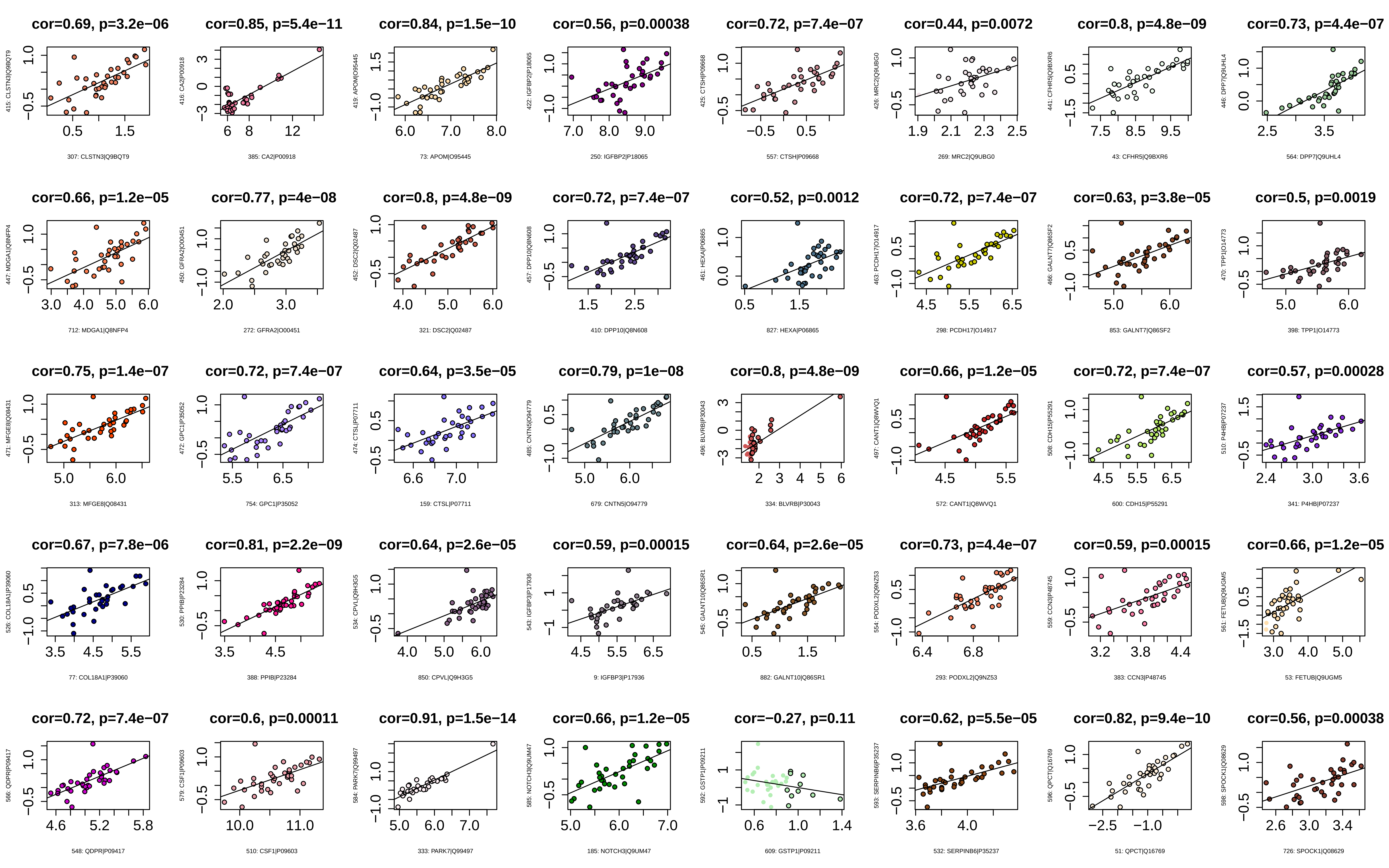
cor=0.69, p=3.2e−06
307: CLSTN3|Q9BQT9
cor=0.85, p=5.4e−11
385: CA2|P00918
cor=0.84, p=1.5e−10
73: APOM|O95445
cor=0.56, p=0.00038
250: IGFBP2|P18065
cor=0.72, p=7.4e−07
557: CTSH|P09668
cor=0.44, p=0.0072
269: MRC2|Q9UBG0
cor=0.8, p=4.8e−09
43: CFHR5|Q9BXR6
cor=0.73, p=4.4e−07
564: DPP7|Q9UHL4
cor=0.66, p=1.2e−05
712: MDGA1|Q8NFP4
cor=0.77, p=4e−08
272: GFRA2|O00451
cor=0.8, p=4.8e−09
321: DSC2|Q02487
cor=0.72, p=7.4e−07
410: DPP10|Q8N608
cor=0.52, p=0.0012
827: HEXA|P06865
cor=0.72, p=7.4e−07
298: PCDH17|O14917
cor=0.63, p=3.8e−05
853: GALNT7|Q86SF2
cor=0.5, p=0.0019
398: TPP1|O14773
cor=0.75, p=1.4e−07
313: MFGE8|Q08431
cor=0.72, p=7.4e−07
754: GPC1|P35052
cor=0.64, p=3.5e−05
159: CTSL|P07711
cor=0.79, p=1e−08
679: CNTN5|O94779
cor=0.8, p=4.8e−09
334: BLVRB|P30043
cor=0.66, p=1.2e−05
572: CANT1|Q8WVQ1
cor=0.72, p=7.4e−07
600: CDH15|P55291
cor=0.57, p=0.00028
341: P4HB|P07237
cor=0.67, p=7.8e−06
77: COL18A1|P39060
cor=0.81, p=2.2e−09
388: PPIB|P23284
cor=0.64, p=2.6e−05
850: CPVL|Q9H3G5
cor=0.59, p=0.00015
9: IGFBP3|P17936
cor=0.64, p=2.6e−05
882: GALNT10|Q86SR1
cor=0.73, p=4.4e−07
293: PODXL2|Q9NZ53
cor=0.59, p=0.00015
383: CCN3|P48745
cor=0.66, p=1.2e−05
53: FETUB|Q9UGM5
cor=0.72, p=7.4e−07
548: QDPR|P09417
cor=0.6, p=0.00011
510: CSF1|P09603
cor=0.91, p=1.5e−14
333: PARK7|Q99497
cor=0.66, p=1.2e−05
185: NOTCH3|Q9UM47
cor=−0.27, p=0.11
609: GSTP1|P09211
cor=0.62, p=5.5e−05
532: SERPINB6|P35237
cor=0.82, p=9.4e−10
51: QPCT|Q16769
cor=0.56, p=0.00038
726: SPOCK1|Q08629

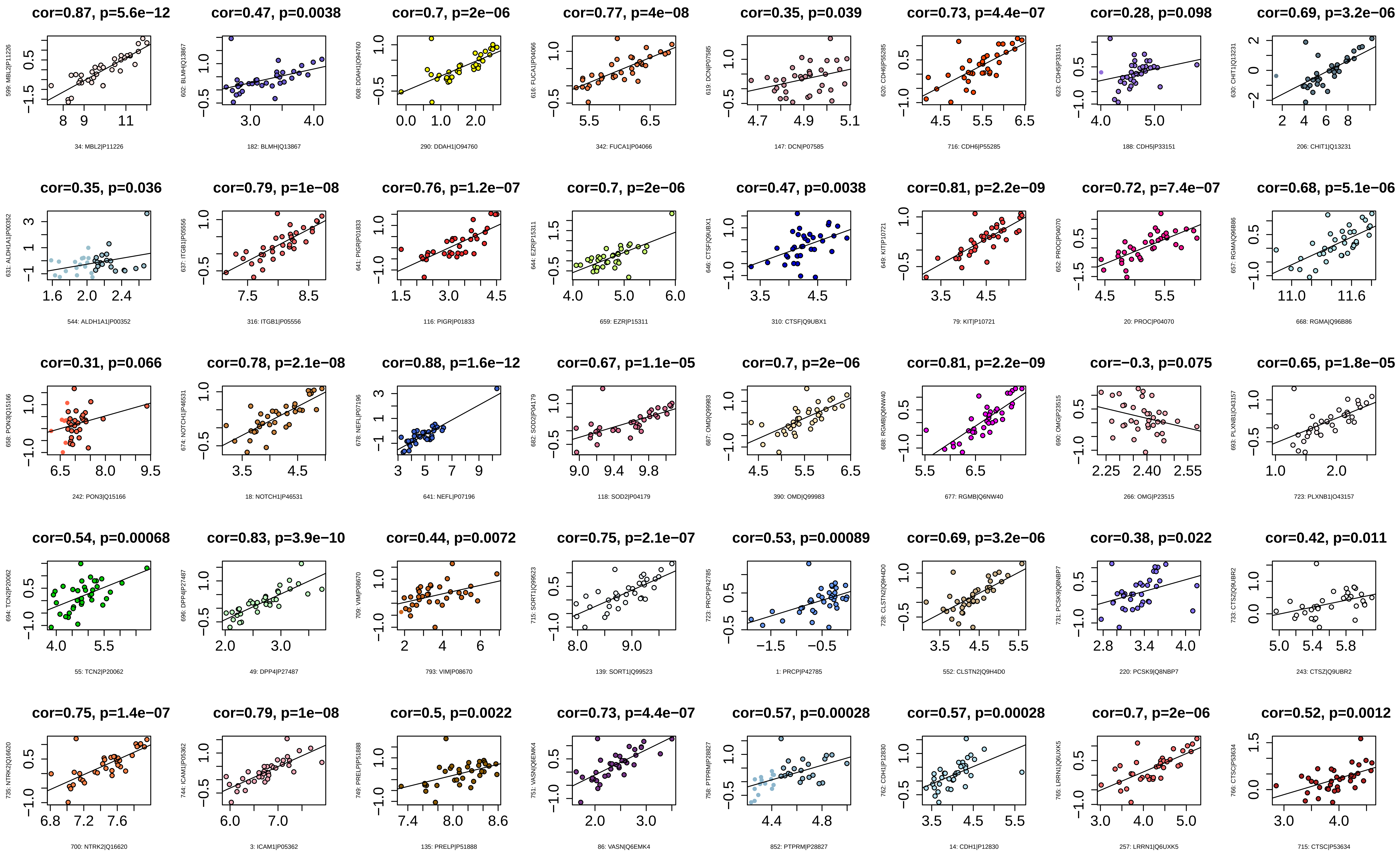
cor=0.87, p=5.6e−12
34: MBL2|P11226
cor=0.47, p=0.0038
182: BLMH|Q13867
cor=0.7, p=2e−06
290: DDAH1|O94760
cor=0.77, p=4e−08
342: FUCA1|P04066
cor=0.35, p=0.039
147: DCN|P07585
cor=0.73, p=4.4e−07
716: CDH6|P55285
cor=0.28, p=0.098
188: CDH5|P33151
cor=0.69, p=3.2e−06
206: CHIT1|Q13231
cor=0.35, p=0.036
544: ALDH1A1|P00352
cor=0.79, p=1e−08
316: ITGB1|P05556
cor=0.76, p=1.2e−07
116: PIGR|P01833
cor=0.7, p=2e−06
659: EZR|P15311
cor=0.47, p=0.0038
310: CTSF|Q9UBX1
cor=0.81, p=2.2e−09
79: KIT|P10721
cor=0.72, p=7.4e−07
20: PROC|P04070
cor=0.68, p=5.1e−06
668: RGMA|Q96B86
cor=0.31, p=0.066
242: PON3|Q15166
cor=0.78, p=2.1e−08
18: NOTCH1|P46531
cor=0.88, p=1.6e−12
641: NEFL|P07196
cor=0.67, p=1.1e−05
118: SOD2|P04179
cor=0.7, p=2e−06
390: OMD|Q99983
cor=0.81, p=2.2e−09
677: RGMB|Q6NW40
cor=−0.3, p=0.075
266: OMG|P23515
cor=0.65, p=1.8e−05
723: PLXNB1|O43157
cor=0.54, p=0.00068
55: TCN2|P20062
cor=0.83, p=3.9e−10
49: DPP4|P27487
cor=0.44, p=0.0072
793: VIM|P08670
cor=0.75, p=2.1e−07
139: SORT1|Q99523
cor=0.53, p=0.00089
1: PRCP|P42785
cor=0.69, p=3.2e−06
552: CLSTN2|Q9H4D0
cor=0.38, p=0.022
220: PCSK9|Q8NBP7
cor=0.42, p=0.011
243: CTSZ|Q9UBR2
cor=0.75, p=1.4e−07
700: NTRK2|Q16620
cor=0.79, p=1e−08
3: ICAM1|P05362
cor=0.5, p=0.0022
135: PRELP|P51888
cor=0.73, p=4.4e−07
86: VASN|Q6EMK4
cor=0.57, p=0.00028
852: PTPRM|P28827
cor=0.57, p=0.00028
14: CDH1|P12830
cor=0.7, p=2e−06
257: LRRN1|Q6UXK5
cor=0.52, p=0.0012
715: CTSC|P53634

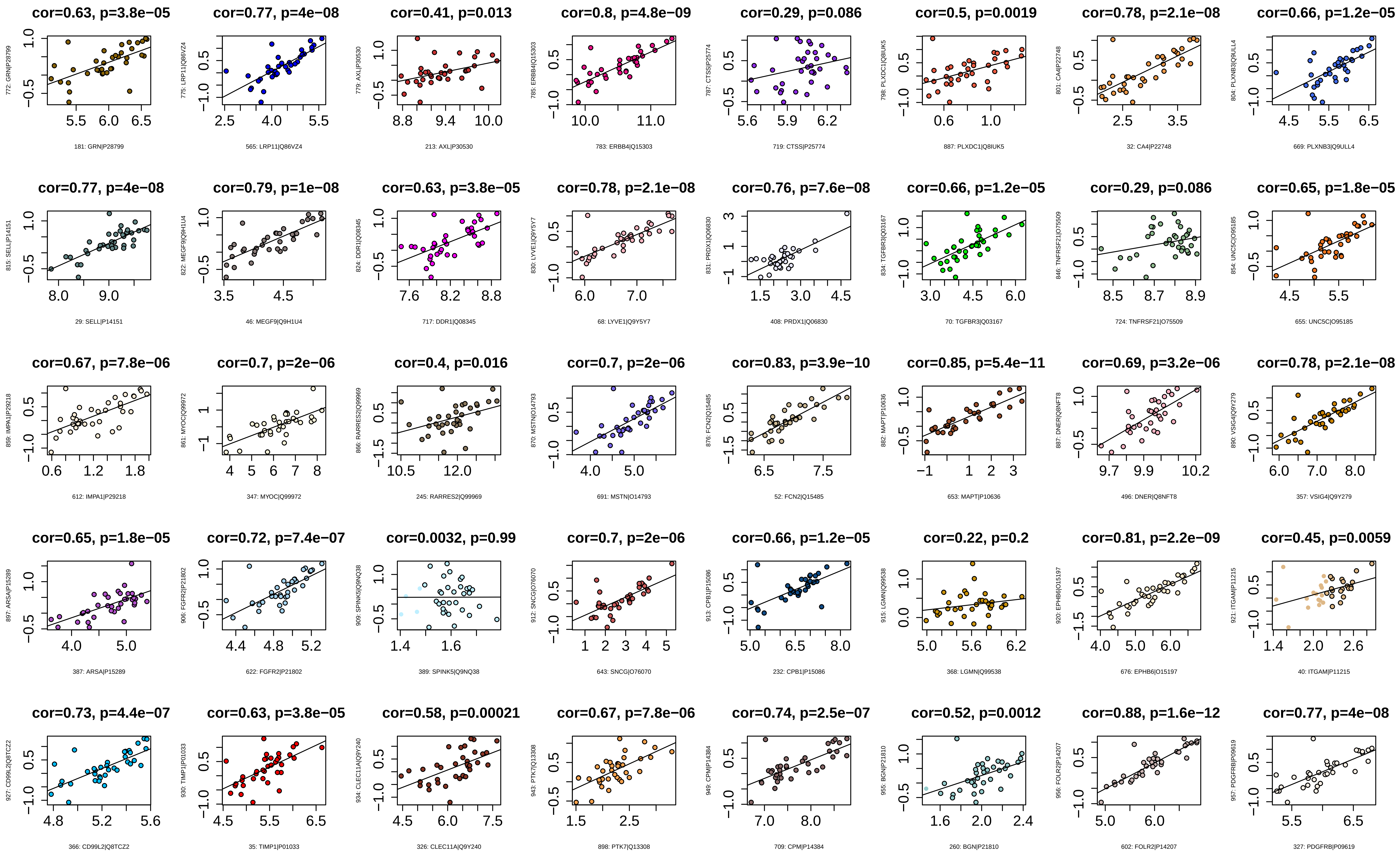
cor=0.63, p=3.8e−05
772: GRN|P28799
181: GRN|P28799
cor=0.77, p=4e−08
775: LRP11|Q86VZ4
565: LRP11|Q86VZ4
cor=0.41, p=0.013
779: AXL|P30530
213: AXL|P30530
cor=0.8, p=4.8e−09
785: ERBB4|Q15303
783: ERBB4|Q15303
cor=0.29, p=0.086
787: CTSS|P25774
719: CTSS|P25774
cor=0.5, p=0.0019
798: PLXDC1|Q8IUK5
887: PLXDC1|Q8IUK5
cor=0.78, p=2.1e−08
801: CA4|P22748
32: CA4|P22748
cor=0.66, p=1.2e−05
804: PLXNB3|Q9ULL4
669: PLXNB3|Q9ULL4
cor=0.77, p=4e−08
815: SELL|P14151
29: SELL|P14151
cor=0.79, p=1e−08
822: MEGF9|Q9H1U4
46: MEGF9|Q9H1U4
cor=0.63, p=3.8e−05
824: DDR1|Q08345
717: DDR1|Q08345
cor=0.78, p=2.1e−08
830: LYVE1|Q9Y5Y7
68: LYVE1|Q9Y5Y7
cor=0.76, p=7.6e−08
831: PRDX1|Q06830
408: PRDX1|Q06830
cor=0.66, p=1.2e−05
834: TGFBR3|Q03167
70: TGFBR3|Q03167
cor=0.29, p=0.086
846: TNFRSF21|O75509
724: TNFRSF21|O75509
cor=0.65, p=1.8e−05
854: UNC5C|O95185
655: UNC5C|O95185
cor=0.67, p=7.8e−06
859: IMPA1|P29218
612: IMPA1|P29218
cor=0.7, p=2e−06
861: MYOC|Q99972
347: MYOC|Q99972
cor=0.4, p=0.016
866: RARRES2|Q99969
245: RARRES2|Q99969
cor=0.7, p=2e−06
870: MSTN|O14793
691: MSTN|O14793
cor=0.83, p=3.9e−10
876: FCN2|Q15485
52: FCN2|Q15485
cor=0.85, p=5.4e−11
882: MAPT|P10636
653: MAPT|P10636
cor=0.69, p=3.2e−06
887: DNER|Q8NFT8
496: DNER|Q8NFT8
cor=0.78, p=2.1e−08
890: VSIG4|Q9Y279
357: VSIG4|Q9Y279
cor=0.65, p=1.8e−05
897: ARSA|P15289
387: ARSA|P15289
cor=0.72, p=7.4e−07
906: FGFR2|P21802
622: FGFR2|P21802
cor=0.0032, p=0.99
909: SPINK5|Q9NQ38
389: SPINK5|Q9NQ38
cor=0.7, p=2e−06
912: SNCG|O76070
643: SNCG|O76070
cor=0.66, p=1.2e−05
913: CPB1|P15086
232: CPB1|P15086
cor=0.22, p=0.2
915: LGMN|Q99538
368: LGMN|Q99538
cor=0.81, p=2.2e−09
920: EPHB6|O15197
676: EPHB6|O15197
cor=0.45, p=0.0059
921: ITGAM|P11215
40: ITGAM|P11215
cor=0.73, p=4.4e−07
927: CD99L2|Q8TCZ2
366: CD99L2|Q8TCZ2
cor=0.63, p=3.8e−05
930: TIMP1|P01033
35: TIMP1|P01033
cor=0.58, p=0.00021
934: CLEC11A|Q9Y240
326: CLEC11A|Q9Y240
cor=0.67, p=7.8e−06
943: PTK7|Q13308
898: PTK7|Q13308
cor=0.74, p=2.5e−07
949: CPM|P14384
709: CPM|P14384
cor=0.52, p=0.0012
955: BGN|P21810
260: BGN|P21810
cor=0.88, p=1.6e−12
956: FOLR2|P14207
602: FOLR2|P14207
cor=0.77, p=4e−08
957: PDGFRB|P09619
327: PDGFRB|P09619

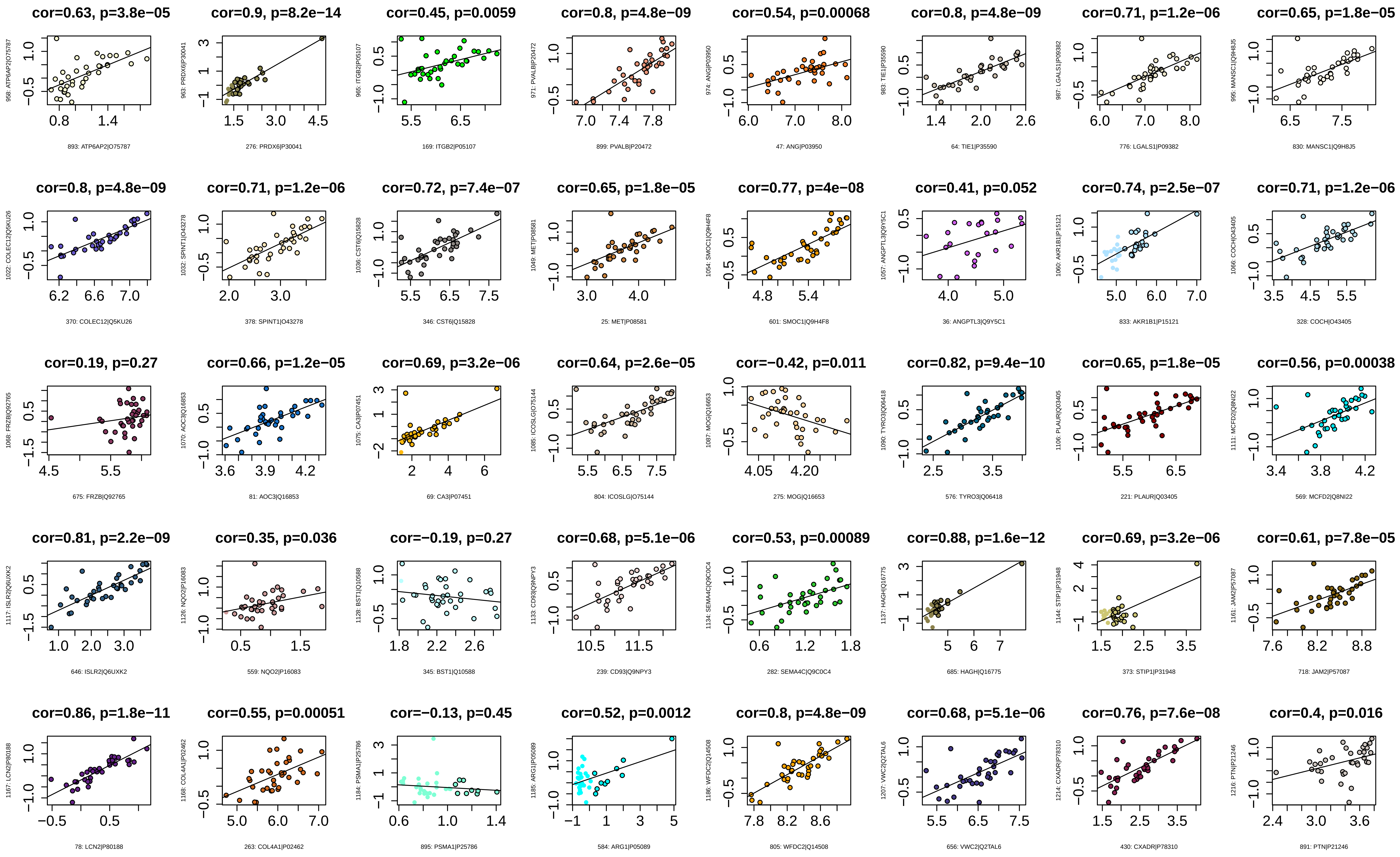
cor=0.63, p=3.8e−05
893: ATP6AP2|O75787
cor=0.9, p=8.2e−14
276: PRDX6|P30041
cor=0.45, p=0.0059
169: ITGB2|P05107
cor=0.8, p=4.8e−09
899: PVALB|P20472
cor=0.54, p=0.00068
47: ANG|P03950
cor=0.8, p=4.8e−09
64: TIE1|P35590
cor=0.71, p=1.2e−06
776: LGALS1|P09382
cor=0.65, p=1.8e−05
830: MANSC1|Q9H8J5
cor=0.8, p=4.8e−09
370: COLEC12|Q5KU26
cor=0.71, p=1.2e−06
378: SPINT1|O43278
cor=0.72, p=7.4e−07
346: CST6|Q15828
cor=0.65, p=1.8e−05
25: MET|P08581
cor=0.77, p=4e−08
601: SMOC1|Q9H4F8
cor=0.41, p=0.052
36: ANGPTL3|Q9Y5C1
cor=0.74, p=2.5e−07
833: AKR1B1|P15121
cor=0.71, p=1.2e−06
328: COCH|O43405
cor=0.19, p=0.27
675: FRZB|Q92765
cor=0.66, p=1.2e−05
81: AOC3|Q16853
cor=0.69, p=3.2e−06
69: CA3|P07451
cor=0.64, p=2.6e−05
804: ICOSLG|O75144
cor=−0.42, p=0.011
275: MOG|Q16653
cor=0.82, p=9.4e−10
576: TYRO3|Q06418
cor=0.65, p=1.8e−05
221: PLAUR|Q03405
cor=0.56, p=0.00038
569: MCFD2|Q8NI22
cor=0.81, p=2.2e−09
646: ISLR2|Q6UXK2
cor=0.35, p=0.036
559: NQO2|P16083
cor=−0.19, p=0.27
345: BST1|Q10588
cor=0.68, p=5.1e−06
239: CD93|Q9NPY3
cor=0.53, p=0.00089
282: SEMA4C|Q9C0C4
cor=0.88, p=1.6e−12
685: HAGH|Q16775
cor=0.69, p=3.2e−06
373: STIP1|P31948
cor=0.61, p=7.8e−05
718: JAM2|P57087
cor=0.86, p=1.8e−11
78: LCN2|P80188
cor=0.55, p=0.00051
263: COL4A1|P02462
cor=−0.13, p=0.45
895: PSMA1|P25786
cor=0.52, p=0.0012
584: ARG1|P05089
cor=0.8, p=4.8e−09
805: WFDC2|Q14508
cor=0.68, p=5.1e−06
656: VWC2|Q2TAL6
cor=0.76, p=7.6e−08
430: CXADR|P78310
cor=0.4, p=0.016
891: PTN|P21246

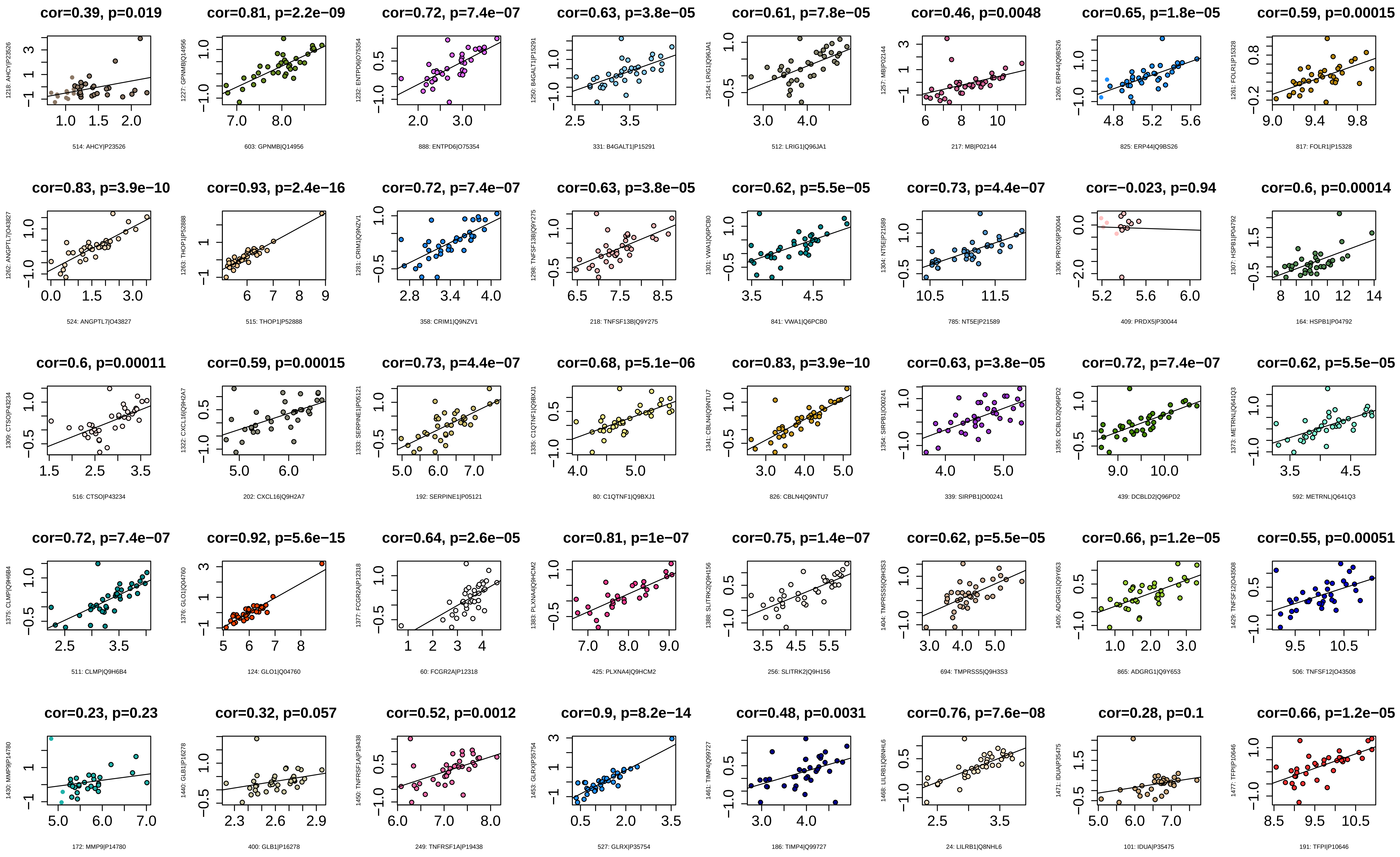
cor=0.39, p=0.019
514: AHCY|P23526
cor=0.81, p=2.2e−09
603: GPNMB|Q14956
cor=0.72, p=7.4e−07
888: ENTPD6|O75354
cor=0.63, p=3.8e−05
331: B4GALT1|P15291
cor=0.61, p=7.8e−05
512: LRIG1|Q96JA1
cor=0.46, p=0.0048
217: MB|P02144
cor=0.65, p=1.8e−05
825: ERP44|Q9BS26
cor=0.59, p=0.00015
817: FOLR1|P15328
cor=0.83, p=3.9e−10
524: ANGPTL7|O43827
cor=0.93, p=2.4e−16
515: THOP1|P52888
cor=0.72, p=7.4e−07
358: CRIM1|Q9NZV1
cor=0.63, p=3.8e−05
218: TNFSF13B|Q9Y275
cor=0.62, p=5.5e−05
841: VWA1|Q6PCB0
cor=0.73, p=4.4e−07
785: NT5E|P21589
cor=−0.023, p=0.94
409: PRDX5|P30044
cor=0.6, p=0.00014
164: HSPB1|P04792
cor=0.6, p=0.00011
516: CTSO|P43234
cor=0.59, p=0.00015
202: CXCL16|Q9H2A7
cor=0.73, p=4.4e−07
192: SERPINE1|P05121
cor=0.68, p=5.1e−06
80: C1QTNF1|Q9BXJ1
cor=0.83, p=3.9e−10
826: CBLN4|Q9NTU7
cor=0.63, p=3.8e−05
339: SIRPB1|O00241
cor=0.72, p=7.4e−07
439: DCBLD2|Q96PD2
cor=0.62, p=5.5e−05
592: METRNL|Q641Q3
cor=0.72, p=7.4e−07
511: CLMP|Q9H6B4
cor=0.92, p=5.6e−15
124: GLO1|Q04760
cor=0.64, p=2.6e−05
60: FCGR2A|P12318
cor=0.81, p=1e−07
425: PLXNA4|Q9HCM2
cor=0.75, p=1.4e−07
256: SLITRK2|Q9H156
cor=0.62, p=5.5e−05
694: TMPRSS5|Q9H3S3
cor=0.66, p=1.2e−05
865: ADGRG1|Q9Y653
cor=0.55, p=0.00051
506: TNFSF12|O43508
cor=0.23, p=0.23
172: MMP9|P14780
cor=0.32, p=0.057
400: GLB1|P16278
cor=0.52, p=0.0012
249: TNFRSF1A|P19438
cor=0.9, p=8.2e−14
527: GLRX|P35754
cor=0.48, p=0.0031
186: TIMP4|Q99727
cor=0.76, p=7.6e−08
24: LILRB1|Q8NHL6
cor=0.28, p=0.1
101: IDUA|P35475
cor=0.66, p=1.2e−05
191: TFPI|P10646

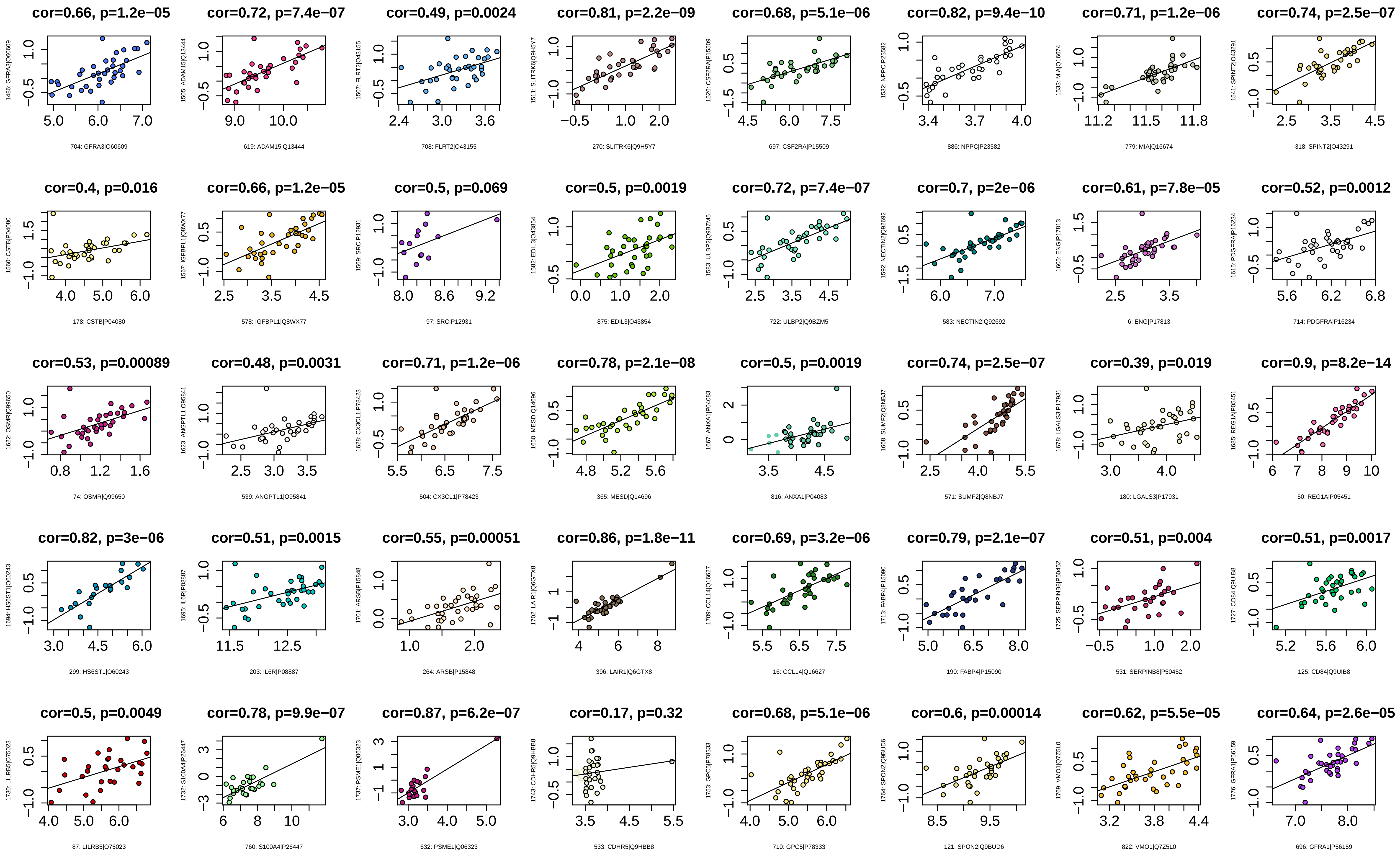
cor=0.66, p=1.2e−05
1486: GFRA3|O60609
704: GFRA3|O60609
cor=0.72, p=7.4e−07
1505: ADAM15|Q13444
619: ADAM15|Q13444
cor=0.49, p=0.0024
1507: FLRT2|O43155
708: FLRT2|O43155
cor=0.81, p=2.2e−09
1511: SLITRK6|Q9H5Y7
270: SLITRK6|Q9H5Y7
cor=0.68, p=5.1e−06
1526: CSF2RA|P15509
697: CSF2RA|P15509
cor=0.82, p=9.4e−10
1532: NPPC|P23582
886: NPPC|P23582
cor=0.71, p=1.2e−06
1533: MIA|Q16674
779: MIA|Q16674
cor=0.74, p=2.5e−07
1541: SPINT2|O43291
318: SPINT2|O43291
cor=0.4, p=0.016
1560: CSTB|P04080
178: CSTB|P04080
cor=0.66, p=1.2e−05
1567: IGFBPL1|Q8WX77
578: IGFBPL1|Q8WX77
cor=0.5, p=0.069
1569: SRC|P12931
97: SRC|P12931
cor=0.5, p=0.0019
1582: EDIL3|O43854
875: EDIL3|O43854
cor=0.72, p=7.4e−07
1583: ULBP2|Q9BZM5
722: ULBP2|Q9BZM5
cor=0.7, p=2e−06
1592: NECTIN2|Q92692
583: NECTIN2|Q92692
cor=0.61, p=7.8e−05
1605: ENG|P17813
6: ENG|P17813
cor=0.52, p=0.0012
1615: PDGFRA|P16234
714: PDGFRA|P16234
cor=0.53, p=0.00089
1622: OSMR|Q99650
74: OSMR|Q99650
cor=0.48, p=0.0031
1623: ANGPTL1|O95841
539: ANGPTL1|O95841
cor=0.71, p=1.2e−06
1628: CX3CL1|P78423
504: CX3CL1|P78423
cor=0.78, p=2.1e−08
1650: MESD|Q14696
365: MESD|Q14696
cor=0.5, p=0.0019
1667: ANXA1|P04083
816: ANXA1|P04083
cor=0.74, p=2.5e−07
1668: SUMF2|Q8NBJ7
571: SUMF2|Q8NBJ7
cor=0.39, p=0.019
1678: LGALS3|P17931
180: LGALS3|P17931
cor=0.9, p=8.2e−14
1685: REG1A|P05451
50: REG1A|P05451
cor=0.82, p=3e−06
1694: HS6ST1|O60243
299: HS6ST1|O60243
cor=0.51, p=0.0015
1695: IL6R|P08887
203: IL6R|P08887
cor=0.55, p=0.00051
1701: ARSB|P15848
264: ARSB|P15848
cor=0.86, p=1.8e−11
1702: LAIR1|Q6GTX8
396: LAIR1|Q6GTX8
cor=0.69, p=3.2e−06
1709: CCL14|Q16627
16: CCL14|Q16627
cor=0.79, p=2.1e−07
1713: FABP4|P15090
190: FABP4|P15090
cor=0.51, p=0.004
1725: SERPINB8|P50452
531: SERPINB8|P50452
cor=0.51, p=0.0017
1727: CD84|Q9UIB8
125: CD84|Q9UIB8
cor=0.5, p=0.0049
1730: LILRB5|O75023
87: LILRB5|O75023
cor=0.78, p=9.9e−07
1732: S100A4|P26447
760: S100A4|P26447
cor=0.87, p=6.2e−07
1737: PSME1|Q06323
632: PSME1|Q06323
cor=0.17, p=0.32
1743: CDHR5|Q9HBB8
533: CDHR5|Q9HBB8
cor=0.68, p=5.1e−06
1753: GPC5|P78333
710: GPC5|P78333
cor=0.6, p=0.00014
1764: SPON2|Q9BUD6
121: SPON2|Q9BUD6
cor=0.62, p=5.5e−05
1769: VMO1|Q7Z5L0
822: VMO1|Q7Z5L0
cor=0.64, p=2.6e−05
1776: GFRA1|P56159
696: GFRA1|P56159

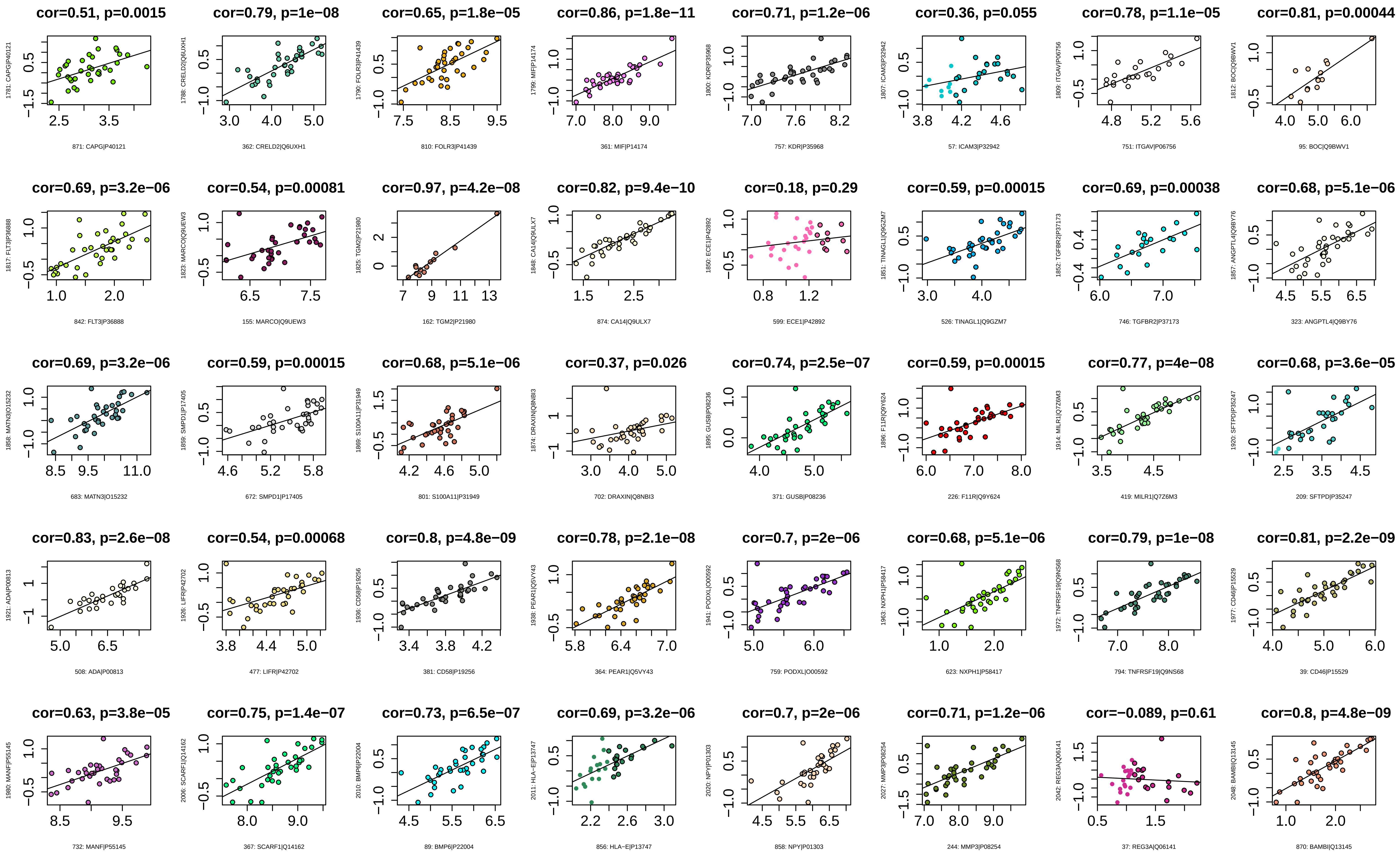
cor=0.51, p=0.0015
1781: CAPG|P40121
871: CAPG|P40121
cor=0.79, p=1e−08
1788: CRELD2|Q6UXH1
362: CRELD2|Q6UXH1
cor=0.65, p=1.8e−05
1790: FOLR3|P41439
810: FOLR3|P41439
cor=0.86, p=1.8e−11
1799: MIF|P14174
361: MIF|P14174
cor=0.71, p=1.2e−06
1800: KDR|P35968
757: KDR|P35968
cor=0.36, p=0.055
1807: ICAM3|P32942
57: ICAM3|P32942
cor=0.78, p=1.1e−05
1809: ITGAV|P06756
751: ITGAV|P06756
cor=0.81, p=0.00044
1812: BOC|Q9BWV1
95: BOC|Q9BWV1
cor=0.69, p=3.2e−06
1817: FLT3|P36888
842: FLT3|P36888
cor=0.54, p=0.00081
1823: MARCO|Q9UEW3
155: MARCO|Q9UEW3
cor=0.97, p=4.2e−08
1825: TGM2|P21980
162: TGM2|P21980
cor=0.82, p=9.4e−10
1848: CA14|Q9ULX7
874: CA14|Q9ULX7
cor=0.18, p=0.29
1850: ECE1|P42892
599: ECE1|P42892
cor=0.59, p=0.00015
1851: TINAGL1|Q9GZM7
526: TINAGL1|Q9GZM7
cor=0.69, p=0.00038
1852: TGFBR2|P37173
746: TGFBR2|P37173
cor=0.68, p=5.1e−06
1857: ANGPTL4|Q9BY76
323: ANGPTL4|Q9BY76
cor=0.69, p=3.2e−06
1858: MATN3|O15232
683: MATN3|O15232
cor=0.59, p=0.00015
1859: SMPD1|P17405
672: SMPD1|P17405
cor=0.68, p=5.1e−06
1869: S100A11|P31949
801: S100A11|P31949
cor=0.37, p=0.026
1874: DRAXIN|Q8NBI3
702: DRAXIN|Q8NBI3
cor=0.74, p=2.5e−07
1895: GUSB|P08236
371: GUSB|P08236
cor=0.59, p=0.00015
1896: F11R|Q9Y624
226: F11R|Q9Y624
cor=0.77, p=4e−08
1914: MILR1|Q7Z6M3
419: MILR1|Q7Z6M3
cor=0.68, p=3.6e−05
1920: SFTPD|P35247
209: SFTPD|P35247
cor=0.83, p=2.6e−08
1921: ADA|P00813
508: ADA|P00813
cor=0.54, p=0.00068
1926: LIFR|P42702
477: LIFR|P42702
cor=0.8, p=4.8e−09
1936: CD58|P19256
381: CD58|P19256
cor=0.78, p=2.1e−08
1938: PEAR1|Q5VY43
364: PEAR1|Q5VY43
cor=0.7, p=2e−06
1941: PODXL|O00592
759: PODXL|O00592
cor=0.68, p=5.1e−06
1963: NXPH1|P58417
623: NXPH1|P58417
cor=0.79, p=1e−08
1972: TNFRSF19|Q9NS68
794: TNFRSF19|Q9NS68
cor=0.81, p=2.2e−09
1977: CD46|P15529
39: CD46|P15529
cor=0.63, p=3.8e−05
1980: MANF|P55145
732: MANF|P55145
cor=0.75, p=1.4e−07
2006: SCARF1|Q14162
367: SCARF1|Q14162
cor=0.73, p=6.5e−07
2010: BMP6|P22004
89: BMP6|P22004
cor=0.69, p=3.2e−06
2011: HLA−E|P13747
856: HLA−E|P13747
cor=0.7, p=2e−06
2020: NPY|P01303
858: NPY|P01303
cor=0.71, p=1.2e−06
2027: MMP3|P08254
244: MMP3|P08254
cor=−0.089, p=0.61
2042: REG3A|Q06141
37: REG3A|Q06141
cor=0.8, p=4.8e−09
2048: BAMBI|Q13145
870: BAMBI|Q13145

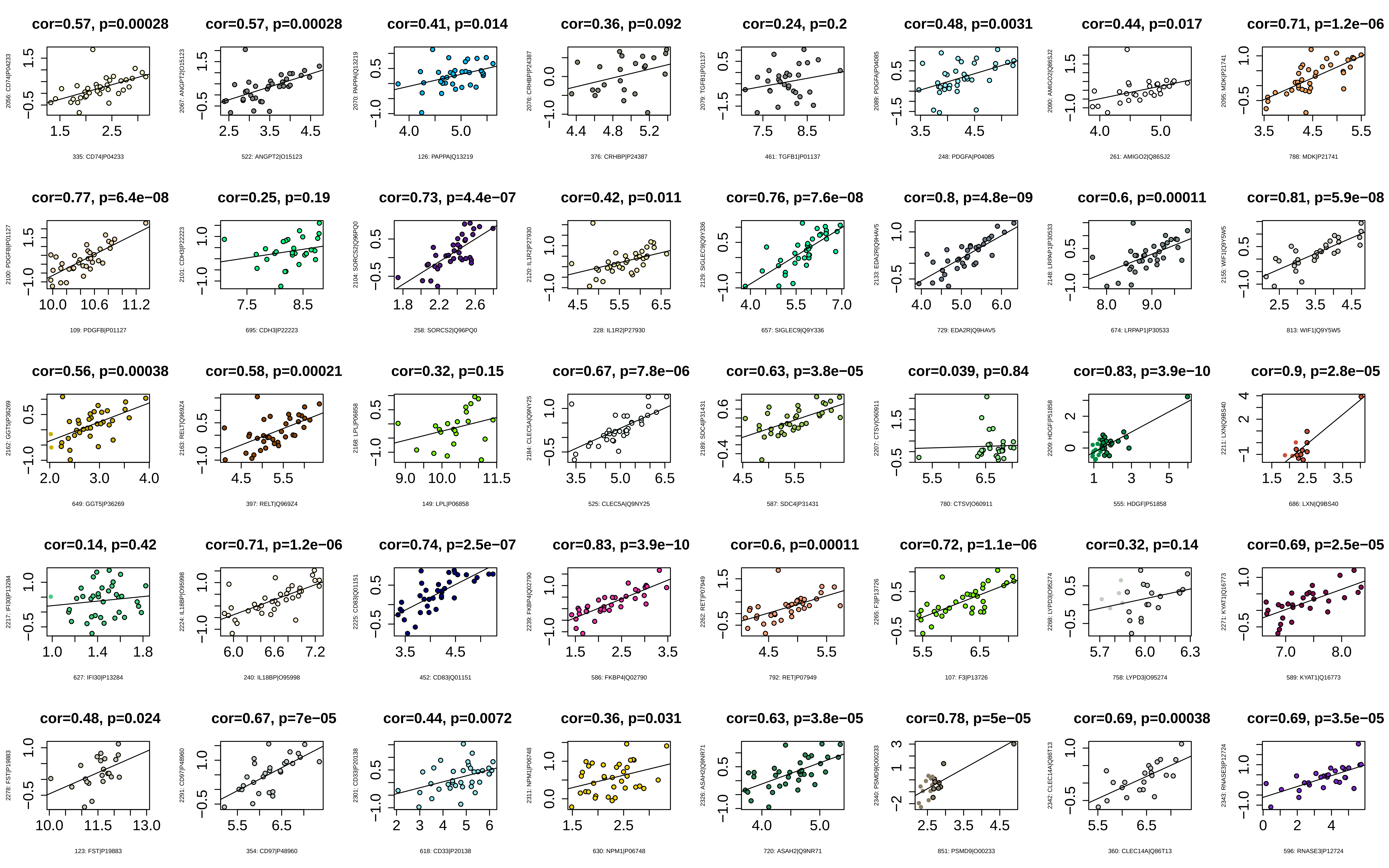
cor=0.57, p=0.00028
335: CD74|P04233
cor=0.57, p=0.00028
522: ANGPT2|O15123
cor=0.41, p=0.014
126: PAPPA|Q13219
cor=0.36, p=0.092
376: CRHBP|P24387
cor=0.24, p=0.2
461: TGFB1|P01137
cor=0.48, p=0.0031
248: PDGFA|P04085
cor=0.44, p=0.017
261: AMIGO2|Q86SJ2
cor=0.71, p=1.2e−06
788: MDK|P21741
cor=0.77, p=6.4e−08
109: PDGFB|P01127
cor=0.25, p=0.19
695: CDH3|P22223
cor=0.73, p=4.4e−07
258: SORCS2|Q96PQ0
cor=0.42, p=0.011
228: IL1R2|P27930
cor=0.76, p=7.6e−08
657: SIGLEC9|Q9Y336
cor=0.8, p=4.8e−09
729: EDA2R|Q9HAV5
cor=0.6, p=0.00011
674: LRPAP1|P30533
cor=0.81, p=5.9e−08
813: WIF1|Q9Y5W5
cor=0.56, p=0.00038
649: GGT5|P36269
cor=0.58, p=0.00021
397: RELT|Q969Z4
cor=0.32, p=0.15
149: LPL|P06858
cor=0.67, p=7.8e−06
525: CLEC5A|Q9NY25
cor=0.63, p=3.8e−05
587: SDC4|P31431
cor=0.039, p=0.84
780: CTSV|O60911
cor=0.83, p=3.9e−10
555: HDGF|P51858
cor=0.9, p=2.8e−05
686: LXN|Q9BS40
cor=0.14, p=0.42
627: IFI30|P13284
cor=0.71, p=1.2e−06
240: IL18BP|O95998
cor=0.74, p=2.5e−07
452: CD83|Q01151
cor=0.83, p=3.9e−10
586: FKBP4|Q02790
cor=0.6, p=0.00011
792: RET|P07949
cor=0.72, p=1.1e−06
107: F3|P13726
cor=0.32, p=0.14
758: LYPD3|O95274
cor=0.69, p=2.5e−05
589: KYAT1|Q16773
cor=0.48, p=0.024
123: FST|P19883
cor=0.67, p=7e−05
354: CD97|P48960
cor=0.44, p=0.0072
618: CD33|P20138
cor=0.36, p=0.031
630: NPM1|P06748
cor=0.63, p=3.8e−05
720: ASAH2|Q9NR71
cor=0.78, p=5e−05
851: PSMD9|O00233
cor=0.69, p=0.00038
360: CLEC14A|Q86T13
cor=0.69, p=3.5e−05
596: RNASE3|P12724

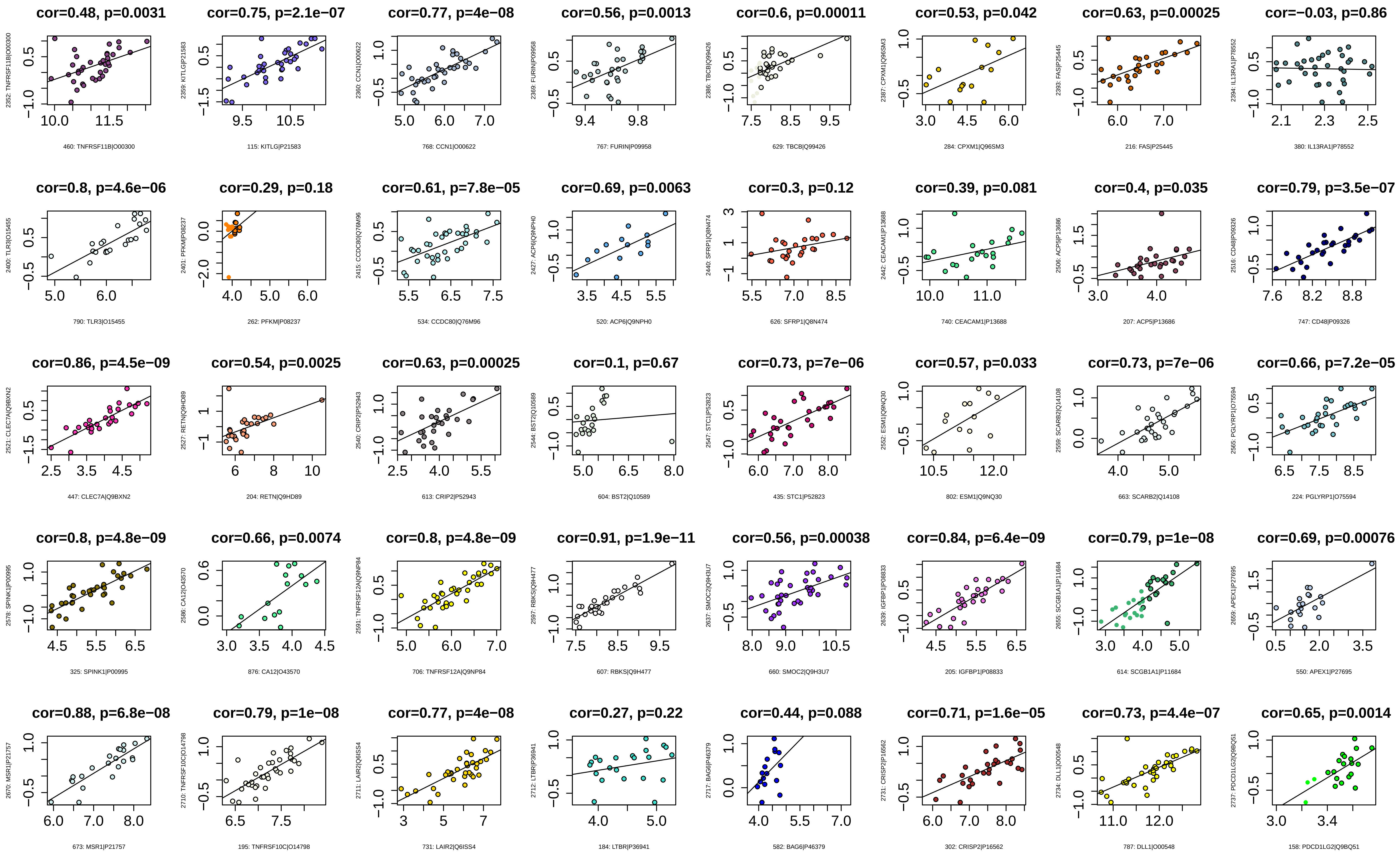

cor=0.48, p=0.0031
2352: TNFRSF11B|O00300
460: TNFRSF11B|O00300
cor=0.75, p=2.1e−07
2359: KITLG|P21583
115: KITLG|P21583
cor=0.77, p=4e−08
2360: CCN1|O00622
768: CCN1|O00622
cor=0.56, p=0.0013
2369: FURIN|P09958
767: FURIN|P09958
cor=0.6, p=0.00011
2386: TBCB|Q99426
629: TBCB|Q99426
cor=0.53, p=0.042
2387: CPXM1|Q96SM3
284: CPXM1|Q96SM3
cor=0.63, p=0.00025
2393: FAS|P25445
216: FAS|P25445
cor=−0.03, p=0.86
2394: IL13RA1|P78552
380: IL13RA1|P78552
cor=0.8, p=4.6e−06
2400: TLR3|O15455
790: TLR3|O15455
cor=0.29, p=0.18
2401: PFKM|P08237
262: PFKM|P08237
cor=0.61, p=7.8e−05
2415: CCDC80|Q76M96
534: CCDC80|Q76M96
cor=0.69, p=0.0063
2427: ACP6|Q9NPH0
520: ACP6|Q9NPH0
cor=0.3, p=0.12
2440: SFRP1|Q8N474
626: SFRP1|Q8N474
cor=0.39, p=0.081
2442: CEACAM1|P13688
740: CEACAM1|P13688
cor=0.4, p=0.035
2506: ACP5|P13686
207: ACP5|P13686
cor=0.79, p=3.5e−07
2516: CD48|P09326
747: CD48|P09326
cor=0.86, p=4.5e−09
2521: CLEC7A|Q9BXN2
447: CLEC7A|Q9BXN2
cor=0.54, p=0.0025
2527: RETN|Q9HD89
204: RETN|Q9HD89
cor=0.63, p=0.00025
2540: CRIP2|P52943
613: CRIP2|P52943
cor=0.1, p=0.67
2544: BST2|Q10589
604: BST2|Q10589
cor=0.73, p=7e−06
2547: STC1|P52823
435: STC1|P52823
cor=0.57, p=0.033
2552: ESM1|Q9NQ30
802: ESM1|Q9NQ30
cor=0.73, p=7e−06
2559: SCARB2|Q14108
663: SCARB2|Q14108
cor=0.66, p=7.2e−05
2565: PGLYRP1|O75594
224: PGLYRP1|O75594
cor=0.8, p=4.8e−09
2578: SPINK1|P00995
325: SPINK1|P00995
cor=0.66, p=0.0074
2586: CA12|O43570
876: CA12|O43570
cor=0.8, p=4.8e−09
2591: TNFRSF12A|Q9NP84
706: TNFRSF12A|Q9NP84
cor=0.91, p=1.9e−11
2597: RBKS|Q9H477
607: RBKS|Q9H477
cor=0.56, p=0.00038
2637: SMOC2|Q9H3U7
660: SMOC2|Q9H3U7
cor=0.84, p=6.4e−09
2639: IGFBP1|P08833
205: IGFBP1|P08833
cor=0.79, p=1e−08
2655: SCGB1A1|P11684
614: SCGB1A1|P11684
cor=0.69, p=0.00076
2659: APEX1|P27695
550: APEX1|P27695
cor=0.88, p=6.8e−08
2670: MSR1|P21757
673: MSR1|P21757
cor=0.79, p=1e−08
2710: TNFRSF10C|O14798
195: TNFRSF10C|O14798
cor=0.77, p=4e−08
2711: LAIR2|Q6ISS4
731: LAIR2|Q6ISS4
cor=0.27, p=0.22
2712: LTBR|P36941
184: LTBR|P36941
cor=0.44, p=0.088
2717: BAG6|P46379
582: BAG6|P46379
cor=0.71, p=1.6e−05
2731: CRISP2|P16562
302: CRISP2|P16562
cor=0.73, p=4.4e−07
2734: DLL1|O00548
787: DLL1|O00548
cor=0.65, p=0.0014
2737: PDCD1LG2|Q9BQ51
158: PDCD1LG2|Q9BQ51

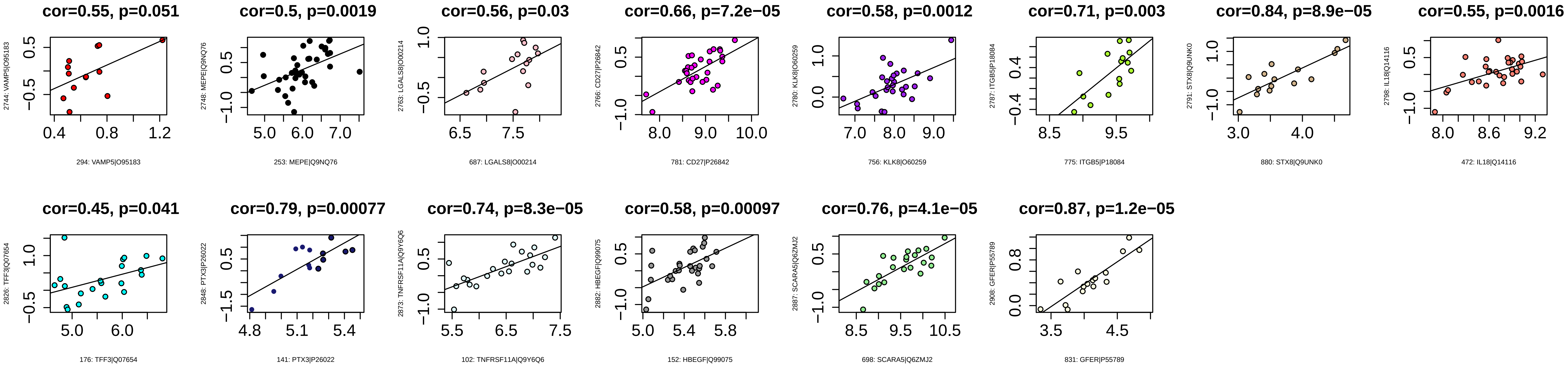

cor=0.55, p=0.051
2744: VAMP5|O95183
294: VAMP5|O95183
cor=0.5, p=0.0019
2748: MEPE|Q9NQ76
253: MEPE|Q9NQ76
cor=0.56, p=0.03
2763: LGALS8|O00214
687: LGALS8|O00214
cor=0.66, p=7.2e−05
2766: CD27|P26842
781: CD27|P26842
cor=0.58, p=0.0012
2780: KLK8|O60259
756: KLK8|O60259
cor=0.71, p=0.003
2787: ITGB5|P18084
775: ITGB5|P18084
cor=0.84, p=8.9e−05
2791: STX8|Q9UNK0
880: STX8|Q9UNK0
cor=0.55, p=0.0016
2798: IL18|Q14116
472: IL18|Q14116
cor=0.45, p=0.041
2826: TFF3|Q07654
176: TFF3|Q07654
cor=0.79, p=0.00077
2848: PTX3|P26022
141: PTX3|P26022
cor=0.74, p=8.3e−05
2873: TNFRSF11A|Q9Y6Q6
102: TNFRSF11A|Q9Y6Q6
cor=0.58, p=0.00097
2882: HBEGF|Q99075
152: HBEGF|Q99075
cor=0.76, p=4.1e−05
2887: SCARA5|Q6ZMJ2
698: SCARA5|Q6ZMJ2
cor=0.87, p=1.2e−05
2908: GFER|P55789
831: GFER|P55789

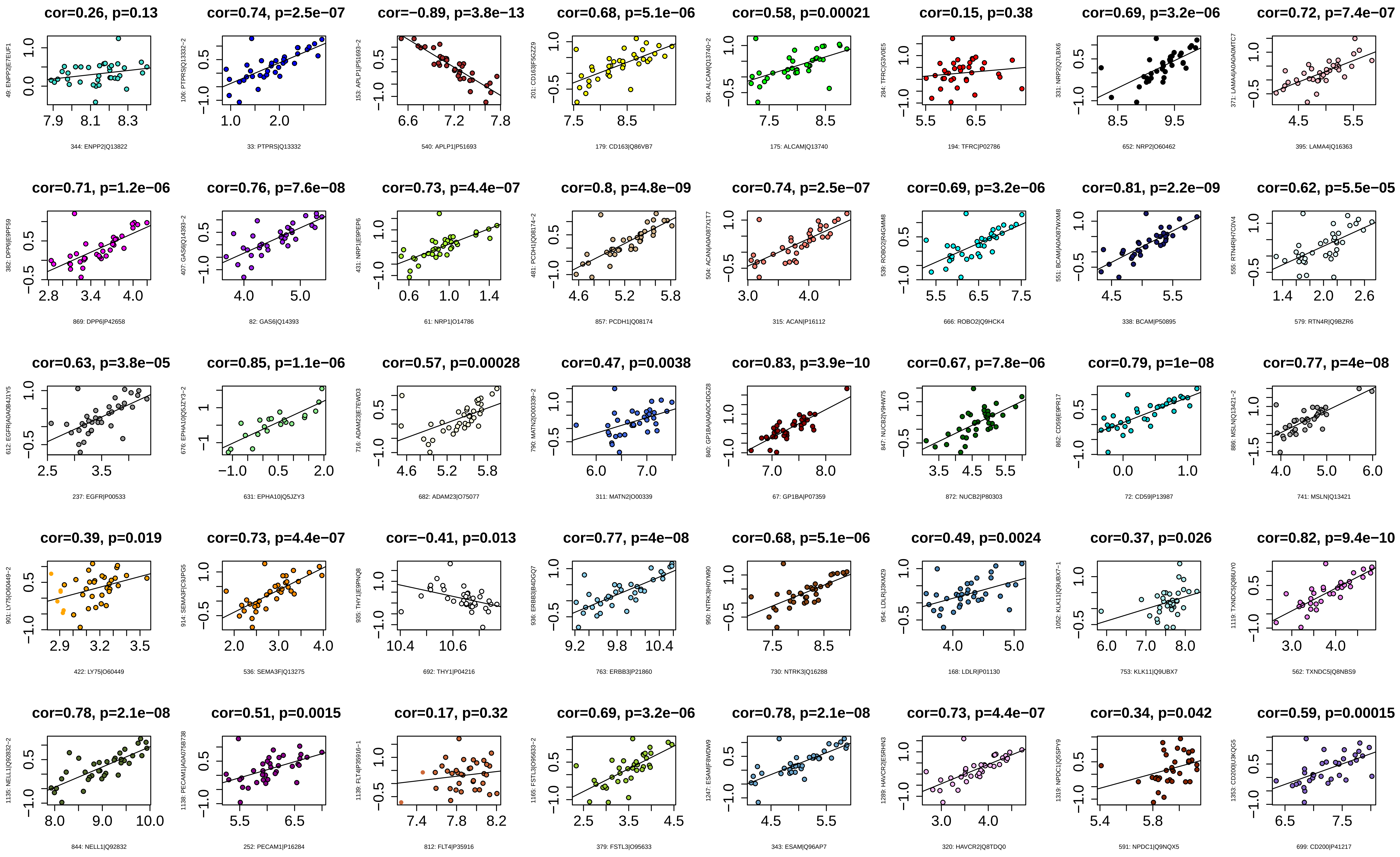

cor=0.26, p=0.13
344: ENPP2|Q13822
cor=0.74, p=2.5e−07
33: PTPRS|Q13332
cor=−0.89, p=3.8e−13
540: APLP1|P51693
cor=0.68, p=5.1e−06
179: CD163|Q86VB7
cor=0.58, p=0.00021
175: ALCAM|Q13740
cor=0.15, p=0.38
194: TFRC|P02786
cor=0.69, p=3.2e−06
652: NRP2|O60462
cor=0.72, p=7.4e−07
395: LAMA4|Q16363
cor=0.71, p=1.2e−06
869: DPP6|P42658
cor=0.76, p=7.6e−08
82: GAS6|Q14393
cor=0.73, p=4.4e−07
61: NRP1|O14786
cor=0.8, p=4.8e−09
857: PCDH1|Q08174
cor=0.74, p=2.5e−07
315: ACAN|P16112
cor=0.69, p=3.2e−06
666: ROBO2|Q9HCK4
cor=0.81, p=2.2e−09
338: BCAM|P50895
cor=0.62, p=5.5e−05
579: RTN4R|Q9BZR6
cor=0.63, p=3.8e−05
237: EGFR|P00533
cor=0.85, p=1.1e−06
631: EPHA10|Q5JZY3
cor=0.57, p=0.00028
682: ADAM23|O75077
cor=0.47, p=0.0038
311: MATN2|O00339
cor=0.83, p=3.9e−10
67: GP1BA|P07359
cor=0.67, p=7.8e−06
872: NUCB2|P80303
cor=0.79, p=1e−08
72: CD59|P13987
cor=0.77, p=4e−08
741: MSLN|Q13421
cor=0.39, p=0.019
422: LY75|O60449
cor=0.73, p=4.4e−07
536: SEMA3F|Q13275
cor=−0.41, p=0.013
692: THY1|P04216
cor=0.77, p=4e−08
763: ERBB3|P21860
cor=0.68, p=5.1e−06
730: NTRK3|Q16288
cor=0.49, p=0.0024
168: LDLR|P01130
cor=0.37, p=0.026
753: KLK11|Q9UBX7
cor=0.82, p=9.4e−10
562: TXNDC5|Q8NBS9
cor=0.78, p=2.1e−08
844: NELL1|Q92832
cor=0.51, p=0.0015
252: PECAM1|P16284
cor=0.17, p=0.32
812: FLT4|P35916
cor=0.69, p=3.2e−06
379: FSTL3|O95633
cor=0.78, p=2.1e−08
343: ESAM|Q96AP7
cor=0.73, p=4.4e−07
320: HAVCR2|Q8TDQ0
cor=0.34, p=0.042
591: NPDC1|Q9NQX5
cor=0.59, p=0.00015
699: CD200|P41217

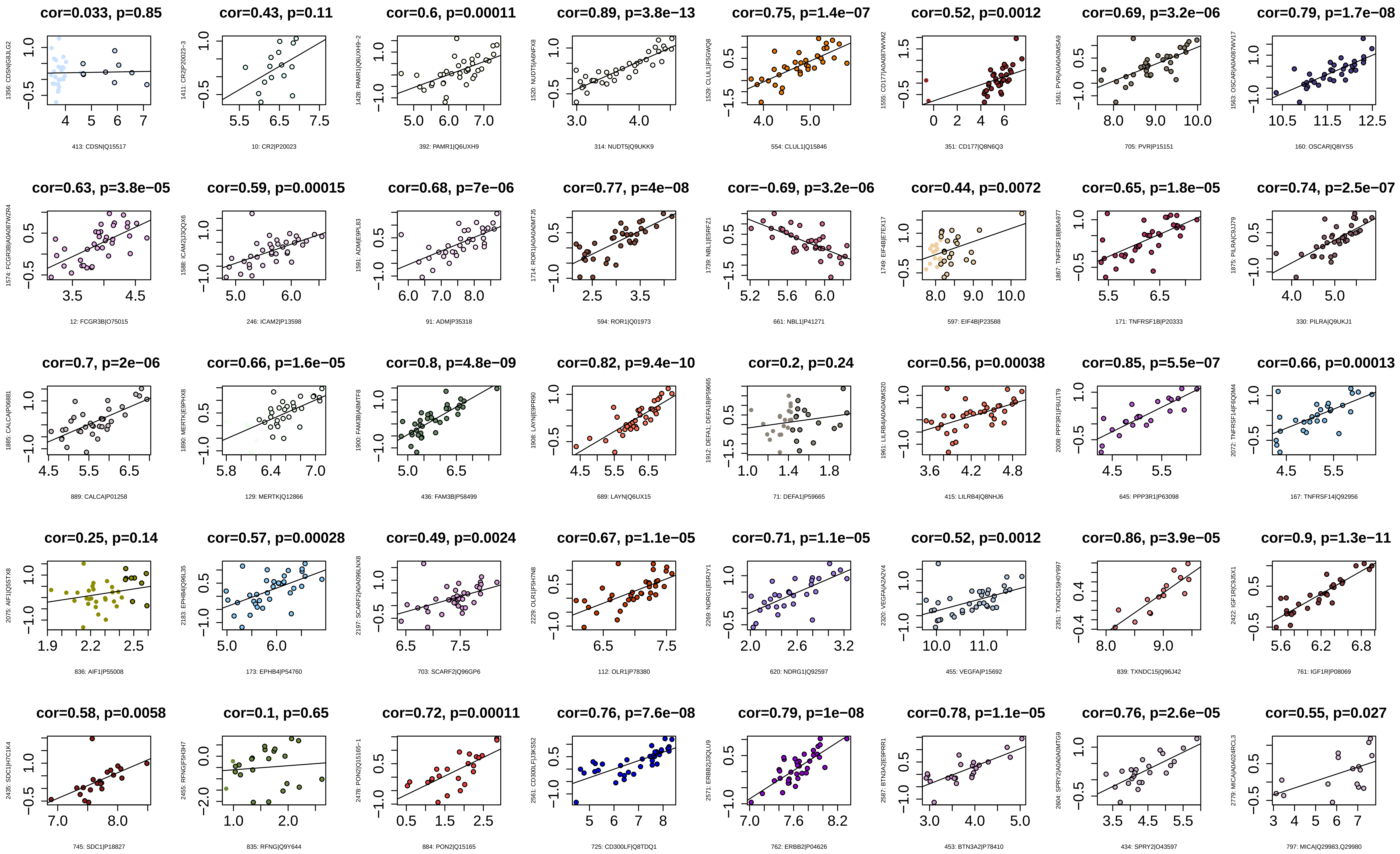
cor=0.033, p=0.85
413: CDSN|Q15517
cor=0.43, p=0.11
10: CR2|P20023
cor=0.6, p=0.00011
392: PAMR1|Q6UXH9
cor=0.89, p=3.8e−13
314: NUDT5|Q9UKK9
cor=0.75, p=1.4e−07
554: CLUL1|Q15846
cor=0.52, p=0.0012
351: CD177|Q8N6Q3
cor=0.69, p=3.2e−06
705: PVR|P15151
cor=0.79, p=1.7e−08
160: OSCAR|Q8IYS5
cor=0.63, p=3.8e−05
12: FCGR3B|O75015
cor=0.59, p=0.00015
246: ICAM2|P13598
cor=0.68, p=7e−06
91: ADM|P35318
cor=0.77, p=4e−08
594: ROR1|Q01973
cor=−0.69, p=3.2e−06
661: NBL1|P41271
cor=0.44, p=0.0072
597: EIF4B|P23588
cor=0.65, p=1.8e−05
171: TNFRSF1B|P20333
cor=0.74, p=2.5e−07
330: PILRA|Q9UKJ1
cor=0.7, p=2e−06
889: CALCA|P01258
cor=0.66, p=1.6e−05
129: MERTK|Q12866
cor=0.8, p=4.8e−09
436: FAM3B|P58499
cor=0.82, p=9.4e−10
689: LAYN|Q6UX15
cor=0.2, p=0.24
71: DEFA1|P59665
cor=0.56, p=0.00038
415: LILRB4|Q8NHJ6
cor=0.85, p=5.5e−07
645: PPP3R1|P63098
cor=0.66, p=0.00013
167: TNFRSF14|Q92956
cor=0.25, p=0.14
836: AIF1|P55008
cor=0.57, p=0.00028
173: EPHB4|P54760
cor=0.49, p=0.0024
703: SCARF2|Q96GP6
cor=0.67, p=1.1e−05
112: OLR1|P78380
cor=0.71, p=1.1e−05
620: NDRG1|Q92597
cor=0.52, p=0.0012
455: VEGFA|P15692
cor=0.86, p=3.9e−05
839: TXNDC15|Q96J42
cor=0.9, p=1.3e−11
761: IGF1R|P08069
cor=0.58, p=0.0058
745: SDC1|P18827
cor=0.1, p=0.65
835: RFNG|Q9Y644
cor=0.72, p=0.00011
884: PON2|Q15165
cor=0.76, p=7.6e−08
725: CD300LF|Q8TDQ1
cor=0.79, p=1e−08
762: ERBB2|P04626
cor=0.78, p=1.1e−05
453: BTN3A2|P78410
cor=0.76, p=2.6e−05
434: SPRY2|O43597
cor=0.55, p=0.027
797: MICA|Q29983,Q29980

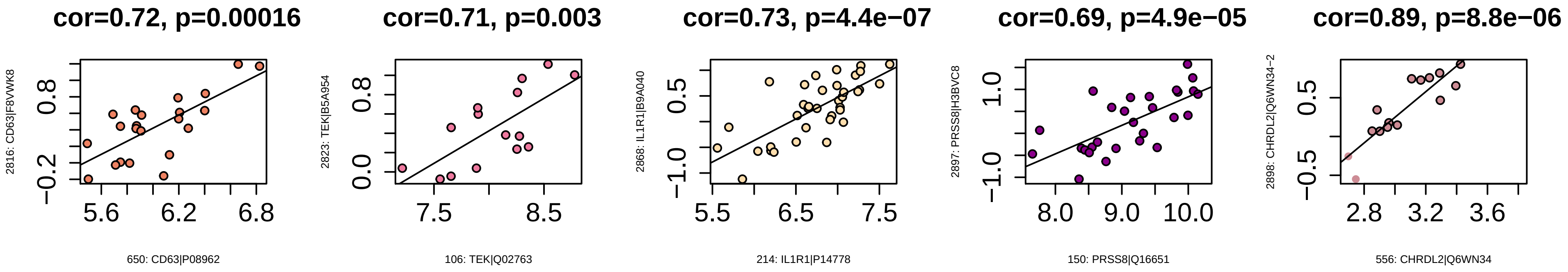
cor=0.72, p=0.00016
2816: CD63|F8VWK8
650: CD63|P08962
cor=0.71, p=0.003
2823: TEK|B5A954
106: TEK|Q02763
cor=0.73, p=4.4e−07
2868: IL1R1|B9A040
214: IL1R1|P14778
cor=0.69, p=4.9e−05
2897: PRSS8|H3BVC8
150: PRSS8|Q16651
cor=0.89, p=8.8e−06
2898: CHRDL2|Q6WN34−2
556: CHRDL2|Q6WN34

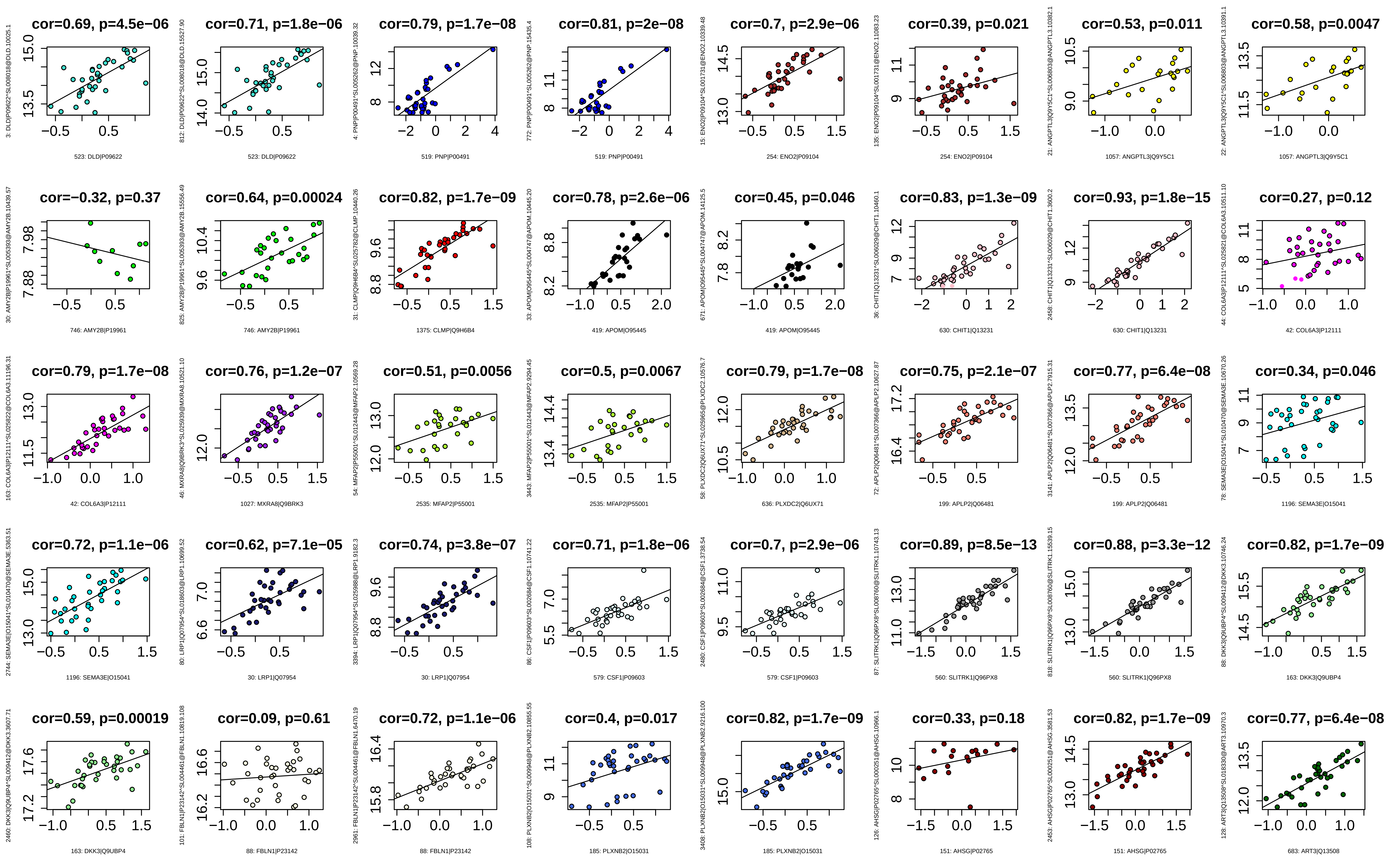
cor=0.69, p=4.5e−06
523: DLD|P09622
cor=0.71, p=1.8e−06
523: DLD|P09622
cor=0.79, p=1.7e−08
519: PNP|P00491
cor=0.81, p=2e−08
519: PNP|P00491
cor=0.7, p=2.9e−06
254: ENO2|P09104
cor=0.39, p=0.021
254: ENO2|P09104
cor=0.53, p=0.011
1057: ANGPTL3|Q9Y5C1
cor=0.58, p=0.0047
1057: ANGPTL3|Q9Y5C1
cor=−0.32, p=0.37
746: AMY2B|P19961
cor=0.64, p=0.00024
746: AMY2B|P19961
cor=0.82, p=1.7e−09
1375: CLMP|Q9H6B4
cor=0.78, p=2.6e−06
419: APOM|O95445
cor=0.45, p=0.046
419: APOM|O95445
cor=0.83, p=1.3e−09
630: CHIT1|Q13231
cor=0.93, p=1.8e−15
630: CHIT1|Q13231
cor=0.27, p=0.12
42: COL6A3|P12111
cor=0.79, p=1.7e−08
42: COL6A3|P12111
cor=0.76, p=1.2e−07
1027: MXRA8|Q9BRK3
cor=0.51, p=0.0056
2535: MFAP2|P55001
cor=0.5, p=0.0067
2535: MFAP2|P55001
cor=0.79, p=1.7e−08
636: PLXDC2|Q6UX71
cor=0.75, p=2.1e−07
199: APLP2|Q06481
cor=0.77, p=6.4e−08
199: APLP2|Q06481
cor=0.34, p=0.046
1196: SEMA3E|O15041
cor=0.72, p=1.1e−06
1196: SEMA3E|O15041
cor=0.62, p=7.1e−05
30: LRP1|Q07954
cor=0.74, p=3.8e−07
30: LRP1|Q07954
cor=0.71, p=1.8e−06
579: CSF1|P09603
cor=0.7, p=2.9e−06
579: CSF1|P09603
cor=0.89, p=8.5e−13
560: SLITRK1|Q96PX8
cor=0.88, p=3.3e−12
560: SLITRK1|Q96PX8
cor=0.82, p=1.7e−09
163: DKK3|Q9UBP4
cor=0.59, p=0.00019
163: DKK3|Q9UBP4
cor=0.09, p=0.61
88: FBLN1|P23142
cor=0.72, p=1.1e−06
88: FBLN1|P23142
cor=0.4, p=0.017
185: PLXNB2|O15031
cor=0.82, p=1.7e−09
185: PLXNB2|O15031
cor=0.33, p=0.18
151: AHSG|P02765
cor=0.82, p=1.7e−09
151: AHSG|P02765
cor=0.77, p=6.4e−08
683: ART3|Q13508

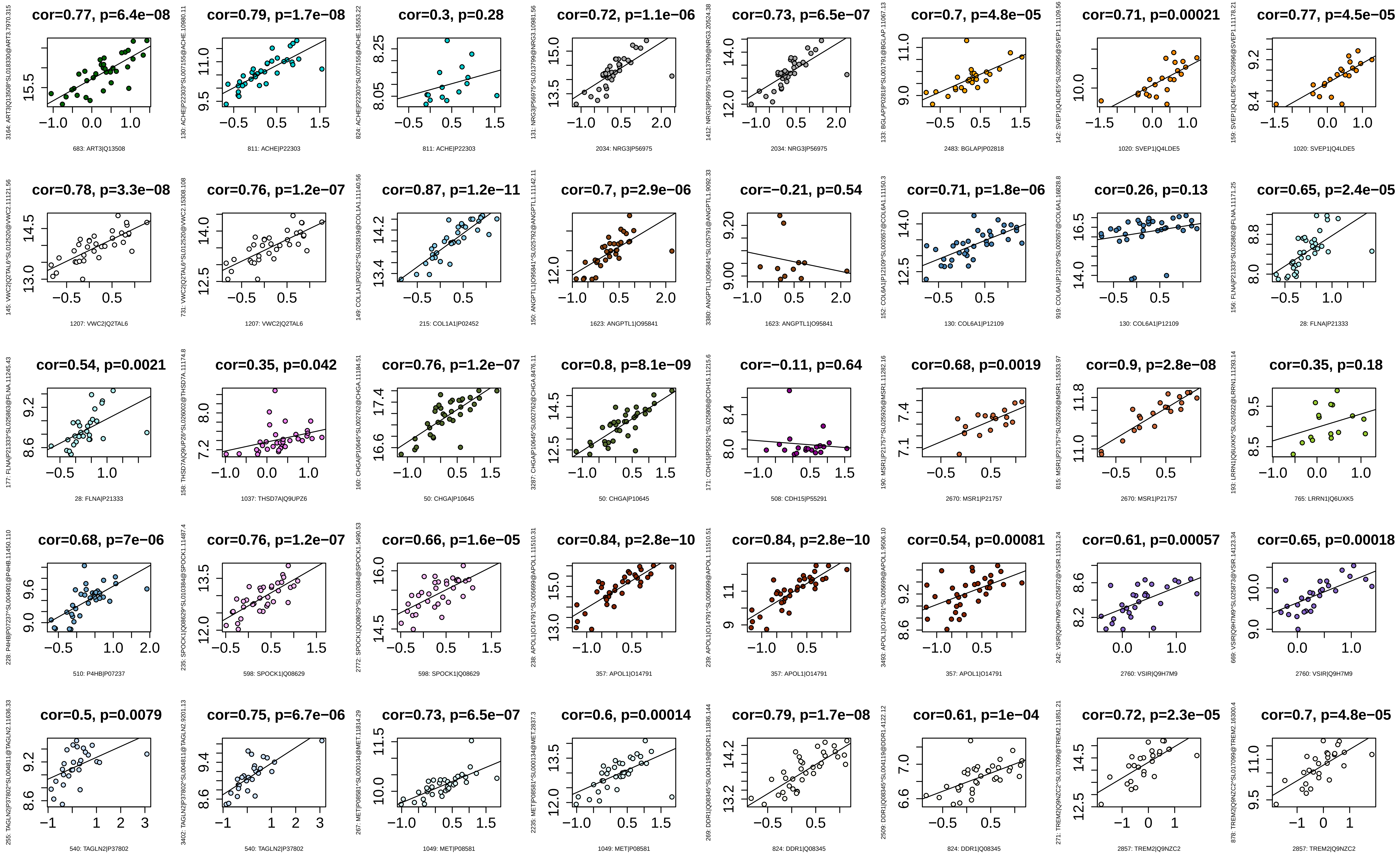
cor=0.77, p=6.4e−08
3164: ART3|Q13508^SL018330@ART3.7970.315
683: ART3|Q13508
cor=0.79, p=1.7e−08
130: ACHE|P22303^SL007155@ACHE.10980.11
811: ACHE|P22303
cor=0.3, p=0.28
824: ACHE|P22303^SL007155@ACHE.15553.22
811: ACHE|P22303
cor=0.72, p=1.1e−06
131: NRG3|P56975^SL013799@NRG3.10981.56
2034: NRG3|P56975
cor=0.73, p=6.5e−07
1412: NRG3|P56975^SL013799@NRG3.20524.38
2034: NRG3|P56975
cor=0.7, p=4.8e−05
133: BGLAP|P02818^SL001791@BGLAP.11067.13
2483: BGLAP|P02818
cor=0.71, p=0.00021
142: SVEP1|Q4LDE5^SL025995@SVEP1.11109.56
1020: SVEP1|Q4LDE5
cor=0.77, p=4.5e−05
159: SVEP1|Q4LDE5^SL025996@SVEP1.11178.21
1020: SVEP1|Q4LDE5
cor=0.78, p=3.3e−08
145: VWC2|Q2TAL6^SL012520@VWC2.11121.56
1207: VWC2|Q2TAL6
cor=0.76, p=1.2e−07
731: VWC2|Q2TAL6^SL012520@VWC2.15308.108
1207: VWC2|Q2TAL6
cor=0.87, p=1.2e−11
149: COL1A1|P02452^SL025819@COL1A1.11140.56
215: COL1A1|P02452
cor=0.7, p=2.9e−06
150: ANGPTL1|O95841^SL025792@ANGPTL1.11142.11
1623: ANGPTL1|O95841
cor=−0.21, p=0.54
3380: ANGPTL1|O95841^SL025791@ANGPTL1.9092.33
1623: ANGPTL1|O95841
cor=0.71, p=1.8e−06
152: COL6A1|P12109^SL002657@COL6A1.11150.3
130: COL6A1|P12109
cor=0.26, p=0.13
919: COL6A1|P12109^SL002657@COL6A1.16828.8
130: COL6A1|P12109
cor=0.65, p=2.4e−05
156: FLNA|P21333^SL025862@FLNA.11171.25
28: FLNA|P21333
cor=0.54, p=0.0021
177: FLNA|P21333^SL025863@FLNA.11245.43
28: FLNA|P21333
cor=0.35, p=0.042
158: THSD7A|Q9UPZ6^SL026002@THSD7A.11174.8
1037: THSD7A|Q9UPZ6
cor=0.76, p=1.2e−07
160: CHGA|P10645^SL002762@CHGA.11184.51
50: CHGA|P10645
cor=0.8, p=8.1e−09
3287: CHGA|P10645^SL002762@CHGA.8476.11
50: CHGA|P10645
cor=−0.11, p=0.64
171: CDH15|P55291^SL025808@CDH15.11215.6
508: CDH15|P55291
cor=0.68, p=0.0019
190: MSR1|P21757^SL025926@MSR1.11282.16
2670: MSR1|P21757
cor=0.9, p=2.8e−08
815: MSR1|P21757^SL025926@MSR1.15533.97
2670: MSR1|P21757
cor=0.35, p=0.18
193: LRRN1|Q6UXK5^SL025922@LRRN1.11293.14
765: LRRN1|Q6UXK5
cor=0.68, p=7e−06
228: P4HB|P07237^SL004901@P4HB.11450.110
510: P4HB|P07237
cor=0.76, p=1.2e−07
235: SPOCK1|Q08629^SL010384@SPOCK1.11487.4
598: SPOCK1|Q08629
cor=0.66, p=1.6e−05
2772: SPOCK1|Q08629^SL010384@SPOCK1.5490.53
598: SPOCK1|Q08629
cor=0.84, p=2.8e−10
238: APOL1|O14791^SL005699@APOL1.11510.31
357: APOL1|O14791
cor=0.84, p=2.8e−10
239: APOL1|O14791^SL005699@APOL1.11510.51
357: APOL1|O14791
cor=0.54, p=0.00081
3493: APOL1|O14791^SL005699@APOL1.9506.10
357: APOL1|O14791
cor=0.61, p=0.00057
242: VSIR|Q9H7M9^SL025872@VSIR.11531.24
2760: VSIR|Q9H7M9
cor=0.65, p=0.00018
669: VSIR|Q9H7M9^SL025873@VSIR.14123.34
2760: VSIR|Q9H7M9
cor=0.5, p=0.0079
255: TAGLN2|P37802^SL004811@TAGLN2.11636.33
540: TAGLN2|P37802
cor=0.75, p=6.7e−06
3402: TAGLN2|P37802^SL004811@TAGLN2.9201.13
540: TAGLN2|P37802
cor=0.73, p=6.5e−07
267: MET|P08581^SL000134@MET.11814.29
1049: MET|P08581
cor=0.6, p=0.00014
2235: MET|P08581^SL000134@MET.2837.3
1049: MET|P08581
cor=0.79, p=1.7e−08
269: DDR1|Q08345^SL004119@DDR1.11836.144
824: DDR1|Q08345
cor=0.61, p=1e−04
2509: DDR1|Q08345^SL004119@DDR1.4122.12
824: DDR1|Q08345
cor=0.72, p=2.3e−05
271: TREM2|Q9NZC2^SL017099@TREM2.11851.21
2857: TREM2|Q9NZC2
cor=0.7, p=4.8e−05
878: TREM2|Q9NZC2^SL017099@TREM2.16300.4
2857: TREM2|Q9NZC2

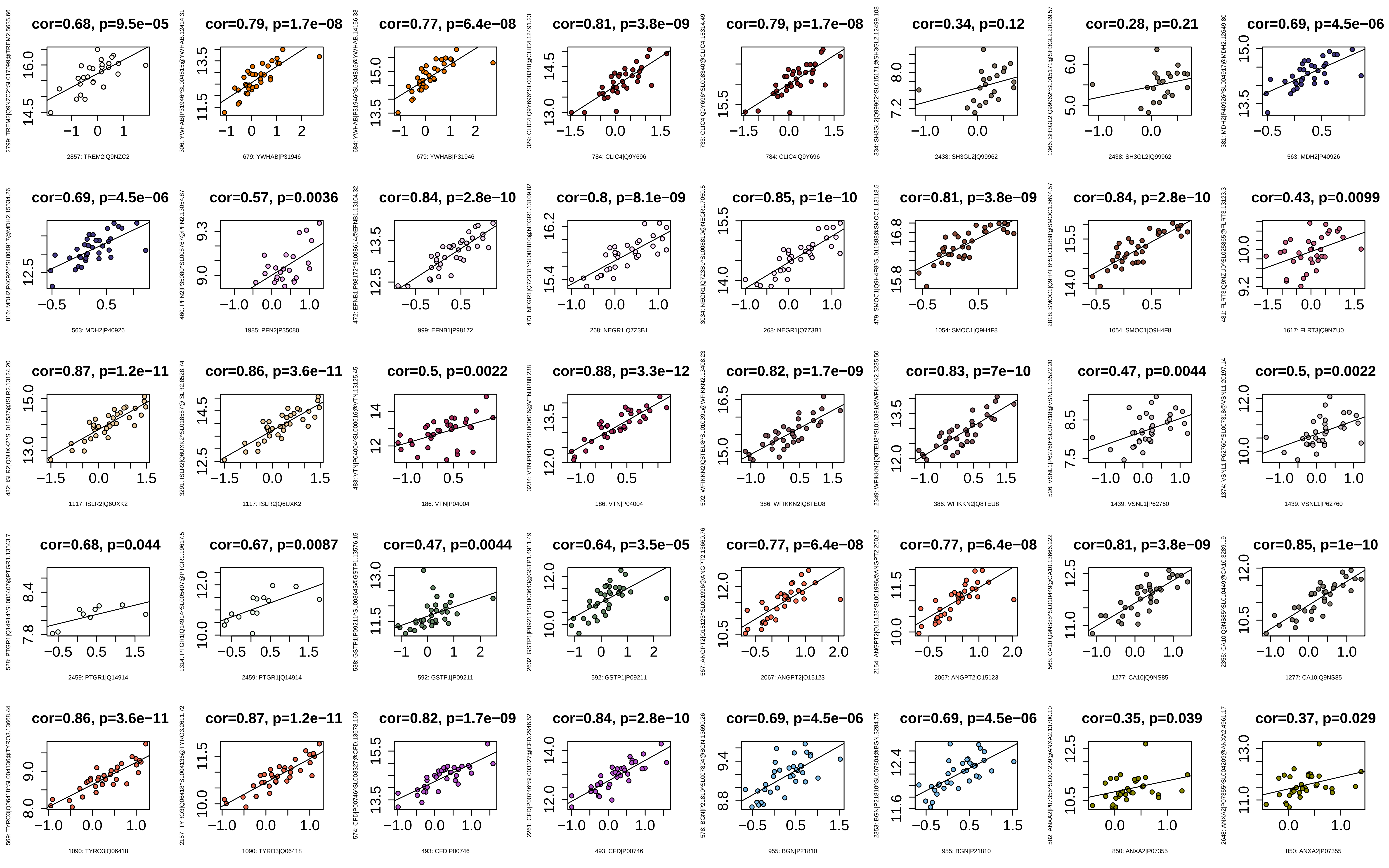

cor=0.68, p=9.5e−05
cor=0.79, p=1.7e−08
cor=0.77, p=6.4e−08
cor=0.81, p=3.8e−09
cor=0.79, p=1.7e−08
cor=0.34, p=0.12
cor=0.28, p=0.21
cor=0.69, p=4.5e−06
2857: TREM2|Q9NZC2
679: YWHAB|P31946
679: YWHAB|P31946
784: CLIC4|Q9Y696
784: CLIC4|Q9Y696
2438: SH3GL2|Q99962
2438: SH3GL2|Q99962
563: MDH2|P40926
cor=0.69, p=4.5e−06
cor=0.57, p=0.0036
cor=0.84, p=2.8e−10
cor=0.8, p=8.1e−09
cor=0.85, p=1e−10
cor=0.81, p=3.8e−09
cor=0.84, p=2.8e−10
cor=0.43, p=0.0099
563: MDH2|P40926
1985: PFN2|P35080
999: EFNB1|P98172
268: NEGR1|Q7Z3B1
268: NEGR1|Q7Z3B1
1054: SMOC1|Q9H4F8
1054: SMOC1|Q9H4F8
1617: FLRT3|Q9NZU0
cor=0.87, p=1.2e−11
cor=0.86, p=3.6e−11
cor=0.5, p=0.0022
cor=0.88, p=3.3e−12
cor=0.82, p=1.7e−09
cor=0.83, p=7e−10
cor=0.47, p=0.0044
cor=0.5, p=0.0022
1117: ISLR2|Q6UXK2
1117: ISLR2|Q6UXK2
186: VTN|P04004
186: VTN|P04004
386: WFIKKN2|Q8TEU8
386: WFIKKN2|Q8TEU8
1439: VSNL1|P62760
1439: VSNL1|P62760
cor=0.68, p=0.044
cor=0.67, p=0.0087
cor=0.47, p=0.0044
cor=0.64, p=3.5e−05
cor=0.77, p=6.4e−08
cor=0.77, p=6.4e−08
cor=0.81, p=3.8e−09
cor=0.85, p=1e−10
2459: PTGR1|Q14914
2459: PTGR1|Q14914
592: GSTP1|P09211
592: GSTP1|P09211
2067: ANGPT2|O15123
2067: ANGPT2|O15123
1277: CA10|Q9NS85
1277: CA10|Q9NS85
cor=0.86, p=3.6e−11
cor=0.87, p=1.2e−11
cor=0.82, p=1.7e−09
cor=0.84, p=2.8e−10
cor=0.69, p=4.5e−06
cor=0.69, p=4.5e−06
cor=0.35, p=0.039
cor=0.37, p=0.029
1090: TYRO3|Q06418
1090: TYRO3|Q06418
493: CFD|P00746
493: CFD|P00746
955: BGN|P21810
955: BGN|P21810
850: ANXA2|P07355
850: ANXA2|P07355

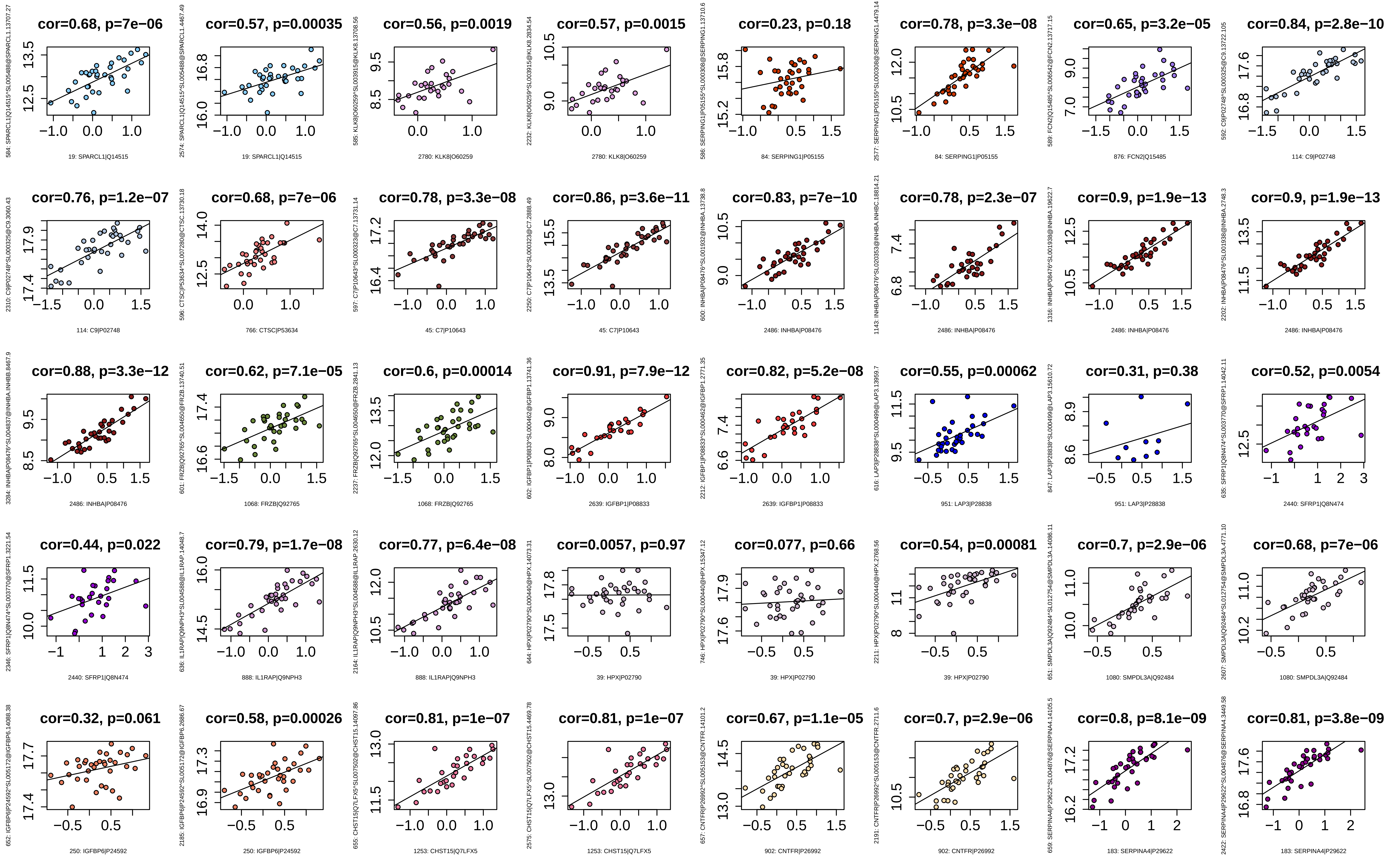

cor=0.68, p=7e−06
19: SPARCL1|Q14515
cor=0.57, p=0.00035
19: SPARCL1|Q14515
cor=0.56, p=0.0019
2780: KLK8|O60259
cor=0.57, p=0.0015
2780: KLK8|O60259
cor=0.23, p=0.18
84: SERPING1|P05155
cor=0.78, p=3.3e−08
84: SERPING1|P05155
cor=0.65, p=3.2e−05
876: FCN2|Q15485
cor=0.84, p=2.8e−10
114: C9|P02748
cor=0.76, p=1.2e−07
114: C9|P02748
cor=0.68, p=7e−06
766: CTSC|P53634
cor=0.78, p=3.3e−08
45: C7|P10643
cor=0.86, p=3.6e−11
45: C7|P10643
cor=0.83, p=7e−10
2486: INHBA|P08476
cor=0.78, p=2.3e−07
2486: INHBA|P08476
cor=0.9, p=1.9e−13
2486: INHBA|P08476
cor=0.9, p=1.9e−13
2486: INHBA|P08476
cor=0.88, p=3.3e−12
2486: INHBA|P08476
cor=0.62, p=7.1e−05
1068: FRZB|Q92765
cor=0.6, p=0.00014
1068: FRZB|Q92765
cor=0.91, p=7.9e−12
2639: IGFBP1|P08833
cor=0.82, p=5.2e−08
2639: IGFBP1|P08833
cor=0.55, p=0.00062
951: LAP3|P28838
cor=0.31, p=0.38
951: LAP3|P28838
cor=0.52, p=0.0054
2440: SFRP1|Q8N474
cor=0.44, p=0.022
2440: SFRP1|Q8N474
cor=0.79, p=1.7e−08
888: IL1RAP|Q9NPH3
cor=0.77, p=6.4e−08
888: IL1RAP|Q9NPH3
cor=0.0057, p=0.97
39: HPX|P02790
cor=0.077, p=0.66
39: HPX|P02790
cor=0.54, p=0.00081
39: HPX|P02790
cor=0.7, p=2.9e−06
1080: SMPDL3A|Q92484
cor=0.68, p=7e−06
1080: SMPDL3A|Q92484
cor=0.32, p=0.061
250: IGFBP6|P24592
cor=0.58, p=0.00026
250: IGFBP6|P24592
cor=0.81, p=1e−07
1253: CHST15|Q7LFX5
cor=0.81, p=1e−07
1253: CHST15|Q7LFX5
cor=0.67, p=1.1e−05
902: CNTFR|P26992
cor=0.7, p=2.9e−06
902: CNTFR|P26992
cor=0.8, p=8.1e−09
183: SERPINA4|P29622
cor=0.81, p=3.8e−09
183: SERPINA4|P29622

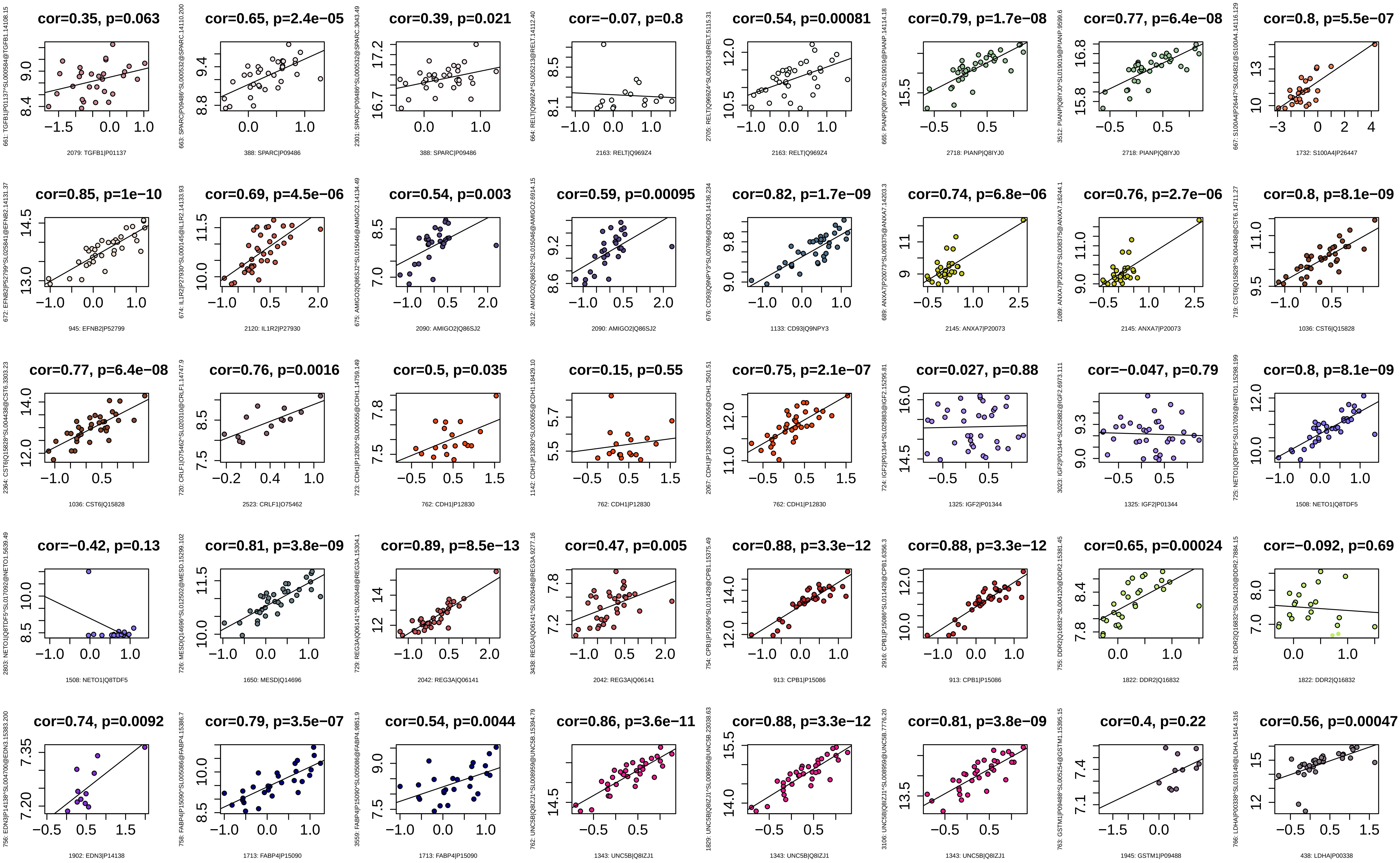
cor=0.35, p=0.063
661: TGFB1|P01137^SL000584@TGFB1.14108.15
2079: TGFB1|P01137
cor=0.65, p=2.4e−05
663: SPARC|P09486^SL000532@SPARC.14110.200
388: SPARC|P09486
cor=0.39, p=0.021
2301: SPARC|P09486^SL000532@SPARC.3043.49
388: SPARC|P09486
cor=−0.07, p=0.8
664: RELT|Q969Z4^SL005213@RELT.14112.40
2163: RELT|Q969Z4
cor=0.54, p=0.00081
2705: RELT|Q969Z4^SL005213@RELT.5115.31
2163: RELT|Q969Z4
cor=0.79, p=1.7e−08
665: PIANP|Q8IYJ0^SL019019@PIANP.14114.18
2718: PIANP|Q8IYJ0
cor=0.77, p=6.4e−08
3512: PIANP|Q8IYJ0^SL019019@PIANP.9599.6
2718: PIANP|Q8IYJ0
cor=0.8, p=5.5e−07
667: S100A4|P26447^SL004821@S100A4.14116.129
1732: S100A4|P26447
cor=0.85, p=1e−10
672: EFNB2|P52799^SL025841@EFNB2.14131.37
945: EFNB2|P52799
cor=0.69, p=4.5e−06
674: IL1R2|P27930^SL000145@IL1R2.14133.93
2120: IL1R2|P27930
cor=0.54, p=0.003
675: AMIGO2|Q86SJ2^SL015046@AMIGO2.14134.49
2090: AMIGO2|Q86SJ2
cor=0.59, p=0.00095
3012: AMIGO2|Q86SJ2^SL015046@AMIGO2.6914.15
2090: AMIGO2|Q86SJ2
cor=0.82, p=1.7e−09
676: CD93|Q9NPY3^SL007696@CD93.14136.234
1133: CD93|Q9NPY3
cor=0.74, p=6.8e−06
689: ANXA7|P20073^SL008375@ANXA7.14203.3
2145: ANXA7|P20073
cor=0.76, p=2.7e−06
1089: ANXA7|P20073^SL008375@ANXA7.18244.1
2145: ANXA7|P20073
cor=0.8, p=8.1e−09
719: CST6|Q15828^SL004438@CST6.14711.27
1036: CST6|Q15828
cor=0.77, p=6.4e−08
2364: CST6|Q15828^SL004438@CST6.3303.23
1036: CST6|Q15828
cor=0.76, p=0.0016
720: CRLF1|O75462^SL020310@CRLF1.14747.9
2523: CRLF1|O75462
cor=0.5, p=0.035
723: CDH1|P12830^SL000055@CDH1.14759.149
762: CDH1|P12830
cor=0.15, p=0.55
1142: CDH1|P12830^SL000055@CDH1.18429.10
762: CDH1|P12830
cor=0.75, p=2.1e−07
2067: CDH1|P12830^SL000055@CDH1.2501.51
762: CDH1|P12830
cor=0.027, p=0.88
724: IGF2|P01344^SL025883@IGF2.15295.81
1325: IGF2|P01344
cor=−0.047, p=0.79
3023: IGF2|P01344^SL025882@IGF2.6973.111
1325: IGF2|P01344
cor=0.8, p=8.1e−09
725: NETO1|Q8TDF5^SL017092@NETO1.15298.199
1508: NETO1|Q8TDF5
cor=−0.42, p=0.13
2803: NETO1|Q8TDF5^SL017092@NETO1.5639.49
1508: NETO1|Q8TDF5
cor=0.81, p=3.8e−09
726: MESD|Q14696^SL012502@MESD.15299.102
1650: MESD|Q14696
cor=0.89, p=8.5e−13
729: REG3A|Q06141^SL002648@REG3A.15304.1
2042: REG3A|Q06141
cor=0.47, p=0.005
3438: REG3A|Q06141^SL002648@REG3A.9277.16
2042: REG3A|Q06141
cor=0.88, p=3.3e−12
754: CPB1|P15086^SL011428@CPB1.15375.49
913: CPB1|P15086
cor=0.88, p=3.3e−12
2916: CPB1|P15086^SL011428@CPB1.6356.3
913: CPB1|P15086
cor=0.65, p=0.00024
755: DDR2|Q16832^SL004120@DDR2.15381.45
1822: DDR2|Q16832
cor=−0.092, p=0.69
3134: DDR2|Q16832^SL004120@DDR2.7884.15
1822: DDR2|Q16832
cor=0.74, p=0.0092
756: EDN3|P14138^SL004700@EDN3.15383.200
1902: EDN3|P14138
cor=0.79, p=3.5e−07
758: FABP4|P15090^SL005086@FABP4.15386.7
1713: FABP4|P15090
cor=0.54, p=0.0044
3559: FABP4|P15090^SL005086@FABP4.9851.9
1713: FABP4|P15090
cor=0.86, p=3.6e−11
762: UNC5B|Q8IZJ1^SL008959@UNC5B.15394.79
1343: UNC5B|Q8IZJ1
cor=0.88, p=3.3e−12
1829: UNC5B|Q8IZJ1^SL008959@UNC5B.23038.63
1343: UNC5B|Q8IZJ1
cor=0.81, p=3.8e−09
3106: UNC5B|Q8IZJ1^SL008959@UNC5B.7776.20
1343: UNC5B|Q8IZJ1
cor=0.4, p=0.22
763: GSTM1|P09488^SL005254@GSTM1.15395.15
1945: GSTM1|P09488
cor=0.56, p=0.00047
766: LDHA|P00338^SL019149@LDHA.15414.316
438: LDHA|P00338

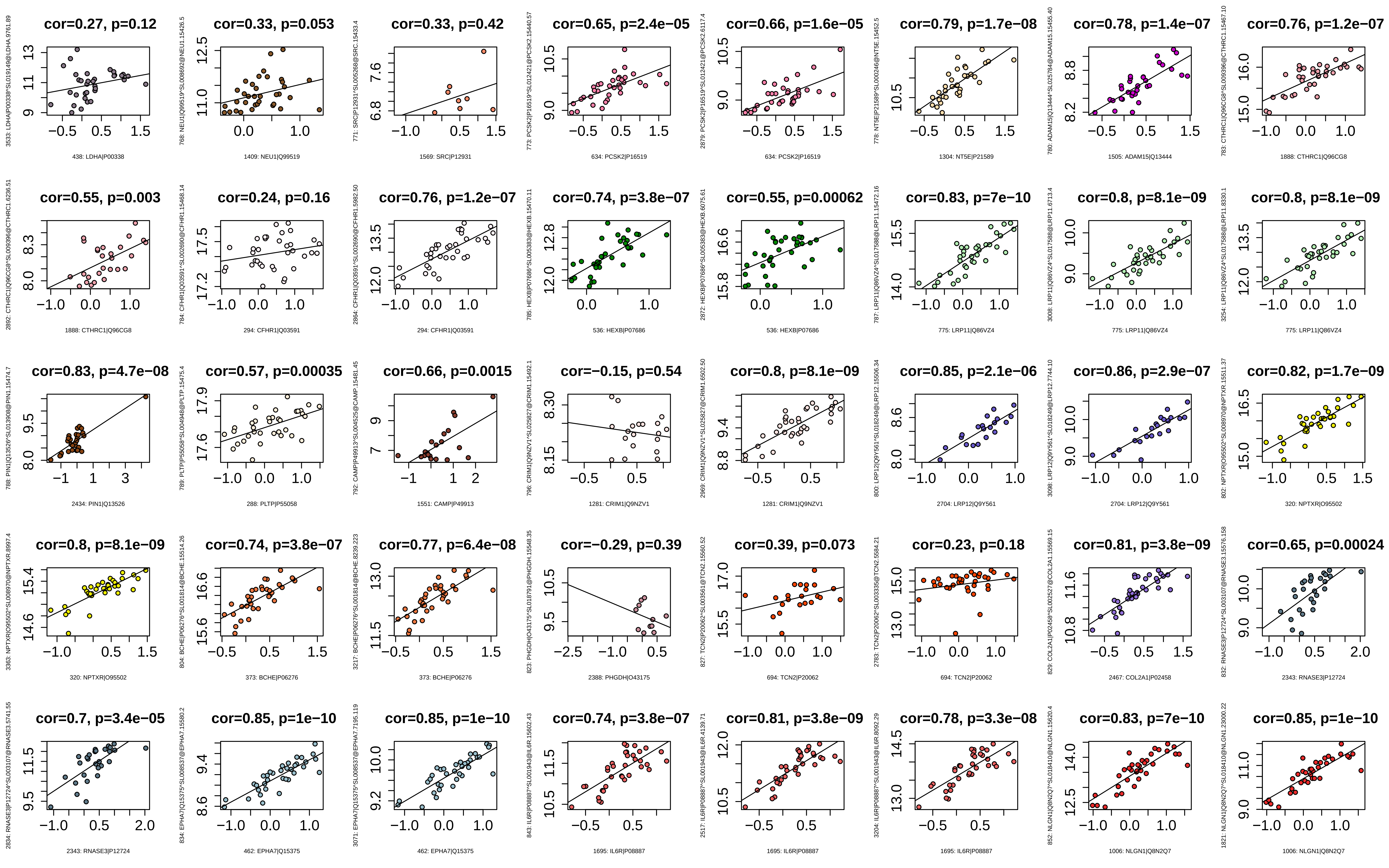
cor=0.27, p=0.12
438: LDHA|P00338
cor=0.33, p=0.053
1409: NEU1|Q99519
cor=0.33, p=0.42
1569: SRC|P12931
cor=0.65, p=2.4e−05
634: PCSK2|P16519
cor=0.66, p=1.6e−05
634: PCSK2|P16519
cor=0.79, p=1.7e−08
1304: NT5E|P21589
cor=0.78, p=1.4e−07
1505: ADAM15|Q13444
cor=0.76, p=1.2e−07
1888: CTHRC1|Q96CG8
cor=0.55, p=0.003
1888: CTHRC1|Q96CG8
cor=0.24, p=0.16
294: CFHR1|Q03591
cor=0.76, p=1.2e−07
294: CFHR1|Q03591
cor=0.74, p=3.8e−07
536: HEXB|P07686
cor=0.55, p=0.00062
536: HEXB|P07686
cor=0.83, p=7e−10
775: LRP11|Q86VZ4
cor=0.8, p=8.1e−09
775: LRP11|Q86VZ4
cor=0.8, p=8.1e−09
775: LRP11|Q86VZ4
cor=0.83, p=4.7e−08
2434: PIN1|Q13526
cor=0.57, p=0.00035
288: PLTP|P55058
cor=0.66, p=0.0015
1551: CAMP|P49913
cor=−0.15, p=0.54
1281: CRIM1|Q9NZV1
cor=0.8, p=8.1e−09
1281: CRIM1|Q9NZV1
cor=0.85, p=2.1e−06
2704: LRP12|Q9Y561
cor=0.86, p=2.9e−07
2704: LRP12|Q9Y561
cor=0.82, p=1.7e−09
320: NPTXR|O95502
cor=0.8, p=8.1e−09
320: NPTXR|O95502
cor=0.74, p=3.8e−07
373: BCHE|P06276
cor=0.77, p=6.4e−08
373: BCHE|P06276
cor=−0.29, p=0.39
2388: PHGDH|O43175
cor=0.39, p=0.073
694: TCN2|P20062
cor=0.23, p=0.18
694: TCN2|P20062
cor=0.81, p=3.8e−09
2467: COL2A1|P02458
cor=0.65, p=0.00024
2343: RNASE3|P12724
cor=0.7, p=3.4e−05
2343: RNASE3|P12724
cor=0.85, p=1e−10
462: EPHA7|Q15375
cor=0.85, p=1e−10
462: EPHA7|Q15375
cor=0.74, p=3.8e−07
1695: IL6R|P08887
cor=0.81, p=3.8e−09
1695: IL6R|P08887
cor=0.78, p=3.3e−08
1695: IL6R|P08887
cor=0.83, p=7e−10
1006: NLGN1|Q8N2Q7
cor=0.85, p=1e−10
1006: NLGN1|Q8N2Q7

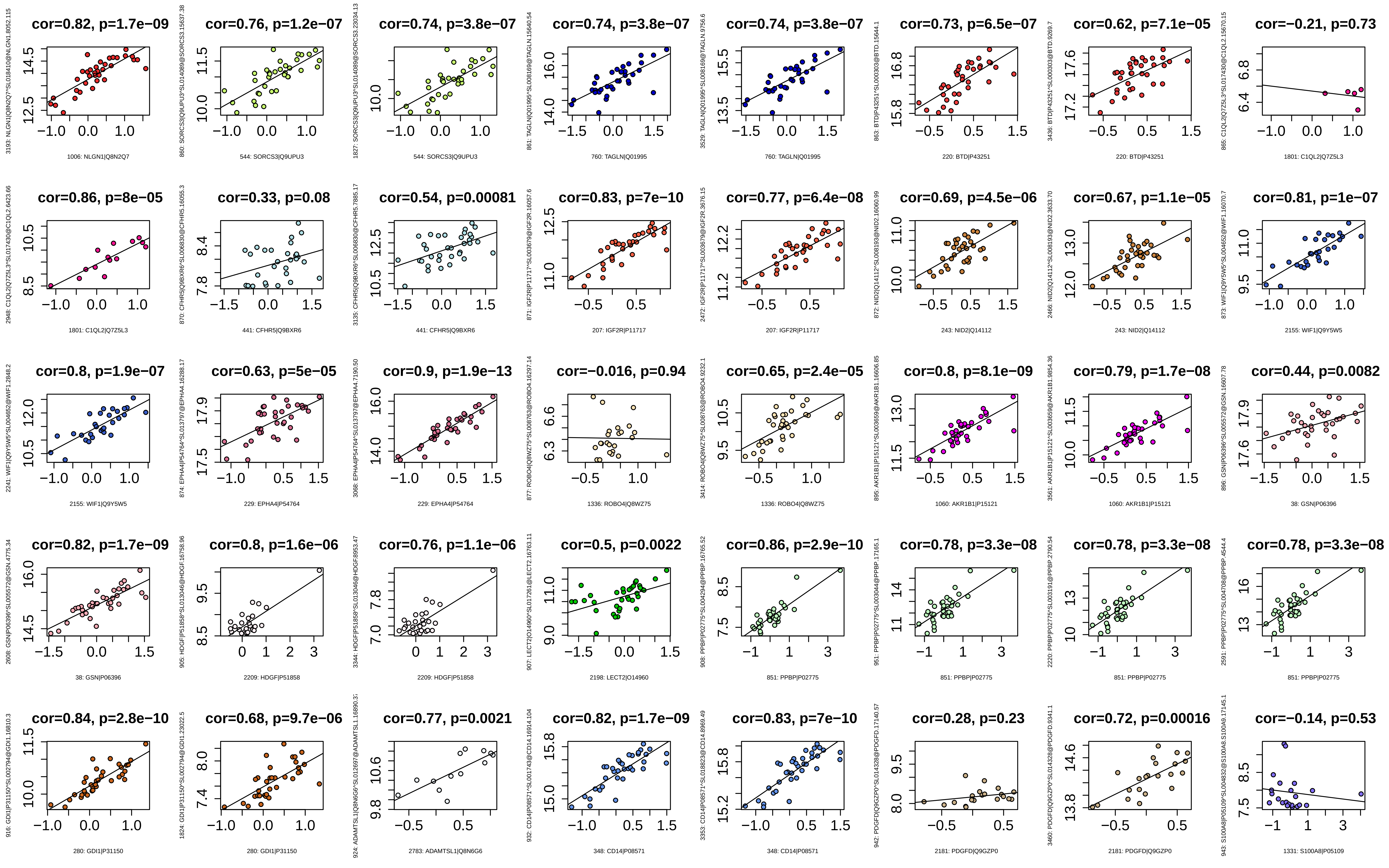
cor=0.82, p=1.7e−09
1006: NLGN1|Q8N2Q7
cor=0.76, p=1.2e−07
544: SORCS3|Q9UPU3
cor=0.74, p=3.8e−07
544: SORCS3|Q9UPU3
cor=0.74, p=3.8e−07
760: TAGLN|Q01995
cor=0.74, p=3.8e−07
760: TAGLN|Q01995
cor=0.73, p=6.5e−07
220: BTD|P43251
cor=0.62, p=7.1e−05
220: BTD|P43251
cor=−0.21, p=0.73
1801: C1QL2|Q7Z5L3
cor=0.86, p=8e−05
1801: C1QL2|Q7Z5L3
cor=0.33, p=0.08
441: CFHR5|Q9BXR6
cor=0.54, p=0.00081
441: CFHR5|Q9BXR6
cor=0.83, p=7e−10
207: IGF2R|P11717
cor=0.77, p=6.4e−08
207: IGF2R|P11717
cor=0.69, p=4.5e−06
243: NID2|Q14112
cor=0.67, p=1.1e−05
243: NID2|Q14112
cor=0.81, p=1e−07
2155: WIF1|Q9Y5W5
cor=0.8, p=1.9e−07
2155: WIF1|Q9Y5W5
cor=0.63, p=5e−05
229: EPHA4|P54764
cor=0.9, p=1.9e−13
229: EPHA4|P54764
cor=−0.016, p=0.94
1336: ROBO4|Q8WZ75
cor=0.65, p=2.4e−05
1336: ROBO4|Q8WZ75
cor=0.8, p=8.1e−09
1060: AKR1B1|P15121
cor=0.79, p=1.7e−08
1060: AKR1B1|P15121
cor=0.44, p=0.0082
38: GSN|P06396
cor=0.82, p=1.7e−09
38: GSN|P06396
cor=0.8, p=1.6e−06
2209: HDGF|P51858
cor=0.76, p=1.1e−06
2209: HDGF|P51858
cor=0.5, p=0.0022
2198: LECT2|O14960
cor=0.86, p=2.9e−10
851: PPBP|P02775
cor=0.78, p=3.3e−08
851: PPBP|P02775
cor=0.78, p=3.3e−08
851: PPBP|P02775
cor=0.78, p=3.3e−08
851: PPBP|P02775
cor=0.84, p=2.8e−10
280: GDI1|P31150
cor=0.68, p=9.7e−06
280: GDI1|P31150
cor=0.77, p=0.0021
2783: ADAMTSL1|Q8N6G6
cor=0.82, p=1.7e−09
348: CD14|P08571
cor=0.83, p=7e−10
348: CD14|P08571
cor=0.28, p=0.23
2181: PDGFD|Q9GZP0
cor=0.72, p=0.00016
2181: PDGFD|Q9GZP0
cor=−0.14, p=0.53
1331: S100A8|P05109

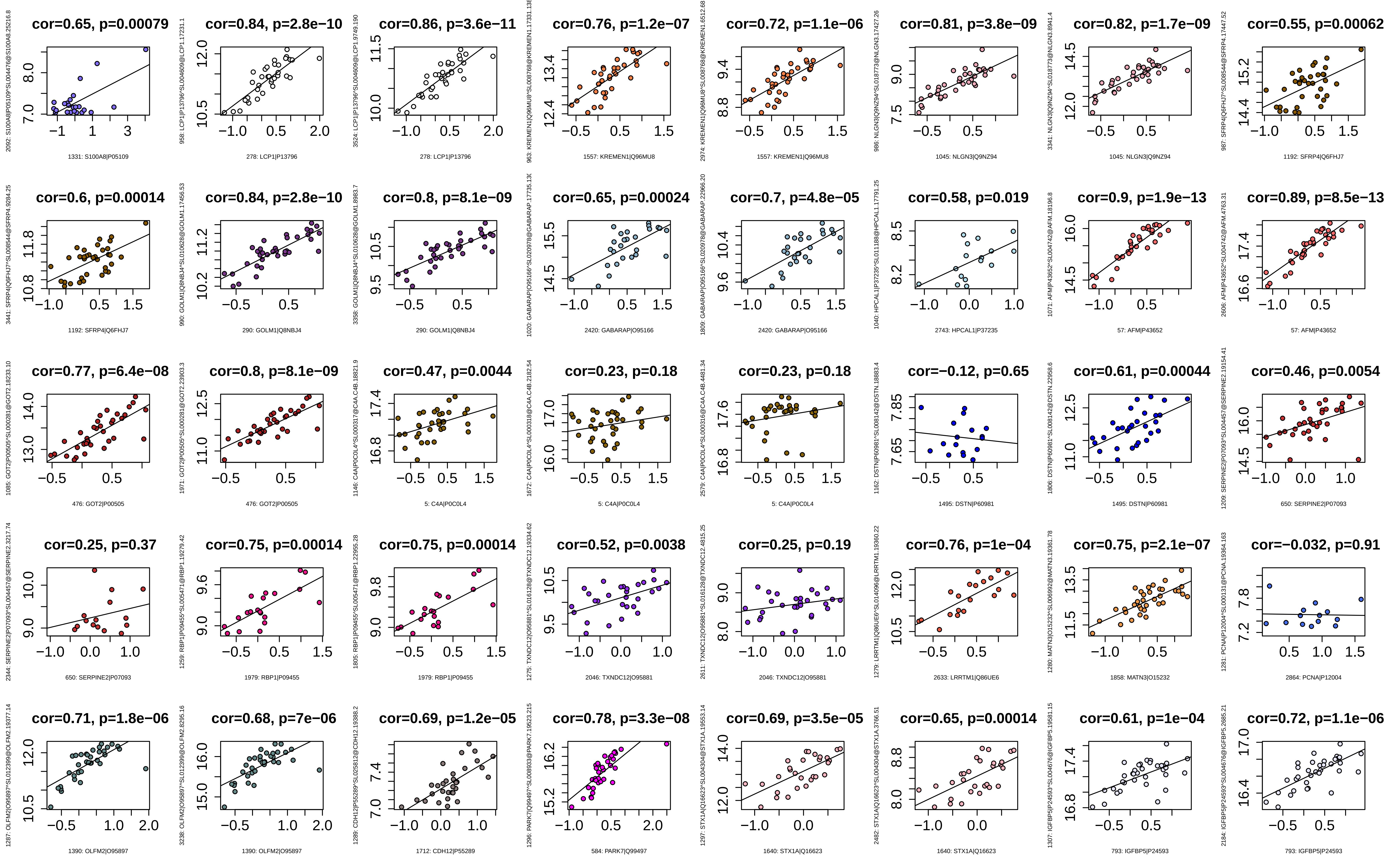
cor=0.65, p=0.00079
1331: S100A8|P05109
cor=0.84, p=2.8e−10
278: LCP1|P13796
cor=0.86, p=3.6e−11
278: LCP1|P13796
cor=0.76, p=1.2e−07
1557: KREMEN1|Q96MU8
cor=0.72, p=1.1e−06
1557: KREMEN1|Q96MU8
cor=0.81, p=3.8e−09
1045: NLGN3|Q9NZ94
cor=0.82, p=1.7e−09
1045: NLGN3|Q9NZ94
cor=0.55, p=0.00062
1192: SFRP4|Q6FHJ7
cor=0.6, p=0.00014
1192: SFRP4|Q6FHJ7
cor=0.84, p=2.8e−10
290: GOLM1|Q8NBJ4
cor=0.8, p=8.1e−09
290: GOLM1|Q8NBJ4
cor=0.65, p=0.00024
2420: GABARAP|O95166
cor=0.7, p=4.8e−05
2420: GABARAP|O95166
cor=0.58, p=0.019
2743: HPCAL1|P37235
cor=0.9, p=1.9e−13
57: AFM|P43652
cor=0.89, p=8.5e−13
57: AFM|P43652
cor=0.77, p=6.4e−08
476: GOT2|P00505
cor=0.8, p=8.1e−09
476: GOT2|P00505
cor=0.47, p=0.0044
5: C4A|P0C0L4
cor=0.23, p=0.18
5: C4A|P0C0L4
cor=0.23, p=0.18
5: C4A|P0C0L4
cor=−0.12, p=0.65
1495: DSTN|P60981
cor=0.61, p=0.00044
1495: DSTN|P60981
cor=0.46, p=0.0054
650: SERPINE2|P07093
cor=0.25, p=0.37
650: SERPINE2|P07093
cor=0.75, p=0.00014
1979: RBP1|P09455
cor=0.75, p=0.00014
1979: RBP1|P09455
cor=0.52, p=0.0038
2046: TXNDC12|O95881
cor=0.25, p=0.19
2046: TXNDC12|O95881
cor=0.76, p=1e−04
2633: LRRTM1|Q86UE6
cor=0.75, p=2.1e−07
1858: MATN3|O15232
cor=−0.032, p=0.91
2864: PCNA|P12004
cor=0.71, p=1.8e−06
1390: OLFM2|O95897
cor=0.68, p=7e−06
1390: OLFM2|O95897
cor=0.69, p=1.2e−05
1712: CDH12|P55289
cor=0.78, p=3.3e−08
584: PARK7|Q99497
cor=0.69, p=3.5e−05
1640: STX1A|Q16623
cor=0.65, p=0.00014
1640: STX1A|Q16623
cor=0.61, p=1e−04
793: IGFBP5|P24593
cor=0.72, p=1.1e−06
793: IGFBP5|P24593

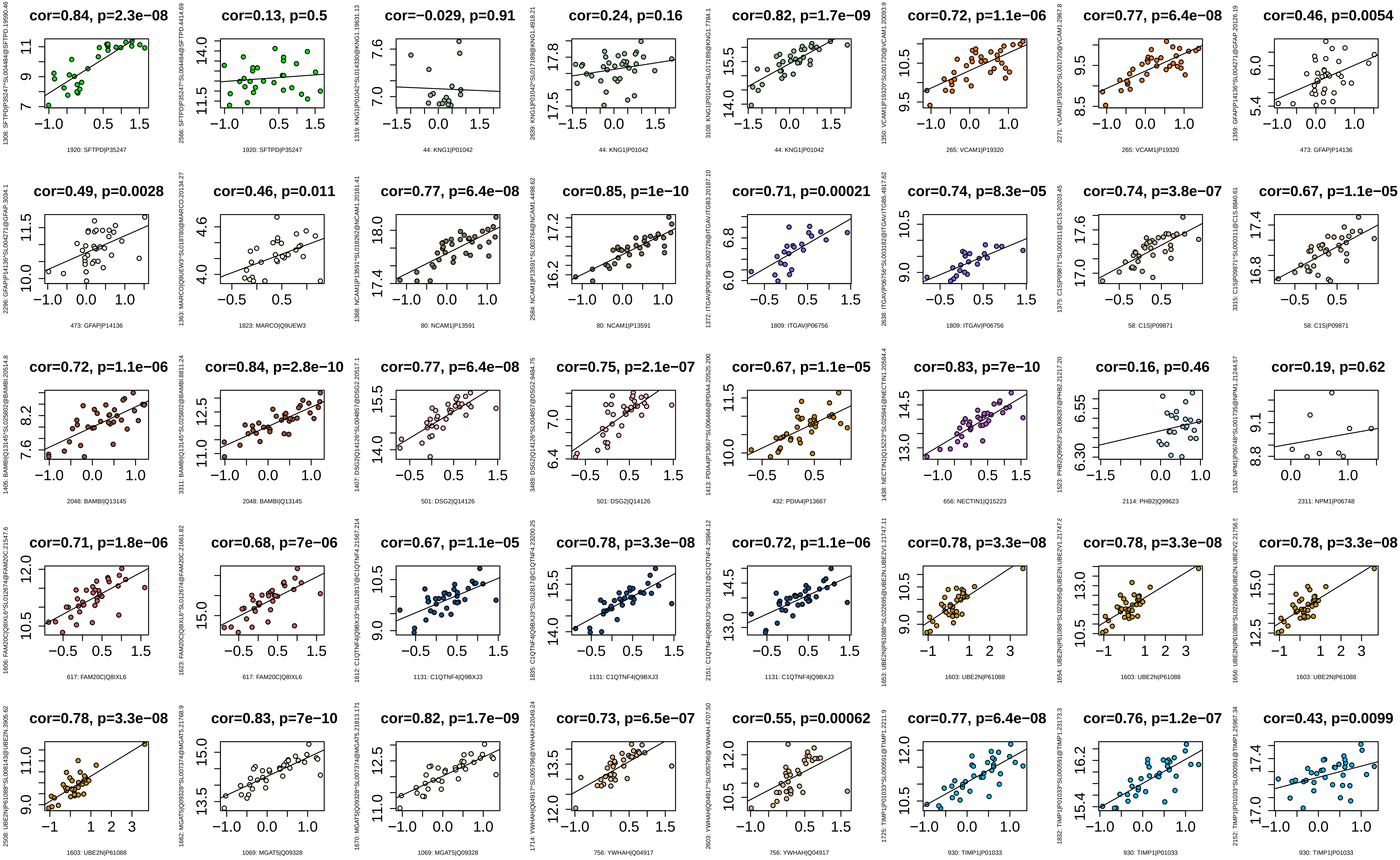
cor=0.84, p=2.3e−08
1920: SFTPD|P35247
cor=0.13, p=0.5
1920: SFTPD|P35247
cor=−0.029, p=0.91
44: KNG1|P01042
cor=0.24, p=0.16
44: KNG1|P01042
cor=0.82, p=1.7e−09
44: KNG1|P01042
cor=0.72, p=1.1e−06
265: VCAM1|P19320
cor=0.77, p=6.4e−08
265: VCAM1|P19320
cor=0.46, p=0.0054
473: GFAP|P14136
cor=0.49, p=0.0028
473: GFAP|P14136
cor=0.46, p=0.011
1823: MARCO|Q9UEW3
cor=0.77, p=6.4e−08
80: NCAM1|P13591
cor=0.85, p=1e−10
80: NCAM1|P13591
cor=0.71, p=0.00021
1809: ITGAV|P06756
cor=0.74, p=8.3e−05
1809: ITGAV|P06756
cor=0.74, p=3.8e−07
58: C1S|P09871
cor=0.67, p=1.1e−05
58: C1S|P09871
cor=0.72, p=1.1e−06
2048: BAMBI|Q13145
cor=0.84, p=2.8e−10
2048: BAMBI|Q13145
cor=0.77, p=6.4e−08
501: DSG2|Q14126
cor=0.75, p=2.1e−07
501: DSG2|Q14126
cor=0.67, p=1.1e−05
432: PDIA4|P13667
cor=0.83, p=7e−10
656: NECTIN1|Q15223
cor=0.16, p=0.46
2114: PHB2|Q99623
cor=0.19, p=0.62
2311: NPM1|P06748
cor=0.71, p=1.8e−06
617: FAM20C|Q8IXL6
cor=0.68, p=7e−06
617: FAM20C|Q8IXL6
cor=0.67, p=1.1e−05
1131: C1QTNF4|Q9BXJ3
cor=0.78, p=3.3e−08
1131: C1QTNF4|Q9BXJ3
cor=0.72, p=1.1e−06
1131: C1QTNF4|Q9BXJ3
cor=0.78, p=3.3e−08
1603: UBE2N|P61088
cor=0.78, p=3.3e−08
1603: UBE2N|P61088
cor=0.78, p=3.3e−08
1603: UBE2N|P61088
cor=0.78, p=3.3e−08
1603: UBE2N|P61088
cor=0.83, p=7e−10
1069: MGAT5|Q09328
cor=0.82, p=1.7e−09
1069: MGAT5|Q09328
cor=0.73, p=6.5e−07
756: YWHAH|Q04917
cor=0.55, p=0.00062
756: YWHAH|Q04917
cor=0.77, p=6.4e−08
930: TIMP1|P01033
cor=0.76, p=1.2e−07
930: TIMP1|P01033
cor=0.43, p=0.0099
930: TIMP1|P01033

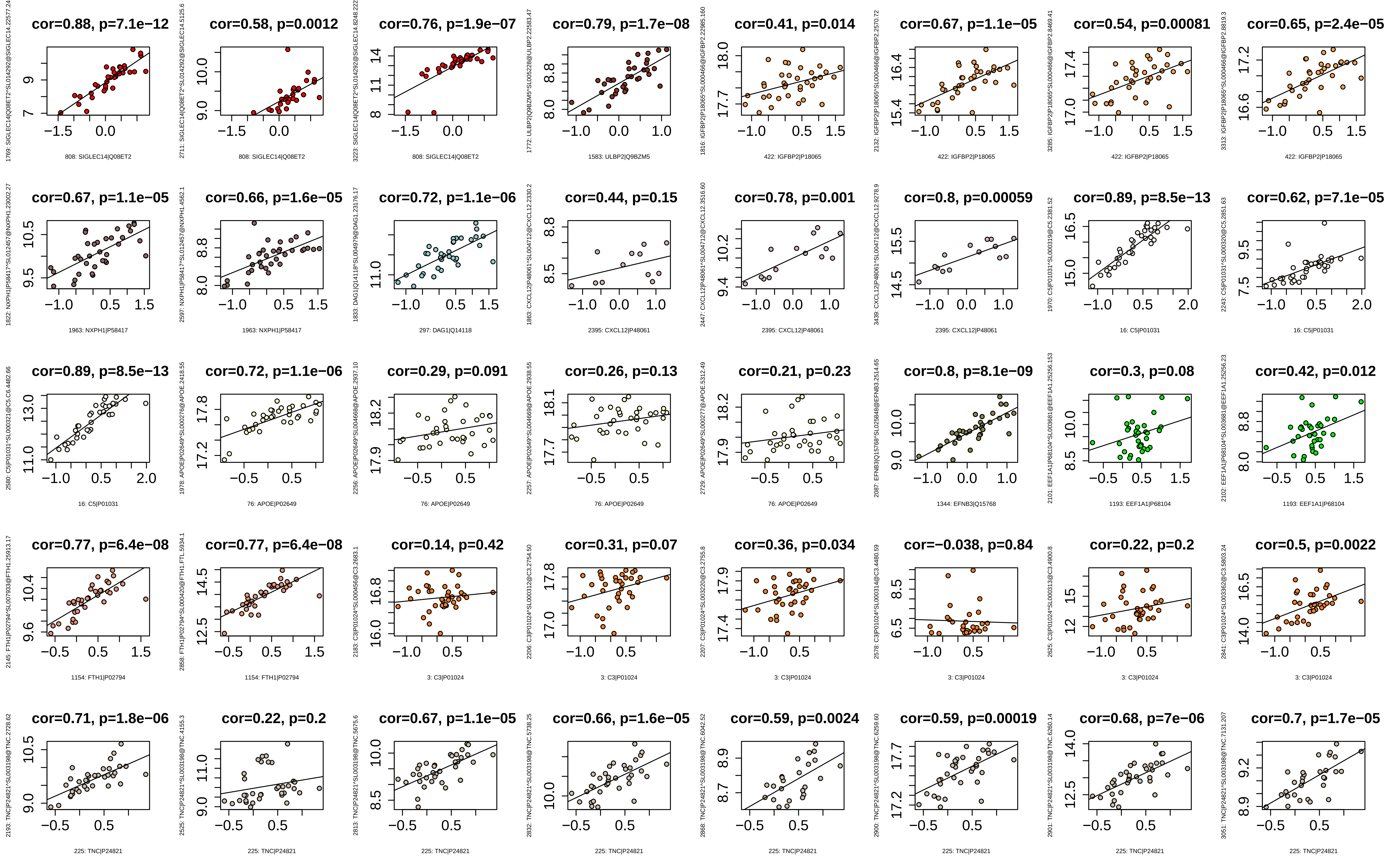
cor=0.88, p=7.1e−12
cor=0.58, p=0.0012
cor=0.76, p=1.9e−07
cor=0.79, p=1.7e−08
cor=0.41, p=0.014
cor=0.67, p=1.1e−05
cor=0.54, p=0.00081
cor=0.65, p=2.4e−05
808: SIGLEC14|Q08ET2
808: SIGLEC14|Q08ET2
808: SIGLEC14|Q08ET2
1583: ULBP2|Q9BZM5
422: IGFBP2|P18065
422: IGFBP2|P18065
422: IGFBP2|P18065
422: IGFBP2|P18065
cor=0.67, p=1.1e−05
cor=0.66, p=1.6e−05
cor=0.72, p=1.1e−06
cor=0.44, p=0.15
cor=0.78, p=0.001
cor=0.8, p=0.00059
cor=0.89, p=8.5e−13
cor=0.62, p=7.1e−05
1963: NXPH1|P58417
1963: NXPH1|P58417
297: DAG1|Q14118
2395: CXCL12|P48061
2395: CXCL12|P48061
2395: CXCL12|P48061
16: C5|P01031
16: C5|P01031
cor=0.89, p=8.5e−13
cor=0.72, p=1.1e−06
cor=0.29, p=0.091
cor=0.26, p=0.13
cor=0.21, p=0.23
cor=0.8, p=8.1e−09
cor=0.3, p=0.08
cor=0.42, p=0.012
16: C5|P01031
76: APOE|P02649
76: APOE|P02649
76: APOE|P02649
76: APOE|P02649
1344: EFNB3|Q15768
1193: EEF1A1|P68104
1193: EEF1A1|P68104
cor=0.77, p=6.4e−08
cor=0.77, p=6.4e−08
cor=0.14, p=0.42
cor=0.31, p=0.07
cor=0.36, p=0.034
cor=−0.038, p=0.84
cor=0.22, p=0.2
cor=0.5, p=0.0022
1154: FTH1|P02794
1154: FTH1|P02794
3: C3|P01024
3: C3|P01024
3: C3|P01024
3: C3|P01024
3: C3|P01024
3: C3|P01024
cor=0.71, p=1.8e−06
cor=0.22, p=0.2
cor=0.67, p=1.1e−05
cor=0.66, p=1.6e−05
cor=0.59, p=0.0024
cor=0.59, p=0.00019
cor=0.68, p=7e−06
cor=0.7, p=1.7e−05
225: TNC|P24821
225: TNC|P24821
225: TNC|P24821
225: TNC|P24821
225: TNC|P24821
225: TNC|P24821
225: TNC|P24821
225: TNC|P24821

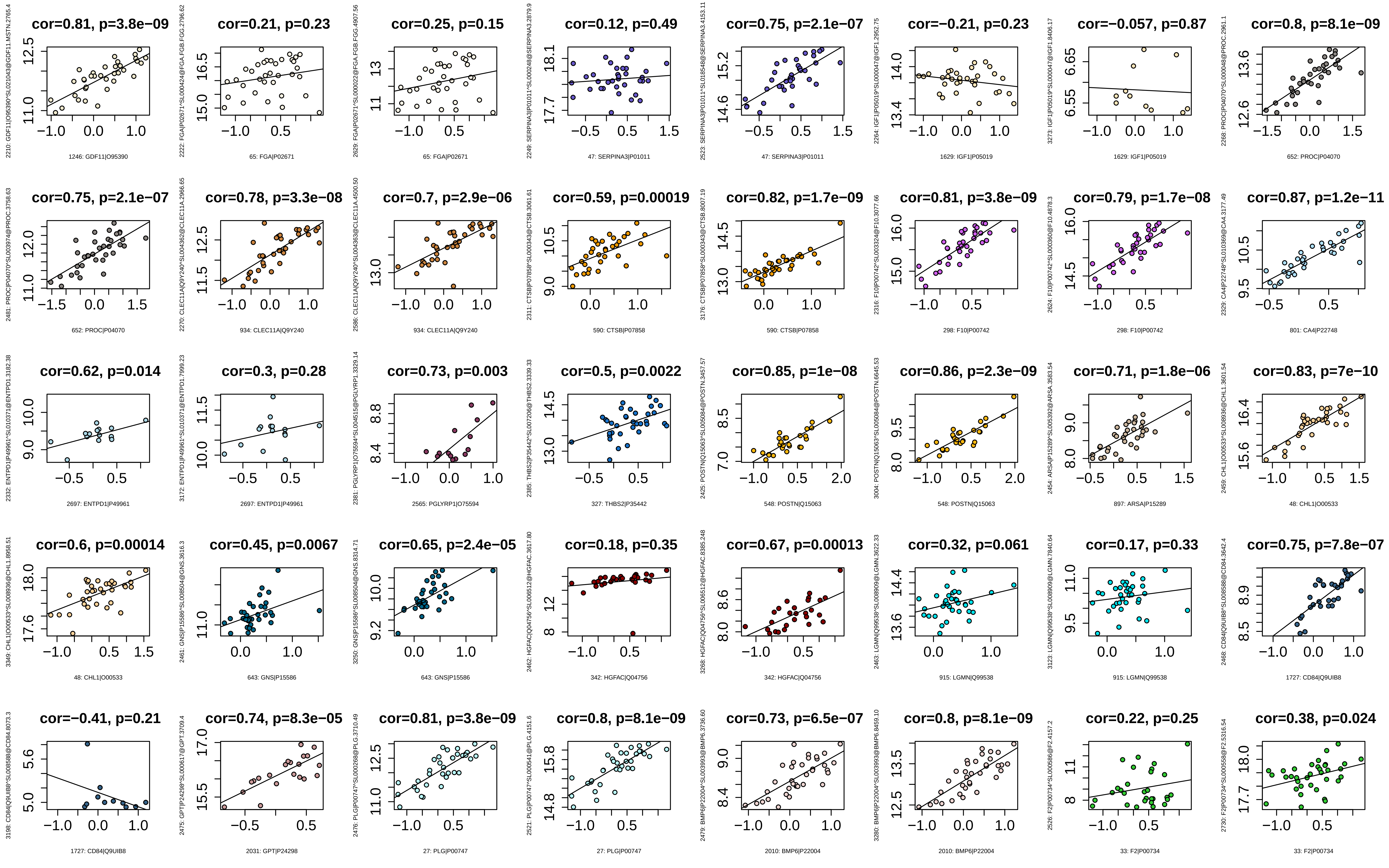
cor=0.81, p=3.8e−09
1246: GDF11|O95390
cor=0.21, p=0.23
65: FGA|P02671
cor=0.25, p=0.15
65: FGA|P02671
cor=0.12, p=0.49
47: SERPINA3|P01011
cor=0.75, p=2.1e−07
47: SERPINA3|P01011
cor=−0.21, p=0.23
1629: IGF1|P05019
cor=−0.057, p=0.87
1629: IGF1|P05019
cor=0.8, p=8.1e−09
652: PROC|P04070
cor=0.75, p=2.1e−07
652: PROC|P04070
cor=0.78, p=3.3e−08
934: CLEC11A|Q9Y240
cor=0.7, p=2.9e−06
934: CLEC11A|Q9Y240
cor=0.59, p=0.00019
590: CTSB|P07858
cor=0.82, p=1.7e−09
590: CTSB|P07858
cor=0.81, p=3.8e−09
298: F10|P00742
cor=0.79, p=1.7e−08
298: F10|P00742
cor=0.87, p=1.2e−11
801: CA4|P22748
cor=0.62, p=0.014
2697: ENTPD1|P49961
cor=0.3, p=0.28
2697: ENTPD1|P49961
cor=0.73, p=0.003
2565: PGLYRP1|O75594
cor=0.5, p=0.0022
327: THBS2|P35442
cor=0.85, p=1e−08
548: POSTN|Q15063
cor=0.86, p=2.3e−09
548: POSTN|Q15063
cor=0.71, p=1.8e−06
897: ARSA|P15289
cor=0.83, p=7e−10
48: CHL1|O00533
cor=0.6, p=0.00014
48: CHL1|O00533
cor=0.45, p=0.0067
643: GNS|P15586
cor=0.65, p=2.4e−05
643: GNS|P15586
cor=0.18, p=0.35
342: HGFAC|Q04756
cor=0.67, p=0.00013
342: HGFAC|Q04756
cor=0.32, p=0.061
915: LGMN|Q99538
cor=0.17, p=0.33
915: LGMN|Q99538
cor=0.75, p=7.8e−07
1727: CD84|Q9UIB8
cor=−0.41, p=0.21
1727: CD84|Q9UIB8
cor=0.74, p=8.3e−05
2031: GPT|P24298
cor=0.81, p=3.8e−09
27: PLG|P00747
cor=0.8, p=8.1e−09
27: PLG|P00747
cor=0.73, p=6.5e−07
2010: BMP6|P22004
cor=0.8, p=8.1e−09
2010: BMP6|P22004
cor=0.22, p=0.25
33: F2|P00734
cor=0.38, p=0.024
33: F2|P00734

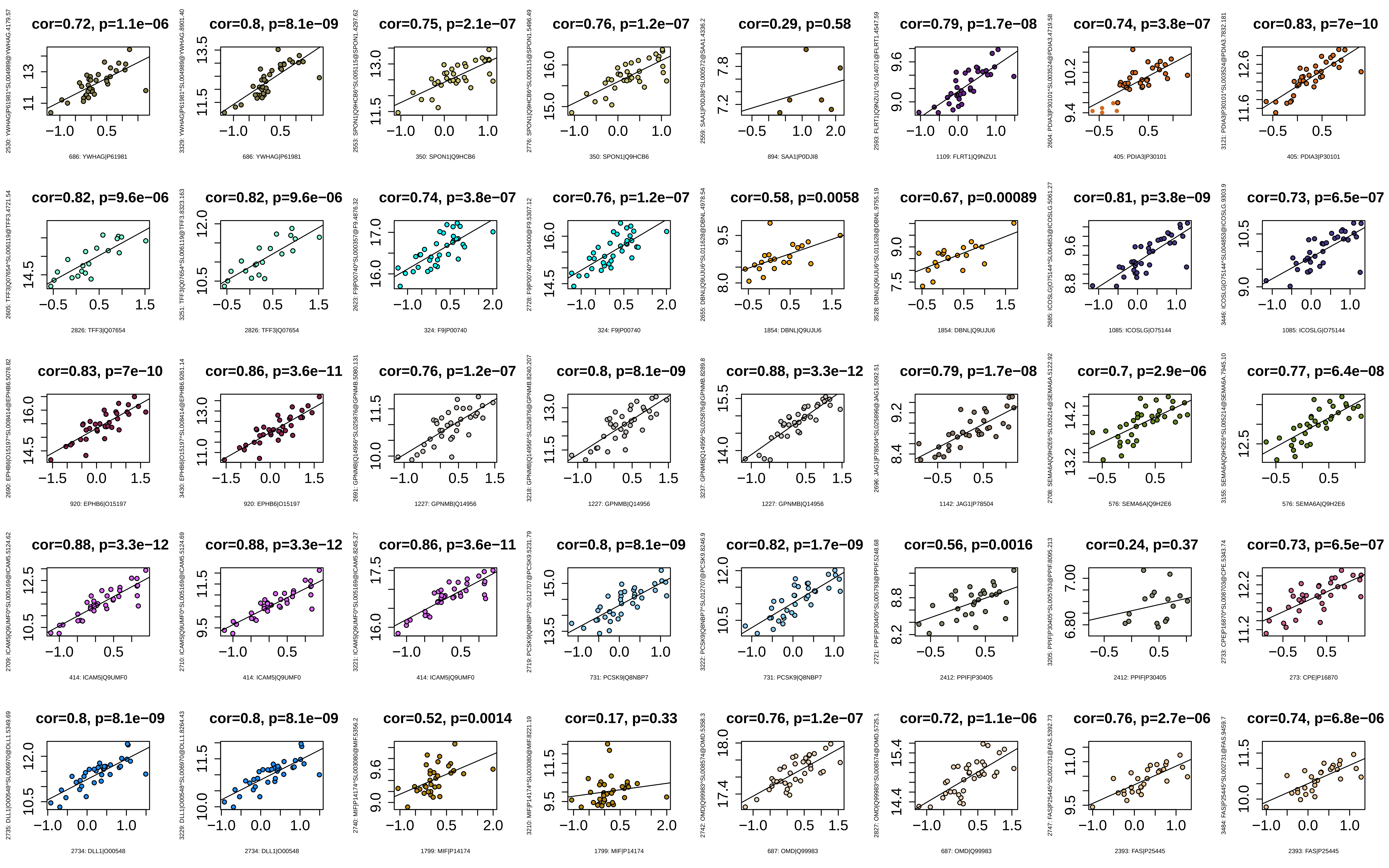
cor=0.72, p=1.1e−06
686: YWHAG|P61981
cor=0.8, p=8.1e−09
686: YWHAG|P61981
cor=0.75, p=2.1e−07
350: SPON1|Q9HCB6
cor=0.76, p=1.2e−07
350: SPON1|Q9HCB6
cor=0.29, p=0.58
894: SAA1|P0DJI8
cor=0.79, p=1.7e−08
1109: FLRT1|Q9NZU1
cor=0.74, p=3.8e−07
405: PDIA3|P30101
cor=0.83, p=7e−10
405: PDIA3|P30101
cor=0.82, p=9.6e−06
2826: TFF3|Q07654
cor=0.82, p=9.6e−06
2826: TFF3|Q07654
cor=0.74, p=3.8e−07
324: F9|P00740
cor=0.76, p=1.2e−07
324: F9|P00740
cor=0.58, p=0.0058
1854: DBNL|Q9UJU6
cor=0.67, p=0.00089
1854: DBNL|Q9UJU6
cor=0.81, p=3.8e−09
1085: ICOSLG|O75144
cor=0.73, p=6.5e−07
1085: ICOSLG|O75144
cor=0.83, p=7e−10
920: EPHB6|O15197
cor=0.86, p=3.6e−11
920: EPHB6|O15197
cor=0.76, p=1.2e−07
1227: GPNMB|Q14956
cor=0.8, p=8.1e−09
1227: GPNMB|Q14956
cor=0.88, p=3.3e−12
1227: GPNMB|Q14956
cor=0.79, p=1.7e−08
1142: JAG1|P78504
cor=0.7, p=2.9e−06
576: SEMA6A|Q9H2E6
cor=0.77, p=6.4e−08
576: SEMA6A|Q9H2E6
cor=0.88, p=3.3e−12
414: ICAM5|Q9UMF0
cor=0.88, p=3.3e−12
414: ICAM5|Q9UMF0
cor=0.86, p=3.6e−11
414: ICAM5|Q9UMF0
cor=0.8, p=8.1e−09
731: PCSK9|Q8NBP7
cor=0.82, p=1.7e−09
731: PCSK9|Q8NBP7
cor=0.56, p=0.0016
2412: PPIF|P30405
cor=0.24, p=0.37
2412: PPIF|P30405
cor=0.73, p=6.5e−07
273: CPE|P16870
cor=0.8, p=8.1e−09
2734: DLL1|O00548
cor=0.8, p=8.1e−09
2734: DLL1|O00548
cor=0.52, p=0.0014
1799: MIF|P14174
cor=0.17, p=0.33
1799: MIF|P14174
cor=0.76, p=1.2e−07
687: OMD|Q99983
cor=0.72, p=1.1e−06
687: OMD|Q99983
cor=0.76, p=2.7e−06
2393: FAS|P25445
cor=0.74, p=6.8e−06
2393: FAS|P25445

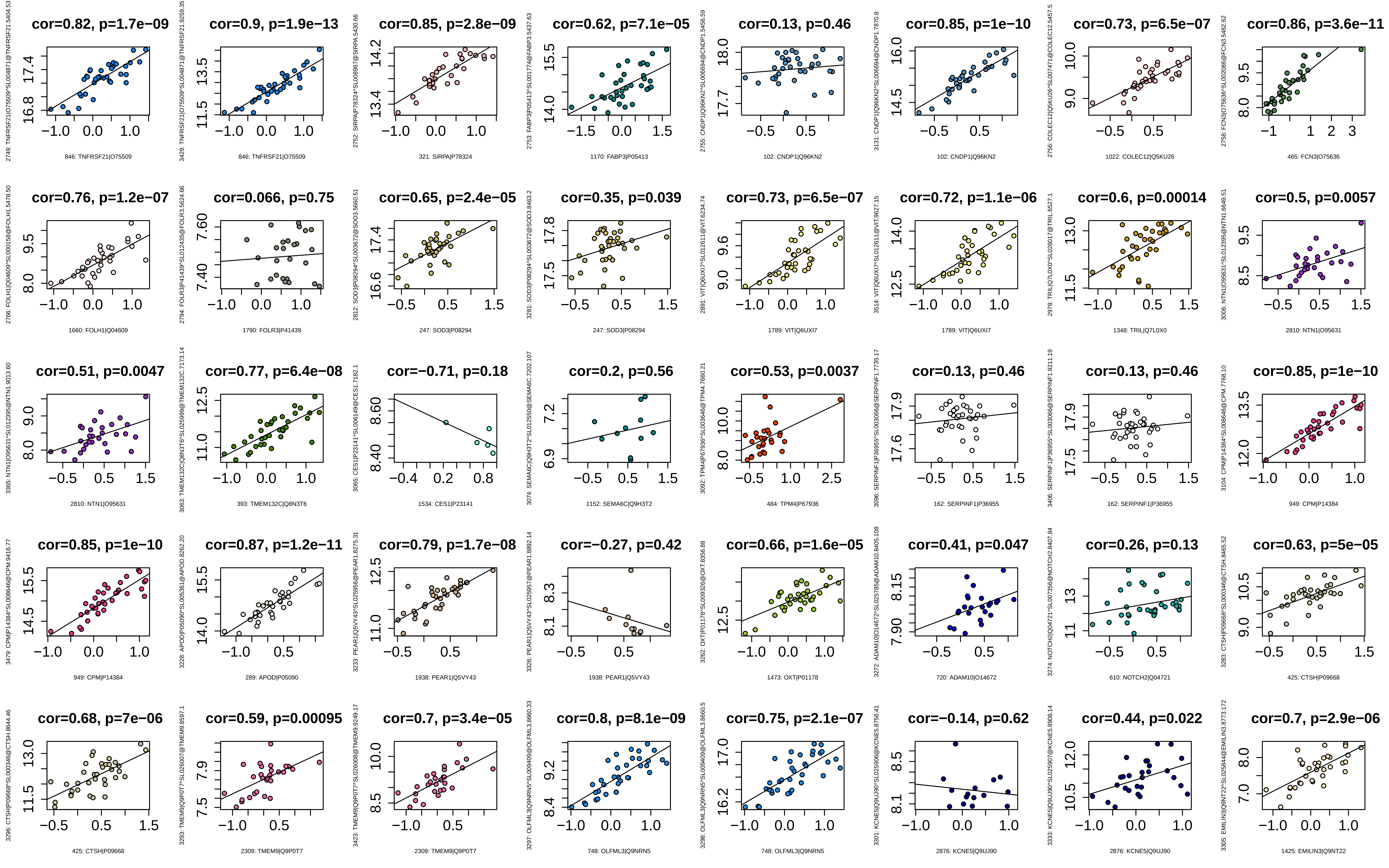
cor=0.82, p=1.7e−09
846: TNFRSF21|O75509
cor=0.9, p=1.9e−13
846: TNFRSF21|O75509
cor=0.85, p=2.8e−09
321: SIRPA|P78324
cor=0.62, p=7.1e−05
1170: FABP3|P05413
cor=0.13, p=0.46
102: CNDP1|Q96KN2
cor=0.85, p=1e−10
102: CNDP1|Q96KN2
cor=0.73, p=6.5e−07
1022: COLEC12|Q5KU26
cor=0.86, p=3.6e−11
465: FCN3|O75636
cor=0.76, p=1.2e−07
1660: FOLH1|Q04609
cor=0.066, p=0.75
1790: FOLR3|P41439
cor=0.65, p=2.4e−05
247: SOD3|P08294
cor=0.35, p=0.039
247: SOD3|P08294
cor=0.73, p=6.5e−07
1789: VIT|Q6UXI7
cor=0.72, p=1.1e−06
1789: VIT|Q6UXI7
cor=0.6, p=0.00014
1348: TRIL|Q7L0X0
cor=0.5, p=0.0057
2810: NTN1|O95631
cor=0.51, p=0.0047
2810: NTN1|O95631
cor=0.77, p=6.4e−08
393: TMEM132C|Q8N3T6
cor=−0.71, p=0.18
1534: CES1|P23141
cor=0.2, p=0.56
1152: SEMA6C|Q9H3T2
cor=0.53, p=0.0037
484: TPM4|P67936
cor=0.13, p=0.46
162: SERPINF1|P36955
cor=0.13, p=0.46
162: SERPINF1|P36955
cor=0.85, p=1e−10
949: CPM|P14384
cor=0.85, p=1e−10
949: CPM|P14384
cor=0.87, p=1.2e−11
289: APOD|P05090
cor=0.79, p=1.7e−08
1938: PEAR1|Q5VY43
cor=−0.27, p=0.42
1938: PEAR1|Q5VY43
cor=0.66, p=1.6e−05
1473: OXT|P01178
cor=0.41, p=0.047
720: ADAM10|O14672
cor=0.26, p=0.13
610: NOTCH2|Q04721
cor=0.63, p=5e−05
425: CTSH|P09668
cor=0.68, p=7e−06
425: CTSH|P09668
cor=0.59, p=0.00095
2309: TMEM9|Q9P0T7
cor=0.7, p=3.4e−05
2309: TMEM9|Q9P0T7
cor=0.8, p=8.1e−09
748: OLFML3|Q9NRN5
cor=0.75, p=2.1e−07
748: OLFML3|Q9NRN5
cor=−0.14, p=0.62
2876: KCNE5|Q9UJ90
cor=0.44, p=0.022
2876: KCNE5|Q9UJ90
cor=0.7, p=2.9e−06
1425: EMILIN3|Q9NT22

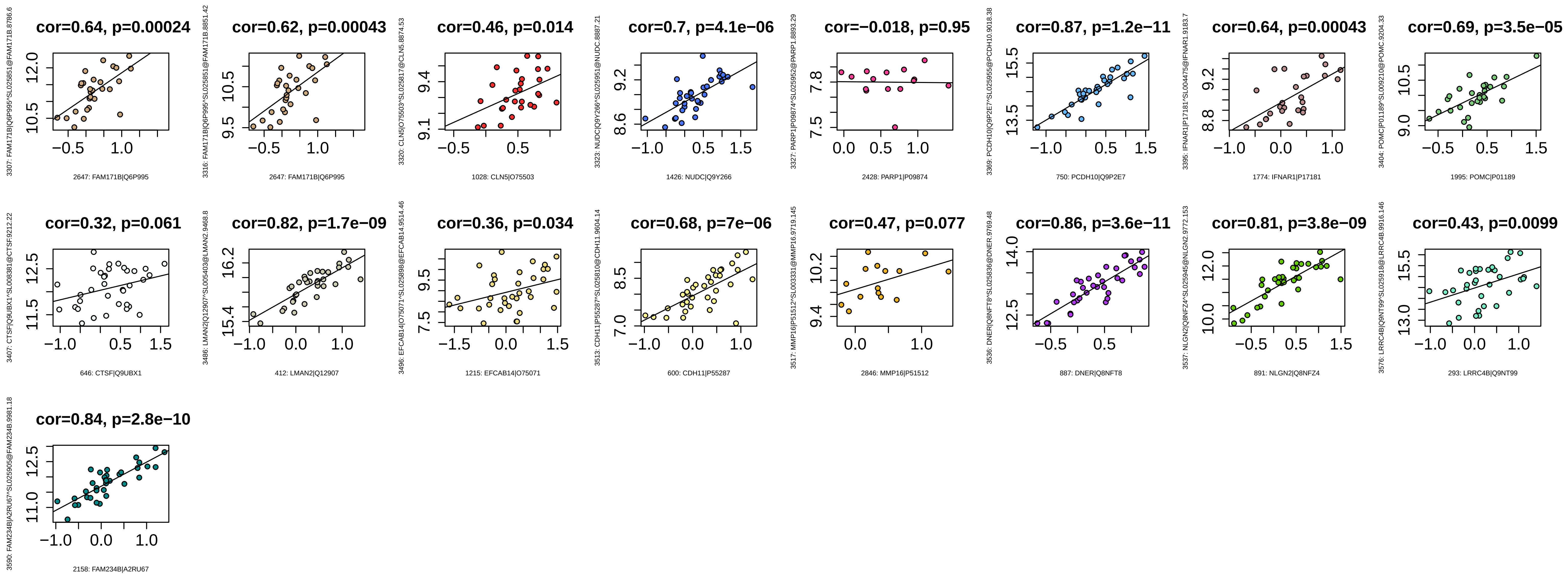
cor=0.64, p=0.00024
3307: FAM171B|Q6P995^SL025851@FAM171B.8786.6
2647: FAM171B|Q6P995
cor=0.62, p=0.00043
3316: FAM171B|Q6P995^SL025851@FAM171B.8851.42
2647: FAM171B|Q6P995
cor=0.46, p=0.014
3320: CLN5|O75503^SL025817@CLN5.8874.53
1028: CLN5|O75503
cor=0.7, p=4.1e−06
3323: NUDC|Q9Y266^SL025951@NUDC.8887.21
1426: NUDC|Q9Y266
cor=−0.018, p=0.95
3327: PARP1|P09874^SL025952@PARP1.8893.29
2428: PARP1|P09874
cor=0.87, p=1.2e−11
3369: PCDH10|Q9P2E7^SL025955@PCDH10.9018.38
750: PCDH10|Q9P2E7
cor=0.64, p=0.00043
3395: IFNAR1|P17181^SL004475@IFNAR1.9183.7
1774: IFNAR1|P17181
cor=0.69, p=3.5e−05
3404: POMC|P01189^SL009210@POMC.9204.33
1995: POMC|P01189
cor=0.32, p=0.061
3407: CTSF|Q9UBX1^SL008381@CTSF.9212.22
646: CTSF|Q9UBX1
cor=0.82, p=1.7e−09
3486: LMAN2|Q12907^SL005403@LMAN2.9468.8
412: LMAN2|Q12907
cor=0.36, p=0.034
3496: EFCAB14|O75071^SL025898@EFCAB14.9514.46
1215: EFCAB14|O75071
cor=0.68, p=7e−06
3513: CDH11|P55287^SL025810@CDH11.9604.14
600: CDH11|P55287
cor=0.47, p=0.077
3517: MMP16|P51512^SL003331@MMP16.9719.145
2846: MMP16|P51512
cor=0.86, p=3.6e−11
3536: DNER|Q8NFT8^SL025836@DNER.9769.48
887: DNER|Q8NFT8
cor=0.81, p=3.8e−09
3537: NLGN2|Q8NFZ4^SL025945@NLGN2.9772.153
891: NLGN2|Q8NFZ4
cor=0.43, p=0.0099
3576: LRRC4B|Q9NT99^SL025918@LRRC4B.9916.146
293: LRRC4B|Q9NT99
cor=0.84, p=2.8e−10
3590: FAM234B|A2RU67^SL025905@FAM234B.9981.18
2158: FAM234B|A2RU67

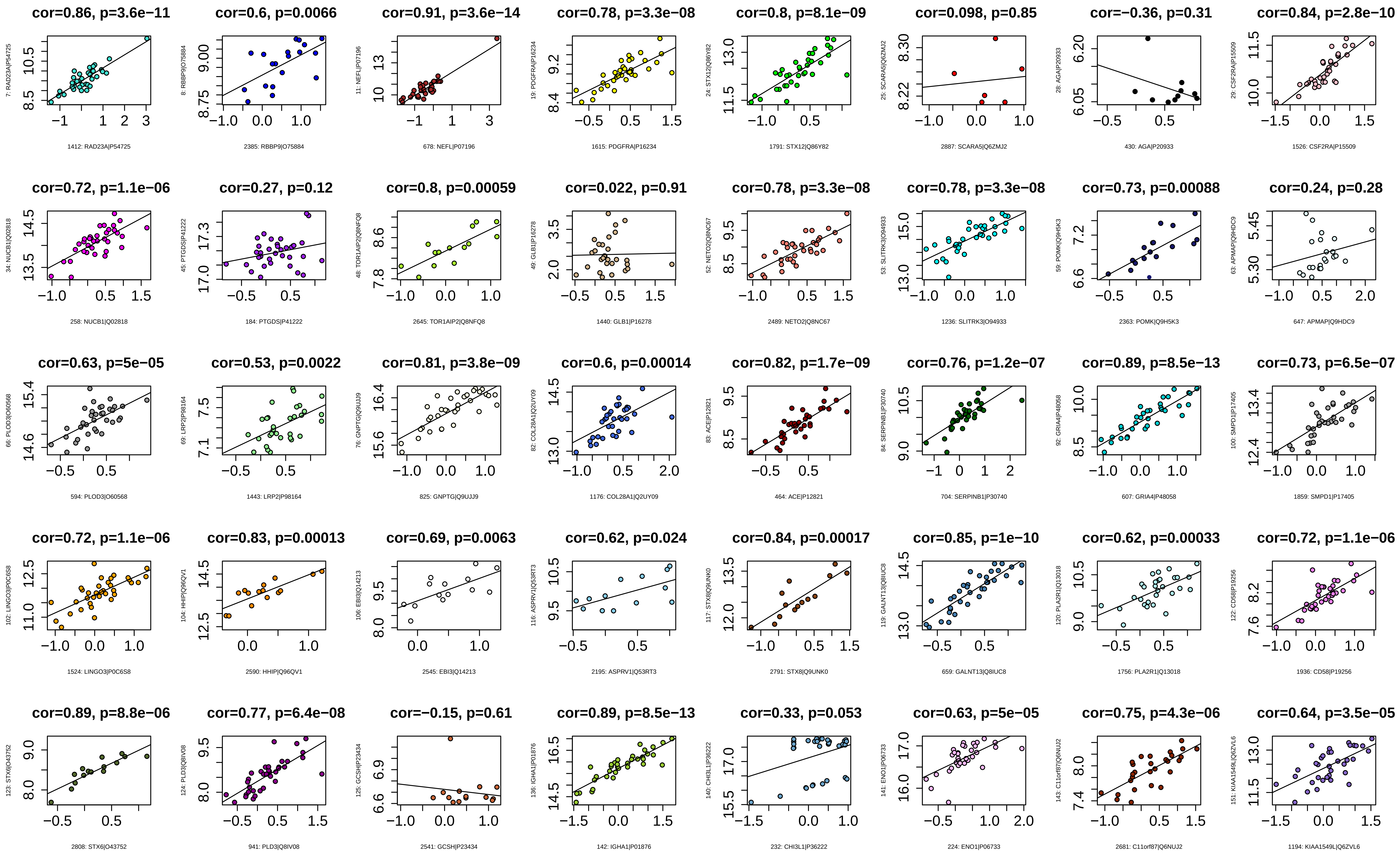
cor=0.86, p=3.6e−11
1412: RAD23A|P54725
cor=0.6, p=0.0066
2385: RBBP9|O75884
cor=0.91, p=3.6e−14
678: NEFL|P07196
cor=0.78, p=3.3e−08
1615: PDGFRA|P16234
cor=0.8, p=8.1e−09
1791: STX12|Q86Y82
cor=0.098, p=0.85
2887: SCARA5|Q6ZMJ2
cor=−0.36, p=0.31
430: AGA|P20933
cor=0.84, p=2.8e−10
1526: CSF2RA|P15509
cor=0.72, p=1.1e−06
258: NUCB1|Q02818
cor=0.27, p=0.12
184: PTGDS|P41222
cor=0.8, p=0.00059
2645: TOR1AIP2|Q8NFQ8
cor=0.022, p=0.91
1440: GLB1|P16278
cor=0.78, p=3.3e−08
2489: NETO2|Q8NC67
cor=0.78, p=3.3e−08
1236: SLITRK3|O94933
cor=0.73, p=0.00088
2363: POMK|Q9H5K3
cor=0.24, p=0.28
647: APMAP|Q9HDC9
cor=0.63, p=5e−05
594: PLOD3|O60568
cor=0.53, p=0.0022
1443: LRP2|P98164
cor=0.81, p=3.8e−09
825: GNPTG|Q9UJJ9
cor=0.6, p=0.00014
1176: COL28A1|Q2UY09
cor=0.82, p=1.7e−09
464: ACE|P12821
cor=0.76, p=1.2e−07
704: SERPINB1|P30740
cor=0.89, p=8.5e−13
607: GRIA4|P48058
cor=0.73, p=6.5e−07
1859: SMPD1|P17405
cor=0.72, p=1.1e−06
1524: LINGO3|P0C6S8
cor=0.83, p=0.00013
2590: HHIP|Q96QV1
cor=0.69, p=0.0063
2545: EBI3|Q14213
cor=0.62, p=0.024
2195: ASPRV1|Q53RT3
cor=0.84, p=0.00017
2791: STX8|Q9UNK0
cor=0.85, p=1e−10
659: GALNT13|Q8IUC8
cor=0.62, p=0.00033
1756: PLA2R1|Q13018
cor=0.72, p=1.1e−06
1936: CD58|P19256
cor=0.89, p=8.8e−06
2808: STX6|O43752
cor=0.77, p=6.4e−08
941: PLD3|Q8IV08
cor=−0.15, p=0.61
2541: GCSH|P23434
cor=0.89, p=8.5e−13
142: IGHA1|P01876
cor=0.33, p=0.053
232: CHI3L1|P36222
cor=0.63, p=5e−05
224: ENO1|P06733
cor=0.75, p=4.3e−06
2681: C11orf87|Q6NUJ2
cor=0.64, p=3.5e−05
1194: KIAA1549L|Q6ZVL6

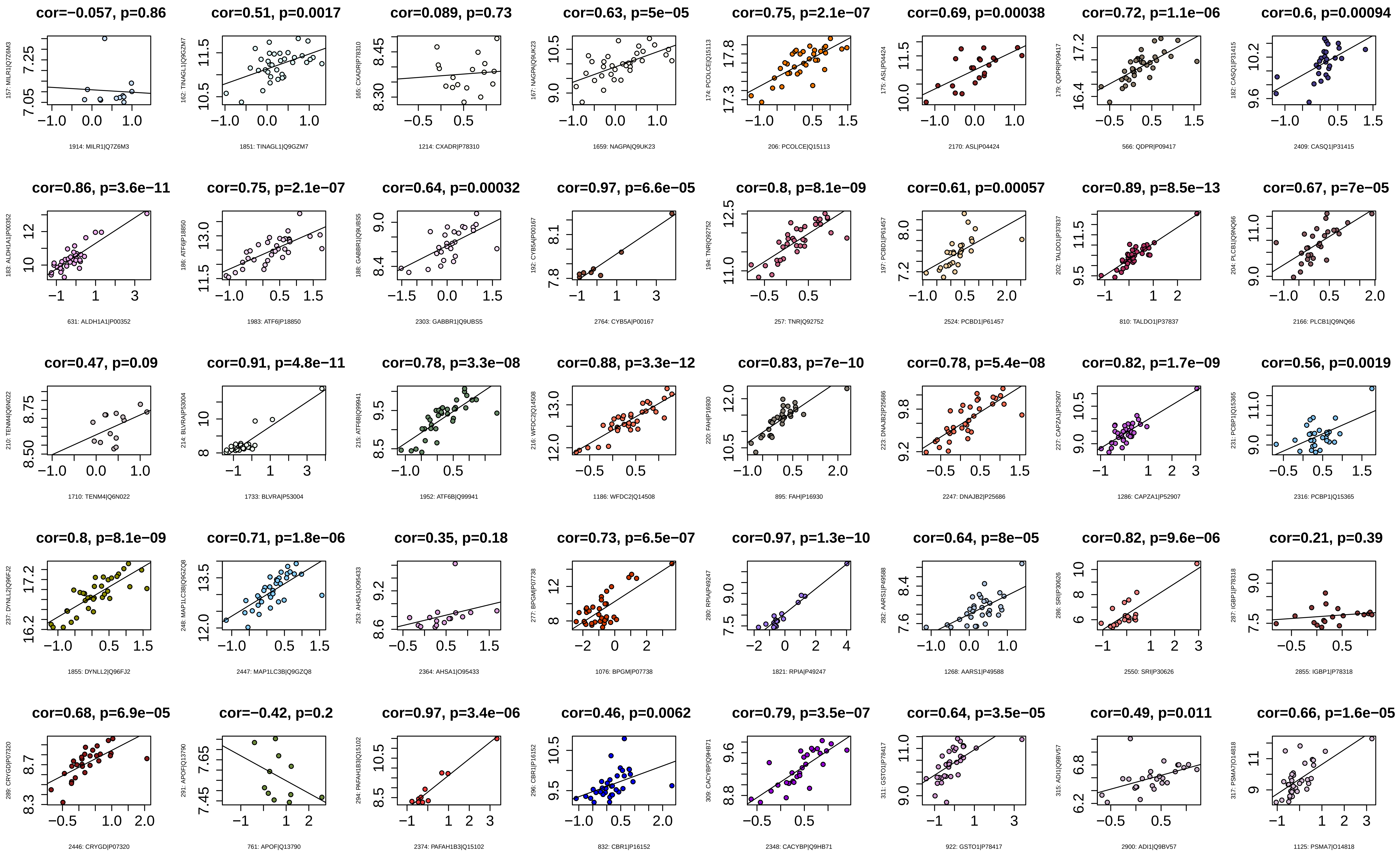
cor=−0.057, p=0.86
157: MILR1|Q7Z6M3
1914: MILR1|Q7Z6M3
cor=0.51, p=0.0017
162: TINAGL1|Q9GZM7
1851: TINAGL1|Q9GZM7
cor=0.089, p=0.73
165: CXADR|P78310
1214: CXADR|P78310
cor=0.63, p=5e−05
167: NAGPA|Q9UK23
1659: NAGPA|Q9UK23
cor=0.75, p=2.1e−07
174: PCOLCE|Q15113
206: PCOLCE|Q15113
cor=0.69, p=0.00038
175: ASL|P04424
2170: ASL|P04424
cor=0.72, p=1.1e−06
179: QDPR|P09417
566: QDPR|P09417
cor=0.6, p=0.00094
182: CASQ1|P31415
2409: CASQ1|P31415
cor=0.86, p=3.6e−11
183: ALDH1A1|P00352
631: ALDH1A1|P00352
cor=0.75, p=2.1e−07
186: ATF6|P18850
1983: ATF6|P18850
cor=0.64, p=0.00032
188: GABBR1|Q9UBS5
2303: GABBR1|Q9UBS5
cor=0.97, p=6.6e−05
192: CYB5A|P00167
2764: CYB5A|P00167
cor=0.8, p=8.1e−09
194: TNR|Q92752
257: TNR|Q92752
cor=0.61, p=0.00057
197: PCBD1|P61457
2524: PCBD1|P61457
cor=0.89, p=8.5e−13
202: TALDO1|P37837
810: TALDO1|P37837
cor=0.67, p=7e−05
204: PLCB1|Q9NQ66
2166: PLCB1|Q9NQ66
cor=0.47, p=0.09
210: TENM4|Q6N022
1710: TENM4|Q6N022
cor=0.91, p=4.8e−11
214: BLVRA|P53004
1733: BLVRA|P53004
cor=0.78, p=3.3e−08
215: ATF6B|Q99941
1952: ATF6B|Q99941
cor=0.88, p=3.3e−12
216: WFDC2|Q14508
1186: WFDC2|Q14508
cor=0.83, p=7e−10
220: FAH|P16930
895: FAH|P16930
cor=0.78, p=5.4e−08
223: DNAJB2|P25686
2247: DNAJB2|P25686
cor=0.82, p=1.7e−09
227: CAPZA1|P52907
1286: CAPZA1|P52907
cor=0.56, p=0.0019
231: PCBP1|Q15365
2316: PCBP1|Q15365
cor=0.8, p=8.1e−09
237: DYNLL2|Q96FJ2
1855: DYNLL2|Q96FJ2
cor=0.71, p=1.8e−06
248: MAP1LC3B|Q9GZQ8
2447: MAP1LC3B|Q9GZQ8
cor=0.35, p=0.18
253: AHSA1|O95433
2364: AHSA1|O95433
cor=0.73, p=6.5e−07
277: BPGM|P07738
1076: BPGM|P07738
cor=0.97, p=1.3e−10
280: RPIA|P49247
1821: RPIA|P49247
cor=0.64, p=8e−05
282: AARS1|P49588
1268: AARS1|P49588
cor=0.82, p=9.6e−06
286: SRI|P30626
2550: SRI|P30626
cor=0.21, p=0.39
287: IGBP1|P78318
2855: IGBP1|P78318
cor=0.68, p=6.9e−05
289: CRYGD|P07320
2446: CRYGD|P07320
cor=−0.42, p=0.2
291: APOF|Q13790
761: APOF|Q13790
cor=0.97, p=3.4e−06
294: PAFAH1B3|Q15102
2374: PAFAH1B3|Q15102
cor=0.46, p=0.0062
296: CBR1|P16152
832: CBR1|P16152
cor=0.79, p=3.5e−07
309: CACYBP|Q9HB71
2348: CACYBP|Q9HB71
cor=0.64, p=3.5e−05
311: GSTO1|P78417
922: GSTO1|P78417
cor=0.49, p=0.011
315: ADI1|Q9BV57
2900: ADI1|Q9BV57
cor=0.66, p=1.6e−05
317: PSMA7|O14818
1125: PSMA7|O14818

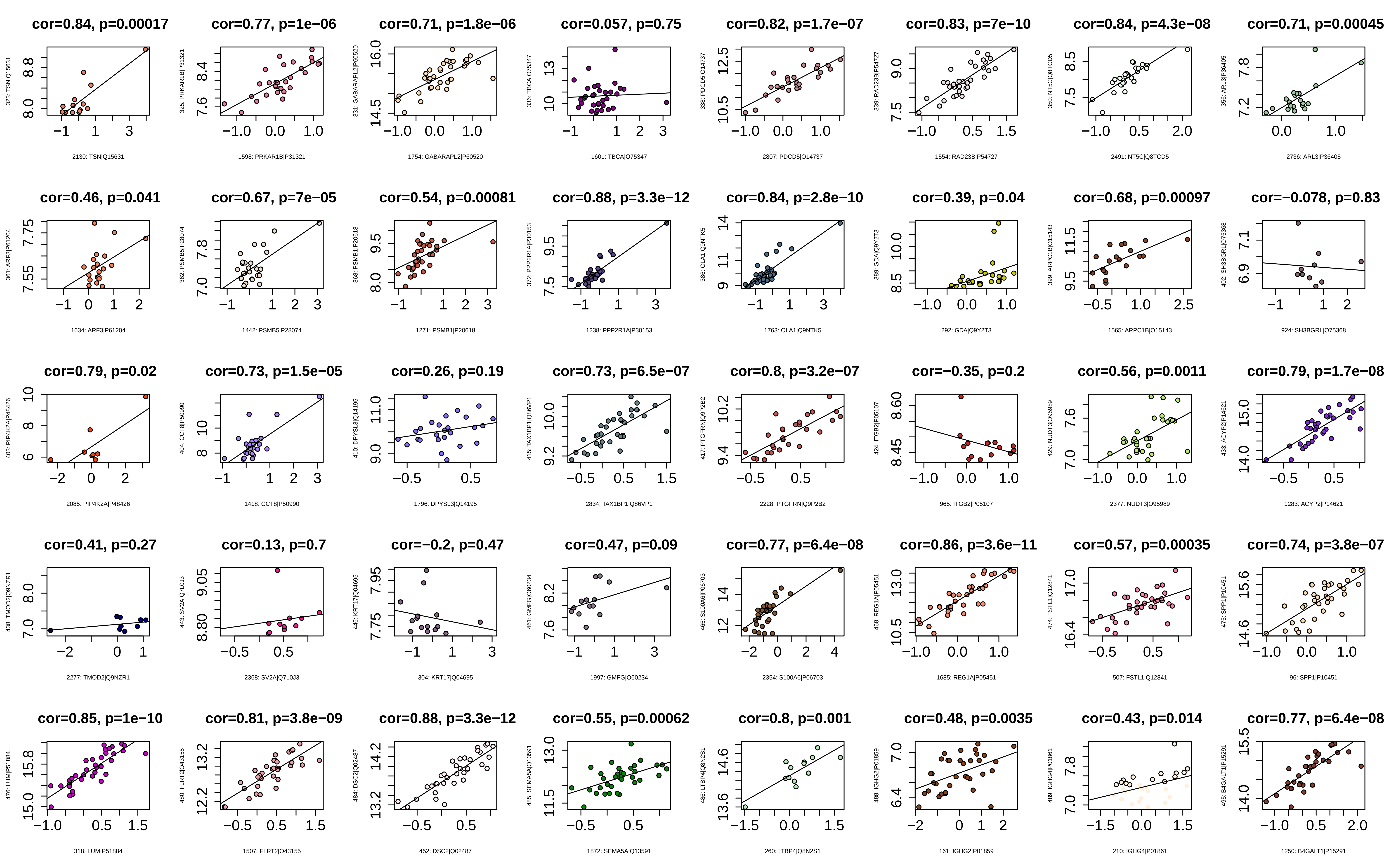

cor=0.84, p=0.00017
323: TSN|Q15631
2130: TSN|Q15631
cor=0.77, p=1e−06
325: PRKAR1B|P31321
1598: PRKAR1B|P31321
cor=0.71, p=1.8e−06
331: GABARAPL2|P60520
1754: GABARAPL2|P60520
cor=0.057, p=0.75
336: TBCA|O75347
1601: TBCA|O75347
cor=0.82, p=1.7e−07
338: PDCD5|O14737
2807: PDCD5|O14737
cor=0.83, p=7e−10
339: RAD23B|P54727
1554: RAD23B|P54727
cor=0.84, p=4.3e−08
350: NT5C|Q8TCD5
2491: NT5C|Q8TCD5
cor=0.71, p=0.00045
356: ARL3|P36405
2736: ARL3|P36405
cor=0.46, p=0.041
361: ARF3|P61204
1634: ARF3|P61204
cor=0.67, p=7e−05
362: PSMB5|P28074
1442: PSMB5|P28074
cor=0.54, p=0.00081
368: PSMB1|P20618
1271: PSMB1|P20618
cor=0.88, p=3.3e−12
372: PPP2R1A|P30153
1238: PPP2R1A|P30153
cor=0.84, p=2.8e−10
386: OLA1|Q9NTK5
1763: OLA1|Q9NTK5
cor=0.39, p=0.04
389: GDA|Q9Y2T3
292: GDA|Q9Y2T3
cor=0.68, p=0.00097
399: ARPC1B|O15143
1565: ARPC1B|O15143
cor=−0.078, p=0.83
402: SH3BGRL|O75368
924: SH3BGRL|O75368
cor=0.79, p=0.02
403: PIP4K2A|P48426
2085: PIP4K2A|P48426
cor=0.73, p=1.5e−05
404: CCT8|P50990
1418: CCT8|P50990
cor=0.26, p=0.19
410: DPYSL3|Q14195
1796: DPYSL3|Q14195
cor=0.73, p=6.5e−07
415: TAX1BP1|Q86VP1
2834: TAX1BP1|Q86VP1
cor=0.8, p=3.2e−07
417: PTGFRN|Q9P2B2
2228: PTGFRN|Q9P2B2
cor=−0.35, p=0.2
424: ITGB2|P05107
965: ITGB2|P05107
cor=0.56, p=0.0011
429: NUDT3|O95989
2377: NUDT3|O95989
cor=0.79, p=1.7e−08
433: ACYP2|P14621
1283: ACYP2|P14621
cor=0.41, p=0.27
438: TMOD2|Q9NZR1
2277: TMOD2|Q9NZR1
cor=0.13, p=0.7
443: SV2A|Q7L0J3
2368: SV2A|Q7L0J3
cor=−0.2, p=0.47
446: KRT17|Q04695
304: KRT17|Q04695
cor=0.47, p=0.09
461: GMFG|O60234
1997: GMFG|O60234
cor=0.77, p=6.4e−08
465: S100A6|P06703
2354: S100A6|P06703
cor=0.86, p=3.6e−11
468: REG1A|P05451
1685: REG1A|P05451
cor=0.57, p=0.00035
474: FSTL1|Q12841
507: FSTL1|Q12841
cor=0.74, p=3.8e−07
475: SPP1|P10451
96: SPP1|P10451
cor=0.85, p=1e−10
476: LUM|P51884
318: LUM|P51884
cor=0.81, p=3.8e−09
480: FLRT2|O43155
1507: FLRT2|O43155
cor=0.88, p=3.3e−12
484: DSC2|Q02487
452: DSC2|Q02487
cor=0.55, p=0.00062
485: SEMA5A|Q13591
1872: SEMA5A|Q13591
cor=0.8, p=0.001
486: LTBP4|Q8N2S1
260: LTBP4|Q8N2S1
cor=0.48, p=0.0035
488: IGHG2|P01859
161: IGHG2|P01859
cor=0.43, p=0.014
489: IGHG4|P01861
210: IGHG4|P01861
cor=0.77, p=6.4e−08
495: B4GALT1|P15291
1250: B4GALT1|P15291

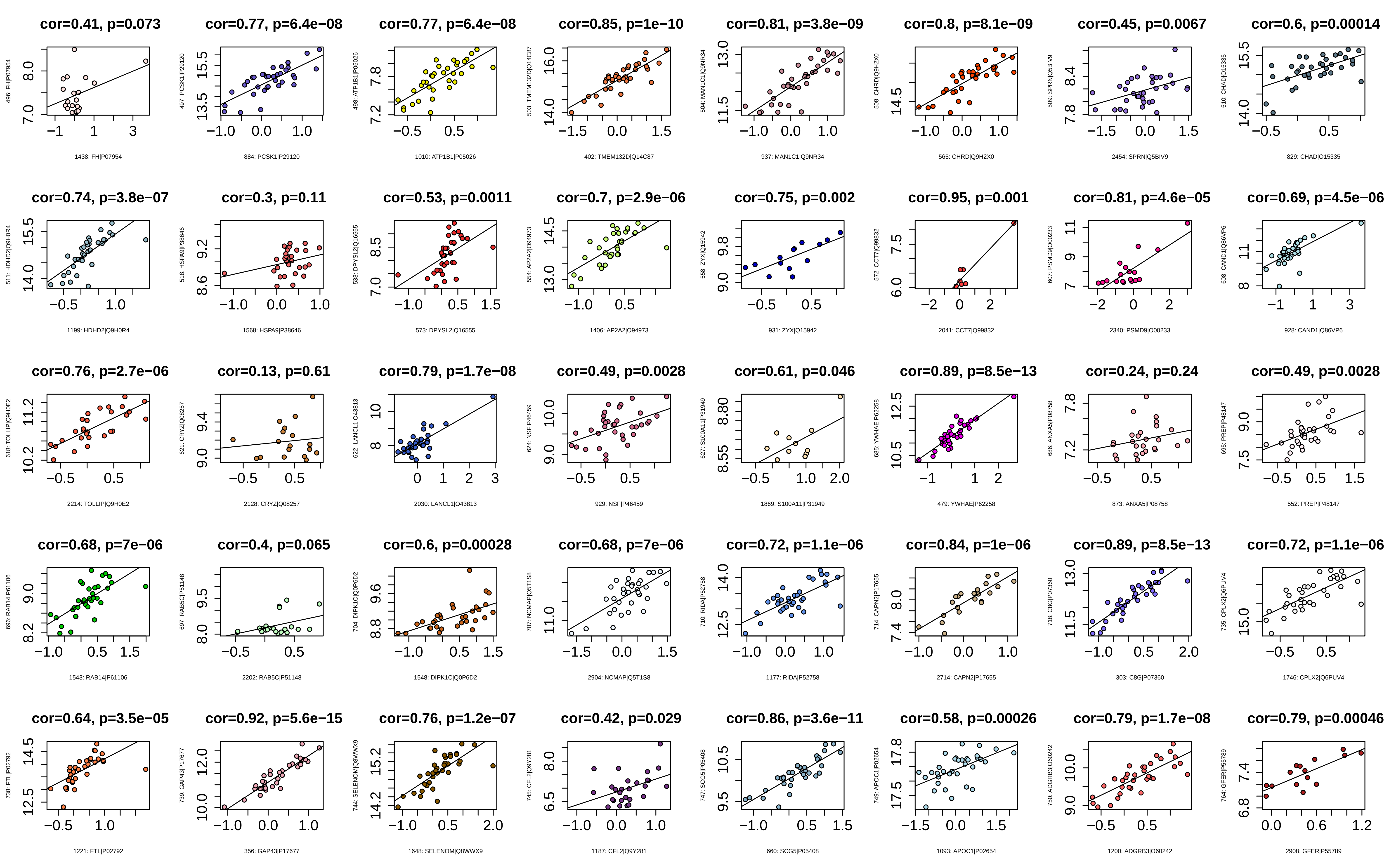
cor=0.41, p=0.073
496: FH|P07954
1438: FH|P07954
cor=0.77, p=6.4e−08
497: PCSK1|P29120
884: PCSK1|P29120
cor=0.77, p=6.4e−08
498: ATP1B1|P05026
1010: ATP1B1|P05026
cor=0.85, p=1e−10
503: TMEM132D|Q14C87
402: TMEM132D|Q14C87
cor=0.81, p=3.8e−09
504: MAN1C1|Q9NR34
937: MAN1C1|Q9NR34
cor=0.8, p=8.1e−09
508: CHRD|Q9H2X0
565: CHRD|Q9H2X0
cor=0.45, p=0.0067
509: SPRN|Q5BIV9
2454: SPRN|Q5BIV9
cor=0.6, p=0.00014
510: CHAD|O15335
829: CHAD|O15335
cor=0.74, p=3.8e−07
511: HDHD2|Q9H0R4
1199: HDHD2|Q9H0R4
cor=0.3, p=0.11
518: HSPA9|P38646
1568: HSPA9|P38646
cor=0.53, p=0.0011
533: DPYSL2|Q16555
573: DPYSL2|Q16555
cor=0.7, p=2.9e−06
554: AP2A2|O94973
1406: AP2A2|O94973
cor=0.75, p=0.002
558: ZYX|Q15942
931: ZYX|Q15942
cor=0.95, p=0.001
572: CCT7|Q99832
2041: CCT7|Q99832
cor=0.81, p=4.6e−05
607: PSMD9|O00233
2340: PSMD9|O00233
cor=0.69, p=4.5e−06
608: CAND1|Q86VP6
928: CAND1|Q86VP6
cor=0.76, p=2.7e−06
618: TOLLIP|Q9H0E2
2214: TOLLIP|Q9H0E2
cor=0.13, p=0.61
621: CRYZ|Q08257
2128: CRYZ|Q08257
cor=0.79, p=1.7e−08
622: LANCL1|O43813
2030: LANCL1|O43813
cor=0.49, p=0.0028
624: NSF|P46459
929: NSF|P46459
cor=0.61, p=0.046
627: S100A11|P31949
1869: S100A11|P31949
cor=0.89, p=8.5e−13
685: YWHAE|P62258
479: YWHAE|P62258
cor=0.24, p=0.24
686: ANXA5|P08758
873: ANXA5|P08758
cor=0.49, p=0.0028
695: PREP|P48147
552: PREP|P48147
cor=0.68, p=7e−06
696: RAB14|P61106
1543: RAB14|P61106
cor=0.4, p=0.065
697: RAB5C|P51148
2202: RAB5C|P51148
cor=0.6, p=0.00028
704: DIPK1C|Q0P6D2
1548: DIPK1C|Q0P6D2
cor=0.68, p=7e−06
707: NCMAP|Q5T1S8
2904: NCMAP|Q5T1S8
cor=0.72, p=1.1e−06
710: RIDA|P52758
1177: RIDA|P52758
cor=0.84, p=1e−06
714: CAPN2|P17655
2714: CAPN2|P17655
cor=0.89, p=8.5e−13
718: C8G|P07360
303: C8G|P07360
cor=0.72, p=1.1e−06
735: CPLX2|Q6PUV4
1746: CPLX2|Q6PUV4
cor=0.64, p=3.5e−05
738: FTL|P02792
1221: FTL|P02792
cor=0.92, p=5.6e−15
739: GAP43|P17677
356: GAP43|P17677
cor=0.76, p=1.2e−07
744: SELENOM|Q8WWX9
1648: SELENOM|Q8WWX9
cor=0.42, p=0.029
745: CFL2|Q9Y281
1187: CFL2|Q9Y281
cor=0.86, p=3.6e−11
747: SCG5|P05408
660: SCG5|P05408
cor=0.58, p=0.00026
749: APOC1|P02654
1093: APOC1|P02654
cor=0.79, p=1.7e−08
750: ADGRB3|O60242
1200: ADGRB3|O60242
cor=0.79, p=0.00046
764: GFER|P55789
2908: GFER|P55789

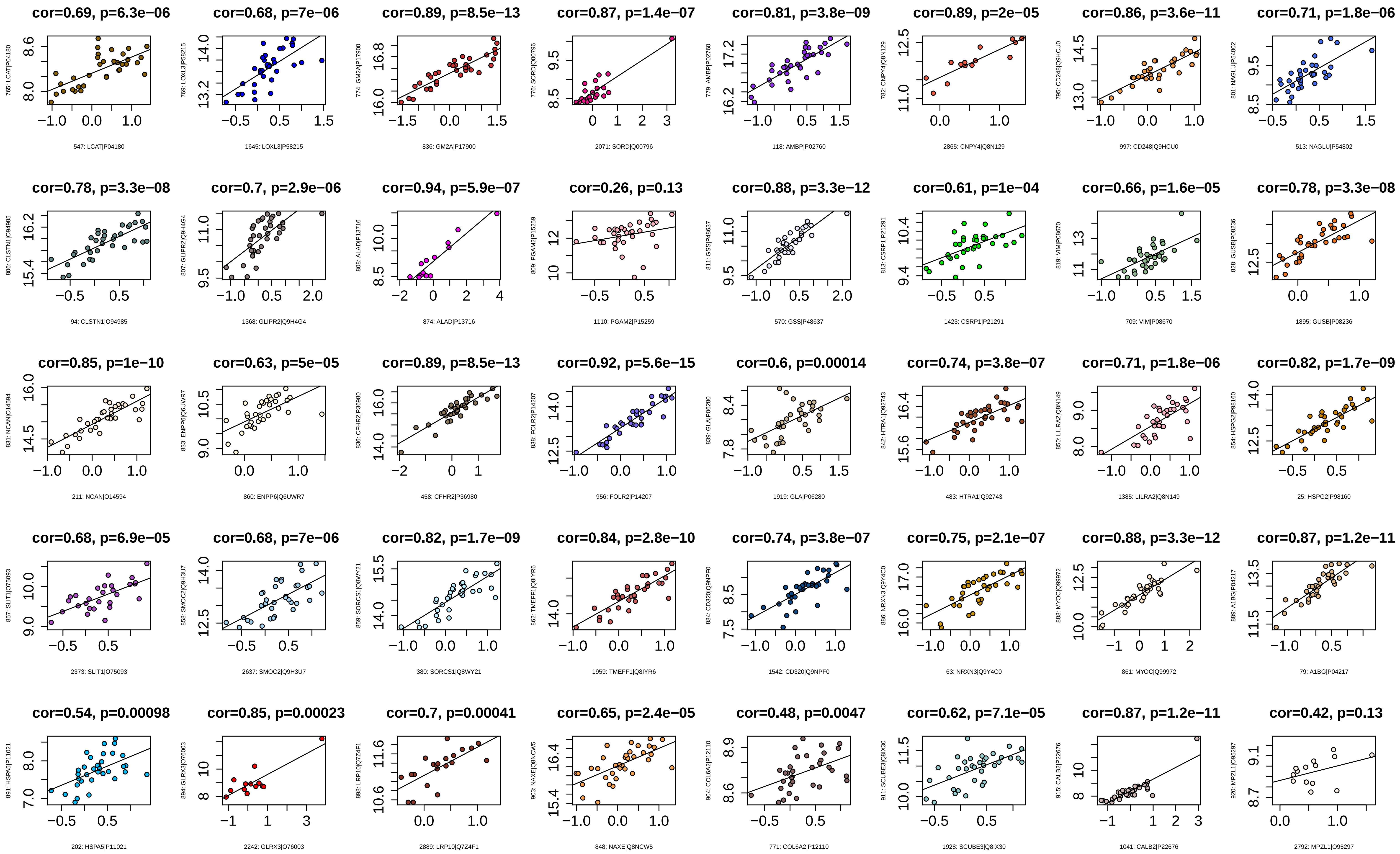
cor=0.69, p=6.3e−06
547: LCAT|P04180
cor=0.68, p=7e−06
1645: LOXL3|P58215
cor=0.89, p=8.5e−13
836: GM2A|P17900
cor=0.87, p=1.4e−07
2071: SORD|Q00796
cor=0.81, p=3.8e−09
118: AMBP|P02760
cor=0.89, p=2e−05
2865: CNPY4|Q8N129
cor=0.86, p=3.6e−11
997: CD248|Q9HCU0
cor=0.71, p=1.8e−06
513: NAGLU|P54802
cor=0.78, p=3.3e−08
94: CLSTN1|O94985
cor=0.7, p=2.9e−06
1368: GLIPR2|Q9H4G4
cor=0.94, p=5.9e−07
874: ALAD|P13716
cor=0.26, p=0.13
1110: PGAM2|P15259
cor=0.88, p=3.3e−12
570: GSS|P48637
cor=0.61, p=1e−04
1423: CSRP1|P21291
cor=0.66, p=1.6e−05
709: VIM|P08670
cor=0.78, p=3.3e−08
1895: GUSB|P08236
cor=0.85, p=1e−10
211: NCAN|O14594
cor=0.63, p=5e−05
860: ENPP6|Q6UWR7
cor=0.89, p=8.5e−13
458: CFHR2|P36980
cor=0.92, p=5.6e−15
956: FOLR2|P14207
cor=0.6, p=0.00014
1919: GLA|P06280
cor=0.74, p=3.8e−07
483: HTRA1|Q92743
cor=0.71, p=1.8e−06
1385: LILRA2|Q8N149
cor=0.82, p=1.7e−09
25: HSPG2|P98160
cor=0.68, p=6.9e−05
2373: SLIT1|O75093
cor=0.68, p=7e−06
2637: SMOC2|Q9H3U7
cor=0.82, p=1.7e−09
380: SORCS1|Q8WY21
cor=0.84, p=2.8e−10
1959: TMEFF1|Q8IYR6
cor=0.74, p=3.8e−07
1542: CD320|Q9NPF0
cor=0.75, p=2.1e−07
63: NRXN3|Q9Y4C0
cor=0.88, p=3.3e−12
861: MYOC|Q99972
cor=0.87, p=1.2e−11
79: A1BG|P04217
cor=0.54, p=0.00098
202: HSPA5|P11021
cor=0.85, p=0.00023
2242: GLRX3|O76003
cor=0.7, p=0.00041
2889: LRP10|Q7Z4F1
cor=0.65, p=2.4e−05
848: NAXE|Q8NCW5
cor=0.48, p=0.0047
771: COL6A2|P12110
cor=0.62, p=7.1e−05
1928: SCUBE3|Q8IX30
cor=0.87, p=1.2e−11
1041: CALB2|P22676
cor=0.42, p=0.13
2792: MPZL1|O95297

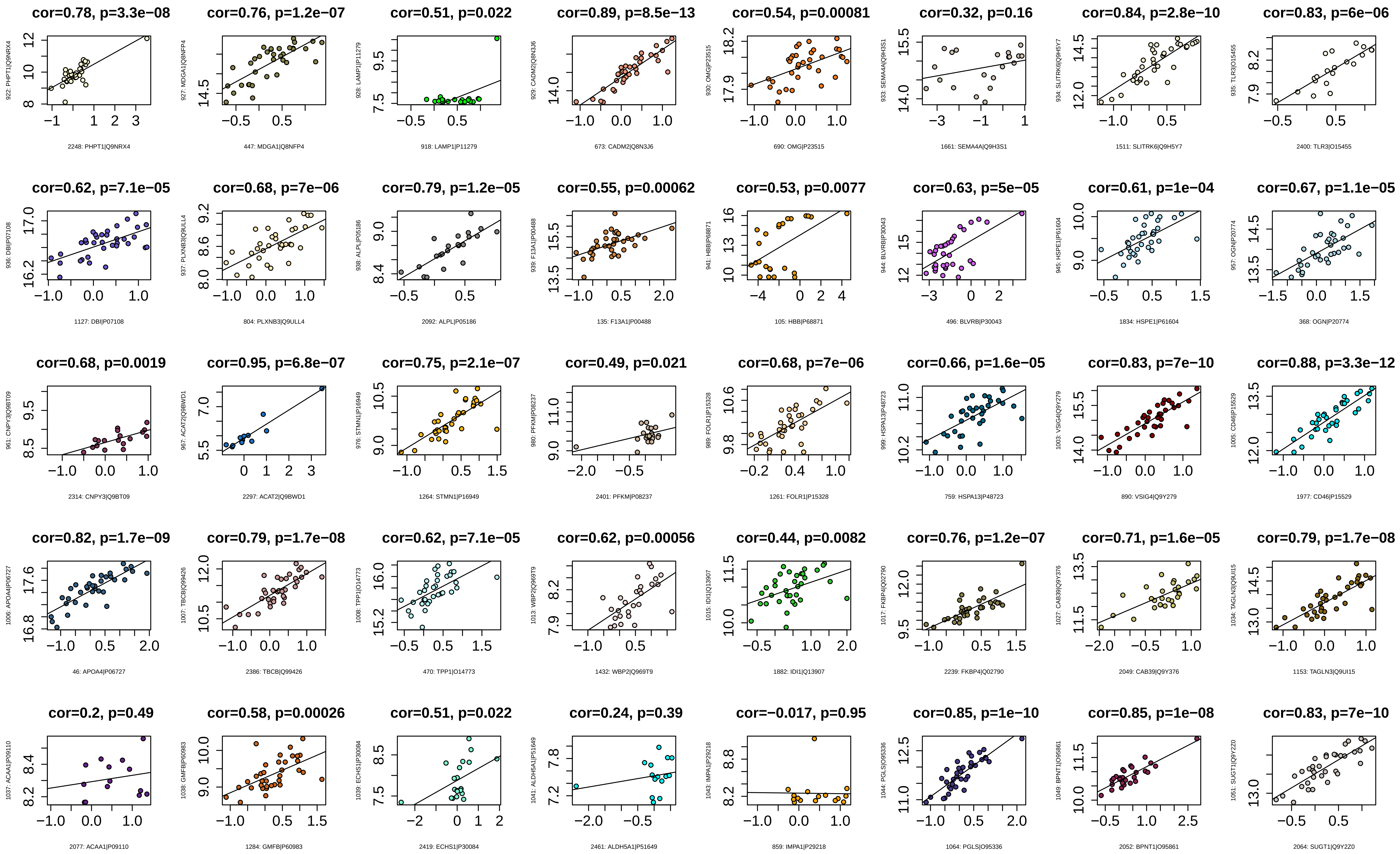
cor=0.78, p=3.3e−08
922: PHPT1|Q9NRX4
2248: PHPT1|Q9NRX4
cor=0.76, p=1.2e−07
927: MDGA1|Q8NFP4
447: MDGA1|Q8NFP4
cor=0.51, p=0.022
928: LAMP1|P11279
918: LAMP1|P11279
cor=0.89, p=8.5e−13
929: CADM2|Q8N3J6
673: CADM2|Q8N3J6
cor=0.54, p=0.00081
930: OMG|P23515
690: OMG|P23515
cor=0.32, p=0.16
933: SEMA4A|Q9H3S1
1661: SEMA4A|Q9H3S1
cor=0.84, p=2.8e−10
934: SLITRK6|Q9H5Y7
1511: SLITRK6|Q9H5Y7
cor=0.83, p=6e−06
935: TLR3|O15455
2400: TLR3|O15455
cor=0.62, p=7.1e−05
936: DBI|P07108
1127: DBI|P07108
cor=0.68, p=7e−06
937: PLXNB3|Q9ULL4
804: PLXNB3|Q9ULL4
cor=0.79, p=1.2e−05
938: ALPL|P05186
2092: ALPL|P05186
cor=0.55, p=0.00062
939: F13A1|P00488
135: F13A1|P00488
cor=0.53, p=0.0077
941: HBB|P68871
105: HBB|P68871
cor=0.63, p=5e−05
944: BLVRB|P30043
496: BLVRB|P30043
cor=0.61, p=1e−04
945: HSPE1|P61604
1834: HSPE1|P61604
cor=0.67, p=1.1e−05
957: OGN|P20774
368: OGN|P20774
cor=0.68, p=0.0019
961: CNPY3|Q9BT09
2314: CNPY3|Q9BT09
cor=0.95, p=6.8e−07
967: ACAT2|Q9BWD1
2297: ACAT2|Q9BWD1
cor=0.75, p=2.1e−07
976: STMN1|P16949
1264: STMN1|P16949
cor=0.49, p=0.021
980: PFKM|P08237
2401: PFKM|P08237
cor=0.68, p=7e−06
989: FOLR1|P15328
1261: FOLR1|P15328
cor=0.66, p=1.6e−05
999: HSPA13|P48723
759: HSPA13|P48723
cor=0.83, p=7e−10
1003: VSIG4|Q9Y279
890: VSIG4|Q9Y279
cor=0.88, p=3.3e−12
1005: CD46|P15529
1977: CD46|P15529
cor=0.82, p=1.7e−09
1006: APOA4|P06727
46: APOA4|P06727
cor=0.79, p=1.7e−08
1007: TBCB|Q99426
2386: TBCB|Q99426
cor=0.62, p=7.1e−05
1008: TPP1|O14773
470: TPP1|O14773
cor=0.62, p=0.00056
1013: WBP2|Q969T9
1432: WBP2|Q969T9
cor=0.44, p=0.0082
1015: IDI1|Q13907
1882: IDI1|Q13907
cor=0.76, p=1.2e−07
1017: FKBP4|Q02790
2239: FKBP4|Q02790
cor=0.71, p=1.6e−05
1027: CAB39|Q9Y376
2049: CAB39|Q9Y376
cor=0.79, p=1.7e−08
1034: TAGLN3|Q9UI15
1153: TAGLN3|Q9UI15
cor=0.2, p=0.49
1037: ACAA1|P09110
2077: ACAA1|P09110
cor=0.58, p=0.00026
1038: GMFB|P60983
1284: GMFB|P60983
cor=0.51, p=0.022
1039: ECHS1|P30084
2419: ECHS1|P30084
cor=0.24, p=0.39
1041: ALDH5A1|P51649
2461: ALDH5A1|P51649
cor=−0.017, p=0.95
1043: IMPA1|P29218
859: IMPA1|P29218
cor=0.85, p=1e−10
1044: PGLS|O95336
1064: PGLS|O95336
cor=0.85, p=1e−08
1049: BPNT1|O95861
2052: BPNT1|O95861
cor=0.83, p=7e−10
1051: SUGT1|Q9Y2Z0
2064: SUGT1|Q9Y2Z0

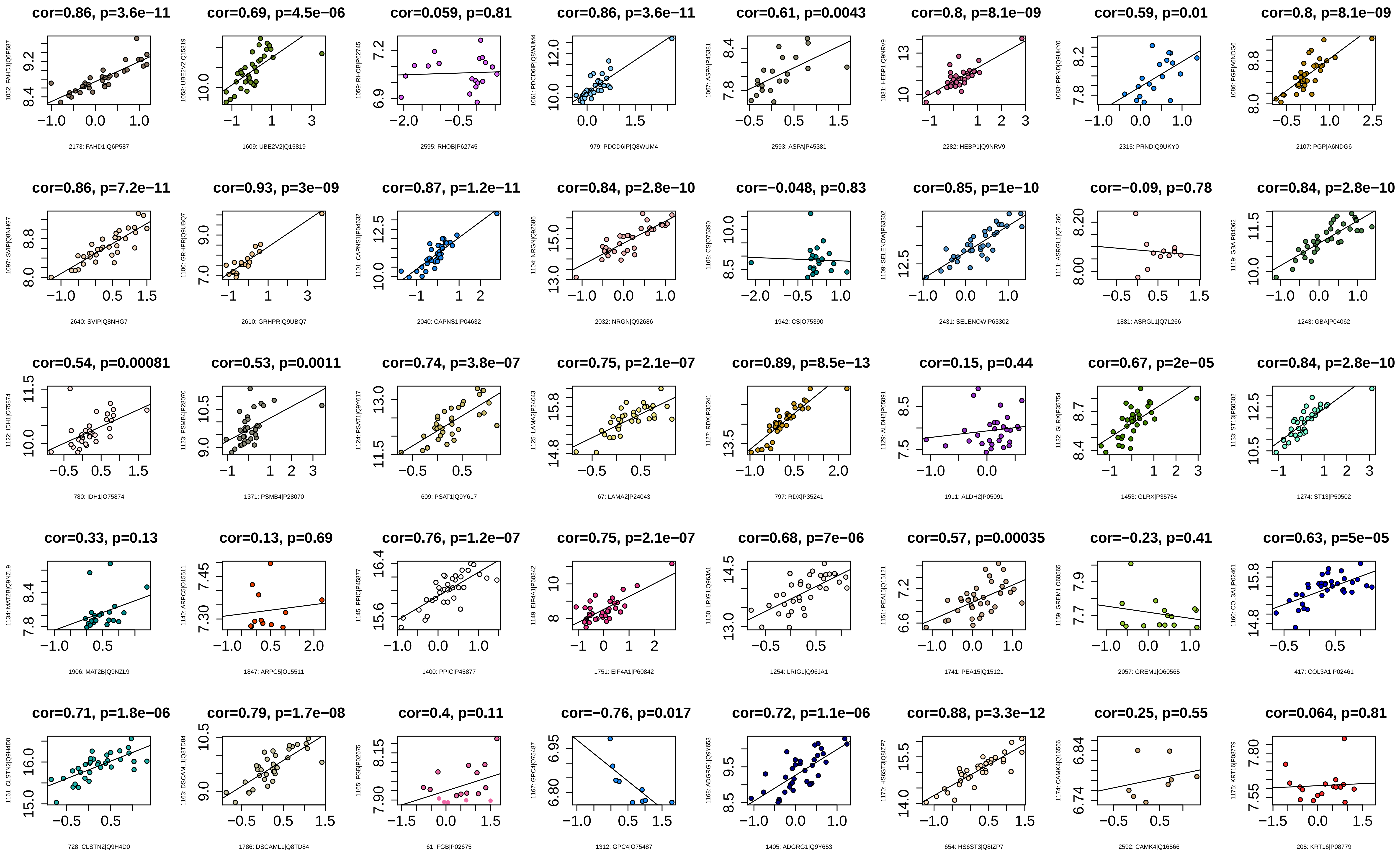

cor=0.86, p=3.6e−11
1052: FAHD1|Q6P587
2173: FAHD1|Q6P587
cor=0.69, p=4.5e−06
1058: UBE2V2|Q15819
1609: UBE2V2|Q15819
cor=0.059, p=0.81
1059: RHOB|P62745
2595: RHOB|P62745
cor=0.86, p=3.6e−11
1061: PDCD6IP|Q8WUM4
979: PDCD6IP|Q8WUM4
cor=0.61, p=0.0043
1067: ASPA|P45381
2593: ASPA|P45381
cor=0.8, p=8.1e−09
1081: HEBP1|Q9NRV9
2282: HEBP1|Q9NRV9
cor=0.59, p=0.01
1083: PRND|Q9UKY0
2315: PRND|Q9UKY0
cor=0.8, p=8.1e−09
1086: PGP|A6NDG6
2107: PGP|A6NDG6
cor=0.86, p=7.2e−11
1097: SVIP|Q8NHG7
2640: SVIP|Q8NHG7
cor=0.93, p=3e−09
1100: GRHPR|Q9UBQ7
2610: GRHPR|Q9UBQ7
cor=0.87, p=1.2e−11
1101: CAPNS1|P04632
2040: CAPNS1|P04632
cor=0.84, p=2.8e−10
1104: NRGN|Q92686
2032: NRGN|Q92686
cor=−0.048, p=0.83
1108: CS|O75390
1942: CS|O75390
cor=0.85, p=1e−10
1109: SELENOW|P63302
2431: SELENOW|P63302
cor=−0.09, p=0.78
1111: ASRGL1|Q7L266
1881: ASRGL1|Q7L266
cor=0.84, p=2.8e−10
1119: GBA|P04062
1243: GBA|P04062
cor=0.54, p=0.00081
1122: IDH1|O75874
780: IDH1|O75874
cor=0.53, p=0.0011
1123: PSMB4|P28070
1371: PSMB4|P28070
cor=0.74, p=3.8e−07
1124: PSAT1|Q9Y617
609: PSAT1|Q9Y617
cor=0.75, p=2.1e−07
1125: LAMA2|P24043
67: LAMA2|P24043
cor=0.89, p=8.5e−13
1127: RDX|P35241
797: RDX|P35241
cor=0.15, p=0.44
1129: ALDH2|P05091
1911: ALDH2|P05091
cor=0.67, p=2e−05
1132: GLRX|P35754
1453: GLRX|P35754
cor=0.84, p=2.8e−10
1133: ST13|P50502
1274: ST13|P50502
cor=0.33, p=0.13
1134: MAT2B|Q9NZL9
1906: MAT2B|Q9NZL9
cor=0.13, p=0.69
1140: ARPC5|O15511
1847: ARPC5|O15511
cor=0.76, p=1.2e−07
1145: PPIC|P45877
1400: PPIC|P45877
cor=0.75, p=2.1e−07
1149: EIF4A1|P60842
1751: EIF4A1|P60842
cor=0.68, p=7e−06
1150: LRIG1|Q96JA1
1254: LRIG1|Q96JA1
cor=0.57, p=0.00035
1151: PEA15|Q15121
1741: PEA15|Q15121
cor=−0.23, p=0.41
1159: GREM1|O60565
2057: GREM1|O60565
cor=0.63, p=5e−05
1160: COL3A1|P02461
417: COL3A1|P02461
cor=0.71, p=1.8e−06
1161: CLSTN2|Q9H4D0
728: CLSTN2|Q9H4D0
cor=0.79, p=1.7e−08
1163: DSCAML1|Q8TD84
1786: DSCAML1|Q8TD84
cor=0.4, p=0.11
1165: FGB|P02675
61: FGB|P02675
cor=−0.76, p=0.017
1167: GPC4|O75487
1312: GPC4|O75487
cor=0.72, p=1.1e−06
1168: ADGRG1|Q9Y653
1405: ADGRG1|Q9Y653
cor=0.88, p=3.3e−12
1170: HS6ST3|Q8IZP7
654: HS6ST3|Q8IZP7
cor=0.25, p=0.55
1174: CAMK4|Q16566
2592: CAMK4|Q16566
cor=0.064, p=0.81
1175: KRT16|P08779
205: KRT16|P08779

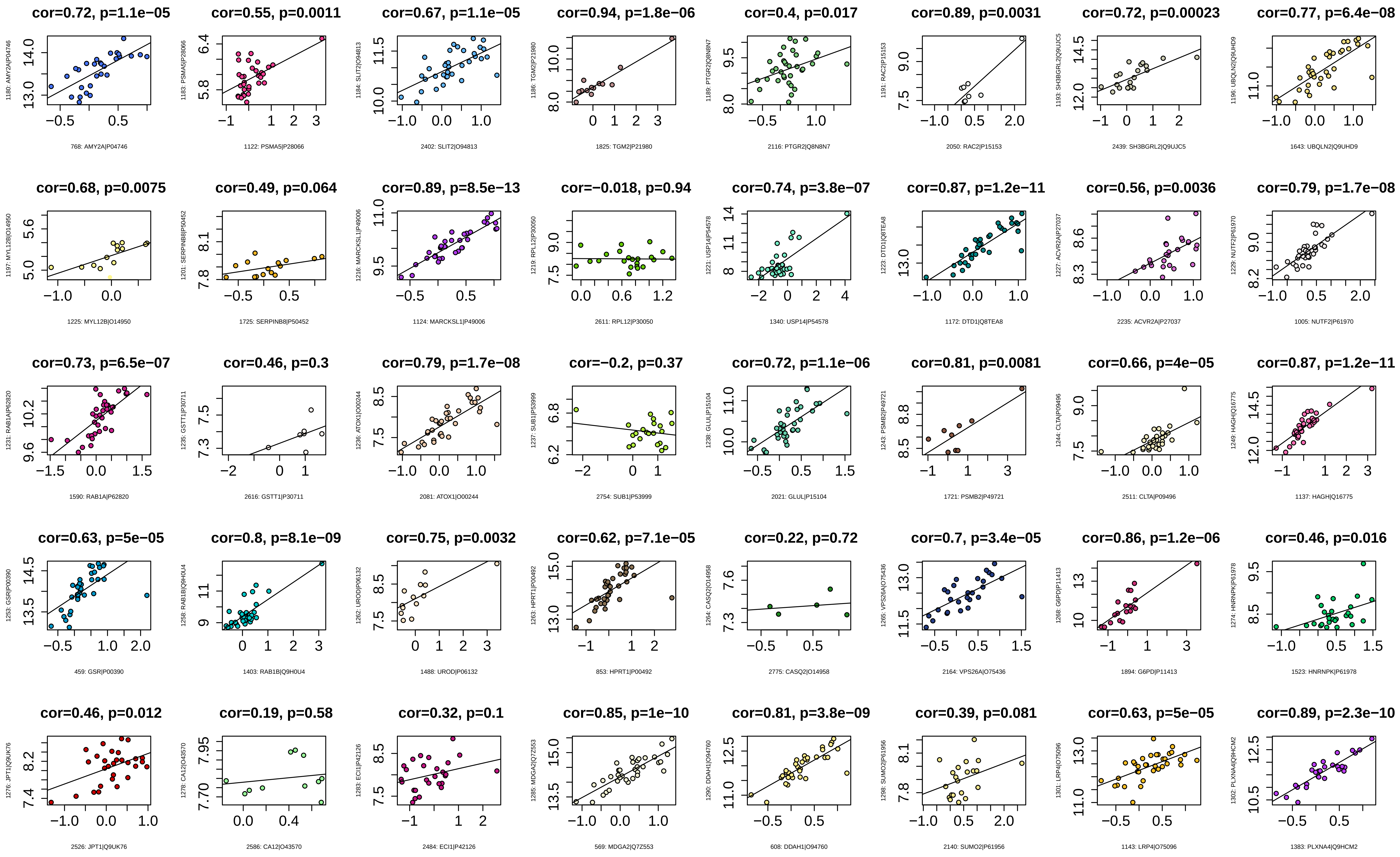

cor=0.72, p=1.1e−05
768: AMY2A|P04746
cor=0.55, p=0.0011
1122: PSMA5|P28066
cor=0.67, p=1.1e−05
2402: SLIT2|O94813
cor=0.94, p=1.8e−06
1825: TGM2|P21980
cor=0.4, p=0.017
2116: PTGR2|Q8N8N7
cor=0.89, p=0.0031
2050: RAC2|P15153
cor=0.72, p=0.00023
2439: SH3BGRL2|Q9UJC5
cor=0.77, p=6.4e−08
1643: UBQLN2|Q9UHD9
cor=0.68, p=0.0075
1225: MYL12B|O14950
cor=0.49, p=0.064
1725: SERPINB8|P50452
cor=0.89, p=8.5e−13
1124: MARCKSL1|P49006
cor=−0.018, p=0.94
2611: RPL12|P30050
cor=0.74, p=3.8e−07
1340: USP14|P54578
cor=0.87, p=1.2e−11
1172: DTD1|Q8TEA8
cor=0.56, p=0.0036
2235: ACVR2A|P27037
cor=0.79, p=1.7e−08
1005: NUTF2|P61970
cor=0.73, p=6.5e−07
1590: RAB1A|P62820
cor=0.46, p=0.3
2616: GSTT1|P30711
cor=0.79, p=1.7e−08
2081: ATOX1|O00244
cor=−0.2, p=0.37
2754: SUB1|P53999
cor=0.72, p=1.1e−06
2021: GLUL|P15104
cor=0.81, p=0.0081
1721: PSMB2|P49721
cor=0.66, p=4e−05
2511: CLTA|P09496
cor=0.87, p=1.2e−11
1137: HAGH|Q16775
cor=0.63, p=5e−05
459: GSR|P00390
cor=0.8, p=8.1e−09
1403: RAB1B|Q9H0U4
cor=0.75, p=0.0032
1488: UROD|P06132
cor=0.62, p=7.1e−05
853: HPRT1|P00492
cor=0.22, p=0.72
2775: CASQ2|O14958
cor=0.7, p=3.4e−05
2164: VPS26A|O75436
cor=0.86, p=1.2e−06
1894: G6PD|P11413
cor=0.46, p=0.016
1523: HNRNPK|P61978
cor=0.46, p=0.012
2526: JPT1|Q9UK76
cor=0.19, p=0.58
2586: CA12|O43570
cor=0.32, p=0.1
2484: ECI1|P42126
cor=0.85, p=1e−10
569: MDGA2|Q7Z553
cor=0.81, p=3.8e−09
608: DDAH1|O94760
cor=0.39, p=0.081
2140: SUMO2|P61956
cor=0.63, p=5e−05
1143: LRP4|O75096
cor=0.89, p=2.3e−10
1383: PLXNA4|Q9HCM2

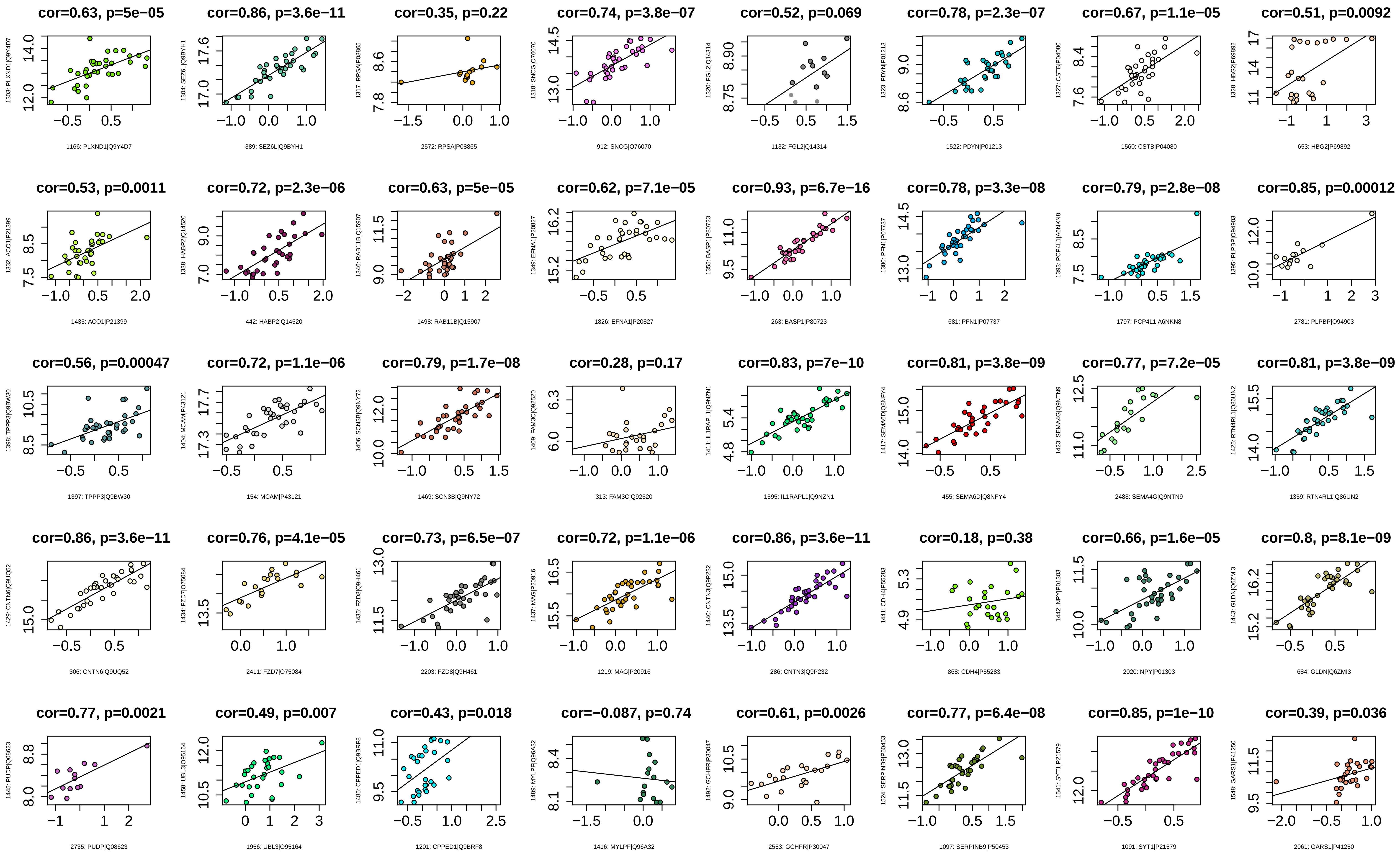
cor=0.63, p=5e−05
1166: PLXND1|Q9Y4D7
cor=0.86, p=3.6e−11
389: SEZ6L|Q9BYH1
cor=0.35, p=0.22
2572: RPSA|P08865
cor=0.74, p=3.8e−07
912: SNCG|O76070
cor=0.52, p=0.069
1132: FGL2|Q14314
cor=0.78, p=2.3e−07
1522: PDYN|P01213
cor=0.67, p=1.1e−05
1560: CSTB|P04080
cor=0.51, p=0.0092
653: HBG2|P69892
cor=0.53, p=0.0011
1435: ACO1|P21399
cor=0.72, p=2.3e−06
442: HABP2|Q14520
cor=0.63, p=5e−05
1498: RAB11B|Q15907
cor=0.62, p=7.1e−05
1826: EFNA1|P20827
cor=0.93, p=6.7e−16
263: BASP1|P80723
cor=0.78, p=3.3e−08
681: PFN1|P07737
cor=0.79, p=2.8e−08
1797: PCP4L1|A6NKN8
cor=0.85, p=0.00012
2781: PLPBP|O94903
cor=0.56, p=0.00047
1397: TPPP3|Q9BW30
cor=0.72, p=1.1e−06
154: MCAM|P43121
cor=0.79, p=1.7e−08
1469: SCN3B|Q9NY72
cor=0.28, p=0.17
313: FAM3C|Q92520
cor=0.83, p=7e−10
1595: IL1RAPL1|Q9NZN1
cor=0.81, p=3.8e−09
455: SEMA6D|Q8NFY4
cor=0.77, p=7.2e−05
2488: SEMA4G|Q9NTN9
cor=0.81, p=3.8e−09
1359: RTN4RL1|Q86UN2
cor=0.86, p=3.6e−11
306: CNTN6|Q9UQ52
cor=0.76, p=4.1e−05
2411: FZD7|O75084
cor=0.73, p=6.5e−07
2203: FZD8|Q9H461
cor=0.72, p=1.1e−06
1219: MAG|P20916
cor=0.86, p=3.6e−11
286: CNTN3|Q9P232
cor=0.18, p=0.38
868: CDH4|P55283
cor=0.66, p=1.6e−05
2020: NPY|P01303
cor=0.8, p=8.1e−09
684: GLDN|Q6ZMI3
cor=0.77, p=0.0021
2735: PUDP|Q08623
cor=0.49, p=0.007
1956: UBL3|O95164
cor=0.43, p=0.018
1201: CPPED1|Q9BRF8
cor=−0.087, p=0.74
1416: MYLPF|Q96A32
cor=0.61, p=0.0026
2553: GCHFR|P30047
cor=0.77, p=6.4e−08
1097: SERPINB9|P50453
cor=0.85, p=1e−10
1091: SYT1|P21579
cor=0.39, p=0.036
2061: GARS1|P41250

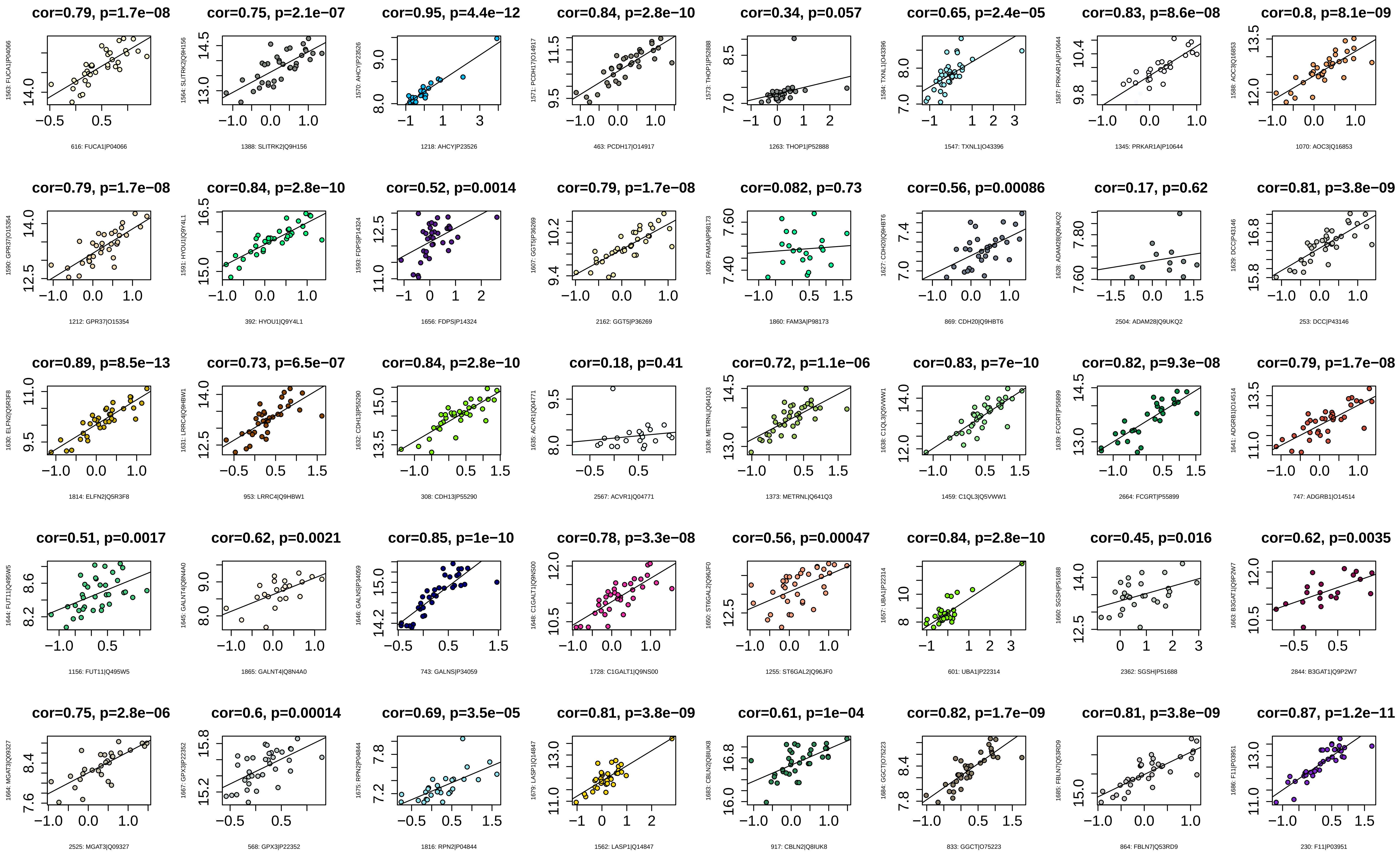
cor=0.79, p=1.7e−08
1563: FUCA1|P04066
616: FUCA1|P04066
cor=0.75, p=2.1e−07
1564: SLITRK2|Q9H156
1388: SLITRK2|Q9H156
cor=0.95, p=4.4e−12
1570: AHCY|P23526
1218: AHCY|P23526
cor=0.84, p=2.8e−10
1571: PCDH17|O14917
463: PCDH17|O14917
cor=0.34, p=0.057
1573: THOP1|P52888
1263: THOP1|P52888
cor=0.65, p=2.4e−05
1584: TXNL1|O43396
1547: TXNL1|O43396
cor=0.83, p=8.6e−08
1587: PRKAR1A|P10644
1345: PRKAR1A|P10644
cor=0.8, p=8.1e−09
1588: AOC3|Q16853
1070: AOC3|Q16853
cor=0.79, p=1.7e−08
1590: GPR37|O15354
1212: GPR37|O15354
cor=0.84, p=2.8e−10
1591: HYOU1|Q9Y4L1
392: HYOU1|Q9Y4L1
cor=0.52, p=0.0014
1593: FDPS|P14324
1656: FDPS|P14324
cor=0.79, p=1.7e−08
1607: GGT5|P36269
2162: GGT5|P36269
cor=0.082, p=0.73
1609: FAM3A|P98173
1860: FAM3A|P98173
cor=0.56, p=0.00086
1627: CDH20|Q9HBT6
869: CDH20|Q9HBT6
cor=0.17, p=0.62
1628: ADAM28|Q9UKQ2
2504: ADAM28|Q9UKQ2
cor=0.81, p=3.8e−09
1629: DCC|P43146
253: DCC|P43146
cor=0.89, p=8.5e−13
1630: ELFN2|Q5R3F8
1814: ELFN2|Q5R3F8
cor=0.73, p=6.5e−07
1631: LRRC4|Q9HBW1
953: LRRC4|Q9HBW1
cor=0.84, p=2.8e−10
1632: CDH13|P55290
308: CDH13|P55290
cor=0.18, p=0.41
1635: ACVR1|Q04771
2567: ACVR1|Q04771
cor=0.72, p=1.1e−06
1636: METRNL|Q641Q3
1373: METRNL|Q641Q3
cor=0.83, p=7e−10
1638: C1QL3|Q5VWW1
1459: C1QL3|Q5VWW1
cor=0.82, p=9.3e−08
1639: FCGRT|P55899
2664: FCGRT|P55899
cor=0.79, p=1.7e−08
1641: ADGRB1|O14514
747: ADGRB1|O14514
cor=0.51, p=0.0017
1644: FUT11|Q495W5
1156: FUT11|Q495W5
cor=0.62, p=0.0021
1645: GALNT4|Q8N4A0
1865: GALNT4|Q8N4A0
cor=0.85, p=1e−10
1646: GALNS|P34059
743: GALNS|P34059
cor=0.78, p=3.3e−08
1648: C1GALT1|Q9NS00
1728: C1GALT1|Q9NS00
cor=0.56, p=0.00047
1650: ST6GAL2|Q96JF0
1255: ST6GAL2|Q96JF0
cor=0.84, p=2.8e−10
1657: UBA1|P22314
601: UBA1|P22314
cor=0.45, p=0.016
1660: SGSH|P51688
2362: SGSH|P51688
cor=0.62, p=0.0035
1663: B3GAT1|Q9P2W7
2844: B3GAT1|Q9P2W7
cor=0.75, p=2.8e−06
1664: MGAT3|Q09327
2525: MGAT3|Q09327
cor=0.6, p=0.00014
1667: GPX3|P22352
568: GPX3|P22352
cor=0.69, p=3.5e−05
1675: RPN2|P04844
1816: RPN2|P04844
cor=0.81, p=3.8e−09
1679: LASP1|Q14847
1562: LASP1|Q14847
cor=0.61, p=1e−04
1683: CBLN2|Q8IUK8
917: CBLN2|Q8IUK8
cor=0.82, p=1.7e−09
1684: GGCT|O75223
833: GGCT|O75223
cor=0.81, p=3.8e−09
1685: FBLN7|Q53RD9
864: FBLN7|Q53RD9
cor=0.87, p=1.2e−11
1686: F11|P03951
230: F11|P03951

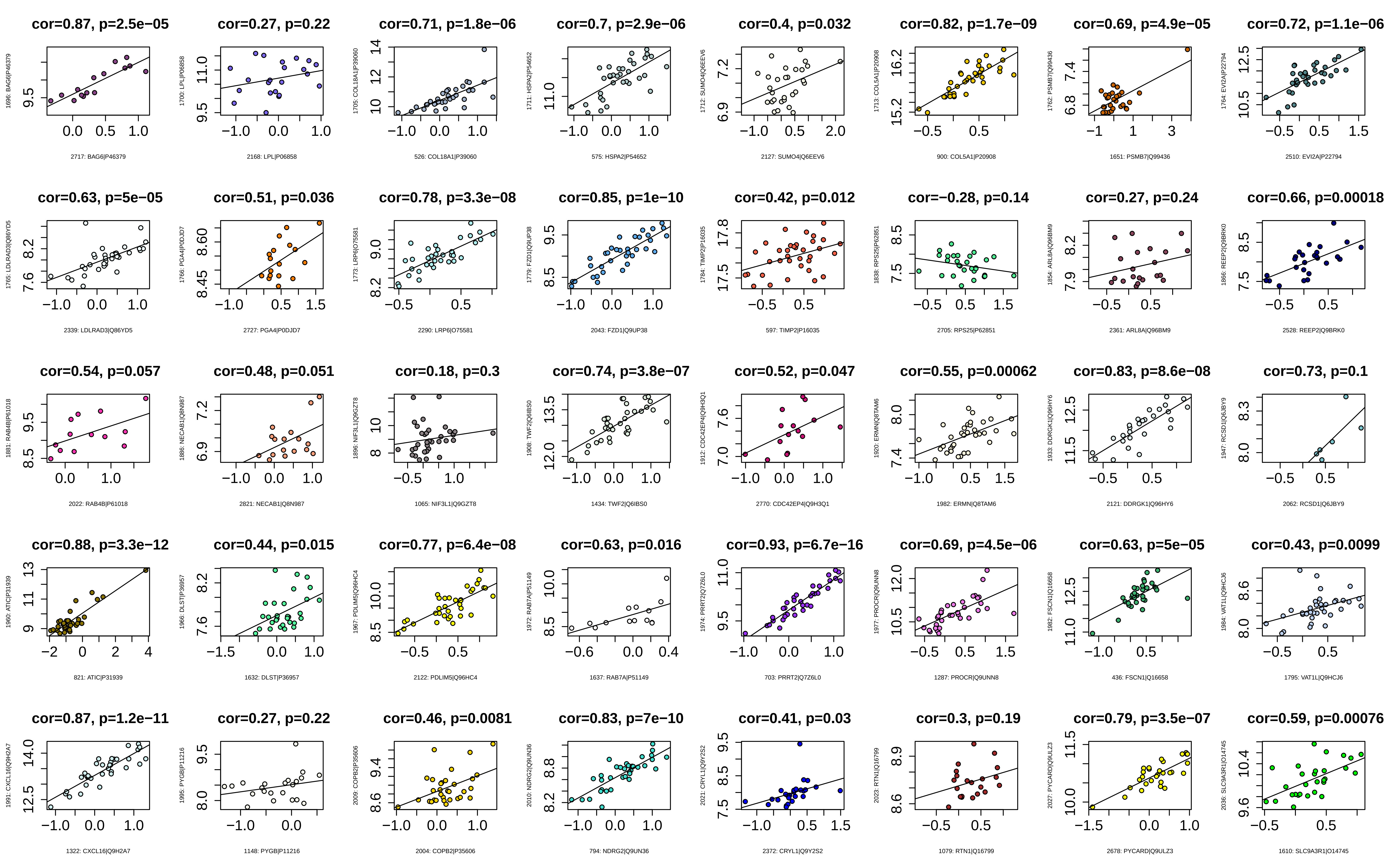
cor=0.87, p=2.5e−05
2717: BAG6|P46379
cor=0.27, p=0.22
2168: LPL|P06858
cor=0.71, p=1.8e−06
526: COL18A1|P39060
cor=0.7, p=2.9e−06
575: HSPA2|P54652
cor=0.4, p=0.032
2127: SUMO4|Q6EEV6
cor=0.82, p=1.7e−09
900: COL5A1|P20908
cor=0.69, p=4.9e−05
1651: PSMB7|Q99436
cor=0.72, p=1.1e−06
2510: EVI2A|P22794
cor=0.63, p=5e−05
2339: LDLRAD3|Q86YD5
cor=0.51, p=0.036
2727: PGA4|P0DJD7
cor=0.78, p=3.3e−08
2290: LRP6|O75581
cor=0.85, p=1e−10
2043: FZD1|Q9UP38
cor=0.42, p=0.012
597: TIMP2|P16035
cor=−0.28, p=0.14
2705: RPS25|P62851
cor=0.27, p=0.24
2361: ARL8A|Q96BM9
cor=0.66, p=0.00018
2528: REEP2|Q9BRK0
cor=0.54, p=0.057
2022: RAB4B|P61018
cor=0.48, p=0.051
2821: NECAB1|Q8N987
cor=0.18, p=0.3
1065: NIF3L1|Q9GZT8
cor=0.74, p=3.8e−07
1434: TWF2|Q6IBS0
cor=0.52, p=0.047
2770: CDC42EP4|Q9H3Q1
cor=0.55, p=0.00062
1982: ERMN|Q8TAM6
cor=0.83, p=8.6e−08
2121: DDRGK1|Q96HY6
cor=0.73, p=0.1
2062: RCSD1|Q6JBY9
cor=0.88, p=3.3e−12
821: ATIC|P31939
cor=0.44, p=0.015
1632: DLST|P36957
cor=0.77, p=6.4e−08
2122: PDLIM5|Q96HC4
cor=0.63, p=0.016
1637: RAB7A|P51149
cor=0.93, p=6.7e−16
703: PRRT2|Q7Z6L0
cor=0.69, p=4.5e−06
1287: PROCR|Q9UNN8
cor=0.63, p=5e−05
436: FSCN1|Q16658
cor=0.43, p=0.0099
1795: VAT1L|Q9HCJ6
cor=0.87, p=1.2e−11
1322: CXCL16|Q9H2A7
cor=0.27, p=0.22
1148: PYGB|P11216
cor=0.46, p=0.0081
2004: COPB2|P35606
cor=0.83, p=7e−10
794: NDRG2|Q9UN36
cor=0.41, p=0.03
2372: CRYL1|Q9Y2S2
cor=0.3, p=0.19
1079: RTN1|Q16799
cor=0.79, p=3.5e−07
2678: PYCARD|Q9ULZ3
cor=0.59, p=0.00076
1610: SLC9A3R1|O14745

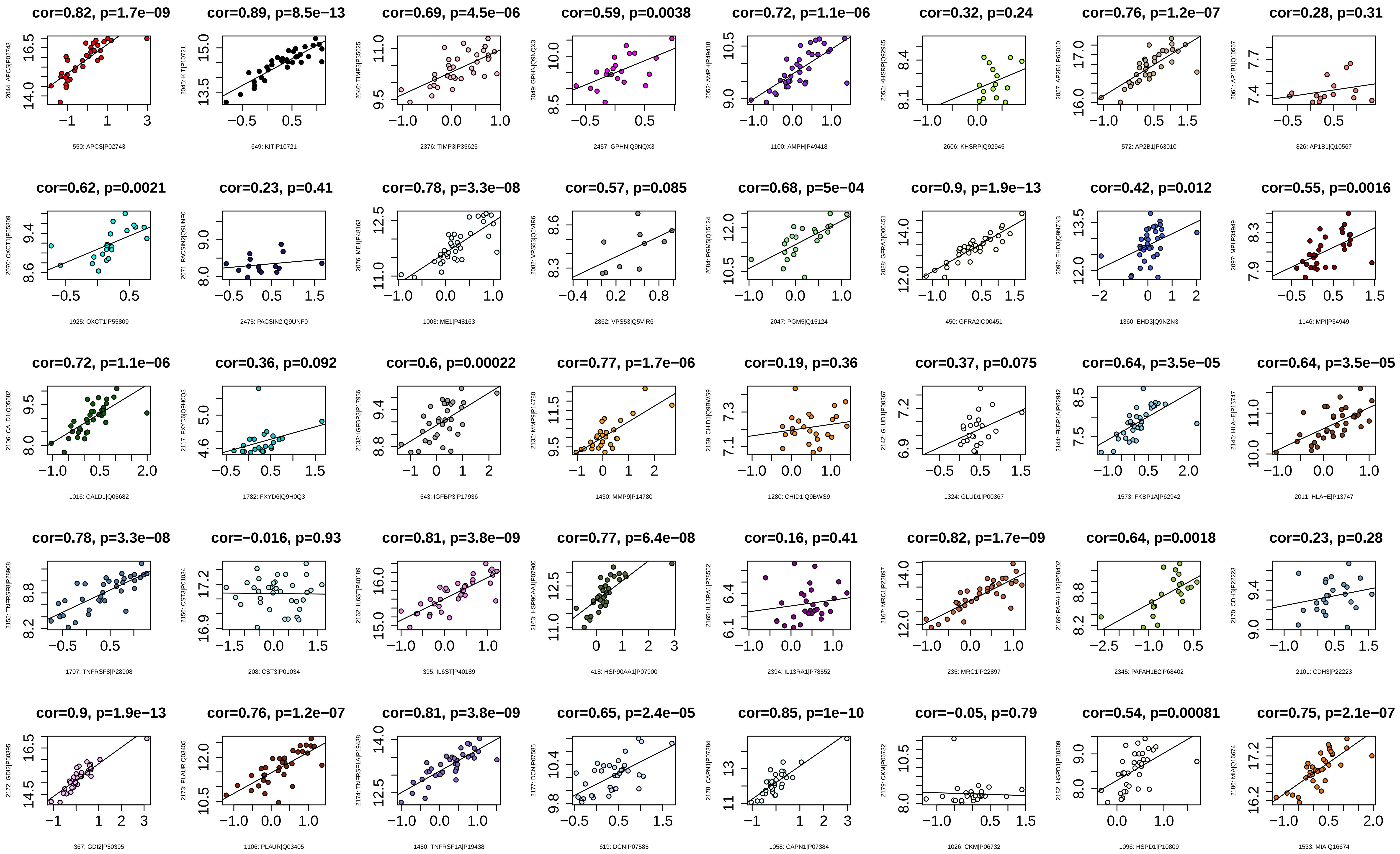
cor=0.82, p=1.7e−09
2044: APCS|P02743
550: APCS|P02743
cor=0.89, p=8.5e−13
2045: KIT|P10721
649: KIT|P10721
cor=0.69, p=4.5e−06
2046: TIMP3|P35625
2376: TIMP3|P35625
cor=0.59, p=0.0038
2049: GPHN|Q9NQX3
2457: GPHN|Q9NQX3
cor=0.72, p=1.1e−06
2052: AMPH|P49418
1100: AMPH|P49418
cor=0.32, p=0.24
2055: KHSRP|Q92945
2606: KHSRP|Q92945
cor=0.76, p=1.2e−07
2057: AP2B1|P63010
572: AP2B1|P63010
cor=0.28, p=0.31
2061: AP1B1|Q10567
826: AP1B1|Q10567
cor=0.62, p=0.0021
2070: OXCT1|P55809
1925: OXCT1|P55809
cor=0.23, p=0.41
2071: PACSIN2|Q9UNF0
2475: PACSIN2|Q9UNF0
cor=0.78, p=3.3e−08
2076: ME1|P48163
1003: ME1|P48163
cor=0.57, p=0.085
2082: VPS53|Q5VIR6
2862: VPS53|Q5VIR6
cor=0.68, p=5e−04
2084: PGM5|Q15124
2047: PGM5|Q15124
cor=0.9, p=1.9e−13
2088: GFRA2|O00451
450: GFRA2|O00451
cor=0.42, p=0.012
2096: EHD3|Q9NZN3
1360: EHD3|Q9NZN3
cor=0.55, p=0.0016
2097: MPI|P34949
1146: MPI|P34949
cor=0.72, p=1.1e−06
2106: CALD1|Q05682
1016: CALD1|Q05682
cor=0.36, p=0.092
2117: FXYD6|Q9H0Q3
1782: FXYD6|Q9H0Q3
cor=0.6, p=0.00022
2133: IGFBP3|P17936
543: IGFBP3|P17936
cor=0.77, p=1.7e−06
2135: MMP9|P14780
1430: MMP9|P14780
cor=0.19, p=0.36
2139: CHID1|Q9BWS9
1280: CHID1|Q9BWS9
cor=0.37, p=0.075
2142: GLUD1|P00367
1324: GLUD1|P00367
cor=0.64, p=3.5e−05
2144: FKBP1A|P62942
1573: FKBP1A|P62942
cor=0.64, p=3.5e−05
2146: HLA−E|P13747
2011: HLA−E|P13747
cor=0.78, p=3.3e−08
2155: TNFRSF8|P28908
1707: TNFRSF8|P28908
cor=−0.016, p=0.93
2156: CST3|P01034
208: CST3|P01034
cor=0.81, p=3.8e−09
2162: IL6ST|P40189
395: IL6ST|P40189
cor=0.77, p=6.4e−08
2163: HSP90AA1|P07900
418: HSP90AA1|P07900
cor=0.16, p=0.41
2165: IL13RA1|P78552
2394: IL13RA1|P78552
cor=0.82, p=1.7e−09
2167: MRC1|P22897
235: MRC1|P22897
cor=0.64, p=0.0018
2169: PAFAH1B2|P68402
2345: PAFAH1B2|P68402
cor=0.23, p=0.28
2170: CDH3|P22223
2101: CDH3|P22223
cor=0.9, p=1.9e−13
2172: GDI2|P50395
367: GDI2|P50395
cor=0.76, p=1.2e−07
2173: PLAUR|Q03405
1106: PLAUR|Q03405
cor=0.81, p=3.8e−09
2174: TNFRSF1A|P19438
1450: TNFRSF1A|P19438
cor=0.65, p=2.4e−05
2177: DCN|P07585
619: DCN|P07585
cor=0.85, p=1e−10
2178: CAPN1|P07384
1058: CAPN1|P07384
cor=−0.05, p=0.79
2179: CKM|P06732
1026: CKM|P06732
cor=0.54, p=0.00081
2182: HSPD1|P10809
1096: HSPD1|P10809
cor=0.75, p=2.1e−07
2186: MIA|Q16674
1533: MIA|Q16674

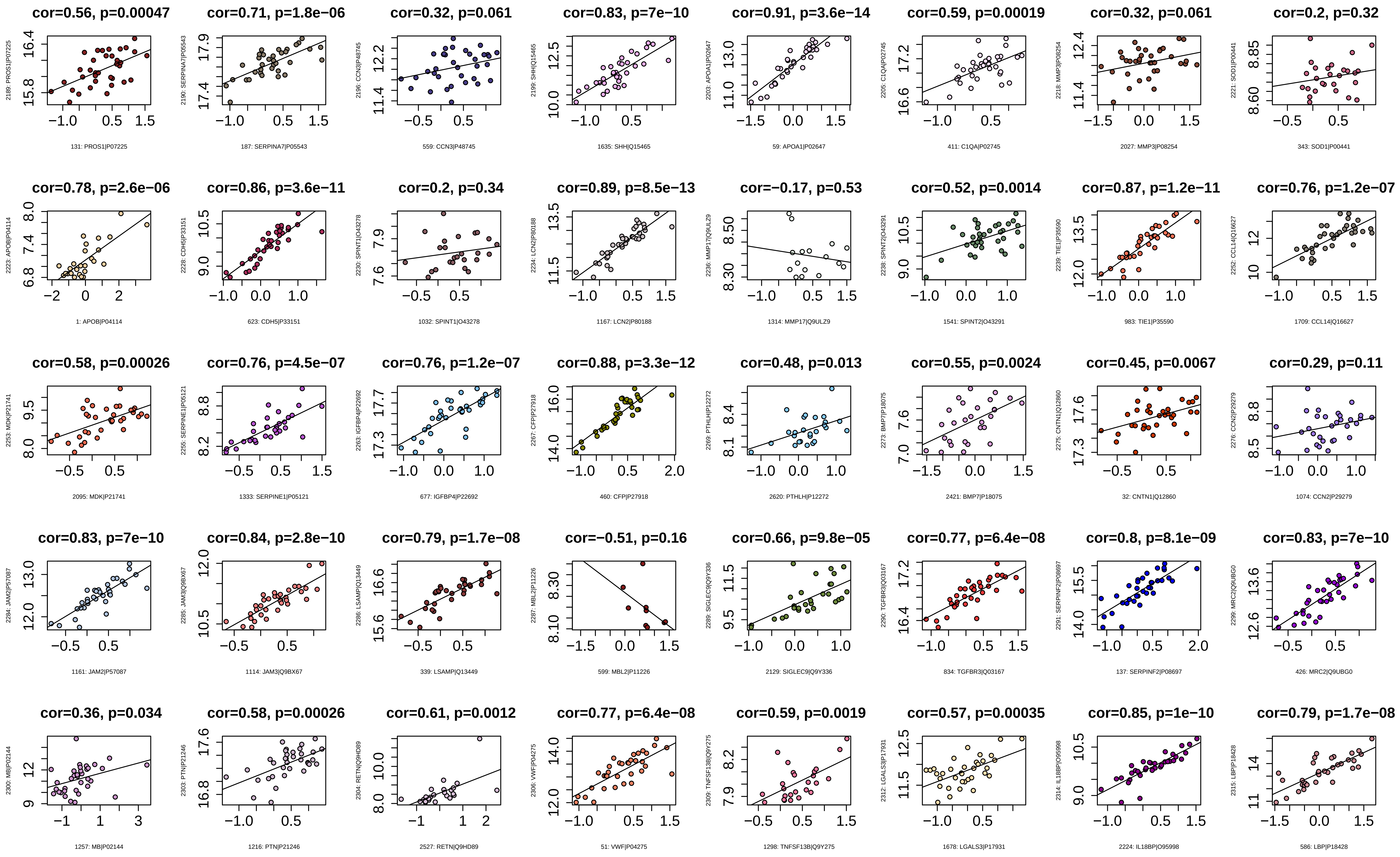
cor=0.56, p=0.00047
2189: PROS1|P07225
131: PROS1|P07225
cor=0.71, p=1.8e−06
2190: SERPINA7|P05543
187: SERPINA7|P05543
cor=0.32, p=0.061
2196: CCN3|P48745
559: CCN3|P48745
cor=0.83, p=7e−10
2199: SHH|Q15465
1635: SHH|Q15465
cor=0.91, p=3.6e−14
2203: APOA1|P02647
59: APOA1|P02647
cor=0.59, p=0.00019
2205: C1QA|P02745
411: C1QA|P02745
cor=0.32, p=0.061
2218: MMP3|P08254
2027: MMP3|P08254
cor=0.2, p=0.32
2221: SOD1|P00441
343: SOD1|P00441
cor=0.78, p=2.6e−06
2223: APOB|P04114
1: APOB|P04114
cor=0.86, p=3.6e−11
2228: CDH5|P33151
623: CDH5|P33151
cor=0.2, p=0.34
2230: SPINT1|O43278
1032: SPINT1|O43278
cor=0.89, p=8.5e−13
2234: LCN2|P80188
1167: LCN2|P80188
cor=−0.17, p=0.53
2236: MMP17|Q9ULZ9
1314: MMP17|Q9ULZ9
cor=0.52, p=0.0014
2238: SPINT2|O43291
1541: SPINT2|O43291
cor=0.87, p=1.2e−11
2239: TIE1|P35590
983: TIE1|P35590
cor=0.76, p=1.2e−07
2252: CCL14|Q16627
1709: CCL14|Q16627
cor=0.58, p=0.00026
2253: MDK|P21741
2095: MDK|P21741
cor=0.76, p=4.5e−07
2255: SERPINE1|P05121
1333: SERPINE1|P05121
cor=0.76, p=1.2e−07
2263: IGFBP4|P22692
677: IGFBP4|P22692
cor=0.88, p=3.3e−12
2267: CFP|P27918
460: CFP|P27918
cor=0.48, p=0.013
2269: PTHLH|P12272
2620: PTHLH|P12272
cor=0.55, p=0.0024
2273: BMP7|P18075
2421: BMP7|P18075
cor=0.45, p=0.0067
2275: CNTN1|Q12860
32: CNTN1|Q12860
cor=0.29, p=0.11
2276: CCN2|P29279
1074: CCN2|P29279
cor=0.83, p=7e−10
2284: JAM2|P57087
1161: JAM2|P57087
cor=0.84, p=2.8e−10
2285: JAM3|Q9BX67
1114: JAM3|Q9BX67
cor=0.79, p=1.7e−08
2286: LSAMP|Q13449
339: LSAMP|Q13449
cor=−0.51, p=0.16
2287: MBL2|P11226
599: MBL2|P11226
cor=0.66, p=9.8e−05
2289: SIGLEC9|Q9Y336
2129: SIGLEC9|Q9Y336
cor=0.77, p=6.4e−08
2290: TGFBR3|Q03167
834: TGFBR3|Q03167
cor=0.8, p=8.1e−09
2291: SERPINF2|P08697
137: SERPINF2|P08697
cor=0.83, p=7e−10
2299: MRC2|Q9UBG0
426: MRC2|Q9UBG0
cor=0.36, p=0.034
2300: MB|P02144
1257: MB|P02144
cor=0.58, p=0.00026
2303: PTN|P21246
1216: PTN|P21246
cor=0.61, p=0.0012
2304: RETN|Q9HD89
2527: RETN|Q9HD89
cor=0.77, p=6.4e−08
2306: VWF|P04275
51: VWF|P04275
cor=0.59, p=0.0019
2309: TNFSF13B|Q9Y275
1298: TNFSF13B|Q9Y275
cor=0.57, p=0.00035
2312: LGALS3|P17931
1678: LGALS3|P17931
cor=0.85, p=1e−10
2314: IL18BP|O95998
2224: IL18BP|O95998
cor=0.79, p=1.7e−08
2315: LBP|P18428
586: LBP|P18428

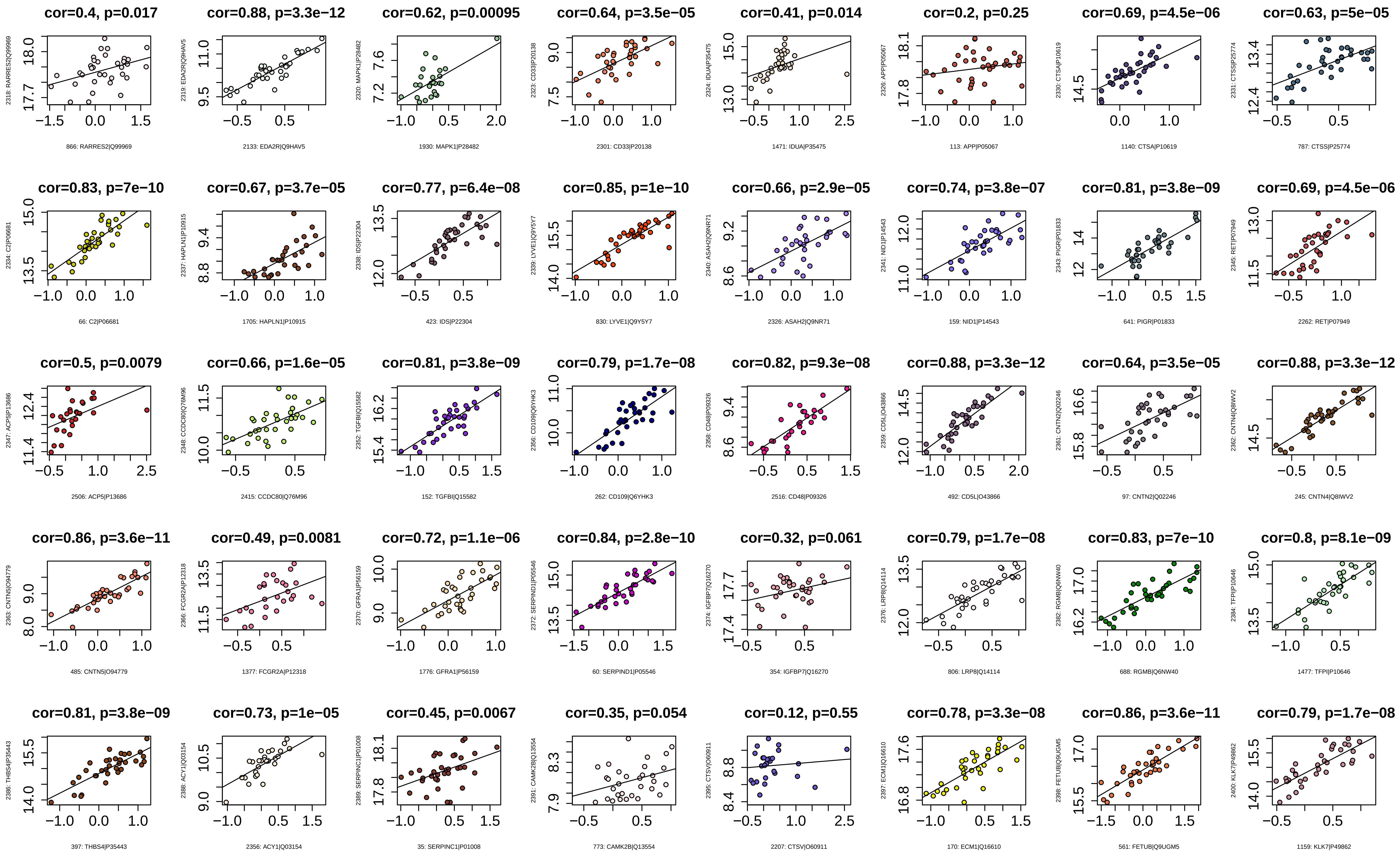
cor=0.4, p=0.017
2318: RARRES2|Q99969
866: RARRES2|Q99969
cor=0.88, p=3.3e−12
2319: EDA2R|Q9HAV5
2133: EDA2R|Q9HAV5
cor=0.62, p=0.00095
2320: MAPK1|P28482
1930: MAPK1|P28482
cor=0.64, p=3.5e−05
2323: CD33|P20138
2301: CD33|P20138
cor=0.41, p=0.014
2324: IDUA|P35475
1471: IDUA|P35475
cor=0.2, p=0.25
2326: APP|P05067
113: APP|P05067
cor=0.69, p=4.5e−06
2330: CTSA|P10619
1140: CTSA|P10619
cor=0.63, p=5e−05
2331: CTSS|P25774
787: CTSS|P25774
cor=0.83, p=7e−10
2334: C2|P06681
66: C2|P06681
cor=0.67, p=3.7e−05
2337: HAPLN1|P10915
1705: HAPLN1|P10915
cor=0.77, p=6.4e−08
2338: IDS|P22304
423: IDS|P22304
cor=0.85, p=1e−10
2339: LYVE1|Q9Y5Y7
830: LYVE1|Q9Y5Y7
cor=0.66, p=2.9e−05
2340: ASAH2|Q9NR71
2326: ASAH2|Q9NR71
cor=0.74, p=3.8e−07
2341: NID1|P14543
159: NID1|P14543
cor=0.81, p=3.8e−09
2343: PIGR|P01833
641: PIGR|P01833
cor=0.69, p=4.5e−06
2345: RET|P07949
2262: RET|P07949
cor=0.5, p=0.0079
2347: ACP5|P13686
2506: ACP5|P13686
cor=0.66, p=1.6e−05
2348: CCDC80|Q76M96
2415: CCDC80|Q76M96
cor=0.81, p=3.8e−09
2352: TGFBI|Q15582
152: TGFBI|Q15582
cor=0.79, p=1.7e−08
2356: CD109|Q6YHK3
262: CD109|Q6YHK3
cor=0.82, p=9.3e−08
2358: CD48|P09326
2516: CD48|P09326
cor=0.88, p=3.3e−12
2359: CD5L|O43866
492: CD5L|O43866
cor=0.64, p=3.5e−05
2361: CNTN2|Q02246
97: CNTN2|Q02246
cor=0.88, p=3.3e−12
2362: CNTN4|Q8IWV2
245: CNTN4|Q8IWV2
cor=0.86, p=3.6e−11
2363: CNTN5|O94779
485: CNTN5|O94779
cor=0.49, p=0.0081
2366: FCGR2A|P12318
1377: FCGR2A|P12318
cor=0.72, p=1.1e−06
2370: GFRA1|P56159
1776: GFRA1|P56159
cor=0.84, p=2.8e−10
2372: SERPIND1|P05546
60: SERPIND1|P05546
cor=0.32, p=0.061
2374: IGFBP7|Q16270
354: IGFBP7|Q16270
cor=0.79, p=1.7e−08
2376: LRP8|Q14114
806: LRP8|Q14114
cor=0.83, p=7e−10
2382: RGMB|Q6NW40
688: RGMB|Q6NW40
cor=0.8, p=8.1e−09
2384: TFPI|P10646
1477: TFPI|P10646
cor=0.81, p=3.8e−09
2386: THBS4|P35443
397: THBS4|P35443
cor=0.73, p=1e−05
2388: ACY1|Q03154
2356: ACY1|Q03154
cor=0.45, p=0.0067
2389: SERPINC1|P01008
35: SERPINC1|P01008
cor=0.35, p=0.054
2391: CAMK2B|Q13554
773: CAMK2B|Q13554
cor=0.12, p=0.55
2395: CTSV|O60911
2207: CTSV|O60911
cor=0.78, p=3.3e−08
2397: ECM1|Q16610
170: ECM1|Q16610
cor=0.86, p=3.6e−11
2398: FETUB|Q9UGM5
561: FETUB|Q9UGM5
cor=0.79, p=1.7e−08
2400: KLK7|P49862
1159: KLK7|P49862

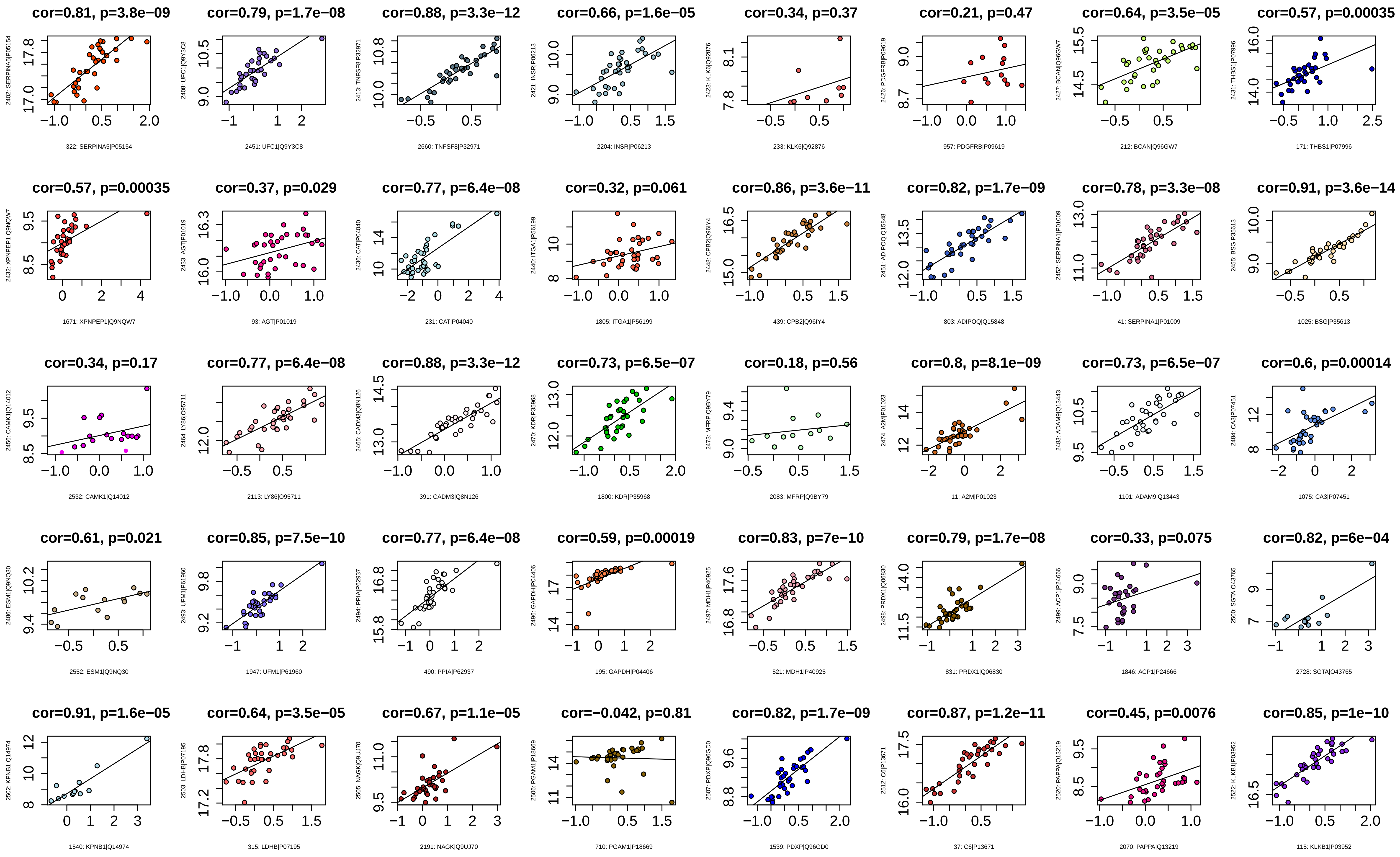
cor=0.81, p=3.8e−09
2402: SERPINA5|P05154
322: SERPINA5|P05154
cor=0.79, p=1.7e−08
2408: UFC1|Q9Y3C8
2451: UFC1|Q9Y3C8
cor=0.88, p=3.3e−12
2413: TNFSF8|P32971
2660: TNFSF8|P32971
cor=0.66, p=1.6e−05
2421: INSR|P06213
2204: INSR|P06213
cor=0.34, p=0.37
2423: KLK6|Q92876
233: KLK6|Q92876
cor=0.21, p=0.47
2426: PDGFRB|P09619
957: PDGFRB|P09619
cor=0.64, p=3.5e−05
2427: BCAN|Q96GW7
212: BCAN|Q96GW7
cor=0.57, p=0.00035
2431: THBS1|P07996
171: THBS1|P07996
cor=0.57, p=0.00035
2432: XPNPEP1|Q9NQW7
1671: XPNPEP1|Q9NQW7
cor=0.37, p=0.029
2433: AGT|P01019
93: AGT|P01019
cor=0.77, p=6.4e−08
2436: CAT|P04040
231: CAT|P04040
cor=0.32, p=0.061
2440: ITGA1|P56199
1805: ITGA1|P56199
cor=0.86, p=3.6e−11
2448: CPB2|Q96IY4
439: CPB2|Q96IY4
cor=0.82, p=1.7e−09
2451: ADIPOQ|Q15848
803: ADIPOQ|Q15848
cor=0.78, p=3.3e−08
2452: SERPINA1|P01009
41: SERPINA1|P01009
cor=0.91, p=3.6e−14
2455: BSG|P35613
1025: BSG|P35613
cor=0.34, p=0.17
2456: CAMK1|Q14012
2532: CAMK1|Q14012
cor=0.77, p=6.4e−08
2464: LY86|O95711
2113: LY86|O95711
cor=0.88, p=3.3e−12
2465: CADM3|Q8N126
391: CADM3|Q8N126
cor=0.73, p=6.5e−07
2470: KDR|P35968
1800: KDR|P35968
cor=0.18, p=0.56
2473: MFRP|Q9BY79
2083: MFRP|Q9BY79
cor=0.8, p=8.1e−09
2474: A2M|P01023
11: A2M|P01023
cor=0.73, p=6.5e−07
2483: ADAM9|Q13443
1101: ADAM9|Q13443
cor=0.6, p=0.00014
2484: CA3|P07451
1075: CA3|P07451
cor=0.61, p=0.021
2486: ESM1|Q9NQ30
2552: ESM1|Q9NQ30
cor=0.85, p=7.5e−10
2493: UFM1|P61960
1947: UFM1|P61960
cor=0.77, p=6.4e−08
2494: PPIA|P62937
490: PPIA|P62937
cor=0.59, p=0.00019
2496: GAPDH|P04406
195: GAPDH|P04406
cor=0.83, p=7e−10
2497: MDH1|P40925
521: MDH1|P40925
cor=0.79, p=1.7e−08
2498: PRDX1|Q06830
831: PRDX1|Q06830
cor=0.33, p=0.075
2499: ACP1|P24666
1846: ACP1|P24666
cor=0.82, p=6e−04
2500: SGTA|O43765
2728: SGTA|O43765
cor=0.91, p=1.6e−05
2502: KPNB1|Q14974
1540: KPNB1|Q14974
cor=0.64, p=3.5e−05
2503: LDHB|P07195
315: LDHB|P07195
cor=0.67, p=1.1e−05
2505: NAGK|Q9UJ70
2191: NAGK|Q9UJ70
cor=−0.042, p=0.81
2506: PGAM1|P18669
710: PGAM1|P18669
cor=0.82, p=1.7e−09
2507: PDXP|Q96GD0
1539: PDXP|Q96GD0
cor=0.87, p=1.2e−11
2512: C6|P13671
37: C6|P13671
cor=0.45, p=0.0076
2520: PAPPA|Q13219
2070: PAPPA|Q13219
cor=0.85, p=1e−10
2522: KLKB1|P03952
115: KLKB1|P03952

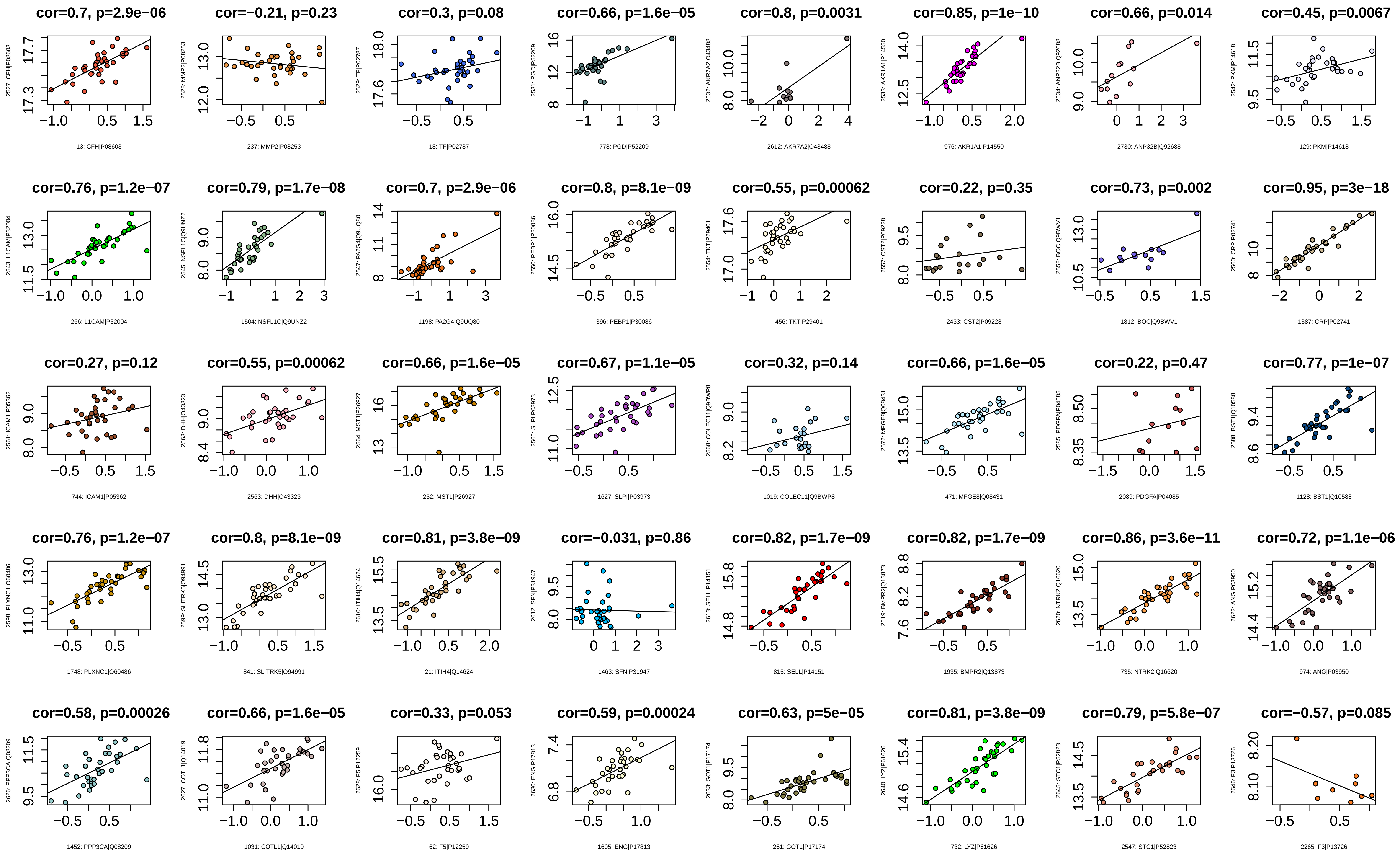
cor=0.7, p=2.9e−06
2527: CFH|P08603
13: CFH|P08603
cor=−0.21, p=0.23
2528: MMP2|P08253
237: MMP2|P08253
cor=0.3, p=0.08
2529: TF|P02787
18: TF|P02787
cor=0.66, p=1.6e−05
2531: PGD|P52209
778: PGD|P52209
cor=0.8, p=0.0031
2532: AKR7A2|O43488
2612: AKR7A2|O43488
cor=0.85, p=1e−10
2533: AKR1A1|P14550
976: AKR1A1|P14550
cor=0.66, p=0.014
2534: ANP32B|Q92688
2730: ANP32B|Q92688
cor=0.45, p=0.0067
2542: PKM|P14618
129: PKM|P14618
cor=0.76, p=1.2e−07
2543: L1CAM|P32004
266: L1CAM|P32004
cor=0.79, p=1.7e−08
2545: NSFL1C|Q9UNZ2
1504: NSFL1C|Q9UNZ2
cor=0.7, p=2.9e−06
2547: PA2G4|Q9UQ80
1198: PA2G4|Q9UQ80
cor=0.8, p=8.1e−09
2550: PEBP1|P30086
396: PEBP1|P30086
cor=0.55, p=0.00062
2554: TKT|P29401
456: TKT|P29401
cor=0.22, p=0.35
2557: CST2|P09228
2433: CST2|P09228
cor=0.73, p=0.002
2558: BOC|Q9BWV1
1812: BOC|Q9BWV1
cor=0.95, p=3e−18
2560: CRP|P02741
1387: CRP|P02741
cor=0.27, p=0.12
2561: ICAM1|P05362
744: ICAM1|P05362
cor=0.55, p=0.00062
2563: DHH|O43323
2563: DHH|O43323
cor=0.66, p=1.6e−05
2564: MST1|P26927
252: MST1|P26927
cor=0.67, p=1.1e−05
2565: SLPI|P03973
1627: SLPI|P03973
cor=0.32, p=0.14
2568: COLEC11|Q9BWP8
1019: COLEC11|Q9BWP8
cor=0.66, p=1.6e−05
2572: MFGE8|Q08431
471: MFGE8|Q08431
cor=0.22, p=0.47
2585: PDGFA|P04085
2089: PDGFA|P04085
cor=0.77, p=1e−07
2588: BST1|Q10588
1128: BST1|Q10588
cor=0.76, p=1.2e−07
2598: PLXNC1|O60486
1748: PLXNC1|O60486
cor=0.8, p=8.1e−09
2599: SLITRK5|O94991
841: SLITRK5|O94991
cor=0.81, p=3.8e−09
2610: ITIH4|Q14624
21: ITIH4|Q14624
cor=−0.031, p=0.86
2612: SFN|P31947
1463: SFN|P31947
cor=0.82, p=1.7e−09
2613: SELL|P14151
815: SELL|P14151
cor=0.82, p=1.7e−09
2619: BMPR2|Q13873
1935: BMPR2|Q13873
cor=0.86, p=3.6e−11
2620: NTRK2|Q16620
735: NTRK2|Q16620
cor=0.72, p=1.1e−06
2622: ANG|P03950
974: ANG|P03950
cor=0.58, p=0.00026
2626: PPP3CA|Q08209
1452: PPP3CA|Q08209
cor=0.66, p=1.6e−05
2627: COTL1|Q14019
1031: COTL1|Q14019
cor=0.33, p=0.053
2628: F5|P12259
62: F5|P12259
cor=0.59, p=0.00024
2630: ENG|P17813
1605: ENG|P17813
cor=0.63, p=5e−05
2633: GOT1|P17174
261: GOT1|P17174
cor=0.81, p=3.8e−09
2640: LYZ|P61626
732: LYZ|P61626
cor=0.79, p=5.8e−07
2645: STC1|P52823
2547: STC1|P52823
cor=−0.57, p=0.085
2646: F3|P13726
2265: F3|P13726

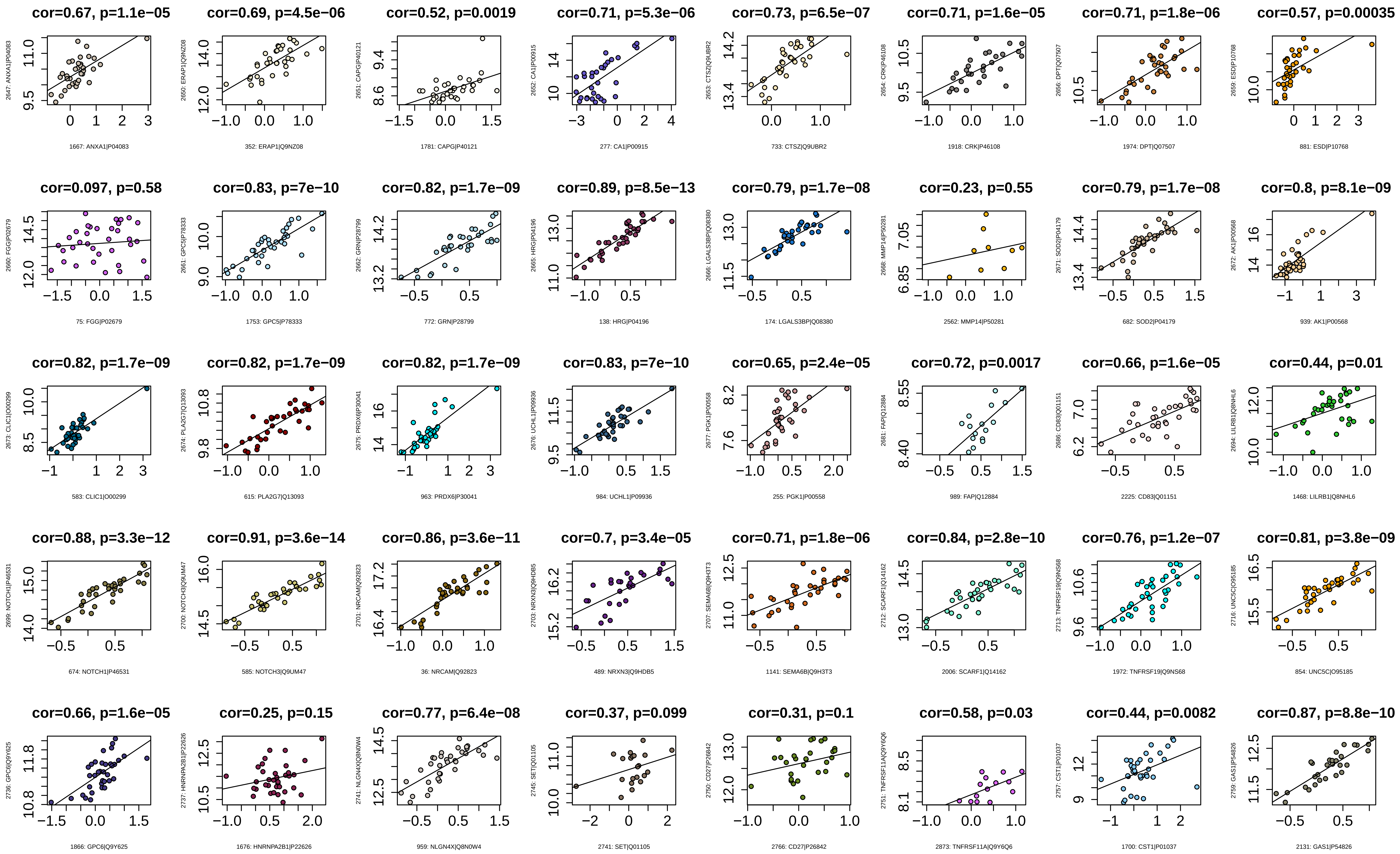

cor=0.67, p=1.1e−05
1667: ANXA1|P04083
cor=0.69, p=4.5e−06
352: ERAP1|Q9NZ08
cor=0.52, p=0.0019
1781: CAPG|P40121
cor=0.71, p=5.3e−06
277: CA1|P00915
cor=0.73, p=6.5e−07
733: CTSZ|Q9UBR2
cor=0.71, p=1.6e−05
1918: CRK|P46108
cor=0.71, p=1.8e−06
1974: DPT|Q07507
cor=0.57, p=0.00035
881: ESD|P10768
cor=0.097, p=0.58
75: FGG|P02679
cor=0.83, p=7e−10
1753: GPC5|P78333
cor=0.82, p=1.7e−09
772: GRN|P28799
cor=0.89, p=8.5e−13
138: HRG|P04196
cor=0.79, p=1.7e−08
174: LGALS3BP|Q08380
cor=0.23, p=0.55
2562: MMP14|P50281
cor=0.79, p=1.7e−08
682: SOD2|P04179
cor=0.8, p=8.1e−09
939: AK1|P00568
cor=0.82, p=1.7e−09
583: CLIC1|O00299
cor=0.82, p=1.7e−09
615: PLA2G7|Q13093
cor=0.82, p=1.7e−09
963: PRDX6|P30041
cor=0.83, p=7e−10
984: UCHL1|P09936
cor=0.65, p=2.4e−05
255: PGK1|P00558
cor=0.72, p=0.0017
989: FAP|Q12884
cor=0.66, p=1.6e−05
2225: CD83|Q01151
cor=0.44, p=0.01
1468: LILRB1|Q8NHL6
cor=0.88, p=3.3e−12
674: NOTCH1|P46531
cor=0.91, p=3.6e−14
585: NOTCH3|Q9UM47
cor=0.86, p=3.6e−11
36: NRCAM|Q92823
cor=0.7, p=3.4e−05
489: NRXN3|Q9HDB5
cor=0.71, p=1.8e−06
1141: SEMA6B|Q9H3T3
cor=0.84, p=2.8e−10
2006: SCARF1|Q14162
cor=0.76, p=1.2e−07
1972: TNFRSF19|Q9NS68
cor=0.81, p=3.8e−09
854: UNC5C|O95185
cor=0.66, p=1.6e−05
1866: GPC6|Q9Y625
cor=0.25, p=0.15
1676: HNRNPA2B1|P22626
cor=0.77, p=6.4e−08
959: NLGN4X|Q8N0W4
cor=0.37, p=0.099
2741: SET|Q01105
cor=0.31, p=0.1
2766: CD27|P26842
cor=0.58, p=0.03
2873: TNFRSF11A|Q9Y6Q6
cor=0.44, p=0.0082
1700: CST1|P01037
cor=0.87, p=8.8e−10
2131: GAS1|P54826

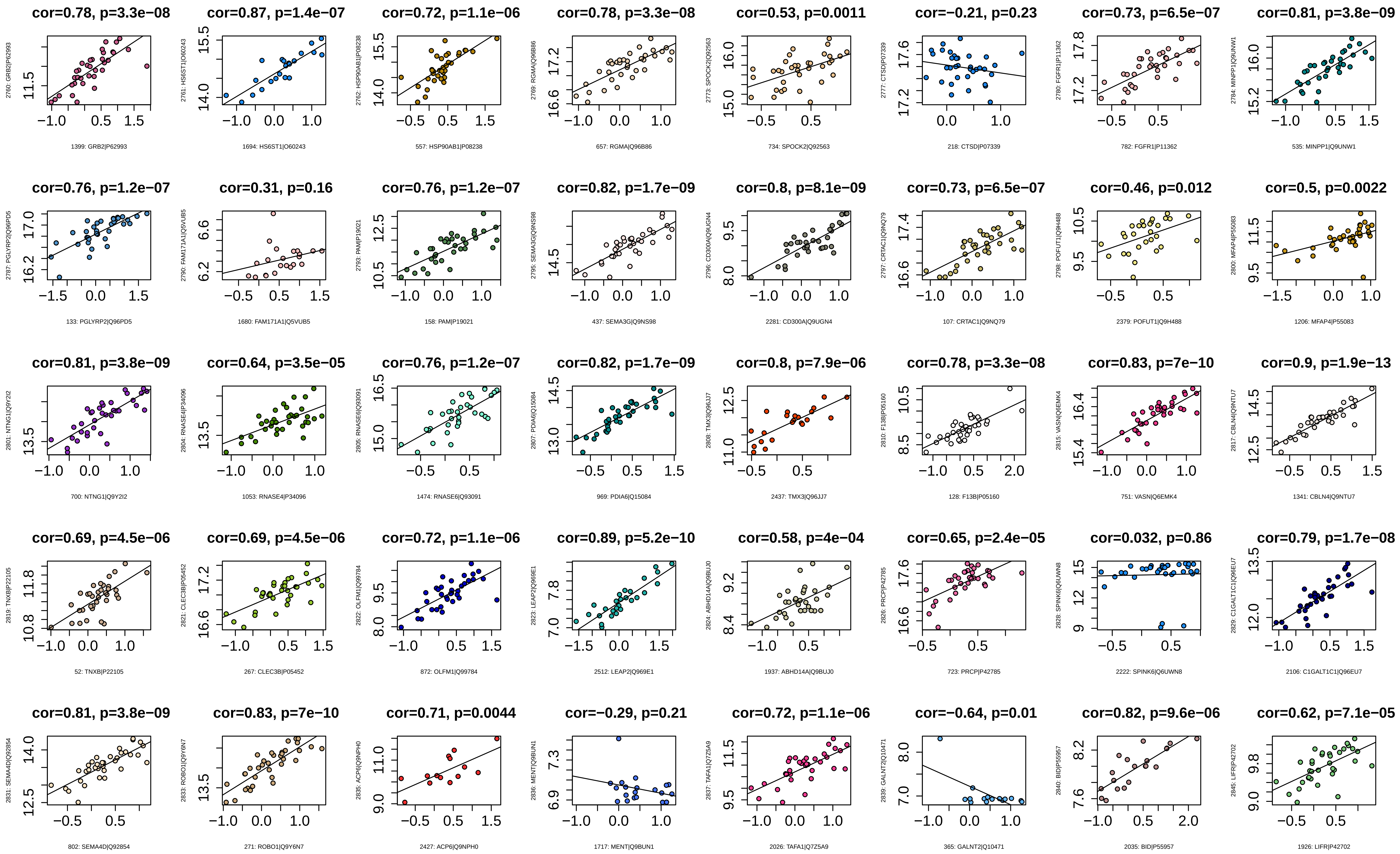
cor=0.78, p=3.3e−08
1399: GRB2|P62993
cor=0.87, p=1.4e−07
1694: HS6ST1|O60243
cor=0.72, p=1.1e−06
557: HSP90AB1|P08238
cor=0.78, p=3.3e−08
657: RGMA|Q96B86
cor=0.53, p=0.0011
734: SPOCK2|Q92563
cor=−0.21, p=0.23
218: CTSD|P07339
cor=0.73, p=6.5e−07
782: FGFR1|P11362
cor=0.81, p=3.8e−09
535: MINPP1|Q9UNW1
cor=0.76, p=1.2e−07
133: PGLYRP2|Q96PD5
cor=0.31, p=0.16
1680: FAM171A1|Q5VUB5
cor=0.76, p=1.2e−07
158: PAM|P19021
cor=0.82, p=1.7e−09
437: SEMA3G|Q9NS98
cor=0.8, p=8.1e−09
2281: CD300A|Q9UGN4
cor=0.73, p=6.5e−07
107: CRTAC1|Q9NQ79
cor=0.46, p=0.012
2379: POFUT1|Q9H488
cor=0.5, p=0.0022
1206: MFAP4|P55083
cor=0.81, p=3.8e−09
700: NTNG1|Q9Y2I2
cor=0.64, p=3.5e−05
1053: RNASE4|P34096
cor=0.76, p=1.2e−07
1474: RNASE6|Q93091
cor=0.82, p=1.7e−09
969: PDIA6|Q15084
cor=0.8, p=7.9e−06
2437: TMX3|Q96JJ7
cor=0.78, p=3.3e−08
128: F13B|P05160
cor=0.83, p=7e−10
751: VASN|Q6EMK4
cor=0.9, p=1.9e−13
1341: CBLN4|Q9NTU7
cor=0.69, p=4.5e−06
52: TNXB|P22105
cor=0.69, p=4.5e−06
267: CLEC3B|P05452
cor=0.72, p=1.1e−06
872: OLFM1|Q99784
cor=0.89, p=5.2e−10
2512: LEAP2|Q969E1
cor=0.58, p=4e−04
1937: ABHD14A|Q9BUJ0
cor=0.65, p=2.4e−05
723: PRCP|P42785
cor=0.032, p=0.86
2222: SPINK6|Q6UWN8
cor=0.79, p=1.7e−08
2106: C1GALT1C1|Q96EU7
cor=0.81, p=3.8e−09
802: SEMA4D|Q92854
cor=0.83, p=7e−10
271: ROBO1|Q9Y6N7
cor=0.71, p=0.0044
2427: ACP6|Q9NPH0
cor=−0.29, p=0.21
1717: MENT|Q9BUN1
cor=0.72, p=1.1e−06
2026: TAFA1|Q7Z5A9
cor=−0.64, p=0.01
365: GALNT2|Q10471
cor=0.82, p=9.6e−06
2035: BID|P55957
cor=0.62, p=7.1e−05
1926: LIFR|P42702

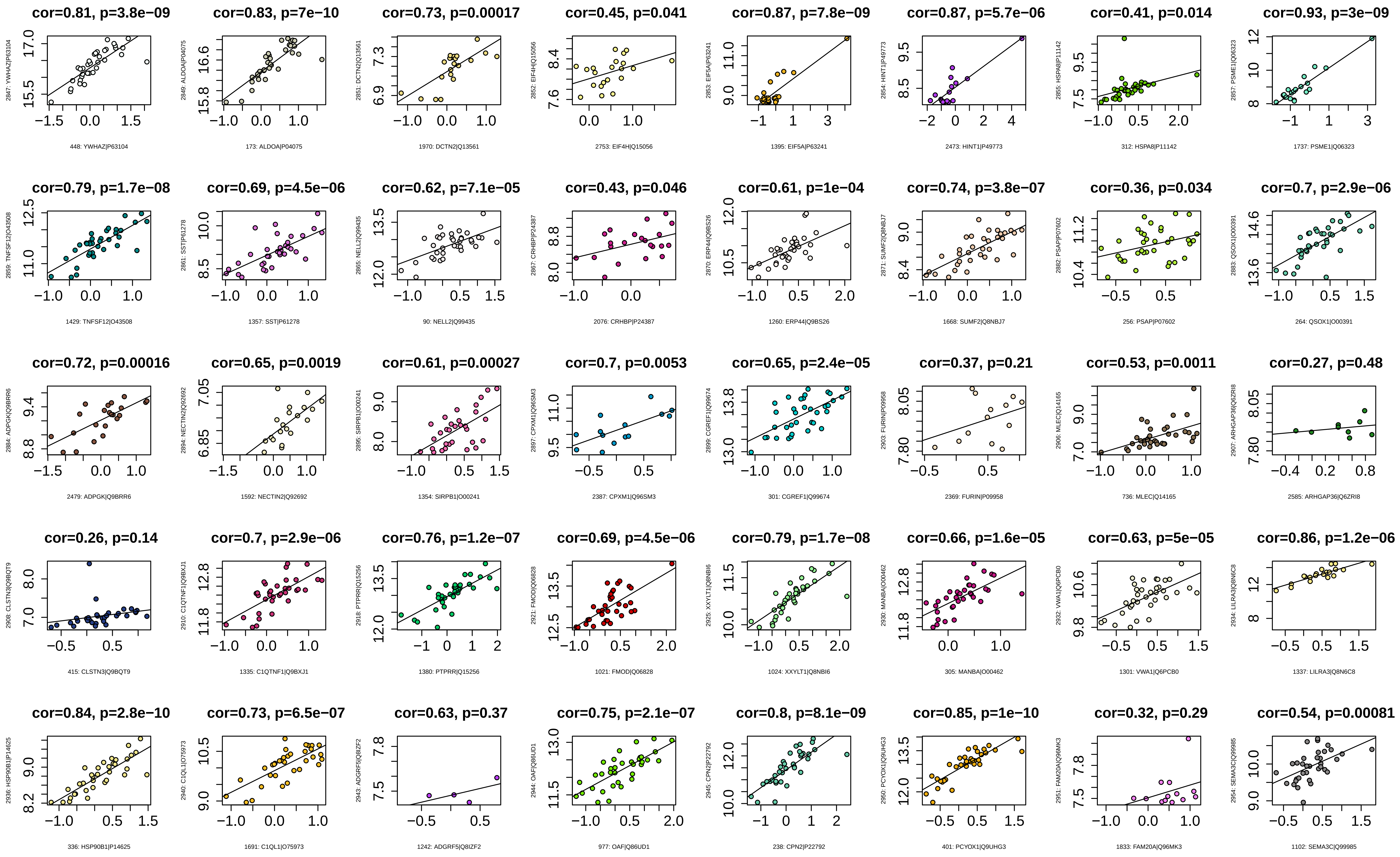
cor=0.81, p=3.8e−09
2847: YWHAZ|P63104
448: YWHAZ|P63104
cor=0.83, p=7e−10
2849: ALDOA|P04075
173: ALDOA|P04075
cor=0.73, p=0.00017
2851: DCTN2|Q13561
1970: DCTN2|Q13561
cor=0.45, p=0.041
2852: EIF4H|Q15056
2753: EIF4H|Q15056
cor=0.87, p=7.8e−09
2853: EIF5A|P63241
1395: EIF5A|P63241
cor=0.87, p=5.7e−06
2854: HINT1|P49773
2473: HINT1|P49773
cor=0.41, p=0.014
2855: HSPA8|P11142
312: HSPA8|P11142
cor=0.93, p=3e−09
2857: PSME1|Q06323
1737: PSME1|Q06323
cor=0.79, p=1.7e−08
2859: TNFSF12|O43508
1429: TNFSF12|O43508
cor=0.69, p=4.5e−06
2861: SST|P61278
1357: SST|P61278
cor=0.62, p=7.1e−05
2865: NELL2|Q99435
90: NELL2|Q99435
cor=0.43, p=0.046
2867: CRHBP|P24387
2076: CRHBP|P24387
cor=0.61, p=1e−04
2870: ERP44|Q9BS26
1260: ERP44|Q9BS26
cor=0.74, p=3.8e−07
2871: SUMF2|Q8NBJ7
1668: SUMF2|Q8NBJ7
cor=0.36, p=0.034
2882: PSAP|P07602
256: PSAP|P07602
cor=0.7, p=2.9e−06
2883: QSOX1|O00391
264: QSOX1|O00391
cor=0.72, p=0.00016
2884: ADPGK|Q9BRR6
2479: ADPGK|Q9BRR6
cor=0.65, p=0.0019
2894: NECTIN2|Q92692
1592: NECTIN2|Q92692
cor=0.61, p=0.00027
2895: SIRPB1|O00241
1354: SIRPB1|O00241
cor=0.7, p=0.0053
2897: CPXM1|Q96SM3
2387: CPXM1|Q96SM3
cor=0.65, p=2.4e−05
2899: CGREF1|Q99674
301: CGREF1|Q99674
cor=0.37, p=0.21
2903: FURIN|P09958
2369: FURIN|P09958
cor=0.53, p=0.0011
2906: MLEC|Q14165
736: MLEC|Q14165
cor=0.27, p=0.48
2907: ARHGAP36|Q6ZRI8
2585: ARHGAP36|Q6ZRI8
cor=0.26, p=0.14
2908: CLSTN3|Q9BQT9
415: CLSTN3|Q9BQT9
cor=0.7, p=2.9e−06
2910: C1QTNF1|Q9BXJ1
1335: C1QTNF1|Q9BXJ1
cor=0.76, p=1.2e−07
2918: PTPRR|Q15256
1380: PTPRR|Q15256
cor=0.69, p=4.5e−06
2921: FMOD|Q06828
1021: FMOD|Q06828
cor=0.79, p=1.7e−08
2925: XXYLT1|Q8NBI6
1024: XXYLT1|Q8NBI6
cor=0.66, p=1.6e−05
2930: MANBA|O00462
305: MANBA|O00462
cor=0.63, p=5e−05
2932: VWA1|Q6PCB0
1301: VWA1|Q6PCB0
cor=0.86, p=1.2e−06
2934: LILRA3|Q8N6C8
1337: LILRA3|Q8N6C8
cor=0.84, p=2.8e−10
2936: HSP90B1|P14625
336: HSP90B1|P14625
cor=0.73, p=6.5e−07
2940: C1QL1|O75973
1691: C1QL1|O75973
cor=0.63, p=0.37
2943: ADGRF5|Q8IZF2
1242: ADGRF5|Q8IZF2
cor=0.75, p=2.1e−07
2944: OAF|Q86UD1
977: OAF|Q86UD1
cor=0.8, p=8.1e−09
2945: CPN2|P22792
238: CPN2|P22792
cor=0.85, p=1e−10
2950: PCYOX1|Q9UHG3
401: PCYOX1|Q9UHG3
cor=0.32, p=0.29
2951: FAM20A|Q96MK3
1833: FAM20A|Q96MK3
cor=0.54, p=0.00081
2954: SEMA3C|Q99985
1102: SEMA3C|Q99985

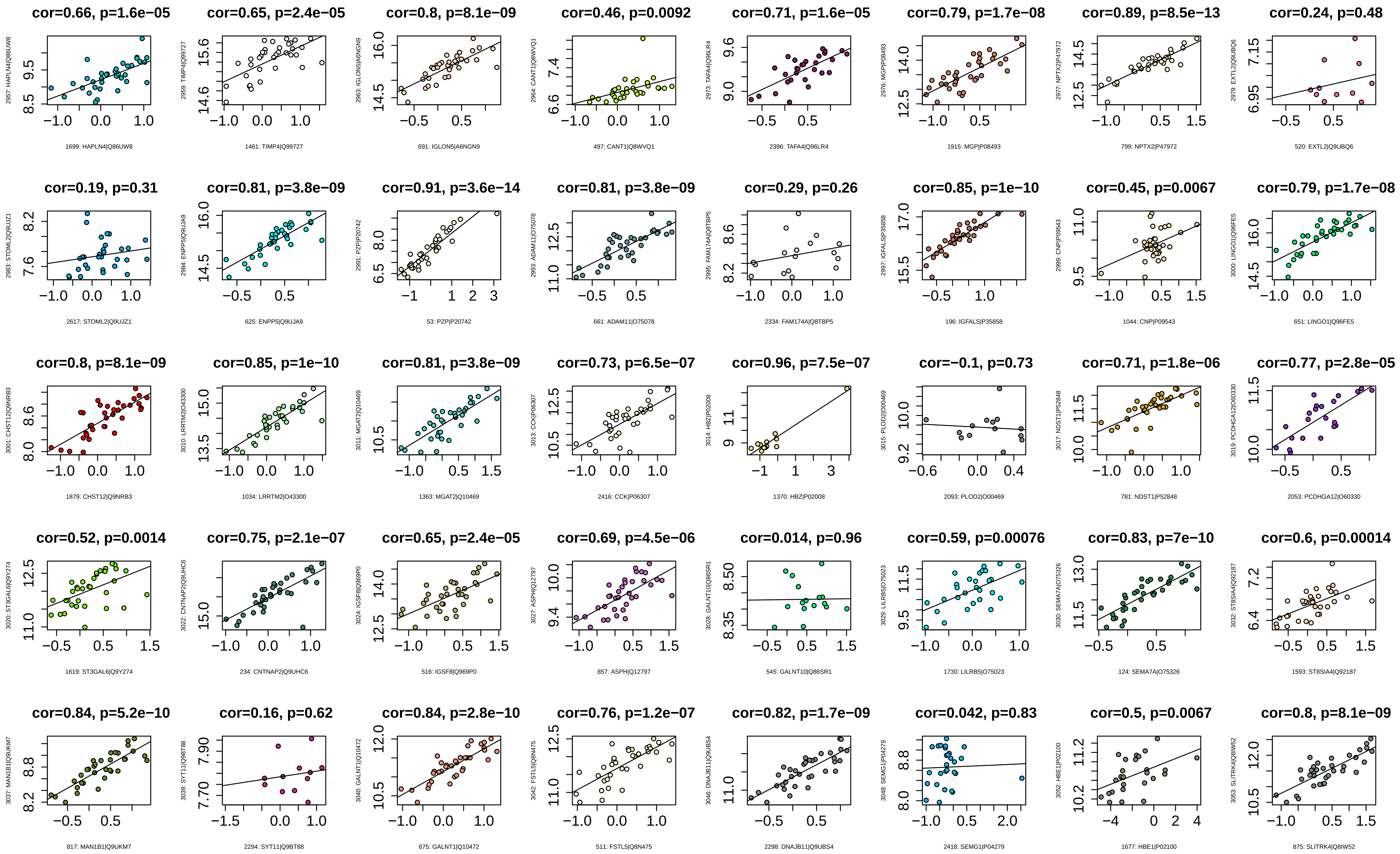
cor=0.66, p=1.6e−05
2957: HAPLN4|Q86UW8
1699: HAPLN4|Q86UW8
cor=0.65, p=2.4e−05
2959: TIMP4|Q99727
1461: TIMP4|Q99727
cor=0.8, p=8.1e−09
2963: IGLON5|A6NGN9
691: IGLON5|A6NGN9
cor=0.46, p=0.0092
2964: CANT1|Q8WVQ1
497: CANT1|Q8WVQ1
cor=0.71, p=1.6e−05
2973: TAFA4|Q96LR4
2396: TAFA4|Q96LR4
cor=0.79, p=1.7e−08
2976: MGP|P08493
1915: MGP|P08493
cor=0.89, p=8.5e−13
2977: NPTX2|P47972
799: NPTX2|P47972
cor=0.24, p=0.48
2979: EXTL2|Q9UBQ6
520: EXTL2|Q9UBQ6
cor=0.19, p=0.31
2983: STOML2|Q9UJZ1
2617: STOML2|Q9UJZ1
cor=0.81, p=3.8e−09
2984: ENPP5|Q9UJA9
625: ENPP5|Q9UJA9
cor=0.91, p=3.6e−14
2991: PZP|P20742
53: PZP|P20742
cor=0.81, p=3.8e−09
2993: ADAM11|O75078
661: ADAM11|O75078
cor=0.29, p=0.26
2995: FAM174A|Q8TBP5
2334: FAM174A|Q8TBP5
cor=0.85, p=1e−10
2997: IGFALS|P35858
196: IGFALS|P35858
cor=0.45, p=0.0067
2999: CNP|P09543
1044: CNP|P09543
cor=0.79, p=1.7e−08
3000: LINGO1|Q96FE5
651: LINGO1|Q96FE5
cor=0.8, p=8.1e−09
3001: CHST12|Q9NRB3
1879: CHST12|Q9NRB3
cor=0.85, p=1e−10
3010: LRRTM2|O43300
1034: LRRTM2|O43300
cor=0.81, p=3.8e−09
3011: MGAT2|Q10469
1363: MGAT2|Q10469
cor=0.73, p=6.5e−07
3013: CCK|P06307
2416: CCK|P06307
cor=0.96, p=7.5e−07
3014: HBZ|P02008
1370: HBZ|P02008
cor=−0.1, p=0.73
3015: PLOD2|O00469
2093: PLOD2|O00469
cor=0.71, p=1.8e−06
3017: NDST1|P52848
781: NDST1|P52848
cor=0.77, p=2.8e−05
3019: PCDHGA12|O60330
2053: PCDHGA12|O60330
cor=0.52, p=0.0014
3020: ST3GAL6|Q9Y274
1619: ST3GAL6|Q9Y274
cor=0.75, p=2.1e−07
3022: CNTNAP2|Q9UHC6
234: CNTNAP2|Q9UHC6
cor=0.65, p=2.4e−05
3024: IGSF8|Q969P0
516: IGSF8|Q969P0
cor=0.69, p=4.5e−06
3027: ASPH|Q12797
857: ASPH|Q12797
cor=0.014, p=0.96
3028: GALNT10|Q86SR1
545: GALNT10|Q86SR1
cor=0.59, p=0.00076
3029: LILRB5|O75023
1730: LILRB5|O75023
cor=0.83, p=7e−10
3030: SEMA7A|O75326
124: SEMA7A|O75326
cor=0.6, p=0.00014
3032: ST8SIA4|Q92187
1593: ST8SIA4|Q92187
cor=0.84, p=5.2e−10
3037: MAN1B1|Q9UKM7
817: MAN1B1|Q9UKM7
cor=0.16, p=0.62
3039: SYT11|Q9BT88
2294: SYT11|Q9BT88
cor=0.84, p=2.8e−10
3040: GALNT1|Q10472
675: GALNT1|Q10472
cor=0.76, p=1.2e−07
3042: FSTL5|Q8N475
511: FSTL5|Q8N475
cor=0.82, p=1.7e−09
3046: DNAJB11|Q9UBS4
2298: DNAJB11|Q9UBS4
cor=0.042, p=0.83
3048: SEMG1|P04279
2418: SEMG1|P04279
cor=0.5, p=0.0067
3052: HBE1|P02100
1677: HBE1|P02100
cor=0.8, p=8.1e−09
3053: SLITRK4|Q8IW52
875: SLITRK4|Q8IW52

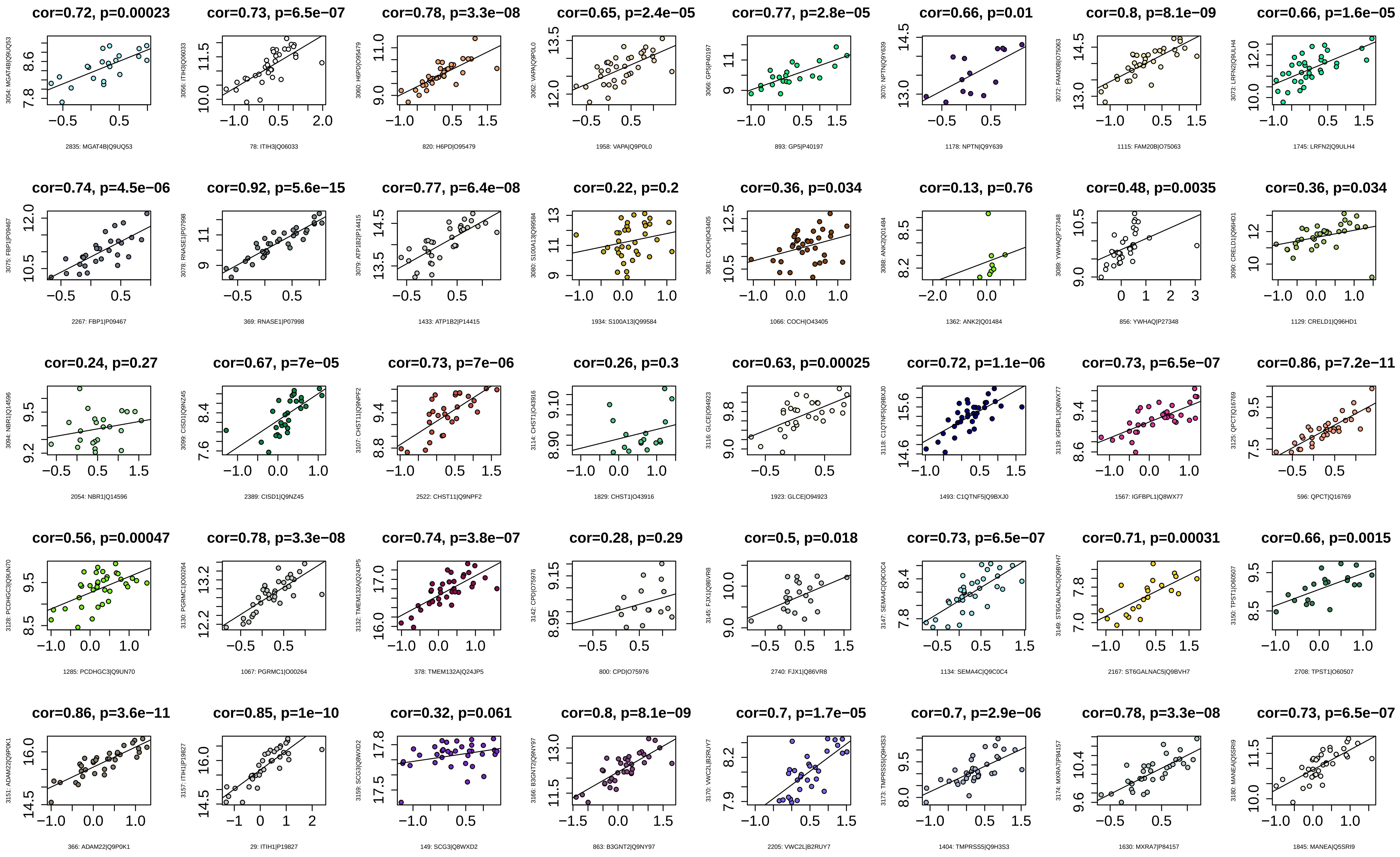
cor=0.72, p=0.00023
3054: MGAT4B|Q9UQ53
2835: MGAT4B|Q9UQ53
cor=0.73, p=6.5e−07
3056: ITIH3|Q06033
78: ITIH3|Q06033
cor=0.78, p=3.3e−08
3060: H6PD|O95479
820: H6PD|O95479
cor=0.65, p=2.4e−05
3062: VAPA|Q9P0L0
1958: VAPA|Q9P0L0
cor=0.77, p=2.8e−05
3066: GP5|P40197
893: GP5|P40197
cor=0.66, p=0.01
3070: NPTN|Q9Y639
1178: NPTN|Q9Y639
cor=0.8, p=8.1e−09
3072: FAM20B|O75063
1115: FAM20B|O75063
cor=0.66, p=1.6e−05
3073: LRFN2|Q9ULH4
1745: LRFN2|Q9ULH4
cor=0.74, p=4.5e−06
3075: FBP1|P09467
2267: FBP1|P09467
cor=0.92, p=5.6e−15
3078: RNASE1|P07998
369: RNASE1|P07998
cor=0.77, p=6.4e−08
3079: ATP1B2|P14415
1433: ATP1B2|P14415
cor=0.22, p=0.2
3080: S100A13|Q99584
1934: S100A13|Q99584
cor=0.36, p=0.034
3081: COCH|O43405
1066: COCH|O43405
cor=0.13, p=0.76
3088: ANK2|Q01484
1362: ANK2|Q01484
cor=0.48, p=0.0035
3089: YWHAQ|P27348
856: YWHAQ|P27348
cor=0.36, p=0.034
3090: CRELD1|Q96HD1
1129: CRELD1|Q96HD1
cor=0.24, p=0.27
3094: NBR1|Q14596
2054: NBR1|Q14596
cor=0.67, p=7e−05
3099: CISD1|Q9NZ45
2389: CISD1|Q9NZ45
cor=0.73, p=7e−06
3107: CHST11|Q9NPF2
2522: CHST11|Q9NPF2
cor=0.26, p=0.3
3114: CHST1|O43916
1829: CHST1|O43916
cor=0.63, p=0.00025
3116: GLCE|O94923
1923: GLCE|O94923
cor=0.72, p=1.1e−06
3118: C1QTNF5|Q9BXJ0
1493: C1QTNF5|Q9BXJ0
cor=0.73, p=6.5e−07
3119: IGFBPL1|Q8WX77
1567: IGFBPL1|Q8WX77
cor=0.86, p=7.2e−11
3125: QPCT|Q16769
596: QPCT|Q16769
cor=0.56, p=0.00047
3128: PCDHGC3|Q9UN70
1285: PCDHGC3|Q9UN70
cor=0.78, p=3.3e−08
3130: PGRMC1|O00264
1067: PGRMC1|O00264
cor=0.74, p=3.8e−07
3132: TMEM132A|Q24JP5
378: TMEM132A|Q24JP5
cor=0.28, p=0.29
3142: CPD|O75976
800: CPD|O75976
cor=0.5, p=0.018
3145: FJX1|Q86VR8
2740: FJX1|Q86VR8
cor=0.73, p=6.5e−07
3147: SEMA4C|Q9C0C4
1134: SEMA4C|Q9C0C4
cor=0.71, p=0.00031
3149: ST6GALNAC5|Q9BVH7
2167: ST6GALNAC5|Q9BVH7
cor=0.66, p=0.0015
3150: TPST1|O60507
2708: TPST1|O60507
cor=0.86, p=3.6e−11
3151: ADAM22|Q9P0K1
366: ADAM22|Q9P0K1
cor=0.85, p=1e−10
3157: ITIH1|P19827
29: ITIH1|P19827
cor=0.32, p=0.061
3159: SCG3|Q8WXD2
149: SCG3|Q8WXD2
cor=0.8, p=8.1e−09
3166: B3GNT2|Q9NY97
863: B3GNT2|Q9NY97
cor=0.7, p=1.7e−05
3170: VWC2L|B2RUY7
2205: VWC2L|B2RUY7
cor=0.7, p=2.9e−06
3173: TMPRSS5|Q9H3S3
1404: TMPRSS5|Q9H3S3
cor=0.78, p=3.3e−08
3174: MXRA7|P84157
1630: MXRA7|P84157
cor=0.73, p=6.5e−07
3180: MANEA|Q5SRI9
1845: MANEA|Q5SRI9

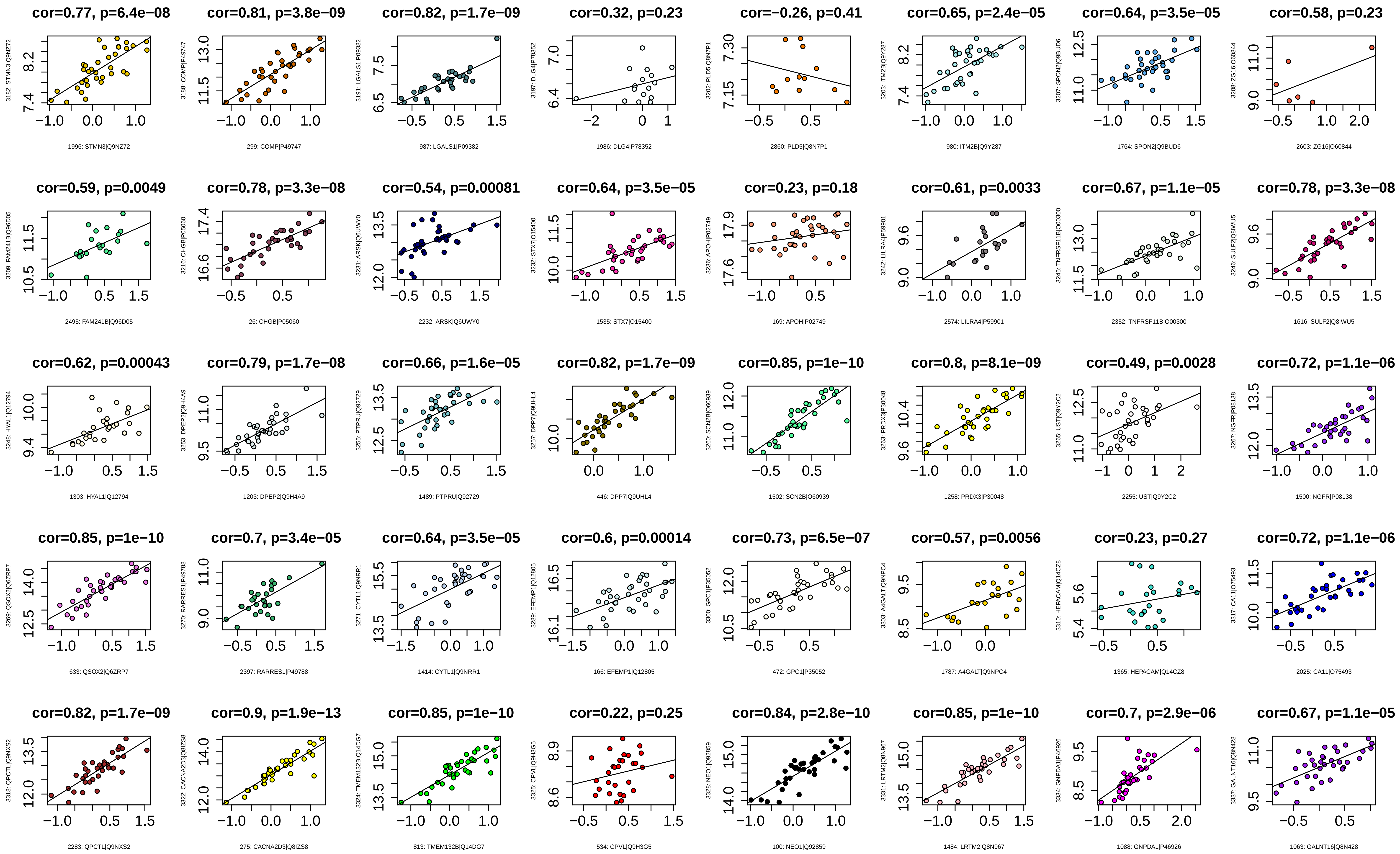
cor=0.77, p=6.4e−08
1996: STMN3|Q9NZ72
cor=0.81, p=3.8e−09
299: COMP|P49747
cor=0.82, p=1.7e−09
987: LGALS1|P09382
cor=0.32, p=0.23
1986: DLG4|P78352
cor=−0.26, p=0.41
2860: PLD5|Q8N7P1
cor=0.65, p=2.4e−05
980: ITM2B|Q9Y287
cor=0.64, p=3.5e−05
1764: SPON2|Q9BUD6
cor=0.58, p=0.23
2603: ZG16|O60844
cor=0.59, p=0.0049
2495: FAM241B|Q96D05
cor=0.78, p=3.3e−08
26: CHGB|P05060
cor=0.54, p=0.00081
2232: ARSK|Q6UWY0
cor=0.64, p=3.5e−05
1535: STX7|O15400
cor=0.23, p=0.18
169: APOH|P02749
cor=0.61, p=0.0033
2574: LILRA4|P59901
cor=0.67, p=1.1e−05
2352: TNFRSF11B|O00300
cor=0.78, p=3.3e−08
1616: SULF2|Q8IWU5
cor=0.62, p=0.00043
1303: HYAL1|Q12794
cor=0.79, p=1.7e−08
1203: DPEP2|Q9H4A9
cor=0.66, p=1.6e−05
1489: PTPRU|Q92729
cor=0.82, p=1.7e−09
446: DPP7|Q9UHL4
cor=0.85, p=1e−10
1502: SCN2B|O60939
cor=0.8, p=8.1e−09
1258: PRDX3|P30048
cor=0.49, p=0.0028
2255: UST|Q9Y2C2
cor=0.72, p=1.1e−06
1500: NGFR|P08138
cor=0.85, p=1e−10
633: QSOX2|Q6ZRP7
cor=0.7, p=3.4e−05
2397: RARRES1|P49788
cor=0.64, p=3.5e−05
1414: CYTL1|Q9NRR1
cor=0.6, p=0.00014
166: EFEMP1|Q12805
cor=0.73, p=6.5e−07
472: GPC1|P35052
cor=0.57, p=0.0056
1787: A4GALT|Q9NPC4
cor=0.23, p=0.27
1365: HEPACAM|Q14CZ8
cor=0.72, p=1.1e−06
2025: CA11|O75493
cor=0.82, p=1.7e−09
2283: QPCTL|Q9NXS2
cor=0.9, p=1.9e−13
275: CACNA2D3|Q8IZS8
cor=0.85, p=1e−10
813: TMEM132B|Q14DG7
cor=0.22, p=0.25
534: CPVL|Q9H3G5
cor=0.84, p=2.8e−10
100: NEO1|Q92859
cor=0.85, p=1e−10
1484: LRTM2|Q8N967
cor=0.7, p=2.9e−06
1088: GNPDA1|P46926
cor=0.67, p=1.1e−05
1063: GALNT16|Q8N428

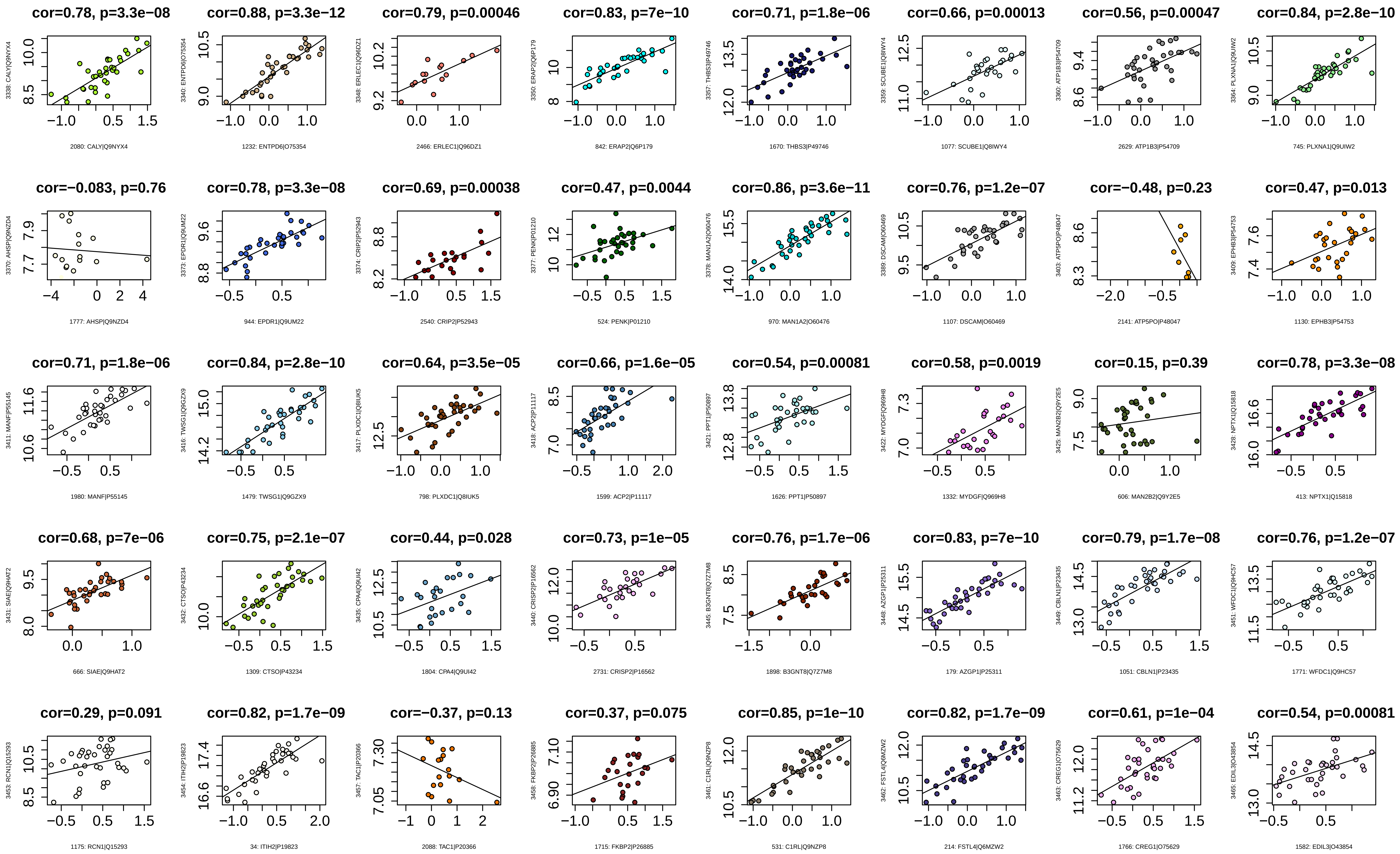
cor=0.78, p=3.3e−08
2080: CALY|Q9NYX4
cor=0.88, p=3.3e−12
1232: ENTPD6|O75354
cor=0.79, p=0.00046
2466: ERLEC1|Q96DZ1
cor=0.83, p=7e−10
842: ERAP2|Q6P179
cor=0.71, p=1.8e−06
1670: THBS3|P49746
cor=0.66, p=0.00013
1077: SCUBE1|Q8IWY4
cor=0.56, p=0.00047
2629: ATP1B3|P54709
cor=0.84, p=2.8e−10
745: PLXNA1|Q9UIW2
cor=−0.083, p=0.76
1777: AHSP|Q9NZD4
cor=0.78, p=3.3e−08
944: EPDR1|Q9UM22
cor=0.69, p=0.00038
2540: CRIP2|P52943
cor=0.47, p=0.0044
524: PENK|P01210
cor=0.86, p=3.6e−11
970: MAN1A2|O60476
cor=0.76, p=1.2e−07
1107: DSCAM|O60469
cor=−0.48, p=0.23
2141: ATP5PO|P48047
cor=0.47, p=0.013
1130: EPHB3|P54753
cor=0.71, p=1.8e−06
1980: MANF|P55145
cor=0.84, p=2.8e−10
1479: TWSG1|Q9GZX9
cor=0.64, p=3.5e−05
798: PLXDC1|Q8IUK5
cor=0.66, p=1.6e−05
1599: ACP2|P11117
cor=0.54, p=0.00081
1626: PPT1|P50897
cor=0.58, p=0.0019
1332: MYDGF|Q969H8
cor=0.15, p=0.39
606: MAN2B2|Q9Y2E5
cor=0.78, p=3.3e−08
413: NPTX1|Q15818
cor=0.68, p=7e−06
666: SIAE|Q9HAT2
cor=0.75, p=2.1e−07
1309: CTSO|P43234
cor=0.44, p=0.028
1804: CPA4|Q9UI42
cor=0.73, p=1e−05
2731: CRISP2|P16562
cor=0.76, p=1.7e−06
1898: B3GNT8|Q7Z7M8
cor=0.83, p=7e−10
179: AZGP1|P25311
cor=0.79, p=1.7e−08
1051: CBLN1|P23435
cor=0.76, p=1.2e−07
1771: WFDC1|Q9HC57
cor=0.29, p=0.091
1175: RCN1|Q15293
cor=0.82, p=1.7e−09
34: ITIH2|P19823
cor=−0.37, p=0.13
2088: TAC1|P20366
cor=0.37, p=0.075
1715: FKBP2|P26885
cor=0.85, p=1e−10
531: C1RL|Q9NZP8
cor=0.82, p=1.7e−09
214: FSTL4|Q6MZW2
cor=0.61, p=1e−04
1766: CREG1|O75629
cor=0.54, p=0.00081
1582: EDIL3|O43854

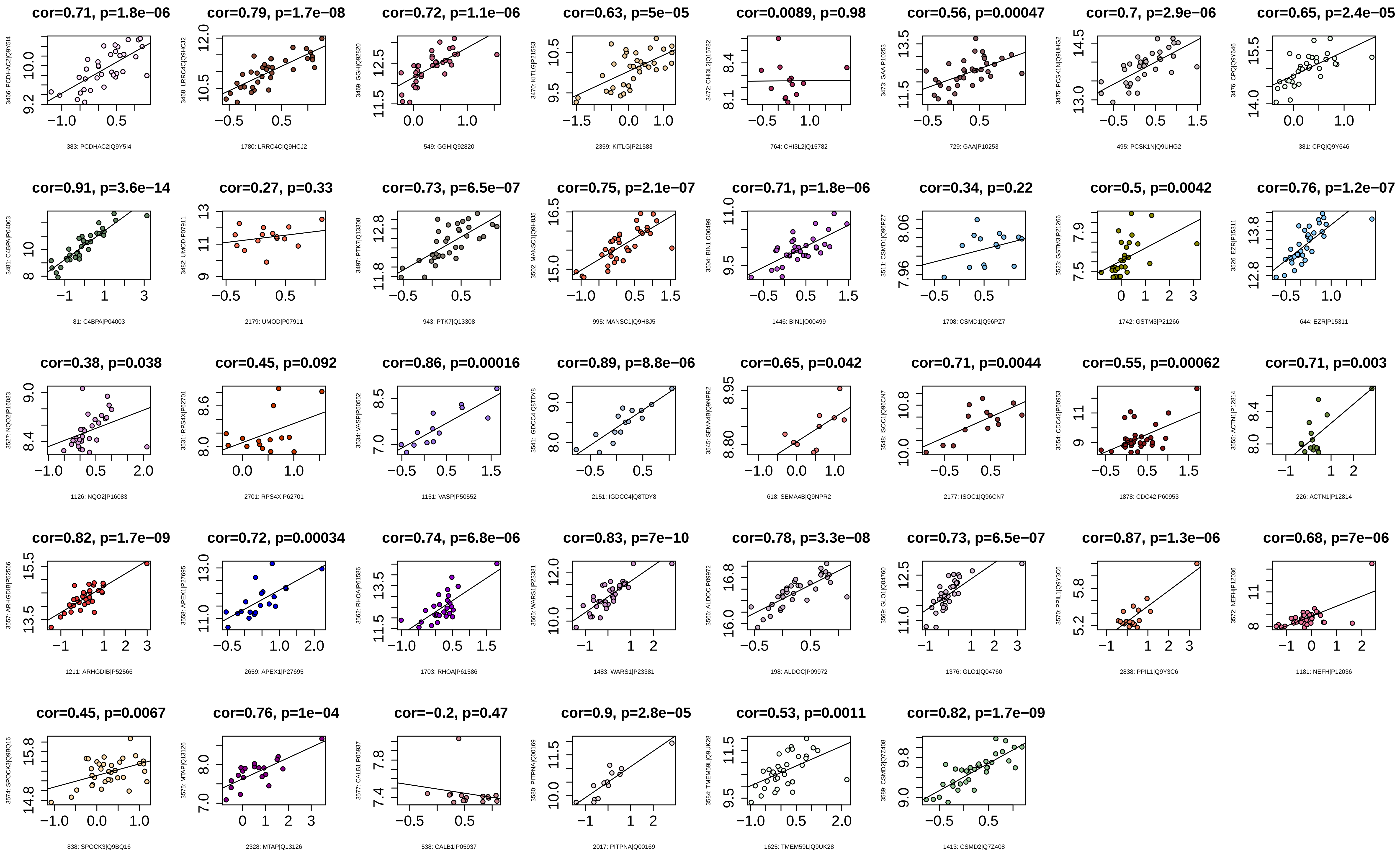
cor=0.71, p=1.8e−06
383: PCDHAC2|Q9Y5I4
cor=0.79, p=1.7e−08
1780: LRRC4C|Q9HCJ2
cor=0.72, p=1.1e−06
549: GGH|Q92820
cor=0.63, p=5e−05
2359: KITLG|P21583
cor=0.0089, p=0.98
764: CHI3L2|Q15782
cor=0.56, p=0.00047
729: GAA|P10253
cor=0.7, p=2.9e−06
495: PCSK1N|Q9UHG2
cor=0.65, p=2.4e−05
381: CPQ|Q9Y646
cor=0.91, p=3.6e−14
81: C4BPA|P04003
cor=0.27, p=0.33
2179: UMOD|P07911
cor=0.73, p=6.5e−07
943: PTK7|Q13308
cor=0.75, p=2.1e−07
995: MANSC1|Q9H8J5
cor=0.71, p=1.8e−06
1446: BIN1|O00499
cor=0.34, p=0.22
1708: CSMD1|Q96PZ7
cor=0.5, p=0.0042
1742: GSTM3|P21266
cor=0.76, p=1.2e−07
644: EZR|P15311
cor=0.38, p=0.038
1126: NQO2|P16083
cor=0.45, p=0.092
2701: RPS4X|P62701
cor=0.86, p=0.00016
1151: VASP|P50552
cor=0.89, p=8.8e−06
2151: IGDCC4|Q8TDY8
cor=0.65, p=0.042
618: SEMA4B|Q9NPR2
cor=0.71, p=0.0044
2177: ISOC1|Q96CN7
cor=0.55, p=0.00062
1878: CDC42|P60953
cor=0.71, p=0.003
226: ACTN1|P12814
cor=0.82, p=1.7e−09
1211: ARHGDIB|P52566
cor=0.72, p=0.00034
2659: APEX1|P27695
cor=0.74, p=6.8e−06
1703: RHOA|P61586
cor=0.83, p=7e−10
1483: WARS1|P23381
cor=0.78, p=3.3e−08
198: ALDOC|P09972
cor=0.73, p=6.5e−07
1376: GLO1|Q04760
cor=0.87, p=1.3e−06
2838: PPIL1|Q9Y3C6
cor=0.68, p=7e−06
1181: NEFH|P12036
cor=0.45, p=0.0067
838: SPOCK3|Q9BQ16
cor=0.76, p=1e−04
2328: MTAP|Q13126
cor=−0.2, p=0.47
538: CALB1|P05937
cor=0.9, p=2.8e−05
2017: PITPNA|Q00169
cor=0.53, p=0.0011
1625: TMEM59L|Q9UK28
cor=0.82, p=1.7e−09
1413: CSMD2|Q7Z408

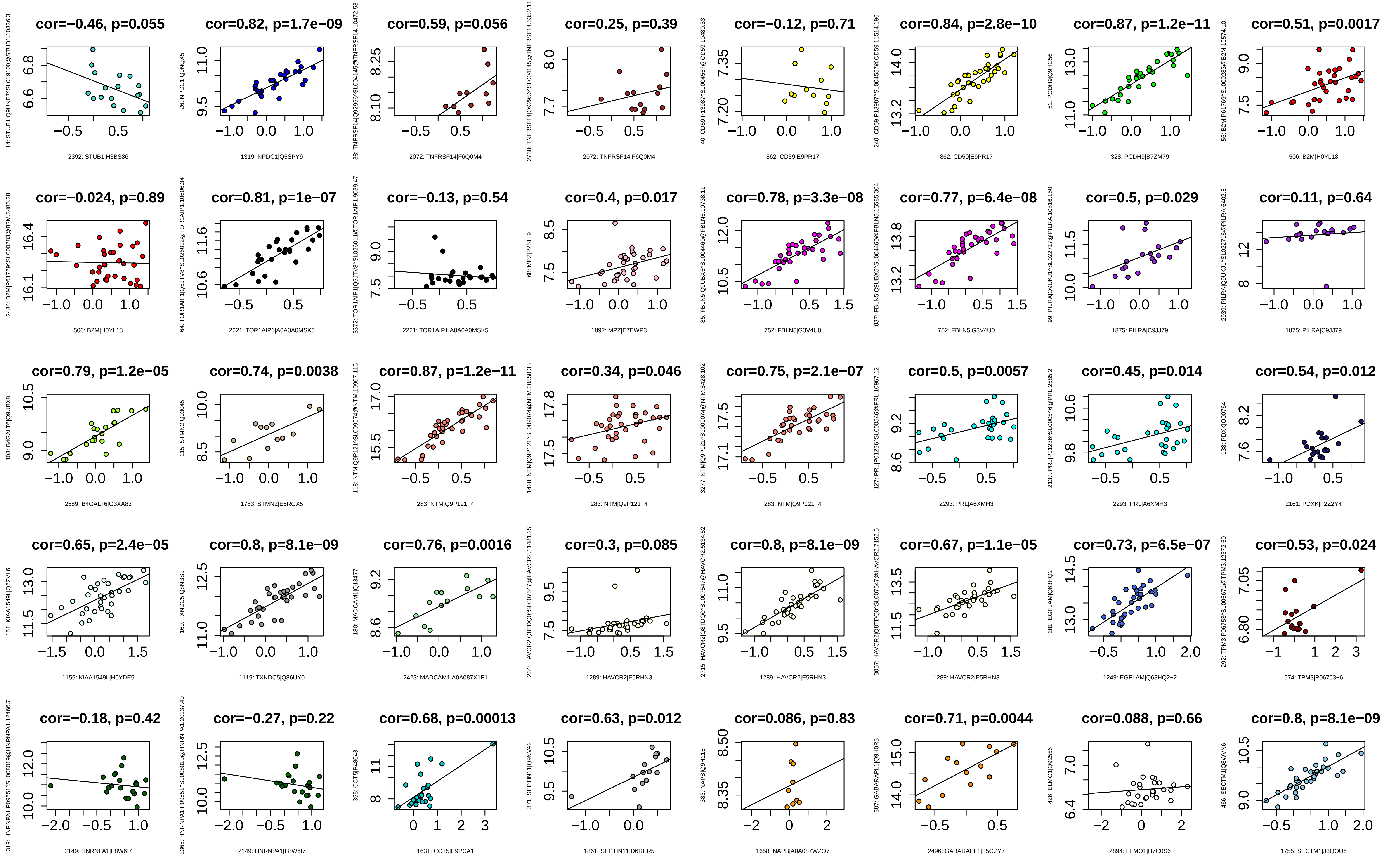
cor=−0.46, p=0.055
14: STUB1|Q9UNE7^SL019100@STUB1.10336.3
2392: STUB1|H3BS86
cor=0.82, p=1.7e−09
26: NPDC1|Q9NQX5
1319: NPDC1|Q5SPY9
cor=0.59, p=0.056
38: TNFRSF14|Q92956^SL004145@TNFRSF14.10472.53
2072: TNFRSF14|F6Q0M4
cor=0.25, p=0.39
2738: TNFRSF14|Q92956^SL004145@TNFRSF14.5352.11
2072: TNFRSF14|F6Q0M4
cor=−0.12, p=0.71
40: CD59|P13987^SL004557@CD59.10480.33
862: CD59|E9PR17
cor=0.84, p=2.8e−10
240: CD59|P13987^SL004557@CD59.11514.196
862: CD59|E9PR17
cor=0.87, p=1.2e−11
51: PCDH9|Q9HC56
328: PCDH9|B7ZM79
cor=0.51, p=0.0017
56: B2M|P61769^SL000283@B2M.10574.10
506: B2M|H0YL18
cor=−0.024, p=0.89
2434: B2M|P61769^SL000283@B2M.3485.28
506: B2M|H0YL18
cor=0.81, p=1e−07
64: TOR1AIP1|Q5JTV8^SL026012@TOR1AIP1.10606.34
2221: TOR1AIP1|A0A0A0MSK5
cor=−0.13, p=0.54
3372: TOR1AIP1|Q5JTV8^SL026011@TOR1AIP1.9039.47
2221: TOR1AIP1|A0A0A0MSK5
cor=0.4, p=0.017
68: MPZ|P25189
1892: MPZ|E7EWP3
cor=0.78, p=3.3e−08
85: FBLN5|Q9UBX5^SL004460@FBLN5.10738.11
752: FBLN5|G3V4U0
cor=0.77, p=6.4e−08
837: FBLN5|Q9UBX5^SL004460@FBLN5.15585.304
752: FBLN5|G3V4U0
cor=0.5, p=0.029
99: PILRA|Q9UKJ1^SL022717@PILRA.10816.150
1875: PILRA|C9JJ79
cor=0.11, p=0.64
2939: PILRA|Q9UKJ1^SL022716@PILRA.6402.8
1875: PILRA|C9JJ79
cor=0.79, p=1.2e−05
103: B4GALT6|Q9UBX8
2589: B4GALT6|G3XA83
cor=0.74, p=0.0038
115: STMN2|Q93045
1783: STMN2|E5RGX5
cor=0.87, p=1.2e−11
118: NTM|Q9P121^SL009074@NTM.10907.116
283: NTM|Q9P121−4
cor=0.34, p=0.046
1428: NTM|Q9P121^SL009074@NTM.20550.38
283: NTM|Q9P121−4
cor=0.75, p=2.1e−07
3277: NTM|Q9P121^SL009074@NTM.8428.102
283: NTM|Q9P121−4
cor=0.5, p=0.0057
127: PRL|P01236^SL000546@PRL.10967.12
2293: PRL|A6XMH3
cor=0.45, p=0.014
2137: PRL|P01236^SL000546@PRL.2585.2
2293: PRL|A6XMH3
cor=0.54, p=0.012
138: PDXK|O00764
2161: PDXK|F2Z2Y4
cor=0.65, p=2.4e−05
151: KIAA1549L|Q6ZVL6
1155: KIAA1549L|H0YDE5
cor=0.8, p=8.1e−09
169: TXNDC5|Q8NBS9
1119: TXNDC5|Q86UY0
cor=0.76, p=0.0016
180: MADCAM1|Q13477
2423: MADCAM1|A0A087X1F1
cor=0.3, p=0.085
234: HAVCR2|Q8TDQ0^SL007547@HAVCR2.11481.25
1289: HAVCR2|E5RHN3
cor=0.8, p=8.1e−09
2715: HAVCR2|Q8TDQ0^SL007547@HAVCR2.5134.52
1289: HAVCR2|E5RHN3
cor=0.67, p=1.1e−05
3057: HAVCR2|Q8TDQ0^SL007547@HAVCR2.7152.5
1289: HAVCR2|E5RHN3
cor=0.73, p=6.5e−07
281: EGFLAM|Q63HQ2
1249: EGFLAM|Q63HQ2−2
cor=0.53, p=0.024
292: TPM3|P06753^SL005671@TPM3.12372.50
574: TPM3|P06753−6
cor=−0.18, p=0.42
319: HNRNPA1|P09651^SL008019@HNRNPA1.12466.7
2149: HNRNPA1|F8W6I7
cor=−0.27, p=0.22
1365: HNRNPA1|P09651^SL008019@HNRNPA1.20137.49
2149: HNRNPA1|F8W6I7
cor=0.68, p=0.00013
355: CCT5|P48643
1631: CCT5|E9PCA1
cor=0.63, p=0.012
371: SEPTIN11|Q9NVA2
1861: SEPTIN11|D6RER5
cor=0.086, p=0.83
383: NAPB|Q9H115
1658: NAPB|A0A087WZQ7
cor=0.71, p=0.0044
387: GABARAPL1|Q9H0R8
2496: GABARAPL1|F5GZY7
cor=0.088, p=0.66
426: ELMO1|Q92556
2894: ELMO1|H7C0S6
cor=0.8, p=8.1e−09
466: SECTM1|Q8WVN6
1755: SECTM1|J3QQU6

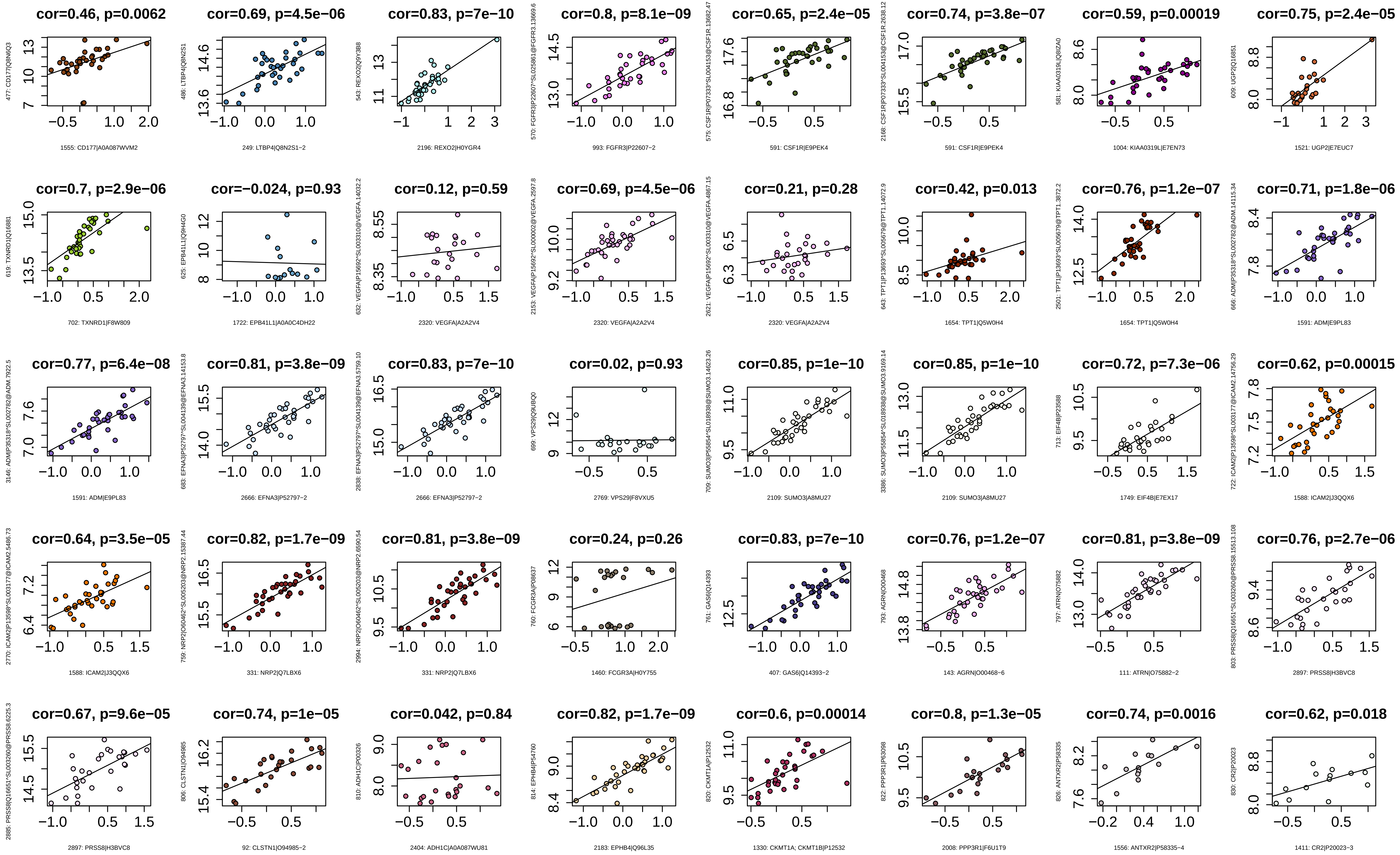
cor=0.46, p=0.0062
477: CD177|Q8N6Q3
1555: CD177|A0A087WVM2
cor=0.69, p=4.5e−06
486: LTBP4|Q8N2S1
249: LTBP4|Q8N2S1−2
cor=0.83, p=7e−10
543: REXO2|Q9Y3B8
2196: REXO2|H0YGR4
cor=0.8, p=8.1e−09
570: FGFR3|P22607^SL025861@FGFR3.13669.6
993: FGFR3|P22607−2
cor=0.65, p=2.4e−05
575: CSF1R|P07333^SL004153@CSF1R.13682.47
591: CSF1R|E9PEK4
cor=0.74, p=3.8e−07
2168: CSF1R|P07333^SL004153@CSF1R.2638.12
591: CSF1R|E9PEK4
cor=0.59, p=0.00019
581: KIAA0319L|Q8IZA0
1004: KIAA0319L|E7EN73
cor=0.75, p=2.4e−05
609: UGP2|Q16851
1521: UGP2|E7EUC7
cor=0.7, p=2.9e−06
619: TXNRD1|Q16881
702: TXNRD1|F8W809
cor=−0.024, p=0.93
625: EPB41L1|Q9H4G0
1722: EPB41L1|A0A0C4DH22
cor=0.12, p=0.59
632: VEGFA|P15692^SL003310@VEGFA.14032.2
2320: VEGFA|A2A2V4
cor=0.69, p=4.5e−06
2153: VEGFA|P15692^SL000002@VEGFA.2597.8
2320: VEGFA|A2A2V4
cor=0.21, p=0.28
2621: VEGFA|P15692^SL003310@VEGFA.4867.15
2320: VEGFA|A2A2V4
cor=0.42, p=0.013
643: TPT1|P13693^SL005679@TPT1.14072.9
1654: TPT1|Q5W0H4
cor=0.76, p=1.2e−07
2501: TPT1|P13693^SL005679@TPT1.3872.2
1654: TPT1|Q5W0H4
cor=0.71, p=1.8e−06
666: ADM|P35318^SL002782@ADM.14115.34
1591: ADM|E9PL83
cor=0.77, p=6.4e−08
3146: ADM|P35318^SL002782@ADM.7922.5
1591: ADM|E9PL83
cor=0.81, p=3.8e−09
683: EFNA3|P52797^SL004139@EFNA3.14153.8
2666: EFNA3|P52797−2
cor=0.83, p=7e−10
2838: EFNA3|P52797^SL004139@EFNA3.5759.10
2666: EFNA3|P52797−2
cor=0.02, p=0.93
699: VPS29|Q9UBQ0
2769: VPS29|F8VXU5
cor=0.85, p=1e−10
709: SUMO3|P55854^SL018938@SUMO3.14623.26
2109: SUMO3|A8MU27
cor=0.85, p=1e−10
3386: SUMO3|P55854^SL018938@SUMO3.9169.14
2109: SUMO3|A8MU27
cor=0.72, p=7.3e−06
713: EIF4B|P23588
1749: EIF4B|E7EX17
cor=0.62, p=0.00015
722: ICAM2|P13598^SL003177@ICAM2.14756.29
1588: ICAM2|J3QQX6
cor=0.64, p=3.5e−05
2770: ICAM2|P13598^SL003177@ICAM2.5486.73
1588: ICAM2|J3QQX6
cor=0.82, p=1.7e−09
759: NRP2|O60462^SL005203@NRP2.15387.44
331: NRP2|Q7LBX6
cor=0.81, p=3.8e−09
2994: NRP2|O60462^SL005203@NRP2.6590.54
331: NRP2|Q7LBX6
cor=0.24, p=0.26
760: FCGR3A|P08637
1460: FCGR3A|H0Y755
cor=0.83, p=7e−10
761: GAS6|Q14393
407: GAS6|Q14393−2
cor=0.76, p=1.2e−07
793: AGRN|O00468
143: AGRN|O00468−6
cor=0.81, p=3.8e−09
797: ATRN|O75882
111: ATRN|O75882−2
cor=0.76, p=2.7e−06
803: PRSS8|Q16651^SL003260@PRSS8.15513.108
2897: PRSS8|H3BVC8
cor=0.67, p=9.6e−05
2885: PRSS8|Q16651^SL003260@PRSS8.6225.3
2897: PRSS8|H3BVC8
cor=0.74, p=1e−05
806: CLSTN1|O94985
92: CLSTN1|O94985−2
cor=0.042, p=0.84
810: ADH1C|P00326
2404: ADH1C|A0A087WU81
cor=0.82, p=1.7e−09
814: EPHB4|P54760
2183: EPHB4|Q96L35
cor=0.6, p=0.00014
820: CKMT1A|P12532
1330: CKMT1A; CKMT1B|P12532
cor=0.8, p=1.3e−05
822: PPP3R1|P63098
2008: PPP3R1|F6U1T9
cor=0.74, p=0.0016
826: ANTXR2|P58335
1556: ANTXR2|P58335−4
cor=0.62, p=0.018
830: CR2|P20023
1411: CR2|P20023−3

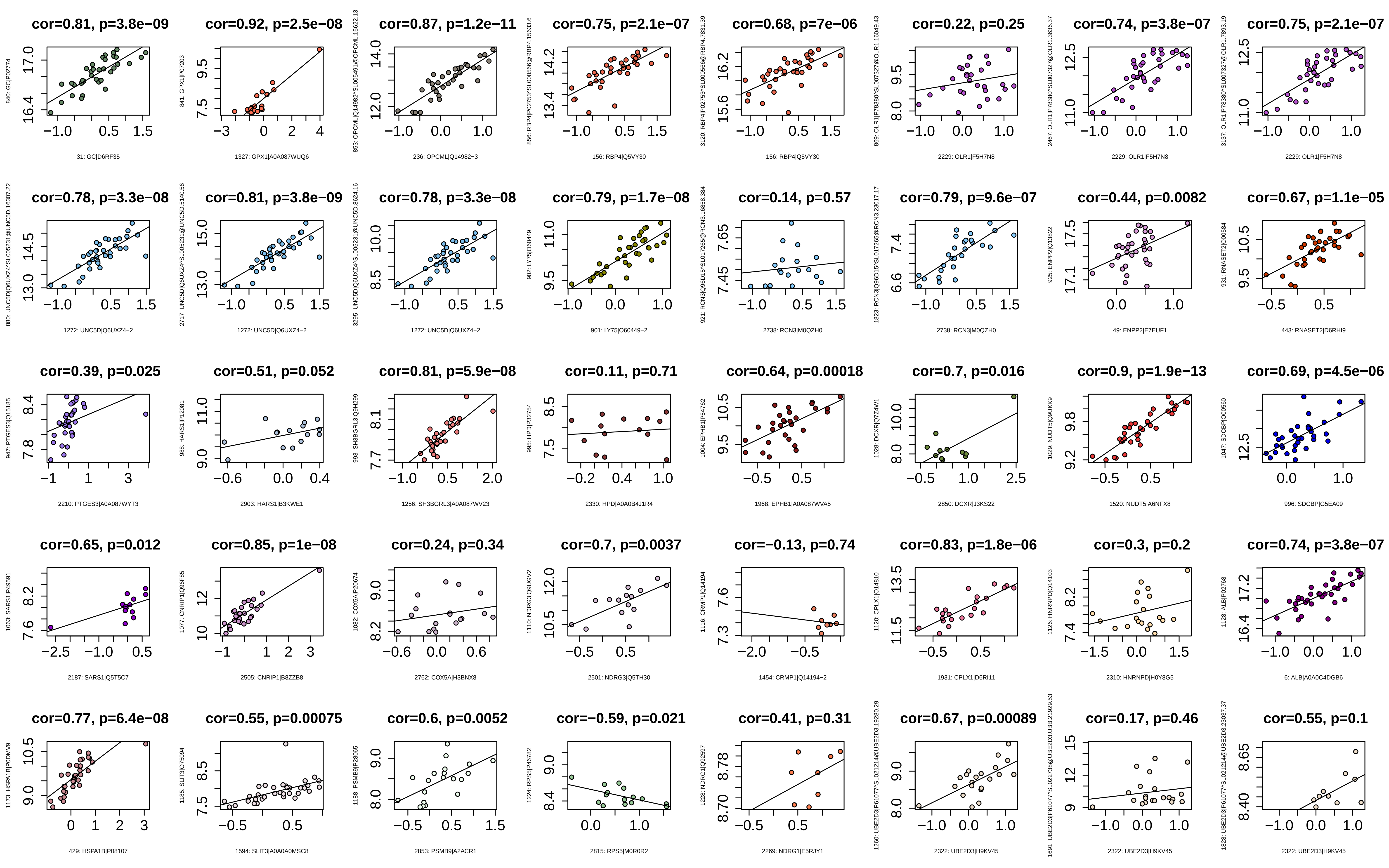

cor=0.81, p=3.8e−09
840: GC|P02774
31: GC|D6RF35
cor=0.92, p=2.5e−08
841: GPX1|P07203
1327: GPX1|A0A087WUQ6
cor=0.87, p=1.2e−11
853: OPCML|Q14982*SL005491@OPCML.15622.13
236: OPCML|Q14982−3
cor=0.75, p=2.1e−07
856: RBP4|P02753*SL000566@RBP4.15633.6
156: RBP4|Q5VY30
cor=0.68, p=7e−06
3120: RBP4|P02753*SL000566@RBP4.7831.39
156: RBP4|Q5VY30
cor=0.22, p=0.25
869: OLR1|P78380*SL007327@OLR1.16049.43
2229: OLR1|F5H7N8
cor=0.74, p=3.8e−07
2467: OLR1|P78380*SL007327@OLR1.3636.37
2229: OLR1|F5H7N8
cor=0.75, p=2.1e−07
3137: OLR1|P78380*SL007327@OLR1.7893.19
2229: OLR1|F5H7N8
cor=0.78, p=3.3e−08
880: UNC5D|Q6UXZ4*SL005231@UNC5D.16307.22
1272: UNC5D|Q6UXZ4−2
cor=0.81, p=3.8e−09
2717: UNC5D|Q6UXZ4*SL005231@UNC5D.5140.56
1272: UNC5D|Q6UXZ4−2
cor=0.78, p=3.3e−08
3295: UNC5D|Q6UXZ4*SL005231@UNC5D.8624.16
1272: UNC5D|Q6UXZ4−2
cor=0.79, p=1.7e−08
902: LY75|O60449
901: LY75|O60449−2
cor=0.14, p=0.57
921: RCN3|Q96D15*SL017265@RCN3.16858.384
2738: RCN3|M0QZH0
cor=0.79, p=9.6e−07
1823: RCN3|Q96D15*SL017265@RCN3.23017.17
2738: RCN3|M0QZH0
cor=0.44, p=0.0082
925: ENPP2|Q13822
49: ENPP2|E7EUF1
cor=0.67, p=1.1e−05
931: RNASET2|O00584
443: RNASET2|D6RHI9
cor=0.39, p=0.025
947: PTGES3|Q15185
2210: PTGES3|A0A087WYT3
cor=0.51, p=0.052
988: HARS1|P12081
2903: HARS1|B3KWE1
cor=0.81, p=5.9e−08
993: SH3BGRL3|Q9H299
1256: SH3BGRL3|A0A087WV23
cor=0.11, p=0.71
995: HPD|P32754
2330: HPD|A0A0B4J1R4
cor=0.64, p=0.00018
1004: EPHB1|P54762
1968: EPHB1|A0A087WVA5
cor=0.7, p=0.016
1028: DCXR|Q7Z4W1
2850: DCXR|J3KS22
cor=0.9, p=1.9e−13
1029: NUDT5|Q9UKK9
1520: NUDT5|A6NFX8
cor=0.69, p=4.5e−06
1047: SDCBP|O00560
996: SDCBP|G5EA09
cor=0.65, p=0.012
1063: SARS1|P49591
2187: SARS1|Q5T5C7
cor=0.85, p=1e−08
1077: CNRIP1|Q96F85
2505: CNRIP1|B8ZZB8
cor=0.24, p=0.34
1082: COX5A|P20674
2762: COX5A|H3BNX8
cor=0.7, p=0.0037
1110: NDRG3|Q9UGV2
2501: NDRG3|Q5TH30
cor=−0.13, p=0.74
1116: CRMP1|Q14194
1454: CRMP1|Q14194−2
cor=0.83, p=1.8e−06
1120: CPLX1|O14810
1931: CPLX1|D6RI11
cor=0.3, p=0.2
1126: HNRNPD|Q14103
2310: HNRNPD|H0Y8G5
cor=0.74, p=3.8e−07
1128: ALB|P02768
6: ALB|A0A0C4DGB6
cor=0.77, p=6.4e−08
1173: HSPA1B|P0DMV9
429: HSPA1B|P08107
cor=0.55, p=0.00075
1185: SLIT3|O75094
1594: SLIT3|A0A0A0MSC8
cor=0.6, p=0.0052
1188: PSMB9|P28065
2853: PSMB9|A2ACR1
cor=−0.59, p=0.021
1224: RPS5|P46782
2815: RPS5|M0R0R2
cor=0.41, p=0.31
1228: NDRG1|Q92597
2269: NDRG1|E5RJY1
cor=0.67, p=0.00089
1260: UBE2D3|P61077*SL021214@UBE2D3.19280.29
2322: UBE2D3|H9KV45
cor=0.17, p=0.46
1691: UBE2D3|P61077*.21929.53
2322: UBE2D3|H9KV45
cor=0.55, p=0.1
1828: UBE2D3|P61077*SL021214@UBE2D3.23037.37
2322: UBE2D3|H9KV45

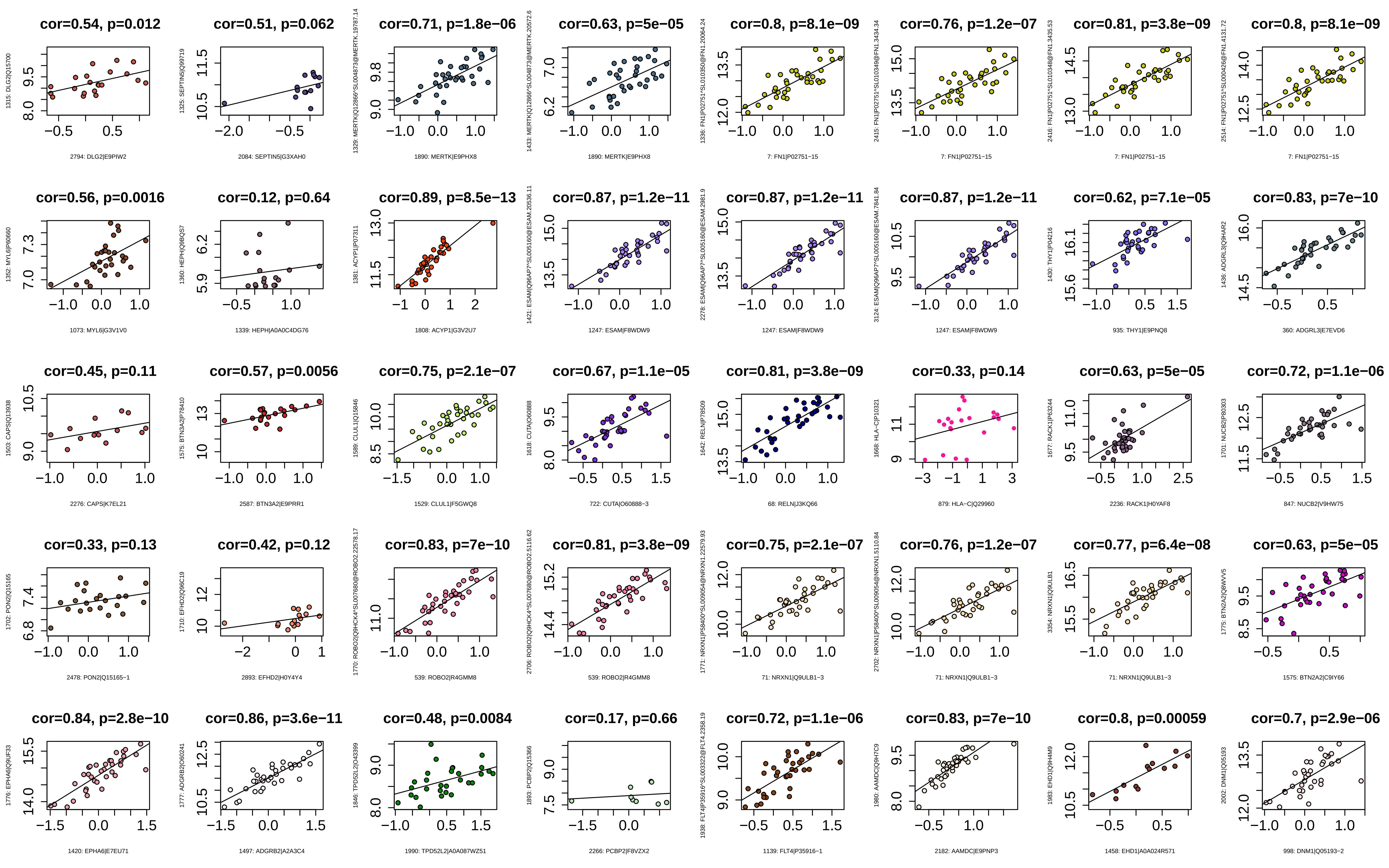
cor=0.54, p=0.012
1315: DLG2|Q15700
2794: DLG2|E9PIW2
cor=0.51, p=0.062
1325: SEPTIN5|Q99719
2084: SEPTIN5|G3XAH0
cor=0.71, p=1.8e−06
1329: MERTK|Q12866^SL004873@MERTK.19787.14
1890: MERTK|E9PHX8
cor=0.63, p=5e−05
1433: MERTK|Q12866^SL004873@MERTK.20572.6
1890: MERTK|E9PHX8
cor=0.8, p=8.1e−09
1336: FN1|P02751^SL010350@FN1.20064.24
7: FN1|P02751−15
cor=0.76, p=1.2e−07
2415: FN1|P02751^SL010349@FN1.3434.34
7: FN1|P02751−15
cor=0.81, p=3.8e−09
2416: FN1|P02751^SL010348@FN1.3435.53
7: FN1|P02751−15
cor=0.8, p=8.1e−09
2514: FN1|P02751^SL000426@FN1.4131.72
7: FN1|P02751−15
cor=0.56, p=0.0016
1352: MYL6|P60660
1073: MYL6|G3V1V0
cor=0.12, p=0.64
1360: HEPH|Q9BQS7
1339: HEPH|A0A0C4DG76
cor=0.89, p=8.5e−13
1381: ACYP1|P07311
1808: ACYP1|G3V2U7
cor=0.87, p=1.2e−11
1421: ESAM|Q96AP7^SL005160@ESAM.20536.11
1247: ESAM|F8WDW9
cor=0.87, p=1.2e−11
2278: ESAM|Q96AP7^SL005160@ESAM.2981.9
1247: ESAM|F8WDW9
cor=0.87, p=1.2e−11
3124: ESAM|Q96AP7^SL005160@ESAM.7841.84
1247: ESAM|F8WDW9
cor=0.62, p=7.1e−05
1430: THY1|P04216
935: THY1|E9PNQ8
cor=0.83, p=7e−10
1436: ADGRL3|Q9HAR2
360: ADGRL3|E7EVD6
cor=0.45, p=0.11
1503: CAPS|Q13938
2276: CAPS|K7EL21
cor=0.57, p=0.0056
1575: BTN3A2|P78410
2587: BTN3A2|E9PRR1
cor=0.75, p=2.1e−07
1589: CLUL1|Q15846
1529: CLUL1|F5GWQ8
cor=0.67, p=1.1e−05
1618: CUTA|O60888
722: CUTA|O60888−3
cor=0.81, p=3.8e−09
1642: RELN|P78509
68: RELN|J3KQ66
cor=0.33, p=0.14
1668: HLA−C|P10321
879: HLA−C|Q29960
cor=0.63, p=5e−05
1677: RACK1|P63244
2236: RACK1|H0YAF8
cor=0.72, p=1.1e−06
1701: NUCB2|P80303
847: NUCB2|V9HW75
cor=0.33, p=0.13
1702: PON2|Q15165
2478: PON2|Q15165−1
cor=0.42, p=0.12
1710: EFHD2|Q96C19
2893: EFHD2|H0Y4Y4
cor=0.83, p=7e−10
1770: ROBO2|Q9HCK4^SL007680@ROBO2.22578.17
539: ROBO2|R4GMM8
cor=0.81, p=3.8e−09
2706: ROBO2|Q9HCK4^SL007680@ROBO2.5116.62
539: ROBO2|R4GMM8
cor=0.75, p=2.1e−07
1771: NRXN1|P58400^SL009054@NRXN1.22579.93
71: NRXN1|Q9ULB1−3
cor=0.76, p=1.2e−07
2702: NRXN1|P58400^SL009054@NRXN1.5110.84
71: NRXN1|Q9ULB1−3
cor=0.77, p=6.4e−08
3354: NRXN1|Q9ULB1
71: NRXN1|Q9ULB1−3
cor=0.63, p=5e−05
1775: BTN2A2|Q8WVV5
1575: BTN2A2|C9IY66
cor=0.84, p=2.8e−10
1776: EPHA6|Q9UF33
1420: EPHA6|E7EU71
cor=0.86, p=3.6e−11
1777: ADGRB2|O60241
1497: ADGRB2|A2A3C4
cor=0.48, p=0.0084
1846: TPD52L2|O43399
1990: TPD52L2|A0A087WZ51
cor=0.17, p=0.66
1893: PCBP2|Q15366
2266: PCBP2|F8VZX2
cor=0.72, p=1.1e−06
1938: FLT4|P35916^SL003322@FLT4.2358.19
1139: FLT4|P35916−1
cor=0.83, p=7e−10
1980: AAMDC|Q9H7C9
2182: AAMDC|E9PNP3
cor=0.8, p=0.00059
1983: EHD1|Q9H4M9
1458: EHD1|A0A024R571
cor=0.7, p=2.9e−06
2002: DNM1|Q05193
998: DNM1|Q05193−2

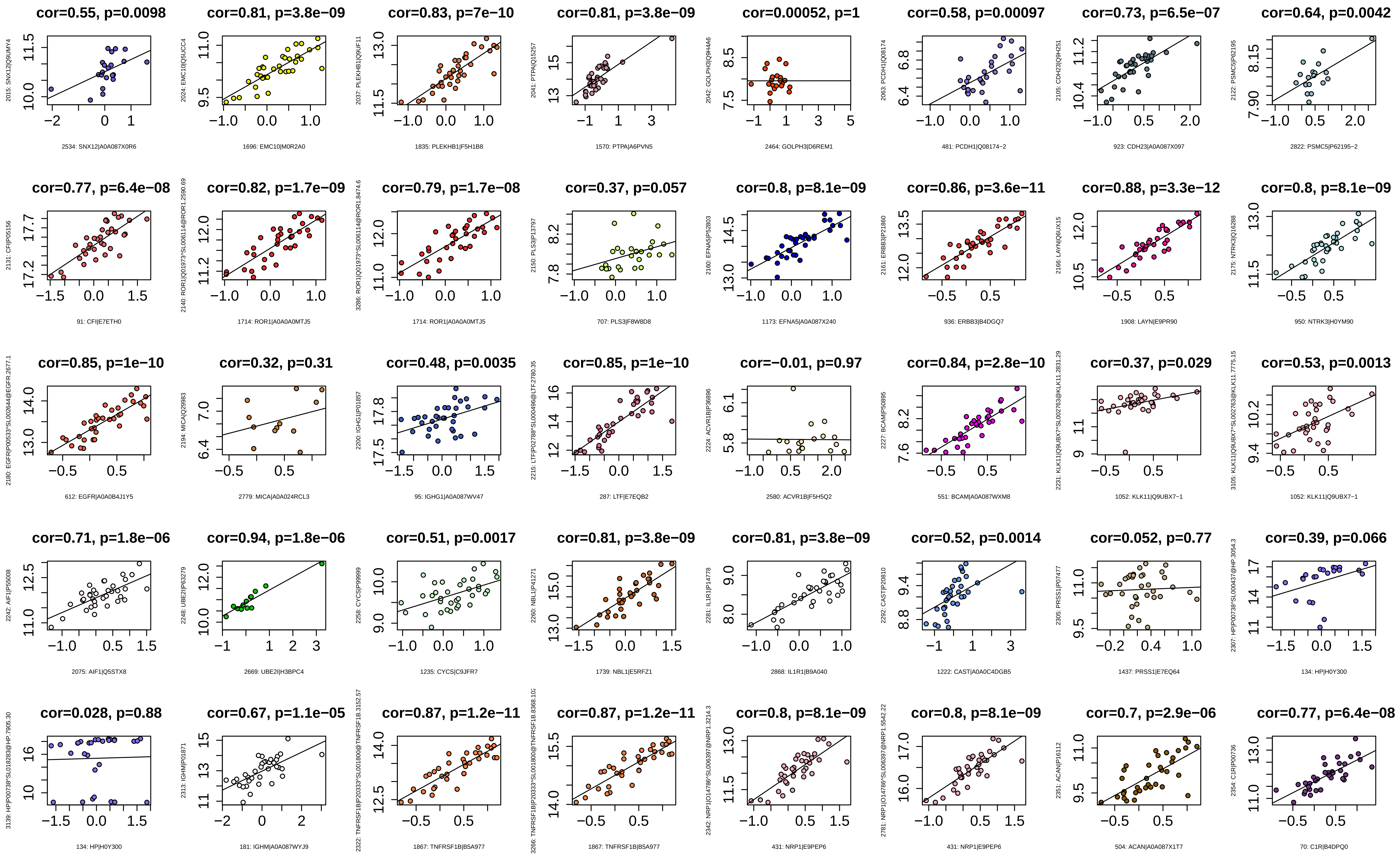
cor=0.55, p=0.0098
2015: SNX12|Q9UMY4
2534: SNX12|A0A087X0R6
cor=0.81, p=3.8e−09
2024: EMC10|Q5UCC4
1696: EMC10|M0R2A0
cor=0.83, p=7e−10
2037: PLEKHB1|Q9UF11
1835: PLEKHB1|F5H1B8
cor=0.81, p=3.8e−09
2041: PTPA|Q15257
1570: PTPA|A6PVN5
cor=0.00052, p=1
2042: GOLPH3|Q9H4A6
2464: GOLPH3|D6REM1
cor=0.58, p=0.00097
2063: PCDH1|Q08174
481: PCDH1|Q08174−2
cor=0.73, p=6.5e−07
2105: CDH23|Q9H251
923: CDH23|A0A087X097
cor=0.64, p=0.0042
2122: PSMC5|P62195
2822: PSMC5|P62195−2
cor=0.77, p=6.4e−08
2131: CFI|P05156
91: CFI|E7ETH0
cor=0.82, p=1.7e−09
2140: ROR1|Q01973^SL006114@ROR1.2590.69
1714: ROR1|A0A0A0MTJ5
cor=0.79, p=1.7e−08
3286: ROR1|Q01973^SL006114@ROR1.8474.6
1714: ROR1|A0A0A0MTJ5
cor=0.37, p=0.057
2150: PLS3|P13797
707: PLS3|F8W8D8
cor=0.8, p=8.1e−09
2160: EFNA5|P52803
1173: EFNA5|A0A087X240
cor=0.86, p=3.6e−11
2161: ERBB3|P21860
936: ERBB3|B4DGQ7
cor=0.88, p=3.3e−12
2166: LAYN|Q6UX15
1908: LAYN|E9PR90
cor=0.8, p=8.1e−09
2175: NTRK3|Q16288
950: NTRK3|H0YM90
cor=0.85, p=1e−10
2180: EGFR|P00533^SL002644@EGFR.2677.1
612: EGFR|A0A0B4J1Y5
cor=0.32, p=0.31
2194: MICA|Q29983
2779: MICA|A0A024RCL3
cor=0.48, p=0.0035
2200: IGHG1|P01857
95: IGHG1|A0A087WV47
cor=0.85, p=1e−10
2215: LTF|P02788^SL000496@LTF.2780.35
287: LTF|E7EQB2
cor=−0.01, p=0.97
2224: ACVR1B|P36896
2580: ACVR1B|F5H5Q2
cor=0.84, p=2.8e−10
2227: BCAM|P50895
551: BCAM|A0A087WXM8
cor=0.37, p=0.029
2231: KLK11|Q9UBX7^SL002763@KLK11.2831.29
1052: KLK11|Q9UBX7−1
cor=0.53, p=0.0013
3105: KLK11|Q9UBX7^SL002763@KLK11.7775.15
1052: KLK11|Q9UBX7−1
cor=0.71, p=1.8e−06
2242: AIF1|P55008
2075: AIF1|Q5STX8
cor=0.94, p=1.8e−06
2248: UBE2I|P63279
2669: UBE2I|H3BPC4
cor=0.51, p=0.0017
2258: CYCS|P99999
1235: CYCS|C9JFR7
cor=0.81, p=3.8e−09
2260: NBL1|P41271
1739: NBL1|E5RFZ1
cor=0.81, p=3.8e−09
2281: IL1R1|P14778
2868: IL1R1|B9A040
cor=0.52, p=0.0014
2292: CAST|P20810
1222: CAST|A0A0C4DGB5
cor=0.052, p=0.77
2305: PRSS1|P07477
1437: PRSS1|E7EQ64
cor=0.39, p=0.066
2307: HP|P00738^SL000437@HP.3054.3
134: HP|H0Y300
cor=0.028, p=0.88
3139: HP|P00738^SL018283@HP.7905.30
134: HP|H0Y300
cor=0.67, p=1.1e−05
2313: IGHM|P01871
181: IGHM|A0A087WYJ9
cor=0.87, p=1.2e−11
2322: TNFRSF1B|P20333^SL001800@TNFRSF18.3152.57
1867: TNFRSF1B|B5A977
cor=0.87, p=1.2e−11
3266: TNFRSF1B|P20333^SL001800@TNFRSF18.8368.102
1867: TNFRSF1B|B5A977
cor=0.8, p=8.1e−09
2342: NRP1|O14786^SL006397@NRP1.3214.3
431: NRP1|E9PEP6
cor=0.8, p=8.1e−09
2781: NRP1|O14786^SL006397@NRP1.5542.22
431: NRP1|E9PEP6
cor=0.7, p=2.9e−06
2351: ACAN|P16112
504: ACAN|A0A087X1T7
cor=0.77, p=6.4e−08
2354: C1R|P00736
70: C1R|B4DPQ0

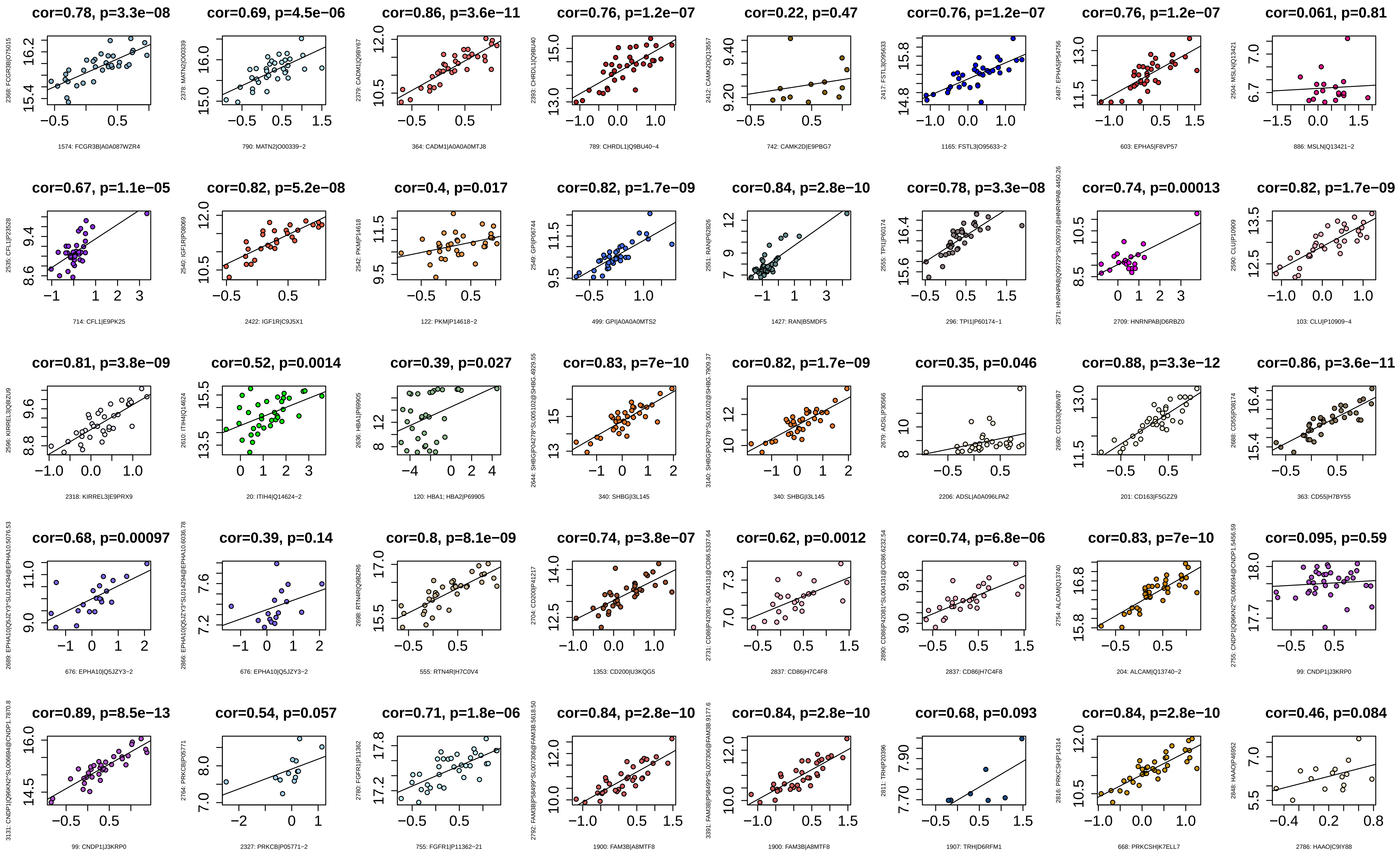
cor=0.78, p=3.3e−08
1574: FCGR3B|A0A087WZR4
cor=0.69, p=4.5e−06
790: MATN2|O00339−2
cor=0.86, p=3.6e−11
364: CADM1|A0A0A0MTJ8
cor=0.76, p=1.2e−07
789: CHRDL1|Q9BU40−4
cor=0.22, p=0.47
742: CAMK2D|E9PBG7
cor=0.76, p=1.2e−07
1165: FSTL3|O95633−2
cor=0.76, p=1.2e−07
603: EPHA5|F8VP57
cor=0.061, p=0.81
886: MSLN|Q13421−2
cor=0.67, p=1.1e−05
714: CFL1|E9PK25
cor=0.82, p=5.2e−08
2422: IGF1R|C9J5X1
cor=0.4, p=0.017
122: PKM|P14618−2
cor=0.82, p=1.7e−09
499: GPI|A0A0A0MTS2
cor=0.84, p=2.8e−10
1427: RAN|B5MDF5
cor=0.78, p=3.3e−08
296: TPI1|P60174−1
cor=0.74, p=0.00013
2709: HNRNPAB|D6RBZ0
cor=0.82, p=1.7e−09
103: CLU|P10909−4
cor=0.81, p=3.8e−09
2318: KIRREL3|E9PRX9
cor=0.52, p=0.0014
20: ITIH4|Q14624−2
cor=0.39, p=0.027
120: HBA1; HBA2|P69905
cor=0.83, p=7e−10
340: SHBG|I3L145
cor=0.82, p=1.7e−09
340: SHBG|I3L145
cor=0.35, p=0.046
2206: ADSL|A0A096LPA2
cor=0.88, p=3.3e−12
201: CD163|F5GZZ9
cor=0.86, p=3.6e−11
363: CD55|H7BY55
cor=0.68, p=0.00097
676: EPHA10|Q5JZY3−2
cor=0.39, p=0.14
676: EPHA10|Q5JZY3−2
cor=0.8, p=8.1e−09
555: RTN4R|H7C0V4
cor=0.74, p=3.8e−07
1353: CD200|U3KQG5
cor=0.62, p=0.0012
2837: CD86|H7C4F8
cor=0.74, p=6.8e−06
2837: CD86|H7C4F8
cor=0.83, p=7e−10
204: ALCAM|Q13740−2
cor=0.095, p=0.59
99: CNDP1|J3KRP0
cor=0.89, p=8.5e−13
99: CNDP1|J3KRP0
cor=0.54, p=0.057
2327: PRKCB|P05771−2
cor=0.71, p=1.8e−06
755: FGFR1|P11362−21
cor=0.84, p=2.8e−10
1900: FAM3B|A8MTF8
cor=0.84, p=2.8e−10
1900: FAM3B|A8MTF8
cor=0.68, p=0.093
1907: TRH|D6RFM1
cor=0.84, p=2.8e−10
668: PRKCSH|K7ELL7
cor=0.46, p=0.084
2786: HAAO|C9IY88

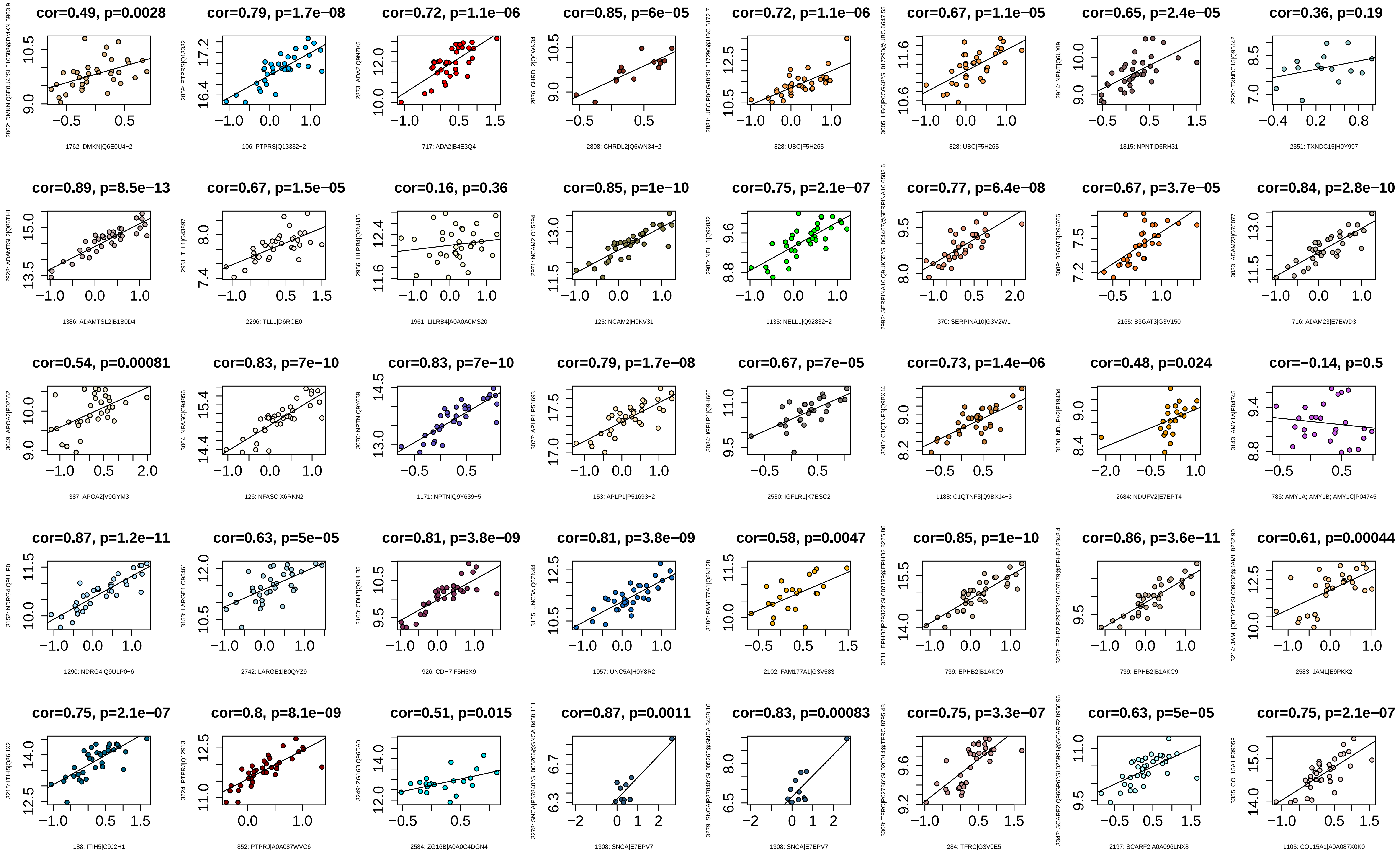

cor=0.49, p=0.0028
2862: DMKN|Q6E0U4*SL010588@DMKN.5963.9
1762: DMKN|Q6E0U4−2
cor=0.79, p=1.7e−08
2869: PTPRS|Q13332
106: PTPRS|Q13332−2
cor=0.72, p=1.1e−06
2873: ADA2|Q9NZK5
717: ADA2|B4E3Q4
cor=0.85, p=6e−05
2876: CHRDL2|Q6WN34
2898: CHRDL2|Q6WN34−2
cor=0.72, p=1.1e−06
2881: UBC|P0CG48*SL017290@UBC.6172.7
828: UBC|F5H265
cor=0.67, p=1.1e−05
3005: UBC|P0CG48*SL017290@UBC.6647.55
828: UBC|F5H265
cor=0.65, p=2.4e−05
2914: NPNT|Q6UXI9
1815: NPNT|D6RH31
cor=0.36, p=0.19
2920: TXNDC15|Q96J42
2351: TXNDC15|H0Y997
cor=0.89, p=8.5e−13
2928: ADAMTSL2|Q86TH1
1386: ADAMTSL2|B1B0D4
cor=0.67, p=1.5e−05
2931: TLL1|O43897
2296: TLL1|D6RCE0
cor=0.16, p=0.36
2956: LILRB4|Q8NHJ6
1961: LILRB4|A0A0A0MS20
cor=0.85, p=1e−10
2971: NCAM2|O15394
125: NCAM2|H9KV31
cor=0.75, p=2.1e−07
2980: NELL1|Q92832
1135: NELL1|Q92832−2
cor=0.77, p=6.4e−08
2992: SERPINA10|Q9UK55*SL004467@SERPINA10.6583.6
370: SERPINA10|G3V2W1
cor=0.67, p=3.7e−05
3009: B3GAT3|O94766
2165: B3GAT3|G3V150
cor=0.84, p=2.8e−10
3033: ADAM23|O75077
716: ADAM23|E7EWD3
cor=0.54, p=0.00081
3049: APOA2|P02652
387: APOA2|V9GYM3
cor=0.83, p=7e−10
3064: NFASC|O94856
126: NFASC|X6RKN2
cor=0.83, p=7e−10
3070: NPTN|Q9Y639
1171: NPTN|Q9Y639−5
cor=0.79, p=1.7e−08
3077: APLP1|P51693
153: APLP1|P51693−2
cor=0.67, p=7e−05
3084: IGFLR1|Q9H665
2530: IGFLR1|K7ESC2
cor=0.73, p=1.4e−06
3085: C1QTNF3|Q9BXJ4
1188: C1QTNF3|Q9BXJ4−3
cor=0.48, p=0.024
3100: NDUFV2|P19404
2684: NDUFV2|E7EPT4
cor=−0.14, p=0.5
3143: AMY1A|P04745
786: AMY1A; AMY1B; AMY1C|P04745
cor=0.87, p=1.2e−11
3152: NDRG4|Q9ULP0
1290: NDRG4|Q9ULP0−6
cor=0.63, p=5e−05
3153: LARGE1|O95461
2742: LARGE1|B0QYZ9
cor=0.81, p=3.8e−09
3160: CDH7|Q9ULB5
926: CDH7|F5H5X9
cor=0.81, p=3.8e−09
3165: UNC5A|Q6ZN44
1957: UNC5A|H0Y8R2
cor=0.58, p=0.0047
3186: FAM177A1|Q8N128
2102: FAM177A1|G3V583
cor=0.85, p=1e−10
3211: EPHB2|P29323*SL007179@EPHB2.8225.86
739: EPHB2|B1AKC9
cor=0.86, p=3.6e−11
3258: EPHB2|P29323*SL007179@EPHB2.8348.4
739: EPHB2|B1AKC9
cor=0.61, p=0.00044
3214: JAML|Q86YT9*SL009202@JAML.8232.90
2583: JAML|E9PKK2
cor=0.75, p=2.1e−07
3215: ITIH5|Q86UX2
188: ITIH5|C9J2H1
cor=0.8, p=8.1e−09
3224: PTPRJ|Q12913
852: PTPRJ|A0A087WVC6
cor=0.51, p=0.015
3249: ZG16B|Q96DA0
2584: ZG16B|A0A0C4DGN4
cor=0.87, p=0.0011
3278: SNCA|P37840*SL005266@SNCA.8458.111
1308: SNCA|E7EPV7
cor=0.83, p=0.00083
3279: SNCA|P37840*SL005266@SNCA.8458.16
1308: SNCA|E7EPV7
cor=0.75, p=3.3e−07
3308: TFRC|P02786*SL002614@TFRC.8795.48
284: TFRC|G3V0E5
cor=0.63, p=5e−05
3347: SCARF2|Q96GP6*SL025991@SCARF2.8956.96
2197: SCARF2|A0A096LNX8
cor=0.75, p=2.1e−07
3355: COL15A1|P39059
1105: COL15A1|A0A087X0K0

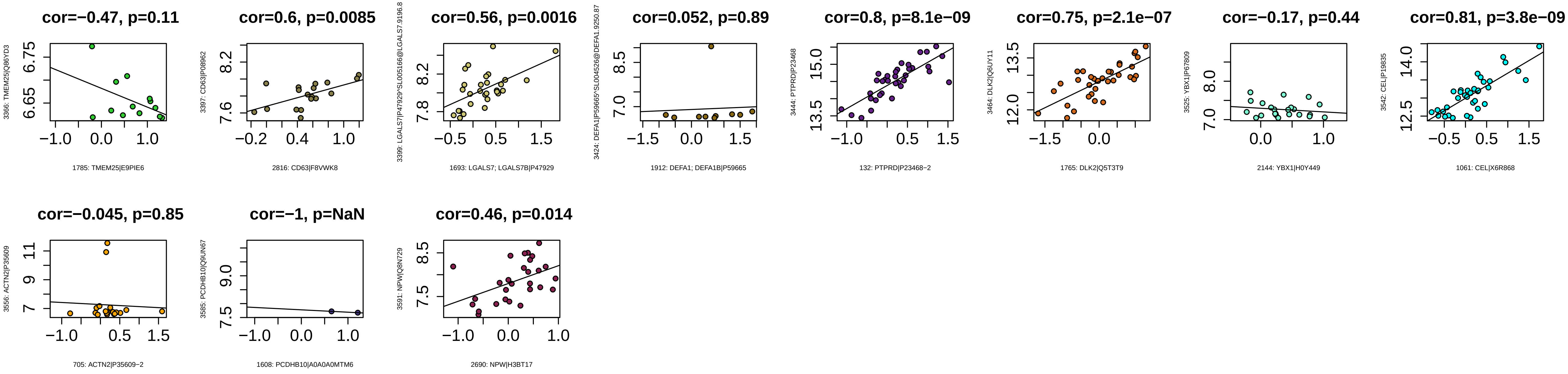

cor=−0.47, p=0.11
3366: TMEM25|Q86YD3
1785: TMEM25|E9PIE6
cor=0.6, p=0.0085
3397: CD63|P08962
2816: CD63|F8VWK8
cor=0.56, p=0.0016
3399: LGALS7|P47929^SL005166@LGALS7.9196.8
1693: LGALS7; LGALS7B|P47929
cor=0.052, p=0.89
3424: DEFA1|P59665^SL004526@DEFA1.9250.87
1912: DEFA1; DEFA1B|P59665
cor=0.8, p=8.1e−09
3444: PTPRD|P23468
132: PTPRD|P23468−2
cor=0.75, p=2.1e−07
3464: DLK2|Q6UY11
1765: DLK2|Q5T3T9
cor=−0.17, p=0.44
3525: YBX1|P67809
2144: YBX1|H0Y449
cor=0.81, p=3.8e−09
3542: CEL|P19835
1061: CEL|X6R868
cor=−0.045, p=0.85
3556: ACTN2|P35609
705: ACTN2|P35609−2
cor=−1, p=NaN
3585: PCDHB10|Q9UN67
1608: PCDHB10|A0A0A0MTM6
cor=0.46, p=0.014
3591: NPW|Q8N729
2690: NPW|H3BT17

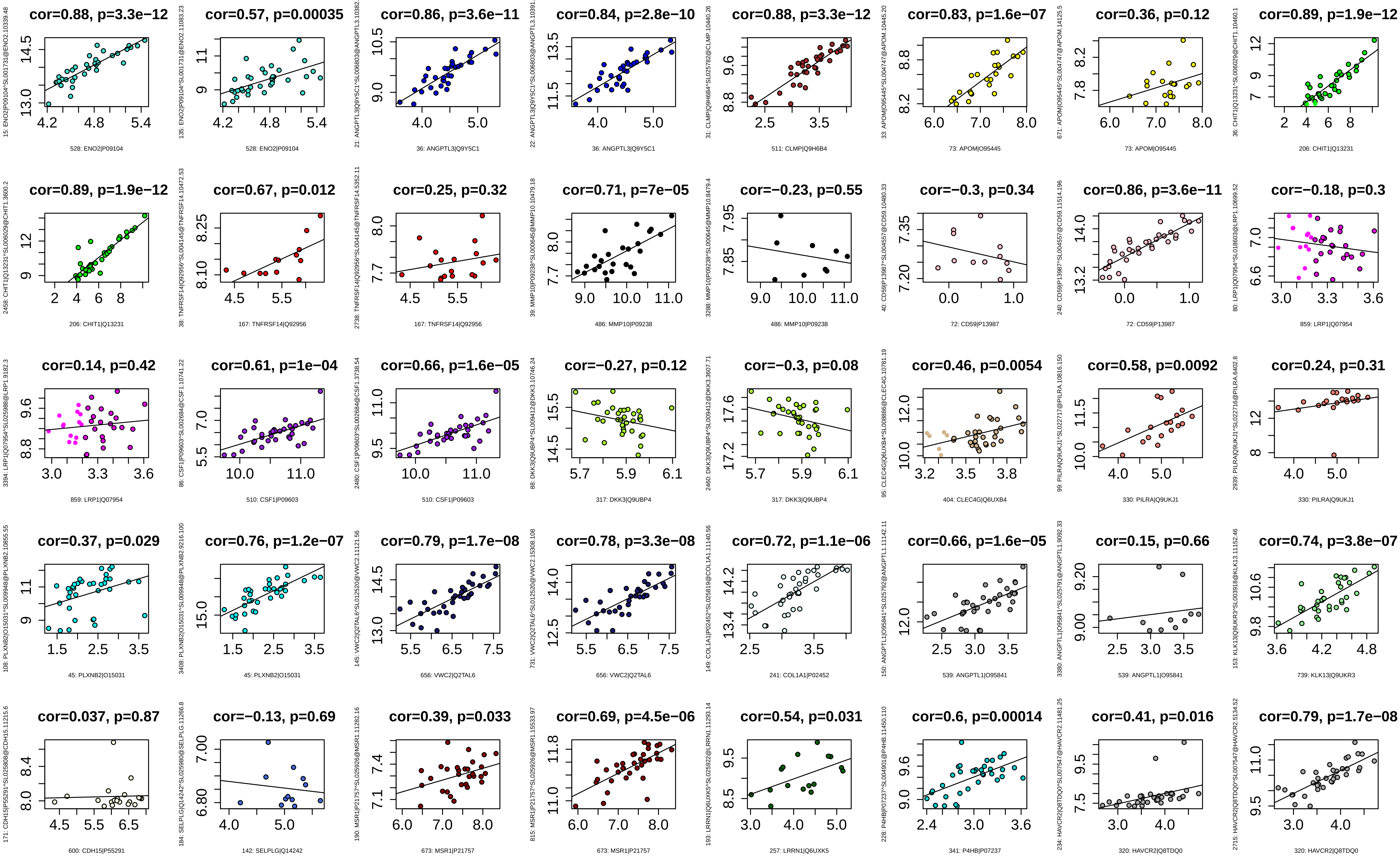

cor=0.88, p=3.3e−12
528: ENO2|P09104
cor=0.57, p=0.00035
528: ENO2|P09104
cor=0.86, p=3.6e−11
36: ANGPTL3|Q9Y5C1
cor=0.84, p=2.8e−10
36: ANGPTL3|Q9Y5C1
cor=0.88, p=3.3e−12
511: CLMP|Q9H6B4
cor=0.83, p=1.6e−07
73: APOM|O95445
cor=0.36, p=0.12
73: APOM|O95445
cor=0.89, p=1.9e−12
206: CHIT1|Q13231
cor=0.89, p=1.9e−12
206: CHIT1|Q13231
cor=0.67, p=0.012
167: TNFRSF14|Q92956
cor=0.25, p=0.32
167: TNFRSF14|Q92956
cor=0.71, p=7e−05
486: MMP10|P09238
cor=−0.23, p=0.55
486: MMP10|P09238
cor=−0.3, p=0.34
72: CD59|P13987
cor=0.86, p=3.6e−11
72: CD59|P13987
cor=−0.18, p=0.3
859: LRP1|Q07954
cor=0.14, p=0.42
859: LRP1|Q07954
cor=0.61, p=1e−04
510: CSF1|P09603
cor=0.66, p=1.6e−05
510: CSF1|P09603
cor=−0.27, p=0.12
317: DKK3|Q9UBP4
cor=−0.3, p=0.08
317: DKK3|Q9UBP4
cor=0.46, p=0.0054
404: CLEC4G|Q6UXB4
cor=0.58, p=0.0092
330: PILRA|Q9UKJ1
cor=0.24, p=0.31
330: PILRA|Q9UKJ1
cor=0.37, p=0.029
45: PLXNB2|O15031
cor=0.76, p=1.2e−07
45: PLXNB2|O15031
cor=0.79, p=1.7e−08
656: VWC2|Q2TAL6
cor=0.78, p=3.3e−08
656: VWC2|Q2TAL6
cor=0.72, p=1.1e−06
241: COL1A1|P02452
cor=0.66, p=1.6e−05
539: ANGPTL1|O95841
cor=0.15, p=0.66
539: ANGPTL1|O95841
cor=0.74, p=3.8e−07
739: KLK13|Q9UKR3
cor=0.037, p=0.87
600: CDH15|P55291
cor=−0.13, p=0.69
142: SELPLG|Q14242
cor=0.39, p=0.033
673: MSR1|P21757
cor=0.69, p=4.5e−06
673: MSR1|P21757
cor=0.54, p=0.031
257: LRRN1|Q6UXK5
cor=0.6, p=0.00014
341: P4HB|P07237
cor=0.41, p=0.016
320: HAVCR2|Q8TDQ0
cor=0.79, p=1.7e−08
320: HAVCR2|Q8TDQ0

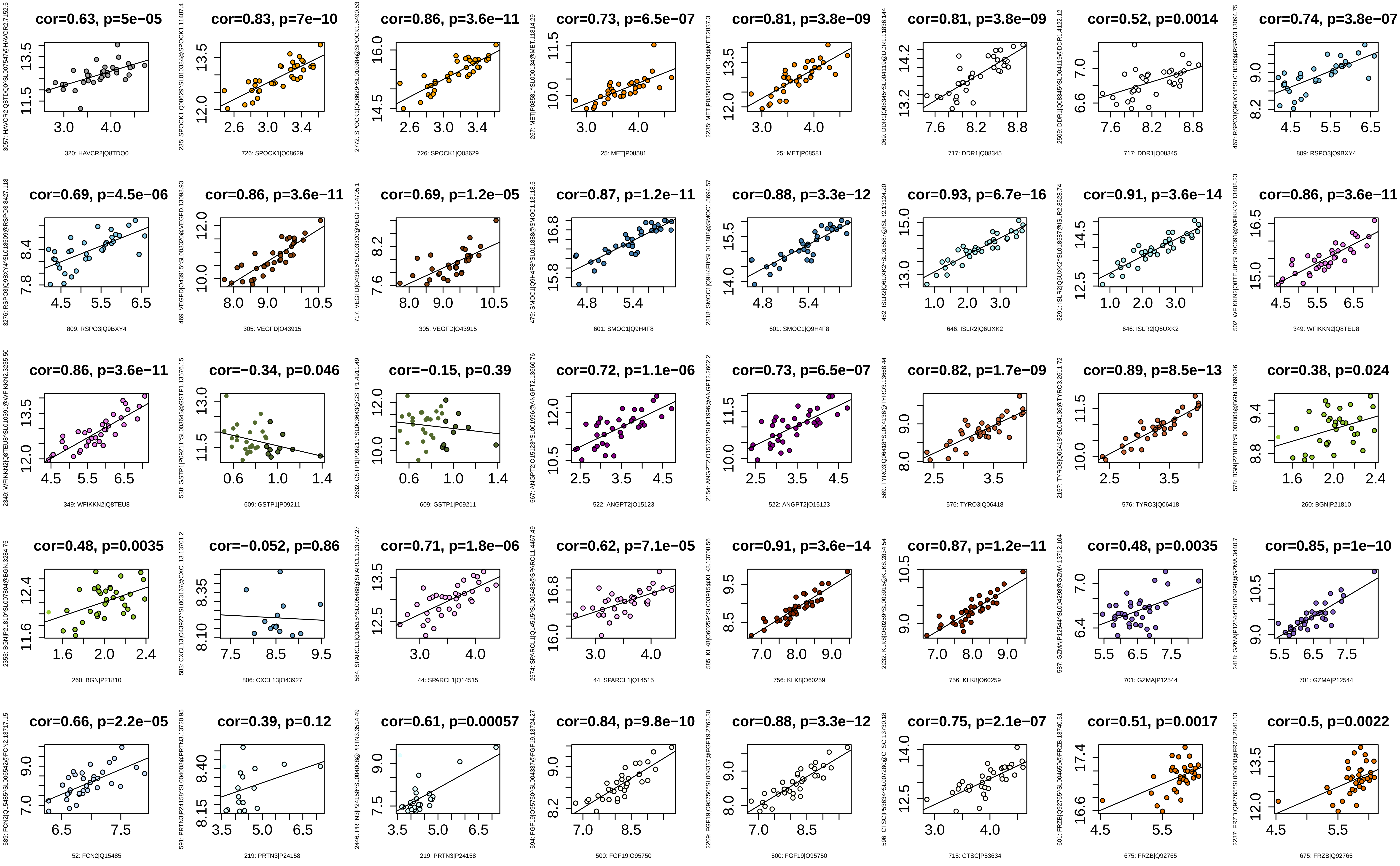
cor=0.63, p=5e−05
320: HAVCR2|Q8TDQ0
cor=0.83, p=7e−10
726: SPOCK1|Q08629
cor=0.86, p=3.6e−11
726: SPOCK1|Q08629
cor=0.73, p=6.5e−07
25: MET|P08581
cor=0.81, p=3.8e−09
25: MET|P08581
cor=0.81, p=3.8e−09
717: DDR1|Q08345
cor=0.52, p=0.0014
717: DDR1|Q08345
cor=0.74, p=3.8e−07
809: RSPO3|Q9BXY4
cor=0.69, p=4.5e−06
809: RSPO3|Q9BXY4
cor=0.86, p=3.6e−11
305: VEGFD|O43915
cor=0.69, p=1.2e−05
305: VEGFD|O43915
cor=0.87, p=1.2e−11
601: SMOC1|Q9H4F8
cor=0.88, p=3.3e−12
601: SMOC1|Q9H4F8
cor=0.93, p=6.7e−16
646: ISLR2|Q6UXK2
cor=0.91, p=3.6e−14
646: ISLR2|Q6UXK2
cor=0.86, p=3.6e−11
349: WFIKKN2|Q8TEU8
cor=0.86, p=3.6e−11
349: WFIKKN2|Q8TEU8
cor=−0.34, p=0.046
609: GSTP1|P09211
cor=−0.15, p=0.39
609: GSTP1|P09211
cor=0.72, p=1.1e−06
522: ANGPT2|O15123
cor=0.73, p=6.5e−07
522: ANGPT2|O15123
cor=0.82, p=1.7e−09
576: TYRO3|Q06418
cor=0.89, p=8.5e−13
576: TYRO3|Q06418
cor=0.38, p=0.024
260: BGN|P21810
cor=0.48, p=0.0035
260: BGN|P21810
cor=−0.052, p=0.86
806: CXCL13|O43927
cor=0.71, p=1.8e−06
44: SPARCL1|Q14515
cor=0.62, p=7.1e−05
44: SPARCL1|Q14515
cor=0.91, p=3.6e−14
756: KLK8|O60259
cor=0.87, p=1.2e−11
756: KLK8|O60259
cor=0.48, p=0.0035
701: GZMA|P12544
cor=0.85, p=1e−10
701: GZMA|P12544
cor=0.66, p=2.2e−05
52: FCN2|Q15485
cor=0.39, p=0.12
219: PRTN3|P24158
cor=0.61, p=0.00057
219: PRTN3|P24158
cor=0.84, p=9.8e−10
500: FGF19|O95750
cor=0.88, p=3.3e−12
500: FGF19|O95750
cor=0.75, p=2.1e−07
715: CTSC|P53634
cor=0.51, p=0.0017
675: FRZB|Q92765
cor=0.5, p=0.0022
675: FRZB|Q92765

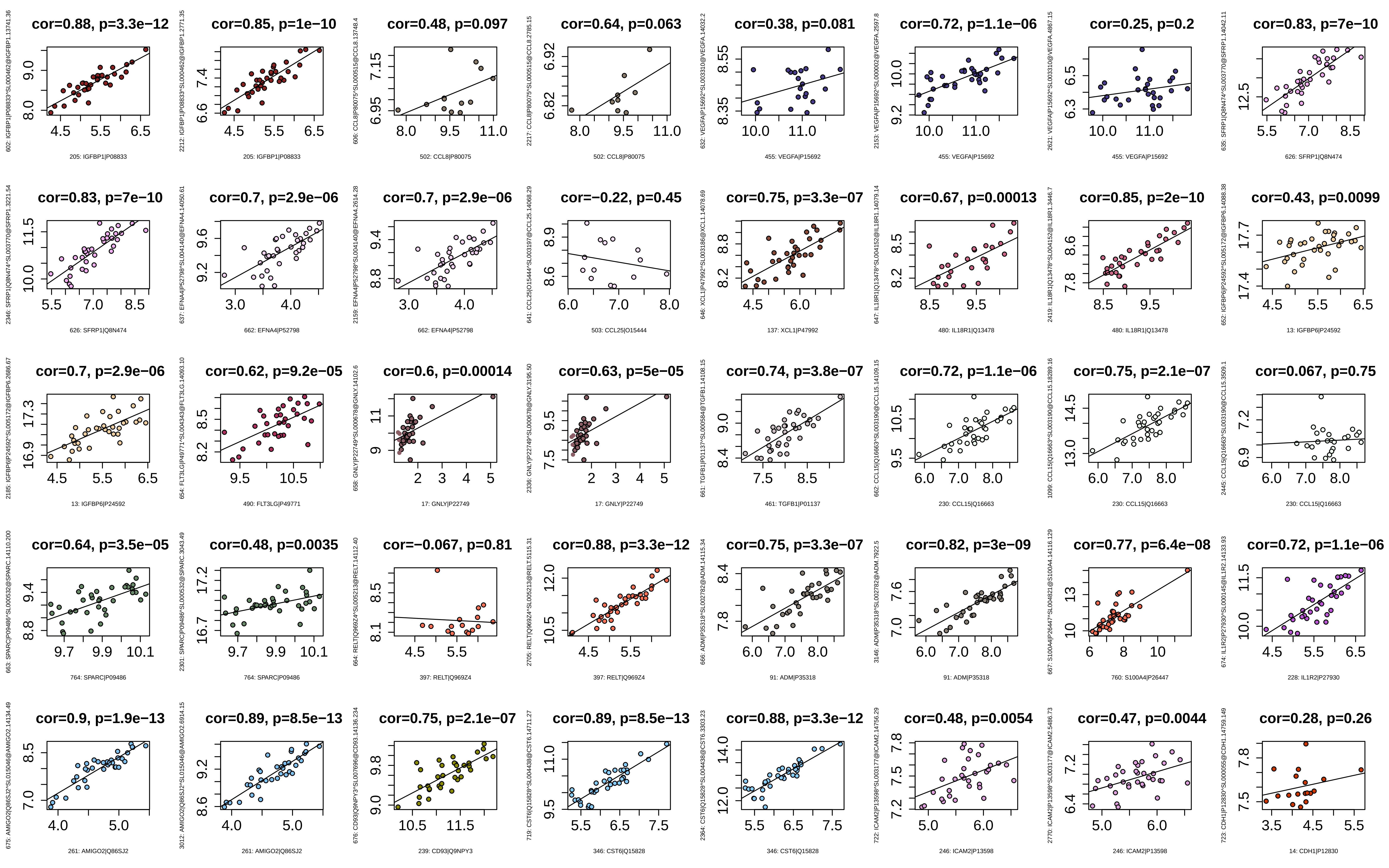

cor=0.88, p=3.3e−12
205: IGFBP1|P08833
cor=0.85, p=1e−10
205: IGFBP1|P08833
cor=0.48, p=0.097
502: CCL8|P80075
cor=0.64, p=0.063
502: CCL8|P80075
cor=0.38, p=0.081
455: VEGFA|P15692
cor=0.72, p=1.1e−06
455: VEGFA|P15692
cor=0.25, p=0.2
455: VEGFA|P15692
cor=0.83, p=7e−10
626: SFRP1|Q8N474
cor=0.83, p=7e−10
626: SFRP1|Q8N474
cor=0.7, p=2.9e−06
662: EFNA4|P52798
cor=0.7, p=2.9e−06
662: EFNA4|P52798
cor=−0.22, p=0.45
503: CCL25|O15444
cor=0.75, p=3.3e−07
137: XCL1|P47992
cor=0.67, p=0.00013
480: IL18R1|Q13478
cor=0.85, p=2e−10
480: IL18R1|Q13478
cor=0.43, p=0.0099
13: IGFBP6|P24592
cor=0.7, p=2.9e−06
13: IGFBP6|P24592
cor=0.62, p=9.2e−05
490: FLT3LG|P49771
cor=0.6, p=0.00014
17: GNLY|P22749
cor=0.63, p=5e−05
17: GNLY|P22749
cor=0.74, p=3.8e−07
461: TGFB1|P01137
cor=0.72, p=1.1e−06
230: CCL15|Q16663
cor=0.75, p=2.1e−07
230: CCL15|Q16663
cor=0.067, p=0.75
230: CCL15|Q16663
cor=0.64, p=3.5e−05
764: SPARC|P09486
cor=0.48, p=0.0035
764: SPARC|P09486
cor=−0.067, p=0.81
397: RELT|Q969Z4
cor=0.88, p=3.3e−12
397: RELT|Q969Z4
cor=0.75, p=3.3e−07
91: ADM|P35318
cor=0.82, p=3e−09
91: ADM|P35318
cor=0.77, p=6.4e−08
760: S100A4|P26447
cor=0.72, p=1.1e−06
228: IL1R2|P27930
cor=0.9, p=1.9e−13
261: AMIGO2|Q86SJ2
cor=0.89, p=8.5e−13
261: AMIGO2|Q86SJ2
cor=0.75, p=2.1e−07
239: CD93|Q9NPY3
cor=0.89, p=8.5e−13
346: CST6|Q15828
cor=0.88, p=3.3e−12
346: CST6|Q15828
cor=0.48, p=0.0054
246: ICAM2|P13598
cor=0.47, p=0.0044
246: ICAM2|P13598
cor=0.28, p=0.26
14: CDH1|P12830

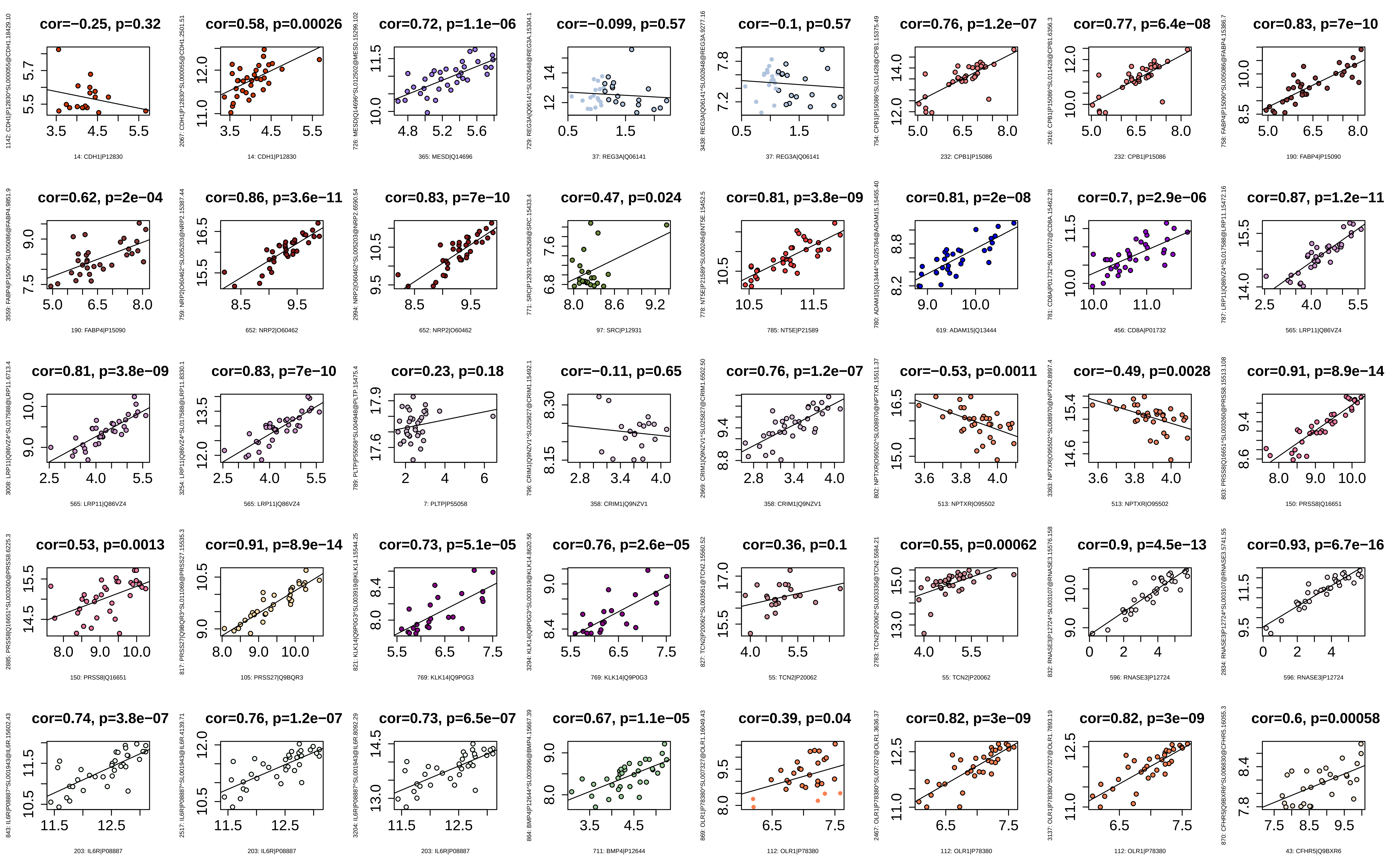
cor=−0.25, p=0.32
14: CDH1|P12830
cor=0.58, p=0.00026
14: CDH1|P12830
cor=0.72, p=1.1e−06
365: MESD|Q14696
cor=−0.099, p=0.57
37: REG3A|Q06141
cor=−0.1, p=0.57
37: REG3A|Q06141
cor=0.76, p=1.2e−07
232: CPB1|P15086
cor=0.77, p=6.4e−08
232: CPB1|P15086
cor=0.83, p=7e−10
190: FABP4|P15090
cor=0.62, p=2e−04
190: FABP4|P15090
cor=0.86, p=3.6e−11
652: NRP2|O60462
cor=0.83, p=7e−10
652: NRP2|O60462
cor=0.47, p=0.024
97: SRC|P12931
cor=0.81, p=3.8e−09
785: NT5E|P21589
cor=0.81, p=2e−08
619: ADAM15|Q13444
cor=0.7, p=2.9e−06
456: CD8A|P01732
cor=0.87, p=1.2e−11
565: LRP11|Q86VZ4
cor=0.81, p=3.8e−09
565: LRP11|Q86VZ4
cor=0.83, p=7e−10
565: LRP11|Q86VZ4
cor=0.23, p=0.18
7: PLTP|P55058
cor=−0.11, p=0.65
358: CRIM1|Q9NZV1
cor=0.76, p=1.2e−07
358: CRIM1|Q9NZV1
cor=−0.53, p=0.0011
513: NPTXR|O95502
cor=−0.49, p=0.0028
513: NPTXR|O95502
cor=0.91, p=8.9e−14
150: PRSS8|Q16651
cor=0.53, p=0.0013
150: PRSS8|Q16651
cor=0.91, p=8.9e−14
105: PRSS27|Q9BQR3
cor=0.73, p=5.1e−05
769: KLK14|Q9P0G3
cor=0.76, p=2.6e−05
769: KLK14|Q9P0G3
cor=0.36, p=0.1
55: TCN2|P20062
cor=0.55, p=0.00062
55: TCN2|P20062
cor=0.9, p=4.5e−13
596: RNASE3|P12724
cor=0.93, p=6.7e−16
596: RNASE3|P12724
cor=0.74, p=3.8e−07
203: IL6R|P08887
cor=0.76, p=1.2e−07
203: IL6R|P08887
cor=0.73, p=6.5e−07
203: IL6R|P08887
cor=0.67, p=1.1e−05
711: BMP4|P12644
cor=0.39, p=0.04
112: OLR1|P78380
cor=0.82, p=3e−09
112: OLR1|P78380
cor=0.82, p=3e−09
112: OLR1|P78380
cor=0.6, p=0.00058
43: CFHR5|Q9BXR6

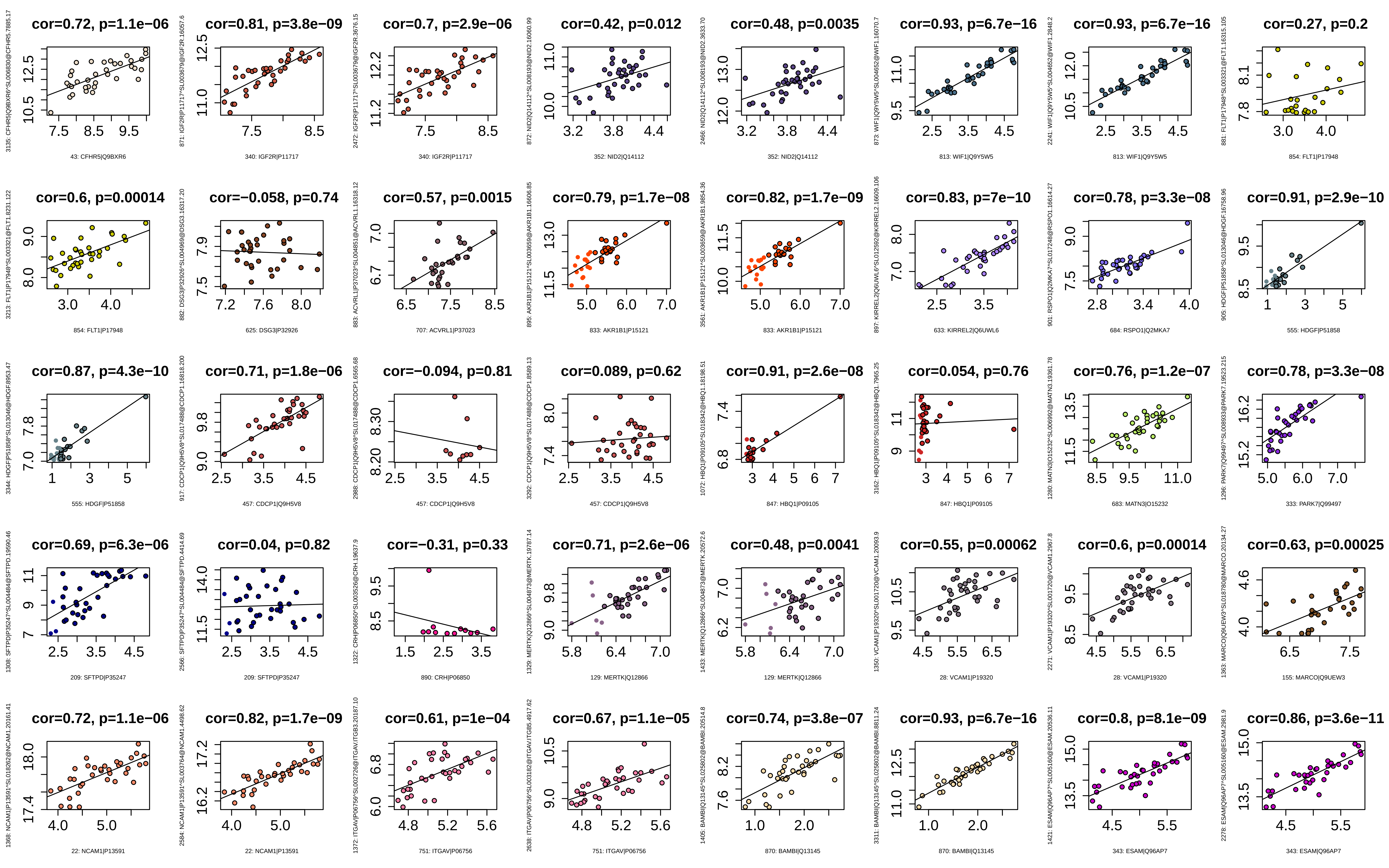
cor=0.72, p=1.1e−06
43: CFHR5|Q9BXR6
cor=0.81, p=3.8e−09
340: IGF2R|P11717
cor=0.7, p=2.9e−06
340: IGF2R|P11717
cor=0.42, p=0.012
352: NID2|Q14112
cor=0.48, p=0.0035
352: NID2|Q14112
cor=0.93, p=6.7e−16
813: WIF1|Q9Y5W5
cor=0.93, p=6.7e−16
813: WIF1|Q9Y5W5
cor=0.27, p=0.2
854: FLT1|P17948
cor=0.6, p=0.00014
854: FLT1|P17948
cor=−0.058, p=0.74
625: DSG3|P32926
cor=0.57, p=0.0015
707: ACVRL1|P37023
cor=0.79, p=1.7e−08
833: AKR1B1|P15121
cor=0.82, p=1.7e−09
833: AKR1B1|P15121
cor=0.83, p=7e−10
633: KIRREL2|Q6UWL6
cor=0.78, p=3.3e−08
684: RSPO1|Q2MKA7
cor=0.91, p=2.9e−10
555: HDGF|P51858
cor=0.87, p=4.3e−10
555: HDGF|P51858
cor=0.71, p=1.8e−06
457: CDCP1|Q9H5V8
cor=−0.094, p=0.81
457: CDCP1|Q9H5V8
cor=0.089, p=0.62
457: CDCP1|Q9H5V8
cor=0.91, p=2.6e−08
847: HBQ1|P09105
cor=0.054, p=0.76
847: HBQ1|P09105
cor=0.76, p=1.2e−07
683: MATN3|O15232
cor=0.78, p=3.3e−08
333: PARK7|Q99497
cor=0.69, p=6.3e−06
209: SFTPD|P35247
cor=0.04, p=0.82
209: SFTPD|P35247
cor=−0.31, p=0.33
890: CRH|P06850
cor=0.71, p=2.6e−06
129: MERTK|Q12866
cor=0.48, p=0.0041
129: MERTK|Q12866
cor=0.55, p=0.00062
28: VCAM1|P19320
cor=0.6, p=0.00014
28: VCAM1|P19320
cor=0.63, p=0.00025
155: MARCO|Q9UEW3
cor=0.72, p=1.1e−06
22: NCAM1|P13591
cor=0.82, p=1.7e−09
22: NCAM1|P13591
cor=0.61, p=1e−04
751: ITGAV|P06756
cor=0.67, p=1.1e−05
751: ITGAV|P06756
cor=0.74, p=3.8e−07
870: BAMBI|Q13145
cor=0.93, p=6.7e−16
870: BAMBI|Q13145
cor=0.8, p=8.1e−09
343: ESAM|Q96AP7
cor=0.86, p=3.6e−11
343: ESAM|Q96AP7

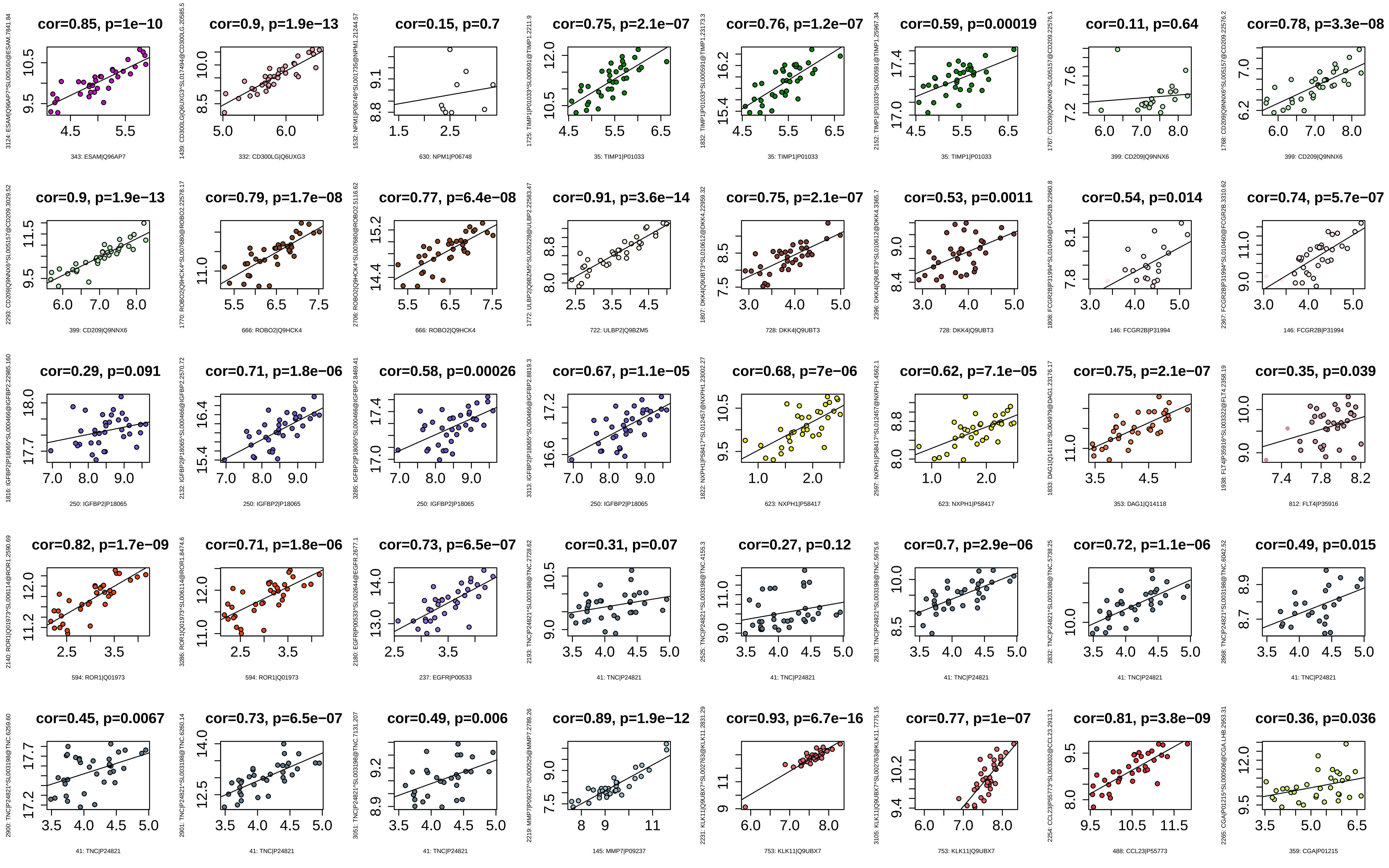
cor=0.85, p=1e−10
343: ESAM|Q96AP7
cor=0.9, p=1.9e−13
332: CD300LG|Q6UXG3
cor=0.15, p=0.7
630: NPM1|P06748
cor=0.75, p=2.1e−07
35: TIMP1|P01033
cor=0.76, p=1.2e−07
35: TIMP1|P01033
cor=0.59, p=0.00019
35: TIMP1|P01033
cor=0.11, p=0.64
399: CD209|Q9NNX6
cor=0.78, p=3.3e−08
399: CD209|Q9NNX6
cor=0.9, p=1.9e−13
399: CD209|Q9NNX6
cor=0.79, p=1.7e−08
666: ROBO2|Q9HCK4
cor=0.77, p=6.4e−08
666: ROBO2|Q9HCK4
cor=0.91, p=3.6e−14
722: ULBP2|Q9BZM5
cor=0.75, p=2.1e−07
728: DKK4|Q9UBT3
cor=0.53, p=0.0011
728: DKK4|Q9UBT3
cor=0.54, p=0.014
146: FCGR2B|P31994
cor=0.74, p=5.7e−07
146: FCGR2B|P31994
cor=0.29, p=0.091
250: IGFBP2|P18065
cor=0.71, p=1.8e−06
250: IGFBP2|P18065
cor=0.58, p=0.00026
250: IGFBP2|P18065
cor=0.67, p=1.1e−05
250: IGFBP2|P18065
cor=0.68, p=7e−06
623: NXPH1|P58417
cor=0.62, p=7.1e−05
623: NXPH1|P58417
cor=0.75, p=2.1e−07
353: DAG1|Q14118
cor=0.35, p=0.039
812: FLT4|P35916
cor=0.82, p=1.7e−09
594: ROR1|Q01973
cor=0.71, p=1.8e−06
594: ROR1|Q01973
cor=0.73, p=6.5e−07
237: EGFR|P00533
cor=0.31, p=0.07
41: TNC|P24821
cor=0.27, p=0.12
41: TNC|P24821
cor=0.7, p=2.9e−06
41: TNC|P24821
cor=0.72, p=1.1e−06
41: TNC|P24821
cor=0.49, p=0.015
41: TNC|P24821
cor=0.45, p=0.0067
41: TNC|P24821
cor=0.73, p=6.5e−07
41: TNC|P24821
cor=0.49, p=0.006
41: TNC|P24821
cor=0.89, p=1.9e−12
145: MMP7|P09237
cor=0.93, p=6.7e−16
753: KLK11|Q9UBX7
cor=0.77, p=1e−07
753: KLK11|Q9UBX7
cor=0.81, p=3.8e−09
488: CCL23|P55773
cor=0.36, p=0.036
359: CGA|P01215

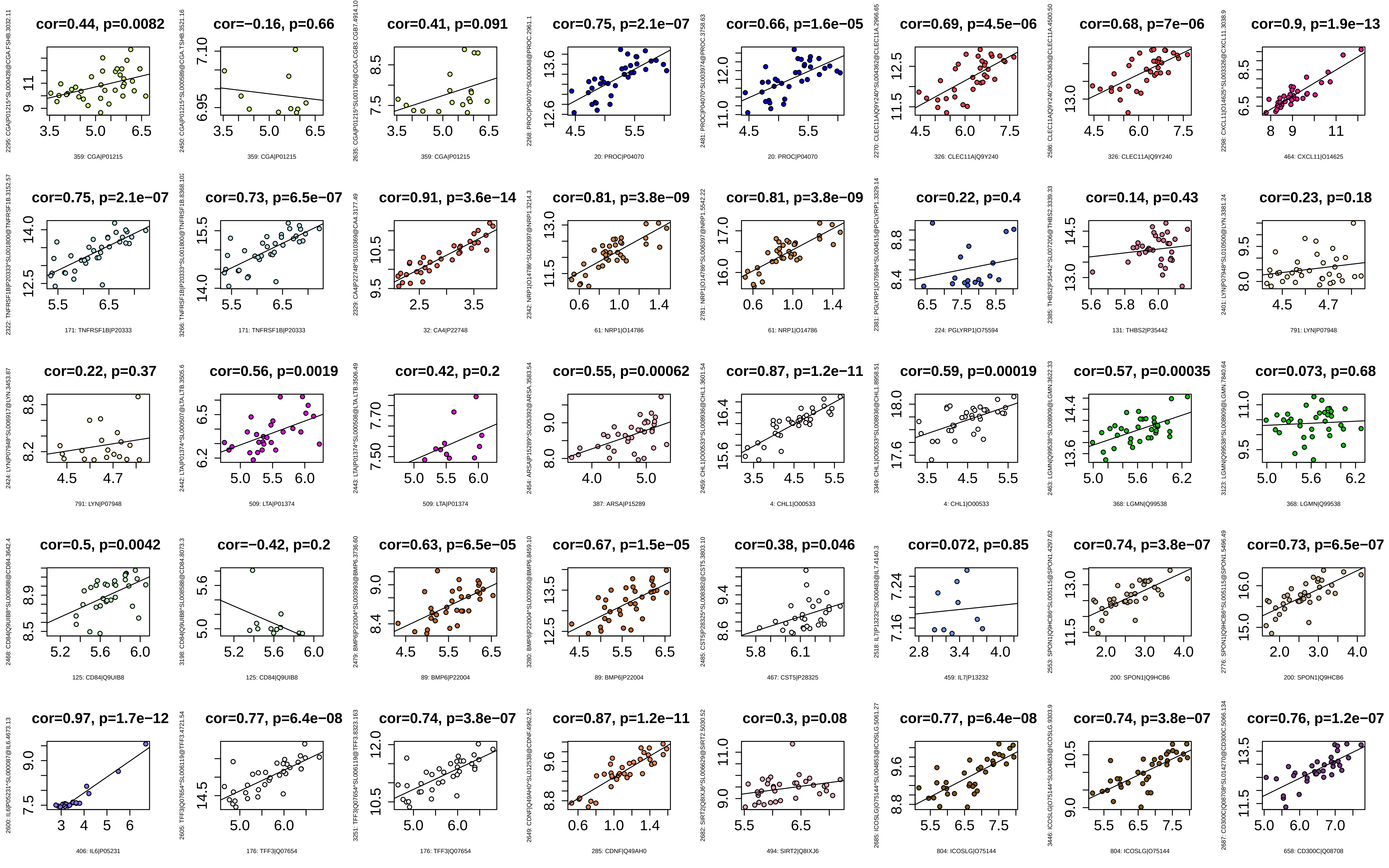
cor=0.44, p=0.0082
359: CGA|P01215
cor=−0.16, p=0.66
359: CGA|P01215
cor=0.41, p=0.091
359: CGA|P01215
cor=0.75, p=2.1e−07
20: PROC|P04070
cor=0.66, p=1.6e−05
20: PROC|P04070
cor=0.69, p=4.5e−06
326: CLEC11A|Q9Y240
cor=0.68, p=7e−06
326: CLEC11A|Q9Y240
cor=0.9, p=1.9e−13
464: CXCL11|O14625
cor=0.75, p=2.1e−07
171: TNFRSF1B|P20333
cor=0.73, p=6.5e−07
171: TNFRSF1B|P20333
cor=0.91, p=3.6e−14
32: CA4|P22748
cor=0.81, p=3.8e−09
61: NRP1|O14786
cor=0.81, p=3.8e−09
61: NRP1|O14786
cor=0.22, p=0.4
224: PGLYRP1|O75594
cor=0.14, p=0.43
131: THBS2|P35442
cor=0.23, p=0.18
791: LYN|P07948
cor=0.22, p=0.37
791: LYN|P07948
cor=0.56, p=0.0019
509: LTA|P01374
cor=0.42, p=0.2
509: LTA|P01374
cor=0.55, p=0.00062
387: ARSA|P15289
cor=0.87, p=1.2e−11
4: CHL1|O00533
cor=0.59, p=0.00019
4: CHL1|O00533
cor=0.57, p=0.00035
368: LGMN|Q99538
cor=0.073, p=0.68
368: LGMN|Q99538
cor=0.5, p=0.0042
125: CD84|Q9UIB8
cor=−0.42, p=0.2
125: CD84|Q9UIB8
cor=0.63, p=6.5e−05
89: BMP6|P22004
cor=0.67, p=1.5e−05
89: BMP6|P22004
cor=0.38, p=0.046
467: CST5|P28325
cor=0.072, p=0.85
459: IL7|P13232
cor=0.74, p=3.8e−07
200: SPON1|Q9HCB6
cor=0.73, p=6.5e−07
200: SPON1|Q9HCB6
cor=0.97, p=1.7e−12
406: IL6|P05231
cor=0.77, p=6.4e−08
176: TFF3|Q07654
cor=0.74, p=3.8e−07
176: TFF3|Q07654
cor=0.87, p=1.2e−11
285: CDNF|Q49AH0
cor=0.3, p=0.08
494: SIRT2|Q8IXJ6
cor=0.77, p=6.4e−08
804: ICOSLG|O75144
cor=0.74, p=3.8e−07
804: ICOSLG|O75144
cor=0.76, p=1.2e−07
658: CD300C|Q08708

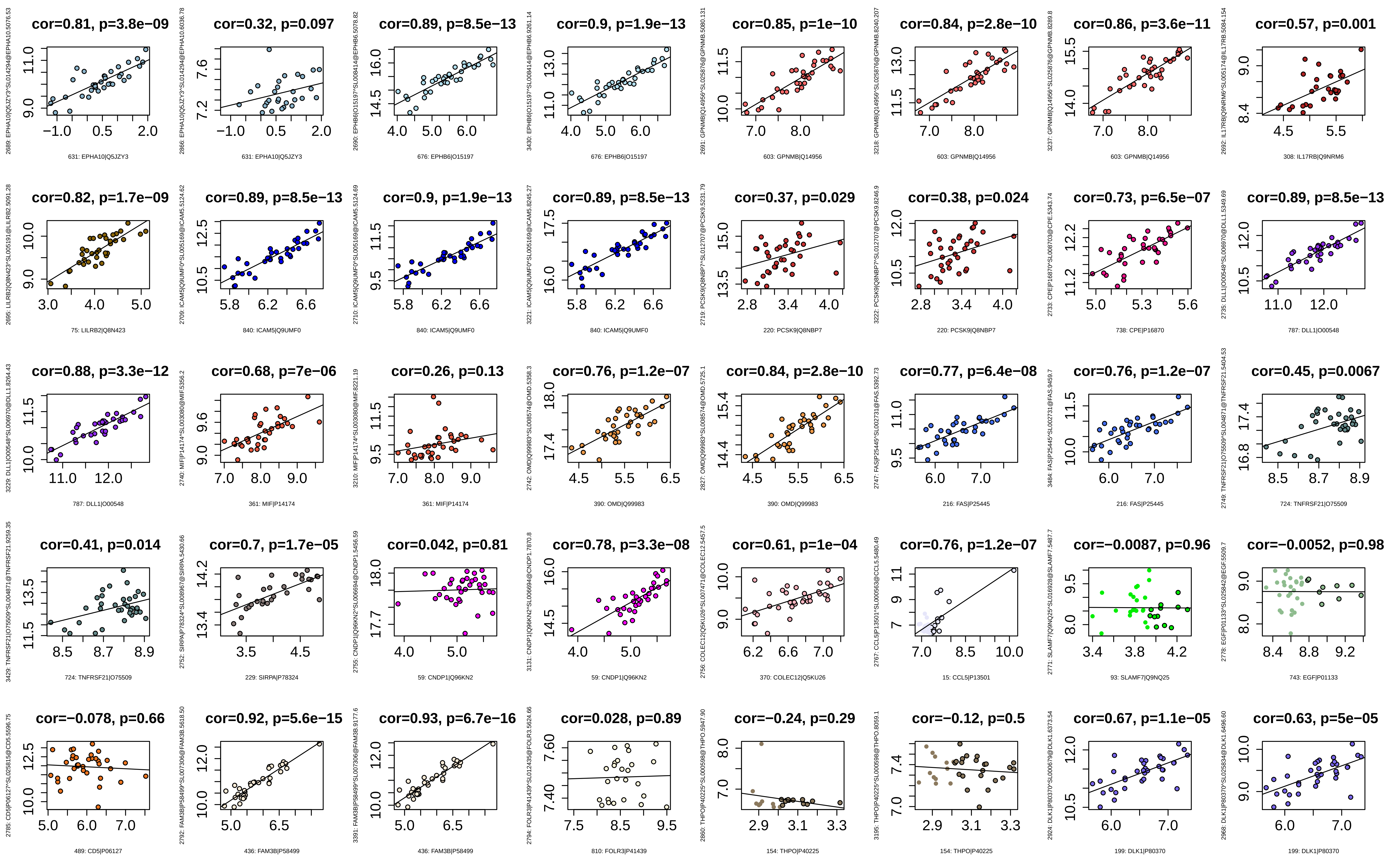
cor=0.81, p=3.8e−09
631: EPHA10|Q5JZY3
cor=0.32, p=0.097
631: EPHA10|Q5JZY3
cor=0.89, p=8.5e−13
676: EPHB6|O15197
cor=0.9, p=1.9e−13
676: EPHB6|O15197
cor=0.85, p=1e−10
603: GPNMB|Q14956
cor=0.84, p=2.8e−10
603: GPNMB|Q14956
cor=0.86, p=3.6e−11
603: GPNMB|Q14956
cor=0.57, p=0.001
308: IL17RB|Q9NRM6
cor=0.82, p=1.7e−09
75: LILRB2|Q8N423
cor=0.89, p=8.5e−13
840: ICAM5|Q9UMF0
cor=0.9, p=1.9e−13
840: ICAM5|Q9UMF0
cor=0.89, p=8.5e−13
840: ICAM5|Q9UMF0
cor=0.37, p=0.029
220: PCSK9|Q8NBP7
cor=0.38, p=0.024
220: PCSK9|Q8NBP7
cor=0.73, p=6.5e−07
738: CPE|P16870
cor=0.89, p=8.5e−13
787: DLL1|O00548
cor=0.88, p=3.3e−12
787: DLL1|O00548
cor=0.68, p=7e−06
361: MIF|P14174
cor=0.26, p=0.13
361: MIF|P14174
cor=0.76, p=1.2e−07
390: OMD|Q99983
cor=0.84, p=2.8e−10
390: OMD|Q99983
cor=0.77, p=6.4e−08
216: FAS|P25445
cor=0.76, p=1.2e−07
216: FAS|P25445
cor=0.45, p=0.0067
724: TNFRSF21|O75509
cor=0.41, p=0.014
724: TNFRSF21|O75509
cor=0.7, p=1.7e−05
229: SIRPA|P78324
cor=0.042, p=0.81
59: CNDP1|Q96KN2
cor=0.78, p=3.3e−08
59: CNDP1|Q96KN2
cor=0.61, p=1e−04
370: COLEC12|Q5KU26
cor=0.76, p=1.2e−07
15: CCL5|P13501
cor=−0.0087, p=0.96
93: SLAMF7|Q9NQ25
cor=−0.0052, p=0.98
743: EGF|P01133
cor=−0.078, p=0.66
489: CD5|P06127
cor=0.92, p=5.6e−15
436: FAM3B|P58499
cor=0.93, p=6.7e−16
436: FAM3B|P58499
cor=0.028, p=0.89
810: FOLR3|P41439
cor=−0.24, p=0.29
154: THPO|P40225
cor=−0.12, p=0.5
154: THPO|P40225
cor=0.67, p=1.1e−05
199: DLK1|P80370
cor=0.63, p=5e−05
199: DLK1|P80370

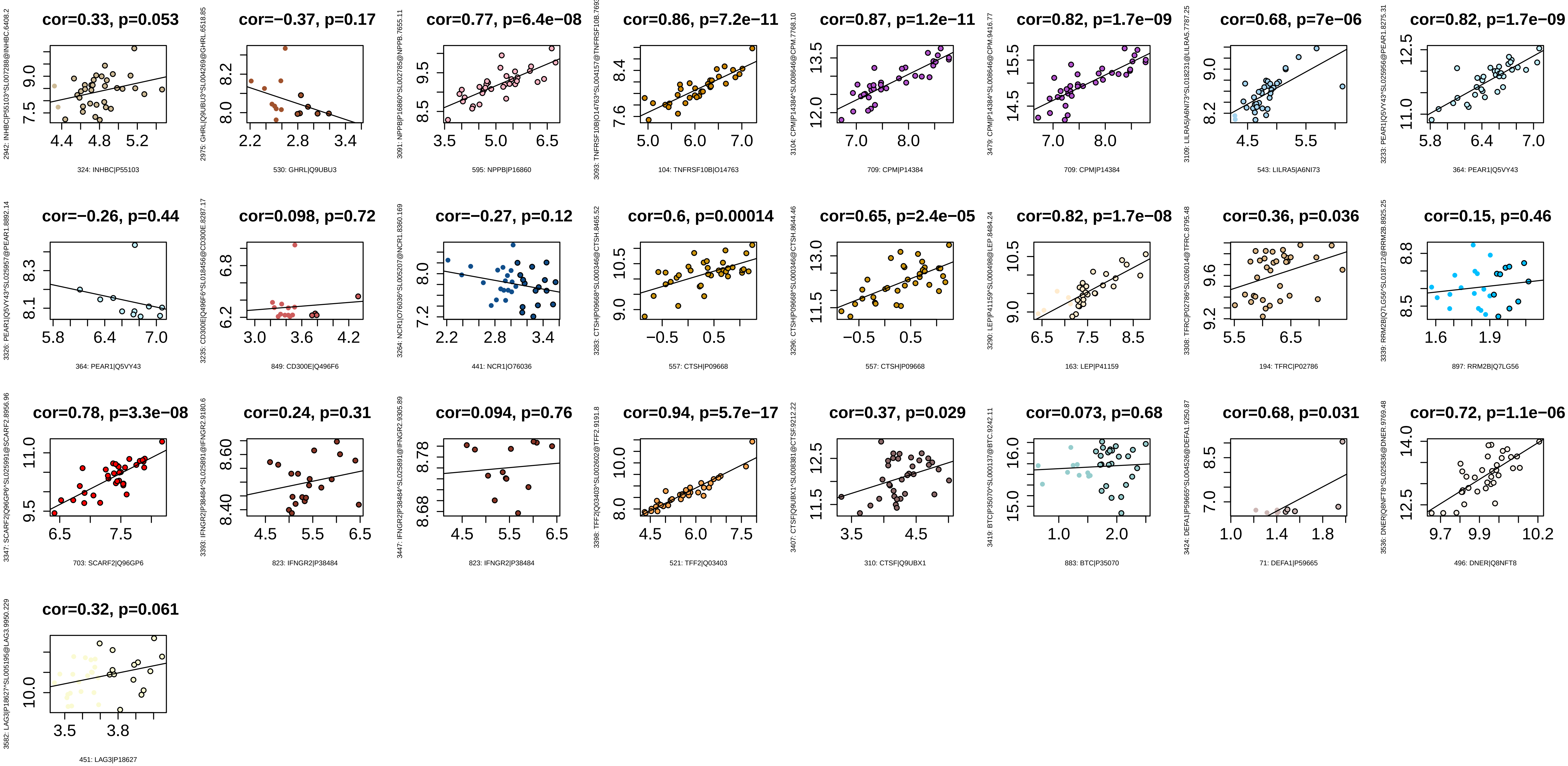
cor=0.33, p=0.053
324: INHBC|P55103
cor=−0.37, p=0.17
530: GHRL|Q9UBU3
cor=0.77, p=6.4e−08
595: NPPB|P16860
cor=0.86, p=7.2e−11
104: TNFRSF10B|O14763
cor=0.87, p=1.2e−11
709: CPM|P14384
cor=0.82, p=1.7e−09
709: CPM|P14384
cor=0.68, p=7e−06
543: LILRA5|A6NI73
cor=0.82, p=1.7e−09
364: PEAR1|Q5VY43
cor=−0.26, p=0.44
364: PEAR1|Q5VY43
cor=0.098, p=0.72
849: CD300E|Q496F6
cor=−0.27, p=0.12
441: NCR1|O76036
cor=0.6, p=0.00014
557: CTSH|P09668
cor=0.65, p=2.4e−05
557: CTSH|P09668
cor=0.82, p=1.7e−08
163: LEP|P41159
cor=0.36, p=0.036
194: TFRC|P02786
cor=0.15, p=0.46
897: RRM2B|Q7LG56
cor=0.78, p=3.3e−08
703: SCARF2|Q96GP6
cor=0.24, p=0.31
823: IFNGR2|P38484
cor=0.094, p=0.76
823: IFNGR2|P38484
cor=0.94, p=5.7e−17
521: TFF2|Q03403
cor=0.37, p=0.029
310: CTSF|Q9UBX1
cor=0.073, p=0.68
883: BTC|P35070
cor=0.68, p=0.031
71: DEFA1|P59665
cor=0.72, p=1.1e−06
496: DNER|Q8NFT8
cor=0.32, p=0.061
451: LAG3|P18627

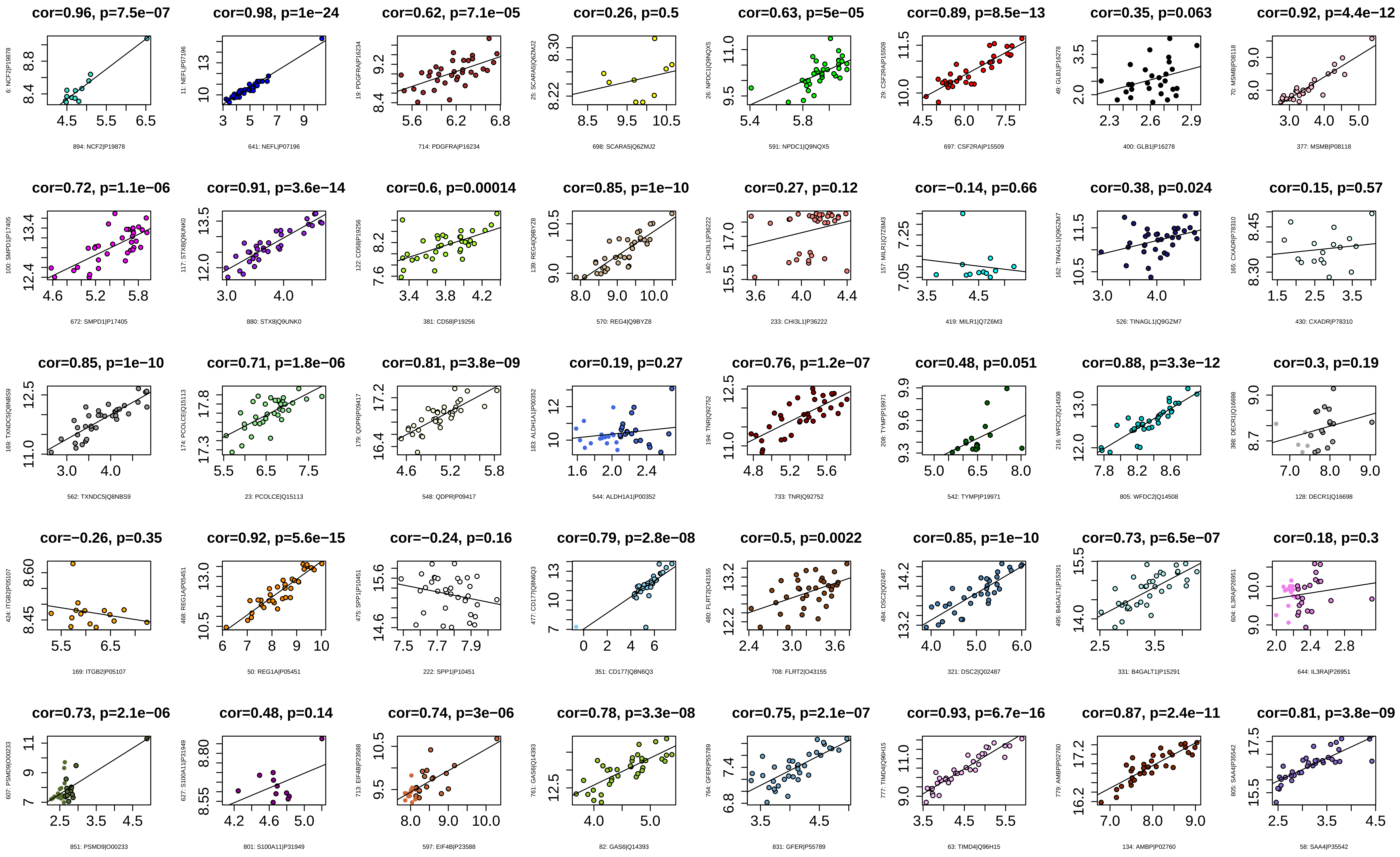
cor=0.96, p=7.5e−07
894: NCF2|P19878
cor=0.98, p=1e−24
641: NEFL|P07196
cor=0.62, p=7.1e−05
714: PDGFRA|P16234
cor=0.26, p=0.5
698: SCARA5|Q6ZMJ2
cor=0.63, p=5e−05
591: NPDC1|Q9NQX5
cor=0.89, p=8.5e−13
697: CSF2RA|P15509
cor=0.35, p=0.063
400: GLB1|P16278
cor=0.92, p=4.4e−12
377: MSMB|P08118
cor=0.72, p=1.1e−06
672: SMPD1|P17405
cor=0.91, p=3.6e−14
880: STX8|Q9UNK0
cor=0.6, p=0.00014
381: CD58|P19256
cor=0.85, p=1e−10
570: REG4|Q9BYZ8
cor=0.27, p=0.12
233: CHI3L1|P36222
cor=−0.14, p=0.66
419: MILR1|Q7Z6M3
cor=0.38, p=0.024
526: TINAGL1|Q9GZM7
cor=0.15, p=0.57
430: CXADR|P78310
cor=0.85, p=1e−10
562: TXNDC5|Q8NBS9
cor=0.71, p=1.8e−06
23: PCOLCE|Q15113
cor=0.81, p=3.8e−09
548: QDPR|P09417
cor=0.19, p=0.27
544: ALDH1A1|P00352
cor=0.76, p=1.2e−07
733: TNR|Q92752
cor=0.48, p=0.051
542: TYMP|P19971
cor=0.88, p=3.3e−12
805: WFDC2|Q14508
cor=0.3, p=0.19
128: DECR1|Q16698
cor=−0.26, p=0.35
169: ITGB2|P05107
cor=0.92, p=5.6e−15
50: REG1A|P05451
cor=−0.24, p=0.16
222: SPP1|P10451
cor=0.79, p=2.8e−08
351: CD177|Q8N6Q3
cor=0.5, p=0.0022
708: FLRT2|O43155
cor=0.85, p=1e−10
321: DSC2|Q02487
cor=0.73, p=6.5e−07
331: B4GALT1|P15291
cor=0.18, p=0.3
644: IL3RA|P26951
cor=0.73, p=2.1e−06
851: PSMD9|O00233
cor=0.48, p=0.14
801: S100A11|P31949
cor=0.74, p=3e−06
597: EIF4B|P23588
cor=0.78, p=3.3e−08
82: GAS6|Q14393
cor=0.75, p=2.1e−07
831: GFER|P55789
cor=0.93, p=6.7e−16
63: TIMD4|Q96H15
cor=0.87, p=2.4e−11
134: AMBP|P02760
cor=0.81, p=3.8e−09
58: SAA4|P35542

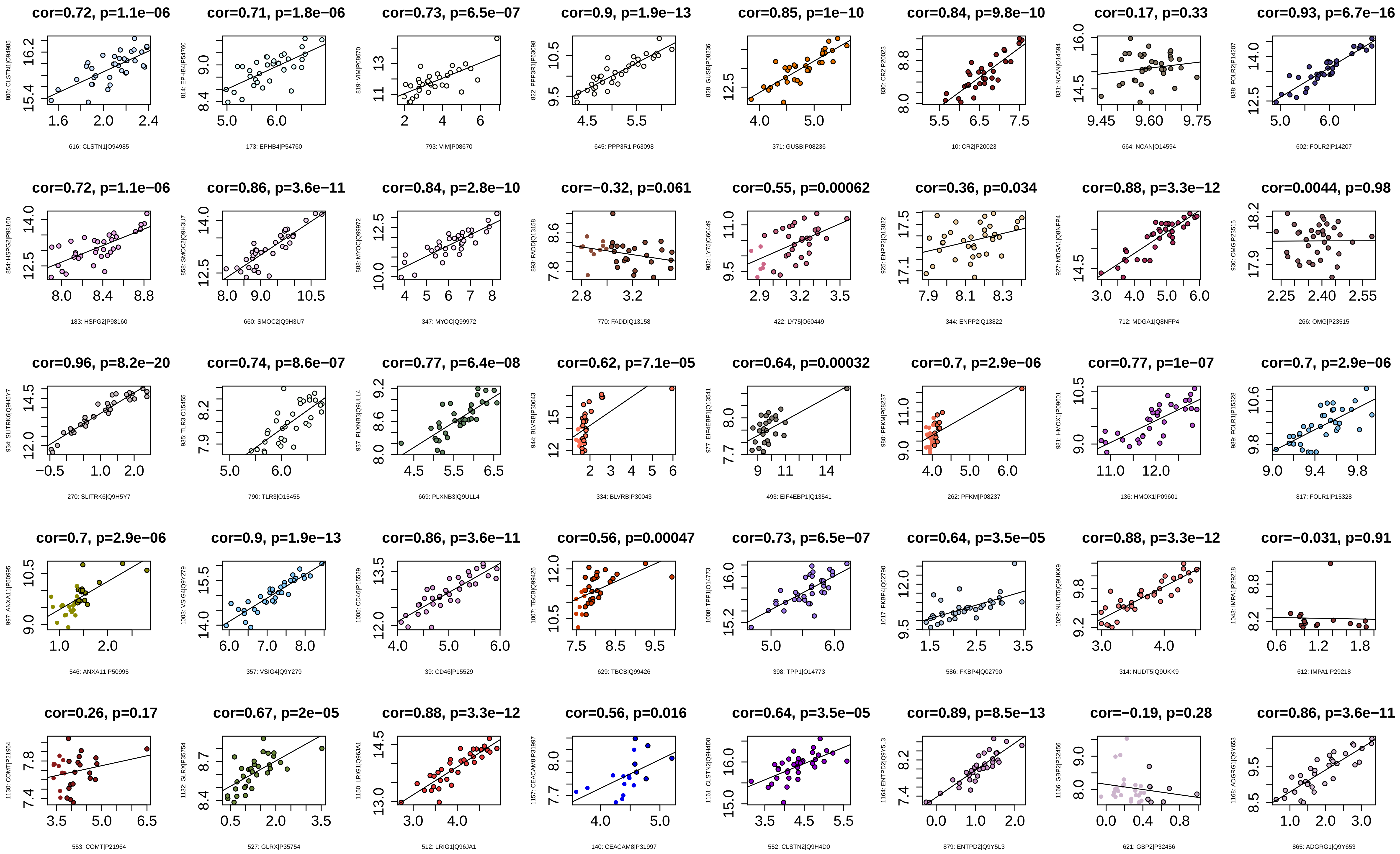
cor=0.72, p=1.1e−06
616: CLSTN1|O94985
cor=0.71, p=1.8e−06
173: EPHB4|P54760
cor=0.73, p=6.5e−07
793: VIM|P08670
cor=0.9, p=1.9e−13
645: PPP3R1|P63098
cor=0.85, p=1e−10
371: GUSB|P08236
cor=0.84, p=9.8e−10
10: CR2|P20023
cor=0.17, p=0.33
664: NCAN|O14594
cor=0.93, p=6.7e−16
602: FOLR2|P14207
cor=0.72, p=1.1e−06
183: HSPG2|P98160
cor=0.86, p=3.6e−11
660: SMOC2|Q9H3U7
cor=0.84, p=2.8e−10
347: MYOC|Q99972
cor=−0.32, p=0.061
770: FADD|Q13158
cor=0.55, p=0.00062
422: LY75|O60449
cor=0.36, p=0.034
344: ENPP2|Q13822
cor=0.88, p=3.3e−12
712: MDGA1|Q8NFP4
cor=0.0044, p=0.98
266: OMG|P23515
cor=0.96, p=8.2e−20
270: SLITRK6|Q9H5Y7
cor=0.74, p=8.6e−07
790: TLR3|O15455
cor=0.77, p=6.4e−08
669: PLXNB3|Q9ULL4
cor=0.62, p=7.1e−05
334: BLVRB|P30043
cor=0.64, p=0.00032
493: EIF4EBP1|Q13541
cor=0.7, p=2.9e−06
262: PFKM|P08237
cor=0.77, p=1e−07
136: HMOX1|P09601
cor=0.7, p=2.9e−06
817: FOLR1|P15328
cor=0.7, p=2.9e−06
546: ANXA11|P50995
cor=0.9, p=1.9e−13
357: VSIG4|Q9Y279
cor=0.86, p=3.6e−11
39: CD46|P15529
cor=0.56, p=0.00047
629: TBCB|Q99426
cor=0.73, p=6.5e−07
398: TPP1|O14773
cor=0.64, p=3.5e−05
586: FKBP4|Q02790
cor=0.88, p=3.3e−12
314: NUDT5|Q9UKK9
cor=−0.031, p=0.91
612: IMPA1|P29218
cor=0.26, p=0.17
553: COMT|P21964
cor=0.67, p=2e−05
527: GLRX|P35754
cor=0.88, p=3.3e−12
512: LRIG1|Q96JA1
cor=0.56, p=0.016
140: CEACAM8|P31997
cor=0.64, p=3.5e−05
552: CLSTN2|Q9H4D0
cor=0.89, p=8.5e−13
879: ENTPD2|Q9Y5L3
cor=−0.19, p=0.28
621: GBP2|P32456
cor=0.86, p=3.6e−11
865: ADGRG1|Q9Y653

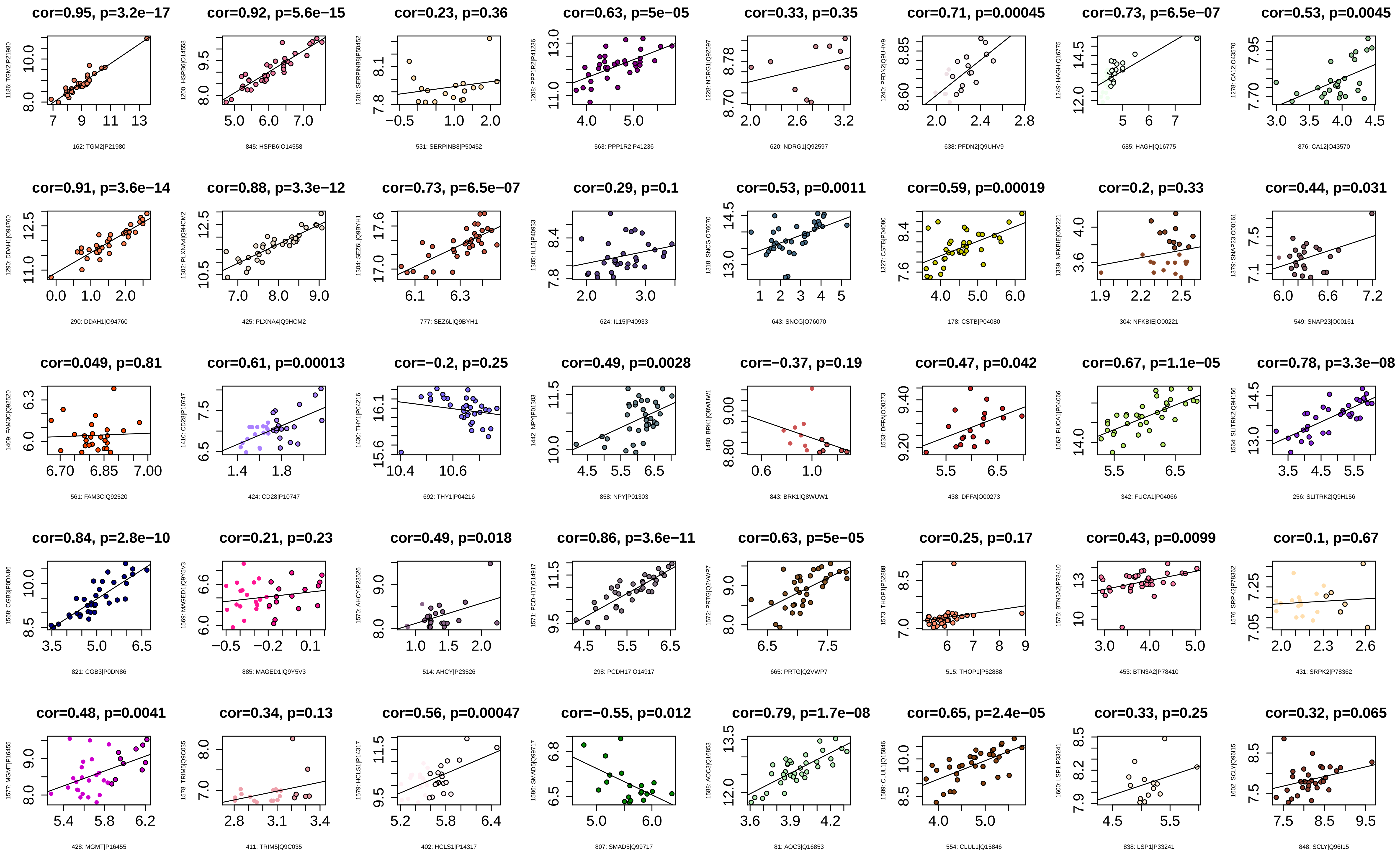
cor=0.95, p=3.2e−17
162: TGM2|P21980
cor=0.92, p=5.6e−15
845: HSPB6|O14558
cor=0.23, p=0.36
531: SERPINB8|P50452
cor=0.63, p=5e−05
563: PPP1R2|P41236
cor=0.33, p=0.35
620: NDRG1|Q92597
cor=0.71, p=0.00045
638: PFDN2|Q9UHV9
cor=0.73, p=6.5e−07
685: HAGH|Q16775
cor=0.53, p=0.0045
876: CA12|O43570
cor=0.91, p=3.6e−14
290: DDAH1|O94760
cor=0.88, p=3.3e−12
425: PLXNA4|Q9HCM2
cor=0.73, p=6.5e−07
777: SEZ6L|Q9BYH1
cor=0.29, p=0.1
624: IL15|P40933
cor=0.53, p=0.0011
643: SNCG|O76070
cor=0.59, p=0.00019
178: CSTB|P04080
cor=0.2, p=0.33
304: NFKBIE|O00221
cor=0.44, p=0.031
549: SNAP23|O00161
cor=0.049, p=0.81
561: FAM3C|Q92520
cor=0.61, p=0.00013
424: CD28|P10747
cor=−0.2, p=0.25
692: THY1|P04216
cor=0.49, p=0.0028
858: NPY|P01303
cor=−0.37, p=0.19
843: BRK1|Q8WUW1
cor=0.47, p=0.042
438: DFFA|O00273
cor=0.67, p=1.1e−05
342: FUCA1|P04066
cor=0.78, p=3.3e−08
256: SLITRK2|Q9H156
cor=0.84, p=2.8e−10
821: CGB3|P0DN86
cor=0.21, p=0.23
885: MAGED1|Q9Y5V3
cor=0.49, p=0.018
514: AHCY|P23526
cor=0.86, p=3.6e−11
298: PCDH17|O14917
cor=0.63, p=5e−05
665: PRTG|Q2VWP7
cor=0.25, p=0.17
515: THOP1|P52888
cor=0.43, p=0.0099
453: BTN3A2|P78410
cor=0.1, p=0.67
431: SRPK2|P78362
cor=0.48, p=0.0041
428: MGMT|P16455
cor=0.34, p=0.13
411: TRIM5|Q9C035
cor=0.56, p=0.00047
402: HCLS1|P14317
cor=−0.55, p=0.012
807: SMAD5|Q99717
cor=0.79, p=1.7e−08
81: AOC3|Q16853
cor=0.65, p=2.4e−05
554: CLUL1|Q15846
cor=0.33, p=0.25
838: LSP1|P33241
cor=0.32, p=0.065
848: SCLY|Q96I15

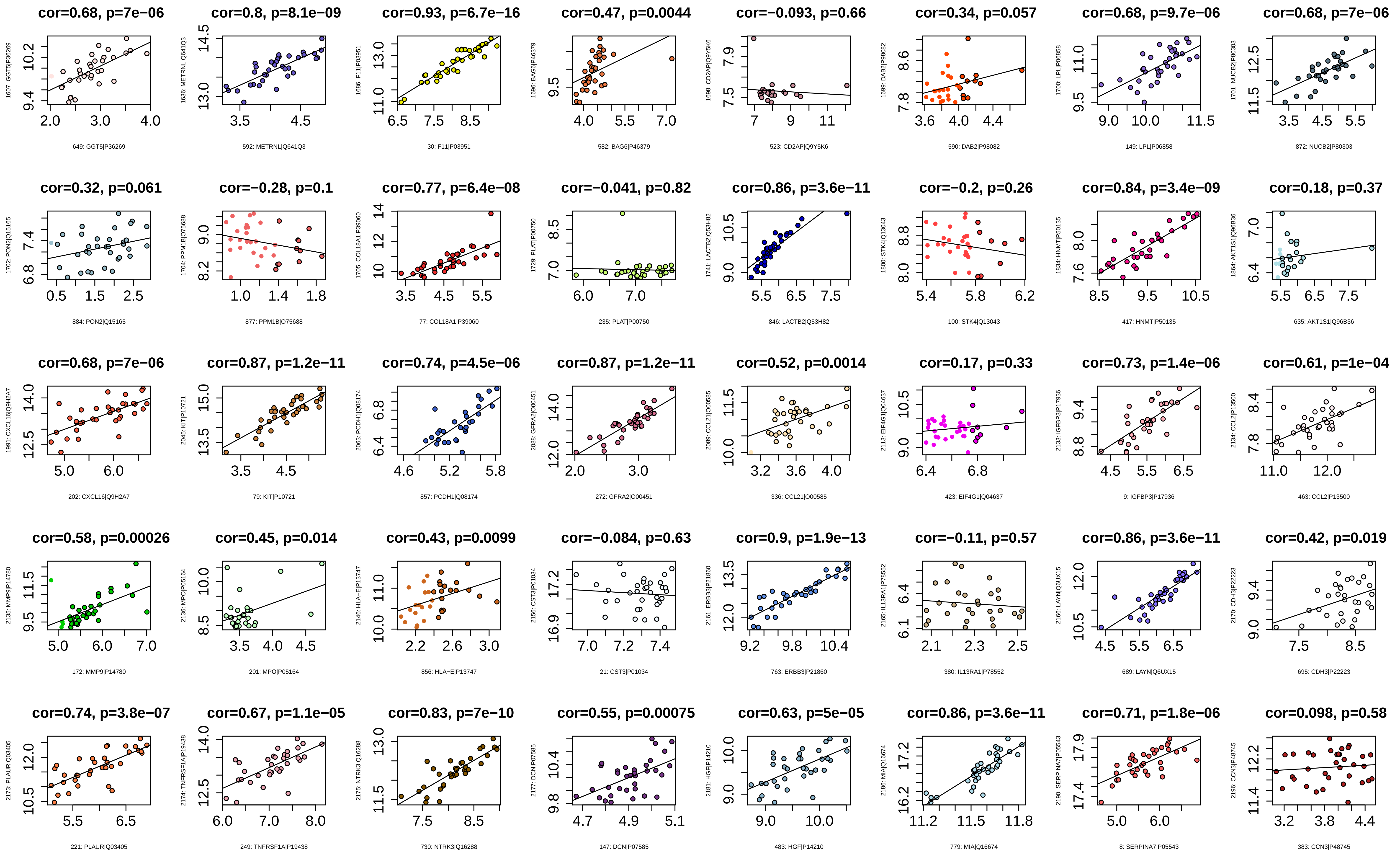

cor=0.68, p=7e−06
1607: GGT5|P36269
649: GGT5|P36269
cor=0.8, p=8.1e−09
1636: METRNL|Q641Q3
592: METRNL|Q641Q3
cor=0.93, p=6.7e−16
1686: F11|P03951
30: F11|P03951
cor=0.47, p=0.0044
1696: BAG6|P46379
582: BAG6|P46379
cor=−0.093, p=0.66
1698: CD2AP|Q9Y5K6
523: CD2AP|Q9Y5K6
cor=0.34, p=0.057
1699: DAB2|P98082
590: DAB2|P98082
cor=0.68, p=9.7e−06
1700: LPL|P06858
149: LPL|P06858
cor=0.68, p=7e−06
1701: NUCB2|P80303
872: NUCB2|P80303
cor=0.32, p=0.061
1702: PON2|Q15165
884: PON2|Q15165
cor=−0.28, p=0.1
1704: PPM1B|O75688
877: PPM1B|O75688
cor=0.77, p=6.4e−08
1705: COL18A1|P39060
77: COL18A1|P39060
cor=−0.041, p=0.82
1729: PLAT|P00750
235: PLAT|P00750
cor=0.86, p=3.6e−11
1741: LACTB2|Q53H82
846: LACTB2|Q53H82
cor=−0.2, p=0.26
1800: STK4|Q13043
100: STK4|Q13043
cor=0.84, p=3.4e−09
1834: HNMT|P50135
417: HNMT|P50135
cor=0.18, p=0.37
1864: AKT1S1|Q96B36
635: AKT1S1|Q96B36
cor=0.68, p=7e−06
1991: CXCL16|Q9H2A7
202: CXCL16|Q9H2A7
cor=0.87, p=1.2e−11
2045: KIT|P10721
79: KIT|P10721
cor=0.74, p=4.5e−06
2063: PCDH1|Q08174
857: PCDH1|Q08174
cor=0.87, p=1.2e−11
2088: GFRA2|O00451
272: GFRA2|O00451
cor=0.52, p=0.0014
2089: CCL21|O00585
336: CCL21|O00585
cor=0.17, p=0.33
2113: EIF4G1|Q04637
423: EIF4G1|Q04637
cor=0.73, p=1.4e−06
2133: IGFBP3|P17936
9: IGFBP3|P17936
cor=0.61, p=1e−04
2134: CCL2|P13500
463: CCL2|P13500
cor=0.58, p=0.00026
2135: MMP9|P14780
172: MMP9|P14780
cor=0.45, p=0.014
2136: MPO|P05164
201: MPO|P05164
cor=0.43, p=0.0099
2146: HLA−E|P13747
856: HLA−E|P13747
cor=−0.084, p=0.63
2156: CST3|P01034
21: CST3|P01034
cor=0.9, p=1.9e−13
2161: ERBB3|P21860
763: ERBB3|P21860
cor=−0.11, p=0.57
2165: IL13RA1|P78552
380: IL13RA1|P78552
cor=0.86, p=3.6e−11
2166: LAYN|Q6UX15
689: LAYN|Q6UX15
cor=0.42, p=0.019
2170: CDH3|P22223
695: CDH3|P22223
cor=0.74, p=3.8e−07
2173: PLAUR|Q03405
221: PLAUR|Q03405
cor=0.67, p=1.1e−05
2174: TNFRSF1A|P19438
249: TNFRSF1A|P19438
cor=0.83, p=7e−10
2175: NTRK3|Q16288
730: NTRK3|Q16288
cor=0.55, p=0.00075
2177: DCN|P07585
147: DCN|P07585
cor=0.63, p=5e−05
2181: HGF|P14210
483: HGF|P14210
cor=0.86, p=3.6e−11
2186: MIA|Q16674
779: MIA|Q16674
cor=0.71, p=1.8e−06
2190: SERPINA7|P05543
8: SERPINA7|P05543
cor=0.098, p=0.58
2196: CCN3|P48745
383: CCN3|P48745

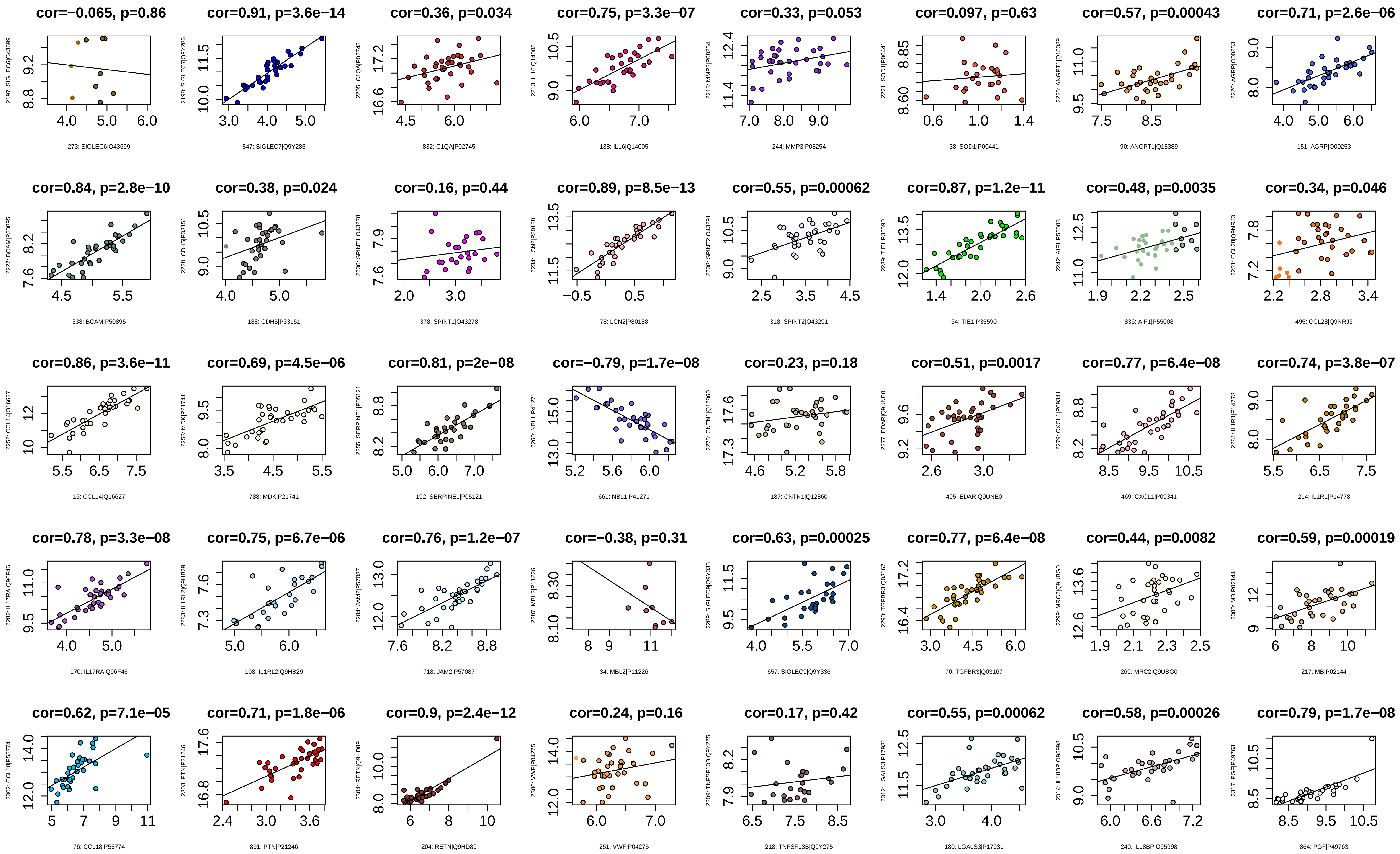
cor=−0.065, p=0.86
273: SIGLEC6|O43699
cor=0.91, p=3.6e−14
547: SIGLEC7|Q9Y286
cor=0.36, p=0.034
832: C1QA|P02745
cor=0.75, p=3.3e−07
138: IL16|Q14005
cor=0.33, p=0.053
244: MMP3|P08254
cor=0.097, p=0.63
38: SOD1|P00441
cor=0.57, p=0.00043
90: ANGPT1|Q15389
cor=0.71, p=2.6e−06
151: AGRP|O00253
cor=0.84, p=2.8e−10
338: BCAM|P50895
cor=0.38, p=0.024
188: CDH5|P33151
cor=0.16, p=0.44
378: SPINT1|O43278
cor=0.89, p=8.5e−13
78: LCN2|P80188
cor=0.55, p=0.00062
318: SPINT2|O43291
cor=0.87, p=1.2e−11
64: TIE1|P35590
cor=0.48, p=0.0035
836: AIF1|P55008
cor=0.34, p=0.046
495: CCL28|Q9NRJ3
cor=0.86, p=3.6e−11
16: CCL14|Q16627
cor=0.69, p=4.5e−06
788: MDK|P21741
cor=0.81, p=2e−08
192: SERPINE1|P05121
cor=−0.79, p=1.7e−08
661: NBL1|P41271
cor=0.23, p=0.18
187: CNTN1|Q12860
cor=0.51, p=0.0017
405: EDAR|Q9UNE0
cor=0.77, p=6.4e−08
469: CXCL1|P09341
cor=0.74, p=3.8e−07
214: IL1R1|P14778
cor=0.78, p=3.3e−08
170: IL17RA|Q96F46
cor=0.75, p=6.7e−06
108: IL1RL2|Q9HB29
cor=0.76, p=1.2e−07
718: JAM2|P57087
cor=−0.38, p=0.31
34: MBL2|P11226
cor=0.63, p=0.00025
657: SIGLEC9|Q9Y336
cor=0.77, p=6.4e−08
70: TGFBR3|Q03167
cor=0.44, p=0.0082
269: MRC2|Q9UBG0
cor=0.59, p=0.00019
217: MB|P02144
cor=0.62, p=7.1e−05
76: CCL18|P55774
cor=0.71, p=1.8e−06
891: PTN|P21246
cor=0.9, p=2.4e−12
204: RETN|Q9HD89
cor=0.24, p=0.16
251: VWF|P04275
cor=0.17, p=0.42
218: TNFSF13B|Q9Y275
cor=0.55, p=0.00062
180: LGALS3|P17931
cor=0.58, p=0.00026
240: IL18BP|O95998
cor=0.79, p=1.7e−08
864: PGF|P49763

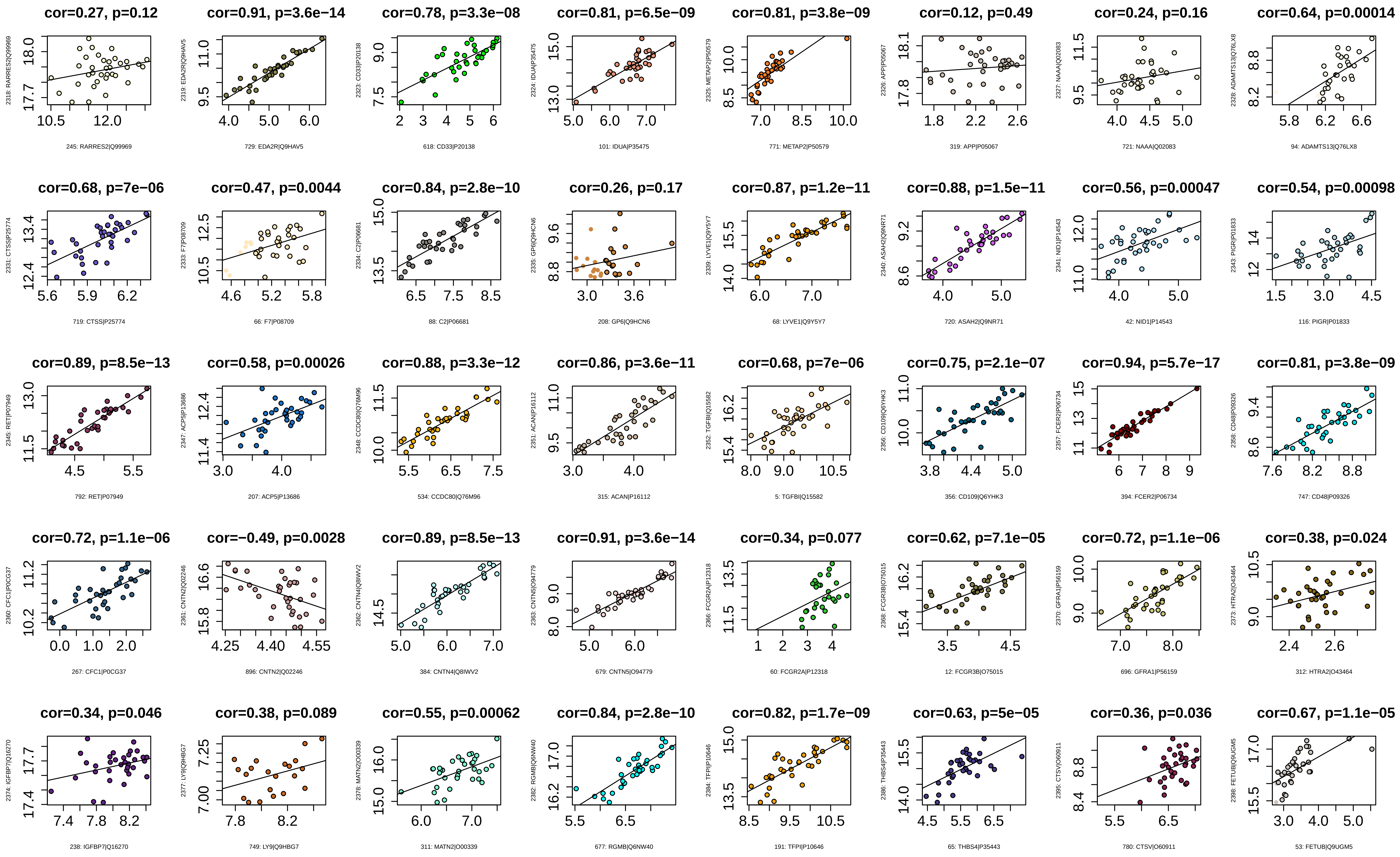
cor=0.27, p=0.12
245: RARRES2|Q99969
cor=0.91, p=3.6e−14
729: EDA2R|Q9HAV5
cor=0.78, p=3.3e−08
618: CD33|P20138
cor=0.81, p=6.5e−09
101: IDUA|P35475
cor=0.81, p=3.8e−09
771: METAP2|P50579
cor=0.12, p=0.49
319: APP|P05067
cor=0.24, p=0.16
721: NAAA|Q02083
cor=0.64, p=0.00014
94: ADAMTS13|Q76LX8
cor=0.68, p=7e−06
719: CTSS|P25774
cor=0.47, p=0.0044
66: F7|P08709
cor=0.84, p=2.8e−10
88: C2|P06681
cor=0.26, p=0.17
208: GP6|Q9HCN6
cor=0.87, p=1.2e−11
68: LYVE1|Q9Y5Y7
cor=0.88, p=1.5e−11
720: ASAH2|Q9NR71
cor=0.56, p=0.00047
42: NID1|P14543
cor=0.54, p=0.00098
116: PIGR|P01833
cor=0.89, p=8.5e−13
792: RET|P07949
cor=0.58, p=0.00026
207: ACP5|P13686
cor=0.88, p=3.3e−12
534: CCDC80|Q76M96
cor=0.86, p=3.6e−11
315: ACAN|P16112
cor=0.68, p=7e−06
5: TGFBI|Q15582
cor=0.75, p=2.1e−07
356: CD109|Q6YHK3
cor=0.94, p=5.7e−17
394: FCER2|P06734
cor=0.81, p=3.8e−09
747: CD48|P09326
cor=0.72, p=1.1e−06
267: CFC1|P0CG37
cor=−0.49, p=0.0028
896: CNTN2|Q02246
cor=0.89, p=8.5e−13
384: CNTN4|Q8IWV2
cor=0.91, p=3.6e−14
679: CNTN5|O94779
cor=0.34, p=0.077
60: FCGR2A|P12318
cor=0.62, p=7.1e−05
12: FCGR3B|O75015
cor=0.72, p=1.1e−06
696: GFRA1|P56159
cor=0.38, p=0.024
312: HTRA2|O43464
cor=0.34, p=0.046
238: IGFBP7|Q16270
cor=0.38, p=0.089
749: LY9|Q9HBG7
cor=0.55, p=0.00062
311: MATN2|O00339
cor=0.84, p=2.8e−10
677: RGMB|Q6NW40
cor=0.82, p=1.7e−09
191: TFPI|P10646
cor=0.63, p=5e−05
65: THBS4|P35443
cor=0.36, p=0.036
780: CTSV|O60911
cor=0.67, p=1.1e−05
53: FETUB|Q9UGM5

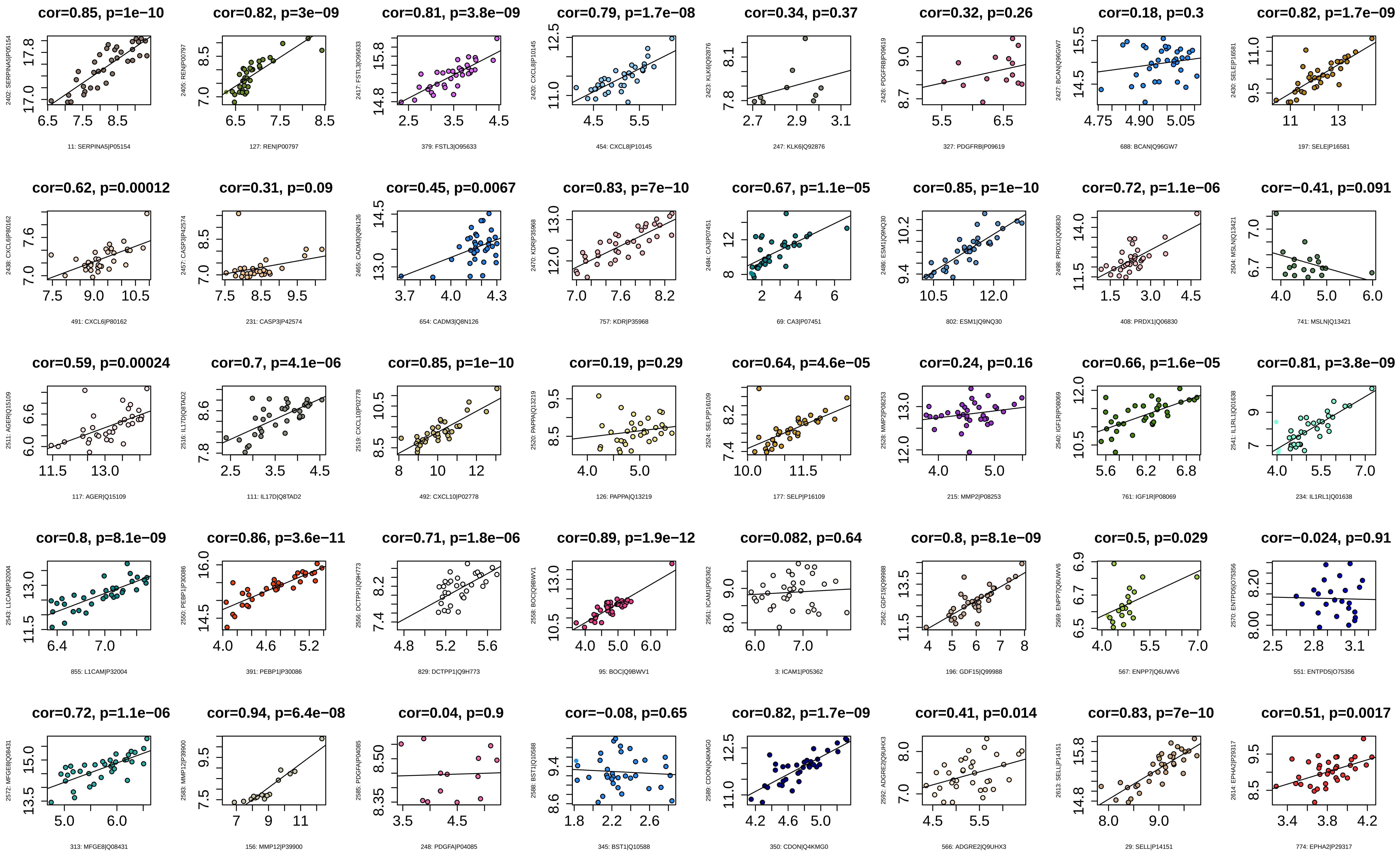
cor=0.85, p=1e−10
2402: SERPINA5|P05154
11: SERPINA5|P05154
cor=0.82, p=3e−09
2405: REN|P00797
127: REN|P00797
cor=0.81, p=3.8e−09
2417: FSTL3|O95633
379: FSTL3|O95633
cor=0.79, p=1.7e−08
2420: CXCL8|P10145
454: CXCL8|P10145
cor=0.34, p=0.37
2423: KLK6|Q92876
247: KLK6|Q92876
cor=0.32, p=0.26
2426: PDGFRB|P09619
327: PDGFRB|P09619
cor=0.18, p=0.3
2427: BCAN|Q96GW7
688: BCAN|Q96GW7
cor=0.82, p=1.7e−09
2430: SELE|P16581
197: SELE|P16581
cor=0.62, p=0.00012
2438: CXCL6|P80162
491: CXCL6|P80162
cor=0.31, p=0.09
2457: CASP3|P42574
231: CASP3|P42574
cor=0.45, p=0.0067
2465: CADM3|Q8N126
654: CADM3|Q8N126
cor=0.83, p=7e−10
2470: KDR|P35968
757: KDR|P35968
cor=0.67, p=1.1e−05
2484: CA3|P07451
69: CA3|P07451
cor=0.85, p=1e−10
2486: ESM1|Q9NQ30
802: ESM1|Q9NQ30
cor=0.72, p=1.1e−06
2498: PRDX1|Q06830
408: PRDX1|Q06830
cor=−0.41, p=0.091
2504: MSLN|Q13421
741: MSLN|Q13421
cor=0.59, p=0.00024
2511: AGER|Q15109
117: AGER|Q15109
cor=0.7, p=4.1e−06
2516: IL17D|Q8TAD2
111: IL17D|Q8TAD2
cor=0.85, p=1e−10
2519: CXCL10|P02778
492: CXCL10|P02778
cor=0.19, p=0.29
2520: PAPPA|Q13219
126: PAPPA|Q13219
cor=0.64, p=4.6e−05
2524: SELP|P16109
177: SELP|P16109
cor=0.24, p=0.16
2528: MMP2|P08253
215: MMP2|P08253
cor=0.66, p=1.6e−05
2540: IGF1R|P08069
761: IGF1R|P08069
cor=0.81, p=3.8e−09
2541: IL1RL1|Q01638
234: IL1RL1|Q01638
cor=0.8, p=8.1e−09
2543: L1CAM|P32004
855: L1CAM|P32004
cor=0.86, p=3.6e−11
2550: PEBP1|P30086
391: PEBP1|P30086
cor=0.71, p=1.8e−06
2556: DCTPP1|Q9H773
829: DCTPP1|Q9H773
cor=0.89, p=1.9e−12
2558: BOC|Q9BWV1
95: BOC|Q9BWV1
cor=0.082, p=0.64
2561: ICAM1|P05362
3: ICAM1|P05362
cor=0.8, p=8.1e−09
2562: GDF15|Q99988
196: GDF15|Q99988
cor=0.5, p=0.029
2569: ENPP7|Q6UWV6
567: ENPP7|Q6UWV6
cor=−0.024, p=0.91
2570: ENTPD5|O75356
551: ENTPD5|O75356
cor=0.72, p=1.1e−06
2572: MFGE8|Q08431
313: MFGE8|Q08431
cor=0.94, p=6.4e−08
2583: MMP12|P39900
156: MMP12|P39900
cor=0.04, p=0.9
2585: PDGFA|P04085
248: PDGFA|P04085
cor=−0.08, p=0.65
2588: BST1|Q10588
345: BST1|Q10588
cor=0.82, p=1.7e−09
2589: CDON|Q4KMG0
350: CDON|Q4KMG0
cor=0.41, p=0.014
2592: ADGRE2|Q9UHX3
566: ADGRE2|Q9UHX3
cor=0.83, p=7e−10
2613: SELL|P14151
29: SELL|P14151
cor=0.51, p=0.0017
2614: EPHA2|P29317
774: EPHA2|P29317

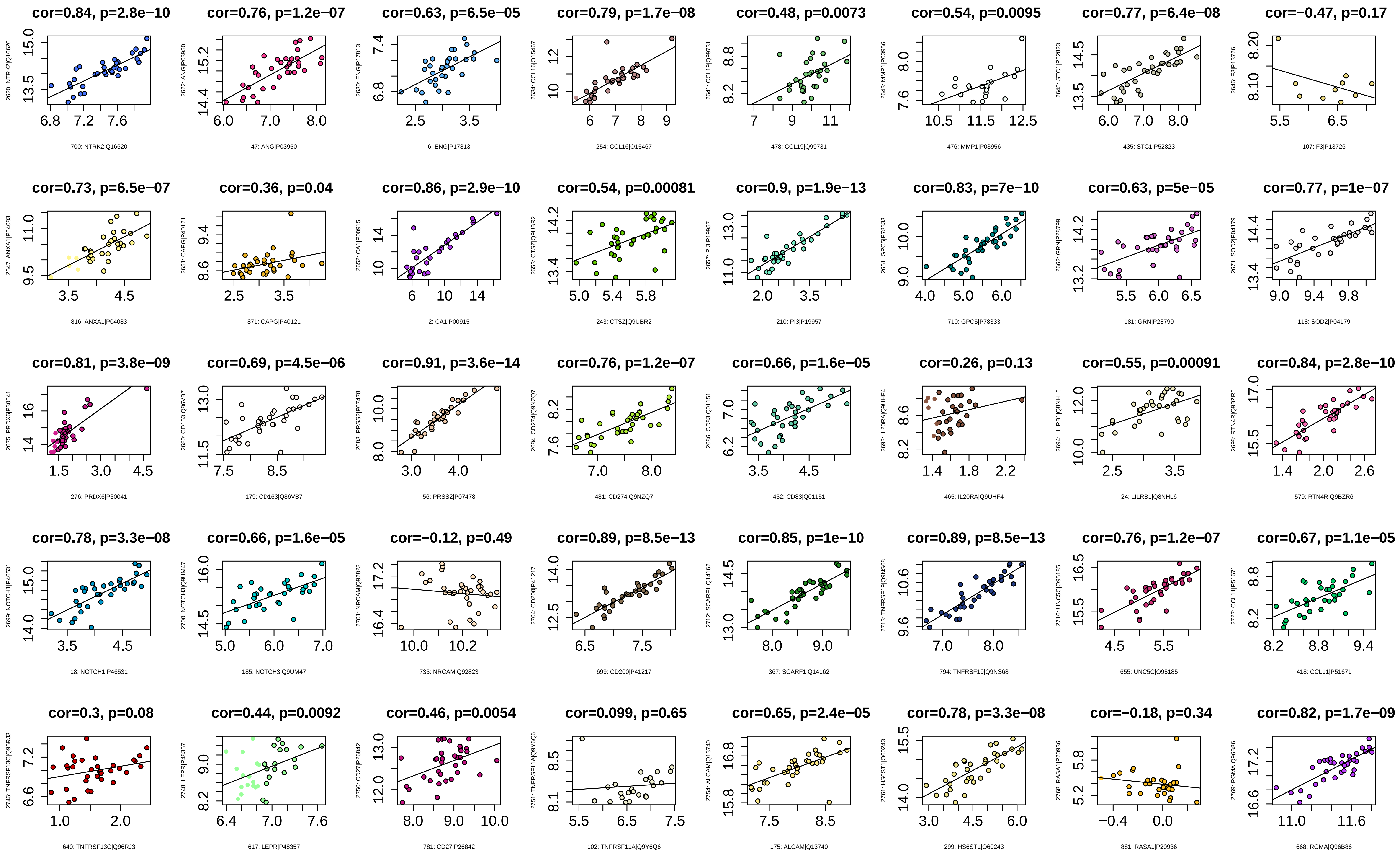
cor=0.84, p=2.8e−10
700: NTRK2|Q16620
cor=0.76, p=1.2e−07
47: ANG|P03950
cor=0.63, p=6.5e−05
6: ENG|P17813
cor=0.79, p=1.7e−08
254: CCL16|O15467
cor=0.48, p=0.0073
478: CCL19|Q99731
cor=0.54, p=0.0095
476: MMP1|P03956
cor=0.77, p=6.4e−08
435: STC1|P52823
cor=−0.47, p=0.17
107: F3|P13726
cor=0.73, p=6.5e−07
816: ANXA1|P04083
cor=0.36, p=0.04
871: CAPG|P40121
cor=0.86, p=2.9e−10
2: CA1|P00915
cor=0.54, p=0.00081
243: CTSZ|Q9UBR2
cor=0.9, p=1.9e−13
210: PI3|P19957
cor=0.83, p=7e−10
710: GPC5|P78333
cor=0.63, p=5e−05
181: GRN|P28799
cor=0.77, p=1e−07
118: SOD2|P04179
cor=0.81, p=3.8e−09
276: PRDX6|P30041
cor=0.69, p=4.5e−06
179: CD163|Q86VB7
cor=0.91, p=3.6e−14
56: PRSS2|P07478
cor=0.76, p=1.2e−07
481: CD274|Q9NZQ7
cor=0.66, p=1.6e−05
452: CD83|Q01151
cor=0.26, p=0.13
465: IL20RA|Q9UHF4
cor=0.55, p=0.00091
24: LILRB1|Q8NHL6
cor=0.84, p=2.8e−10
579: RTN4R|Q9BZR6
cor=0.78, p=3.3e−08
18: NOTCH1|P46531
cor=0.66, p=1.6e−05
185: NOTCH3|Q9UM47
cor=−0.12, p=0.49
735: NRCAM|Q92823
cor=0.89, p=8.5e−13
699: CD200|P41217
cor=0.85, p=1e−10
367: SCARF1|Q14162
cor=0.89, p=8.5e−13
794: TNFRSF19|Q9NS68
cor=0.76, p=1.2e−07
655: UNC5C|O95185
cor=0.67, p=1.1e−05
418: CCL11|P51671
cor=0.3, p=0.08
640: TNFRSF13C|Q96RJ3
cor=0.44, p=0.0092
617: LEPR|P48357
cor=0.46, p=0.0054
781: CD27|P26842
cor=0.099, p=0.65
102: TNFRSF11A|Q9Y6Q6
cor=0.65, p=2.4e−05
175: ALCAM|Q13740
cor=0.78, p=3.3e−08
299: HS6ST1|O60243
cor=−0.18, p=0.34
881: RASA1|P20936
cor=0.82, p=1.7e−09
668: RGMA|Q96B86

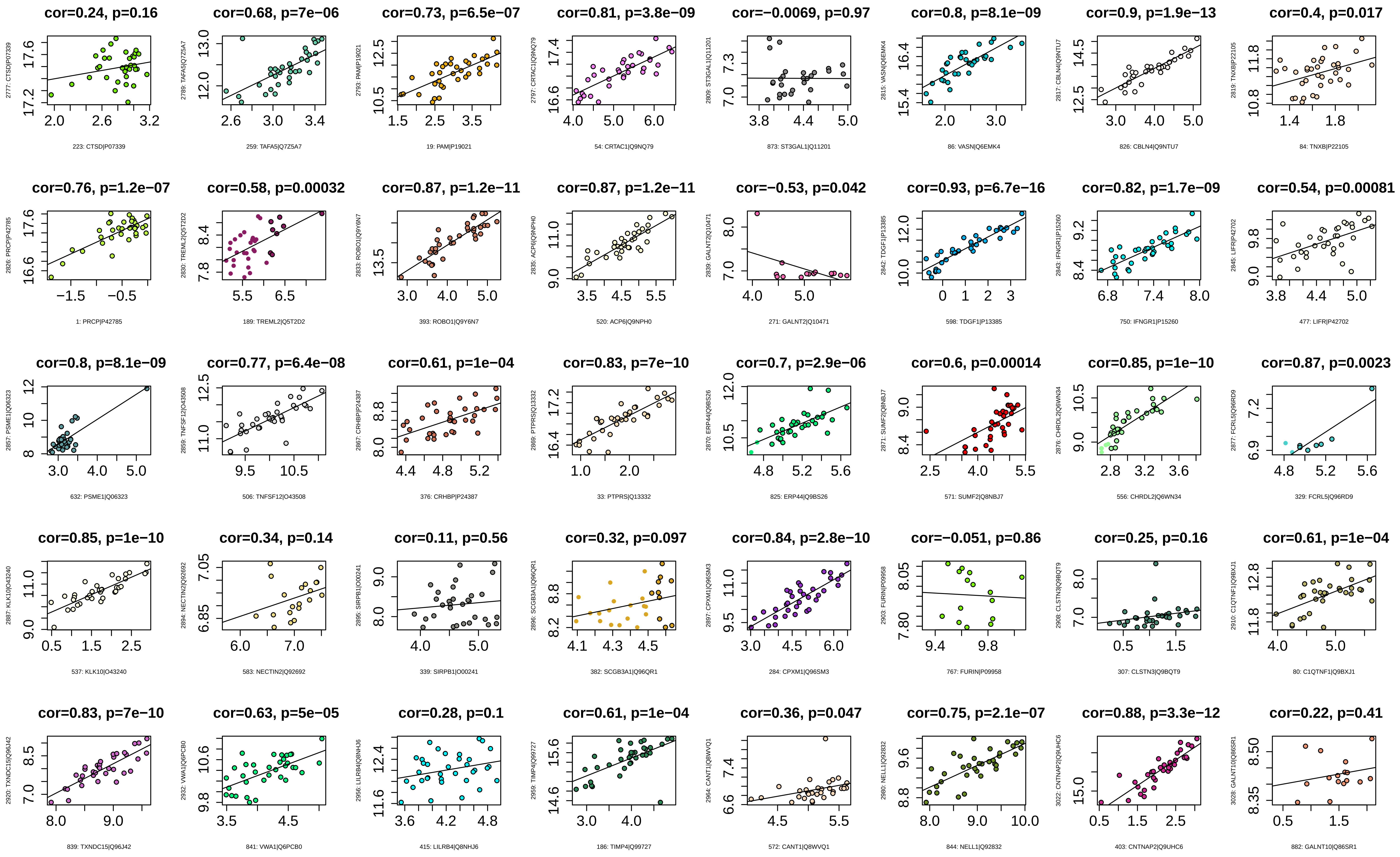
cor=0.24, p=0.16
2777: CTSD|P07339
223: CTSD|P07339
cor=0.68, p=7e−06
2789: TAFA5|Q7Z5A7
259: TAFA5|Q7Z5A7
cor=0.73, p=6.5e−07
2793: PAM|P19021
19: PAM|P19021
cor=0.81, p=3.8e−09
2797: CRTAC1|Q9NQ79
54: CRTAC1|Q9NQ79
cor=−0.0069, p=0.97
2809: ST3GAL1|Q11201
873: ST3GAL1|Q11201
cor=0.8, p=8.1e−09
2815: VASN|Q6EMK4
86: VASN|Q6EMK4
cor=0.9, p=1.9e−13
2817: CBLN4|Q9NTU7
826: CBLN4|Q9NTU7
cor=0.4, p=0.017
2819: TNXB|P22105
84: TNXB|P22105
cor=0.76, p=1.2e−07
2826: PRCP|P42785
1: PRCP|P42785
cor=0.58, p=0.00032
2830: TREML2|Q5T2D2
189: TREML2|Q5T2D2
cor=0.87, p=1.2e−11
2833: ROBO1|Q9Y6N7
393: ROBO1|Q9Y6N7
cor=0.87, p=1.2e−11
2835: ACP6|Q9NPH0
520: ACP6|Q9NPH0
cor=−0.53, p=0.042
2839: GALNT2|Q10471
271: GALNT2|Q10471
cor=0.93, p=6.7e−16
2842: TDGF1|P13385
598: TDGF1|P13385
cor=0.82, p=1.7e−09
2843: IFNGR1|P15260
750: IFNGR1|P15260
cor=0.54, p=0.00081
2845: LIFR|P42702
477: LIFR|P42702
cor=0.8, p=8.1e−09
2857: PSME1|Q06323
632: PSME1|Q06323
cor=0.77, p=6.4e−08
2859: TNFSF12|O43508
506: TNFSF12|O43508
cor=0.61, p=1e−04
2867: CRHBP|P24387
376: CRHBP|P24387
cor=0.83, p=7e−10
2869: PTPRS|Q13332
33: PTPRS|Q13332
cor=0.7, p=2.9e−06
2870: ERP44|Q9BS26
825: ERP44|Q9BS26
cor=0.6, p=0.00014
2871: SUMF2|Q8NBJ7
571: SUMF2|Q8NBJ7
cor=0.85, p=1e−10
2876: CHRDL2|Q6WN34
556: CHRDL2|Q6WN34
cor=0.87, p=0.0023
2877: FCRL5|Q96RD9
329: FCRL5|Q96RD9
cor=0.85, p=1e−10
2887: KLK10|O43240
537: KLK10|O43240
cor=0.34, p=0.14
2894: NECTIN2|Q92692
583: NECTIN2|Q92692
cor=0.11, p=0.56
2895: SIRPB1|O00241
339: SIRPB1|O00241
cor=0.32, p=0.097
2896: SCGB3A1|Q96QR1
382: SCGB3A1|Q96QR1
cor=0.84, p=2.8e−10
2897: CPXM1|Q96SM3
284: CPXM1|Q96SM3
cor=−0.051, p=0.86
2903: FURIN|P09958
767: FURIN|P09958
cor=0.25, p=0.16
2908: CLSTN3|Q9BQT9
307: CLSTN3|Q9BQT9
cor=0.61, p=1e−04
2910: C1QTNF1|Q9BXJ1
80: C1QTNF1|Q9BXJ1
cor=0.83, p=7e−10
2920: TXNDC15|Q96J42
839: TXNDC15|Q96J42
cor=0.63, p=5e−05
2932: VWA1|Q6PCB0
841: VWA1|Q6PCB0
cor=0.28, p=0.1
2956: LILRB4|Q8NHJ6
415: LILRB4|Q8NHJ6
cor=0.61, p=1e−04
2959: TIMP4|Q99727
186: TIMP4|Q99727
cor=0.36, p=0.047
2964: CANT1|Q8WVQ1
572: CANT1|Q8WVQ1
cor=0.75, p=2.1e−07
2980: NELL1|Q92832
844: NELL1|Q92832
cor=0.88, p=3.3e−12
3022: CNTNAP2|Q9UHC6
403: CNTNAP2|Q9UHC6
cor=0.22, p=0.41
3028: GALNT10|Q86SR1
882: GALNT10|Q86SR1

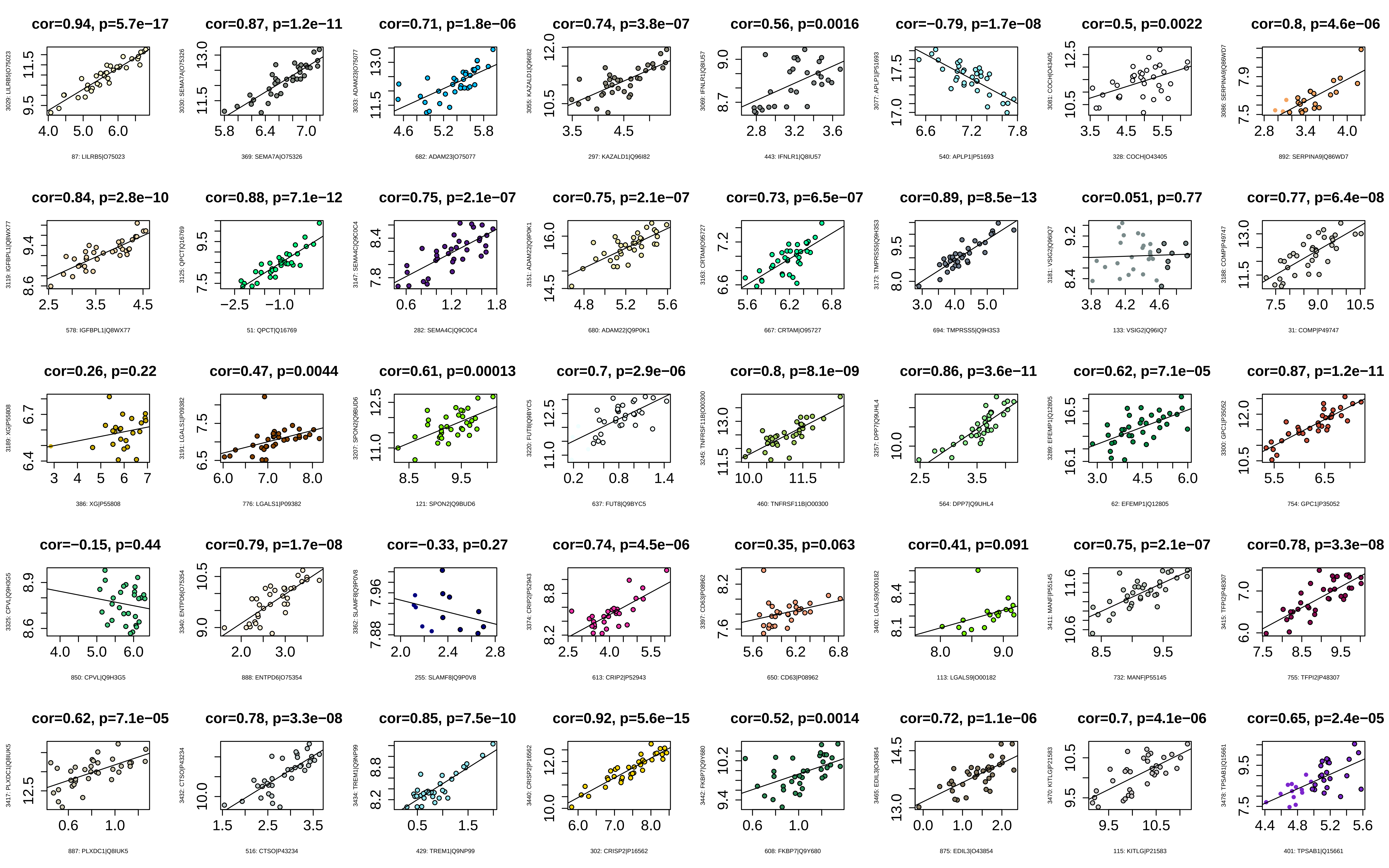
cor=0.94, p=5.7e−17
87: LILRB5|O75023
cor=0.87, p=1.2e−11
369: SEMA7A|O75326
cor=0.71, p=1.8e−06
682: ADAM23|O75077
cor=0.74, p=3.8e−07
297: KAZALD1|Q96I82
cor=0.56, p=0.0016
443: IFNLR1|Q8IU57
cor=−0.79, p=1.7e−08
540: APLP1|P51693
cor=0.5, p=0.0022
328: COCH|O43405
cor=0.8, p=4.6e−06
892: SERPINA9|Q86WD7
cor=0.84, p=2.8e−10
578: IGFBPL1|Q8WX77
cor=0.88, p=7.1e−12
51: QPCT|Q16769
cor=0.75, p=2.1e−07
282: SEMA4C|Q9C0C4
cor=0.75, p=2.1e−07
680: ADAM22|Q9P0K1
cor=0.73, p=6.5e−07
667: CRTAM|O95727
cor=0.89, p=8.5e−13
694: TMPRSS5|Q9H3S3
cor=0.051, p=0.77
133: VSIG2|Q96IQ7
cor=0.77, p=6.4e−08
31: COMP|P49747
cor=0.26, p=0.22
386: XG|P55808
cor=0.47, p=0.0044
776: LGALS1|P09382
cor=0.61, p=0.00013
121: SPON2|Q9BUD6
cor=0.7, p=2.9e−06
637: FUT8|Q9BYC5
cor=0.8, p=8.1e−09
460: TNFRSF11B|O00300
cor=0.86, p=3.6e−11
564: DPP7|Q9UHL4
cor=0.62, p=7.1e−05
62: EFEMP1|Q12805
cor=0.87, p=1.2e−11
754: GPC1|P35052
cor=−0.15, p=0.44
850: CPVL|Q9H3G5
cor=0.79, p=1.7e−08
888: ENTPD6|O75354
cor=−0.33, p=0.27
255: SLAMF8|Q9P0V8
cor=0.74, p=4.5e−06
613: CRIP2|P52943
cor=0.35, p=0.063
650: CD63|P08962
cor=0.41, p=0.091
113: LGALS9|O00182
cor=0.75, p=2.1e−07
732: MANF|P55145
cor=0.78, p=3.3e−08
755: TFPI2|P48307
cor=0.62, p=7.1e−05
887: PLXDC1|Q8IUK5
cor=0.78, p=3.3e−08
516: CTSO|P43234
cor=0.85, p=7.5e−10
429: TREM1|Q9NP99
cor=0.92, p=5.6e−15
302: CRISP2|P16562
cor=0.52, p=0.0014
608: FKBP7|Q9Y680
cor=0.72, p=1.1e−06
875: EDIL3|O43854
cor=0.7, p=4.1e−06
115: KITLG|P21583
cor=0.65, p=2.4e−05
401: TPSAB1|Q15661

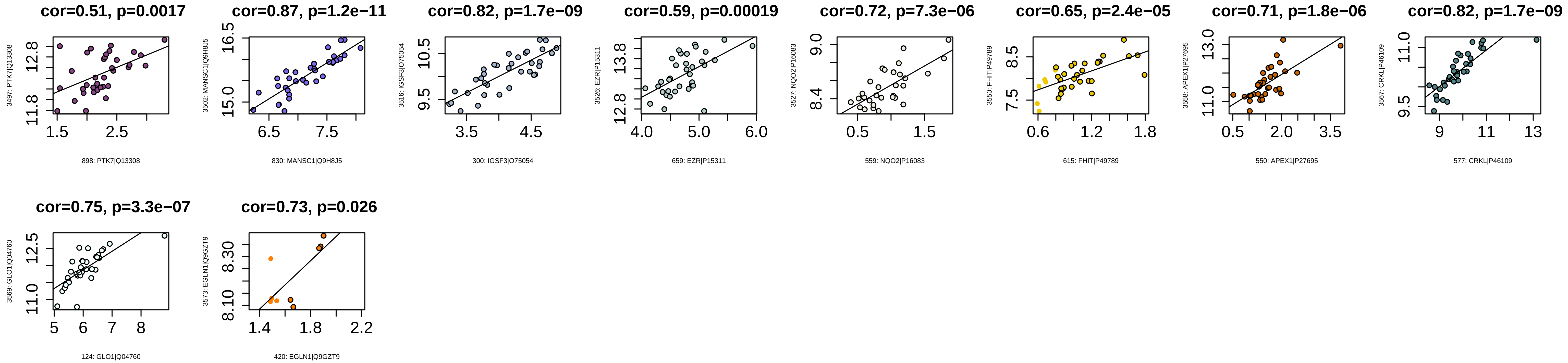
cor=0.51, p=0.0017
3497: PTK7|Q13308
898: PTK7|Q13308
cor=0.87, p=1.2e−11
3502: MANSC1|Q9H8J5
830: MANSC1|Q9H8J5
cor=0.82, p=1.7e−09
3516: IGSF3|O75054
300: IGSF3|O75054
cor=0.59, p=0.00019
3526: EZR|P15311
659: EZR|P15311
cor=0.72, p=7.3e−06
3527: NQO2|P16083
559: NQO2|P16083
cor=0.65, p=2.4e−05
3550: FHIT|P49789
615: FHIT|P49789
cor=0.71, p=1.8e−06
3558: APEX1|P27695
550: APEX1|P27695
cor=0.82, p=1.7e−09
3567: CRKL|P46109
577: CRKL|P46109
cor=0.75, p=3.3e−07
3569: GLO1|Q04760
124: GLO1|Q04760
cor=0.73, p=0.026
3573: EGLN1|Q9GZT9
420: EGLN1|Q9GZT9

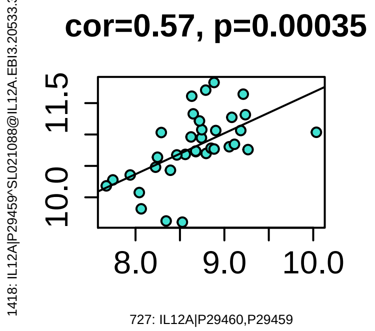
cor=0.57, p=0.00035
1418: IL12A|P29459^SL021088@IL12A.EBI3.20533.39
10.0
11.5
8.0
9.0
10.0
727: IL12A|P29460,P29459

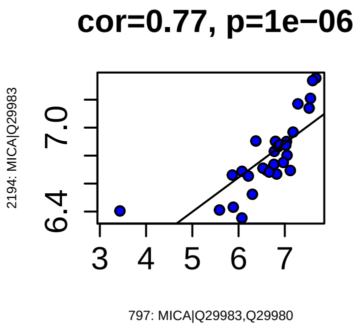
cor=0.77, p=1e−06
2194: MICA|Q29983
6.4
7.0
3
4
5
6
7
797: MICA|Q29983,Q29980

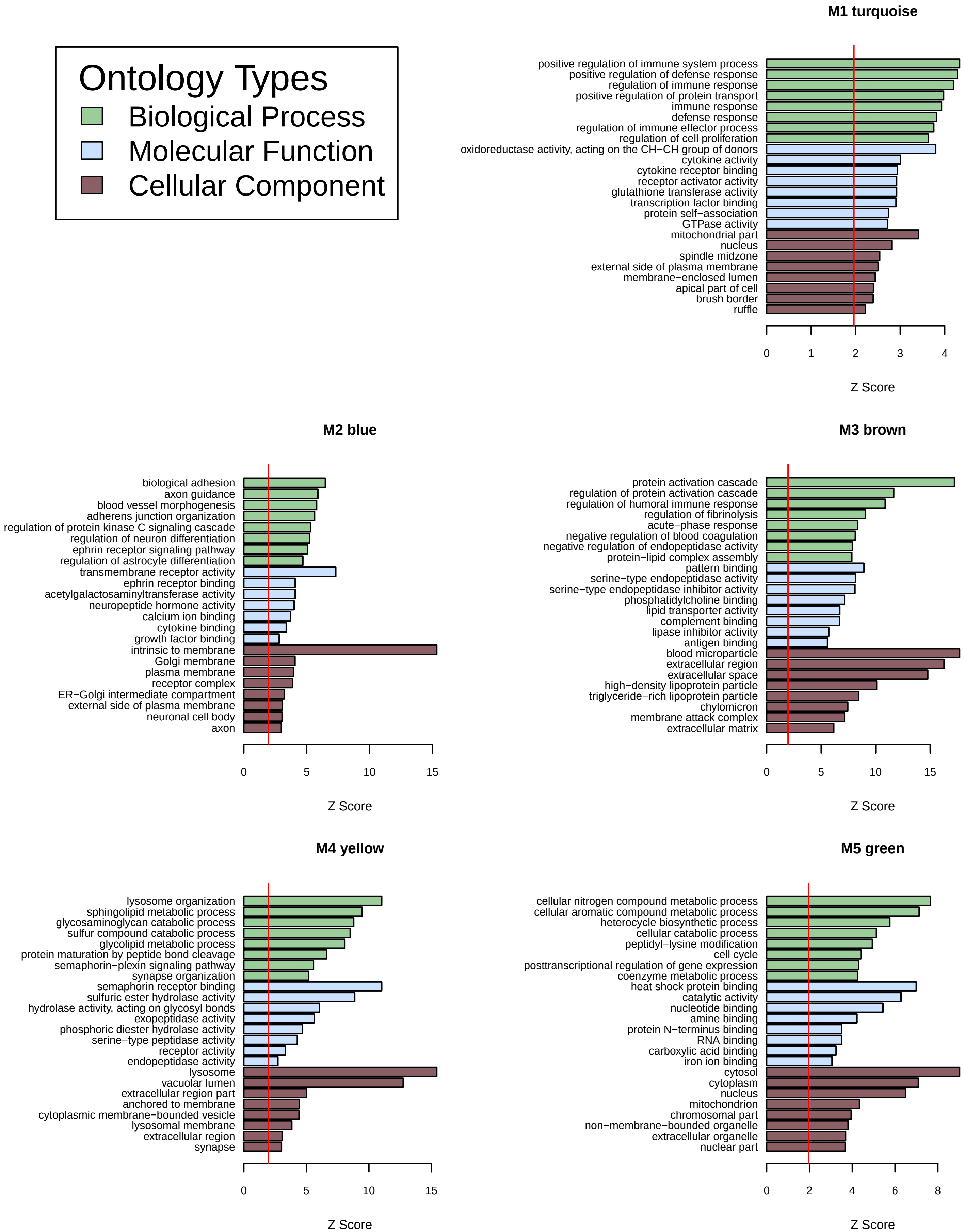
Ontology Types
Biological Process
Molecular Function
Cellular Component
M1 turquoise
positive regulation of immune system process
positive regulation of defense response
regulation of immune response
positive regulation of protein transport
immune response
defense response
regulation of immune effector process
regulation of cell proliferation
oxidoreductase activity, acting on the CH−CH group of donors
cytokine activity
cytokine receptor binding
receptor activator activity
glutathione transferase activity
transcription factor binding
protein self−association
GTPase activity
mitochondrial part
nucleus
spindle midzone
external side of plasma membrane
membrane−enclosed lumen
apical part of cell
brush border
ruffle
0
1
2
3
4
Z Score
M2 blue
biological adhesion
axon guidance
blood vessel morphogenesis
adherens junction organization
regulation of protein kinase C signaling cascade
regulation of neuron differentiation
ephrin receptor signaling pathway
regulation of astrocyte differentiation
transmembrane receptor activity
ephrin receptor binding
acetylgalactosaminyltransferase activity
neuropeptide hormone activity
calcium ion binding
cytokine binding
growth factor binding
intrinsic to membrane
Golgi membrane
plasma membrane
receptor complex
ER−Golgi intermediate compartment
external side of plasma membrane
neuronal cell body
axon
0
5
10
15
Z Score
M3 brown
protein activation cascade
regulation of protein activation cascade
regulation of humoral immune response
regulation of fibrinolysis
acute−phase response
negative regulation of blood coagulation
negative regulation of endopeptidase activity
protein−lipid complex assembly
pattern binding
serine−type endopeptidase activity
serine−type endopeptidase inhibitor activity
phosphatidylcholine binding
lipid transporter activity
complement binding
lipase inhibitor activity
antigen binding
blood microparticle
extracellular region
extracellular space
high−density lipoprotein particle
triglyceride−rich lipoprotein particle
chylomicron
membrane attack complex
extracellular matrix
0
5
10
15
Z Score
M4 yellow
lysosome organization
sphingolipid metabolic process
glycosaminoglycan catabolic process
sulfur compound catabolic process
glycolipid metabolic process
protein maturation by peptide bond cleavage
semaphorin−plexin signaling pathway
synapse organization
semaphorin receptor binding
sulfuric ester hydrolase activity
hydrolase activity, acting on glycosyl bonds
exopeptidase activity
phosphoric diester hydrolase activity
serine−type peptidase activity
receptor activity
endopeptidase activity
lysosome
vacuolar lumen
extracellular region part
anchored to membrane
cytoplasmic membrane−bounded vesicle
lysosomal membrane
extracellular region
synapse
0
5
10
15
Z Score
M5 green
cellular nitrogen compound metabolic process
cellular aromatic compound metabolic process
heterocycle biosynthetic process
cellular catabolic process
peptidyl−lysine modification
cell cycle
posttranscriptional regulation of gene expression
coenzyme metabolic process
heat shock protein binding
catalytic activity
nucleotide binding
amine binding
protein N−terminus binding
RNA binding
carboxylic acid binding
iron ion binding
cytosol
cytoplasm
nucleus
mitochondrion
chromosomal part
non−membrane−bounded organelle
extracellular organelle
nuclear part
0
2
4
6
8
Z Score

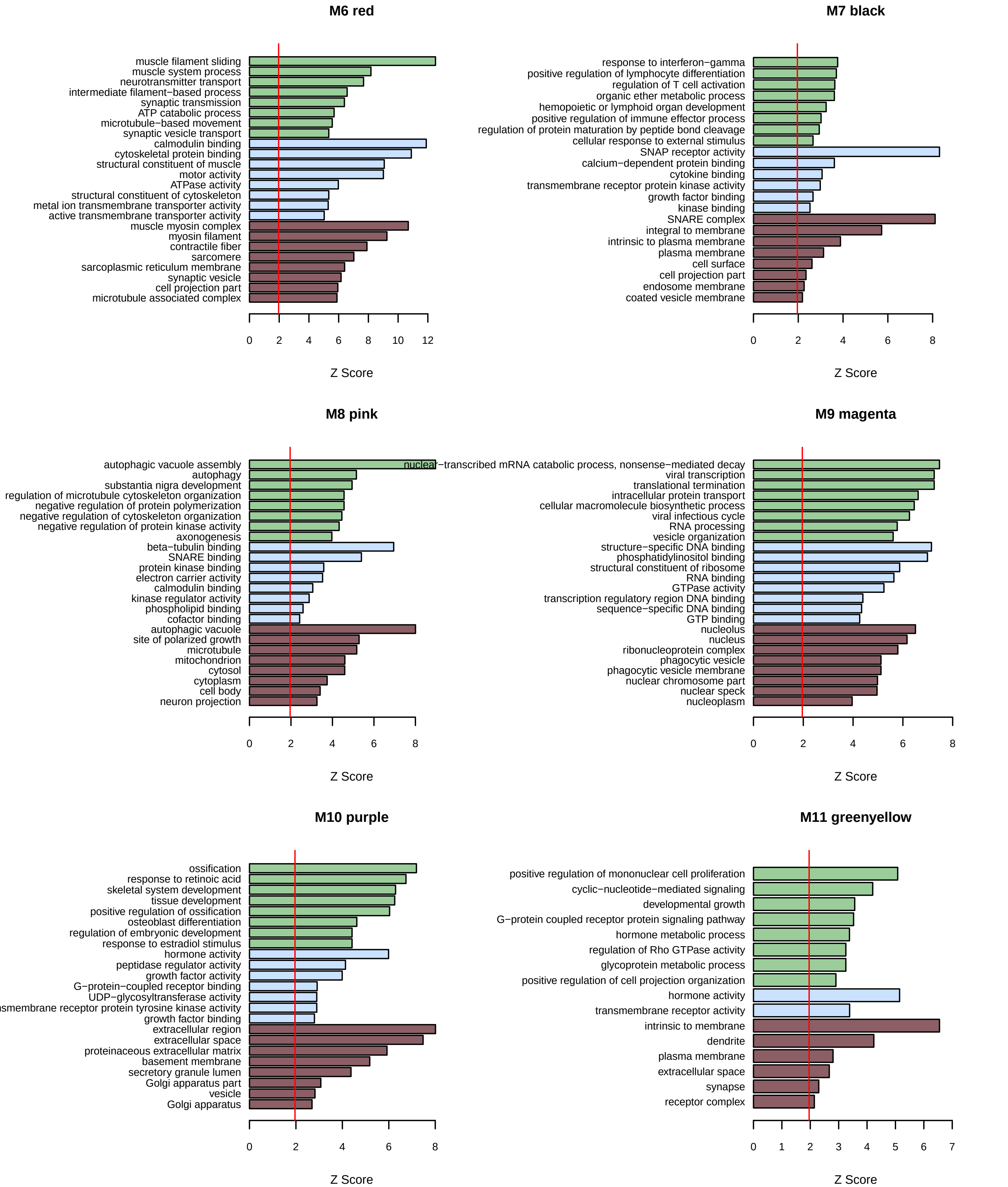
M6 red
muscle filament sliding
muscle system process
neurotransmitter transport
intermediate filament−based process
synaptic transmission
ATP catabolic process
microtubule−based movement
synaptic vesicle transport
calmodulin binding
cytoskeletal protein binding
structural constituent of muscle
motor activity
ATPase activity
structural constituent of cytoskeleton
metal ion transmembrane transporter activity
active transmembrane transporter activity
muscle myosin complex
myosin filament
contractile fiber
sarcomere
sarcoplasmic reticulum membrane
synaptic vesicle
cell projection part
microtubule associated complex
0 2 4 6 8 10 12
Z Score
M7 black
response to interferon−gamma
positive regulation of lymphocyte differentiation
regulation of T cell activation
organic ether metabolic process
hemopoietic or lymphoid organ development
positive regulation of immune effector process
regulation of protein maturation by peptide bond cleavage
cellular response to external stimulus
SNAP receptor activity
calcium−dependent protein binding
cytokine binding
transmembrane receptor protein kinase activity
growth factor binding
kinase binding
SNARE complex
integral to membrane
intrinsic to plasma membrane
plasma membrane
cell surface
cell projection part
endosome membrane
coated vesicle membrane
0 2 4 6 8
Z Score
M8 pink
autophagic vacuole assembly
autophagy
substantia nigra development
regulation of microtubule cytoskeleton organization
negative regulation of protein polymerization
negative regulation of cytoskeleton organization
negative regulation of protein kinase activity
axonogenesis
beta−tubulin binding
SNARE binding
protein kinase binding
electron carrier activity
calmodulin binding
kinase regulator activity
phospholipid binding
cofactor binding
autophagic vacuole
site of polarized growth
microtubule
mitochondrion
cytosol
cytoplasm
cell body
neuron projection
0 2 4 6 8
Z Score
M9 magenta
nuclear−transcribed mRNA catabolic process, nonsense−mediated decay
viral transcription
translational termination
intracellular protein transport
cellular macromolecule biosynthetic process
viral infectious cycle
RNA processing
vesicle organization
structure−specific DNA binding
phosphatidylinositol binding
structural constituent of ribosome
RNA binding
GTPase activity
transcription regulatory region DNA binding
sequence−specific DNA binding
GTP binding
nucleolus
nucleus
ribonucleoprotein complex
phagocytic vesicle
phagocytic vesicle membrane
nuclear chromosome part
nuclear speck
nucleoplasm
0 2 4 6 8
Z Score
M10 purple
ossification
response to retinoic acid
skeletal system development
tissue development
positive regulation of ossification
osteoblast differentiation
regulation of embryonic development
response to estradiol stimulus
hormone activity
peptidase regulator activity
growth factor activity
G−protein−coupled receptor binding
UDP−glycosyltransferase activity
nsmembrane receptor protein tyrosine kinase activity
growth factor binding
extracellular region
extracellular space
proteinaceous extracellular matrix
basement membrane
secretory granule lumen
Golgi apparatus part
vesicle
Golgi apparatus
0 2 4 6 8
Z Score
M11 greenyellow
positive regulation of mononuclear cell proliferation
cyclic−nucleotide−mediated signaling
developmental growth
G−protein coupled receptor protein signaling pathway
hormone metabolic process
regulation of Rho GTPase activity
glycoprotein metabolic process
positive regulation of cell projection organization
hormone activity
transmembrane receptor activity
intrinsic to membrane
dendrite
plasma membrane
extracellular space
synapse
receptor complex
0 1 2 3 4 5 6 7
Z Score

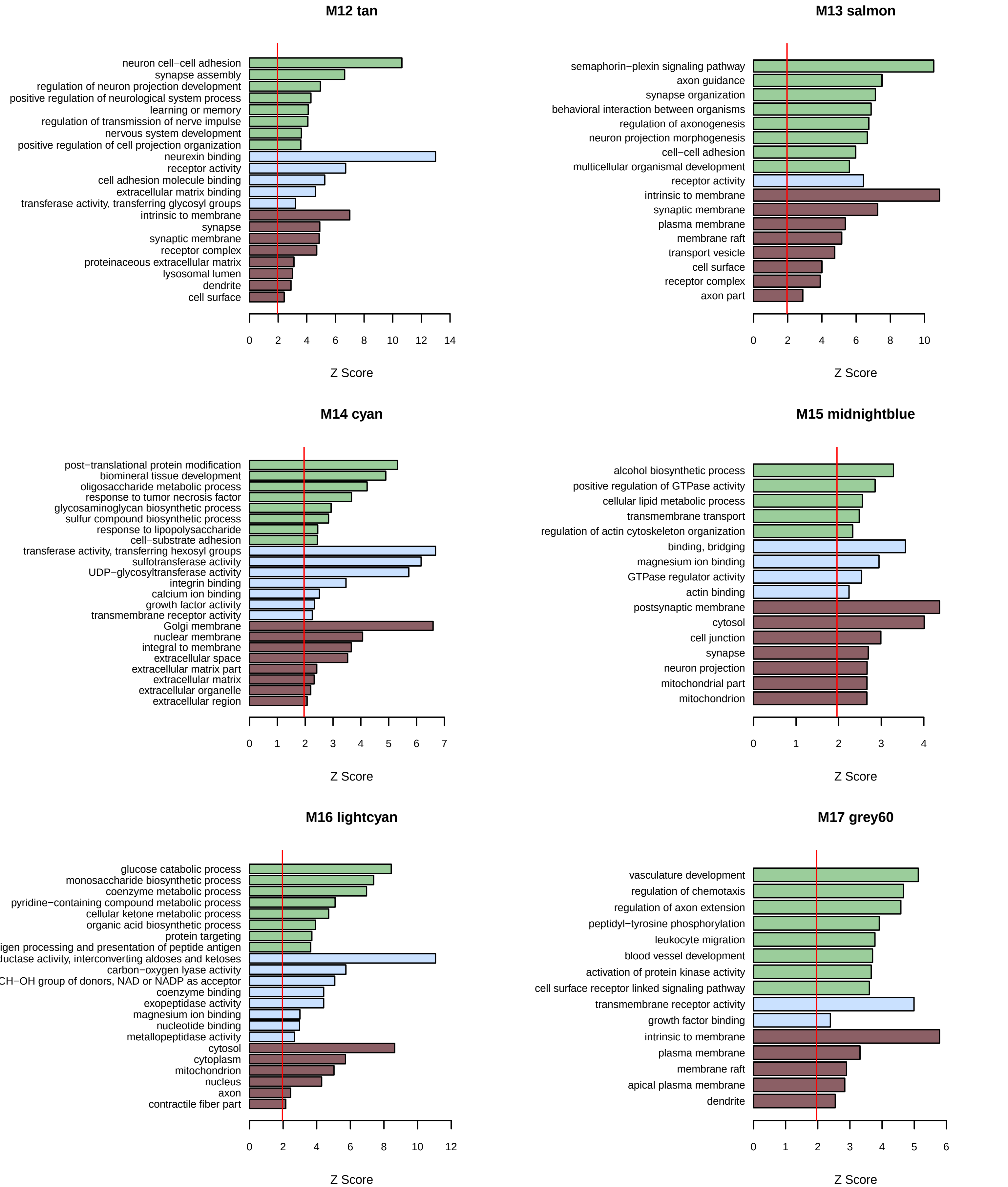
M12 tan
neuron cell−cell adhesion
synapse assembly
regulation of neuron projection development
positive regulation of neurological system process
learning or memory
regulation of transmission of nerve impulse
nervous system development
positive regulation of cell projection organization
neurexin binding
receptor activity
cell adhesion molecule binding
extracellular matrix binding
transferase activity, transferring glycosyl groups
intrinsic to membrane
synapse
synaptic membrane
receptor complex
proteinaceous extracellular matrix
lysosomal lumen
dendrite
cell surface
0 2 4 6 8 10 12 14
Z Score
M13 salmon
semaphorin−plexin signaling pathway
axon guidance
synapse organization
behavioral interaction between organisms
regulation of axonogenesis
neuron projection morphogenesis
cell−cell adhesion
multicellular organismal development
receptor activity
intrinsic to membrane
synaptic membrane
plasma membrane
membrane raft
transport vesicle
cell surface
receptor complex
axon part
0 2 4 6 8 10
Z Score
M14 cyan
post−translational protein modification
biomineral tissue development
oligosaccharide metabolic process
response to tumor necrosis factor
glycosaminoglycan biosynthetic process
sulfur compound biosynthetic process
response to lipopolysaccharide
cell−substrate adhesion
transferase activity, transferring hexosyl groups
sulfotransferase activity
UDP−glycosyltransferase activity
integrin binding
calcium ion binding
growth factor activity
transmembrane receptor activity
Golgi membrane
nuclear membrane
integral to membrane
extracellular space
extracellular matrix part
extracellular matrix
extracellular organelle
extracellular region
0 1 2 3 4 5 6 7
Z Score
M15 midnightblue
alcohol biosynthetic process
positive regulation of GTPase activity
cellular lipid metabolic process
transmembrane transport
regulation of actin cytoskeleton organization
binding, bridging
magnesium ion binding
GTPase regulator activity
actin binding
postsynaptic membrane
cytosol
cell junction
synapse
neuron projection
mitochondrial part
mitochondrion
0 1 2 3 4
Z Score
M16 lightcyan
glucose catabolic process
monosaccharide biosynthetic process
coenzyme metabolic process
pyridine−containing compound metabolic process
cellular ketone metabolic process
organic acid biosynthetic process
protein targeting
igen processing and presentation of peptide antigen
ductase activity, interconverting aldoses and ketoses
carbon−oxygen lyase activity
CH−OH group of donors, NAD or NADP as acceptor
coenzyme binding
exopeptidase activity
magnesium ion binding
nucleotide binding
metallopeptidase activity
cytosol
cytoplasm
mitochondrion
nucleus
axon
contractile fiber part
0 2 4 6 8 10 12
Z Score
M17 grey60
vasculature development
regulation of chemotaxis
regulation of axon extension
peptidyl−tyrosine phosphorylation
leukocyte migration
blood vessel development
activation of protein kinase activity
cell surface receptor linked signaling pathway
transmembrane receptor activity
growth factor binding
intrinsic to membrane
plasma membrane
membrane raft
apical plasma membrane
dendrite
0 1 2 3 4 5 6
Z Score

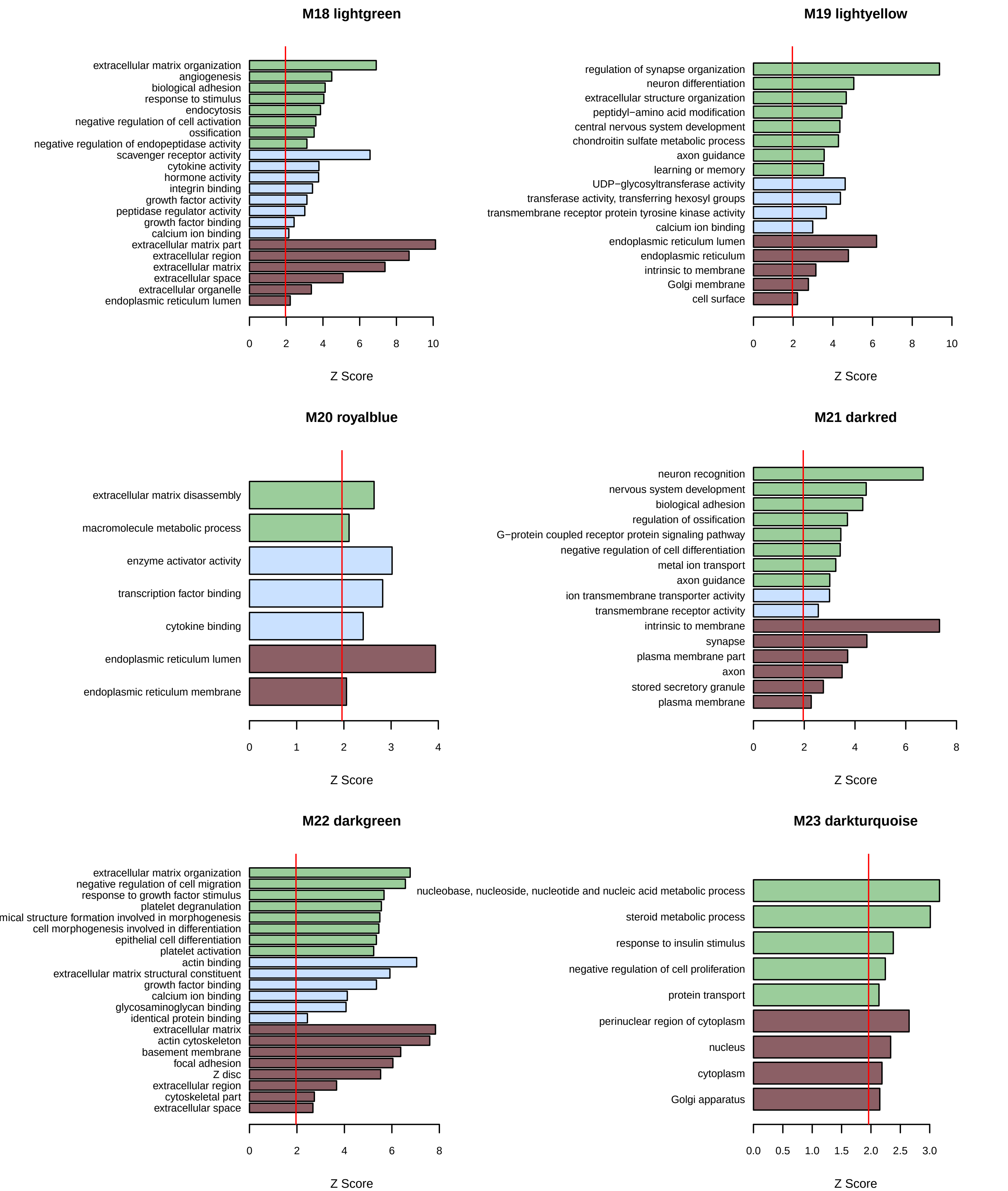
M18 lightgreen
extracellular matrix organization
angiogenesis
biological adhesion
response to stimulus
endocytosis
negative regulation of cell activation
ossification
negative regulation of endopeptidase activity
scavenger receptor activity
cytokine activity
hormone activity
integrin binding
growth factor activity
peptidase regulator activity
growth factor binding
calcium ion binding
extracellular matrix part
extracellular region
extracellular matrix
extracellular space
extracellular organelle
endoplasmic reticulum lumen
0 2 4 6 8 10
Z Score
M19 lightyellow
regulation of synapse organization
neuron differentiation
extracellular structure organization
peptidyl−amino acid modification
central nervous system development
chondroitin sulfate metabolic process
axon guidance
learning or memory
UDP−glycosyltransferase activity
transferase activity, transferring hexosyl groups
transmembrane receptor protein tyrosine kinase activity
calcium ion binding
endoplasmic reticulum lumen
endoplasmic reticulum
intrinsic to membrane
Golgi membrane
cell surface
0 2 4 6 8 10
Z Score
M20 royalblue
extracellular matrix disassembly
macromolecule metabolic process
enzyme activator activity
transcription factor binding
cytokine binding
endoplasmic reticulum lumen
endoplasmic reticulum membrane
0 1 2 3 4
Z Score
M21 darkred
neuron recognition
nervous system development
biological adhesion
regulation of ossification
G−protein coupled receptor protein signaling pathway
negative regulation of cell differentiation
metal ion transport
axon guidance
ion transmembrane transporter activity
transmembrane receptor activity
intrinsic to membrane
synapse
plasma membrane part
axon
stored secretory granule
plasma membrane
0 2 4 6 8
Z Score
M22 darkgreen
extracellular matrix organization
negative regulation of cell migration
response to growth factor stimulus
platelet degranulation
mical structure formation involved in morphogenesis
cell morphogenesis involved in differentiation
epithelial cell differentiation
platelet activation
actin binding
extracellular matrix structural constituent
growth factor binding
calcium ion binding
glycosaminoglycan binding
identical protein binding
extracellular matrix
actin cytoskeleton
basement membrane
focal adhesion
Z disc
extracellular region
cytoskeletal part
extracellular space
0 2 4 6 8
Z Score
M23 darkturquoise
nucleobase, nucleoside, nucleotide and nucleic acid metabolic process
steroid metabolic process
response to insulin stimulus
negative regulation of cell proliferation
protein transport
perinuclear region of cytoplasm
nucleus
cytoplasm
Golgi apparatus
0.0 0.5 1.0 1.5 2.0 2.5 3.0
Z Score

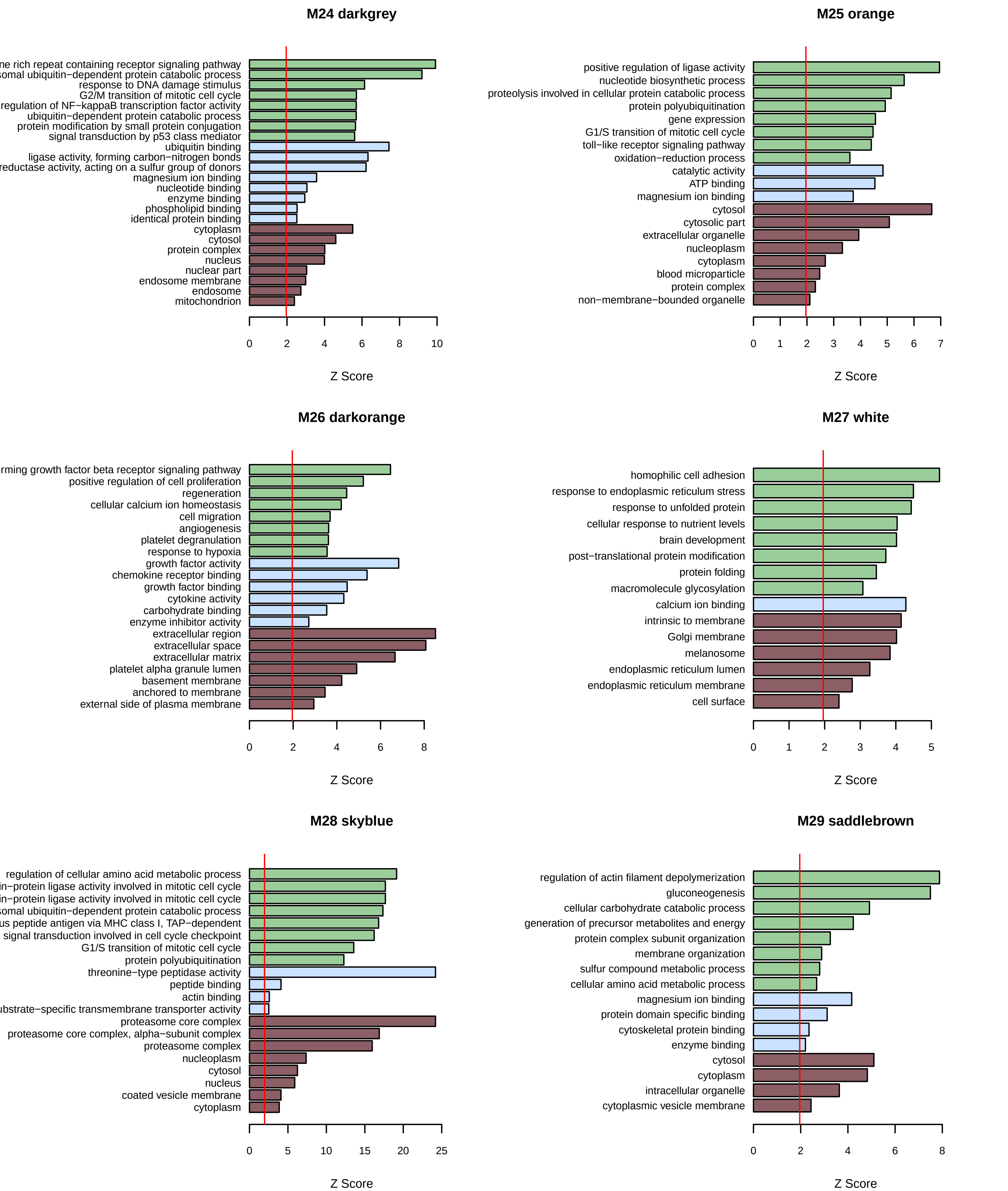
M24 darkgrey
ne rich repeat containing receptor signaling pathway
somal ubiquitin−dependent protein catabolic process
response to DNA damage stimulus
G2/M transition of mitotic cell cycle
regulation of NF−kappaB transcription factor activity
ubiquitin−dependent protein catabolic process
protein modification by small protein conjugation
signal transduction by p53 class mediator
ubiquitin binding
ligase activity, forming carbon−nitrogen bonds
reductase activity, acting on a sulfur group of donors
magnesium ion binding
nucleotide binding
enzyme binding
phospholipid binding
identical protein binding
cytoplasm
cytosol
protein complex
nucleus
nuclear part
endosome membrane
endosome
mitochondrion
0 2 4 6 8 10
Z Score
M25 orange
positive regulation of ligase activity
nucleotide biosynthetic process
proteolysis involved in cellular protein catabolic process
protein polyubiquitination
gene expression
G1/S transition of mitotic cell cycle
toll−like receptor signaling pathway
oxidation−reduction process
catalytic activity
ATP binding
magnesium ion binding
cytosol
cytosolic part
extracellular organelle
nucleoplasm
cytoplasm
blood microparticle
protein complex
non−membrane−bounded organelle
0 1 2 3 4 5 6 7
Z Score
M26 darkorange
rming growth factor beta receptor signaling pathway
positive regulation of cell proliferation
regeneration
cellular calcium ion homeostasis
cell migration
angiogenesis
platelet degranulation
response to hypoxia
growth factor activity
chemokine receptor binding
growth factor binding
cytokine activity
carbohydrate binding
enzyme inhibitor activity
extracellular region
extracellular space
extracellular matrix
platelet alpha granule lumen
basement membrane
anchored to membrane
external side of plasma membrane
0 2 4 6 8
Z Score
M27 white
homophilic cell adhesion
response to endoplasmic reticulum stress
response to unfolded protein
cellular response to nutrient levels
brain development
post−translational protein modification
protein folding
macromolecule glycosylation
calcium ion binding
intrinsic to membrane
Golgi membrane
melanosome
endoplasmic reticulum lumen
endoplasmic reticulum membrane
cell surface
0 1 2 3 4 5
Z Score
M28 skyblue
regulation of cellular amino acid metabolic process
in−protein ligase activity involved in mitotic cell cycle
in−protein ligase activity involved in mitotic cell cycle
somal ubiquitin−dependent protein catabolic process
us peptide antigen via MHC class I, TAP−dependent
signal transduction involved in cell cycle checkpoint
G1/S transition of mitotic cell cycle
protein polyubiquitination
threonine−type peptidase activity
peptide binding
actin binding
ubstrate−specific transmembrane transporter activity
proteasome core complex
proteasome core complex, alpha−subunit complex
proteasome complex
nucleoplasm
cytosol
nucleus
coated vesicle membrane
cytoplasm
0 5 10 15 20 25
Z Score
M29 saddlebrown
regulation of actin filament depolymerization
gluconeogenesis
cellular carbohydrate catabolic process
generation of precursor metabolites and energy
protein complex subunit organization
membrane organization
sulfur compound metabolic process
cellular amino acid metabolic process
magnesium ion binding
protein domain specific binding
cytoskeletal protein binding
enzyme binding
cytosol
cytoplasm
intracellular organelle
cytoplasmic vesicle membrane
0 2 4 6 8
Z Score

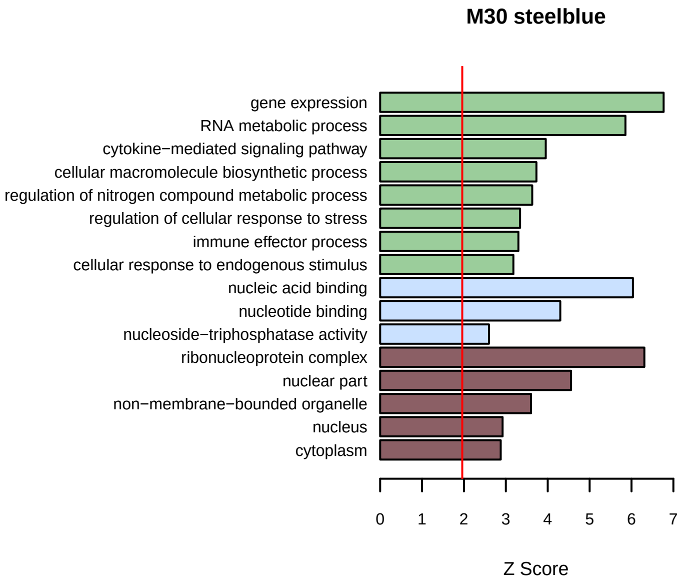
M30 steelblue
gene expression
RNA metabolic process
cytokine−mediated signaling pathway
cellular macromolecule biosynthetic process
regulation of nitrogen compound metabolic process
regulation of cellular response to stress
immune effector process
cellular response to endogenous stimulus
nucleic acid binding
nucleotide binding
nucleoside−triphosphatase activity
ribonucleoprotein complex
nuclear part
non−membrane−bounded organelle
nucleus
cytoplasm
0 1 2 3 4 5 6 7
Z Score

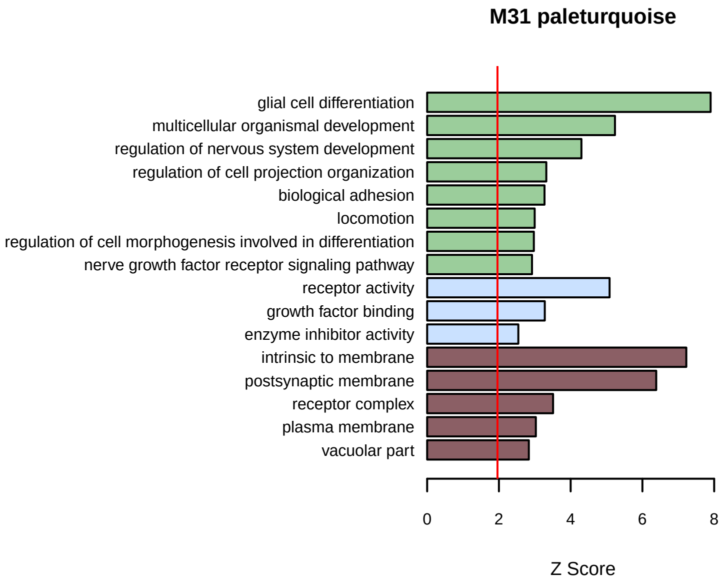
M31 paleturquoise
glial cell differentiation
multicellular organismal development
regulation of nervous system development
regulation of cell projection organization
biological adhesion
locomotion
regulation of cell morphogenesis involved in differentiation
nerve growth factor receptor signaling pathway
receptor activity
growth factor binding
enzyme inhibitor activity
intrinsic to membrane
postsynaptic membrane
receptor complex
plasma membrane
vacuolar part
0 2 4 6 8
Z Score

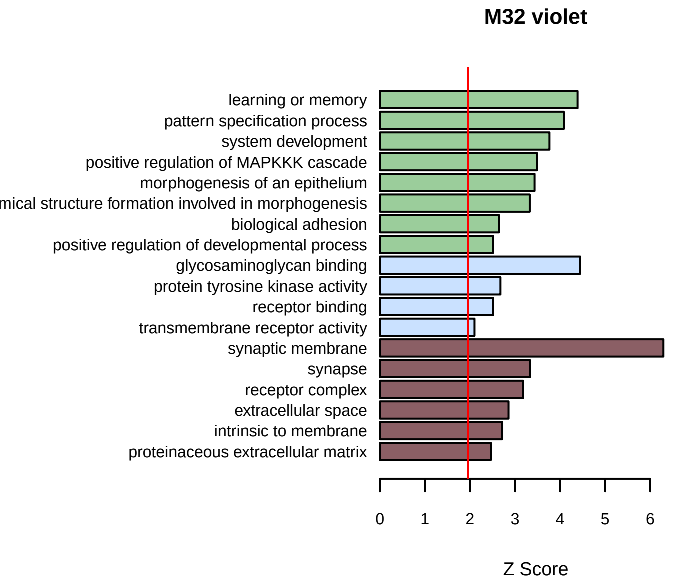
M32 violet
learning or memory
pattern specification process
system development
positive regulation of MAPKKK cascade
morphogenesis of an epithelium
mical structure formation involved in morphogenesis
biological adhesion
positive regulation of developmental process
glycosaminoglycan binding
protein tyrosine kinase activity
receptor binding
transmembrane receptor activity
synaptic membrane
synapse
receptor complex
extracellular space
intrinsic to membrane
proteinaceous extracellular matrix
0 1 2 3 4 5 6
Z Score

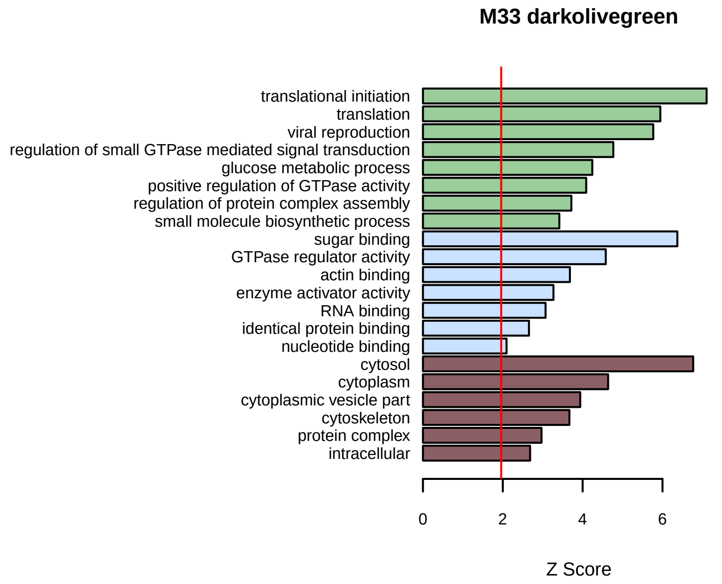
M33 darkolivegreen
translational initiation
translation
viral reproduction
regulation of small GTPase mediated signal transduction
glucose metabolic process
positive regulation of GTPase activity
regulation of protein complex assembly
small molecule biosynthetic process
sugar binding
GTPase regulator activity
actin binding
enzyme activator activity
RNA binding
identical protein binding
nucleotide binding
cytosol
cytoplasm
cytoplasmic vesicle part
cytoskeleton
protein complex
intracellular
0 2 4 6
Z Score

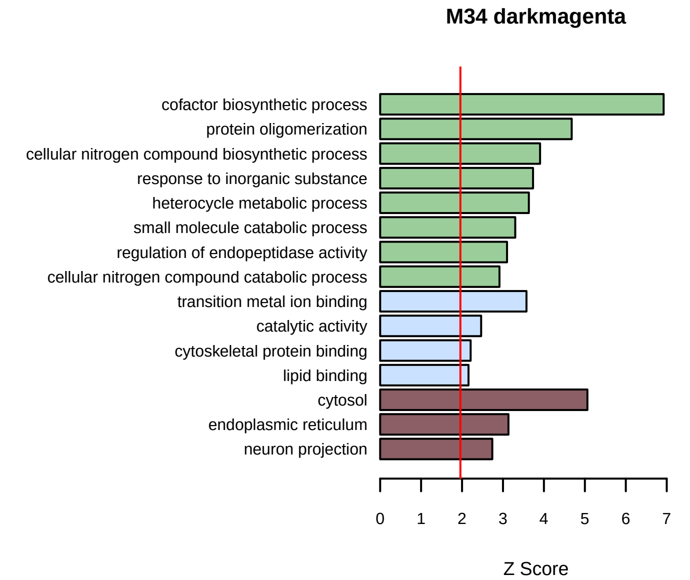
M34 darkmagenta
cofactor biosynthetic process
protein oligomerization
cellular nitrogen compound biosynthetic process
response to inorganic substance
heterocycle metabolic process
small molecule catabolic process
regulation of endopeptidase activity
cellular nitrogen compound catabolic process
transition metal ion binding
catalytic activity
cytoskeletal protein binding
lipid binding
cytosol
endoplasmic reticulum
neuron projection
0 1 2 3 4 5 6 7
Z Score

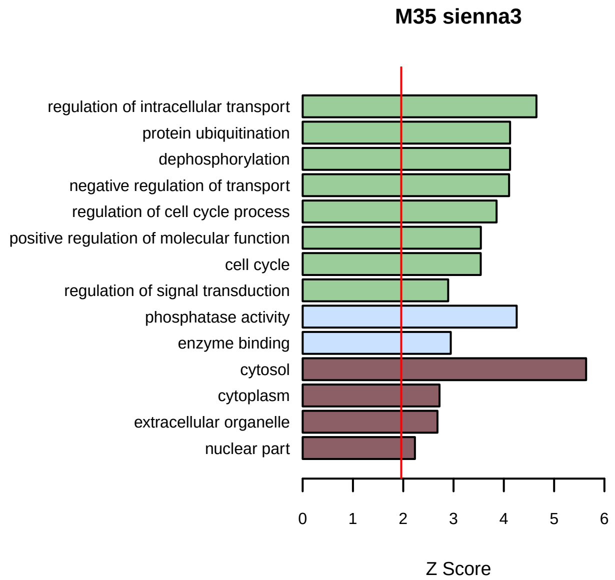
M35 sienna3
regulation of intracellular transport
protein ubiquitination
dephosphorylation
negative regulation of transport
regulation of cell cycle process
positive regulation of molecular function
cell cycle
regulation of signal transduction
phosphatase activity
enzyme binding
cytosol
cytoplasm
extracellular organelle
nuclear part
0 1 2 3 4 5 6
Z Score

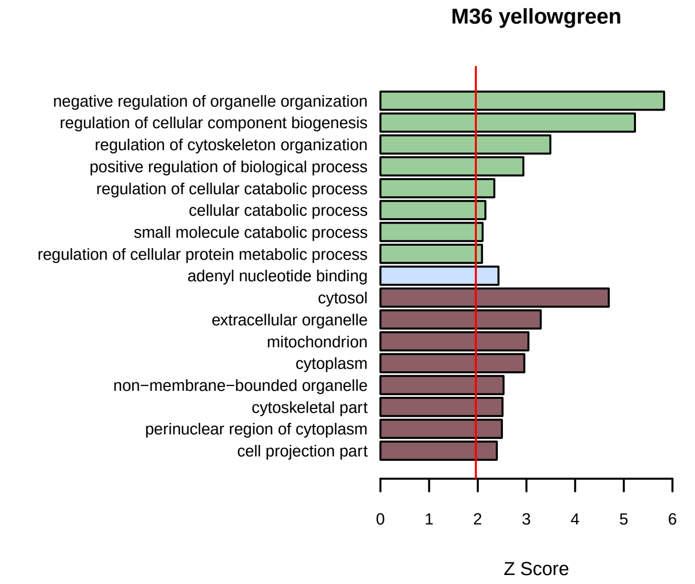
M36 yellowgreen
negative regulation of organelle organization
regulation of cellular component biogenesis
regulation of cytoskeleton organization
positive regulation of biological process
regulation of cellular catabolic process
cellular catabolic process
small molecule catabolic process
regulation of cellular protein metabolic process
adenyl nucleotide binding
cytosol
extracellular organelle
mitochondrion
cytoplasm
non−membrane−bounded organelle
cytoskeletal part
perinuclear region of cytoplasm
cell projection part
0 1 2 3 4 5 6
Z Score

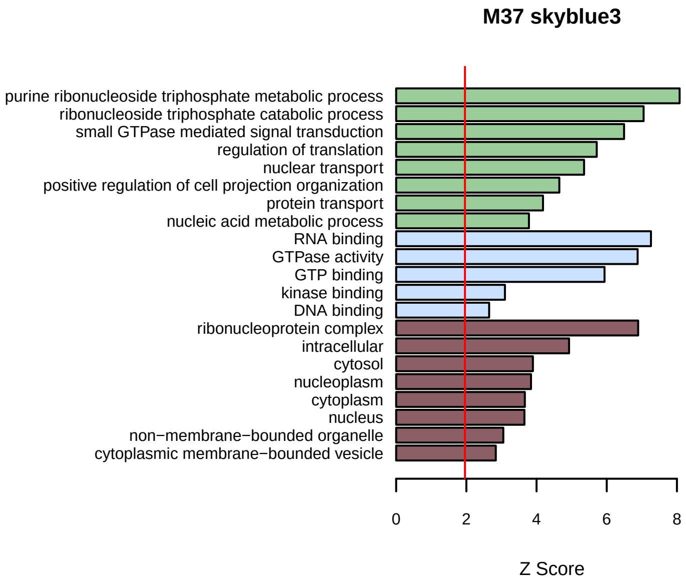
M37 skyblue3
purine ribonucleoside triphosphate metabolic process
ribonucleoside triphosphate catabolic process
small GTPase mediated signal transduction
regulation of translation
nuclear transport
positive regulation of cell projection organization
protein transport
nucleic acid metabolic process
RNA binding
GTPase activity
GTP binding
kinase binding
DNA binding
ribonucleoprotein complex
intracellular
cytosol
nucleoplasm
cytoplasm
nucleus
non−membrane−bounded organelle
cytoplasmic membrane−bounded vesicle
0 2 4 6 8
Z Score

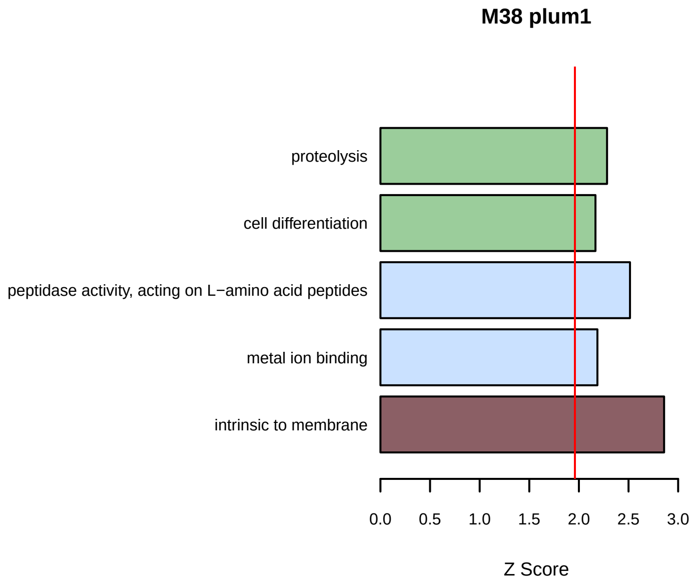
M38 plum1
proteolysis
cell differentiation
peptidase activity, acting on L−amino acid peptides
metal ion binding
intrinsic to membrane
0.0 0.5 1.0 1.5 2.0 2.5 3.0
Z Score

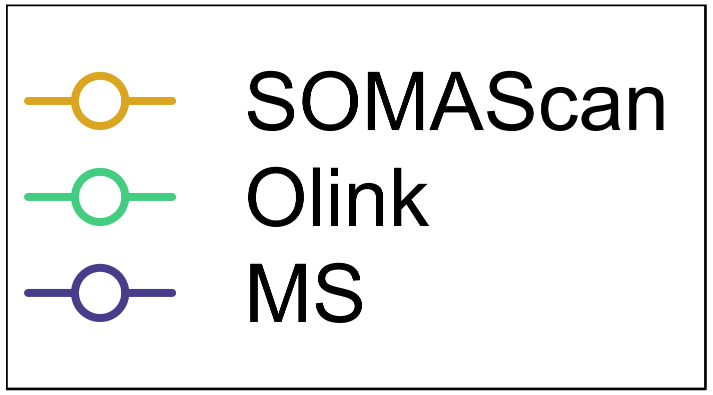
SOMAScan
Olink
MS

**M11 greenyellow module: Ambiguous**

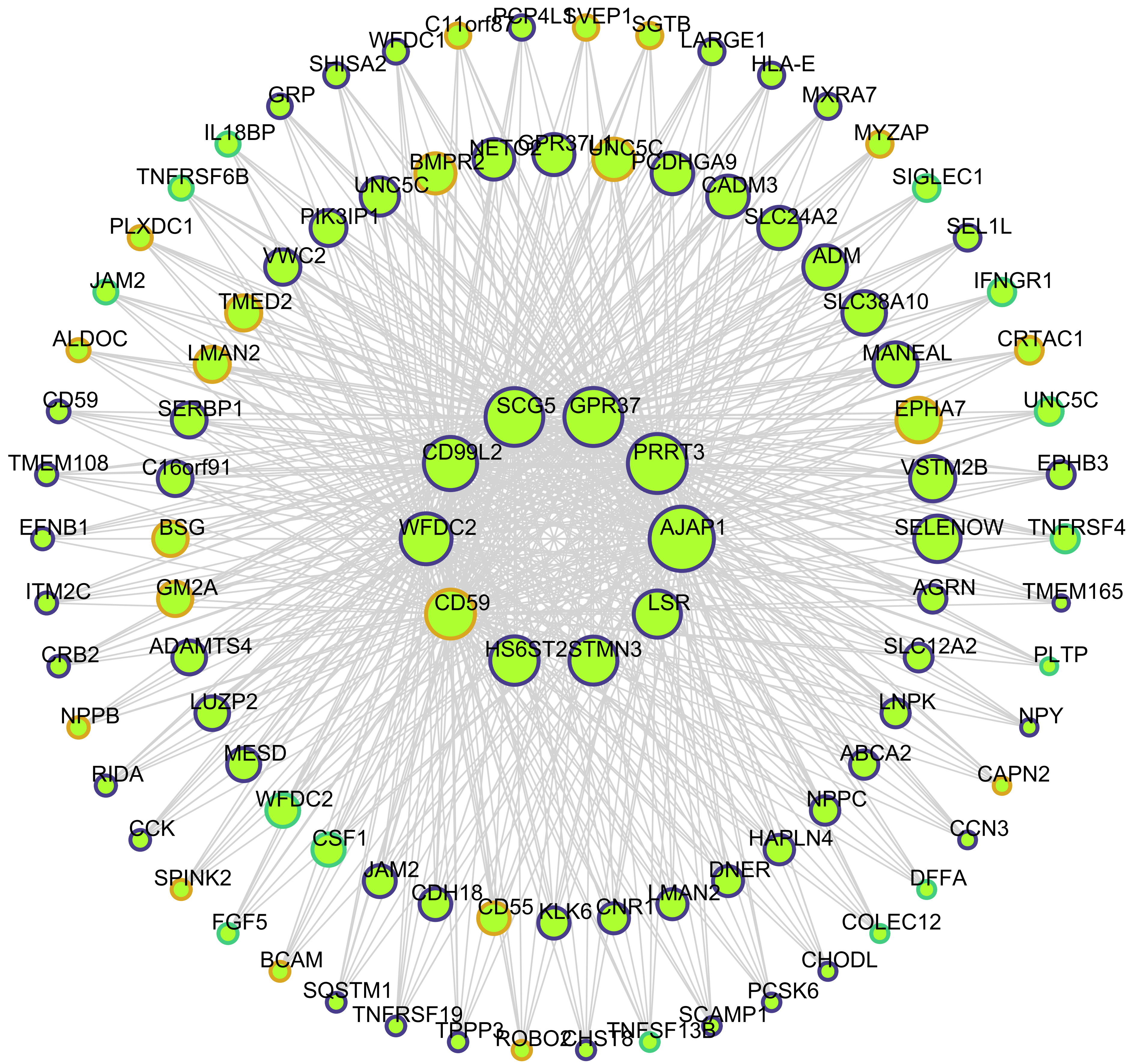

**M21 darkred module: Neuron Recognition**

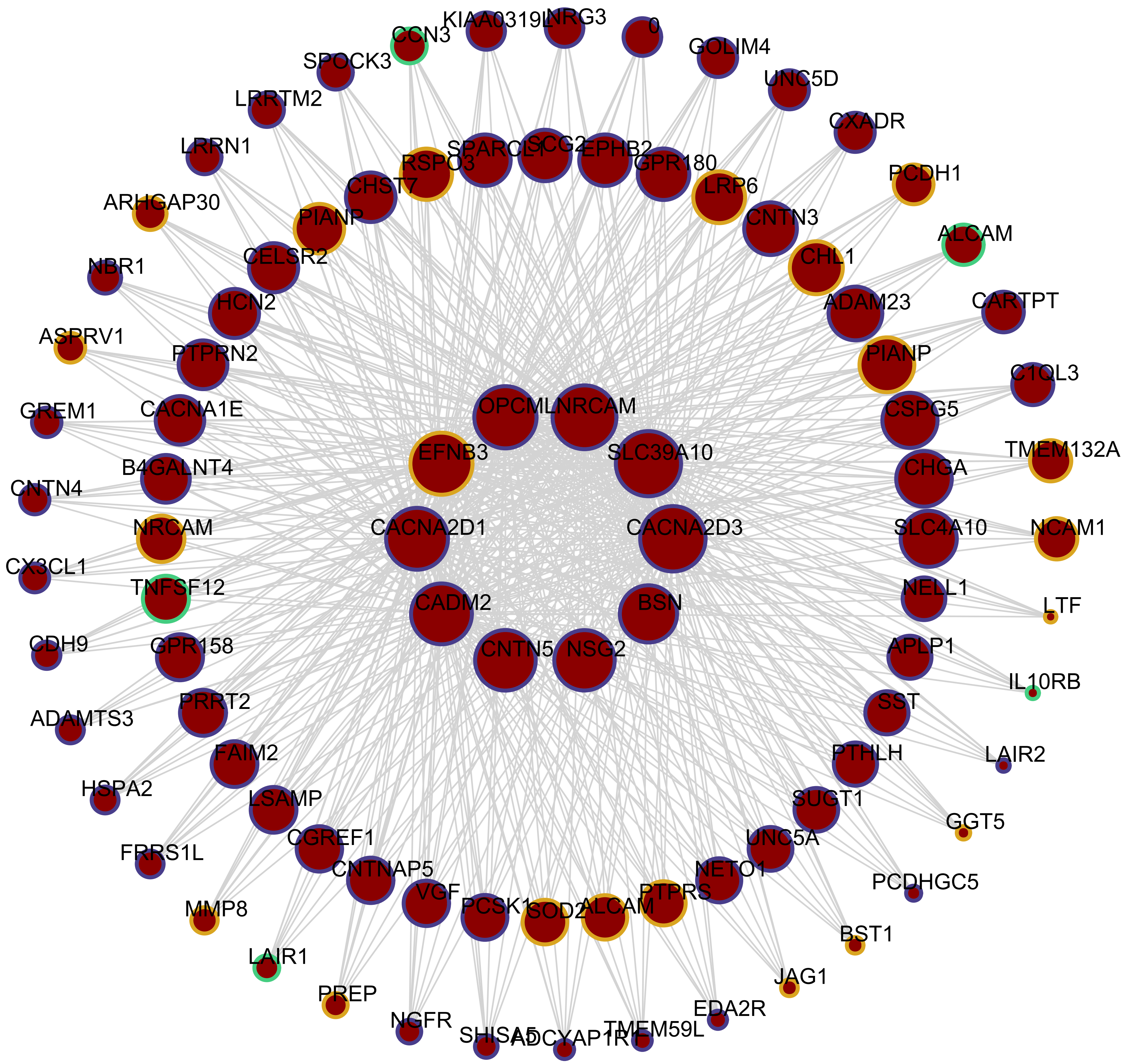

**M17 grey60 module: Ambiguous**

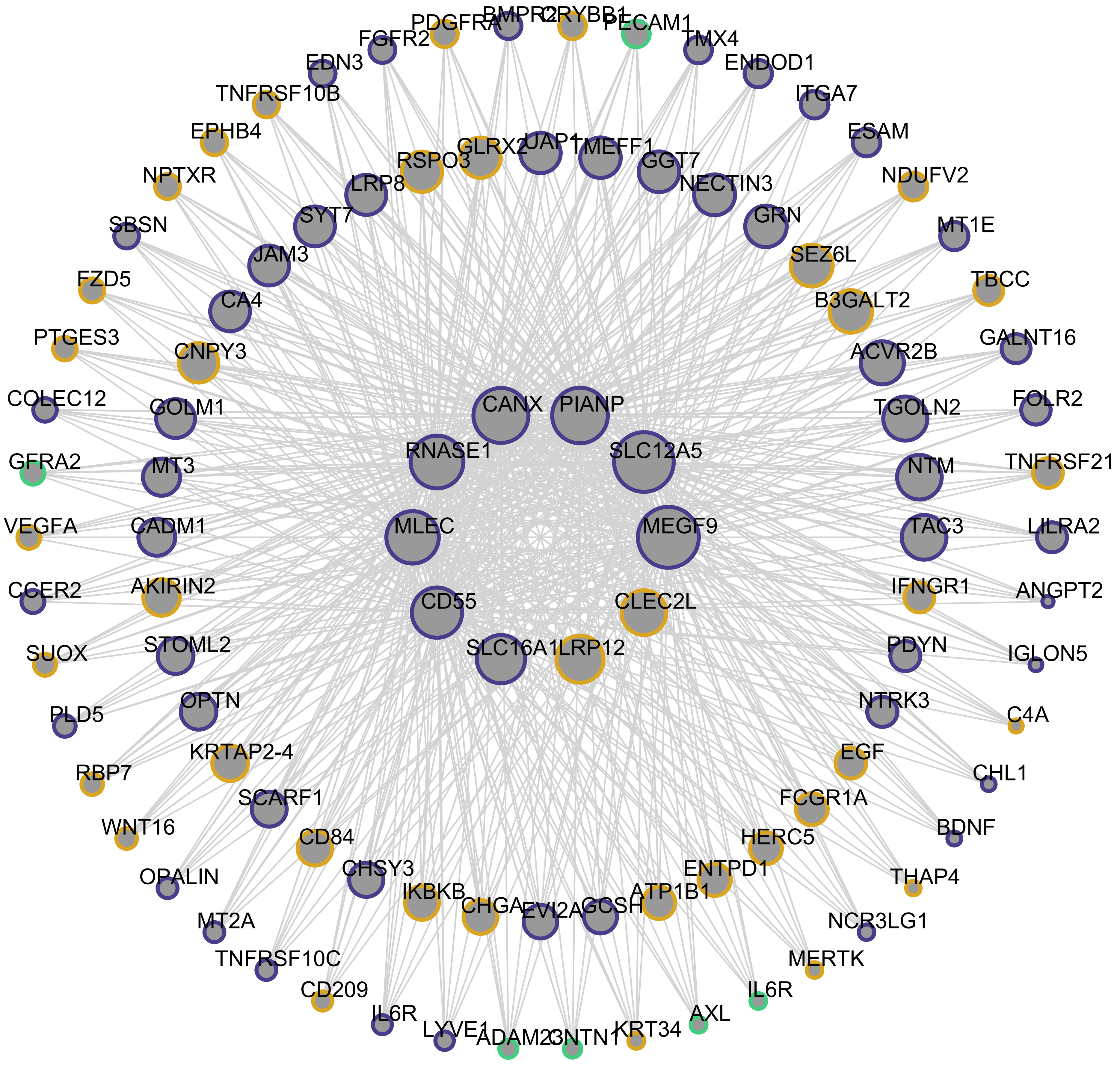

**M6 red module: Muscle/Neurotransmitter Transport**

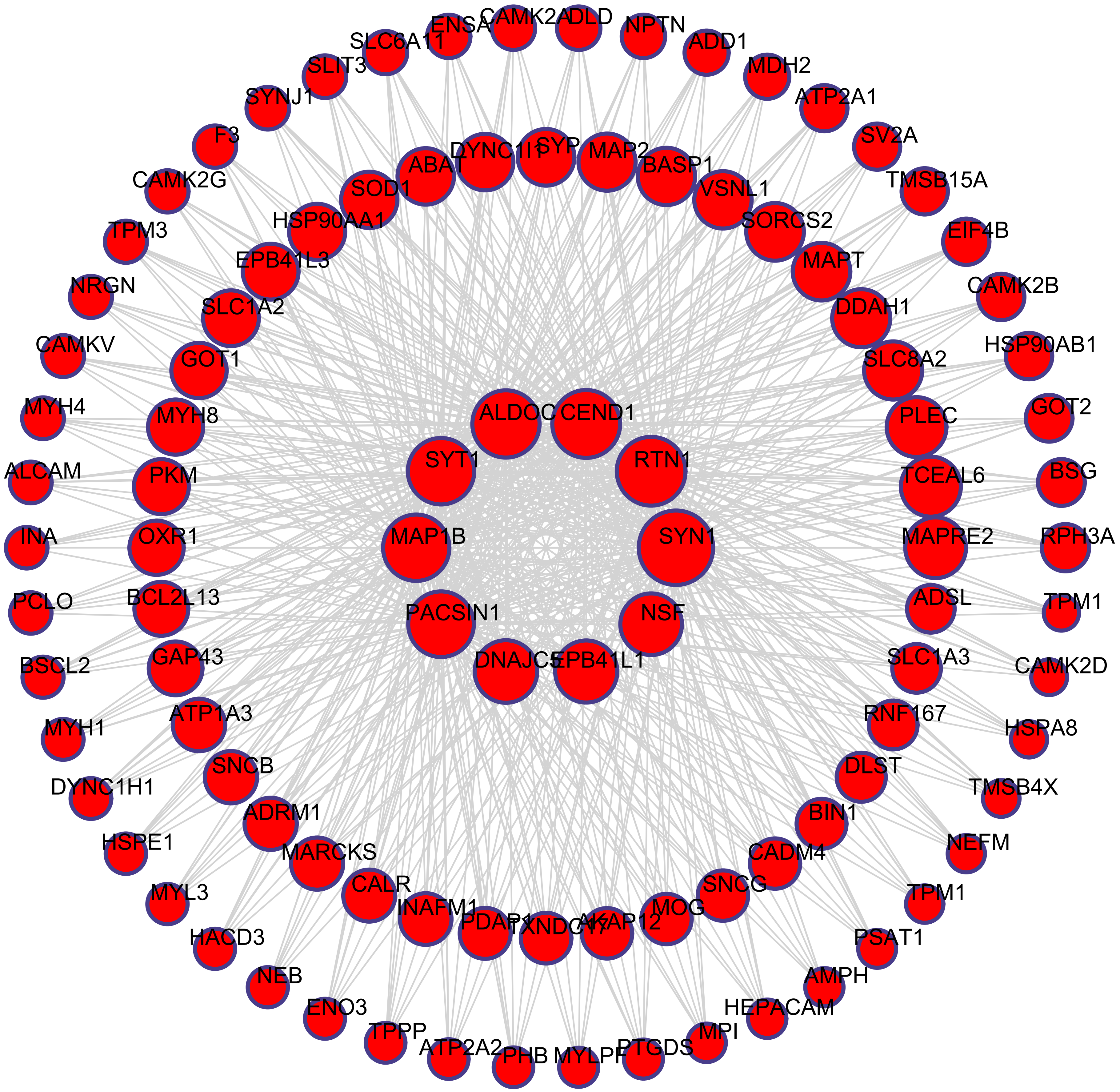

**M15 midnightblue module: Post-Synaptic Membrane**

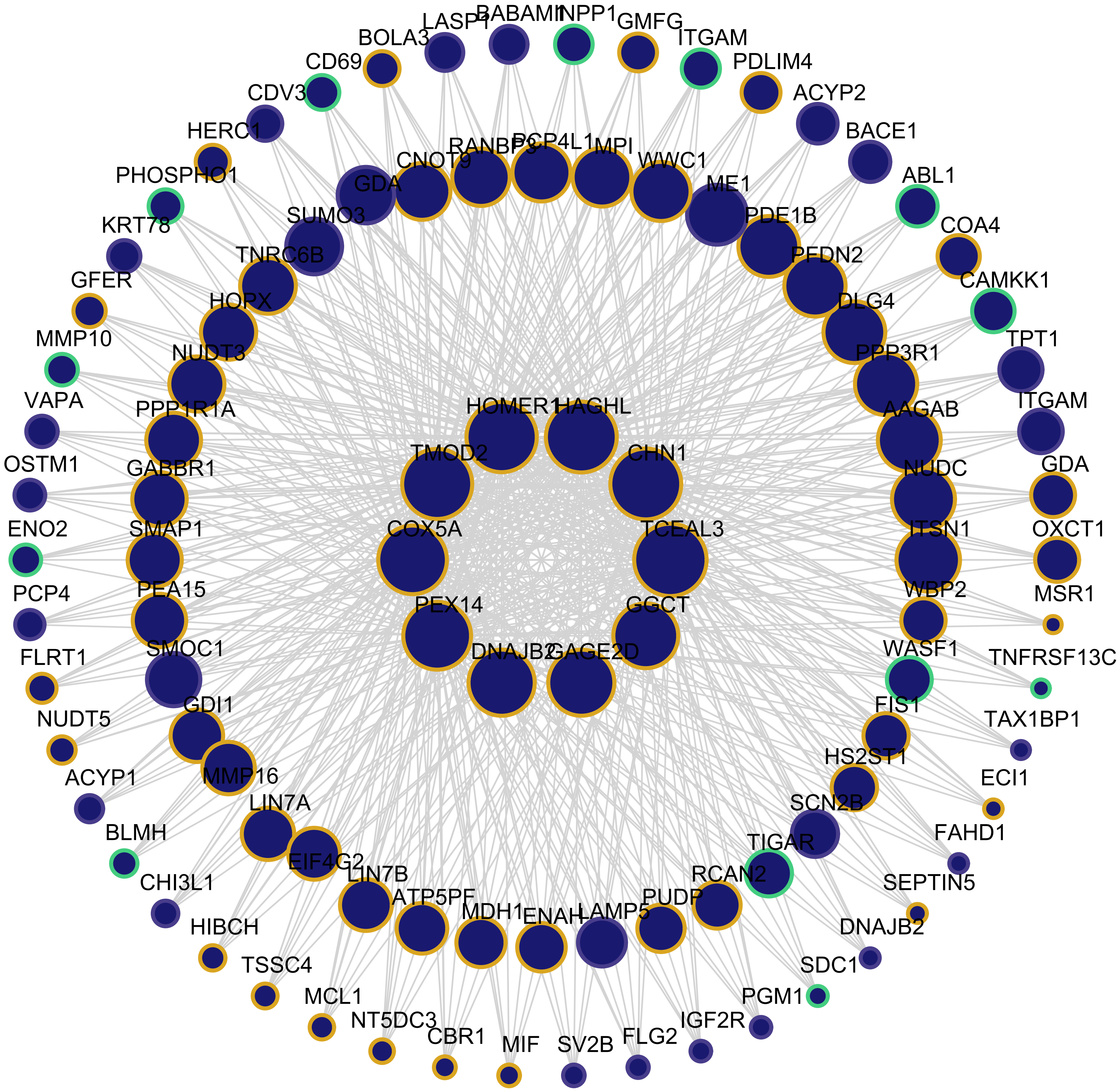

**M8 pink module: Autophagy**

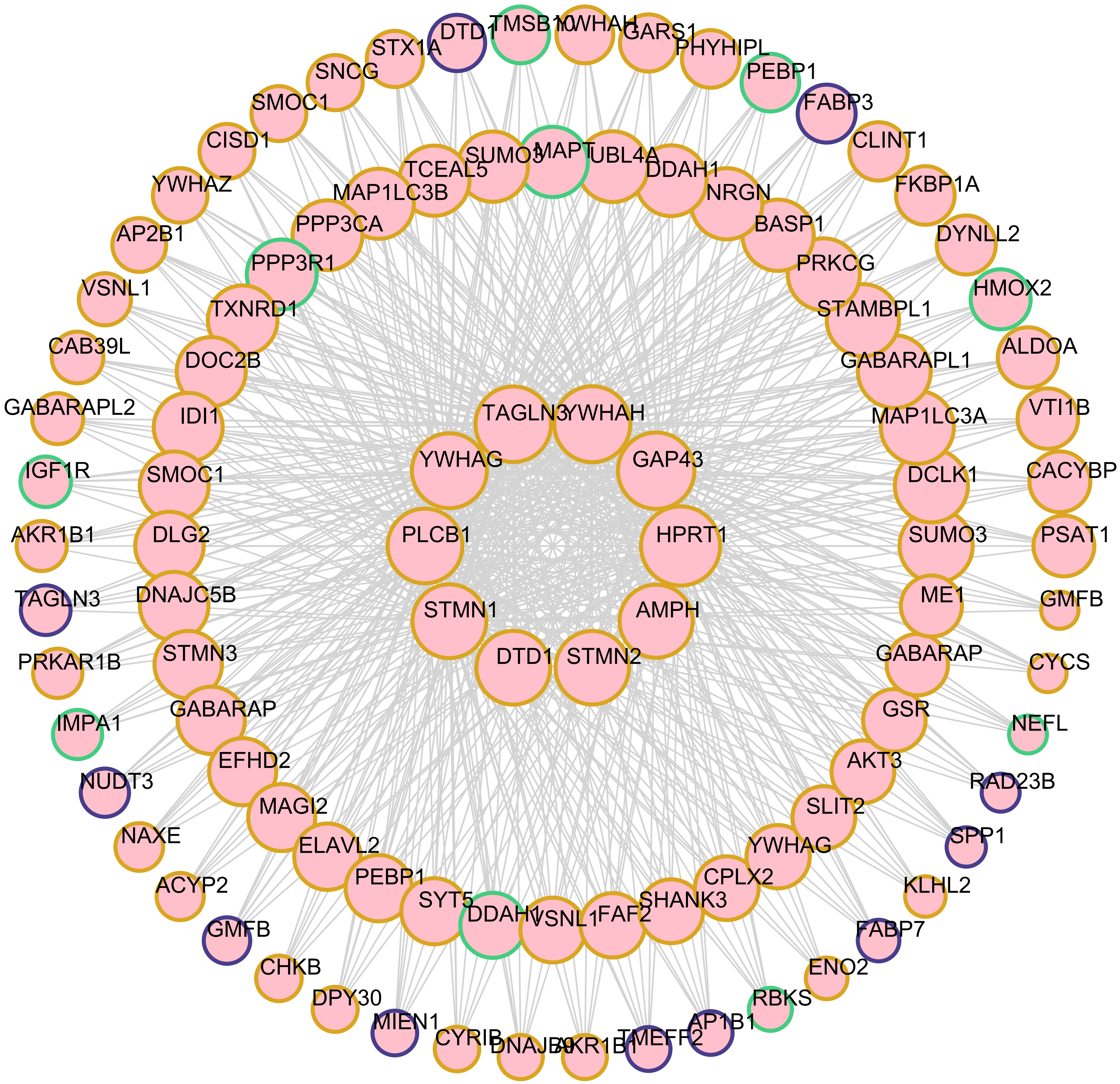

**M19 lightyellow module: Synapse Organization**

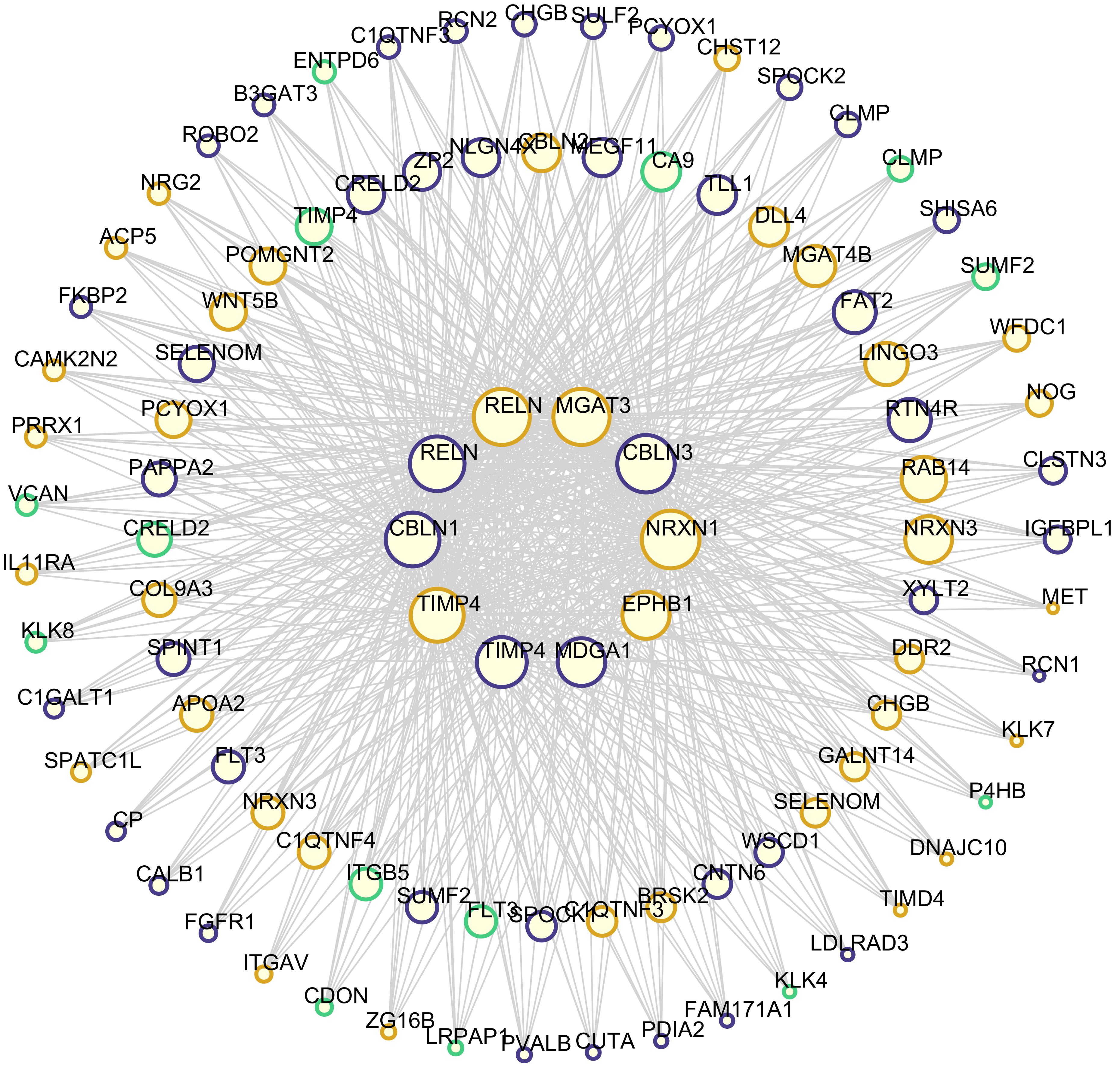

**M38 plum1 module: Ambiguous**

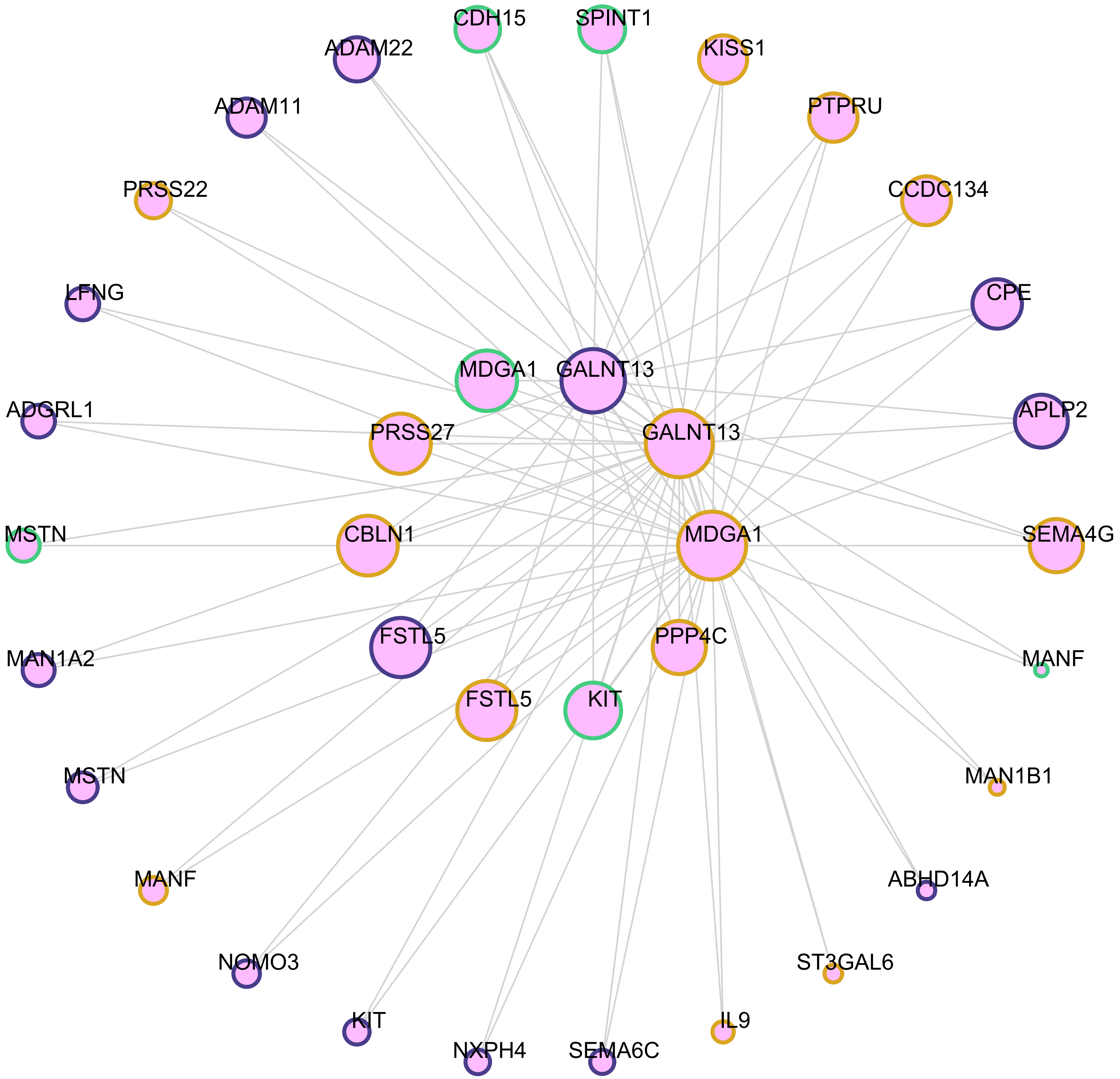

**M7 black module: SNAP Receptor/SNARE Complex**

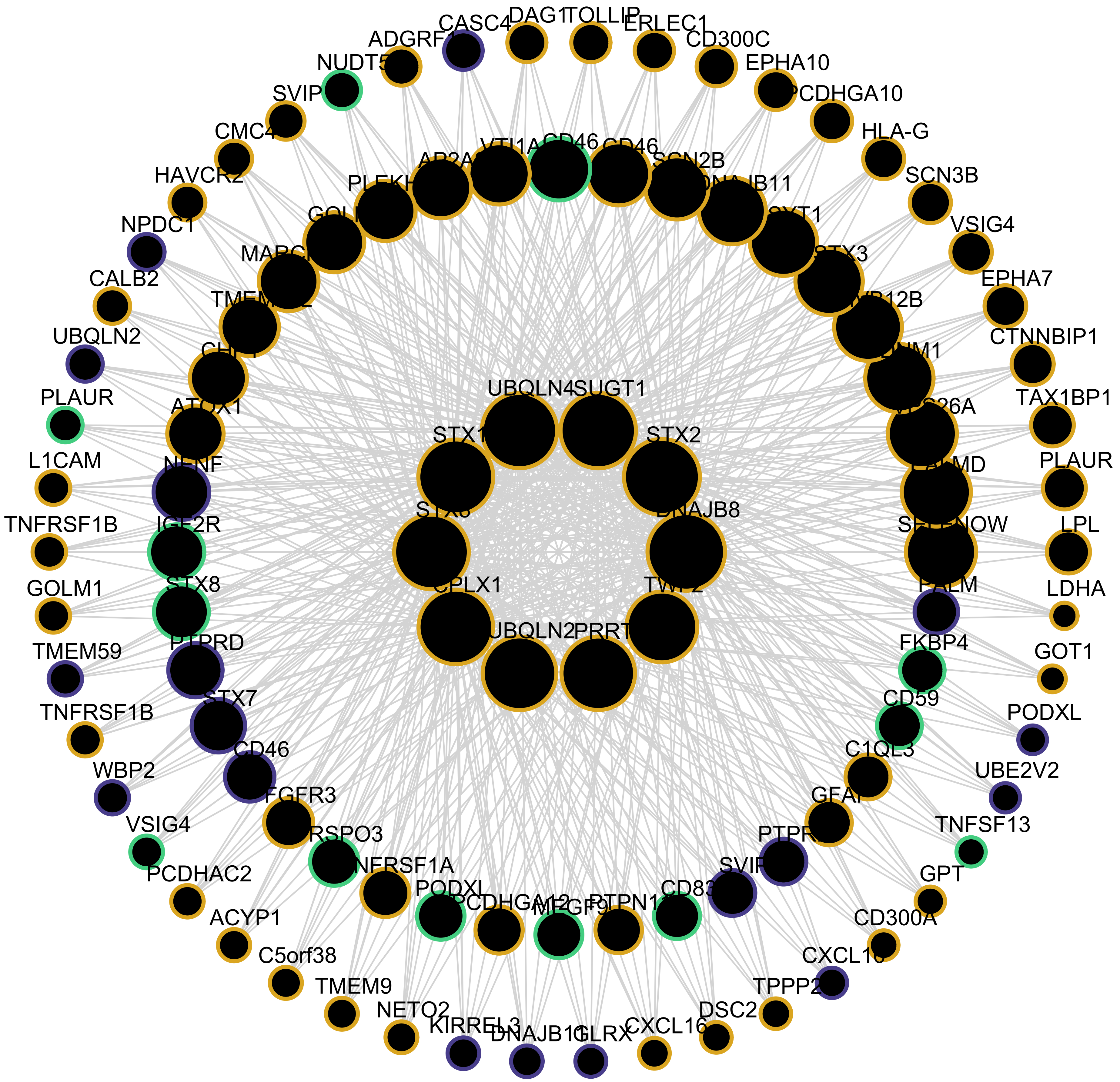

**M2 blue module: Neuronal/Axon Development**

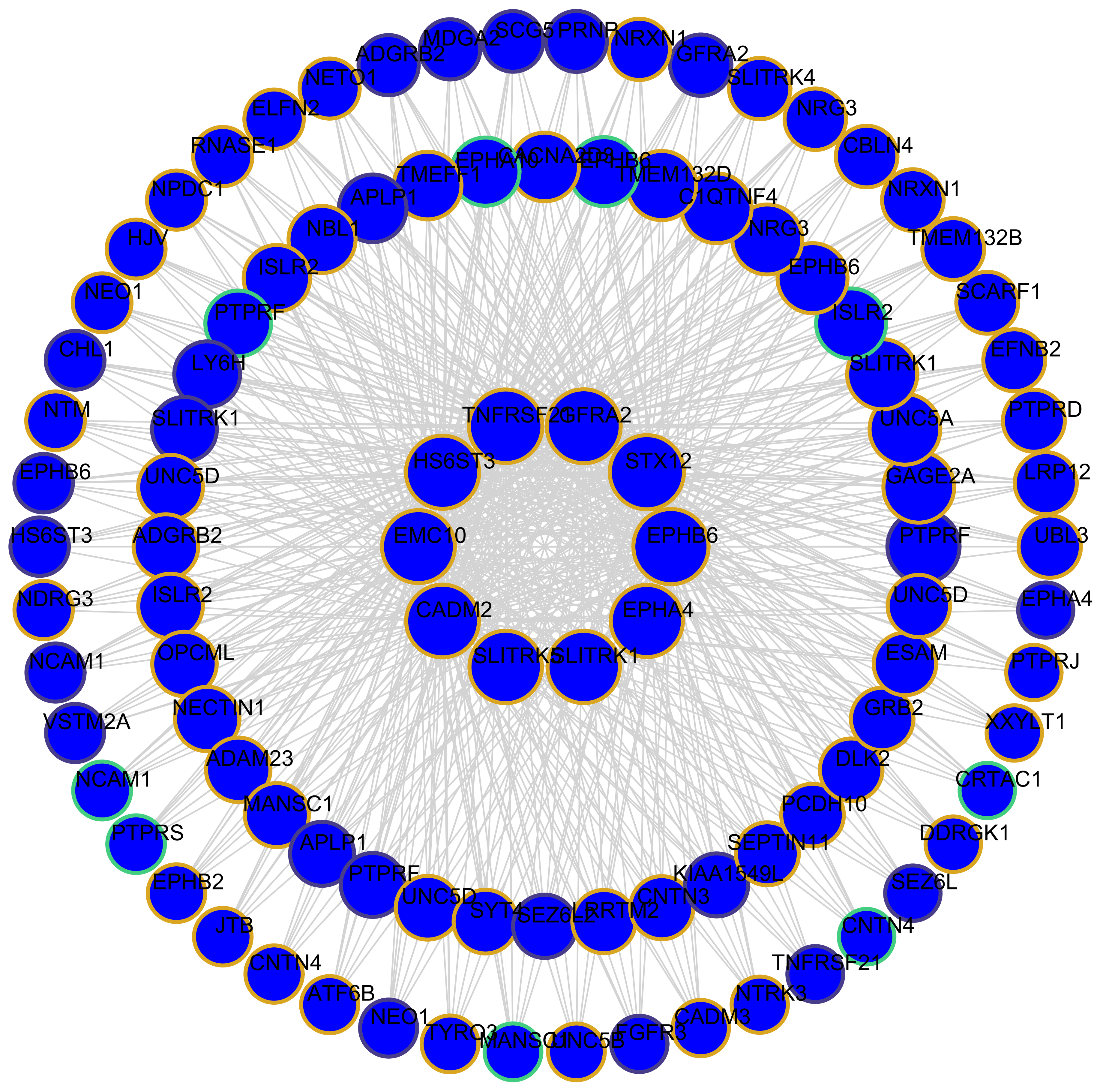

**M14 cyan module: Golgi/Glycosylation**

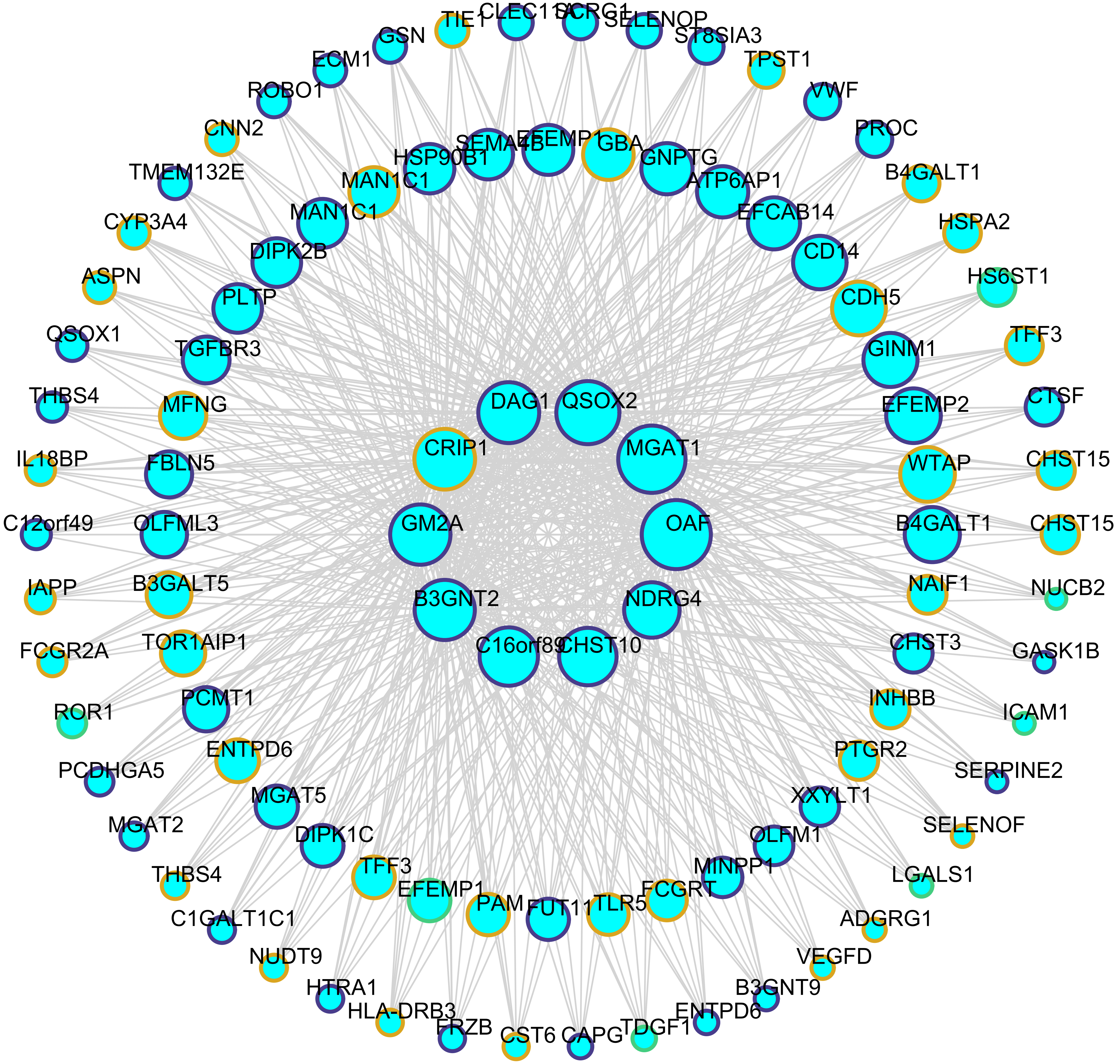

**M27 white module: Ambiguous**

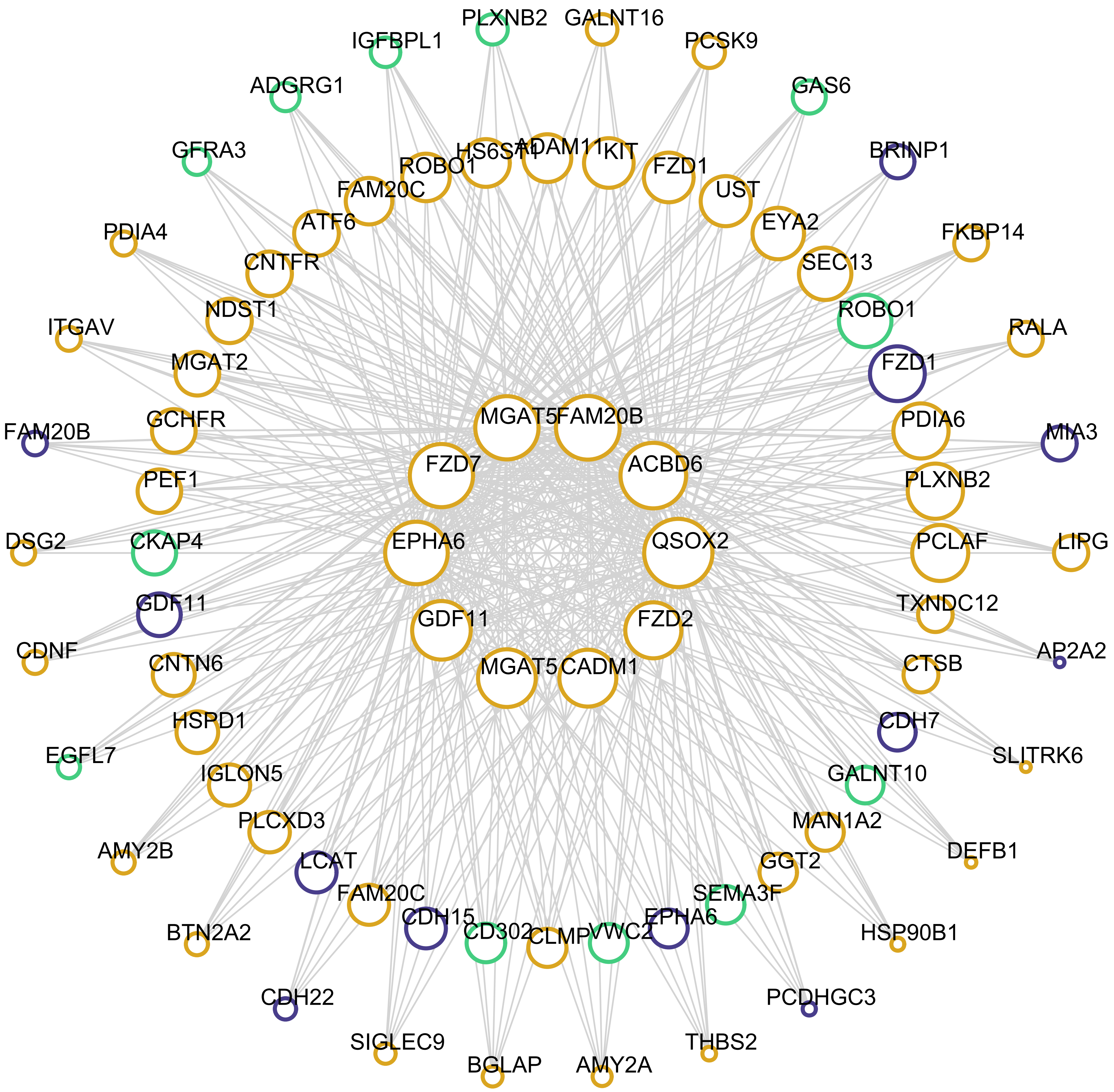

**M31 paleturquoise module: Glial Cell Differentiation**

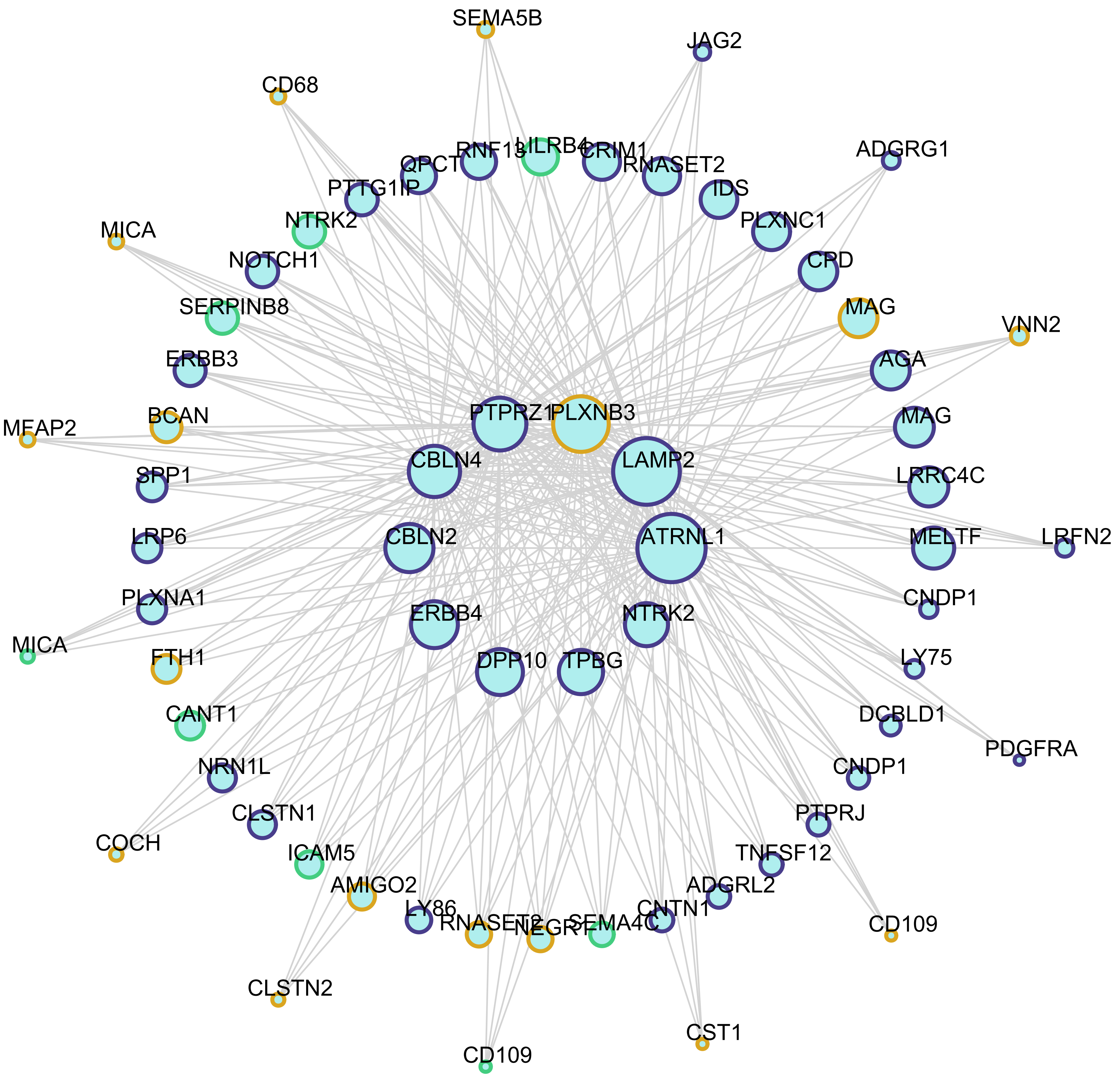

**M32 violet module: Synaptic Membrane/Matrisome**

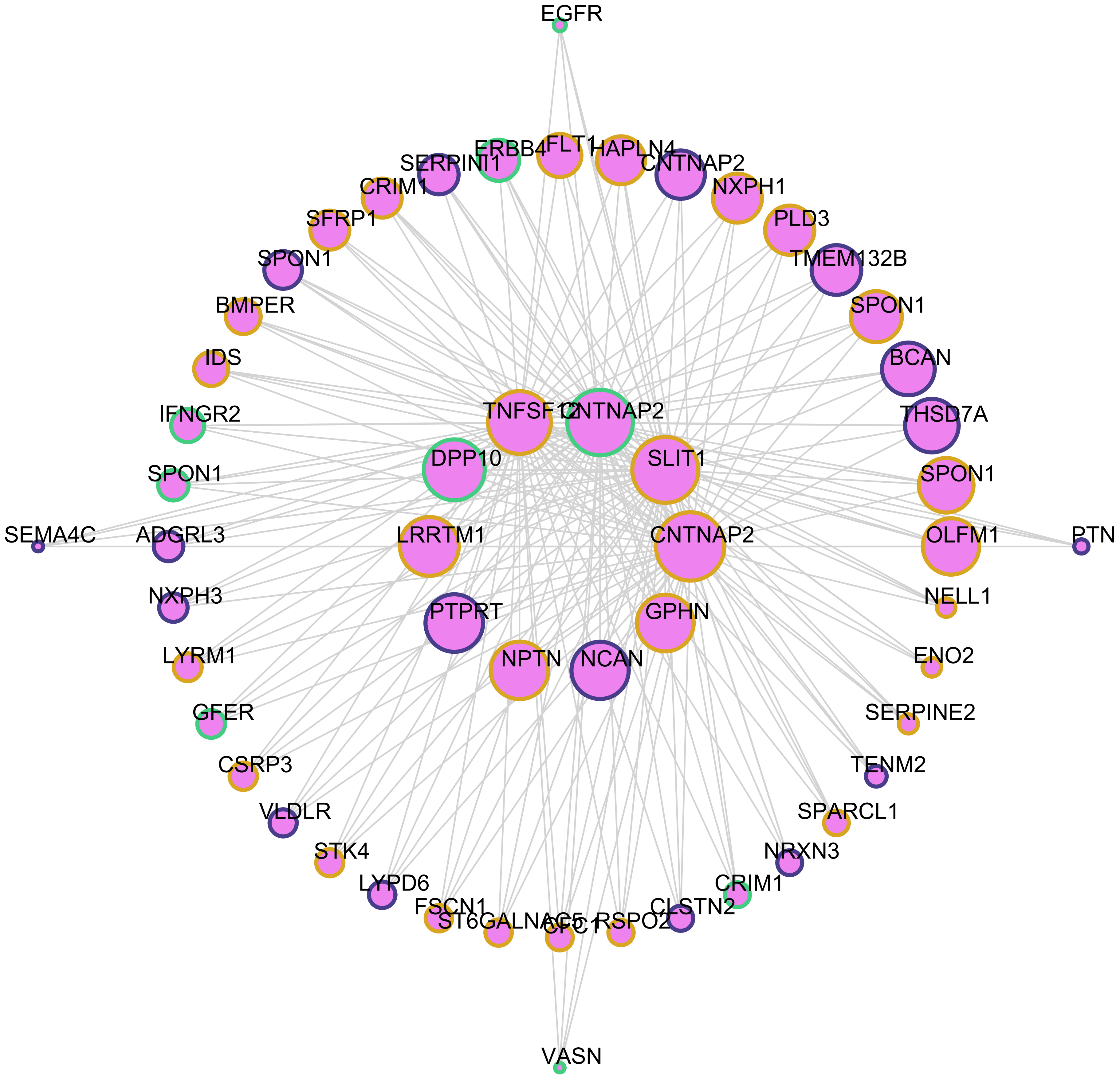

**M13 salmon module: Semaphorin-Plexin/Axon Guidance**

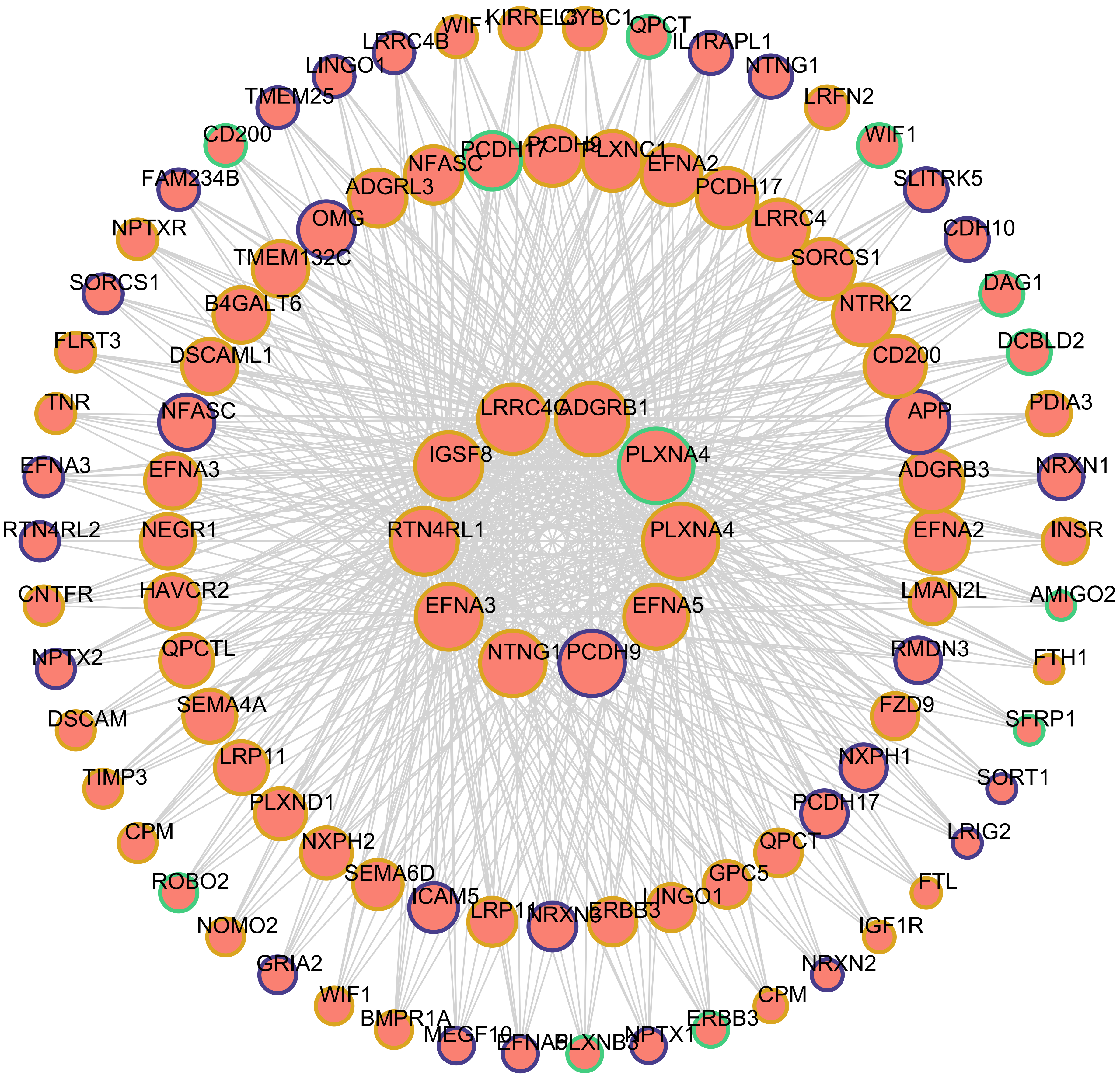

**M12 tan module: Neurexin/Neuronal Adhesion**

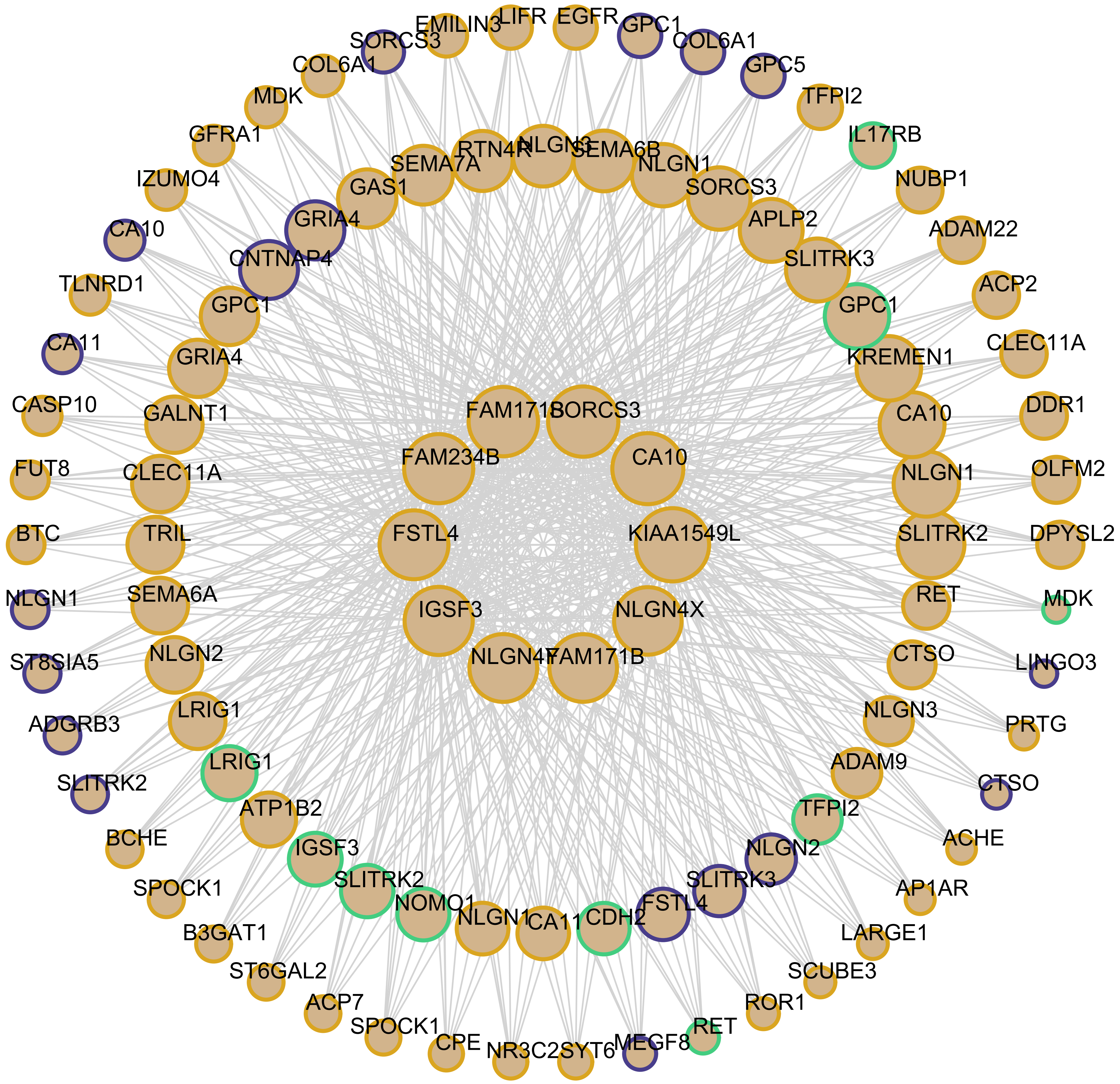

**M9 magenta module: Ambiguous**

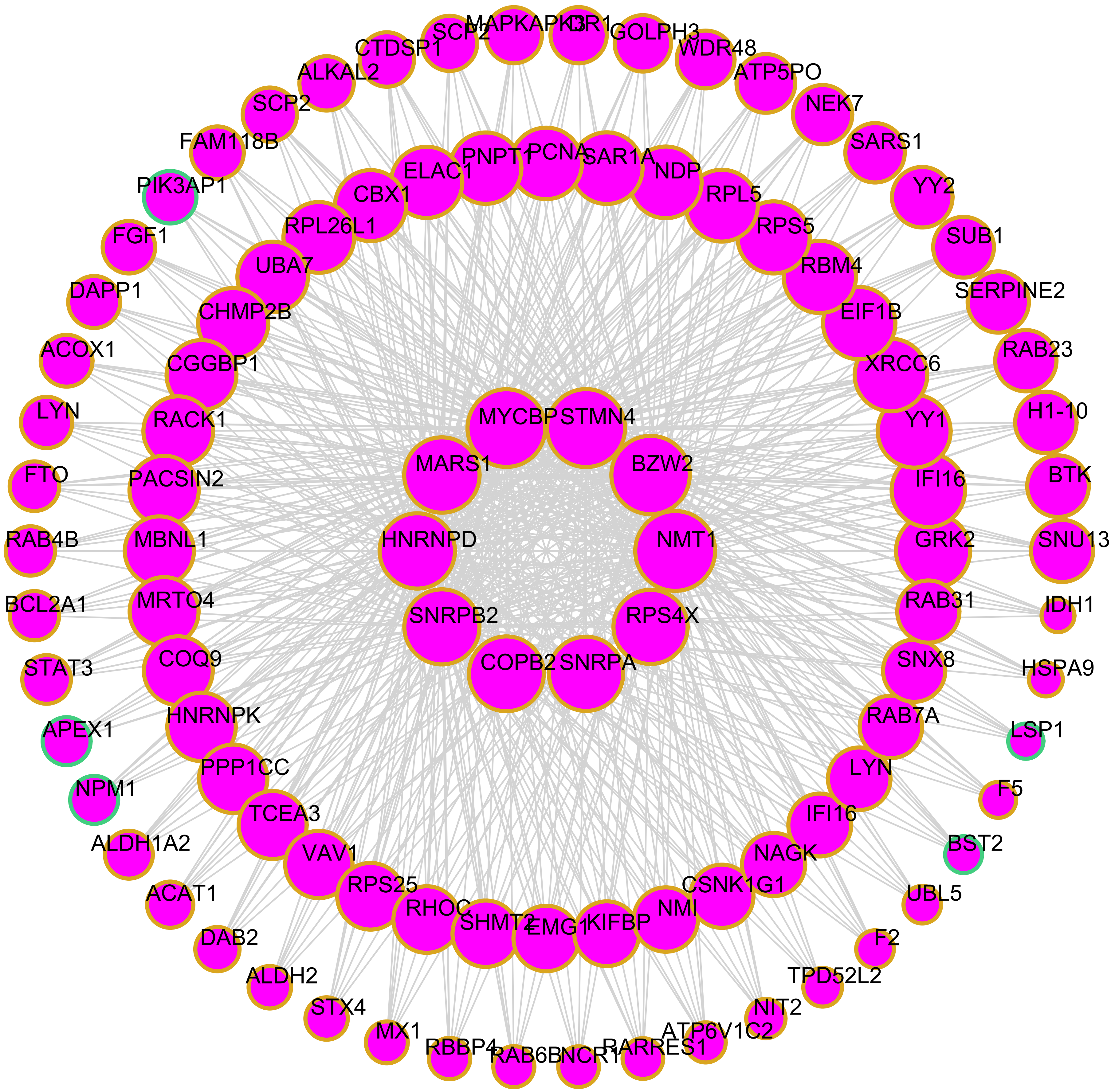

**M30 steelblue module: Ribonucleoprotein Complex**

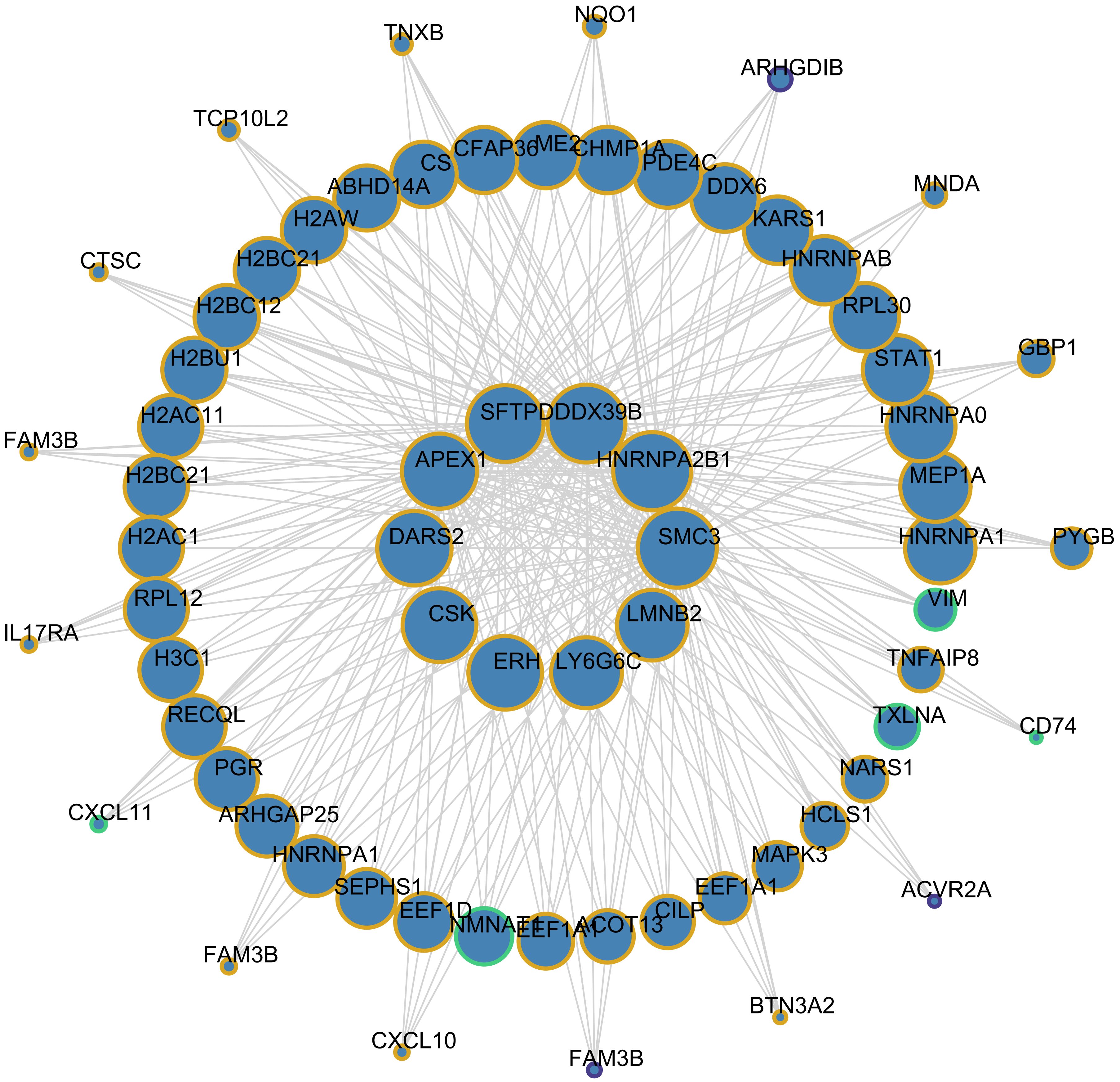

**M25 orange module: Ambiguous**

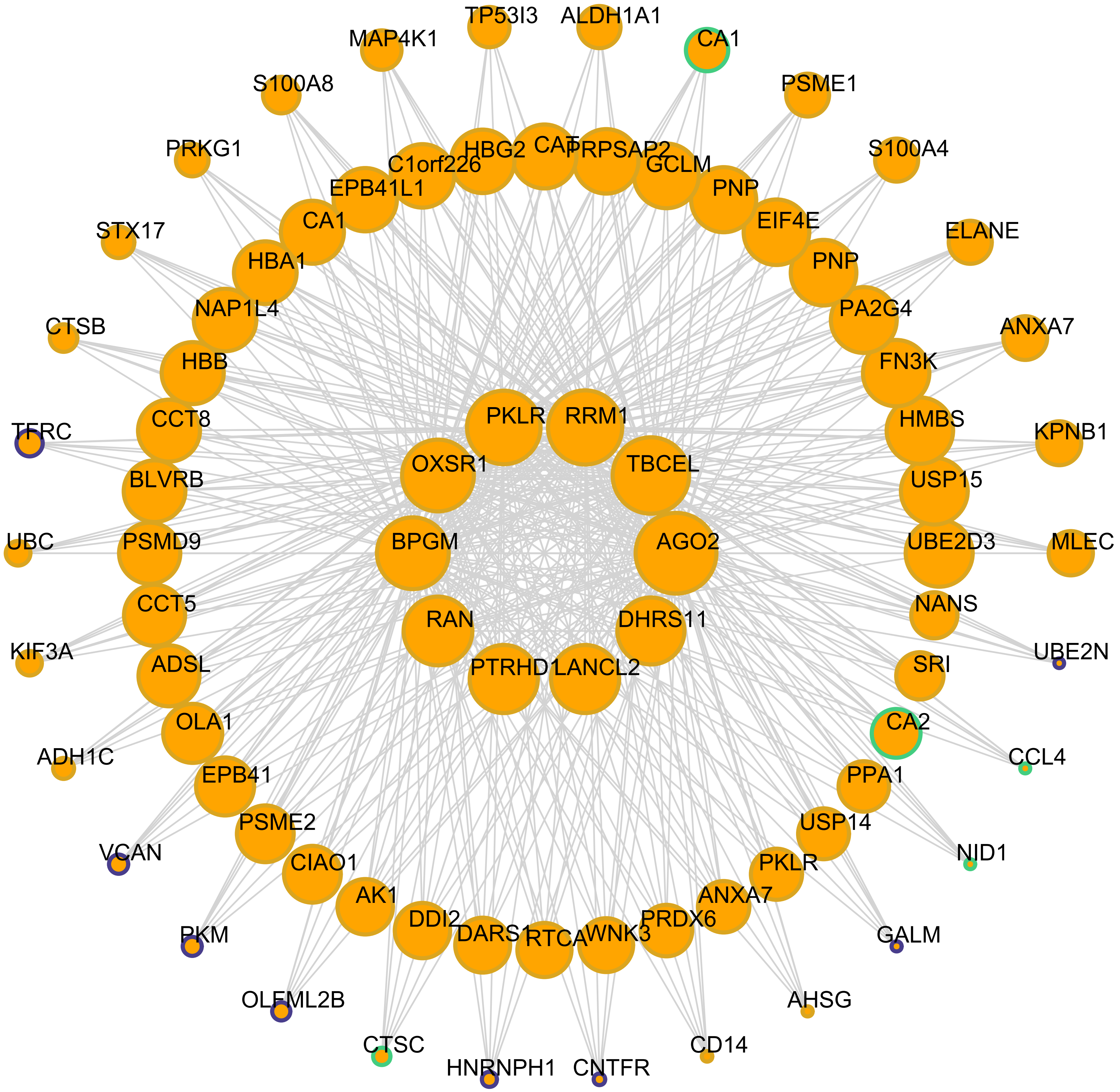

M5 green module: Compound Metabolism/Heat Shock Protein Binding

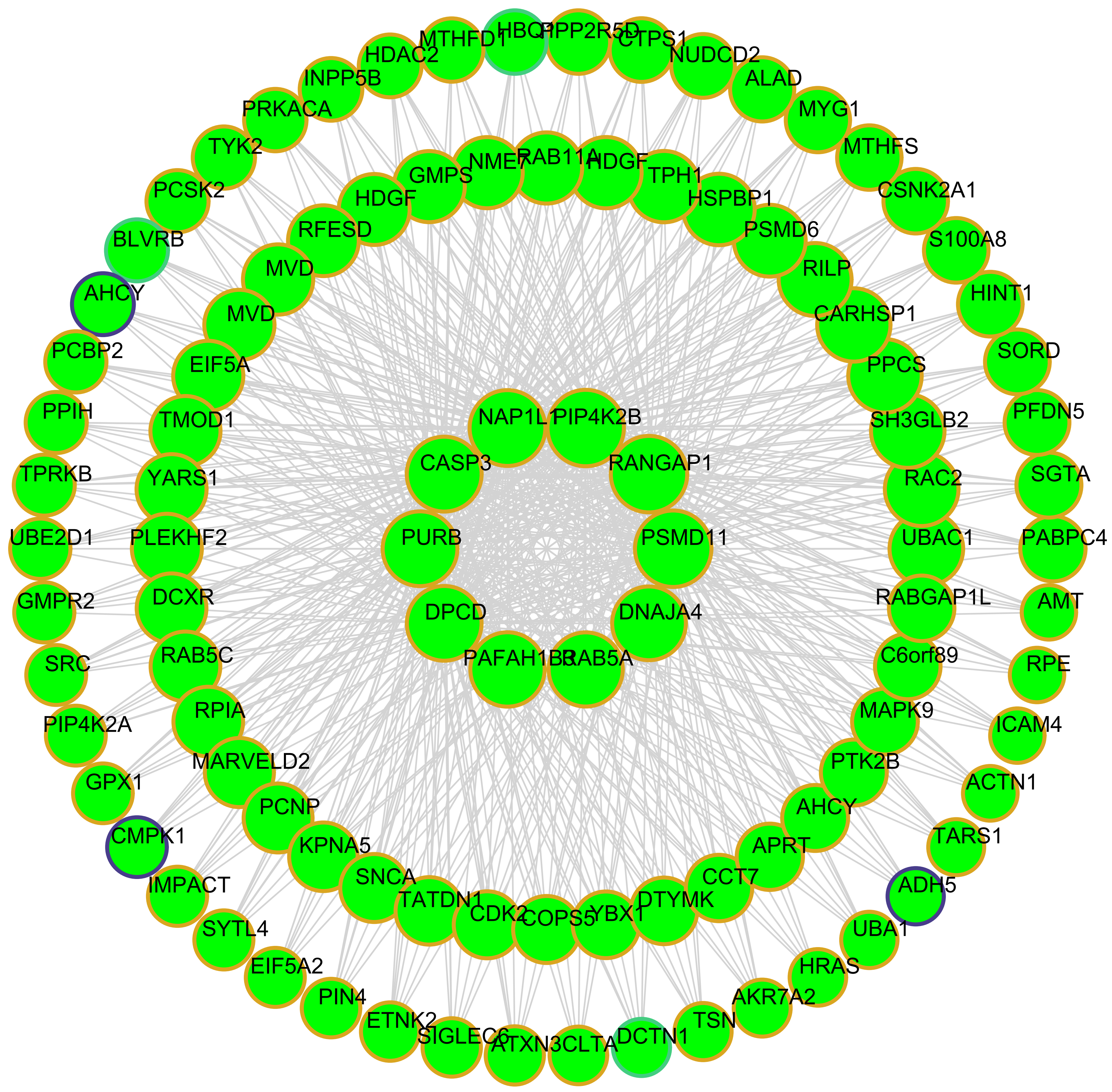

**M37 skyblue3 module: RNA Binding/Metabolism**

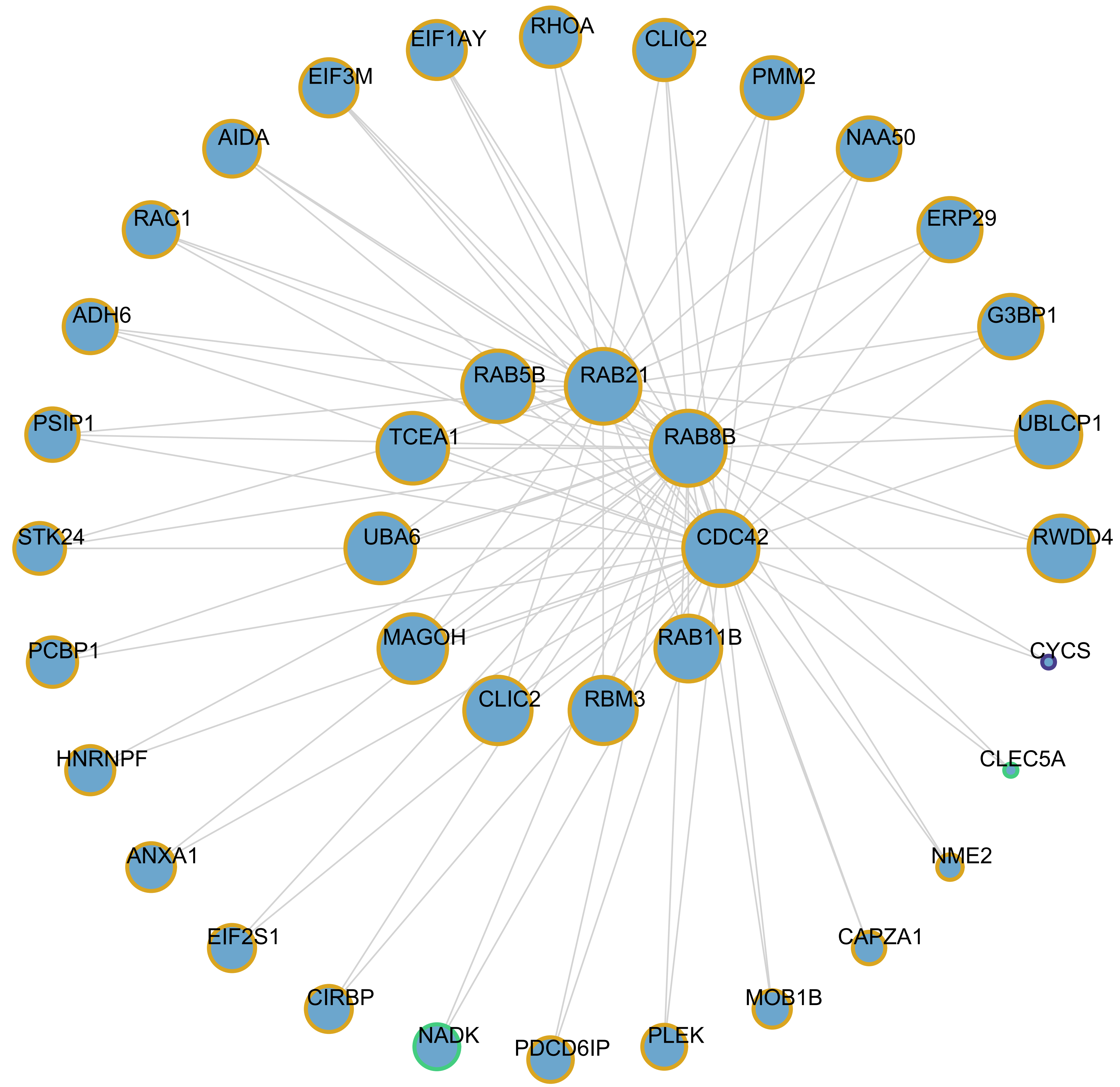

M34 darkmagenta module: Cofactor Biosynthesis

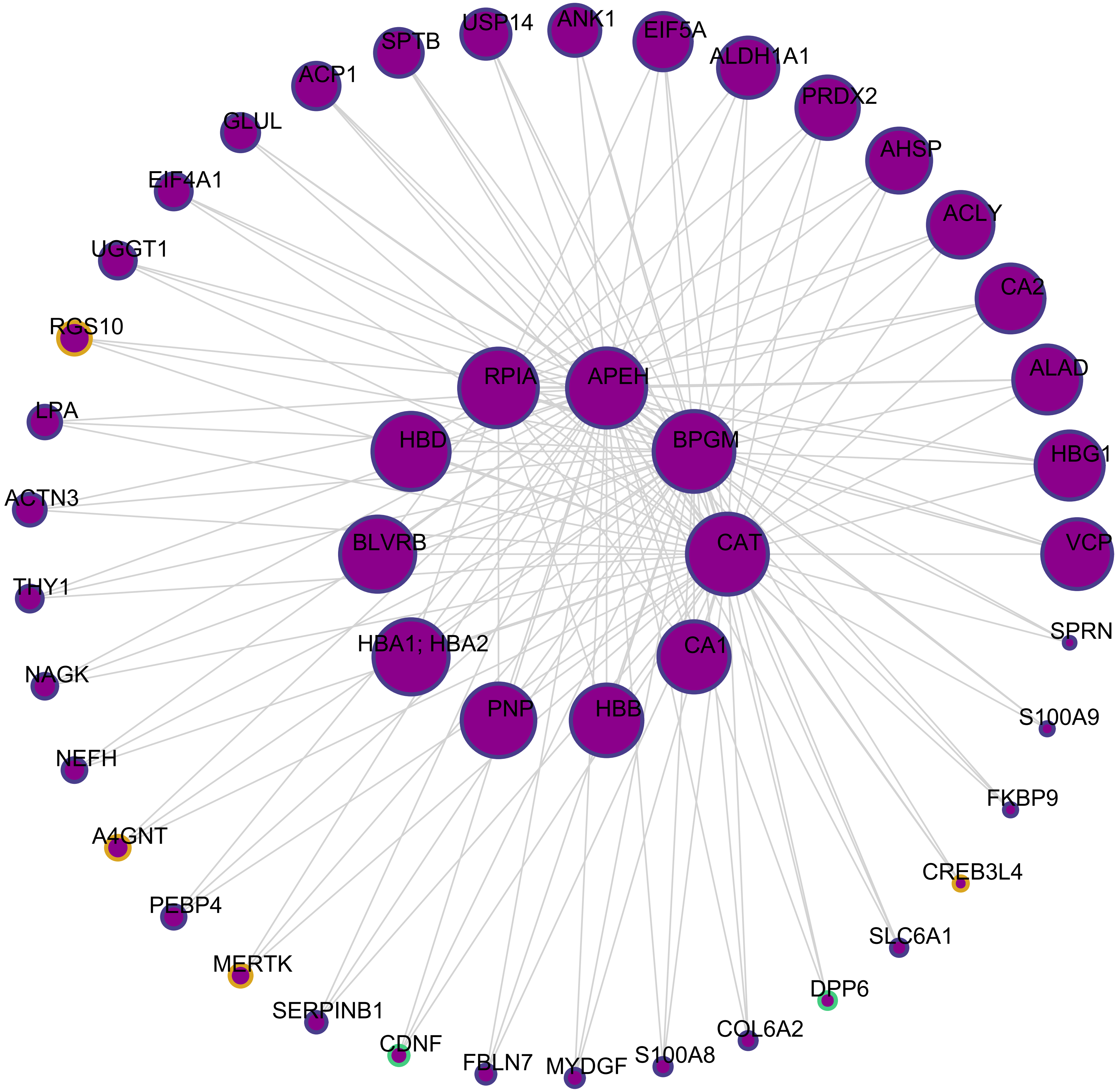

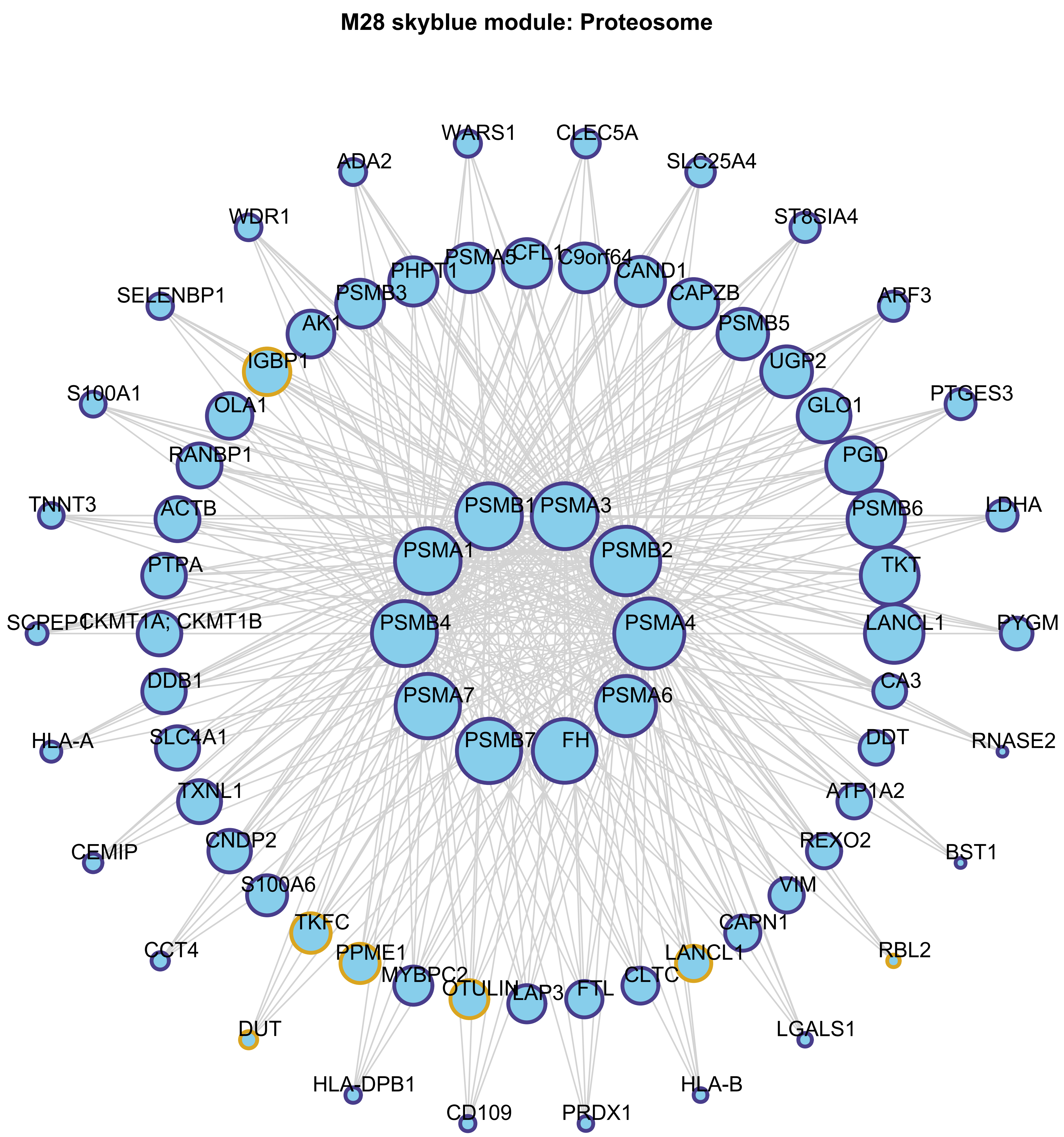

M28 skyblue module: Proteosome
WARS1
CLEC5A
ADA2
SLC25A4
WDR1
ST8SIA4
PSMA5
CFL1
C9orf64
PHPT1
CAND1
SELENBP1
PSMB3
CAPZB
ARF3
AK1
PSMB5
IGBP1
UGP2
S100A1
OLA1
GLO1
PTGES3
RANBP1
PGD
TNNT3
ACTB
PSMB1
PSMA3
PSMB6
LDHA
PSMA1
PSMB2
PTPA
TKT
SCPEP1
CKMT1A; CKMT1B
PSMB4
PSMA4
LANCL1
PYGM
DDB1
CA3
PSMA7
PSMA6
HLA-A
SLC4A1
PSMB7
FH
DDT
RNASE2
TXNL1
ATP1A2
CEMIP
CNDP2
REXO2
BST1
S100A6
VIM
TKFC
CAPN1
CCT4
PPME1
LANCL1
RBL2
MYBPC2
OTULIN
LAP3
FTL
CLTC
DUT
LGALS1
HLA-DPB1
HLA-B
CD109
PRDX1

#### M16 lightcyan module: Sugar Metabolism

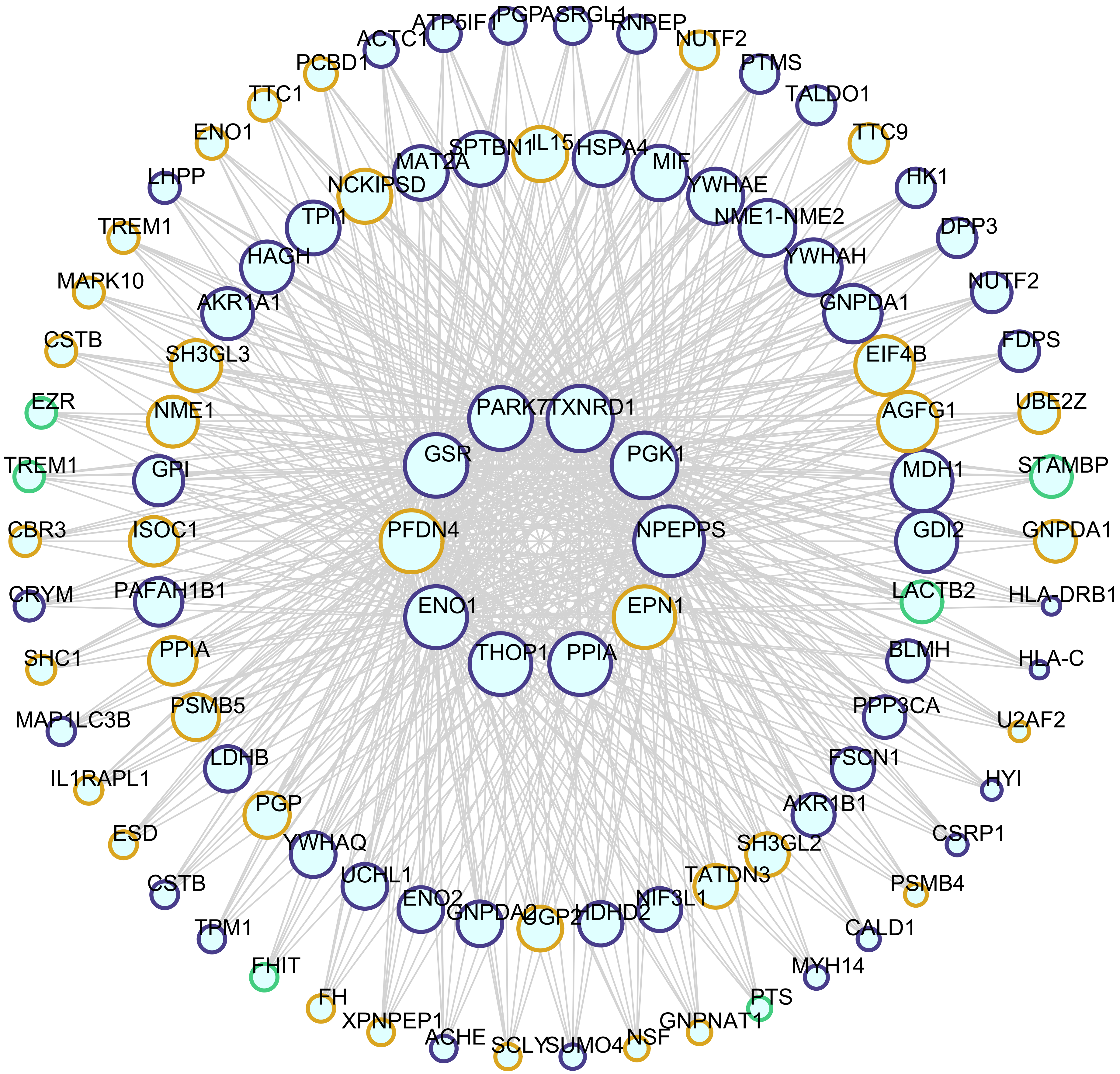

**M29 saddlebrown module: Sugar Metabolism/Actin Depolymerization**

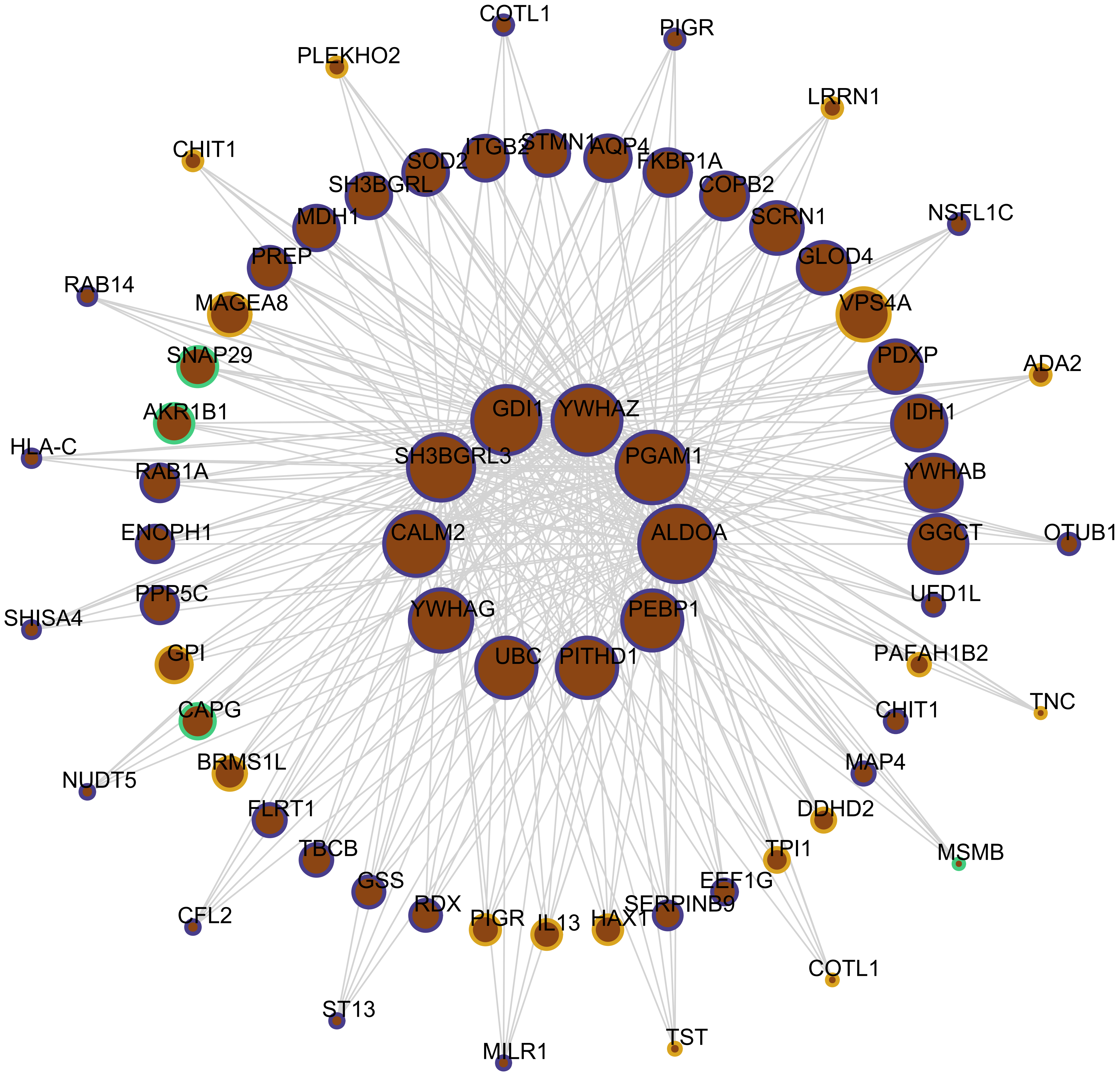

**M24 darkgrey module: Ubiquitination**

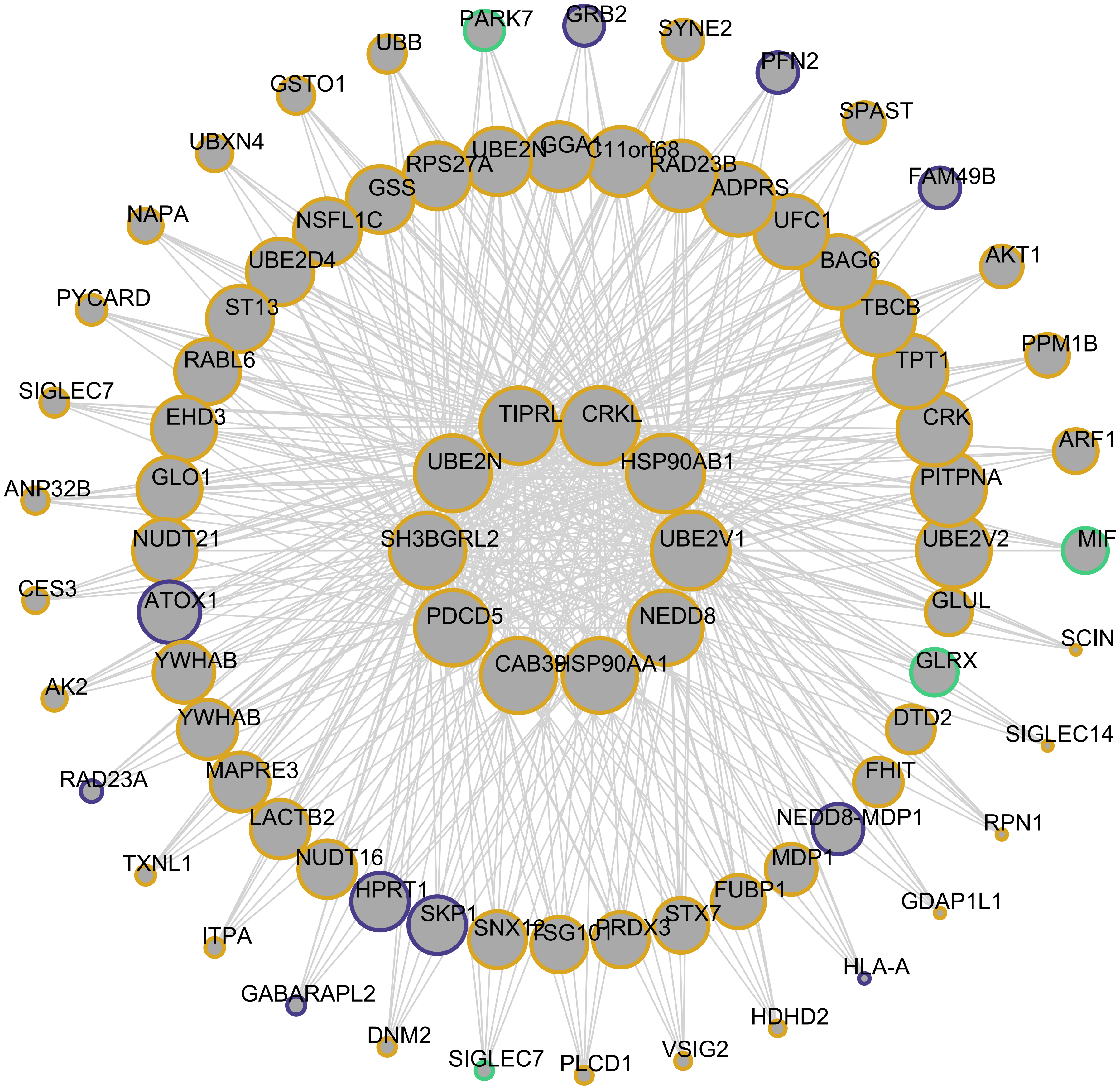

M35 sienna3 module: Ambiguous

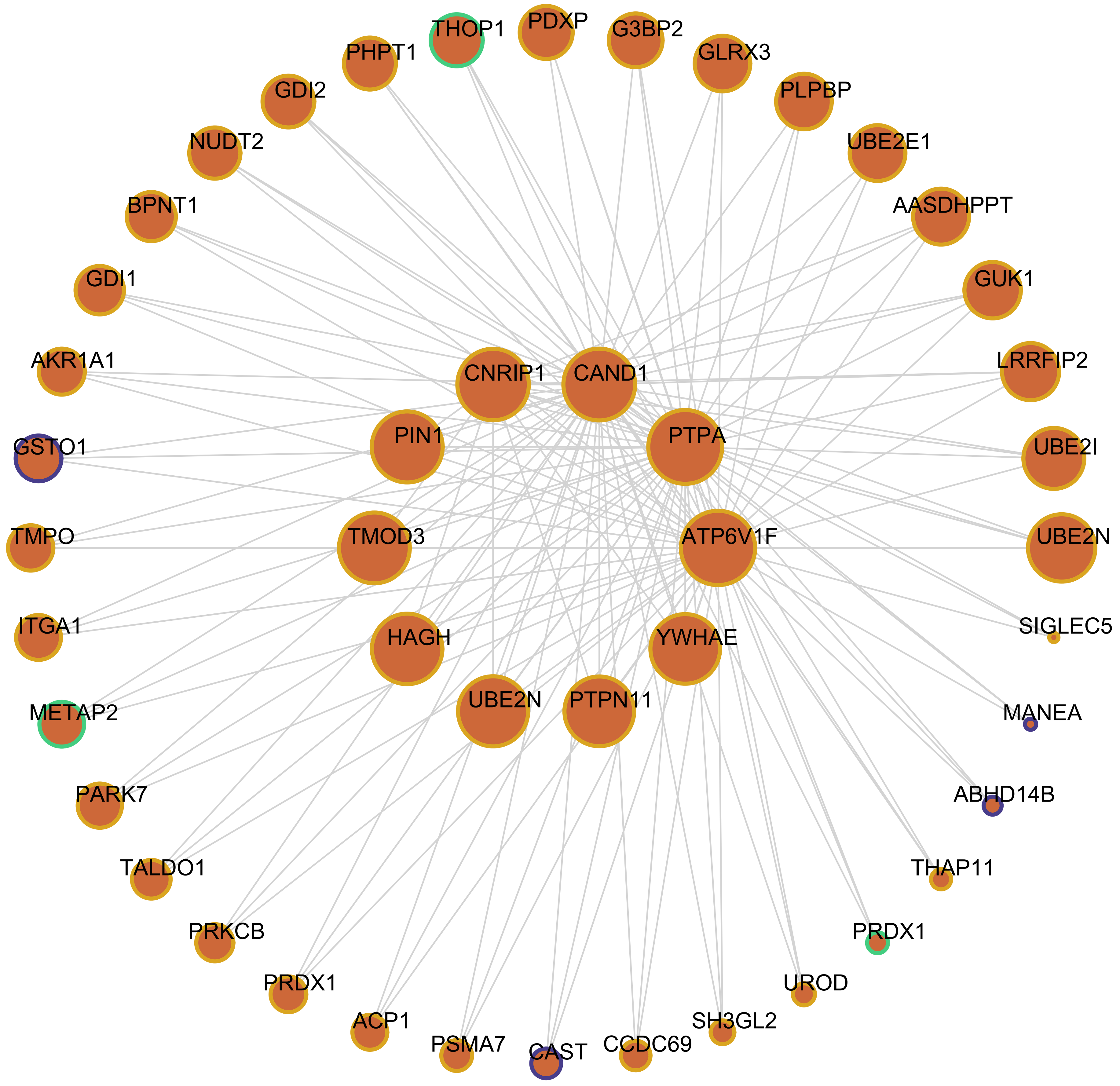

M33 darkolivegreen module: Translation/Sugar Binding

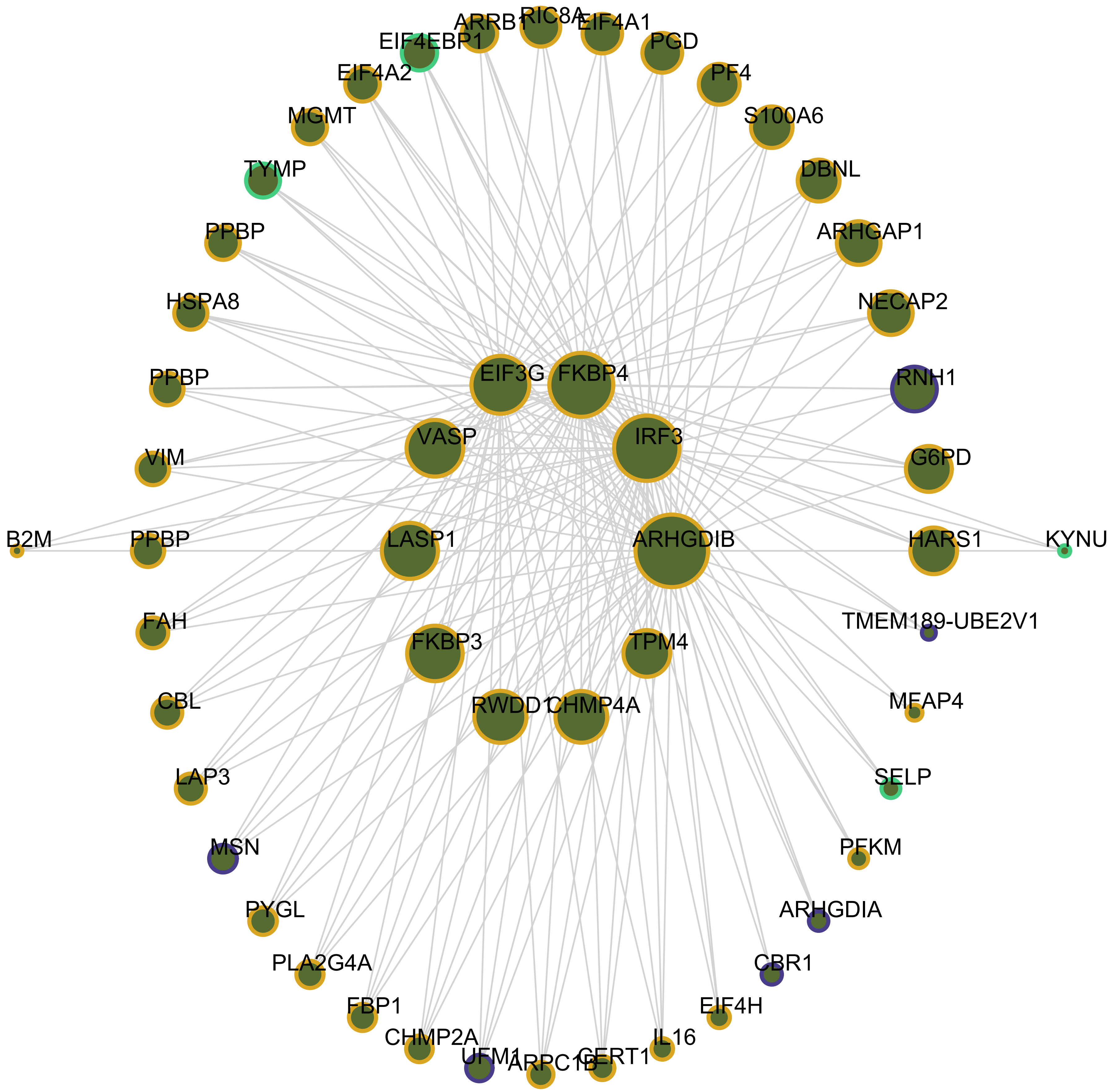

**M36 yellowgreen module: Organelle Organization/Biogenesis**

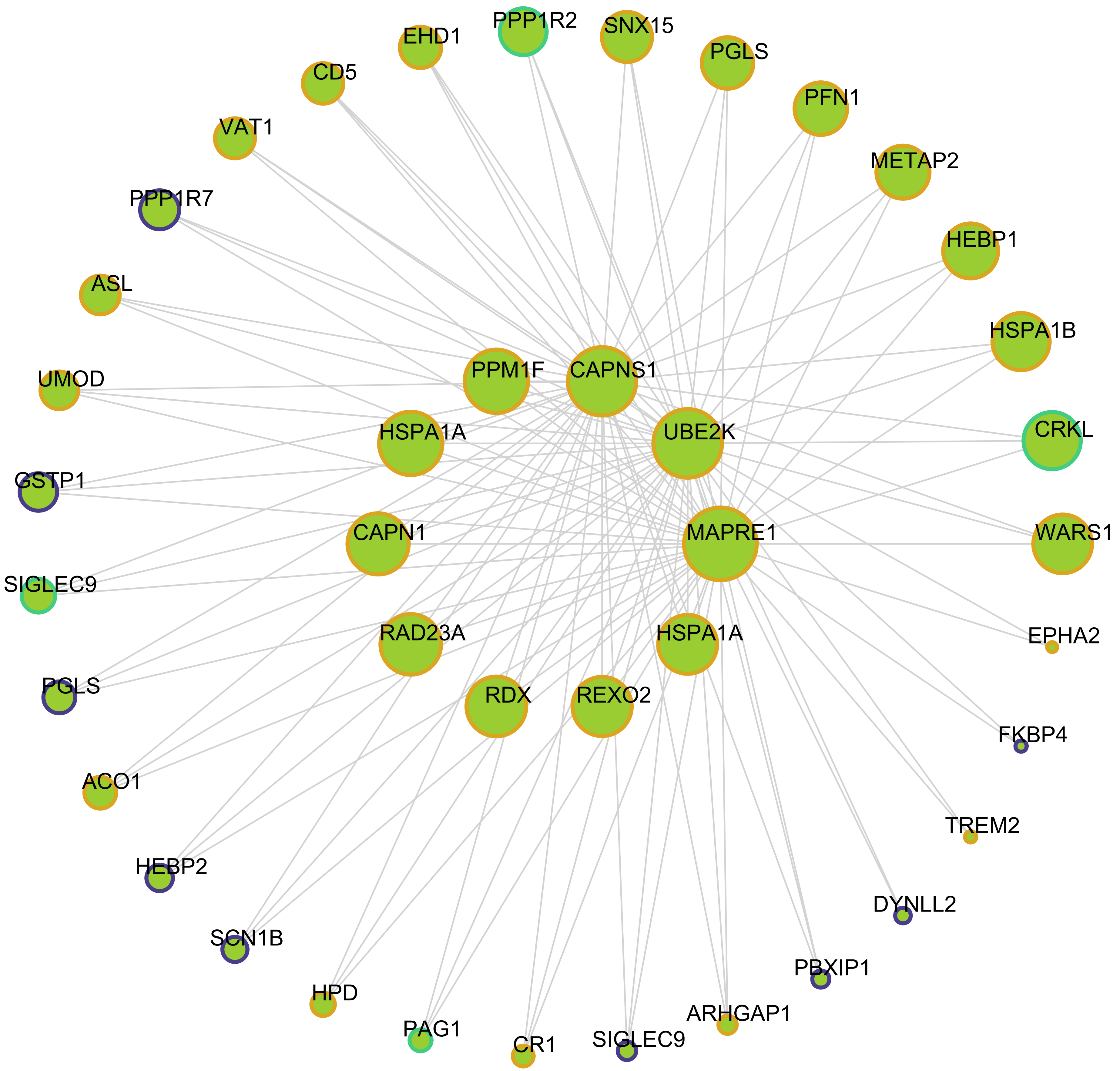

**M1 turquoise module: Immune Response**

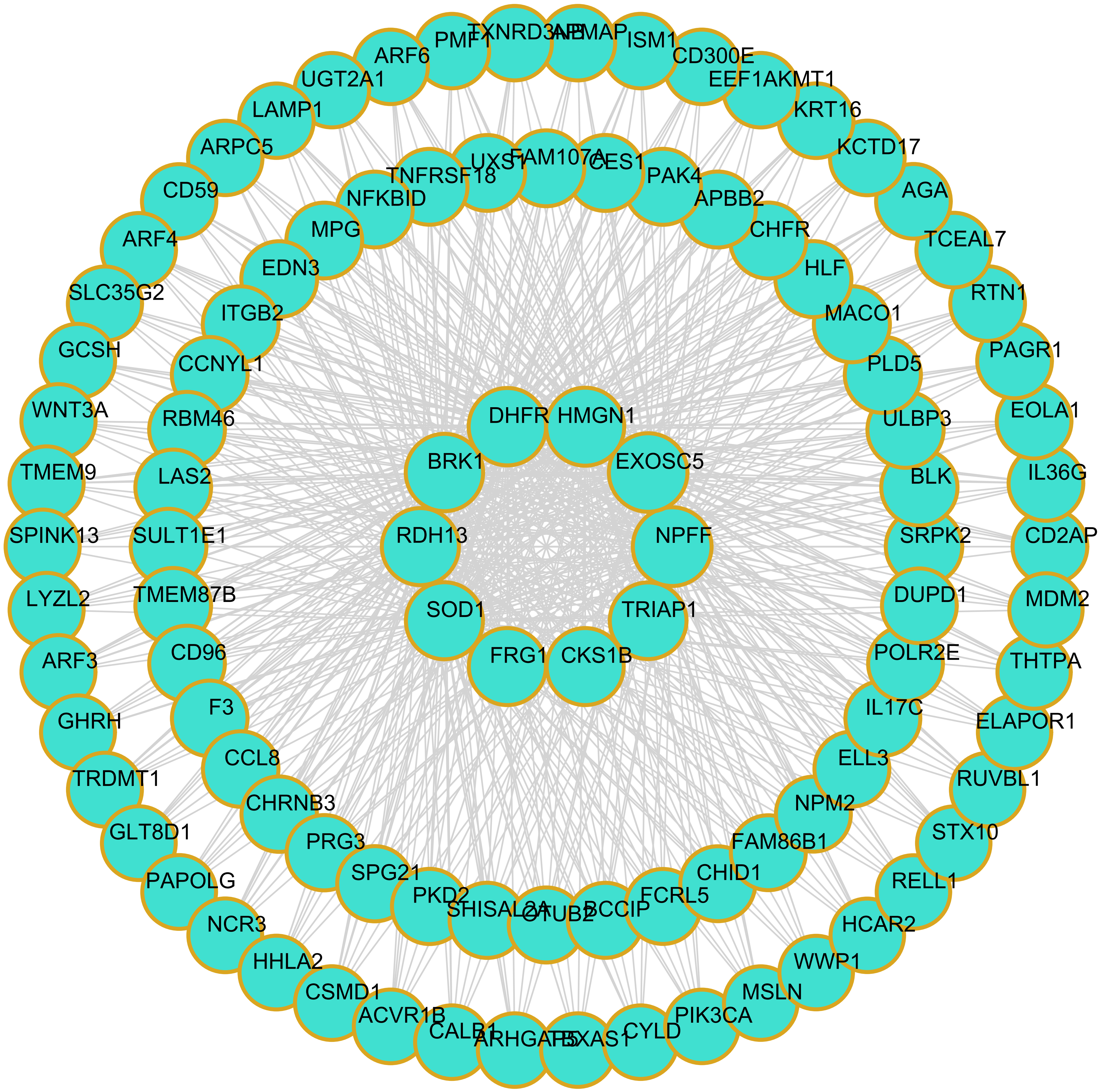

#### M4 yellow module: Lysosome

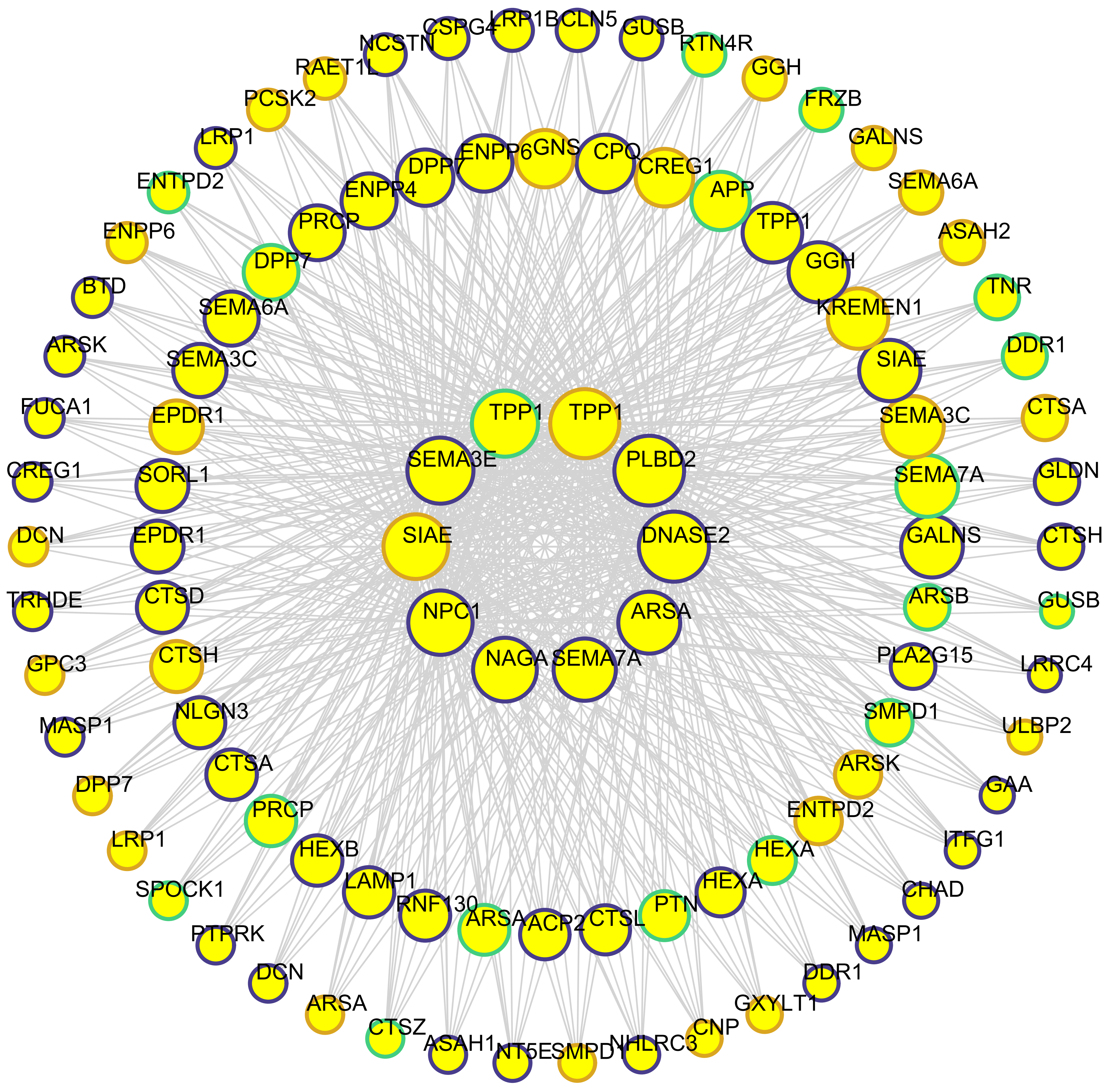

**M23 darkturquoise module: Nucleic Acid/Steroid Metabolism**

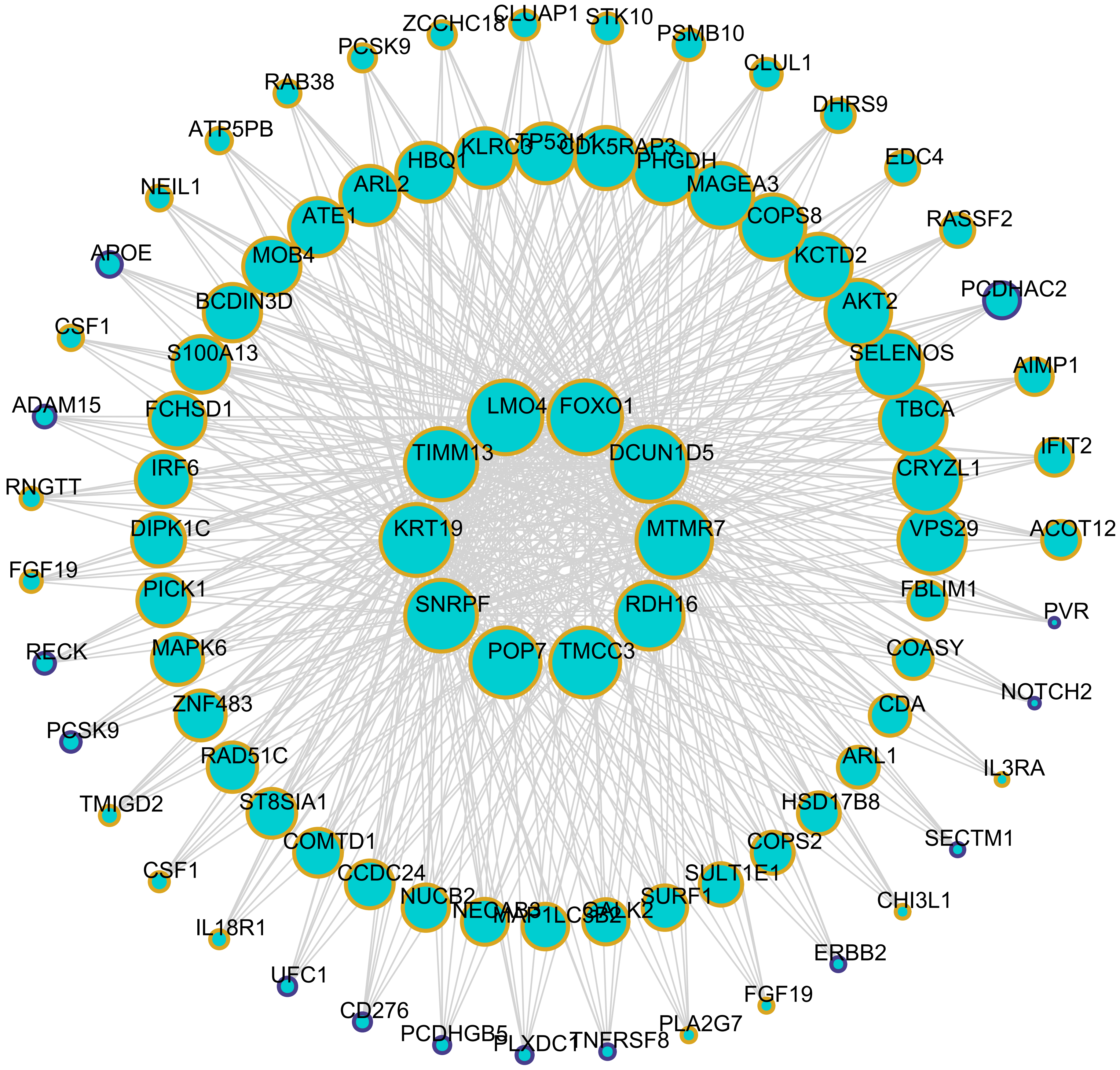

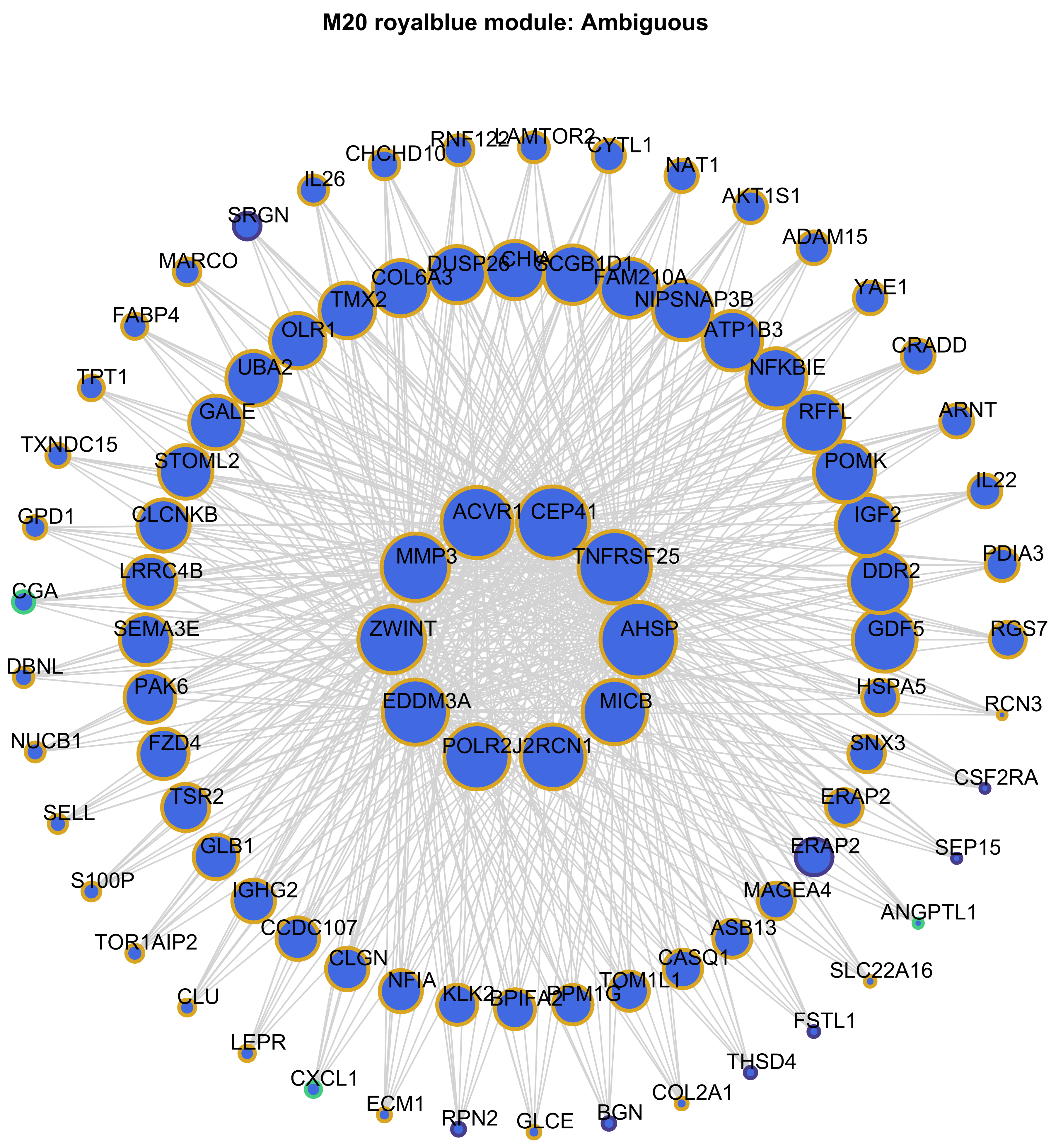
M20 royalblue module: Ambiguous
RNF122
LAMTOR2
CYTL1
CHCHD10
NAT1
IL26
AKT1S1
SRGN
ADAM15
MARCO
DUSP26
CHIA
SCGB1D1
COL6A3
FAM210A
YAE1
TMX2
NIPSNAP3B
FABP4
OLR1
ATP1B3
CRADD
UBA2
NFKBIE
TPT1
GALE
RFFL
ARNT
TXNDC15
STOML2
POMK
IL22
GPD1
CLCNKB
ACVR1
CEP41
IGF2
MMP3
TNFRSF25
PDIA3
LRRC4B
DDR2
CGA
SEMA3E
ZWINT
AHSP
GDF5
RGS7
DBNL
PAK6
HSPA5
EDDM3A
MICB
RCN3
NUCB1
FZD4
POLR2J2
RCN1
SNX3
CSF2RA
SELL
TSR2
ERAP2
GLB1
ERAP2
SEP15
S100P
IGHG2
MAGEA4
ANGPTL1
CCDC107
ASB13
TOR1AIP2
CLGN
CASQ1
SLC22A16
NFIA
TOM1L1
CLU
KLK2
BPIFA2
PPM1G
FSTL1
LEPR
THSD4
CXCL1
COL2A1
ECM1
RPN2
GLCE
BGN

**M22 darkgreen module: ECM/Actin Binding**

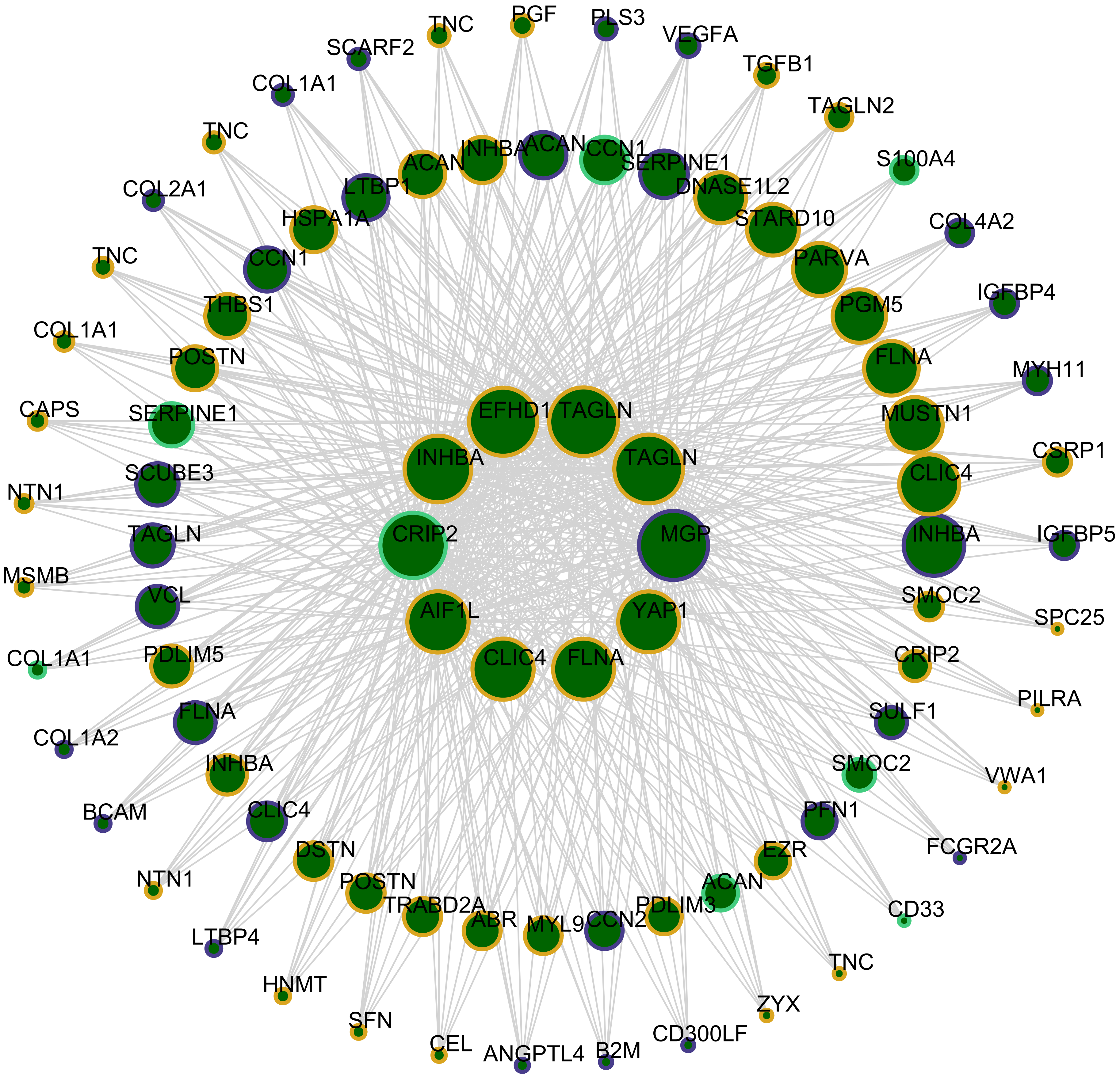

**M26 darkorange module: TGF-? Signaling**

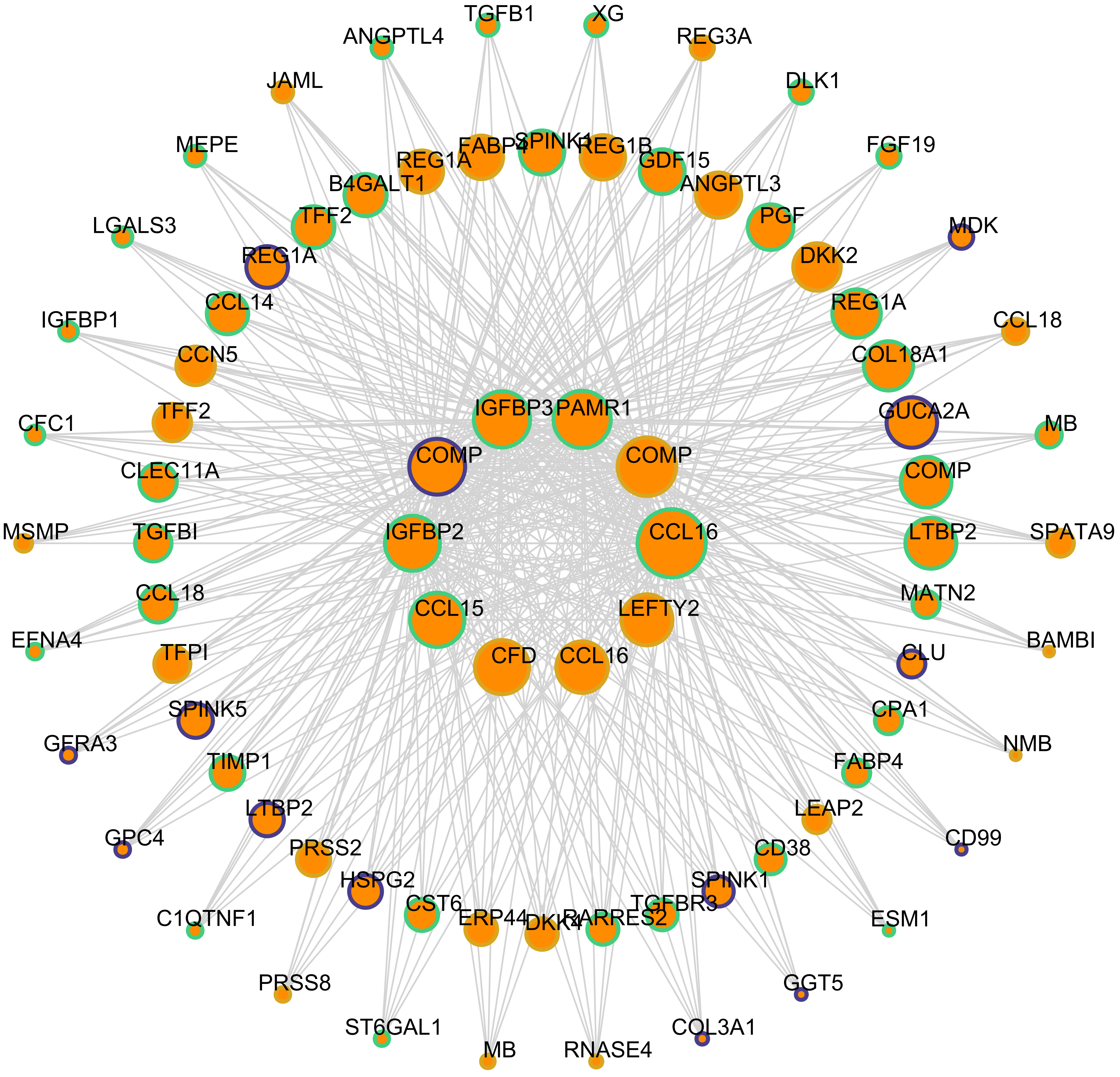

**M18 lightgreen module: ECM Organization/Scavenger Receptor**

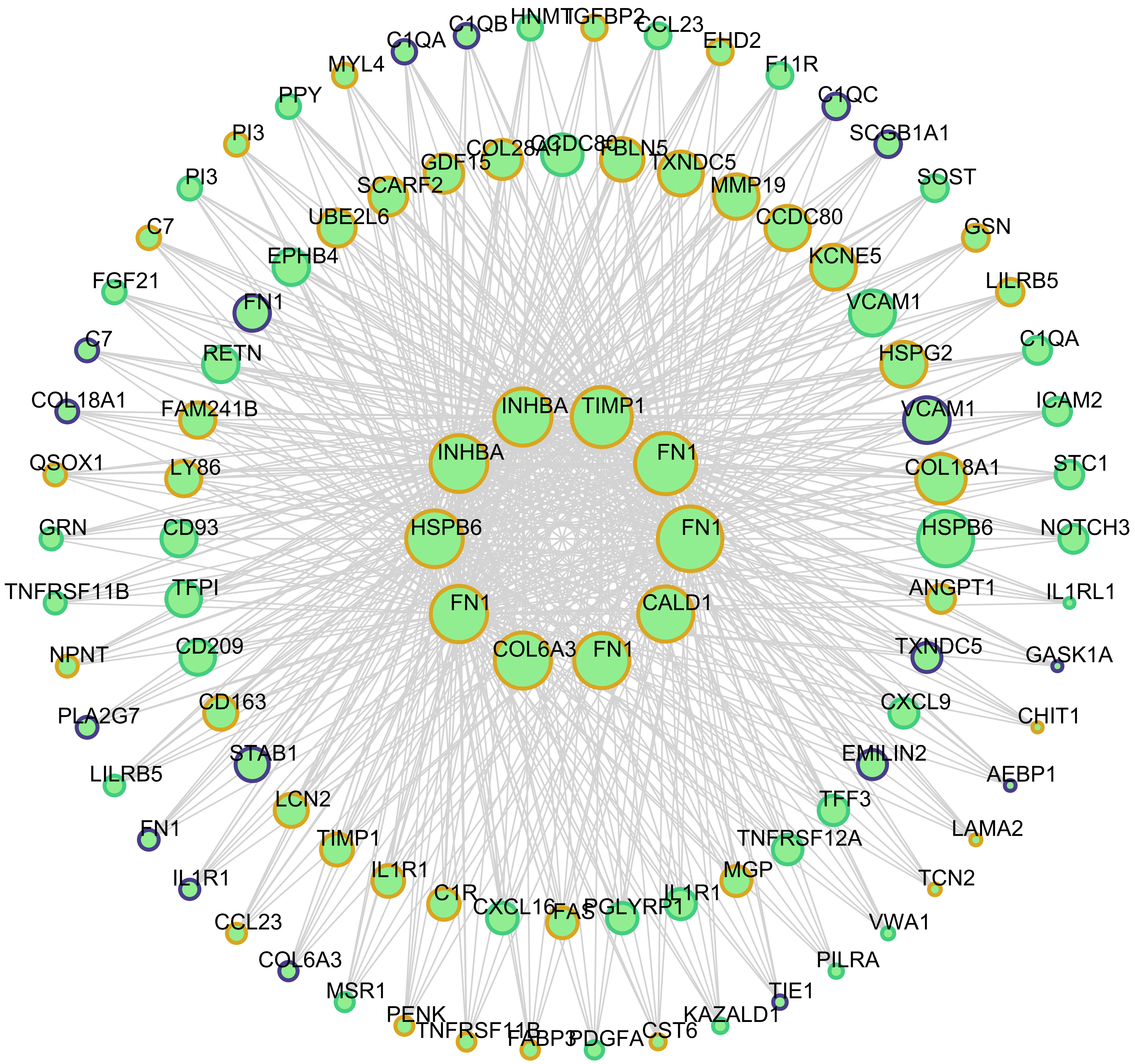

**M3 brown module: Complement/Protein Activation Cascade**

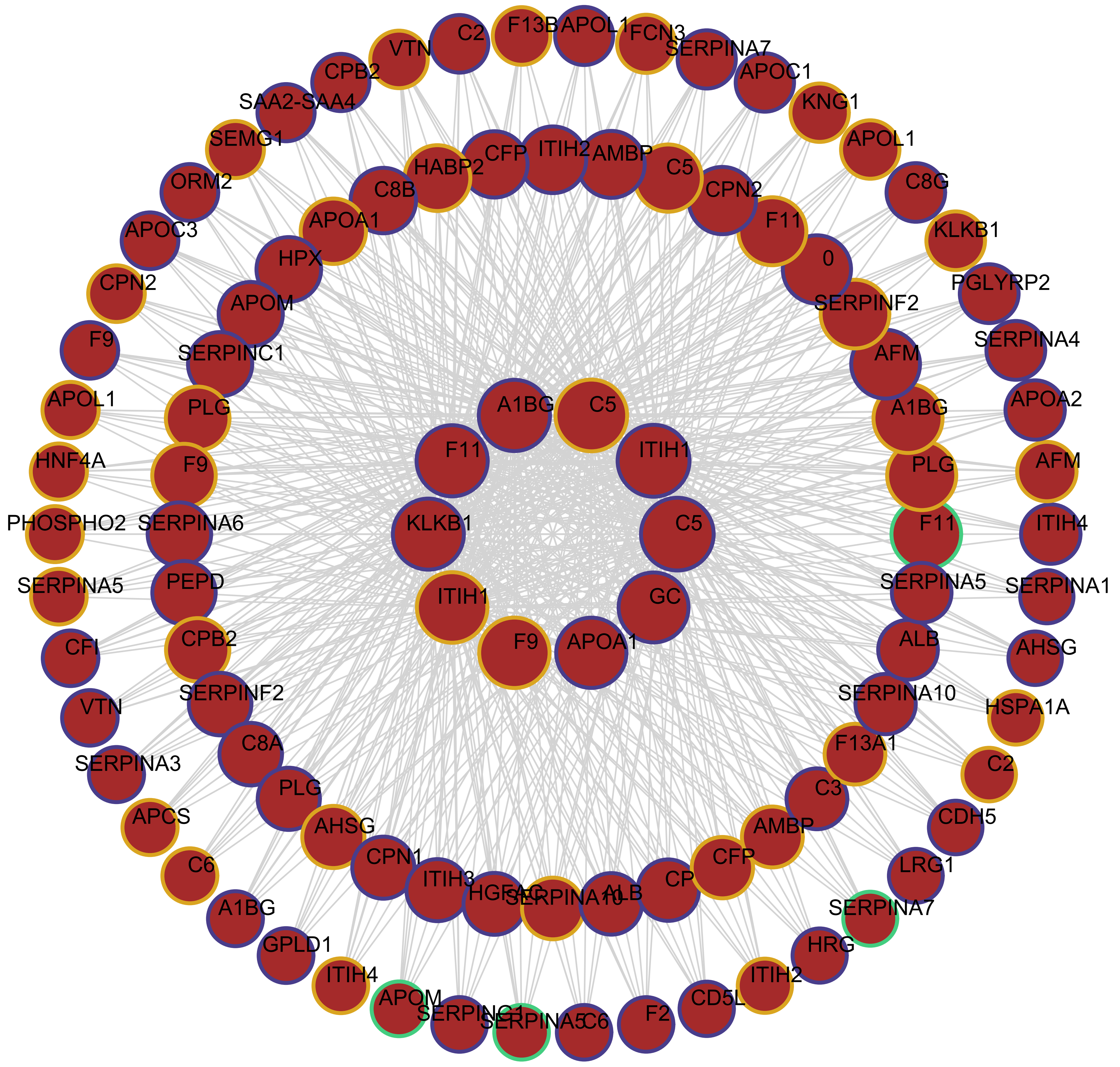

**M10 purple module: Ossification/Bone Development**

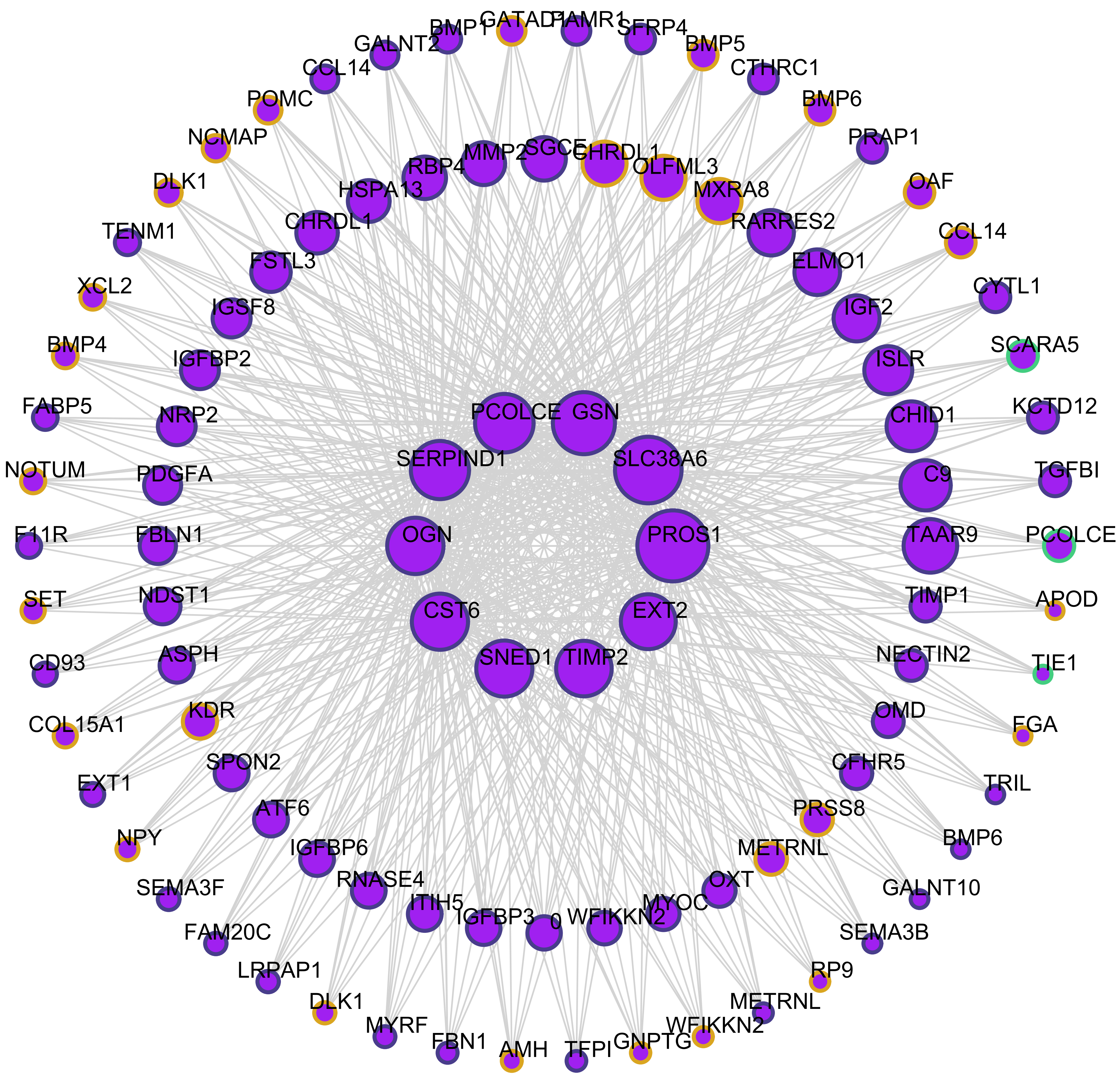

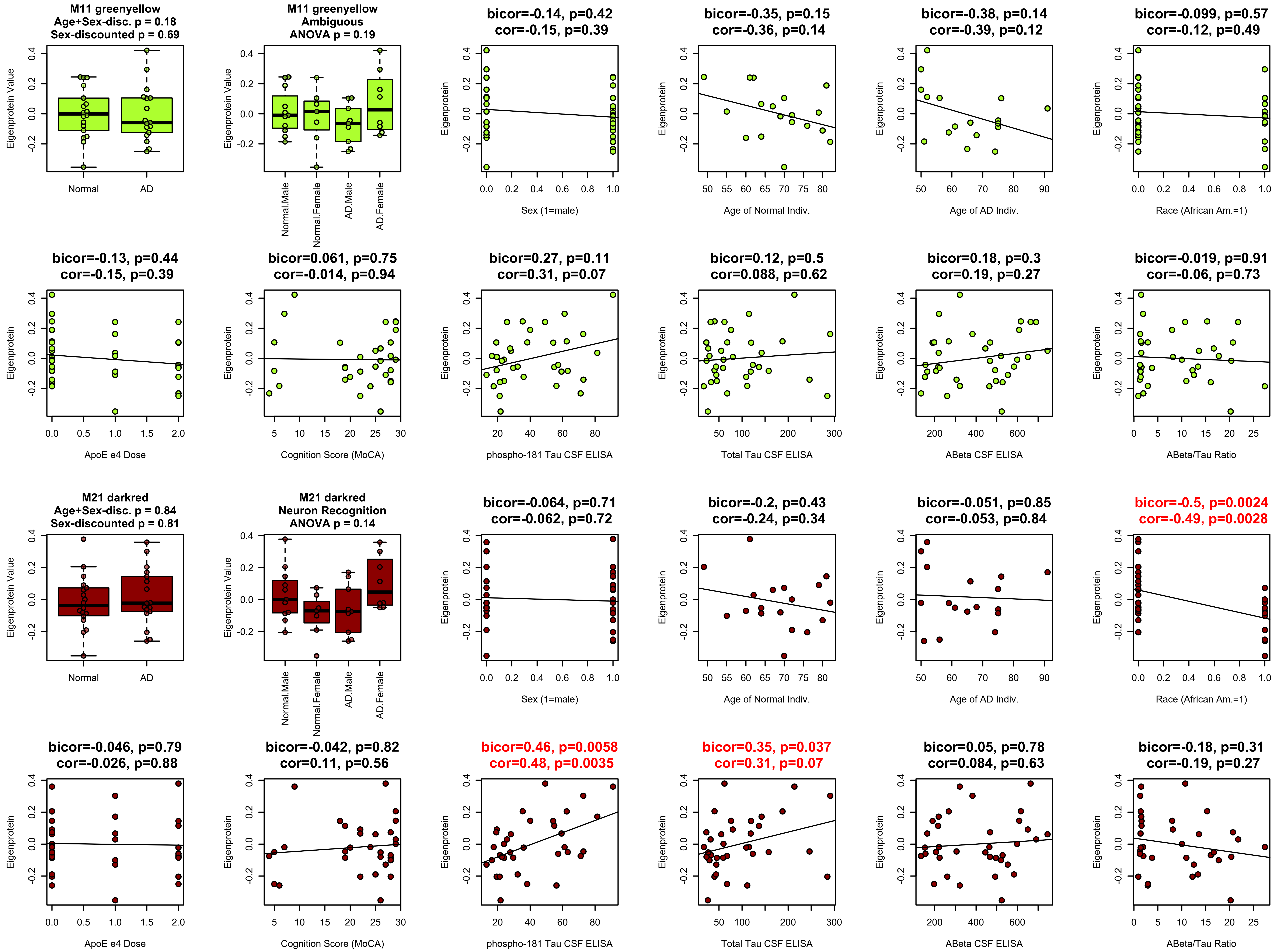

M11 greenyellow
Age+Sex-disc. p = 0.18
Sex-discounted p = 0.69
Eigenprotein Value
Normal
AD
M11 greenyellow
Ambiguous
ANOVA p = 0.19
Eigenprotein Value
Normal.Male
Normal.Female
AD.Male
AD.Female
bicor=-0.14, p=0.42
cor=-0.15, p=0.39
Eigenprotein
Sex (1=male)
bicor=-0.35, p=0.15
cor=-0.36, p=0.14
Eigenprotein
Age of Normal Indiv.
bicor=-0.38, p=0.14
cor=-0.39, p=0.12
Eigenprotein
Age of AD Indiv.
bicor=-0.099, p=0.57
cor=-0.12, p=0.49
Eigenprotein
Race (African Am.=1)
bicor=-0.13, p=0.44
cor=-0.15, p=0.39
Eigenprotein
ApoE e4 Dose
bicor=0.061, p=0.75
cor=-0.014, p=0.94
Eigenprotein
Cognition Score (MoCA)
bicor=0.27, p=0.11
cor=0.31, p=0.07
Eigenprotein
phospho-181 Tau CSF ELISA
bicor=0.12, p=0.5
cor=0.088, p=0.62
Eigenprotein
Total Tau CSF ELISA
bicor=0.18, p=0.3
cor=0.19, p=0.27
Eigenprotein
ABeta CSF ELISA
bicor=-0.019, p=0.91
cor=-0.06, p=0.73
Eigenprotein
ABeta/Tau Ratio
M21 darkred
Age+Sex-disc. p = 0.84
Sex-discounted p = 0.81
Eigenprotein Value
Normal
AD
M21 darkred
Neuron Recognition
ANOVA p = 0.14
Eigenprotein Value
Normal.Male
Normal.Female
AD.Male
AD.Female
bicor=-0.064, p=0.71
cor=-0.062, p=0.72
Eigenprotein
Sex (1=male)
bicor=-0.2, p=0.43
cor=-0.24, p=0.34
Eigenprotein
Age of Normal Indiv.
bicor=-0.051, p=0.85
cor=-0.053, p=0.84
Eigenprotein
Age of AD Indiv.
bicor=-0.5, p=0.0024
cor=-0.49, p=0.0028
Eigenprotein
Race (African Am.=1)
bicor=-0.046, p=0.79
cor=-0.026, p=0.88
Eigenprotein
ApoE e4 Dose
bicor=-0.042, p=0.82
cor=0.11, p=0.56
Eigenprotein
Cognition Score (MoCA)
bicor=0.46, p=0.0058
cor=0.48, p=0.0035
Eigenprotein
phospho-181 Tau CSF ELISA
bicor=0.35, p=0.037
cor=0.31, p=0.07
Eigenprotein
Total Tau CSF ELISA
bicor=0.05, p=0.78
cor=0.084, p=0.63
Eigenprotein
ABeta CSF ELISA
bicor=-0.18, p=0.31
cor=-0.19, p=0.27
Eigenprotein
ABeta/Tau Ratio

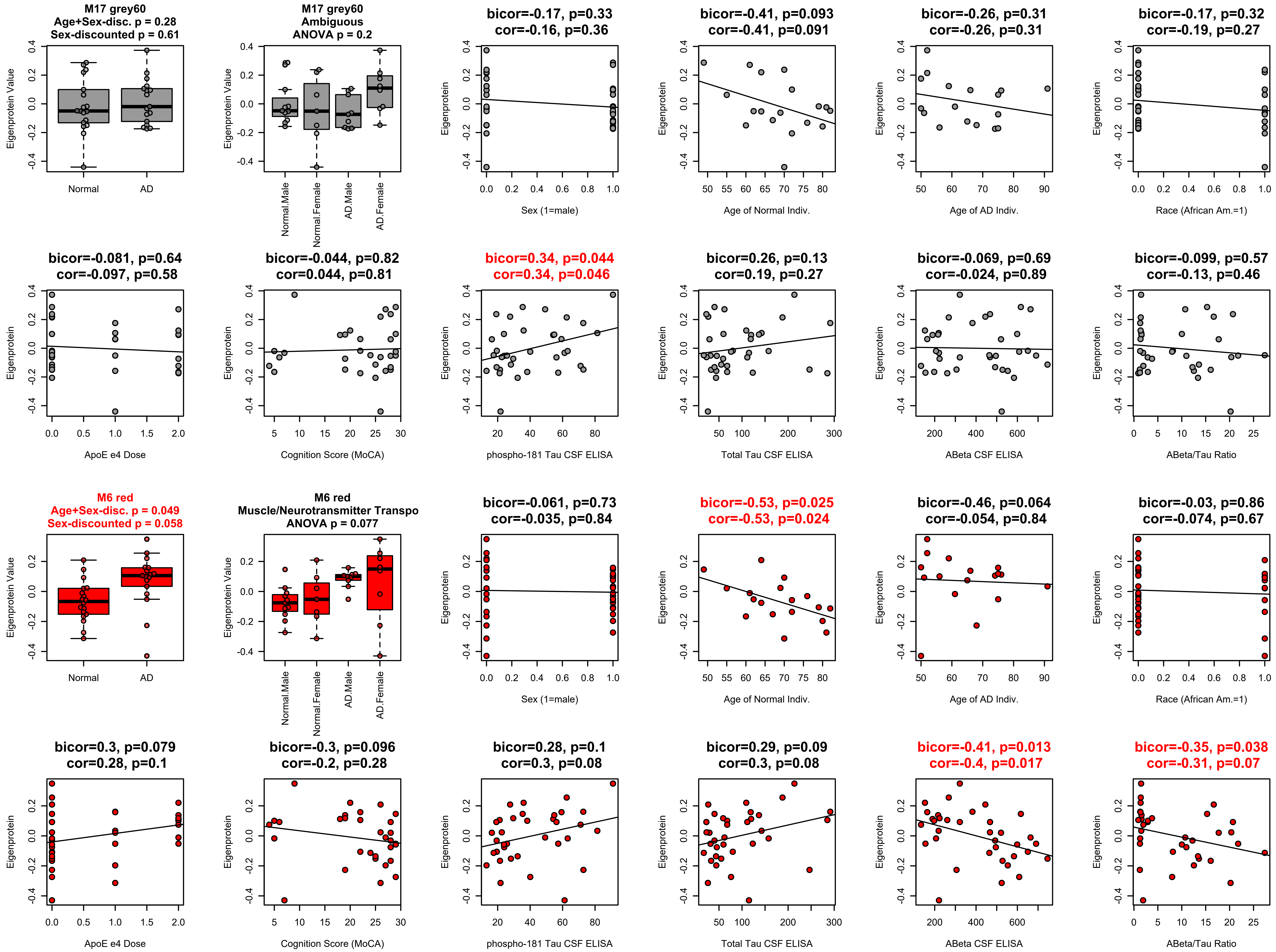

M17 grey60
Age+Sex-disc. p = 0.28
Sex-discounted p = 0.61
M17 grey60
Ambiguous
ANOVA p = 0.2
bicor=-0.17, p=0.33
cor=-0.16, p=0.36
bicor=-0.41, p=0.093
cor=-0.41, p=0.091
bicor=-0.26, p=0.31
cor=-0.26, p=0.31
bicor=-0.17, p=0.32
cor=-0.19, p=0.27
Eigenprotein Value
Eigenprotein
Normal
AD
Normal.Male
Normal.Female
AD.Male
AD.Female
Sex (1=male)
Age of Normal Indiv.
Age of AD Indiv.
Race (African Am.=1)
bicor=-0.081, p=0.64
cor=-0.097, p=0.58
bicor=-0.044, p=0.82
cor=0.044, p=0.81
bicor=0.34, p=0.044
cor=0.34, p=0.046
bicor=0.26, p=0.13
cor=0.19, p=0.27
bicor=-0.069, p=0.69
cor=-0.024, p=0.89
bicor=-0.099, p=0.57
cor=-0.13, p=0.46
ApoE e4 Dose
Cognition Score (MoCA)
phospho-181 Tau CSF ELISA
Total Tau CSF ELISA
ABeta CSF ELISA
ABeta/Tau Ratio
M6 red
Age+Sex-disc. p = 0.049
Sex-discounted p = 0.058
M6 red
Muscle/Neurotransmitter Transpo
ANOVA p = 0.077
bicor=-0.061, p=0.73
cor=-0.035, p=0.84
bicor=-0.53, p=0.025
cor=-0.53, p=0.024
bicor=-0.46, p=0.064
cor=-0.054, p=0.84
bicor=-0.03, p=0.86
cor=-0.074, p=0.67
bicor=0.3, p=0.079
cor=0.28, p=0.1
bicor=-0.3, p=0.096
cor=-0.2, p=0.28
bicor=0.28, p=0.1
cor=0.3, p=0.08
bicor=0.29, p=0.09
cor=0.3, p=0.08
bicor=-0.41, p=0.013
cor=-0.4, p=0.017
bicor=-0.35, p=0.038
cor=-0.31, p=0.07

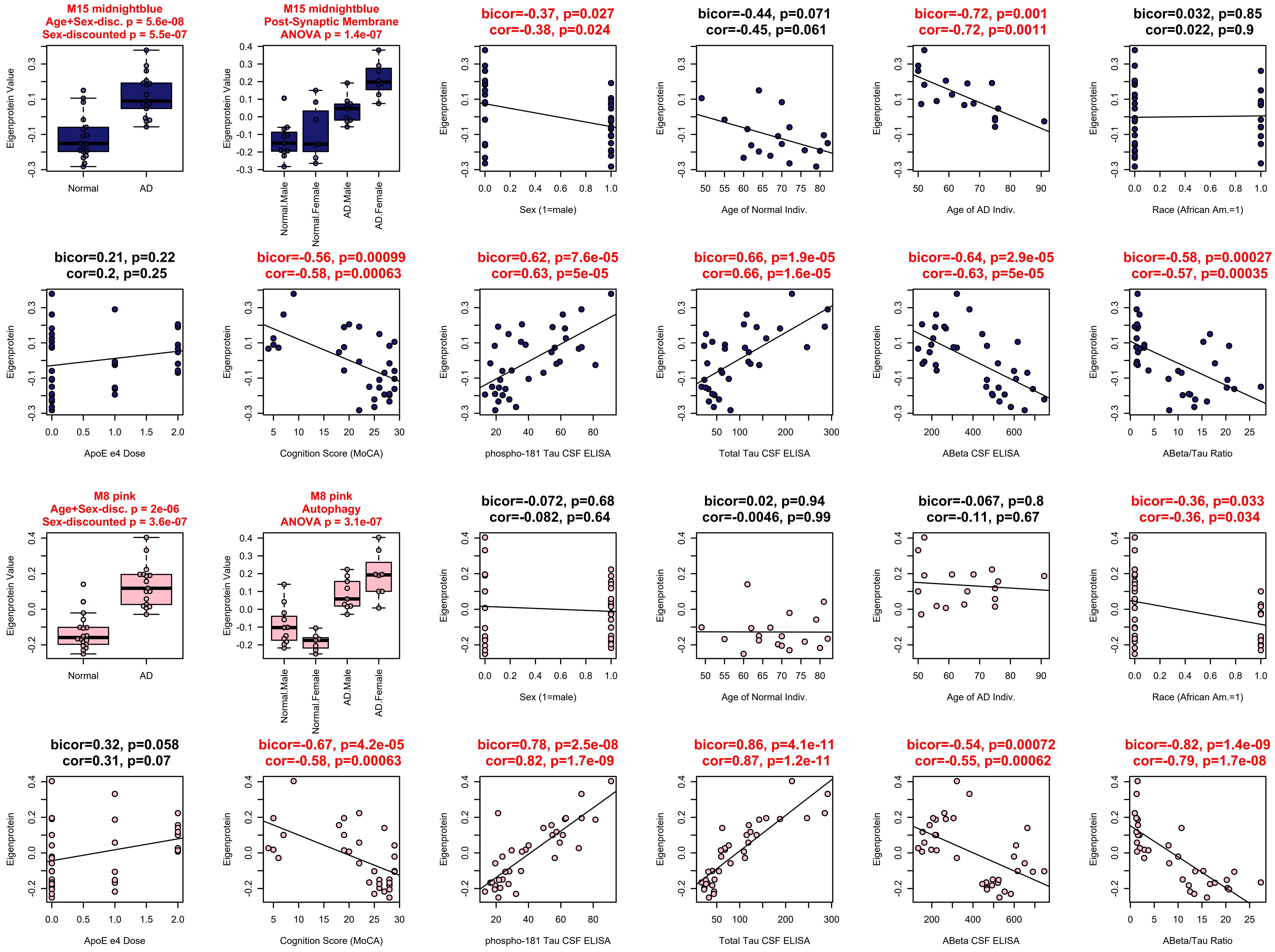
M15 midnightblue
Age+Sex-disc. p = 5.6e-08
Sex-discounted p = 5.5e-07
Eigenprotein Value
Normal
AD
M15 midnightblue
Post-Synaptic Membrane
ANOVA p = 1.4e-07
Eigenprotein Value
Normal.Male
Normal.Female
AD.Male
AD.Female
bicor=-0.37, p=0.027
cor=-0.38, p=0.024
Eigenprotein
Sex (1=male)
bicor=-0.44, p=0.071
cor=-0.45, p=0.061
Eigenprotein
Age of Normal Indiv.
bicor=-0.72, p=0.001
cor=-0.72, p=0.0011
Eigenprotein
Age of AD Indiv.
bicor=0.032, p=0.85
cor=0.022, p=0.9
Eigenprotein
Race (African Am.=1)
bicor=0.21, p=0.22
cor=0.2, p=0.25
Eigenprotein
ApoE e4 Dose
bicor=-0.56, p=0.00099
cor=-0.58, p=0.00063
Eigenprotein
Cognition Score (MoCA)
bicor=0.62, p=7.6e-05
cor=0.63, p=5e-05
Eigenprotein
phospho-181 Tau CSF ELISA
bicor=0.66, p=1.9e-05
cor=0.66, p=1.6e-05
Eigenprotein
Total Tau CSF ELISA
bicor=-0.64, p=2.9e-05
cor=-0.63, p=5e-05
Eigenprotein
ABeta CSF ELISA
bicor=-0.58, p=0.00027
cor=-0.57, p=0.00035
Eigenprotein
ABeta/Tau Ratio
M8 pink
Age+Sex-disc. p = 2e-06
Sex-discounted p = 3.6e-07
Eigenprotein Value
Normal
AD
M8 pink
Autophagy
ANOVA p = 3.1e-07
Eigenprotein Value
Normal.Male
Normal.Female
AD.Male
AD.Female
bicor=-0.072, p=0.68
cor=-0.082, p=0.64
Eigenprotein
Sex (1=male)
bicor=0.02, p=0.94
cor=-0.0046, p=0.99
Eigenprotein
Age of Normal Indiv.
bicor=-0.067, p=0.8
cor=-0.11, p=0.67
Eigenprotein
Age of AD Indiv.
bicor=-0.36, p=0.033
cor=-0.36, p=0.034
Eigenprotein
Race (African Am.=1)
bicor=0.32, p=0.058
cor=0.31, p=0.07
Eigenprotein
ApoE e4 Dose
bicor=-0.67, p=4.2e-05
cor=-0.58, p=0.00063
Eigenprotein
Cognition Score (MoCA)
bicor=0.78, p=2.5e-08
cor=0.82, p=1.7e-09
Eigenprotein
phospho-181 Tau CSF ELISA
bicor=0.86, p=4.1e-11
cor=0.87, p=1.2e-11
Eigenprotein
Total Tau CSF ELISA
bicor=-0.54, p=0.00072
cor=-0.55, p=0.00062
Eigenprotein
ABeta CSF ELISA
bicor=-0.82, p=1.4e-09
cor=-0.79, p=1.7e-08
Eigenprotein
ABeta/Tau Ratio

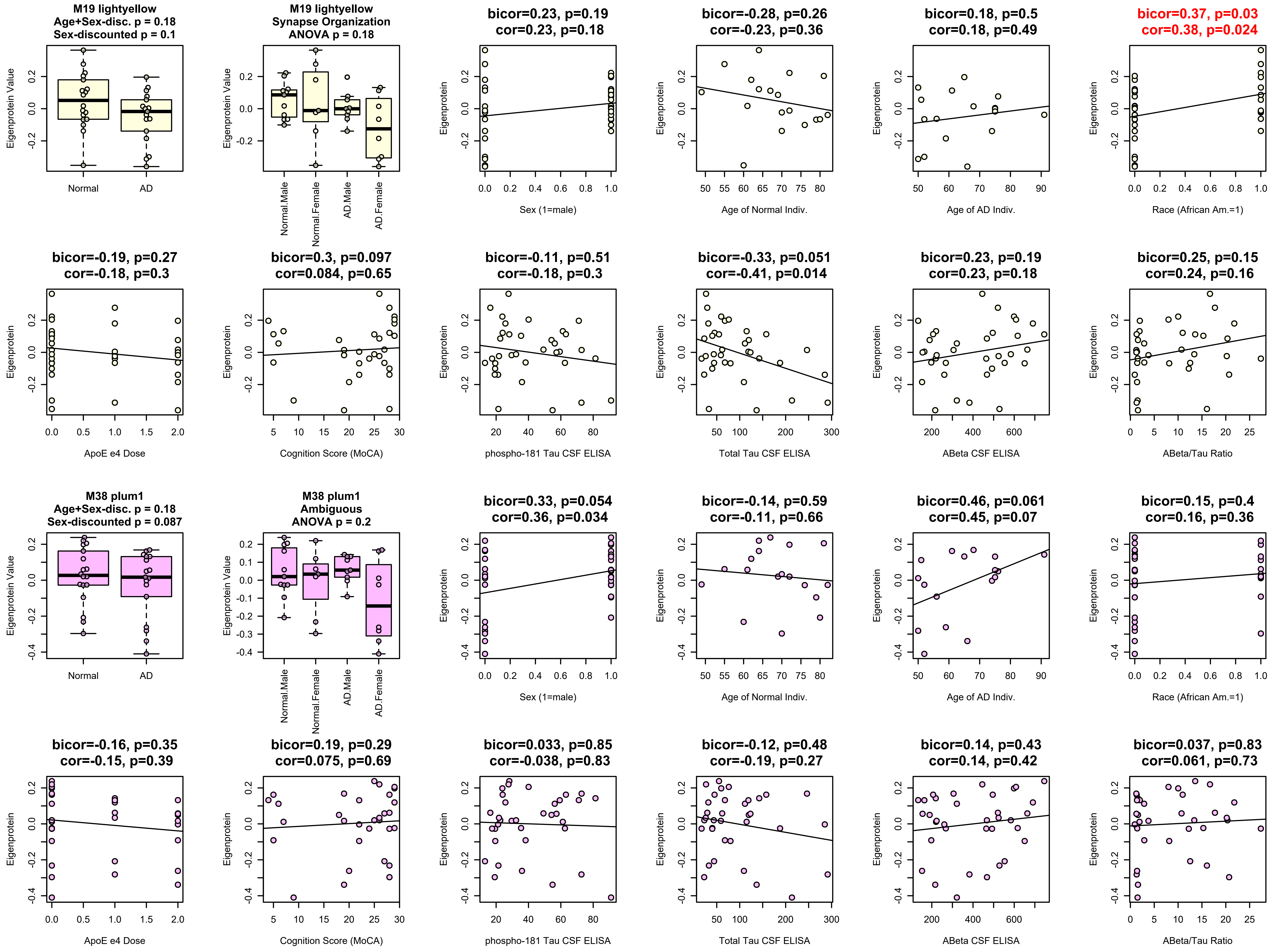
M19 lightyellow
Age+Sex-disc. p = 0.18
Sex-discounted p = 0.1
Eigenprotein Value
Normal
AD
M19 lightyellow
Synapse Organization
ANOVA p = 0.18
Normal.Male
Normal.Female
AD.Male
AD.Female
bicor=0.23, p=0.19
cor=0.23, p=0.18
Sex (1=male)
bicor=-0.28, p=0.26
cor=-0.23, p=0.36
Age of Normal Indiv.
bicor=0.18, p=0.5
cor=0.18, p=0.49
Age of AD Indiv.
bicor=0.37, p=0.03
cor=0.38, p=0.024
Race (African Am.=1)
bicor=-0.19, p=0.27
cor=-0.18, p=0.3
ApoE e4 Dose
bicor=0.3, p=0.097
cor=0.084, p=0.65
Cognition Score (MoCA)
bicor=-0.11, p=0.51
cor=-0.18, p=0.3
phospho-181 Tau CSF ELISA
bicor=-0.33, p=0.051
cor=-0.41, p=0.014
Total Tau CSF ELISA
bicor=0.23, p=0.19
cor=0.23, p=0.18
ABeta CSF ELISA
bicor=0.25, p=0.15
cor=0.24, p=0.16
ABeta/Tau Ratio
M38 plum1
Age+Sex-disc. p = 0.18
Sex-discounted p = 0.087
M38 plum1
Ambiguous
ANOVA p = 0.2
bicor=0.33, p=0.054
cor=0.36, p=0.034
bicor=-0.14, p=0.59
cor=-0.11, p=0.66
bicor=0.46, p=0.061
cor=0.45, p=0.07
bicor=0.15, p=0.4
cor=0.16, p=0.36
bicor=-0.16, p=0.35
cor=-0.15, p=0.39
bicor=0.19, p=0.29
cor=0.075, p=0.69
bicor=0.033, p=0.85
cor=-0.038, p=0.83
bicor=-0.12, p=0.48
cor=-0.19, p=0.27
bicor=0.14, p=0.43
cor=0.14, p=0.42
bicor=0.037, p=0.83
cor=0.061, p=0.73
Eigenprotein

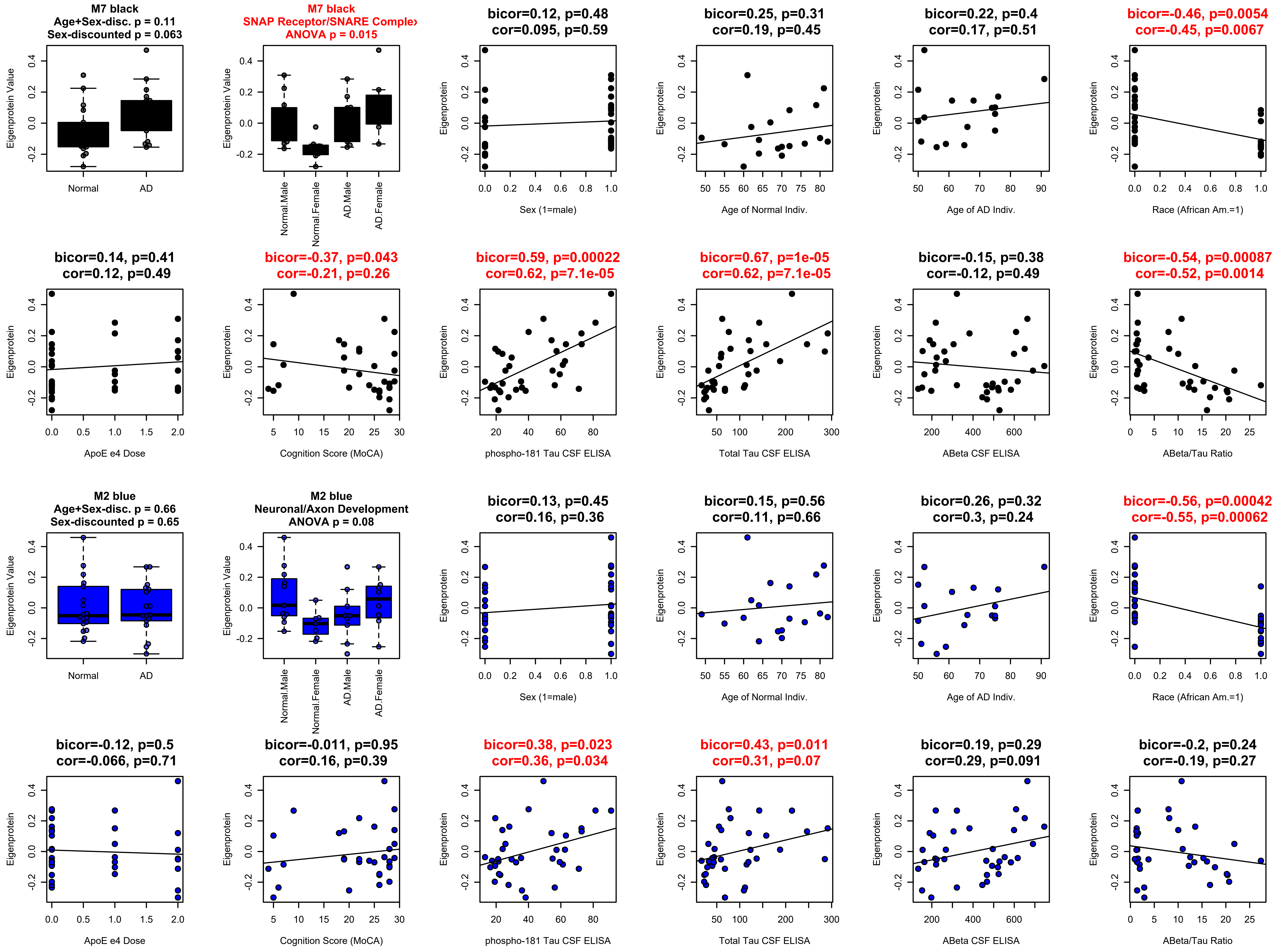

M7 black
Age+Sex-disc. p = 0.11
Sex-discounted p = 0.063
M7 black
SNAP Receptor/SNARE Complex
ANOVA p = 0.015
bicor=0.12, p=0.48
cor=0.095, p=0.59
bicor=0.25, p=0.31
cor=0.19, p=0.45
bicor=0.22, p=0.4
cor=0.17, p=0.51
bicor=-0.46, p=0.0054
cor=-0.45, p=0.0067
Eigenprotein Value
Eigenprotein
Normal
AD
Normal.Male
Normal.Female
AD.Male
AD.Female
Sex (1=male)
Age of Normal Indiv.
Age of AD Indiv.
Race (African Am.=1)
bicor=0.14, p=0.41
cor=0.12, p=0.49
bicor=-0.37, p=0.043
cor=-0.21, p=0.26
bicor=0.59, p=0.00022
cor=0.62, p=7.1e-05
bicor=0.67, p=1e-05
cor=0.62, p=7.1e-05
bicor=-0.15, p=0.38
cor=-0.12, p=0.49
bicor=-0.54, p=0.00087
cor=-0.52, p=0.0014
ApoE e4 Dose
Cognition Score (MoCA)
phospho-181 Tau CSF ELISA
Total Tau CSF ELISA
ABeta CSF ELISA
ABeta/Tau Ratio
M2 blue
Age+Sex-disc. p = 0.66
Sex-discounted p = 0.65
M2 blue
Neuronal/Axon Development
ANOVA p = 0.08
bicor=0.13, p=0.45
cor=0.16, p=0.36
bicor=0.15, p=0.56
cor=0.11, p=0.66
bicor=0.26, p=0.32
cor=0.3, p=0.24
bicor=-0.56, p=0.00042
cor=-0.55, p=0.00062
bicor=-0.12, p=0.5
cor=-0.066, p=0.71
bicor=-0.011, p=0.95
cor=0.16, p=0.39
bicor=0.38, p=0.023
cor=0.36, p=0.034
bicor=0.43, p=0.011
cor=0.31, p=0.07
bicor=0.19, p=0.29
cor=0.29, p=0.091
bicor=-0.2, p=0.24
cor=-0.19, p=0.27

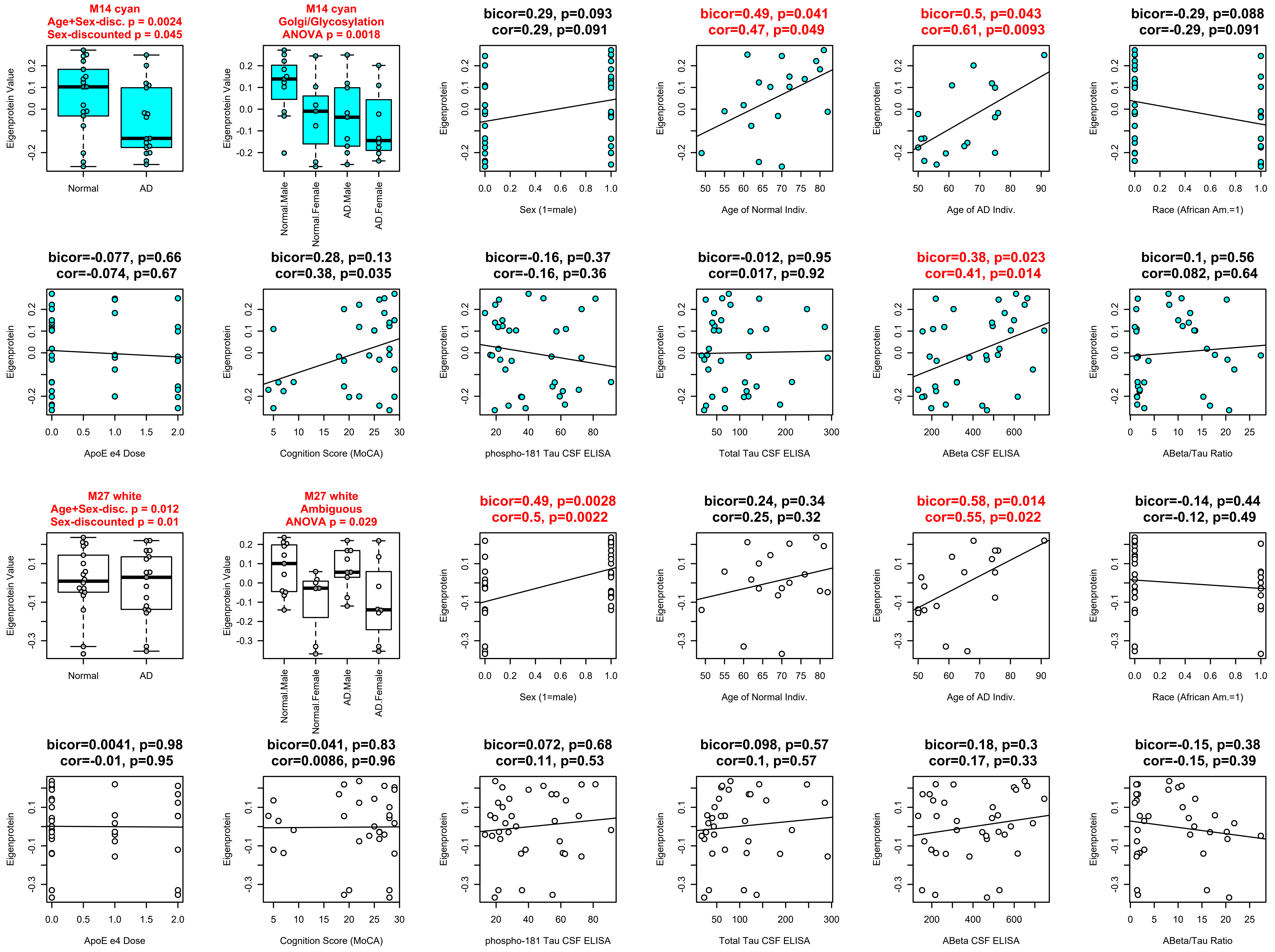
M14 cyan
Age+Sex-disc. p = 0.0024
Sex-discounted p = 0.045
Eigenprotein Value
Normal
AD
M14 cyan
Golgi/Glycosylation
ANOVA p = 0.0018
Eigenprotein Value
Normal.Male
Normal.Female
AD.Male
AD.Female
bicor=0.29, p=0.093
cor=0.29, p=0.091
Eigenprotein
Sex (1=male)
bicor=0.49, p=0.041
cor=0.47, p=0.049
Eigenprotein
Age of Normal Indiv.
bicor=0.5, p=0.043
cor=0.61, p=0.0093
Eigenprotein
Age of AD Indiv.
bicor=-0.29, p=0.088
cor=-0.29, p=0.091
Eigenprotein
Race (African Am.=1)
bicor=-0.077, p=0.66
cor=-0.074, p=0.67
Eigenprotein
ApoE e4 Dose
bicor=0.28, p=0.13
cor=0.38, p=0.035
Eigenprotein
Cognition Score (MoCA)
bicor=-0.16, p=0.37
cor=-0.16, p=0.36
Eigenprotein
phospho-181 Tau CSF ELISA
bicor=-0.012, p=0.95
cor=0.017, p=0.92
Eigenprotein
Total Tau CSF ELISA
bicor=0.38, p=0.023
cor=0.41, p=0.014
Eigenprotein
ABeta CSF ELISA
bicor=0.1, p=0.56
cor=0.082, p=0.64
Eigenprotein
ABeta/Tau Ratio
M27 white
Age+Sex-disc. p = 0.012
Sex-discounted p = 0.01
Eigenprotein Value
Normal
AD
M27 white
Ambiguous
ANOVA p = 0.029
Eigenprotein Value
Normal.Male
Normal.Female
AD.Male
AD.Female
bicor=0.49, p=0.0028
cor=0.5, p=0.0022
Eigenprotein
Sex (1=male)
bicor=0.24, p=0.34
cor=0.25, p=0.32
Eigenprotein
Age of Normal Indiv.
bicor=0.58, p=0.014
cor=0.55, p=0.022
Eigenprotein
Age of AD Indiv.
bicor=-0.14, p=0.44
cor=-0.12, p=0.49
Eigenprotein
Race (African Am.=1)
bicor=0.0041, p=0.98
cor=-0.01, p=0.95
Eigenprotein
ApoE e4 Dose
bicor=0.041, p=0.83
cor=0.0086, p=0.96
Eigenprotein
Cognition Score (MoCA)
bicor=0.072, p=0.68
cor=0.11, p=0.53
Eigenprotein
phospho-181 Tau CSF ELISA
bicor=0.098, p=0.57
cor=0.1, p=0.57
Eigenprotein
Total Tau CSF ELISA
bicor=0.18, p=0.3
cor=0.17, p=0.33
Eigenprotein
ABeta CSF ELISA
bicor=-0.15, p=0.38
cor=-0.15, p=0.39
Eigenprotein
ABeta/Tau Ratio

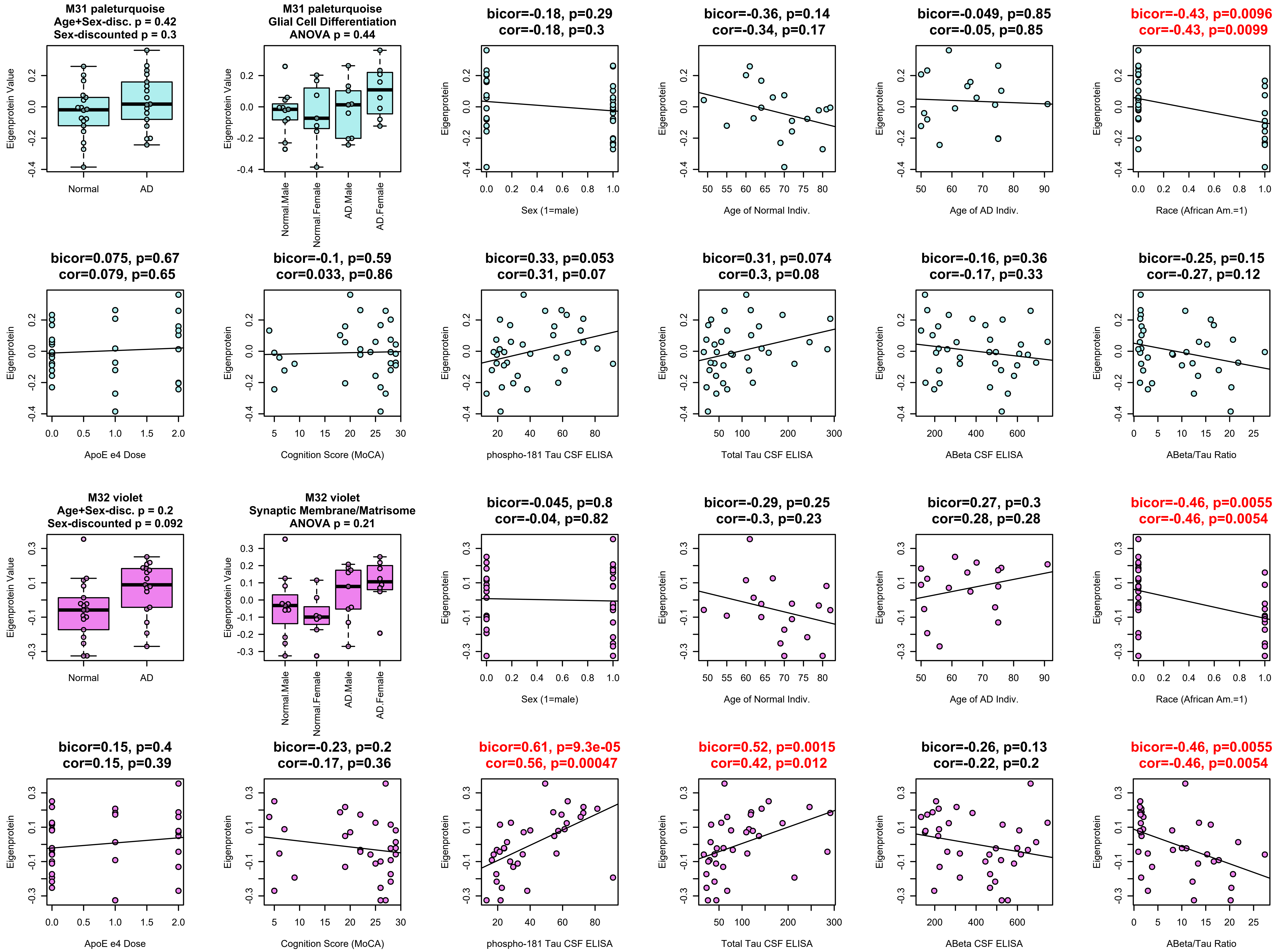
M31 paleturquoise
Age+Sex-disc. p = 0.42
Sex-discounted p = 0.3
Eigenprotein Value
Normal
AD
M31 paleturquoise
Glial Cell Differentiation
ANOVA p = 0.44
Eigenprotein Value
Normal.Male
Normal.Female
AD.Male
AD.Female
bicor=-0.18, p=0.29
cor=-0.18, p=0.3
Eigenprotein
Sex (1=male)
bicor=-0.36, p=0.14
cor=-0.34, p=0.17
Eigenprotein
Age of Normal Indiv.
bicor=-0.049, p=0.85
cor=-0.05, p=0.85
Eigenprotein
Age of AD Indiv.
bicor=-0.43, p=0.0096
cor=-0.43, p=0.0099
Eigenprotein
Race (African Am.=1)
bicor=0.075, p=0.67
cor=0.079, p=0.65
Eigenprotein
ApoE e4 Dose
bicor=-0.1, p=0.59
cor=0.033, p=0.86
Eigenprotein
Cognition Score (MoCA)
bicor=0.33, p=0.053
cor=0.31, p=0.07
Eigenprotein
phospho-181 Tau CSF ELISA
bicor=0.31, p=0.074
cor=0.3, p=0.08
Eigenprotein
Total Tau CSF ELISA
bicor=-0.16, p=0.36
cor=-0.17, p=0.33
Eigenprotein
ABeta CSF ELISA
bicor=-0.25, p=0.15
cor=-0.27, p=0.12
Eigenprotein
ABeta/Tau Ratio
M32 violet
Age+Sex-disc. p = 0.2
Sex-discounted p = 0.092
Eigenprotein Value
Normal
AD
M32 violet
Synaptic Membrane/Matrisome
ANOVA p = 0.21
Eigenprotein Value
Normal.Male
Normal.Female
AD.Male
AD.Female
bicor=-0.045, p=0.8
cor=-0.04, p=0.82
Eigenprotein
Sex (1=male)
bicor=-0.29, p=0.25
cor=-0.3, p=0.23
Eigenprotein
Age of Normal Indiv.
bicor=0.27, p=0.3
cor=0.28, p=0.28
Eigenprotein
Age of AD Indiv.
bicor=-0.46, p=0.0055
cor=-0.46, p=0.0054
Eigenprotein
Race (African Am.=1)
bicor=0.15, p=0.4
cor=0.15, p=0.39
Eigenprotein
ApoE e4 Dose
bicor=-0.23, p=0.2
cor=-0.17, p=0.36
Eigenprotein
Cognition Score (MoCA)
bicor=0.61, p=9.3e-05
cor=0.56, p=0.00047
Eigenprotein
phospho-181 Tau CSF ELISA
bicor=0.52, p=0.0015
cor=0.42, p=0.012
Eigenprotein
Total Tau CSF ELISA
bicor=-0.26, p=0.13
cor=-0.22, p=0.2
Eigenprotein
ABeta CSF ELISA
bicor=-0.46, p=0.0055
cor=-0.46, p=0.0054
Eigenprotein
ABeta/Tau Ratio

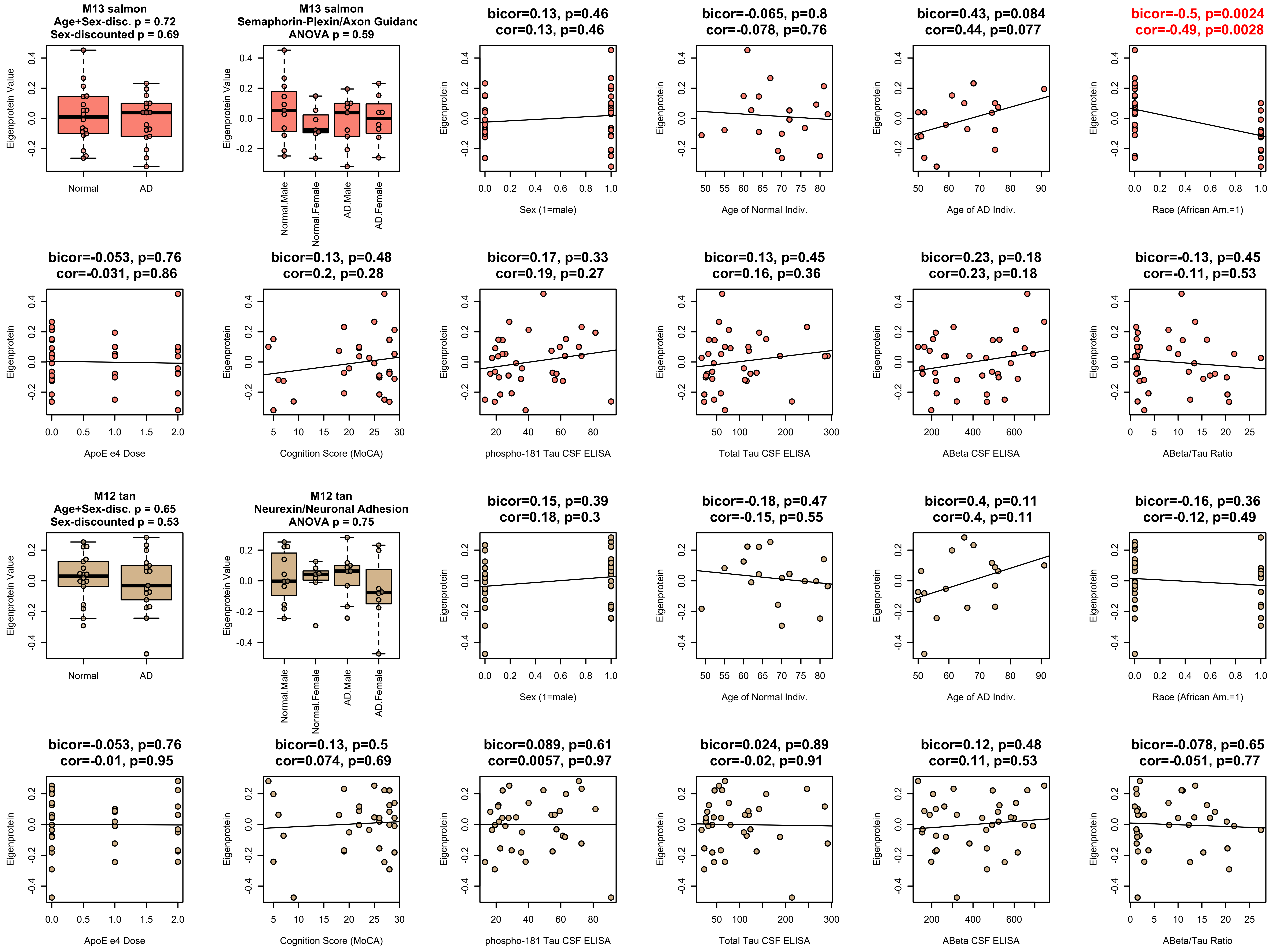
M13 salmon
Age+Sex-disc. p = 0.72
Sex-discounted p = 0.69
Eigenprotein Value
Normal
AD
M13 salmon
Semaphorin-Plexin/Axon Guidanc
ANOVA p = 0.59
Eigenprotein Value
Normal.Male
Normal.Female
AD.Male
AD.Female
bicor=0.13, p=0.46
cor=0.13, p=0.46
Eigenprotein
Sex (1=male)
bicor=-0.065, p=0.8
cor=-0.078, p=0.76
Eigenprotein
Age of Normal Indiv.
bicor=0.43, p=0.084
cor=0.44, p=0.077
Eigenprotein
Age of AD Indiv.
bicor=-0.5, p=0.0024
cor=-0.49, p=0.0028
Eigenprotein
Race (African Am.=1)
bicor=-0.053, p=0.76
cor=-0.031, p=0.86
Eigenprotein
ApoE e4 Dose
bicor=0.13, p=0.48
cor=0.2, p=0.28
Eigenprotein
Cognition Score (MoCA)
bicor=0.17, p=0.33
cor=0.19, p=0.27
Eigenprotein
phospho-181 Tau CSF ELISA
bicor=0.13, p=0.45
cor=0.16, p=0.36
Eigenprotein
Total Tau CSF ELISA
bicor=0.23, p=0.18
cor=0.23, p=0.18
Eigenprotein
ABeta CSF ELISA
bicor=-0.13, p=0.45
cor=-0.11, p=0.53
Eigenprotein
ABeta/Tau Ratio
M12 tan
Age+Sex-disc. p = 0.65
Sex-discounted p = 0.53
Eigenprotein Value
Normal
AD
M12 tan
Neurexin/Neuronal Adhesion
ANOVA p = 0.75
Eigenprotein Value
Normal.Male
Normal.Female
AD.Male
AD.Female
bicor=0.15, p=0.39
cor=0.18, p=0.3
Eigenprotein
Sex (1=male)
bicor=-0.18, p=0.47
cor=-0.15, p=0.55
Eigenprotein
Age of Normal Indiv.
bicor=0.4, p=0.11
cor=0.4, p=0.11
Eigenprotein
Age of AD Indiv.
bicor=-0.16, p=0.36
cor=-0.12, p=0.49
Eigenprotein
Race (African Am.=1)
bicor=-0.053, p=0.76
cor=-0.01, p=0.95
Eigenprotein
ApoE e4 Dose
bicor=0.13, p=0.5
cor=0.074, p=0.69
Eigenprotein
Cognition Score (MoCA)
bicor=0.089, p=0.61
cor=0.0057, p=0.97
Eigenprotein
phospho-181 Tau CSF ELISA
bicor=0.024, p=0.89
cor=-0.02, p=0.91
Eigenprotein
Total Tau CSF ELISA
bicor=0.12, p=0.48
cor=0.11, p=0.53
Eigenprotein
ABeta CSF ELISA
bicor=-0.078, p=0.65
cor=-0.051, p=0.77
Eigenprotein
ABeta/Tau Ratio

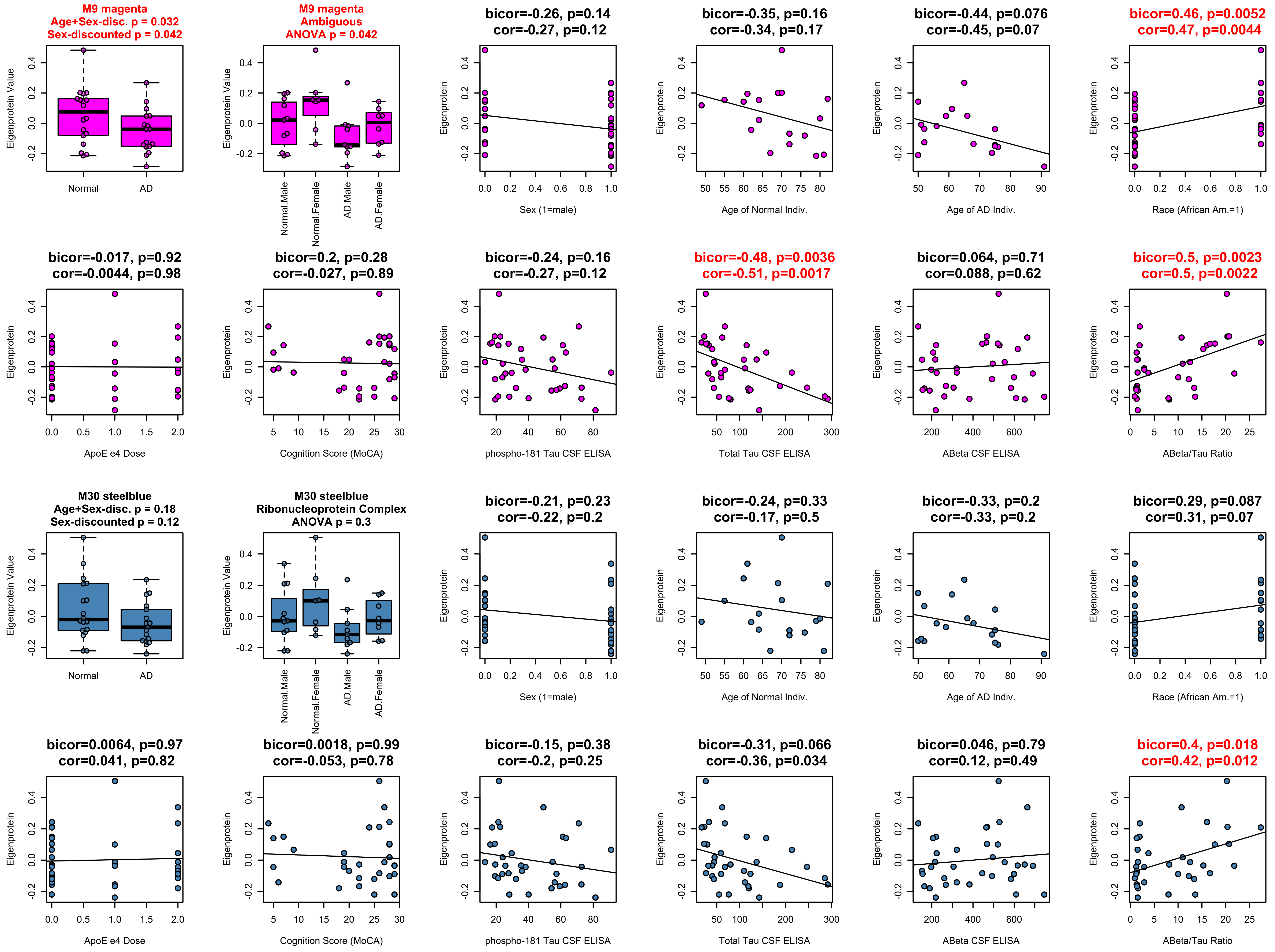
M9 magenta
Age+Sex-disc. p = 0.032
Sex-discounted p = 0.042
Eigenprotein Value
Normal
AD
M9 magenta
Ambiguous
ANOVA p = 0.042
Eigenprotein Value
Normal.Male
Normal.Female
AD.Male
AD.Female
bicor=-0.26, p=0.14
cor=-0.27, p=0.12
Eigenprotein
Sex (1=male)
bicor=-0.35, p=0.16
cor=-0.34, p=0.17
Eigenprotein
Age of Normal Indiv.
bicor=-0.44, p=0.076
cor=-0.45, p=0.07
Eigenprotein
Age of AD Indiv.
bicor=0.46, p=0.0052
cor=0.47, p=0.0044
Eigenprotein
Race (African Am.=1)
bicor=-0.017, p=0.92
cor=-0.0044, p=0.98
Eigenprotein
ApoE e4 Dose
bicor=0.2, p=0.28
cor=-0.027, p=0.89
Eigenprotein
Cognition Score (MoCA)
bicor=-0.24, p=0.16
cor=-0.27, p=0.12
Eigenprotein
phospho-181 Tau CSF ELISA
bicor=-0.48, p=0.0036
cor=-0.51, p=0.0017
Eigenprotein
Total Tau CSF ELISA
bicor=0.064, p=0.71
cor=0.088, p=0.62
Eigenprotein
ABeta CSF ELISA
bicor=0.5, p=0.0023
cor=0.5, p=0.0022
Eigenprotein
ABeta/Tau Ratio
M30 steelblue
Age+Sex-disc. p = 0.18
Sex-discounted p = 0.12
Eigenprotein Value
Normal
AD
M30 steelblue
Ribonucleoprotein Complex
ANOVA p = 0.3
Eigenprotein Value
Normal.Male
Normal.Female
AD.Male
AD.Female
bicor=-0.21, p=0.23
cor=-0.22, p=0.2
Eigenprotein
Sex (1=male)
bicor=-0.24, p=0.33
cor=-0.17, p=0.5
Eigenprotein
Age of Normal Indiv.
bicor=-0.33, p=0.2
cor=-0.33, p=0.2
Eigenprotein
Age of AD Indiv.
bicor=0.29, p=0.087
cor=0.31, p=0.07
Eigenprotein
Race (African Am.=1)
bicor=0.0064, p=0.97
cor=0.041, p=0.82
Eigenprotein
ApoE e4 Dose
bicor=0.0018, p=0.99
cor=-0.053, p=0.78
Eigenprotein
Cognition Score (MoCA)
bicor=-0.15, p=0.38
cor=-0.2, p=0.25
Eigenprotein
phospho-181 Tau CSF ELISA
bicor=-0.31, p=0.066
cor=-0.36, p=0.034
Eigenprotein
Total Tau CSF ELISA
bicor=0.046, p=0.79
cor=0.12, p=0.49
Eigenprotein
ABeta CSF ELISA
bicor=0.4, p=0.018
cor=0.42, p=0.012
Eigenprotein
ABeta/Tau Ratio

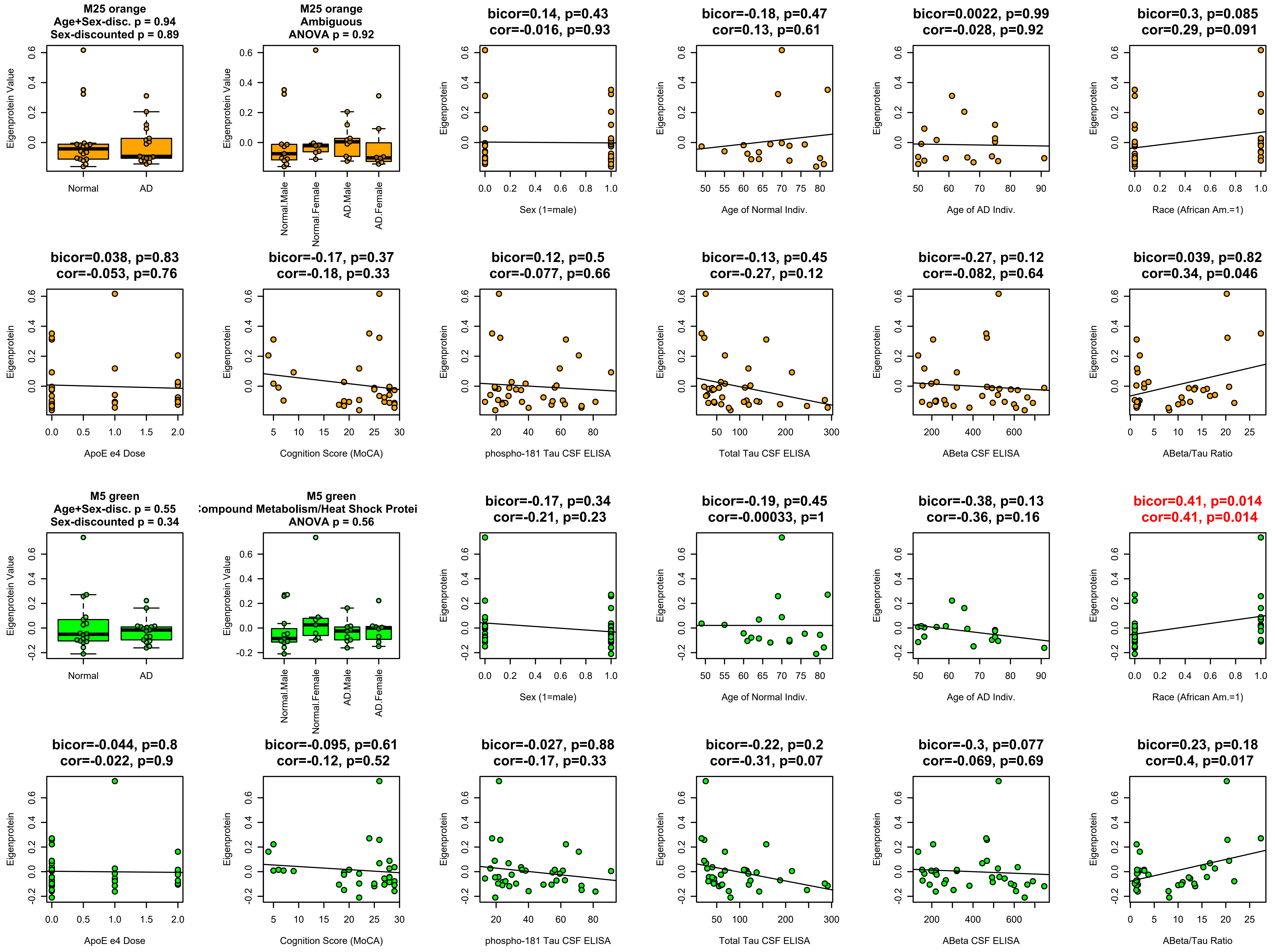

M25 orange
Age+Sex-disc. p = 0.94
Sex-discounted p = 0.89
Eigenprotein Value
Normal
AD
M25 orange
Ambiguous
ANOVA p = 0.92
Eigenprotein Value
Normal.Male
Normal.Female
AD.Male
AD.Female
bicor=0.14, p=0.43
cor=-0.016, p=0.93
Eigenprotein
Sex (1=male)
bicor=-0.18, p=0.47
cor=0.13, p=0.61
Eigenprotein
Age of Normal Indiv.
bicor=0.0022, p=0.99
cor=-0.028, p=0.92
Eigenprotein
Age of AD Indiv.
bicor=0.3, p=0.085
cor=0.29, p=0.091
Eigenprotein
Race (African Am.=1)
bicor=0.038, p=0.83
cor=-0.053, p=0.76
Eigenprotein
ApoE e4 Dose
bicor=-0.17, p=0.37
cor=-0.18, p=0.33
Eigenprotein
Cognition Score (MoCA)
bicor=0.12, p=0.5
cor=-0.077, p=0.66
Eigenprotein
phospho-181 Tau CSF ELISA
bicor=-0.13, p=0.45
cor=-0.27, p=0.12
Eigenprotein
Total Tau CSF ELISA
bicor=-0.27, p=0.12
cor=-0.082, p=0.64
Eigenprotein
ABeta CSF ELISA
bicor=0.039, p=0.82
cor=0.34, p=0.046
Eigenprotein
ABeta/Tau Ratio
M5 green
Age+Sex-disc. p = 0.55
Sex-discounted p = 0.34
Eigenprotein Value
Normal
AD
M5 green
ompound Metabolism/Heat Shock Protei
ANOVA p = 0.56
Eigenprotein Value
Normal.Male
Normal.Female
AD.Male
AD.Female
bicor=-0.17, p=0.34
cor=-0.21, p=0.23
Eigenprotein
Sex (1=male)
bicor=-0.19, p=0.45
cor=-0.00033, p=1
Eigenprotein
Age of Normal Indiv.
bicor=-0.38, p=0.13
cor=-0.36, p=0.16
Eigenprotein
Age of AD Indiv.
bicor=0.41, p=0.014
cor=0.41, p=0.014
Eigenprotein
Race (African Am.=1)
bicor=-0.044, p=0.8
cor=-0.022, p=0.9
Eigenprotein
ApoE e4 Dose
bicor=-0.095, p=0.61
cor=-0.12, p=0.52
Eigenprotein
Cognition Score (MoCA)
bicor=-0.027, p=0.88
cor=-0.17, p=0.33
Eigenprotein
phospho-181 Tau CSF ELISA
bicor=-0.22, p=0.2
cor=-0.31, p=0.07
Eigenprotein
Total Tau CSF ELISA
bicor=-0.3, p=0.077
cor=-0.069, p=0.69
Eigenprotein
ABeta CSF ELISA
bicor=0.23, p=0.18
cor=0.4, p=0.017
Eigenprotein
ABeta/Tau Ratio

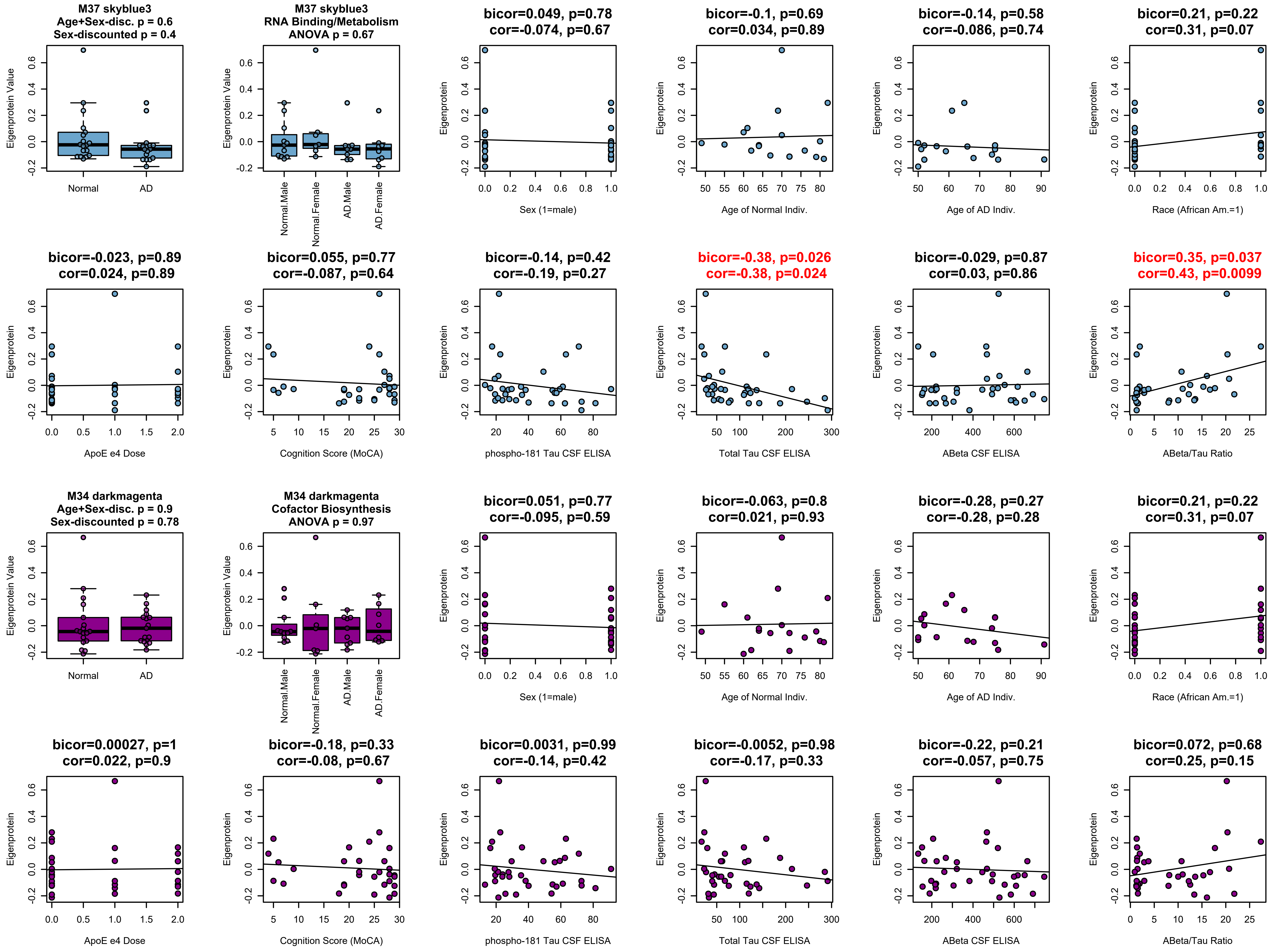
M37 skyblue3
Age+Sex-disc. p = 0.6
Sex-discounted p = 0.4
Eigenprotein Value
Normal
AD
M37 skyblue3
RNA Binding/Metabolism
ANOVA p = 0.67
Eigenprotein Value
Normal.Male
Normal.Female
AD.Male
AD.Female
bicor=0.049, p=0.78
cor=-0.074, p=0.67
Eigenprotein
Sex (1=male)
bicor=-0.1, p=0.69
cor=0.034, p=0.89
Eigenprotein
Age of Normal Indiv.
bicor=-0.14, p=0.58
cor=-0.086, p=0.74
Eigenprotein
Age of AD Indiv.
bicor=0.21, p=0.22
cor=0.31, p=0.07
Eigenprotein
Race (African Am.=1)
bicor=-0.023, p=0.89
cor=0.024, p=0.89
Eigenprotein
ApoE e4 Dose
bicor=0.055, p=0.77
cor=-0.087, p=0.64
Eigenprotein
Cognition Score (MoCA)
bicor=-0.14, p=0.42
cor=-0.19, p=0.27
Eigenprotein
phospho-181 Tau CSF ELISA
bicor=-0.38, p=0.026
cor=-0.38, p=0.024
Eigenprotein
Total Tau CSF ELISA
bicor=-0.029, p=0.87
cor=0.03, p=0.86
Eigenprotein
ABeta CSF ELISA
bicor=0.35, p=0.037
cor=0.43, p=0.0099
Eigenprotein
ABeta/Tau Ratio
M34 darkmagenta
Age+Sex-disc. p = 0.9
Sex-discounted p = 0.78
Eigenprotein Value
Normal
AD
M34 darkmagenta
Cofactor Biosynthesis
ANOVA p = 0.97
Eigenprotein Value
Normal.Male
Normal.Female
AD.Male
AD.Female
bicor=0.051, p=0.77
cor=-0.095, p=0.59
Eigenprotein
Sex (1=male)
bicor=-0.063, p=0.8
cor=0.021, p=0.93
Eigenprotein
Age of Normal Indiv.
bicor=-0.28, p=0.27
cor=-0.28, p=0.28
Eigenprotein
Age of AD Indiv.
bicor=0.21, p=0.22
cor=0.31, p=0.07
Eigenprotein
Race (African Am.=1)
bicor=0.00027, p=1
cor=0.022, p=0.9
Eigenprotein
ApoE e4 Dose
bicor=-0.18, p=0.33
cor=-0.08, p=0.67
Eigenprotein
Cognition Score (MoCA)
bicor=0.0031, p=0.99
cor=-0.14, p=0.42
Eigenprotein
phospho-181 Tau CSF ELISA
bicor=-0.0052, p=0.98
cor=-0.17, p=0.33
Eigenprotein
Total Tau CSF ELISA
bicor=-0.22, p=0.21
cor=-0.057, p=0.75
Eigenprotein
ABeta CSF ELISA
bicor=0.072, p=0.68
cor=0.25, p=0.15
Eigenprotein
ABeta/Tau Ratio

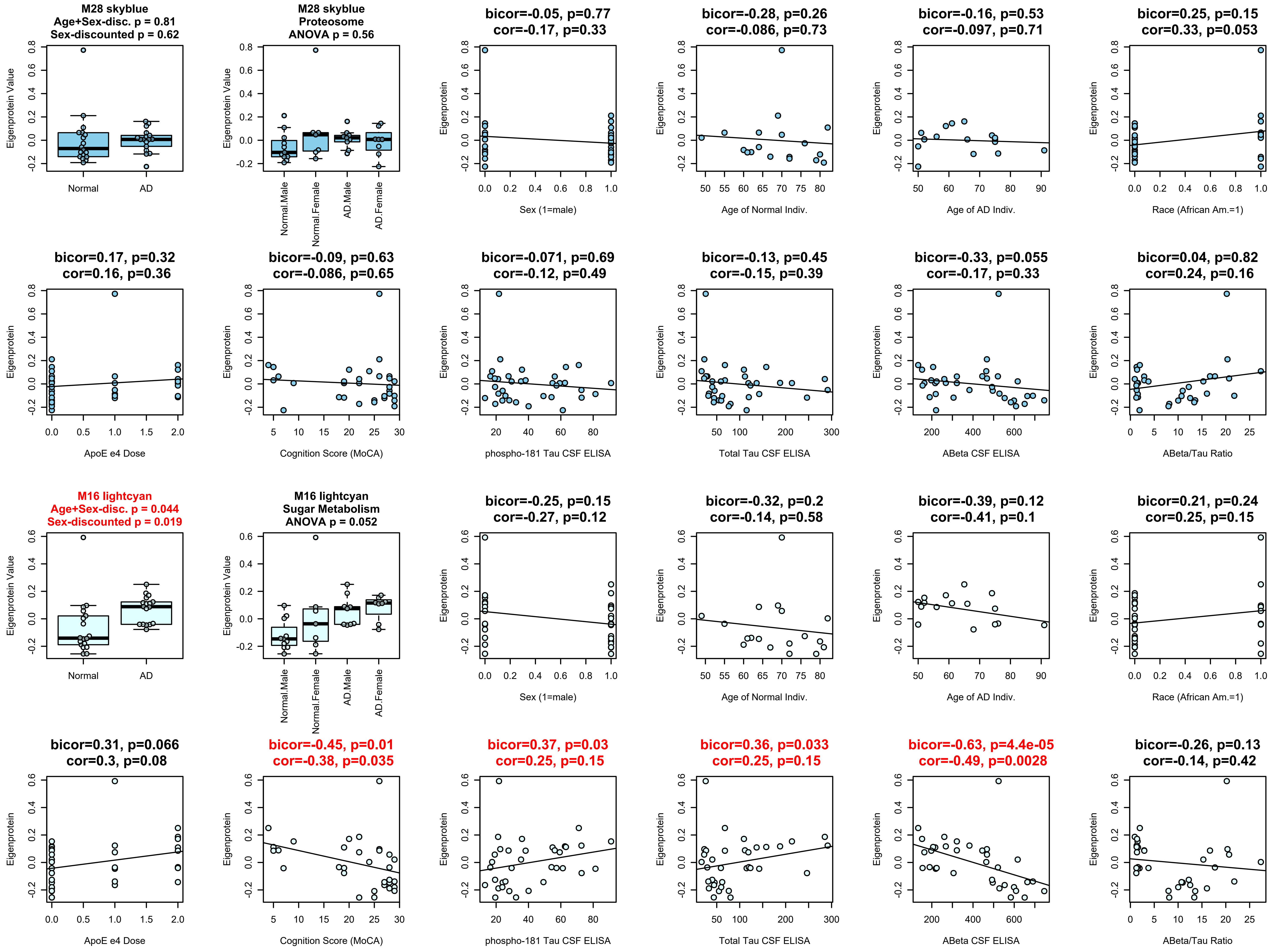
M28 skyblue
Age+Sex-disc. p = 0.81
Sex-discounted p = 0.62
Eigenprotein Value
Normal
AD
M28 skyblue
Proteosome
ANOVA p = 0.56
Eigenprotein Value
Normal.Male
Normal.Female
AD.Male
AD.Female
bicor=-0.05, p=0.77
cor=-0.17, p=0.33
Eigenprotein
Sex (1=male)
bicor=-0.28, p=0.26
cor=-0.086, p=0.73
Eigenprotein
Age of Normal Indiv.
bicor=-0.16, p=0.53
cor=-0.097, p=0.71
Eigenprotein
Age of AD Indiv.
bicor=0.25, p=0.15
cor=0.33, p=0.053
Eigenprotein
Race (African Am.=1)
bicor=0.17, p=0.32
cor=0.16, p=0.36
Eigenprotein
ApoE e4 Dose
bicor=-0.09, p=0.63
cor=-0.086, p=0.65
Eigenprotein
Cognition Score (MoCA)
bicor=-0.071, p=0.69
cor=-0.12, p=0.49
Eigenprotein
phospho-181 Tau CSF ELISA
bicor=-0.13, p=0.45
cor=-0.15, p=0.39
Eigenprotein
Total Tau CSF ELISA
bicor=-0.33, p=0.055
cor=-0.17, p=0.33
Eigenprotein
ABeta CSF ELISA
bicor=0.04, p=0.82
cor=0.24, p=0.16
Eigenprotein
ABeta/Tau Ratio
M16 lightcyan
Age+Sex-disc. p = 0.044
Sex-discounted p = 0.019
Eigenprotein Value
Normal
AD
M16 lightcyan
Sugar Metabolism
ANOVA p = 0.052
Eigenprotein Value
Normal.Male
Normal.Female
AD.Male
AD.Female
bicor=-0.25, p=0.15
cor=-0.27, p=0.12
Eigenprotein
Sex (1=male)
bicor=-0.32, p=0.2
cor=-0.14, p=0.58
Eigenprotein
Age of Normal Indiv.
bicor=-0.39, p=0.12
cor=-0.41, p=0.1
Eigenprotein
Age of AD Indiv.
bicor=0.21, p=0.24
cor=0.25, p=0.15
Eigenprotein
Race (African Am.=1)
bicor=0.31, p=0.066
cor=0.3, p=0.08
Eigenprotein
ApoE e4 Dose
bicor=-0.45, p=0.01
cor=-0.38, p=0.035
Eigenprotein
Cognition Score (MoCA)
bicor=0.37, p=0.03
cor=0.25, p=0.15
Eigenprotein
phospho-181 Tau CSF ELISA
bicor=0.36, p=0.033
cor=0.25, p=0.15
Eigenprotein
Total Tau CSF ELISA
bicor=-0.63, p=4.4e-05
cor=-0.49, p=0.0028
Eigenprotein
ABeta CSF ELISA
bicor=-0.26, p=0.13
cor=-0.14, p=0.42
Eigenprotein
ABeta/Tau Ratio

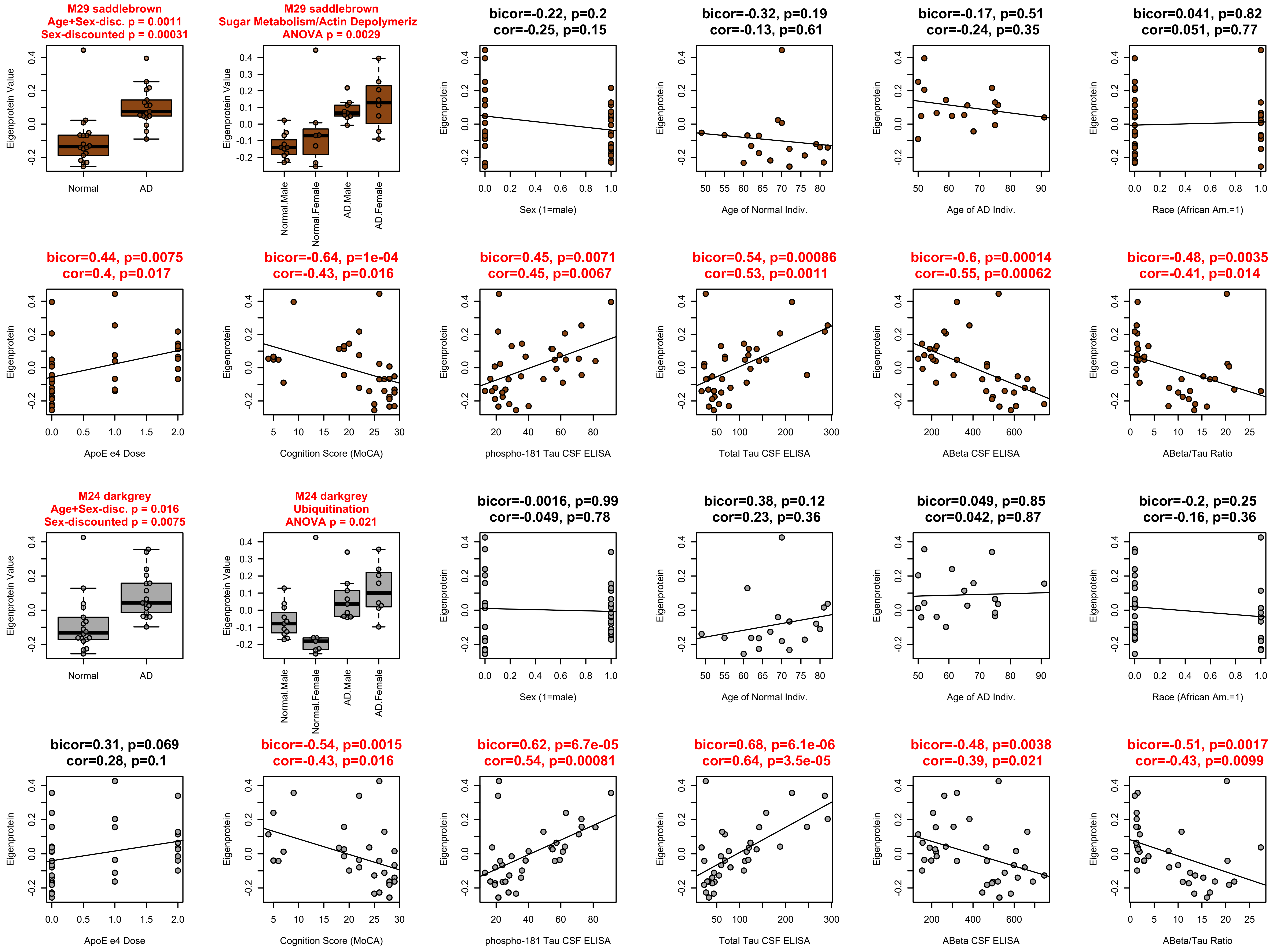
M29 saddlebrown
Age+Sex-disc. p = 0.0011
Sex-discounted p = 0.00031
Eigenprotein Value
Normal
AD
M29 saddlebrown
Sugar Metabolism/Actin Depolymeriz
ANOVA p = 0.0029
Eigenprotein Value
Normal.Male
Normal.Female
AD.Male
AD.Female
bicor=-0.22, p=0.2
cor=-0.25, p=0.15
Eigenprotein
Sex (1=male)
bicor=-0.32, p=0.19
cor=-0.13, p=0.61
Eigenprotein
Age of Normal Indiv.
bicor=-0.17, p=0.51
cor=-0.24, p=0.35
Eigenprotein
Age of AD Indiv.
bicor=0.041, p=0.82
cor=0.051, p=0.77
Eigenprotein
Race (African Am.=1)
bicor=0.44, p=0.0075
cor=0.4, p=0.017
Eigenprotein
ApoE e4 Dose
bicor=-0.64, p=1e-04
cor=-0.43, p=0.016
Eigenprotein
Cognition Score (MoCA)
bicor=0.45, p=0.0071
cor=0.45, p=0.0067
Eigenprotein
phospho-181 Tau CSF ELISA
bicor=0.54, p=0.00086
cor=0.53, p=0.0011
Eigenprotein
Total Tau CSF ELISA
bicor=-0.6, p=0.00014
cor=-0.55, p=0.00062
Eigenprotein
ABeta CSF ELISA
bicor=-0.48, p=0.0035
cor=-0.41, p=0.014
Eigenprotein
ABeta/Tau Ratio
M24 darkgrey
Age+Sex-disc. p = 0.016
Sex-discounted p = 0.0075
Eigenprotein Value
Normal
AD
M24 darkgrey
Ubiquitination
ANOVA p = 0.021
Eigenprotein Value
Normal.Male
Normal.Female
AD.Male
AD.Female
bicor=-0.0016, p=0.99
cor=-0.049, p=0.78
Eigenprotein
Sex (1=male)
bicor=0.38, p=0.12
cor=0.23, p=0.36
Eigenprotein
Age of Normal Indiv.
bicor=0.049, p=0.85
cor=0.042, p=0.87
Eigenprotein
Age of AD Indiv.
bicor=-0.2, p=0.25
cor=-0.16, p=0.36
Eigenprotein
Race (African Am.=1)
bicor=0.31, p=0.069
cor=0.28, p=0.1
Eigenprotein
ApoE e4 Dose
bicor=-0.54, p=0.0015
cor=-0.43, p=0.016
Eigenprotein
Cognition Score (MoCA)
bicor=0.62, p=6.7e-05
cor=0.54, p=0.00081
Eigenprotein
phospho-181 Tau CSF ELISA
bicor=0.68, p=6.1e-06
cor=0.64, p=3.5e-05
Eigenprotein
Total Tau CSF ELISA
bicor=-0.48, p=0.0038
cor=-0.39, p=0.021
Eigenprotein
ABeta CSF ELISA
bicor=-0.51, p=0.0017
cor=-0.43, p=0.0099
Eigenprotein
ABeta/Tau Ratio

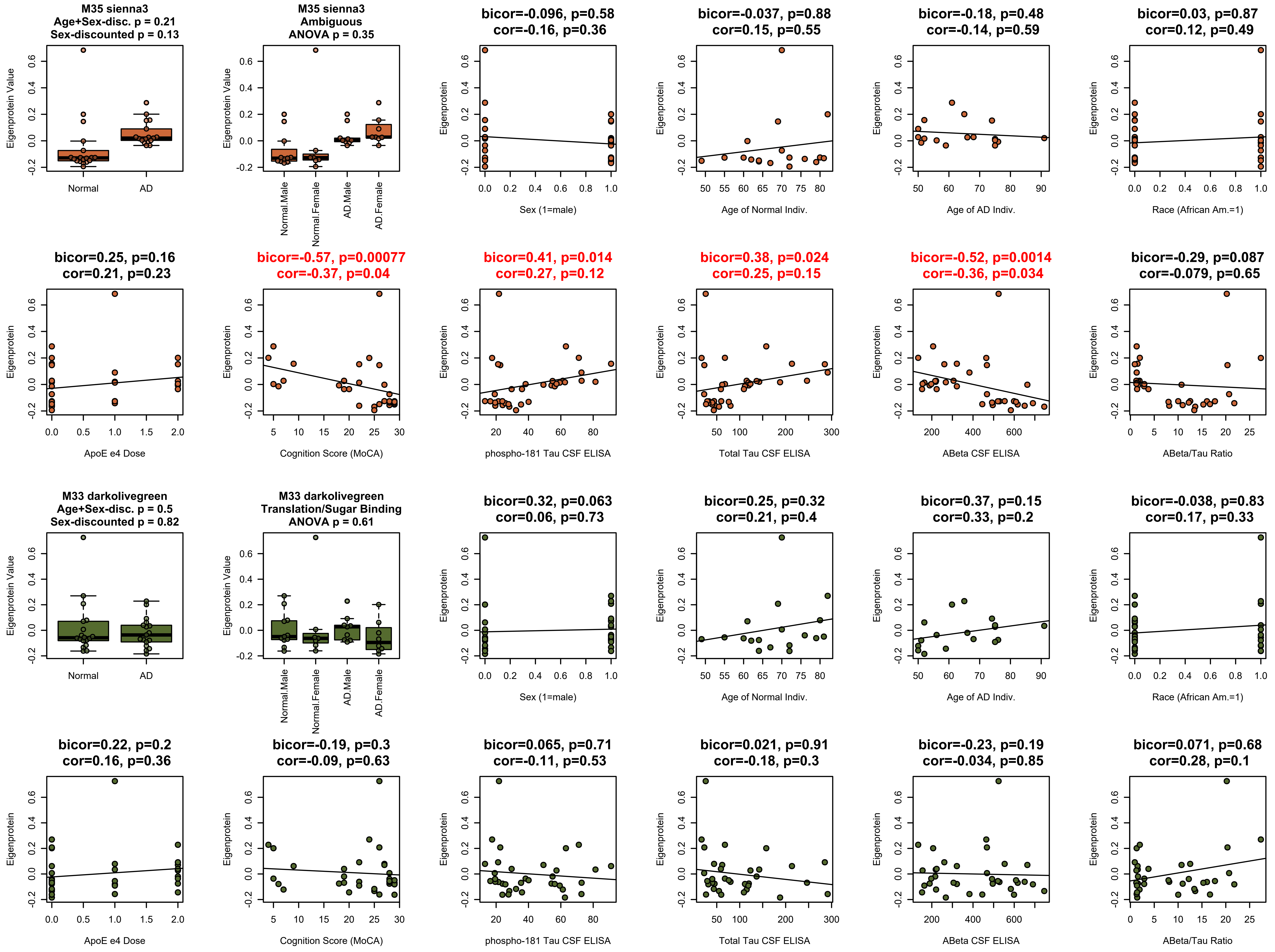

M35 sienna3
Age+Sex-disc. p = 0.21
Sex-discounted p = 0.13
Eigenprotein Value
Normal
AD
M35 sienna3
Ambiguous
ANOVA p = 0.35
Eigenprotein Value
Normal.Male
Normal.Female
AD.Male
AD.Female
bicor=-0.096, p=0.58
cor=-0.16, p=0.36
Eigenprotein
Sex (1=male)
bicor=-0.037, p=0.88
cor=0.15, p=0.55
Eigenprotein
Age of Normal Indiv.
bicor=-0.18, p=0.48
cor=-0.14, p=0.59
Eigenprotein
Age of AD Indiv.
bicor=0.03, p=0.87
cor=0.12, p=0.49
Eigenprotein
Race (African Am.=1)
bicor=0.25, p=0.16
cor=0.21, p=0.23
Eigenprotein
ApoE e4 Dose
bicor=-0.57, p=0.00077
cor=-0.37, p=0.04
Eigenprotein
Cognition Score (MoCA)
bicor=0.41, p=0.014
cor=0.27, p=0.12
Eigenprotein
phospho-181 Tau CSF ELISA
bicor=0.38, p=0.024
cor=0.25, p=0.15
Eigenprotein
Total Tau CSF ELISA
bicor=-0.52, p=0.0014
cor=-0.36, p=0.034
Eigenprotein
ABeta CSF ELISA
bicor=-0.29, p=0.087
cor=-0.079, p=0.65
Eigenprotein
ABeta/Tau Ratio
M33 darkolivegreen
Age+Sex-disc. p = 0.5
Sex-discounted p = 0.82
Eigenprotein Value
Normal
AD
M33 darkolivegreen
Translation/Sugar Binding
ANOVA p = 0.61
Eigenprotein Value
Normal.Male
Normal.Female
AD.Male
AD.Female
bicor=0.32, p=0.063
cor=0.06, p=0.73
Eigenprotein
Sex (1=male)
bicor=0.25, p=0.32
cor=0.21, p=0.4
Eigenprotein
Age of Normal Indiv.
bicor=0.37, p=0.15
cor=0.33, p=0.2
Eigenprotein
Age of AD Indiv.
bicor=-0.038, p=0.83
cor=0.17, p=0.33
Eigenprotein
Race (African Am.=1)
bicor=0.22, p=0.2
cor=0.16, p=0.36
Eigenprotein
ApoE e4 Dose
bicor=-0.19, p=0.3
cor=-0.09, p=0.63
Eigenprotein
Cognition Score (MoCA)
bicor=0.065, p=0.71
cor=-0.11, p=0.53
Eigenprotein
phospho-181 Tau CSF ELISA
bicor=0.021, p=0.91
cor=-0.18, p=0.3
Eigenprotein
Total Tau CSF ELISA
bicor=-0.23, p=0.19
cor=-0.034, p=0.85
Eigenprotein
ABeta CSF ELISA
bicor=0.071, p=0.68
cor=0.28, p=0.1
Eigenprotein
ABeta/Tau Ratio

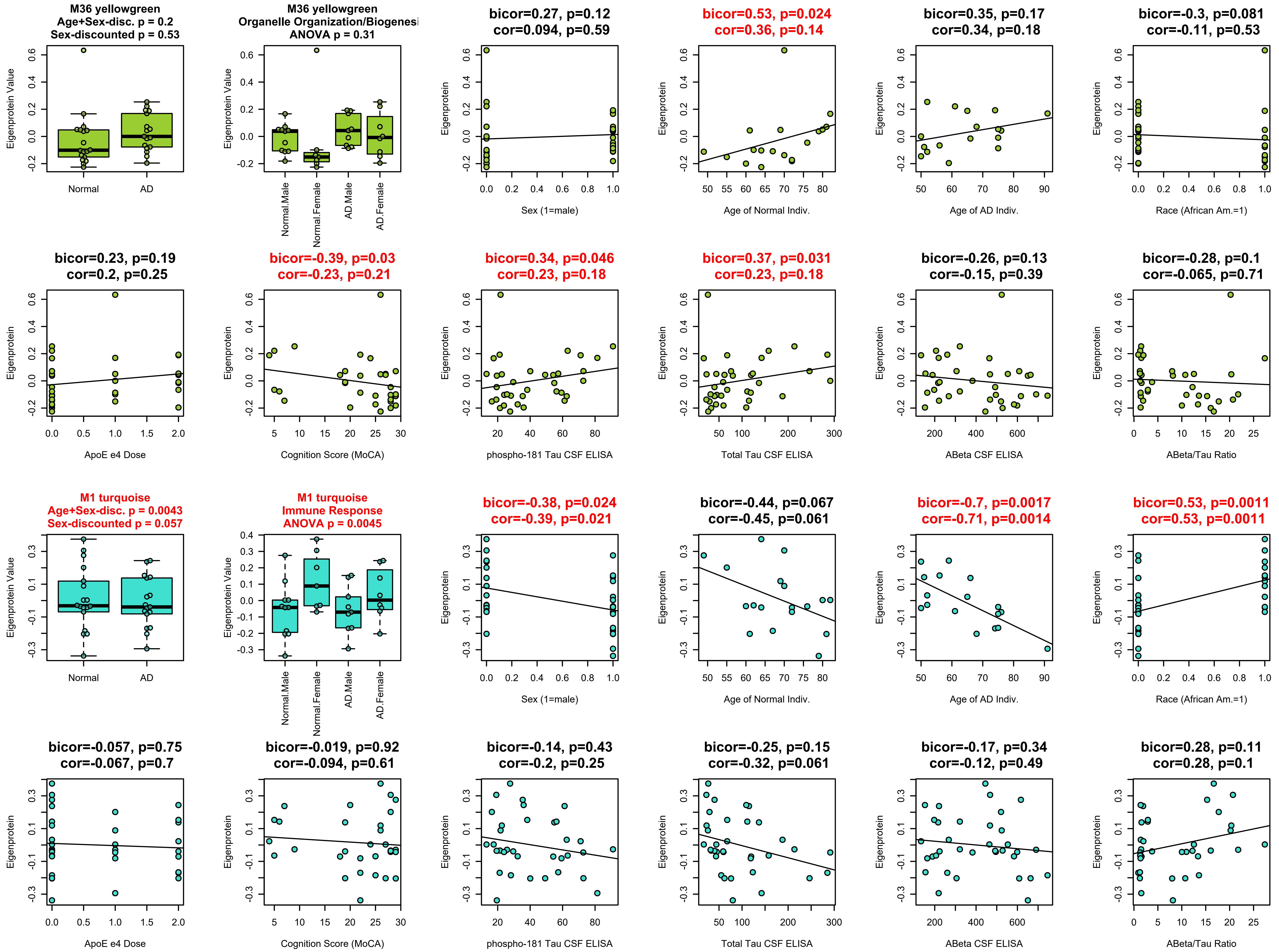
M36 yellowgreen
Age+Sex-disc. p = 0.2
Sex-discounted p = 0.53
Eigenprotein Value
Normal
AD
M36 yellowgreen
Organelle Organization/Biogenesi
ANOVA p = 0.31
Normal.Male
Normal.Female
AD.Male
AD.Female
bicor=0.27, p=0.12
cor=0.094, p=0.59
Sex (1=male)
bicor=0.53, p=0.024
cor=0.36, p=0.14
Age of Normal Indiv.
bicor=0.35, p=0.17
cor=0.34, p=0.18
Age of AD Indiv.
bicor=-0.3, p=0.081
cor=-0.11, p=0.53
Race (African Am.=1)
bicor=0.23, p=0.19
cor=0.2, p=0.25
ApoE e4 Dose
bicor=-0.39, p=0.03
cor=-0.23, p=0.21
Cognition Score (MoCA)
bicor=0.34, p=0.046
cor=0.23, p=0.18
phospho-181 Tau CSF ELISA
bicor=0.37, p=0.031
cor=0.23, p=0.18
Total Tau CSF ELISA
bicor=-0.26, p=0.13
cor=-0.15, p=0.39
ABeta CSF ELISA
bicor=-0.28, p=0.1
cor=-0.065, p=0.71
ABeta/Tau Ratio
Eigenprotein
M1 turquoise
Age+Sex-disc. p = 0.0043
Sex-discounted p = 0.057
Eigenprotein Value
Normal
AD
M1 turquoise
Immune Response
ANOVA p = 0.0045
Normal.Male
Normal.Female
AD.Male
AD.Female
bicor=-0.38, p=0.024
cor=-0.39, p=0.021
Sex (1=male)
bicor=-0.44, p=0.067
cor=-0.45, p=0.061
Age of Normal Indiv.
bicor=-0.7, p=0.0017
cor=-0.71, p=0.0014
Age of AD Indiv.
bicor=0.53, p=0.0011
cor=0.53, p=0.0011
Race (African Am.=1)
bicor=-0.057, p=0.75
cor=-0.067, p=0.7
ApoE e4 Dose
bicor=-0.019, p=0.92
cor=-0.094, p=0.61
Cognition Score (MoCA)
bicor=-0.14, p=0.43
cor=-0.2, p=0.25
phospho-181 Tau CSF ELISA
bicor=-0.25, p=0.15
cor=-0.32, p=0.061
Total Tau CSF ELISA
bicor=-0.17, p=0.34
cor=-0.12, p=0.49
ABeta CSF ELISA
bicor=0.28, p=0.11
cor=0.28, p=0.1
ABeta/Tau Ratio
Eigenprotein

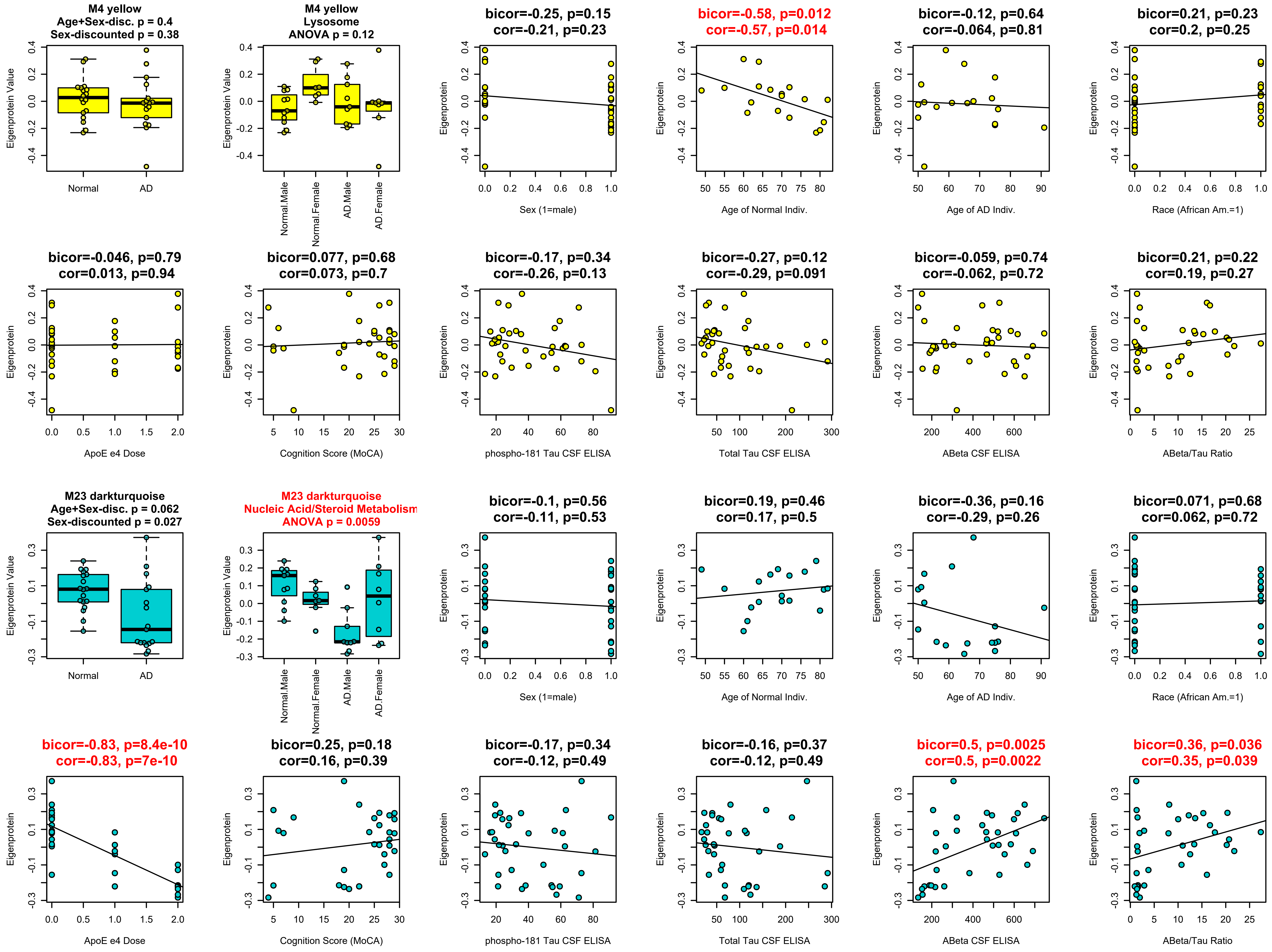
M4 yellow
Age+Sex-disc. p = 0.4
Sex-discounted p = 0.38
Eigenprotein Value
Normal
AD
M4 yellow
Lysosome
ANOVA p = 0.12
Eigenprotein Value
Normal.Male
Normal.Female
AD.Male
AD.Female
bicor=-0.25, p=0.15
cor=-0.21, p=0.23
Eigenprotein
Sex (1=male)
bicor=-0.58, p=0.012
cor=-0.57, p=0.014
Eigenprotein
Age of Normal Indiv.
bicor=-0.12, p=0.64
cor=-0.064, p=0.81
Eigenprotein
Age of AD Indiv.
bicor=0.21, p=0.23
cor=0.2, p=0.25
Eigenprotein
Race (African Am.=1)
bicor=-0.046, p=0.79
cor=0.013, p=0.94
Eigenprotein
ApoE e4 Dose
bicor=0.077, p=0.68
cor=0.073, p=0.7
Eigenprotein
Cognition Score (MoCA)
bicor=-0.17, p=0.34
cor=-0.26, p=0.13
Eigenprotein
phospho-181 Tau CSF ELISA
bicor=-0.27, p=0.12
cor=-0.29, p=0.091
Eigenprotein
Total Tau CSF ELISA
bicor=-0.059, p=0.74
cor=-0.062, p=0.72
Eigenprotein
ABeta CSF ELISA
bicor=0.21, p=0.22
cor=0.19, p=0.27
Eigenprotein
ABeta/Tau Ratio
M23 darkturquoise
Age+Sex-disc. p = 0.062
Sex-discounted p = 0.027
Eigenprotein Value
Normal
AD
M23 darkturquoise
Nucleic Acid/Steroid Metabolism
ANOVA p = 0.0059
Eigenprotein Value
Normal.Male
Normal.Female
AD.Male
AD.Female
bicor=-0.1, p=0.56
cor=-0.11, p=0.53
Eigenprotein
Sex (1=male)
bicor=0.19, p=0.46
cor=0.17, p=0.5
Eigenprotein
Age of Normal Indiv.
bicor=-0.36, p=0.16
cor=-0.29, p=0.26
Eigenprotein
Age of AD Indiv.
bicor=0.071, p=0.68
cor=0.062, p=0.72
Eigenprotein
Race (African Am.=1)
bicor=-0.83, p=8.4e-10
cor=-0.83, p=7e-10
Eigenprotein
ApoE e4 Dose
bicor=0.25, p=0.18
cor=0.16, p=0.39
Eigenprotein
Cognition Score (MoCA)
bicor=-0.17, p=0.34
cor=-0.12, p=0.49
Eigenprotein
phospho-181 Tau CSF ELISA
bicor=-0.16, p=0.37
cor=-0.12, p=0.49
Eigenprotein
Total Tau CSF ELISA
bicor=0.5, p=0.0025
cor=0.5, p=0.0022
Eigenprotein
ABeta CSF ELISA
bicor=0.36, p=0.036
cor=0.35, p=0.039
Eigenprotein
ABeta/Tau Ratio

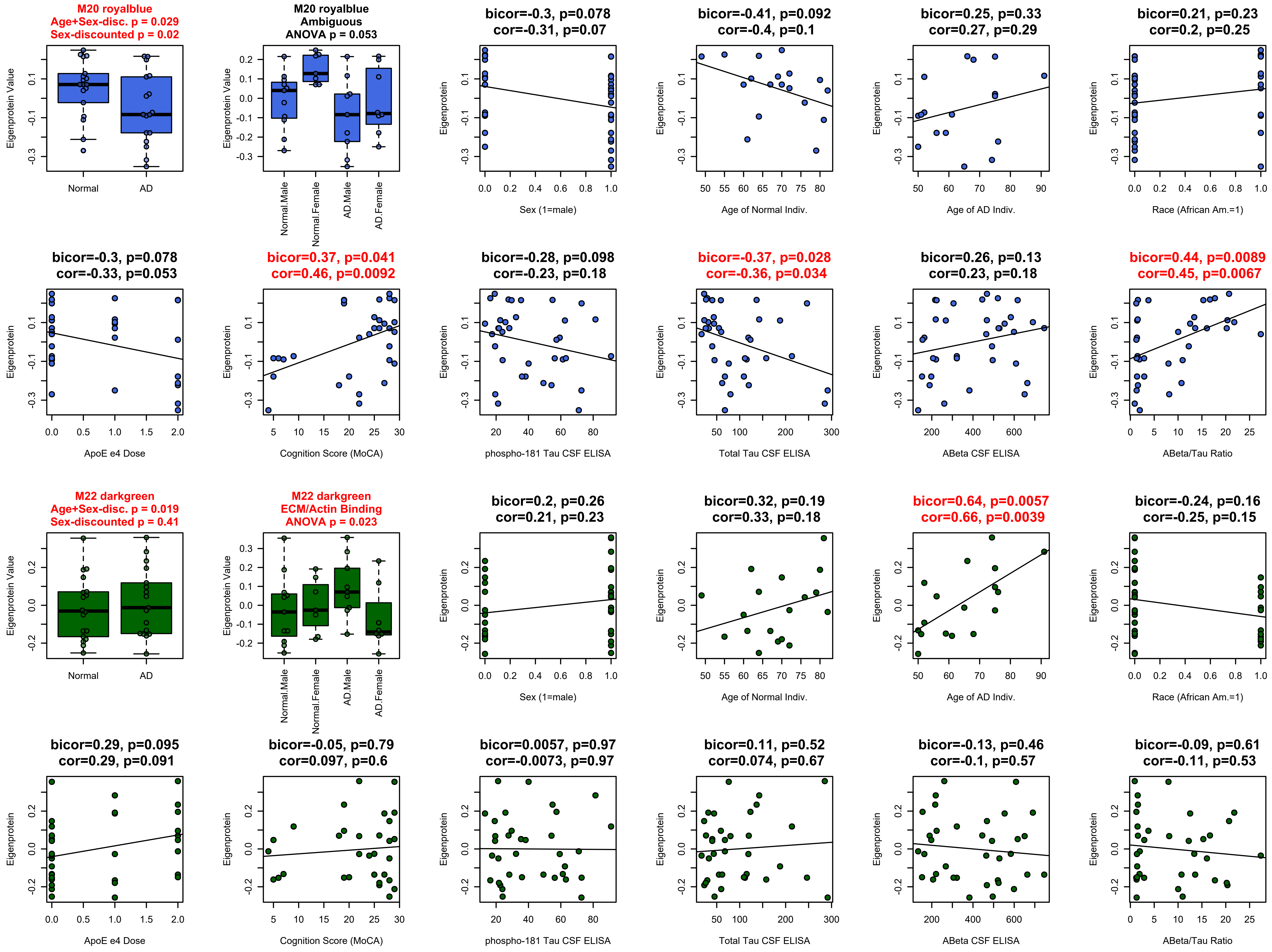
M20 royalblue
Age+Sex-disc. p = 0.029
Sex-discounted p = 0.02
Eigenprotein Value
Normal
AD
M20 royalblue
Ambiguous
ANOVA p = 0.053
Normal.Male
Normal.Female
AD.Male
AD.Female
bicor=-0.3, p=0.078
cor=-0.31, p=0.07
Sex (1=male)
bicor=-0.41, p=0.092
cor=-0.4, p=0.1
Age of Normal Indiv.
bicor=0.25, p=0.33
cor=0.27, p=0.29
Age of AD Indiv.
bicor=0.21, p=0.23
cor=0.2, p=0.25
Race (African Am.=1)
bicor=-0.3, p=0.078
cor=-0.33, p=0.053
ApoE e4 Dose
bicor=0.37, p=0.041
cor=0.46, p=0.0092
Cognition Score (MoCA)
bicor=-0.28, p=0.098
cor=-0.23, p=0.18
phospho-181 Tau CSF ELISA
bicor=-0.37, p=0.028
cor=-0.36, p=0.034
Total Tau CSF ELISA
bicor=0.26, p=0.13
cor=0.23, p=0.18
ABeta CSF ELISA
bicor=0.44, p=0.0089
cor=0.45, p=0.0067
ABeta/Tau Ratio
Eigenprotein
M22 darkgreen
Age+Sex-disc. p = 0.019
Sex-discounted p = 0.41
M22 darkgreen
ECM/Actin Binding
ANOVA p = 0.023
bicor=0.2, p=0.26
cor=0.21, p=0.23
bicor=0.32, p=0.19
cor=0.33, p=0.18
bicor=0.64, p=0.0057
cor=0.66, p=0.0039
bicor=-0.24, p=0.16
cor=-0.25, p=0.15
bicor=0.29, p=0.095
cor=0.29, p=0.091
bicor=-0.05, p=0.79
cor=0.097, p=0.6
bicor=0.0057, p=0.97
cor=-0.0073, p=0.97
bicor=0.11, p=0.52
cor=0.074, p=0.67
bicor=-0.13, p=0.46
cor=-0.1, p=0.57
bicor=-0.09, p=0.61
cor=-0.11, p=0.53

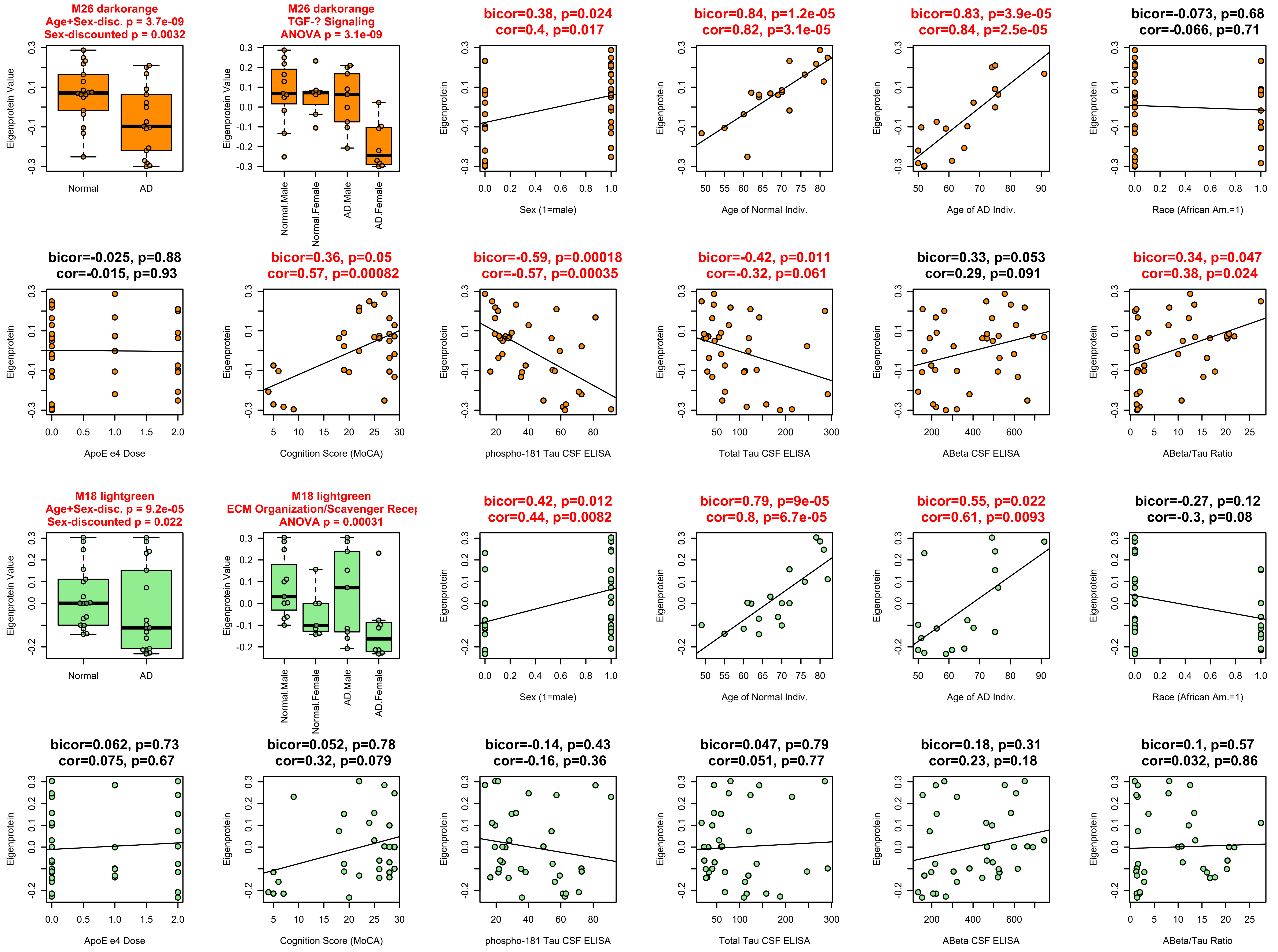

M26 darkorange
Age+Sex-disc. p = 3.7e-09
Sex-discounted p = 0.0032
Eigenprotein Value
Normal
AD
M26 darkorange
TGF-? Signaling
ANOVA p = 3.1e-09
Normal.Male
Normal.Female
AD.Male
AD.Female
bicor=0.38, p=0.024
cor=0.4, p=0.017
Sex (1=male)
bicor=0.84, p=1.2e-05
cor=0.82, p=3.1e-05
Age of Normal Indiv.
bicor=0.83, p=3.9e-05
cor=0.84, p=2.5e-05
Age of AD Indiv.
bicor=-0.073, p=0.68
cor=-0.066, p=0.71
Race (African Am.=1)
bicor=-0.025, p=0.88
cor=-0.015, p=0.93
ApoE e4 Dose
bicor=0.36, p=0.05
cor=0.57, p=0.00082
Cognition Score (MoCA)
bicor=-0.59, p=0.00018
cor=-0.57, p=0.00035
phospho-181 Tau CSF ELISA
bicor=-0.42, p=0.011
cor=-0.32, p=0.061
Total Tau CSF ELISA
bicor=0.33, p=0.053
cor=0.29, p=0.091
ABeta CSF ELISA
bicor=0.34, p=0.047
cor=0.38, p=0.024
ABeta/Tau Ratio
M18 lightgreen
Age+Sex-disc. p = 9.2e-05
Sex-discounted p = 0.022
M18 lightgreen
ECM Organization/Scavenger Recep
ANOVA p = 0.00031
bicor=0.42, p=0.012
cor=0.44, p=0.0082
bicor=0.79, p=9e-05
cor=0.8, p=6.7e-05
bicor=0.55, p=0.022
cor=0.61, p=0.0093
bicor=-0.27, p=0.12
cor=-0.3, p=0.08
bicor=0.062, p=0.73
cor=0.075, p=0.67
bicor=0.052, p=0.78
cor=0.32, p=0.079
bicor=-0.14, p=0.43
cor=-0.16, p=0.36
bicor=0.047, p=0.79
cor=0.051, p=0.77
bicor=0.18, p=0.31
cor=0.23, p=0.18
bicor=0.1, p=0.57
cor=0.032, p=0.86
Eigenprotein

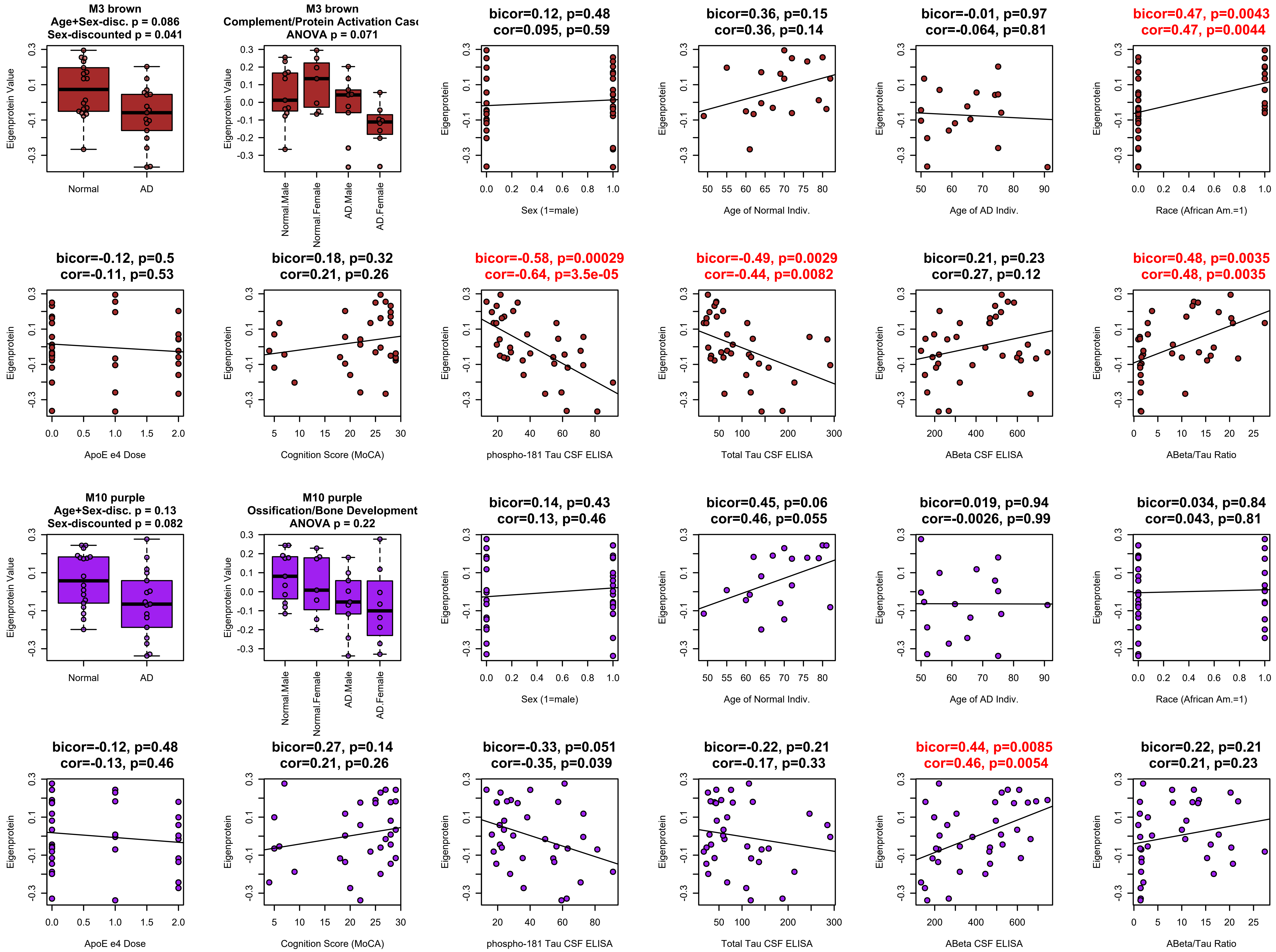
M3 brown
Age+Sex-disc. p = 0.086
Sex-discounted p = 0.041
Eigenprotein Value
Normal
AD
M3 brown
Complement/Protein Activation Cas
ANOVA p = 0.071
Normal.Male
Normal.Female
AD.Male
AD.Female
bicor=0.12, p=0.48
cor=0.095, p=0.59
Eigenprotein
Sex (1=male)
bicor=0.36, p=0.15
cor=0.36, p=0.14
Age of Normal Indiv.
bicor=-0.01, p=0.97
cor=-0.064, p=0.81
Age of AD Indiv.
bicor=0.47, p=0.0043
cor=0.47, p=0.0044
Race (African Am.=1)
bicor=-0.12, p=0.5
cor=-0.11, p=0.53
ApoE e4 Dose
bicor=0.18, p=0.32
cor=0.21, p=0.26
Cognition Score (MoCA)
bicor=-0.58, p=0.00029
cor=-0.64, p=3.5e-05
phospho-181 Tau CSF ELISA
bicor=-0.49, p=0.0029
cor=-0.44, p=0.0082
Total Tau CSF ELISA
bicor=0.21, p=0.23
cor=0.27, p=0.12
ABeta CSF ELISA
bicor=0.48, p=0.0035
cor=0.48, p=0.0035
ABeta/Tau Ratio
M10 purple
Age+Sex-disc. p = 0.13
Sex-discounted p = 0.082
Eigenprotein Value
Normal
AD
M10 purple
Ossification/Bone Development
ANOVA p = 0.22
Normal.Male
Normal.Female
AD.Male
AD.Female
bicor=0.14, p=0.43
cor=0.13, p=0.46
Sex (1=male)
bicor=0.45, p=0.06
cor=0.46, p=0.055
Age of Normal Indiv.
bicor=0.019, p=0.94
cor=-0.0026, p=0.99
Age of AD Indiv.
bicor=0.034, p=0.84
cor=0.043, p=0.81
Race (African Am.=1)
bicor=-0.12, p=0.48
cor=-0.13, p=0.46
ApoE e4 Dose
bicor=0.27, p=0.14
cor=0.21, p=0.26
Cognition Score (MoCA)
bicor=-0.33, p=0.051
cor=-0.35, p=0.039
phospho-181 Tau CSF ELISA
bicor=-0.22, p=0.21
cor=-0.17, p=0.33
Total Tau CSF ELISA
bicor=0.44, p=0.0085
cor=0.46, p=0.0054
ABeta CSF ELISA
bicor=0.22, p=0.21
cor=0.21, p=0.23
ABeta/Tau Ratio

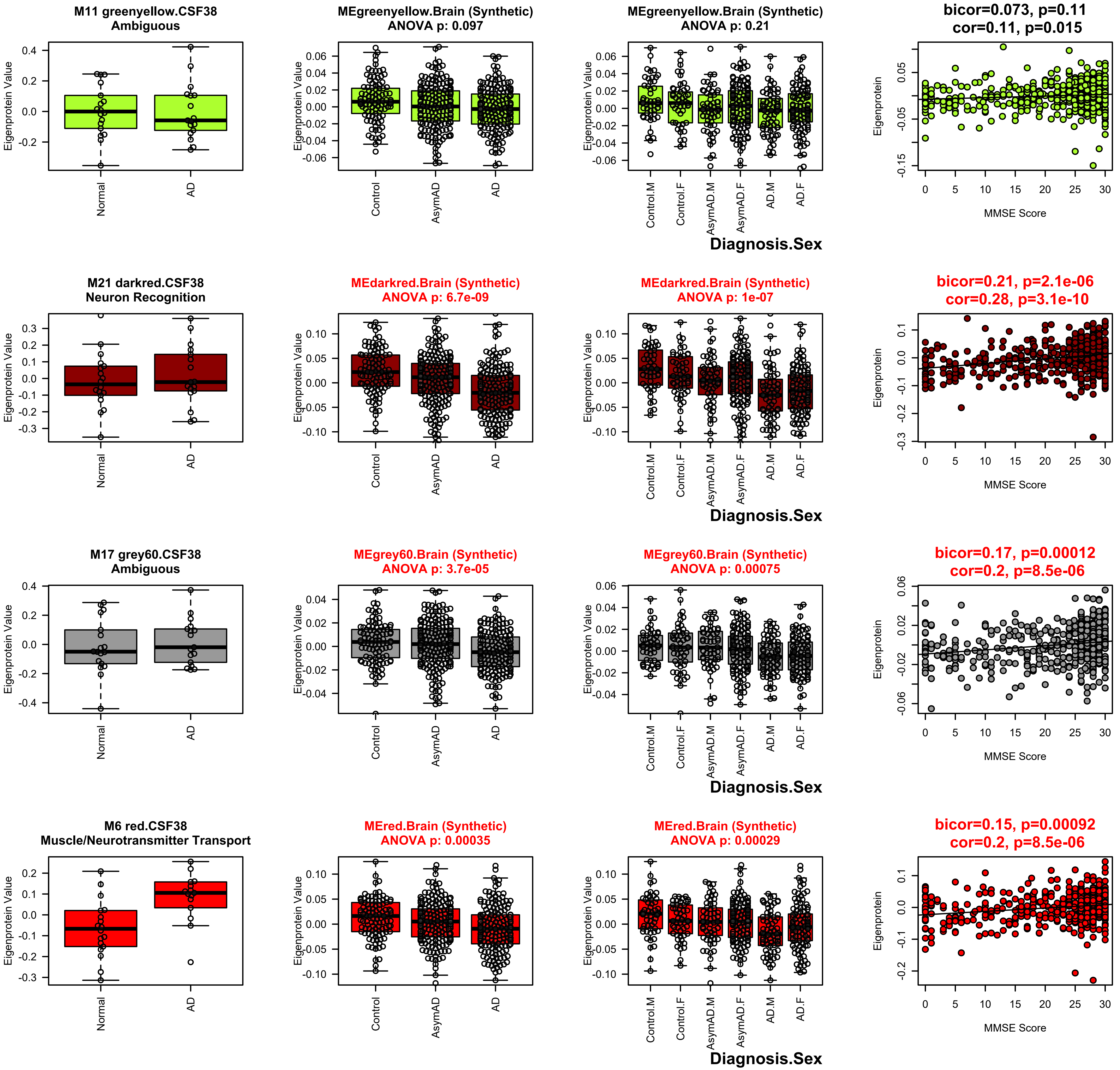
M11 greenyellow.CSF38
Ambiguous
Eigenprotein Value
Normal
AD
MEgreenyellow.Brain (Synthetic)
ANOVA p: 0.097
Control
AsymAD
AD
MEgreenyellow.Brain (Synthetic)
ANOVA p: 0.21
Control.M
Control.F
AsymAD.M
AsymAD.F
AD.M
AD.F
Diagnosis.Sex
bicor=0.073, p=0.11
cor=0.11, p=0.015
Eigenprotein
MMSE Score
M21 darkred.CSF38
Neuron Recognition
MEdarkred.Brain (Synthetic)
ANOVA p: 6.7e-09
MEdarkred.Brain (Synthetic)
ANOVA p: 1e-07
bicor=0.21, p=2.1e-06
cor=0.28, p=3.1e-10
M17 grey60.CSF38
Ambiguous
MEgrey60.Brain (Synthetic)
ANOVA p: 3.7e-05
MEgrey60.Brain (Synthetic)
ANOVA p: 0.00075
bicor=0.17, p=0.00012
cor=0.2, p=8.5e-06
M6 red.CSF38
Muscle/Neurotransmitter Transport
MEred.Brain (Synthetic)
ANOVA p: 0.00035
MEred.Brain (Synthetic)
ANOVA p: 0.00029
bicor=0.15, p=0.00092
cor=0.2, p=8.5e-06

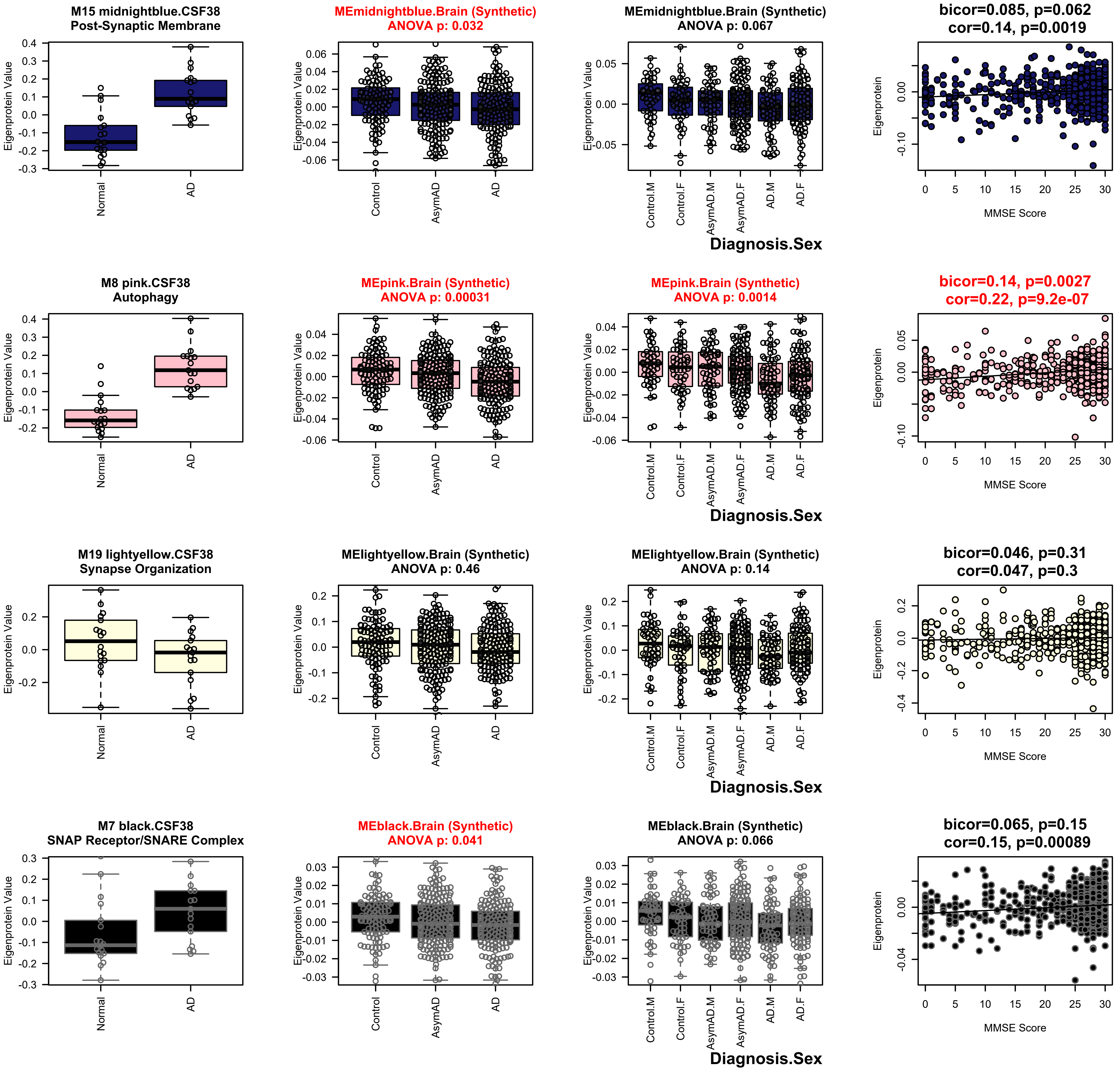

M15 midnightblue.CSF38
Post-Synaptic Membrane
MEmidnightblue.Brain (Synthetic)
ANOVA p: 0.032
MEmidnightblue.Brain (Synthetic)
ANOVA p: 0.067
bicor=0.085, p=0.062
cor=0.14, p=0.0019
M8 pink.CSF38
Autophagy
MEpink.Brain (Synthetic)
ANOVA p: 0.00031
MEpink.Brain (Synthetic)
ANOVA p: 0.0014
bicor=0.14, p=0.0027
cor=0.22, p=9.2e-07
M19 lightyellow.CSF38
Synapse Organization
MElightyellow.Brain (Synthetic)
ANOVA p: 0.46
MElightyellow.Brain (Synthetic)
ANOVA p: 0.14
bicor=0.046, p=0.31
cor=0.047, p=0.3
M7 black.CSF38
SNAP Receptor/SNARE Complex
MEblack.Brain (Synthetic)
ANOVA p: 0.041
MEblack.Brain (Synthetic)
ANOVA p: 0.066
bicor=0.065, p=0.15
cor=0.15, p=0.00089
Eigenprotein Value
Eigenprotein
Normal
AD
Control
AsymAD
Control.M
Control.F
AsymAD.M
AsymAD.F
AD.M
AD.F
Diagnosis.Sex
MMSE Score

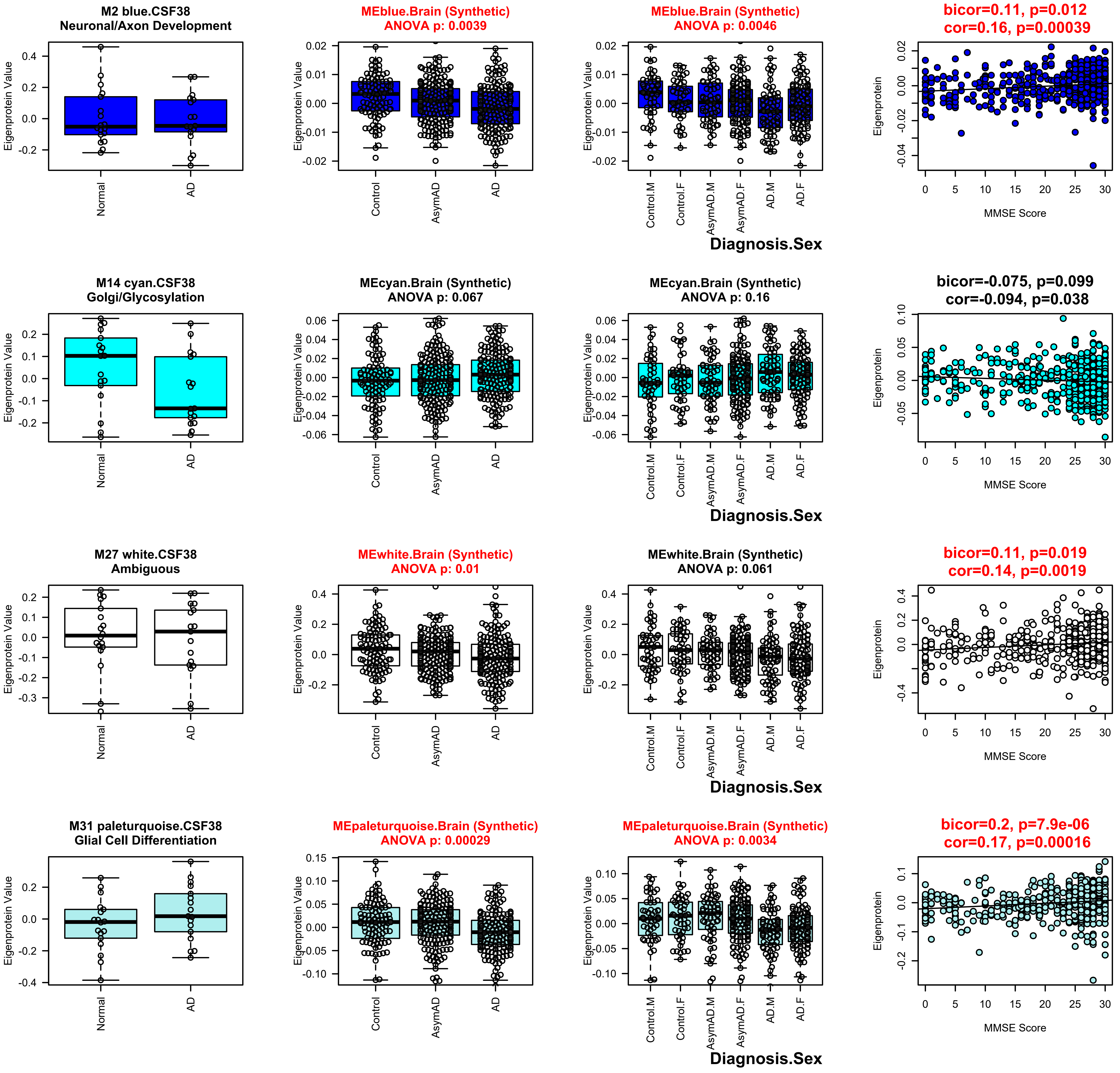
M2 blue.CSF38
Neuronal/Axon Development
MEblue.Brain (Synthetic)
ANOVA p: 0.0039
MEblue.Brain (Synthetic)
ANOVA p: 0.0046
bicor=0.11, p=0.012
cor=0.16, p=0.00039
M14 cyan.CSF38
Golgi/Glycosylation
MEcyan.Brain (Synthetic)
ANOVA p: 0.067
MEcyan.Brain (Synthetic)
ANOVA p: 0.16
bicor=-0.075, p=0.099
cor=-0.094, p=0.038
M27 white.CSF38
Ambiguous
MEwhite.Brain (Synthetic)
ANOVA p: 0.01
MEwhite.Brain (Synthetic)
ANOVA p: 0.061
bicor=0.11, p=0.019
cor=0.14, p=0.0019
M31 paleturquoise.CSF38
Glial Cell Differentiation
MEpaleturquoise.Brain (Synthetic)
ANOVA p: 0.00029
MEpaleturquoise.Brain (Synthetic)
ANOVA p: 0.0034
bicor=0.2, p=7.9e-06
cor=0.17, p=0.00016
Eigenprotein Value
Eigenprotein
Normal
AD
Control
AsymAD
Control.M
Control.F
AsymAD.M
AsymAD.F
AD.M
AD.F
Diagnosis.Sex
MMSE Score

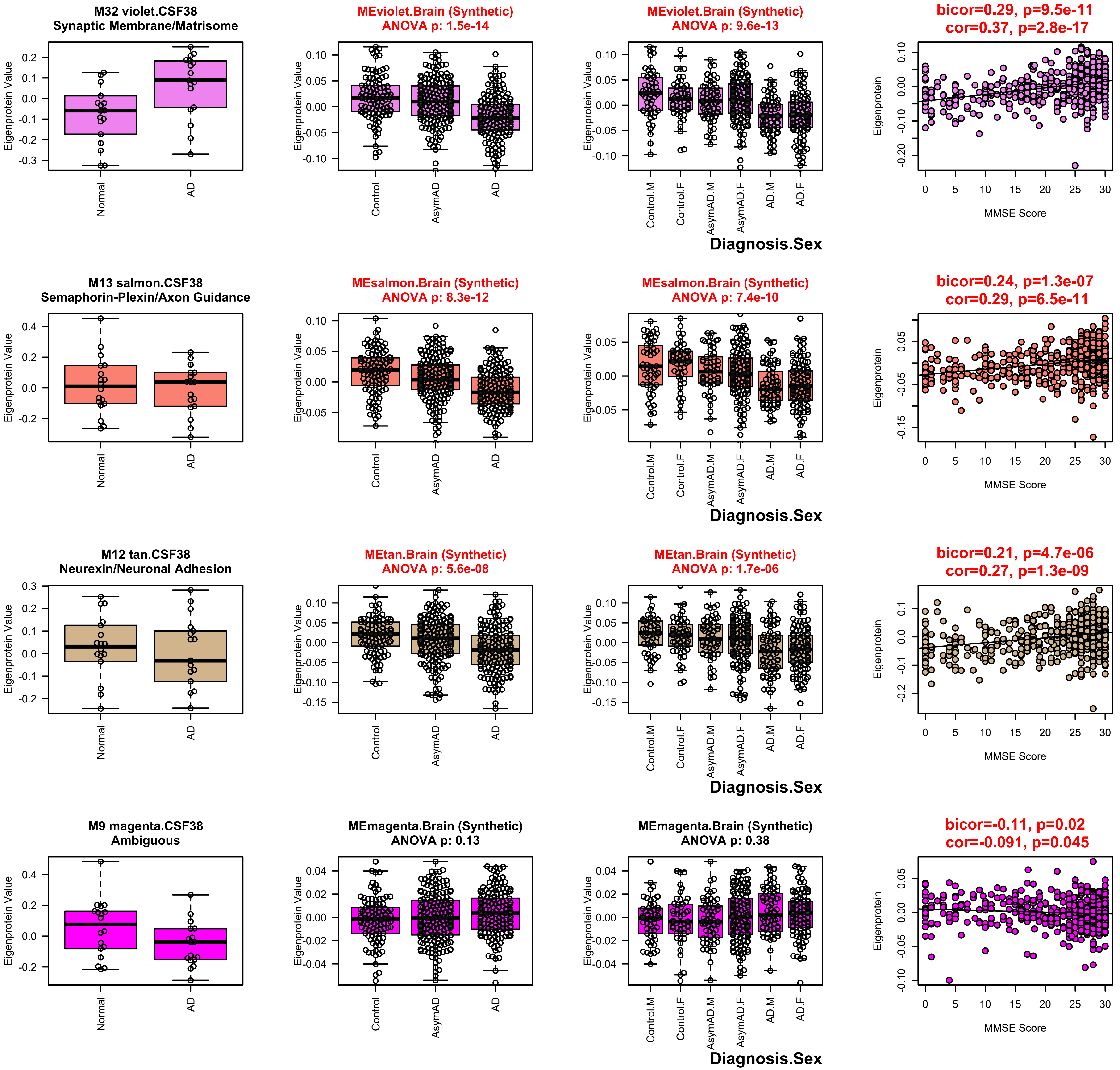
M32 violet.CSF38
Synaptic Membrane/Matrisome
MEviolet.Brain (Synthetic)
ANOVA p: 1.5e-14
MEviolet.Brain (Synthetic)
ANOVA p: 9.6e-13
bicor=0.29, p=9.5e-11
cor=0.37, p=2.8e-17
M13 salmon.CSF38
Semaphorin-Plexin/Axon Guidance
MEsalmon.Brain (Synthetic)
ANOVA p: 8.3e-12
MEsalmon.Brain (Synthetic)
ANOVA p: 7.4e-10
bicor=0.24, p=1.3e-07
cor=0.29, p=6.5e-11
M12 tan.CSF38
Neurexin/Neuronal Adhesion
MEtan.Brain (Synthetic)
ANOVA p: 5.6e-08
MEtan.Brain (Synthetic)
ANOVA p: 1.7e-06
bicor=0.21, p=4.7e-06
cor=0.27, p=1.3e-09
M9 magenta.CSF38
Ambiguous
MEmagenta.Brain (Synthetic)
ANOVA p: 0.13
MEmagenta.Brain (Synthetic)
ANOVA p: 0.38
bicor=-0.11, p=0.02
cor=-0.091, p=0.045
Eigenprotein Value
Eigenprotein
Normal
AD
Control
AsymAD
Control.M
Control.F
AsymAD.M
AsymAD.F
AD.M
AD.F
Diagnosis.Sex
MMSE Score

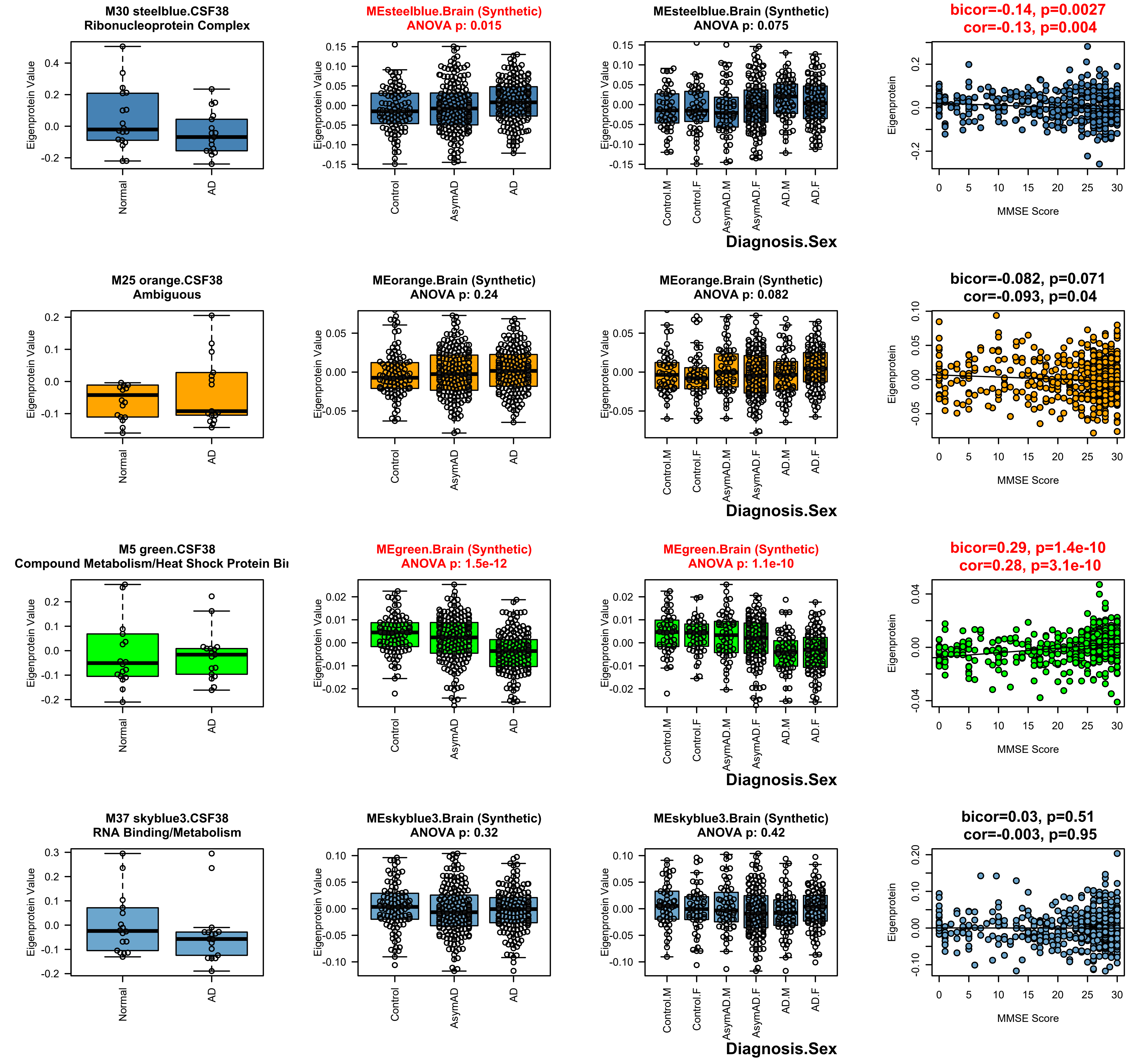
M30 steelblue.CSF38
Ribonucleoprotein Complex
MEsteelblue.Brain (Synthetic)
ANOVA p: 0.015
MEsteelblue.Brain (Synthetic)
ANOVA p: 0.075
bicor=-0.14, p=0.0027
cor=-0.13, p=0.004
M25 orange.CSF38
Ambiguous
MEorange.Brain (Synthetic)
ANOVA p: 0.24
MEorange.Brain (Synthetic)
ANOVA p: 0.082
bicor=-0.082, p=0.071
cor=-0.093, p=0.04
M5 green.CSF38
Compound Metabolism/Heat Shock Protein Bi
MEgreen.Brain (Synthetic)
ANOVA p: 1.5e-12
MEgreen.Brain (Synthetic)
ANOVA p: 1.1e-10
bicor=0.29, p=1.4e-10
cor=0.28, p=3.1e-10
M37 skyblue3.CSF38
RNA Binding/Metabolism
MEskyblue3.Brain (Synthetic)
ANOVA p: 0.32
MEskyblue3.Brain (Synthetic)
ANOVA p: 0.42
bicor=0.03, p=0.51
cor=-0.003, p=0.95
Eigenprotein Value
Eigenprotein
Normal
AD
Control
AsymAD
Control.M
Control.F
AsymAD.M
AsymAD.F
AD.M
AD.F
Diagnosis.Sex
MMSE Score

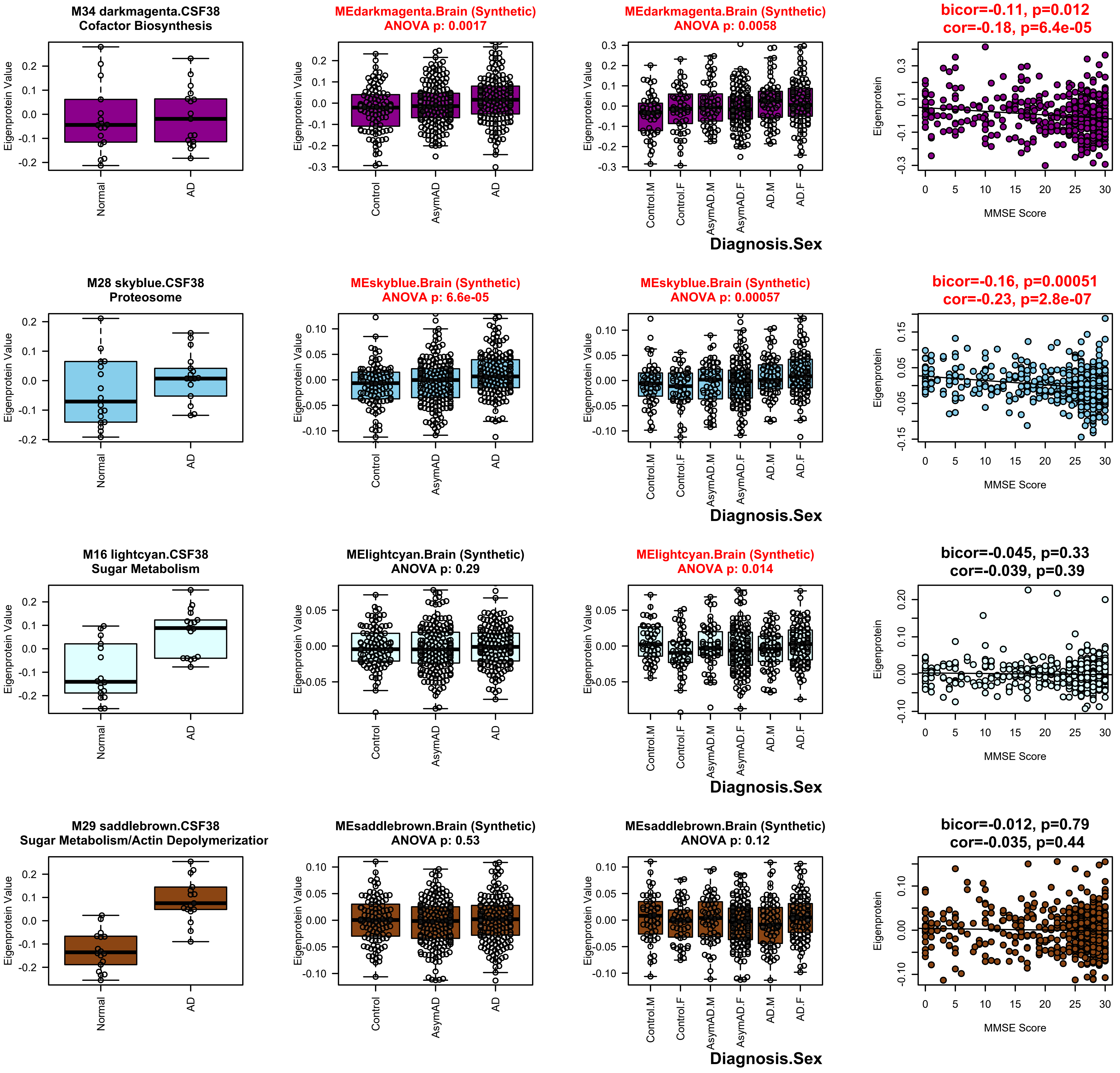

M34 darkmagenta.CSF38
Cofactor Biosynthesis
MEdarkmagenta.Brain (Synthetic)
ANOVA p: 0.0017
MEdarkmagenta.Brain (Synthetic)
ANOVA p: 0.0058
bicor=-0.11, p=0.012
cor=-0.18, p=6.4e-05
M28 skyblue.CSF38
Proteosome
MEskyblue.Brain (Synthetic)
ANOVA p: 6.6e-05
MEskyblue.Brain (Synthetic)
ANOVA p: 0.00057
bicor=-0.16, p=0.00051
cor=-0.23, p=2.8e-07
M16 lightcyan.CSF38
Sugar Metabolism
MElightcyan.Brain (Synthetic)
ANOVA p: 0.29
MElightcyan.Brain (Synthetic)
ANOVA p: 0.014
bicor=-0.045, p=0.33
cor=-0.039, p=0.39
M29 saddlebrown.CSF38
Sugar Metabolism/Actin Depolymerizatior
MEsaddlebrown.Brain (Synthetic)
ANOVA p: 0.53
MEsaddlebrown.Brain (Synthetic)
ANOVA p: 0.12
bicor=-0.012, p=0.79
cor=-0.035, p=0.44
Eigenprotein Value
Eigenprotein
Normal
AD
Control
AsymAD
Control.M
Control.F
AsymAD.M
AsymAD.F
AD.M
AD.F
Diagnosis.Sex
MMSE Score

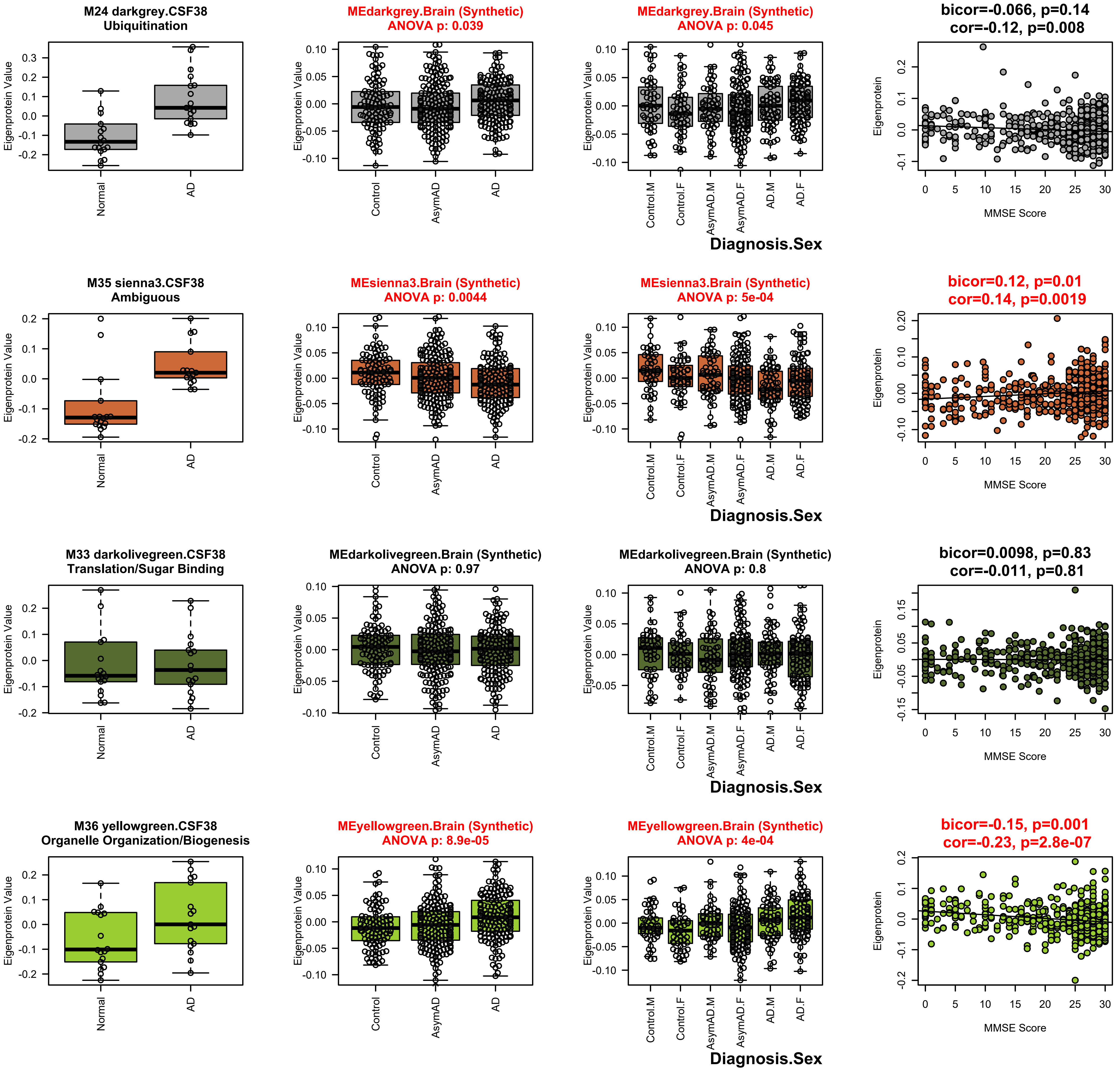
M24 darkgrey.CSF38
Ubiquitination
MEdarkgrey.Brain (Synthetic)
ANOVA p: 0.039
MEdarkgrey.Brain (Synthetic)
ANOVA p: 0.045
bicor=-0.066, p=0.14
cor=-0.12, p=0.008
M35 sienna3.CSF38
Ambiguous
MEsienna3.Brain (Synthetic)
ANOVA p: 0.0044
MEsienna3.Brain (Synthetic)
ANOVA p: 5e-04
bicor=0.12, p=0.01
cor=0.14, p=0.0019
M33 darkolivegreen.CSF38
Translation/Sugar Binding
MEdarkolivegreen.Brain (Synthetic)
ANOVA p: 0.97
MEdarkolivegreen.Brain (Synthetic)
ANOVA p: 0.8
bicor=0.0098, p=0.83
cor=-0.011, p=0.81
M36 yellowgreen.CSF38
Organelle Organization/Biogenesis
MEyellowgreen.Brain (Synthetic)
ANOVA p: 8.9e-05
MEyellowgreen.Brain (Synthetic)
ANOVA p: 4e-04
bicor=-0.15, p=0.001
cor=-0.23, p=2.8e-07
Eigenprotein Value
Eigenprotein
Normal
AD
Control
AsymAD
Control.M
Control.F
AsymAD.M
AsymAD.F
AD.M
AD.F
Diagnosis.Sex
MMSE Score

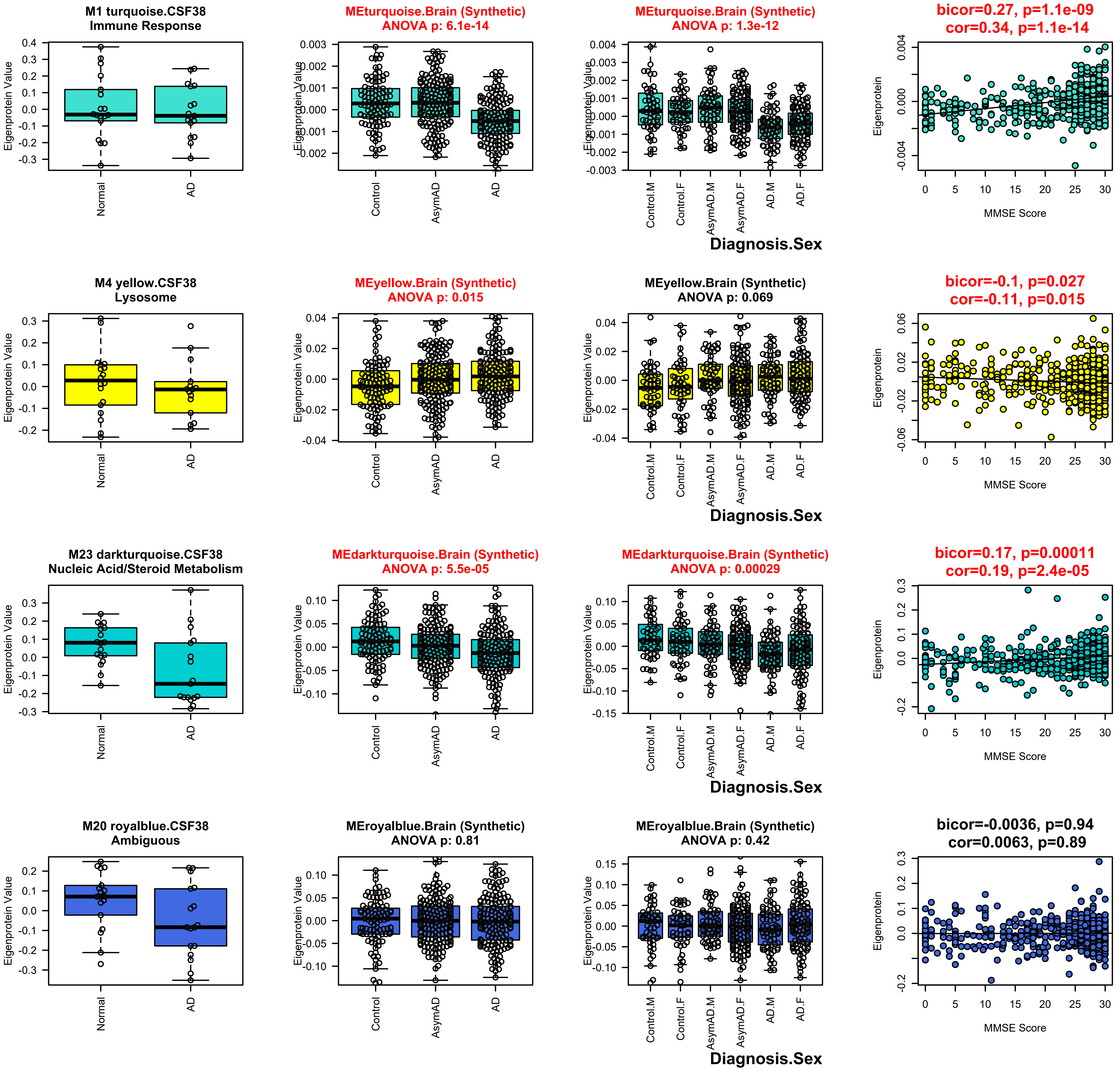
M1 turquoise.CSF38
Immune Response
MEturquoise.Brain (Synthetic)
ANOVA p: 6.1e-14
MEturquoise.Brain (Synthetic)
ANOVA p: 1.3e-12
bicor=0.27, p=1.1e-09
cor=0.34, p=1.1e-14
M4 yellow.CSF38
Lysosome
MEyellow.Brain (Synthetic)
ANOVA p: 0.015
MEyellow.Brain (Synthetic)
ANOVA p: 0.069
bicor=-0.1, p=0.027
cor=-0.11, p=0.015
M23 darkturquoise.CSF38
Nucleic Acid/Steroid Metabolism
MEdarkturquoise.Brain (Synthetic)
ANOVA p: 5.5e-05
MEdarkturquoise.Brain (Synthetic)
ANOVA p: 0.00029
bicor=0.17, p=0.00011
cor=0.19, p=2.4e-05
M20 royalblue.CSF38
Ambiguous
MEroyalblue.Brain (Synthetic)
ANOVA p: 0.81
MEroyalblue.Brain (Synthetic)
ANOVA p: 0.42
bicor=-0.0036, p=0.94
cor=0.0063, p=0.89
Eigenprotein Value
Eigenprotein
Normal
AD
Control
AsymAD
Control.M
Control.F
AsymAD.M
AsymAD.F
AD.M
AD.F
Diagnosis.Sex
MMSE Score

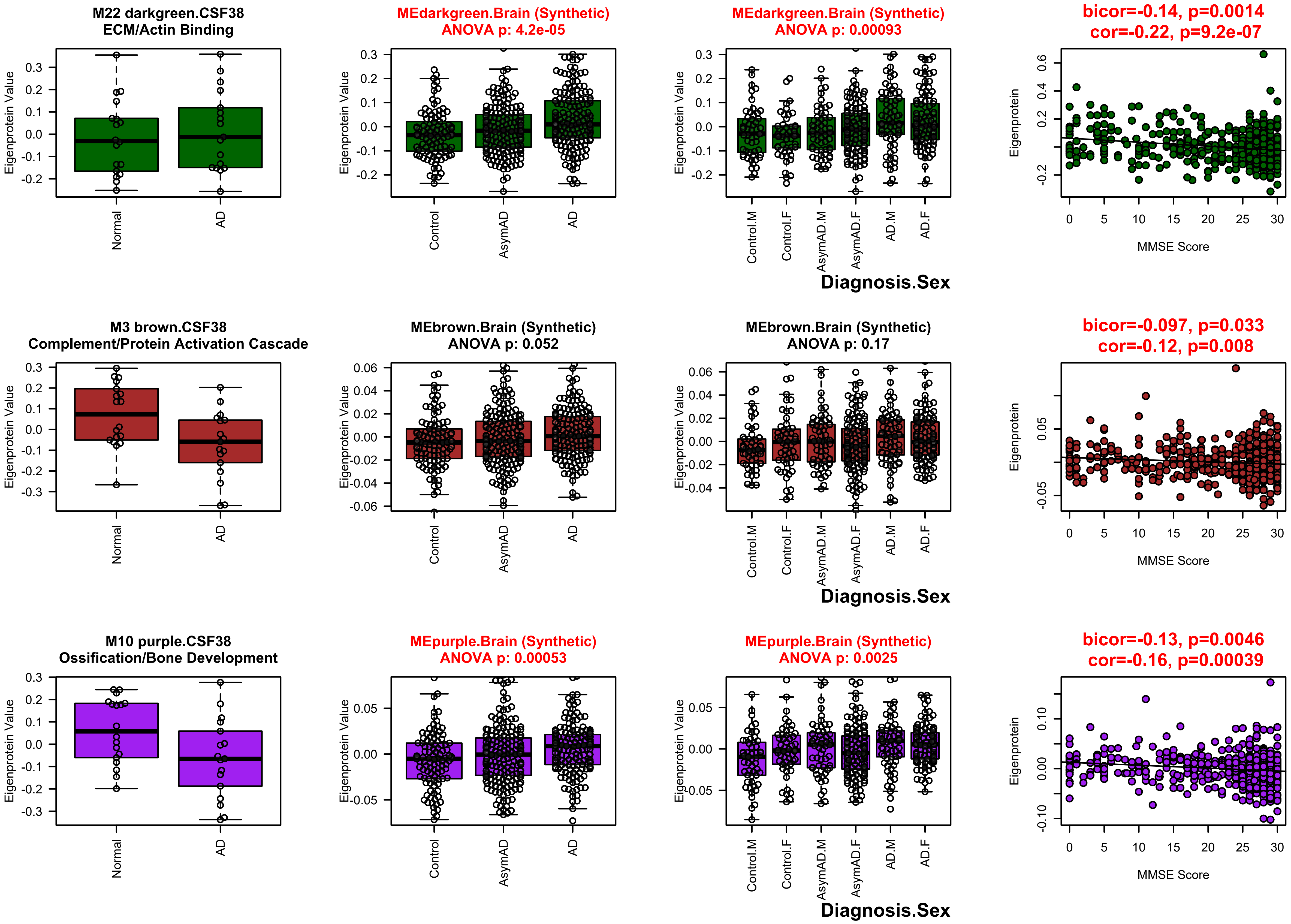
M22 darkgreen.CSF38
ECM/Actin Binding
Eigenprotein Value
Normal
AD
MEdarkgreen.Brain (Synthetic)
ANOVA p: 4.2e-05
Control
AsymAD
AD
MEdarkgreen.Brain (Synthetic)
ANOVA p: 0.00093
Control.M
Control.F
AsymAD.M
AsymAD.F
AD.M
AD.F
Diagnosis.Sex
bicor=-0.14, p=0.0014
cor=-0.22, p=9.2e-07
Eigenprotein
MMSE Score
M3 brown.CSF38
Complement/Protein Activation Cascade
MEbrown.Brain (Synthetic)
ANOVA p: 0.052
MEbrown.Brain (Synthetic)
ANOVA p: 0.17
bicor=-0.097, p=0.033
cor=-0.12, p=0.008
M10 purple.CSF38
Ossification/Bone Development
MEpurple.Brain (Synthetic)
ANOVA p: 0.00053
MEpurple.Brain (Synthetic)
ANOVA p: 0.0025
bicor=-0.13, p=0.0046
cor=-0.16, p=0.00039

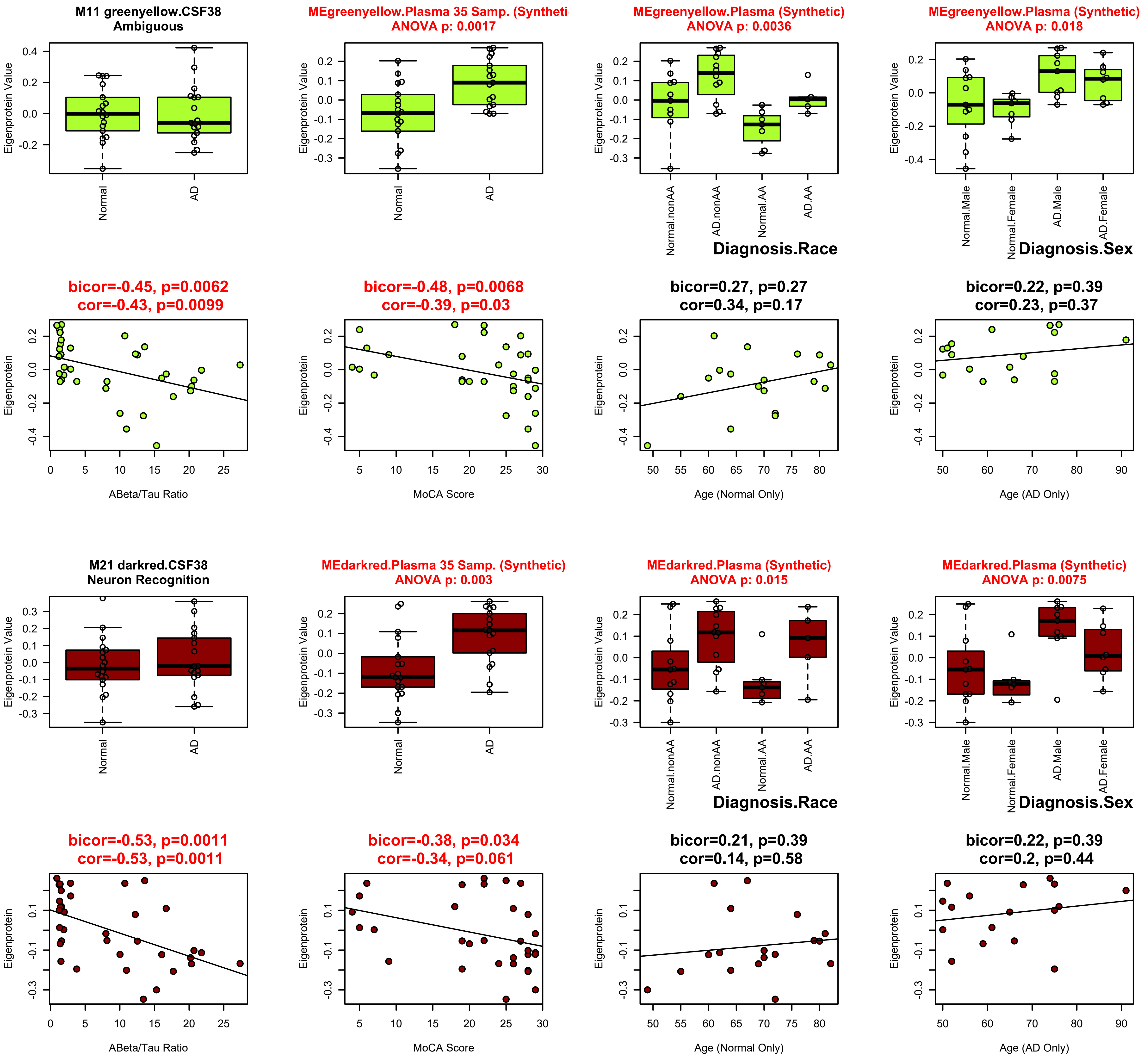
M11 greenyellow.CSF38
Ambiguous
MEgreenyellow.Plasma 35 Samp. (Syntheti
ANOVA p: 0.0017
MEgreenyellow.Plasma (Synthetic)
ANOVA p: 0.0036
MEgreenyellow.Plasma (Synthetic)
ANOVA p: 0.018
Eigenprotein Value
Normal
AD
Normal.nonAA
AD.nonAA
Normal.AA
AD.AA
Diagnosis.Race
Normal.Male
Normal.Female
AD.Male
AD.Female
Diagnosis.Sex
bicor=-0.45, p=0.0062
cor=-0.43, p=0.0099
bicor=-0.48, p=0.0068
cor=-0.39, p=0.03
bicor=0.27, p=0.27
cor=0.34, p=0.17
bicor=0.22, p=0.39
cor=0.23, p=0.37
Eigenprotein
ABeta/Tau Ratio
MoCA Score
Age (Normal Only)
Age (AD Only)
M21 darkred.CSF38
Neuron Recognition
MEdarkred.Plasma 35 Samp. (Synthetic)
ANOVA p: 0.003
MEdarkred.Plasma (Synthetic)
ANOVA p: 0.015
MEdarkred.Plasma (Synthetic)
ANOVA p: 0.0075
Eigenprotein Value
Normal
AD
Normal.nonAA
AD.nonAA
Normal.AA
AD.AA
Diagnosis.Race
Normal.Male
Normal.Female
AD.Male
AD.Female
Diagnosis.Sex
bicor=-0.53, p=0.0011
cor=-0.53, p=0.0011
bicor=-0.38, p=0.034
cor=-0.34, p=0.061
bicor=0.21, p=0.39
cor=0.14, p=0.58
bicor=0.22, p=0.39
cor=0.2, p=0.44
Eigenprotein
ABeta/Tau Ratio
MoCA Score
Age (Normal Only)
Age (AD Only)

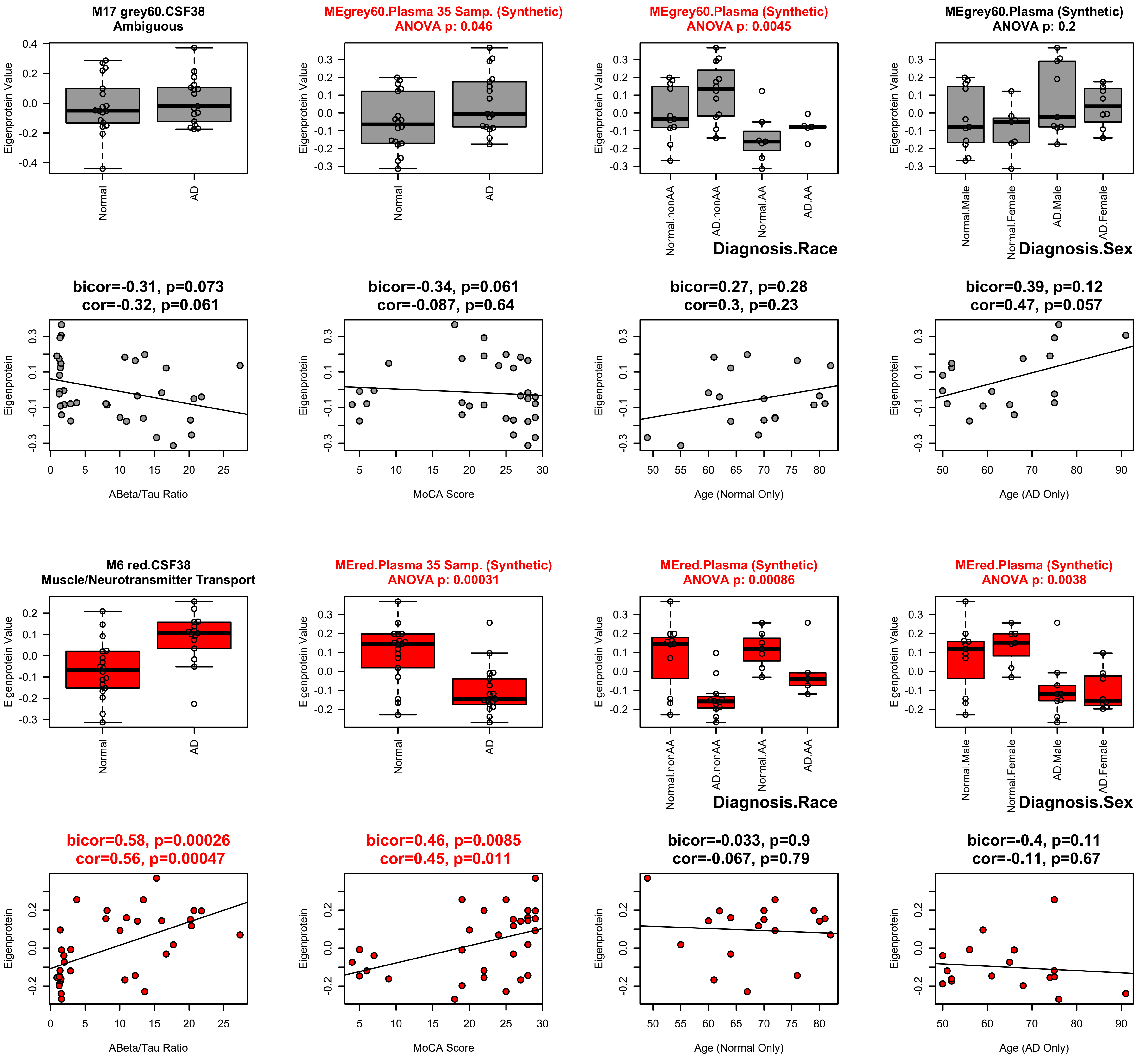

M17 grey60.CSF38
Ambiguous
Eigenprotein Value
Normal
AD
MEgrey60.Plasma 35 Samp. (Synthetic)
ANOVA p: 0.046
MEgrey60.Plasma (Synthetic)
ANOVA p: 0.0045
Normal.nonAA
AD.nonAA
Normal.AA
AD.AA
Diagnosis.Race
MEgrey60.Plasma (Synthetic)
ANOVA p: 0.2
Normal.Male
Normal.Female
AD.Male
AD.Female
Diagnosis.Sex
bicor=-0.31, p=0.073
cor=-0.32, p=0.061
Eigenprotein
ABeta/Tau Ratio
bicor=-0.34, p=0.061
cor=-0.087, p=0.64
MoCA Score
bicor=0.27, p=0.28
cor=0.3, p=0.23
Age (Normal Only)
bicor=0.39, p=0.12
cor=0.47, p=0.057
Age (AD Only)
M6 red.CSF38
Muscle/Neurotransmitter Transport
MEred.Plasma 35 Samp. (Synthetic)
ANOVA p: 0.00031
MEred.Plasma (Synthetic)
ANOVA p: 0.00086
MEred.Plasma (Synthetic)
ANOVA p: 0.0038
bicor=0.58, p=0.00026
cor=0.56, p=0.00047
bicor=0.46, p=0.0085
cor=0.45, p=0.011
bicor=-0.033, p=0.9
cor=-0.067, p=0.79
bicor=-0.4, p=0.11
cor=-0.11, p=0.67

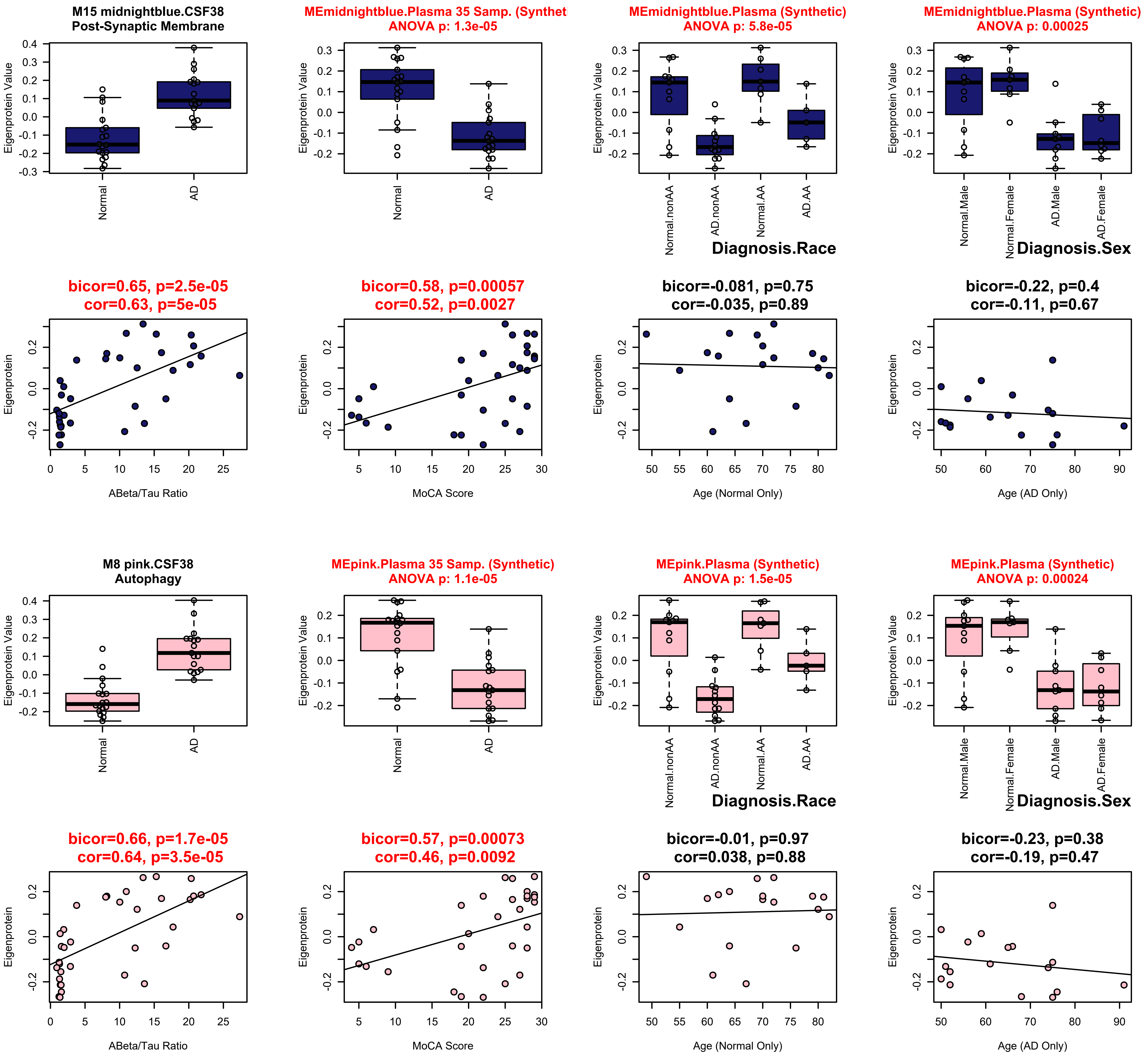
M15 midnightblue.CSF38
Post-Synaptic Membrane
Eigenprotein Value
Normal
AD
MEmidnightblue.Plasma 35 Samp. (Syntheti
ANOVA p: 1.3e-05
MEmidnightblue.Plasma (Synthetic)
ANOVA p: 5.8e-05
Normal.nonAA
AD.nonAA
Normal.AA
AD.AA
Diagnosis.Race
MEmidnightblue.Plasma (Synthetic)
ANOVA p: 0.00025
Normal.Male
Normal.Female
AD.Male
AD.Female
Diagnosis.Sex
bicor=0.65, p=2.5e-05
cor=0.63, p=5e-05
Eigenprotein
ABeta/Tau Ratio
bicor=0.58, p=0.00057
cor=0.52, p=0.0027
MoCA Score
bicor=-0.081, p=0.75
cor=-0.035, p=0.89
Age (Normal Only)
bicor=-0.22, p=0.4
cor=-0.11, p=0.67
Age (AD Only)
M8 pink.CSF38
Autophagy
MEpink.Plasma 35 Samp. (Synthetic)
ANOVA p: 1.1e-05
MEpink.Plasma (Synthetic)
ANOVA p: 1.5e-05
MEpink.Plasma (Synthetic)
ANOVA p: 0.00024
bicor=0.66, p=1.7e-05
cor=0.64, p=3.5e-05
bicor=0.57, p=0.00073
cor=0.46, p=0.0092
bicor=-0.01, p=0.97
cor=0.038, p=0.88
bicor=-0.23, p=0.38
cor=-0.19, p=0.47

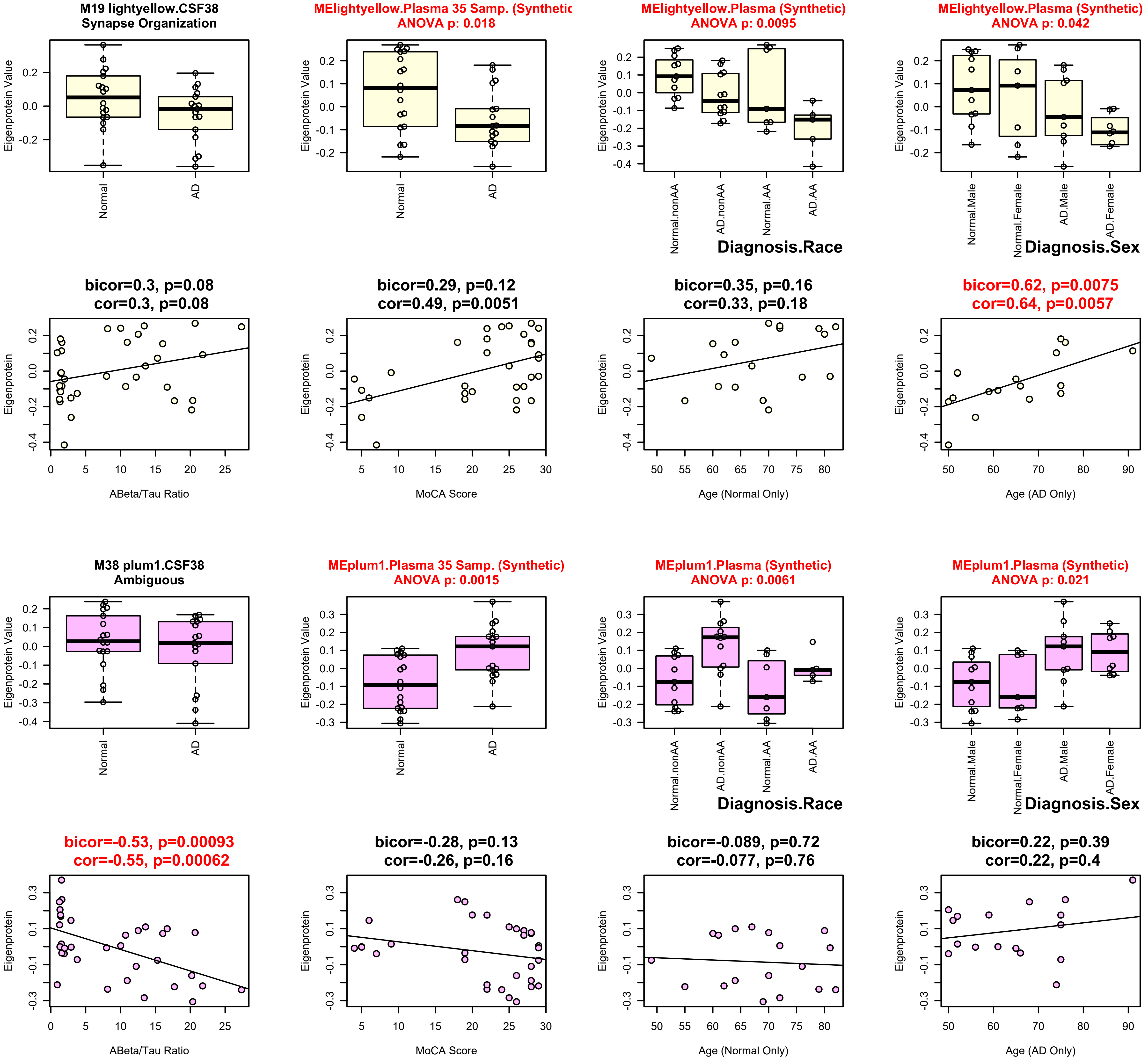

M19 lightyellow.CSF38
Synapse Organization
Eigenprotein Value
Normal
AD
MElightyellow.Plasma 35 Samp. (Synthetic
ANOVA p: 0.018
MElightyellow.Plasma (Synthetic)
ANOVA p: 0.0095
Normal.nonAA
AD.nonAA
Normal.AA
AD.AA
Diagnosis.Race
MElightyellow.Plasma (Synthetic)
ANOVA p: 0.042
Normal.Male
Normal.Female
AD.Male
AD.Female
Diagnosis.Sex
bicor=0.3, p=0.08
cor=0.3, p=0.08
Eigenprotein
ABeta/Tau Ratio
bicor=0.29, p=0.12
cor=0.49, p=0.0051
MoCA Score
bicor=0.35, p=0.16
cor=0.33, p=0.18
Age (Normal Only)
bicor=0.62, p=0.0075
cor=0.64, p=0.0057
Age (AD Only)
M38 plum1.CSF38
Ambiguous
MEplum1.Plasma 35 Samp. (Synthetic)
ANOVA p: 0.0015
MEplum1.Plasma (Synthetic)
ANOVA p: 0.0061
Diagnosis.Race
MEplum1.Plasma (Synthetic)
ANOVA p: 0.021
Diagnosis.Sex
bicor=-0.53, p=0.00093
cor=-0.55, p=0.00062
ABeta/Tau Ratio
bicor=-0.28, p=0.13
cor=-0.26, p=0.16
MoCA Score
bicor=-0.089, p=0.72
cor=-0.077, p=0.76
Age (Normal Only)
bicor=0.22, p=0.39
cor=0.22, p=0.4
Age (AD Only)

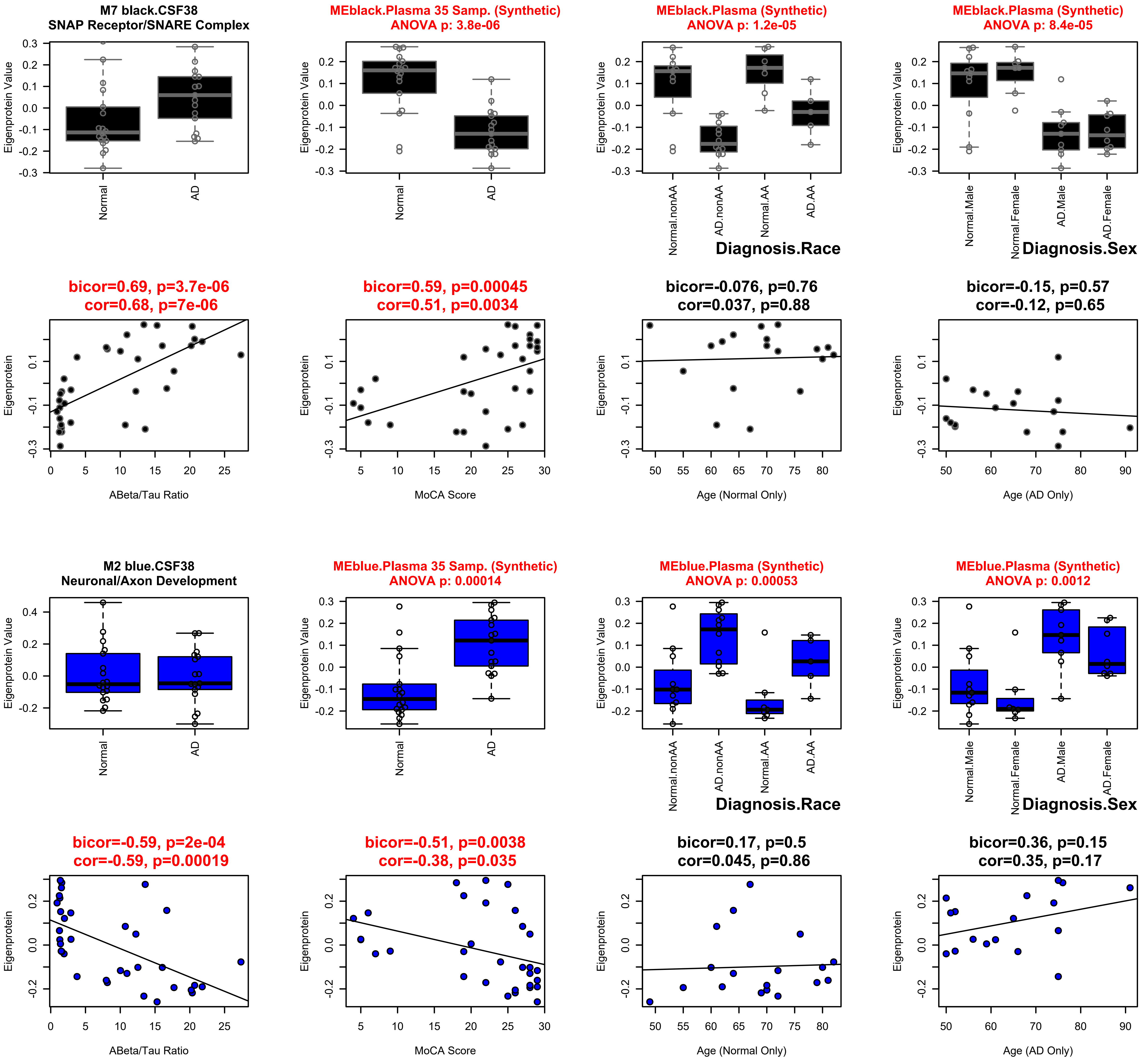

M7 black.CSF38
SNAP Receptor/SNARE Complex
MEblack.Plasma 35 Samp. (Synthetic)
ANOVA p: 3.8e-06
MEblack.Plasma (Synthetic)
ANOVA p: 1.2e-05
MEblack.Plasma (Synthetic)
ANOVA p: 8.4e-05
Eigenprotein Value
Normal
AD
Normal.nonAA
AD.nonAA
Normal.AA
AD.AA
Diagnosis.Race
Normal.Male
Normal.Female
AD.Male
AD.Female
Diagnosis.Sex
bicor=0.69, p=3.7e-06
cor=0.68, p=7e-06
bicor=0.59, p=0.00045
cor=0.51, p=0.0034
bicor=-0.076, p=0.76
cor=0.037, p=0.88
bicor=-0.15, p=0.57
cor=-0.12, p=0.65
Eigenprotein
ABeta/Tau Ratio
MoCA Score
Age (Normal Only)
Age (AD Only)
M2 blue.CSF38
Neuronal/Axon Development
MEblue.Plasma 35 Samp. (Synthetic)
ANOVA p: 0.00014
MEblue.Plasma (Synthetic)
ANOVA p: 0.00053
MEblue.Plasma (Synthetic)
ANOVA p: 0.0012
bicor=-0.59, p=2e-04
cor=-0.59, p=0.00019
bicor=-0.51, p=0.0038
cor=-0.38, p=0.035
bicor=0.17, p=0.5
cor=0.045, p=0.86
bicor=0.36, p=0.15
cor=0.35, p=0.17

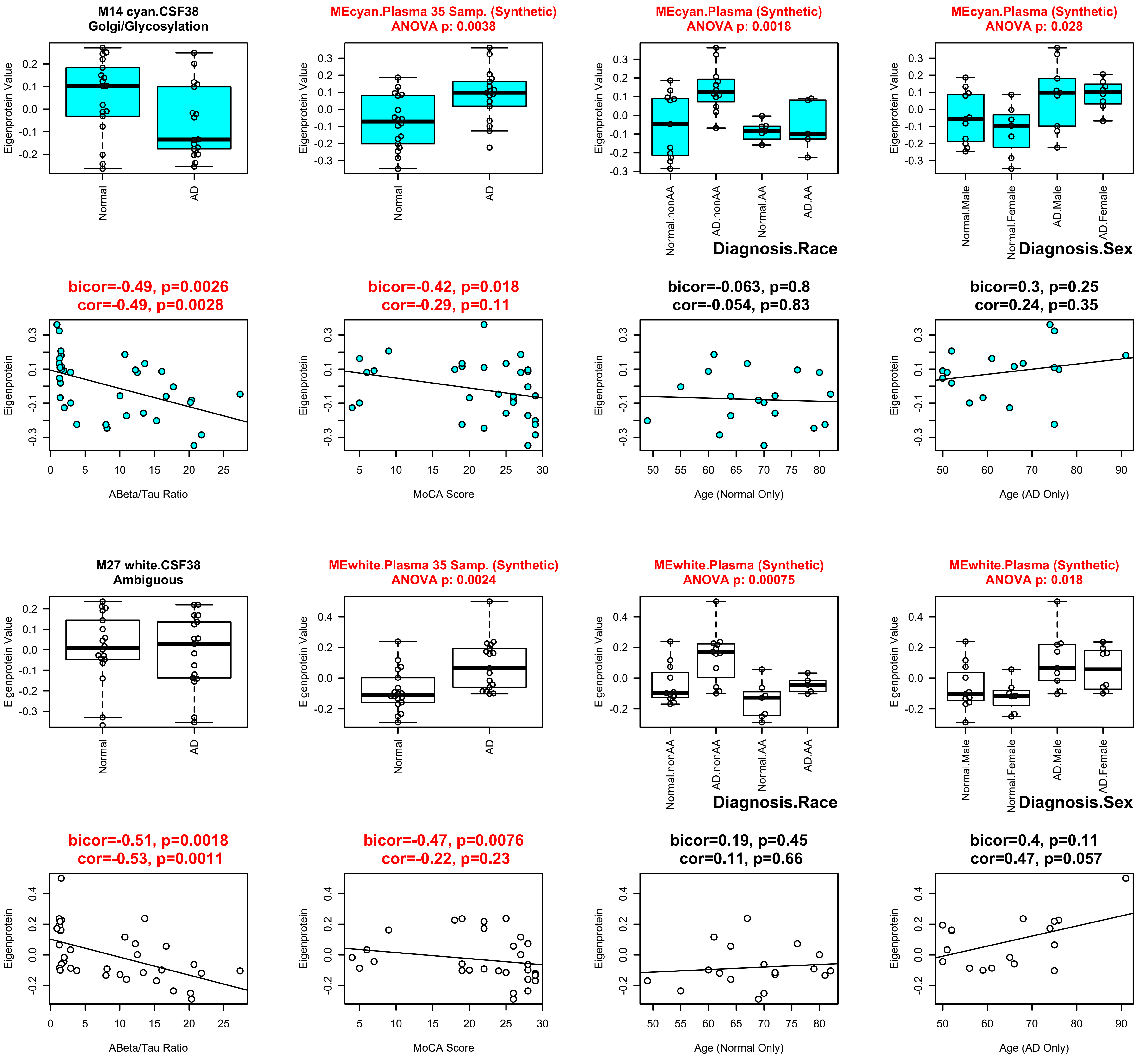
M14 cyan.CSF38
Golgi/Glycosylation
MEcyan.Plasma 35 Samp. (Synthetic)
ANOVA p: 0.0038
MEcyan.Plasma (Synthetic)
ANOVA p: 0.0018
MEcyan.Plasma (Synthetic)
ANOVA p: 0.028
Eigenprotein Value
Normal
AD
Normal.nonAA
AD.nonAA
Normal.AA
AD.AA
Diagnosis.Race
Normal.Male
Normal.Female
AD.Male
AD.Female
Diagnosis.Sex
bicor=-0.49, p=0.0026
cor=-0.49, p=0.0028
bicor=-0.42, p=0.018
cor=-0.29, p=0.11
bicor=-0.063, p=0.8
cor=-0.054, p=0.83
bicor=0.3, p=0.25
cor=0.24, p=0.35
Eigenprotein
ABeta/Tau Ratio
MoCA Score
Age (Normal Only)
Age (AD Only)
M27 white.CSF38
Ambiguous
MEwhite.Plasma 35 Samp. (Synthetic)
ANOVA p: 0.0024
MEwhite.Plasma (Synthetic)
ANOVA p: 0.00075
MEwhite.Plasma (Synthetic)
ANOVA p: 0.018
bicor=-0.51, p=0.0018
cor=-0.53, p=0.0011
bicor=-0.47, p=0.0076
cor=-0.22, p=0.23
bicor=0.19, p=0.45
cor=0.11, p=0.66
bicor=0.4, p=0.11
cor=0.47, p=0.057

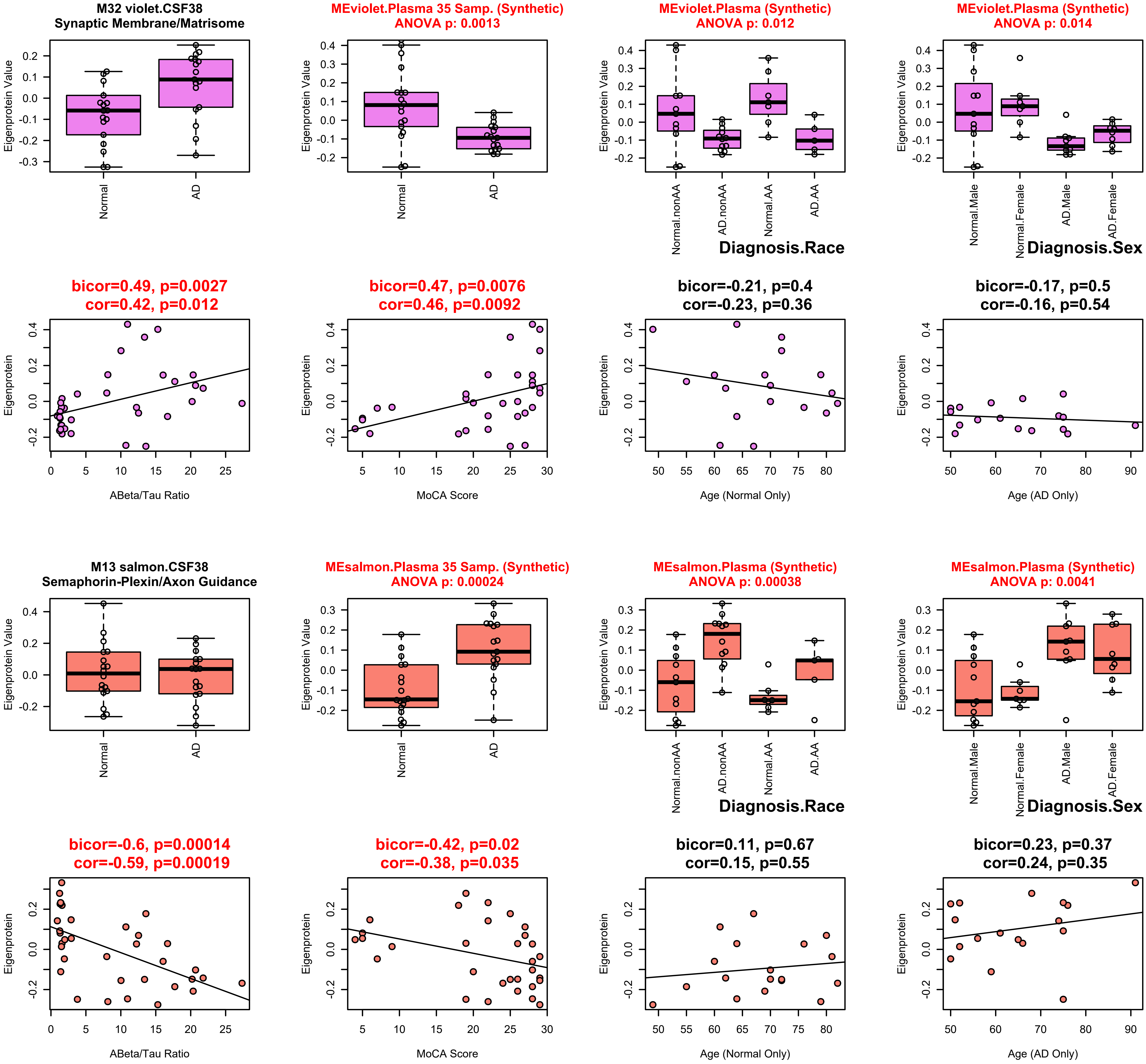
M32 violet.CSF38
Synaptic Membrane/Matrisome
Eigenprotein Value
Normal
AD
MEviolet.Plasma 35 Samp. (Synthetic)
ANOVA p: 0.0013
MEviolet.Plasma (Synthetic)
ANOVA p: 0.012
Normal.nonAA
AD.nonAA
Normal.AA
AD.AA
Diagnosis.Race
MEviolet.Plasma (Synthetic)
ANOVA p: 0.014
Normal.Male
Normal.Female
AD.Male
AD.Female
Diagnosis.Sex
bicor=0.49, p=0.0027
cor=0.42, p=0.012
Eigenprotein
ABeta/Tau Ratio
bicor=0.47, p=0.0076
cor=0.46, p=0.0092
MoCA Score
bicor=-0.21, p=0.4
cor=-0.23, p=0.36
Age (Normal Only)
bicor=-0.17, p=0.5
cor=-0.16, p=0.54
Age (AD Only)
M13 salmon.CSF38
Semaphorin-Plexin/Axon Guidance
MEsalmon.Plasma 35 Samp. (Synthetic)
ANOVA p: 0.00024
MEsalmon.Plasma (Synthetic)
ANOVA p: 0.00038
MEsalmon.Plasma (Synthetic)
ANOVA p: 0.0041
bicor=-0.6, p=0.00014
cor=-0.59, p=0.00019
bicor=-0.42, p=0.02
cor=-0.38, p=0.035
bicor=0.11, p=0.67
cor=0.15, p=0.55
bicor=0.23, p=0.37
cor=0.24, p=0.35

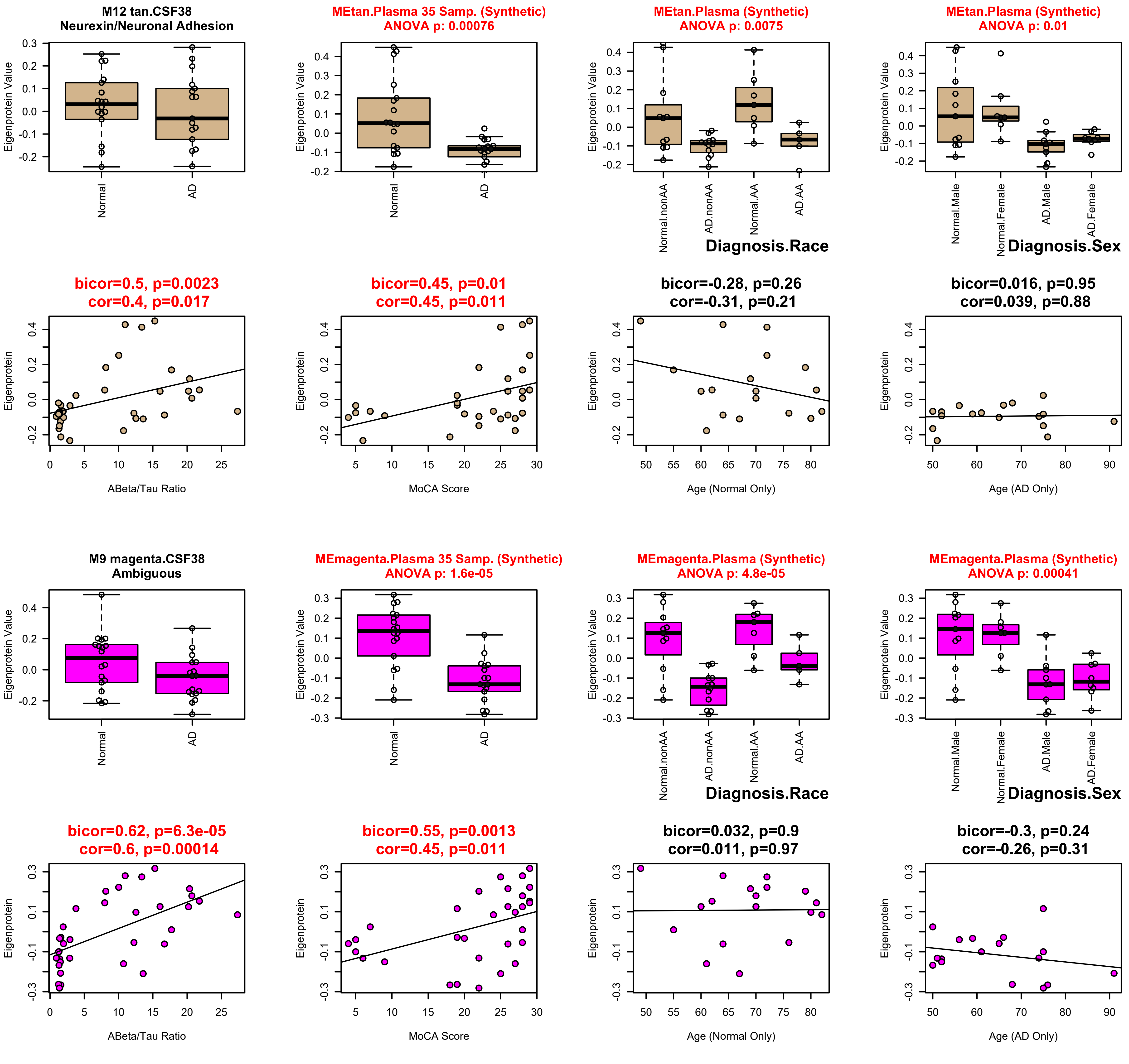
M12 tan.CSF38
Neurexin/Neuronal Adhesion
Eigenprotein Value
Normal
AD
MEtan.Plasma 35 Samp. (Synthetic)
ANOVA p: 0.00076
Eigenprotein Value
Normal
AD
MEtan.Plasma (Synthetic)
ANOVA p: 0.0075
Eigenprotein Value
Normal.nonAA
AD.nonAA
Normal.AA
AD.AA
Diagnosis.Race
MEtan.Plasma (Synthetic)
ANOVA p: 0.01
Eigenprotein Value
Normal.Male
Normal.Female
AD.Male
AD.Female
Diagnosis.Sex
bicor=0.5, p=0.0023
cor=0.4, p=0.017
Eigenprotein
ABeta/Tau Ratio
bicor=0.45, p=0.01
cor=0.45, p=0.011
Eigenprotein
MoCA Score
bicor=-0.28, p=0.26
cor=-0.31, p=0.21
Eigenprotein
Age (Normal Only)
bicor=0.016, p=0.95
cor=0.039, p=0.88
Eigenprotein
Age (AD Only)
M9 magenta.CSF38
Ambiguous
Eigenprotein Value
Normal
AD
MEmagenta.Plasma 35 Samp. (Synthetic)
ANOVA p: 1.6e-05
Eigenprotein Value
Normal
AD
MEmagenta.Plasma (Synthetic)
ANOVA p: 4.8e-05
Eigenprotein Value
Normal.nonAA
AD.nonAA
Normal.AA
AD.AA
Diagnosis.Race
MEmagenta.Plasma (Synthetic)
ANOVA p: 0.00041
Eigenprotein Value
Normal.Male
Normal.Female
AD.Male
AD.Female
Diagnosis.Sex
bicor=0.62, p=6.3e-05
cor=0.6, p=0.00014
Eigenprotein
ABeta/Tau Ratio
bicor=0.55, p=0.0013
cor=0.45, p=0.011
Eigenprotein
MoCA Score
bicor=0.032, p=0.9
cor=0.011, p=0.97
Eigenprotein
Age (Normal Only)
bicor=-0.3, p=0.24
cor=-0.26, p=0.31
Eigenprotein
Age (AD Only)

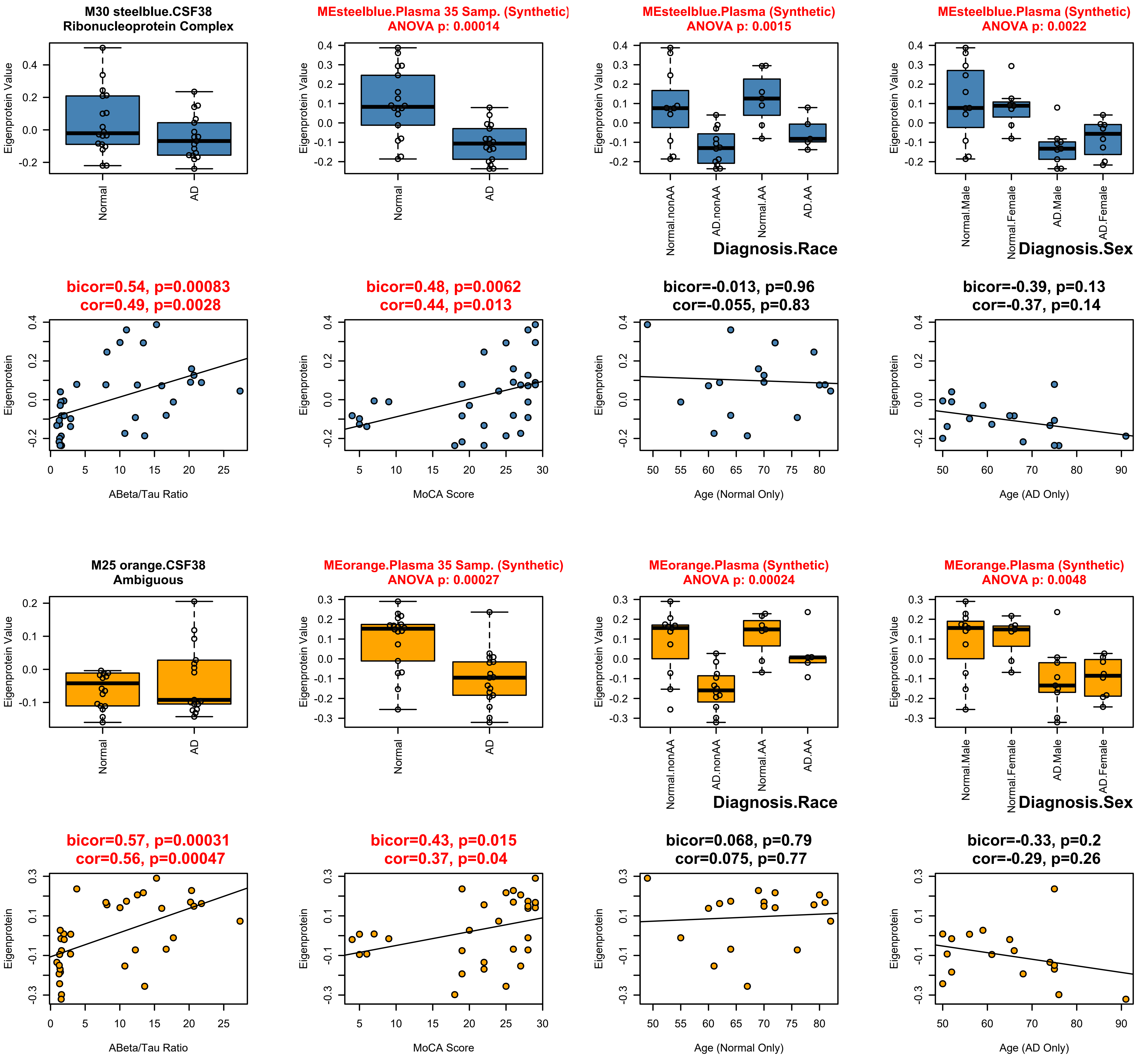

M30 steelblue.CSF38
Ribonucleoprotein Complex
MEsteelblue.Plasma 35 Samp. (Synthetic)
ANOVA p: 0.00014
MEsteelblue.Plasma (Synthetic)
ANOVA p: 0.0015
MEsteelblue.Plasma (Synthetic)
ANOVA p: 0.0022
Eigenprotein Value
Normal
AD
Normal.nonAA
AD.nonAA
Normal.AA
AD.AA
Diagnosis.Race
Normal.Male
Normal.Female
AD.Male
AD.Female
Diagnosis.Sex
bicor=0.54, p=0.00083
cor=0.49, p=0.0028
bicor=0.48, p=0.0062
cor=0.44, p=0.013
bicor=-0.013, p=0.96
cor=-0.055, p=0.83
bicor=-0.39, p=0.13
cor=-0.37, p=0.14
Eigenprotein
ABeta/Tau Ratio
MoCA Score
Age (Normal Only)
Age (AD Only)
M25 orange.CSF38
Ambiguous
MEorange.Plasma 35 Samp. (Synthetic)
ANOVA p: 0.00027
MEorange.Plasma (Synthetic)
ANOVA p: 0.00024
MEorange.Plasma (Synthetic)
ANOVA p: 0.0048
bicor=0.57, p=0.00031
cor=0.56, p=0.00047
bicor=0.43, p=0.015
cor=0.37, p=0.04
bicor=0.068, p=0.79
cor=0.075, p=0.77
bicor=-0.33, p=0.2
cor=-0.29, p=0.26

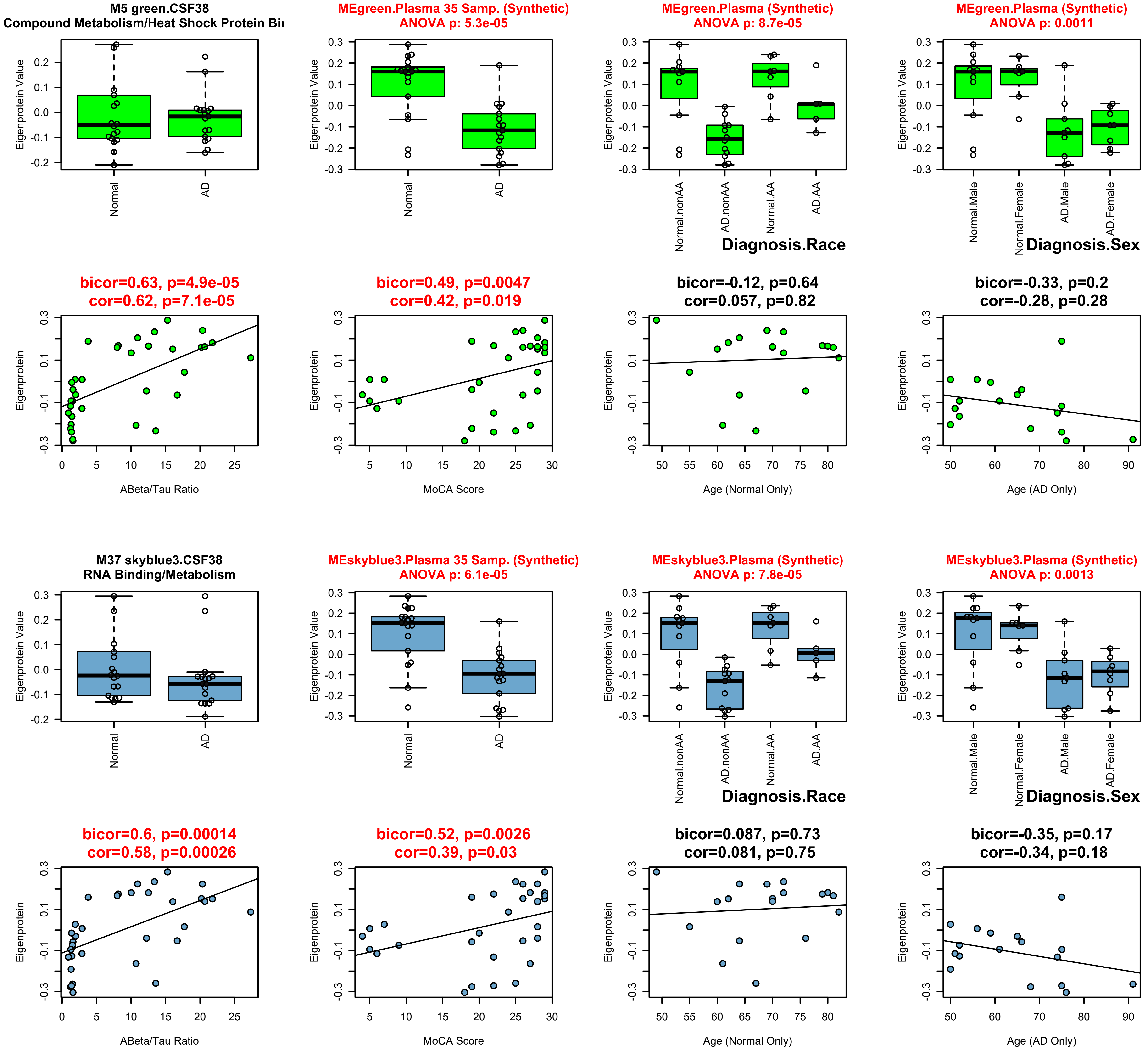

M5 green.CSF38
Compound Metabolism/Heat Shock Protein Bi
Eigenprotein Value
Normal
AD
MEgreen.Plasma 35 Samp. (Synthetic)
ANOVA p: 5.3e-05
MEgreen.Plasma (Synthetic)
ANOVA p: 8.7e-05
Normal.nonAA
AD.nonAA
Normal.AA
AD.AA
Diagnosis.Race
MEgreen.Plasma (Synthetic)
ANOVA p: 0.0011
Normal.Male
Normal.Female
AD.Male
AD.Female
Diagnosis.Sex
bicor=0.63, p=4.9e-05
cor=0.62, p=7.1e-05
Eigenprotein
ABeta/Tau Ratio
bicor=0.49, p=0.0047
cor=0.42, p=0.019
MoCA Score
bicor=-0.12, p=0.64
cor=0.057, p=0.82
Age (Normal Only)
bicor=-0.33, p=0.2
cor=-0.28, p=0.28
Age (AD Only)
M37 skyblue3.CSF38
RNA Binding/Metabolism
MEskyblue3.Plasma 35 Samp. (Synthetic)
ANOVA p: 6.1e-05
MEskyblue3.Plasma (Synthetic)
ANOVA p: 7.8e-05
MEskyblue3.Plasma (Synthetic)
ANOVA p: 0.0013
bicor=0.6, p=0.00014
cor=0.58, p=0.00026
bicor=0.52, p=0.0026
cor=0.39, p=0.03
bicor=0.087, p=0.73
cor=0.081, p=0.75
bicor=-0.35, p=0.17
cor=-0.34, p=0.18

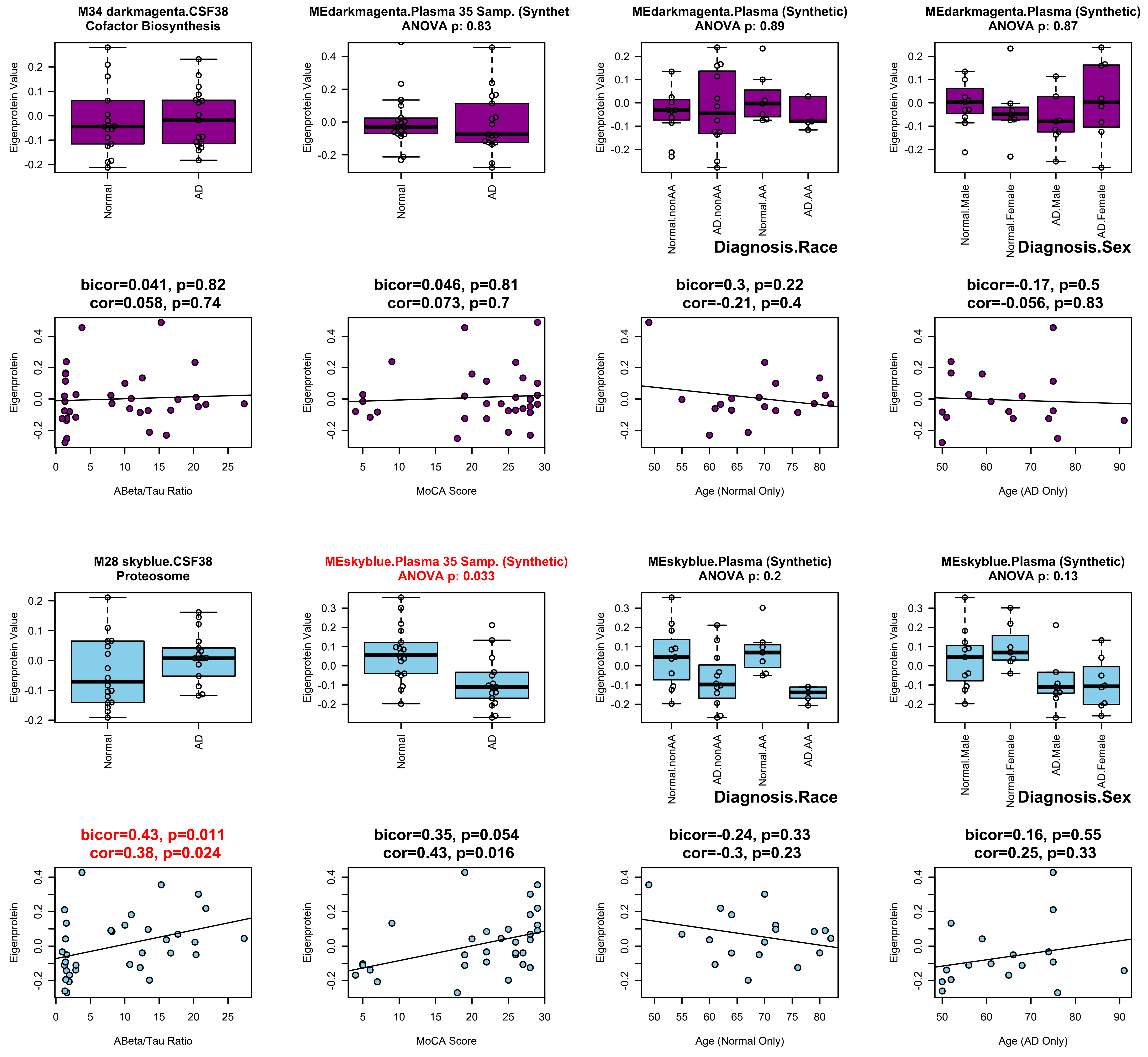
M34 darkmagenta.CSF38
Cofactor Biosynthesis
MEdarkmagenta.Plasma 35 Samp. (Syntheti
ANOVA p: 0.83
MEdarkmagenta.Plasma (Synthetic)
ANOVA p: 0.89
MEdarkmagenta.Plasma (Synthetic)
ANOVA p: 0.87
Eigenprotein Value
Normal
AD
Normal.nonAA
AD.nonAA
Normal.AA
AD.AA
Diagnosis.Race
Normal.Male
Normal.Female
AD.Male
AD.Female
Diagnosis.Sex
bicor=0.041, p=0.82
cor=0.058, p=0.74
bicor=0.046, p=0.81
cor=0.073, p=0.7
bicor=0.3, p=0.22
cor=-0.21, p=0.4
bicor=-0.17, p=0.5
cor=-0.056, p=0.83
Eigenprotein
ABeta/Tau Ratio
MoCA Score
Age (Normal Only)
Age (AD Only)
M28 skyblue.CSF38
Proteosome
MEskyblue.Plasma 35 Samp. (Synthetic)
ANOVA p: 0.033
MEskyblue.Plasma (Synthetic)
ANOVA p: 0.2
MEskyblue.Plasma (Synthetic)
ANOVA p: 0.13
Eigenprotein Value
Normal
AD
Normal.nonAA
AD.nonAA
Normal.AA
AD.AA
Diagnosis.Race
Normal.Male
Normal.Female
AD.Male
AD.Female
Diagnosis.Sex
bicor=0.43, p=0.011
cor=0.38, p=0.024
bicor=0.35, p=0.054
cor=0.43, p=0.016
bicor=-0.24, p=0.33
cor=-0.3, p=0.23
bicor=0.16, p=0.55
cor=0.25, p=0.33
Eigenprotein
ABeta/Tau Ratio
MoCA Score
Age (Normal Only)
Age (AD Only)

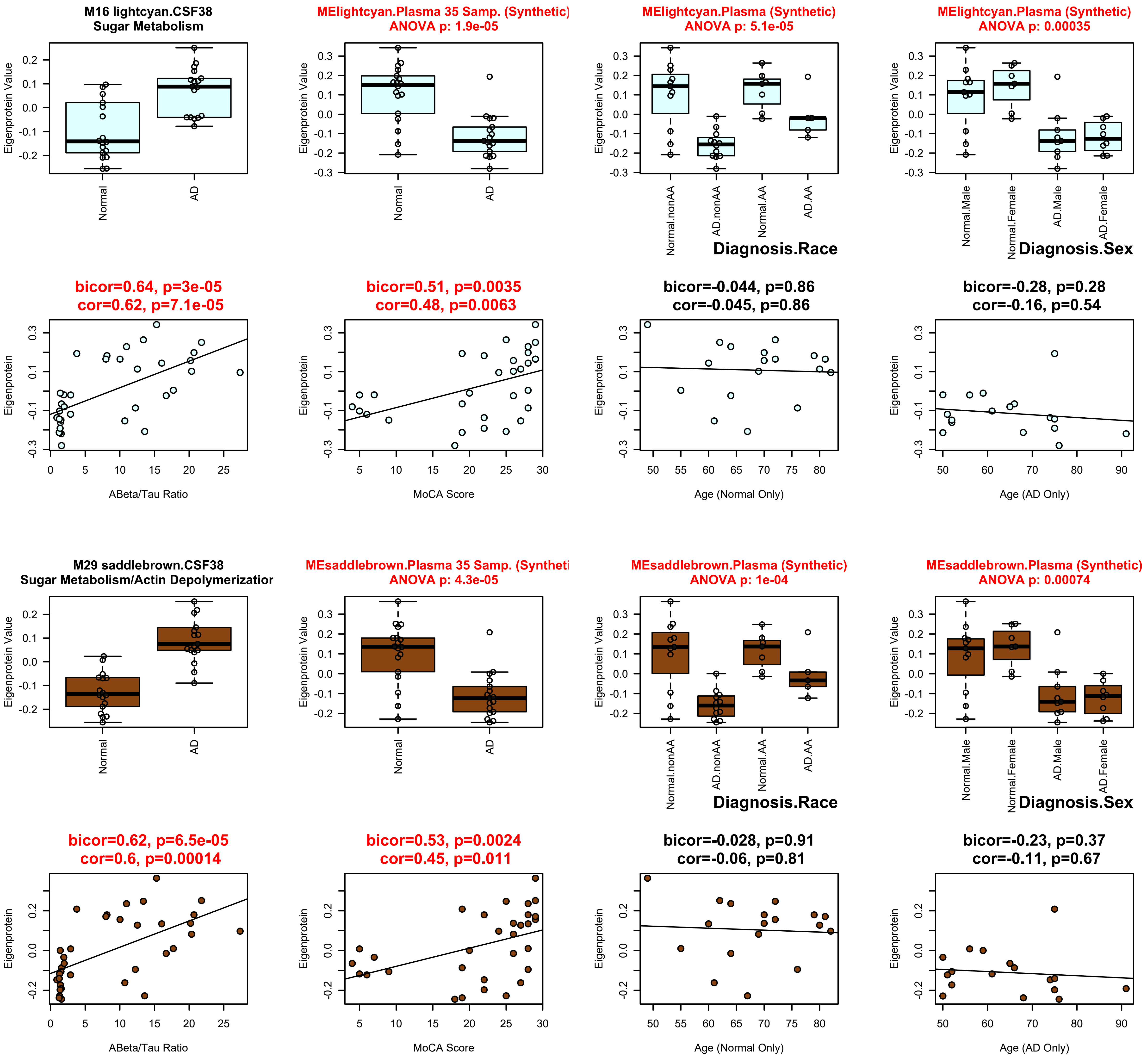

M16 lightcyan.CSF38
Sugar Metabolism
MElightcyan.Plasma 35 Samp. (Synthetic)
ANOVA p: 1.9e-05
MElightcyan.Plasma (Synthetic)
ANOVA p: 5.1e-05
MElightcyan.Plasma (Synthetic)
ANOVA p: 0.00035
Eigenprotein Value
Normal
AD
Normal.nonAA
AD.nonAA
Normal.AA
AD.AA
Diagnosis.Race
Normal.Male
Normal.Female
AD.Male
AD.Female
Diagnosis.Sex
bicor=0.64, p=3e-05
cor=0.62, p=7.1e-05
bicor=0.51, p=0.0035
cor=0.48, p=0.0063
bicor=-0.044, p=0.86
cor=-0.045, p=0.86
bicor=-0.28, p=0.28
cor=-0.16, p=0.54
Eigenprotein
ABeta/Tau Ratio
MoCA Score
Age (Normal Only)
Age (AD Only)
M29 saddlebrown.CSF38
Sugar Metabolism/Actin Depolymerizatior
MEsaddlebrown.Plasma 35 Samp. (Syntheti
ANOVA p: 4.3e-05
MEsaddlebrown.Plasma (Synthetic)
ANOVA p: 1e-04
MEsaddlebrown.Plasma (Synthetic)
ANOVA p: 0.00074
Eigenprotein Value
Normal
AD
Normal.nonAA
AD.nonAA
Normal.AA
AD.AA
Diagnosis.Race
Normal.Male
Normal.Female
AD.Male
AD.Female
Diagnosis.Sex
bicor=0.62, p=6.5e-05
cor=0.6, p=0.00014
bicor=0.53, p=0.0024
cor=0.45, p=0.011
bicor=-0.028, p=0.91
cor=-0.06, p=0.81
bicor=-0.23, p=0.37
cor=-0.11, p=0.67
Eigenprotein
ABeta/Tau Ratio
MoCA Score
Age (Normal Only)
Age (AD Only)

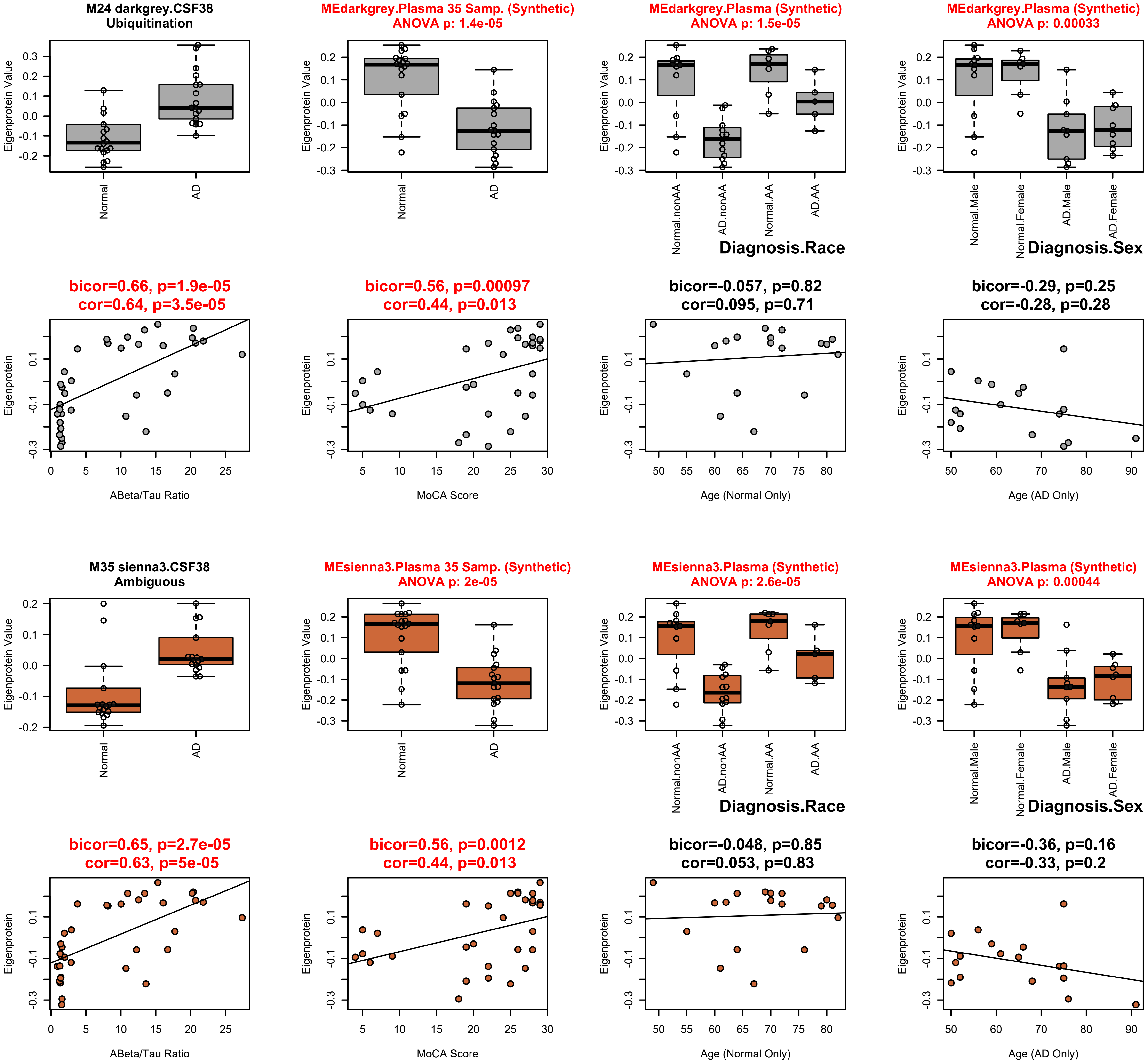
M24 darkgrey.CSF38
Ubiquitination
Eigenprotein Value
Normal
AD
MEdarkgrey.Plasma 35 Samp. (Synthetic)
ANOVA p: 1.4e-05
Eigenprotein Value
Normal
AD
MEdarkgrey.Plasma (Synthetic)
ANOVA p: 1.5e-05
Eigenprotein Value
Normal.nonAA
AD.nonAA
Normal.AA
AD.AA
Diagnosis.Race
MEdarkgrey.Plasma (Synthetic)
ANOVA p: 0.00033
Eigenprotein Value
Normal.Male
Normal.Female
AD.Male
AD.Female
Diagnosis.Sex
bicor=0.66, p=1.9e-05
cor=0.64, p=3.5e-05
Eigenprotein
ABeta/Tau Ratio
bicor=0.56, p=0.00097
cor=0.44, p=0.013
Eigenprotein
MoCA Score
bicor=-0.057, p=0.82
cor=0.095, p=0.71
Eigenprotein
Age (Normal Only)
bicor=-0.29, p=0.25
cor=-0.28, p=0.28
Eigenprotein
Age (AD Only)
M35 sienna3.CSF38
Ambiguous
Eigenprotein Value
Normal
AD
MEsienna3.Plasma 35 Samp. (Synthetic)
ANOVA p: 2e-05
Eigenprotein Value
Normal
AD
MEsienna3.Plasma (Synthetic)
ANOVA p: 2.6e-05
Eigenprotein Value
Normal.nonAA
AD.nonAA
Normal.AA
AD.AA
Diagnosis.Race
MEsienna3.Plasma (Synthetic)
ANOVA p: 0.00044
Eigenprotein Value
Normal.Male
Normal.Female
AD.Male
AD.Female
Diagnosis.Sex
bicor=0.65, p=2.7e-05
cor=0.63, p=5e-05
Eigenprotein
ABeta/Tau Ratio
bicor=0.56, p=0.0012
cor=0.44, p=0.013
Eigenprotein
MoCA Score
bicor=-0.048, p=0.85
cor=0.053, p=0.83
Eigenprotein
Age (Normal Only)
bicor=-0.36, p=0.16
cor=-0.33, p=0.2
Eigenprotein
Age (AD Only)

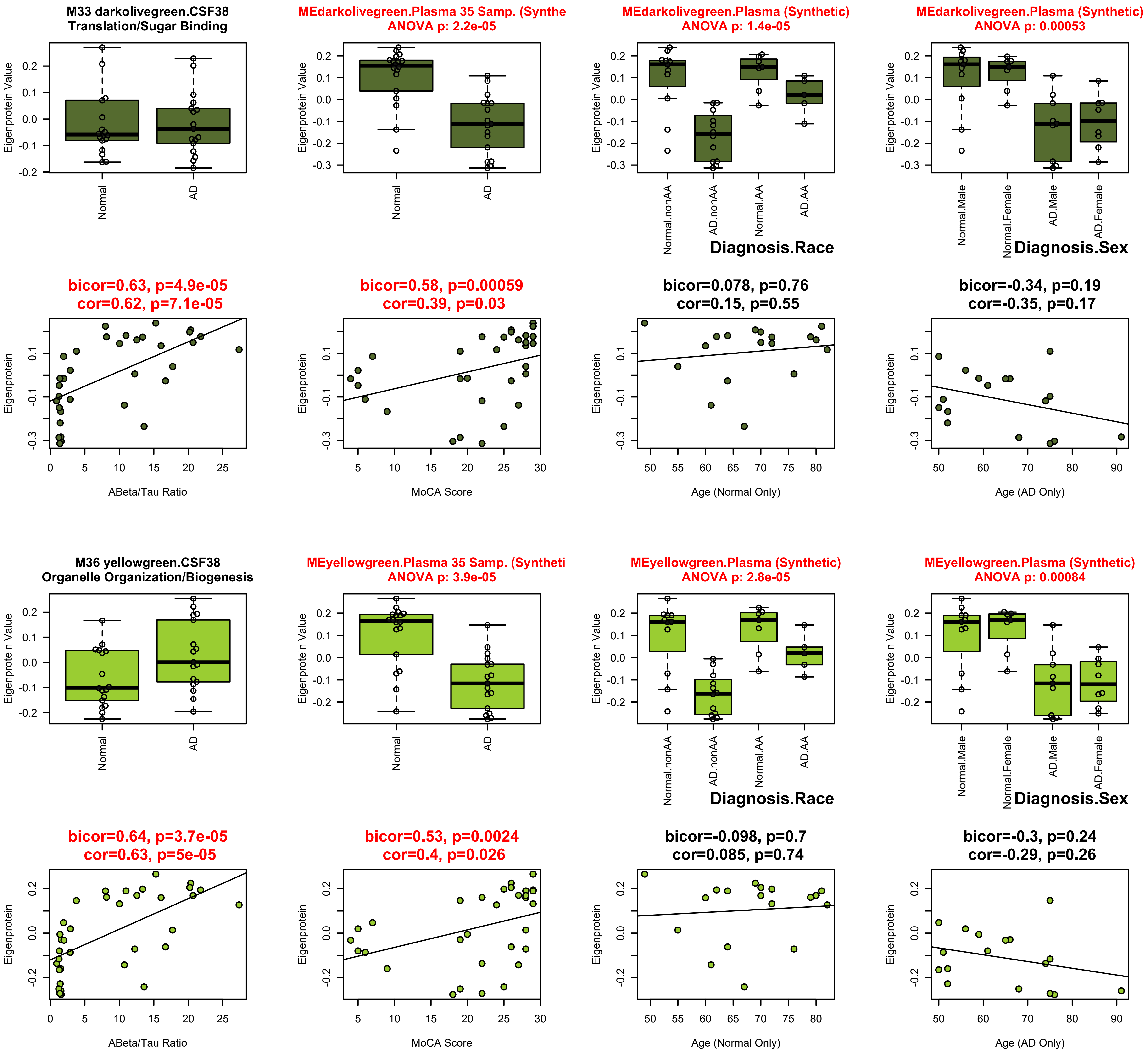

M33 darkolivegreen.CSF38
Translation/Sugar Binding
MEdarkolivegreen.Plasma 35 Samp. (Synthe
ANOVA p: 2.2e-05
MEdarkolivegreen.Plasma (Synthetic)
ANOVA p: 1.4e-05
MEdarkolivegreen.Plasma (Synthetic)
ANOVA p: 0.00053
Eigenprotein Value
Normal
AD
Normal.nonAA
AD.nonAA
Normal.AA
AD.AA
Diagnosis.Race
Normal.Male
Normal.Female
AD.Male
AD.Female
Diagnosis.Sex
bicor=0.63, p=4.9e-05
cor=0.62, p=7.1e-05
bicor=0.58, p=0.00059
cor=0.39, p=0.03
bicor=0.078, p=0.76
cor=0.15, p=0.55
bicor=-0.34, p=0.19
cor=-0.35, p=0.17
Eigenprotein
ABeta/Tau Ratio
MoCA Score
Age (Normal Only)
Age (AD Only)
M36 yellowgreen.CSF38
Organelle Organization/Biogenesis
MEyellowgreen.Plasma 35 Samp. (Syntheti
ANOVA p: 3.9e-05
MEyellowgreen.Plasma (Synthetic)
ANOVA p: 2.8e-05
MEyellowgreen.Plasma (Synthetic)
ANOVA p: 0.00084
bicor=0.64, p=3.7e-05
cor=0.63, p=5e-05
bicor=0.53, p=0.0024
cor=0.4, p=0.026
bicor=-0.098, p=0.7
cor=0.085, p=0.74
bicor=-0.3, p=0.24
cor=-0.29, p=0.26

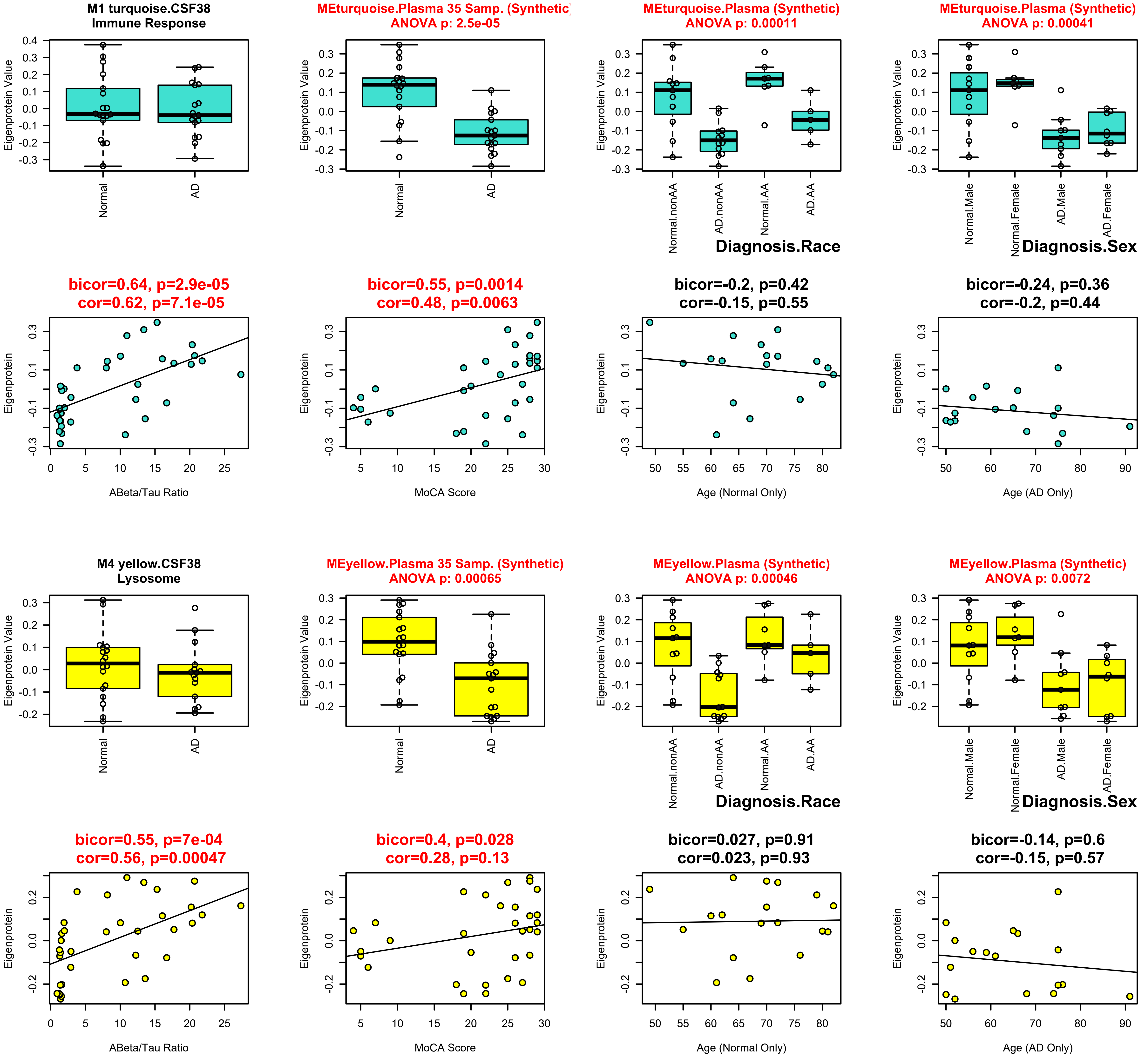
M1 turquoise.CSF38
Immune Response
Eigenprotein Value
Normal
AD
MEturquoise.Plasma 35 Samp. (Synthetic)
ANOVA p: 2.5e-05
Eigenprotein Value
Normal
AD
MEturquoise.Plasma (Synthetic)
ANOVA p: 0.00011
Eigenprotein Value
Normal.nonAA
AD.nonAA
Normal.AA
AD.AA
Diagnosis.Race
MEturquoise.Plasma (Synthetic)
ANOVA p: 0.00041
Eigenprotein Value
Normal.Male
Normal.Female
AD.Male
AD.Female
Diagnosis.Sex
bicor=0.64, p=2.9e-05
cor=0.62, p=7.1e-05
Eigenprotein
ABeta/Tau Ratio
bicor=0.55, p=0.0014
cor=0.48, p=0.0063
Eigenprotein
MoCA Score
bicor=-0.2, p=0.42
cor=-0.15, p=0.55
Eigenprotein
Age (Normal Only)
bicor=-0.24, p=0.36
cor=-0.2, p=0.44
Eigenprotein
Age (AD Only)
M4 yellow.CSF38
Lysosome
Eigenprotein Value
Normal
AD
MEyellow.Plasma 35 Samp. (Synthetic)
ANOVA p: 0.00065
Eigenprotein Value
Normal
AD
MEyellow.Plasma (Synthetic)
ANOVA p: 0.00046
Eigenprotein Value
Normal.nonAA
AD.nonAA
Normal.AA
AD.AA
Diagnosis.Race
MEyellow.Plasma (Synthetic)
ANOVA p: 0.0072
Eigenprotein Value
Normal.Male
Normal.Female
AD.Male
AD.Female
Diagnosis.Sex
bicor=0.55, p=7e-04
cor=0.56, p=0.00047
Eigenprotein
ABeta/Tau Ratio
bicor=0.4, p=0.028
cor=0.28, p=0.13
Eigenprotein
MoCA Score
bicor=0.027, p=0.91
cor=0.023, p=0.93
Eigenprotein
Age (Normal Only)
bicor=-0.14, p=0.6
cor=-0.15, p=0.57
Eigenprotein
Age (AD Only)

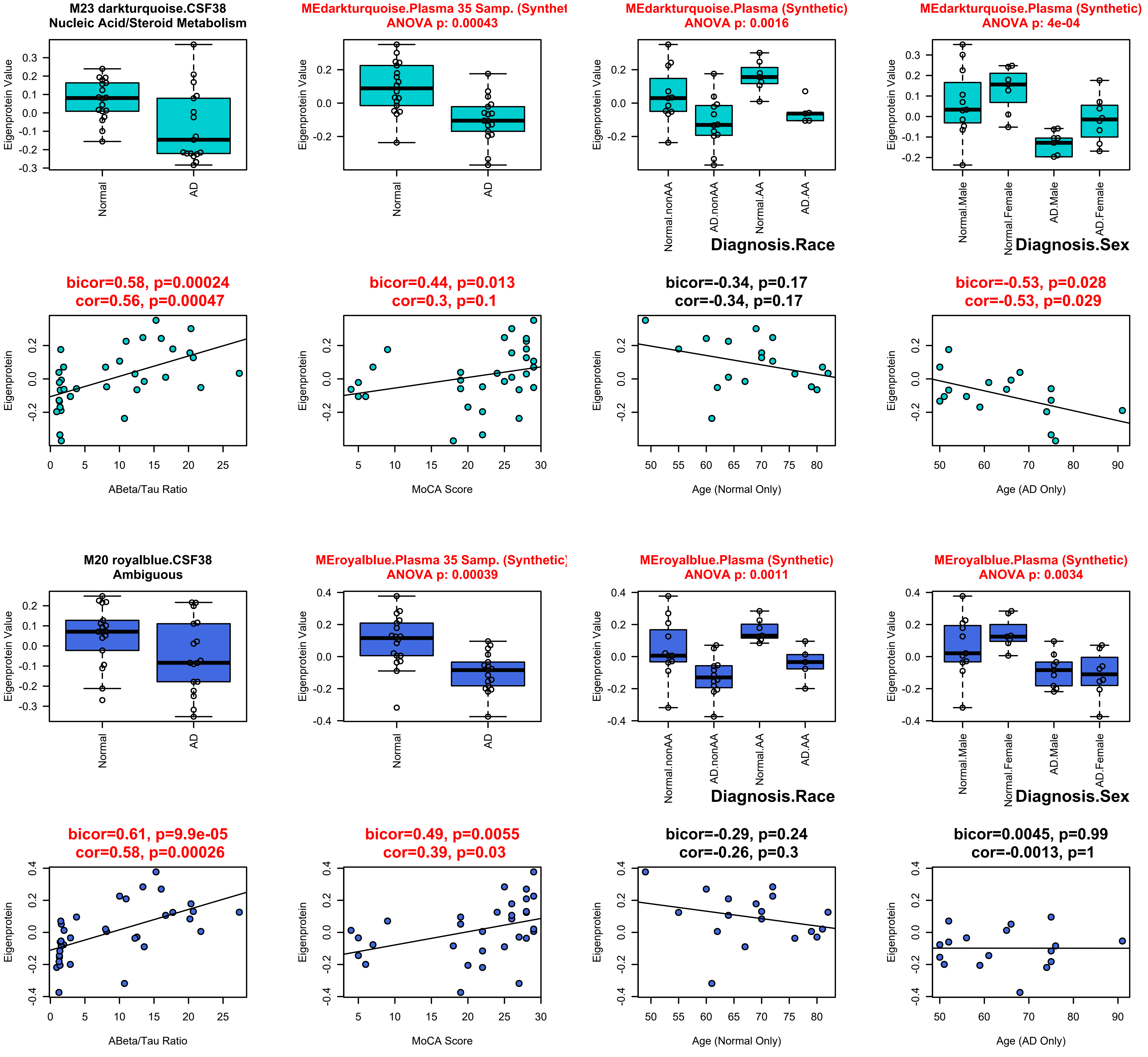
M23 darkturquoise.CSF38
Nucleic Acid/Steroid Metabolism
MEdarkturquoise.Plasma 35 Samp. (Synthet
ANOVA p: 0.00043
MEdarkturquoise.Plasma (Synthetic)
ANOVA p: 0.0016
MEdarkturquoise.Plasma (Synthetic)
ANOVA p: 4e-04
Eigenprotein Value
Normal
AD
Normal.nonAA
AD.nonAA
Normal.AA
AD.AA
Diagnosis.Race
Normal.Male
Normal.Female
AD.Male
AD.Female
Diagnosis.Sex
bicor=0.58, p=0.00024
cor=0.56, p=0.00047
bicor=0.44, p=0.013
cor=0.3, p=0.1
bicor=-0.34, p=0.17
cor=-0.34, p=0.17
bicor=-0.53, p=0.028
cor=-0.53, p=0.029
Eigenprotein
ABeta/Tau Ratio
MoCA Score
Age (Normal Only)
Age (AD Only)
M20 royalblue.CSF38
Ambiguous
MEroyalblue.Plasma 35 Samp. (Synthetic)
ANOVA p: 0.00039
MEroyalblue.Plasma (Synthetic)
ANOVA p: 0.0011
MEroyalblue.Plasma (Synthetic)
ANOVA p: 0.0034
bicor=0.61, p=9.9e-05
cor=0.58, p=0.00026
bicor=0.49, p=0.0055
cor=0.39, p=0.03
bicor=-0.29, p=0.24
cor=-0.26, p=0.3
bicor=0.0045, p=0.99
cor=-0.0013, p=1

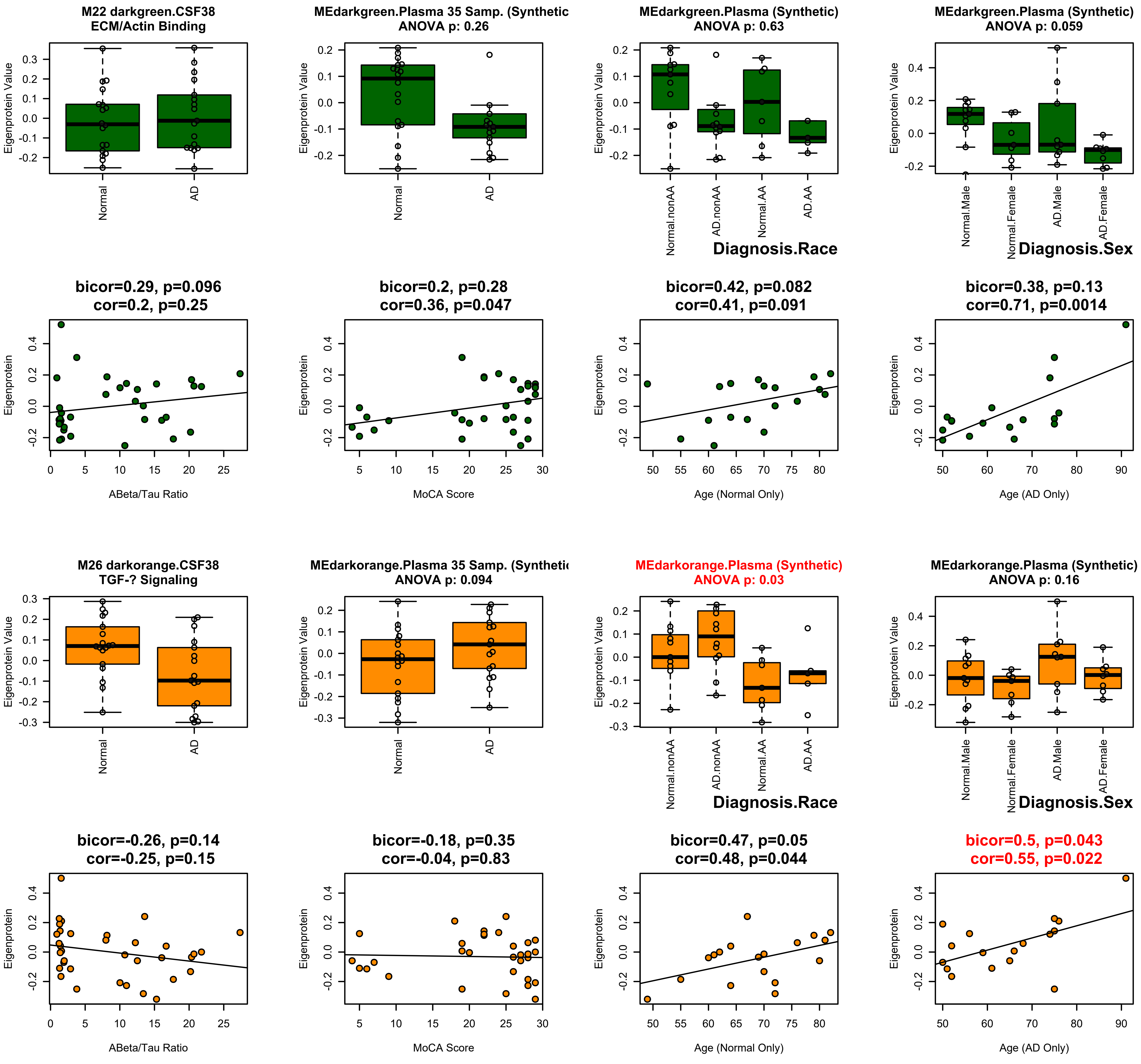
M22 darkgreen.CSF38
ECM/Actin Binding
Eigenprotein Value
Normal
AD
MEdarkgreen.Plasma 35 Samp. (Synthetic
ANOVA p: 0.26
Eigenprotein Value
Normal
AD
MEdarkgreen.Plasma (Synthetic)
ANOVA p: 0.63
Eigenprotein Value
Normal.nonAA
AD.nonAA
Normal.AA
AD.AA
Diagnosis.Race
MEdarkgreen.Plasma (Synthetic)
ANOVA p: 0.059
Eigenprotein Value
Normal.Male
Normal.Female
AD.Male
AD.Female
Diagnosis.Sex
bicor=0.29, p=0.096
cor=0.2, p=0.25
Eigenprotein
ABeta/Tau Ratio
bicor=0.2, p=0.28
cor=0.36, p=0.047
Eigenprotein
MoCA Score
bicor=0.42, p=0.082
cor=0.41, p=0.091
Eigenprotein
Age (Normal Only)
bicor=0.38, p=0.13
cor=0.71, p=0.0014
Eigenprotein
Age (AD Only)
M26 darkorange.CSF38
TGF-? Signaling
Eigenprotein Value
Normal
AD
MEdarkorange.Plasma 35 Samp. (Synthetic
ANOVA p: 0.094
Eigenprotein Value
Normal
AD
MEdarkorange.Plasma (Synthetic)
ANOVA p: 0.03
Eigenprotein Value
Normal.nonAA
AD.nonAA
Normal.AA
AD.AA
Diagnosis.Race
MEdarkorange.Plasma (Synthetic)
ANOVA p: 0.16
Eigenprotein Value
Normal.Male
Normal.Female
AD.Male
AD.Female
Diagnosis.Sex
bicor=-0.26, p=0.14
cor=-0.25, p=0.15
Eigenprotein
ABeta/Tau Ratio
bicor=-0.18, p=0.35
cor=-0.04, p=0.83
Eigenprotein
MoCA Score
bicor=0.47, p=0.05
cor=0.48, p=0.044
Eigenprotein
Age (Normal Only)
bicor=0.5, p=0.043
cor=0.55, p=0.022
Eigenprotein
Age (AD Only)

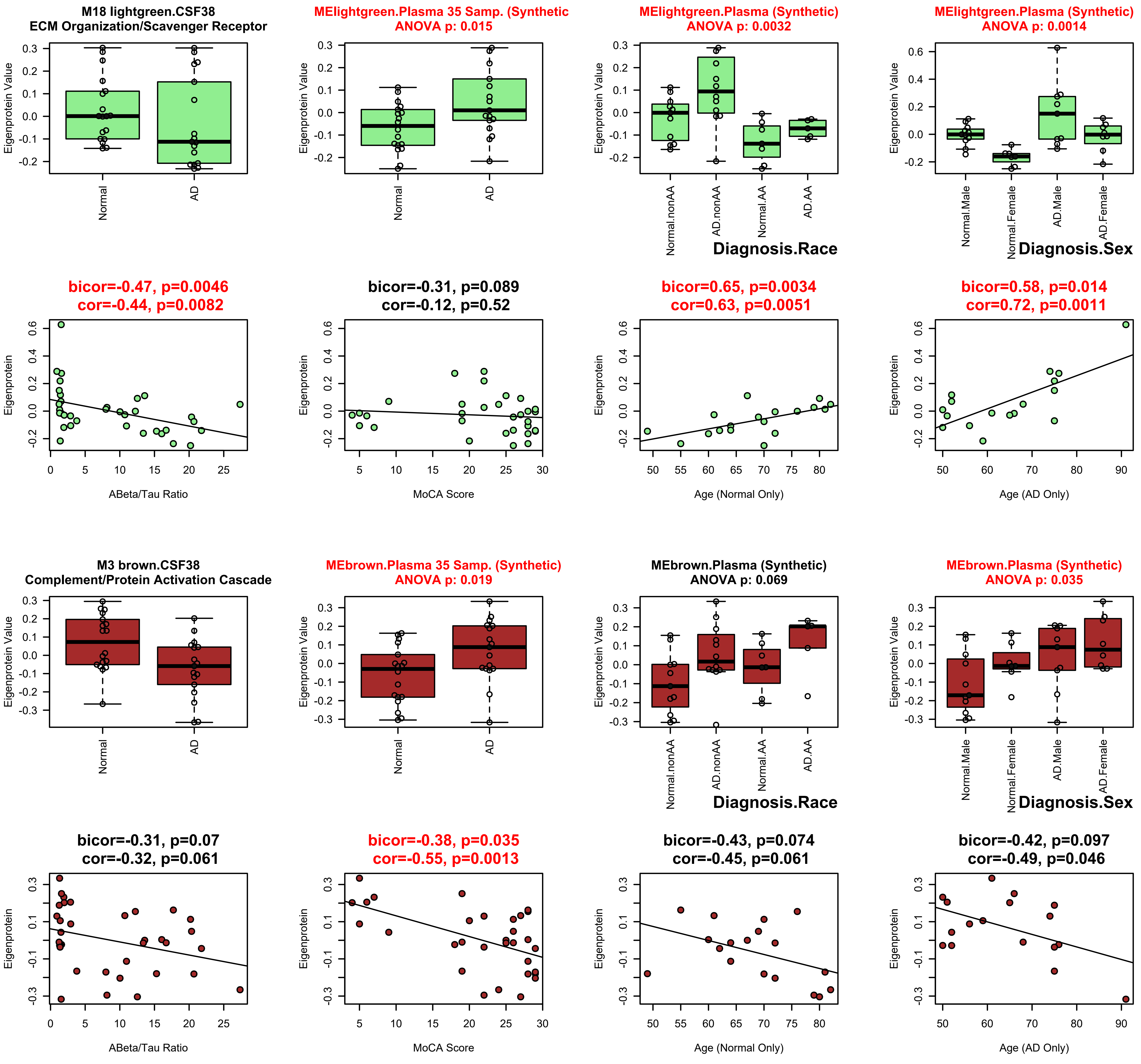
M18 lightgreen.CSF38
ECM Organization/Scavenger Receptor
MElightgreen.Plasma 35 Samp. (Synthetic)
ANOVA p: 0.015
MElightgreen.Plasma (Synthetic)
ANOVA p: 0.0032
MElightgreen.Plasma (Synthetic)
ANOVA p: 0.0014
Eigenprotein Value
Normal
AD
Normal.nonAA
AD.nonAA
Normal.AA
AD.AA
Diagnosis.Race
Normal.Male
Normal.Female
AD.Male
AD.Female
Diagnosis.Sex
bicor=-0.47, p=0.0046
cor=-0.44, p=0.0082
bicor=-0.31, p=0.089
cor=-0.12, p=0.52
bicor=0.65, p=0.0034
cor=0.63, p=0.0051
bicor=0.58, p=0.014
cor=0.72, p=0.0011
Eigenprotein
ABeta/Tau Ratio
MoCA Score
Age (Normal Only)
Age (AD Only)
M3 brown.CSF38
Complement/Protein Activation Cascade
MEbrown.Plasma 35 Samp. (Synthetic)
ANOVA p: 0.019
MEbrown.Plasma (Synthetic)
ANOVA p: 0.069
MEbrown.Plasma (Synthetic)
ANOVA p: 0.035
bicor=-0.31, p=0.07
cor=-0.32, p=0.061
bicor=-0.38, p=0.035
cor=-0.55, p=0.0013
bicor=-0.43, p=0.074
cor=-0.45, p=0.061
bicor=-0.42, p=0.097
cor=-0.49, p=0.046

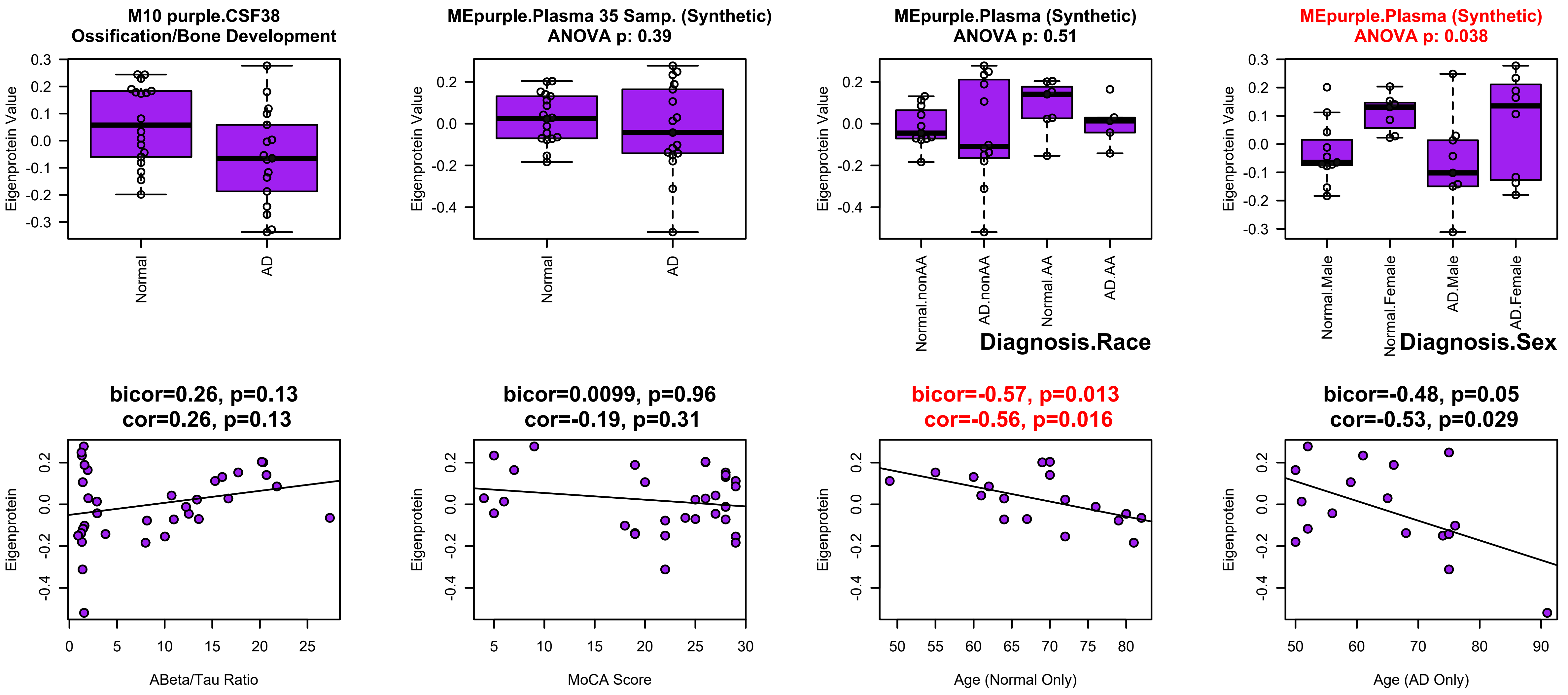

M10 purple.CSF38
Ossification/Bone Development
Eigenprotein Value
Normal
AD
MEpurple.Plasma 35 Samp. (Synthetic)
ANOVA p: 0.39
Eigenprotein Value
Normal
AD
MEpurple.Plasma (Synthetic)
ANOVA p: 0.51
Eigenprotein Value
Normal.nonAA
AD.nonAA
Normal.AA
AD.AA
Diagnosis.Race
MEpurple.Plasma (Synthetic)
ANOVA p: 0.038
Eigenprotein Value
Normal.Male
Normal.Female
AD.Male
AD.Female
Diagnosis.Sex
bicor=0.26, p=0.13
cor=0.26, p=0.13
Eigenprotein
ABeta/Tau Ratio
bicor=0.0099, p=0.96
cor=-0.19, p=0.31
Eigenprotein
MoCA Score
bicor=-0.57, p=0.013
cor=-0.56, p=0.016
Eigenprotein
Age (Normal Only)
bicor=-0.48, p=0.05
cor=-0.53, p=0.029
Eigenprotein
Age (AD Only)

### Data Subset:

### CSF 37 of 38 modules as template

### M8 pink: Autophagy

slopeDiff(AD−CT): −0.49

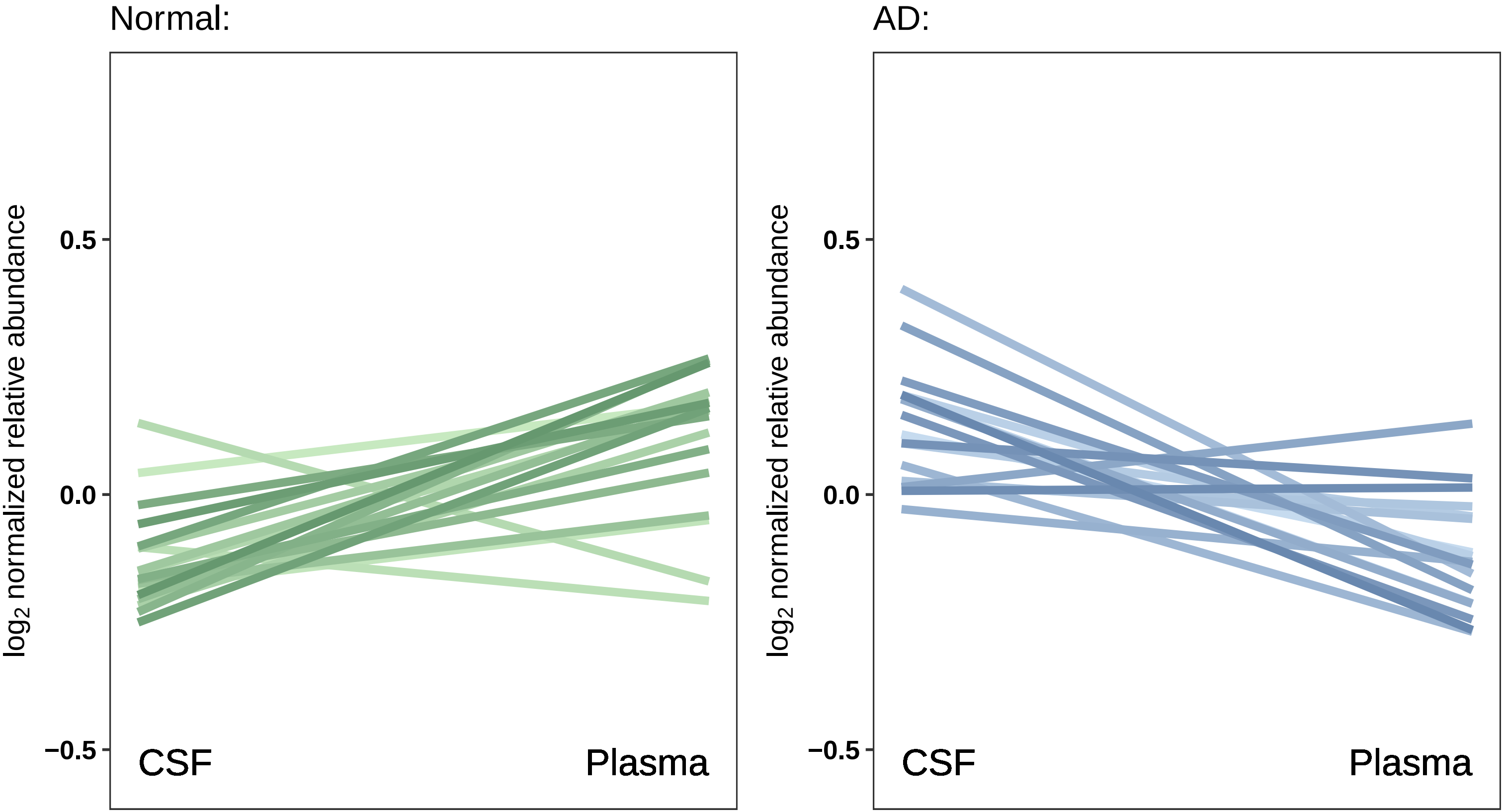

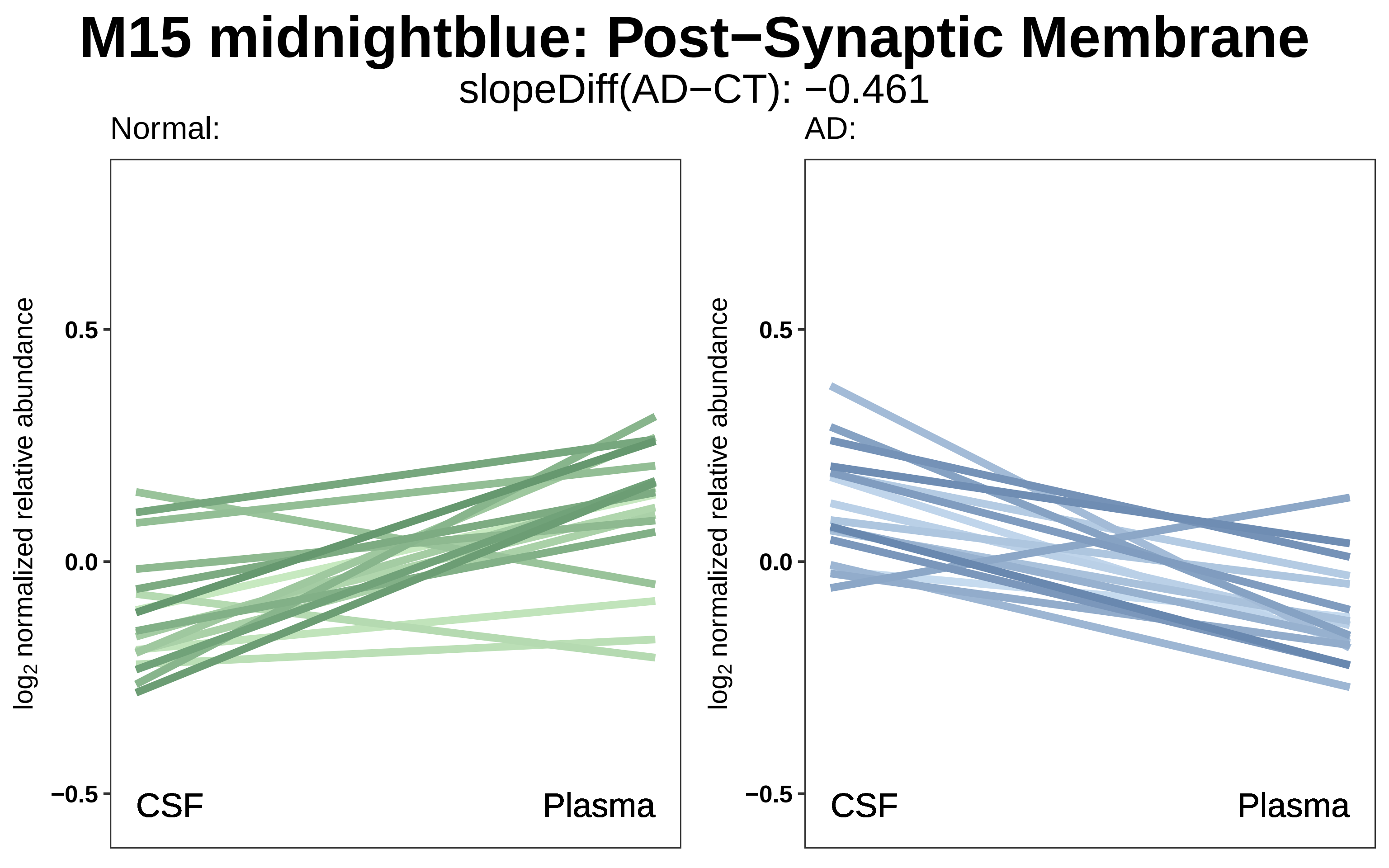
M15 midnightblue: Post−Synaptic Membrane
slopeDiff(AD−CT): −0.461
Normal:
AD:
log2 normalized relative abundance
0.5
0.0
−0.5
CSF
Plasma
log2 normalized relative abundance
0.5
0.0
−0.5
CSF
Plasma

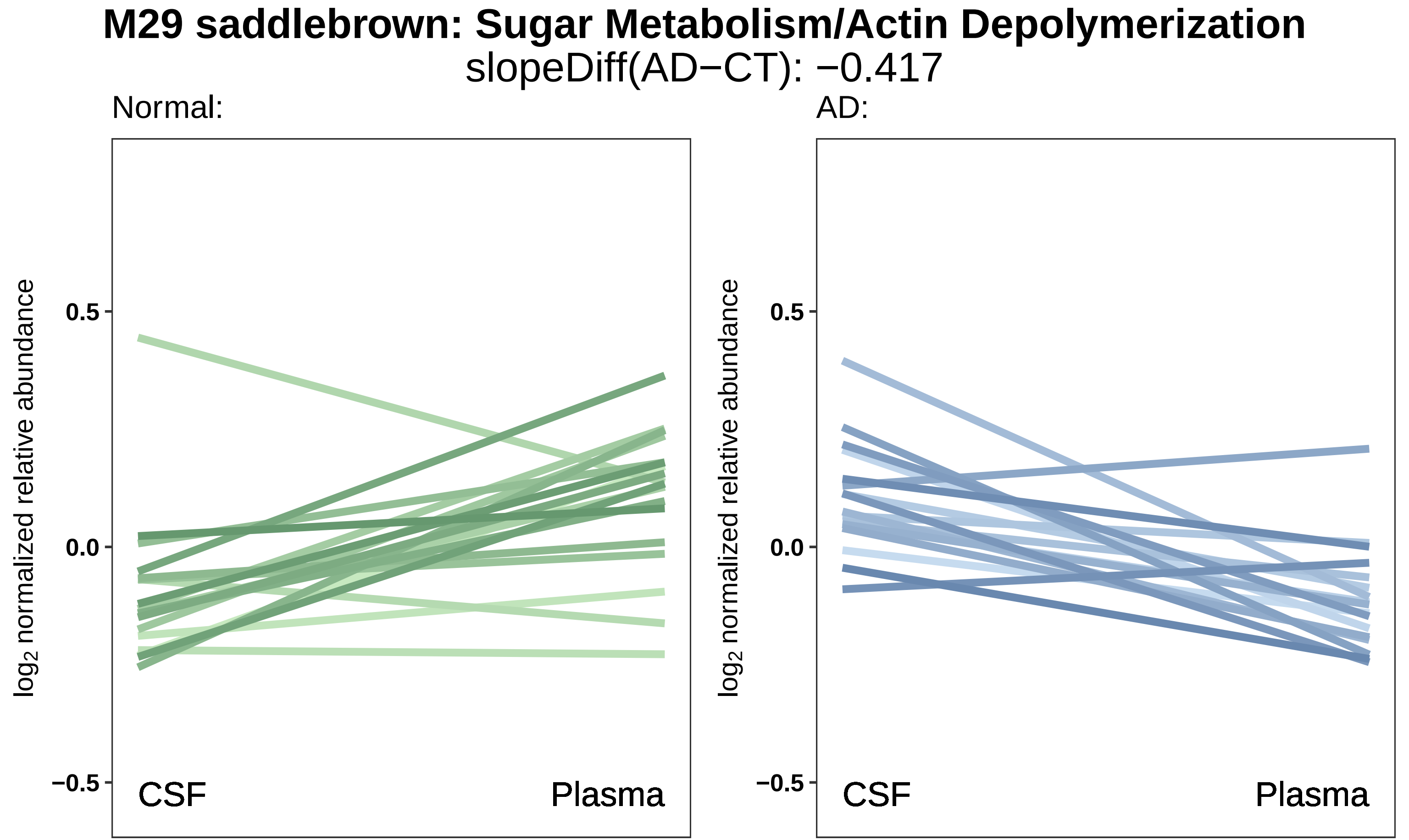

M29 saddlebrown: Sugar Metabolism/Actin Depolymerization
slopeDiff(AD−CT): −0.417
Normal:
AD:
log2 normalized relative abundance
0.5
0.0
−0.5
CSF
Plasma

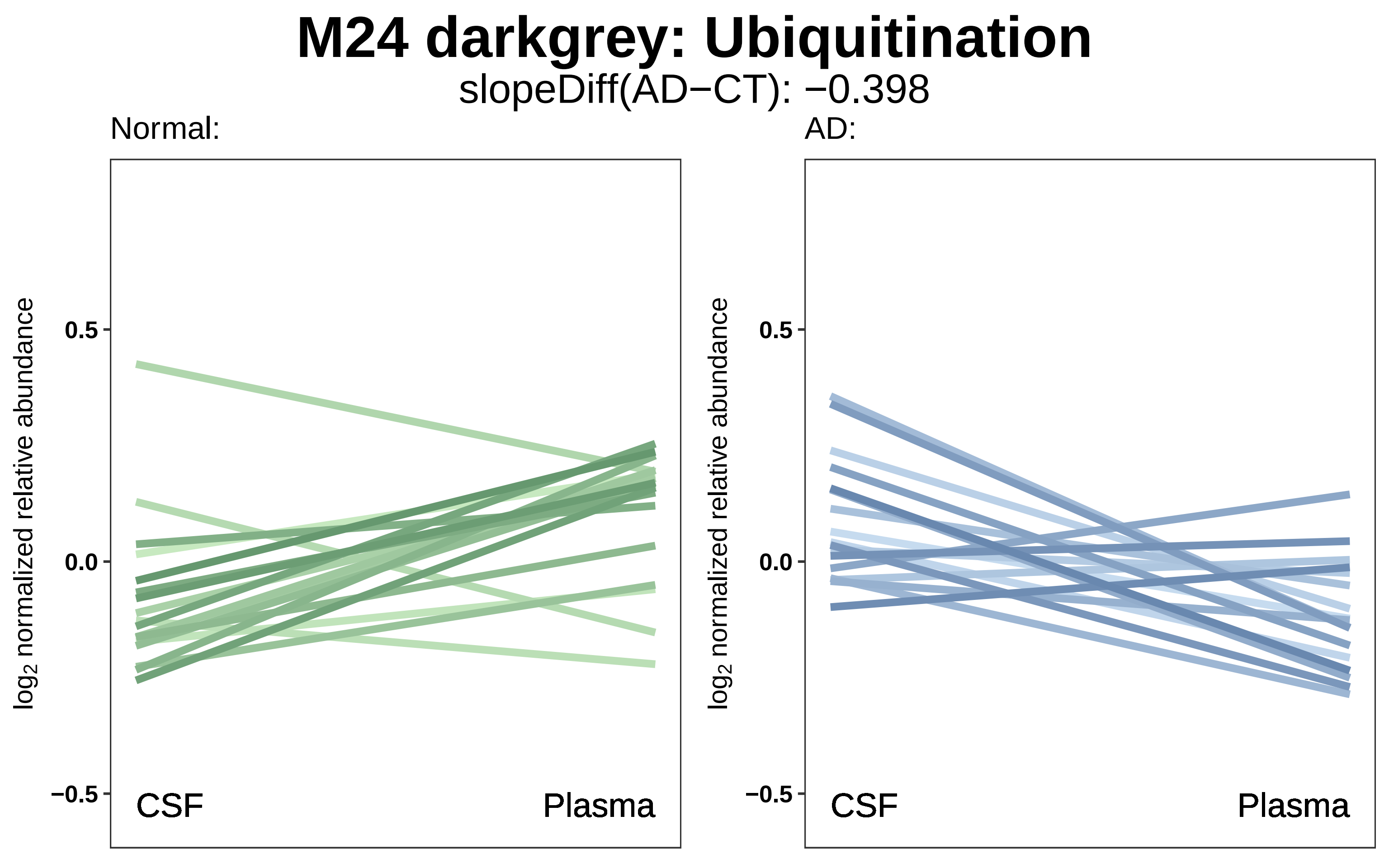

M24 darkgrey: Ubiquitination
slopeDiff(AD−CT): −0.398
Normal:
AD:
log2 normalized relative abundance
0.5
0.0
−0.5
CSF
Plasma

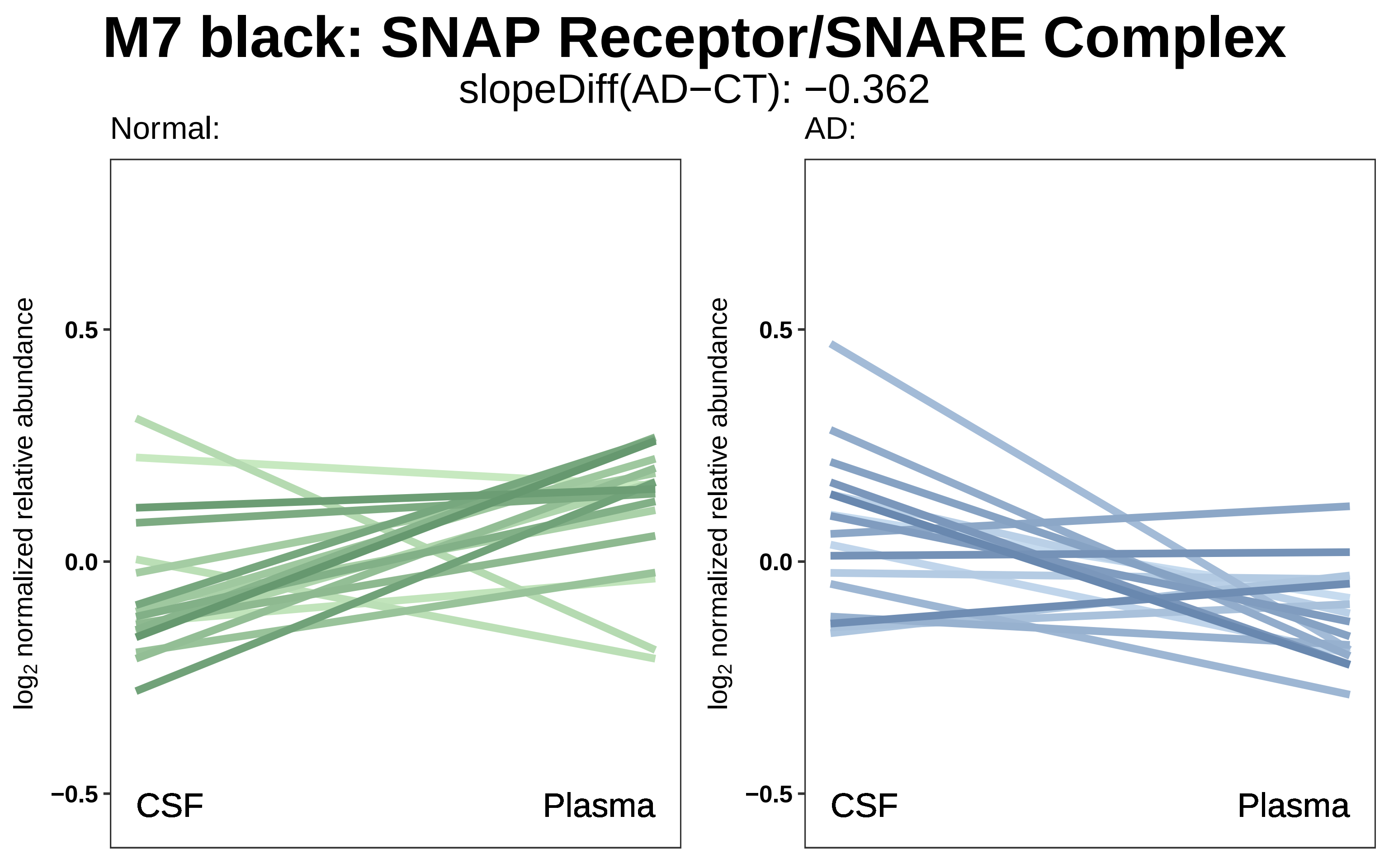

M7 black: SNAP Receptor/SNARE Complex
slopeDiff(AD−CT): −0.362
Normal:
AD:
log2 normalized relative abundance
0.5
0.0
−0.5
CSF
Plasma

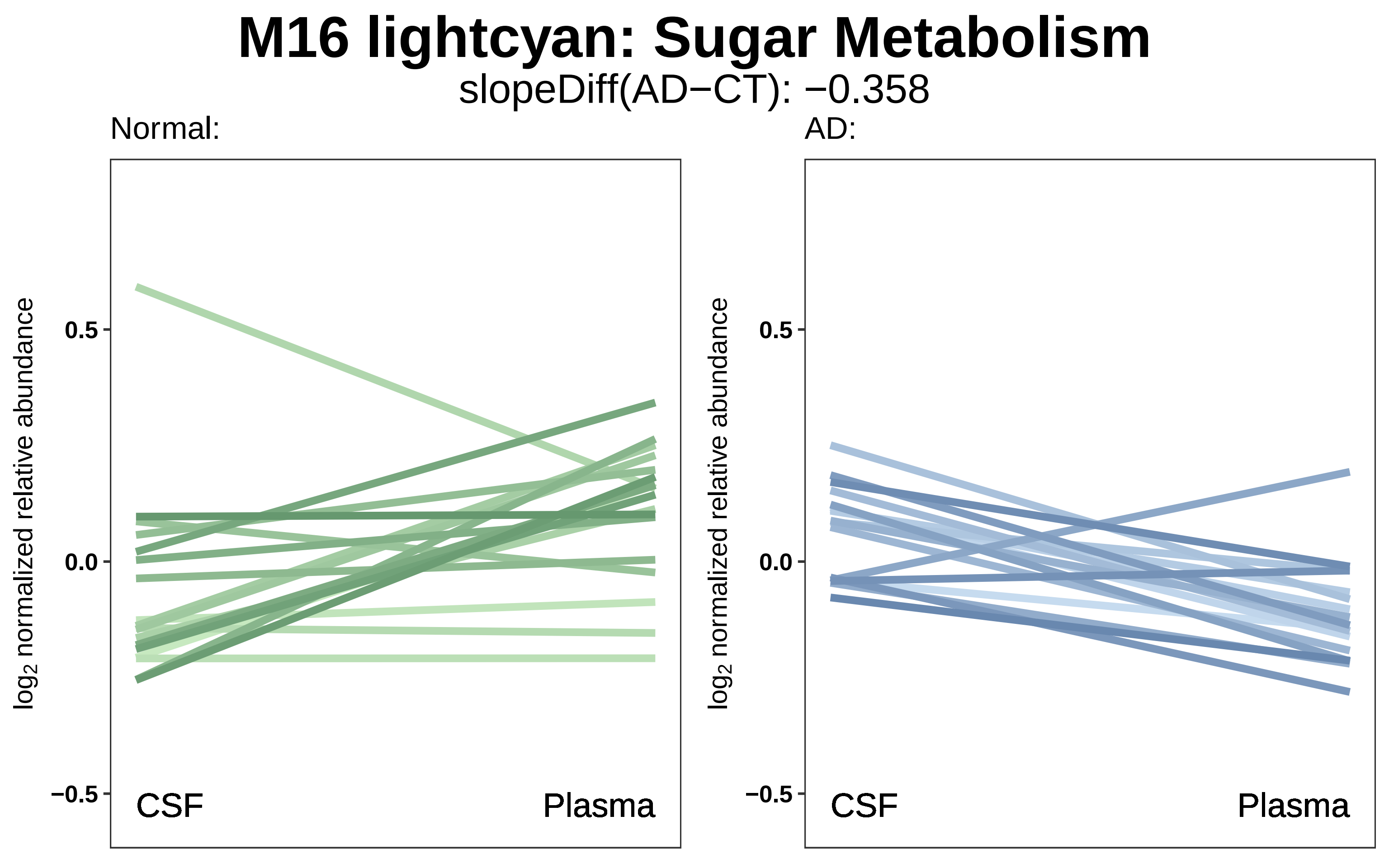

M16 lightcyan: Sugar Metabolism
slopeDiff(AD−CT): −0.358
Normal:
AD:
log2 normalized relative abundance
0.5
0.0
−0.5
CSF
Plasma

### M6 red: Muscle/Neurotransmitter Transport

slopeDiff(AD−CT): −0.331

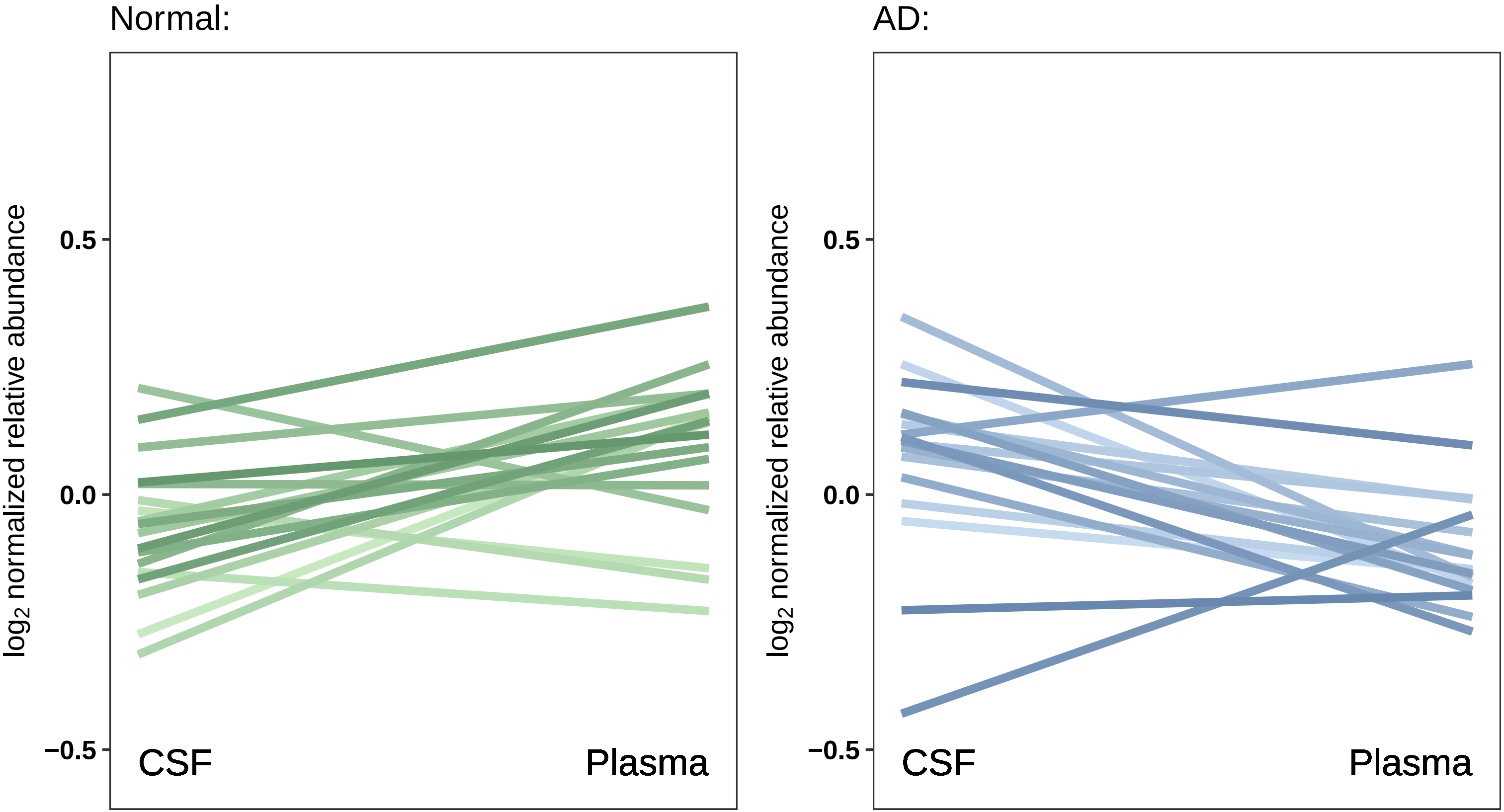

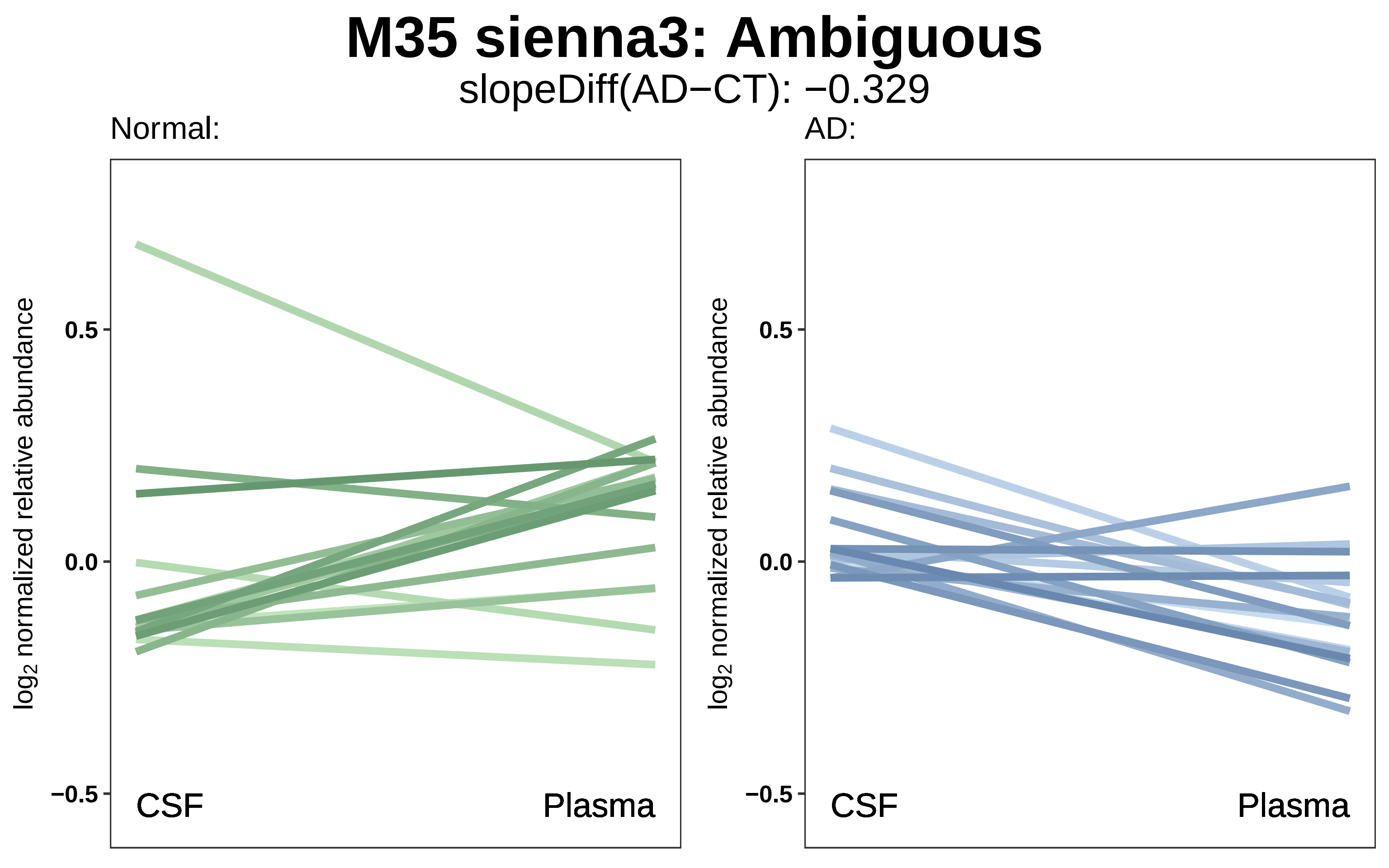

M35 sienna3: Ambiguous
slopeDiff(AD−CT): −0.329
Normal:
AD:
log2 normalized relative abundance
0.5
0.0
−0.5
CSF
Plasma

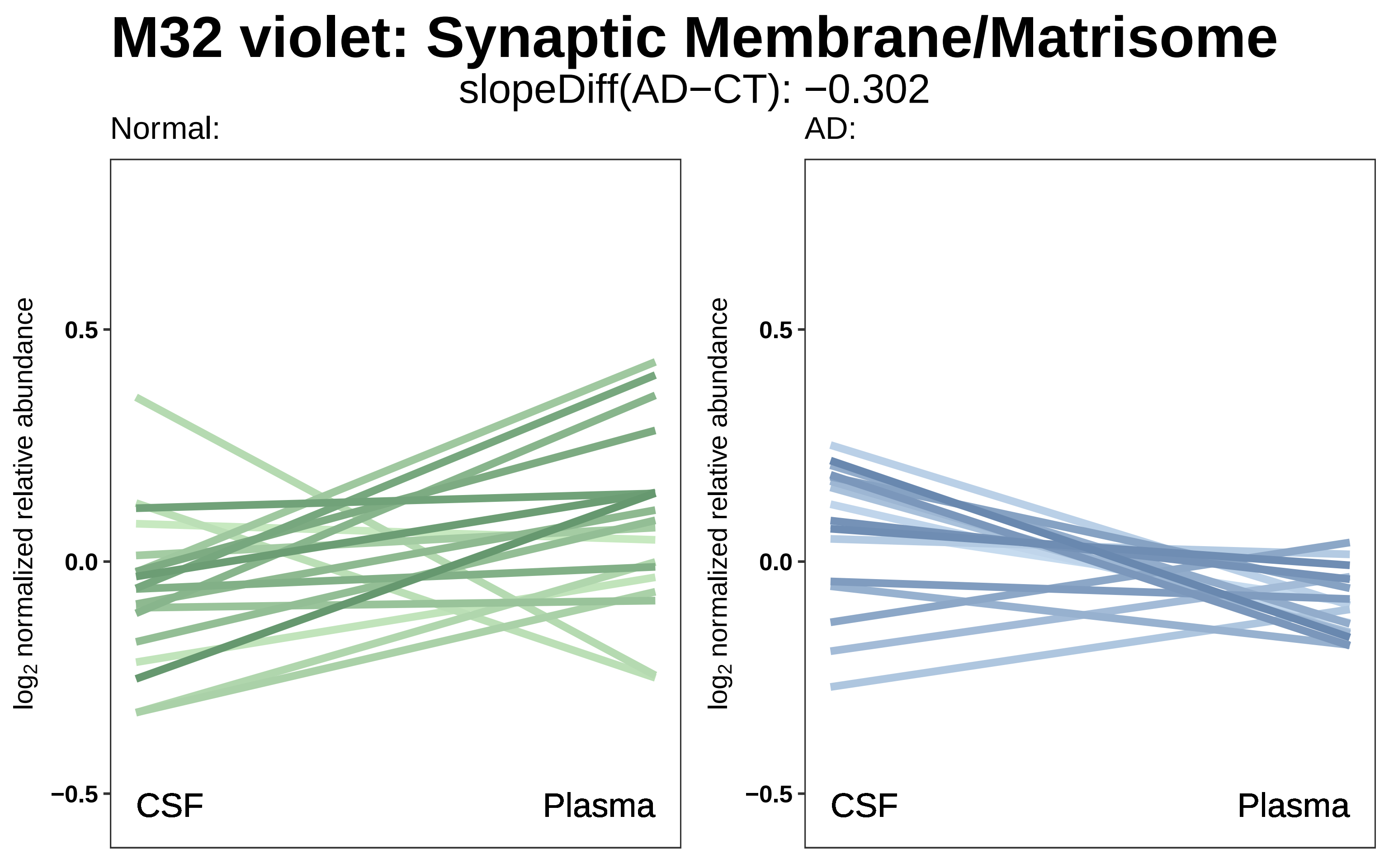

M32 violet: Synaptic Membrane/Matrisome
slopeDiff(AD−CT): −0.302
Normal:
AD:
log2 normalized relative abundance
0.5
0.0
−0.5
CSF
Plasma
log2 normalized relative abundance
0.5
0.0
−0.5
CSF
Plasma

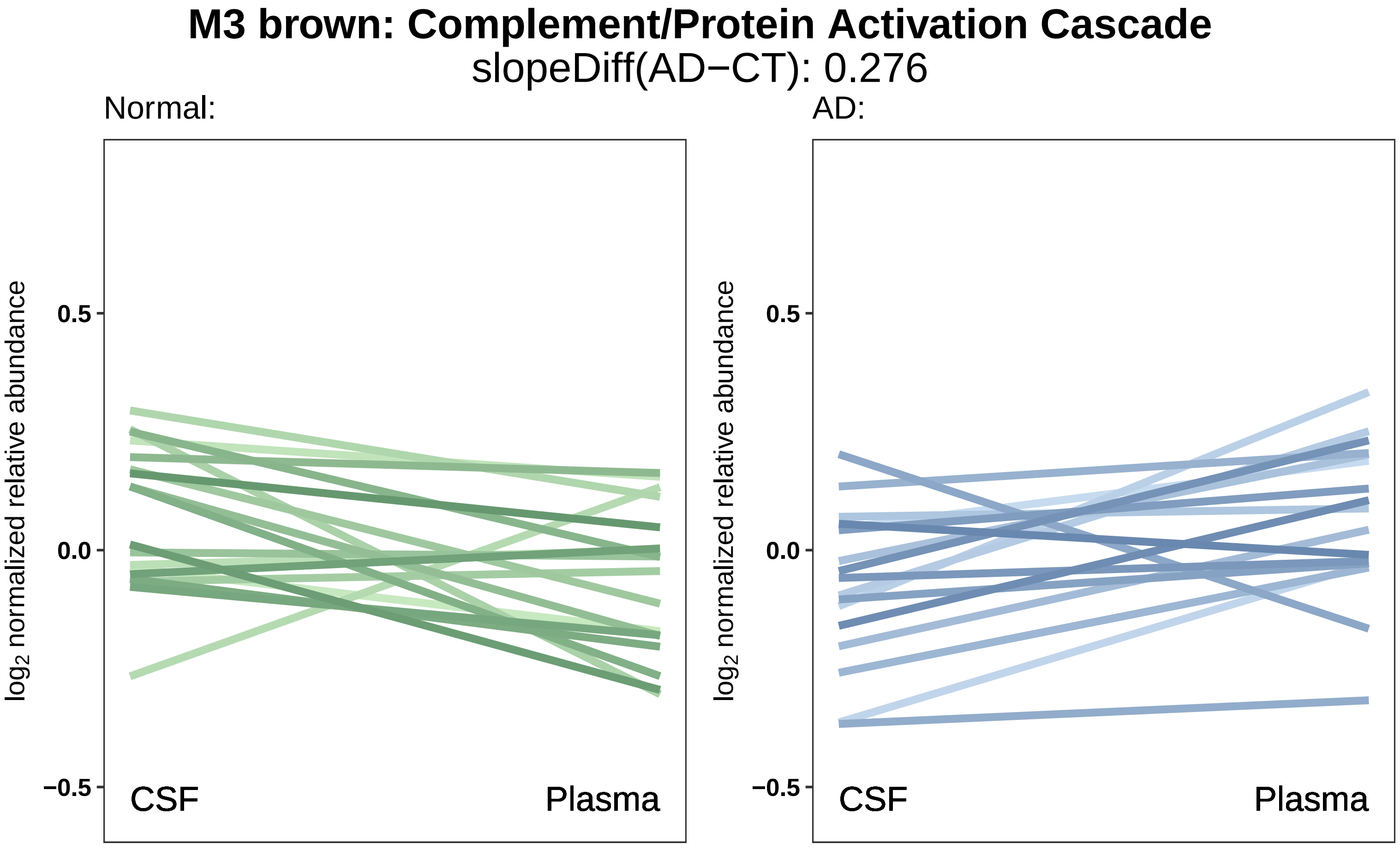

M3 brown: Complement/Protein Activation Cascade
slopeDiff(AD−CT): 0.276
Normal:
AD:
log2 normalized relative abundance
0.5
0.0
−0.5
CSF
Plasma
log2 normalized relative abundance
0.5
0.0
−0.5
CSF
Plasma

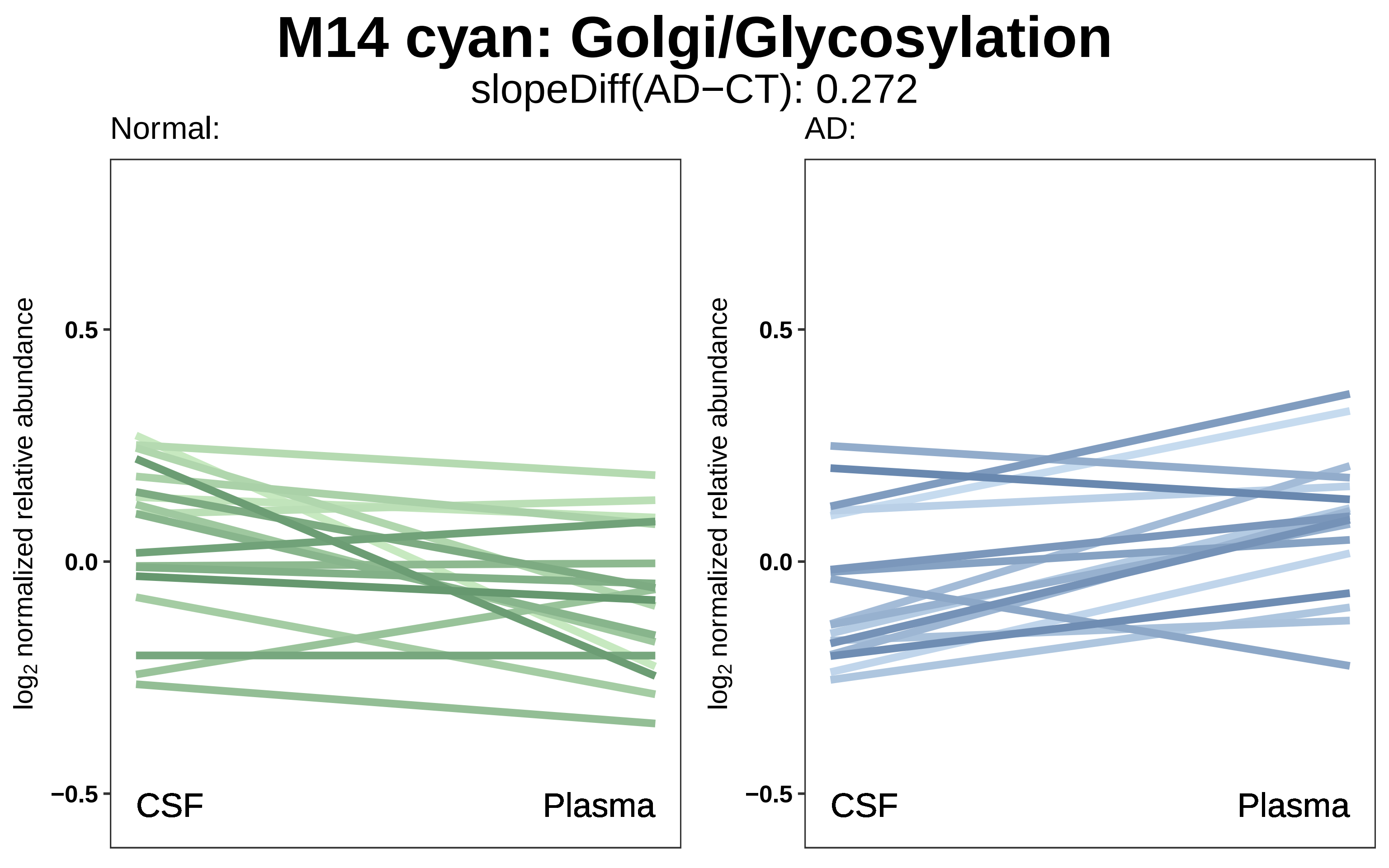

M14 cyan: Golgi/Glycosylation
slopeDiff(AD−CT): 0.272
Normal:
AD:
log2 normalized relative abundance
0.5
0.0
−0.5
CSF
Plasma
log2 normalized relative abundance
0.5
0.0
−0.5
CSF
Plasma

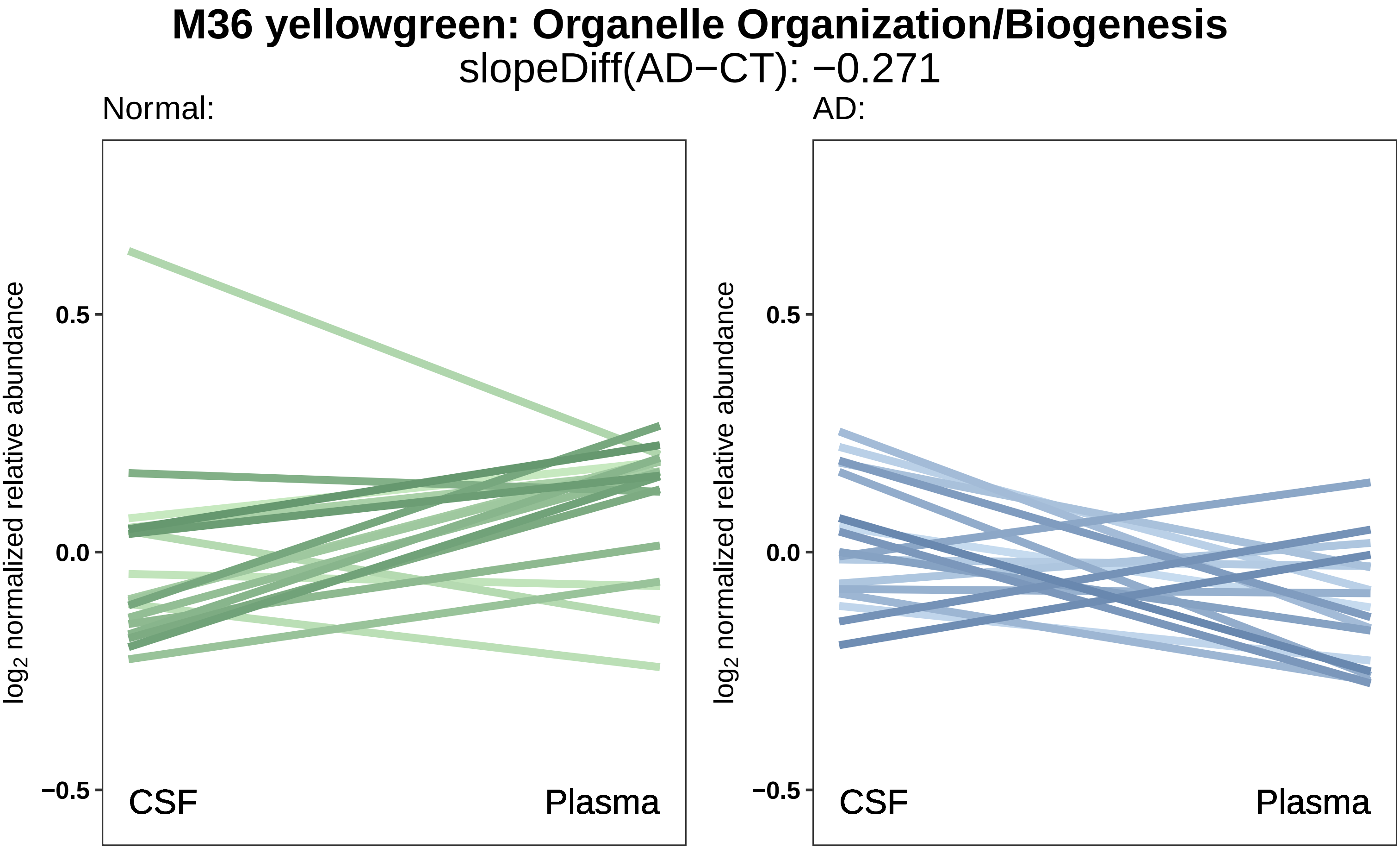
M36 yellowgreen: Organelle Organization/Biogenesis
slopeDiff(AD−CT): −0.271
Normal:
AD:
log2 normalized relative abundance
0.5
0.0
−0.5
CSF
Plasma
log2 normalized relative abundance
0.5
0.0
−0.5
CSF
Plasma

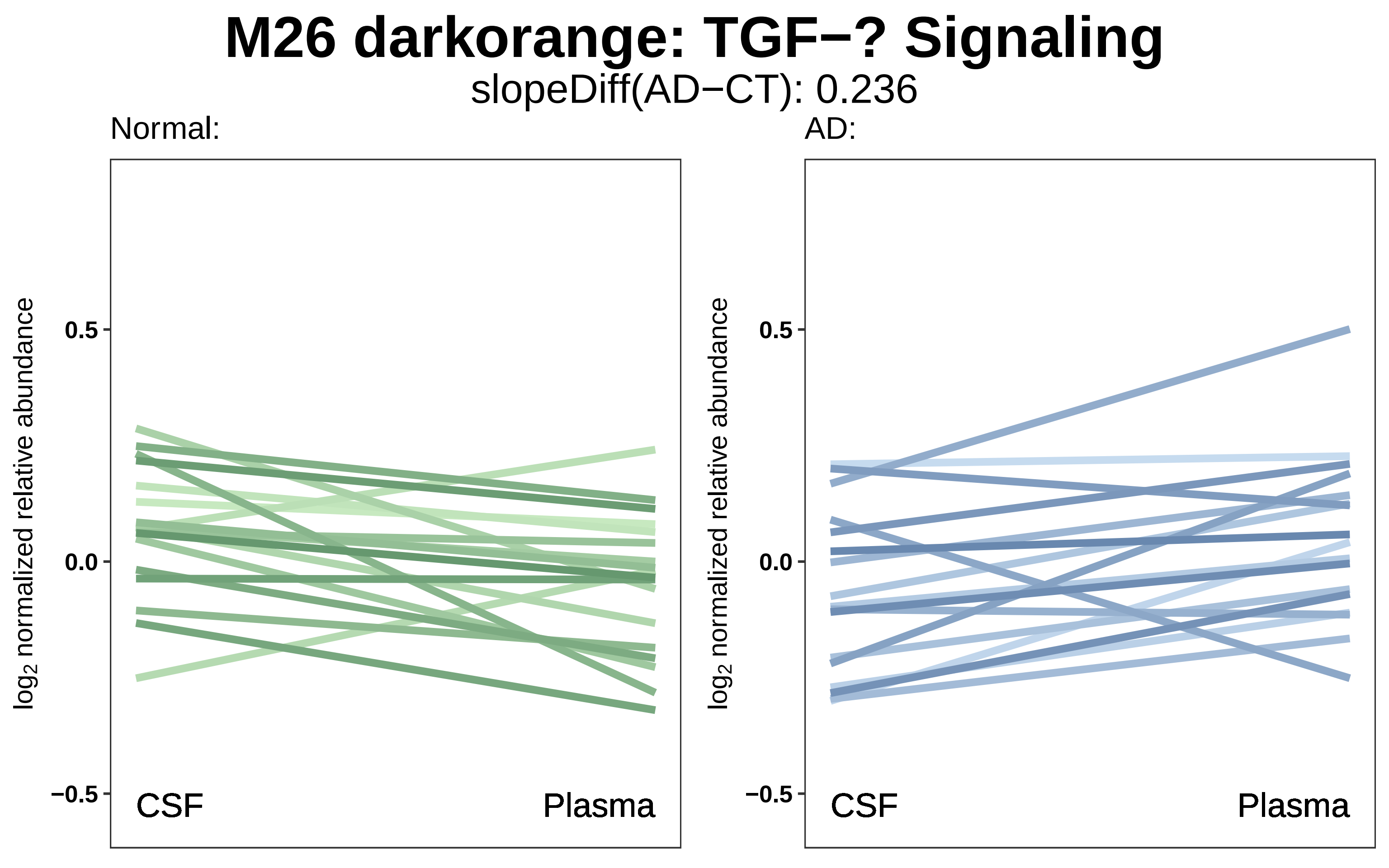

M26 darkorange: TGF−? Signaling
slopeDiff(AD−CT): 0.236
Normal:
AD:
log2 normalized relative abundance
0.5
0.0
−0.5
CSF
Plasma

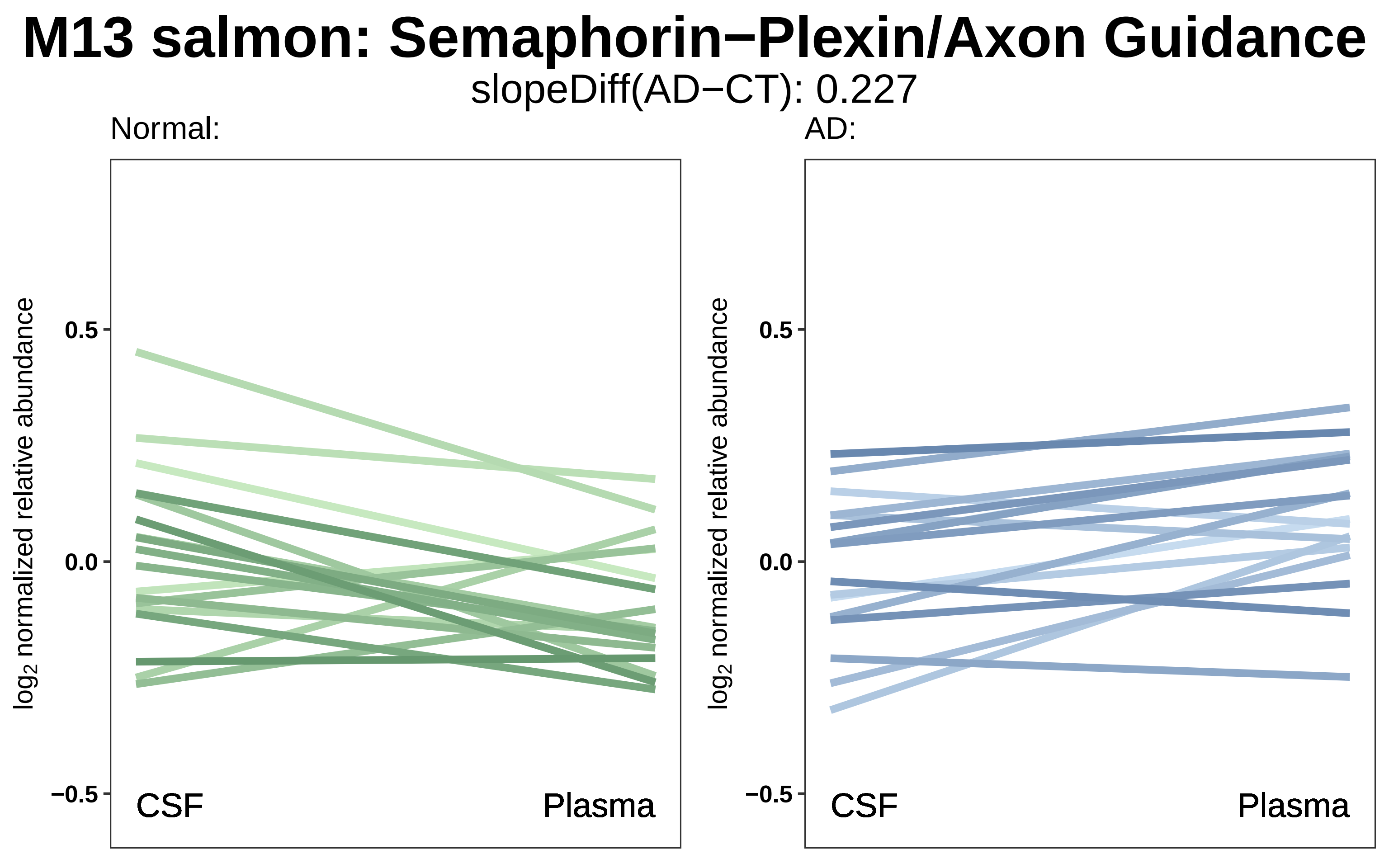
M13 salmon: Semaphorin−Plexin/Axon Guidance
slopeDiff(AD−CT): 0.227
Normal:
AD:
log2 normalized relative abundance
0.5
0.0
−0.5
CSF
Plasma

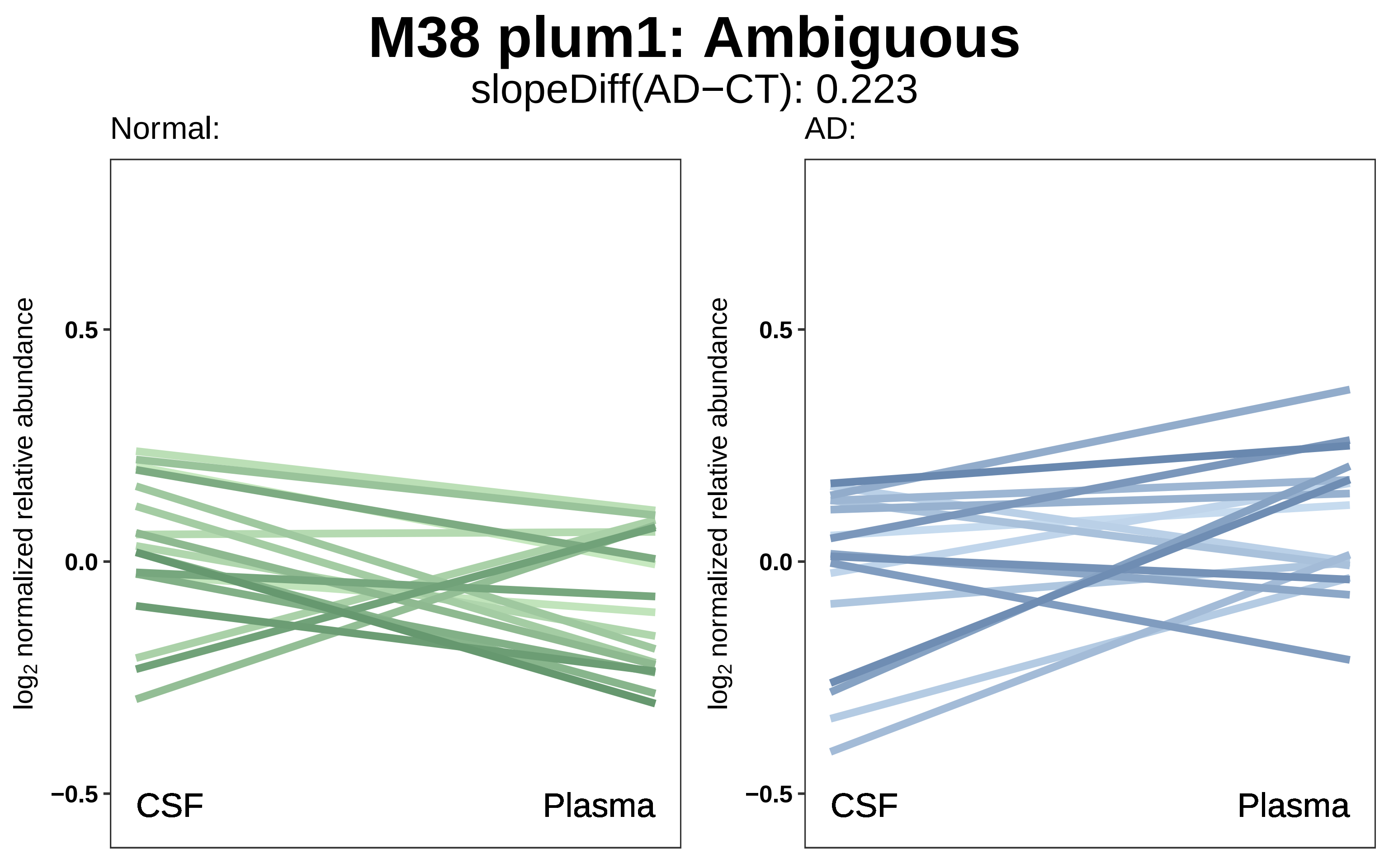

M38 plum1: Ambiguous
slopeDiff(AD−CT): 0.223
Normal:
AD:
log2 normalized relative abundance
0.5
0.0
−0.5
CSF
Plasma

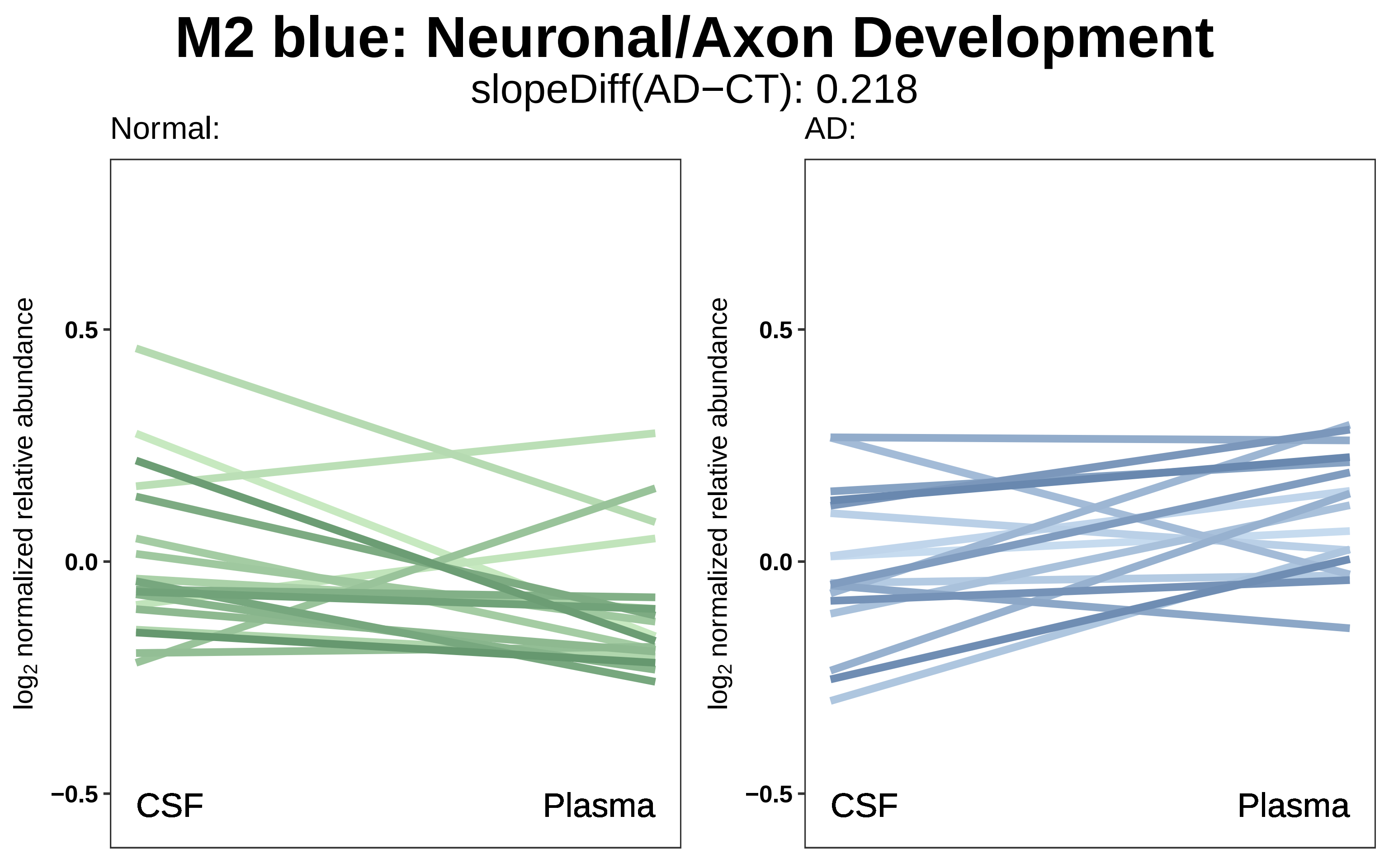
M2 blue: Neuronal/Axon Development
slopeDiff(AD−CT): 0.218
Normal:
AD:
log2 normalized relative abundance
0.5
0.0
−0.5
CSF
Plasma

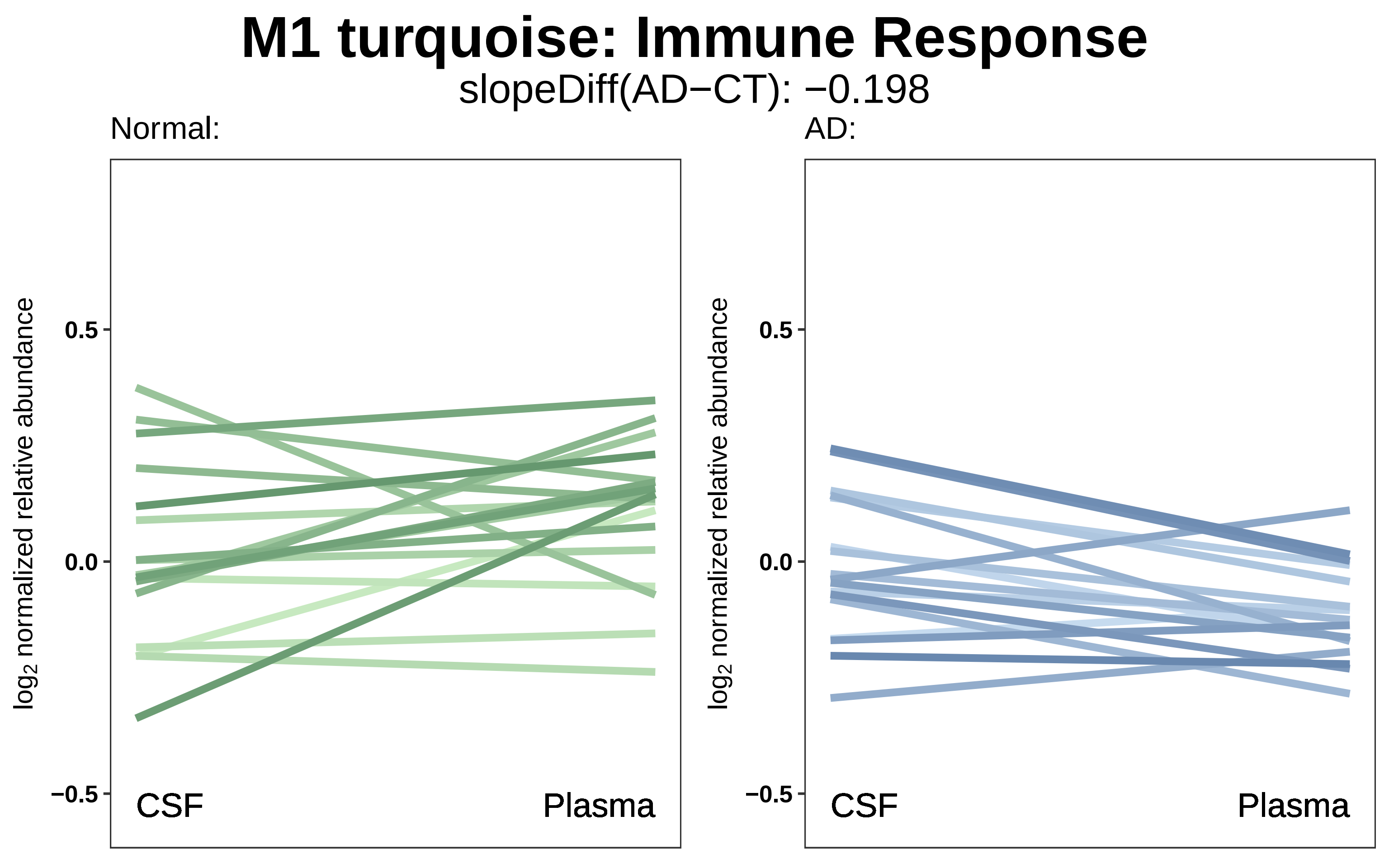
M1 turquoise: Immune Response
slopeDiff(AD−CT): −0.198
Normal:
AD:
log2 normalized relative abundance
0.5
0.0
−0.5
CSF
Plasma
log2 normalized relative abundance
0.5
0.0
−0.5
CSF
Plasma

### M18 lightgreen: ECM Organization/Scavenger Receptor
slopeDiff(AD−CT): 0.195

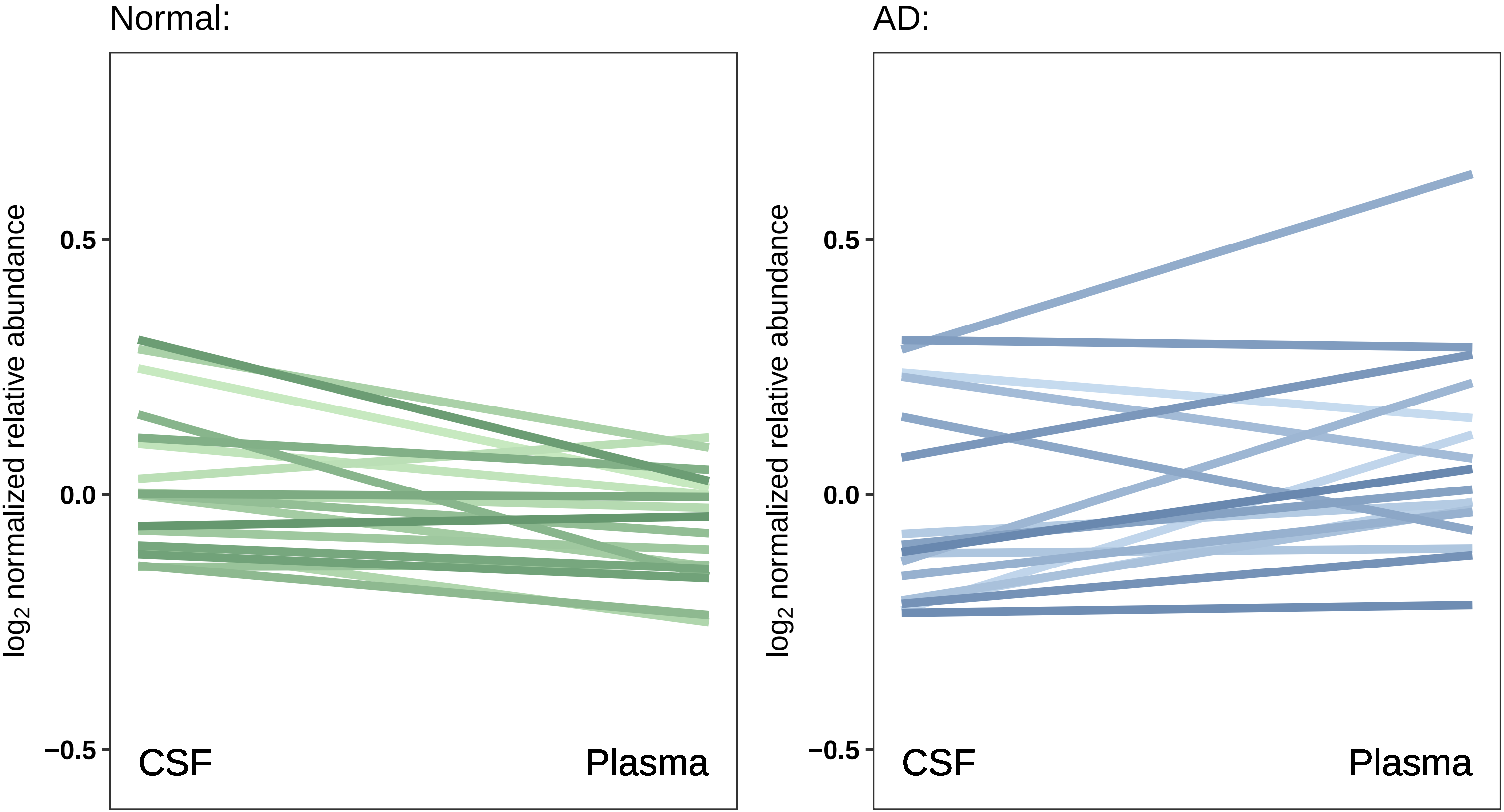

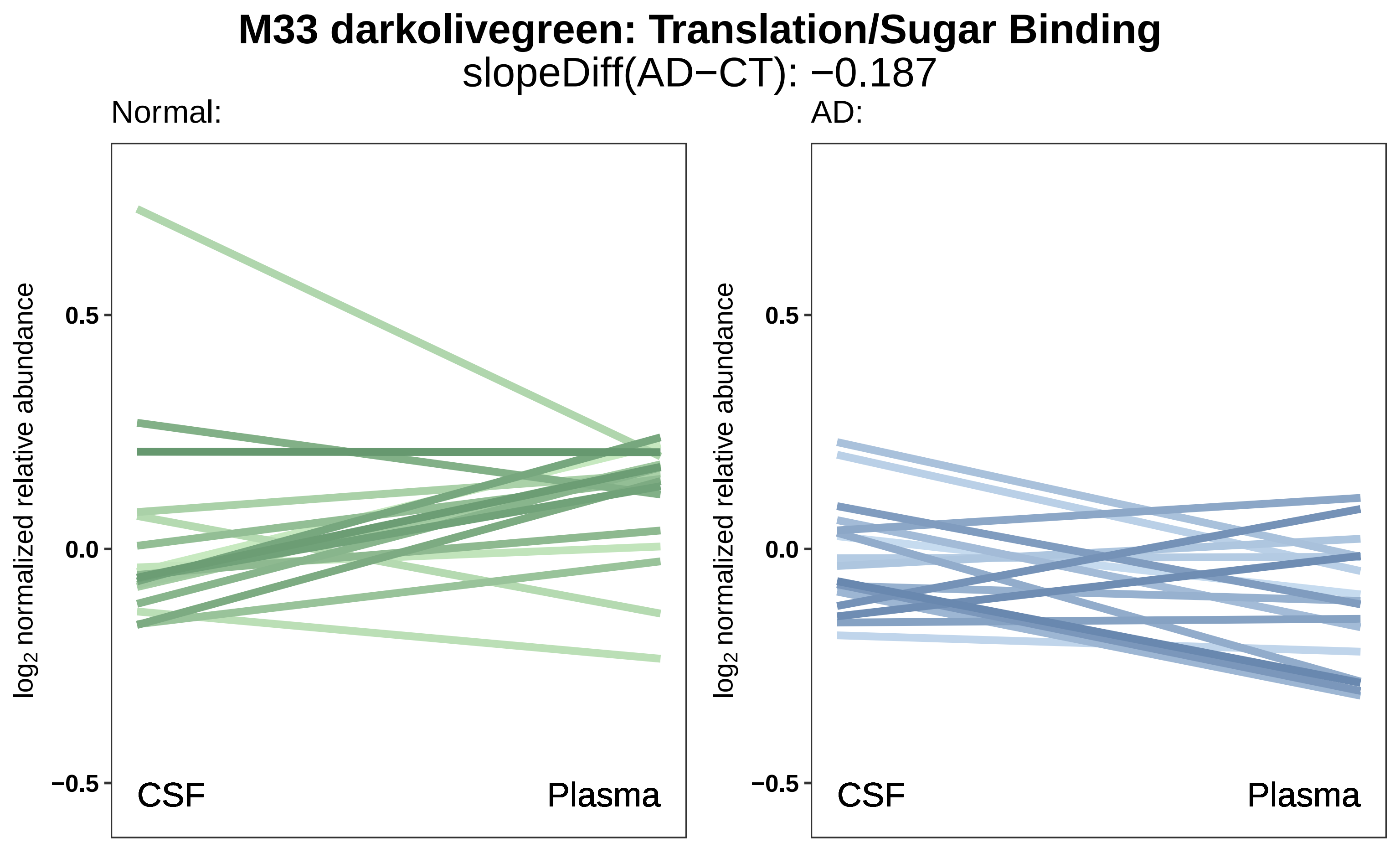
M33 darkolivegreen: Translation/Sugar Binding
slopeDiff(AD−CT): −0.187
Normal:
AD:
log2 normalized relative abundance
0.5
0.0
−0.5
CSF
Plasma
log2 normalized relative abundance
0.5
0.0
−0.5
CSF
Plasma

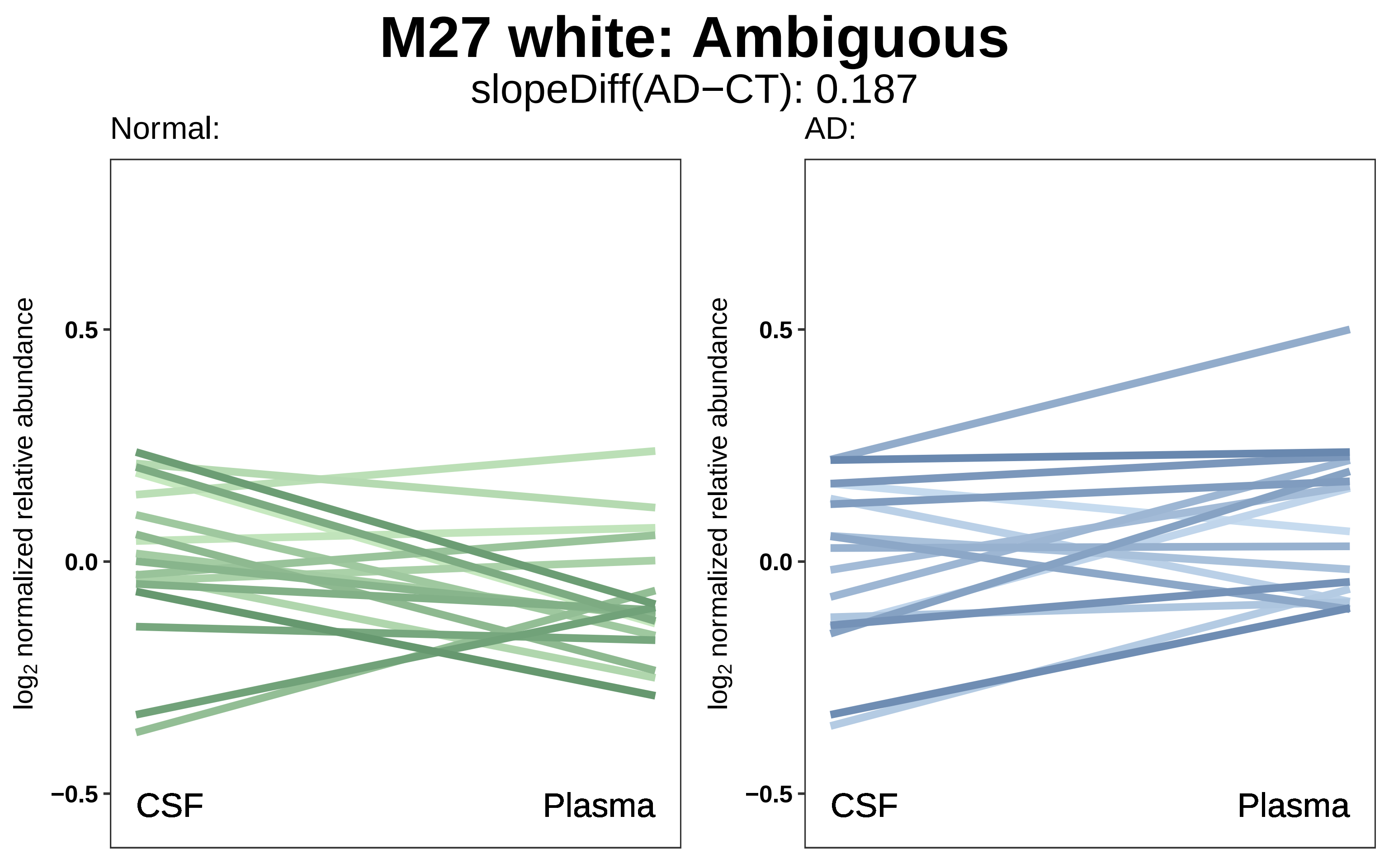

M27 white: Ambiguous
slopeDiff(AD−CT): 0.187
Normal:
AD:
log2 normalized relative abundance
0.5
0.0
−0.5
CSF
Plasma
log2 normalized relative abundance
0.5
0.0
−0.5
CSF
Plasma

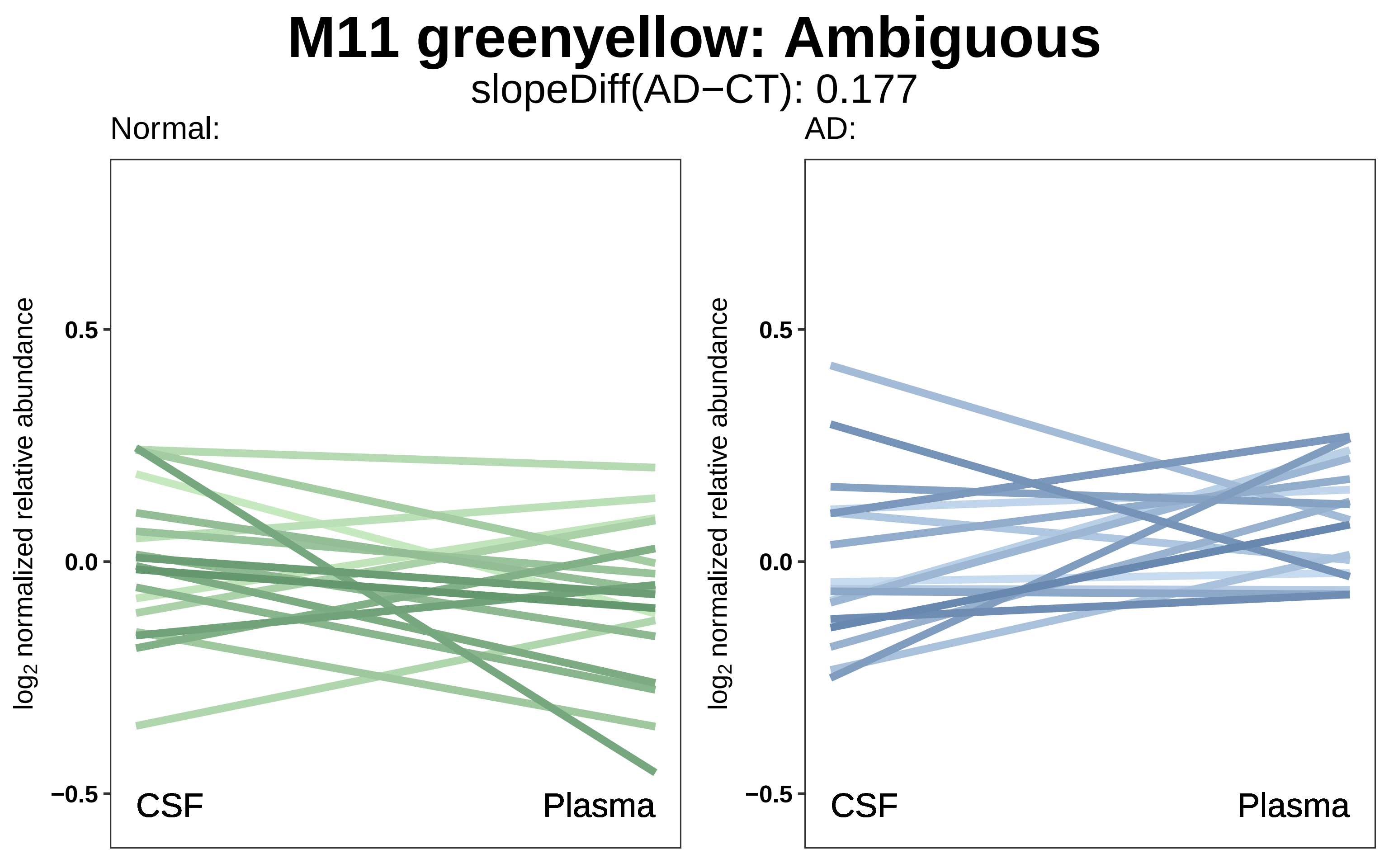
M11 greenyellow: Ambiguous
slopeDiff(AD−CT): 0.177
Normal:
AD:
log2 normalized relative abundance
0.5
0.0
−0.5
CSF
Plasma
log2 normalized relative abundance
0.5
0.0
−0.5
CSF
Plasma

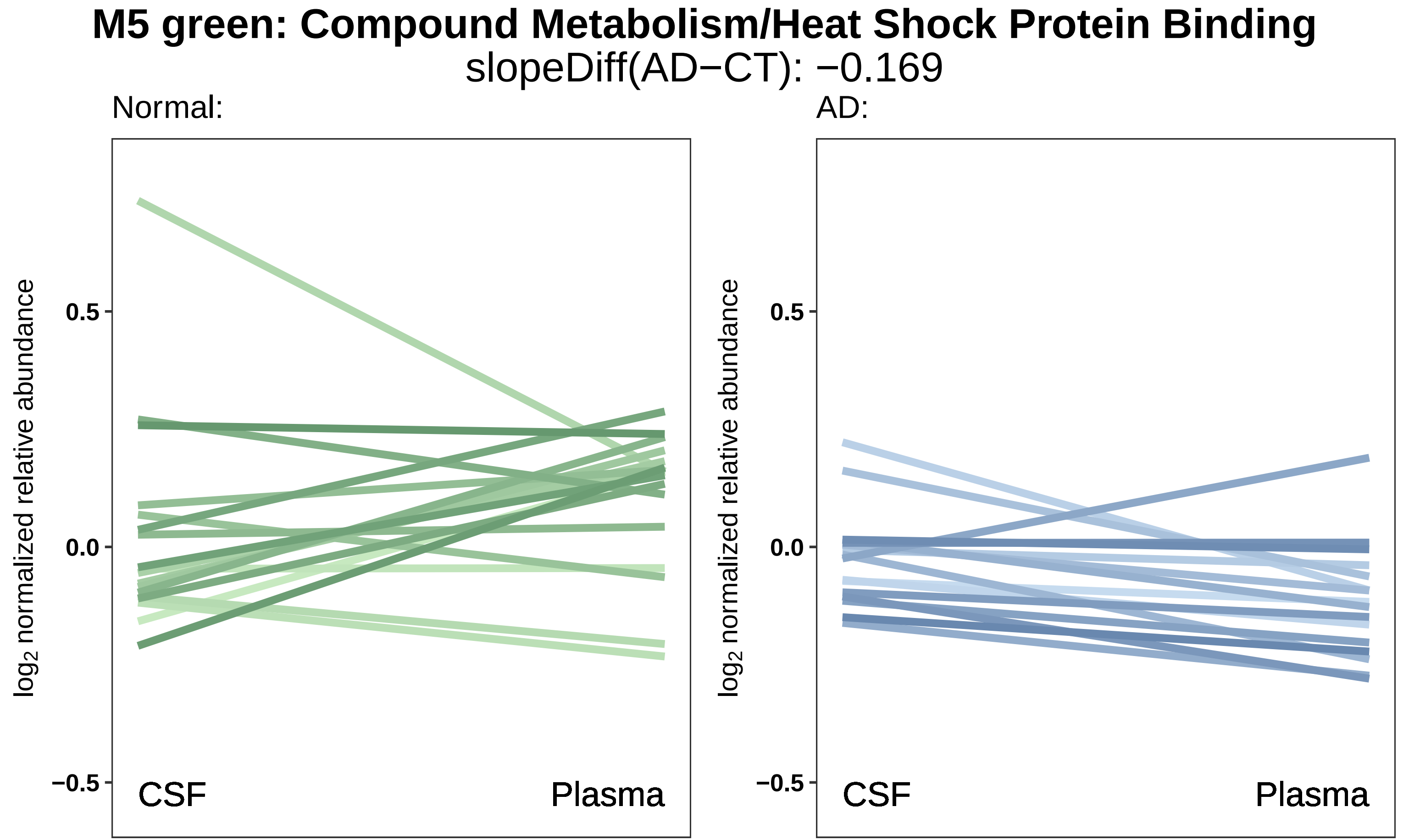

M5 green: Compound Metabolism/Heat Shock Protein Binding
slopeDiff(AD−CT): −0.169
Normal:
AD:
log2 normalized relative abundance
0.5
0.0
−0.5
CSF
Plasma

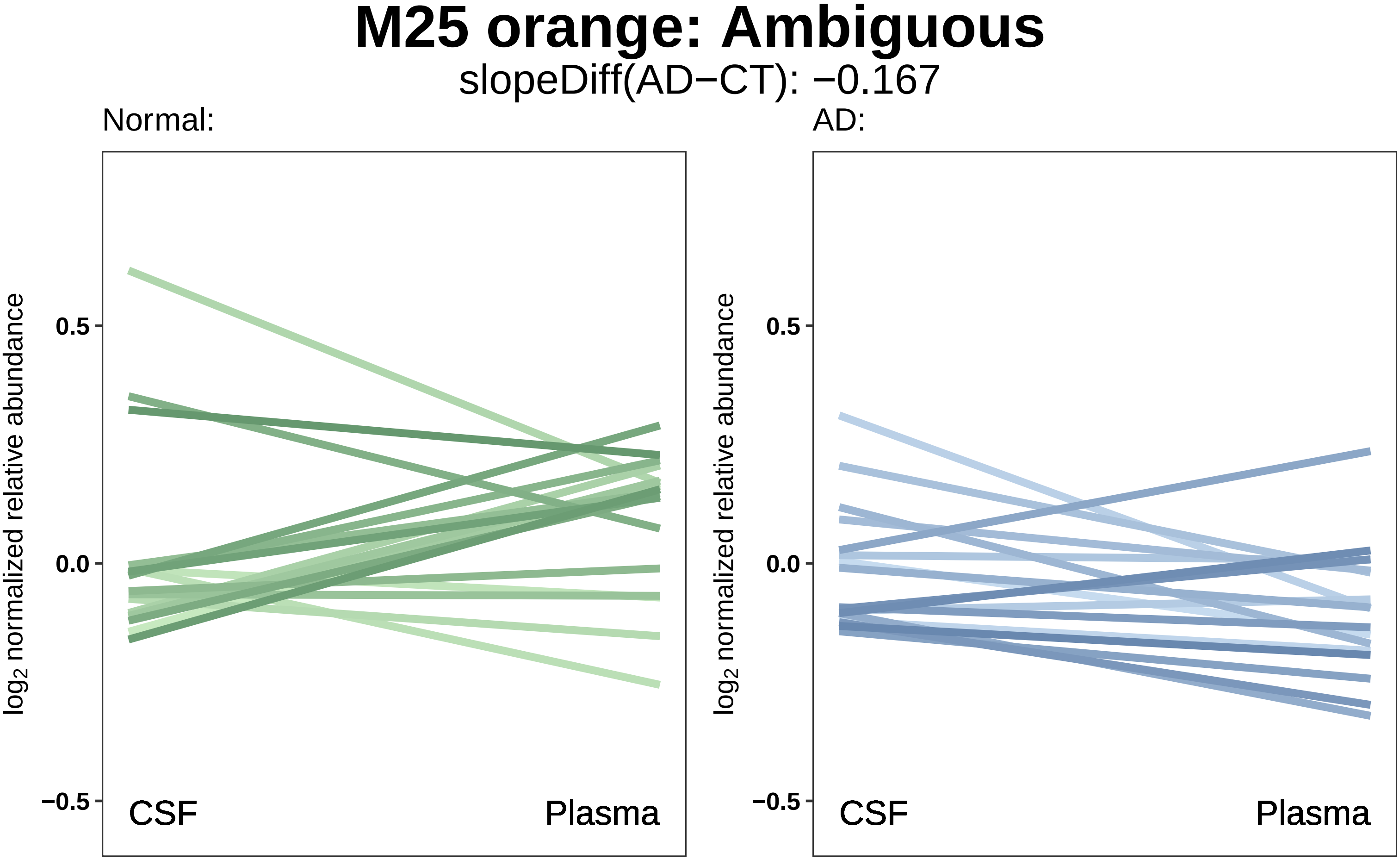

M25 orange: Ambiguous
slopeDiff(AD−CT): −0.167
Normal:
AD:
log2 normalized relative abundance
0.5
0.0
−0.5
CSF
Plasma
log2 normalized relative abundance
0.5
0.0
−0.5
CSF
Plasma

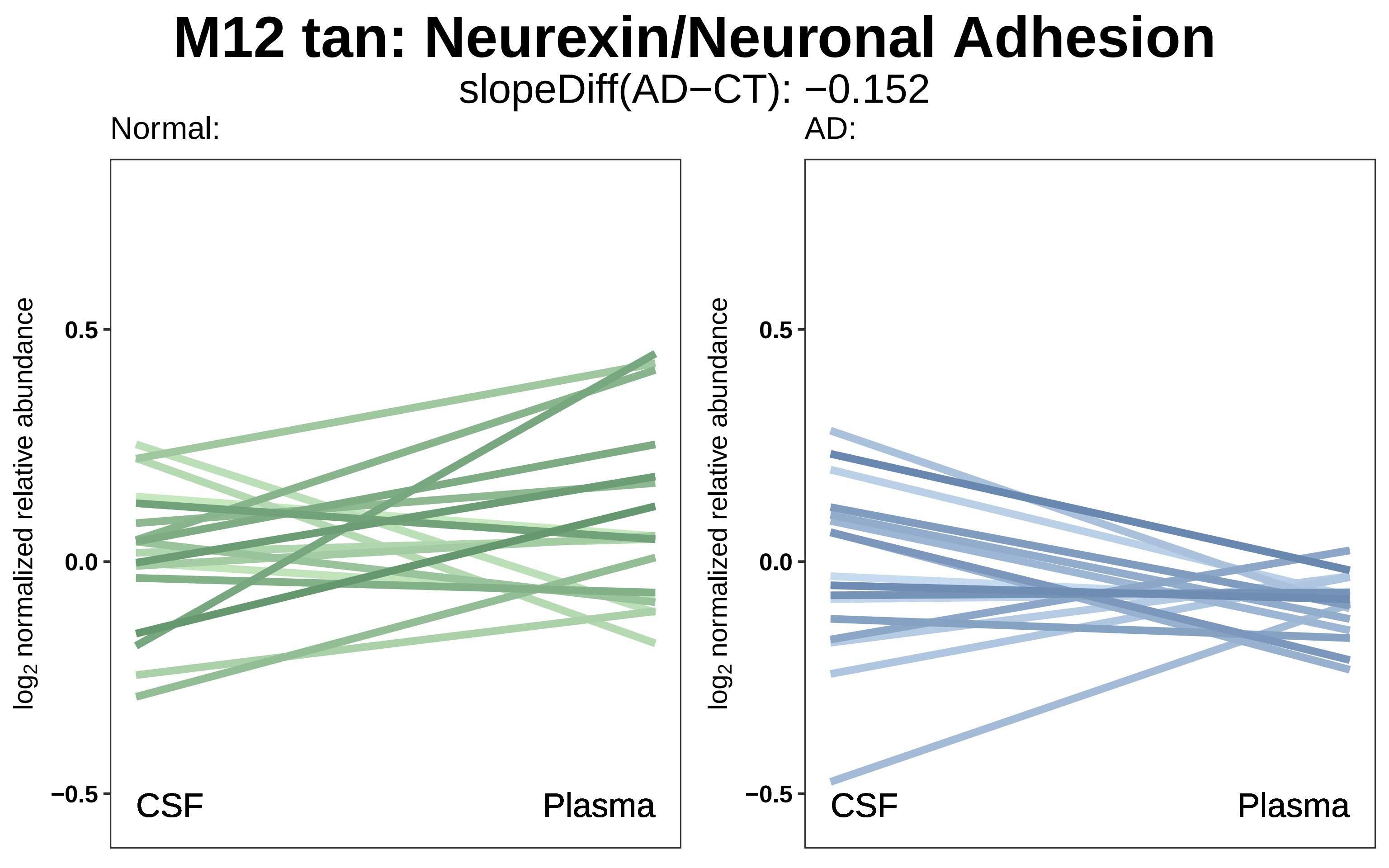
M12 tan: Neurexin/Neuronal Adhesion
slopeDiff(AD−CT): −0.152
Normal:
AD:
log2 normalized relative abundance
0.5
0.0
−0.5
CSF
Plasma

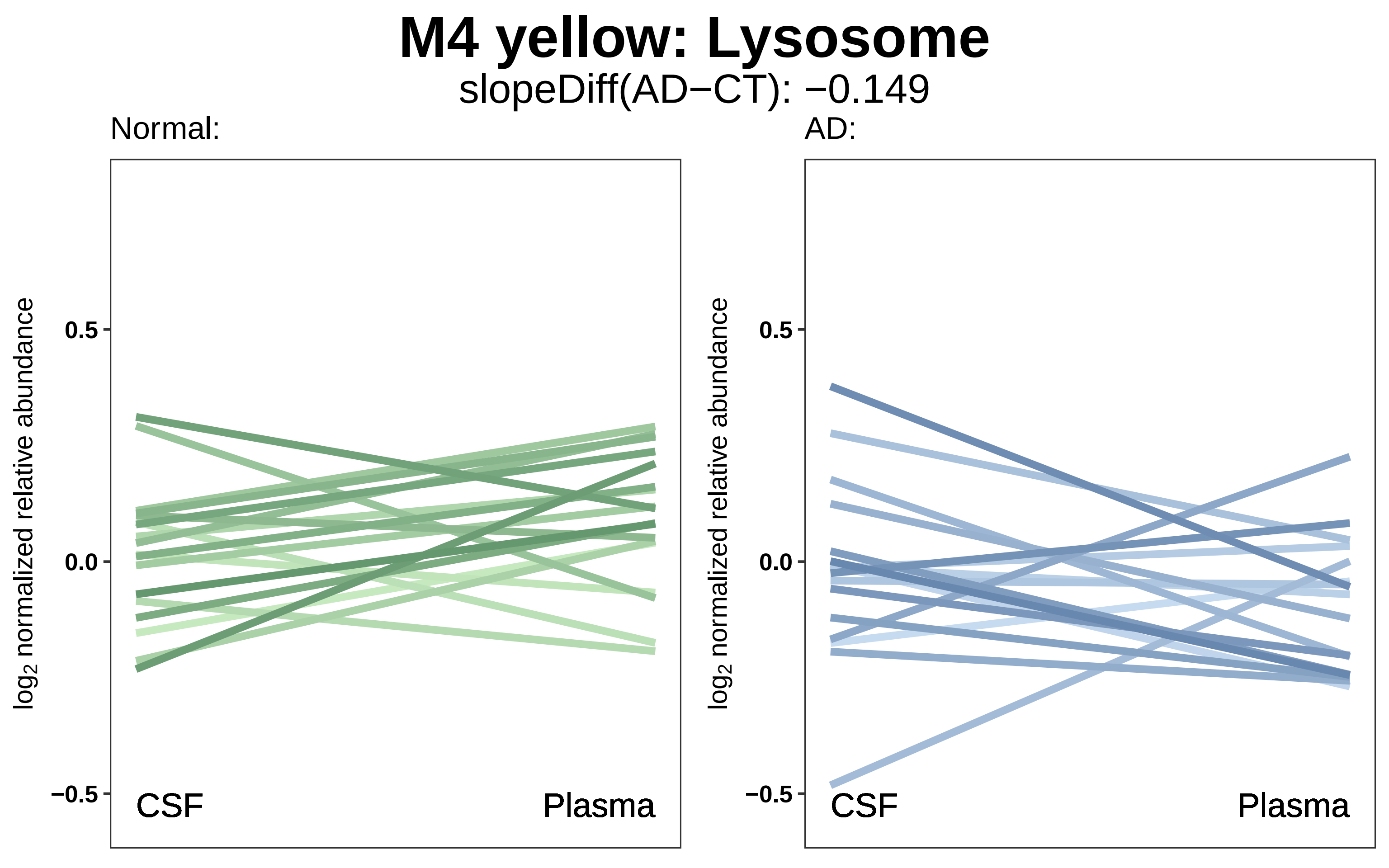
M4 yellow: Lysosome
slopeDiff(AD−CT): −0.149
Normal:
AD:
log2 normalized relative abundance
0.5
0.0
−0.5
CSF
Plasma
log2 normalized relative abundance
0.5
0.0
−0.5
CSF
Plasma

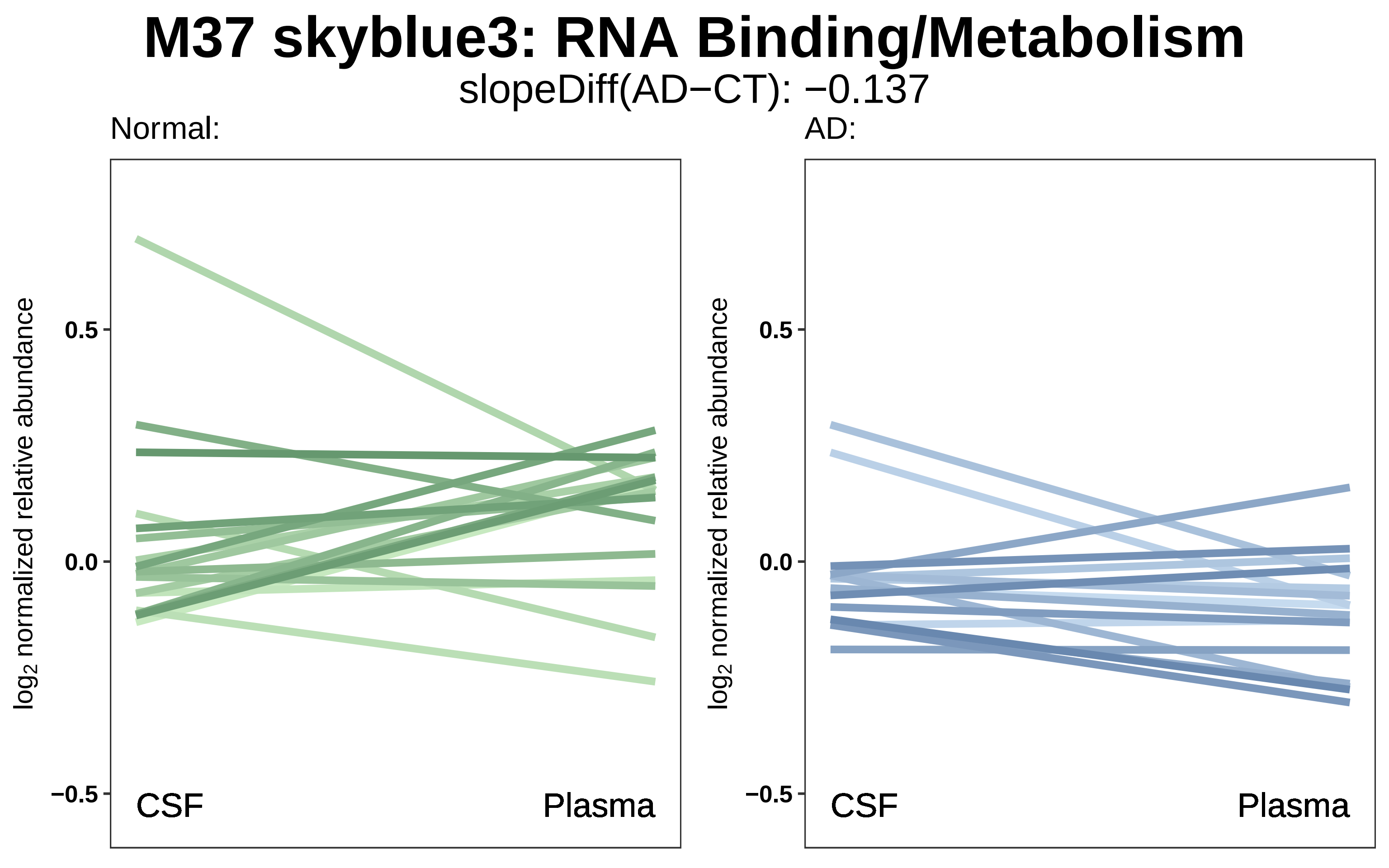
M37 skyblue3: RNA Binding/Metabolism
slopeDiff(AD−CT): −0.137
Normal:
AD:
log2 normalized relative abundance
0.5
0.0
−0.5
CSF
Plasma
log2 normalized relative abundance
0.5
0.0
−0.5
CSF
Plasma

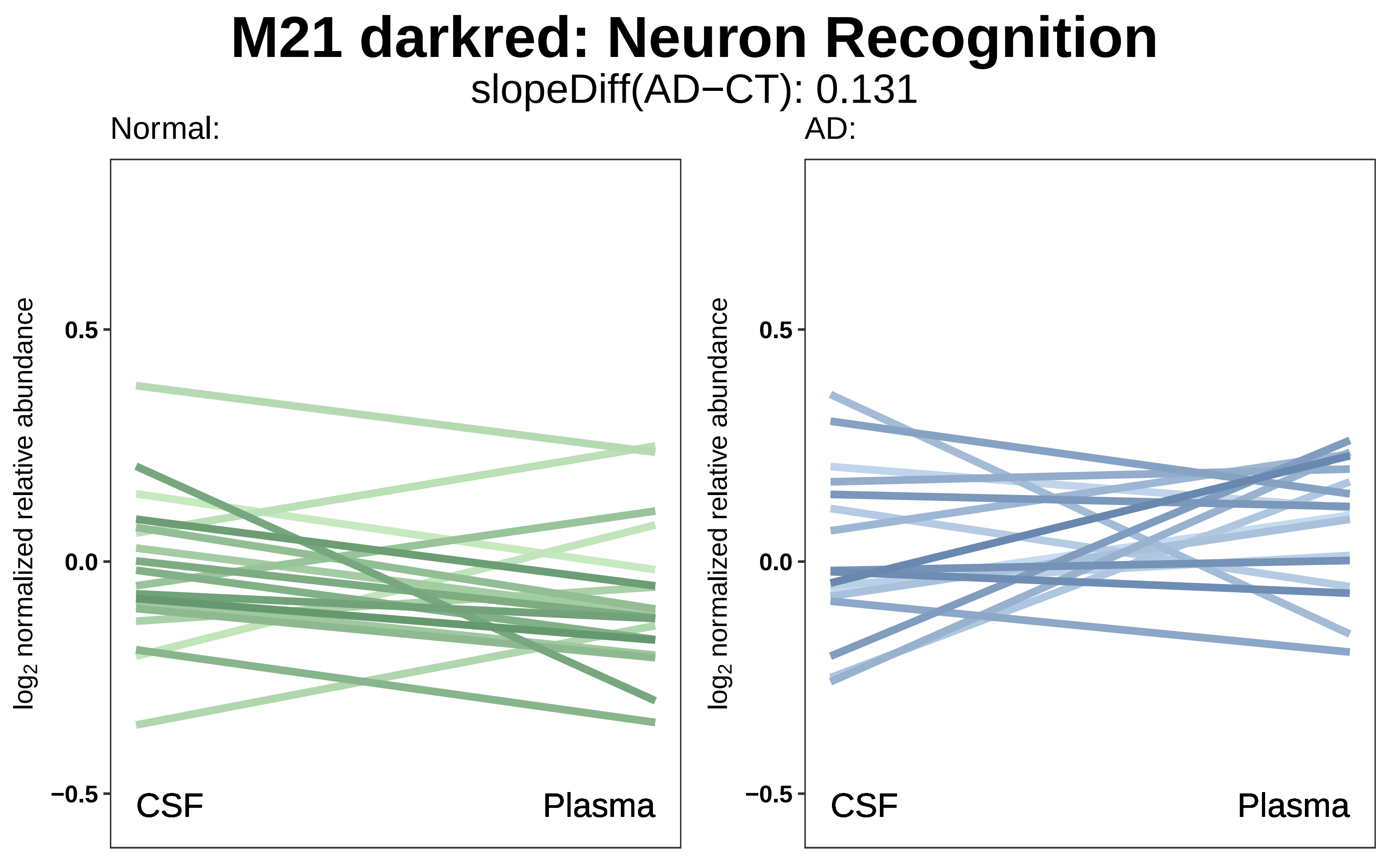

M21 darkred: Neuron Recognition
slopeDiff(AD−CT): 0.131
Normal:
AD:
log2 normalized relative abundance
0.5
0.0
−0.5
CSF
Plasma

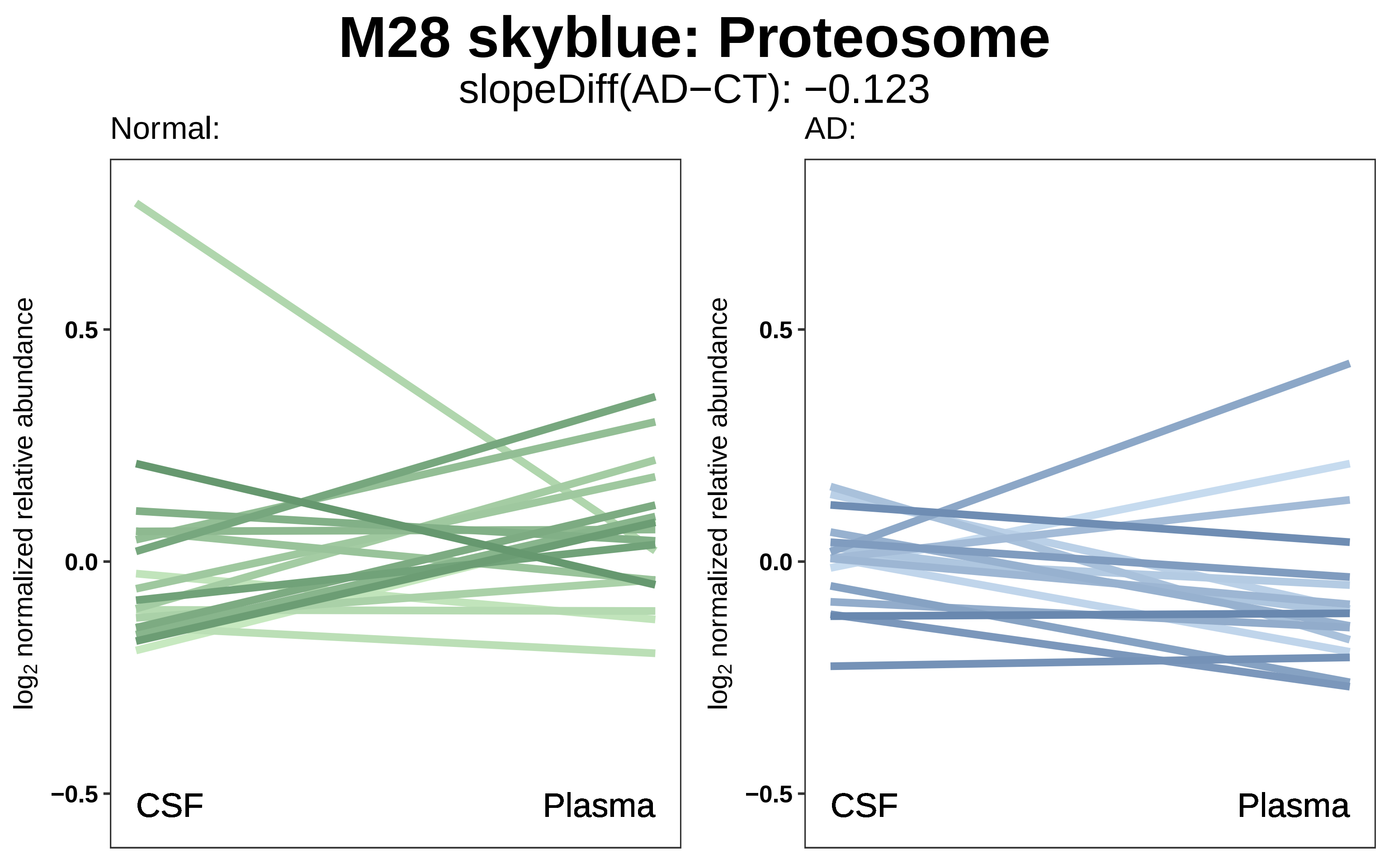
M28 skyblue: Proteosome
slopeDiff(AD−CT): −0.123
Normal:
AD:
log2 normalized relative abundance
0.5
0.0
−0.5
CSF
Plasma

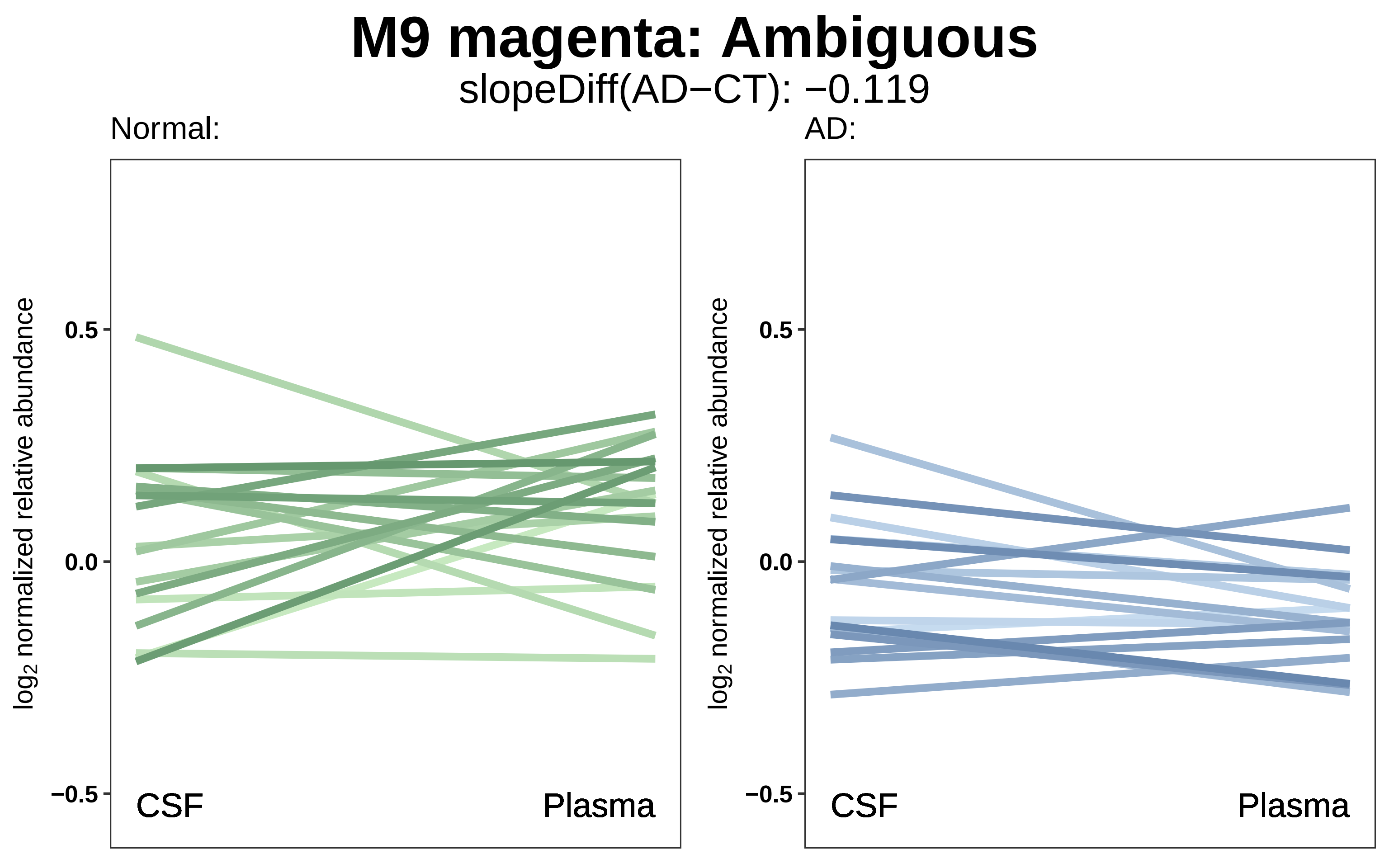

M9 magenta: Ambiguous
slopeDiff(AD−CT): −0.119
Normal:
AD:
log2 normalized relative abundance
0.5
0.0
−0.5
CSF
Plasma
log2 normalized relative abundance
0.5
0.0
−0.5
CSF
Plasma

### M30 steelblue: Ribonucleoprotein Complex

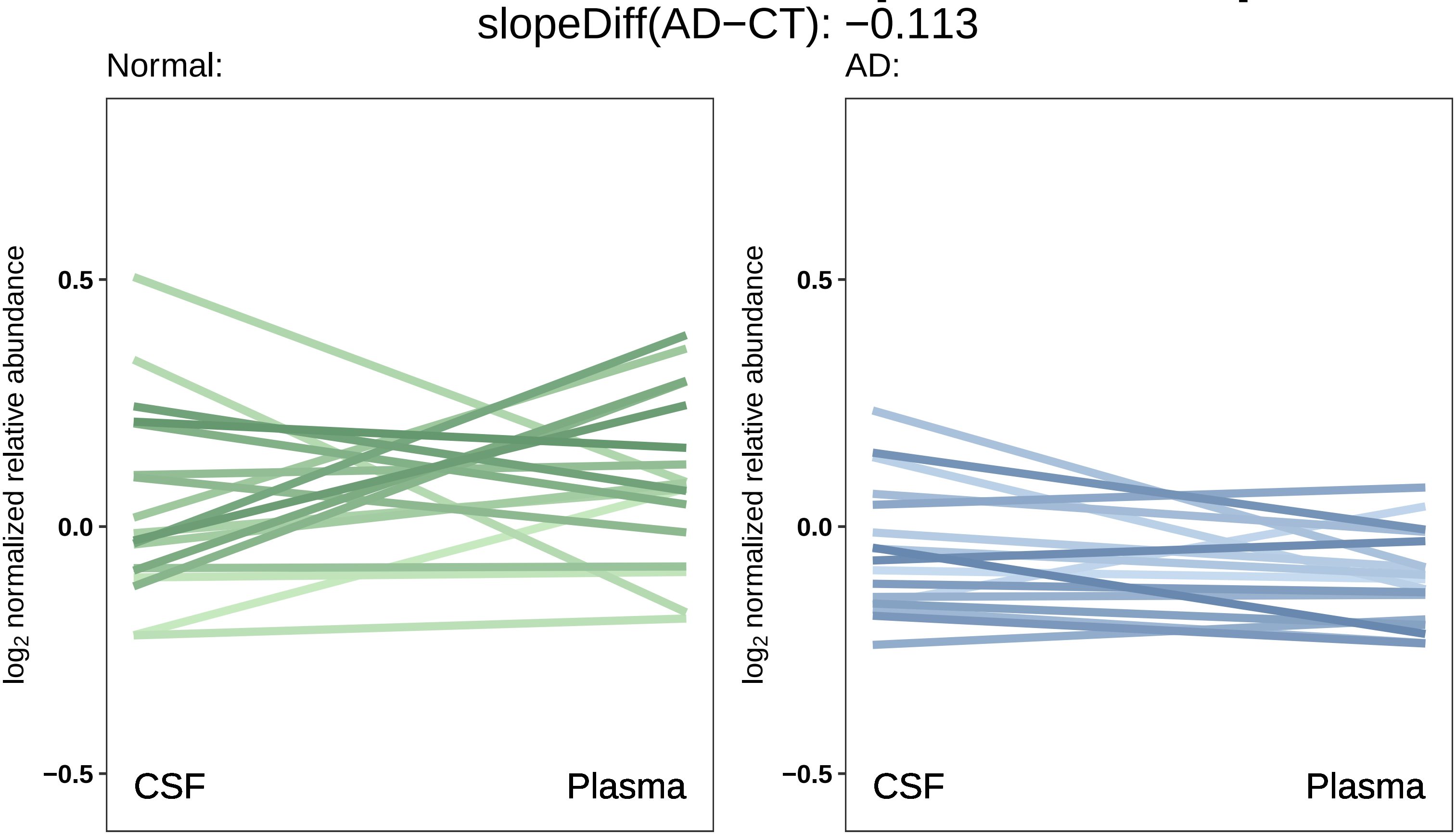

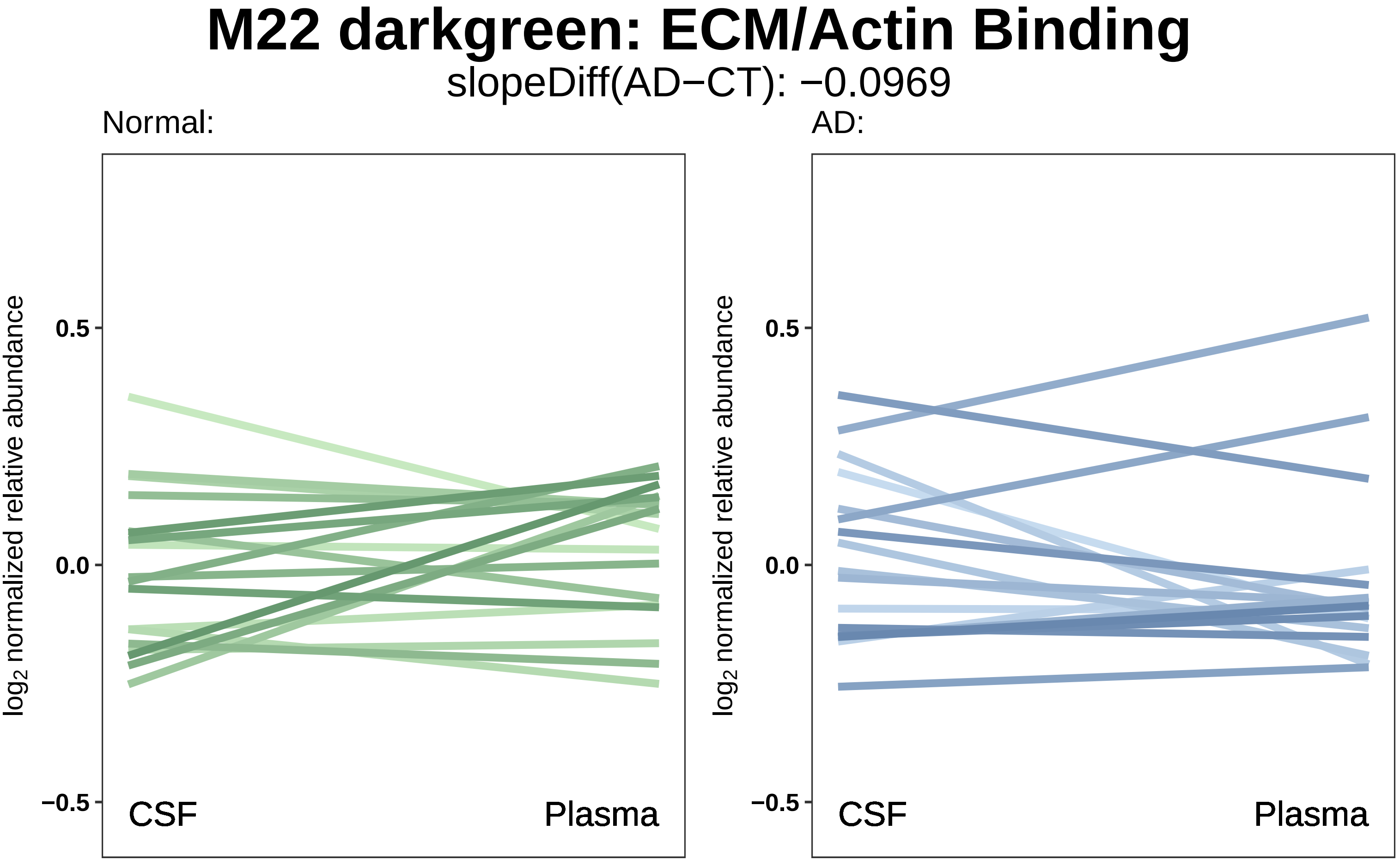
M22 darkgreen: ECM/Actin Binding
slopeDiff(AD−CT): −0.0969
Normal:
AD:
log2 normalized relative abundance
0.5
0.0
−0.5
CSF
Plasma

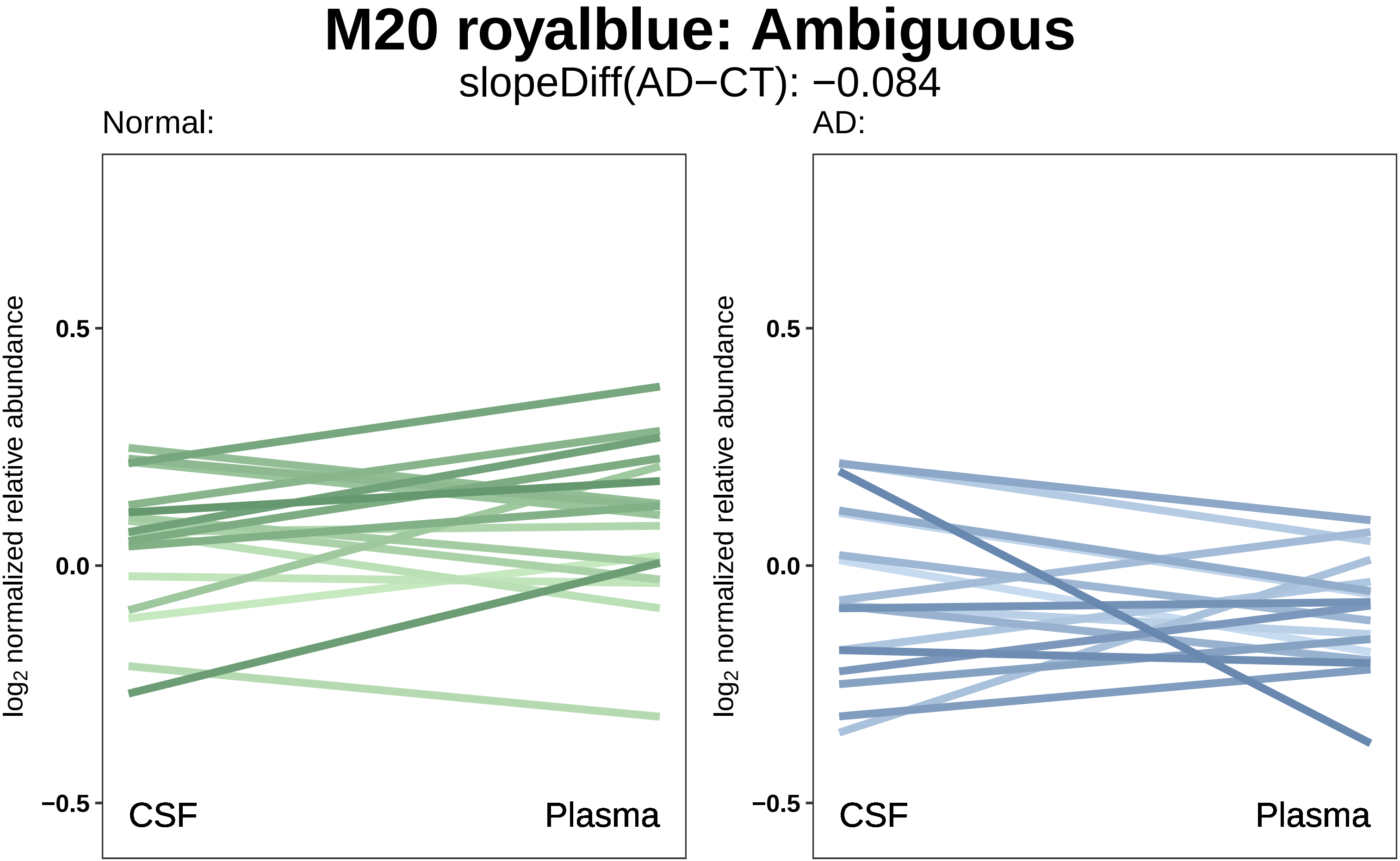
M20 royalblue: Ambiguous
slopeDiff(AD−CT): −0.084
Normal:
AD:
log2 normalized relative abundance
0.5
0.0
−0.5
CSF
Plasma

### M17 grey60: Ambiguous

slopeDiff(AD−CT): 0.0829

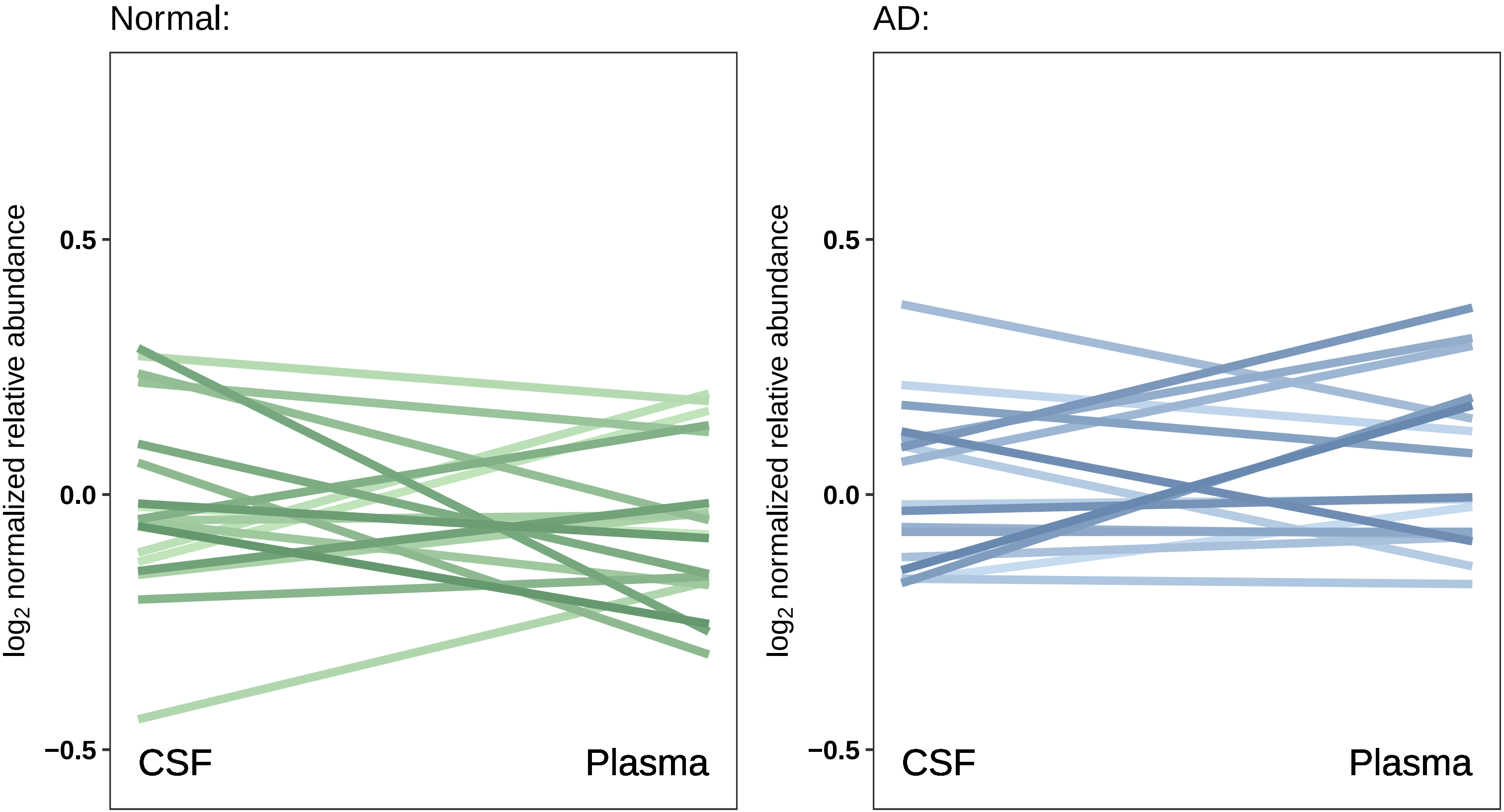

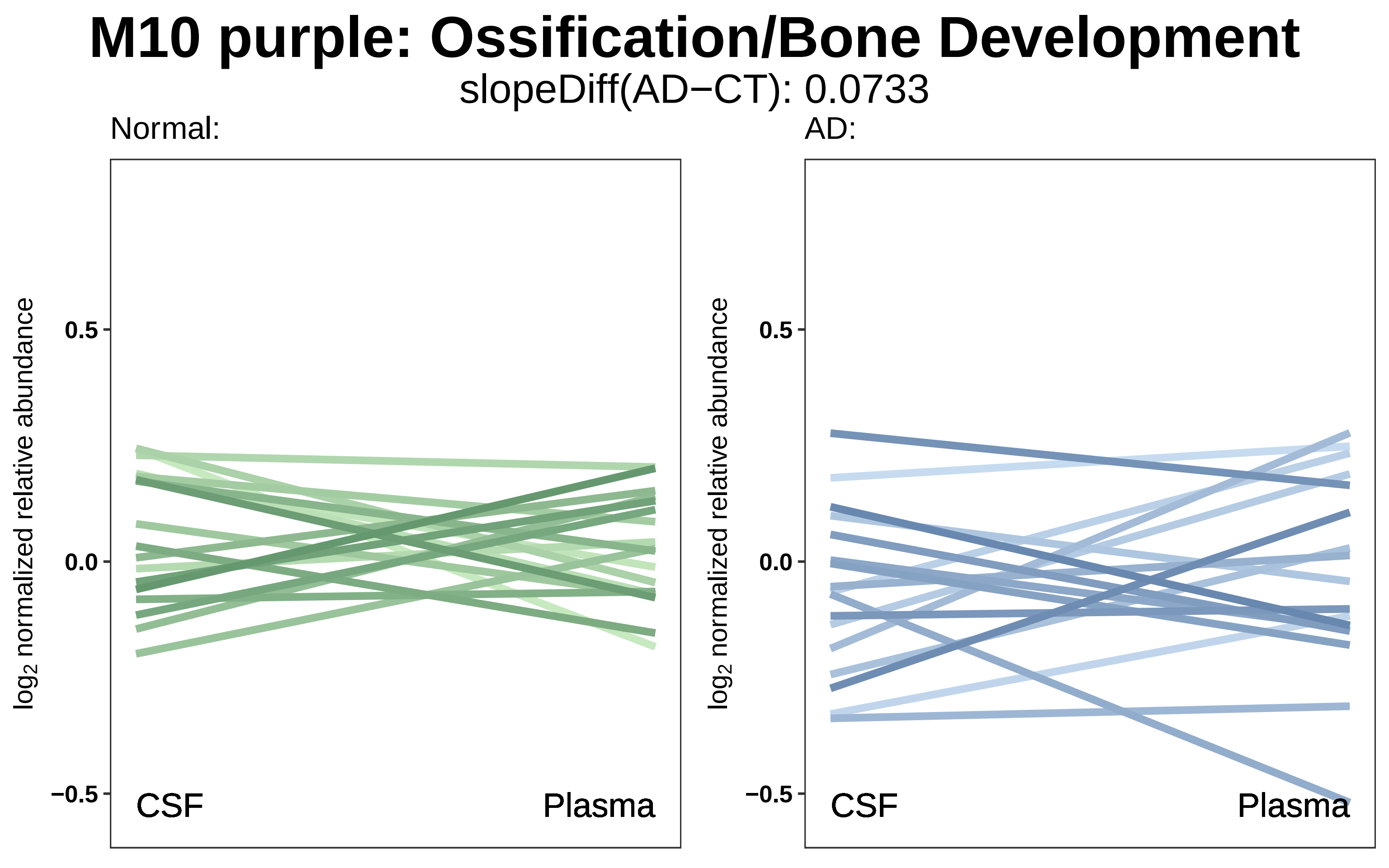

M10 purple: Ossification/Bone Development
slopeDiff(AD−CT): 0.0733
Normal:
AD:
log2 normalized relative abundance
0.5
0.0
−0.5
CSF
Plasma
log2 normalized relative abundance
0.5
0.0
−0.5
CSF
Plasma

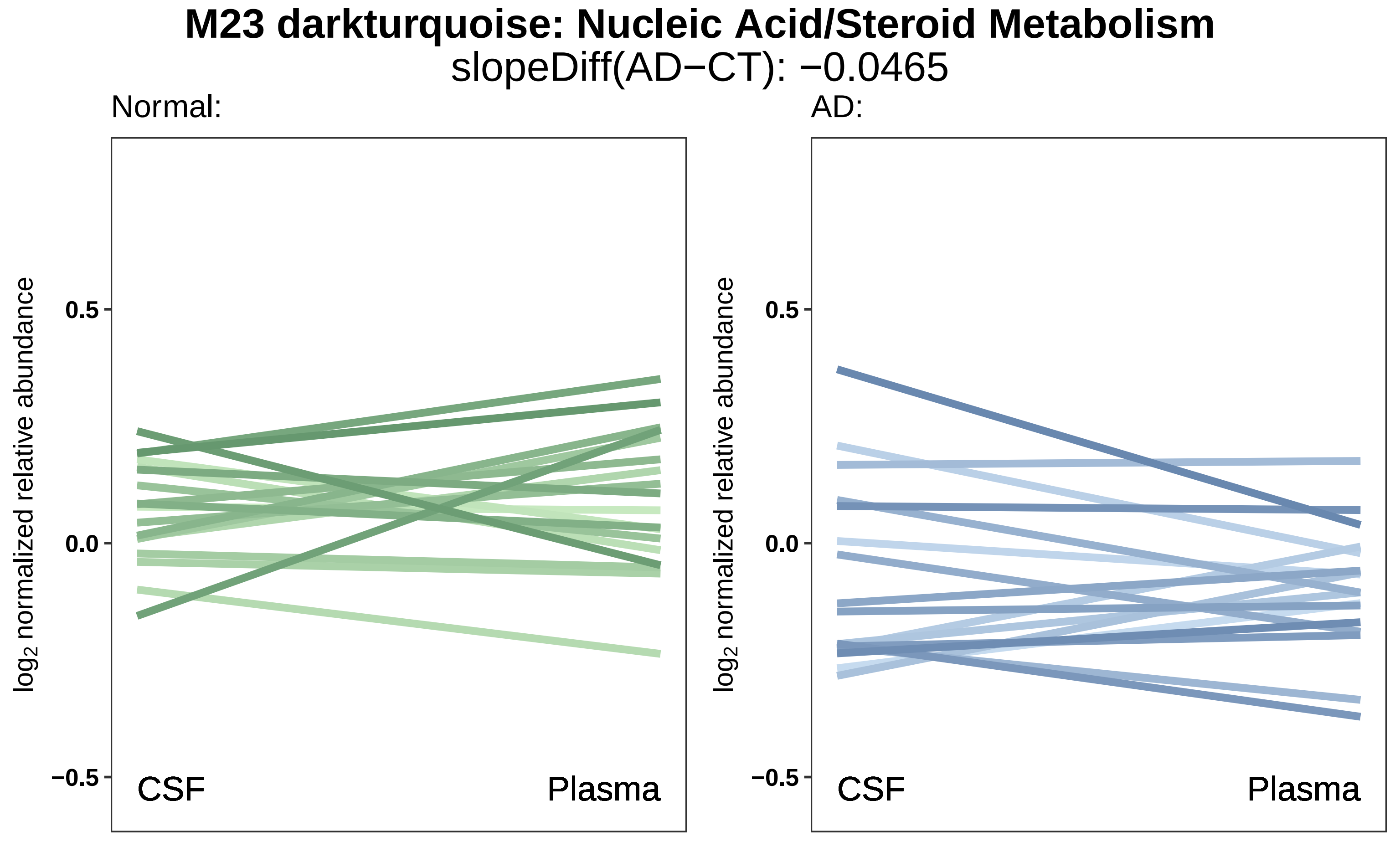
M23 darkturquoise: Nucleic Acid/Steroid Metabolism
slopeDiff(AD−CT): −0.0465
Normal:
AD:
log2 normalized relative abundance
0.5
0.0
−0.5
CSF
Plasma
log2 normalized relative abundance
0.5
0.0
−0.5
CSF
Plasma

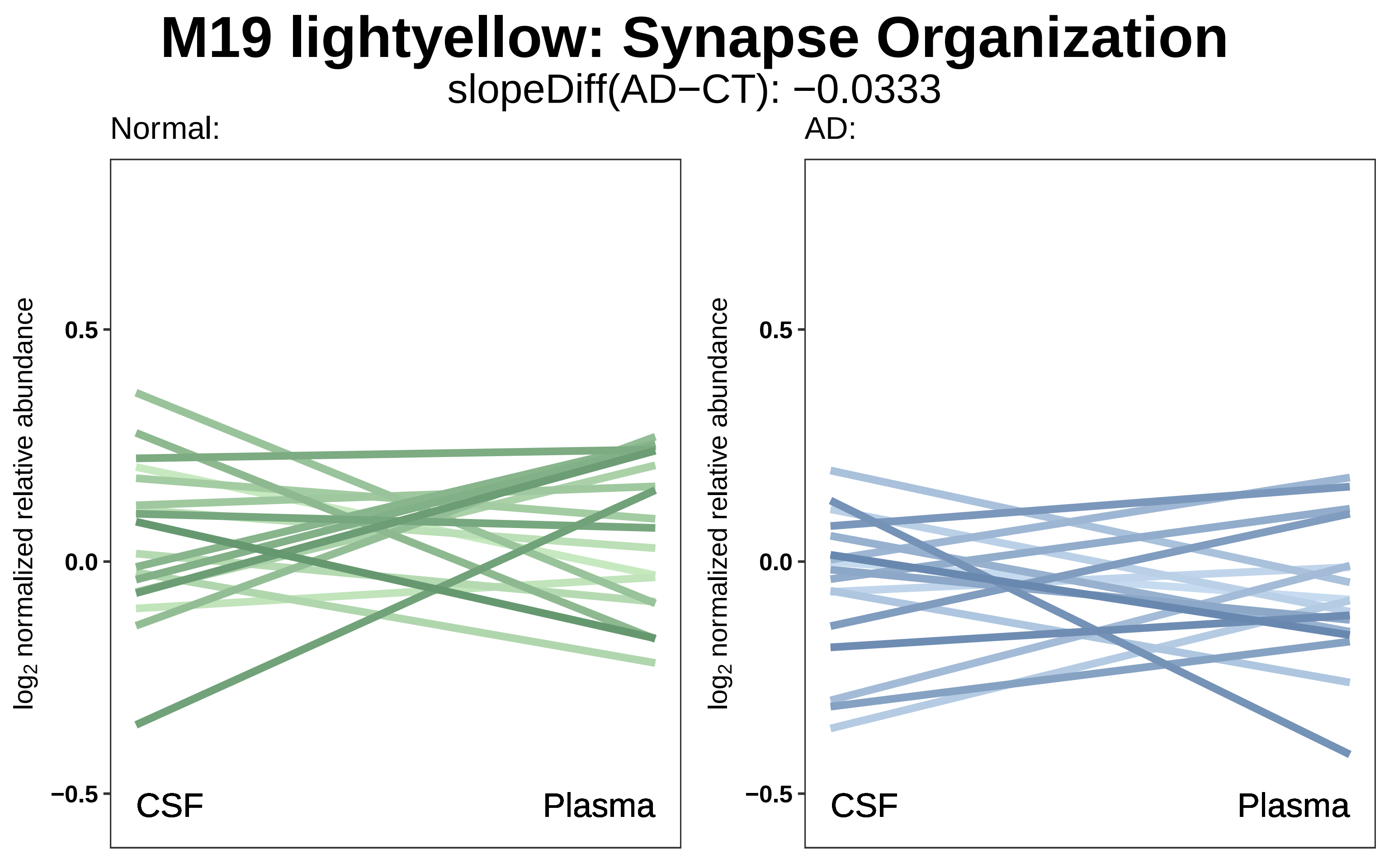
M19 lightyellow: Synapse Organization
slopeDiff(AD−CT): −0.0333
Normal:
AD:
log2 normalized relative abundance
0.5
0.0
−0.5
CSF
Plasma
log2 normalized relative abundance
0.5
0.0
−0.5
CSF
Plasma

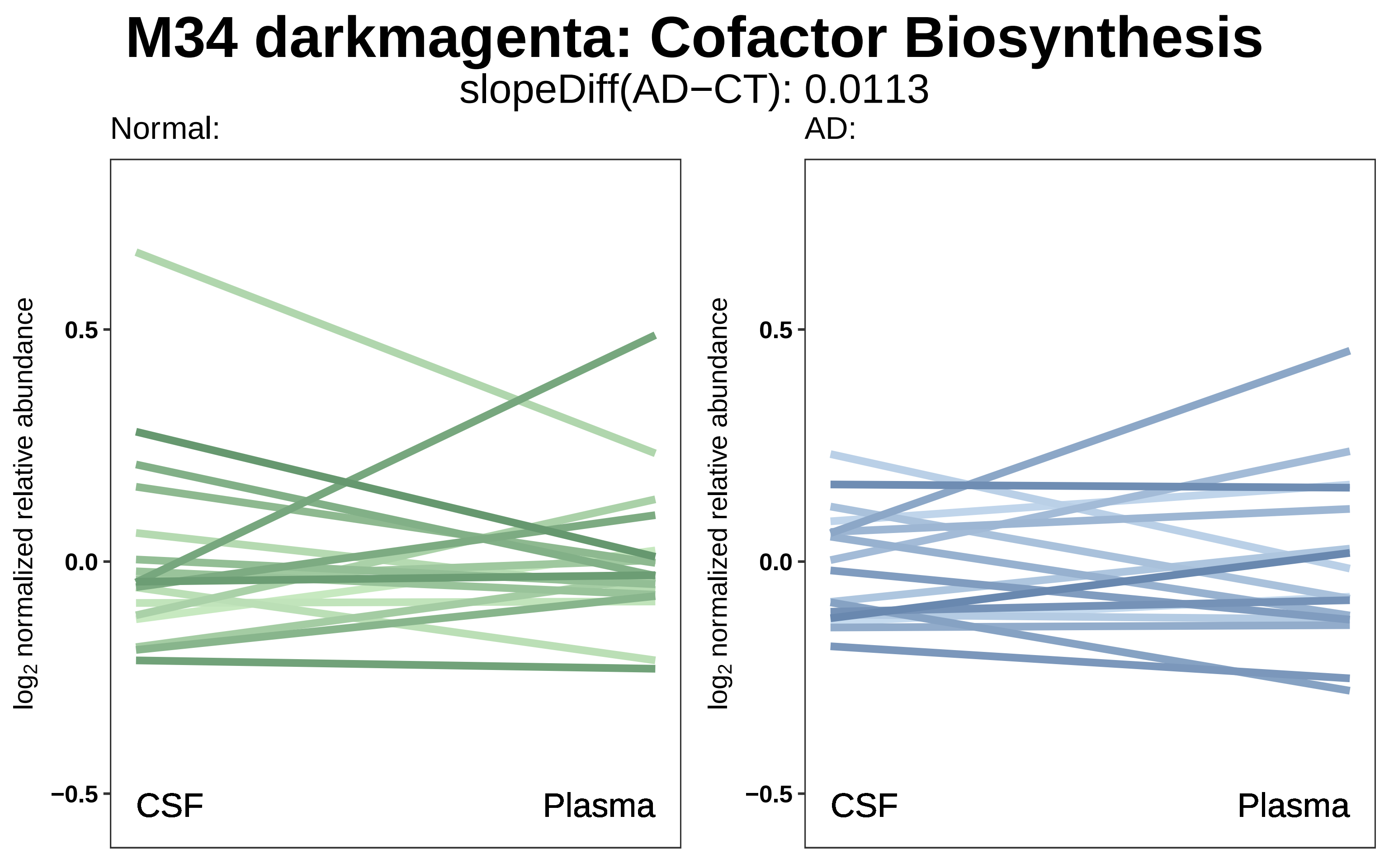
M34 darkmagenta: Cofactor Biosynthesis
slopeDiff(AD−CT): 0.0113
Normal:
AD:
log2 normalized relative abundance
0.5
0.0
−0.5
CSF
Plasma
log2 normalized relative abundance
0.5
0.0
−0.5
CSF
Plasma

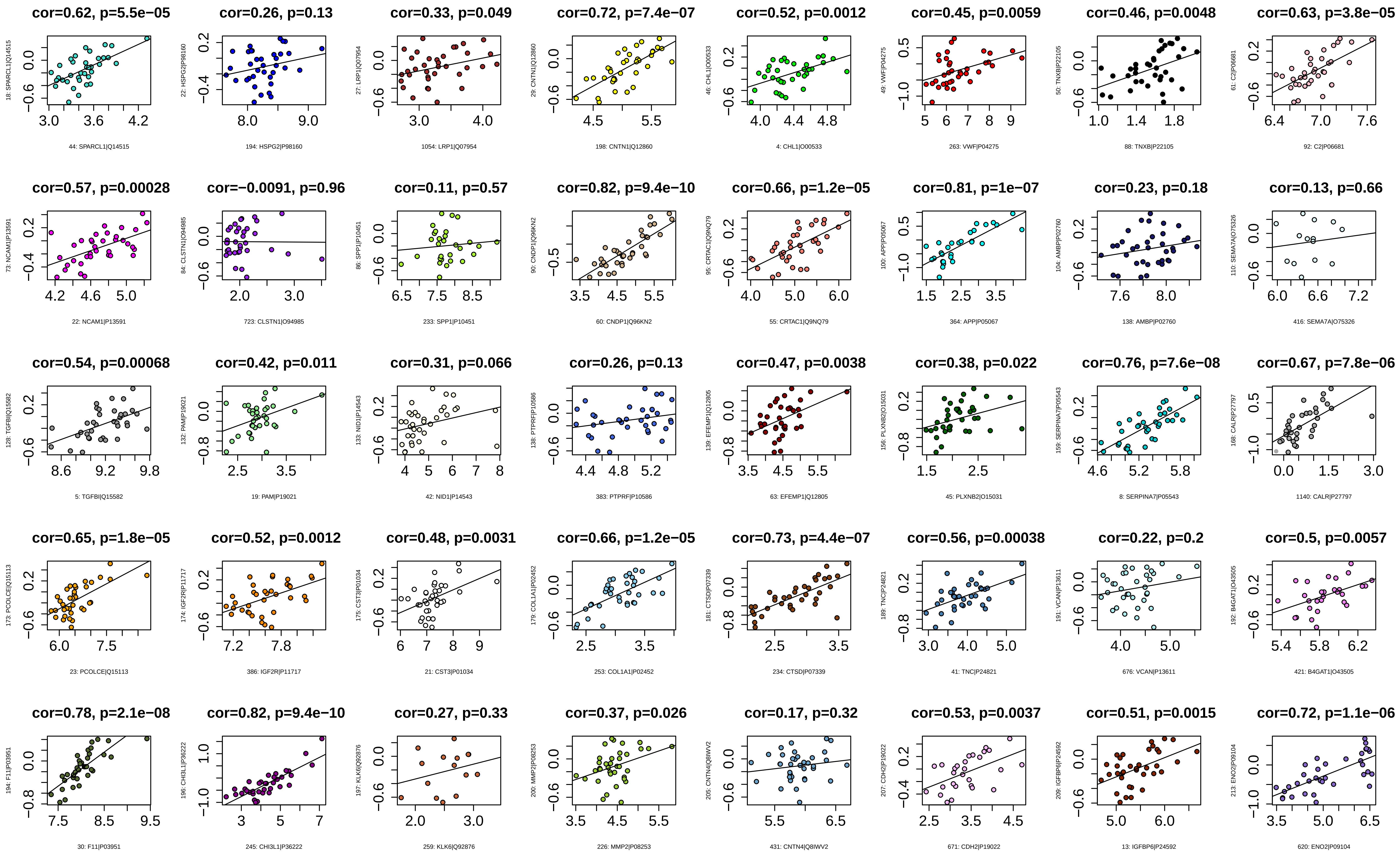
cor=0.62, p=5.5e−05
18: SPARCL1|Q14515
44: SPARCL1|Q14515
cor=0.26, p=0.13
22: HSPG2|P98160
194: HSPG2|P98160
cor=0.33, p=0.049
27: LRP1|Q07954
1054: LRP1|Q07954
cor=0.72, p=7.4e−07
29: CNTN1|Q12860
198: CNTN1|Q12860
cor=0.52, p=0.0012
46: CHL1|O00533
4: CHL1|O00533
cor=0.45, p=0.0059
49: VWF|P04275
263: VWF|P04275
cor=0.46, p=0.0048
50: TNXB|P22105
88: TNXB|P22105
cor=0.63, p=3.8e−05
61: C2|P06681
92: C2|P06681
cor=0.57, p=0.00028
73: NCAM1|P13591
22: NCAM1|P13591
cor=−0.0091, p=0.96
84: CLSTN1|O94985
723: CLSTN1|O94985
cor=0.11, p=0.57
86: SPP1|P10451
233: SPP1|P10451
cor=0.82, p=9.4e−10
90: CNDP1|Q96KN2
60: CNDP1|Q96KN2
cor=0.66, p=1.2e−05
95: CRTAC1|Q9NQ79
55: CRTAC1|Q9NQ79
cor=0.81, p=1e−07
100: APP|P05067
364: APP|P05067
cor=0.23, p=0.18
104: AMBP|P02760
138: AMBP|P02760
cor=0.13, p=0.66
110: SEMA7A|O75326
416: SEMA7A|O75326
cor=0.54, p=0.00068
128: TGFBI|Q15582
5: TGFBI|Q15582
cor=0.42, p=0.011
132: PAM|P19021
19: PAM|P19021
cor=0.31, p=0.066
133: NID1|P14543
42: NID1|P14543
cor=0.26, p=0.13
138: PTPRF|P10586
383: PTPRF|P10586
cor=0.47, p=0.0038
139: EFEMP1|Q12805
63: EFEMP1|Q12805
cor=0.38, p=0.022
156: PLXNB2|O15031
45: PLXNB2|O15031
cor=0.76, p=7.6e−08
159: SERPINA7|P05543
8: SERPINA7|P05543
cor=0.67, p=7.8e−06
168: CALR|P27797
1140: CALR|P27797
cor=0.65, p=1.8e−05
173: PCOLCE|Q15113
23: PCOLCE|Q15113
cor=0.52, p=0.0012
174: IGF2R|P11717
386: IGF2R|P11717
cor=0.48, p=0.0031
175: CST3|P01034
21: CST3|P01034
cor=0.66, p=1.2e−05
179: COL1A1|P02452
253: COL1A1|P02452
cor=0.73, p=4.4e−07
181: CTSD|P07339
234: CTSD|P07339
cor=0.56, p=0.00038
189: TNC|P24821
41: TNC|P24821
cor=0.22, p=0.2
191: VCAN|P13611
676: VCAN|P13611
cor=0.5, p=0.0057
192: B4GAT1|O43505
421: B4GAT1|O43505
cor=0.78, p=2.1e−08
194: F11|P03951
30: F11|P03951
cor=0.82, p=9.4e−10
196: CHI3L1|P36222
245: CHI3L1|P36222
cor=0.27, p=0.33
197: KLK6|Q92876
259: KLK6|Q92876
cor=0.37, p=0.026
200: MMP2|P08253
226: MMP2|P08253
cor=0.17, p=0.32
205: CNTN4|Q8IWV2
431: CNTN4|Q8IWV2
cor=0.53, p=0.0037
207: CDH2|P19022
671: CDH2|P19022
cor=0.51, p=0.0015
209: IGFBP6|P24592
13: IGFBP6|P24592
cor=0.72, p=1.1e−06
213: ENO2|P09104
620: ENO2|P09104

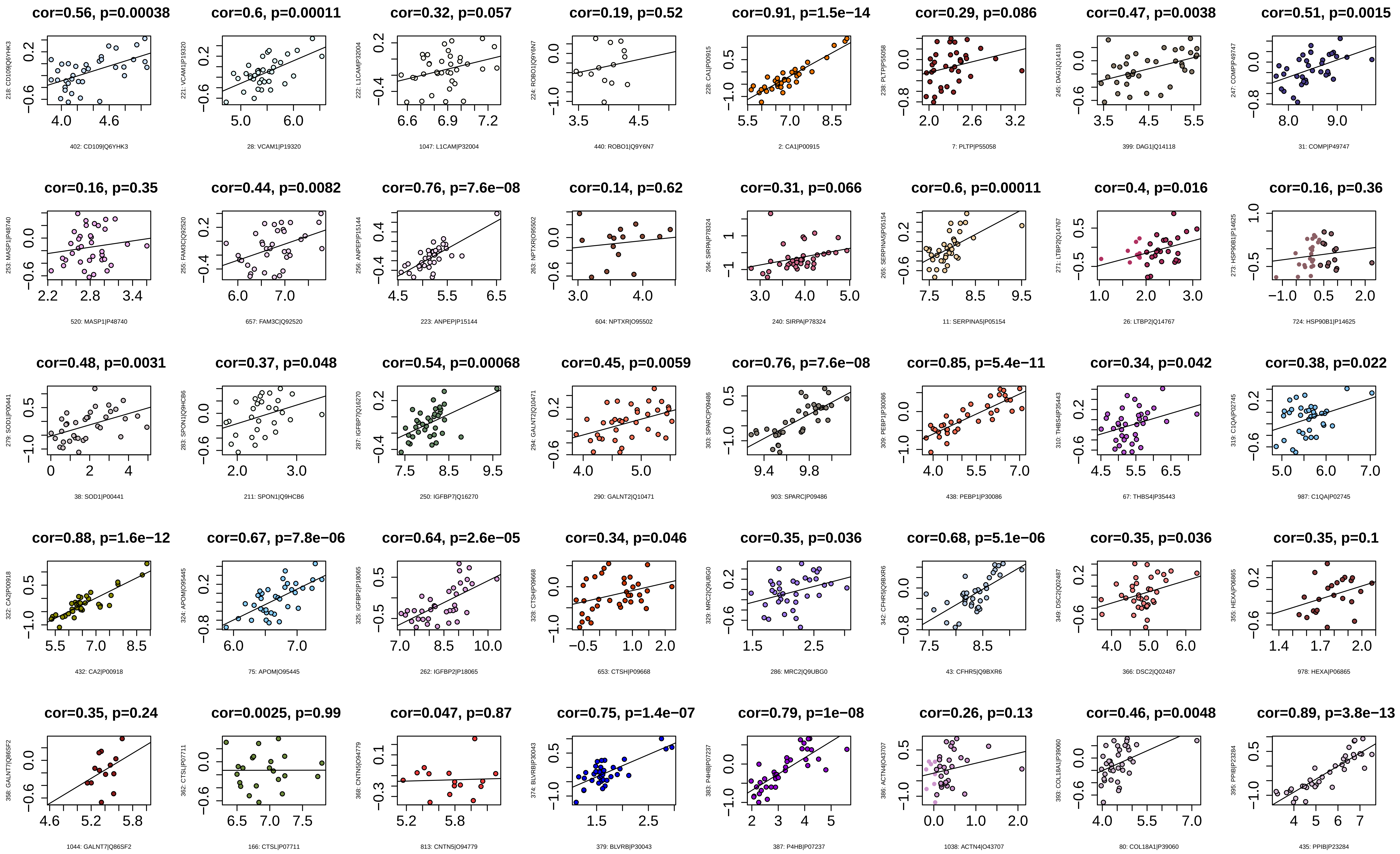
cor=0.56, p=0.00038
218: CD109|Q6YHK3
402: CD109|Q6YHK3
cor=0.6, p=0.00011
221: VCAM1|P19320
28: VCAM1|P19320
cor=0.32, p=0.057
222: L1CAM|P32004
1047: L1CAM|P32004
cor=0.19, p=0.52
224: ROBO1|Q9Y6N7
440: ROBO1|Q9Y6N7
cor=0.91, p=1.5e−14
228: CA1|P00915
2: CA1|P00915
cor=0.29, p=0.086
238: PLTP|P55058
7: PLTP|P55058
cor=0.47, p=0.0038
245: DAG1|Q14118
399: DAG1|Q14118
cor=0.51, p=0.0015
247: COMP|P49747
31: COMP|P49747
cor=0.16, p=0.35
253: MASP1|P48740
520: MASP1|P48740
cor=0.44, p=0.0082
255: FAM3C|Q92520
657: FAM3C|Q92520
cor=0.76, p=7.6e−08
256: ANPEP|P15144
223: ANPEP|P15144
cor=0.14, p=0.62
263: NPTXR|O95502
604: NPTXR|O95502
cor=0.31, p=0.066
264: SIRPA|P78324
240: SIRPA|P78324
cor=0.6, p=0.00011
265: SERPINA5|P05154
11: SERPINA5|P05154
cor=0.4, p=0.016
271: LTBP2|Q14767
26: LTBP2|Q14767
cor=0.16, p=0.36
273: HSP90B1|P14625
724: HSP90B1|P14625
cor=0.48, p=0.0031
279: SOD1|P00441
38: SOD1|P00441
cor=0.37, p=0.048
283: SPON1|Q9HCB6
211: SPON1|Q9HCB6
cor=0.54, p=0.00068
287: IGFBP7|Q16270
250: IGFBP7|Q16270
cor=0.45, p=0.0059
294: GALNT2|Q10471
290: GALNT2|Q10471
cor=0.76, p=7.6e−08
303: SPARC|P09486
903: SPARC|P09486
cor=0.85, p=5.4e−11
309: PEBP1|P30086
438: PEBP1|P30086
cor=0.34, p=0.042
310: THBS4|P35443
67: THBS4|P35443
cor=0.38, p=0.022
319: C1QA|P02745
987: C1QA|P02745
cor=0.88, p=1.6e−12
322: CA2|P00918
432: CA2|P00918
cor=0.67, p=7.8e−06
324: APOM|O95445
75: APOM|O95445
cor=0.64, p=2.6e−05
325: IGFBP2|P18065
262: IGFBP2|P18065
cor=0.34, p=0.046
328: CTSH|P09668
653: CTSH|P09668
cor=0.35, p=0.036
329: MRC2|Q9UBG0
286: MRC2|Q9UBG0
cor=0.68, p=5.1e−06
342: CFHR5|Q9BXR6
43: CFHR5|Q9BXR6
cor=0.35, p=0.036
349: DSC2|Q02487
366: DSC2|Q02487
cor=0.35, p=0.1
355: HEXA|P06865
978: HEXA|P06865
cor=0.35, p=0.24
358: GALNT7|Q86SF2
1044: GALNT7|Q86SF2
cor=0.0025, p=0.99
362: CTSL|P07711
166: CTSL|P07711
cor=0.047, p=0.87
368: CNTN5|O94779
813: CNTN5|O94779
cor=0.75, p=1.4e−07
374: BLVRB|P30043
379: BLVRB|P30043
cor=0.79, p=1e−08
383: P4HB|P07237
387: P4HB|P07237
cor=0.26, p=0.13
386: ACTN4|O43707
1038: ACTN4|O43707
cor=0.46, p=0.0048
393: COL18A1|P39060
80: COL18A1|P39060
cor=0.89, p=3.8e−13
395: PPIB|P23284
435: PPIB|P23284

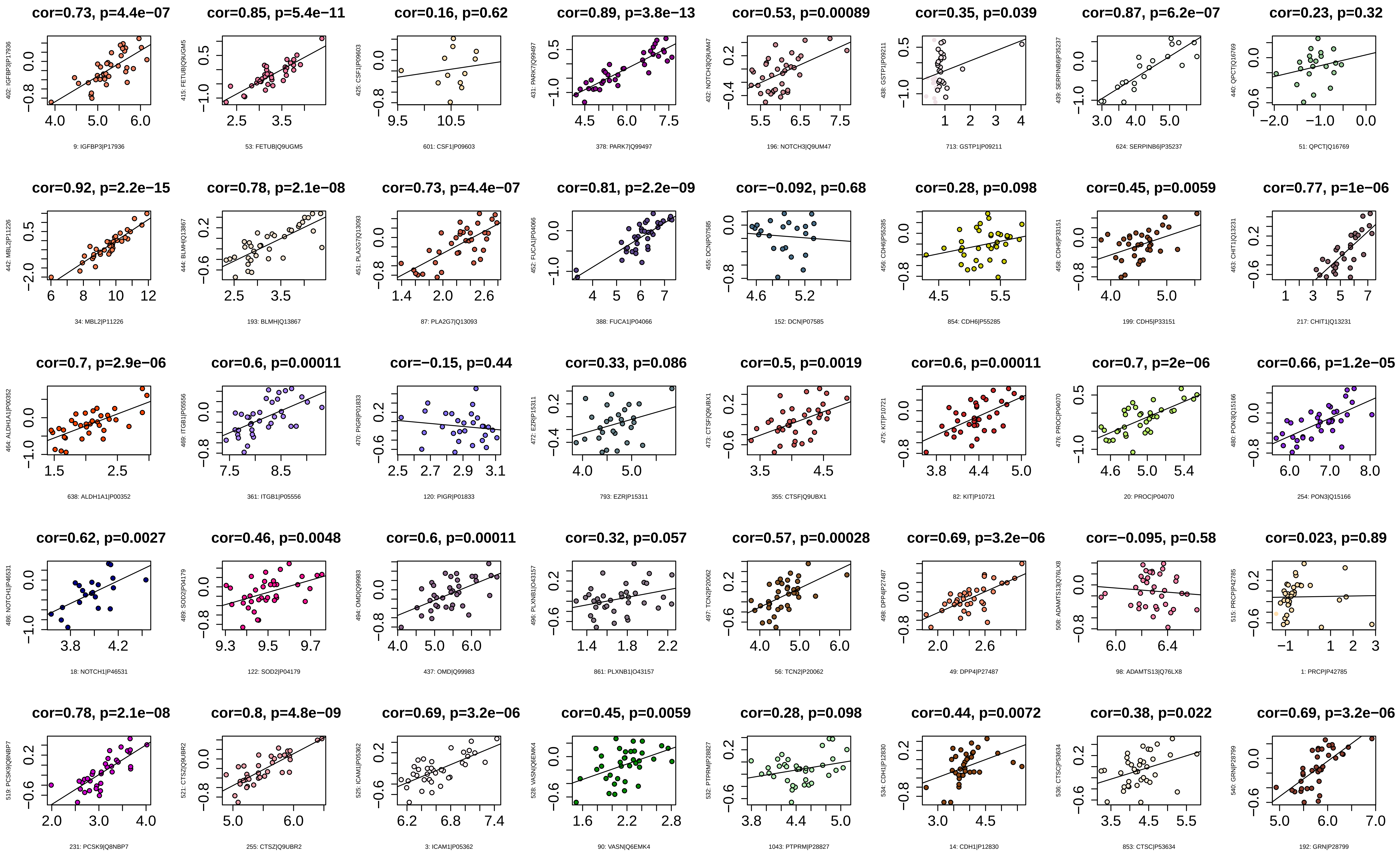
cor=0.73, p=4.4e−07
402: IGFBP3|P17936
9: IGFBP3|P17936
cor=0.85, p=5.4e−11
415: FETUB|Q9UGM5
53: FETUB|Q9UGM5
cor=0.16, p=0.62
425: CSF1|P09603
601: CSF1|P09603
cor=0.89, p=3.8e−13
431: PARK7|Q99497
378: PARK7|Q99497
cor=0.53, p=0.00089
432: NOTCH3|Q9UM47
196: NOTCH3|Q9UM47
cor=0.35, p=0.039
438: GSTP1|P09211
713: GSTP1|P09211
cor=0.87, p=6.2e−07
439: SERPINB6|P35237
624: SERPINB6|P35237
cor=0.23, p=0.32
440: QPCT|Q16769
51: QPCT|Q16769
cor=0.92, p=2.2e−15
442: MBL2|P11226
34: MBL2|P11226
cor=0.78, p=2.1e−08
444: BLMH|Q13867
193: BLMH|Q13867
cor=0.73, p=4.4e−07
451: PLA2G7|Q13093
87: PLA2G7|Q13093
cor=0.81, p=2.2e−09
452: FUCA1|P04066
388: FUCA1|P04066
cor=−0.092, p=0.68
455: DCN|P07585
152: DCN|P07585
cor=0.28, p=0.098
456: CDH6|P55285
854: CDH6|P55285
cor=0.45, p=0.0059
458: CDH5|P33151
199: CDH5|P33151
cor=0.77, p=1e−06
463: CHIT1|Q13231
217: CHIT1|Q13231
cor=0.7, p=2.9e−06
464: ALDH1A1|P00352
638: ALDH1A1|P00352
cor=0.6, p=0.00011
469: ITGB1|P05556
361: ITGB1|P05556
cor=−0.15, p=0.44
470: PIGR|P01833
120: PIGR|P01833
cor=0.33, p=0.086
472: EZR|P15311
793: EZR|P15311
cor=0.5, p=0.0019
473: CTSF|Q9UBX1
355: CTSF|Q9UBX1
cor=0.6, p=0.00011
475: KIT|P10721
82: KIT|P10721
cor=0.7, p=2e−06
476: PROC|P04070
20: PROC|P04070
cor=0.66, p=1.2e−05
480: PON3|Q15166
254: PON3|Q15166
cor=0.62, p=0.0027
486: NOTCH1|P46531
18: NOTCH1|P46531
cor=0.46, p=0.0048
489: SOD2|P04179
122: SOD2|P04179
cor=0.6, p=0.00011
494: OMD|Q99983
437: OMD|Q99983
cor=0.32, p=0.057
496: PLXNB1|O43157
861: PLXNB1|O43157
cor=0.57, p=0.00028
497: TCN2|P20062
56: TCN2|P20062
cor=0.69, p=3.2e−06
498: DPP4|P27487
49: DPP4|P27487
cor=−0.095, p=0.58
508: ADAMTS13|Q76LX8
98: ADAMTS13|Q76LX8
cor=0.023, p=0.89
515: PRCP|P42785
1: PRCP|P42785
cor=0.78, p=2.1e−08
519: PCSK9|Q8NBP7
231: PCSK9|Q8NBP7
cor=0.8, p=4.8e−09
521: CTSZ|Q9UBR2
255: CTSZ|Q9UBR2
cor=0.69, p=3.2e−06
525: ICAM1|P05362
3: ICAM1|P05362
cor=0.45, p=0.0059
528: VASN|Q6EMK4
90: VASN|Q6EMK4
cor=0.28, p=0.098
532: PTPRM|P28827
1043: PTPRM|P28827
cor=0.44, p=0.0072
534: CDH1|P12830
14: CDH1|P12830
cor=0.38, p=0.022
536: CTSC|P53634
853: CTSC|P53634
cor=0.69, p=3.2e−06
540: GRN|P28799
192: GRN|P28799

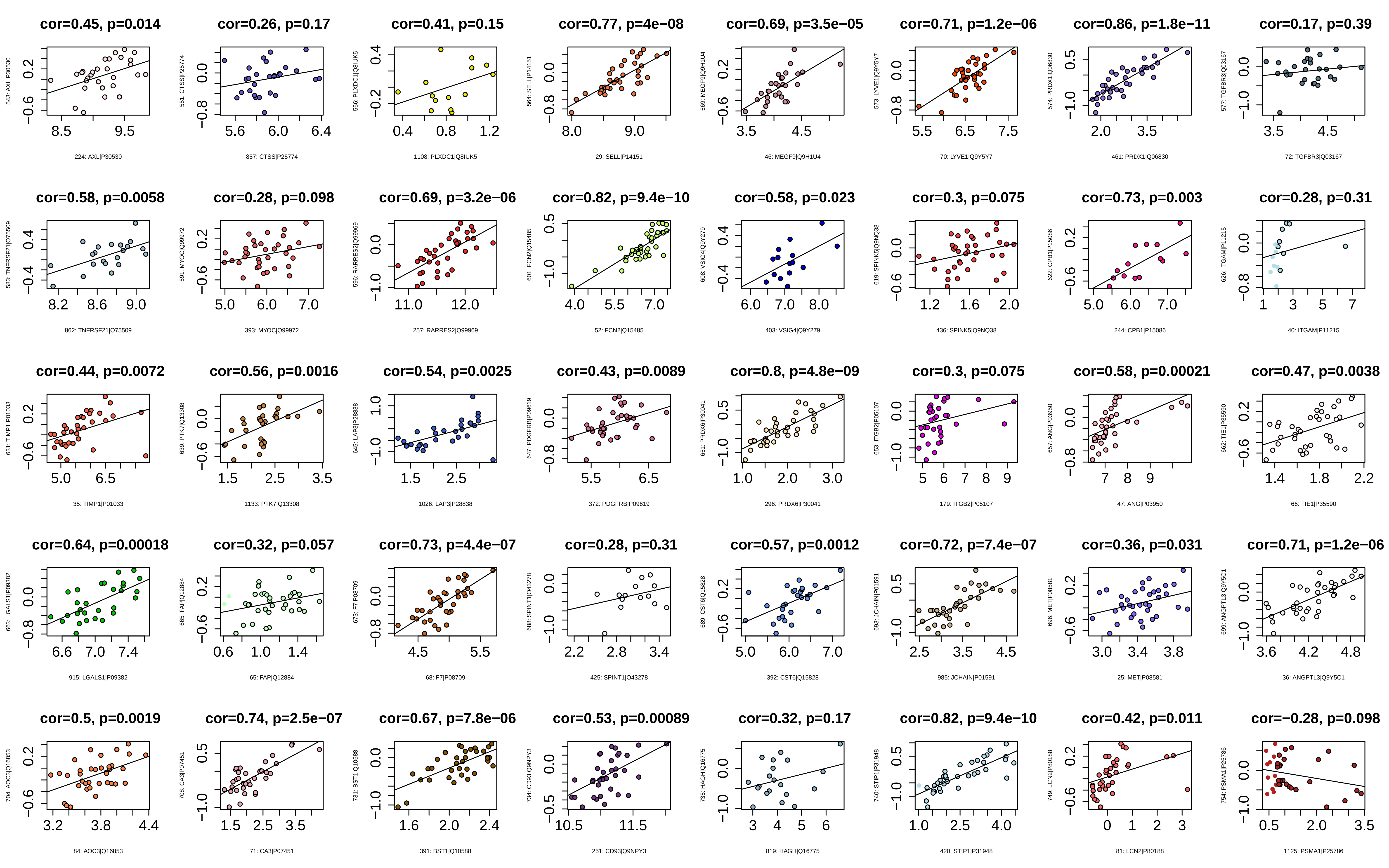
cor=0.45, p=0.014
224: AXL|P30530
cor=0.26, p=0.17
857: CTSS|P25774
cor=0.41, p=0.15
1108: PLXDC1|Q8IUK5
cor=0.77, p=4e−08
29: SELL|P14151
cor=0.69, p=3.5e−05
46: MEGF9|Q9H1U4
cor=0.71, p=1.2e−06
70: LYVE1|Q9Y5Y7
cor=0.86, p=1.8e−11
461: PRDX1|Q06830
cor=0.17, p=0.39
72: TGFBR3|Q03167
cor=0.58, p=0.0058
862: TNFRSF21|O75509
cor=0.28, p=0.098
393: MYOC|Q99972
cor=0.69, p=3.2e−06
257: RARRES2|Q99969
cor=0.82, p=9.4e−10
52: FCN2|Q15485
cor=0.58, p=0.023
403: VSIG4|Q9Y279
cor=0.3, p=0.075
436: SPINK5|Q9NQ38
cor=0.73, p=0.003
244: CPB1|P15086
cor=0.28, p=0.31
40: ITGAM|P11215
cor=0.44, p=0.0072
35: TIMP1|P01033
cor=0.56, p=0.0016
1133: PTK7|Q13308
cor=0.54, p=0.0025
1026: LAP3|P28838
cor=0.43, p=0.0089
372: PDGFRB|P09619
cor=0.8, p=4.8e−09
296: PRDX6|P30041
cor=0.3, p=0.075
179: ITGB2|P05107
cor=0.58, p=0.00021
47: ANG|P03950
cor=0.47, p=0.0038
66: TIE1|P35590
cor=0.64, p=0.00018
915: LGALS1|P09382
cor=0.32, p=0.057
65: FAP|Q12884
cor=0.73, p=4.4e−07
68: F7|P08709
cor=0.28, p=0.31
425: SPINT1|O43278
cor=0.57, p=0.0012
392: CST6|Q15828
cor=0.72, p=7.4e−07
985: JCHAIN|P01591
cor=0.36, p=0.031
25: MET|P08581
cor=0.71, p=1.2e−06
36: ANGPTL3|Q9Y5C1
cor=0.5, p=0.0019
84: AOC3|Q16853
cor=0.74, p=2.5e−07
71: CA3|P07451
cor=0.67, p=7.8e−06
391: BST1|Q10588
cor=0.53, p=0.00089
251: CD93|Q9NPY3
cor=0.32, p=0.17
819: HAGH|Q16775
cor=0.82, p=9.4e−10
420: STIP1|P31948
cor=0.42, p=0.011
81: LCN2|P80188
cor=−0.28, p=0.098
1125: PSMA1|P25786

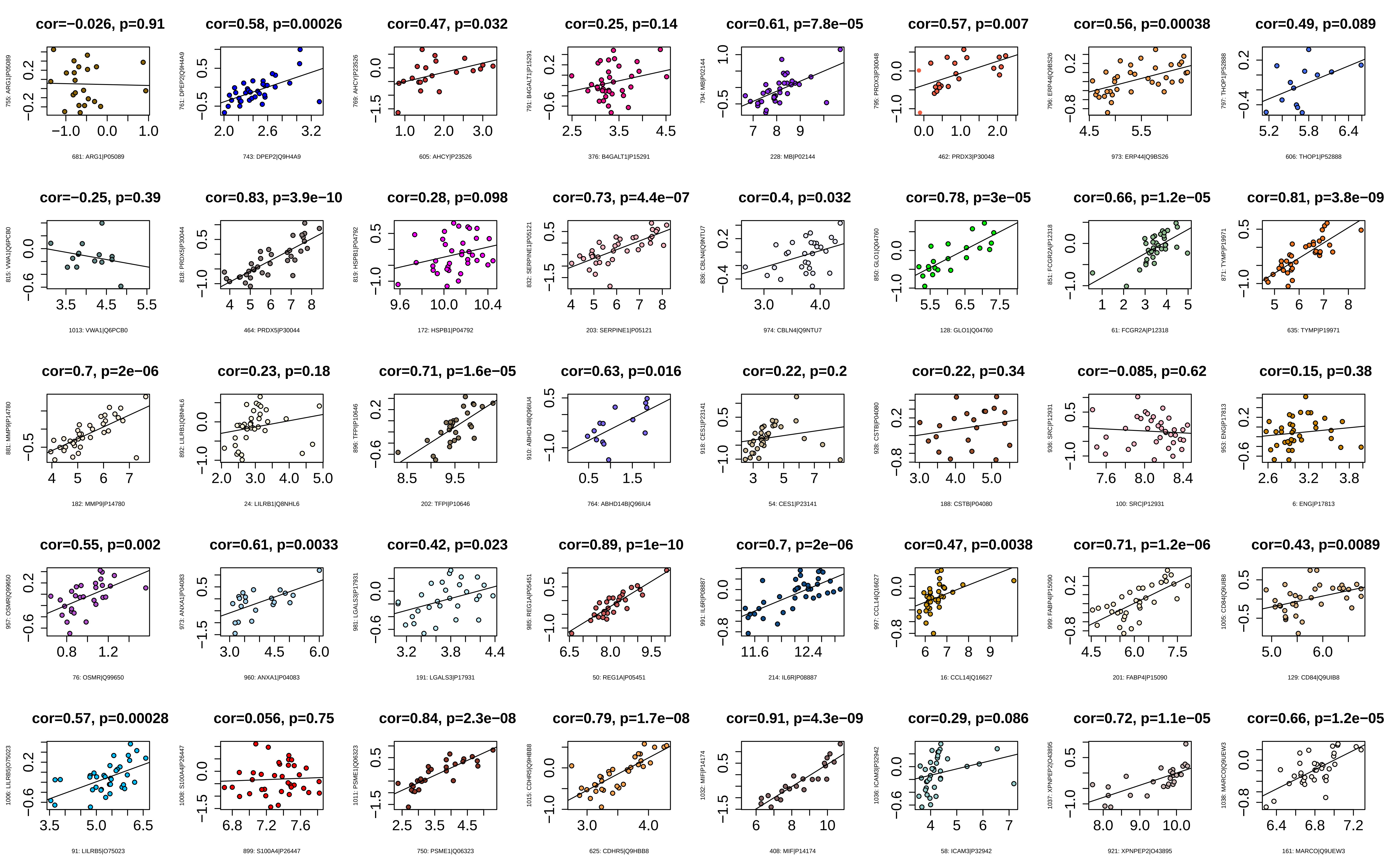
cor=−0.026, p=0.91
755: ARG1|P05089
681: ARG1|P05089
cor=0.58, p=0.00026
761: DPEP2|Q9H4A9
743: DPEP2|Q9H4A9
cor=0.47, p=0.032
769: AHCY|P23526
605: AHCY|P23526
cor=0.25, p=0.14
791: B4GALT1|P15291
376: B4GALT1|P15291
cor=0.61, p=7.8e−05
794: MB|P02144
228: MB|P02144
cor=0.57, p=0.007
795: PRDX3|P30048
462: PRDX3|P30048
cor=0.56, p=0.00038
796: ERP44|Q9BS26
973: ERP44|Q9BS26
cor=0.49, p=0.089
797: THOP1|P52888
606: THOP1|P52888
cor=−0.25, p=0.39
815: VWA1|Q6PCB0
1013: VWA1|Q6PCB0
cor=0.83, p=3.9e−10
818: PRDX5|P30044
464: PRDX5|P30044
cor=0.28, p=0.098
819: HSPB1|P04792
172: HSPB1|P04792
cor=0.73, p=4.4e−07
832: SERPINE1|P05121
203: SERPINE1|P05121
cor=0.4, p=0.032
836: CBLN4|Q9NTU7
974: CBLN4|Q9NTU7
cor=0.78, p=3e−05
850: GLO1|Q04760
128: GLO1|Q04760
cor=0.66, p=1.2e−05
851: FCGR2A|P12318
61: FCGR2A|P12318
cor=0.81, p=3.8e−09
871: TYMP|P19971
635: TYMP|P19971
cor=0.7, p=2e−06
881: MMP9|P14780
182: MMP9|P14780
cor=0.23, p=0.18
892: LILRB1|Q8NHL6
24: LILRB1|Q8NHL6
cor=0.71, p=1.6e−05
896: TFPI|P10646
202: TFPI|P10646
cor=0.63, p=0.016
910: ABHD14B|Q96IU4
764: ABHD14B|Q96IU4
cor=0.22, p=0.2
918: CES1|P23141
54: CES1|P23141
cor=0.22, p=0.34
928: CSTB|P04080
188: CSTB|P04080
cor=−0.085, p=0.62
936: SRC|P12931
100: SRC|P12931
cor=0.15, p=0.38
953: ENG|P17813
6: ENG|P17813
cor=0.55, p=0.002
957: OSMR|Q99650
76: OSMR|Q99650
cor=0.61, p=0.0033
973: ANXA1|P04083
960: ANXA1|P04083
cor=0.42, p=0.023
981: LGALS3|P17931
191: LGALS3|P17931
cor=0.89, p=1e−10
985: REG1A|P05451
50: REG1A|P05451
cor=0.7, p=2e−06
991: IL6R|P08887
214: IL6R|P08887
cor=0.47, p=0.0038
997: CCL14|Q16627
16: CCL14|Q16627
cor=0.71, p=1.2e−06
999: FABP4|P15090
201: FABP4|P15090
cor=0.43, p=0.0089
1005: CD84|Q9UIB8
129: CD84|Q9UIB8
cor=0.57, p=0.00028
1006: LILRB5|O75023
91: LILRB5|O75023
cor=0.056, p=0.75
1008: S100A4|P26447
899: S100A4|P26447
cor=0.84, p=2.3e−08
1011: PSME1|Q06323
750: PSME1|Q06323
cor=0.79, p=1.7e−08
1015: CDHR5|Q9HBB8
625: CDHR5|Q9HBB8
cor=0.91, p=4.3e−09
1032: MIF|P14174
408: MIF|P14174
cor=0.29, p=0.086
1036: ICAM3|P32942
58: ICAM3|P32942
cor=0.72, p=1.1e−05
1037: XPNPEP2|O43895
921: XPNPEP2|O43895
cor=0.66, p=1.2e−05
1038: MARCO|Q9UEW3
161: MARCO|Q9UEW3

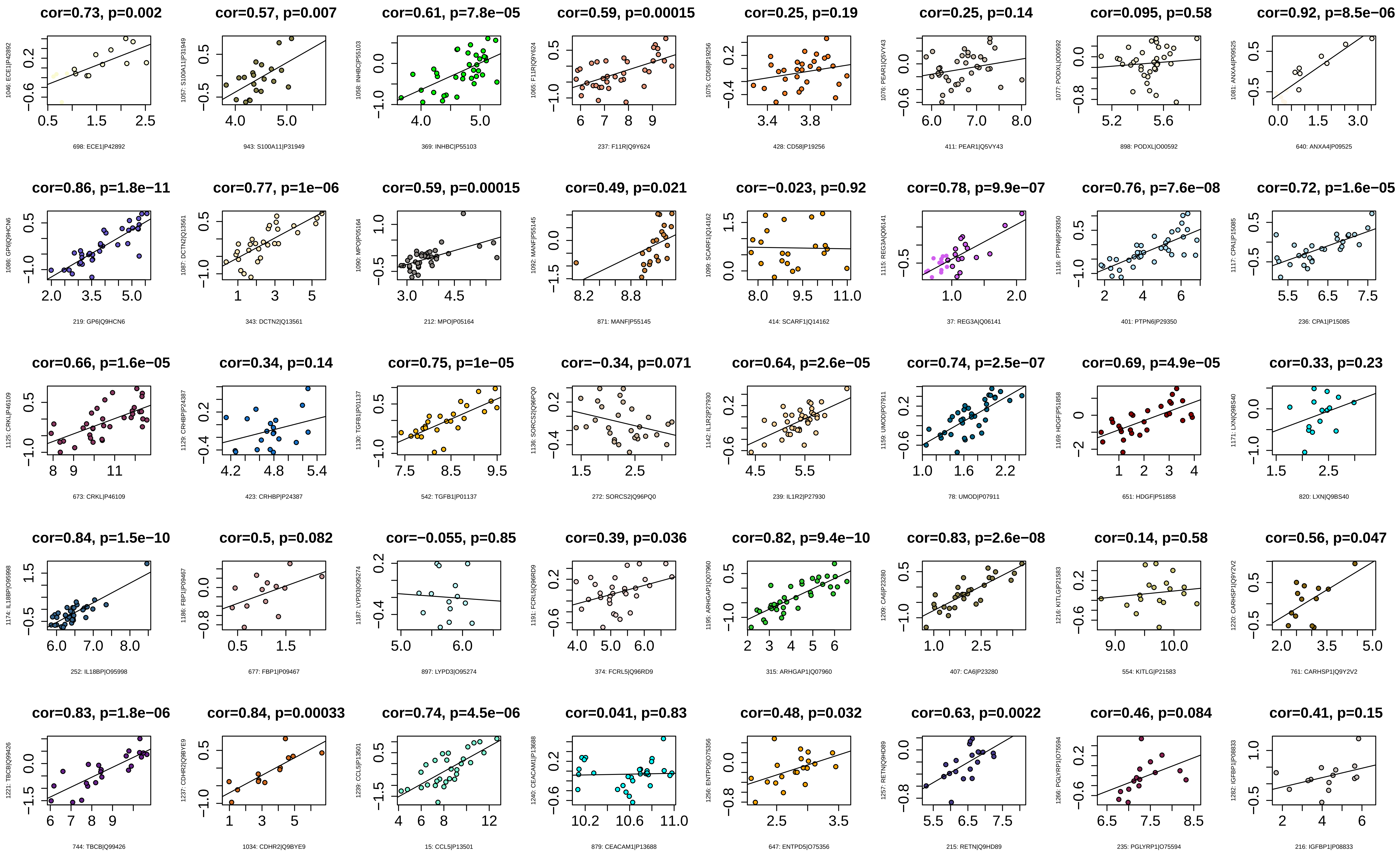
cor=0.73, p=0.002
1046: ECE1|P42892
698: ECE1|P42892
cor=0.57, p=0.007
1057: S100A11|P31949
943: S100A11|P31949
cor=0.61, p=7.8e−05
1058: INHBC|P55103
369: INHBC|P55103
cor=0.59, p=0.00015
1065: F11R|Q9Y624
237: F11R|Q9Y624
cor=0.25, p=0.19
1075: CD58|P19256
428: CD58|P19256
cor=0.25, p=0.14
1076: PEAR1|Q5VY43
411: PEAR1|Q5VY43
cor=0.095, p=0.58
1077: PODXL|O00592
898: PODXL|O00592
cor=0.92, p=8.5e−06
1081: ANXA4|P09525
640: ANXA4|P09525
cor=0.86, p=1.8e−11
1086: GP6|Q9HCN6
219: GP6|Q9HCN6
cor=0.77, p=1e−06
1087: DCTN2|Q13561
343: DCTN2|Q13561
cor=0.59, p=0.00015
1090: MPO|P05164
212: MPO|P05164
cor=0.49, p=0.021
1092: MANF|P55145
871: MANF|P55145
cor=−0.023, p=0.92
1099: SCARF1|Q14162
414: SCARF1|Q14162
cor=0.78, p=9.9e−07
1115: REG3A|Q06141
37: REG3A|Q06141
cor=0.76, p=7.6e−08
1116: PTPN6|P29350
401: PTPN6|P29350
cor=0.72, p=1.6e−05
1117: CPA1|P15085
236: CPA1|P15085
cor=0.66, p=1.6e−05
1125: CRKL|P46109
673: CRKL|P46109
cor=0.34, p=0.14
1129: CRHBP|P24387
423: CRHBP|P24387
cor=0.75, p=1e−05
1130: TGFB1|P01137
542: TGFB1|P01137
cor=−0.34, p=0.071
1136: SORCS2|Q96PQ0
272: SORCS2|Q96PQ0
cor=0.64, p=2.6e−05
1142: IL1R2|P27930
239: IL1R2|P27930
cor=0.74, p=2.5e−07
1159: UMOD|P07911
78: UMOD|P07911
cor=0.69, p=4.9e−05
1169: HDGF|P51858
651: HDGF|P51858
cor=0.33, p=0.23
1171: LXN|Q9BS40
820: LXN|Q9BS40
cor=0.84, p=1.5e−10
1174: IL18BP|O95998
252: IL18BP|O95998
cor=0.5, p=0.082
1186: FBP1|P09467
677: FBP1|P09467
cor=−0.055, p=0.85
1187: LYPD3|O95274
897: LYPD3|O95274
cor=0.39, p=0.036
1191: FCRL5|Q96RD9
374: FCRL5|Q96RD9
cor=0.82, p=9.4e−10
1195: ARHGAP1|Q07960
315: ARHGAP1|Q07960
cor=0.83, p=2.6e−08
1209: CA6|P23280
407: CA6|P23280
cor=0.14, p=0.58
1216: KITLG|P21583
554: KITLG|P21583
cor=0.56, p=0.047
1220: CARHSP1|Q9Y2V2
761: CARHSP1|Q9Y2V2
cor=0.83, p=1.8e−06
1221: TBCB|Q99426
744: TBCB|Q99426
cor=0.84, p=0.00033
1237: CDHR2|Q9BYE9
1034: CDHR2|Q9BYE9
cor=0.74, p=4.5e−06
1239: CCL5|P13501
15: CCL5|P13501
cor=0.041, p=0.83
1240: CEACAM1|P13688
879: CEACAM1|P13688
cor=0.48, p=0.032
1256: ENTPD5|O75356
647: ENTPD5|O75356
cor=0.63, p=0.0022
1257: RETN|Q9HD89
215: RETN|Q9HD89
cor=0.46, p=0.084
1266: PGLYRP1|O75594
235: PGLYRP1|O75594
cor=0.41, p=0.15
1282: IGFBP1|P08833
216: IGFBP1|P08833

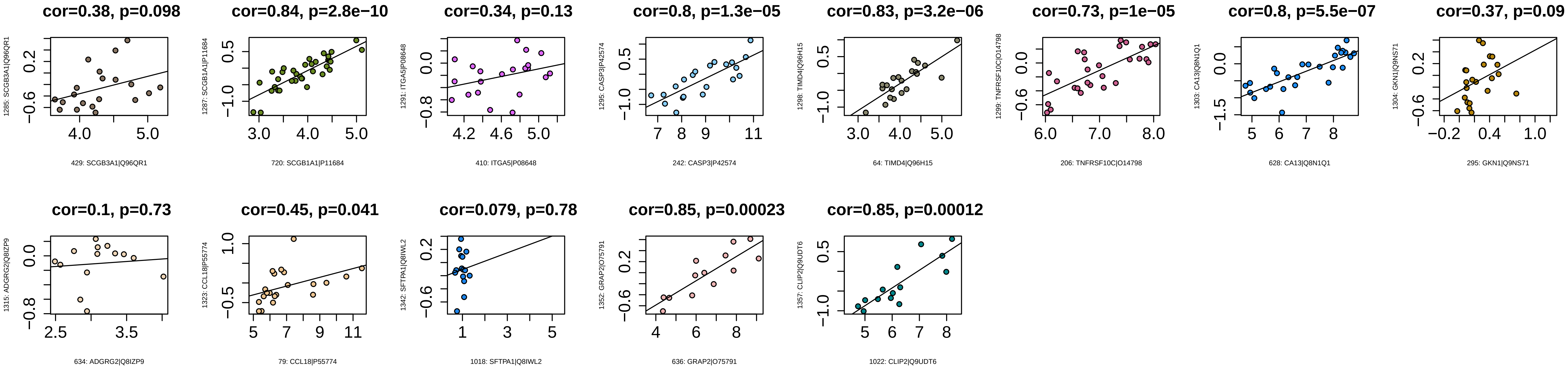
cor=0.38, p=0.098
1285: SCGB3A1|Q96QR1
429: SCGB3A1|Q96QR1
cor=0.84, p=2.8e−10
1287: SCGB1A1|P11684
720: SCGB1A1|P11684
cor=0.34, p=0.13
1291: ITGA5|P08648
410: ITGA5|P08648
cor=0.8, p=1.3e−05
1295: CASP3|P42574
242: CASP3|P42574
cor=0.83, p=3.2e−06
1298: TIMD4|Q96H15
64: TIMD4|Q96H15
cor=0.73, p=1e−05
1299: TNFRSF10C|O14798
206: TNFRSF10C|O14798
cor=0.8, p=5.5e−07
1303: CA13|Q8N1Q1
628: CA13|Q8N1Q1
cor=0.37, p=0.09
1304: GKN1|Q9NS71
295: GKN1|Q9NS71
cor=0.1, p=0.73
1315: ADGRG2|Q8IZP9
634: ADGRG2|Q8IZP9
cor=0.45, p=0.041
1323: CCL18|P55774
79: CCL18|P55774
cor=0.079, p=0.78
1342: SFTPA1|Q8IWL2
1018: SFTPA1|Q8IWL2
cor=0.85, p=0.00023
1352: GRAP2|O75791
636: GRAP2|O75791
cor=0.85, p=0.00012
1357: CLIP2|Q9UDT6
1022: CLIP2|Q9UDT6

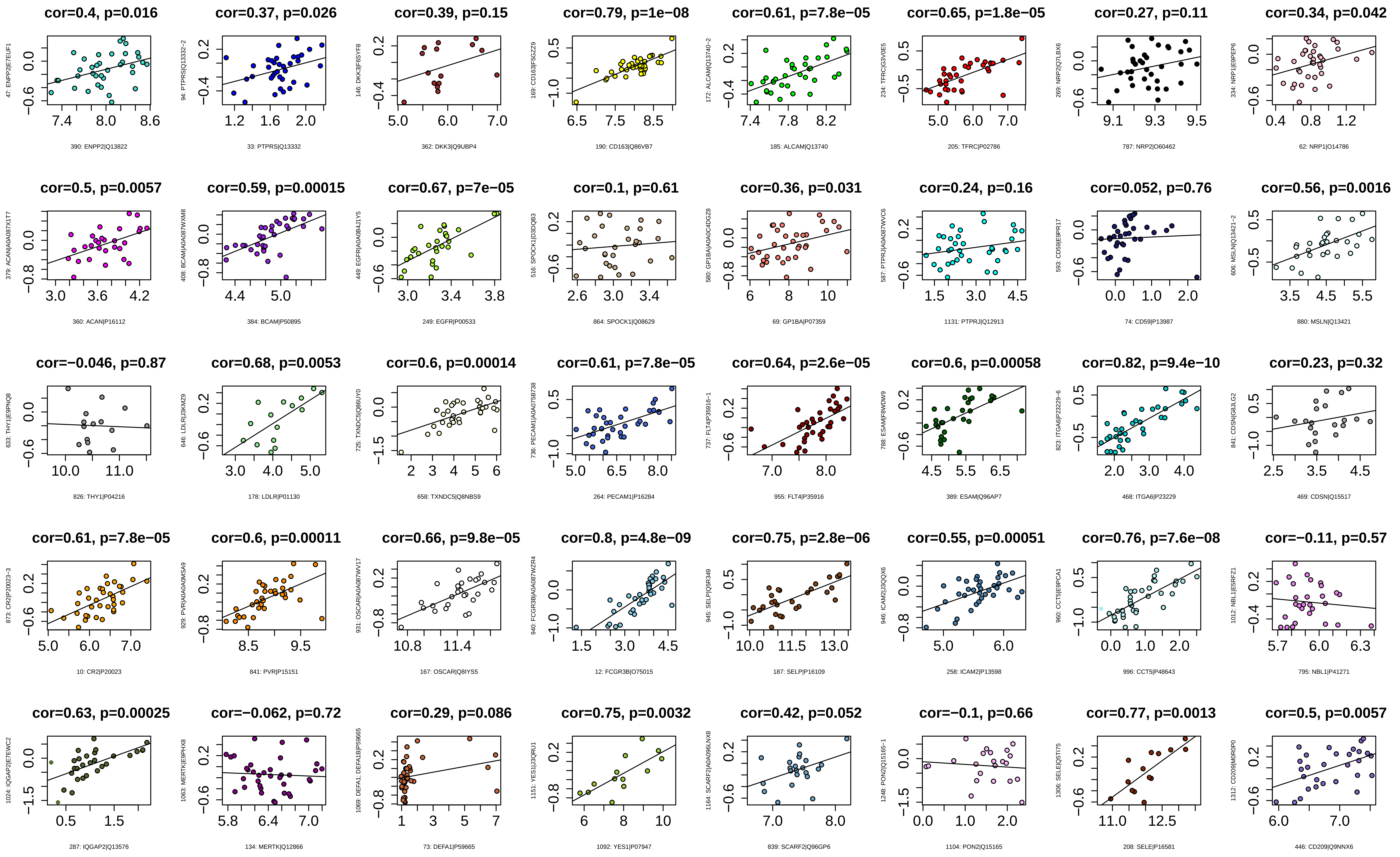
cor=0.4, p=0.016
390: ENPP2|Q13822
cor=0.37, p=0.026
33: PTPRS|Q13332
cor=0.39, p=0.15
362: DKK3|Q9UBP4
cor=0.79, p=1e−08
190: CD163|Q86VB7
cor=0.61, p=7.8e−05
185: ALCAM|Q13740
cor=0.65, p=1.8e−05
205: TFRC|P02786
cor=0.27, p=0.11
787: NRP2|O60462
cor=0.34, p=0.042
62: NRP1|O14786
cor=0.5, p=0.0057
360: ACAN|P16112
cor=0.59, p=0.00015
384: BCAM|P50895
cor=0.67, p=7e−05
249: EGFR|P00533
cor=0.1, p=0.61
864: SPOCK1|Q08629
cor=0.36, p=0.031
69: GP1BA|P07359
cor=0.24, p=0.16
1131: PTPRJ|Q12913
cor=0.052, p=0.76
74: CD59|P13987
cor=0.56, p=0.0016
880: MSLN|Q13421
cor=−0.046, p=0.87
826: THY1|P04216
cor=0.68, p=0.0053
178: LDLR|P01130
cor=0.6, p=0.00014
658: TXNDC5|Q8NBS9
cor=0.61, p=7.8e−05
264: PECAM1|P16284
cor=0.64, p=2.6e−05
955: FLT4|P35916
cor=0.6, p=0.00058
389: ESAM|Q96AP7
cor=0.82, p=9.4e−10
468: ITGA6|P23229
cor=0.23, p=0.32
469: CDSN|Q15517
cor=0.61, p=7.8e−05
10: CR2|P20023
cor=0.6, p=0.00011
841: PVR|P15151
cor=0.66, p=9.8e−05
167: OSCAR|Q8IYS5
cor=0.8, p=4.8e−09
12: FCGR3B|O75015
cor=0.75, p=2.8e−06
187: SELP|P16109
cor=0.55, p=0.00051
258: ICAM2|P13598
cor=0.76, p=7.6e−08
996: CCT5|P48643
cor=−0.11, p=0.57
795: NBL1|P41271
cor=0.63, p=0.00025
287: IQGAP2|Q13576
cor=−0.062, p=0.72
134: MERTK|Q12866
cor=0.29, p=0.086
73: DEFA1|P59665
cor=0.75, p=0.0032
1092: YES1|P07947
cor=0.42, p=0.052
839: SCARF2|Q96GP6
cor=−0.1, p=0.66
1104: PON2|Q15165
cor=0.77, p=0.0013
208: SELE|P16581
cor=0.5, p=0.0057
446: CD209|Q9NNX6

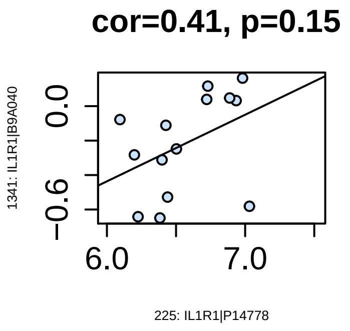
cor=0.41, p=0.15
0.0
−0.6
6.0
7.0
1341: IL1R1|B9A040
225: IL1R1|P14778

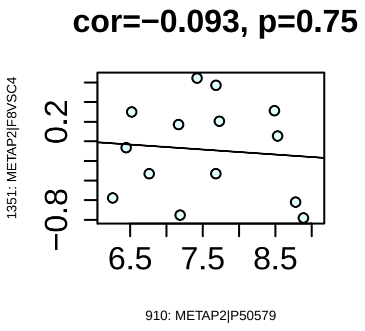
cor=−0.093, p=0.75
0.2
−0.8
6.5
7.5
8.5
1351: METAP2|F8VSC4
910: METAP2|P50579

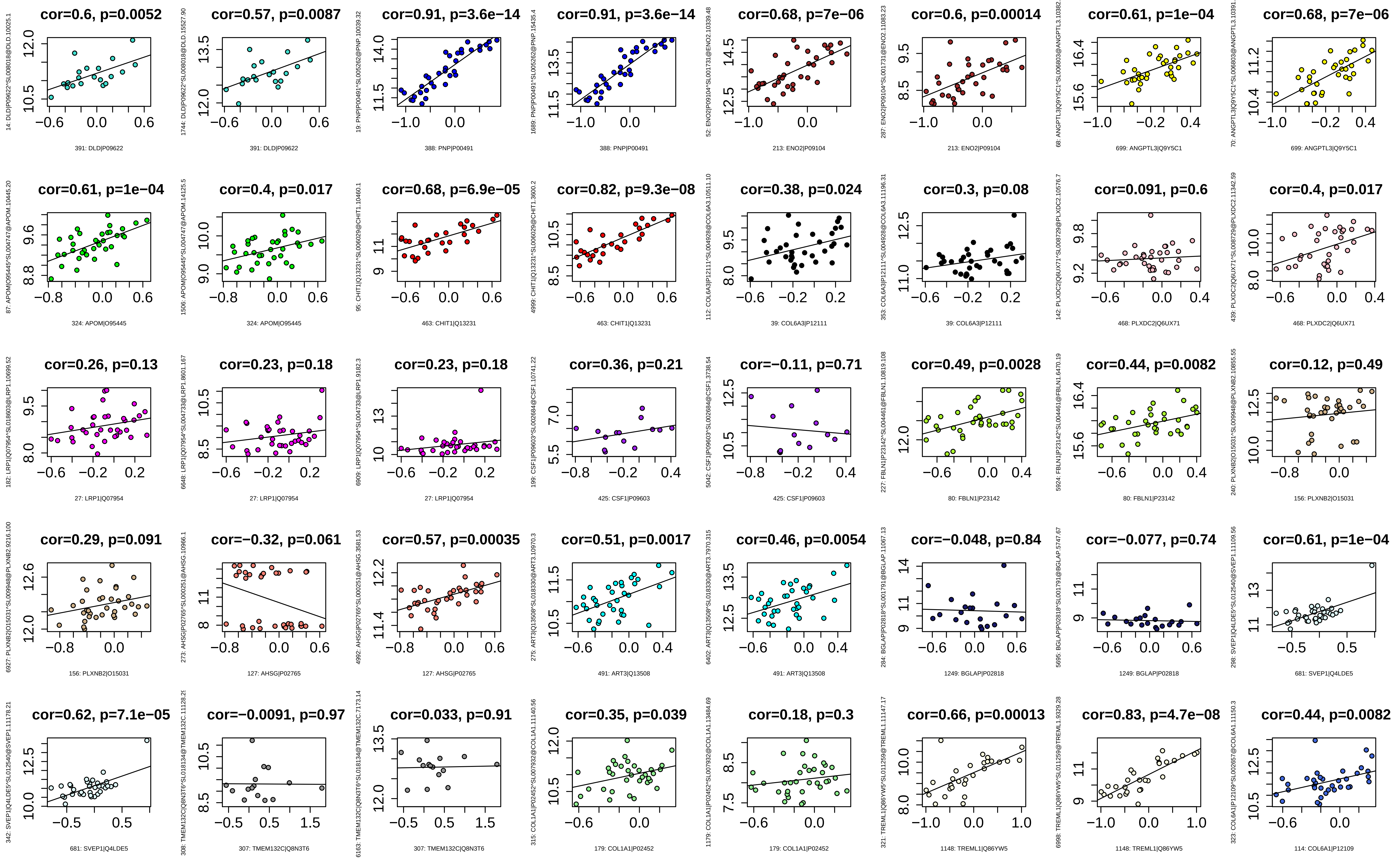

cor=0.6, p=0.0052
391: DLD|P09622
cor=0.57, p=0.0087
391: DLD|P09622
cor=0.91, p=3.6e−14
388: PNP|P00491
cor=0.91, p=3.6e−14
388: PNP|P00491
cor=0.68, p=7e−06
213: ENO2|P09104
cor=0.6, p=0.00014
213: ENO2|P09104
cor=0.61, p=1e−04
699: ANGPTL3|Q9Y5C1
cor=0.68, p=7e−06
699: ANGPTL3|Q9Y5C1
cor=0.61, p=1e−04
324: APOM|O95445
cor=0.4, p=0.017
324: APOM|O95445
cor=0.68, p=6.9e−05
463: CHIT1|Q13231
cor=0.82, p=9.3e−08
463: CHIT1|Q13231
cor=0.38, p=0.024
39: COL6A3|P12111
cor=0.3, p=0.08
39: COL6A3|P12111
cor=0.091, p=0.6
468: PLXDC2|Q6UX71
cor=0.4, p=0.017
468: PLXDC2|Q6UX71
cor=0.26, p=0.13
27: LRP1|Q07954
cor=0.23, p=0.18
27: LRP1|Q07954
cor=0.23, p=0.18
27: LRP1|Q07954
cor=0.36, p=0.21
425: CSF1|P09603
cor=−0.11, p=0.71
425: CSF1|P09603
cor=0.49, p=0.0028
80: FBLN1|P23142
cor=0.44, p=0.0082
80: FBLN1|P23142
cor=0.12, p=0.49
156: PLXNB2|O15031
cor=0.29, p=0.091
156: PLXNB2|O15031
cor=−0.32, p=0.061
127: AHSG|P02765
cor=0.57, p=0.00035
127: AHSG|P02765
cor=0.51, p=0.0017
491: ART3|Q13508
cor=0.46, p=0.0054
491: ART3|Q13508
cor=−0.048, p=0.84
1249: BGLAP|P02818
cor=−0.077, p=0.74
1249: BGLAP|P02818
cor=0.61, p=1e−04
681: SVEP1|Q4LDE5
cor=0.62, p=7.1e−05
681: SVEP1|Q4LDE5
cor=−0.0091, p=0.97
307: TMEM132C|Q8N3T6
cor=0.033, p=0.91
307: TMEM132C|Q8N3T6
cor=0.35, p=0.039
179: COL1A1|P02452
cor=0.18, p=0.3
179: COL1A1|P02452
cor=0.66, p=0.00013
1148: TREML1|Q86YW5
cor=0.83, p=4.7e−08
1148: TREML1|Q86YW5
cor=0.44, p=0.0082
114: COL6A1|P12109

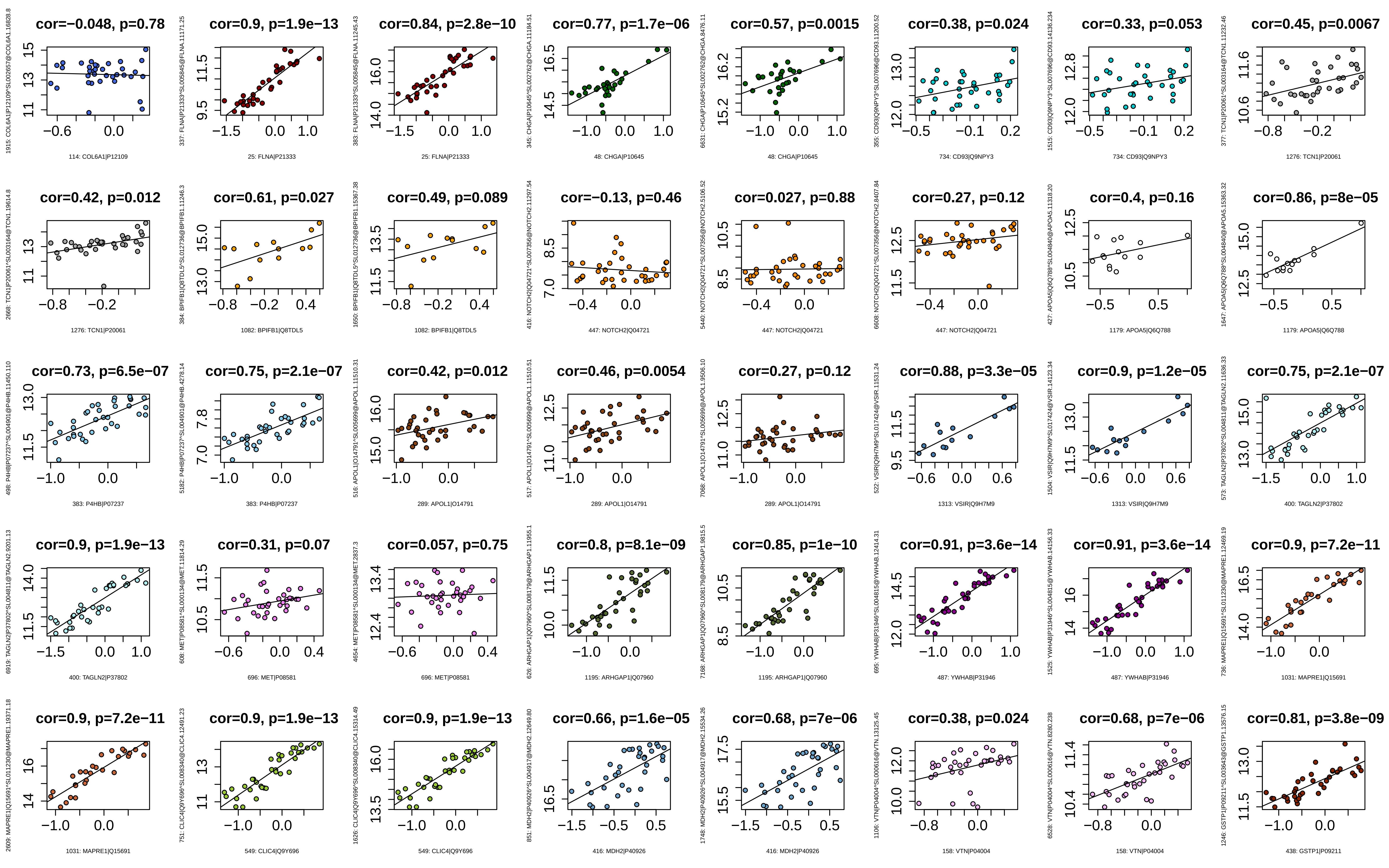
cor=−0.048, p=0.78
114: COL6A1|P12109
cor=0.9, p=1.9e−13
25: FLNA|P21333
cor=0.84, p=2.8e−10
25: FLNA|P21333
cor=0.77, p=1.7e−06
48: CHGA|P10645
cor=0.57, p=0.0015
48: CHGA|P10645
cor=0.38, p=0.024
734: CD93|Q9NPY3
cor=0.33, p=0.053
734: CD93|Q9NPY3
cor=0.45, p=0.0067
1276: TCN1|P20061
cor=0.42, p=0.012
1276: TCN1|P20061
cor=0.61, p=0.027
1082: BPIFB1|Q8TDL5
cor=0.49, p=0.089
1082: BPIFB1|Q8TDL5
cor=−0.13, p=0.46
447: NOTCH2|Q04721
cor=0.027, p=0.88
447: NOTCH2|Q04721
cor=0.27, p=0.12
447: NOTCH2|Q04721
cor=0.4, p=0.16
1179: APOA5|Q6Q788
cor=0.86, p=8e−05
1179: APOA5|Q6Q788
cor=0.73, p=6.5e−07
383: P4HB|P07237
cor=0.75, p=2.1e−07
383: P4HB|P07237
cor=0.42, p=0.012
289: APOL1|O14791
cor=0.46, p=0.0054
289: APOL1|O14791
cor=0.27, p=0.12
289: APOL1|O14791
cor=0.88, p=3.3e−05
1313: VSIR|Q9H7M9
cor=0.9, p=1.2e−05
1313: VSIR|Q9H7M9
cor=0.75, p=2.1e−07
400: TAGLN2|P37802
cor=0.9, p=1.9e−13
400: TAGLN2|P37802
cor=0.31, p=0.07
696: MET|P08581
cor=0.057, p=0.75
696: MET|P08581
cor=0.8, p=8.1e−09
1195: ARHGAP1|Q07960
cor=0.85, p=1e−10
1195: ARHGAP1|Q07960
cor=0.91, p=3.6e−14
487: YWHAB|P31946
cor=0.91, p=3.6e−14
487: YWHAB|P31946
cor=0.9, p=7.2e−11
1031: MAPRE1|Q15691
cor=0.9, p=7.2e−11
1031: MAPRE1|Q15691
cor=0.9, p=1.9e−13
549: CLIC4|Q9Y696
cor=0.9, p=1.9e−13
549: CLIC4|Q9Y696
cor=0.66, p=1.6e−05
416: MDH2|P40926
cor=0.68, p=7e−06
416: MDH2|P40926
cor=0.38, p=0.024
158: VTN|P04004
cor=0.68, p=7e−06
158: VTN|P04004
cor=0.81, p=3.8e−09
438: GSTP1|P09211

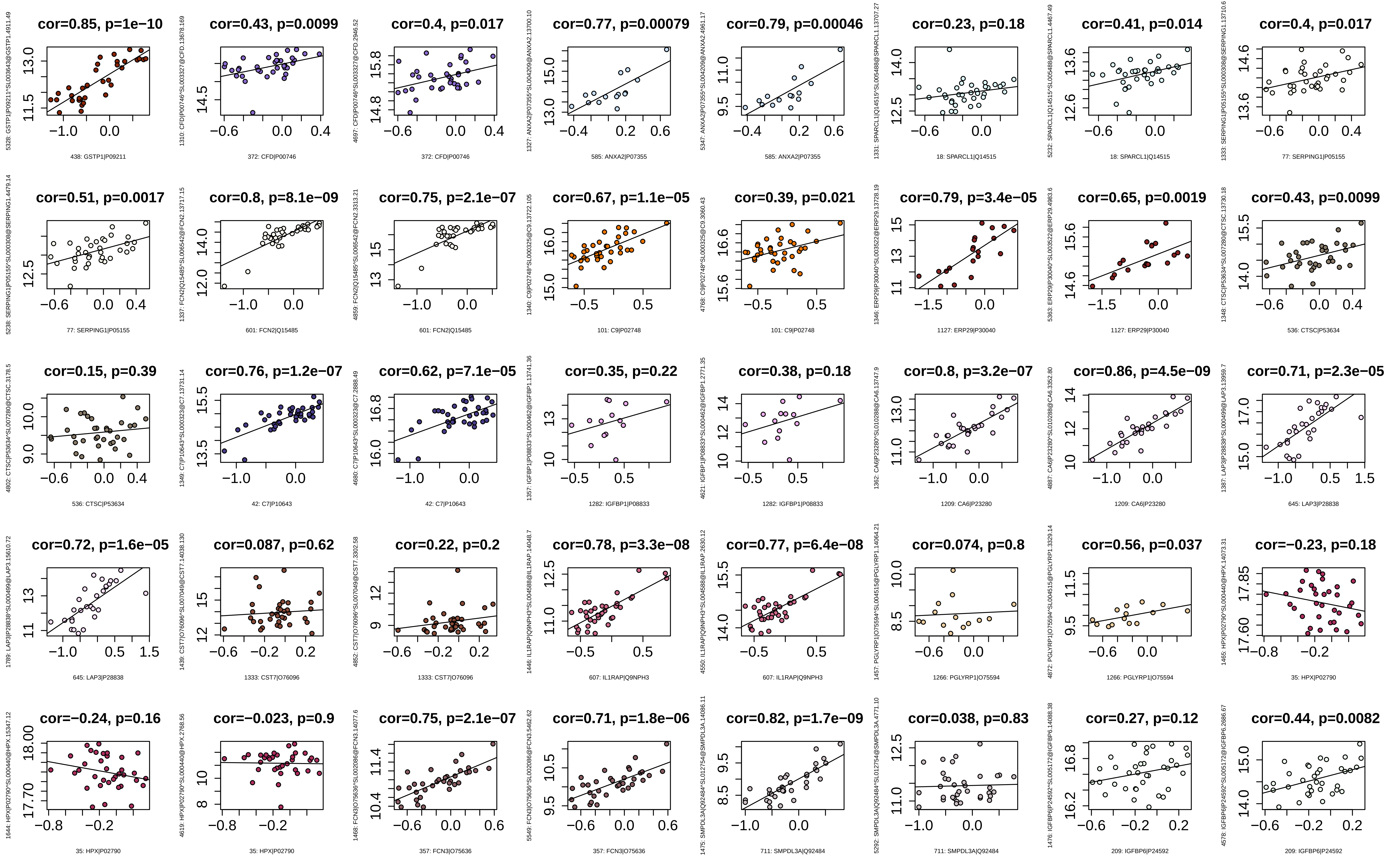

cor=0.85, p=1e−10
438: GSTP1|P09211
cor=0.43, p=0.0099
372: CFD|P00746
cor=0.4, p=0.017
372: CFD|P00746
cor=0.77, p=0.00079
585: ANXA2|P07355
cor=0.79, p=0.00046
585: ANXA2|P07355
cor=0.23, p=0.18
18: SPARCL1|Q14515
cor=0.41, p=0.014
18: SPARCL1|Q14515
cor=0.4, p=0.017
77: SERPING1|P05155
cor=0.51, p=0.0017
77: SERPING1|P05155
cor=0.8, p=8.1e−09
601: FCN2|Q15485
cor=0.75, p=2.1e−07
601: FCN2|Q15485
cor=0.67, p=1.1e−05
101: C9|P02748
cor=0.39, p=0.021
101: C9|P02748
cor=0.79, p=3.4e−05
1127: ERP29|P30040
cor=0.65, p=0.0019
1127: ERP29|P30040
cor=0.43, p=0.0099
536: CTSC|P53634
cor=0.15, p=0.39
536: CTSC|P53634
cor=0.76, p=1.2e−07
42: C7|P10643
cor=0.62, p=7.1e−05
42: C7|P10643
cor=0.35, p=0.22
1282: IGFBP1|P08833
cor=0.38, p=0.18
1282: IGFBP1|P08833
cor=0.8, p=3.2e−07
1209: CA6|P23280
cor=0.86, p=4.5e−09
1209: CA6|P23280
cor=0.71, p=2.3e−05
645: LAP3|P28838
cor=0.72, p=1.6e−05
645: LAP3|P28838
cor=0.087, p=0.62
1333: CST7|O76096
cor=0.22, p=0.2
1333: CST7|O76096
cor=0.78, p=3.3e−08
607: IL1RAP|Q9NPH3
cor=0.77, p=6.4e−08
607: IL1RAP|Q9NPH3
cor=0.074, p=0.8
1266: PGLYRP1|O75594
cor=0.56, p=0.037
1266: PGLYRP1|O75594
cor=−0.23, p=0.18
35: HPX|P02790
cor=−0.24, p=0.16
35: HPX|P02790
cor=−0.023, p=0.9
35: HPX|P02790
cor=0.75, p=2.1e−07
357: FCN3|O75636
cor=0.71, p=1.8e−06
357: FCN3|O75636
cor=0.82, p=1.7e−09
711: SMPDL3A|Q92484
cor=0.038, p=0.83
711: SMPDL3A|Q92484
cor=0.27, p=0.12
209: IGFBP6|P24592
cor=0.44, p=0.0082
209: IGFBP6|P24592

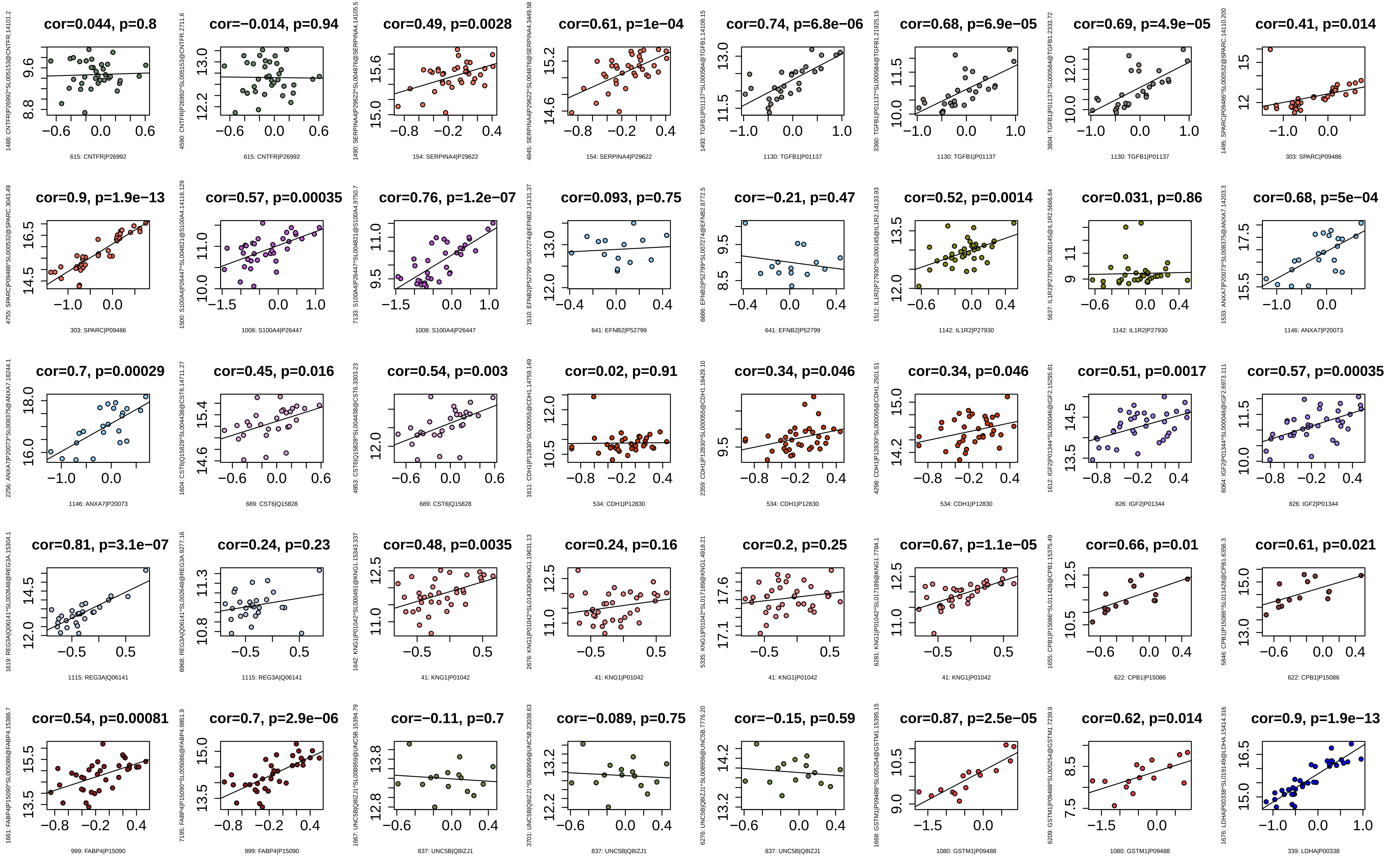
cor=0.044, p=0.8
615: CNTFR|P26992
cor=−0.014, p=0.94
615: CNTFR|P26992
cor=0.49, p=0.0028
154: SERPINA4|P29622
cor=0.61, p=1e−04
154: SERPINA4|P29622
cor=0.74, p=6.8e−06
1130: TGFB1|P01137
cor=0.68, p=6.9e−05
1130: TGFB1|P01137
cor=0.69, p=4.9e−05
1130: TGFB1|P01137
cor=0.41, p=0.014
303: SPARC|P09486
cor=0.9, p=1.9e−13
303: SPARC|P09486
cor=0.57, p=0.00035
1008: S100A4|P26447
cor=0.76, p=1.2e−07
1008: S100A4|P26447
cor=0.093, p=0.75
641: EFNB2|P52799
cor=−0.21, p=0.47
641: EFNB2|P52799
cor=0.52, p=0.0014
1142: IL1R2|P27930
cor=0.031, p=0.86
1142: IL1R2|P27930
cor=0.68, p=5e−04
1146: ANXA7|P20073
cor=0.7, p=0.00029
1146: ANXA7|P20073
cor=0.45, p=0.016
689: CST6|Q15828
cor=0.54, p=0.003
689: CST6|Q15828
cor=0.02, p=0.91
534: CDH1|P12830
cor=0.34, p=0.046
534: CDH1|P12830
cor=0.34, p=0.046
534: CDH1|P12830
cor=0.51, p=0.0017
826: IGF2|P01344
cor=0.57, p=0.00035
826: IGF2|P01344
cor=0.81, p=3.1e−07
1115: REG3A|Q06141
cor=0.24, p=0.23
1115: REG3A|Q06141
cor=0.48, p=0.0035
41: KNG1|P01042
cor=0.24, p=0.16
41: KNG1|P01042
cor=0.2, p=0.25
41: KNG1|P01042
cor=0.67, p=1.1e−05
41: KNG1|P01042
cor=0.66, p=0.01
622: CPB1|P15086
cor=0.61, p=0.021
622: CPB1|P15086
cor=0.54, p=0.00081
999: FABP4|P15090
cor=0.7, p=2.9e−06
999: FABP4|P15090
cor=−0.11, p=0.7
837: UNC5B|Q8IZJ1
cor=−0.089, p=0.75
837: UNC5B|Q8IZJ1
cor=−0.15, p=0.59
837: UNC5B|Q8IZJ1
cor=0.87, p=2.5e−05
1080: GSTM1|P09488
cor=0.62, p=0.014
1080: GSTM1|P09488
cor=0.9, p=1.9e−13
339: LDHA|P00338

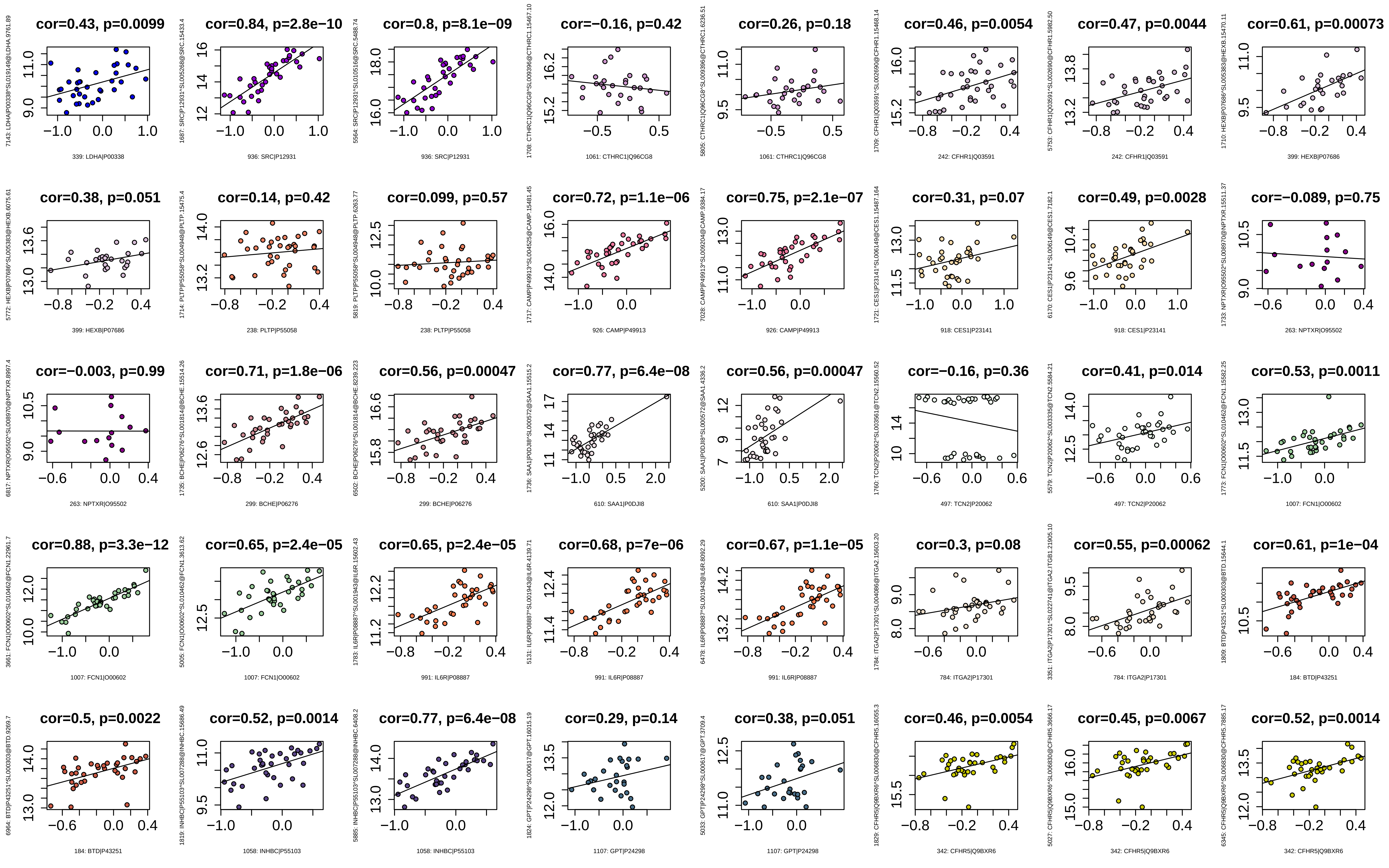
cor=0.43, p=0.0099
339: LDHA|P00338
cor=0.84, p=2.8e−10
936: SRC|P12931
cor=0.8, p=8.1e−09
936: SRC|P12931
cor=−0.16, p=0.42
1061: CTHRC1|Q96CG8
cor=0.26, p=0.18
1061: CTHRC1|Q96CG8
cor=0.46, p=0.0054
242: CFHR1|Q03591
cor=0.47, p=0.0044
242: CFHR1|Q03591
cor=0.61, p=0.00073
399: HEXB|P07686
cor=0.38, p=0.051
399: HEXB|P07686
cor=0.14, p=0.42
238: PLTP|P55058
cor=0.099, p=0.57
238: PLTP|P55058
cor=0.72, p=1.1e−06
926: CAMP|P49913
cor=0.75, p=2.1e−07
926: CAMP|P49913
cor=0.31, p=0.07
918: CES1|P23141
cor=0.49, p=0.0028
918: CES1|P23141
cor=−0.089, p=0.75
263: NPTXR|O95502
cor=−0.003, p=0.99
263: NPTXR|O95502
cor=0.71, p=1.8e−06
299: BCHE|P06276
cor=0.56, p=0.00047
299: BCHE|P06276
cor=0.77, p=6.4e−08
610: SAA1|P0DJI8
cor=0.56, p=0.00047
610: SAA1|P0DJI8
cor=−0.16, p=0.36
497: TCN2|P20062
cor=0.41, p=0.014
497: TCN2|P20062
cor=0.53, p=0.0011
1007: FCN1|O00602
cor=0.88, p=3.3e−12
1007: FCN1|O00602
cor=0.65, p=2.4e−05
1007: FCN1|O00602
cor=0.65, p=2.4e−05
991: IL6R|P08887
cor=0.68, p=7e−06
991: IL6R|P08887
cor=0.67, p=1.1e−05
991: IL6R|P08887
cor=0.3, p=0.08
784: ITGA2|P17301
cor=0.55, p=0.00062
784: ITGA2|P17301
cor=0.61, p=1e−04
184: BTD|P43251
cor=0.5, p=0.0022
184: BTD|P43251
cor=0.52, p=0.0014
1058: INHBC|P55103
cor=0.77, p=6.4e−08
1058: INHBC|P55103
cor=0.29, p=0.14
1107: GPT|P24298
cor=0.38, p=0.051
1107: GPT|P24298
cor=0.46, p=0.0054
342: CFHR5|Q9BXR6
cor=0.45, p=0.0067
342: CFHR5|Q9BXR6
cor=0.52, p=0.0014
342: CFHR5|Q9BXR6

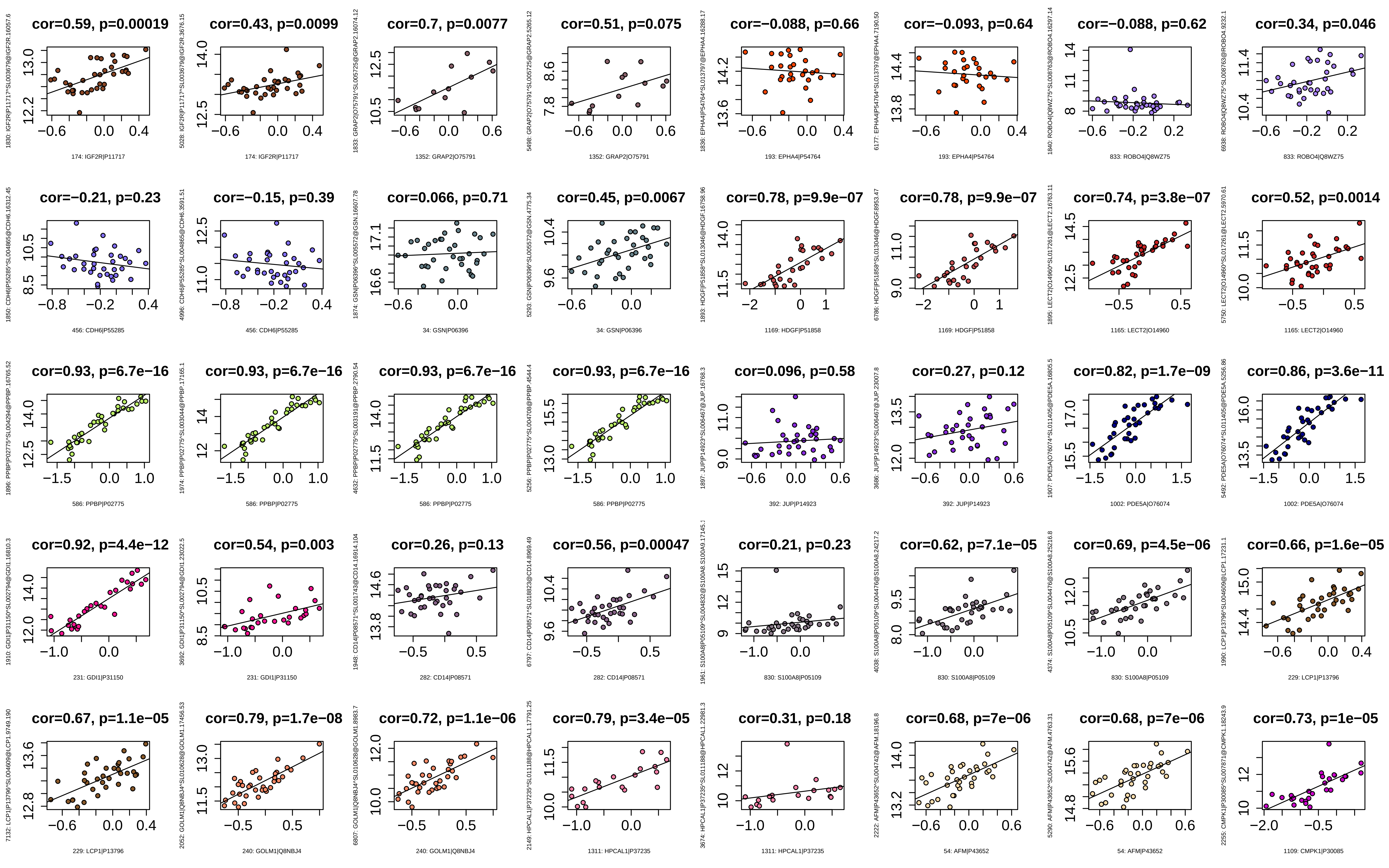
cor=0.59, p=0.00019
174: IGF2R|P11717
cor=0.43, p=0.0099
174: IGF2R|P11717
cor=0.7, p=0.0077
1352: GRAP2|O75791
cor=0.51, p=0.075
1352: GRAP2|O75791
cor=−0.088, p=0.66
193: EPHA4|P54764
cor=−0.093, p=0.64
193: EPHA4|P54764
cor=−0.088, p=0.62
833: ROBO4|Q8WZ75
cor=0.34, p=0.046
833: ROBO4|Q8WZ75
cor=−0.21, p=0.23
456: CDH6|P55285
cor=−0.15, p=0.39
456: CDH6|P55285
cor=0.066, p=0.71
34: GSN|P06396
cor=0.45, p=0.0067
34: GSN|P06396
cor=0.78, p=9.9e−07
1169: HDGF|P51858
cor=0.78, p=9.9e−07
1169: HDGF|P51858
cor=0.74, p=3.8e−07
1165: LECT2|O14960
cor=0.52, p=0.0014
1165: LECT2|O14960
cor=0.93, p=6.7e−16
586: PPBP|P02775
cor=0.93, p=6.7e−16
586: PPBP|P02775
cor=0.93, p=6.7e−16
586: PPBP|P02775
cor=0.93, p=6.7e−16
586: PPBP|P02775
cor=0.096, p=0.58
392: JUP|P14923
cor=0.27, p=0.12
392: JUP|P14923
cor=0.82, p=1.7e−09
1002: PDE5A|O76074
cor=0.86, p=3.6e−11
1002: PDE5A|O76074
cor=0.92, p=4.4e−12
231: GDI1|P31150
cor=0.54, p=0.003
231: GDI1|P31150
cor=0.26, p=0.13
282: CD14|P08571
cor=0.56, p=0.00047
282: CD14|P08571
cor=0.21, p=0.23
830: S100A8|P05109
cor=0.62, p=7.1e−05
830: S100A8|P05109
cor=0.69, p=4.5e−06
830: S100A8|P05109
cor=0.66, p=1.6e−05
229: LCP1|P13796
cor=0.67, p=1.1e−05
229: LCP1|P13796
cor=0.79, p=1.7e−08
240: GOLM1|Q8NBJ4
cor=0.72, p=1.1e−06
240: GOLM1|Q8NBJ4
cor=0.79, p=3.4e−05
1311: HPCAL1|P37235
cor=0.31, p=0.18
1311: HPCAL1|P37235
cor=0.68, p=7e−06
54: AFM|P43652
cor=0.68, p=7e−06
54: AFM|P43652
cor=0.73, p=1e−05
1109: CMPK1|P30085

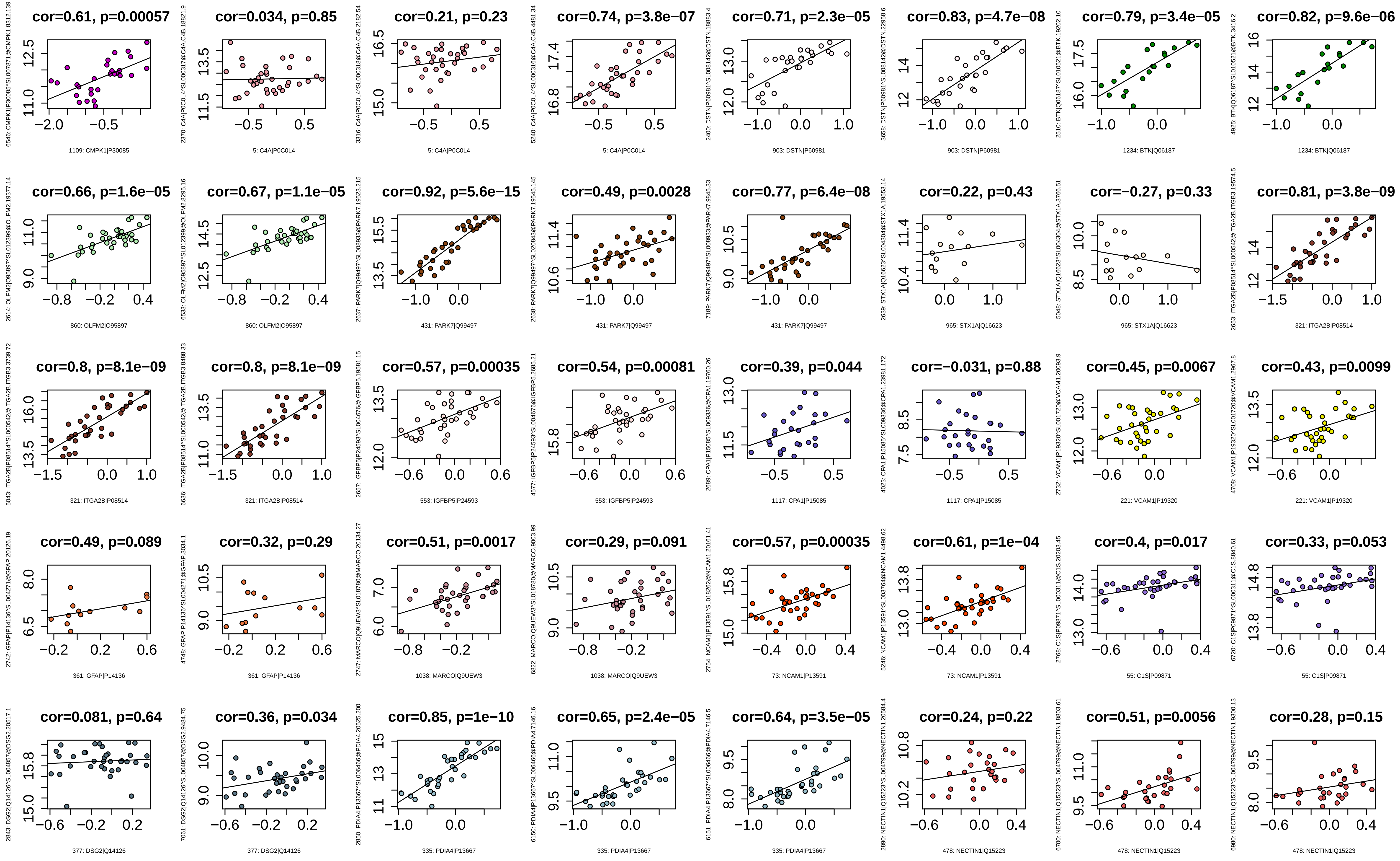
cor=0.61, p=0.00057
1109: CMPK1|P30085
cor=0.034, p=0.85
5: C4A|P0C0L4
cor=0.21, p=0.23
5: C4A|P0C0L4
cor=0.74, p=3.8e−07
5: C4A|P0C0L4
cor=0.71, p=2.3e−05
903: DSTN|P60981
cor=0.83, p=4.7e−08
903: DSTN|P60981
cor=0.79, p=3.4e−05
1234: BTK|Q06187
cor=0.82, p=9.6e−06
1234: BTK|Q06187
cor=0.66, p=1.6e−05
860: OLFM2|O95897
cor=0.67, p=1.1e−05
860: OLFM2|O95897
cor=0.92, p=5.6e−15
431: PARK7|Q99497
cor=0.49, p=0.0028
431: PARK7|Q99497
cor=0.77, p=6.4e−08
431: PARK7|Q99497
cor=0.22, p=0.43
965: STX1A|Q16623
cor=−0.27, p=0.33
965: STX1A|Q16623
cor=0.81, p=3.8e−09
321: ITGA2B|P08514
cor=0.8, p=8.1e−09
321: ITGA2B|P08514
cor=0.8, p=8.1e−09
321: ITGA2B|P08514
cor=0.57, p=0.00035
553: IGFBP5|P24593
cor=0.54, p=0.00081
553: IGFBP5|P24593
cor=0.39, p=0.044
1117: CPA1|P15085
cor=−0.031, p=0.88
1117: CPA1|P15085
cor=0.45, p=0.0067
221: VCAM1|P19320
cor=0.43, p=0.0099
221: VCAM1|P19320
cor=0.49, p=0.089
361: GFAP|P14136
cor=0.32, p=0.29
361: GFAP|P14136
cor=0.51, p=0.0017
1038: MARCO|Q9UEW3
cor=0.29, p=0.091
1038: MARCO|Q9UEW3
cor=0.57, p=0.00035
73: NCAM1|P13591
cor=0.61, p=1e−04
73: NCAM1|P13591
cor=0.4, p=0.017
55: C1S|P09871
cor=0.33, p=0.053
55: C1S|P09871
cor=0.081, p=0.64
377: DSG2|Q14126
cor=0.36, p=0.034
377: DSG2|Q14126
cor=0.85, p=1e−10
335: PDIA4|P13667
cor=0.65, p=2.4e−05
335: PDIA4|P13667
cor=0.64, p=3.5e−05
335: PDIA4|P13667
cor=0.24, p=0.22
478: NECTIN1|Q15223
cor=0.51, p=0.0056
478: NECTIN1|Q15223
cor=0.28, p=0.15
478: NECTIN1|Q15223

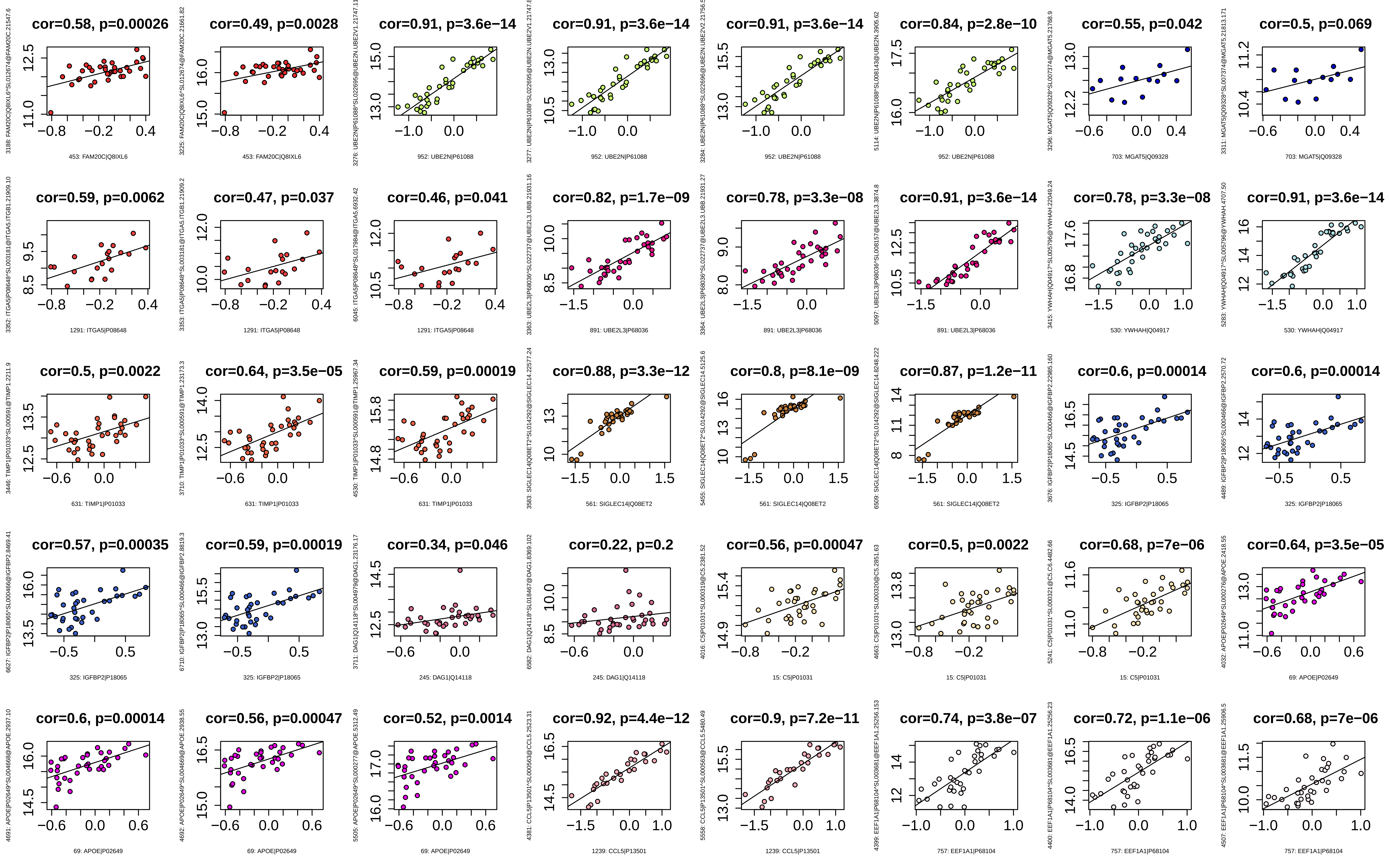
cor=0.58, p=0.00026
453: FAM20C|Q8IXL6
cor=0.49, p=0.0028
453: FAM20C|Q8IXL6
cor=0.91, p=3.6e−14
952: UBE2N|P61088
cor=0.91, p=3.6e−14
952: UBE2N|P61088
cor=0.91, p=3.6e−14
952: UBE2N|P61088
cor=0.84, p=2.8e−10
952: UBE2N|P61088
cor=0.55, p=0.042
703: MGAT5|Q09328
cor=0.5, p=0.069
703: MGAT5|Q09328
cor=0.59, p=0.0062
1291: ITGA5|P08648
cor=0.47, p=0.037
1291: ITGA5|P08648
cor=0.46, p=0.041
1291: ITGA5|P08648
cor=0.82, p=1.7e−09
891: UBE2L3|P68036
cor=0.78, p=3.3e−08
891: UBE2L3|P68036
cor=0.91, p=3.6e−14
891: UBE2L3|P68036
cor=0.78, p=3.3e−08
530: YWHAH|Q04917
cor=0.91, p=3.6e−14
530: YWHAH|Q04917
cor=0.5, p=0.0022
631: TIMP1|P01033
cor=0.64, p=3.5e−05
631: TIMP1|P01033
cor=0.59, p=0.00019
631: TIMP1|P01033
cor=0.88, p=3.3e−12
561: SIGLEC14|Q08ET2
cor=0.8, p=8.1e−09
561: SIGLEC14|Q08ET2
cor=0.87, p=1.2e−11
561: SIGLEC14|Q08ET2
cor=0.6, p=0.00014
325: IGFBP2|P18065
cor=0.6, p=0.00014
325: IGFBP2|P18065
cor=0.57, p=0.00035
325: IGFBP2|P18065
cor=0.59, p=0.00019
325: IGFBP2|P18065
cor=0.34, p=0.046
245: DAG1|Q14118
cor=0.22, p=0.2
245: DAG1|Q14118
cor=0.56, p=0.00047
15: C5|P01031
cor=0.5, p=0.0022
15: C5|P01031
cor=0.68, p=7e−06
15: C5|P01031
cor=0.64, p=3.5e−05
69: APOE|P02649
cor=0.6, p=0.00014
69: APOE|P02649
cor=0.56, p=0.00047
69: APOE|P02649
cor=0.52, p=0.0014
69: APOE|P02649
cor=0.92, p=4.4e−12
1239: CCL5|P13501
cor=0.9, p=7.2e−11
1239: CCL5|P13501
cor=0.74, p=3.8e−07
757: EEF1A1|P68104
cor=0.72, p=1.1e−06
757: EEF1A1|P68104
cor=0.68, p=7e−06
757: EEF1A1|P68104

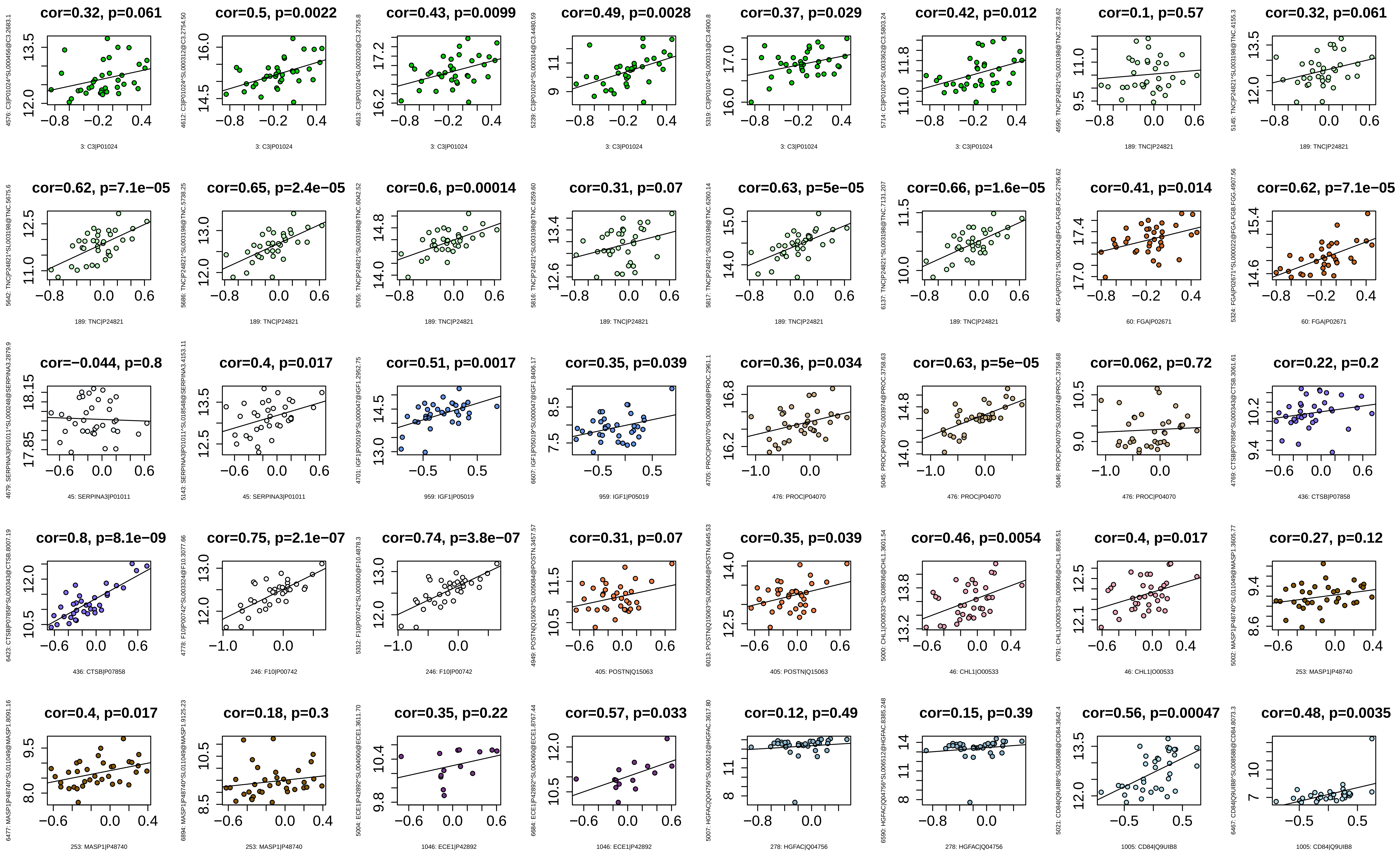

cor=0.32, p=0.061
4576: C3|P01024^SL000456@C3.2683.1
3: C3|P01024
cor=0.5, p=0.0022
4612: C3|P01024^SL000312@C3.2754.50
3: C3|P01024
cor=0.43, p=0.0099
4613: C3|P01024^SL003220@C3.2755.8
3: C3|P01024
cor=0.49, p=0.0028
5239: C3|P01024^SL000314@C3.4480.59
3: C3|P01024
cor=0.37, p=0.029
5319: C3|P01024^SL000313@C3.4900.8
3: C3|P01024
cor=0.42, p=0.012
5714: C3|P01024^SL003362@C3.5803.24
3: C3|P01024
cor=0.1, p=0.57
4595: TNC|P24821^SL003198@TNC.2728.62
189: TNC|P24821
cor=0.32, p=0.061
5145: TNC|P24821^SL003198@TNC.4155.3
189: TNC|P24821
cor=0.62, p=7.1e−05
5642: TNC|P24821^SL003198@TNC.5675.6
189: TNC|P24821
cor=0.65, p=2.4e−05
5686: TNC|P24821^SL003198@TNC.5738.25
189: TNC|P24821
cor=0.6, p=0.00014
5765: TNC|P24821^SL003198@TNC.6042.52
189: TNC|P24821
cor=0.31, p=0.07
5816: TNC|P24821^SL003198@TNC.6259.60
189: TNC|P24821
cor=0.63, p=5e−05
5817: TNC|P24821^SL003198@TNC.6260.14
189: TNC|P24821
cor=0.66, p=1.6e−05
6137: TNC|P24821^SL003198@TNC.7131.207
189: TNC|P24821
cor=0.41, p=0.014
4634: FGA|P02671^.2796.62
60: FGA|P02671
cor=0.62, p=7.1e−05
5324: FGA|P02671^.4907.56
60: FGA|P02671
cor=−0.044, p=0.8
4679: SERPINA3|P01011^SL000248@SERPINA3.2879.9
45: SERPINA3|P01011
cor=0.4, p=0.017
5143: SERPINA3|P01011^SL018548@SERPINA3.4153.11
45: SERPINA3|P01011
cor=0.51, p=0.0017
4701: IGF1|P05019^SL000047@IGF1.2952.75
959: IGF1|P05019
cor=0.35, p=0.039
6607: IGF1|P05019^SL000047@IGF1.8406.17
959: IGF1|P05019
cor=0.36, p=0.034
4705: PROC|P04070^SL000048@PROC.2961.1
476: PROC|P04070
cor=0.63, p=5e−05
5045: PROC|P04070^SL003974@PROC.3758.63
476: PROC|P04070
cor=0.062, p=0.72
5046: PROC|P04070^SL003974@PROC.3758.68
476: PROC|P04070
cor=0.22, p=0.2
4769: CTSB|P07858^SL000343@CTSB.3061.61
436: CTSB|P07858
cor=0.8, p=8.1e−09
6423: CTSB|P07858^SL000343@CTSB.8007.19
436: CTSB|P07858
cor=0.75, p=2.1e−07
4778: F10|P00742^SL003324@F10.3077.66
246: F10|P00742
cor=0.74, p=3.8e−07
5312: F10|P00742^SL000360@F10.4878.3
246: F10|P00742
cor=0.31, p=0.07
4949: POSTN|Q15063^SL005084@POSTN.3457.57
405: POSTN|Q15063
cor=0.35, p=0.039
6013: POSTN|Q15063^SL005084@POSTN.6645.53
405: POSTN|Q15063
cor=0.46, p=0.0054
5000: CHL1|O00533^SL008936@CHL1.3601.54
46: CHL1|O00533
cor=0.4, p=0.017
6791: CHL1|O00533^SL008936@CHL1.8958.51
46: CHL1|O00533
cor=0.27, p=0.12
5002: MASP1|P48740^SL011049@MASP1.3605.77
253: MASP1|P48740
cor=0.4, p=0.017
6477: MASP1|P48740^SL011049@MASP1.8091.16
253: MASP1|P48740
cor=0.18, p=0.3
6894: MASP1|P48740^SL011049@MASP1.9125.23
253: MASP1|P48740
cor=0.35, p=0.22
5004: ECE1|P42892^SL004060@ECE1.3611.70
1046: ECE1|P42892
cor=0.57, p=0.033
6684: ECE1|P42892^SL004060@ECE1.8767.44
1046: ECE1|P42892
cor=0.12, p=0.49
5007: HGFAC|Q04756^SL006512@HGFAC.3617.80
278: HGFAC|Q04756
cor=0.15, p=0.39
6590: HGFAC|Q04756^SL006512@HGFAC.8385.248
278: HGFAC|Q04756
cor=0.56, p=0.00047
5021: CD84|Q9UIB8^SL008588@CD84.3642.4
1005: CD84|Q9UIB8
cor=0.48, p=0.0035
6467: CD84|Q9UIB8^SL008588@CD84.8073.3
1005: CD84|Q9UIB8

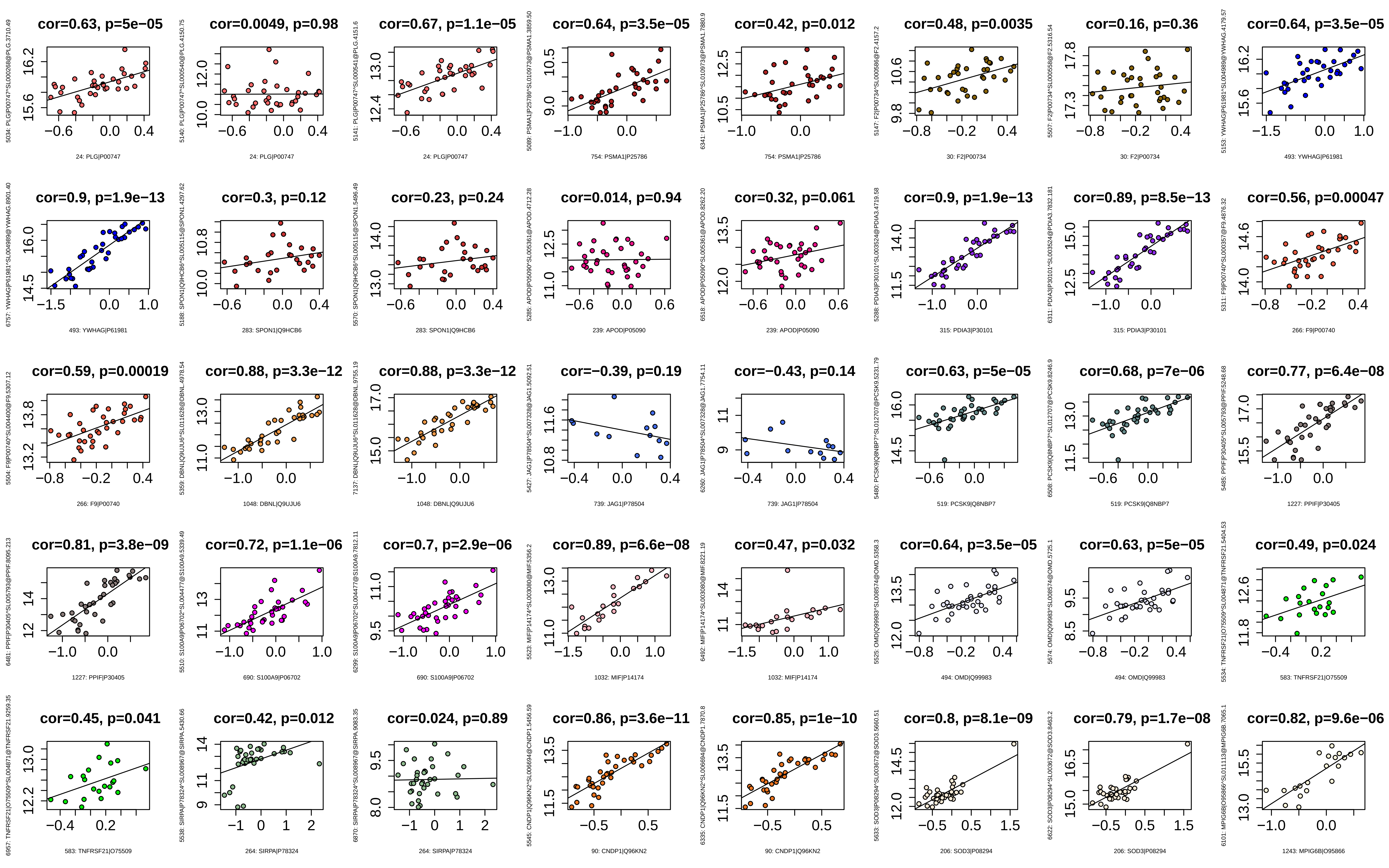
cor=0.63, p=5e−05
24: PLG|P00747
cor=0.0049, p=0.98
24: PLG|P00747
cor=0.67, p=1.1e−05
24: PLG|P00747
cor=0.64, p=3.5e−05
754: PSMA1|P25786
cor=0.42, p=0.012
754: PSMA1|P25786
cor=0.48, p=0.0035
30: F2|P00734
cor=0.16, p=0.36
30: F2|P00734
cor=0.64, p=3.5e−05
493: YWHAG|P61981
cor=0.9, p=1.9e−13
493: YWHAG|P61981
cor=0.3, p=0.12
283: SPON1|Q9HCB6
cor=0.23, p=0.24
283: SPON1|Q9HCB6
cor=0.014, p=0.94
239: APOD|P05090
cor=0.32, p=0.061
239: APOD|P05090
cor=0.9, p=1.9e−13
315: PDIA3|P30101
cor=0.89, p=8.5e−13
315: PDIA3|P30101
cor=0.56, p=0.00047
266: F9|P00740
cor=0.59, p=0.00019
266: F9|P00740
cor=0.88, p=3.3e−12
1048: DBNL|Q9UJU6
cor=0.88, p=3.3e−12
1048: DBNL|Q9UJU6
cor=−0.39, p=0.19
739: JAG1|P78504
cor=−0.43, p=0.14
739: JAG1|P78504
cor=0.63, p=5e−05
519: PCSK9|Q8NBP7
cor=0.68, p=7e−06
519: PCSK9|Q8NBP7
cor=0.77, p=6.4e−08
1227: PPIF|P30405
cor=0.81, p=3.8e−09
1227: PPIF|P30405
cor=0.72, p=1.1e−06
690: S100A9|P06702
cor=0.7, p=2.9e−06
690: S100A9|P06702
cor=0.89, p=6.6e−08
1032: MIF|P14174
cor=0.47, p=0.032
1032: MIF|P14174
cor=0.64, p=3.5e−05
494: OMD|Q99983
cor=0.63, p=5e−05
494: OMD|Q99983
cor=0.49, p=0.024
583: TNFRSF21|O75509
cor=0.45, p=0.041
583: TNFRSF21|O75509
cor=0.42, p=0.012
264: SIRPA|P78324
cor=0.024, p=0.89
264: SIRPA|P78324
cor=0.86, p=3.6e−11
90: CNDP1|Q96KN2
cor=0.85, p=1e−10
90: CNDP1|Q96KN2
cor=0.8, p=8.1e−09
206: SOD3|P08294
cor=0.79, p=1.7e−08
206: SOD3|P08294
cor=0.82, p=9.6e−06
1243: MPIG6B|O95866

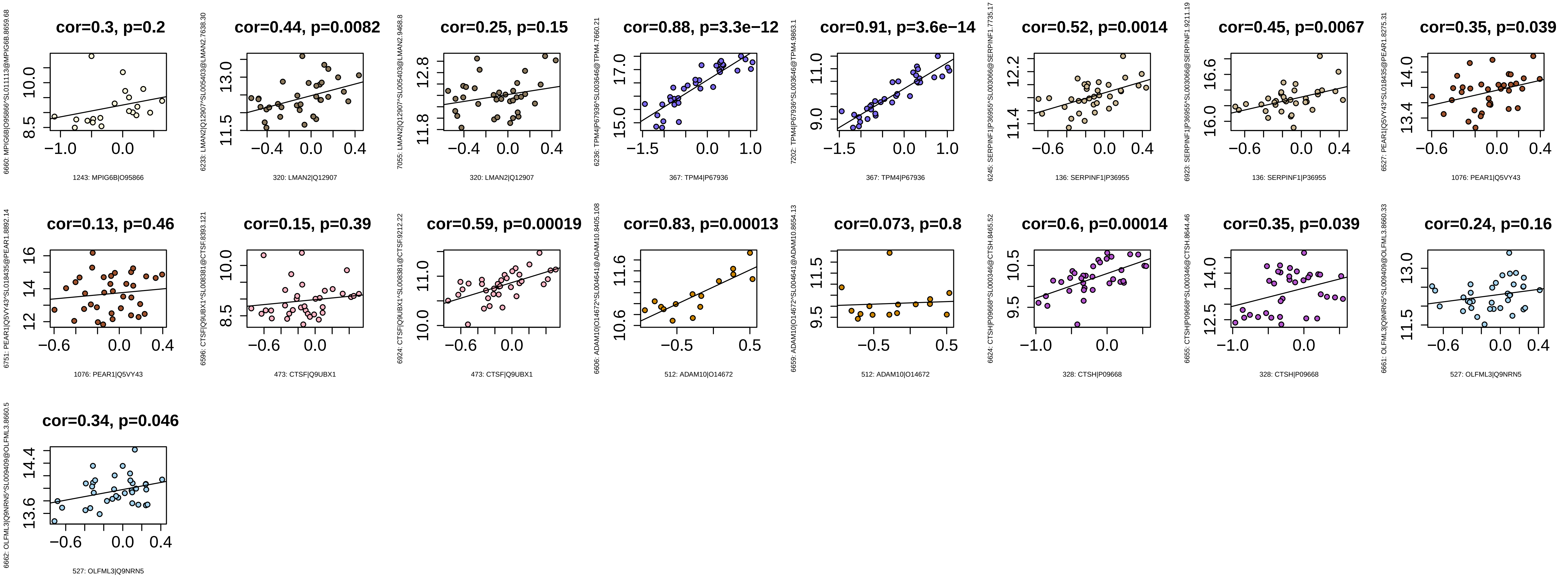
cor=0.3, p=0.2
6660: MPIG6B|O95866^SL011113@MPIG6B.8659.68
1243: MPIG6B|O95866
cor=0.44, p=0.0082
6233: LMAN2|Q12907^SL005403@LMAN2.7638.30
320: LMAN2|Q12907
cor=0.25, p=0.15
7055: LMAN2|Q12907^SL005403@LMAN2.9468.8
320: LMAN2|Q12907
cor=0.88, p=3.3e−12
6236: TPM4|P67936^SL003646@TPM4.7660.21
367: TPM4|P67936
cor=0.91, p=3.6e−14
7202: TPM4|P67936^SL003646@TPM4.9863.1
367: TPM4|P67936
cor=0.52, p=0.0014
6245: SERPINF1|P36955^SL003066@SERPINF1.7735.17
136: SERPINF1|P36955
cor=0.45, p=0.0067
6923: SERPINF1|P36955^SL003066@SERPINF1.9211.19
136: SERPINF1|P36955
cor=0.35, p=0.039
6527: PEAR1|Q5VY43^SL018435@PEAR1.8275.31
1076: PEAR1|Q5VY43
cor=0.13, p=0.46
6751: PEAR1|Q5VY43^SL018435@PEAR1.8892.14
1076: PEAR1|Q5VY43
cor=0.15, p=0.39
6596: CTSF|Q9UBX1^SL008381@CTSF.8393.121
473: CTSF|Q9UBX1
cor=0.59, p=0.00019
6924: CTSF|Q9UBX1^SL008381@CTSF.9212.22
473: CTSF|Q9UBX1
cor=0.83, p=0.00013
6606: ADAM10|O14672^SL004641@ADAM10.8405.108
512: ADAM10|O14672
cor=0.073, p=0.8
6659: ADAM10|O14672^SL004641@ADAM10.8654.13
512: ADAM10|O14672
cor=0.6, p=0.00014
6624: CTSH|P09668^SL000346@CTSH.8465.52
328: CTSH|P09668
cor=0.35, p=0.039
6655: CTSH|P09668^SL000346@CTSH.8644.46
328: CTSH|P09668
cor=0.24, p=0.16
6661: OLFML3|Q9NRN5^SL009409@OLFML3.8660.33
527: OLFML3|Q9NRN5
cor=0.34, p=0.046
6662: OLFML3|Q9NRN5^SL009409@OLFML3.8660.5
527: OLFML3|Q9NRN5

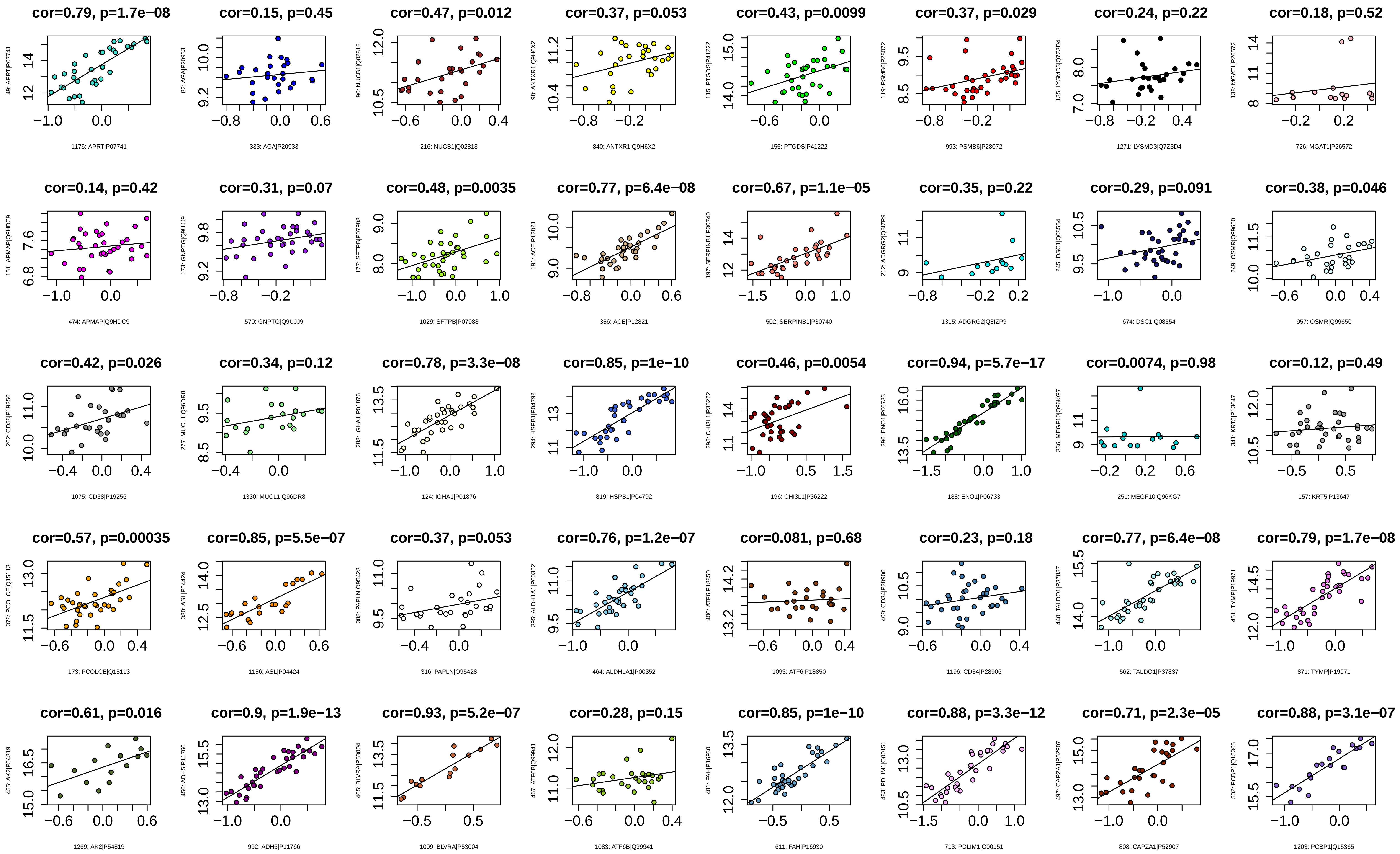
cor=0.79, p=1.7e−08
1176: APRT|P07741
cor=0.15, p=0.45
333: AGA|P20933
cor=0.47, p=0.012
216: NUCB1|Q02818
cor=0.37, p=0.053
840: ANTXR1|Q9H6X2
cor=0.43, p=0.0099
155: PTGDS|P41222
cor=0.37, p=0.029
993: PSMB6|P28072
cor=0.24, p=0.22
1271: LYSMD3|Q7Z3D4
cor=0.18, p=0.52
726: MGAT1|P26572
cor=0.14, p=0.42
474: APMAP|Q9HDC9
cor=0.31, p=0.07
570: GNPTG|Q9UJJ9
cor=0.48, p=0.0035
1029: SFTPB|P07988
cor=0.77, p=6.4e−08
356: ACE|P12821
cor=0.67, p=1.1e−05
502: SERPINB1|P30740
cor=0.35, p=0.22
1315: ADGRG2|Q8IZP9
cor=0.29, p=0.091
674: DSC1|Q08554
cor=0.38, p=0.046
957: OSMR|Q99650
cor=0.42, p=0.026
1075: CD58|P19256
cor=0.34, p=0.12
1330: MUCL1|Q96DR8
cor=0.78, p=3.3e−08
124: IGHA1|P01876
cor=0.85, p=1e−10
819: HSPB1|P04792
cor=0.46, p=0.0054
196: CHI3L1|P36222
cor=0.94, p=5.7e−17
188: ENO1|P06733
cor=0.0074, p=0.98
251: MEGF10|Q96KG7
cor=0.12, p=0.49
157: KRT5|P13647
cor=0.57, p=0.00035
173: PCOLCE|Q15113
cor=0.85, p=5.5e−07
1156: ASL|P04424
cor=0.37, p=0.053
316: PAPLN|O95428
cor=0.76, p=1.2e−07
464: ALDH1A1|P00352
cor=0.081, p=0.68
1093: ATF6|P18850
cor=0.23, p=0.18
1196: CD34|P28906
cor=0.77, p=6.4e−08
562: TALDO1|P37837
cor=0.79, p=1.7e−08
871: TYMP|P19971
cor=0.61, p=0.016
1269: AK2|P54819
cor=0.9, p=1.9e−13
992: ADH5|P11766
cor=0.93, p=5.2e−07
1009: BLVRA|P53004
cor=0.28, p=0.15
1083: ATF6B|Q99941
cor=0.85, p=1e−10
611: FAH|P16930
cor=0.88, p=3.3e−12
713: PDLIM1|O00151
cor=0.71, p=2.3e−05
808: CAPZA1|P52907
cor=0.88, p=3.1e−07
1203: PCBP1|Q15365

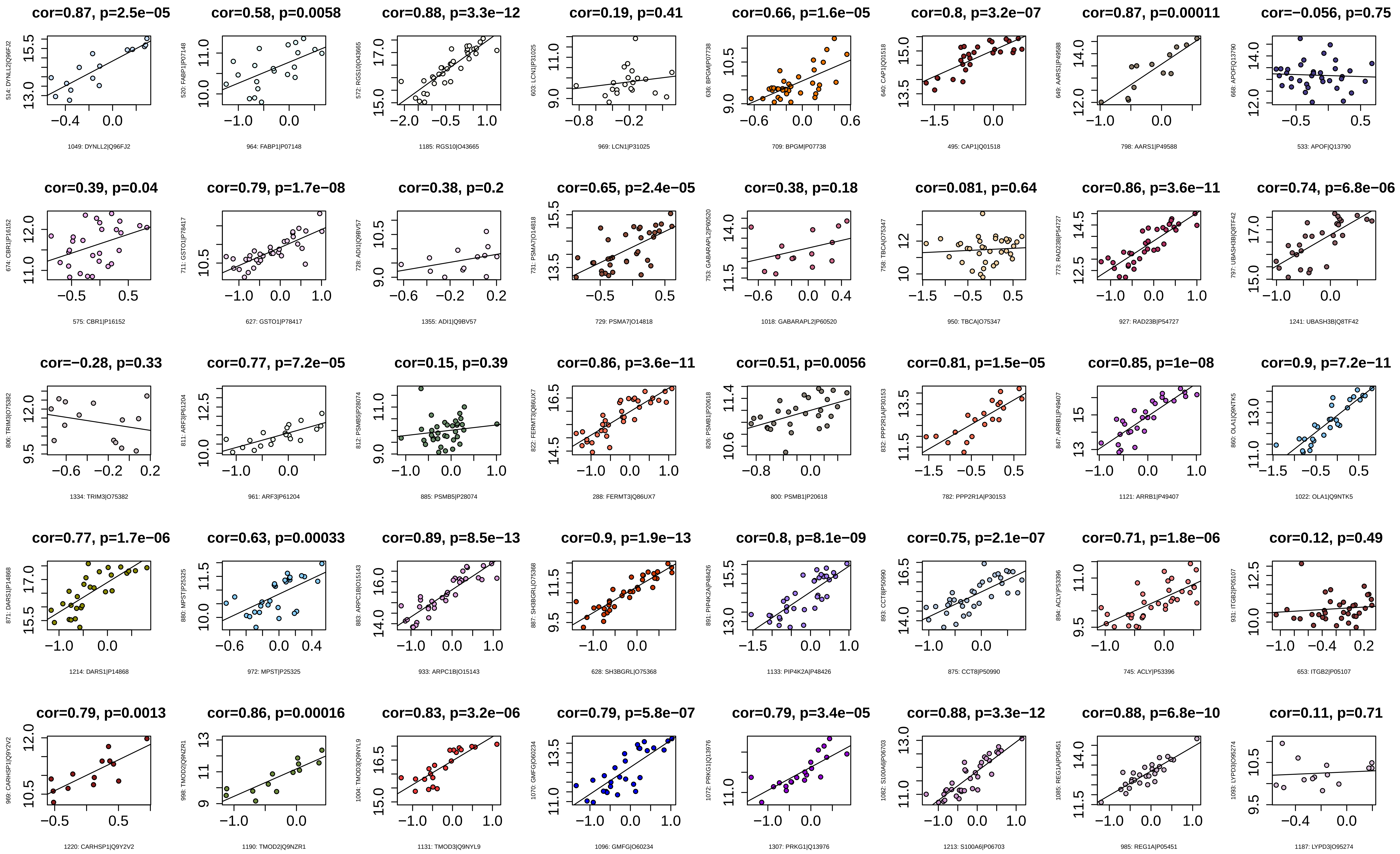
cor=0.87, p=2.5e−05
1049: DYNLL2|Q96FJ2
cor=0.58, p=0.0058
964: FABP1|P07148
cor=0.88, p=3.3e−12
1185: RGS10|O43665
cor=0.19, p=0.41
969: LCN1|P31025
cor=0.66, p=1.6e−05
709: BPGM|P07738
cor=0.8, p=3.2e−07
495: CAP1|Q01518
cor=0.87, p=0.00011
798: AARS1|P49588
cor=−0.056, p=0.75
533: APOF|Q13790
cor=0.39, p=0.04
575: CBR1|P16152
cor=0.79, p=1.7e−08
627: GSTO1|P78417
cor=0.38, p=0.2
1355: ADI1|Q9BV57
cor=0.65, p=2.4e−05
729: PSMA7|O14818
cor=0.38, p=0.18
1018: GABARAPL2|P60520
cor=0.081, p=0.64
950: TBCA|O75347
cor=0.86, p=3.6e−11
927: RAD23B|P54727
cor=0.74, p=6.8e−06
1241: UBASH3B|Q8TF42
cor=−0.28, p=0.33
1334: TRIM3|O75382
cor=0.77, p=7.2e−05
961: ARF3|P61204
cor=0.15, p=0.39
885: PSMB5|P28074
cor=0.86, p=3.6e−11
288: FERMT3|Q86UX7
cor=0.51, p=0.0056
800: PSMB1|P20618
cor=0.81, p=1.5e−05
782: PPP2R1A|P30153
cor=0.85, p=1e−08
1121: ARRB1|P49407
cor=0.9, p=7.2e−11
1022: OLA1|Q9NTK5
cor=0.77, p=1.7e−06
1214: DARS1|P14868
cor=0.63, p=0.00033
972: MPST|P25325
cor=0.89, p=8.5e−13
933: ARPC1B|O15143
cor=0.9, p=1.9e−13
628: SH3BGRL|O75368
cor=0.8, p=8.1e−09
1133: PIP4K2A|P48426
cor=0.75, p=2.1e−07
875: CCT8|P50990
cor=0.71, p=1.8e−06
745: ACLY|P53396
cor=0.12, p=0.49
653: ITGB2|P05107
cor=0.79, p=0.0013
1220: CARHSP1|Q9Y2V2
cor=0.86, p=0.00016
1190: TMOD2|Q9NZR1
cor=0.83, p=3.2e−06
1131: TMOD3|Q9NYL9
cor=0.79, p=5.8e−07
1096: GMFG|O60234
cor=0.79, p=3.4e−05
1307: PRKG1|Q13976
cor=0.88, p=3.3e−12
1213: S100A6|P06703
cor=0.88, p=6.8e−10
985: REG1A|P05451
cor=0.11, p=0.71
1187: LYPD3|O95274

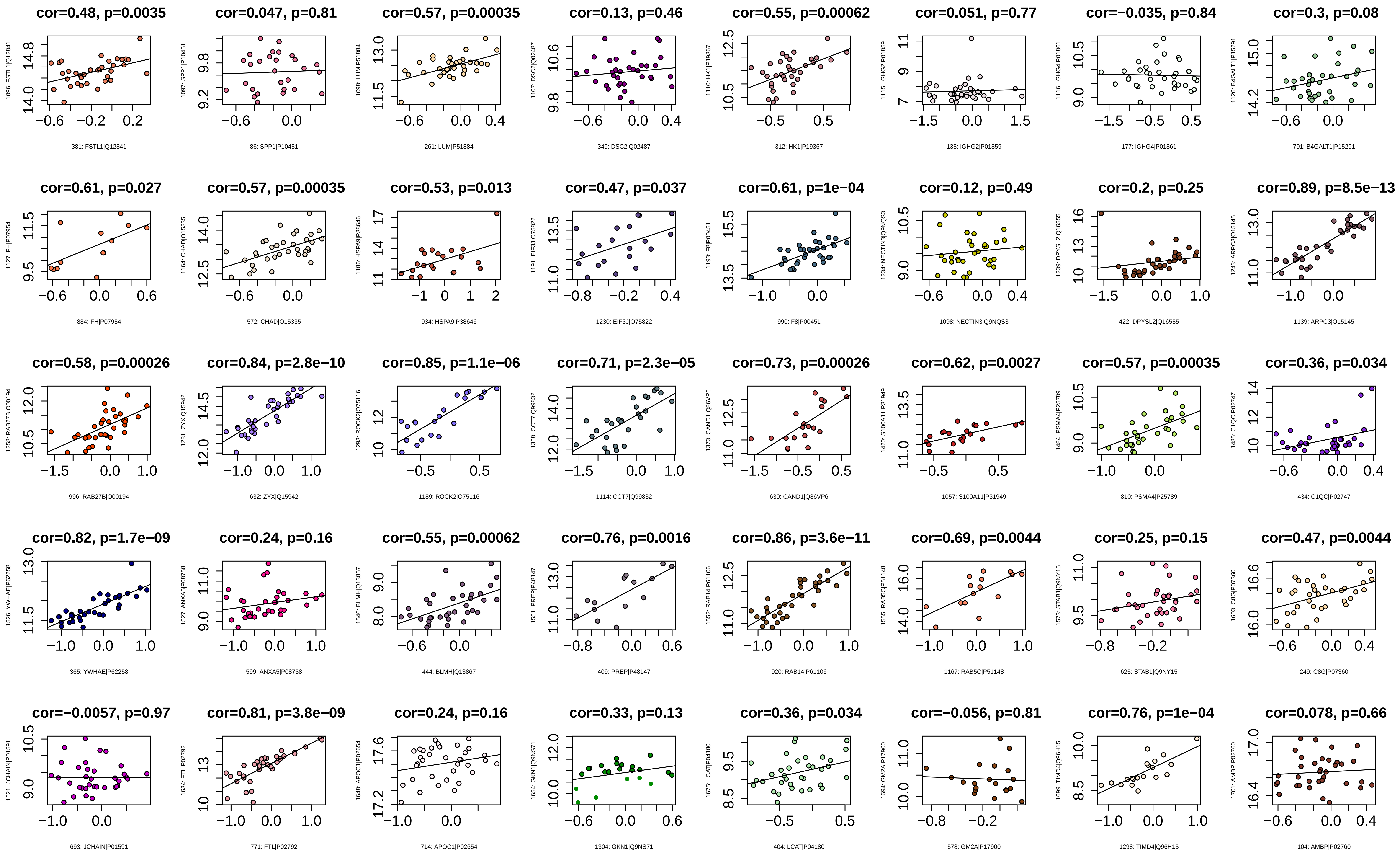
cor=0.48, p=0.0035
381: FSTL1|Q12841
cor=0.047, p=0.81
86: SPP1|P10451
cor=0.57, p=0.00035
261: LUM|P51884
cor=0.13, p=0.46
349: DSC2|Q02487
cor=0.55, p=0.00062
312: HK1|P19367
cor=0.051, p=0.77
135: IGHG2|P01859
cor=−0.035, p=0.84
177: IGHG4|P01861
cor=0.3, p=0.08
791: B4GALT1|P15291
cor=0.61, p=0.027
884: FH|P07954
cor=0.57, p=0.00035
572: CHAD|O15335
cor=0.53, p=0.013
934: HSPA9|P38646
cor=0.47, p=0.037
1230: EIF3J|O75822
cor=0.61, p=1e−04
990: F8|P00451
cor=0.12, p=0.49
1098: NECTIN3|Q9NQS3
cor=0.2, p=0.25
422: DPYSL2|Q16555
cor=0.89, p=8.5e−13
1139: ARPC3|O15145
cor=0.58, p=0.00026
996: RAB27B|O00194
cor=0.84, p=2.8e−10
632: ZYX|Q15942
cor=0.85, p=1.1e−06
1189: ROCK2|O75116
cor=0.71, p=2.3e−05
1114: CCT7|Q99832
cor=0.73, p=0.00026
630: CAND1|Q86VP6
cor=0.62, p=0.0027
1057: S100A11|P31949
cor=0.57, p=0.00035
810: PSMA4|P25789
cor=0.36, p=0.034
434: C1QC|P02747
cor=0.82, p=1.7e−09
365: YWHAE|P62258
cor=0.24, p=0.16
599: ANXA5|P08758
cor=0.55, p=0.00062
444: BLMH|Q13867
cor=0.76, p=0.0016
409: PREP|P48147
cor=0.86, p=3.6e−11
920: RAB14|P61106
cor=0.69, p=0.0044
1167: RAB5C|P51148
cor=0.25, p=0.15
625: STAB1|Q9NY15
cor=0.47, p=0.0044
249: C8G|P07360
cor=−0.0057, p=0.97
693: JCHAIN|P01591
cor=0.81, p=3.8e−09
771: FTL|P02792
cor=0.24, p=0.16
714: APOC1|P02654
cor=0.33, p=0.13
1304: GKN1|Q9NS71
cor=0.36, p=0.034
404: LCAT|P04180
cor=−0.056, p=0.81
578: GM2A|P17900
cor=0.76, p=1e−04
1298: TIMD4|Q96H15
cor=0.078, p=0.66
104: AMBP|P02760

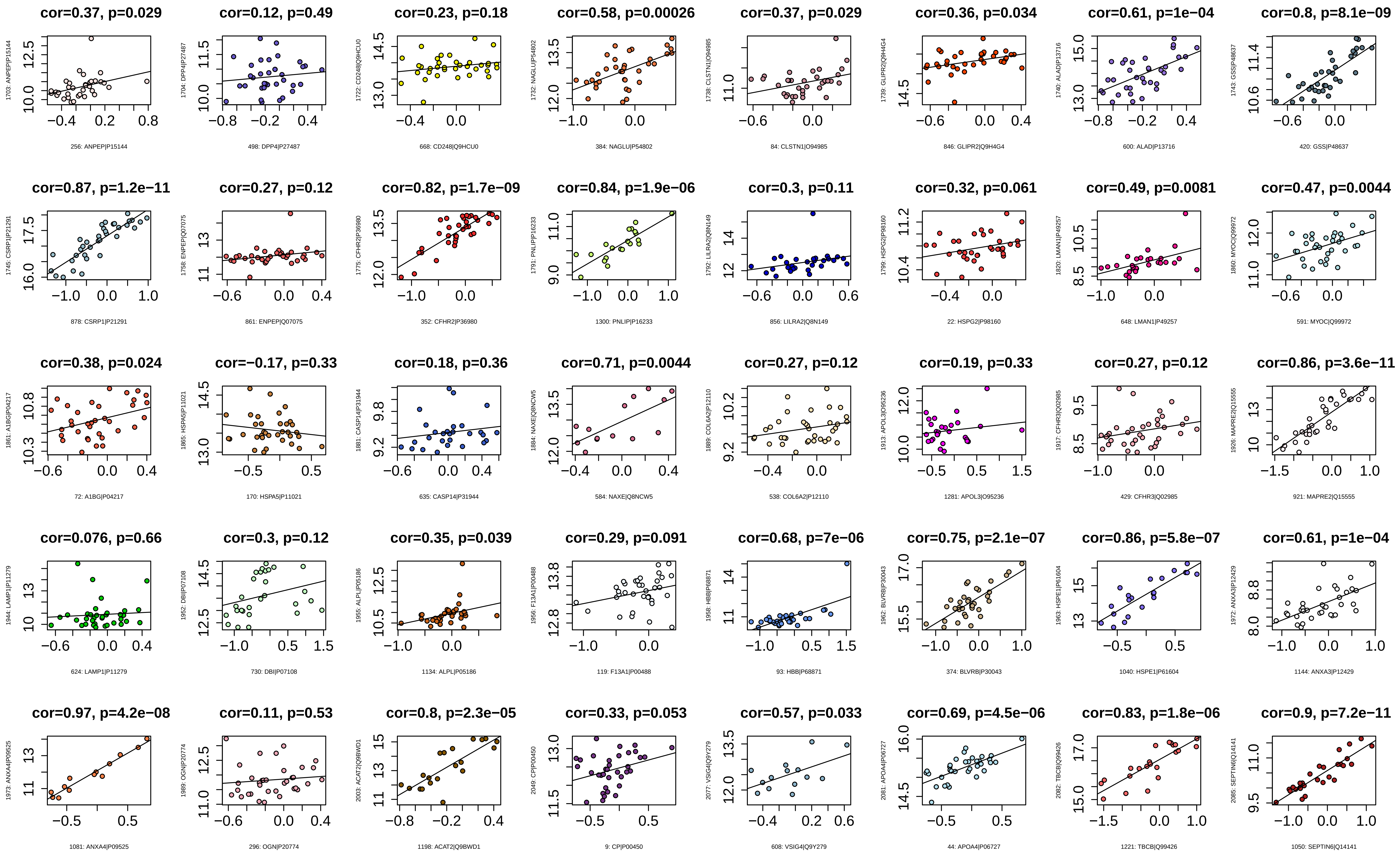

cor=0.37, p=0.029
1703: ANPEP|P15144
256: ANPEP|P15144
cor=0.12, p=0.49
1704: DPP4|P27487
498: DPP4|P27487
cor=0.23, p=0.18
1722: CD248|Q9HCU0
668: CD248|Q9HCU0
cor=0.58, p=0.00026
1732: NAGLU|P54802
384: NAGLU|P54802
cor=0.37, p=0.029
1738: CLSTN1|O94985
84: CLSTN1|O94985
cor=0.36, p=0.034
1739: GLIPR2|Q9H4G4
846: GLIPR2|Q9H4G4
cor=0.61, p=1e−04
1740: ALAD|P13716
600: ALAD|P13716
cor=0.8, p=8.1e−09
1743: GSS|P48637
420: GSS|P48637
cor=0.87, p=1.2e−11
1745: CSRP1|P21291
878: CSRP1|P21291
cor=0.27, p=0.12
1758: ENPEP|Q07075
861: ENPEP|Q07075
cor=0.82, p=1.7e−09
1775: CFHR2|P36980
352: CFHR2|P36980
cor=0.84, p=1.9e−06
1791: PNLIP|P16233
1300: PNLIP|P16233
cor=0.3, p=0.11
1792: LILRA2|Q8N149
856: LILRA2|Q8N149
cor=0.32, p=0.061
1799: HSPG2|P98160
22: HSPG2|P98160
cor=0.49, p=0.0081
1820: LMAN1|P49257
648: LMAN1|P49257
cor=0.47, p=0.0044
1860: MYOC|Q99972
591: MYOC|Q99972
cor=0.38, p=0.024
1861: A1BG|P04217
72: A1BG|P04217
cor=−0.17, p=0.33
1865: HSPA5|P11021
170: HSPA5|P11021
cor=0.18, p=0.36
1881: CASP14|P31944
635: CASP14|P31944
cor=0.71, p=0.0044
1884: NAXE|Q8NCW5
584: NAXE|Q8NCW5
cor=0.27, p=0.12
1889: COL6A2|P12110
538: COL6A2|P12110
cor=0.19, p=0.33
1913: APOL3|O95236
1281: APOL3|O95236
cor=0.27, p=0.12
1917: CFHR3|Q02985
429: CFHR3|Q02985
cor=0.86, p=3.6e−11
1926: MAPRE2|Q15555
921: MAPRE2|Q15555
cor=0.076, p=0.66
1944: LAMP1|P11279
624: LAMP1|P11279
cor=0.3, p=0.12
1952: DBI|P07108
730: DBI|P07108
cor=0.35, p=0.039
1955: ALPL|P05186
1134: ALPL|P05186
cor=0.29, p=0.091
1956: F13A1|P00488
119: F13A1|P00488
cor=0.68, p=7e−06
1958: HBB|P68871
93: HBB|P68871
cor=0.75, p=2.1e−07
1962: BLVRB|P30043
374: BLVRB|P30043
cor=0.86, p=5.8e−07
1963: HSPE1|P61604
1040: HSPE1|P61604
cor=0.61, p=1e−04
1972: ANXA3|P12429
1144: ANXA3|P12429
cor=0.97, p=4.2e−08
1973: ANXA4|P09525
1081: ANXA4|P09525
cor=0.11, p=0.53
1989: OGN|P20774
296: OGN|P20774
cor=0.8, p=2.3e−05
2003: ACAT2|Q9BWD1
1198: ACAT2|Q9BWD1
cor=0.33, p=0.053
2049: CP|P00450
9: CP|P00450
cor=0.57, p=0.033
2077: VSIG4|Q9Y279
608: VSIG4|Q9Y279
cor=0.69, p=4.5e−06
2081: APOA4|P06727
44: APOA4|P06727
cor=0.83, p=1.8e−06
2082: TBCB|Q99426
1221: TBCB|Q99426
cor=0.9, p=7.2e−11
2085: SEPTIN6|Q14141
1050: SEPTIN6|Q14141

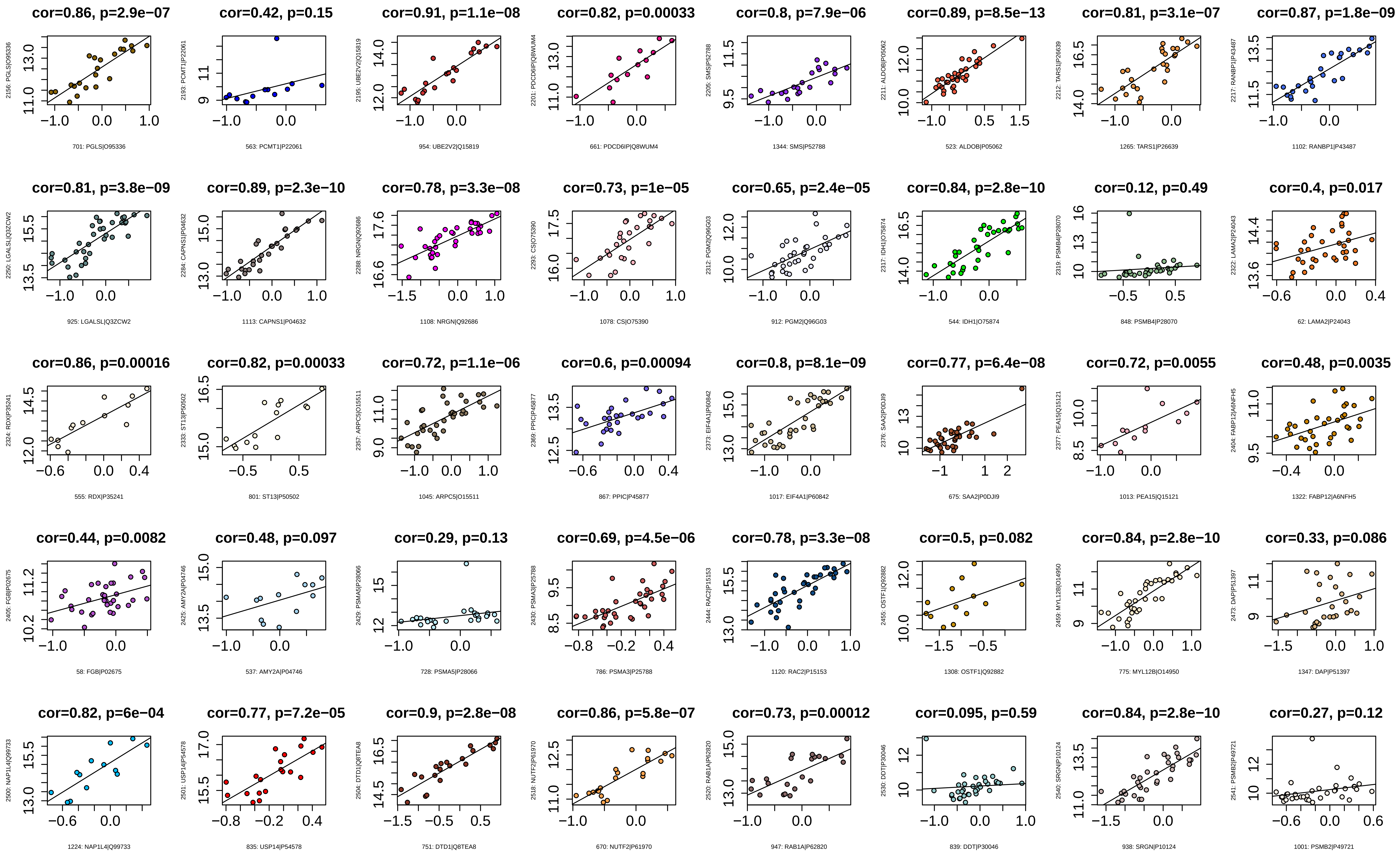
cor=0.86, p=2.9e−07
2156: PGLS|O95336
701: PGLS|O95336
cor=0.42, p=0.15
2193: PCMT1|P22061
563: PCMT1|P22061
cor=0.91, p=1.1e−08
2195: UBE2V2|Q15819
954: UBE2V2|Q15819
cor=0.82, p=0.00033
2201: PDCD6IP|Q8WUM4
661: PDCD6IP|Q8WUM4
cor=0.8, p=7.9e−06
2206: SMS|P52788
1344: SMS|P52788
cor=0.89, p=8.5e−13
2211: ALDOB|P05062
523: ALDOB|P05062
cor=0.81, p=3.1e−07
2212: TARS1|P26639
1265: TARS1|P26639
cor=0.87, p=1.8e−09
2217: RANBP1|P43487
1102: RANBP1|P43487
cor=0.81, p=3.8e−09
2250: LGALSL|Q3ZCW2
925: LGALSL|Q3ZCW2
cor=0.89, p=2.3e−10
2284: CAPNS1|P04632
1113: CAPNS1|P04632
cor=0.78, p=3.3e−08
2288: NRGN|Q92686
1108: NRGN|Q92686
cor=0.73, p=1e−05
2293: CS|O75390
1078: CS|O75390
cor=0.65, p=2.4e−05
2312: PGM2|Q96G03
912: PGM2|Q96G03
cor=0.84, p=2.8e−10
2317: IDH1|O75874
544: IDH1|O75874
cor=0.12, p=0.49
2319: PSMB4|P28070
848: PSMB4|P28070
cor=0.4, p=0.017
2322: LAMA2|P24043
62: LAMA2|P24043
cor=0.86, p=0.00016
2324: RDX|P35241
555: RDX|P35241
cor=0.82, p=0.00033
2333: ST13|P50502
801: ST13|P50502
cor=0.72, p=1.1e−06
2357: ARPC5|O15511
1045: ARPC5|O15511
cor=0.6, p=0.00094
2369: PPIC|P45877
867: PPIC|P45877
cor=0.8, p=8.1e−09
2373: EIF4A1|P60842
1017: EIF4A1|P60842
cor=0.77, p=6.4e−08
2376: SAA2|P0DJI9
675: SAA2|P0DJI9
cor=0.72, p=0.0055
2377: PEA15|Q15121
1013: PEA15|Q15121
cor=0.48, p=0.0035
2404: FABP12|A6NFH5
1322: FABP12|A6NFH5
cor=0.44, p=0.0082
2405: FGB|P02675
58: FGB|P02675
cor=0.48, p=0.097
2425: AMY2A|P04746
537: AMY2A|P04746
cor=0.29, p=0.13
2429: PSMA5|P28066
728: PSMA5|P28066
cor=0.69, p=4.5e−06
2430: PSMA3|P25788
786: PSMA3|P25788
cor=0.78, p=3.3e−08
2444: RAC2|P15153
1120: RAC2|P15153
cor=0.5, p=0.082
2455: OSTF1|Q92882
1308: OSTF1|Q92882
cor=0.84, p=2.8e−10
2459: MYL12B|O14950
775: MYL12B|O14950
cor=0.33, p=0.086
2473: DAP|P51397
1347: DAP|P51397
cor=0.82, p=6e−04
2500: NAP1L4|Q99733
1224: NAP1L4|Q99733
cor=0.77, p=7.2e−05
2501: USP14|P54578
835: USP14|P54578
cor=0.9, p=2.8e−08
2504: DTD1|Q8TEA8
751: DTD1|Q8TEA8
cor=0.86, p=5.8e−07
2518: NUTF2|P61970
670: NUTF2|P61970
cor=0.73, p=0.00012
2520: RAB1A|P62820
947: RAB1A|P62820
cor=0.095, p=0.59
2530: DDT|P30046
839: DDT|P30046
cor=0.84, p=2.8e−10
2540: SRGN|P10124
938: SRGN|P10124
cor=0.27, p=0.12
2541: PSMB2|P49721
1001: PSMB2|P49721

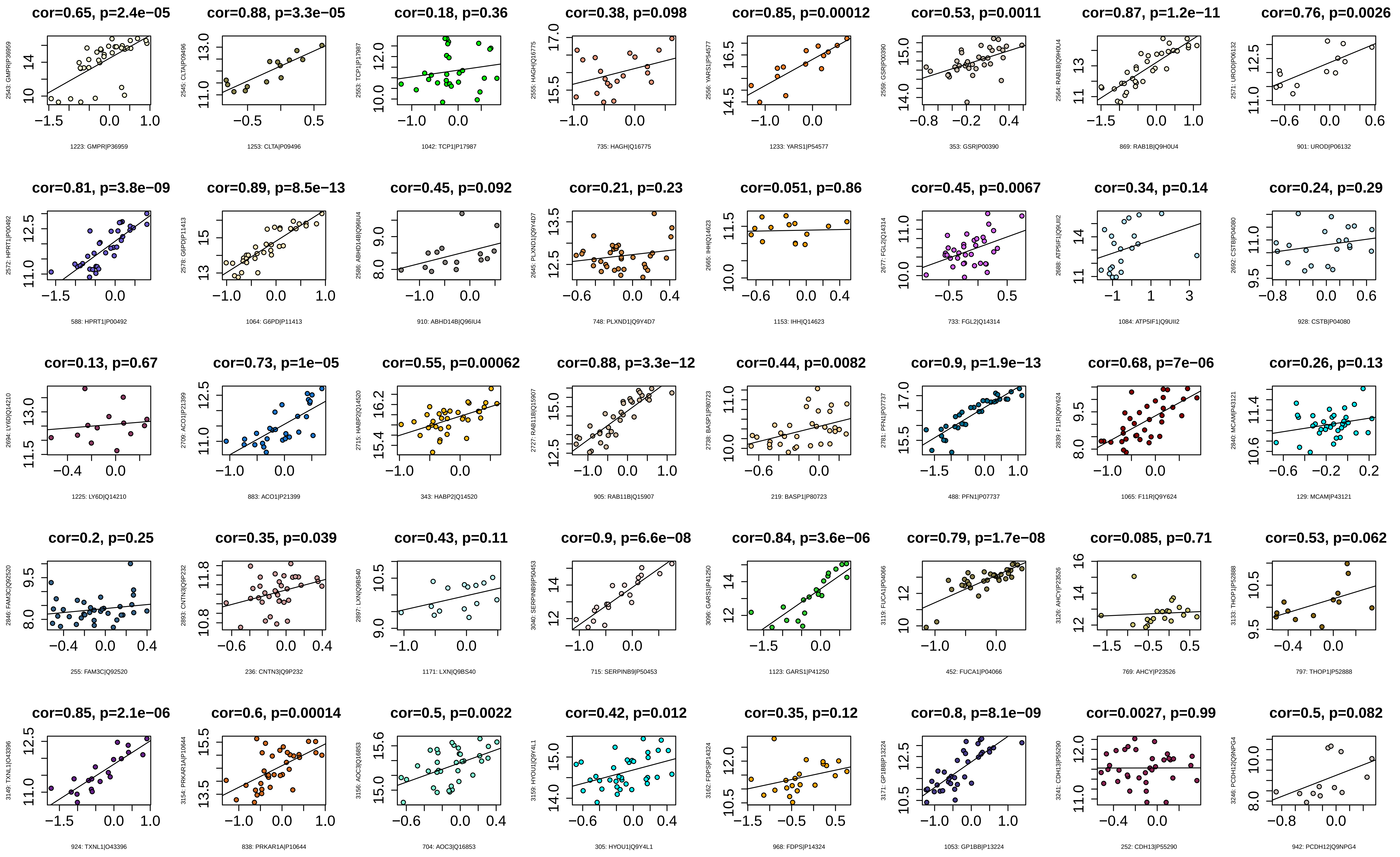
cor=0.65, p=2.4e−05
2543: GMPR|P36959
1223: GMPR|P36959
cor=0.88, p=3.3e−05
2545: CLTA|P09496
1253: CLTA|P09496
cor=0.18, p=0.36
2553: TCP1|P17987
1042: TCP1|P17987
cor=0.38, p=0.098
2555: HAGH|Q16775
735: HAGH|Q16775
cor=0.85, p=0.00012
2556: YARS1|P54577
1233: YARS1|P54577
cor=0.53, p=0.0011
2559: GSR|P00390
353: GSR|P00390
cor=0.87, p=1.2e−11
2564: RAB1B|Q9H0U4
869: RAB1B|Q9H0U4
cor=0.76, p=0.0026
2571: UROD|P06132
901: UROD|P06132
cor=0.81, p=3.8e−09
2572: HPRT1|P00492
588: HPRT1|P00492
cor=0.89, p=8.5e−13
2578: G6PD|P11413
1064: G6PD|P11413
cor=0.45, p=0.092
2586: ABHD14B|Q96IU4
910: ABHD14B|Q96IU4
cor=0.21, p=0.23
2645: PLXND1|Q9Y4D7
748: PLXND1|Q9Y4D7
cor=0.051, p=0.86
2665: IHH|Q14623
1153: IHH|Q14623
cor=0.45, p=0.0067
2677: FGL2|Q14314
733: FGL2|Q14314
cor=0.34, p=0.14
2688: ATP5IF1|Q9UII2
1084: ATP5IF1|Q9UII2
cor=0.24, p=0.29
2692: CSTB|P04080
928: CSTB|P04080
cor=0.13, p=0.67
2694: LY6D|Q14210
1225: LY6D|Q14210
cor=0.73, p=1e−05
2709: ACO1|P21399
883: ACO1|P21399
cor=0.55, p=0.00062
2715: HABP2|Q14520
343: HABP2|Q14520
cor=0.88, p=3.3e−12
2727: RAB11B|Q15907
905: RAB11B|Q15907
cor=0.44, p=0.0082
2738: BASP1|P80723
219: BASP1|P80723
cor=0.9, p=1.9e−13
2781: PFN1|P07737
488: PFN1|P07737
cor=0.68, p=7e−06
2839: F11R|Q9Y624
1065: F11R|Q9Y624
cor=0.26, p=0.13
2840: MCAM|P43121
129: MCAM|P43121
cor=0.2, p=0.25
2846: FAM3C|Q92520
255: FAM3C|Q92520
cor=0.35, p=0.039
2893: CNTN3|Q9P232
236: CNTN3|Q9P232
cor=0.43, p=0.11
2897: LXN|Q9BS40
1171: LXN|Q9BS40
cor=0.9, p=6.6e−08
3040: SERPINB9|P50453
715: SERPINB9|P50453
cor=0.84, p=3.6e−06
3096: GARS1|P41250
1123: GARS1|P41250
cor=0.79, p=1.7e−08
3119: FUCA1|P04066
452: FUCA1|P04066
cor=0.085, p=0.71
3126: AHCY|P23526
769: AHCY|P23526
cor=0.53, p=0.062
3133: THOP1|P52888
797: THOP1|P52888
cor=0.85, p=2.1e−06
3149: TXNL1|O43396
924: TXNL1|O43396
cor=0.6, p=0.00014
3154: PRKAR1A|P10644
838: PRKAR1A|P10644
cor=0.5, p=0.0022
3156: AOC3|Q16853
704: AOC3|Q16853
cor=0.42, p=0.012
3159: HYOU1|Q9Y4L1
305: HYOU1|Q9Y4L1
cor=0.35, p=0.12
3162: FDPS|P14324
968: FDPS|P14324
cor=0.8, p=8.1e−09
3171: GP1BB|P13224
1053: GP1BB|P13224
cor=0.0027, p=0.99
3241: CDH13|P55290
252: CDH13|P55290
cor=0.5, p=0.082
3246: PCDH12|Q9NPG4
942: PCDH12|Q9NPG4

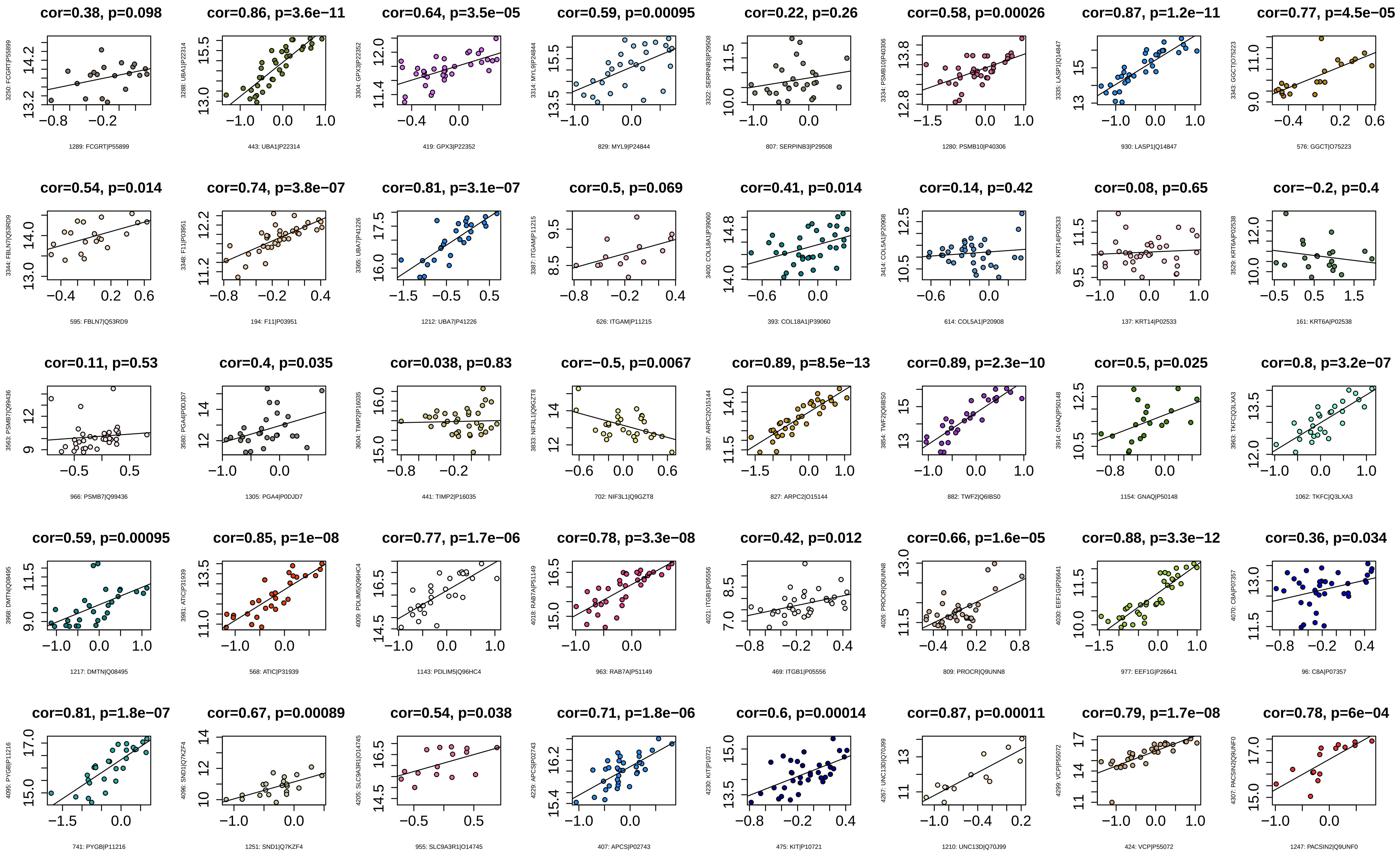
cor=0.38, p=0.098
3250: FCGRT|P55899
1289: FCGRT|P55899
cor=0.86, p=3.6e−11
3288: UBA1|P22314
443: UBA1|P22314
cor=0.64, p=3.5e−05
3304: GPX3|P22352
419: GPX3|P22352
cor=0.59, p=0.00095
3314: MYL9|P24844
829: MYL9|P24844
cor=0.22, p=0.26
3322: SERPINB3|P29508
807: SERPINB3|P29508
cor=0.58, p=0.00026
3334: PSMB10|P40306
1280: PSMB10|P40306
cor=0.87, p=1.2e−11
3335: LASP1|Q14847
930: LASP1|Q14847
cor=0.77, p=4.5e−05
3343: GGCT|O75223
576: GGCT|O75223
cor=0.54, p=0.014
3344: FBLN7|Q53RD9
595: FBLN7|Q53RD9
cor=0.74, p=3.8e−07
3348: F11|P03951
194: F11|P03951
cor=0.81, p=3.1e−07
3365: UBA7|P41226
1212: UBA7|P41226
cor=0.5, p=0.069
3387: ITGAM|P11215
626: ITGAM|P11215
cor=0.41, p=0.014
3400: COL18A1|P39060
393: COL18A1|P39060
cor=0.14, p=0.42
3414: COL5A1|P20908
614: COL5A1|P20908
cor=0.08, p=0.65
3525: KRT14|P02533
137: KRT14|P02533
cor=−0.2, p=0.4
3529: KRT6A|P02538
161: KRT6A|P02538
cor=0.11, p=0.53
3563: PSMB7|Q99436
966: PSMB7|Q99436
cor=0.4, p=0.035
3580: PGA4|P0DJD7
1305: PGA4|P0DJD7
cor=0.038, p=0.83
3604: TIMP2|P16035
441: TIMP2|P16035
cor=−0.5, p=0.0067
3833: NIF3L1|Q9GZT8
702: NIF3L1|Q9GZT8
cor=0.89, p=8.5e−13
3837: ARPC2|O15144
827: ARPC2|O15144
cor=0.89, p=2.3e−10
3854: TWF2|Q6IBS0
882: TWF2|Q6IBS0
cor=0.5, p=0.025
3914: GNAQ|P50148
1154: GNAQ|P50148
cor=0.8, p=3.2e−07
3963: TKFC|Q3LXA3
1062: TKFC|Q3LXA3
cor=0.59, p=0.00095
3968: DMTN|Q08495
1217: DMTN|Q08495
cor=0.85, p=1e−08
3981: ATIC|P31939
568: ATIC|P31939
cor=0.77, p=1.7e−06
4009: PDLIM5|Q96HC4
1143: PDLIM5|Q96HC4
cor=0.78, p=3.3e−08
4018: RAB7A|P51149
963: RAB7A|P51149
cor=0.42, p=0.012
4021: ITGB1|P05556
469: ITGB1|P05556
cor=0.66, p=1.6e−05
4026: PROCR|Q9UNN8
809: PROCR|Q9UNN8
cor=0.88, p=3.3e−12
4030: EEF1G|P26641
977: EEF1G|P26641
cor=0.36, p=0.034
4070: C8A|P07357
96: C8A|P07357
cor=0.81, p=1.8e−07
4095: PYGB|P11216
741: PYGB|P11216
cor=0.67, p=0.00089
4096: SND1|Q7KZF4
1251: SND1|Q7KZF4
cor=0.54, p=0.038
4205: SLC9A3R1|O14745
955: SLC9A3R1|O14745
cor=0.71, p=1.8e−06
4229: APCS|P02743
407: APCS|P02743
cor=0.6, p=0.00014
4230: KIT|P10721
475: KIT|P10721
cor=0.87, p=0.00011
4267: UNC13D|Q70J99
1210: UNC13D|Q70J99
cor=0.79, p=1.7e−08
4299: VCP|P55072
424: VCP|P55072
cor=0.78, p=6e−04
4307: PACSIN2|Q9UNF0
1247: PACSIN2|Q9UNF0

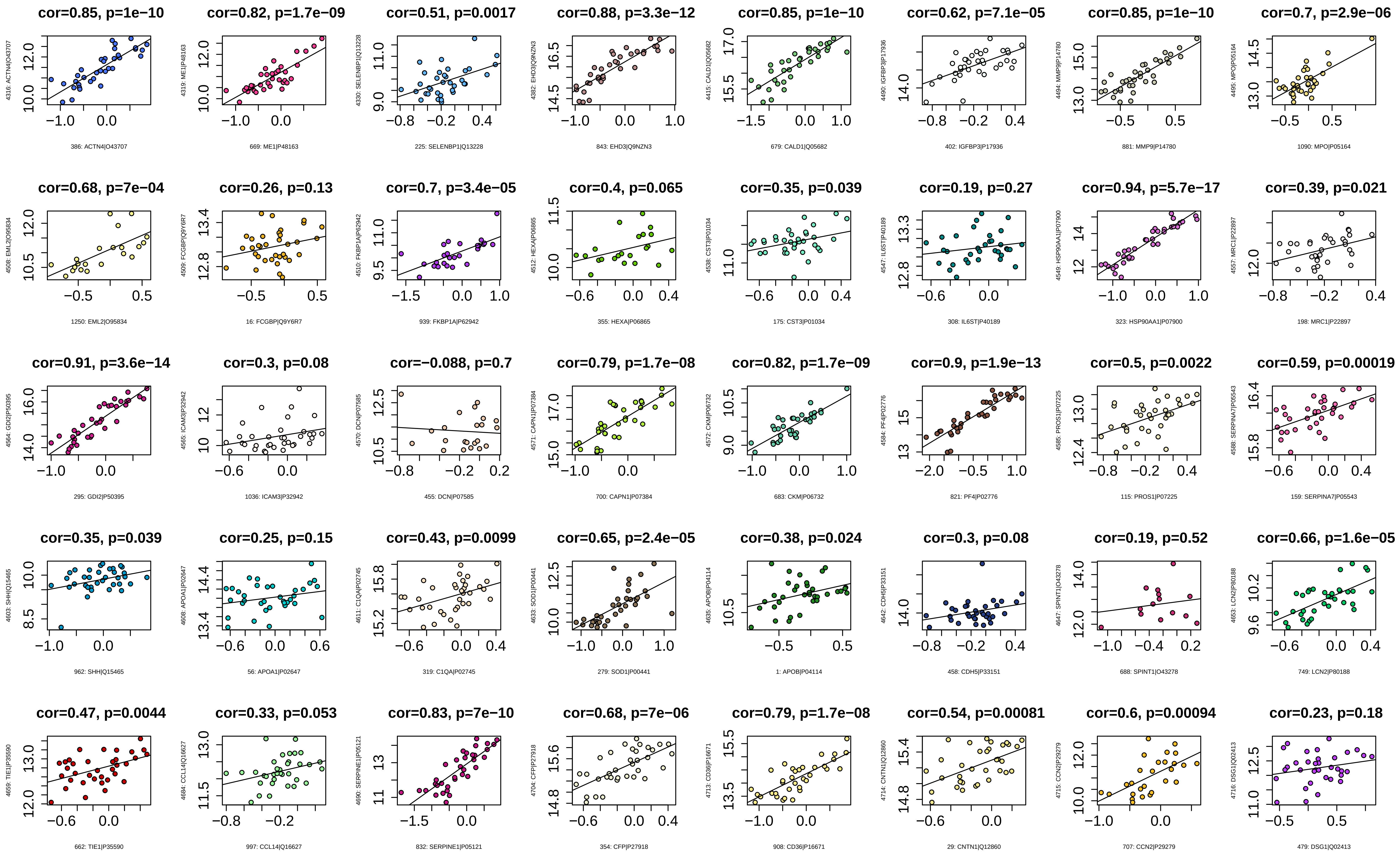
cor=0.85, p=1e−10
4316: ACTN4|O43707
386: ACTN4|O43707
cor=0.82, p=1.7e−09
4319: ME1|P48163
669: ME1|P48163
cor=0.51, p=0.0017
4330: SELENBP1|Q13228
225: SELENBP1|Q13228
cor=0.88, p=3.3e−12
4382: EHD3|Q9NZN3
843: EHD3|Q9NZN3
cor=0.85, p=1e−10
4415: CALD1|Q05682
679: CALD1|Q05682
cor=0.62, p=7.1e−05
4490: IGFBP3|P17936
402: IGFBP3|P17936
cor=0.85, p=1e−10
4494: MMP9|P14780
881: MMP9|P14780
cor=0.7, p=2.9e−06
4495: MPO|P05164
1090: MPO|P05164
cor=0.68, p=7e−04
4508: EML2|O95834
1250: EML2|O95834
cor=0.26, p=0.13
4509: FCGBP|Q9Y6R7
16: FCGBP|Q9Y6R7
cor=0.7, p=3.4e−05
4510: FKBP1A|P62942
939: FKBP1A|P62942
cor=0.4, p=0.065
4512: HEXA|P06865
355: HEXA|P06865
cor=0.35, p=0.039
4538: CST3|P01034
175: CST3|P01034
cor=0.19, p=0.27
4547: IL6ST|P40189
308: IL6ST|P40189
cor=0.94, p=5.7e−17
4549: HSP90AA1|P07900
323: HSP90AA1|P07900
cor=0.39, p=0.021
4557: MRC1|P22897
198: MRC1|P22897
cor=0.91, p=3.6e−14
4564: GDI2|P50395
295: GDI2|P50395
cor=0.3, p=0.08
4565: ICAM3|P32942
1036: ICAM3|P32942
cor=−0.088, p=0.7
4570: DCN|P07585
455: DCN|P07585
cor=0.79, p=1.7e−08
4571: CAPN1|P07384
700: CAPN1|P07384
cor=0.82, p=1.7e−09
4572: CKM|P06732
683: CKM|P06732
cor=0.9, p=1.9e−13
4584: PF4|P02776
821: PF4|P02776
cor=0.5, p=0.0022
4585: PROS1|P07225
115: PROS1|P07225
cor=0.59, p=0.00019
4588: SERPINA7|P05543
159: SERPINA7|P05543
cor=0.35, p=0.039
4603: SHH|Q15465
962: SHH|Q15465
cor=0.25, p=0.15
4608: APOA1|P02647
56: APOA1|P02647
cor=0.43, p=0.0099
4611: C1QA|P02745
319: C1QA|P02745
cor=0.65, p=2.4e−05
4633: SOD1|P00441
279: SOD1|P00441
cor=0.38, p=0.024
4635: APOB|P04114
1: APOB|P04114
cor=0.3, p=0.08
4642: CDH5|P33151
458: CDH5|P33151
cor=0.19, p=0.52
4647: SPINT1|O43278
688: SPINT1|O43278
cor=0.66, p=1.6e−05
4653: LCN2|P80188
749: LCN2|P80188
cor=0.47, p=0.0044
4659: TIE1|P35590
662: TIE1|P35590
cor=0.33, p=0.053
4684: CCL14|Q16627
997: CCL14|Q16627
cor=0.83, p=7e−10
4690: SERPINE1|P05121
832: SERPINE1|P05121
cor=0.68, p=7e−06
4704: CFP|P27918
354: CFP|P27918
cor=0.79, p=1.7e−08
4713: CD36|P16671
908: CD36|P16671
cor=0.54, p=0.00081
4714: CNTN1|Q12860
29: CNTN1|Q12860
cor=0.6, p=0.00094
4715: CCN2|P29279
707: CCN2|P29279
cor=0.23, p=0.18
4716: DSG1|Q02413
479: DSG1|Q02413

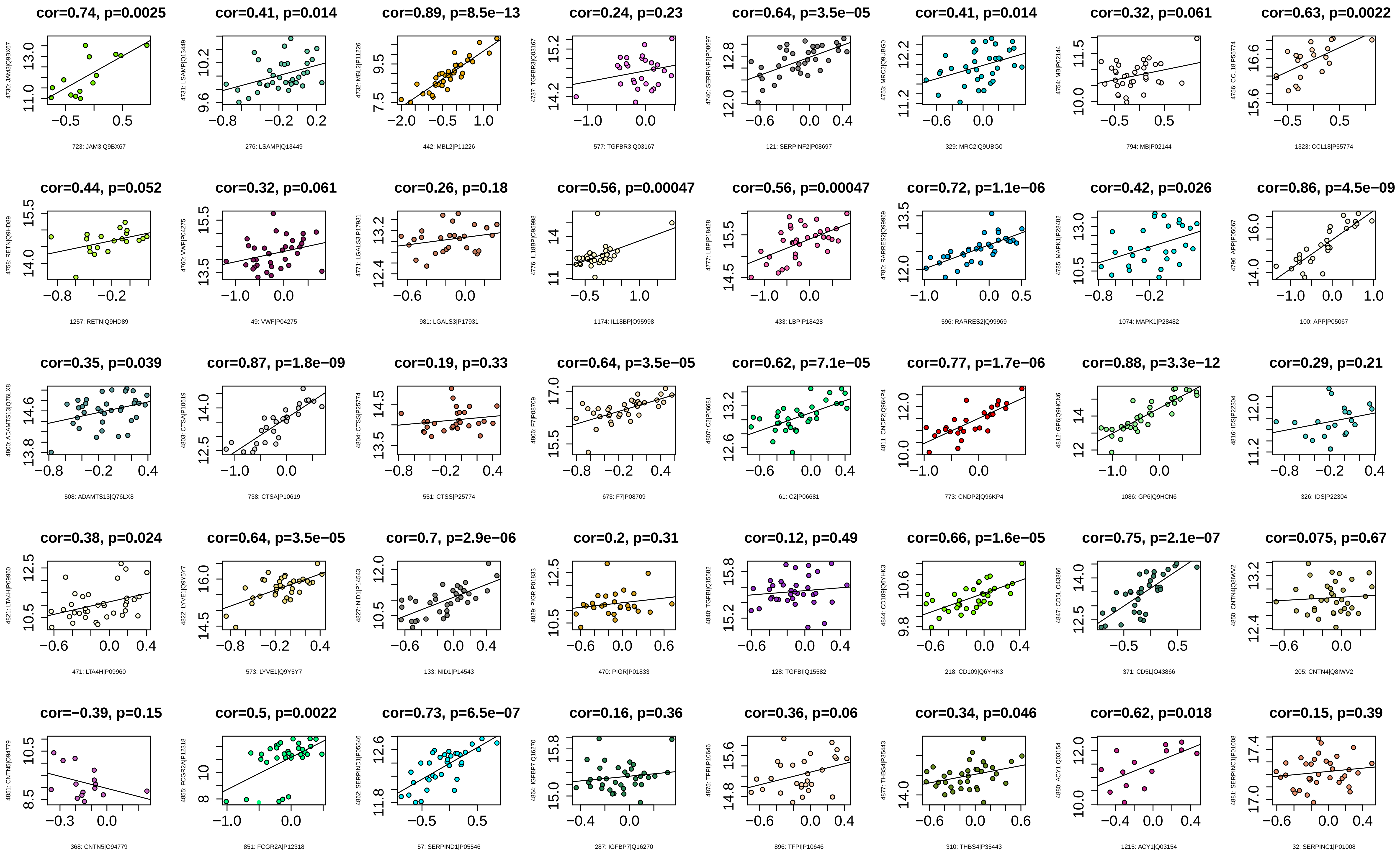
cor=0.74, p=0.0025
4730: JAM3|Q9BX67
723: JAM3|Q9BX67
cor=0.41, p=0.014
4731: LSAMP|Q13449
276: LSAMP|Q13449
cor=0.89, p=8.5e−13
4732: MBL2|P11226
442: MBL2|P11226
cor=0.24, p=0.23
4737: TGFBR3|Q03167
577: TGFBR3|Q03167
cor=0.64, p=3.5e−05
4740: SERPINF2|P08697
121: SERPINF2|P08697
cor=0.41, p=0.014
4753: MRC2|Q9UBG0
329: MRC2|Q9UBG0
cor=0.32, p=0.061
4754: MB|P02144
794: MB|P02144
cor=0.63, p=0.0022
4756: CCL18|P55774
1323: CCL18|P55774
cor=0.44, p=0.052
4758: RETN|Q9HD89
1257: RETN|Q9HD89
cor=0.32, p=0.061
4760: VWF|P04275
49: VWF|P04275
cor=0.26, p=0.18
4771: LGALS3|P17931
981: LGALS3|P17931
cor=0.56, p=0.00047
4776: IL18BP|O95998
1174: IL18BP|O95998
cor=0.56, p=0.00047
4777: LBP|P18428
433: LBP|P18428
cor=0.72, p=1.1e−06
4780: RARRES2|Q99969
596: RARRES2|Q99969
cor=0.42, p=0.026
4785: MAPK1|P28482
1074: MAPK1|P28482
cor=0.86, p=4.5e−09
4796: APP|P05067
100: APP|P05067
cor=0.35, p=0.039
4800: ADAMTS13|Q76LX8
508: ADAMTS13|Q76LX8
cor=0.87, p=1.8e−09
4803: CTSA|P10619
738: CTSA|P10619
cor=0.19, p=0.33
4804: CTSS|P25774
551: CTSS|P25774
cor=0.64, p=3.5e−05
4806: F7|P08709
673: F7|P08709
cor=0.62, p=7.1e−05
4807: C2|P06681
61: C2|P06681
cor=0.77, p=1.7e−06
4811: CNDP2|Q96KP4
773: CNDP2|Q96KP4
cor=0.88, p=3.3e−12
4812: GP6|Q9HCN6
1086: GP6|Q9HCN6
cor=0.29, p=0.21
4816: IDS|P22304
326: IDS|P22304
cor=0.38, p=0.024
4821: LTA4H|P09960
471: LTA4H|P09960
cor=0.64, p=3.5e−05
4822: LYVE1|Q9Y5Y7
573: LYVE1|Q9Y5Y7
cor=0.7, p=2.9e−06
4827: NID1|P14543
133: NID1|P14543
cor=0.2, p=0.31
4829: PIGR|P01833
470: PIGR|P01833
cor=0.12, p=0.49
4840: TGFBI|Q15582
128: TGFBI|Q15582
cor=0.66, p=1.6e−05
4844: CD109|Q6YHK3
218: CD109|Q6YHK3
cor=0.75, p=2.1e−07
4847: CD5L|O43866
371: CD5L|O43866
cor=0.075, p=0.67
4850: CNTN4|Q8IWV2
205: CNTN4|Q8IWV2
cor=−0.39, p=0.15
4851: CNTN5|O94779
368: CNTN5|O94779
cor=0.5, p=0.0022
4855: FCGR2A|P12318
851: FCGR2A|P12318
cor=0.73, p=6.5e−07
4862: SERPIND1|P05546
57: SERPIND1|P05546
cor=0.16, p=0.36
4864: IGFBP7|Q16270
287: IGFBP7|Q16270
cor=0.36, p=0.06
4875: TFPI|P10646
896: TFPI|P10646
cor=0.34, p=0.046
4877: THBS4|P35443
310: THBS4|P35443
cor=0.62, p=0.018
4880: ACY1|Q03154
1215: ACY1|Q03154
cor=0.15, p=0.39
4881: SERPINC1|P01008
32: SERPINC1|P01008

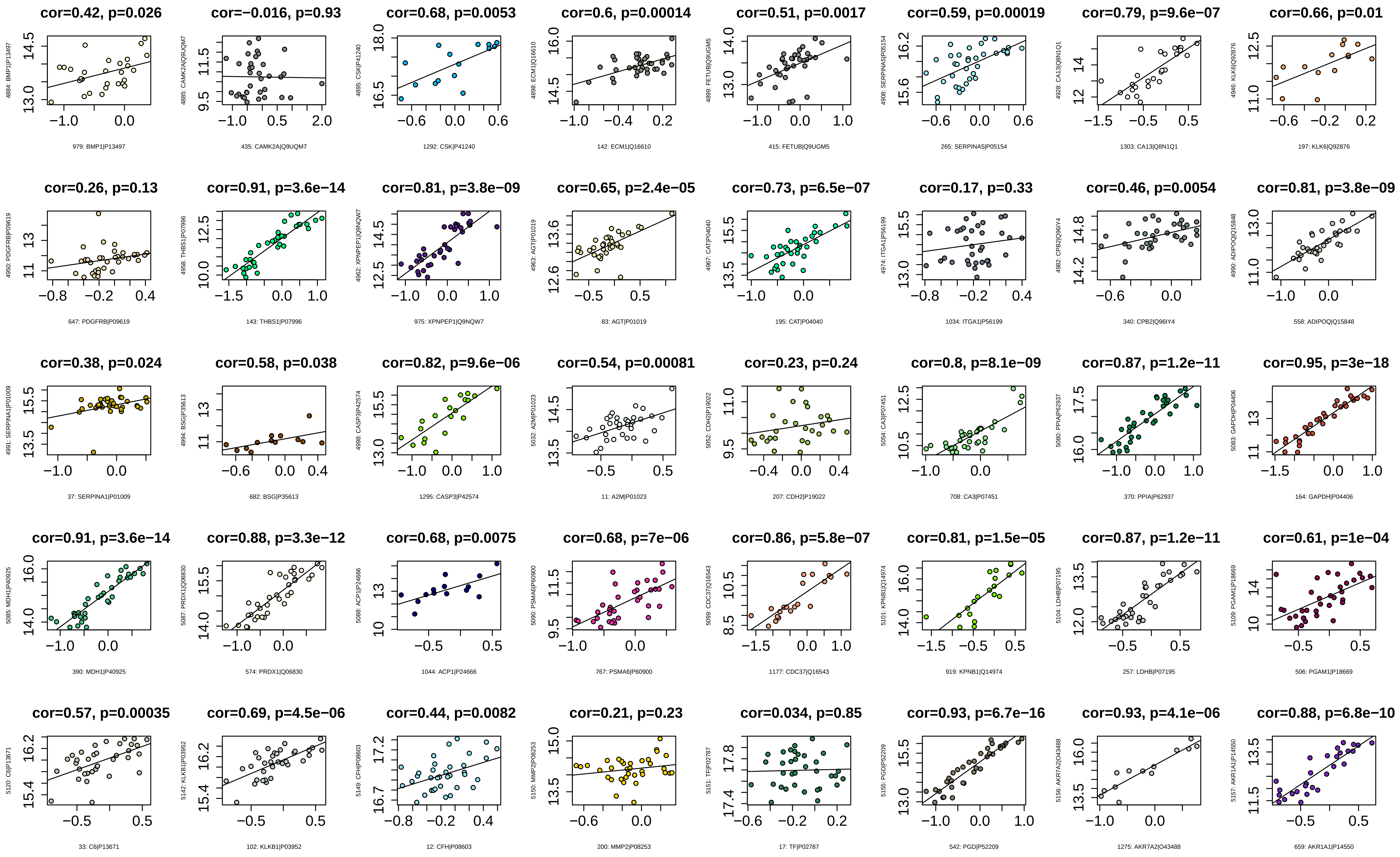
cor=0.42, p=0.026
4884: BMP1|P13497
979: BMP1|P13497
cor=−0.016, p=0.93
4885: CAMK2A|Q9UQM7
435: CAMK2A|Q9UQM7
cor=0.68, p=0.0053
4895: CSK|P41240
1292: CSK|P41240
cor=0.6, p=0.00014
4898: ECM1|Q16610
142: ECM1|Q16610
cor=0.51, p=0.0017
4899: FETUB|Q9UGM5
415: FETUB|Q9UGM5
cor=0.59, p=0.00019
4906: SERPINA5|P05154
265: SERPINA5|P05154
cor=0.79, p=9.6e−07
4928: CA13|Q8N1Q1
1303: CA13|Q8N1Q1
cor=0.66, p=0.01
4946: KLK6|Q92876
197: KLK6|Q92876
cor=0.26, p=0.13
4950: PDGFRB|P09619
647: PDGFRB|P09619
cor=0.91, p=3.6e−14
4958: THBS1|P07996
143: THBS1|P07996
cor=0.81, p=3.8e−09
4962: XPNPEP1|Q9NQW7
975: XPNPEP1|Q9NQW7
cor=0.65, p=2.4e−05
4963: AGT|P01019
83: AGT|P01019
cor=0.73, p=6.5e−07
4967: CAT|P04040
195: CAT|P04040
cor=0.17, p=0.33
4974: ITGA1|P56199
1034: ITGA1|P56199
cor=0.46, p=0.0054
4982: CPB2|Q96IY4
340: CPB2|Q96IY4
cor=0.81, p=3.8e−09
4990: ADIPOQ|Q15848
558: ADIPOQ|Q15848
cor=0.38, p=0.024
4991: SERPINA1|P01009
37: SERPINA1|P01009
cor=0.58, p=0.038
4994: BSG|P35613
682: BSG|P35613
cor=0.82, p=9.6e−06
4998: CASP3|P42574
1295: CASP3|P42574
cor=0.54, p=0.00081
5032: A2M|P01023
11: A2M|P01023
cor=0.23, p=0.24
5052: CDH2|P19022
207: CDH2|P19022
cor=0.8, p=8.1e−09
5054: CA3|P07451
708: CA3|P07451
cor=0.87, p=1.2e−11
5080: PPIA|P62937
370: PPIA|P62937
cor=0.95, p=3e−18
5083: GAPDH|P04406
164: GAPDH|P04406
cor=0.91, p=3.6e−14
5085: MDH1|P40925
390: MDH1|P40925
cor=0.88, p=3.3e−12
5087: PRDX1|Q06830
574: PRDX1|Q06830
cor=0.68, p=0.0075
5088: ACP1|P24666
1044: ACP1|P24666
cor=0.68, p=7e−06
5090: PSMA6|P60900
767: PSMA6|P60900
cor=0.86, p=5.8e−07
5099: CDC37|Q16543
1177: CDC37|Q16543
cor=0.81, p=1.5e−05
5101: KPNB1|Q14974
919: KPNB1|Q14974
cor=0.87, p=1.2e−11
5104: LDHB|P07195
257: LDHB|P07195
cor=0.61, p=1e−04
5109: PGAM1|P18669
506: PGAM1|P18669
cor=0.57, p=0.00035
5120: C6|P13671
33: C6|P13671
cor=0.69, p=4.5e−06
5142: KLKB1|P03952
102: KLKB1|P03952
cor=0.44, p=0.0082
5149: CFH|P08603
12: CFH|P08603
cor=0.21, p=0.23
5150: MMP2|P08253
200: MMP2|P08253
cor=0.034, p=0.85
5151: TF|P02787
17: TF|P02787
cor=0.93, p=6.7e−16
5155: PGD|P52209
542: PGD|P52209
cor=0.93, p=4.1e−06
5156: AKR7A2|O43488
1275: AKR7A2|O43488
cor=0.88, p=6.8e−10
5157: AKR1A1|P14550
659: AKR1A1|P14550

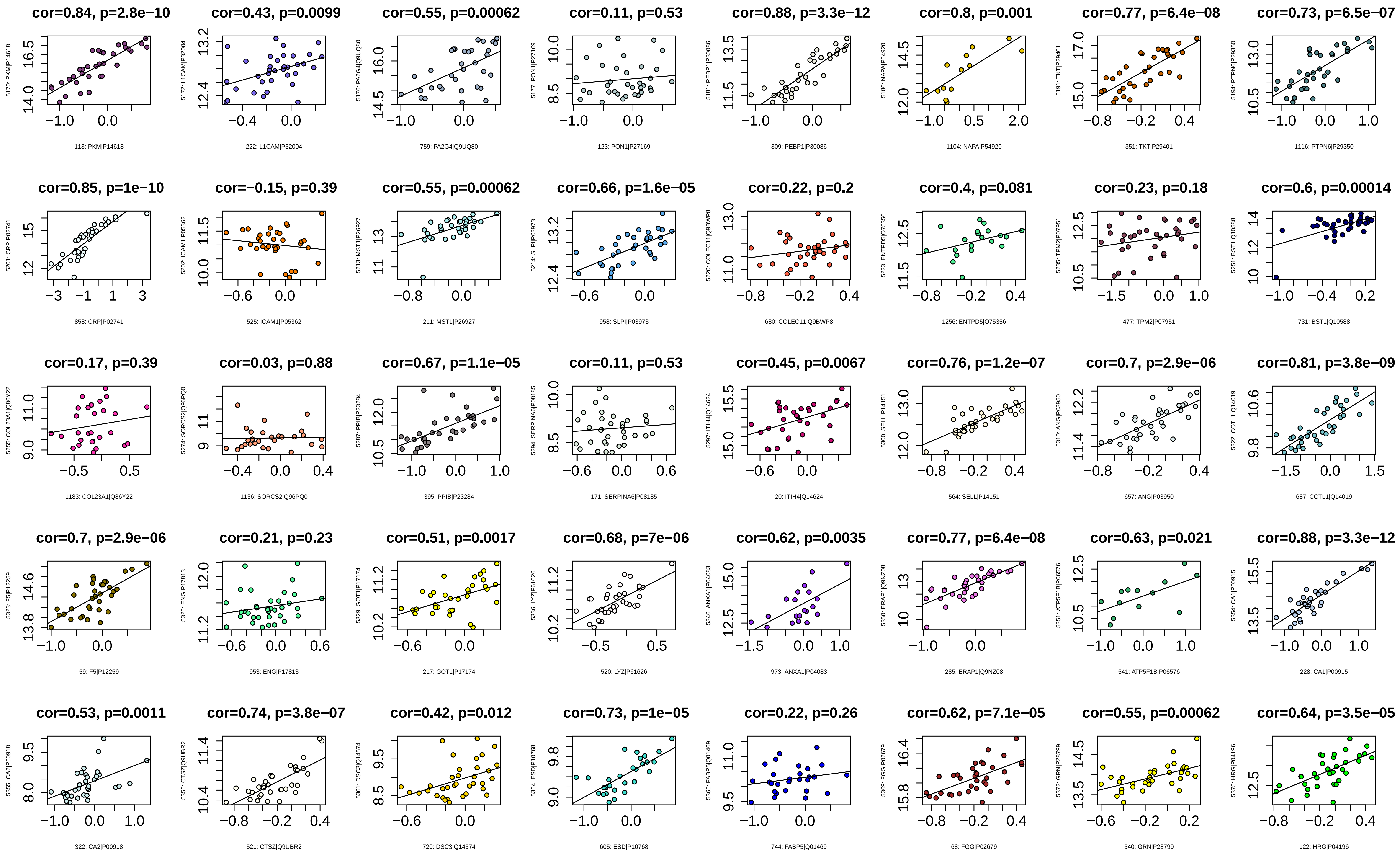
cor=0.84, p=2.8e−10
5170: PKM|P14618
113: PKM|P14618
cor=0.43, p=0.0099
5172: L1CAM|P32004
222: L1CAM|P32004
cor=0.55, p=0.00062
5176: PA2G4|Q9UQ80
759: PA2G4|Q9UQ80
cor=0.11, p=0.53
5177: PON1|P27169
123: PON1|P27169
cor=0.88, p=3.3e−12
5181: PEBP1|P30086
309: PEBP1|P30086
cor=0.8, p=0.001
5186: NAPA|P54920
1104: NAPA|P54920
cor=0.77, p=6.4e−08
5191: TKT|P29401
351: TKT|P29401
cor=0.73, p=6.5e−07
5194: PTPN6|P29350
1116: PTPN6|P29350
cor=0.85, p=1e−10
5201: CRP|P02741
858: CRP|P02741
cor=−0.15, p=0.39
5202: ICAM1|P05362
525: ICAM1|P05362
cor=0.55, p=0.00062
5213: MST1|P26927
211: MST1|P26927
cor=0.66, p=1.6e−05
5214: SLPI|P03973
958: SLPI|P03973
cor=0.22, p=0.2
5220: COLEC11|Q9BWP8
680: COLEC11|Q9BWP8
cor=0.4, p=0.081
5223: ENTPD5|O75356
1256: ENTPD5|O75356
cor=0.23, p=0.18
5235: TPM2|P07951
477: TPM2|P07951
cor=0.6, p=0.00014
5251: BST1|Q10588
731: BST1|Q10588
cor=0.17, p=0.39
5255: COL23A1|Q86Y22
1183: COL23A1|Q86Y22
cor=0.03, p=0.88
5274: SORCS2|Q96PQ0
1136: SORCS2|Q96PQ0
cor=0.67, p=1.1e−05
5287: PPIB|P23284
395: PPIB|P23284
cor=0.11, p=0.53
5294: SERPINA6|P08185
171: SERPINA6|P08185
cor=0.45, p=0.0067
5297: ITIH4|Q14624
20: ITIH4|Q14624
cor=0.76, p=1.2e−07
5300: SELL|P14151
564: SELL|P14151
cor=0.7, p=2.9e−06
5310: ANG|P03950
657: ANG|P03950
cor=0.81, p=3.8e−09
5322: COTL1|Q14019
687: COTL1|Q14019
cor=0.7, p=2.9e−06
5323: F5|P12259
59: F5|P12259
cor=0.21, p=0.23
5325: ENG|P17813
953: ENG|P17813
cor=0.51, p=0.0017
5329: GOT1|P17174
217: GOT1|P17174
cor=0.68, p=7e−06
5336: LYZ|P61626
520: LYZ|P61626
cor=0.62, p=0.0035
5346: ANXA1|P04083
973: ANXA1|P04083
cor=0.77, p=6.4e−08
5350: ERAP1|Q9NZ08
285: ERAP1|Q9NZ08
cor=0.63, p=0.021
5351: ATP5F1B|P06576
541: ATP5F1B|P06576
cor=0.88, p=3.3e−12
5354: CA1|P00915
228: CA1|P00915
cor=0.53, p=0.0011
5355: CA2|P00918
322: CA2|P00918
cor=0.74, p=3.8e−07
5356: CTSZ|Q9UBR2
521: CTSZ|Q9UBR2
cor=0.42, p=0.012
5361: DSC3|Q14574
720: DSC3|Q14574
cor=0.73, p=1e−05
5364: ESD|P10768
605: ESD|P10768
cor=0.22, p=0.26
5365: FABP5|Q01469
744: FABP5|Q01469
cor=0.62, p=7.1e−05
5369: FGG|P02679
68: FGG|P02679
cor=0.55, p=0.00062
5372: GRN|P28799
540: GRN|P28799
cor=0.64, p=3.5e−05
5375: HRG|P04196
122: HRG|P04196

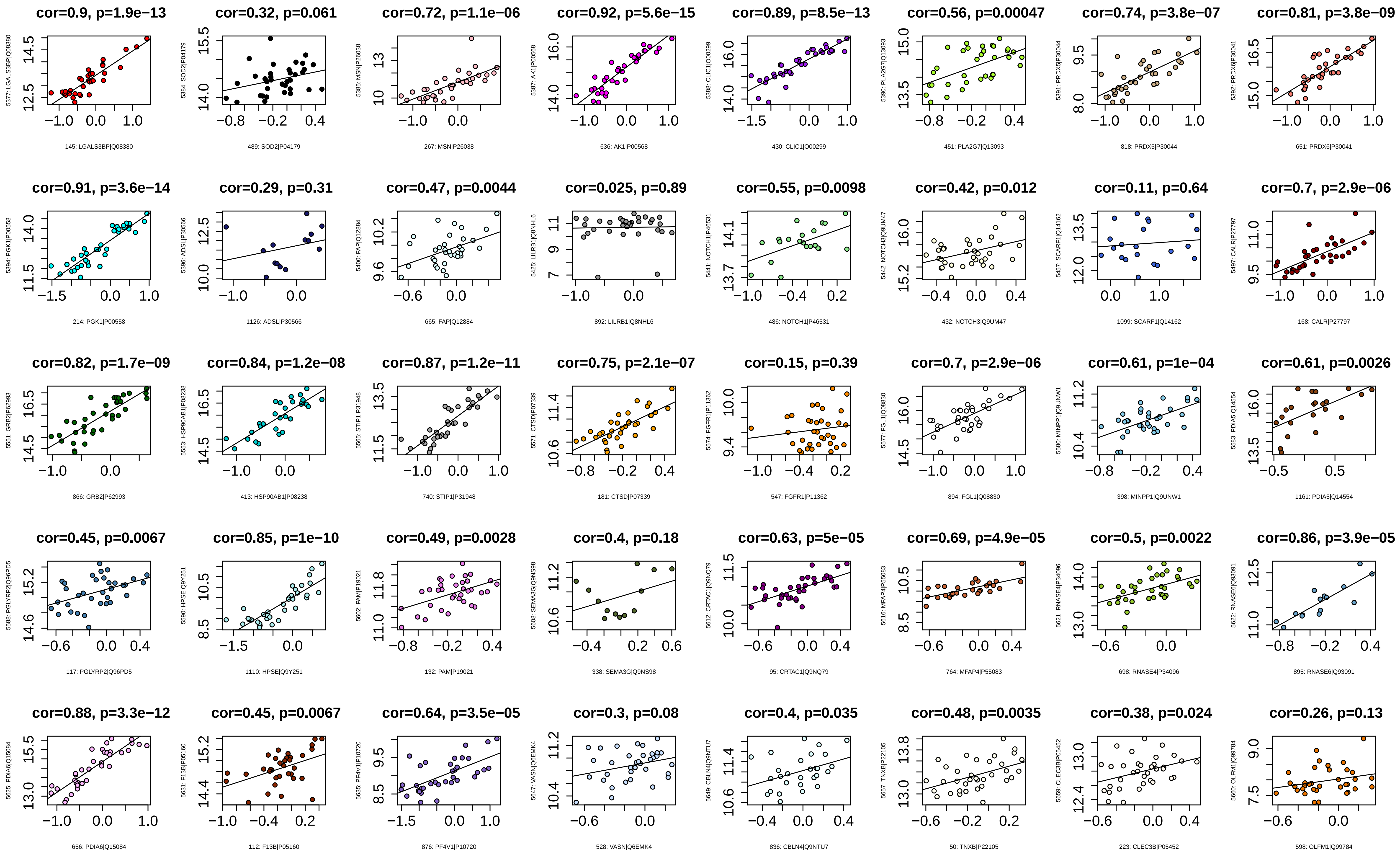

cor=0.9, p=1.9e−13
145: LGALS3BP|Q08380
cor=0.32, p=0.061
489: SOD2|P04179
cor=0.72, p=1.1e−06
267: MSN|P26038
cor=0.92, p=5.6e−15
636: AK1|P00568
cor=0.89, p=8.5e−13
430: CLIC1|O00299
cor=0.56, p=0.00047
451: PLA2G7|Q13093
cor=0.74, p=3.8e−07
818: PRDX5|P30044
cor=0.81, p=3.8e−09
651: PRDX6|P30041
cor=0.91, p=3.6e−14
214: PGK1|P00558
cor=0.29, p=0.31
1126: ADSL|P30566
cor=0.47, p=0.0044
665: FAP|Q12884
cor=0.025, p=0.89
892: LILRB1|Q8NHL6
cor=0.55, p=0.0098
486: NOTCH1|P46531
cor=0.42, p=0.012
432: NOTCH3|Q9UM47
cor=0.11, p=0.64
1099: SCARF1|Q14162
cor=0.7, p=2.9e−06
168: CALR|P27797
cor=0.82, p=1.7e−09
866: GRB2|P62993
cor=0.84, p=1.2e−08
413: HSP90AB1|P08238
cor=0.87, p=1.2e−11
740: STIP1|P31948
cor=0.75, p=2.1e−07
181: CTSD|P07339
cor=0.15, p=0.39
547: FGFR1|P11362
cor=0.7, p=2.9e−06
894: FGL1|Q08830
cor=0.61, p=1e−04
398: MINPP1|Q9UNW1
cor=0.61, p=0.0026
1161: PDIA5|Q14554
cor=0.45, p=0.0067
117: PGLYRP2|Q96PD5
cor=0.85, p=1e−10
1110: HPSE|Q9Y251
cor=0.49, p=0.0028
132: PAM|P19021
cor=0.4, p=0.18
338: SEMA3G|Q9NS98
cor=0.63, p=5e−05
95: CRTAC1|Q9NQ79
cor=0.69, p=4.9e−05
764: MFAP4|P55083
cor=0.5, p=0.0022
698: RNASE4|P34096
cor=0.86, p=3.9e−05
895: RNASE6|Q93091
cor=0.88, p=3.3e−12
656: PDIA6|Q15084
cor=0.45, p=0.0067
112: F13B|P05160
cor=0.64, p=3.5e−05
876: PF4V1|P10720
cor=0.3, p=0.08
528: VASN|Q6EMK4
cor=0.4, p=0.035
836: CBLN4|Q9NTU7
cor=0.48, p=0.0035
50: TNXB|P22105
cor=0.38, p=0.024
223: CLEC3B|P05452
cor=0.26, p=0.13
598: OLFM1|Q99784

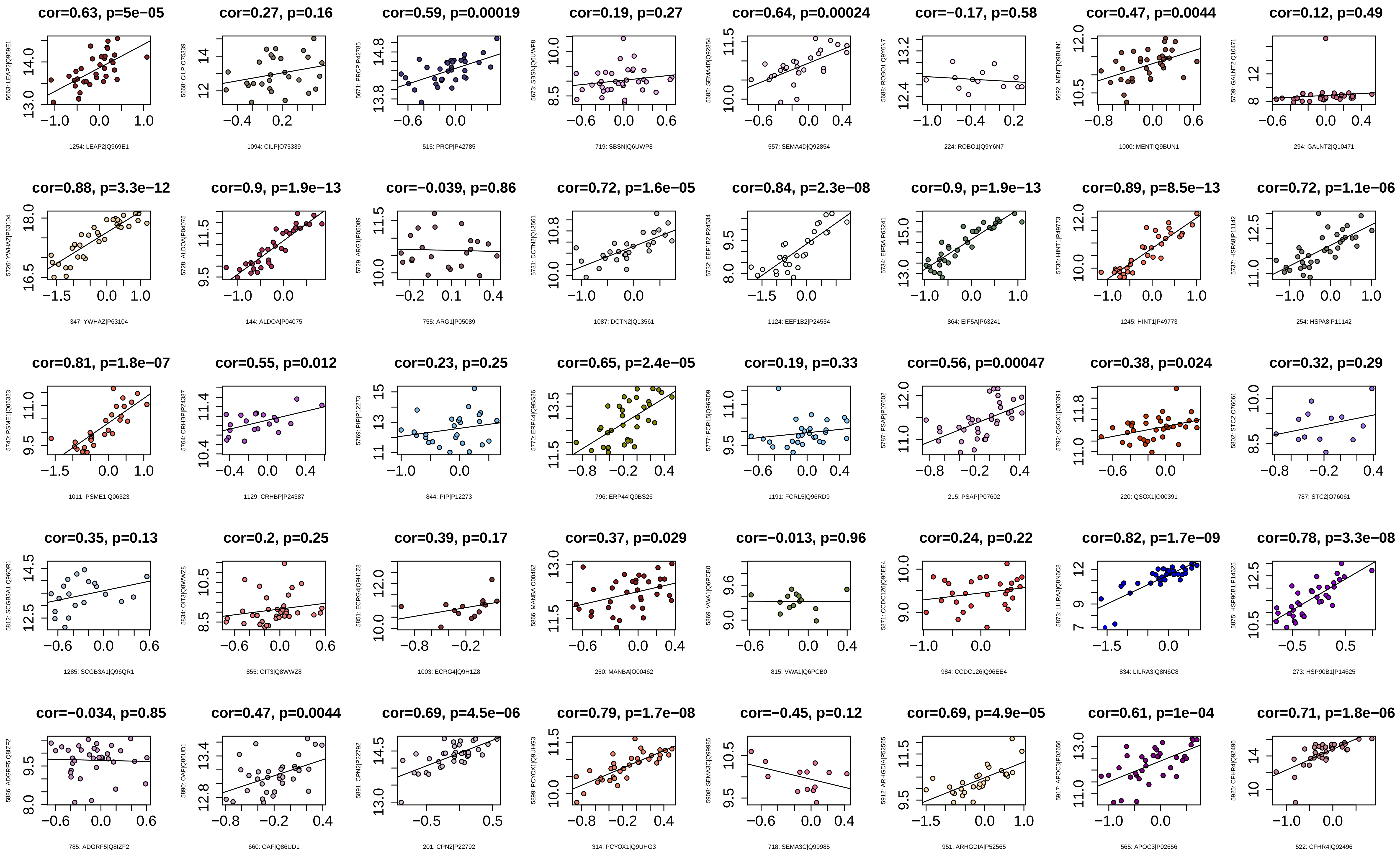
cor=0.63, p=5e−05
5663: LEAP2|Q969E1
1254: LEAP2|Q969E1
cor=0.27, p=0.16
5668: CILP|O75339
1094: CILP|O75339
cor=0.59, p=0.00019
5671: PRCP|P42785
515: PRCP|P42785
cor=0.19, p=0.27
5673: SBSN|Q6UWP8
719: SBSN|Q6UWP8
cor=0.64, p=0.00024
5685: SEMA4D|Q92854
557: SEMA4D|Q92854
cor=−0.17, p=0.58
5688: ROBO1|Q9Y6N7
224: ROBO1|Q9Y6N7
cor=0.47, p=0.0044
5692: MENT|Q9BUN1
1000: MENT|Q9BUN1
cor=0.12, p=0.49
5709: GALNT2|Q10471
294: GALNT2|Q10471
cor=0.88, p=3.3e−12
5726: YWHAZ|P63104
347: YWHAZ|P63104
cor=0.9, p=1.9e−13
5728: ALDOA|P04075
144: ALDOA|P04075
cor=−0.039, p=0.86
5729: ARG1|P05089
755: ARG1|P05089
cor=0.72, p=1.6e−05
5731: DCTN2|Q13561
1087: DCTN2|Q13561
cor=0.84, p=2.3e−08
5732: EEF1B2|P24534
1124: EEF1B2|P24534
cor=0.9, p=1.9e−13
5734: EIF5A|P63241
864: EIF5A|P63241
cor=0.89, p=8.5e−13
5736: HINT1|P49773
1245: HINT1|P49773
cor=0.72, p=1.1e−06
5737: HSPA8|P11142
254: HSPA8|P11142
cor=0.81, p=1.8e−07
5740: PSME1|Q06323
1011: PSME1|Q06323
cor=0.55, p=0.012
5764: CRHBP|P24387
1129: CRHBP|P24387
cor=0.23, p=0.25
5769: PIP|P12273
844: PIP|P12273
cor=0.65, p=2.4e−05
5770: ERP44|Q9BS26
796: ERP44|Q9BS26
cor=0.19, p=0.33
5777: FCRL5|Q96RD9
1191: FCRL5|Q96RD9
cor=0.56, p=0.00047
5787: PSAP|P07602
215: PSAP|P07602
cor=0.38, p=0.024
5792: QSOX1|O00391
220: QSOX1|O00391
cor=0.32, p=0.29
5802: STC2|O76061
787: STC2|O76061
cor=0.35, p=0.13
5812: SCGB3A1|Q96QR1
1285: SCGB3A1|Q96QR1
cor=0.2, p=0.25
5834: OIT3|Q8WWZ8
855: OIT3|Q8WWZ8
cor=0.39, p=0.17
5851: ECRG4|Q9H1Z8
1003: ECRG4|Q9H1Z8
cor=0.37, p=0.029
5866: MANBA|O00462
250: MANBA|O00462
cor=−0.013, p=0.96
5869: VWA1|Q6PCB0
815: VWA1|Q6PCB0
cor=0.24, p=0.22
5871: CCDC126|Q96EE4
984: CCDC126|Q96EE4
cor=0.82, p=1.7e−09
5873: LILRA3|Q8N6C8
834: LILRA3|Q8N6C8
cor=0.78, p=3.3e−08
5875: HSP90B1|P14625
273: HSP90B1|P14625
cor=−0.034, p=0.85
5886: ADGRF5|Q8IZF2
785: ADGRF5|Q8IZF2
cor=0.47, p=0.0044
5890: OAF|Q86UD1
660: OAF|Q86UD1
cor=0.69, p=4.5e−06
5891: CPN2|P22792
201: CPN2|P22792
cor=0.79, p=1.7e−08
5899: PCYOX1|Q9UHG3
314: PCYOX1|Q9UHG3
cor=−0.45, p=0.12
5908: SEMA3C|Q99985
718: SEMA3C|Q99985
cor=0.69, p=4.9e−05
5912: ARHGDIA|P52565
951: ARHGDIA|P52565
cor=0.61, p=1e−04
5917: APOC3|P02656
565: APOC3|P02656
cor=0.71, p=1.8e−06
5925: CFHR4|Q92496
522: CFHR4|Q92496

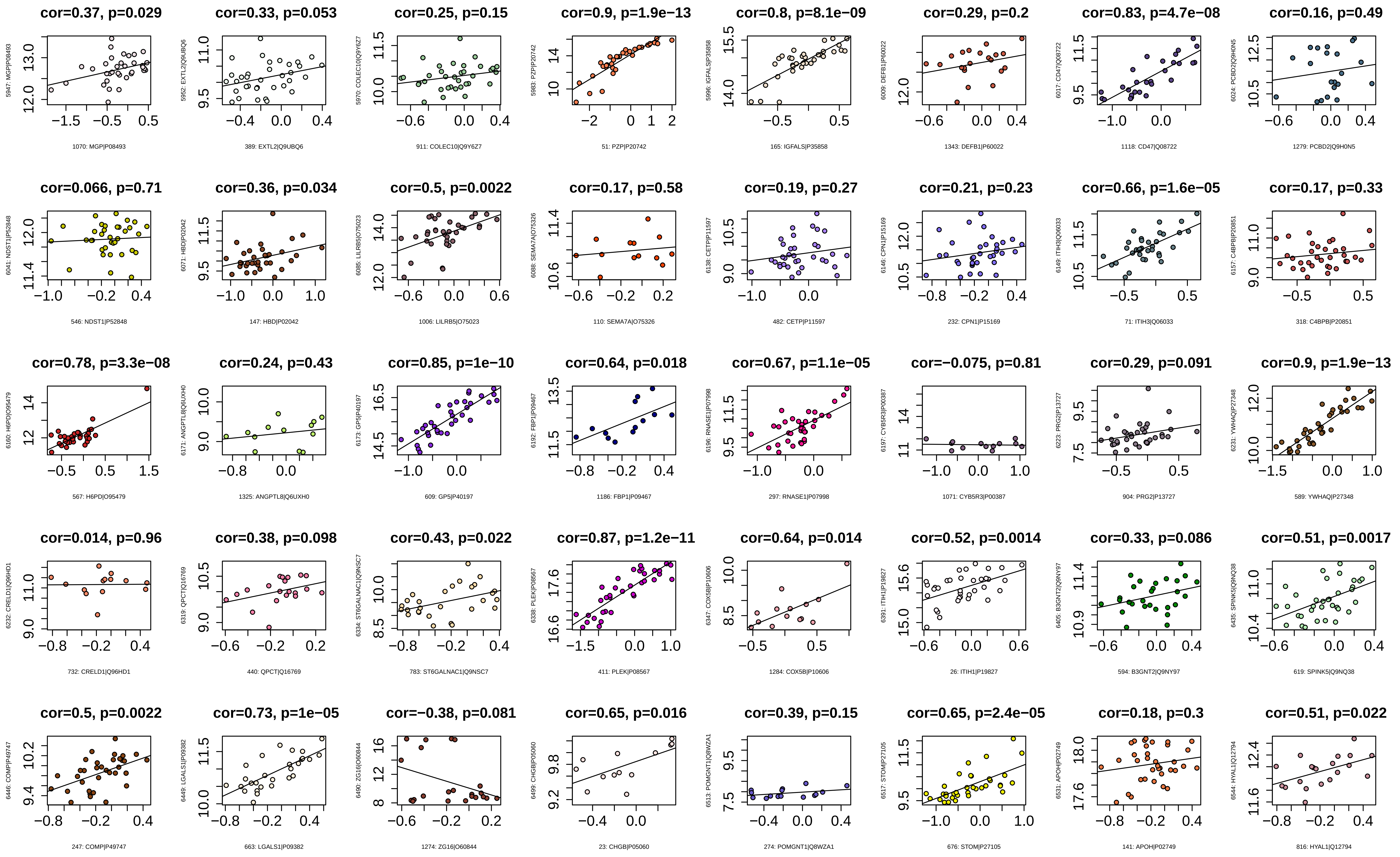
cor=0.37, p=0.029
1070: MGP|P08493
cor=0.33, p=0.053
389: EXTL2|Q9UBQ6
cor=0.25, p=0.15
911: COLEC10|Q9Y6Z7
cor=0.9, p=1.9e−13
51: PZP|P20742
cor=0.8, p=8.1e−09
165: IGFALS|P35858
cor=0.29, p=0.2
1343: DEFB1|P60022
cor=0.83, p=4.7e−08
1118: CD47|Q08722
cor=0.16, p=0.49
1279: PCBD2|Q9H0N5
cor=0.066, p=0.71
546: NDST1|P52848
cor=0.36, p=0.034
147: HBD|P02042
cor=0.5, p=0.0022
1006: LILRB5|O75023
cor=0.17, p=0.58
110: SEMA7A|O75326
cor=0.19, p=0.27
482: CETP|P11597
cor=0.21, p=0.23
232: CPN1|P15169
cor=0.66, p=1.6e−05
71: ITIH3|Q06033
cor=0.17, p=0.33
318: C4BPB|P20851
cor=0.78, p=3.3e−08
567: H6PD|O95479
cor=0.24, p=0.43
1325: ANGPTL8|Q6UXH0
cor=0.85, p=1e−10
609: GP5|P40197
cor=0.64, p=0.018
1186: FBP1|P09467
cor=0.67, p=1.1e−05
297: RNASE1|P07998
cor=−0.075, p=0.81
1071: CYB5R3|P00387
cor=0.29, p=0.091
904: PRG2|P13727
cor=0.9, p=1.9e−13
589: YWHAQ|P27348
cor=0.014, p=0.96
732: CRELD1|Q96HD1
cor=0.38, p=0.098
440: QPCT|Q16769
cor=0.43, p=0.022
783: ST6GALNAC1|Q9NSC7
cor=0.87, p=1.2e−11
411: PLEK|P08567
cor=0.64, p=0.014
1284: COX5B|P10606
cor=0.52, p=0.0014
26: ITIH1|P19827
cor=0.33, p=0.086
594: B3GNT2|Q9NY97
cor=0.51, p=0.0017
619: SPINK5|Q9NQ38
cor=0.5, p=0.0022
247: COMP|P49747
cor=0.73, p=1e−05
663: LGALS1|P09382
cor=−0.38, p=0.081
1274: ZG16|O60844
cor=0.65, p=0.016
23: CHGB|P05060
cor=0.39, p=0.15
274: POMGNT1|Q8WZA1
cor=0.65, p=2.4e−05
676: STOM|P27105
cor=0.18, p=0.3
141: APOH|P02749
cor=0.51, p=0.022
816: HYAL1|Q12794

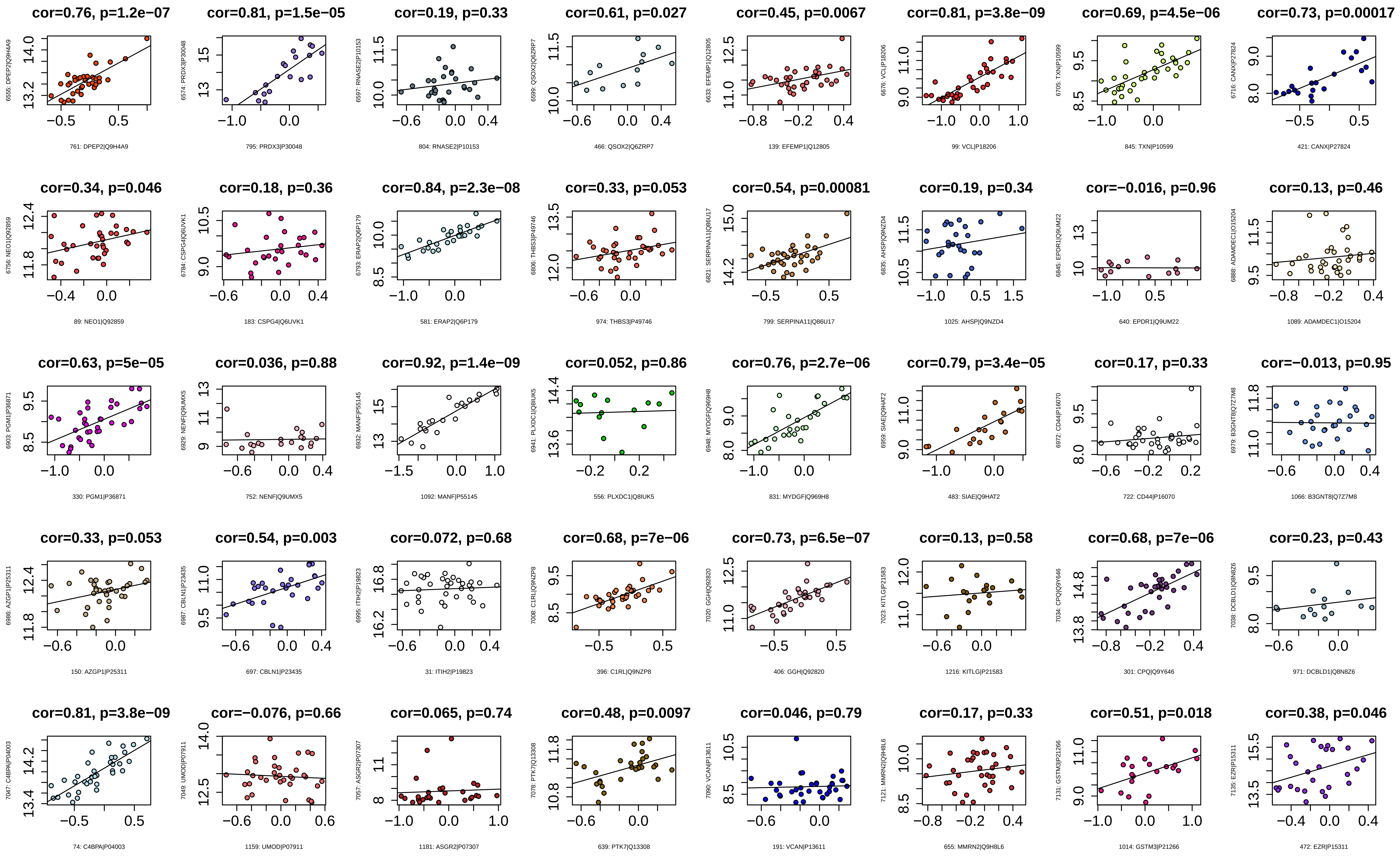
cor=0.76, p=1.2e−07
6555: DPEP2|Q9H4A9
761: DPEP2|Q9H4A9
cor=0.81, p=1.5e−05
6574: PRDX3|P30048
795: PRDX3|P30048
cor=0.19, p=0.33
6597: RNASE2|P10153
804: RNASE2|P10153
cor=0.61, p=0.027
6599: QSOX2|Q6ZRP7
466: QSOX2|Q6ZRP7
cor=0.45, p=0.0067
6633: EFEMP1|Q12805
139: EFEMP1|Q12805
cor=0.81, p=3.8e−09
6676: VCL|P18206
99: VCL|P18206
cor=0.69, p=4.5e−06
6705: TXN|P10599
845: TXN|P10599
cor=0.73, p=0.00017
6716: CANX|P27824
421: CANX|P27824
cor=0.34, p=0.046
6756: NEO1|Q92859
89: NEO1|Q92859
cor=0.18, p=0.36
6784: CSPG4|Q6UVK1
183: CSPG4|Q6UVK1
cor=0.84, p=2.3e−08
6793: ERAP2|Q6P179
581: ERAP2|Q6P179
cor=0.33, p=0.053
6806: THBS3|P49746
974: THBS3|P49746
cor=0.54, p=0.00081
6821: SERPINA11|Q86U17
799: SERPINA11|Q86U17
cor=0.19, p=0.34
6835: AHSP|Q9NZD4
1025: AHSP|Q9NZD4
cor=−0.016, p=0.96
6845: EPDR1|Q9UM22
640: EPDR1|Q9UM22
cor=0.13, p=0.46
6888: ADAMDEC1|O15204
1089: ADAMDEC1|O15204
cor=0.63, p=5e−05
6903: PGM1|P36871
330: PGM1|P36871
cor=0.036, p=0.88
6929: NENF|Q9UMX5
752: NENF|Q9UMX5
cor=0.92, p=1.4e−09
6932: MANF|P55145
1092: MANF|P55145
cor=0.052, p=0.86
6941: PLXDC1|Q8IUK5
556: PLXDC1|Q8IUK5
cor=0.76, p=2.7e−06
6948: MYDGF|Q969H8
831: MYDGF|Q969H8
cor=0.79, p=3.4e−05
6959: SIAE|Q9HAT2
483: SIAE|Q9HAT2
cor=0.17, p=0.33
6972: CD44|P16070
722: CD44|P16070
cor=−0.013, p=0.95
6979: B3GNT8|Q7Z7M8
1066: B3GNT8|Q7Z7M8
cor=0.33, p=0.053
6986: AZGP1|P25311
150: AZGP1|P25311
cor=0.54, p=0.003
6987: CBLN1|P23435
697: CBLN1|P23435
cor=0.072, p=0.68
6995: ITIH2|P19823
31: ITIH2|P19823
cor=0.68, p=7e−06
7008: C1RL|Q9NZP8
396: C1RL|Q9NZP8
cor=0.73, p=6.5e−07
7020: GGH|Q92820
406: GGH|Q92820
cor=0.13, p=0.58
7023: KITLG|P21583
1216: KITLG|P21583
cor=0.68, p=7e−06
7034: CPQ|Q9Y646
301: CPQ|Q9Y646
cor=0.23, p=0.43
7038: DCBLD1|Q8N8Z6
971: DCBLD1|Q8N8Z6
cor=0.81, p=3.8e−09
7047: C4BPA|P04003
74: C4BPA|P04003
cor=−0.076, p=0.66
7049: UMOD|P07911
1159: UMOD|P07911
cor=0.065, p=0.74
7057: ASGR2|P07307
1181: ASGR2|P07307
cor=0.48, p=0.0097
7078: PTK7|Q13308
639: PTK7|Q13308
cor=0.046, p=0.79
7090: VCAN|P13611
191: VCAN|P13611
cor=0.17, p=0.33
7121: MMRN2|Q9H8L6
655: MMRN2|Q9H8L6
cor=0.51, p=0.018
7131: GSTM3|P21266
1014: GSTM3|P21266
cor=0.38, p=0.046
7135: EZR|P15311
472: EZR|P15311

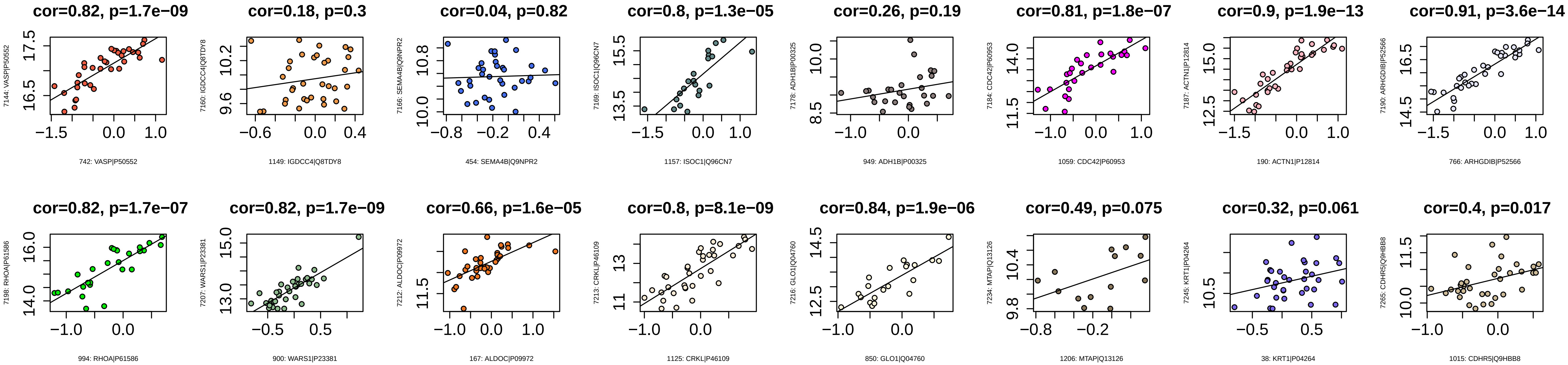

cor=0.82, p=1.7e−09
742: VASP|P50552
7144: VASP|P50552
cor=0.18, p=0.3
1149: IGDCC4|Q8TDY8
7160: IGDCC4|Q8TDY8
cor=0.04, p=0.82
454: SEMA4B|Q9NPR2
7166: SEMA4B|Q9NPR2
cor=0.8, p=1.3e−05
1157: ISOC1|Q96CN7
7169: ISOC1|Q96CN7
cor=0.26, p=0.19
949: ADH1B|P00325
7178: ADH1B|P00325
cor=0.81, p=1.8e−07
1059: CDC42|P60953
7184: CDC42|P60953
cor=0.9, p=1.9e−13
190: ACTN1|P12814
7187: ACTN1|P12814
cor=0.91, p=3.6e−14
766: ARHGDIB|P52566
7190: ARHGDIB|P52566
cor=0.82, p=1.7e−07
994: RHOA|P61586
7198: RHOA|P61586
cor=0.82, p=1.7e−09
900: WARS1|P23381
7207: WARS1|P23381
cor=0.66, p=1.6e−05
167: ALDOC|P09972
7212: ALDOC|P09972
cor=0.8, p=8.1e−09
1125: CRKL|P46109
7213: CRKL|P46109
cor=0.84, p=1.9e−06
850: GLO1|Q04760
7216: GLO1|Q04760
cor=0.49, p=0.075
1206: MTAP|Q13126
7234: MTAP|Q13126
cor=0.32, p=0.061
38: KRT1|P04264
7245: KRT1|P04264
cor=0.4, p=0.017
1015: CDHR5|Q9HBB8
7265: CDHR5|Q9HBB8

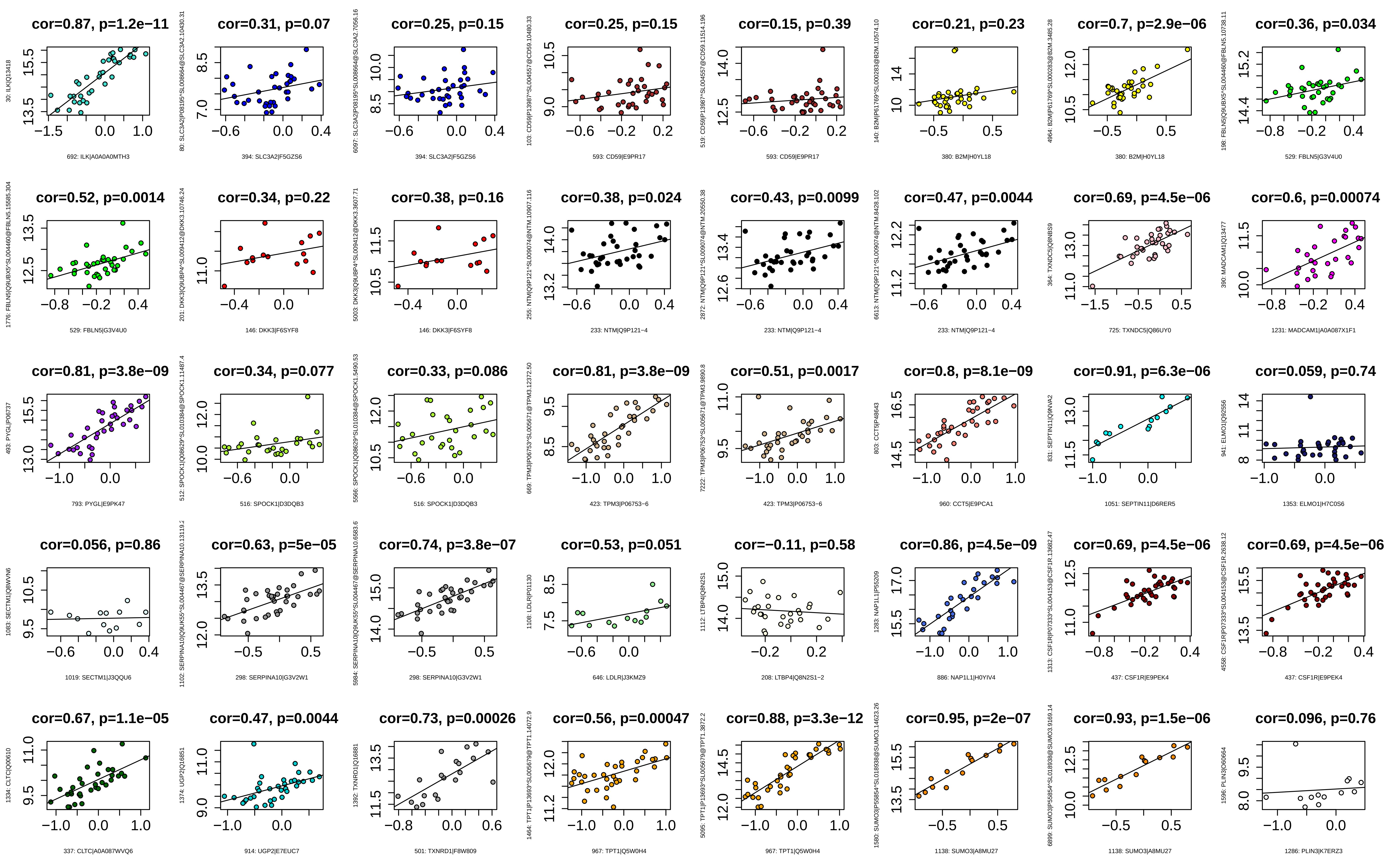
cor=0.87, p=1.2e−11
692: ILK|A0A0A0MTH3
cor=0.31, p=0.07
394: SLC3A2|F5GZS6
cor=0.25, p=0.15
394: SLC3A2|F5GZS6
cor=0.25, p=0.15
593: CD59|E9PR17
cor=0.15, p=0.39
593: CD59|E9PR17
cor=0.21, p=0.23
380: B2M|H0YL18
cor=0.7, p=2.9e−06
380: B2M|H0YL18
cor=0.36, p=0.034
529: FBLN5|G3V4U0
cor=0.52, p=0.0014
529: FBLN5|G3V4U0
cor=0.34, p=0.22
146: DKK3|F6SYF8
cor=0.38, p=0.16
146: DKK3|F6SYF8
cor=0.38, p=0.024
233: NTM|Q9P121−4
cor=0.43, p=0.0099
233: NTM|Q9P121−4
cor=0.47, p=0.0044
233: NTM|Q9P121−4
cor=0.69, p=4.5e−06
725: TXNDC5|Q86UY0
cor=0.6, p=0.00074
1231: MADCAM1|A0A087X1F1
cor=0.81, p=3.8e−09
793: PYGL|E9PK47
cor=0.34, p=0.077
516: SPOCK1|D3DQB3
cor=0.33, p=0.086
516: SPOCK1|D3DQB3
cor=0.81, p=3.8e−09
423: TPM3|P06753−6
cor=0.51, p=0.0017
423: TPM3|P06753−6
cor=0.8, p=8.1e−09
960: CCT5|E9PCA1
cor=0.91, p=6.3e−06
1051: SEPTIN11|D6RER5
cor=0.059, p=0.74
1353: ELMO1|H7C0S6
cor=0.056, p=0.86
1019: SECTM1|J3QQU6
cor=0.63, p=5e−05
298: SERPINA10|G3V2W1
cor=0.74, p=3.8e−07
298: SERPINA10|G3V2W1
cor=0.53, p=0.051
646: LDLR|J3KMZ9
cor=−0.11, p=0.58
208: LTBP4|Q8N2S1−2
cor=0.86, p=4.5e−09
886: NAP1L1|H0YIV4
cor=0.69, p=4.5e−06
437: CSF1R|E9PEK4
cor=0.69, p=4.5e−06
437: CSF1R|E9PEK4
cor=0.67, p=1.1e−05
337: CLTC|A0A087WVQ6
cor=0.47, p=0.0044
914: UGP2|E7EUC7
cor=0.73, p=0.00026
501: TXNRD1|F8W809
cor=0.56, p=0.00047
967: TPT1|Q5W0H4
cor=0.88, p=3.3e−12
967: TPT1|Q5W0H4
cor=0.95, p=2e−07
1138: SUMO3|A8MU27
cor=0.93, p=1.5e−06
1138: SUMO3|A8MU27
cor=0.096, p=0.76
1286: PLIN3|K7ERZ3

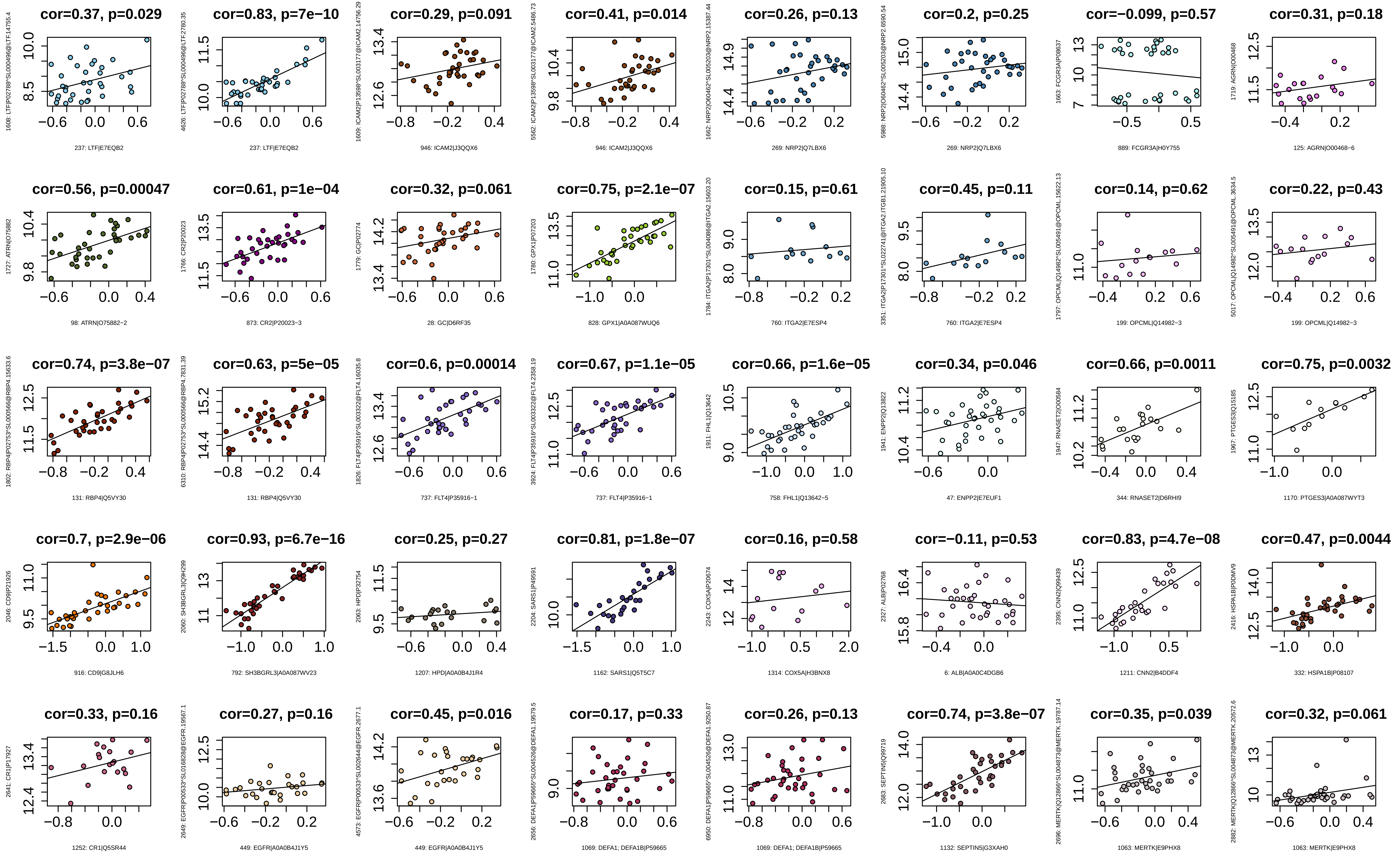

cor=0.37, p=0.029
237: LTF|E7EQB2
cor=0.83, p=7e−10
237: LTF|E7EQB2
cor=0.29, p=0.091
946: ICAM2|J3QQX6
cor=0.41, p=0.014
946: ICAM2|J3QQX6
cor=0.26, p=0.13
269: NRP2|Q7LBX6
cor=0.2, p=0.25
269: NRP2|Q7LBX6
cor=−0.099, p=0.57
889: FCGR3A|H0Y755
cor=0.31, p=0.18
125: AGRN|O00468−6
cor=0.56, p=0.00047
98: ATRN|O75882−2
cor=0.61, p=1e−04
873: CR2|P20023−3
cor=0.32, p=0.061
28: GC|D6RF35
cor=0.75, p=2.1e−07
828: GPX1|A0A087WUQ6
cor=0.15, p=0.61
760: ITGA2|E7ESP4
cor=0.45, p=0.11
760: ITGA2|E7ESP4
cor=0.14, p=0.62
199: OPCML|Q14982−3
cor=0.22, p=0.43
199: OPCML|Q14982−3
cor=0.74, p=3.8e−07
131: RBP4|Q5VY30
cor=0.63, p=5e−05
131: RBP4|Q5VY30
cor=0.6, p=0.00014
737: FLT4|P35916−1
cor=0.67, p=1.1e−05
737: FLT4|P35916−1
cor=0.66, p=1.6e−05
758: FHL1|Q13642−5
cor=0.34, p=0.046
47: ENPP2|E7EUF1
cor=0.66, p=0.0011
344: RNASET2|D6RHI9
cor=0.75, p=0.0032
1170: PTGES3|A0A087WYT3
cor=0.7, p=2.9e−06
916: CD9|G8JLH6
cor=0.93, p=6.7e−16
792: SH3BGRL3|A0A087WV23
cor=0.25, p=0.27
1207: HPD|A0A0B4J1R4
cor=0.81, p=1.8e−07
1162: SARS1|Q5T5C7
cor=0.16, p=0.58
1314: COX5A|H3BNX8
cor=−0.11, p=0.53
6: ALB|A0A0C4DGB6
cor=0.83, p=4.7e−08
1211: CNN2|B4DDF4
cor=0.47, p=0.0044
332: HSPA1B|P08107
cor=0.33, p=0.16
1252: CR1|Q5SR44
cor=0.27, p=0.16
449: EGFR|A0A0B4J1Y5
cor=0.45, p=0.016
449: EGFR|A0A0B4J1Y5
cor=0.17, p=0.33
1069: DEFA1; DEFA1B|P59665
cor=0.26, p=0.13
1069: DEFA1; DEFA1B|P59665
cor=0.74, p=3.8e−07
1132: SEPTIN5|G3XAH0
cor=0.35, p=0.039
1063: MERTK|E9PHX8
cor=0.32, p=0.061
1063: MERTK|E9PHX8

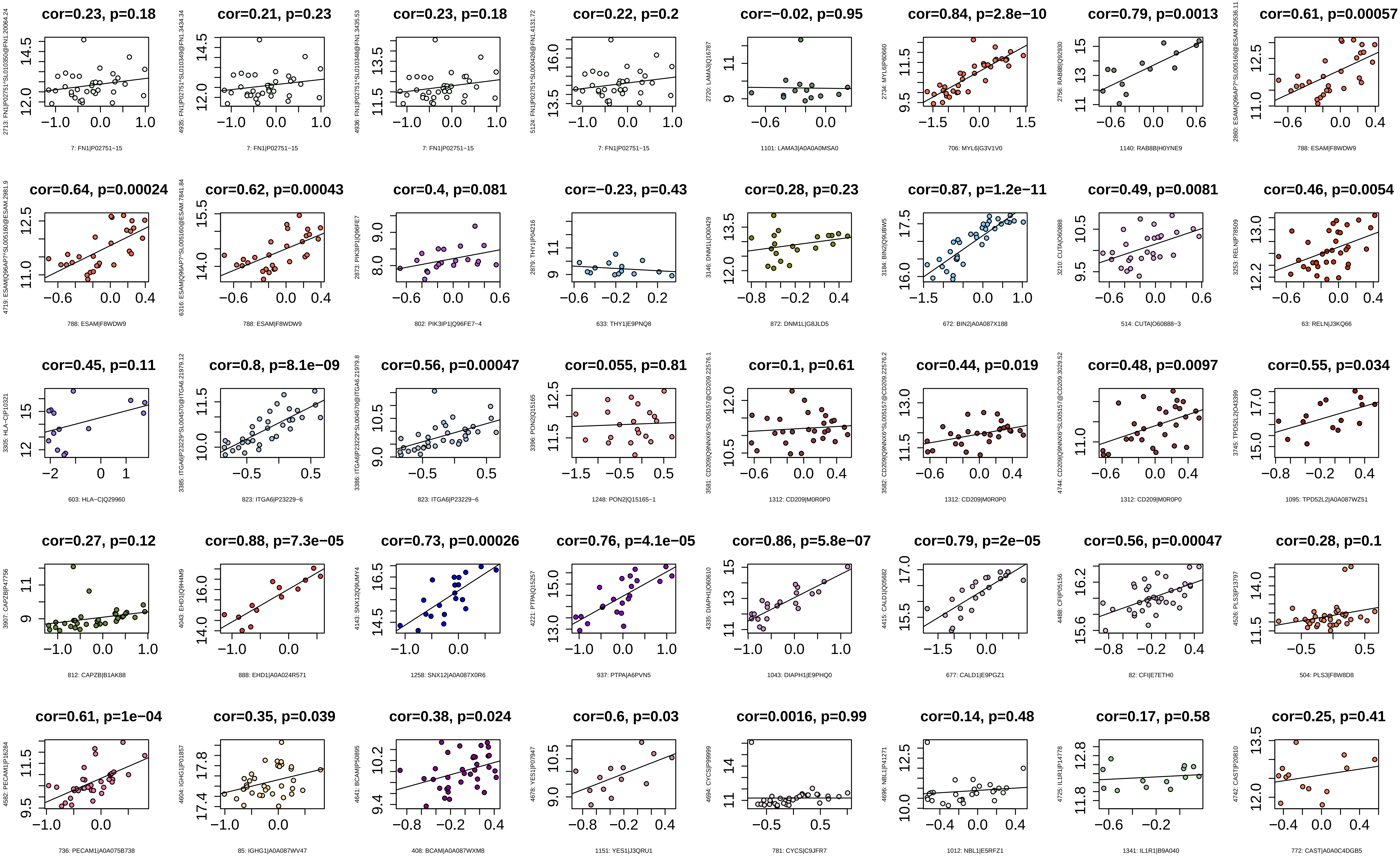
cor=0.23, p=0.18
2713: FN1|P02751^SL010350@FN1.20064.24
7: FN1|P02751−15
cor=0.21, p=0.23
4935: FN1|P02751^SL010349@FN1.3434.34
7: FN1|P02751−15
cor=0.23, p=0.18
4936: FN1|P02751^SL010348@FN1.3435.53
7: FN1|P02751−15
cor=0.22, p=0.2
5124: FN1|P02751^SL000426@FN1.4131.72
7: FN1|P02751−15
cor=−0.02, p=0.95
2720: LAMA3|Q16787
1101: LAMA3|A0A0A0MSA0
cor=0.84, p=2.8e−10
2734: MYL6|P60660
706: MYL6|G3V1V0
cor=0.79, p=0.0013
2756: RAB8B|Q92930
1140: RAB8B|H0YNE9
cor=0.61, p=0.00057
2860: ESAM|Q96AP7^SL005160@ESAM.20536.11
788: ESAM|F8WDW9
cor=0.64, p=0.00024
4719: ESAM|Q96AP7^SL005160@ESAM.2981.9
788: ESAM|F8WDW9
cor=0.62, p=0.00043
6316: ESAM|Q96AP7^SL005160@ESAM.7841.84
788: ESAM|F8WDW9
cor=0.4, p=0.081
2873: PIK3IP1|Q96FE7
802: PIK3IP1|Q96FE7−4
cor=−0.23, p=0.43
2879: THY1|P04216
633: THY1|E9PNQ8
cor=0.28, p=0.23
3146: DNM1L|O00429
872: DNM1L|G8JLD5
cor=0.87, p=1.2e−11
3184: BIN2|Q9UBW5
672: BIN2|A0A087X188
cor=0.49, p=0.0081
3210: CUTA|O60888
514: CUTA|O60888−3
cor=0.46, p=0.0054
3253: RELN|P78509
63: RELN|J3KQ66
cor=0.45, p=0.11
3305: HLA−C|P10321
603: HLA−C|Q29960
cor=0.8, p=8.1e−09
3385: ITGA6|P23229^SL004570@ITGA6.21979.12
823: ITGA6|P23229−6
cor=0.56, p=0.00047
3386: ITGA6|P23229^SL004570@ITGA6.21979.8
823: ITGA6|P23229−6
cor=0.055, p=0.81
3396: PON2|Q15165
1248: PON2|Q15165−1
cor=0.1, p=0.61
3581: CD209|Q9NNX6^SL005157@CD209.22576.1
1312: CD209|M0R0P0
cor=0.44, p=0.019
3582: CD209|Q9NNX6^SL005157@CD209.22576.2
1312: CD209|M0R0P0
cor=0.48, p=0.0097
4744: CD209|Q9NNX6^SL005157@CD209.3029.52
1312: CD209|M0R0P0
cor=0.55, p=0.034
3745: TPD52L2|O43399
1095: TPD52L2|A0A087WZ51
cor=0.27, p=0.12
3907: CAPZB|P47756
812: CAPZB|B1AK88
cor=0.88, p=7.3e−05
4043: EHD1|Q9H4M9
888: EHD1|A0A024R571
cor=0.73, p=0.00026
4143: SNX12|Q9UMY4
1258: SNX12|A0A087X0R6
cor=0.76, p=4.1e−05
4221: PTPA|Q15257
937: PTPA|A6PVN5
cor=0.86, p=5.8e−07
4335: DIAPH1|O60610
1043: DIAPH1|E9PHQ0
cor=0.79, p=2e−05
4415: CALD1|Q05682
677: CALD1|E9PGZ1
cor=0.56, p=0.00047
4488: CFI|P05156
82: CFI|E7ETH0
cor=0.28, p=0.1
4526: PLS3|P13797
504: PLS3|F8W8D8
cor=0.61, p=1e−04
4582: PECAM1|P16284
736: PECAM1|A0A075B738
cor=0.35, p=0.039
4604: IGHG1|P01857
85: IGHG1|A0A087WV47
cor=0.38, p=0.024
4641: BCAM|P50895
408: BCAM|A0A087WXM8
cor=0.6, p=0.03
4678: YES1|P07947
1151: YES1|J3QRU1
cor=0.0016, p=0.99
4694: CYCS|P99999
781: CYCS|C9JFR7
cor=0.14, p=0.48
4696: NBL1|P41271
1012: NBL1|E5RFZ1
cor=0.17, p=0.58
4725: IL1R1|P14778
1341: IL1R1|B9A040
cor=0.25, p=0.41
4742: CAST|P20810
772: CAST|A0A0C4DGB5

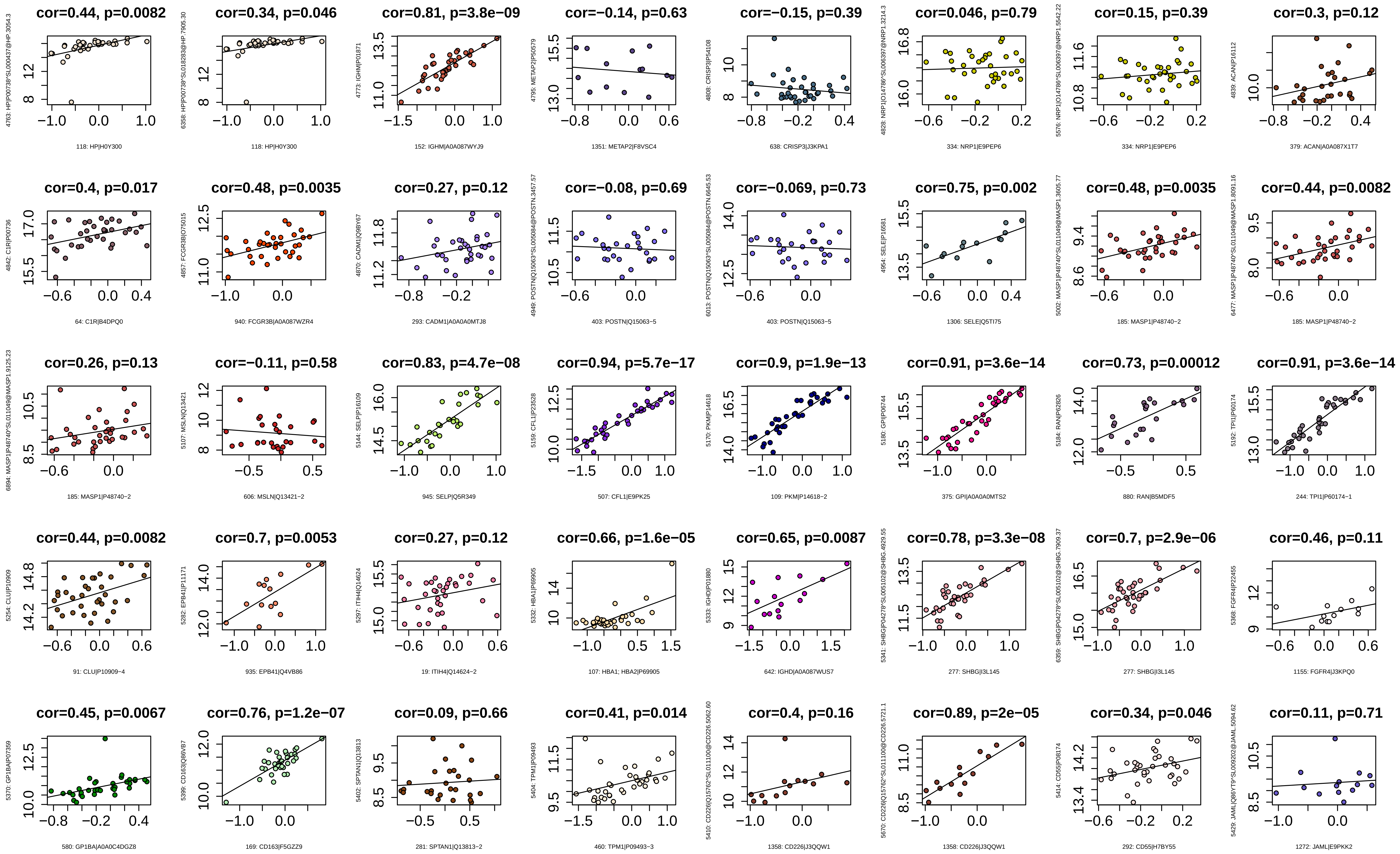
cor=0.44, p=0.0082
4763: HP|P00738^SL000437@HP.3054.3
118: HP|H0Y300
cor=0.34, p=0.046
6358: HP|P00738^SL018283@HP.7905.30
118: HP|H0Y300
cor=0.81, p=3.8e−09
4773: IGHM|P01871
152: IGHM|A0A087WYJ9
cor=−0.14, p=0.63
4795: METAP2|P50579
1351: METAP2|F8VSC4
cor=−0.15, p=0.39
4808: CRISP3|P54108
638: CRISP3|J3KPA1
cor=0.046, p=0.79
4828: NRP1|O14786^SL006397@NRP1.3214.3
334: NRP1|E9PEP6
cor=0.15, p=0.39
5576: NRP1|O14786^SL006397@NRP1.5542.22
334: NRP1|E9PEP6
cor=0.3, p=0.12
4839: ACAN|P16112
379: ACAN|A0A087X1T7
cor=0.4, p=0.017
4842: C1R|P00736
64: C1R|B4DPQ0
cor=0.48, p=0.0035
4857: FCGR3B|O75015
940: FCGR3B|A0A087WZR4
cor=0.27, p=0.12
4870: CADM1|Q9BY67
293: CADM1|A0A0A0MTJ8
cor=−0.08, p=0.69
4949: POSTN|Q15063^SL005084@POSTN.3457.57
403: POSTN|Q15063−5
cor=−0.069, p=0.73
6013: POSTN|Q15063^SL005084@POSTN.6645.53
403: POSTN|Q15063−5
cor=0.75, p=0.002
4954: SELE|P16581
1306: SELE|Q5TI75
cor=0.48, p=0.0035
5002: MASP1|P48740^SL011049@MASP1.3605.77
185: MASP1|P48740−2
cor=0.44, p=0.0082
6477: MASP1|P48740^SL011049@MASP1.8091.16
185: MASP1|P48740−2
cor=0.26, p=0.13
6894: MASP1|P48740^SL011049@MASP1.9125.23
185: MASP1|P48740−2
cor=−0.11, p=0.58
5107: MSLN|Q13421
606: MSLN|Q13421−2
cor=0.83, p=4.7e−08
5144: SELP|P16109
945: SELP|Q5R349
cor=0.94, p=5.7e−17
5159: CFL1|P23528
507: CFL1|E9PK25
cor=0.9, p=1.9e−13
5170: PKM|P14618
109: PKM|P14618−2
cor=0.91, p=3.6e−14
5180: GPI|P06744
375: GPI|A0A0A0MTS2
cor=0.73, p=0.00012
5184: RAN|P62826
880: RAN|B5MDF5
cor=0.91, p=3.6e−14
5192: TPI1|P60174
244: TPI1|P60174−1
cor=0.44, p=0.0082
5254: CLU|P10909
91: CLU|P10909−4
cor=0.7, p=0.0053
5282: EPB41|P11171
935: EPB41|Q4VB86
cor=0.27, p=0.12
5297: ITIH4|Q14624
19: ITIH4|Q14624−2
cor=0.66, p=1.6e−05
5332: HBA1|P69905
107: HBA1; HBA2|P69905
cor=0.65, p=0.0087
5333: IGHD|P01880
642: IGHD|A0A087WUS7
cor=0.78, p=3.3e−08
5341: SHBG|P04278^SL005102@SHBG.4929.55
277: SHBG|I3L145
cor=0.7, p=2.9e−06
6359: SHBG|P04278^SL005102@SHBG.7909.37
277: SHBG|I3L145
cor=0.46, p=0.11
5368: FGFR4|P22455
1155: FGFR4|J3KPQ0
cor=0.45, p=0.0067
5370: GP1BA|P07359
580: GP1BA|A0A0C4DGZ8
cor=0.76, p=1.2e−07
5399: CD163|Q86VB7
169: CD163|F5GZZ9
cor=0.09, p=0.66
5402: SPTAN1|Q13813
281: SPTAN1|Q13813−2
cor=0.41, p=0.014
5404: TPM1|P09493
460: TPM1|P09493−3
cor=0.4, p=0.16
5410: CD226|Q15762^SL011100@CD226.5062.60
1358: CD226|J3QQW1
cor=0.89, p=2e−05
5670: CD226|Q15762^SL011100@CD226.5721.1
1358: CD226|J3QQW1
cor=0.34, p=0.046
5414: CD55|P08174
292: CD55|H7BY55
cor=0.11, p=0.71
5429: JAML|Q86YT9^SL009202@JAML.5094.62
1272: JAML|E9PKK2

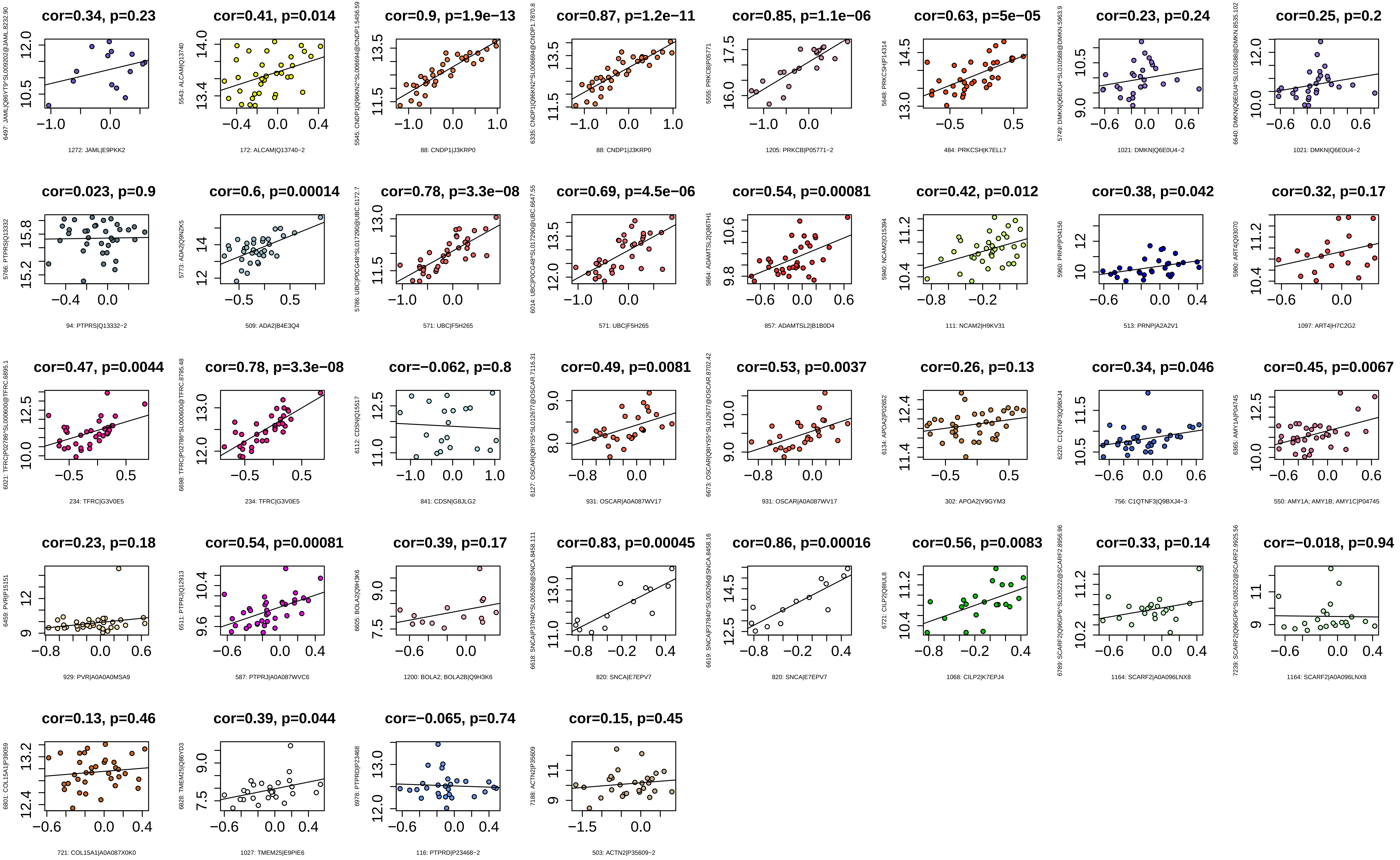
cor=0.34, p=0.23
1272: JAML|E9PKK2
cor=0.41, p=0.014
172: ALCAM|Q13740−2
cor=0.9, p=1.9e−13
88: CNDP1|J3KRP0
cor=0.87, p=1.2e−11
88: CNDP1|J3KRP0
cor=0.85, p=1.1e−06
1205: PRKCB|P05771−2
cor=0.63, p=5e−05
484: PRKCSH|K7ELL7
cor=0.23, p=0.24
1021: DMKN|Q6E0U4−2
cor=0.25, p=0.2
1021: DMKN|Q6E0U4−2
cor=0.023, p=0.9
94: PTPRS|Q13332−2
cor=0.6, p=0.00014
509: ADA2|B4E3Q4
cor=0.78, p=3.3e−08
571: UBC|F5H265
cor=0.69, p=4.5e−06
571: UBC|F5H265
cor=0.54, p=0.00081
857: ADAMTSL2|B1B0D4
cor=0.42, p=0.012
111: NCAM2|H9KV31
cor=0.38, p=0.042
513: PRNP|A2A2V1
cor=0.32, p=0.17
1097: ART4|H7C2G2
cor=0.47, p=0.0044
234: TFRC|G3V0E5
cor=0.78, p=3.3e−08
234: TFRC|G3V0E5
cor=−0.062, p=0.8
841: CDSN|G8JLG2
cor=0.49, p=0.0081
931: OSCAR|A0A087WV17
cor=0.53, p=0.0037
931: OSCAR|A0A087WV17
cor=0.26, p=0.13
302: APOA2|V9GYM3
cor=0.34, p=0.046
756: C1QTNF3|Q9BXJ4−3
cor=0.45, p=0.0067
550: AMY1A; AMY1B; AMY1C|P04745
cor=0.23, p=0.18
929: PVR|A0A0A0MSA9
cor=0.54, p=0.00081
587: PTPRJ|A0A087WVC6
cor=0.39, p=0.17
1200: BOLA2; BOLA2B|Q9H3K6
cor=0.83, p=0.00045
820: SNCA|E7EPV7
cor=0.86, p=0.00016
820: SNCA|E7EPV7
cor=0.56, p=0.0083
1068: CILP2|K7EPJ4
cor=0.33, p=0.14
1164: SCARF2|A0A096LNX8
cor=−0.018, p=0.94
1164: SCARF2|A0A096LNX8
cor=0.13, p=0.46
721: COL15A1|A0A087X0K0
cor=0.39, p=0.044
1027: TMEM25|E9PIE6
cor=−0.065, p=0.74
116: PTPRD|P23468−2
cor=0.15, p=0.45
503: ACTN2|P35609−2

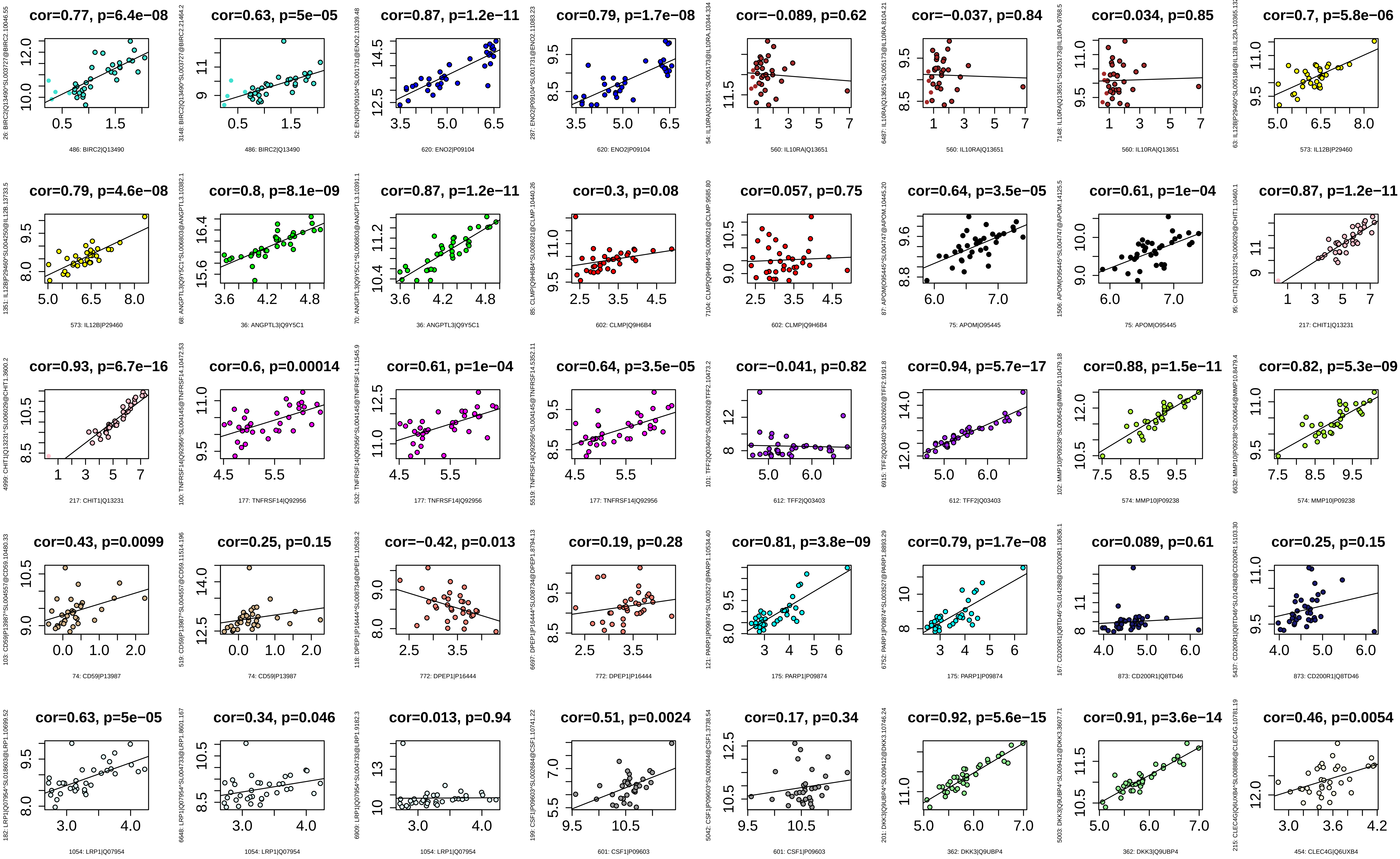
cor=0.77, p=6.4e−08
486: BIRC2|Q13490
cor=0.63, p=5e−05
486: BIRC2|Q13490
cor=0.87, p=1.2e−11
620: ENO2|P09104
cor=0.79, p=1.7e−08
620: ENO2|P09104
cor=−0.089, p=0.62
560: IL10RA|Q13651
cor=−0.037, p=0.84
560: IL10RA|Q13651
cor=0.034, p=0.85
560: IL10RA|Q13651
cor=0.7, p=5.8e−06
573: IL12B|P29460
cor=0.79, p=4.6e−08
573: IL12B|P29460
cor=0.8, p=8.1e−09
36: ANGPTL3|Q9Y5C1
cor=0.87, p=1.2e−11
36: ANGPTL3|Q9Y5C1
cor=0.3, p=0.08
602: CLMP|Q9H6B4
cor=0.057, p=0.75
602: CLMP|Q9H6B4
cor=0.64, p=3.5e−05
75: APOM|O95445
cor=0.61, p=1e−04
75: APOM|O95445
cor=0.87, p=1.2e−11
217: CHIT1|Q13231
cor=0.93, p=6.7e−16
217: CHIT1|Q13231
cor=0.6, p=0.00014
177: TNFRSF14|Q92956
cor=0.61, p=1e−04
177: TNFRSF14|Q92956
cor=0.64, p=3.5e−05
177: TNFRSF14|Q92956
cor=−0.041, p=0.82
612: TFF2|Q03403
cor=0.94, p=5.7e−17
612: TFF2|Q03403
cor=0.88, p=1.5e−11
574: MMP10|P09238
cor=0.82, p=5.3e−09
574: MMP10|P09238
cor=0.43, p=0.0099
74: CD59|P13987
cor=0.25, p=0.15
74: CD59|P13987
cor=−0.42, p=0.013
772: DPEP1|P16444
cor=0.19, p=0.28
772: DPEP1|P16444
cor=0.81, p=3.8e−09
175: PARP1|P09874
cor=0.79, p=1.7e−08
175: PARP1|P09874
cor=0.089, p=0.61
873: CD200R1|Q8TD46
cor=0.25, p=0.15
873: CD200R1|Q8TD46
cor=0.63, p=5e−05
1054: LRP1|Q07954
cor=0.34, p=0.046
1054: LRP1|Q07954
cor=0.013, p=0.94
1054: LRP1|Q07954
cor=0.51, p=0.0024
601: CSF1|P09603
cor=0.17, p=0.34
601: CSF1|P09603
cor=0.92, p=5.6e−15
362: DKK3|Q9UBP4
cor=0.91, p=3.6e−14
362: DKK3|Q9UBP4
cor=0.46, p=0.0054
454: CLEC4G|Q6UXB4

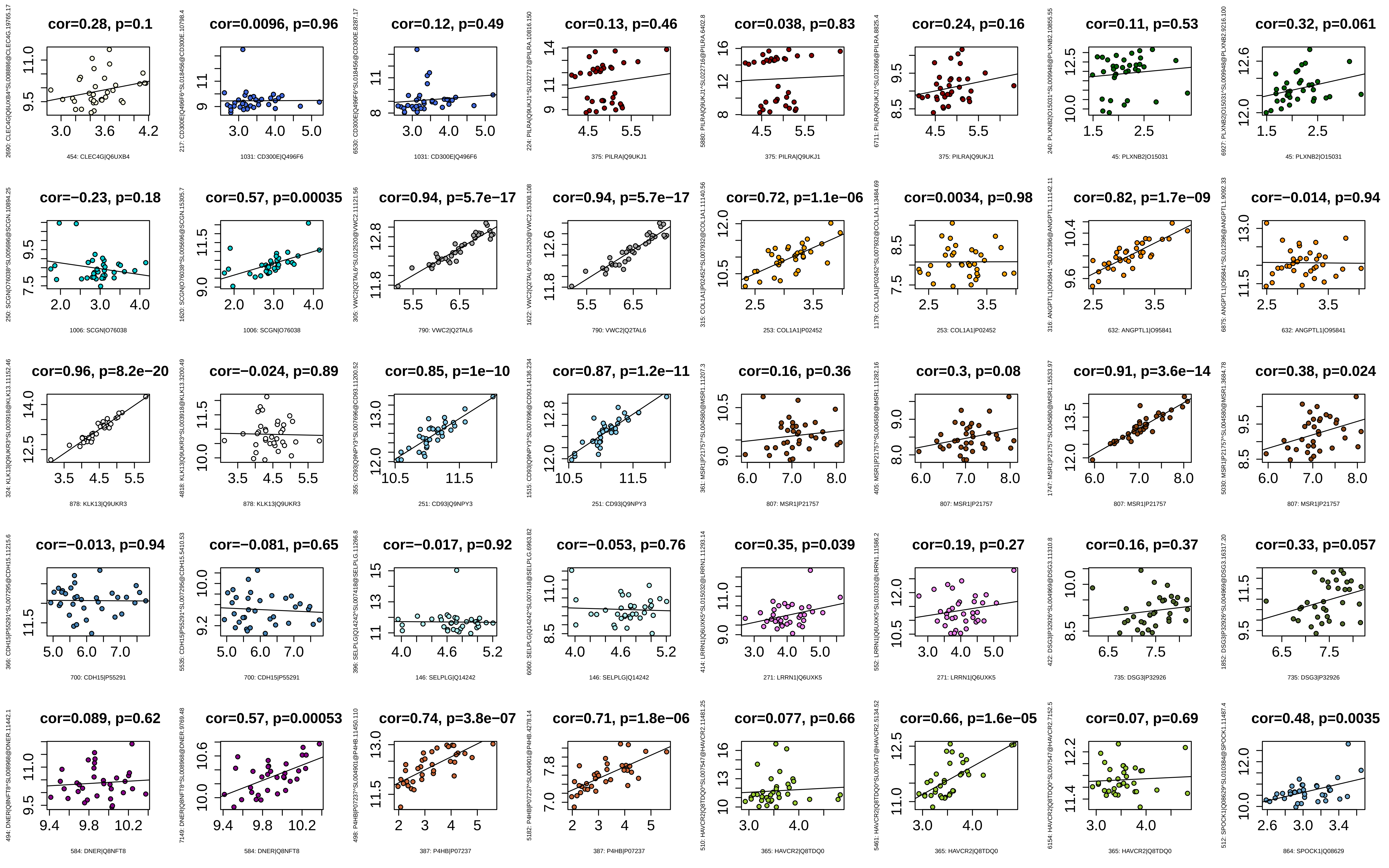
cor=0.28, p=0.1
454: CLEC4G|Q6UXB4
cor=0.0096, p=0.96
1031: CD300E|Q496F6
cor=0.12, p=0.49
1031: CD300E|Q496F6
cor=0.13, p=0.46
375: PILRA|Q9UKJ1
cor=0.038, p=0.83
375: PILRA|Q9UKJ1
cor=0.24, p=0.16
375: PILRA|Q9UKJ1
cor=0.11, p=0.53
45: PLXNB2|O15031
cor=0.32, p=0.061
45: PLXNB2|O15031
cor=−0.23, p=0.18
1006: SCGN|O76038
cor=0.57, p=0.00035
1006: SCGN|O76038
cor=0.94, p=5.7e−17
790: VWC2|Q2TAL6
cor=0.94, p=5.7e−17
790: VWC2|Q2TAL6
cor=0.72, p=1.1e−06
253: COL1A1|P02452
cor=0.0034, p=0.98
253: COL1A1|P02452
cor=0.82, p=1.7e−09
632: ANGPTL1|O95841
cor=−0.014, p=0.94
632: ANGPTL1|O95841
cor=0.96, p=8.2e−20
878: KLK13|Q9UKR3
cor=−0.024, p=0.89
878: KLK13|Q9UKR3
cor=0.85, p=1e−10
251: CD93|Q9NPY3
cor=0.87, p=1.2e−11
251: CD93|Q9NPY3
cor=0.16, p=0.36
807: MSR1|P21757
cor=0.3, p=0.08
807: MSR1|P21757
cor=0.91, p=3.6e−14
807: MSR1|P21757
cor=0.38, p=0.024
807: MSR1|P21757
cor=−0.013, p=0.94
700: CDH15|P55291
cor=−0.081, p=0.65
700: CDH15|P55291
cor=−0.017, p=0.92
146: SELPLG|Q14242
cor=−0.053, p=0.76
146: SELPLG|Q14242
cor=0.35, p=0.039
271: LRRN1|Q6UXK5
cor=0.19, p=0.27
271: LRRN1|Q6UXK5
cor=0.16, p=0.37
735: DSG3|P32926
cor=0.33, p=0.057
735: DSG3|P32926
cor=0.089, p=0.62
584: DNER|Q8NFT8
cor=0.57, p=0.00053
584: DNER|Q8NFT8
cor=0.74, p=3.8e−07
387: P4HB|P07237
cor=0.71, p=1.8e−06
387: P4HB|P07237
cor=0.077, p=0.66
365: HAVCR2|Q8TDQ0
cor=0.66, p=1.6e−05
365: HAVCR2|Q8TDQ0
cor=0.07, p=0.69
365: HAVCR2|Q8TDQ0
cor=0.48, p=0.0035
864: SPOCK1|Q08629

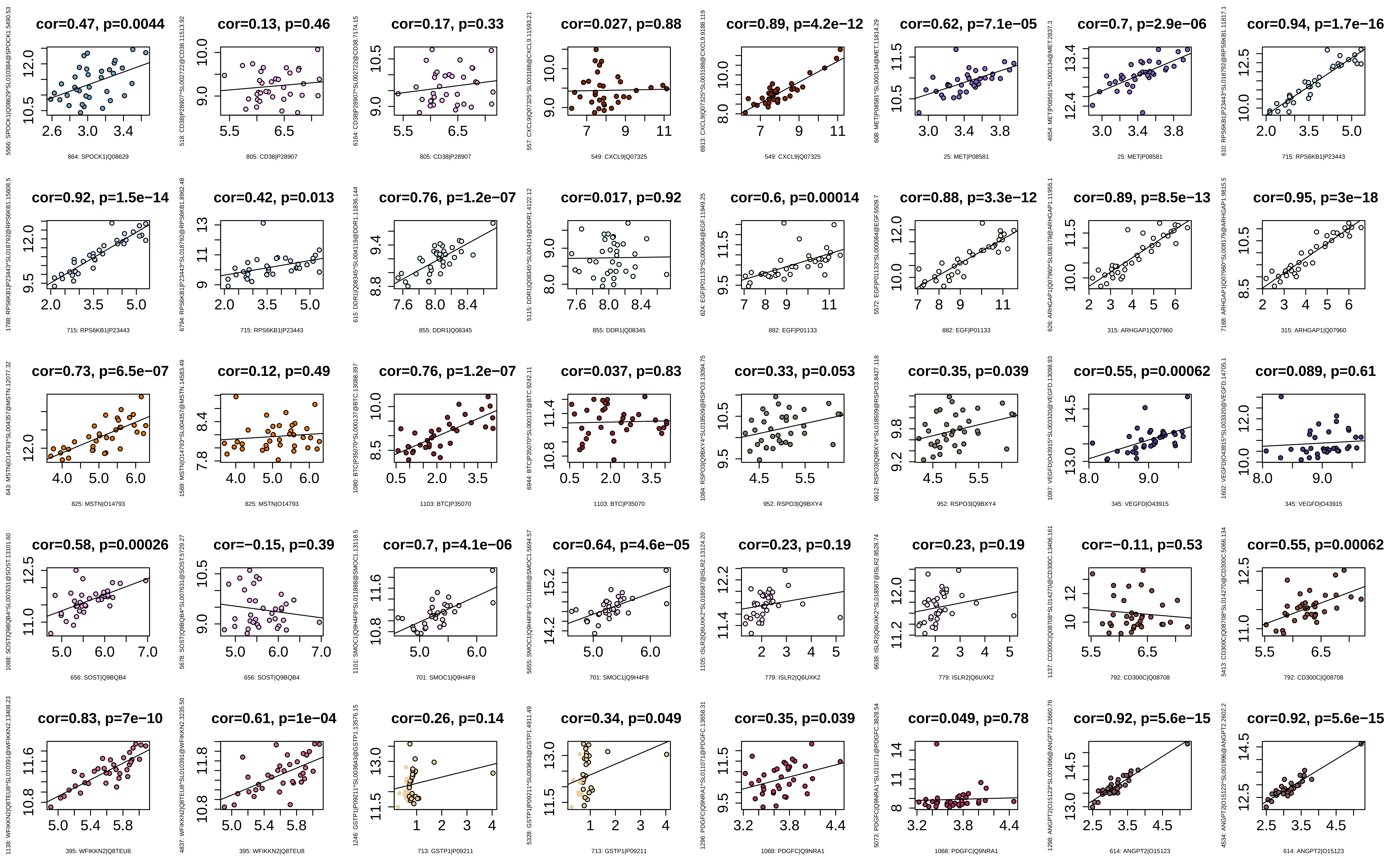
cor=0.47, p=0.0044
864: SPOCK1|Q08629
cor=0.13, p=0.46
805: CD38|P28907
cor=0.17, p=0.33
805: CD38|P28907
cor=0.027, p=0.88
549: CXCL9|Q07325
cor=0.89, p=4.2e−12
549: CXCL9|Q07325
cor=0.62, p=7.1e−05
25: MET|P08581
cor=0.7, p=2.9e−06
25: MET|P08581
cor=0.94, p=1.7e−16
715: RPS6KB1|P23443
cor=0.92, p=1.5e−14
715: RPS6KB1|P23443
cor=0.42, p=0.013
715: RPS6KB1|P23443
cor=0.76, p=1.2e−07
855: DDR1|Q08345
cor=0.017, p=0.92
855: DDR1|Q08345
cor=0.6, p=0.00014
882: EGF|P01133
cor=0.88, p=3.3e−12
882: EGF|P01133
cor=0.89, p=8.5e−13
315: ARHGAP1|Q07960
cor=0.95, p=3e−18
315: ARHGAP1|Q07960
cor=0.73, p=6.5e−07
825: MSTN|O14793
cor=0.12, p=0.49
825: MSTN|O14793
cor=0.76, p=1.2e−07
1103: BTC|P35070
cor=0.037, p=0.83
1103: BTC|P35070
cor=0.33, p=0.053
952: RSPO3|Q9BXY4
cor=0.35, p=0.039
952: RSPO3|Q9BXY4
cor=0.55, p=0.00062
345: VEGFD|O43915
cor=0.089, p=0.61
345: VEGFD|O43915
cor=0.58, p=0.00026
656: SOST|Q9BQB4
cor=−0.15, p=0.39
656: SOST|Q9BQB4
cor=0.7, p=4.1e−06
701: SMOC1|Q9H4F8
cor=0.64, p=4.6e−05
701: SMOC1|Q9H4F8
cor=0.23, p=0.19
779: ISLR2|Q6UXK2
cor=0.23, p=0.19
779: ISLR2|Q6UXK2
cor=−0.11, p=0.53
792: CD300C|Q08708
cor=0.55, p=0.00062
792: CD300C|Q08708
cor=0.83, p=7e−10
395: WFIKKN2|Q8TEU8
cor=0.61, p=1e−04
395: WFIKKN2|Q8TEU8
cor=0.26, p=0.14
713: GSTP1|P09211
cor=0.34, p=0.049
713: GSTP1|P09211
cor=0.35, p=0.039
1068: PDGFC|Q9NRA1
cor=0.049, p=0.78
1068: PDGFC|Q9NRA1
cor=0.92, p=5.6e−15
614: ANGPT2|O15123
cor=0.92, p=5.6e−15
614: ANGPT2|O15123

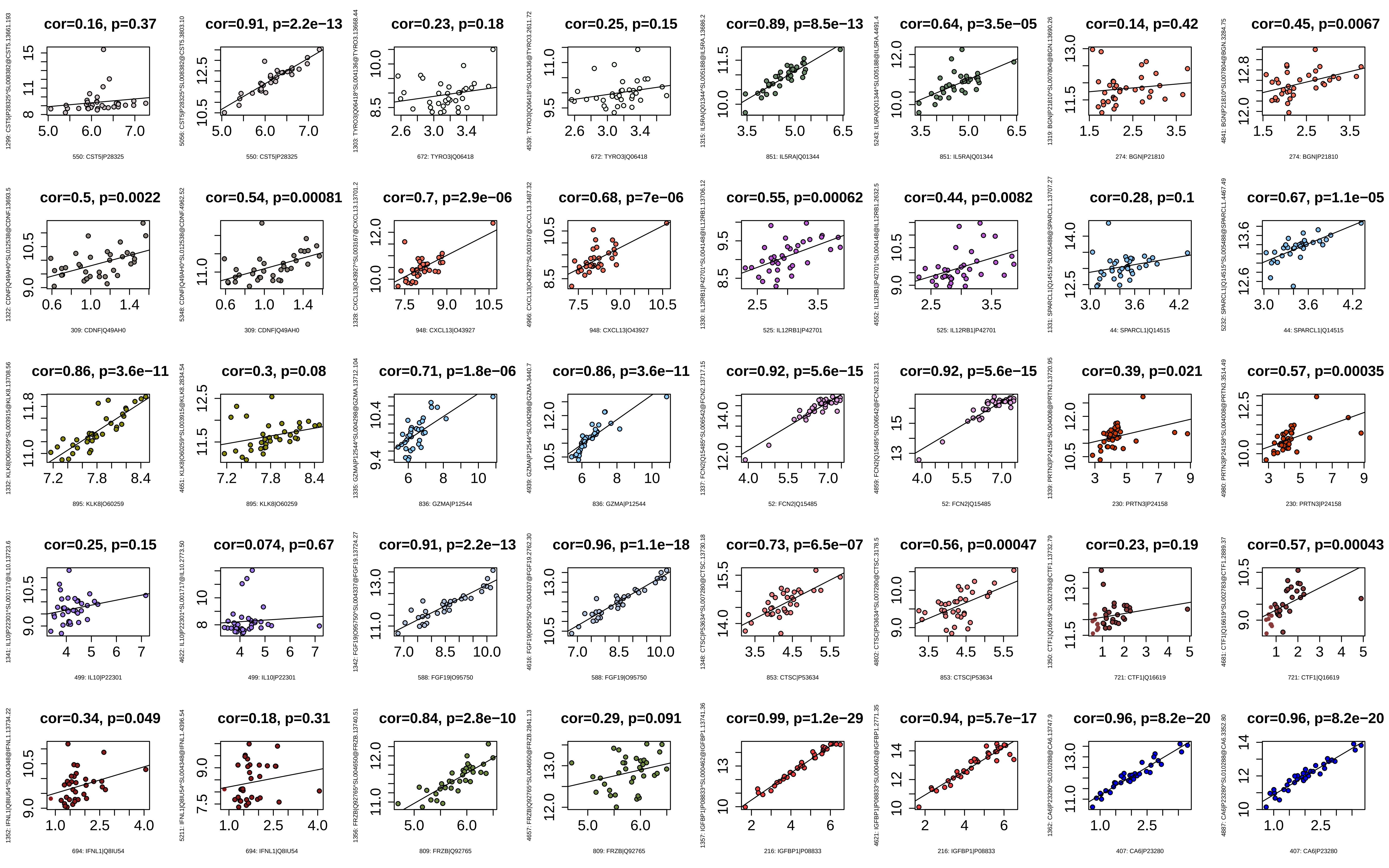
cor=0.16, p=0.37
550: CST5|P28325
cor=0.91, p=2.2e−13
550: CST5|P28325
cor=0.23, p=0.18
672: TYRO3|Q06418
cor=0.25, p=0.15
672: TYRO3|Q06418
cor=0.89, p=8.5e−13
851: IL5RA|Q01344
cor=0.64, p=3.5e−05
851: IL5RA|Q01344
cor=0.14, p=0.42
274: BGN|P21810
cor=0.45, p=0.0067
274: BGN|P21810
cor=0.5, p=0.0022
309: CDNF|Q49AH0
cor=0.54, p=0.00081
309: CDNF|Q49AH0
cor=0.7, p=2.9e−06
948: CXCL13|O43927
cor=0.68, p=7e−06
948: CXCL13|O43927
cor=0.55, p=0.00062
525: IL12RB1|P42701
cor=0.44, p=0.0082
525: IL12RB1|P42701
cor=0.28, p=0.1
44: SPARCL1|Q14515
cor=0.67, p=1.1e−05
44: SPARCL1|Q14515
cor=0.86, p=3.6e−11
895: KLK8|O60259
cor=0.3, p=0.08
895: KLK8|O60259
cor=0.71, p=1.8e−06
836: GZMA|P12544
cor=0.86, p=3.6e−11
836: GZMA|P12544
cor=0.92, p=5.6e−15
52: FCN2|Q15485
cor=0.92, p=5.6e−15
52: FCN2|Q15485
cor=0.39, p=0.021
230: PRTN3|P24158
cor=0.57, p=0.00035
230: PRTN3|P24158
cor=0.25, p=0.15
499: IL10|P22301
cor=0.074, p=0.67
499: IL10|P22301
cor=0.91, p=2.2e−13
588: FGF19|O95750
cor=0.96, p=1.1e−18
588: FGF19|O95750
cor=0.73, p=6.5e−07
853: CTSC|P53634
cor=0.56, p=0.00047
853: CTSC|P53634
cor=0.23, p=0.19
721: CTF1|Q16619
cor=0.57, p=0.00043
721: CTF1|Q16619
cor=0.34, p=0.049
694: IFNL1|Q8IU54
cor=0.18, p=0.31
694: IFNL1|Q8IU54
cor=0.84, p=2.8e−10
809: FRZB|Q92765
cor=0.29, p=0.091
809: FRZB|Q92765
cor=0.99, p=1.2e−29
216: IGFBP1|P08833
cor=0.94, p=5.7e−17
216: IGFBP1|P08833
cor=0.96, p=8.2e−20
407: CA6|P23280
cor=0.96, p=8.2e−20
407: CA6|P23280

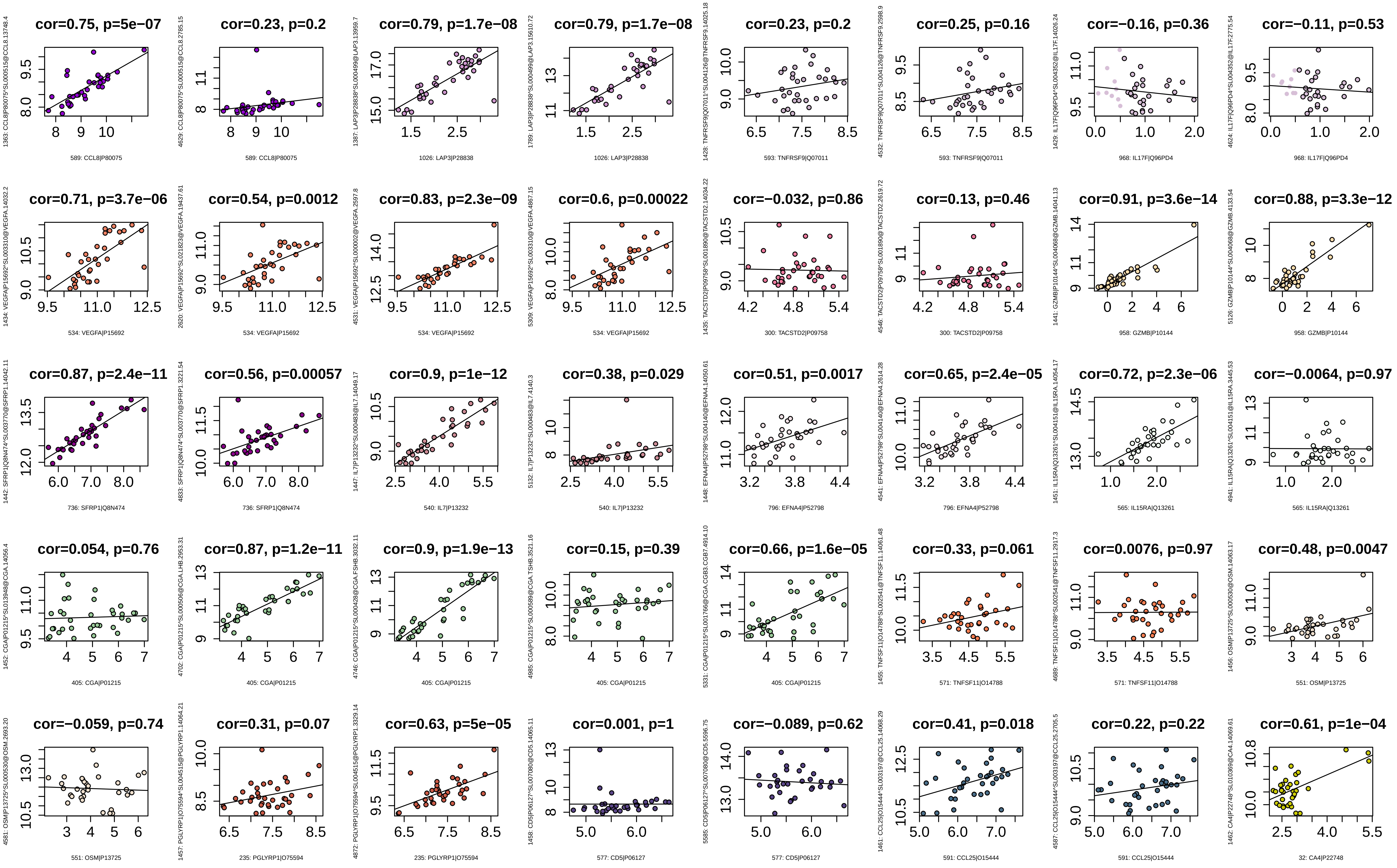
cor=0.75, p=5e−07
589: CCL8|P80075
cor=0.23, p=0.2
589: CCL8|P80075
cor=0.79, p=1.7e−08
1026: LAP3|P28838
cor=0.79, p=1.7e−08
1026: LAP3|P28838
cor=0.23, p=0.2
593: TNFRSF9|Q07011
cor=0.25, p=0.16
593: TNFRSF9|Q07011
cor=−0.16, p=0.36
968: IL17F|Q96PD4
cor=−0.11, p=0.53
968: IL17F|Q96PD4
cor=0.71, p=3.7e−06
534: VEGFA|P15692
cor=0.54, p=0.0012
534: VEGFA|P15692
cor=0.83, p=2.3e−09
534: VEGFA|P15692
cor=0.6, p=0.00022
534: VEGFA|P15692
cor=−0.032, p=0.86
300: TACSTD2|P09758
cor=0.13, p=0.46
300: TACSTD2|P09758
cor=0.91, p=3.6e−14
958: GZMB|P10144
cor=0.88, p=3.3e−12
958: GZMB|P10144
cor=0.87, p=2.4e−11
736: SFRP1|Q8N474
cor=0.56, p=0.00057
736: SFRP1|Q8N474
cor=0.9, p=1e−12
540: IL7|P13232
cor=0.38, p=0.029
540: IL7|P13232
cor=0.51, p=0.0017
796: EFNA4|P52798
cor=0.65, p=2.4e−05
796: EFNA4|P52798
cor=0.72, p=2.3e−06
565: IL15RA|Q13261
cor=−0.0064, p=0.97
565: IL15RA|Q13261
cor=0.054, p=0.76
405: CGA|P01215
cor=0.87, p=1.2e−11
405: CGA|P01215
cor=0.9, p=1.9e−13
405: CGA|P01215
cor=0.15, p=0.39
405: CGA|P01215
cor=0.66, p=1.6e−05
405: CGA|P01215
cor=0.33, p=0.061
571: TNFSF11|O14788
cor=0.0076, p=0.97
571: TNFSF11|O14788
cor=0.48, p=0.0047
551: OSM|P13725
cor=−0.059, p=0.74
551: OSM|P13725
cor=0.31, p=0.07
235: PGLYRP1|O75594
cor=0.63, p=5e−05
235: PGLYRP1|O75594
cor=0.001, p=1
577: CD5|P06127
cor=−0.089, p=0.62
577: CD5|P06127
cor=0.41, p=0.018
591: CCL25|O15444
cor=0.22, p=0.22
591: CCL25|O15444
cor=0.61, p=1e−04
32: CA4|P22748

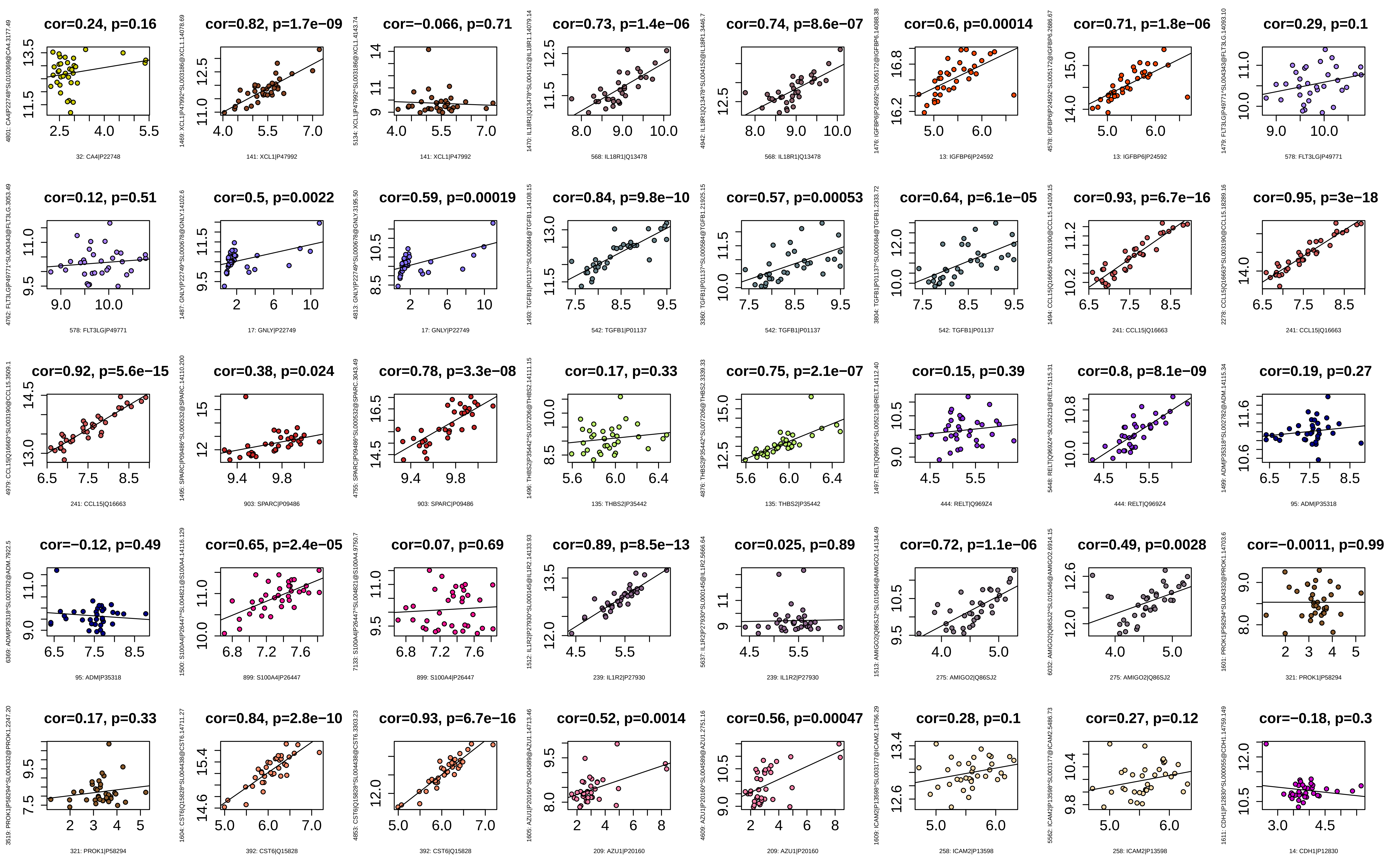
cor=0.24, p=0.16
32: CA4|P22748
cor=0.82, p=1.7e−09
141: XCL1|P47992
cor=−0.066, p=0.71
141: XCL1|P47992
cor=0.73, p=1.4e−06
568: IL18R1|Q13478
cor=0.74, p=8.6e−07
568: IL18R1|Q13478
cor=0.6, p=0.00014
13: IGFBP6|P24592
cor=0.71, p=1.8e−06
13: IGFBP6|P24592
cor=0.29, p=0.1
578: FLT3LG|P49771
cor=0.12, p=0.51
578: FLT3LG|P49771
cor=0.5, p=0.0022
17: GNLY|P22749
cor=0.59, p=0.00019
17: GNLY|P22749
cor=0.84, p=9.8e−10
542: TGFB1|P01137
cor=0.57, p=0.00053
542: TGFB1|P01137
cor=0.64, p=6.1e−05
542: TGFB1|P01137
cor=0.93, p=6.7e−16
241: CCL15|Q16663
cor=0.95, p=3e−18
241: CCL15|Q16663
cor=0.92, p=5.6e−15
241: CCL15|Q16663
cor=0.38, p=0.024
903: SPARC|P09486
cor=0.78, p=3.3e−08
903: SPARC|P09486
cor=0.17, p=0.33
135: THBS2|P35442
cor=0.75, p=2.1e−07
135: THBS2|P35442
cor=0.15, p=0.39
444: RELT|Q969Z4
cor=0.8, p=8.1e−09
444: RELT|Q969Z4
cor=0.19, p=0.27
95: ADM|P35318
cor=−0.12, p=0.49
95: ADM|P35318
cor=0.65, p=2.4e−05
899: S100A4|P26447
cor=0.07, p=0.69
899: S100A4|P26447
cor=0.89, p=8.5e−13
239: IL1R2|P27930
cor=0.025, p=0.89
239: IL1R2|P27930
cor=0.72, p=1.1e−06
275: AMIGO2|Q86SJ2
cor=0.49, p=0.0028
275: AMIGO2|Q86SJ2
cor=−0.0011, p=0.99
321: PROK1|P58294
cor=0.17, p=0.33
321: PROK1|P58294
cor=0.84, p=2.8e−10
392: CST6|Q15828
cor=0.93, p=6.7e−16
392: CST6|Q15828
cor=0.52, p=0.0014
209: AZU1|P20160
cor=0.56, p=0.00047
209: AZU1|P20160
cor=0.28, p=0.1
258: ICAM2|P13598
cor=0.27, p=0.12
258: ICAM2|P13598
cor=−0.18, p=0.3
14: CDH1|P12830

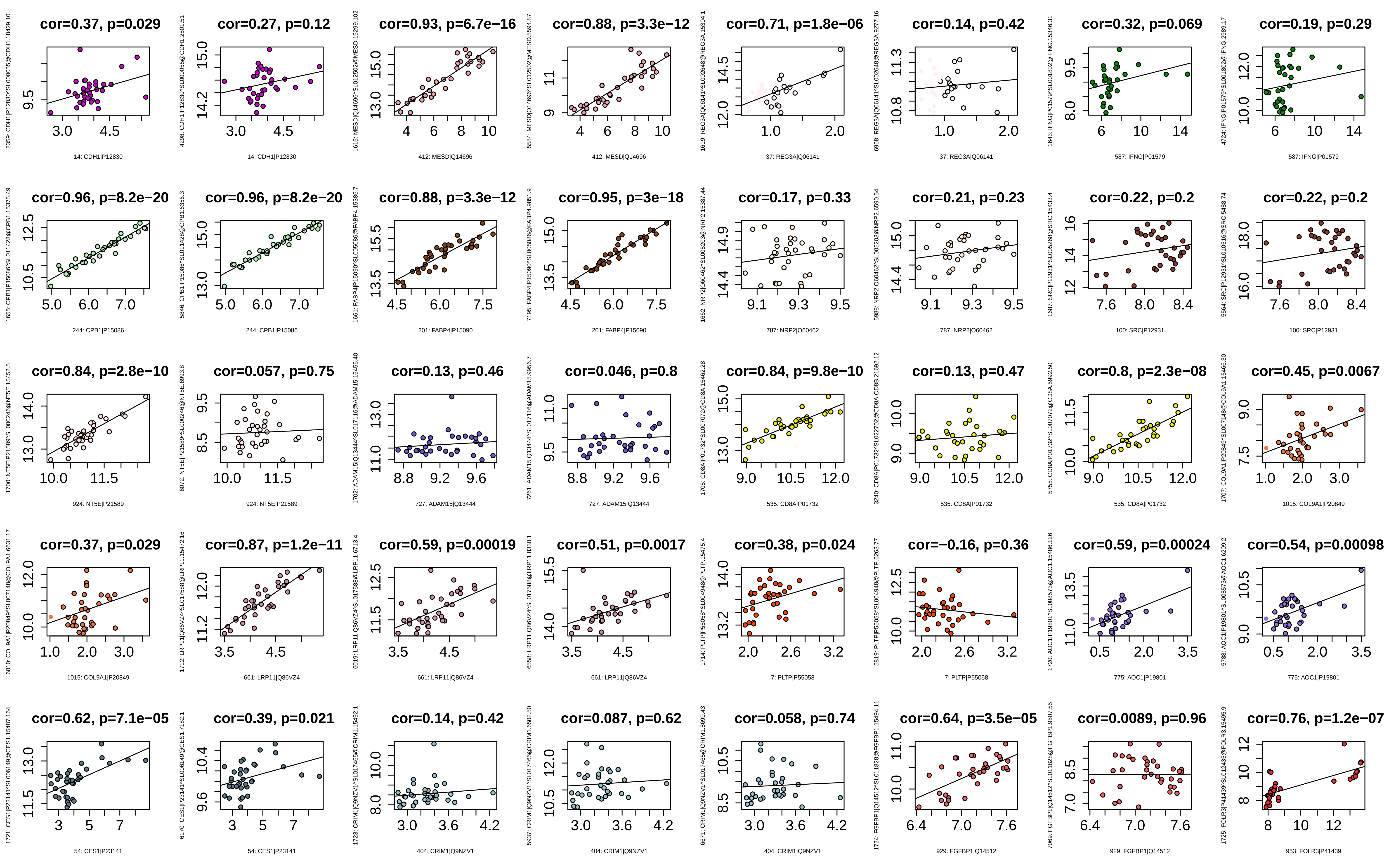

cor=0.37, p=0.029
14: CDH1|P12830
cor=0.27, p=0.12
14: CDH1|P12830
cor=0.93, p=6.7e−16
412: MESD|Q14696
cor=0.88, p=3.3e−12
412: MESD|Q14696
cor=0.71, p=1.8e−06
37: REG3A|Q06141
cor=0.14, p=0.42
37: REG3A|Q06141
cor=0.32, p=0.069
587: IFNG|P01579
cor=0.19, p=0.29
587: IFNG|P01579
cor=0.96, p=8.2e−20
244: CPB1|P15086
cor=0.96, p=8.2e−20
244: CPB1|P15086
cor=0.88, p=3.3e−12
201: FABP4|P15090
cor=0.95, p=3e−18
201: FABP4|P15090
cor=0.17, p=0.33
787: NRP2|O60462
cor=0.21, p=0.23
787: NRP2|O60462
cor=0.22, p=0.2
100: SRC|P12931
cor=0.22, p=0.2
100: SRC|P12931
cor=0.84, p=2.8e−10
924: NT5E|P21589
cor=0.057, p=0.75
924: NT5E|P21589
cor=0.13, p=0.46
727: ADAM15|Q13444
cor=0.046, p=0.8
727: ADAM15|Q13444
cor=0.84, p=9.8e−10
535: CD8A|P01732
cor=0.13, p=0.47
535: CD8A|P01732
cor=0.8, p=2.3e−08
535: CD8A|P01732
cor=0.45, p=0.0067
1015: COL9A1|P20849
cor=0.37, p=0.029
1015: COL9A1|P20849
cor=0.87, p=1.2e−11
661: LRP11|Q86VZ4
cor=0.59, p=0.00019
661: LRP11|Q86VZ4
cor=0.51, p=0.0017
661: LRP11|Q86VZ4
cor=0.38, p=0.024
7: PLTP|P55058
cor=−0.16, p=0.36
7: PLTP|P55058
cor=0.59, p=0.00024
775: AOC1|P19801
cor=0.54, p=0.00098
775: AOC1|P19801
cor=0.62, p=7.1e−05
54: CES1|P23141
cor=0.39, p=0.021
54: CES1|P23141
cor=0.14, p=0.42
404: CRIM1|Q9NZV1
cor=0.087, p=0.62
404: CRIM1|Q9NZV1
cor=0.058, p=0.74
404: CRIM1|Q9NZV1
cor=0.64, p=3.5e−05
929: FGFBP1|Q14512
cor=0.0089, p=0.96
929: FGFBP1|Q14512
cor=0.76, p=1.2e−07
953: FOLR3|P41439

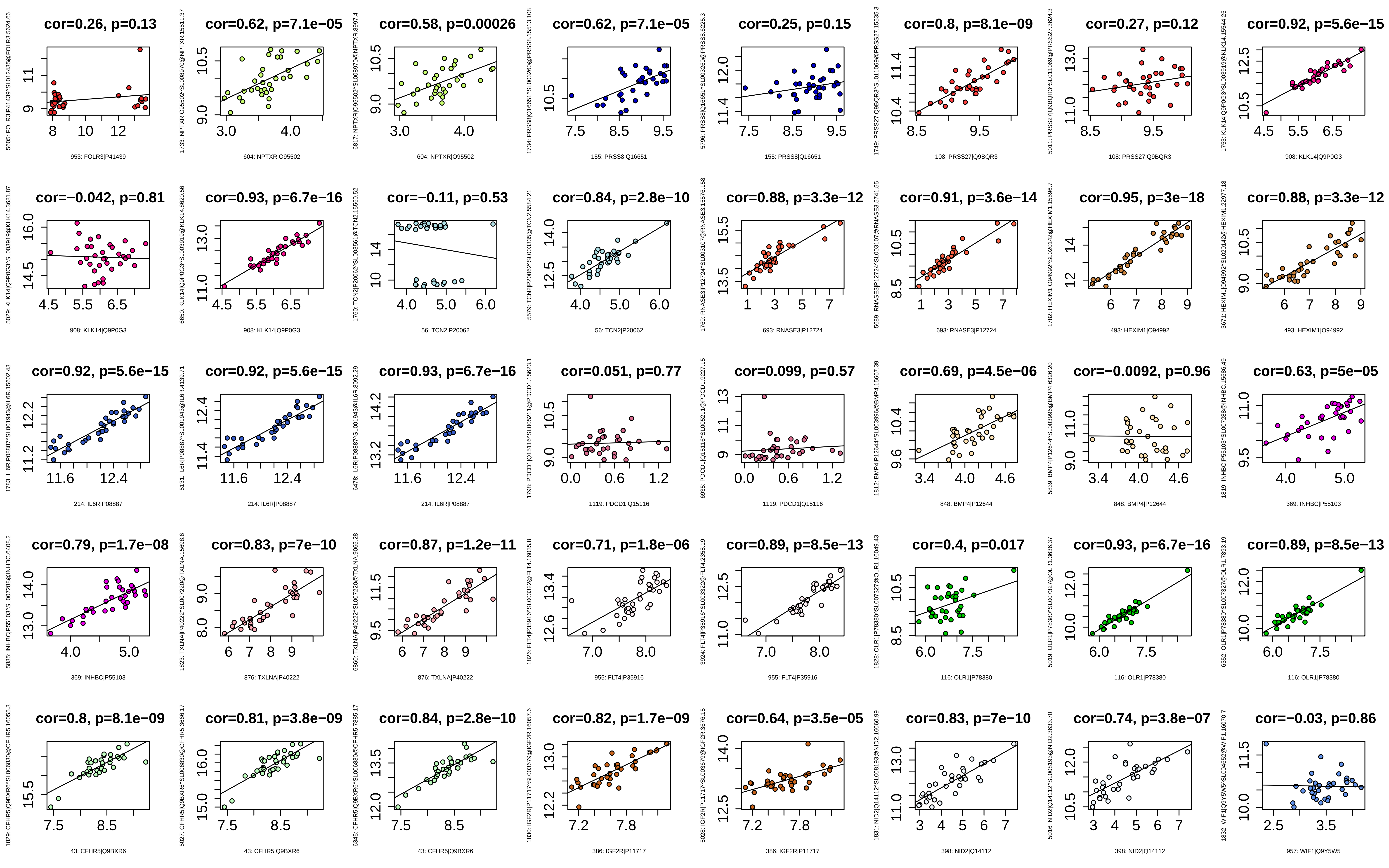
cor=0.26, p=0.13
953: FOLR3|P41439
cor=0.62, p=7.1e−05
604: NPTXR|O95502
cor=0.58, p=0.00026
604: NPTXR|O95502
cor=0.62, p=7.1e−05
155: PRSS8|Q16651
cor=0.25, p=0.15
155: PRSS8|Q16651
cor=0.8, p=8.1e−09
108: PRSS27|Q9BQR3
cor=0.27, p=0.12
108: PRSS27|Q9BQR3
cor=0.92, p=5.6e−15
908: KLK14|Q9P0G3
cor=−0.042, p=0.81
908: KLK14|Q9P0G3
cor=0.93, p=6.7e−16
908: KLK14|Q9P0G3
cor=−0.11, p=0.53
56: TCN2|P20062
cor=0.84, p=2.8e−10
56: TCN2|P20062
cor=0.88, p=3.3e−12
693: RNASE3|P12724
cor=0.91, p=3.6e−14
693: RNASE3|P12724
cor=0.95, p=3e−18
493: HEXIM1|O94992
cor=0.88, p=3.3e−12
493: HEXIM1|O94992
cor=0.92, p=5.6e−15
214: IL6R|P08887
cor=0.92, p=5.6e−15
214: IL6R|P08887
cor=0.93, p=6.7e−16
214: IL6R|P08887
cor=0.051, p=0.77
1119: PDCD1|Q15116
cor=0.099, p=0.57
1119: PDCD1|Q15116
cor=0.69, p=4.5e−06
848: BMP4|P12644
cor=−0.0092, p=0.96
848: BMP4|P12644
cor=0.63, p=5e−05
369: INHBC|P55103
cor=0.79, p=1.7e−08
369: INHBC|P55103
cor=0.83, p=7e−10
876: TXLNA|P40222
cor=0.87, p=1.2e−11
876: TXLNA|P40222
cor=0.71, p=1.8e−06
955: FLT4|P35916
cor=0.89, p=8.5e−13
955: FLT4|P35916
cor=0.4, p=0.017
116: OLR1|P78380
cor=0.93, p=6.7e−16
116: OLR1|P78380
cor=0.89, p=8.5e−13
116: OLR1|P78380
cor=0.8, p=8.1e−09
43: CFHR5|Q9BXR6
cor=0.81, p=3.8e−09
43: CFHR5|Q9BXR6
cor=0.84, p=2.8e−10
43: CFHR5|Q9BXR6
cor=0.82, p=1.7e−09
386: IGF2R|P11717
cor=0.64, p=3.5e−05
386: IGF2R|P11717
cor=0.83, p=7e−10
398: NID2|Q14112
cor=0.74, p=3.8e−07
398: NID2|Q14112
cor=−0.03, p=0.86
957: WIF1|Q9Y5W5

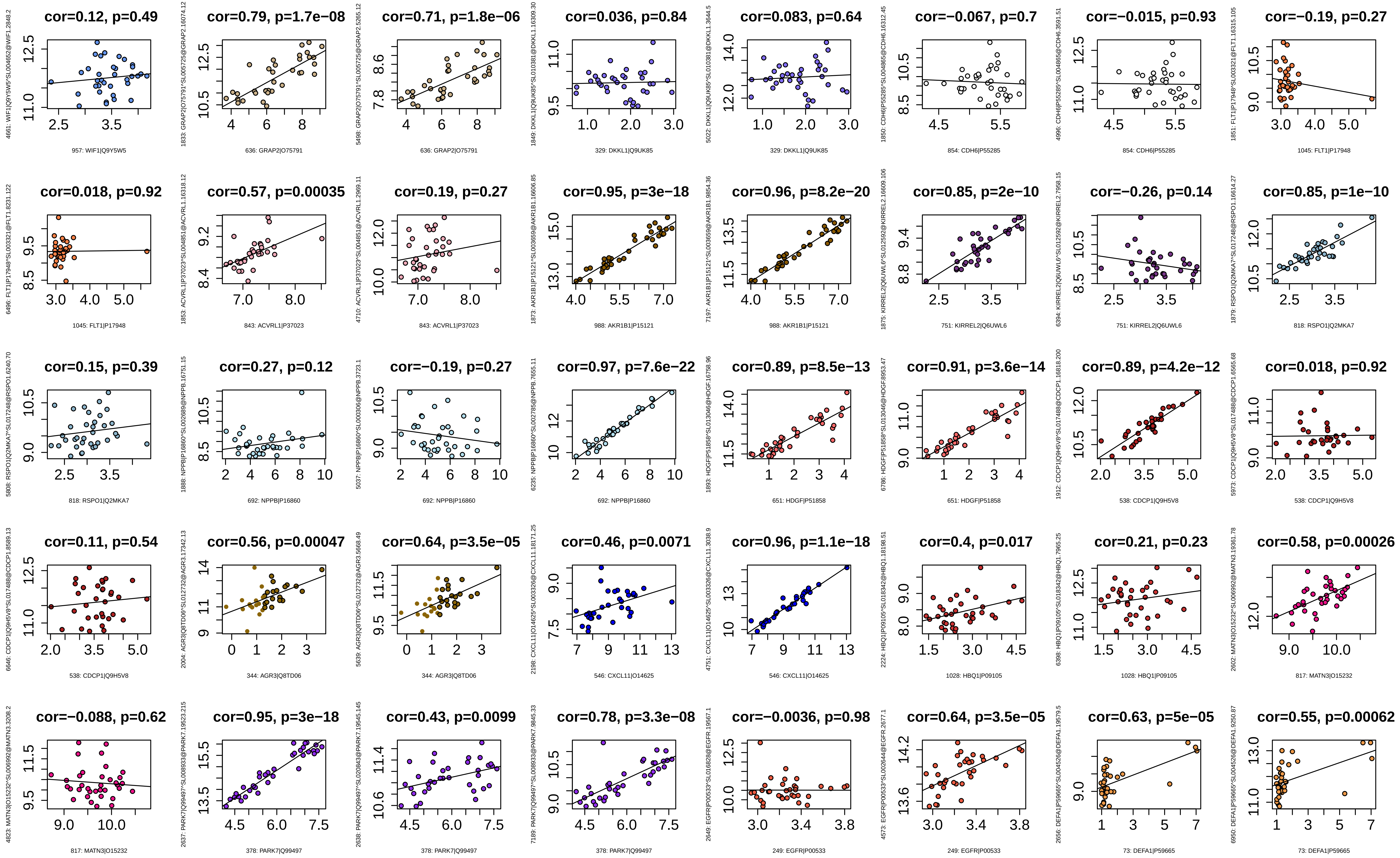
cor=0.12, p=0.49
957: WIF1|Q9Y5W5
cor=0.79, p=1.7e−08
636: GRAP2|O75791
cor=0.71, p=1.8e−06
636: GRAP2|O75791
cor=0.036, p=0.84
329: DKKL1|Q9UK85
cor=0.083, p=0.64
329: DKKL1|Q9UK85
cor=−0.067, p=0.7
854: CDH6|P55285
cor=−0.015, p=0.93
854: CDH6|P55285
cor=−0.19, p=0.27
1045: FLT1|P17948
cor=0.018, p=0.92
1045: FLT1|P17948
cor=0.57, p=0.00035
843: ACVRL1|P37023
cor=0.19, p=0.27
843: ACVRL1|P37023
cor=0.95, p=3e−18
988: AKR1B1|P15121
cor=0.96, p=8.2e−20
988: AKR1B1|P15121
cor=0.85, p=2e−10
751: KIRREL2|Q6UWL6
cor=−0.26, p=0.14
751: KIRREL2|Q6UWL6
cor=0.85, p=1e−10
818: RSPO1|Q2MKA7
cor=0.15, p=0.39
818: RSPO1|Q2MKA7
cor=0.27, p=0.12
692: NPPB|P16860
cor=−0.19, p=0.27
692: NPPB|P16860
cor=0.97, p=7.6e−22
692: NPPB|P16860
cor=0.89, p=8.5e−13
651: HDGF|P51858
cor=0.91, p=3.6e−14
651: HDGF|P51858
cor=0.89, p=4.2e−12
538: CDCP1|Q9H5V8
cor=0.018, p=0.92
538: CDCP1|Q9H5V8
cor=0.11, p=0.54
538: CDCP1|Q9H5V8
cor=0.56, p=0.00047
344: AGR3|Q8TD06
cor=0.64, p=3.5e−05
344: AGR3|Q8TD06
cor=0.46, p=0.0071
546: CXCL11|O14625
cor=0.96, p=1.1e−18
546: CXCL11|O14625
cor=0.4, p=0.017
1028: HBQ1|P09105
cor=0.21, p=0.23
1028: HBQ1|P09105
cor=0.58, p=0.00026
817: MATN3|O15232
cor=−0.088, p=0.62
817: MATN3|O15232
cor=0.95, p=3e−18
378: PARK7|Q99497
cor=0.43, p=0.0099
378: PARK7|Q99497
cor=0.78, p=3.3e−08
378: PARK7|Q99497
cor=−0.0036, p=0.98
249: EGFR|P00533
cor=0.64, p=3.5e−05
249: EGFR|P00533
cor=0.63, p=5e−05
73: DEFA1|P59665
cor=0.55, p=0.00062
73: DEFA1|P59665

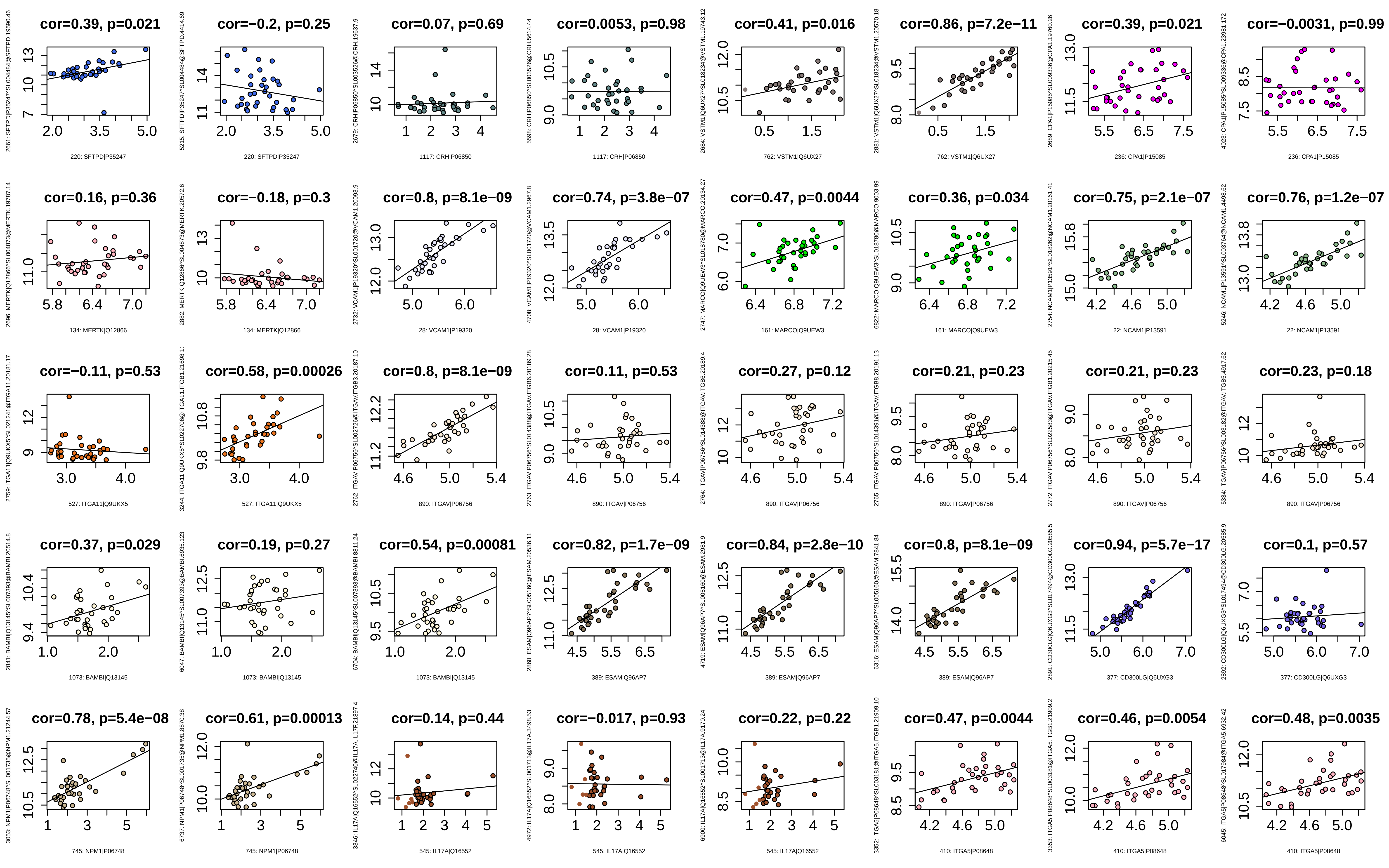

cor=0.39, p=0.021
220: SFTPD|P35247
cor=−0.2, p=0.25
220: SFTPD|P35247
cor=0.07, p=0.69
1117: CRH|P06850
cor=0.0053, p=0.98
1117: CRH|P06850
cor=0.41, p=0.016
762: VSTM1|Q6UX27
cor=0.86, p=7.2e−11
762: VSTM1|Q6UX27
cor=0.39, p=0.021
236: CPA1|P15085
cor=−0.0031, p=0.99
236: CPA1|P15085
cor=0.16, p=0.36
134: MERTK|Q12866
cor=−0.18, p=0.3
134: MERTK|Q12866
cor=0.8, p=8.1e−09
28: VCAM1|P19320
cor=0.74, p=3.8e−07
28: VCAM1|P19320
cor=0.47, p=0.0044
161: MARCO|Q9UEW3
cor=0.36, p=0.034
161: MARCO|Q9UEW3
cor=0.75, p=2.1e−07
22: NCAM1|P13591
cor=0.76, p=1.2e−07
22: NCAM1|P13591
cor=−0.11, p=0.53
527: ITGA11|Q9UKX5
cor=0.58, p=0.00026
527: ITGA11|Q9UKX5
cor=0.8, p=8.1e−09
890: ITGAV|P06756
cor=0.11, p=0.53
890: ITGAV|P06756
cor=0.27, p=0.12
890: ITGAV|P06756
cor=0.21, p=0.23
890: ITGAV|P06756
cor=0.21, p=0.23
890: ITGAV|P06756
cor=0.23, p=0.18
890: ITGAV|P06756
cor=0.37, p=0.029
1073: BAMBI|Q13145
cor=0.19, p=0.27
1073: BAMBI|Q13145
cor=0.54, p=0.00081
1073: BAMBI|Q13145
cor=0.82, p=1.7e−09
389: ESAM|Q96AP7
cor=0.84, p=2.8e−10
389: ESAM|Q96AP7
cor=0.8, p=8.1e−09
389: ESAM|Q96AP7
cor=0.94, p=5.7e−17
377: CD300LG|Q6UXG3
cor=0.1, p=0.57
377: CD300LG|Q6UXG3
cor=0.78, p=5.4e−08
745: NPM1|P06748
cor=0.61, p=0.00013
745: NPM1|P06748
cor=0.14, p=0.44
545: IL17A|Q16552
cor=−0.017, p=0.93
545: IL17A|Q16552
cor=0.22, p=0.22
545: IL17A|Q16552
cor=0.47, p=0.0044
410: ITGA5|P08648
cor=0.46, p=0.0054
410: ITGA5|P08648
cor=0.48, p=0.0035
410: ITGA5|P08648

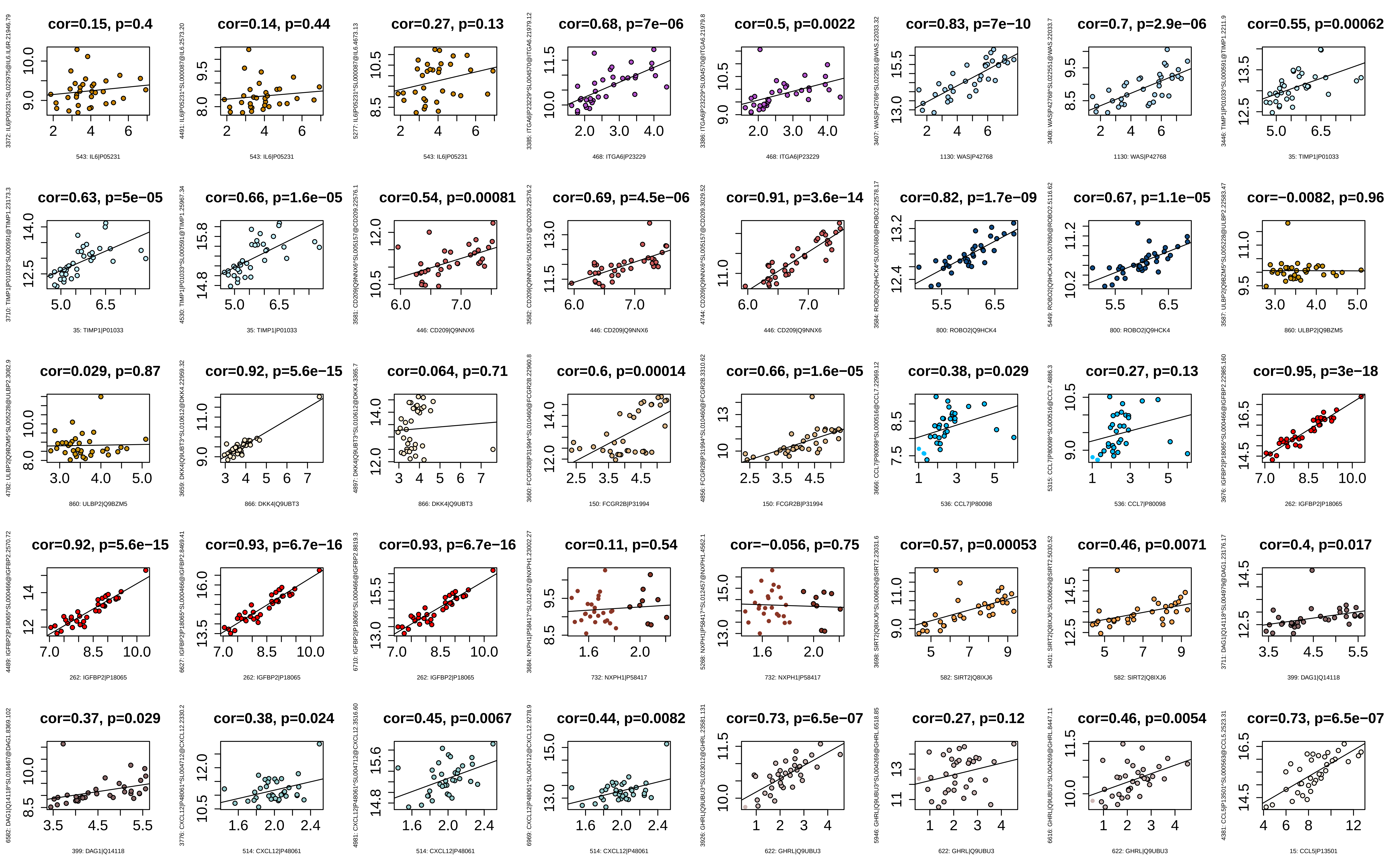
cor=0.15, p=0.4
543: IL6|P05231
cor=0.14, p=0.44
543: IL6|P05231
cor=0.27, p=0.13
543: IL6|P05231
cor=0.68, p=7e−06
468: ITGA6|P23229
cor=0.5, p=0.0022
468: ITGA6|P23229
cor=0.83, p=7e−10
1130: WAS|P42768
cor=0.7, p=2.9e−06
1130: WAS|P42768
cor=0.55, p=0.00062
35: TIMP1|P01033
cor=0.63, p=5e−05
35: TIMP1|P01033
cor=0.66, p=1.6e−05
35: TIMP1|P01033
cor=0.54, p=0.00081
446: CD209|Q9NNX6
cor=0.69, p=4.5e−06
446: CD209|Q9NNX6
cor=0.91, p=3.6e−14
446: CD209|Q9NNX6
cor=0.82, p=1.7e−09
800: ROBO2|Q9HCK4
cor=0.67, p=1.1e−05
800: ROBO2|Q9HCK4
cor=−0.0082, p=0.96
860: ULBP2|Q9BZM5
cor=0.029, p=0.87
860: ULBP2|Q9BZM5
cor=0.92, p=5.6e−15
866: DKK4|Q9UBT3
cor=0.064, p=0.71
866: DKK4|Q9UBT3
cor=0.6, p=0.00014
150: FCGR2B|P31994
cor=0.66, p=1.6e−05
150: FCGR2B|P31994
cor=0.38, p=0.029
536: CCL7|P80098
cor=0.27, p=0.13
536: CCL7|P80098
cor=0.95, p=3e−18
262: IGFBP2|P18065
cor=0.92, p=5.6e−15
262: IGFBP2|P18065
cor=0.93, p=6.7e−16
262: IGFBP2|P18065
cor=0.93, p=6.7e−16
262: IGFBP2|P18065
cor=0.11, p=0.54
732: NXPH1|P58417
cor=−0.056, p=0.75
732: NXPH1|P58417
cor=0.57, p=0.00053
582: SIRT2|Q8IXJ6
cor=0.46, p=0.0071
582: SIRT2|Q8IXJ6
cor=0.4, p=0.017
399: DAG1|Q14118
cor=0.37, p=0.029
399: DAG1|Q14118
cor=0.38, p=0.024
514: CXCL12|P48061
cor=0.45, p=0.0067
514: CXCL12|P48061
cor=0.44, p=0.0082
514: CXCL12|P48061
cor=0.73, p=6.5e−07
622: GHRL|Q9UBU3
cor=0.27, p=0.12
622: GHRL|Q9UBU3
cor=0.46, p=0.0054
622: GHRL|Q9UBU3
cor=0.73, p=6.5e−07
15: CCL5|P13501

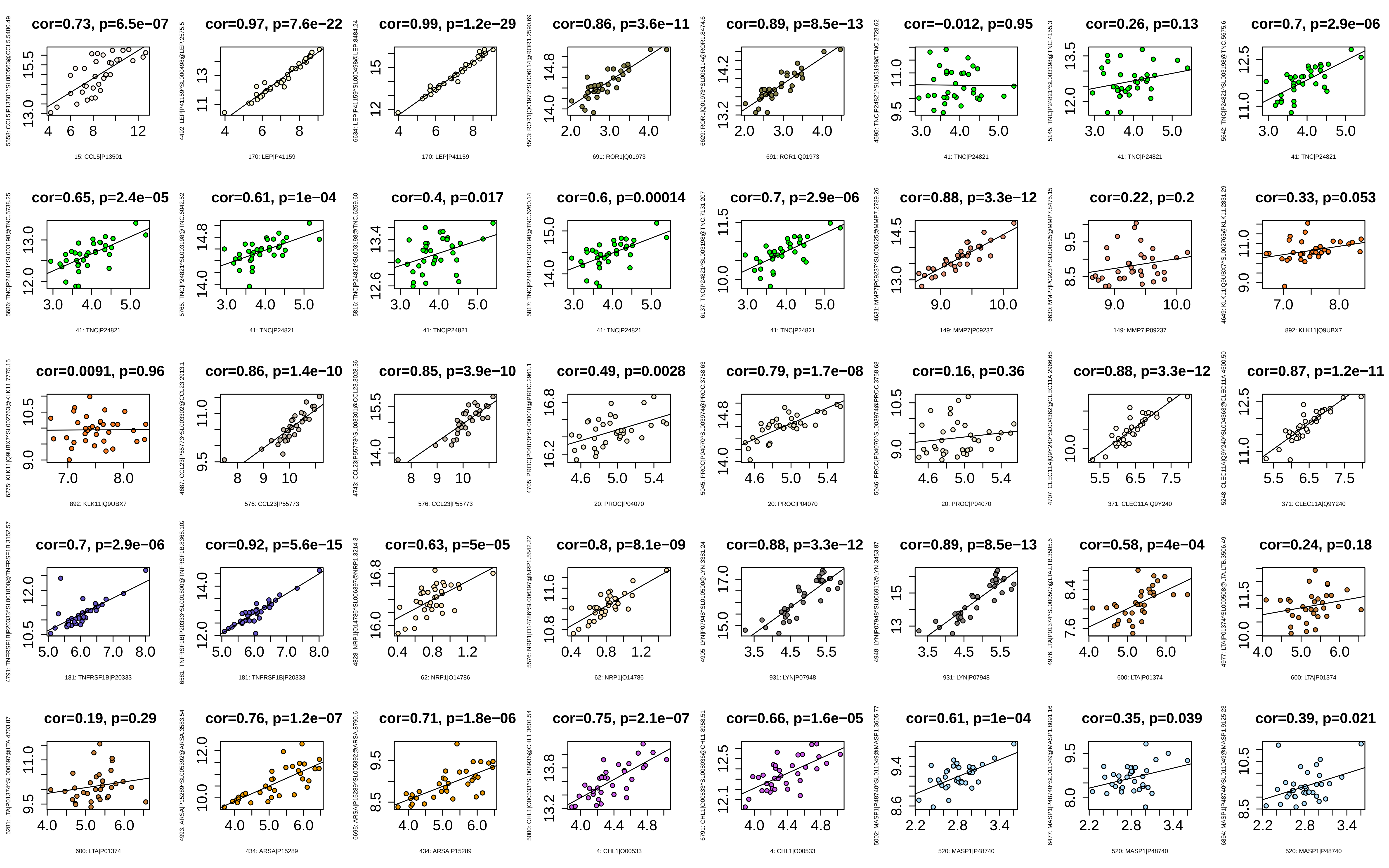

cor=0.73, p=6.5e−07
15: CCL5|P13501
cor=0.97, p=7.6e−22
170: LEP|P41159
cor=0.99, p=1.2e−29
170: LEP|P41159
cor=0.86, p=3.6e−11
691: ROR1|Q01973
cor=0.89, p=8.5e−13
691: ROR1|Q01973
cor=−0.012, p=0.95
41: TNC|P24821
cor=0.26, p=0.13
41: TNC|P24821
cor=0.7, p=2.9e−06
41: TNC|P24821
cor=0.65, p=2.4e−05
41: TNC|P24821
cor=0.61, p=1e−04
41: TNC|P24821
cor=0.4, p=0.017
41: TNC|P24821
cor=0.6, p=0.00014
41: TNC|P24821
cor=0.7, p=2.9e−06
41: TNC|P24821
cor=0.88, p=3.3e−12
149: MMP7|P09237
cor=0.22, p=0.2
149: MMP7|P09237
cor=0.33, p=0.053
892: KLK11|Q9UBX7
cor=0.0091, p=0.96
892: KLK11|Q9UBX7
cor=0.86, p=1.4e−10
576: CCL23|P55773
cor=0.85, p=3.9e−10
576: CCL23|P55773
cor=0.49, p=0.0028
20: PROC|P04070
cor=0.79, p=1.7e−08
20: PROC|P04070
cor=0.16, p=0.36
20: PROC|P04070
cor=0.88, p=3.3e−12
371: CLEC11A|Q9Y240
cor=0.87, p=1.2e−11
371: CLEC11A|Q9Y240
cor=0.7, p=2.9e−06
181: TNFRSF1B|P20333
cor=0.92, p=5.6e−15
181: TNFRSF1B|P20333
cor=0.63, p=5e−05
62: NRP1|O14786
cor=0.8, p=8.1e−09
62: NRP1|O14786
cor=0.88, p=3.3e−12
931: LYN|P07948
cor=0.89, p=8.5e−13
931: LYN|P07948
cor=0.58, p=4e−04
600: LTA|P01374
cor=0.24, p=0.18
600: LTA|P01374
cor=0.19, p=0.29
600: LTA|P01374
cor=0.76, p=1.2e−07
434: ARSA|P15289
cor=0.71, p=1.8e−06
434: ARSA|P15289
cor=0.75, p=2.1e−07
4: CHL1|O00533
cor=0.66, p=1.6e−05
4: CHL1|O00533
cor=0.61, p=1e−04
520: MASP1|P48740
cor=0.35, p=0.039
520: MASP1|P48740
cor=0.39, p=0.021
520: MASP1|P48740

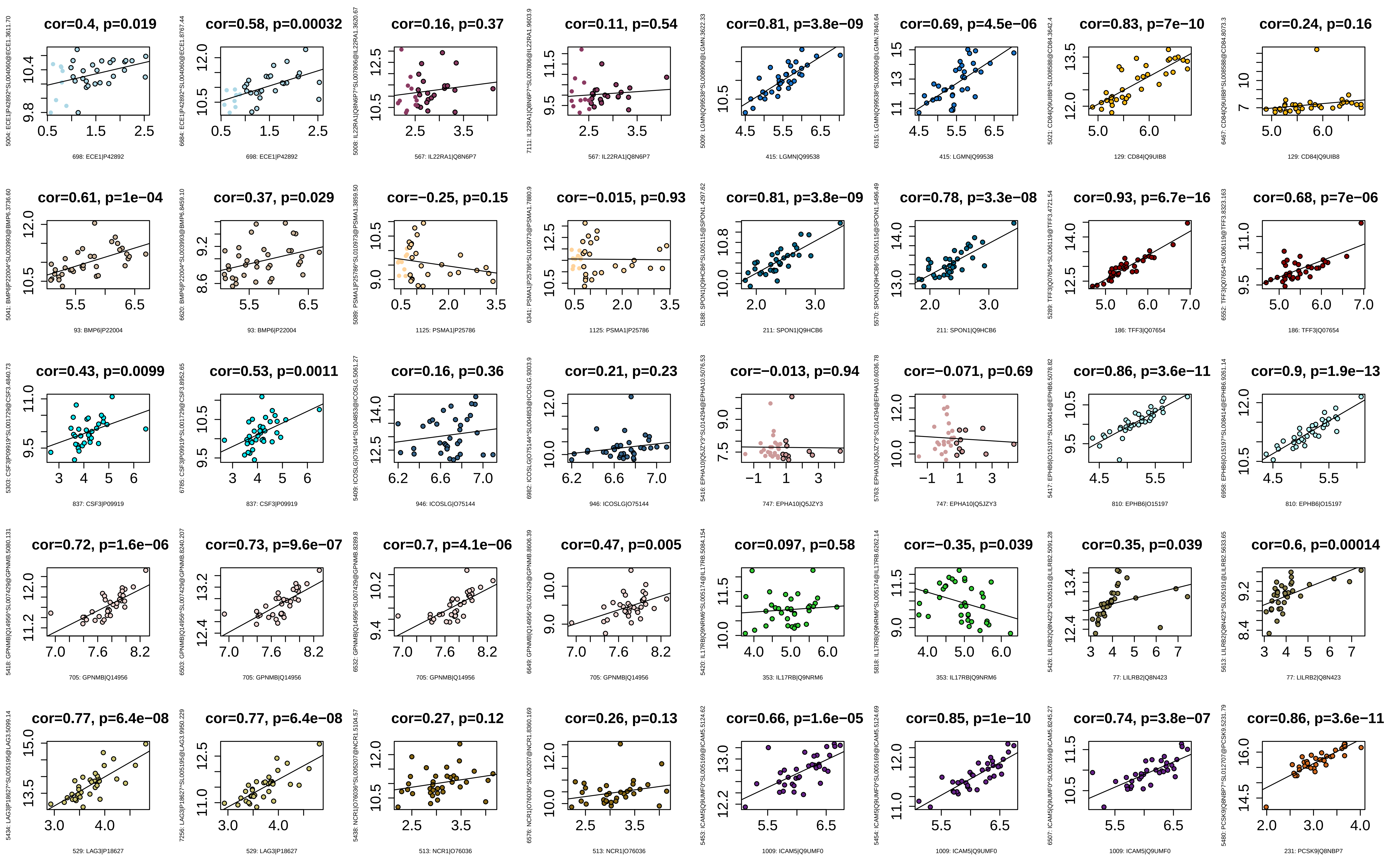

cor=0.4, p=0.019
698: ECE1|P42892
cor=0.58, p=0.00032
698: ECE1|P42892
cor=0.16, p=0.37
567: IL22RA1|Q8N6P7
cor=0.11, p=0.54
567: IL22RA1|Q8N6P7
cor=0.81, p=3.8e−09
415: LGMN|Q99538
cor=0.69, p=4.5e−06
415: LGMN|Q99538
cor=0.83, p=7e−10
129: CD84|Q9UIB8
cor=0.24, p=0.16
129: CD84|Q9UIB8
cor=0.61, p=1e−04
93: BMP6|P22004
cor=0.37, p=0.029
93: BMP6|P22004
cor=−0.25, p=0.15
1125: PSMA1|P25786
cor=−0.015, p=0.93
1125: PSMA1|P25786
cor=0.81, p=3.8e−09
211: SPON1|Q9HCB6
cor=0.78, p=3.3e−08
211: SPON1|Q9HCB6
cor=0.93, p=6.7e−16
186: TFF3|Q07654
cor=0.68, p=7e−06
186: TFF3|Q07654
cor=0.43, p=0.0099
837: CSF3|P09919
cor=0.53, p=0.0011
837: CSF3|P09919
cor=0.16, p=0.36
946: ICOSLG|O75144
cor=0.21, p=0.23
946: ICOSLG|O75144
cor=−0.013, p=0.94
747: EPHA10|Q5JZY3
cor=−0.071, p=0.69
747: EPHA10|Q5JZY3
cor=0.86, p=3.6e−11
810: EPHB6|O15197
cor=0.9, p=1.9e−13
810: EPHB6|O15197
cor=0.72, p=1.6e−06
705: GPNMB|Q14956
cor=0.73, p=9.6e−07
705: GPNMB|Q14956
cor=0.7, p=4.1e−06
705: GPNMB|Q14956
cor=0.47, p=0.005
705: GPNMB|Q14956
cor=0.097, p=0.58
353: IL17RB|Q9NRM6
cor=−0.35, p=0.039
353: IL17RB|Q9NRM6
cor=0.35, p=0.039
77: LILRB2|Q8N423
cor=0.6, p=0.00014
77: LILRB2|Q8N423
cor=0.77, p=6.4e−08
529: LAG3|P18627
cor=0.77, p=6.4e−08
529: LAG3|P18627
cor=0.27, p=0.12
513: NCR1|O76036
cor=0.26, p=0.13
513: NCR1|O76036
cor=0.66, p=1.6e−05
1009: ICAM5|Q9UMF0
cor=0.85, p=1e−10
1009: ICAM5|Q9UMF0
cor=0.74, p=3.8e−07
1009: ICAM5|Q9UMF0
cor=0.86, p=3.6e−11
231: PCSK9|Q8NBP7

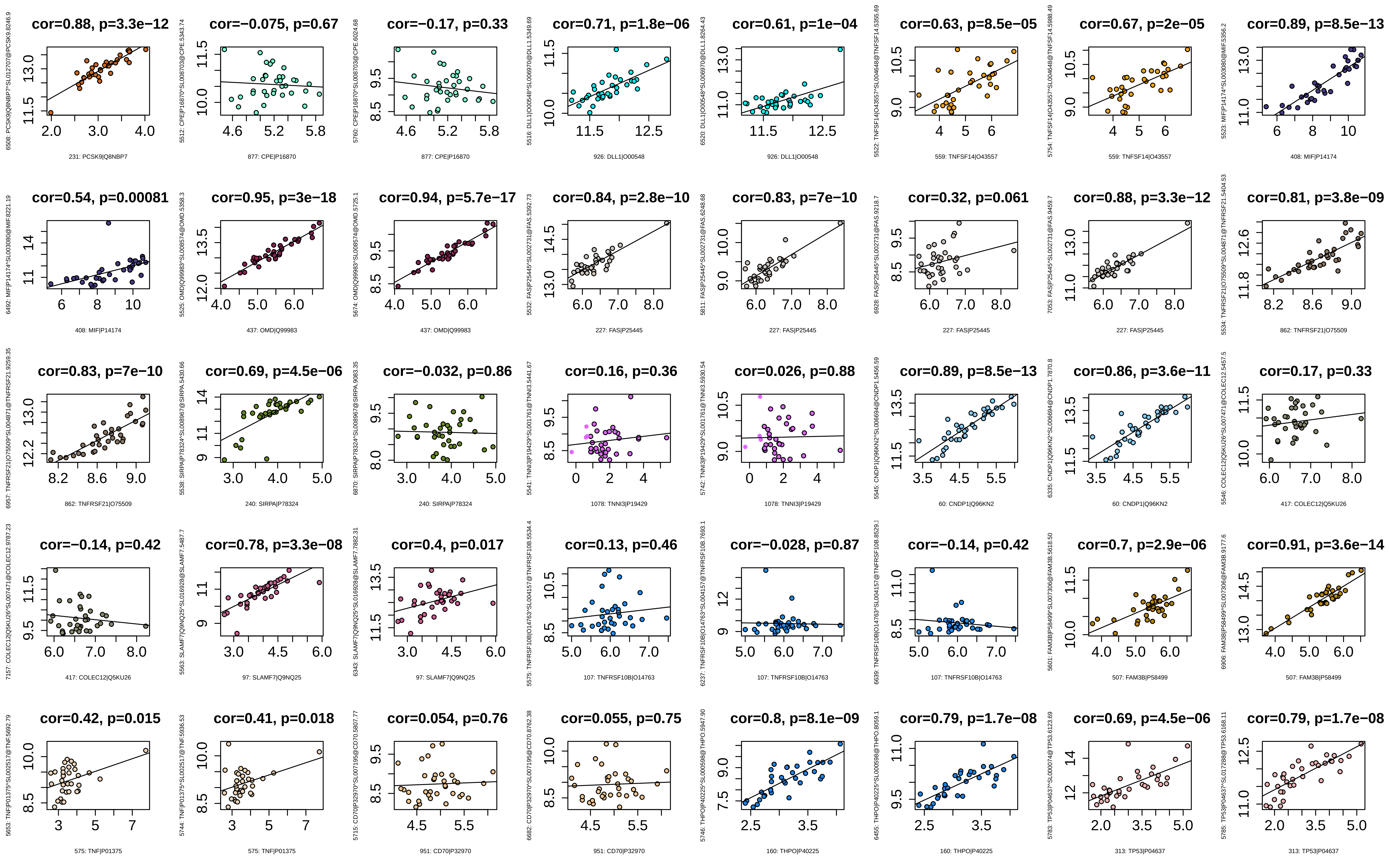
cor=0.88, p=3.3e−12
6508: PCSK9|Q8NBP7^SL012707@PCSK9.8246.9
231: PCSK9|Q8NBP7
cor=−0.075, p=0.67
5512: CPE|P16870^SL008703@CPE.5343.74
877: CPE|P16870
cor=−0.17, p=0.33
5760: CPE|P16870^SL008703@CPE.6024.68
877: CPE|P16870
cor=0.71, p=1.8e−06
5516: DLL1|O00548^SL006970@DLL1.5349.69
926: DLL1|O00548
cor=0.61, p=1e−04
6520: DLL1|O00548^SL006970@DLL1.8264.43
926: DLL1|O00548
cor=0.63, p=8.5e−05
5522: TNFSF14|O43557^SL004648@TNFSF14.5355.69
559: TNFSF14|O43557
cor=0.67, p=2e−05
5754: TNFSF14|O43557^SL004648@TNFSF14.5988.49
559: TNFSF14|O43557
cor=0.89, p=8.5e−13
5523: MIF|P14174^SL003080@MIF.5356.2
408: MIF|P14174
cor=0.54, p=0.00081
6492: MIF|P14174^SL003080@MIF.8221.19
408: MIF|P14174
cor=0.95, p=3e−18
5525: OMD|Q99983^SL008574@OMD.5358.3
437: OMD|Q99983
cor=0.94, p=5.7e−17
5674: OMD|Q99983^SL008574@OMD.5725.1
437: OMD|Q99983
cor=0.84, p=2.8e−10
5532: FAS|P25445^SL002731@FAS.5392.73
227: FAS|P25445
cor=0.83, p=7e−10
5811: FAS|P25445^SL002731@FAS.6248.68
227: FAS|P25445
cor=0.32, p=0.061
6928: FAS|P25445^SL002731@FAS.9218.7
227: FAS|P25445
cor=0.88, p=3.3e−12
7053: FAS|P25445^SL002731@FAS.9459.7
227: FAS|P25445
cor=0.81, p=3.8e−09
5534: TNFRSF21|O75509^SL004871@TNFRSF21.5404.53
862: TNFRSF21|O75509
cor=0.83, p=7e−10
6957: TNFRSF21|O75509^SL004871@TNFRSF21.9259.35
862: TNFRSF21|O75509
cor=0.69, p=4.5e−06
5538: SIRPA|P78324^SL008967@SIRPA.5430.66
240: SIRPA|P78324
cor=−0.032, p=0.86
6870: SIRPA|P78324^SL008967@SIRPA.9083.35
240: SIRPA|P78324
cor=0.16, p=0.36
5541: TNNI3|P19429^SL001761@TNNI3.5441.67
1078: TNNI3|P19429
cor=0.026, p=0.88
5742: TNNI3|P19429^SL001761@TNNI3.5930.54
1078: TNNI3|P19429
cor=0.89, p=8.5e−13
5545: CNDP1|Q96KN2^SL006694@CNDP1.5456.59
60: CNDP1|Q96KN2
cor=0.86, p=3.6e−11
6335: CNDP1|Q96KN2^SL006694@CNDP1.7870.8
60: CNDP1|Q96KN2
cor=0.17, p=0.33
5546: COLEC12|Q5KU26^SL007471@COLEC12.5457.5
417: COLEC12|Q5KU26
cor=−0.14, p=0.42
7157: COLEC12|Q5KU26^SL007471@COLEC12.9787.23
417: COLEC12|Q5KU26
cor=0.78, p=3.3e−08
5563: SLAMF7|Q9NQ25^SL016928@SLAMF7.5487.7
97: SLAMF7|Q9NQ25
cor=0.4, p=0.017
6343: SLAMF7|Q9NQ25^SL016928@SLAMF7.7882.31
97: SLAMF7|Q9NQ25
cor=0.13, p=0.46
5575: TNFRSF10B|O14763^SL004157@TNFRSF10B.5534.4
107: TNFRSF10B|O14763
cor=−0.028, p=0.87
6237: TNFRSF10B|O14763^SL004157@TNFRSF10B.7693.1
107: TNFRSF10B|O14763
cor=−0.14, p=0.42
6639: TNFRSF10B|O14763^SL004157@TNFRSF10B.8529.1
107: TNFRSF10B|O14763
cor=0.7, p=2.9e−06
5601: FAM3B|P58499^SL007306@FAM3B.5618.50
507: FAM3B|P58499
cor=0.91, p=3.6e−14
6906: FAM3B|P58499^SL007306@FAM3B.9177.6
507: FAM3B|P58499
cor=0.42, p=0.015
5653: TNF|P01375^SL002517@TNF.5692.79
575: TNF|P01375
cor=0.41, p=0.018
5744: TNF|P01375^SL002517@TNF.5936.53
575: TNF|P01375
cor=0.054, p=0.76
5715: CD70|P32970^SL007195@CD70.5807.77
951: CD70|P32970
cor=0.055, p=0.75
6682: CD70|P32970^SL007195@CD70.8762.38
951: CD70|P32970
cor=0.8, p=8.1e−09
5746: THPO|P40225^SL000598@THPO.5947.90
160: THPO|P40225
cor=0.79, p=1.7e−08
6455: THPO|P40225^SL000598@THPO.8059.1
160: THPO|P40225
cor=0.69, p=4.5e−06
5783: TP53|P04637^SL000074@TP53.6123.69
313: TP53|P04637
cor=0.79, p=1.7e−08
5785: TP53|P04637^SL017288@TP53.6168.11
313: TP53|P04637

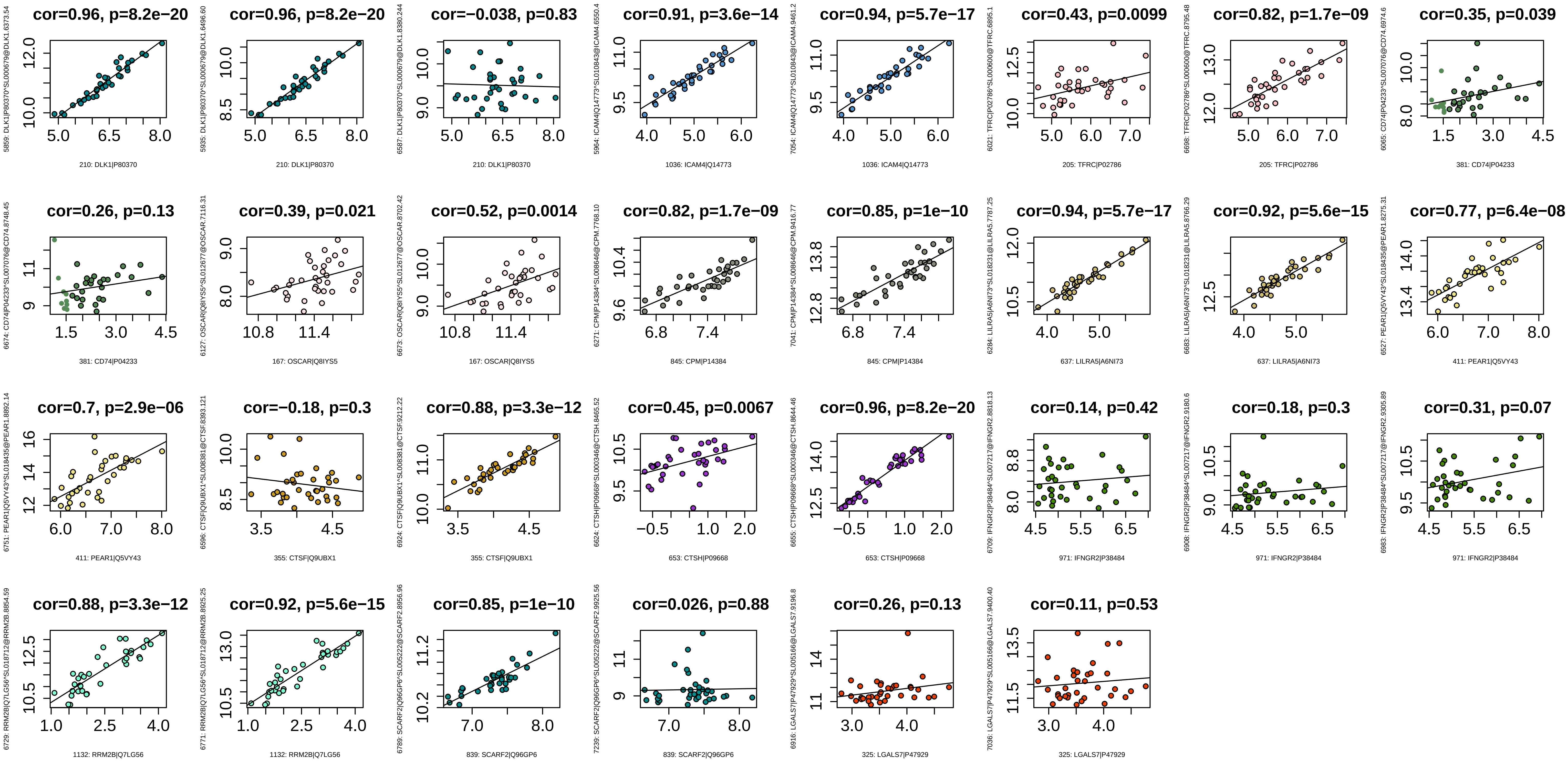
cor=0.96, p=8.2e−20
cor=0.96, p=8.2e−20
cor=−0.038, p=0.83
cor=0.91, p=3.6e−14
cor=0.94, p=5.7e−17
cor=0.43, p=0.0099
cor=0.82, p=1.7e−09
cor=0.35, p=0.039
210: DLK1|P80370
1036: ICAM4|Q14773
205: TFRC|P02786
381: CD74|P04233
cor=0.26, p=0.13
cor=0.39, p=0.021
cor=0.52, p=0.0014
cor=0.82, p=1.7e−09
cor=0.85, p=1e−10
cor=0.94, p=5.7e−17
cor=0.92, p=5.6e−15
cor=0.77, p=6.4e−08
167: OSCAR|Q8IYS5
845: CPM|P14384
637: LILRA5|A6NI73
411: PEAR1|Q5VY43
cor=0.7, p=2.9e−06
cor=−0.18, p=0.3
cor=0.88, p=3.3e−12
cor=0.45, p=0.0067
cor=0.96, p=8.2e−20
cor=0.14, p=0.42
cor=0.18, p=0.3
cor=0.31, p=0.07
355: CTSF|Q9UBX1
653: CTSH|P09668
971: IFNGR2|P38484
cor=0.88, p=3.3e−12
cor=0.92, p=5.6e−15
cor=0.85, p=1e−10
cor=0.026, p=0.88
cor=0.26, p=0.13
cor=0.11, p=0.53
1132: RRM2B|Q7LG56
839: SCARF2|Q96GP6
325: LGALS7|P47929

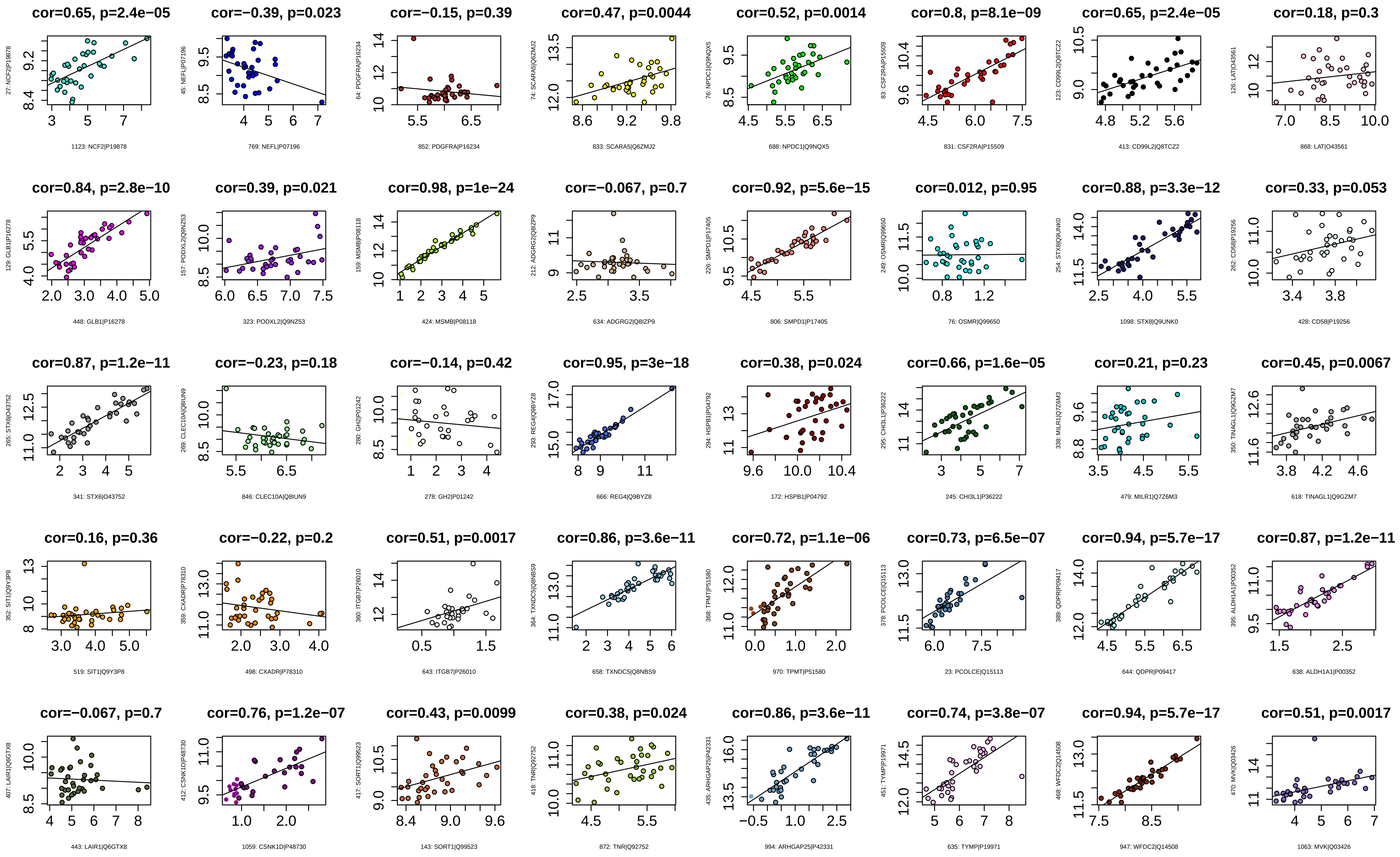

cor=0.65, p=2.4e−05
27: NCF2|P19878
1123: NCF2|P19878
cor=−0.39, p=0.023
45: NEFL|P07196
769: NEFL|P07196
cor=−0.15, p=0.39
64: PDGFRA|P16234
852: PDGFRA|P16234
cor=0.47, p=0.0044
74: SCARA5|Q6ZMJ2
833: SCARA5|Q6ZMJ2
cor=0.52, p=0.0014
76: NPDC1|Q9NQX5
688: NPDC1|Q9NQX5
cor=0.8, p=8.1e−09
83: CSF2RA|P15509
831: CSF2RA|P15509
cor=0.65, p=2.4e−05
123: CD99L2|Q8TCZ2
413: CD99L2|Q8TCZ2
cor=0.18, p=0.3
126: LAT|O43561
868: LAT|O43561
cor=0.84, p=2.8e−10
129: GLB1|P16278
448: GLB1|P16278
cor=0.39, p=0.021
157: PODXL2|Q9NZ53
323: PODXL2|Q9NZ53
cor=0.98, p=1e−24
159: MSMB|P08118
424: MSMB|P08118
cor=−0.067, p=0.7
212: ADGRG2|Q8IZP9
634: ADGRG2|Q8IZP9
cor=0.92, p=5.6e−15
226: SMPD1|P17405
806: SMPD1|P17405
cor=0.012, p=0.95
249: OSMR|Q99650
76: OSMR|Q99650
cor=0.88, p=3.3e−12
254: STX8|Q9UNK0
1098: STX8|Q9UNK0
cor=0.33, p=0.053
262: CD58|P19256
428: CD58|P19256
cor=0.87, p=1.2e−11
265: STX6|O43752
341: STX6|O43752
cor=−0.23, p=0.18
269: CLEC10A|Q8IUN9
846: CLEC10A|Q8IUN9
cor=−0.14, p=0.42
280: GH2|P01242
278: GH2|P01242
cor=0.95, p=3e−18
293: REG4|Q9BYZ8
666: REG4|Q9BYZ8
cor=0.38, p=0.024
294: HSPB1|P04792
172: HSPB1|P04792
cor=0.66, p=1.6e−05
295: CHI3L1|P36222
245: CHI3L1|P36222
cor=0.21, p=0.23
338: MILR1|Q7Z6M3
479: MILR1|Q7Z6M3
cor=0.45, p=0.0067
350: TINAGL1|Q9GZM7
618: TINAGL1|Q9GZM7
cor=0.16, p=0.36
352: SIT1|Q9Y3P8
519: SIT1|Q9Y3P8
cor=−0.22, p=0.2
359: CXADR|P78310
498: CXADR|P78310
cor=0.51, p=0.0017
360: ITGB7|P26010
643: ITGB7|P26010
cor=0.86, p=3.6e−11
364: TXNDC5|Q8NBS9
658: TXNDC5|Q8NBS9
cor=0.72, p=1.1e−06
368: TPMT|P51580
970: TPMT|P51580
cor=0.73, p=6.5e−07
378: PCOLCE|Q15113
23: PCOLCE|Q15113
cor=0.94, p=5.7e−17
389: QDPR|P09417
644: QDPR|P09417
cor=0.87, p=1.2e−11
395: ALDH1A1|P00352
638: ALDH1A1|P00352
cor=−0.067, p=0.7
407: LAIR1|Q6GTX8
443: LAIR1|Q6GTX8
cor=0.76, p=1.2e−07
412: CSNK1D|P48730
1059: CSNK1D|P48730
cor=0.43, p=0.0099
417: SORT1|Q99523
143: SORT1|Q99523
cor=0.38, p=0.024
418: TNR|Q92752
872: TNR|Q92752
cor=0.86, p=3.6e−11
435: ARHGAP25|P42331
994: ARHGAP25|P42331
cor=0.74, p=3.8e−07
451: TYMP|P19971
635: TYMP|P19971
cor=0.94, p=5.7e−17
468: WFDC2|Q14508
947: WFDC2|Q14508
cor=0.51, p=0.0017
470: MVK|Q03426
1063: MVK|Q03426

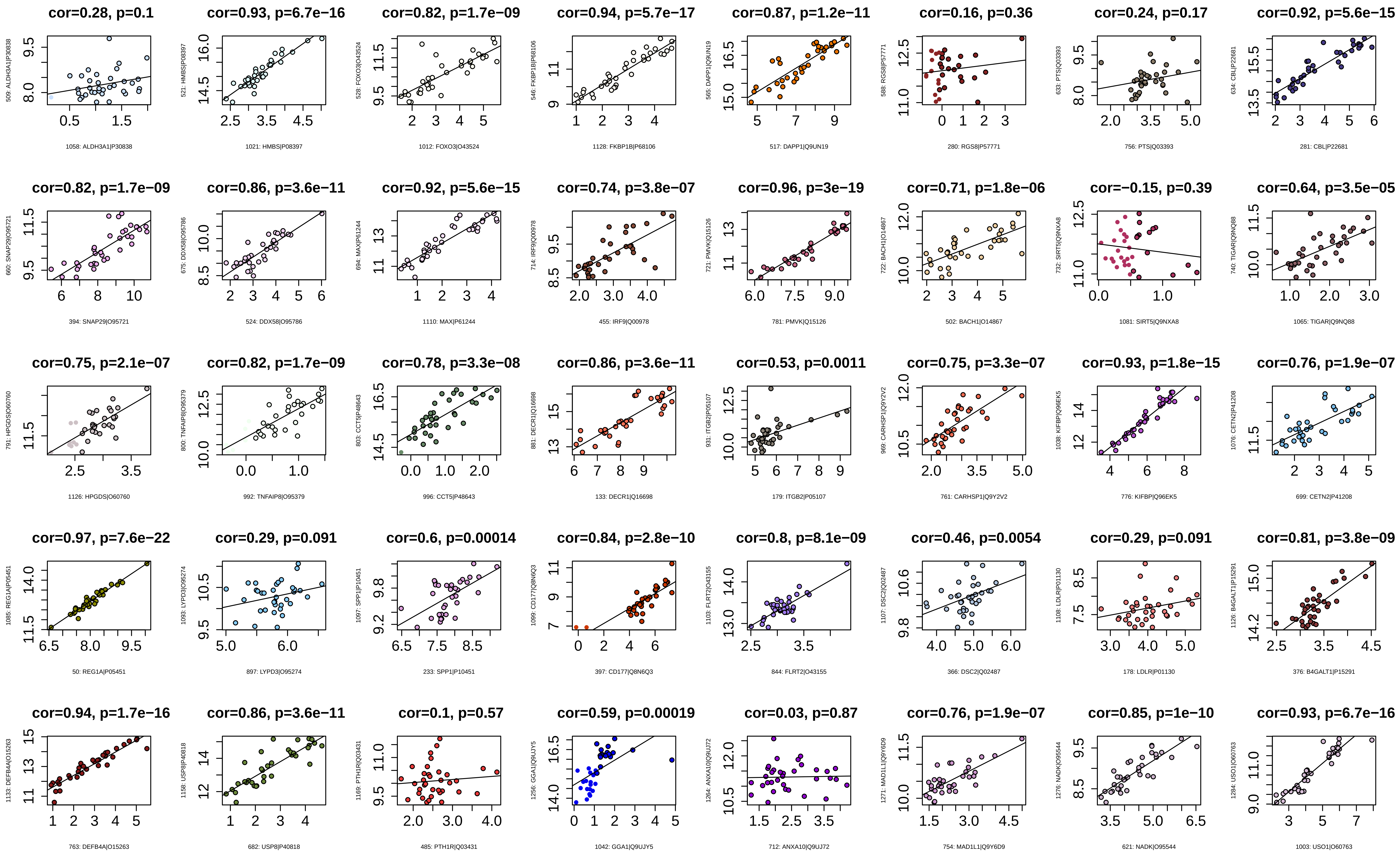
cor=0.28, p=0.1
1058: ALDH3A1|P30838
cor=0.93, p=6.7e−16
1021: HMBS|P08397
cor=0.82, p=1.7e−09
1012: FOXO3|O43524
cor=0.94, p=5.7e−17
1128: FKBP1B|P68106
cor=0.87, p=1.2e−11
517: DAPP1|Q9UN19
cor=0.16, p=0.36
280: RGS8|P57771
cor=0.24, p=0.17
756: PTS|Q03393
cor=0.92, p=5.6e−15
281: CBL|P22681
cor=0.82, p=1.7e−09
394: SNAP29|O95721
cor=0.86, p=3.6e−11
524: DDX58|O95786
cor=0.92, p=5.6e−15
1110: MAX|P61244
cor=0.74, p=3.8e−07
455: IRF9|Q00978
cor=0.96, p=3e−19
781: PMVK|Q15126
cor=0.71, p=1.8e−06
502: BACH1|O14867
cor=−0.15, p=0.39
1081: SIRT5|Q9NXA8
cor=0.64, p=3.5e−05
1065: TIGAR|Q9NQ88
cor=0.75, p=2.1e−07
1126: HPGDS|O60760
cor=0.82, p=1.7e−09
992: TNFAIP8|O95379
cor=0.78, p=3.3e−08
996: CCT5|P48643
cor=0.86, p=3.6e−11
133: DECR1|Q16698
cor=0.53, p=0.0011
179: ITGB2|P05107
cor=0.75, p=3.3e−07
761: CARHSP1|Q9Y2V2
cor=0.93, p=1.8e−15
776: KIFBP|Q96EK5
cor=0.76, p=1.9e−07
699: CETN2|P41208
cor=0.97, p=7.6e−22
50: REG1A|P05451
cor=0.29, p=0.091
897: LYPD3|O95274
cor=0.6, p=0.00014
233: SPP1|P10451
cor=0.84, p=2.8e−10
397: CD177|Q8N6Q3
cor=0.8, p=8.1e−09
844: FLRT2|O43155
cor=0.46, p=0.0054
366: DSC2|Q02487
cor=0.29, p=0.091
178: LDLR|P01130
cor=0.81, p=3.8e−09
376: B4GALT1|P15291
cor=0.94, p=1.7e−16
763: DEFB4A|O15263
cor=0.86, p=3.6e−11
682: USP8|P40818
cor=0.1, p=0.57
485: PTH1R|Q03431
cor=0.59, p=0.00019
1042: GGA1|Q9UJY5
cor=0.03, p=0.87
712: ANXA10|Q9UJ72
cor=0.76, p=1.9e−07
754: MAD1L1|Q9Y6D9
cor=0.85, p=1e−10
621: NADK|O95544
cor=0.93, p=6.7e−16
1003: USO1|O60763

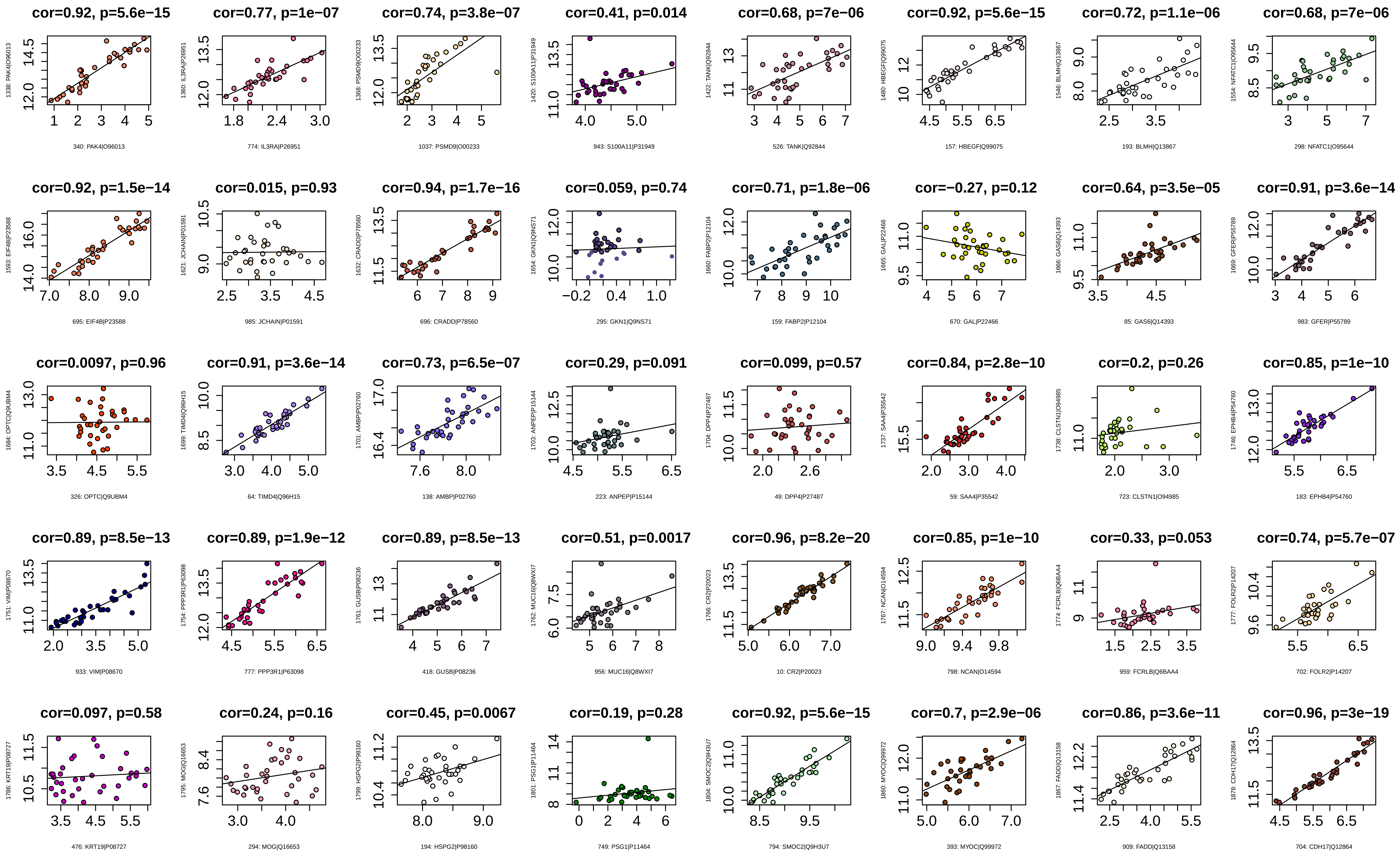
cor=0.92, p=5.6e−15
340: PAK4|O96013
cor=0.77, p=1e−07
774: IL3RA|P26951
cor=0.74, p=3.8e−07
1037: PSMD9|O00233
cor=0.41, p=0.014
943: S100A11|P31949
cor=0.68, p=7e−06
526: TANK|Q92844
cor=0.92, p=5.6e−15
157: HBEGF|Q99075
cor=0.72, p=1.1e−06
193: BLMH|Q13867
cor=0.68, p=7e−06
298: NFATC1|O95644
cor=0.92, p=1.5e−14
695: EIF4B|P23588
cor=0.015, p=0.93
985: JCHAIN|P01591
cor=0.94, p=1.7e−16
696: CRADD|P78560
cor=0.059, p=0.74
295: GKN1|Q9NS71
cor=0.71, p=1.8e−06
159: FABP2|P12104
cor=−0.27, p=0.12
670: GAL|P22466
cor=0.64, p=3.5e−05
85: GAS6|Q14393
cor=0.91, p=3.6e−14
983: GFER|P55789
cor=0.0097, p=0.96
326: OPTC|Q9UBM4
cor=0.91, p=3.6e−14
64: TIMD4|Q96H15
cor=0.73, p=6.5e−07
138: AMBP|P02760
cor=0.29, p=0.091
223: ANPEP|P15144
cor=0.099, p=0.57
49: DPP4|P27487
cor=0.84, p=2.8e−10
59: SAA4|P35542
cor=0.2, p=0.26
723: CLSTN1|O94985
cor=0.85, p=1e−10
183: EPHB4|P54760
cor=0.89, p=8.5e−13
933: VIM|P08670
cor=0.89, p=1.9e−12
777: PPP3R1|P63098
cor=0.89, p=8.5e−13
418: GUSB|P08236
cor=0.51, p=0.0017
956: MUC16|Q8WXI7
cor=0.96, p=8.2e−20
10: CR2|P20023
cor=0.85, p=1e−10
798: NCAN|O14594
cor=0.33, p=0.053
959: FCRLB|Q6BAA4
cor=0.74, p=5.7e−07
702: FOLR2|P14207
cor=0.097, p=0.58
476: KRT19|P08727
cor=0.24, p=0.16
294: MOG|Q16653
cor=0.45, p=0.0067
194: HSPG2|P98160
cor=0.19, p=0.28
749: PSG1|P11464
cor=0.92, p=5.6e−15
794: SMOC2|Q9H3U7
cor=0.7, p=2.9e−06
393: MYOC|Q99972
cor=0.86, p=3.6e−11
909: FADD|Q13158
cor=0.96, p=3e−19
704: CDH17|Q12864

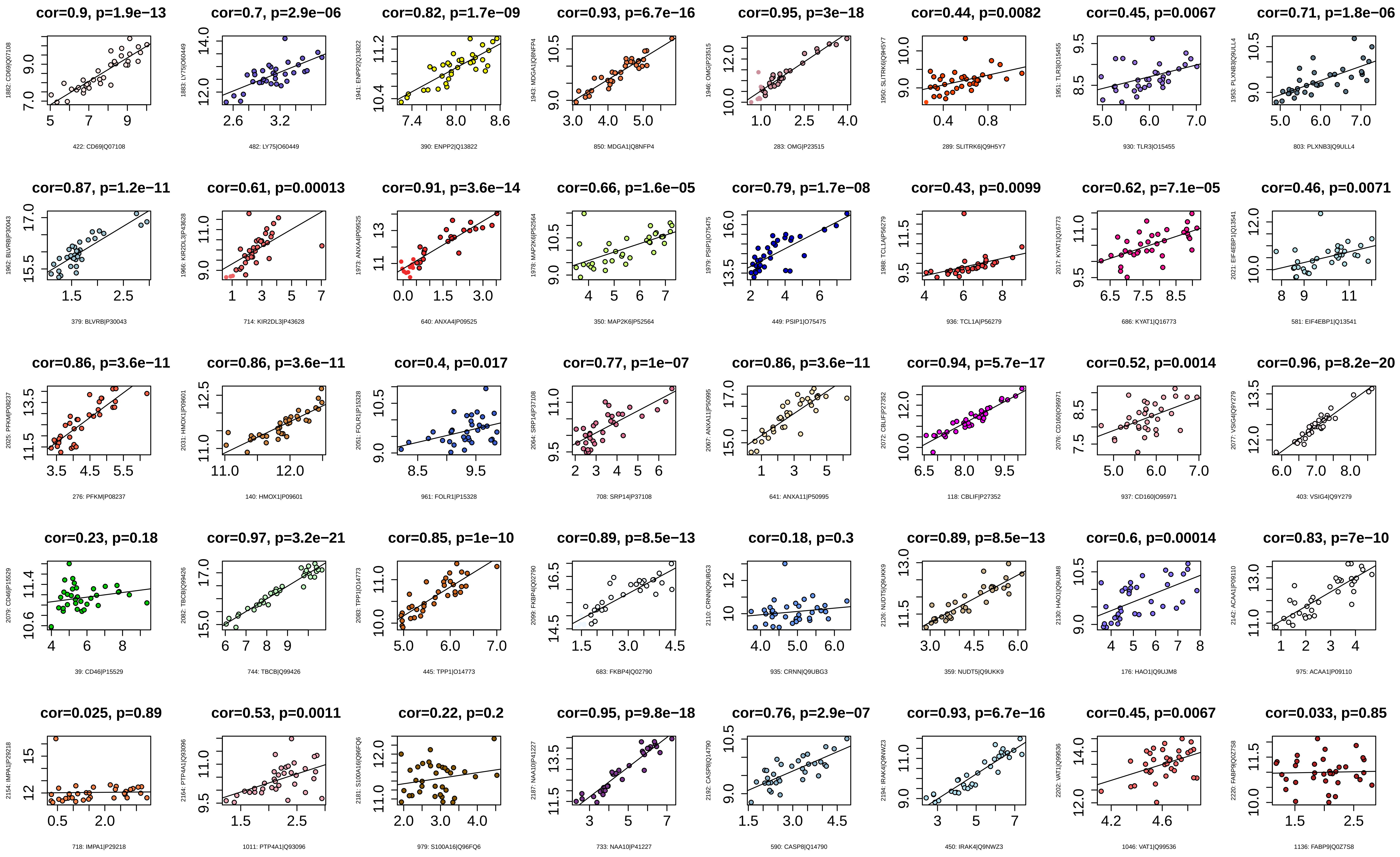
cor=0.9, p=1.9e−13
422: CD69|Q07108
cor=0.7, p=2.9e−06
482: LY75|O60449
cor=0.82, p=1.7e−09
390: ENPP2|Q13822
cor=0.93, p=6.7e−16
850: MDGA1|Q8NFP4
cor=0.95, p=3e−18
283: OMG|P23515
cor=0.44, p=0.0082
289: SLITRK6|Q9H5Y7
cor=0.45, p=0.0067
930: TLR3|O15455
cor=0.71, p=1.8e−06
803: PLXNB3|Q9ULL4
cor=0.87, p=1.2e−11
379: BLVRB|P30043
cor=0.61, p=0.00013
714: KIR2DL3|P43628
cor=0.91, p=3.6e−14
640: ANXA4|P09525
cor=0.66, p=1.6e−05
350: MAP2K6|P52564
cor=0.79, p=1.7e−08
449: PSIP1|O75475
cor=0.43, p=0.0099
936: TCL1A|P56279
cor=0.62, p=7.1e−05
686: KYAT1|Q16773
cor=0.46, p=0.0071
581: EIF4EBP1|Q13541
cor=0.86, p=3.6e−11
276: PFKM|P08237
cor=0.86, p=3.6e−11
140: HMOX1|P09601
cor=0.4, p=0.017
961: FOLR1|P15328
cor=0.77, p=1e−07
708: SRP14|P37108
cor=0.86, p=3.6e−11
641: ANXA11|P50995
cor=0.94, p=5.7e−17
118: CBLIF|P27352
cor=0.52, p=0.0014
937: CD160|O95971
cor=0.96, p=8.2e−20
403: VSIG4|Q9Y279
cor=0.23, p=0.18
39: CD46|P15529
cor=0.97, p=3.2e−21
744: TBCB|Q99426
cor=0.85, p=1e−10
445: TPP1|O14773
cor=0.89, p=8.5e−13
683: FKBP4|Q02790
cor=0.18, p=0.3
935: CRNN|Q9UBG3
cor=0.89, p=8.5e−13
359: NUDT5|Q9UKK9
cor=0.6, p=0.00014
176: HAO1|Q9UJM8
cor=0.83, p=7e−10
975: ACAA1|P09110
cor=0.025, p=0.89
718: IMPA1|P29218
cor=0.53, p=0.0011
1011: PTP4A1|Q93096
cor=0.22, p=0.2
979: S100A16|Q96FQ6
cor=0.95, p=9.8e−18
733: NAA10|P41227
cor=0.76, p=2.9e−07
590: CASP8|Q14790
cor=0.93, p=6.7e−16
450: IRAK4|Q9NWZ3
cor=0.45, p=0.0067
1046: VAT1|Q99536
cor=0.033, p=0.85
1136: FABP9|Q0Z7S8

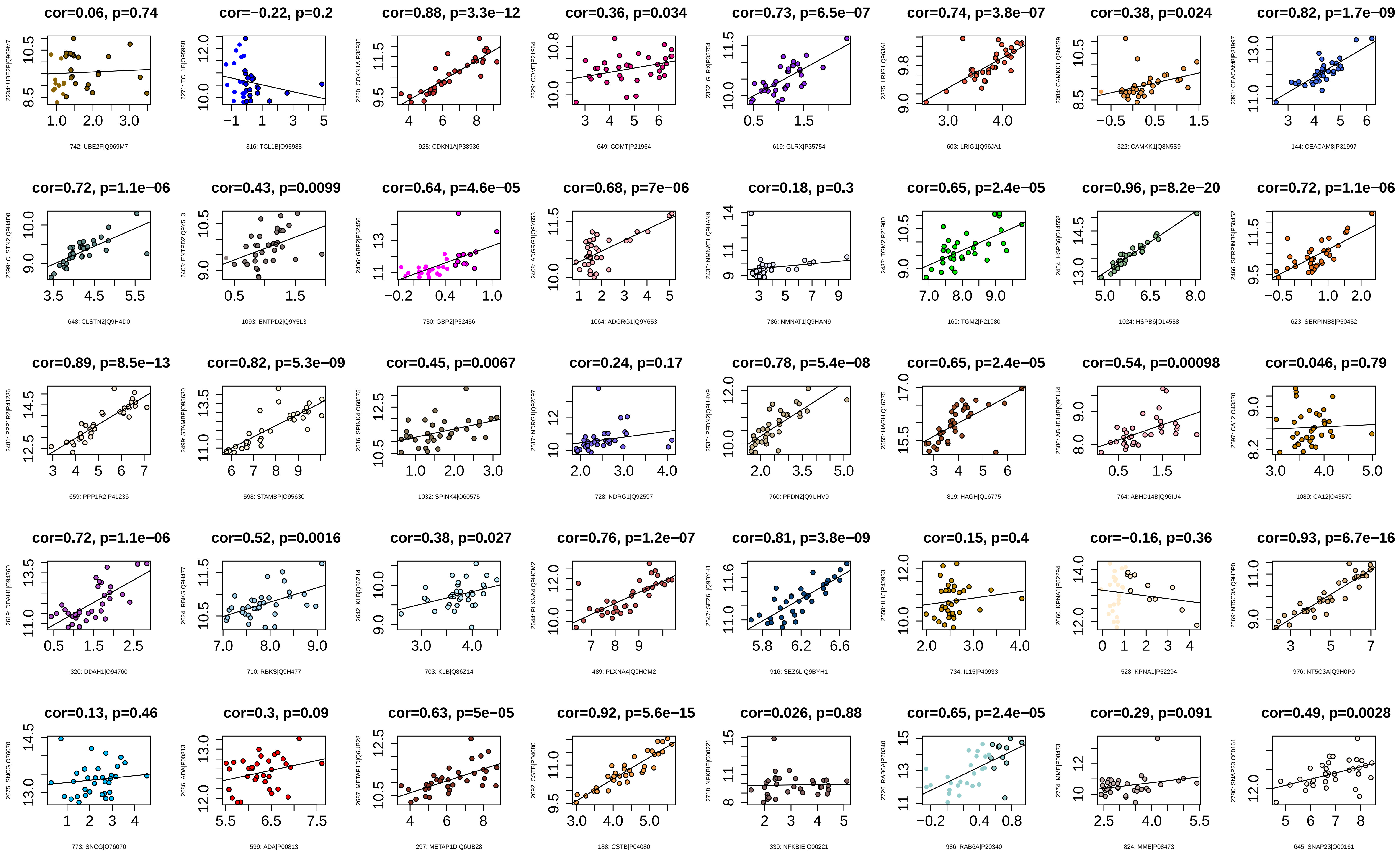
cor=0.06, p=0.74
742: UBE2F|Q969M7
cor=−0.22, p=0.2
316: TCL1B|O95988
cor=0.88, p=3.3e−12
925: CDKN1A|P38936
cor=0.36, p=0.034
649: COMT|P21964
cor=0.73, p=6.5e−07
619: GLRX|P35754
cor=0.74, p=3.8e−07
603: LRIG1|Q96JA1
cor=0.38, p=0.024
322: CAMKK1|Q8N5S9
cor=0.82, p=1.7e−09
144: CEACAM8|P31997
cor=0.72, p=1.1e−06
648: CLSTN2|Q9H4D0
cor=0.43, p=0.0099
1093: ENTPD2|Q9Y5L3
cor=0.64, p=4.6e−05
730: GBP2|P32456
cor=0.68, p=7e−06
1064: ADGRG1|Q9Y653
cor=0.18, p=0.3
786: NMNAT1|Q9HAN9
cor=0.65, p=2.4e−05
169: TGM2|P21980
cor=0.96, p=8.2e−20
1024: HSPB6|O14558
cor=0.72, p=1.1e−06
623: SERPINB8|P50452
cor=0.89, p=8.5e−13
659: PPP1R2|P41236
cor=0.82, p=5.3e−09
598: STAMBP|O95630
cor=0.45, p=0.0067
1032: SPINK4|O60575
cor=0.24, p=0.17
728: NDRG1|Q92597
cor=0.78, p=5.4e−08
760: PFDN2|Q9UHV9
cor=0.65, p=2.4e−05
819: HAGH|Q16775
cor=0.54, p=0.00098
764: ABHD14B|Q96IU4
cor=0.046, p=0.79
1089: CA12|O43570
cor=0.72, p=1.1e−06
320: DDAH1|O94760
cor=0.52, p=0.0016
710: RBKS|Q9H477
cor=0.38, p=0.027
703: KLB|Q86Z14
cor=0.76, p=1.2e−07
489: PLXNA4|Q9HCM2
cor=0.81, p=3.8e−09
916: SEZ6L|Q9BYH1
cor=0.15, p=0.4
734: IL15|P40933
cor=−0.16, p=0.36
528: KPNA1|P52294
cor=0.93, p=6.7e−16
976: NT5C3A|Q9H0P0
cor=0.13, p=0.46
773: SNCG|O76070
cor=0.3, p=0.09
599: ADA|P00813
cor=0.63, p=5e−05
297: METAP1D|Q6UB28
cor=0.92, p=5.6e−15
188: CSTB|P04080
cor=0.026, p=0.88
339: NFKBIE|O00221
cor=0.65, p=2.4e−05
986: RAB6A|P20340
cor=0.29, p=0.091
824: MME|P08473
cor=0.49, p=0.0028
645: SNAP23|O00161

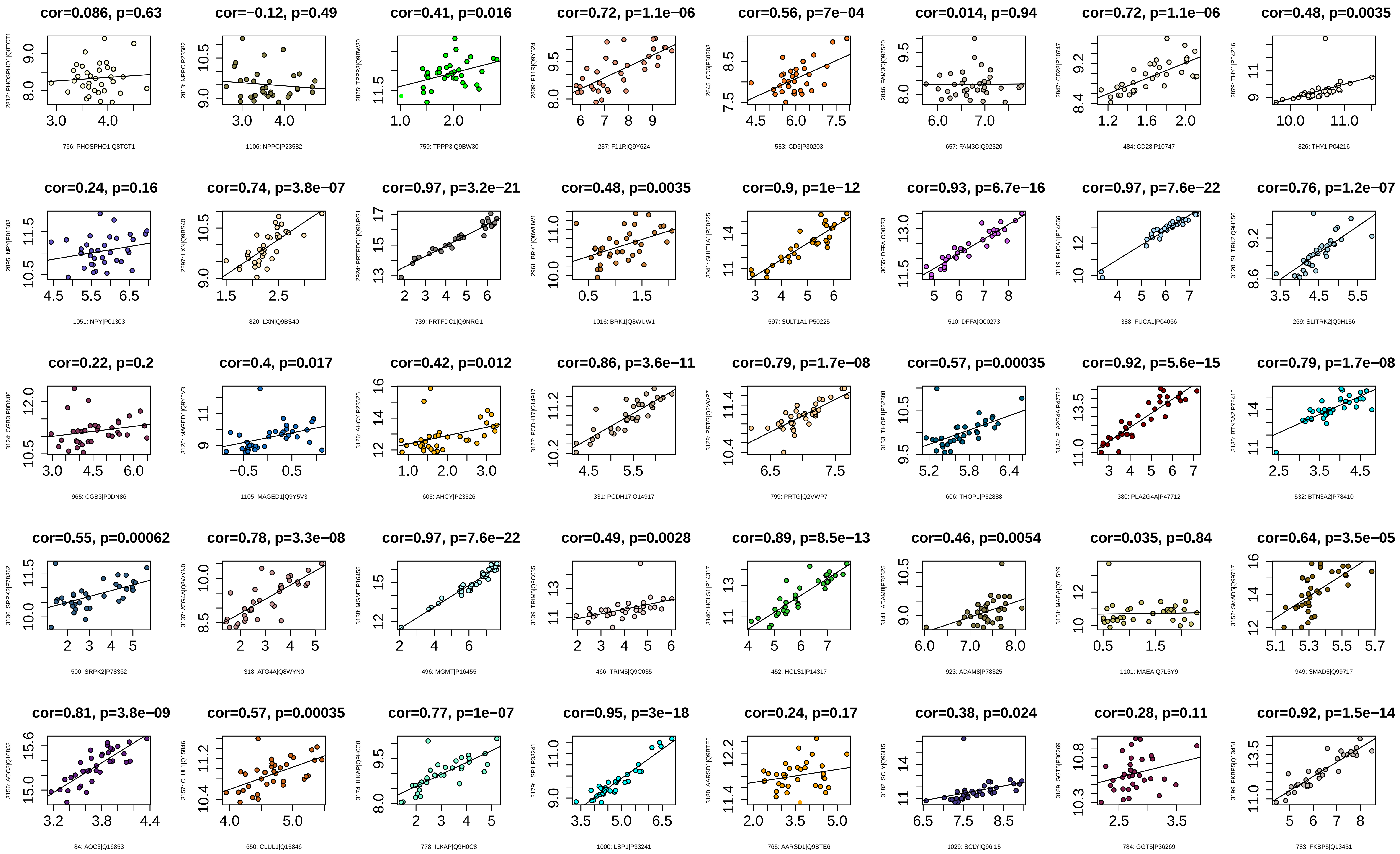
cor=0.086, p=0.63
2812: PHOSPHO1|Q8TCT1
766: PHOSPHO1|Q8TCT1
cor=−0.12, p=0.49
2813: NPPC|P23582
1106: NPPC|P23582
cor=0.41, p=0.016
2825: TPPP3|Q9BW30
759: TPPP3|Q9BW30
cor=0.72, p=1.1e−06
2839: F11R|Q9Y624
237: F11R|Q9Y624
cor=0.56, p=7e−04
2845: CD6|P30203
553: CD6|P30203
cor=0.014, p=0.94
2846: FAM3C|Q92520
657: FAM3C|Q92520
cor=0.72, p=1.1e−06
2847: CD28|P10747
484: CD28|P10747
cor=0.48, p=0.0035
2879: THY1|P04216
826: THY1|P04216
cor=0.24, p=0.16
2895: NPY|P01303
1051: NPY|P01303
cor=0.74, p=3.8e−07
2897: LXN|Q9BS40
820: LXN|Q9BS40
cor=0.97, p=3.2e−21
2924: PRTFDC1|Q9NRG1
739: PRTFDC1|Q9NRG1
cor=0.48, p=0.0035
2961: BRK1|Q8WUW1
1016: BRK1|Q8WUW1
cor=0.9, p=1e−12
3041: SULT1A1|P50225
597: SULT1A1|P50225
cor=0.93, p=6.7e−16
3055: DFFA|O00273
510: DFFA|O00273
cor=0.97, p=7.6e−22
3119: FUCA1|P04066
388: FUCA1|P04066
cor=0.76, p=1.2e−07
3120: SLITRK2|Q9H156
269: SLITRK2|Q9H156
cor=0.22, p=0.2
3124: CGB3|P0DN86
965: CGB3|P0DN86
cor=0.4, p=0.017
3125: MAGED1|Q9Y5V3
1105: MAGED1|Q9Y5V3
cor=0.42, p=0.012
3126: AHCY|P23526
605: AHCY|P23526
cor=0.86, p=3.6e−11
3127: PCDH17|O14917
331: PCDH17|O14917
cor=0.79, p=1.7e−08
3128: PRTG|Q2VWP7
799: PRTG|Q2VWP7
cor=0.57, p=0.00035
3133: THOP1|P52888
606: THOP1|P52888
cor=0.92, p=5.6e−15
3134: PLA2G4A|P47712
380: PLA2G4A|P47712
cor=0.79, p=1.7e−08
3135: BTN3A2|P78410
532: BTN3A2|P78410
cor=0.55, p=0.00062
3136: SRPK2|P78362
500: SRPK2|P78362
cor=0.78, p=3.3e−08
3137: ATG4A|Q8WYN0
318: ATG4A|Q8WYN0
cor=0.97, p=7.6e−22
3138: MGMT|P16455
496: MGMT|P16455
cor=0.49, p=0.0028
3139: TRIM5|Q9C035
466: TRIM5|Q9C035
cor=0.89, p=8.5e−13
3140: HCLS1|P14317
452: HCLS1|P14317
cor=0.46, p=0.0054
3141: ADAM8|P78325
923: ADAM8|P78325
cor=0.035, p=0.84
3151: MAEA|Q7L5Y9
1101: MAEA|Q7L5Y9
cor=0.64, p=3.5e−05
3152: SMAD5|Q99717
949: SMAD5|Q99717
cor=0.81, p=3.8e−09
3156: AOC3|Q16853
84: AOC3|Q16853
cor=0.57, p=0.00035
3157: CLUL1|Q15846
650: CLUL1|Q15846
cor=0.77, p=1e−07
3174: ILKAP|Q9H0C8
778: ILKAP|Q9H0C8
cor=0.95, p=3e−18
3179: LSP1|P33241
1000: LSP1|P33241
cor=0.24, p=0.17
3180: AARSD1|Q9BTE6
765: AARSD1|Q9BTE6
cor=0.38, p=0.024
3182: SCLY|Q96I15
1029: SCLY|Q96I15
cor=0.28, p=0.11
3189: GGT5|P36269
784: GGT5|P36269
cor=0.92, p=1.5e−14
3199: FKBP5|Q13451
783: FKBP5|Q13451

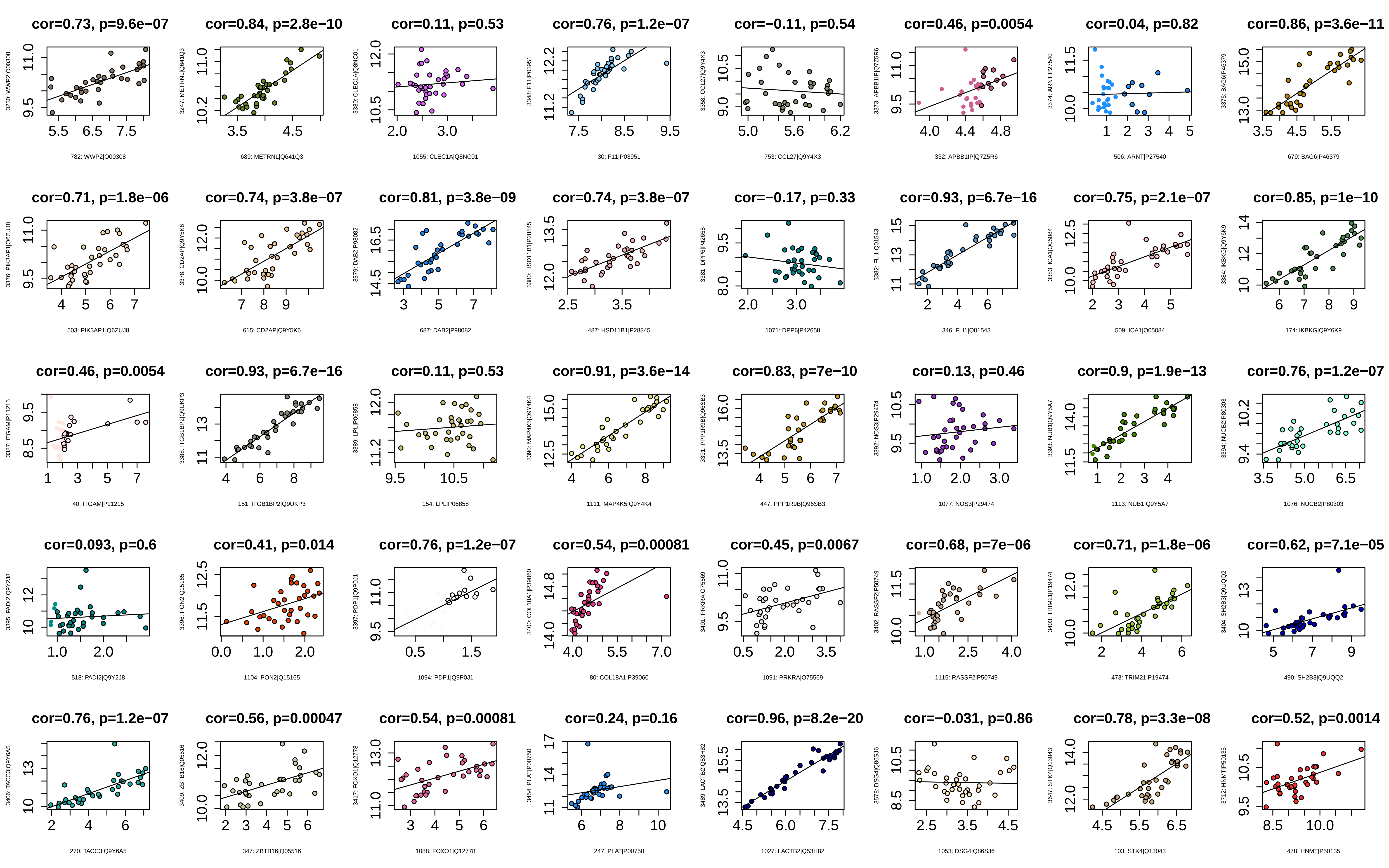

cor=0.73, p=9.6e−07
3230: WWP2|O00308
782: WWP2|O00308
cor=0.84, p=2.8e−10
3247: METRNL|Q641Q3
689: METRNL|Q641Q3
cor=0.11, p=0.53
3330: CLEC1A|Q8NC01
1055: CLEC1A|Q8NC01
cor=0.76, p=1.2e−07
3348: F11|P03951
30: F11|P03951
cor=−0.11, p=0.54
3358: CCL27|Q9Y4X3
753: CCL27|Q9Y4X3
cor=0.46, p=0.0054
3373: APBB1IP|Q7Z5R6
332: APBB1IP|Q7Z5R6
cor=0.04, p=0.82
3374: ARNT|P27540
506: ARNT|P27540
cor=0.86, p=3.6e−11
3375: BAG6|P46379
679: BAG6|P46379
cor=0.71, p=1.8e−06
3376: PIK3AP1|Q6ZUJ8
503: PIK3AP1|Q6ZUJ8
cor=0.74, p=3.8e−07
3378: CD2AP|Q9Y5K6
615: CD2AP|Q9Y5K6
cor=0.81, p=3.8e−09
3379: DAB2|P98082
687: DAB2|P98082
cor=0.74, p=3.8e−07
3380: HSD11B1|P28845
487: HSD11B1|P28845
cor=−0.17, p=0.33
3381: DPP6|P42658
1071: DPP6|P42658
cor=0.93, p=6.7e−16
3382: FLI1|Q01543
346: FLI1|Q01543
cor=0.75, p=2.1e−07
3383: ICA1|Q05084
509: ICA1|Q05084
cor=0.85, p=1e−10
3384: IKBKG|Q9Y6K9
174: IKBKG|Q9Y6K9
cor=0.46, p=0.0054
3387: ITGAM|P11215
40: ITGAM|P11215
cor=0.93, p=6.7e−16
3388: ITGB1BP2|Q9UKP3
151: ITGB1BP2|Q9UKP3
cor=0.11, p=0.53
3389: LPL|P06858
154: LPL|P06858
cor=0.91, p=3.6e−14
3390: MAP4K5|Q9Y4K4
1111: MAP4K5|Q9Y4K4
cor=0.83, p=7e−10
3391: PPP1R9B|Q96SB3
447: PPP1R9B|Q96SB3
cor=0.13, p=0.46
3392: NOS3|P29474
1077: NOS3|P29474
cor=0.9, p=1.9e−13
3393: NUB1|Q9Y5A7
1113: NUB1|Q9Y5A7
cor=0.76, p=1.2e−07
3394: NUCB2|P80303
1076: NUCB2|P80303
cor=0.093, p=0.6
3395: PADI2|Q9Y2J8
518: PADI2|Q9Y2J8
cor=0.41, p=0.014
3396: PON2|Q15165
1104: PON2|Q15165
cor=0.76, p=1.2e−07
3397: PDP1|Q9P0J1
1094: PDP1|Q9P0J1
cor=0.54, p=0.00081
3400: COL18A1|P39060
80: COL18A1|P39060
cor=0.45, p=0.0067
3401: PRKRA|O75569
1091: PRKRA|O75569
cor=0.68, p=7e−06
3402: RASSF2|P50749
1115: RASSF2|P50749
cor=0.71, p=1.8e−06
3403: TRIM21|P19474
473: TRIM21|P19474
cor=0.62, p=7.1e−05
3404: SH2B3|Q9UQQ2
490: SH2B3|Q9UQQ2
cor=0.76, p=1.2e−07
3406: TACC3|Q9Y6A5
270: TACC3|Q9Y6A5
cor=0.56, p=0.00047
3409: ZBTB16|Q05516
347: ZBTB16|Q05516
cor=0.54, p=0.00081
3417: FOXO1|Q12778
1088: FOXO1|Q12778
cor=0.24, p=0.16
3454: PLAT|P00750
247: PLAT|P00750
cor=0.96, p=8.2e−20
3489: LACTB2|Q53H82
1027: LACTB2|Q53H82
cor=−0.031, p=0.86
3578: DSG4|Q86SJ6
1053: DSG4|Q86SJ6
cor=0.78, p=3.3e−08
3647: STK4|Q13043
103: STK4|Q13043
cor=0.52, p=0.0014
3712: HNMT|P50135
478: HNMT|P50135

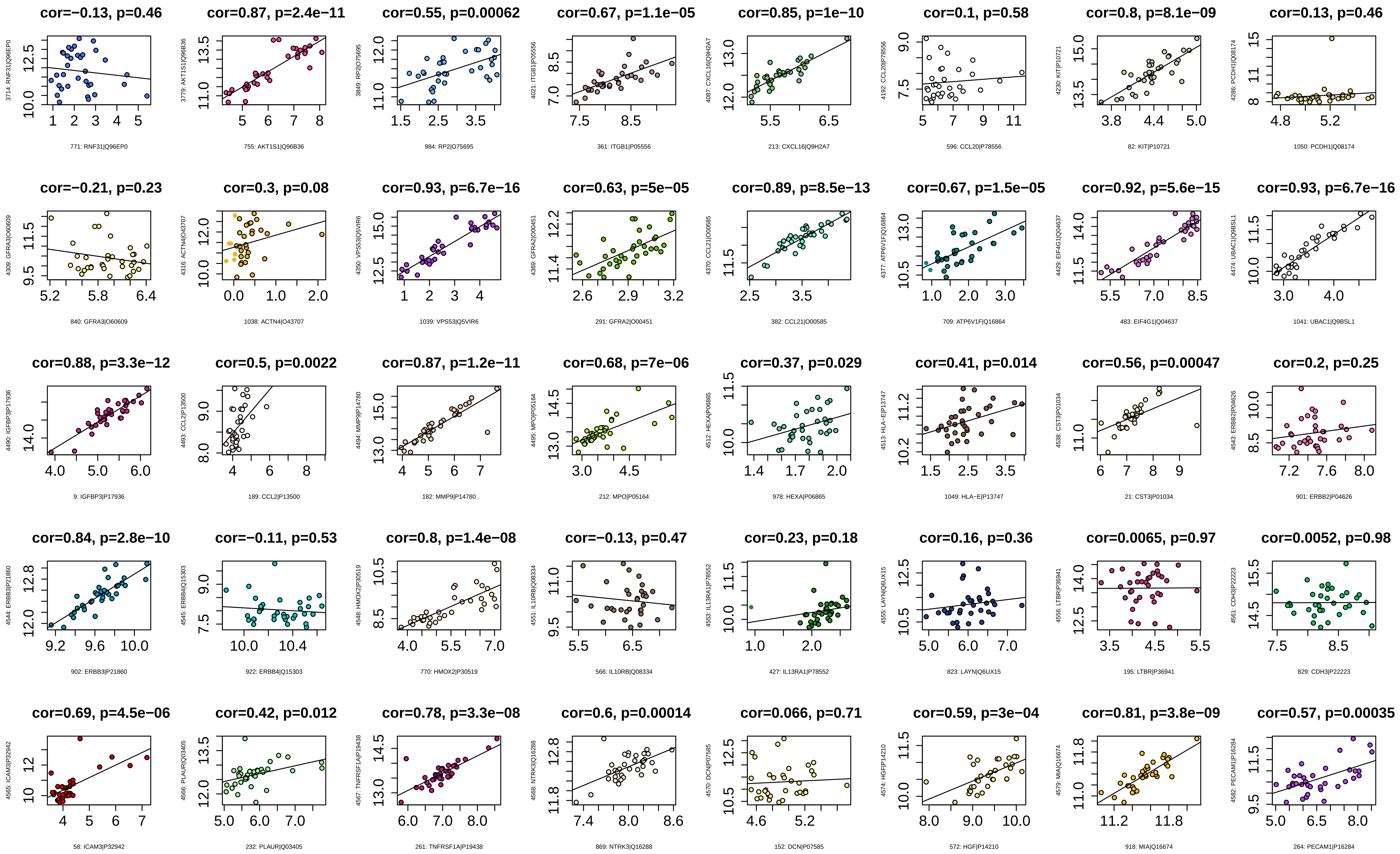

cor=−0.13, p=0.46
3714: RNF31|Q96EP0
771: RNF31|Q96EP0
cor=0.87, p=2.4e−11
3779: AKT1S1|Q96B36
755: AKT1S1|Q96B36
cor=0.55, p=0.00062
3849: RP2|O75695
984: RP2|O75695
cor=0.67, p=1.1e−05
4021: ITGB1|P05556
361: ITGB1|P05556
cor=0.85, p=1e−10
4087: CXCL16|Q9H2A7
213: CXCL16|Q9H2A7
cor=0.1, p=0.58
4192: CCL20|P78556
596: CCL20|P78556
cor=0.8, p=8.1e−09
4230: KIT|P10721
82: KIT|P10721
cor=0.13, p=0.46
4286: PCDH1|Q08174
1050: PCDH1|Q08174
cor=−0.21, p=0.23
4309: GFRA3|O60609
840: GFRA3|O60609
cor=0.3, p=0.08
4316: ACTN4|O43707
1038: ACTN4|O43707
cor=0.93, p=6.7e−16
4350: VPS53|Q5VIR6
1039: VPS53|Q5VIR6
cor=0.63, p=5e−05
4369: GFRA2|O00451
291: GFRA2|O00451
cor=0.89, p=8.5e−13
4370: CCL21|O00585
382: CCL21|O00585
cor=0.67, p=1.5e−05
4377: ATP6V1F|Q16864
709: ATP6V1F|Q16864
cor=0.92, p=5.6e−15
4429: EIF4G1|Q04637
483: EIF4G1|Q04637
cor=0.93, p=6.7e−16
4474: UBAC1|Q9BSL1
1041: UBAC1|Q9BSL1
cor=0.88, p=3.3e−12
4490: IGFBP3|P17936
9: IGFBP3|P17936
cor=0.5, p=0.0022
4493: CCL2|P13500
189: CCL2|P13500
cor=0.87, p=1.2e−11
4494: MMP9|P14780
182: MMP9|P14780
cor=0.68, p=7e−06
4495: MPO|P05164
212: MPO|P05164
cor=0.37, p=0.029
4512: HEXA|P06865
978: HEXA|P06865
cor=0.41, p=0.014
4513: HLA−E|P13747
1049: HLA−E|P13747
cor=0.56, p=0.00047
4538: CST3|P01034
21: CST3|P01034
cor=0.2, p=0.25
4543: ERBB2|P04626
901: ERBB2|P04626
cor=0.84, p=2.8e−10
4544: ERBB3|P21860
902: ERBB3|P21860
cor=−0.11, p=0.53
4545: ERBB4|Q15303
922: ERBB4|Q15303
cor=0.8, p=1.4e−08
4548: HMOX2|P30519
770: HMOX2|P30519
cor=−0.13, p=0.47
4551: IL10RB|Q08334
566: IL10RB|Q08334
cor=0.23, p=0.18
4553: IL13RA1|P78552
427: IL13RA1|P78552
cor=0.16, p=0.36
4555: LAYN|Q6UX15
823: LAYN|Q6UX15
cor=0.0065, p=0.97
4556: LTBR|P36941
195: LTBR|P36941
cor=0.0052, p=0.98
4561: CDH3|P22223
829: CDH3|P22223
cor=0.69, p=4.5e−06
4565: ICAM3|P32942
58: ICAM3|P32942
cor=0.42, p=0.012
4566: PLAUR|Q03405
232: PLAUR|Q03405
cor=0.78, p=3.3e−08
4567: TNFRSF1A|P19438
261: TNFRSF1A|P19438
cor=0.6, p=0.00014
4568: NTRK3|Q16288
869: NTRK3|Q16288
cor=0.066, p=0.71
4570: DCN|P07585
152: DCN|P07585
cor=0.59, p=3e−04
4574: HGF|P14210
572: HGF|P14210
cor=0.81, p=3.8e−09
4579: MIA|Q16674
918: MIA|Q16674
cor=0.57, p=0.00035
4582: PECAM1|P16284
264: PECAM1|P16284

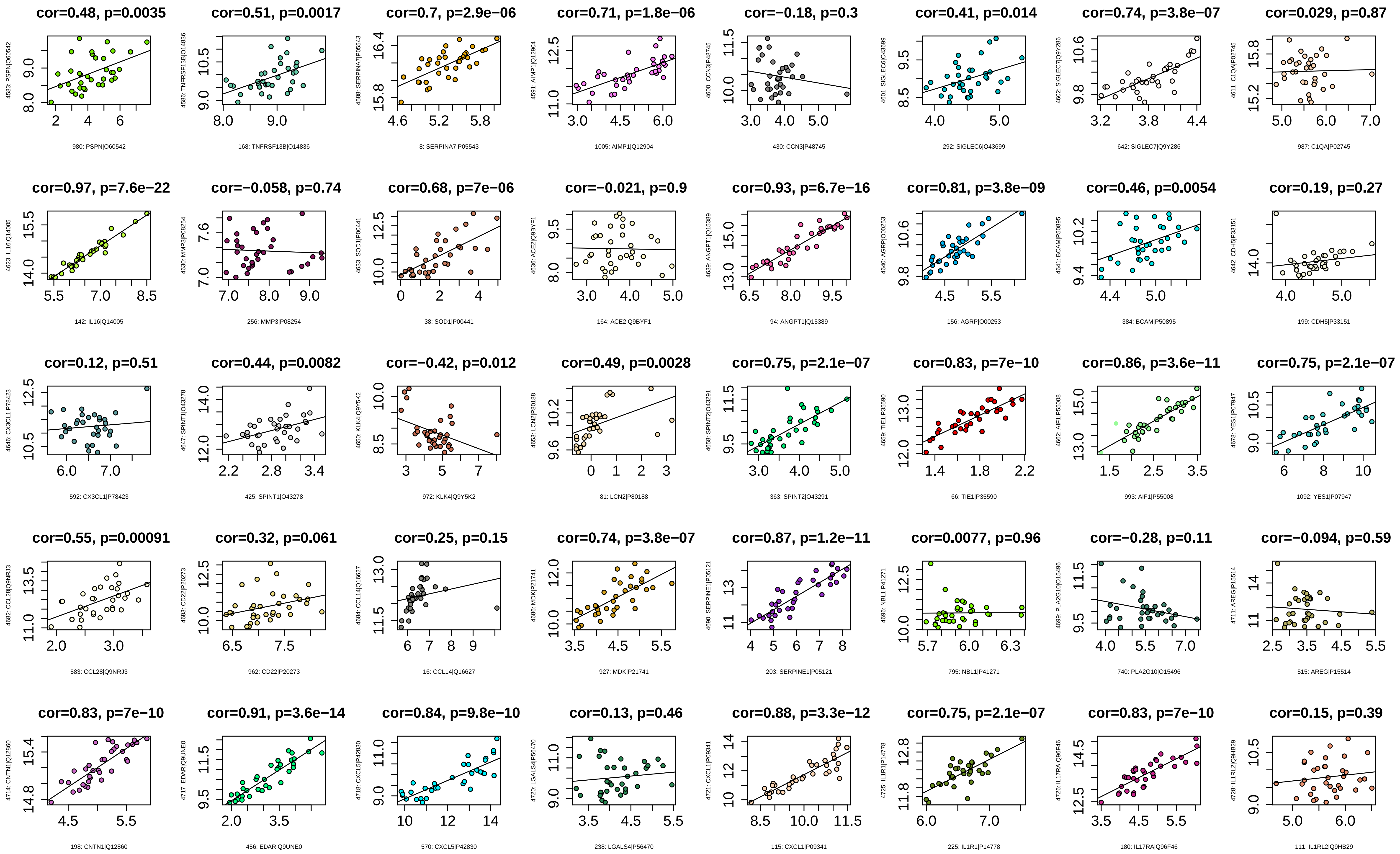

cor=0.48, p=0.0035
4583: PSPN|O60542
980: PSPN|O60542
cor=0.51, p=0.0017
4586: TNFRSF13B|O14836
168: TNFRSF13B|O14836
cor=0.7, p=2.9e−06
4588: SERPINA7|P05543
8: SERPINA7|P05543
cor=0.71, p=1.8e−06
4591: AIMP1|Q12904
1005: AIMP1|Q12904
cor=−0.18, p=0.3
4600: CCN3|P48745
430: CCN3|P48745
cor=0.41, p=0.014
4601: SIGLEC6|O43699
292: SIGLEC6|O43699
cor=0.74, p=3.8e−07
4602: SIGLEC7|Q9Y286
642: SIGLEC7|Q9Y286
cor=0.029, p=0.87
4611: C1QA|P02745
987: C1QA|P02745
cor=0.97, p=7.6e−22
4623: IL16|Q14005
142: IL16|Q14005
cor=−0.058, p=0.74
4630: MMP3|P08254
256: MMP3|P08254
cor=0.68, p=7e−06
4633: SOD1|P00441
38: SOD1|P00441
cor=−0.021, p=0.9
4636: ACE2|Q9BYF1
164: ACE2|Q9BYF1
cor=0.93, p=6.7e−16
4639: ANGPT1|Q15389
94: ANGPT1|Q15389
cor=0.81, p=3.8e−09
4640: AGRP|O00253
156: AGRP|O00253
cor=0.46, p=0.0054
4641: BCAM|P50895
384: BCAM|P50895
cor=0.19, p=0.27
4642: CDH5|P33151
199: CDH5|P33151
cor=0.12, p=0.51
4646: CX3CL1|P78423
592: CX3CL1|P78423
cor=0.44, p=0.0082
4647: SPINT1|O43278
425: SPINT1|O43278
cor=−0.42, p=0.012
4650: KLK4|Q9Y5K2
972: KLK4|Q9Y5K2
cor=0.49, p=0.0028
4653: LCN2|P80188
81: LCN2|P80188
cor=0.75, p=2.1e−07
4658: SPINT2|O43291
363: SPINT2|O43291
cor=0.83, p=7e−10
4659: TIE1|P35590
66: TIE1|P35590
cor=0.86, p=3.6e−11
4662: AIF1|P55008
993: AIF1|P55008
cor=0.75, p=2.1e−07
4678: YES1|P07947
1092: YES1|P07947
cor=0.55, p=0.00091
4682: CCL28|Q9NRJ3
583: CCL28|Q9NRJ3
cor=0.32, p=0.061
4683: CD22|P20273
962: CD22|P20273
cor=0.25, p=0.15
4684: CCL14|Q16627
16: CCL14|Q16627
cor=0.74, p=3.8e−07
4686: MDK|P21741
927: MDK|P21741
cor=0.87, p=1.2e−11
4690: SERPINE1|P05121
203: SERPINE1|P05121
cor=0.0077, p=0.96
4696: NBL1|P41271
795: NBL1|P41271
cor=−0.28, p=0.11
4699: PLA2G10|O15496
740: PLA2G10|O15496
cor=−0.094, p=0.59
4711: AREG|P15514
515: AREG|P15514
cor=0.83, p=7e−10
4714: CNTN1|Q12860
198: CNTN1|Q12860
cor=0.91, p=3.6e−14
4717: EDAR|Q9UNE0
456: EDAR|Q9UNE0
cor=0.84, p=9.8e−10
4718: CXCL5|P42830
570: CXCL5|P42830
cor=0.13, p=0.46
4720: LGALS4|P56470
238: LGALS4|P56470
cor=0.88, p=3.3e−12
4721: CXCL1|P09341
115: CXCL1|P09341
cor=0.75, p=2.1e−07
4725: IL1R1|P14778
225: IL1R1|P14778
cor=0.83, p=7e−10
4726: IL17RA|Q96F46
180: IL17RA|Q96F46
cor=0.15, p=0.39
4728: IL1RL2|Q9HB29
111: IL1RL2|Q9HB29

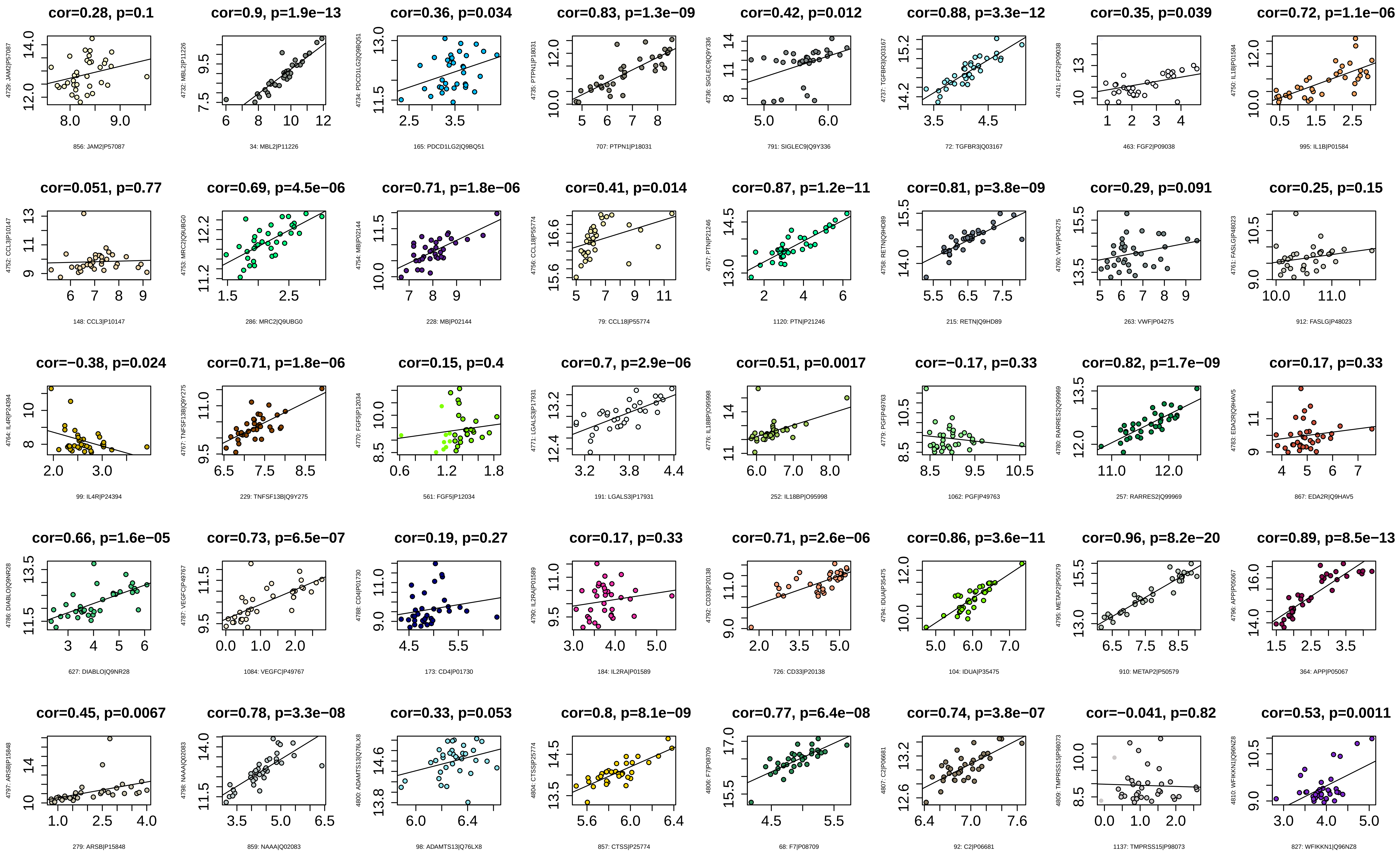
cor=0.28, p=0.1
856: JAM2|P57087
cor=0.9, p=1.9e−13
34: MBL2|P11226
cor=0.36, p=0.034
165: PDCD1LG2|Q9BQ51
cor=0.83, p=1.3e−09
707: PTPN1|P18031
cor=0.42, p=0.012
791: SIGLEC9|Q9Y336
cor=0.88, p=3.3e−12
72: TGFBR3|Q03167
cor=0.35, p=0.039
463: FGF2|P09038
cor=0.72, p=1.1e−06
995: IL1B|P01584
cor=0.051, p=0.77
148: CCL3|P10147
cor=0.69, p=4.5e−06
286: MRC2|Q9UBG0
cor=0.71, p=1.8e−06
228: MB|P02144
cor=0.41, p=0.014
79: CCL18|P55774
cor=0.87, p=1.2e−11
1120: PTN|P21246
cor=0.81, p=3.8e−09
215: RETN|Q9HD89
cor=0.29, p=0.091
263: VWF|P04275
cor=0.25, p=0.15
912: FASLG|P48023
cor=−0.38, p=0.024
99: IL4R|P24394
cor=0.71, p=1.8e−06
229: TNFSF13B|Q9Y275
cor=0.15, p=0.4
561: FGF5|P12034
cor=0.7, p=2.9e−06
191: LGALS3|P17931
cor=0.51, p=0.0017
252: IL18BP|O95998
cor=−0.17, p=0.33
1062: PGF|P49763
cor=0.82, p=1.7e−09
257: RARRES2|Q99969
cor=0.17, p=0.33
867: EDA2R|Q9HAV5
cor=0.66, p=1.6e−05
627: DIABLO|Q9NR28
cor=0.73, p=6.5e−07
1084: VEGFC|P49767
cor=0.19, p=0.27
173: CD4|P01730
cor=0.17, p=0.33
184: IL2RA|P01589
cor=0.71, p=2.6e−06
726: CD33|P20138
cor=0.86, p=3.6e−11
104: IDUA|P35475
cor=0.96, p=8.2e−20
910: METAP2|P50579
cor=0.89, p=8.5e−13
364: APP|P05067
cor=0.45, p=0.0067
279: ARSB|P15848
cor=0.78, p=3.3e−08
859: NAAA|Q02083
cor=0.33, p=0.053
98: ADAMTS13|Q76LX8
cor=0.8, p=8.1e−09
857: CTSS|P25774
cor=0.77, p=6.4e−08
68: F7|P08709
cor=0.74, p=3.8e−07
92: C2|P06681
cor=−0.041, p=0.82
1137: TMPRSS15|P98073
cor=0.53, p=0.0011
827: WFIKKN1|Q96NZ8

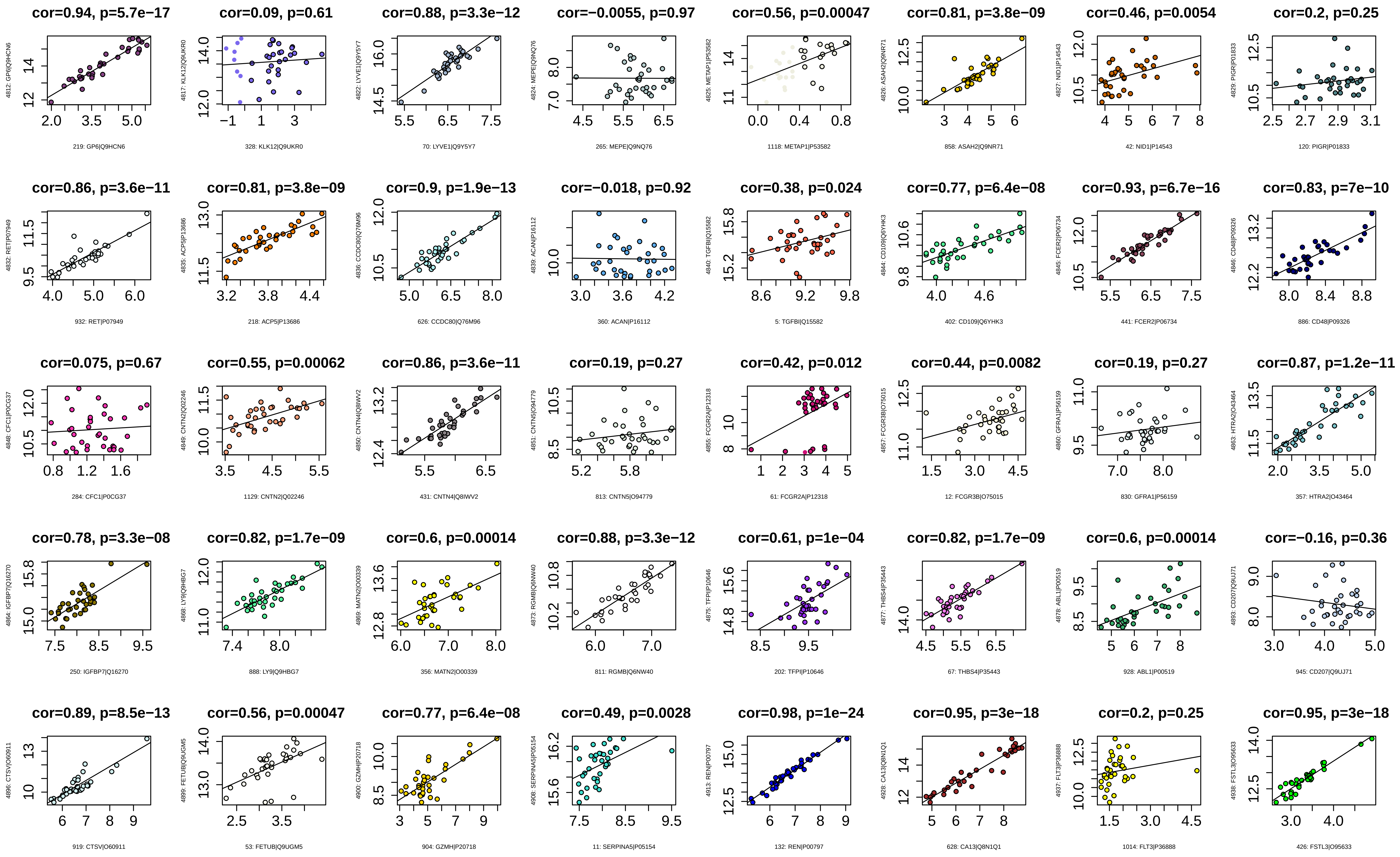
cor=0.94, p=5.7e−17
219: GP6|Q9HCN6
cor=0.09, p=0.61
328: KLK12|Q9UKR0
cor=0.88, p=3.3e−12
70: LYVE1|Q9Y5Y7
cor=−0.0055, p=0.97
265: MEPE|Q9NQ76
cor=0.56, p=0.00047
1118: METAP1|P53582
cor=0.81, p=3.8e−09
858: ASAH2|Q9NR71
cor=0.46, p=0.0054
42: NID1|P14543
cor=0.2, p=0.25
120: PIGR|P01833
cor=0.86, p=3.6e−11
932: RET|P07949
cor=0.81, p=3.8e−09
218: ACP5|P13686
cor=0.9, p=1.9e−13
626: CCDC80|Q76M96
cor=−0.018, p=0.92
360: ACAN|P16112
cor=0.38, p=0.024
5: TGFBI|Q15582
cor=0.77, p=6.4e−08
402: CD109|Q6YHK3
cor=0.93, p=6.7e−16
441: FCER2|P06734
cor=0.83, p=7e−10
886: CD48|P09326
cor=0.075, p=0.67
284: CFC1|P0CG37
cor=0.55, p=0.00062
1129: CNTN2|Q02246
cor=0.86, p=3.6e−11
431: CNTN4|Q8IWV2
cor=0.19, p=0.27
813: CNTN5|O94779
cor=0.42, p=0.012
61: FCGR2A|P12318
cor=0.44, p=0.0082
12: FCGR3B|O75015
cor=0.19, p=0.27
830: GFRA1|P56159
cor=0.87, p=1.2e−11
357: HTRA2|O43464
cor=0.78, p=3.3e−08
250: IGFBP7|Q16270
cor=0.82, p=1.7e−09
888: LY9|Q9HBG7
cor=0.6, p=0.00014
356: MATN2|O00339
cor=0.88, p=3.3e−12
811: RGMB|Q6NW40
cor=0.61, p=1e−04
202: TFPI|P10646
cor=0.82, p=1.7e−09
67: THBS4|P35443
cor=0.6, p=0.00014
928: ABL1|P00519
cor=−0.16, p=0.36
945: CD207|Q9UJ71
cor=0.89, p=8.5e−13
919: CTSV|O60911
cor=0.56, p=0.00047
53: FETUB|Q9UGM5
cor=0.77, p=6.4e−08
904: GZMH|P20718
cor=0.49, p=0.0028
11: SERPINA5|P05154
cor=0.98, p=1e−24
132: REN|P00797
cor=0.95, p=3e−18
628: CA13|Q8N1Q1
cor=0.2, p=0.25
1014: FLT3|P36888
cor=0.95, p=3e−18
426: FSTL3|O95633

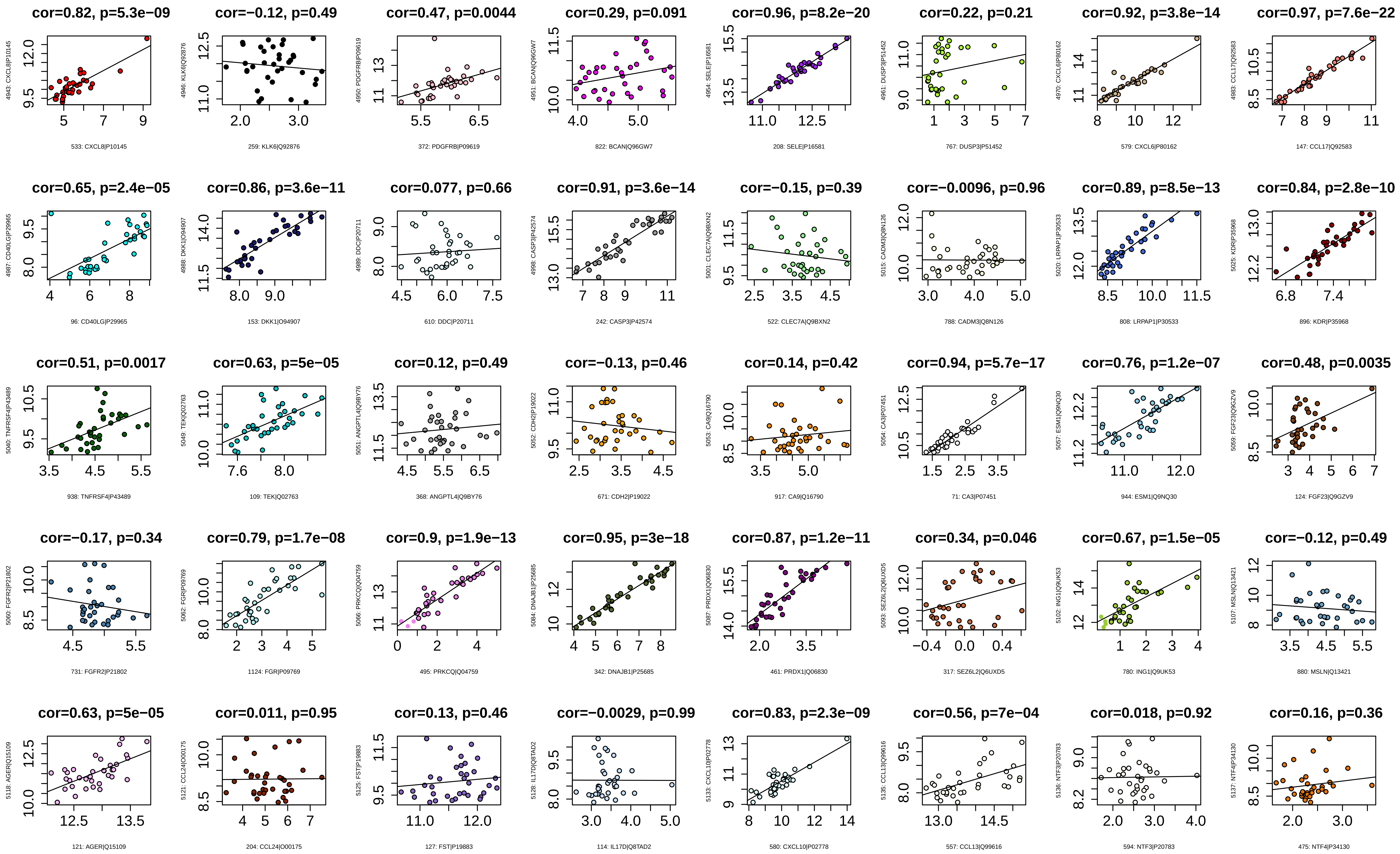

cor=0.82, p=5.3e−09
4943: CXCL8|P10145
533: CXCL8|P10145
cor=−0.12, p=0.49
4946: KLK6|Q92876
259: KLK6|Q92876
cor=0.47, p=0.0044
4950: PDGFRB|P09619
372: PDGFRB|P09619
cor=0.29, p=0.091
4951: BCAN|Q96GW7
822: BCAN|Q96GW7
cor=0.96, p=8.2e−20
4954: SELE|P16581
208: SELE|P16581
cor=0.22, p=0.21
4961: DUSP3|P51452
767: DUSP3|P51452
cor=0.92, p=3.8e−14
4970: CXCL6|P80162
579: CXCL6|P80162
cor=0.97, p=7.6e−22
4983: CCL17|Q92583
147: CCL17|Q92583
cor=0.65, p=2.4e−05
4987: CD40LG|P29965
96: CD40LG|P29965
cor=0.86, p=3.6e−11
4988: DKK1|O94907
153: DKK1|O94907
cor=0.077, p=0.66
4989: DDC|P20711
610: DDC|P20711
cor=0.91, p=3.6e−14
4998: CASP3|P42574
242: CASP3|P42574
cor=−0.15, p=0.39
5001: CLEC7A|Q9BXN2
522: CLEC7A|Q9BXN2
cor=−0.0096, p=0.96
5015: CADM3|Q8N126
788: CADM3|Q8N126
cor=0.89, p=8.5e−13
5020: LRPAP1|P30533
808: LRPAP1|P30533
cor=0.84, p=2.8e−10
5025: KDR|P35968
896: KDR|P35968
cor=0.51, p=0.0017
5040: TNFRSF4|P43489
938: TNFRSF4|P43489
cor=0.63, p=5e−05
5049: TEK|Q02763
109: TEK|Q02763
cor=0.12, p=0.49
5051: ANGPTL4|Q9BY76
368: ANGPTL4|Q9BY76
cor=−0.13, p=0.46
5052: CDH2|P19022
671: CDH2|P19022
cor=0.14, p=0.42
5053: CA9|Q16790
917: CA9|Q16790
cor=0.94, p=5.7e−17
5054: CA3|P07451
71: CA3|P07451
cor=0.76, p=1.2e−07
5057: ESM1|Q9NQ30
944: ESM1|Q9NQ30
cor=0.48, p=0.0035
5059: FGF23|Q9GZV9
124: FGF23|Q9GZV9
cor=−0.17, p=0.34
5060: FGFR2|P21802
731: FGFR2|P21802
cor=0.79, p=1.7e−08
5062: FGR|P09769
1124: FGR|P09769
cor=0.9, p=1.9e−13
5066: PRKCQ|Q04759
495: PRKCQ|Q04759
cor=0.95, p=3e−18
5084: DNAJB1|P25685
342: DNAJB1|P25685
cor=0.87, p=1.2e−11
5087: PRDX1|Q06830
461: PRDX1|Q06830
cor=0.34, p=0.046
5093: SEZ6L2|Q6UXD5
317: SEZ6L2|Q6UXD5
cor=0.67, p=1.5e−05
5102: ING1|Q9UK53
780: ING1|Q9UK53
cor=−0.12, p=0.49
5107: MSLN|Q13421
880: MSLN|Q13421
cor=0.63, p=5e−05
5118: AGER|Q15109
121: AGER|Q15109
cor=0.011, p=0.95
5121: CCL24|O00175
204: CCL24|O00175
cor=0.13, p=0.46
5125: FST|P19883
127: FST|P19883
cor=−0.0029, p=0.99
5128: IL17D|Q8TAD2
114: IL17D|Q8TAD2
cor=0.83, p=2.3e−09
5133: CXCL10|P02778
580: CXCL10|P02778
cor=0.56, p=7e−04
5135: CCL13|Q99616
557: CCL13|Q99616
cor=0.018, p=0.92
5136: NTF3|P20783
594: NTF3|P20783
cor=0.16, p=0.36
5137: NTF4|P34130
475: NTF4|P34130

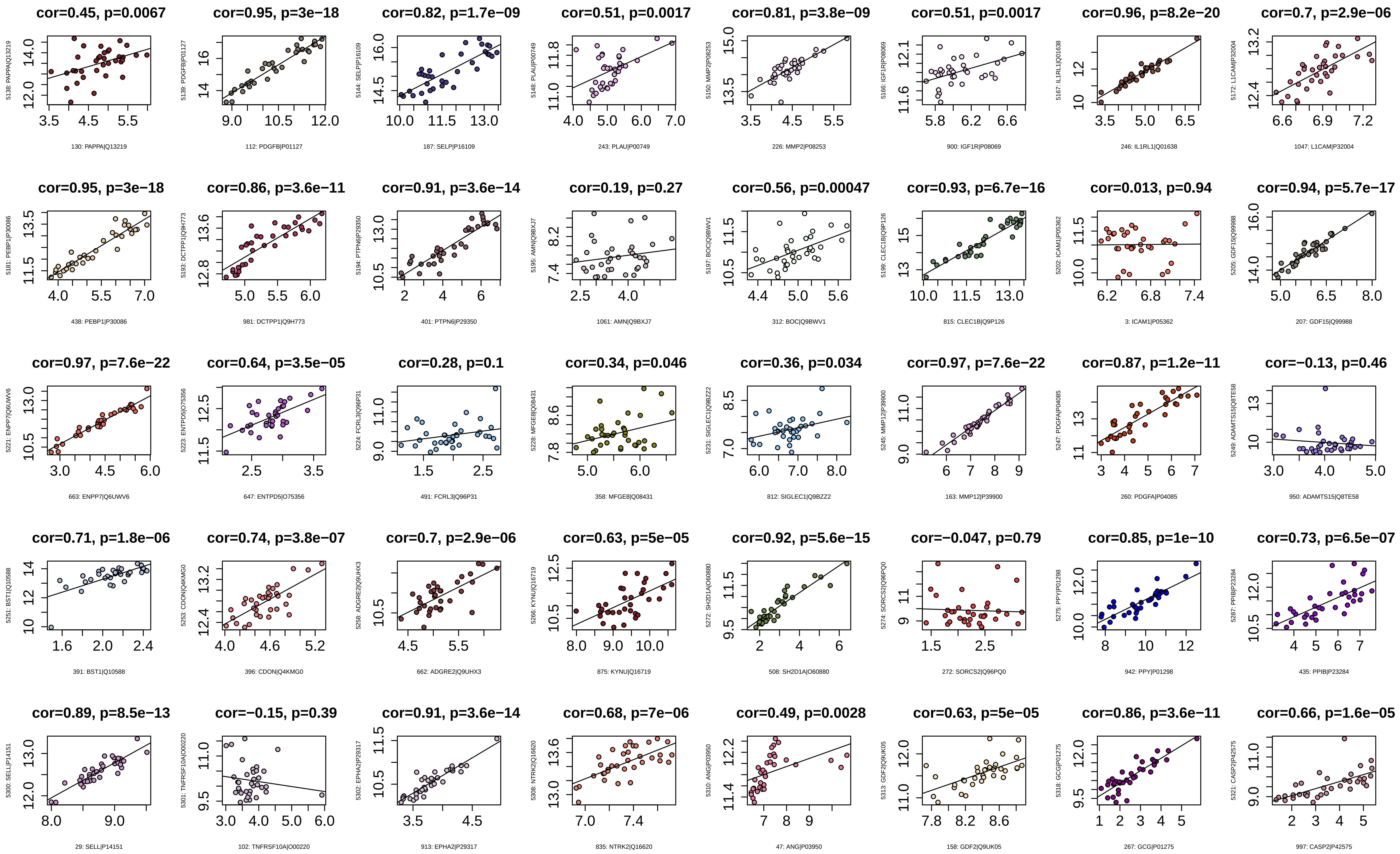

cor=0.45, p=0.0067
5138: PAPPA|Q13219
130: PAPPA|Q13219
cor=0.95, p=3e−18
5139: PDGFB|P01127
112: PDGFB|P01127
cor=0.82, p=1.7e−09
5144: SELP|P16109
187: SELP|P16109
cor=0.51, p=0.0017
5148: PLAU|P00749
243: PLAU|P00749
cor=0.81, p=3.8e−09
5150: MMP2|P08253
226: MMP2|P08253
cor=0.51, p=0.0017
5166: IGF1R|P08069
900: IGF1R|P08069
cor=0.96, p=8.2e−20
5167: IL1RL1|Q01638
246: IL1RL1|Q01638
cor=0.7, p=2.9e−06
5172: L1CAM|P32004
1047: L1CAM|P32004
cor=0.95, p=3e−18
5181: PEBP1|P30086
438: PEBP1|P30086
cor=0.86, p=3.6e−11
5193: DCTPP1|Q9H773
981: DCTPP1|Q9H773
cor=0.91, p=3.6e−14
5194: PTPN6|P29350
401: PTPN6|P29350
cor=0.19, p=0.27
5195: AMN|Q9BXJ7
1061: AMN|Q9BXJ7
cor=0.56, p=0.00047
5197: BOC|Q9BWV1
312: BOC|Q9BWV1
cor=0.93, p=6.7e−16
5199: CLEC1B|Q9P126
815: CLEC1B|Q9P126
cor=0.013, p=0.94
5202: ICAM1|P05362
3: ICAM1|P05362
cor=0.94, p=5.7e−17
5205: GDF15|Q99988
207: GDF15|Q99988
cor=0.97, p=7.6e−22
5221: ENPP7|Q6UWV6
663: ENPP7|Q6UWV6
cor=0.64, p=3.5e−05
5223: ENTPD5|O75356
647: ENTPD5|O75356
cor=0.28, p=0.1
5224: FCRL3|Q96P31
491: FCRL3|Q96P31
cor=0.34, p=0.046
5228: MFGE8|Q08431
358: MFGE8|Q08431
cor=0.36, p=0.034
5231: SIGLEC1|Q9BZZ2
812: SIGLEC1|Q9BZZ2
cor=0.97, p=7.6e−22
5245: MMP12|P39900
163: MMP12|P39900
cor=0.87, p=1.2e−11
5247: PDGFA|P04085
260: PDGFA|P04085
cor=−0.13, p=0.46
5249: ADAMTS15|Q8TE58
950: ADAMTS15|Q8TE58
cor=0.71, p=1.8e−06
5251: BST1|Q10588
391: BST1|Q10588
cor=0.74, p=3.8e−07
5253: CDON|Q4KMG0
396: CDON|Q4KMG0
cor=0.7, p=2.9e−06
5258: ADGRE2|Q9UHX3
662: ADGRE2|Q9UHX3
cor=0.63, p=5e−05
5266: KYNU|Q16719
875: KYNU|Q16719
cor=0.92, p=5.6e−15
5272: SH2D1A|O60880
508: SH2D1A|O60880
cor=−0.047, p=0.79
5274: SORCS2|Q96PQ0
272: SORCS2|Q96PQ0
cor=0.85, p=1e−10
5275: PPY|P01298
942: PPY|P01298
cor=0.73, p=6.5e−07
5287: PPIB|P23284
435: PPIB|P23284
cor=0.89, p=8.5e−13
5300: SELL|P14151
29: SELL|P14151
cor=−0.15, p=0.39
5301: TNFRSF10A|O00220
102: TNFRSF10A|O00220
cor=0.91, p=3.6e−14
5302: EPHA2|P29317
913: EPHA2|P29317
cor=0.68, p=7e−06
5308: NTRK2|Q16620
835: NTRK2|Q16620
cor=0.49, p=0.0028
5310: ANG|P03950
47: ANG|P03950
cor=0.63, p=5e−05
5313: GDF2|Q9UK05
158: GDF2|Q9UK05
cor=0.86, p=3.6e−11
5318: GCG|P01275
267: GCG|P01275
cor=0.66, p=1.6e−05
5321: CASP2|P42575
997: CASP2|P42575

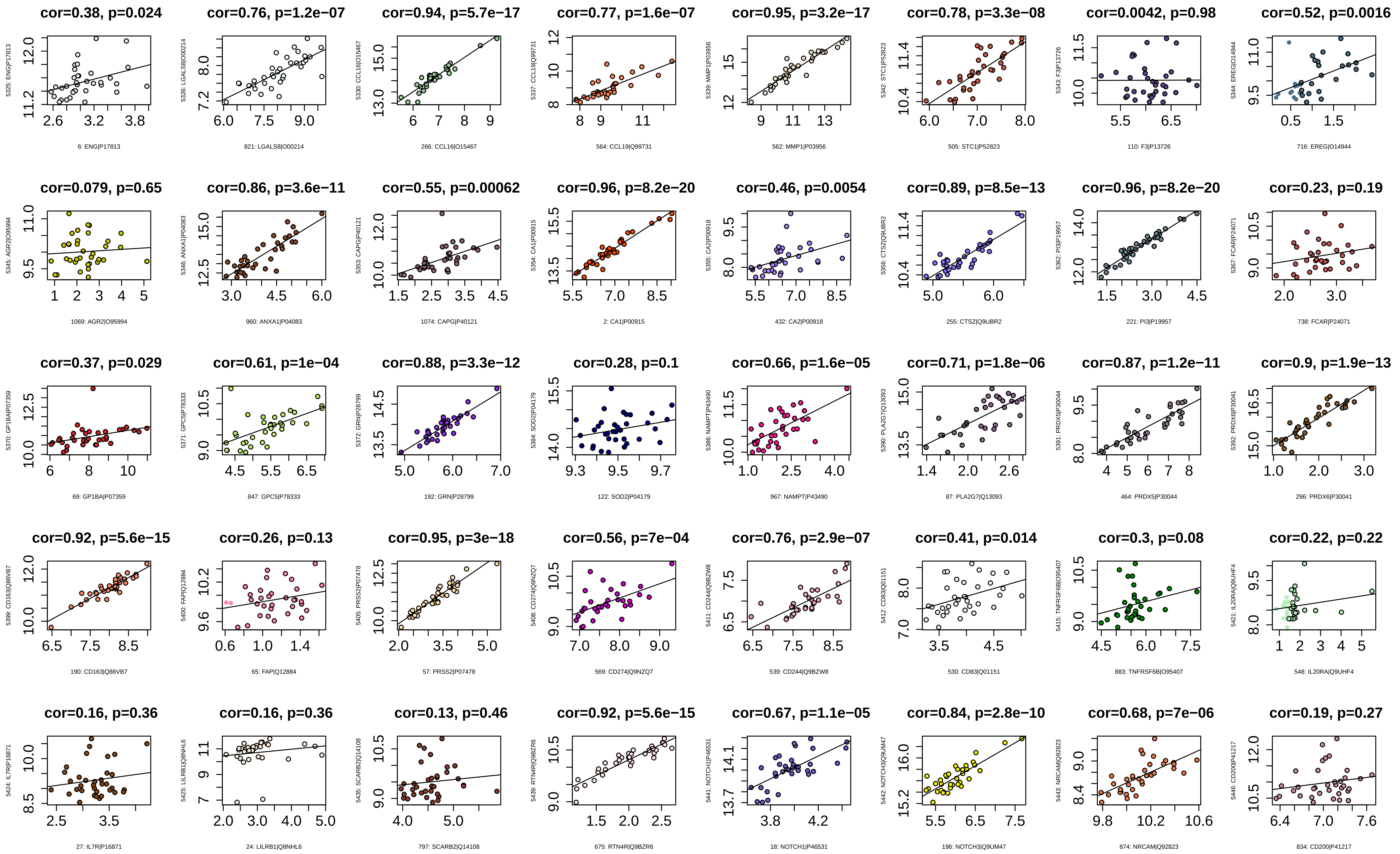

cor=0.38, p=0.024
6: ENG|P17813
cor=0.76, p=1.2e−07
821: LGALS8|O00214
cor=0.94, p=5.7e−17
266: CCL16|O15467
cor=0.77, p=1.6e−07
564: CCL19|Q99731
cor=0.95, p=3.2e−17
562: MMP1|P03956
cor=0.78, p=3.3e−08
505: STC1|P52823
cor=0.0042, p=0.98
110: F3|P13726
cor=0.52, p=0.0016
716: EREG|O14944
cor=0.079, p=0.65
1069: AGR2|O95994
cor=0.86, p=3.6e−11
960: ANXA1|P04083
cor=0.55, p=0.00062
1074: CAPG|P40121
cor=0.96, p=8.2e−20
2: CA1|P00915
cor=0.46, p=0.0054
432: CA2|P00918
cor=0.89, p=8.5e−13
255: CTSZ|Q9UBR2
cor=0.96, p=8.2e−20
221: PI3|P19957
cor=0.23, p=0.19
738: FCAR|P24071
cor=0.37, p=0.029
69: GP1BA|P07359
cor=0.61, p=1e−04
847: GPC5|P78333
cor=0.88, p=3.3e−12
192: GRN|P28799
cor=0.28, p=0.1
122: SOD2|P04179
cor=0.66, p=1.6e−05
967: NAMPT|P43490
cor=0.71, p=1.8e−06
87: PLA2G7|Q13093
cor=0.87, p=1.2e−11
464: PRDX5|P30044
cor=0.9, p=1.9e−13
296: PRDX6|P30041
cor=0.92, p=5.6e−15
190: CD163|Q86VB7
cor=0.26, p=0.13
65: FAP|Q12884
cor=0.95, p=3e−18
57: PRSS2|P07478
cor=0.56, p=7e−04
569: CD274|Q9NZQ7
cor=0.76, p=2.9e−07
539: CD244|Q9BZW8
cor=0.41, p=0.014
530: CD83|Q01151
cor=0.3, p=0.08
883: TNFRSF6B|O95407
cor=0.22, p=0.22
548: IL20RA|Q9UHF4
cor=0.16, p=0.36
27: IL7R|P16871
cor=0.16, p=0.36
24: LILRB1|Q8NHL6
cor=0.13, p=0.46
797: SCARB2|Q14108
cor=0.92, p=5.6e−15
675: RTN4R|Q9BZR6
cor=0.67, p=1.1e−05
18: NOTCH1|P46531
cor=0.84, p=2.8e−10
196: NOTCH3|Q9UM47
cor=0.68, p=7e−06
874: NRCAM|Q92823
cor=0.19, p=0.27
834: CD200|P41217

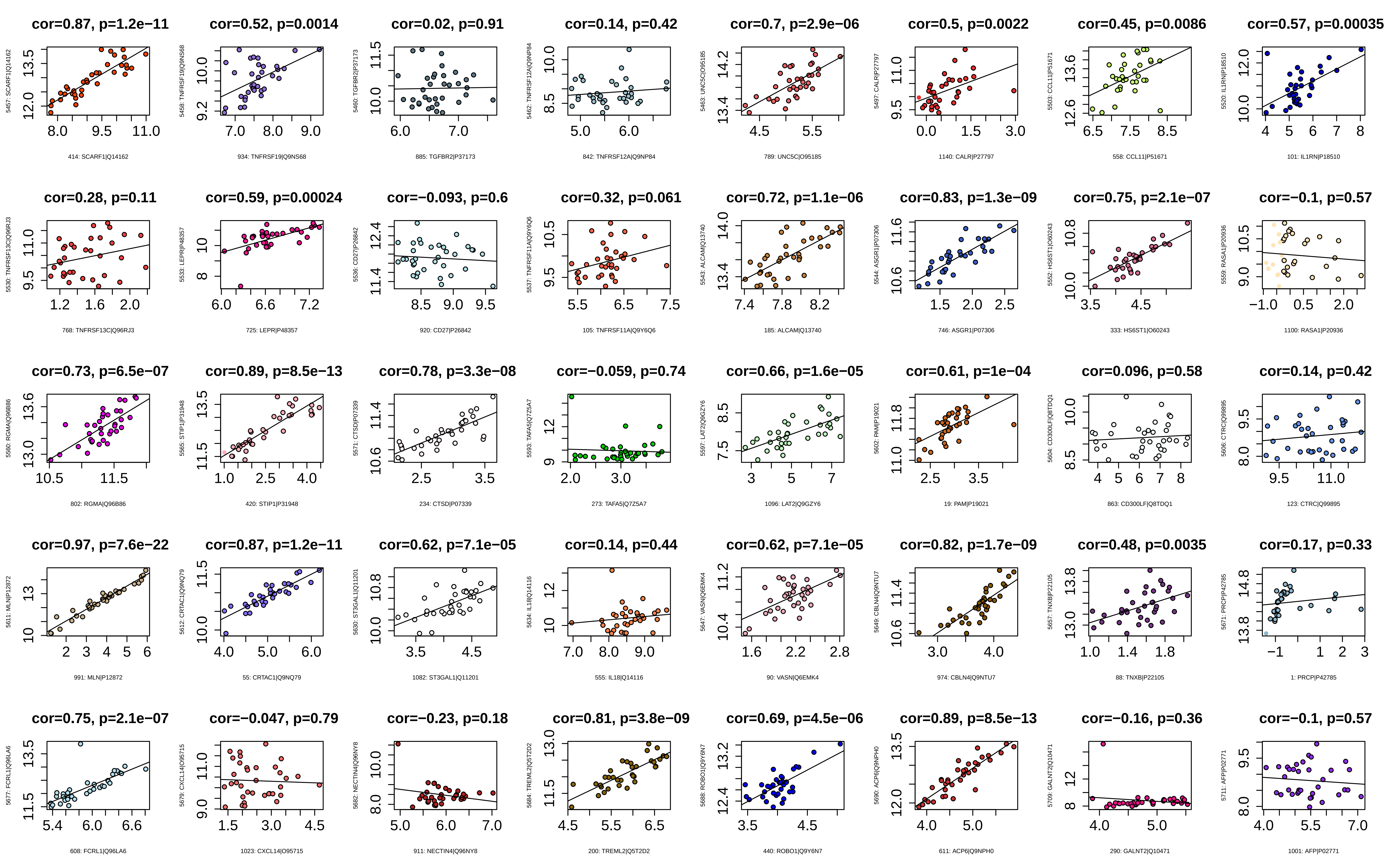
cor=0.87, p=1.2e−11
5457: SCARF1|Q14162
414: SCARF1|Q14162
cor=0.52, p=0.0014
5458: TNFRSF19|Q9NS68
934: TNFRSF19|Q9NS68
cor=0.02, p=0.91
5460: TGFBR2|P37173
885: TGFBR2|P37173
cor=0.14, p=0.42
5462: TNFRSF12A|Q9NP84
842: TNFRSF12A|Q9NP84
cor=0.7, p=2.9e−06
5463: UNC5C|O95185
789: UNC5C|O95185
cor=0.5, p=0.0022
5497: CALR|P27797
1140: CALR|P27797
cor=0.45, p=0.0086
5503: CCL11|P51671
558: CCL11|P51671
cor=0.57, p=0.00035
5520: IL1RN|P18510
101: IL1RN|P18510
cor=0.28, p=0.11
5530: TNFRSF13C|Q96RJ3
768: TNFRSF13C|Q96RJ3
cor=0.59, p=0.00024
5533: LEPR|P48357
725: LEPR|P48357
cor=−0.093, p=0.6
5536: CD27|P26842
920: CD27|P26842
cor=0.32, p=0.061
5537: TNFRSF11A|Q9Y6Q6
105: TNFRSF11A|Q9Y6Q6
cor=0.72, p=1.1e−06
5543: ALCAM|Q13740
185: ALCAM|Q13740
cor=0.83, p=1.3e−09
5544: ASGR1|P07306
746: ASGR1|P07306
cor=0.75, p=2.1e−07
5552: HS6ST1|O60243
333: HS6ST1|O60243
cor=−0.1, p=0.57
5559: RASA1|P20936
1100: RASA1|P20936
cor=0.73, p=6.5e−07
5560: RGMA|Q96B86
802: RGMA|Q96B86
cor=0.89, p=8.5e−13
5565: STIP1|P31948
420: STIP1|P31948
cor=0.78, p=3.3e−08
5571: CTSD|P07339
234: CTSD|P07339
cor=−0.059, p=0.74
5593: TAFA5|Q7Z5A7
273: TAFA5|Q7Z5A7
cor=0.66, p=1.6e−05
5597: LAT2|Q9GZY6
1096: LAT2|Q9GZY6
cor=0.61, p=1e−04
5602: PAM|P19021
19: PAM|P19021
cor=0.096, p=0.58
5604: CD300LF|Q8TDQ1
863: CD300LF|Q8TDQ1
cor=0.14, p=0.42
5606: CTRC|Q99895
123: CTRC|Q99895
cor=0.97, p=7.6e−22
5611: MLN|P12872
991: MLN|P12872
cor=0.87, p=1.2e−11
5612: CRTAC1|Q9NQ79
55: CRTAC1|Q9NQ79
cor=0.62, p=7.1e−05
5630: ST3GAL1|Q11201
1082: ST3GAL1|Q11201
cor=0.14, p=0.44
5634: IL18|Q14116
555: IL18|Q14116
cor=0.62, p=7.1e−05
5647: VASN|Q6EMK4
90: VASN|Q6EMK4
cor=0.82, p=1.7e−09
5649: CBLN4|Q9NTU7
974: CBLN4|Q9NTU7
cor=0.48, p=0.0035
5657: TNXB|P22105
88: TNXB|P22105
cor=0.17, p=0.33
5671: PRCP|P42785
1: PRCP|P42785
cor=0.75, p=2.1e−07
5677: FCRL1|Q96LA6
608: FCRL1|Q96LA6
cor=−0.047, p=0.79
5679: CXCL14|O95715
1023: CXCL14|O95715
cor=−0.23, p=0.18
5682: NECTIN4|Q96NY8
911: NECTIN4|Q96NY8
cor=0.81, p=3.8e−09
5684: TREML2|Q5T2D2
200: TREML2|Q5T2D2
cor=0.69, p=4.5e−06
5688: ROBO1|Q9Y6N7
440: ROBO1|Q9Y6N7
cor=0.89, p=8.5e−13
5690: ACP6|Q9NPH0
611: ACP6|Q9NPH0
cor=−0.16, p=0.36
5709: GALNT2|Q10471
290: GALNT2|Q10471
cor=−0.1, p=0.57
5711: AFP|P02771
1001: AFP|P02771

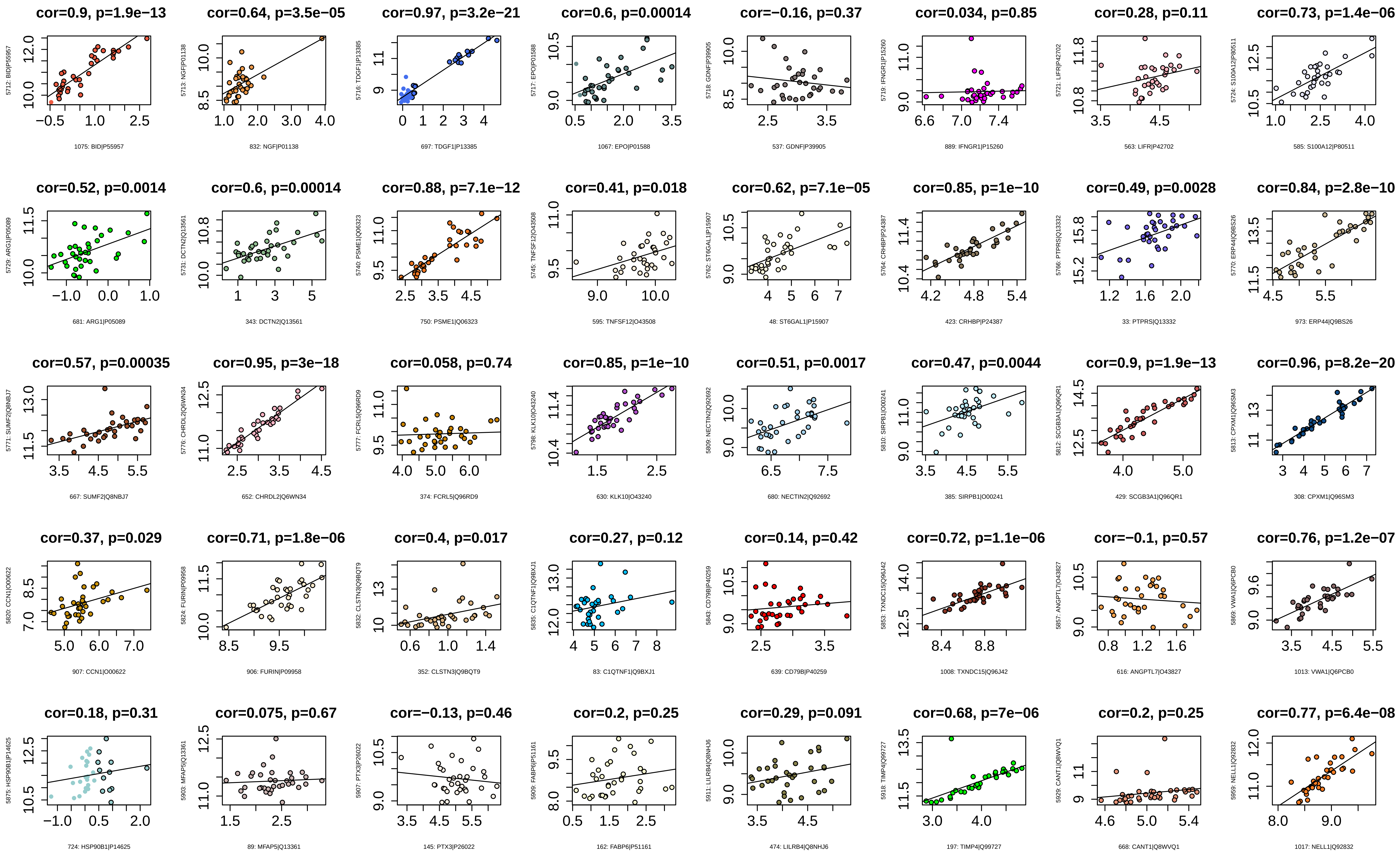

cor=0.9, p=1.9e−13
1075: BID|P55957
5712: BID|P55957
cor=0.64, p=3.5e−05
832: NGF|P01138
5713: NGF|P01138
cor=0.97, p=3.2e−21
697: TDGF1|P13385
5716: TDGF1|P13385
cor=0.6, p=0.00014
1067: EPO|P01588
5717: EPO|P01588
cor=−0.16, p=0.37
537: GDNF|P39905
5718: GDNF|P39905
cor=0.034, p=0.85
889: IFNGR1|P15260
5719: IFNGR1|P15260
cor=0.28, p=0.11
563: LIFR|P42702
5721: LIFR|P42702
cor=0.73, p=1.4e−06
585: S100A12|P80511
5724: S100A12|P80511
cor=0.52, p=0.0014
681: ARG1|P05089
5729: ARG1|P05089
cor=0.6, p=0.00014
343: DCTN2|Q13561
5731: DCTN2|Q13561
cor=0.88, p=7.1e−12
750: PSME1|Q06323
5740: PSME1|Q06323
cor=0.41, p=0.018
595: TNFSF12|O43508
5745: TNFSF12|O43508
cor=0.62, p=7.1e−05
48: ST6GAL1|P15907
5762: ST6GAL1|P15907
cor=0.85, p=1e−10
423: CRHBP|P24387
5764: CRHBP|P24387
cor=0.49, p=0.0028
33: PTPRS|Q13332
5766: PTPRS|Q13332
cor=0.84, p=2.8e−10
973: ERP44|Q9BS26
5770: ERP44|Q9BS26
cor=0.57, p=0.00035
667: SUMF2|Q8NBJ7
5771: SUMF2|Q8NBJ7
cor=0.95, p=3e−18
652: CHRDL2|Q6WN34
5776: CHRDL2|Q6WN34
cor=0.058, p=0.74
374: FCRL5|Q96RD9
5777: FCRL5|Q96RD9
cor=0.85, p=1e−10
630: KLK10|O43240
5798: KLK10|O43240
cor=0.51, p=0.0017
680: NECTIN2|Q92692
5809: NECTIN2|Q92692
cor=0.47, p=0.0044
385: SIRPB1|O00241
5810: SIRPB1|O00241
cor=0.9, p=1.9e−13
429: SCGB3A1|Q96QR1
5812: SCGB3A1|Q96QR1
cor=0.96, p=8.2e−20
308: CPXM1|Q96SM3
5813: CPXM1|Q96SM3
cor=0.37, p=0.029
907: CCN1|O00622
5820: CCN1|O00622
cor=0.71, p=1.8e−06
906: FURIN|P09958
5824: FURIN|P09958
cor=0.4, p=0.017
352: CLSTN3|Q9BQT9
5832: CLSTN3|Q9BQT9
cor=0.27, p=0.12
83: C1QTNF1|Q9BXJ1
5835: C1QTNF1|Q9BXJ1
cor=0.14, p=0.42
639: CD79B|P40259
5843: CD79B|P40259
cor=0.72, p=1.1e−06
1008: TXNDC15|Q96J42
5853: TXNDC15|Q96J42
cor=−0.1, p=0.57
616: ANGPTL7|O43827
5857: ANGPTL7|O43827
cor=0.76, p=1.2e−07
1013: VWA1|Q6PCB0
5869: VWA1|Q6PCB0
cor=0.18, p=0.31
724: HSP90B1|P14625
5875: HSP90B1|P14625
cor=0.075, p=0.67
89: MFAP5|Q13361
5903: MFAP5|Q13361
cor=−0.13, p=0.46
145: PTX3|P26022
5907: PTX3|P26022
cor=0.2, p=0.25
162: FABP6|P51161
5909: FABP6|P51161
cor=0.29, p=0.091
474: LILRB4|Q8NHJ6
5911: LILRB4|Q8NHJ6
cor=0.68, p=7e−06
197: TIMP4|Q99727
5918: TIMP4|Q99727
cor=0.2, p=0.25
668: CANT1|Q8WVQ1
5929: CANT1|Q8WVQ1
cor=0.77, p=6.4e−08
1017: NELL1|Q92832
5959: NELL1|Q92832

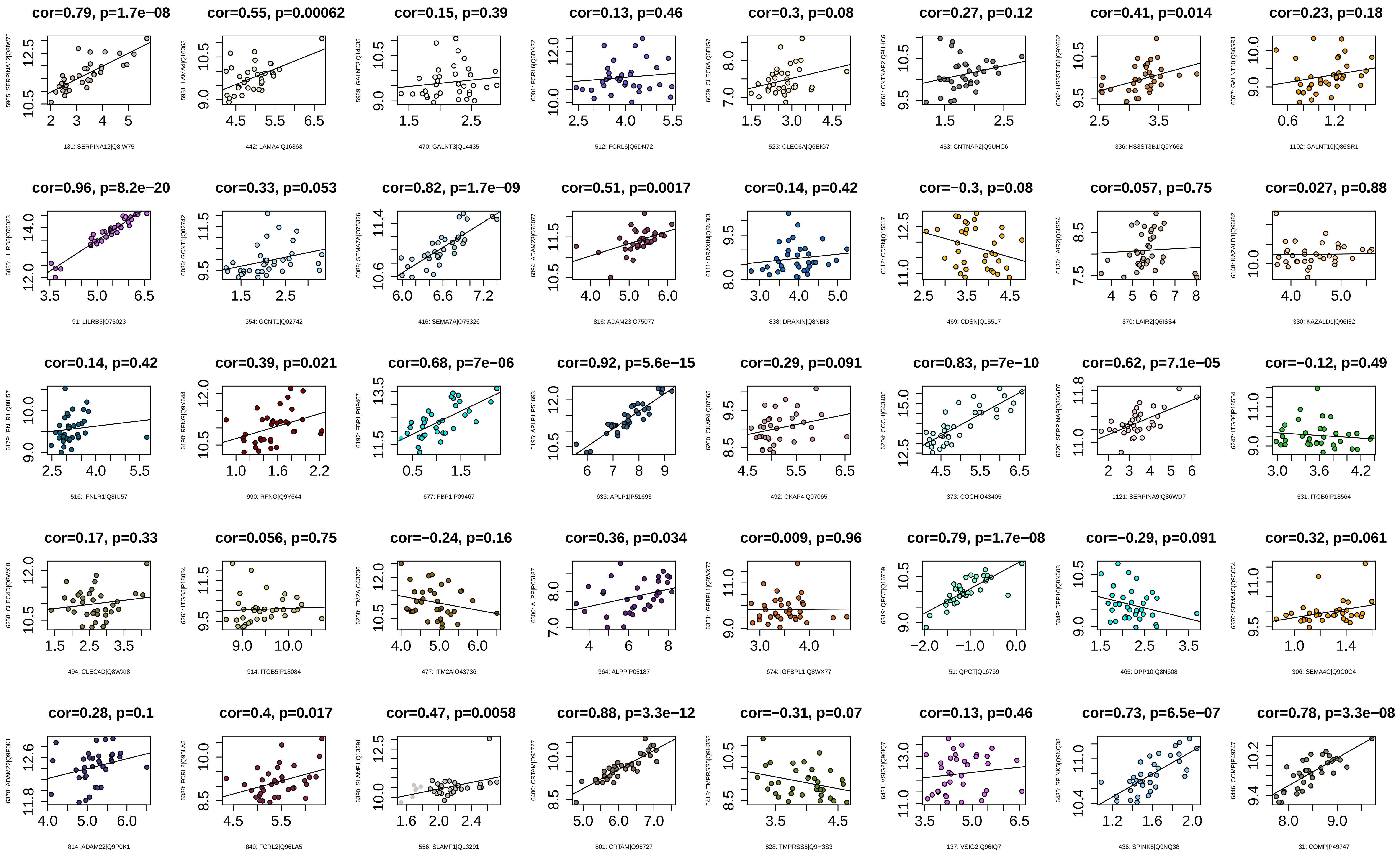
cor=0.79, p=1.7e−08
131: SERPINA12|Q8IW75
cor=0.55, p=0.00062
442: LAMA4|Q16363
cor=0.15, p=0.39
470: GALNT3|Q14435
cor=0.13, p=0.46
512: FCRL6|Q6DN72
cor=0.3, p=0.08
523: CLEC6A|Q6EIG7
cor=0.27, p=0.12
453: CNTNAP2|Q9UHC6
cor=0.41, p=0.014
336: HS3ST3B1|Q9Y662
cor=0.23, p=0.18
1102: GALNT10|Q86SR1
cor=0.96, p=8.2e−20
91: LILRB5|O75023
cor=0.33, p=0.053
354: GCNT1|Q02742
cor=0.82, p=1.7e−09
416: SEMA7A|O75326
cor=0.51, p=0.0017
816: ADAM23|O75077
cor=0.14, p=0.42
838: DRAXIN|Q8NBI3
cor=−0.3, p=0.08
469: CDSN|Q15517
cor=0.057, p=0.75
870: LAIR2|Q6ISS4
cor=0.027, p=0.88
330: KAZALD1|Q96I82
cor=0.14, p=0.42
516: IFNLR1|Q8IU57
cor=0.39, p=0.021
990: RFNG|Q9Y644
cor=0.68, p=7e−06
677: FBP1|P09467
cor=0.92, p=5.6e−15
633: APLP1|P51693
cor=0.29, p=0.091
492: CKAP4|Q07065
cor=0.83, p=7e−10
373: COCH|O43405
cor=0.62, p=7.1e−05
1121: SERPINA9|Q86WD7
cor=−0.12, p=0.49
531: ITGB6|P18564
cor=0.17, p=0.33
494: CLEC4D|Q8WXI8
cor=0.056, p=0.75
914: ITGB5|P18084
cor=−0.24, p=0.16
477: ITM2A|O43736
cor=0.36, p=0.034
964: ALPP|P05187
cor=0.009, p=0.96
674: IGFBPL1|Q8WX77
cor=0.79, p=1.7e−08
51: QPCT|Q16769
cor=−0.29, p=0.091
465: DPP10|Q8N608
cor=0.32, p=0.061
306: SEMA4C|Q9C0C4
cor=0.28, p=0.1
814: ADAM22|Q9P0K1
cor=0.4, p=0.017
849: FCRL2|Q96LA5
cor=0.47, p=0.0058
556: SLAMF1|Q13291
cor=0.88, p=3.3e−12
801: CRTAM|O95727
cor=−0.31, p=0.07
828: TMPRSS5|Q9H3S3
cor=0.13, p=0.46
137: VSIG2|Q96IQ7
cor=0.73, p=6.5e−07
436: SPINK5|Q9NQ38
cor=0.78, p=3.3e−08
31: COMP|P49747

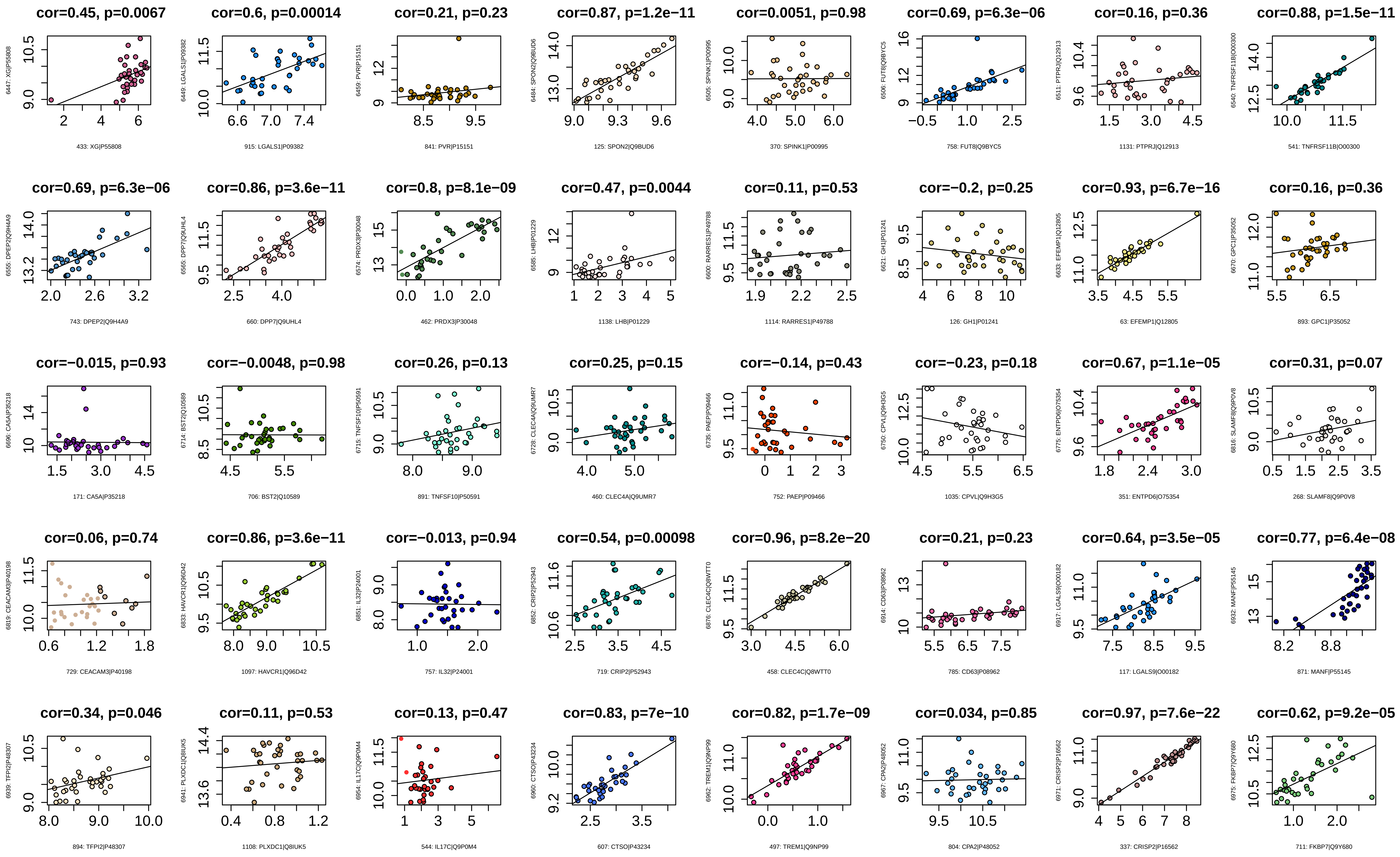

cor=0.45, p=0.0067
6447: XG|P55808
433: XG|P55808
cor=0.6, p=0.00014
6449: LGALS1|P09382
915: LGALS1|P09382
cor=0.21, p=0.23
6459: PVR|P15151
841: PVR|P15151
cor=0.87, p=1.2e−11
6484: SPON2|Q9BUD6
125: SPON2|Q9BUD6
cor=0.0051, p=0.98
6505: SPINK1|P00995
370: SPINK1|P00995
cor=0.69, p=6.3e−06
6506: FUT8|Q9BYC5
758: FUT8|Q9BYC5
cor=0.16, p=0.36
6511: PTPRJ|Q12913
1131: PTPRJ|Q12913
cor=0.88, p=1.5e−11
6540: TNFRSF11B|O00300
541: TNFRSF11B|O00300
cor=0.69, p=6.3e−06
6555: DPEP2|Q9H4A9
743: DPEP2|Q9H4A9
cor=0.86, p=3.6e−11
6565: DPP7|Q9UHL4
660: DPP7|Q9UHL4
cor=0.8, p=8.1e−09
6574: PRDX3|P30048
462: PRDX3|P30048
cor=0.47, p=0.0044
6585: LHB|P01229
1138: LHB|P01229
cor=0.11, p=0.53
6600: RARRES1|P49788
1114: RARRES1|P49788
cor=−0.2, p=0.25
6621: GH1|P01241
126: GH1|P01241
cor=0.93, p=6.7e−16
6633: EFEMP1|Q12805
63: EFEMP1|Q12805
cor=0.16, p=0.36
6670: GPC1|P35052
893: GPC1|P35052
cor=−0.015, p=0.93
6696: CA5A|P35218
171: CA5A|P35218
cor=−0.0048, p=0.98
6714: BST2|Q10589
706: BST2|Q10589
cor=0.26, p=0.13
6715: TNFSF10|P50591
891: TNFSF10|P50591
cor=0.25, p=0.15
6728: CLEC4A|Q9UMR7
460: CLEC4A|Q9UMR7
cor=−0.14, p=0.43
6735: PAEP|P09466
752: PAEP|P09466
cor=−0.23, p=0.18
6750: CPVL|Q9H3G5
1035: CPVL|Q9H3G5
cor=0.67, p=1.1e−05
6775: ENTPD6|O75354
351: ENTPD6|O75354
cor=0.31, p=0.07
6816: SLAMF8|Q9P0V8
268: SLAMF8|Q9P0V8
cor=0.06, p=0.74
6819: CEACAM3|P40198
729: CEACAM3|P40198
cor=0.86, p=3.6e−11
6833: HAVCR1|Q96D42
1097: HAVCR1|Q96D42
cor=−0.013, p=0.94
6851: IL32|P24001
757: IL32|P24001
cor=0.54, p=0.00098
6852: CRIP2|P52943
719: CRIP2|P52943
cor=0.96, p=8.2e−20
6876: CLEC4C|Q8WTT0
458: CLEC4C|Q8WTT0
cor=0.21, p=0.23
6914: CD63|P08962
785: CD63|P08962
cor=0.64, p=3.5e−05
6917: LGALS9|O00182
117: LGALS9|O00182
cor=0.77, p=6.4e−08
6932: MANF|P55145
871: MANF|P55145
cor=0.34, p=0.046
6939: TFPI2|P48307
894: TFPI2|P48307
cor=0.11, p=0.53
6941: PLXDC1|Q8IUK5
1108: PLXDC1|Q8IUK5
cor=0.13, p=0.47
6954: IL17C|Q9P0M4
544: IL17C|Q9P0M4
cor=0.83, p=7e−10
6960: CTSO|P43234
607: CTSO|P43234
cor=0.82, p=1.7e−09
6962: TREM1|Q9NP99
497: TREM1|Q9NP99
cor=0.034, p=0.85
6967: CPA2|P48052
804: CPA2|P48052
cor=0.97, p=7.6e−22
6971: CRISP2|P16562
337: CRISP2|P16562
cor=0.62, p=9.2e−05
6975: FKBP7|Q9Y680
711: FKBP7|Q9Y680

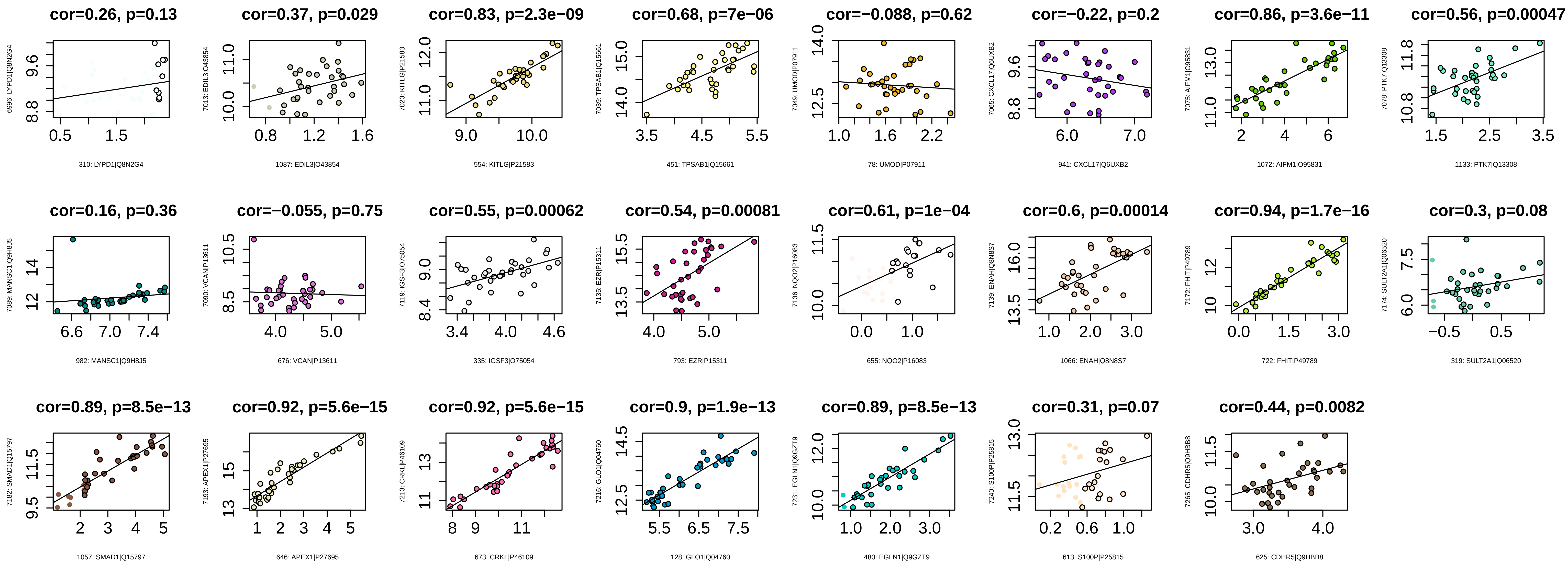
cor=0.26, p=0.13
310: LYPD1|Q8N2G4
6996: LYPD1|Q8N2G4
cor=0.37, p=0.029
1087: EDIL3|O43854
7013: EDIL3|O43854
cor=0.83, p=2.3e−09
554: KITLG|P21583
7023: KITLG|P21583
cor=0.68, p=7e−06
451: TPSAB1|Q15661
7039: TPSAB1|Q15661
cor=−0.088, p=0.62
78: UMOD|P07911
7049: UMOD|P07911
cor=−0.22, p=0.2
941: CXCL17|Q6UXB2
7065: CXCL17|Q6UXB2
cor=0.86, p=3.6e−11
1072: AIFM1|O95831
7075: AIFM1|O95831
cor=0.56, p=0.00047
1133: PTK7|Q13308
7078: PTK7|Q13308
cor=0.16, p=0.36
982: MANSC1|Q9H8J5
7089: MANSC1|Q9H8J5
cor=−0.055, p=0.75
676: VCAN|P13611
7090: VCAN|P13611
cor=0.55, p=0.00062
335: IGSF3|O75054
7119: IGSF3|O75054
cor=0.54, p=0.00081
793: EZR|P15311
7135: EZR|P15311
cor=0.61, p=1e−04
655: NQO2|P16083
7136: NQO2|P16083
cor=0.6, p=0.00014
1066: ENAH|Q8N8S7
7139: ENAH|Q8N8S7
cor=0.94, p=1.7e−16
722: FHIT|P49789
7172: FHIT|P49789
cor=0.3, p=0.08
319: SULT2A1|Q06520
7174: SULT2A1|Q06520
cor=0.89, p=8.5e−13
1057: SMAD1|Q15797
7182: SMAD1|Q15797
cor=0.92, p=5.6e−15
646: APEX1|P27695
7193: APEX1|P27695
cor=0.92, p=5.6e−15
673: CRKL|P46109
7213: CRKL|P46109
cor=0.9, p=1.9e−13
128: GLO1|Q04760
7216: GLO1|Q04760
cor=0.89, p=8.5e−13
480: EGLN1|Q9GZT9
7231: EGLN1|Q9GZT9
cor=0.31, p=0.07
613: S100P|P25815
7240: S100P|P25815
cor=0.44, p=0.0082
625: CDHR5|Q9HBB8
7265: CDHR5|Q9HBB8

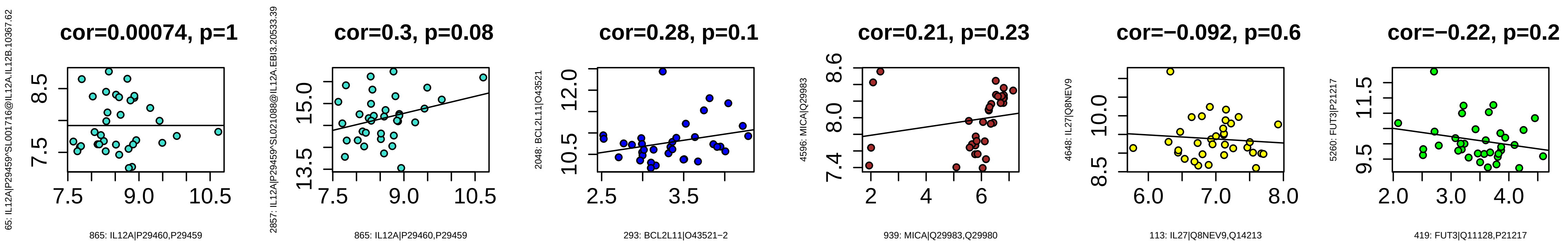
cor=0.00074, p=1
65: IL12A|P29459^SL001716@IL12A.IL12B.10367.62
865: IL12A|P29460,P29459
cor=0.3, p=0.08
2857: IL12A|P29459^SL021088@IL12A.EBI3.20533.39
865: IL12A|P29460,P29459
cor=0.28, p=0.1
2048: BCL2L11|O43521
293: BCL2L11|O43521−2
cor=0.21, p=0.23
4596: MICA|Q29983
939: MICA|Q29983,Q29980
cor=−0.092, p=0.6
4648: IL27|Q8NEV9
113: IL27|Q8NEV9,Q14213
cor=−0.22, p=0.2
5260: FUT3|P21217
419: FUT3|Q11128,P21217

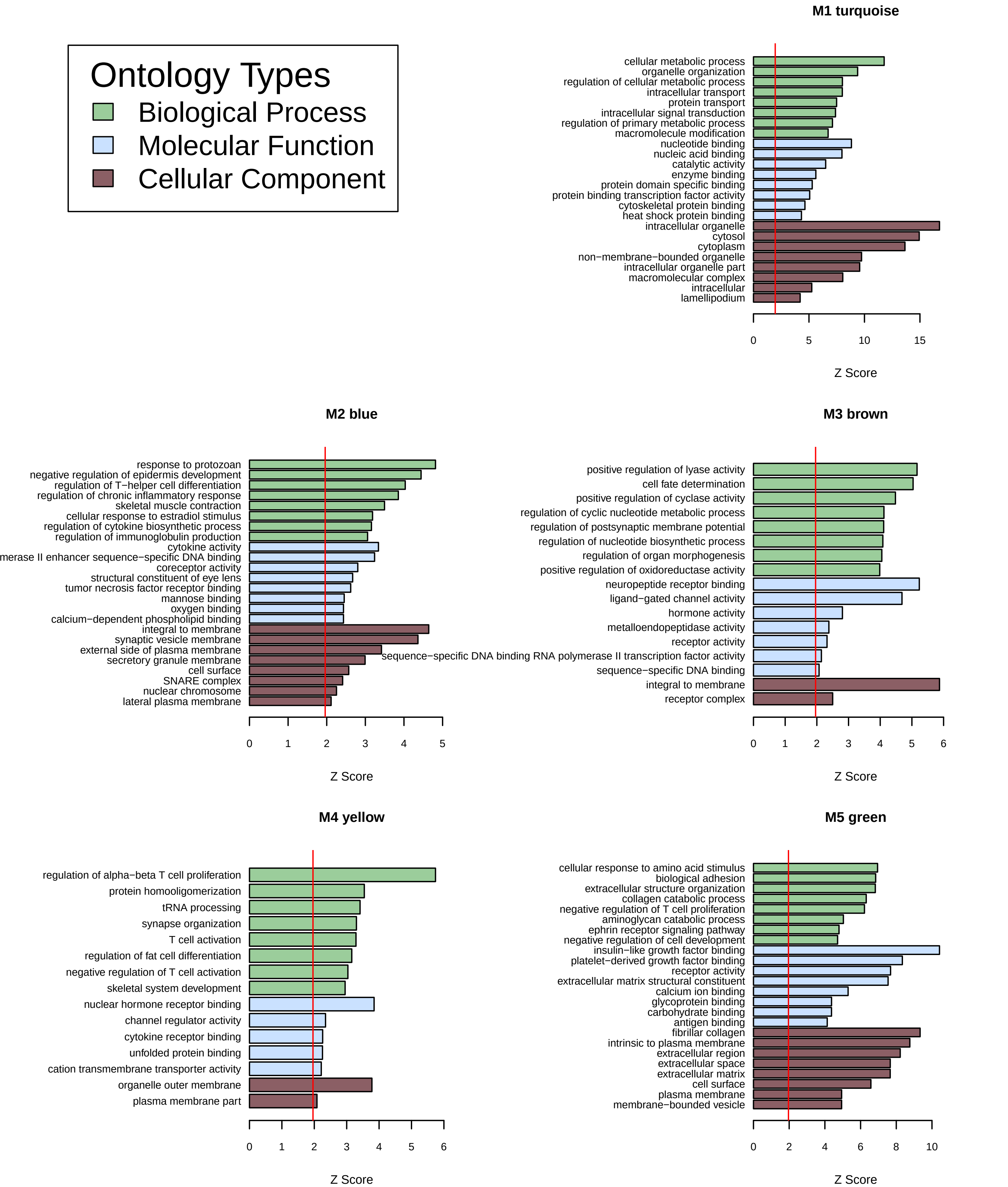

Ontology Types
Biological Process
Molecular Function
Cellular Component
M1 turquoise
cellular metabolic process
organelle organization
regulation of cellular metabolic process
intracellular transport
protein transport
intracellular signal transduction
regulation of primary metabolic process
macromolecule modification
nucleotide binding
nucleic acid binding
catalytic activity
enzyme binding
protein domain specific binding
protein binding transcription factor activity
cytoskeletal protein binding
heat shock protein binding
intracellular organelle
cytosol
cytoplasm
non−membrane−bounded organelle
intracellular organelle part
macromolecular complex
intracellular
lamellipodium
0
5
10
15
Z Score
M2 blue
response to protozoan
negative regulation of epidermis development
regulation of T−helper cell differentiation
regulation of chronic inflammatory response
skeletal muscle contraction
cellular response to estradiol stimulus
regulation of cytokine biosynthetic process
regulation of immunoglobulin production
cytokine activity
merase II enhancer sequence−specific DNA binding
coreceptor activity
structural constituent of eye lens
tumor necrosis factor receptor binding
mannose binding
oxygen binding
calcium−dependent phospholipid binding
integral to membrane
synaptic vesicle membrane
external side of plasma membrane
secretory granule membrane
cell surface
SNARE complex
nuclear chromosome
lateral plasma membrane
0
1
2
3
4
5
Z Score
M3 brown
positive regulation of lyase activity
cell fate determination
positive regulation of cyclase activity
regulation of cyclic nucleotide metabolic process
regulation of postsynaptic membrane potential
regulation of nucleotide biosynthetic process
regulation of organ morphogenesis
positive regulation of oxidoreductase activity
neuropeptide receptor binding
ligand−gated channel activity
hormone activity
metalloendopeptidase activity
receptor activity
sequence−specific DNA binding RNA polymerase II transcription factor activity
sequence−specific DNA binding
integral to membrane
receptor complex
0
1
2
3
4
5
6
Z Score
M4 yellow
regulation of alpha−beta T cell proliferation
protein homooligomerization
tRNA processing
synapse organization
T cell activation
regulation of fat cell differentiation
negative regulation of T cell activation
skeletal system development
nuclear hormone receptor binding
channel regulator activity
cytokine receptor binding
unfolded protein binding
cation transmembrane transporter activity
organelle outer membrane
plasma membrane part
0
1
2
3
4
5
6
Z Score
M5 green
cellular response to amino acid stimulus
biological adhesion
extracellular structure organization
collagen catabolic process
negative regulation of T cell proliferation
aminoglycan catabolic process
ephrin receptor signaling pathway
negative regulation of cell development
insulin−like growth factor binding
platelet−derived growth factor binding
receptor activity
extracellular matrix structural constituent
calcium ion binding
glycoprotein binding
carbohydrate binding
antigen binding
fibrillar collagen
intrinsic to plasma membrane
extracellular region
extracellular space
extracellular matrix
cell surface
plasma membrane
membrane−bounded vesicle
0
2
4
6
8
10
Z Score

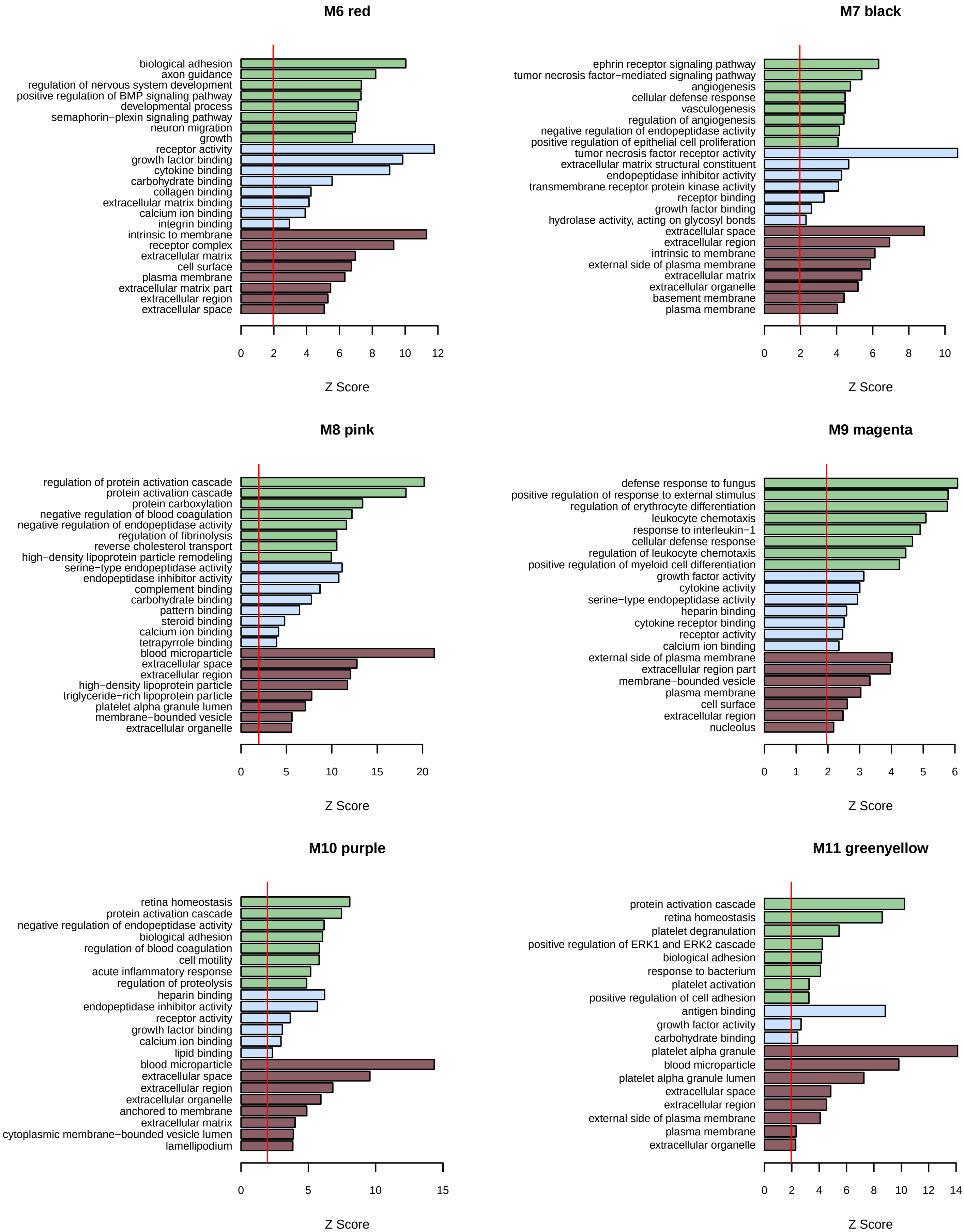

M6 red
biological adhesion
axon guidance
regulation of nervous system development
positive regulation of BMP signaling pathway
developmental process
semaphorin−plexin signaling pathway
neuron migration
growth
receptor activity
growth factor binding
cytokine binding
carbohydrate binding
collagen binding
extracellular matrix binding
calcium ion binding
integrin binding
intrinsic to membrane
receptor complex
extracellular matrix
cell surface
plasma membrane
extracellular matrix part
extracellular region
extracellular space
0 2 4 6 8 10 12
Z Score
M7 black
ephrin receptor signaling pathway
tumor necrosis factor−mediated signaling pathway
angiogenesis
cellular defense response
vasculogenesis
regulation of angiogenesis
negative regulation of endopeptidase activity
positive regulation of epithelial cell proliferation
tumor necrosis factor receptor activity
extracellular matrix structural constituent
endopeptidase inhibitor activity
transmembrane receptor protein kinase activity
receptor binding
growth factor binding
hydrolase activity, acting on glycosyl bonds
extracellular space
extracellular region
intrinsic to membrane
external side of plasma membrane
extracellular matrix
extracellular organelle
basement membrane
plasma membrane
0 2 4 6 8 10
Z Score
M8 pink
regulation of protein activation cascade
protein activation cascade
protein carboxylation
negative regulation of blood coagulation
negative regulation of endopeptidase activity
regulation of fibrinolysis
reverse cholesterol transport
high−density lipoprotein particle remodeling
serine−type endopeptidase activity
endopeptidase inhibitor activity
complement binding
carbohydrate binding
pattern binding
steroid binding
calcium ion binding
tetrapyrrole binding
blood microparticle
extracellular space
extracellular region
high−density lipoprotein particle
triglyceride−rich lipoprotein particle
platelet alpha granule lumen
membrane−bounded vesicle
extracellular organelle
0 5 10 15 20
Z Score
M9 magenta
defense response to fungus
positive regulation of response to external stimulus
regulation of erythrocyte differentiation
leukocyte chemotaxis
response to interleukin−1
cellular defense response
regulation of leukocyte chemotaxis
positive regulation of myeloid cell differentiation
growth factor activity
cytokine activity
serine−type endopeptidase activity
heparin binding
cytokine receptor binding
receptor activity
calcium ion binding
external side of plasma membrane
extracellular region part
membrane−bounded vesicle
plasma membrane
cell surface
extracellular region
nucleolus
0 1 2 3 4 5 6
Z Score
M10 purple
retina homeostasis
protein activation cascade
negative regulation of endopeptidase activity
biological adhesion
regulation of blood coagulation
cell motility
acute inflammatory response
regulation of proteolysis
heparin binding
endopeptidase inhibitor activity
receptor activity
growth factor binding
calcium ion binding
lipid binding
blood microparticle
extracellular space
extracellular region
extracellular organelle
anchored to membrane
extracellular matrix
cytoplasmic membrane−bounded vesicle lumen
lamellipodium
0 5 10 15
Z Score
M11 greenyellow
protein activation cascade
retina homeostasis
platelet degranulation
positive regulation of ERK1 and ERK2 cascade
biological adhesion
response to bacterium
platelet activation
positive regulation of cell adhesion
antigen binding
growth factor activity
carbohydrate binding
platelet alpha granule
blood microparticle
platelet alpha granule lumen
extracellular space
extracellular region
external side of plasma membrane
plasma membrane
extracellular organelle
0 2 4 6 8 10 12 14
Z Score

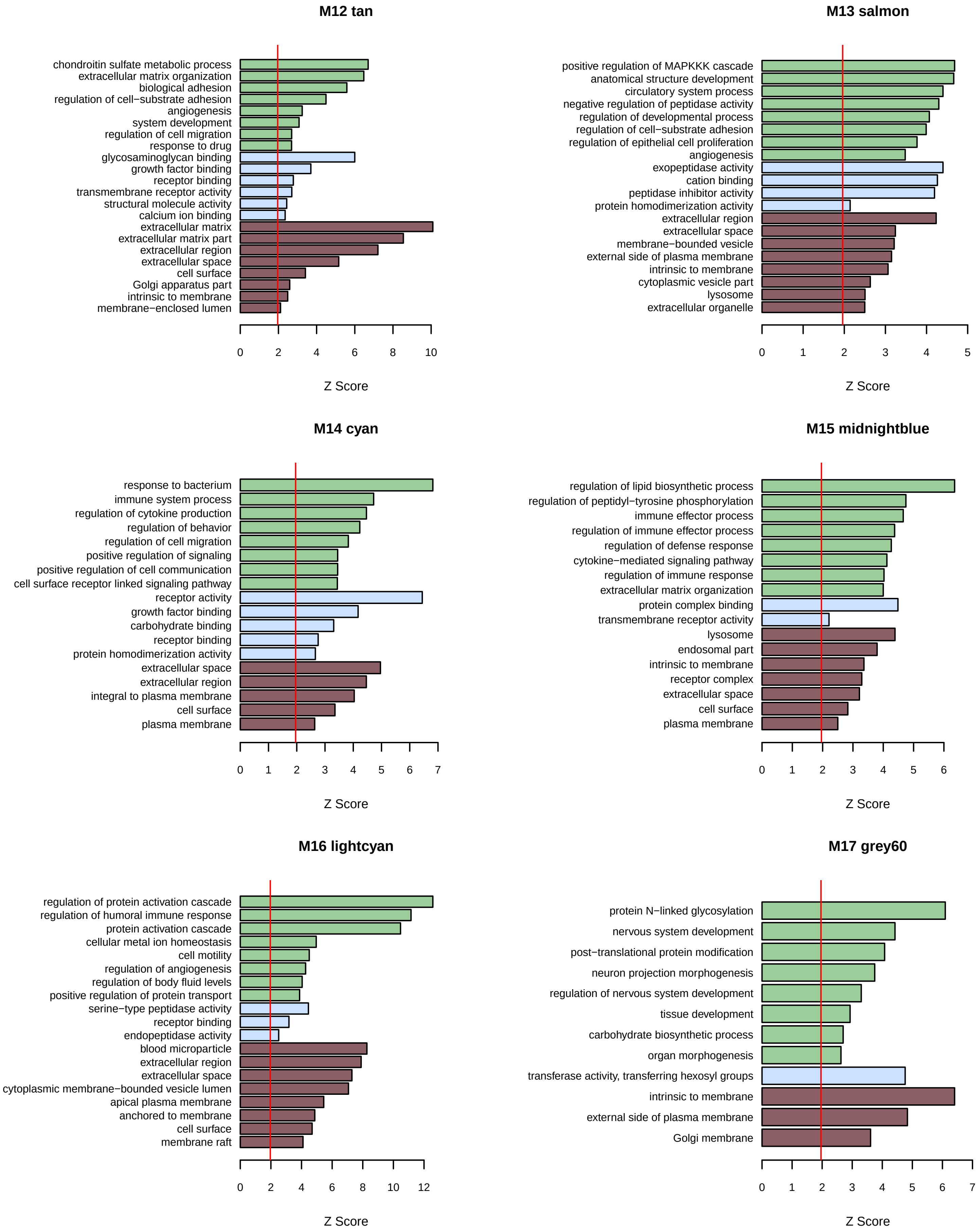
M12 tan
chondroitin sulfate metabolic process
extracellular matrix organization
biological adhesion
regulation of cell−substrate adhesion
angiogenesis
system development
regulation of cell migration
response to drug
glycosaminoglycan binding
growth factor binding
receptor binding
transmembrane receptor activity
structural molecule activity
calcium ion binding
extracellular matrix
extracellular matrix part
extracellular region
extracellular space
cell surface
Golgi apparatus part
intrinsic to membrane
membrane−enclosed lumen
0 2 4 6 8 10
Z Score
M13 salmon
positive regulation of MAPKKK cascade
anatomical structure development
circulatory system process
negative regulation of peptidase activity
regulation of developmental process
regulation of cell−substrate adhesion
regulation of epithelial cell proliferation
angiogenesis
exopeptidase activity
cation binding
peptidase inhibitor activity
protein homodimerization activity
extracellular region
extracellular space
membrane−bounded vesicle
external side of plasma membrane
intrinsic to membrane
cytoplasmic vesicle part
lysosome
extracellular organelle
0 1 2 3 4 5
Z Score
M14 cyan
response to bacterium
immune system process
regulation of cytokine production
regulation of behavior
regulation of cell migration
positive regulation of signaling
positive regulation of cell communication
cell surface receptor linked signaling pathway
receptor activity
growth factor binding
carbohydrate binding
receptor binding
protein homodimerization activity
extracellular space
extracellular region
integral to plasma membrane
cell surface
plasma membrane
0 1 2 3 4 5 6 7
Z Score
M15 midnightblue
regulation of lipid biosynthetic process
regulation of peptidyl−tyrosine phosphorylation
immune effector process
regulation of immune effector process
regulation of defense response
cytokine−mediated signaling pathway
regulation of immune response
extracellular matrix organization
protein complex binding
transmembrane receptor activity
lysosome
endosomal part
intrinsic to membrane
receptor complex
extracellular space
cell surface
plasma membrane
0 1 2 3 4 5 6
Z Score
M16 lightcyan
regulation of protein activation cascade
regulation of humoral immune response
protein activation cascade
cellular metal ion homeostasis
cell motility
regulation of angiogenesis
regulation of body fluid levels
positive regulation of protein transport
serine−type peptidase activity
receptor binding
endopeptidase activity
blood microparticle
extracellular region
extracellular space
cytoplasmic membrane−bounded vesicle lumen
apical plasma membrane
anchored to membrane
cell surface
membrane raft
0 2 4 6 8 10 12
Z Score
M17 grey60
protein N−linked glycosylation
nervous system development
post−translational protein modification
neuron projection morphogenesis
regulation of nervous system development
tissue development
carbohydrate biosynthetic process
organ morphogenesis
transferase activity, transferring hexosyl groups
intrinsic to membrane
external side of plasma membrane
Golgi membrane
0 1 2 3 4 5 6 7
Z Score

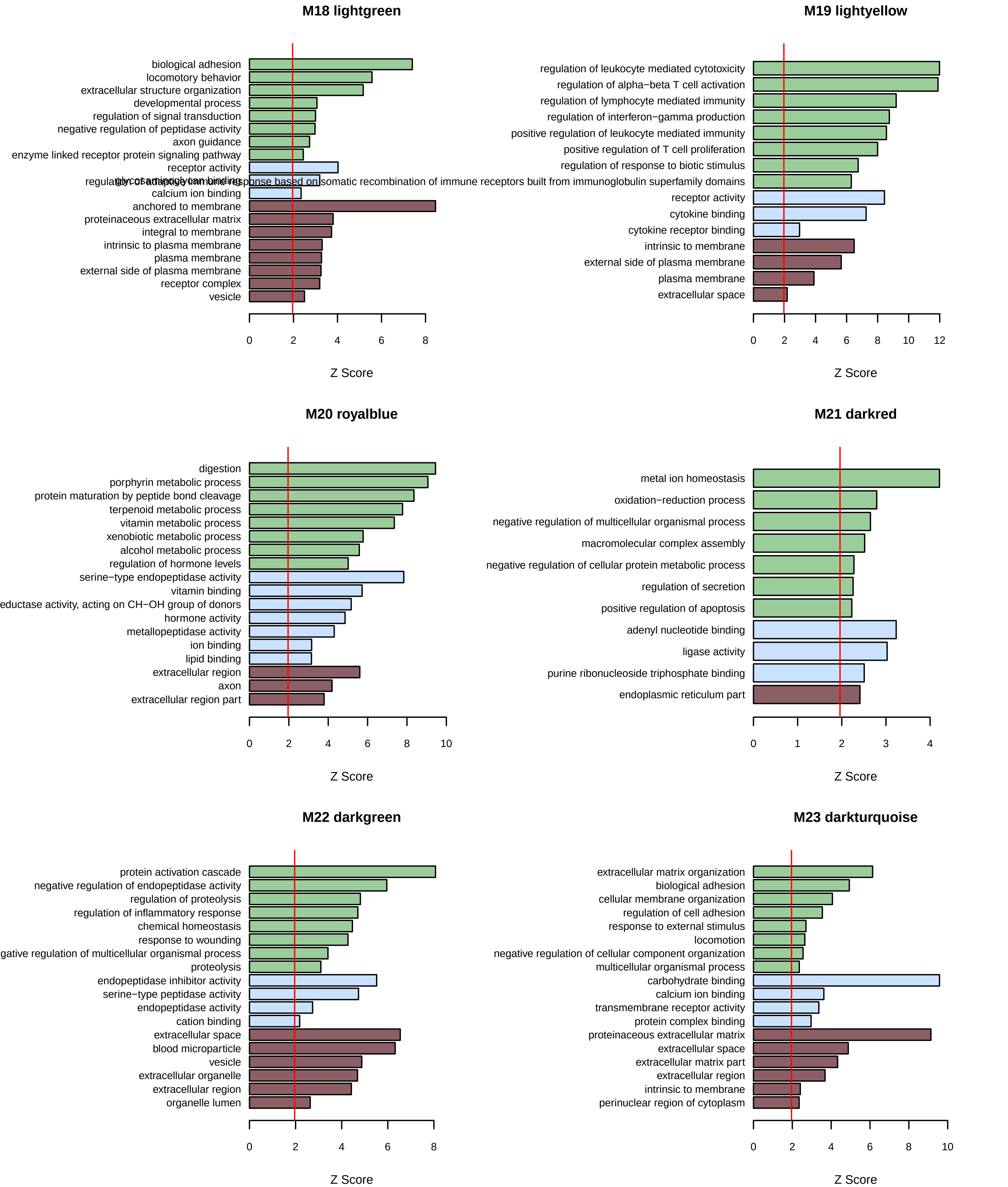
M18 lightgreen
biological adhesion
locomotory behavior
extracellular structure organization
developmental process
regulation of signal transduction
negative regulation of peptidase activity
axon guidance
enzyme linked receptor protein signaling pathway
receptor activity
glycosaminoglycan binding
calcium ion binding
anchored to membrane
proteinaceous extracellular matrix
integral to membrane
intrinsic to plasma membrane
plasma membrane
external side of plasma membrane
receptor complex
vesicle
0 2 4 6 8
Z Score
M19 lightyellow
regulation of leukocyte mediated cytotoxicity
regulation of alpha−beta T cell activation
regulation of lymphocyte mediated immunity
regulation of interferon−gamma production
positive regulation of leukocyte mediated immunity
positive regulation of T cell proliferation
regulation of response to biotic stimulus
regulation of adaptive immune response based on somatic recombination of immune receptors built from immunoglobulin superfamily domains
receptor activity
cytokine binding
cytokine receptor binding
intrinsic to membrane
external side of plasma membrane
plasma membrane
extracellular space
0 2 4 6 8 10 12
Z Score
M20 royalblue
digestion
porphyrin metabolic process
protein maturation by peptide bond cleavage
terpenoid metabolic process
vitamin metabolic process
xenobiotic metabolic process
alcohol metabolic process
regulation of hormone levels
serine−type endopeptidase activity
vitamin binding
eductase activity, acting on CH−OH group of donors
hormone activity
metallopeptidase activity
ion binding
lipid binding
extracellular region
axon
extracellular region part
0 2 4 6 8 10
Z Score
M21 darkred
metal ion homeostasis
oxidation−reduction process
negative regulation of multicellular organismal process
macromolecular complex assembly
negative regulation of cellular protein metabolic process
regulation of secretion
positive regulation of apoptosis
adenyl nucleotide binding
ligase activity
purine ribonucleoside triphosphate binding
endoplasmic reticulum part
0 1 2 3 4
Z Score
M22 darkgreen
protein activation cascade
negative regulation of endopeptidase activity
regulation of proteolysis
regulation of inflammatory response
chemical homeostasis
response to wounding
gative regulation of multicellular organismal process
proteolysis
endopeptidase inhibitor activity
serine−type peptidase activity
endopeptidase activity
cation binding
extracellular space
blood microparticle
vesicle
extracellular organelle
extracellular region
organelle lumen
0 2 4 6 8
Z Score
M23 darkturquoise
extracellular matrix organization
biological adhesion
cellular membrane organization
regulation of cell adhesion
response to external stimulus
locomotion
negative regulation of cellular component organization
multicellular organismal process
carbohydrate binding
calcium ion binding
transmembrane receptor activity
protein complex binding
proteinaceous extracellular matrix
extracellular space
extracellular matrix part
extracellular region
intrinsic to membrane
perinuclear region of cytoplasm
0 2 4 6 8 10
Z Score

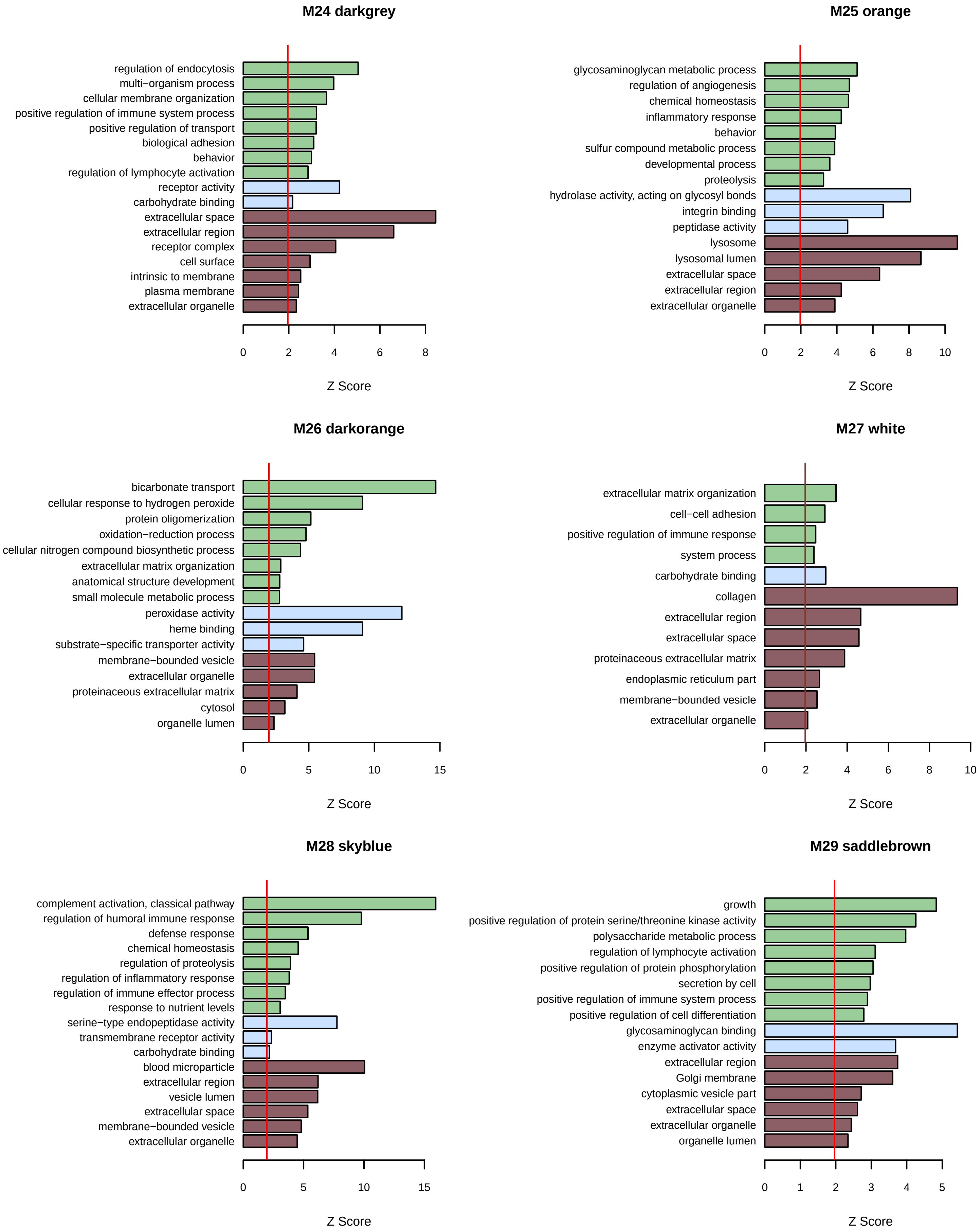

M24 darkgrey
regulation of endocytosis
multi−organism process
cellular membrane organization
positive regulation of immune system process
positive regulation of transport
biological adhesion
behavior
regulation of lymphocyte activation
receptor activity
carbohydrate binding
extracellular space
extracellular region
receptor complex
cell surface
intrinsic to membrane
plasma membrane
extracellular organelle
0 2 4 6 8
Z Score
M25 orange
glycosaminoglycan metabolic process
regulation of angiogenesis
chemical homeostasis
inflammatory response
behavior
sulfur compound metabolic process
developmental process
proteolysis
hydrolase activity, acting on glycosyl bonds
integrin binding
peptidase activity
lysosome
lysosomal lumen
extracellular space
extracellular region
extracellular organelle
0 2 4 6 8 10
Z Score
M26 darkorange
bicarbonate transport
cellular response to hydrogen peroxide
protein oligomerization
oxidation−reduction process
cellular nitrogen compound biosynthetic process
extracellular matrix organization
anatomical structure development
small molecule metabolic process
peroxidase activity
heme binding
substrate−specific transporter activity
membrane−bounded vesicle
extracellular organelle
proteinaceous extracellular matrix
cytosol
organelle lumen
0 5 10 15
Z Score
M27 white
extracellular matrix organization
cell−cell adhesion
positive regulation of immune response
system process
carbohydrate binding
collagen
extracellular region
extracellular space
proteinaceous extracellular matrix
endoplasmic reticulum part
membrane−bounded vesicle
extracellular organelle
0 2 4 6 8 10
Z Score
M28 skyblue
complement activation, classical pathway
regulation of humoral immune response
defense response
chemical homeostasis
regulation of proteolysis
regulation of inflammatory response
regulation of immune effector process
response to nutrient levels
serine−type endopeptidase activity
transmembrane receptor activity
carbohydrate binding
blood microparticle
extracellular region
vesicle lumen
extracellular space
membrane−bounded vesicle
extracellular organelle
0 5 10 15
Z Score
M29 saddlebrown
growth
positive regulation of protein serine/threonine kinase activity
polysaccharide metabolic process
regulation of lymphocyte activation
positive regulation of protein phosphorylation
secretion by cell
positive regulation of immune system process
positive regulation of cell differentiation
glycosaminoglycan binding
enzyme activator activity
extracellular region
Golgi membrane
cytoplasmic vesicle part
extracellular space
extracellular organelle
organelle lumen
0 1 2 3 4 5
Z Score

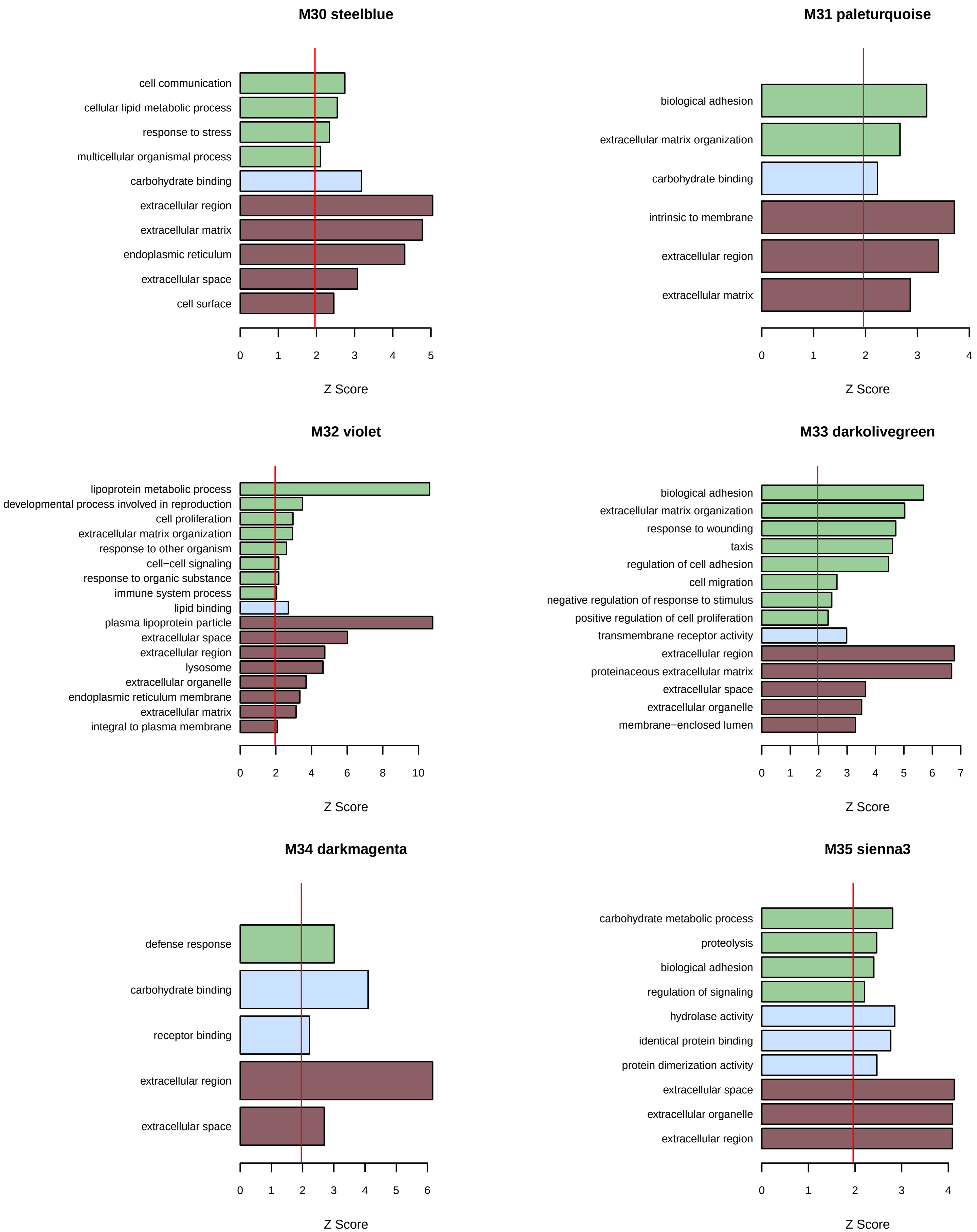
M30 steelblue
cell communication
cellular lipid metabolic process
response to stress
multicellular organismal process
carbohydrate binding
extracellular region
extracellular matrix
endoplasmic reticulum
extracellular space
cell surface
0
1
2
3
4
5
Z Score
M31 paleturquoise
biological adhesion
extracellular matrix organization
carbohydrate binding
intrinsic to membrane
extracellular region
extracellular matrix
0
1
2
3
4
Z Score
M32 violet
lipoprotein metabolic process
developmental process involved in reproduction
cell proliferation
extracellular matrix organization
response to other organism
cell−cell signaling
response to organic substance
immune system process
lipid binding
plasma lipoprotein particle
extracellular space
extracellular region
lysosome
extracellular organelle
endoplasmic reticulum membrane
extracellular matrix
integral to plasma membrane
0
2
4
6
8
10
Z Score
M33 darkolivegreen
biological adhesion
extracellular matrix organization
response to wounding
taxis
regulation of cell adhesion
cell migration
negative regulation of response to stimulus
positive regulation of cell proliferation
transmembrane receptor activity
extracellular region
proteinaceous extracellular matrix
extracellular space
extracellular organelle
membrane−enclosed lumen
0
1
2
3
4
5
6
7
Z Score
M34 darkmagenta
defense response
carbohydrate binding
receptor binding
extracellular region
extracellular space
0
1
2
3
4
5
6
Z Score
M35 sienna3
carbohydrate metabolic process
proteolysis
biological adhesion
regulation of signaling
hydrolase activity
identical protein binding
protein dimerization activity
extracellular space
extracellular organelle
extracellular region
0
1
2
3
4
Z Score

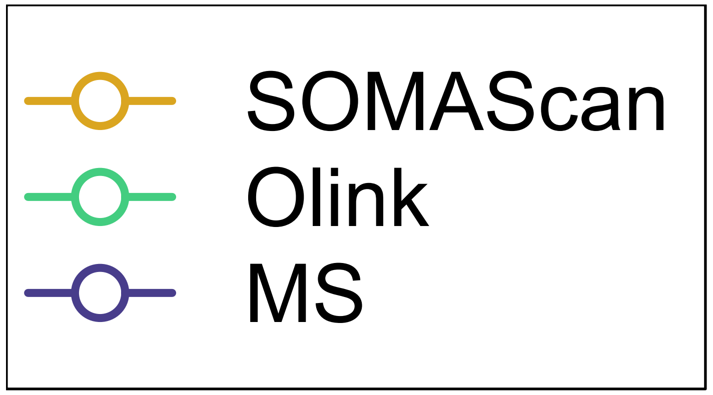
SOMAScan
Olink
MS

**M26 darkorange module: Bicarbonate Transport/Peroxidase**

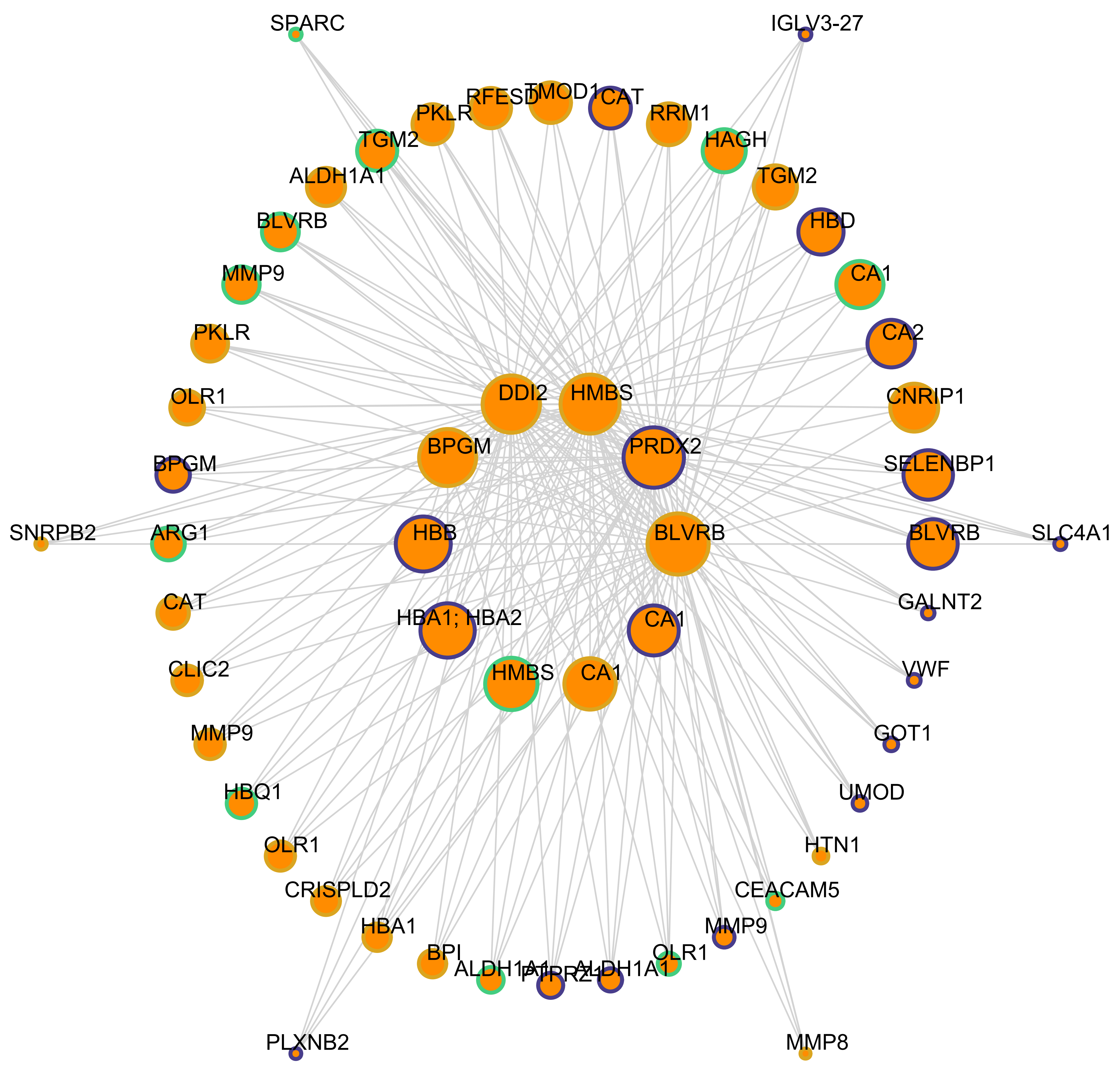

**M4 yellow module: T Cell Regulation**

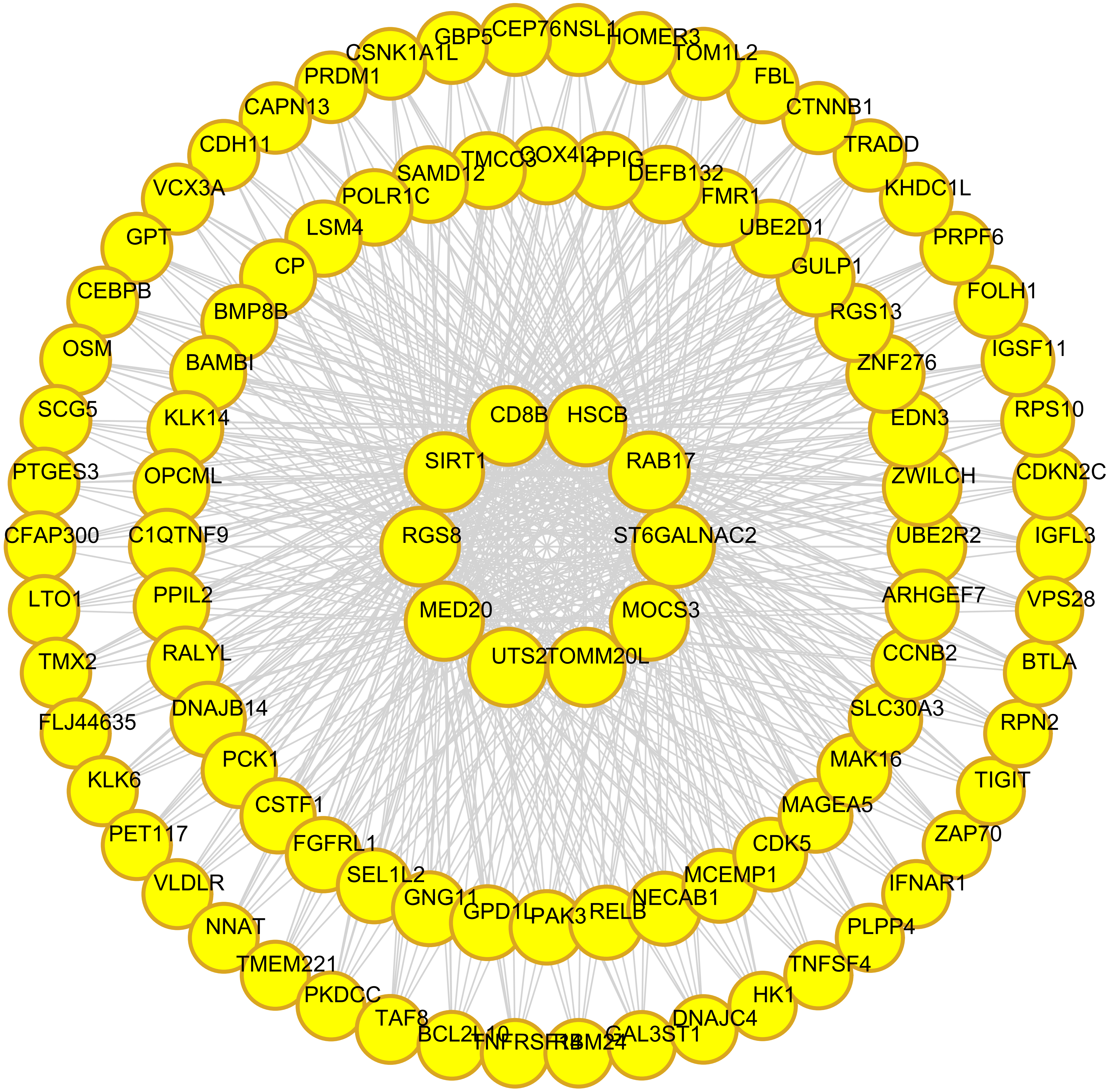

**M9 magenta module: Leukocyte Chemotaxis**

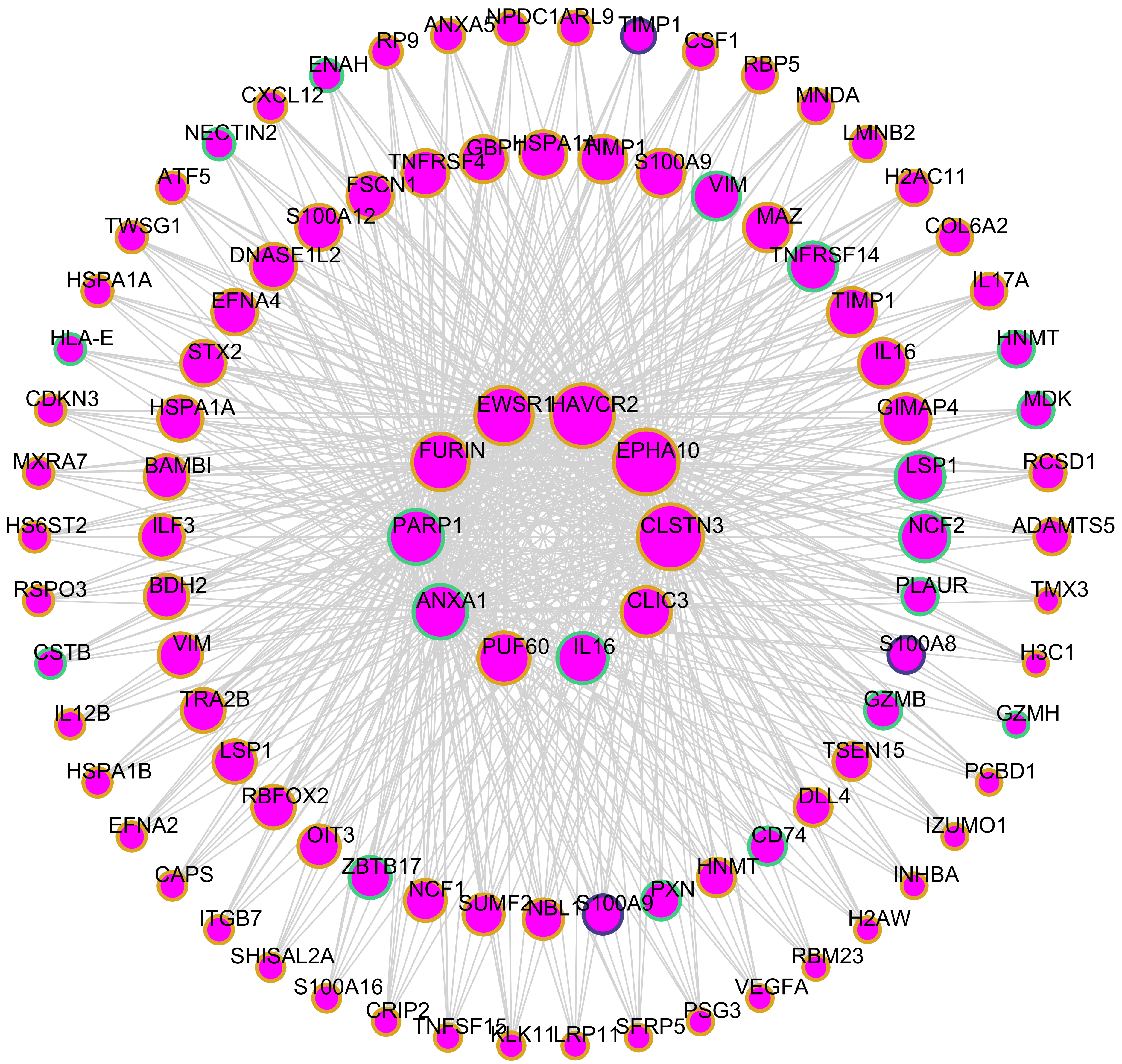

**M1 turquoise module: Cellular Metabolism/Intracellular Transport**

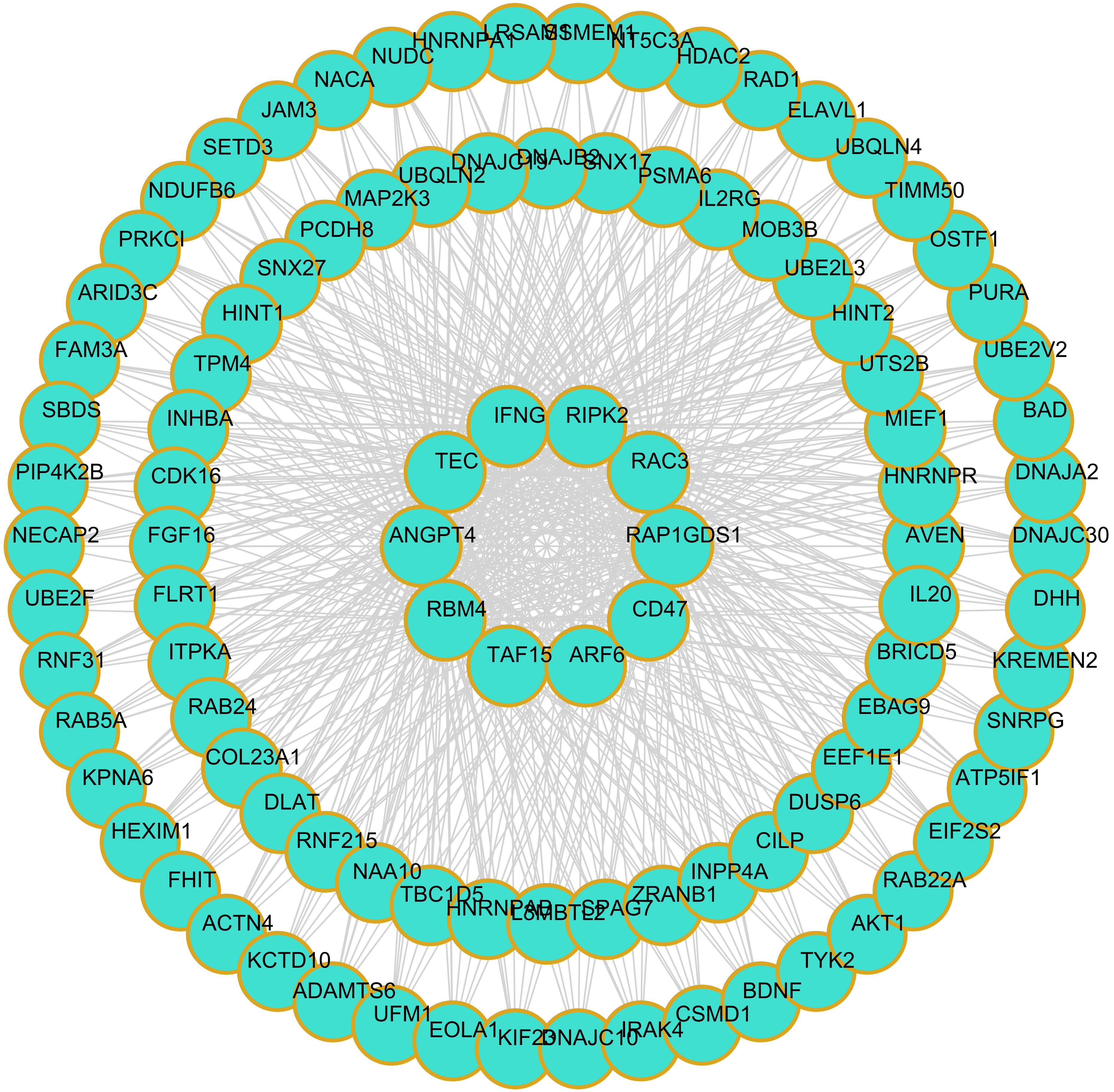

**M2 blue module: Ambiguous**

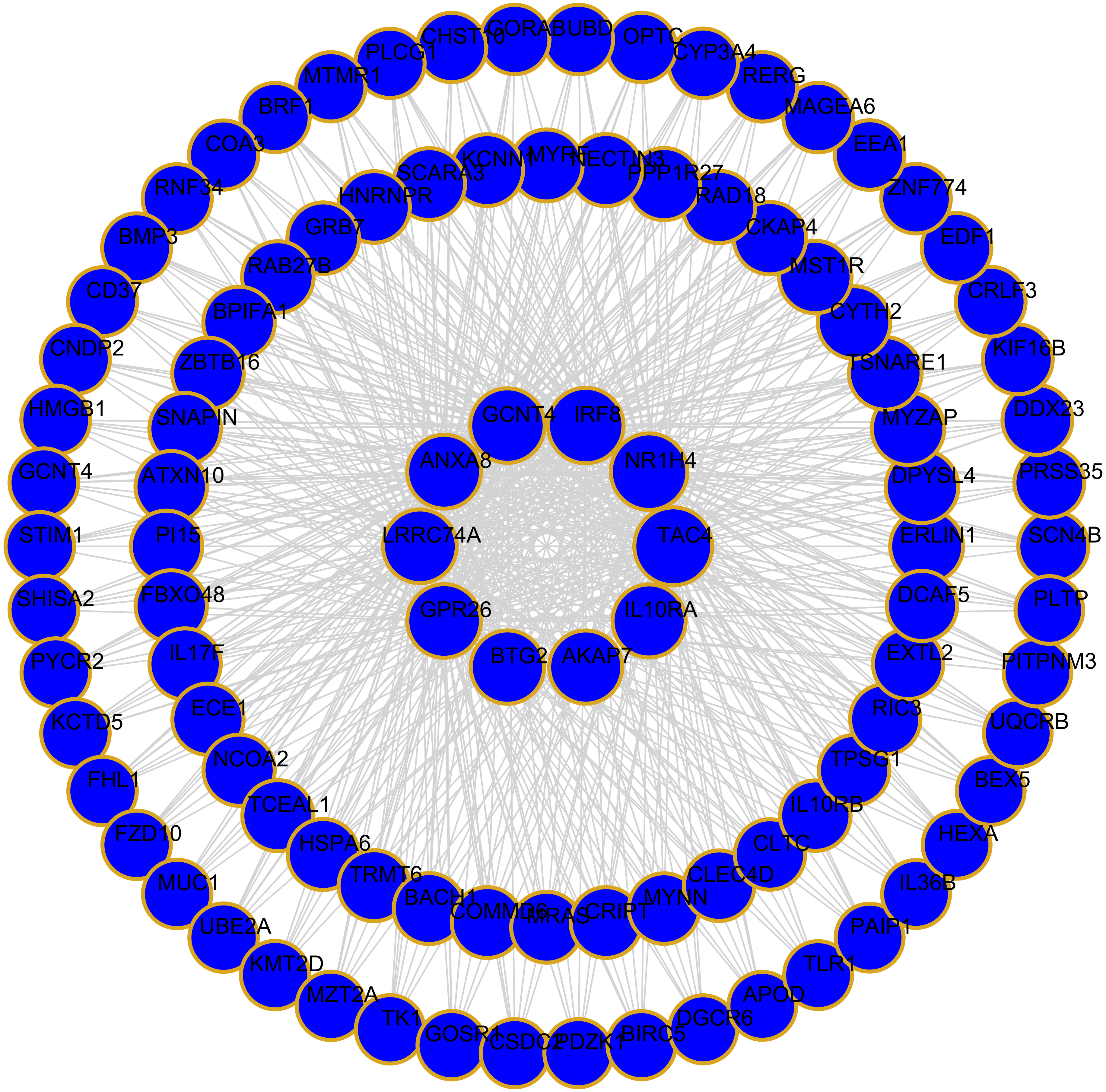

**M3 brown module: Ambiguous**

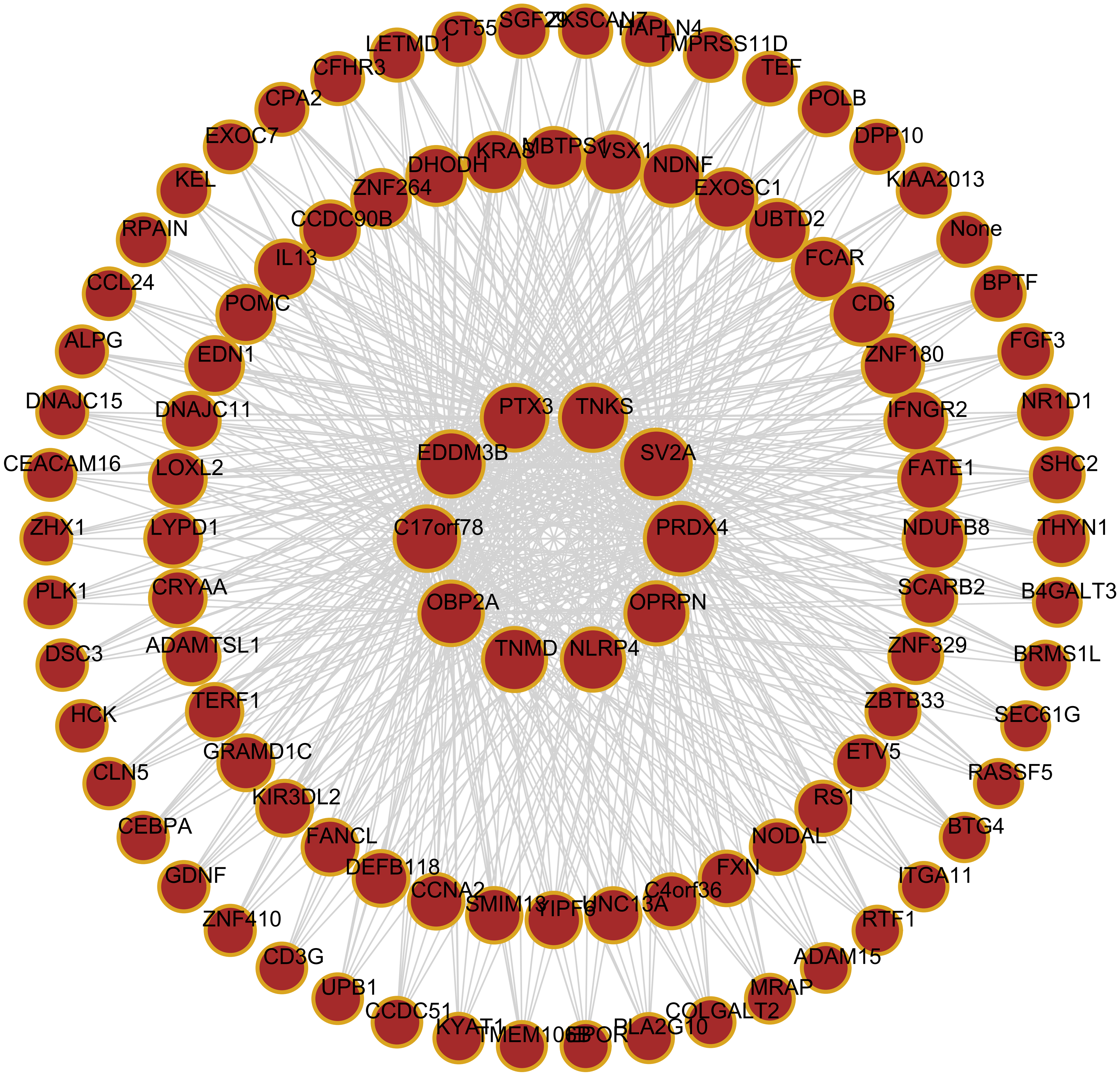

**M31 paleturquoise module: Adhesion/ECM**

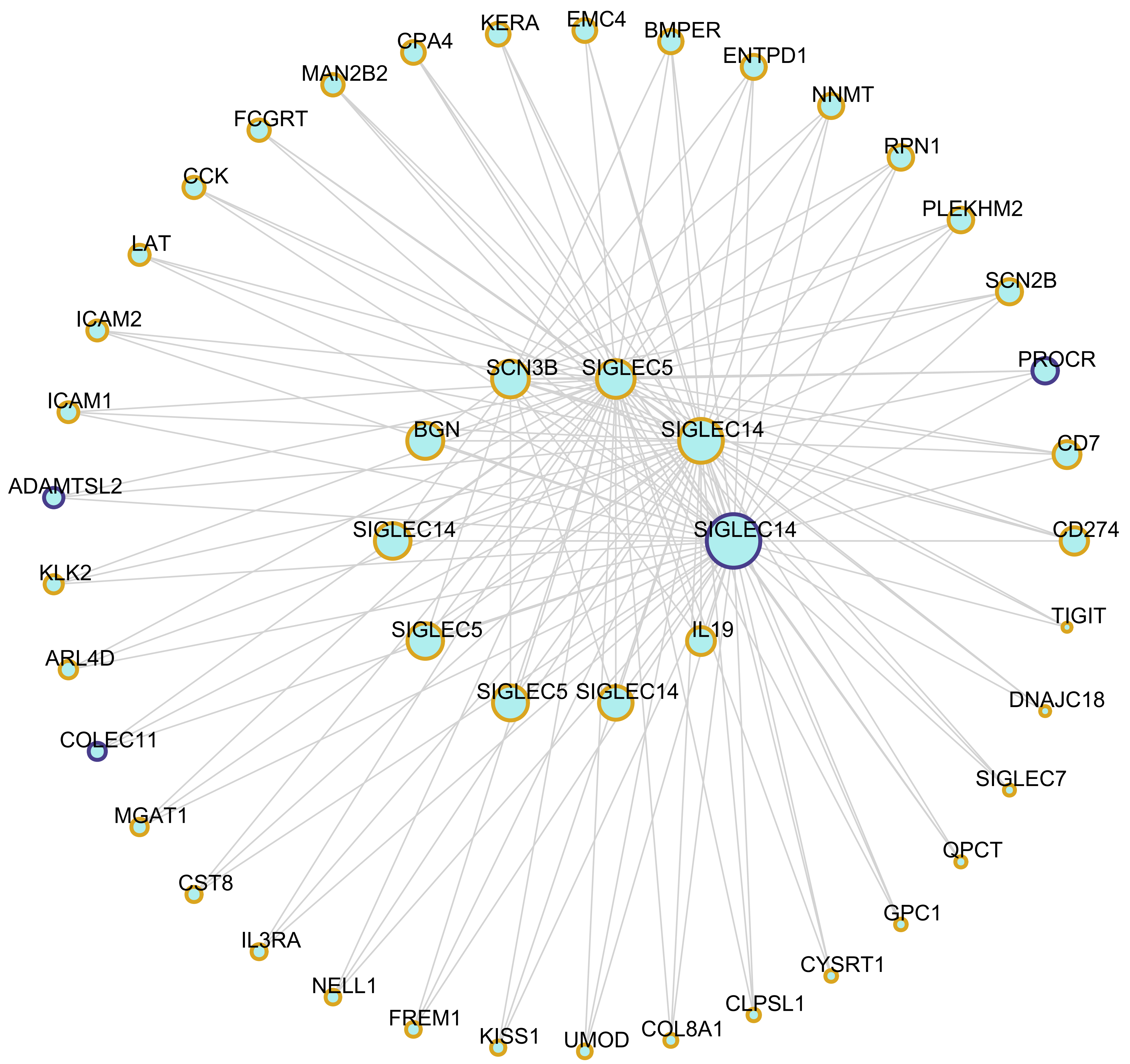

**M14 cyan module: Innate Immune Response**

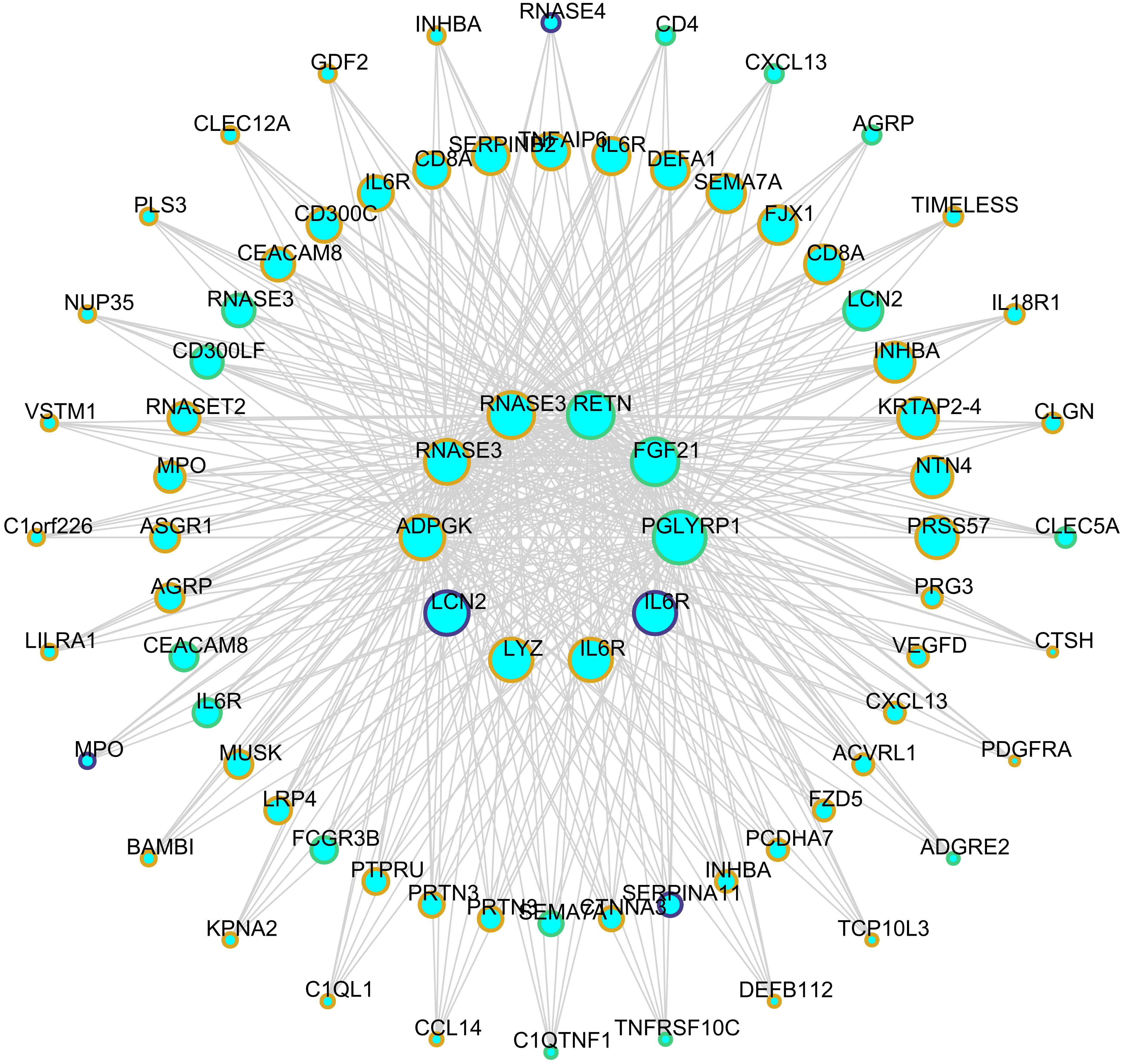

**M19 lightyellow module: Leukocyte Activation**

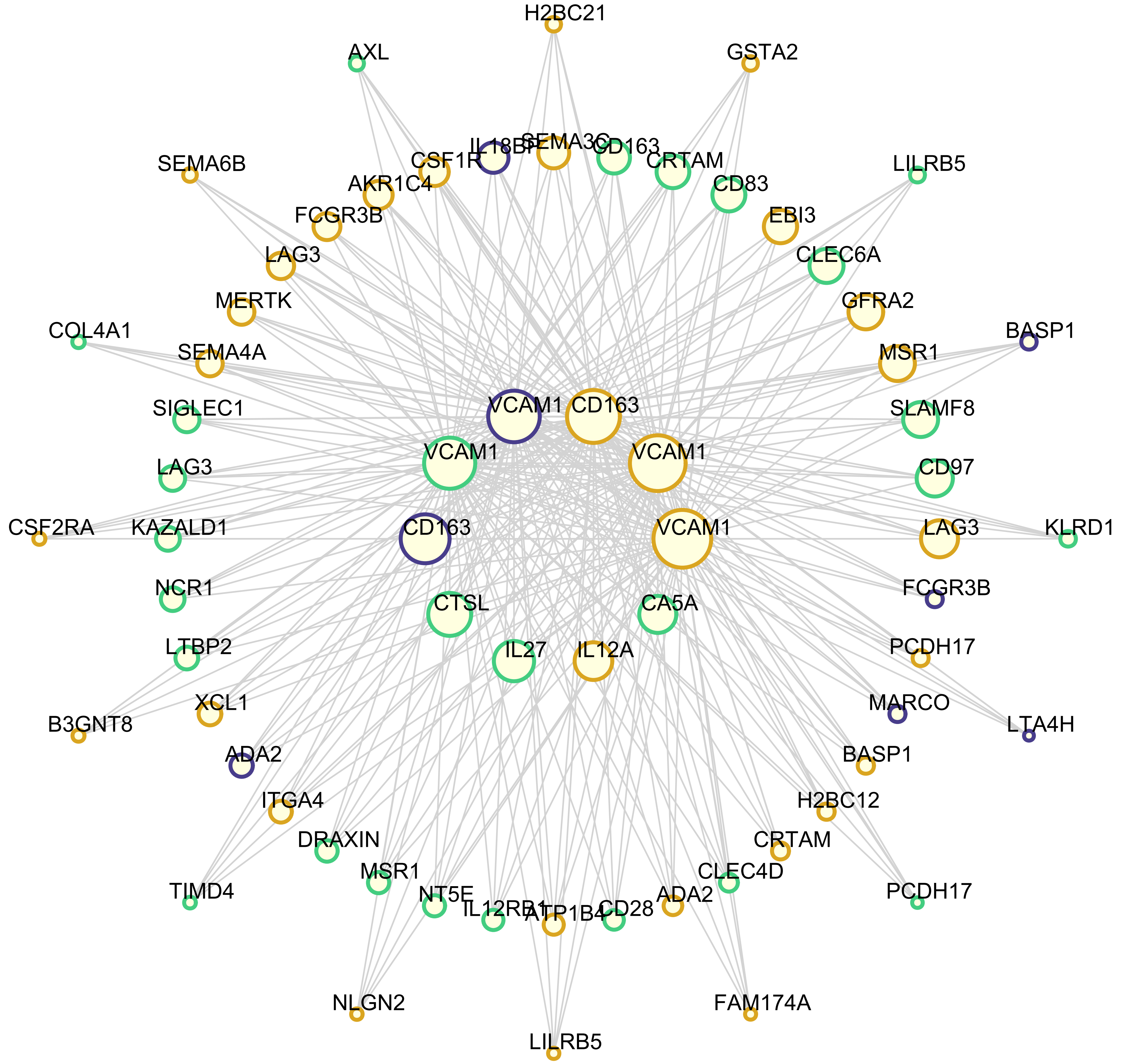

**M15 midnightblue module: Lipid Biosynthesis/Immune Response**

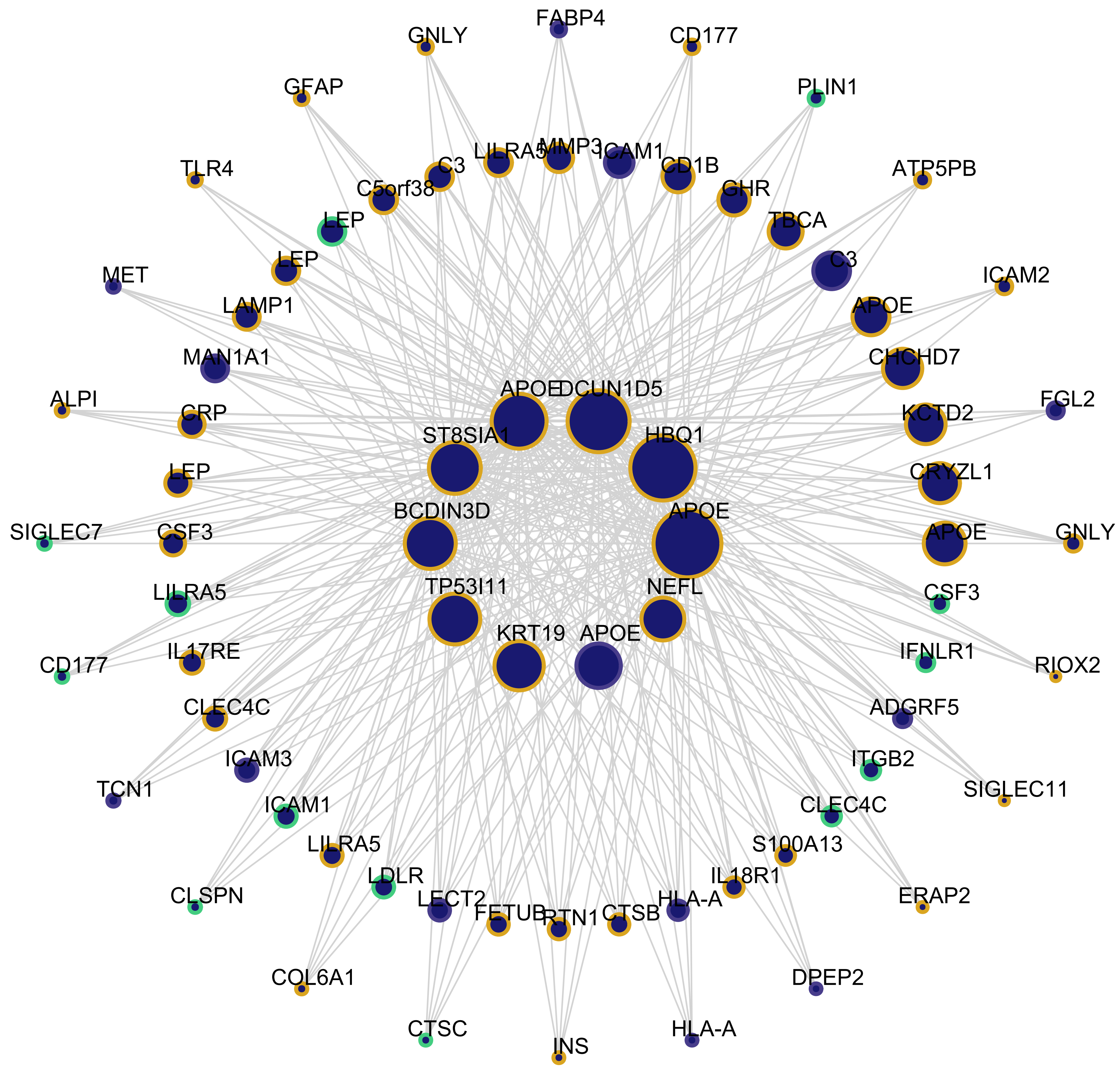

#### M29 saddlebrown module: IGF-Growth/GAG Binding

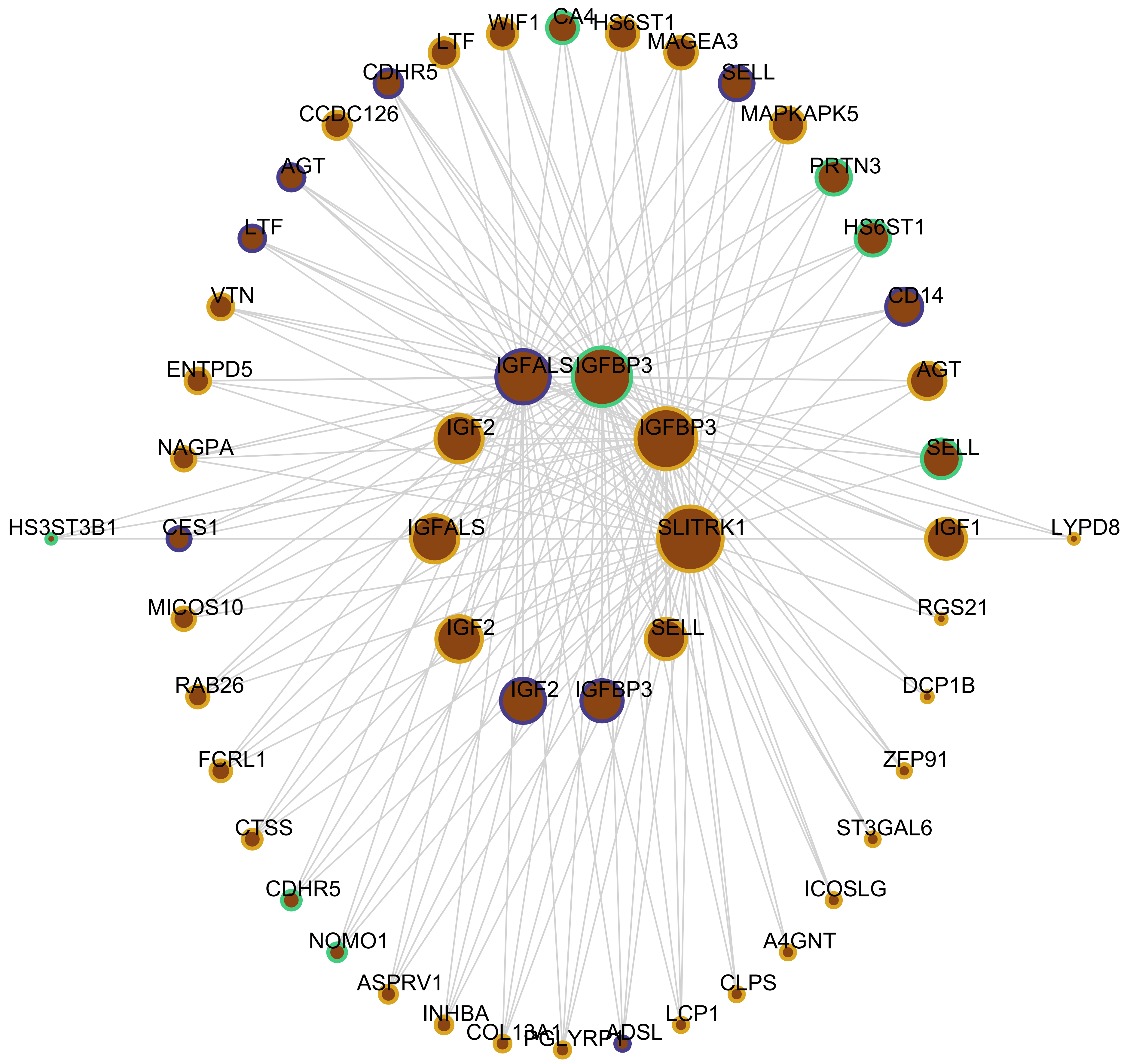

M28 skyblue module: Complement/Humoral Immune Response

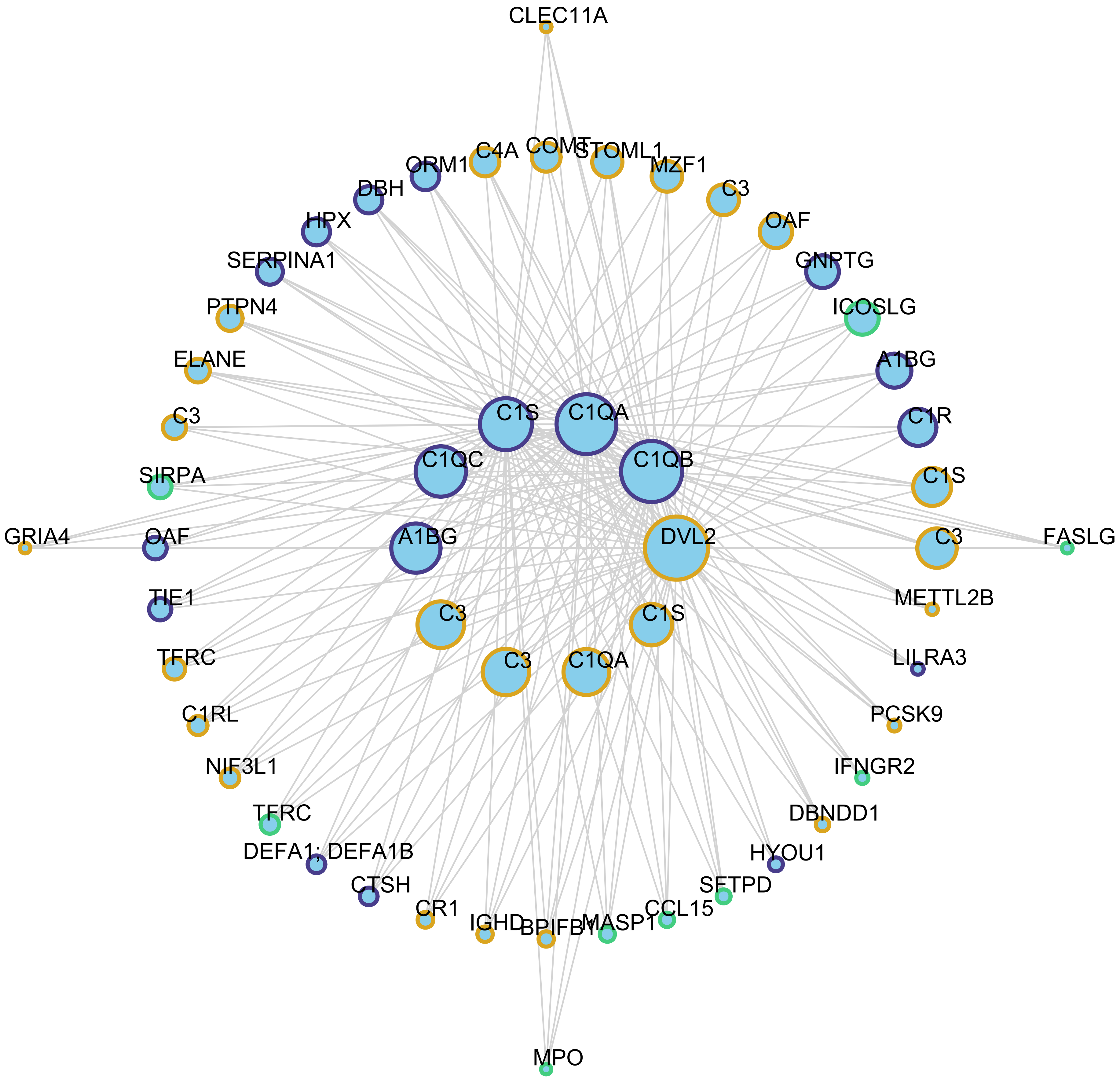

#### M21 darkred module: Metal Ion Homeostasis

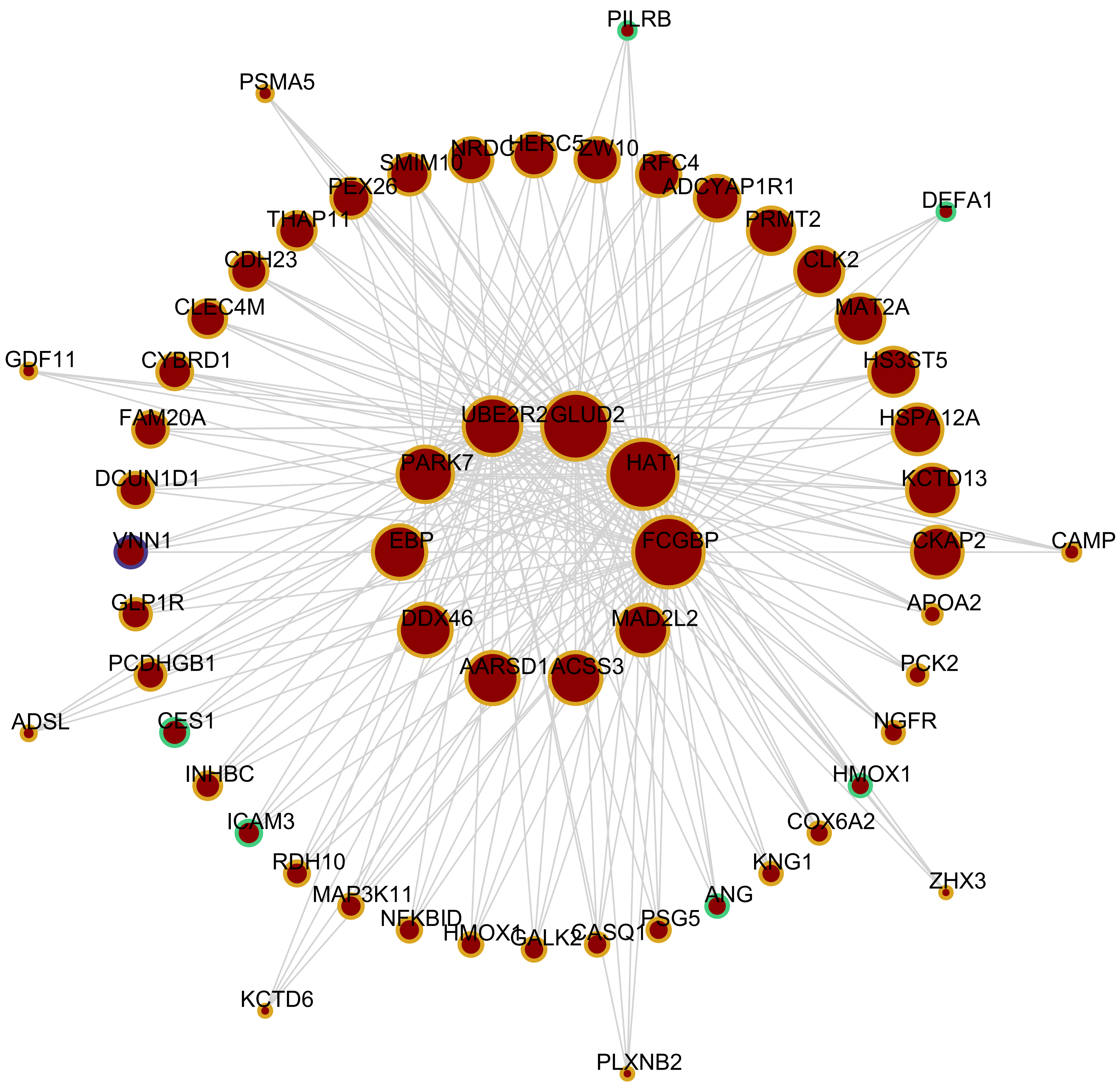

M32 violet module: Lipoprotein Metabolism

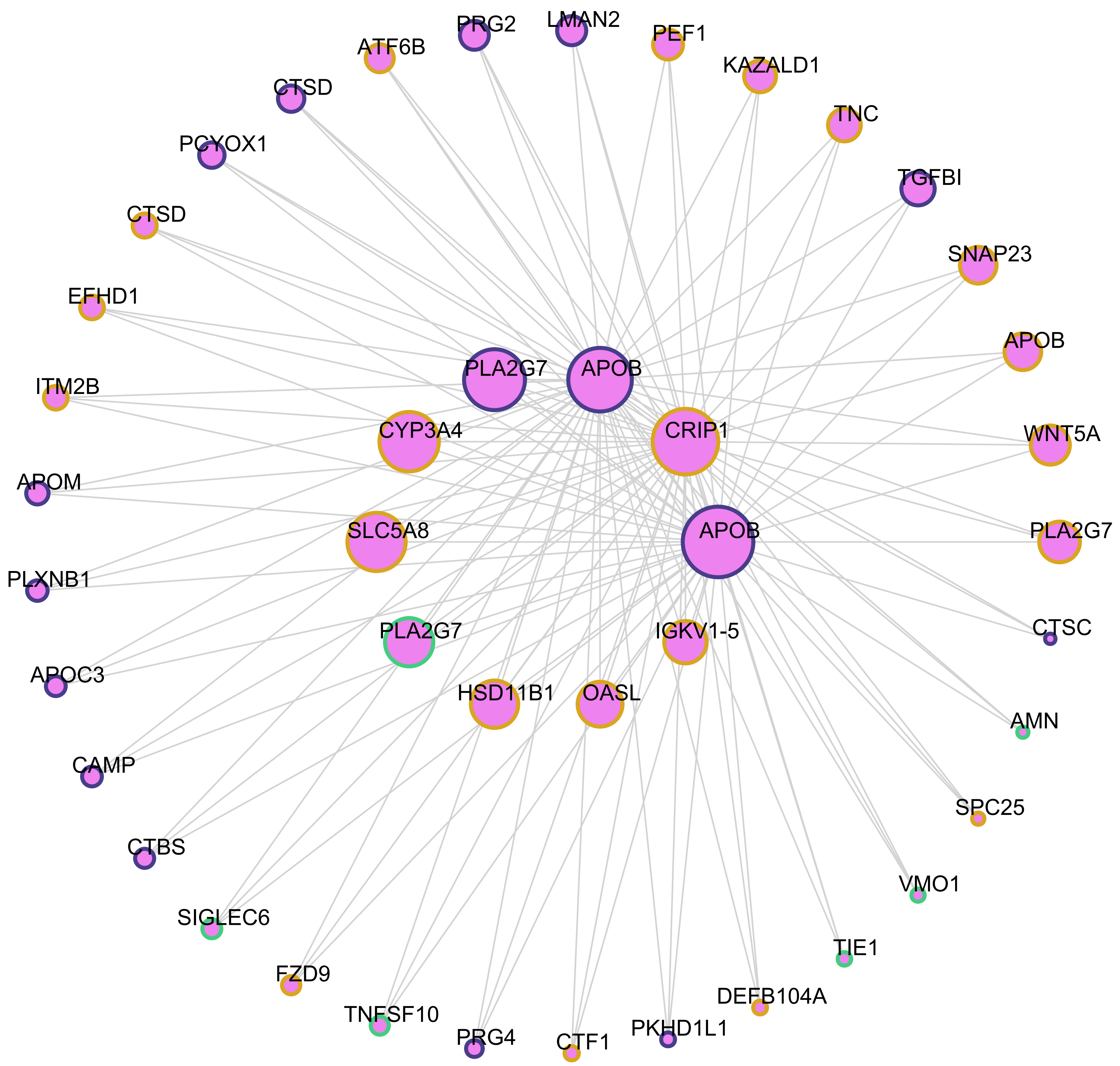

**M11 greenyellow module: Immunoglobulins/Coagulation Cascade**

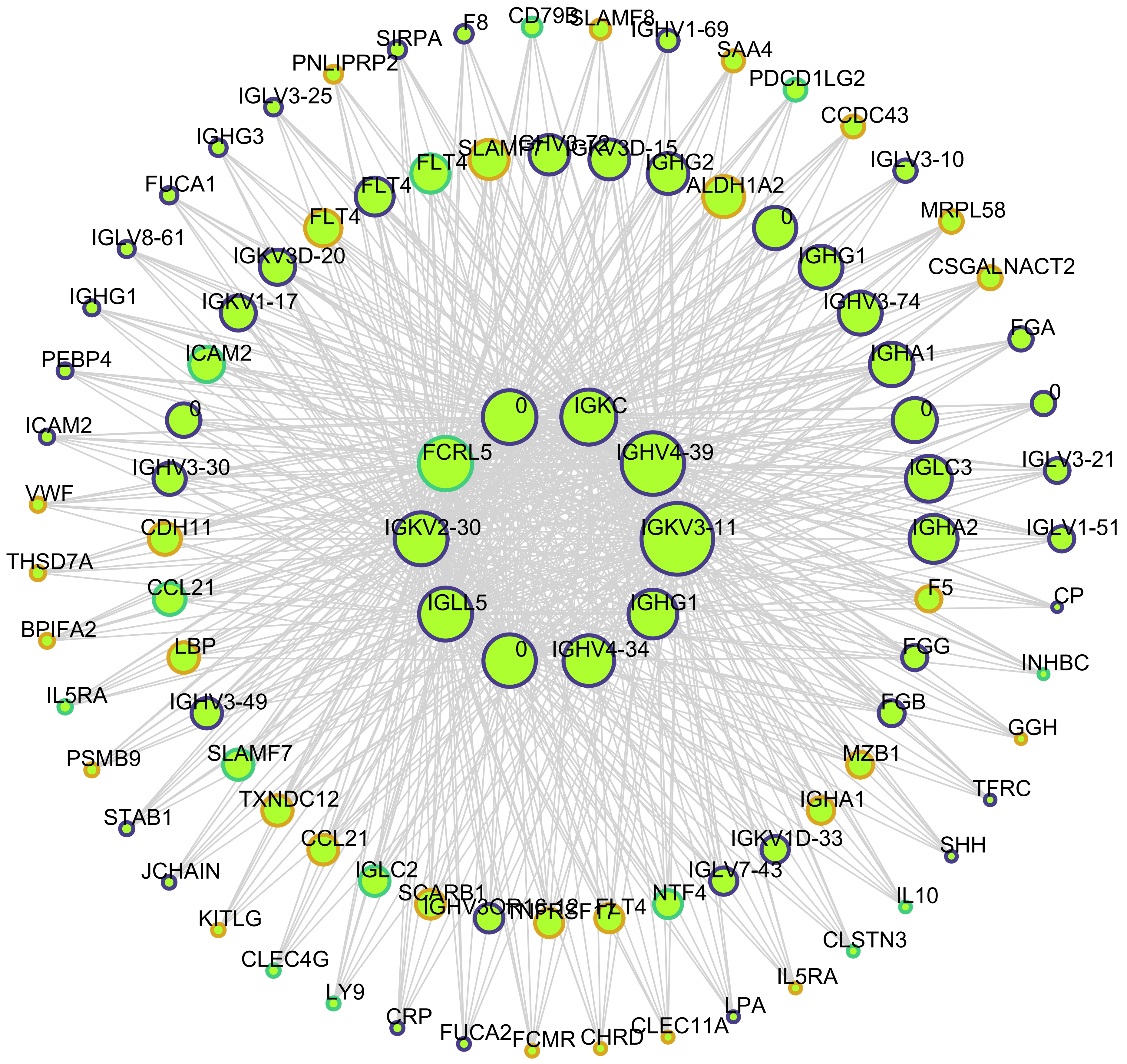

**M27 white module: Collagen/ECM**

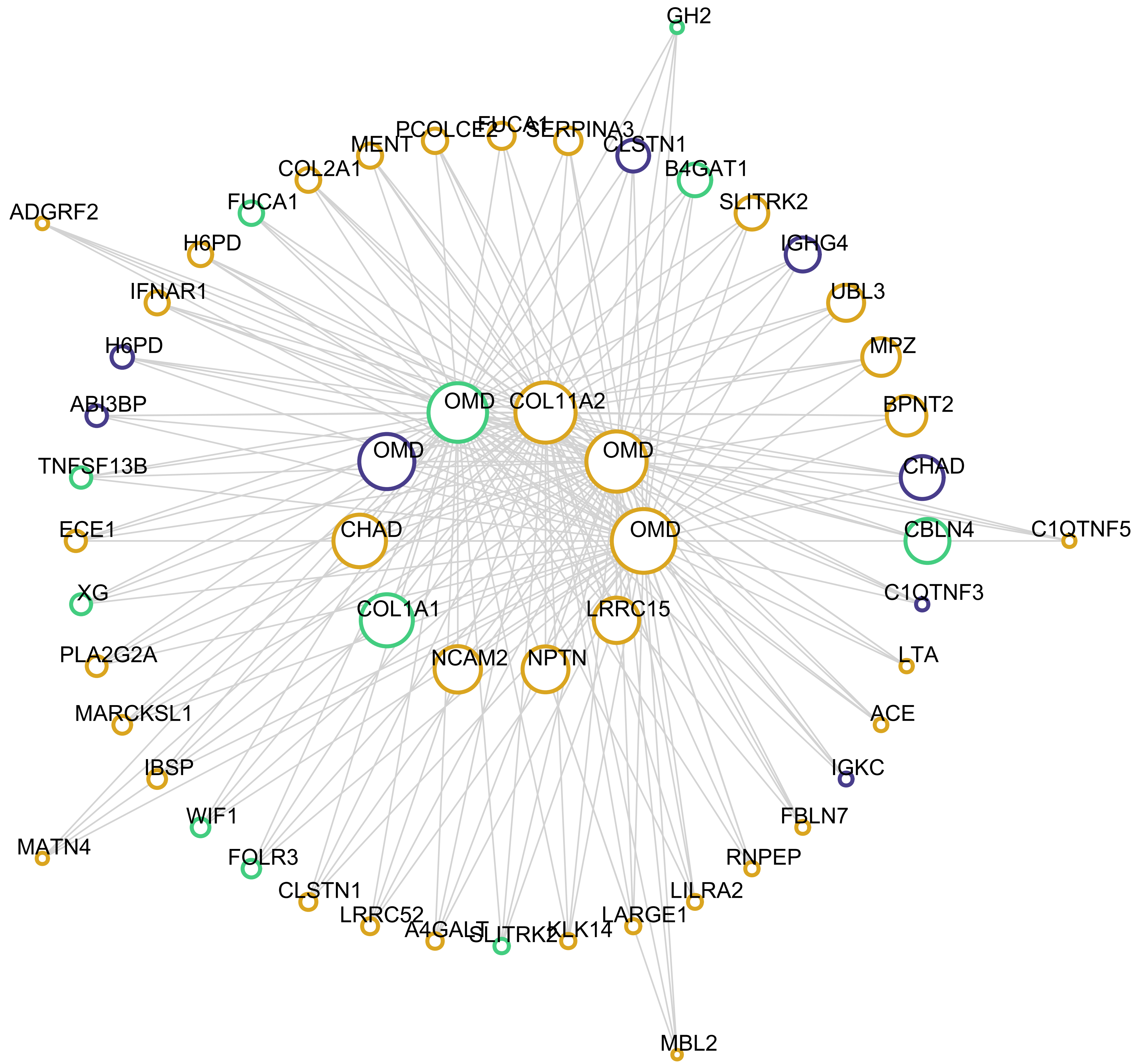

**M25 orange module: Lysosome**

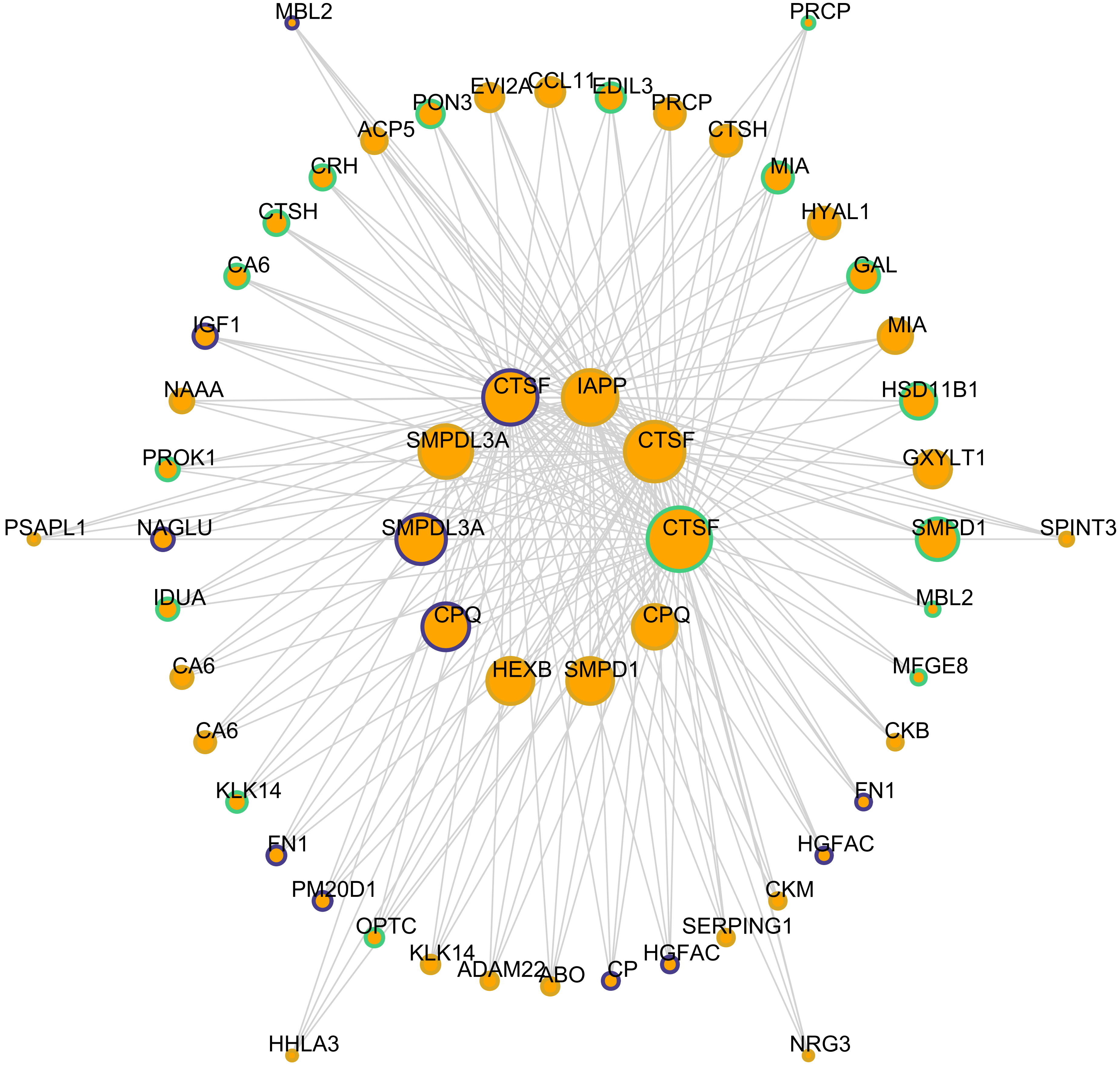

**M13 salmon module: Ambiguous**

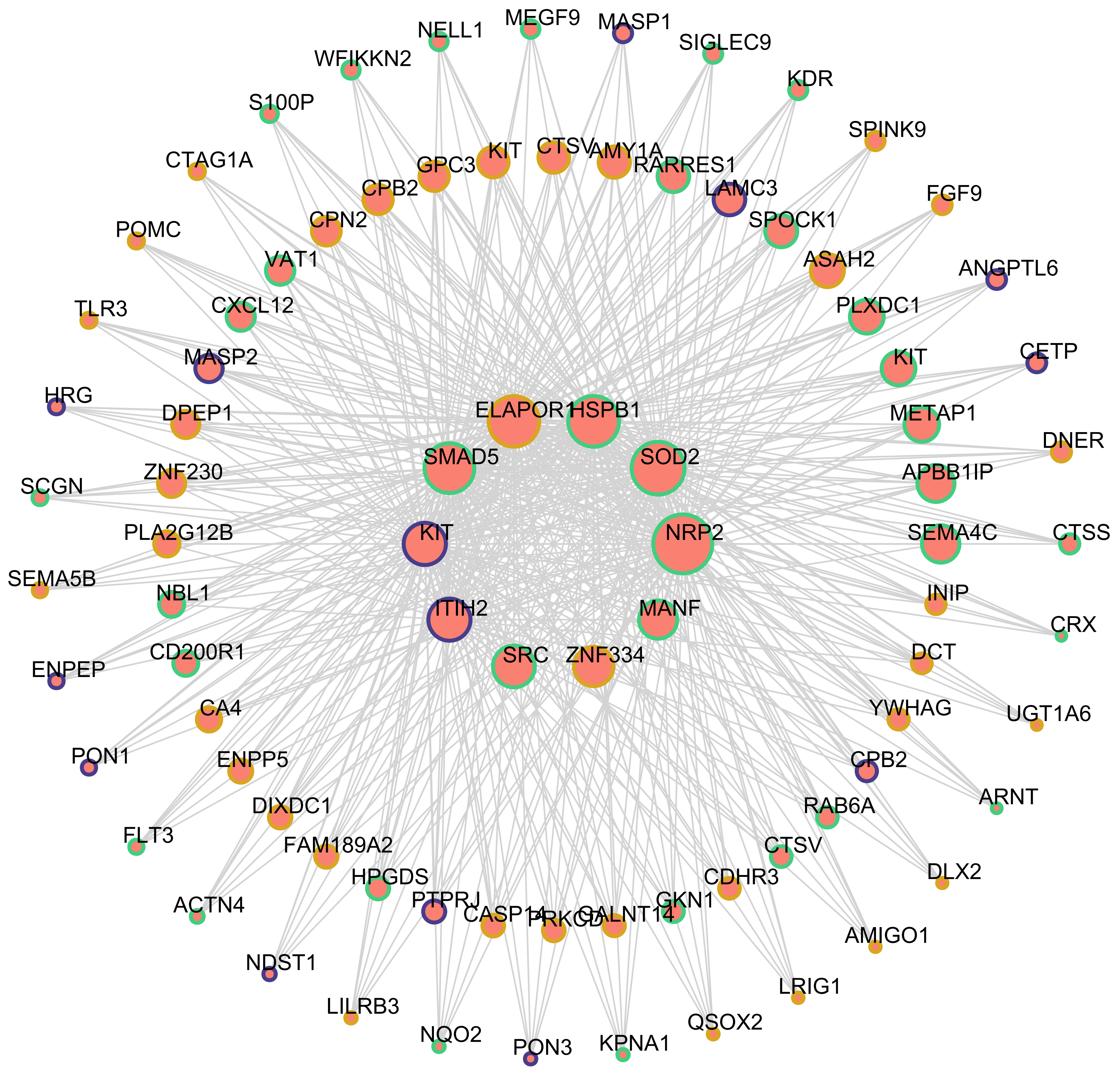

**M22 darkgreen module: Protein Activation Cascade/Proteolysis**

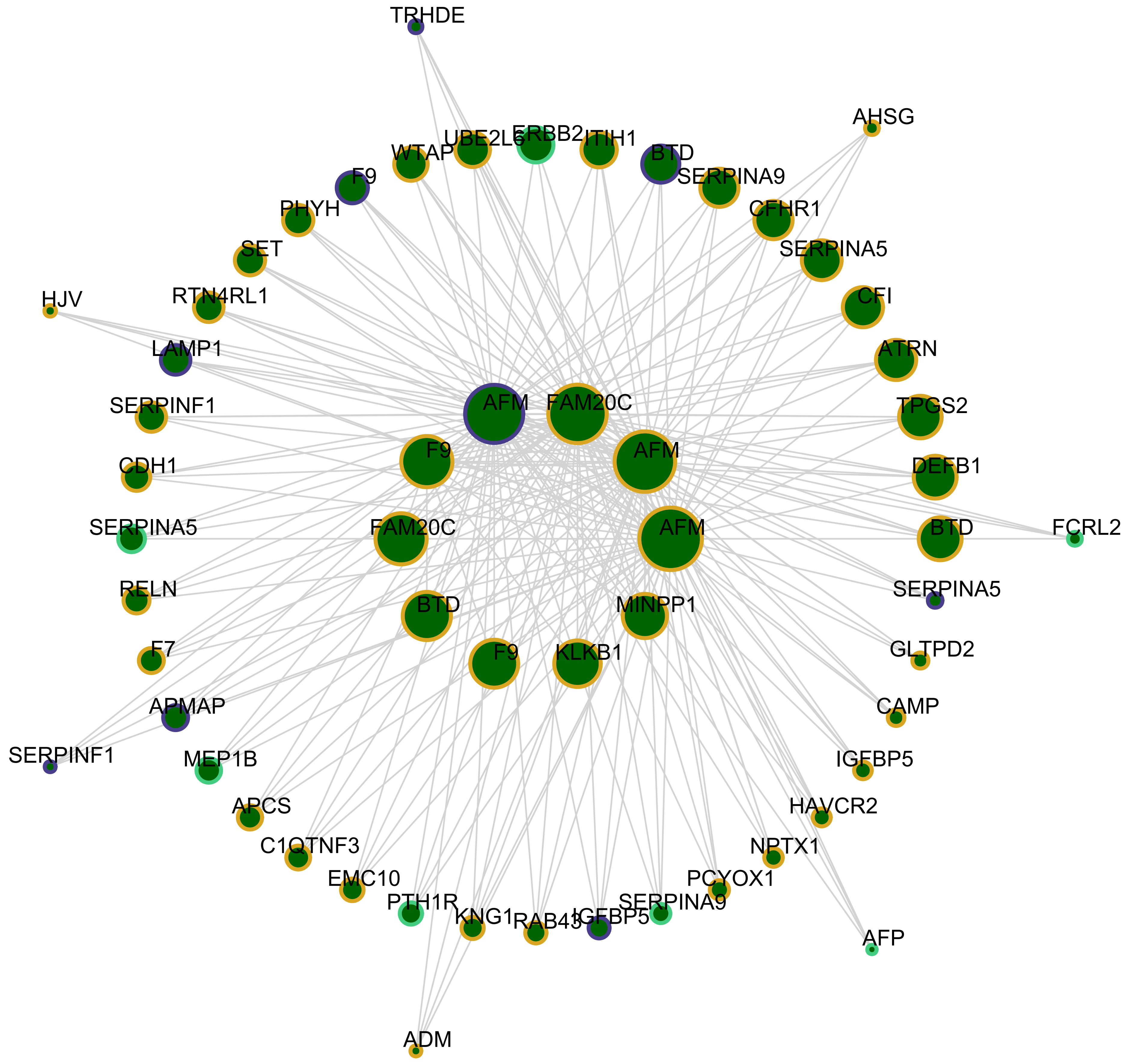

**M35 sienna3 module: Ambiguous**

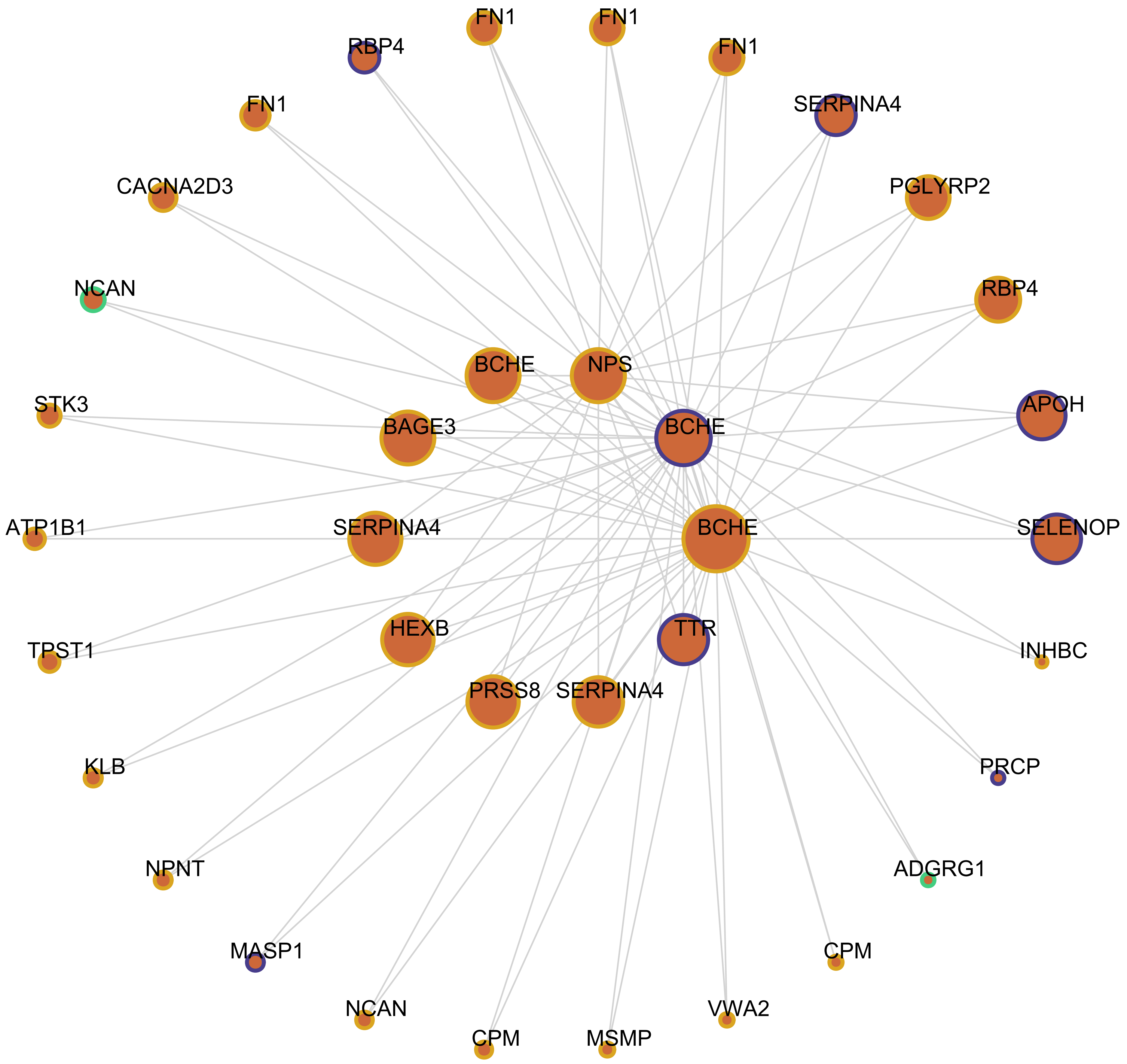

**M16 lightcyan module: Complement/Protein Activation Cascade**

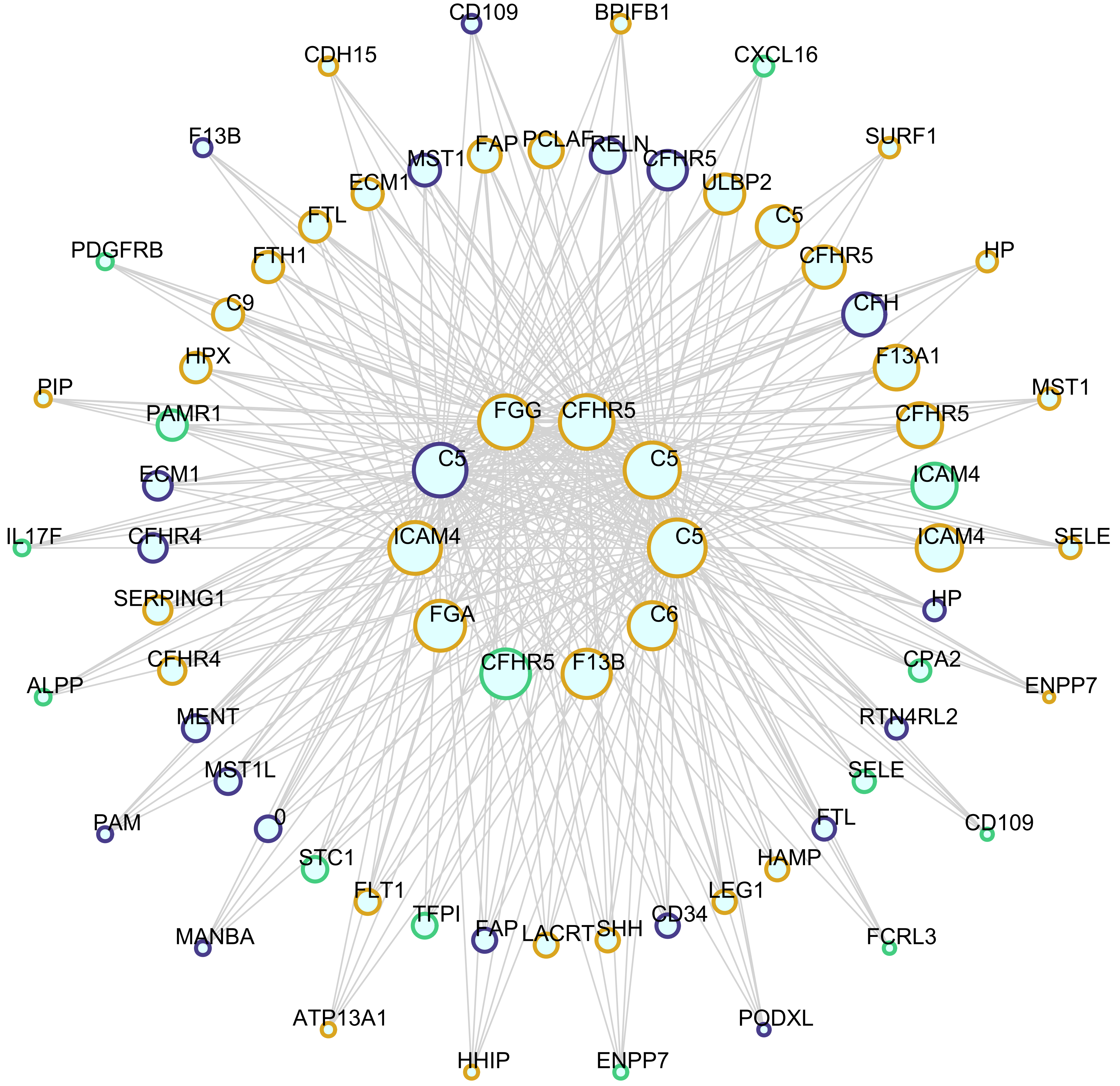

**M8 pink module: Protein Activation Cascade**

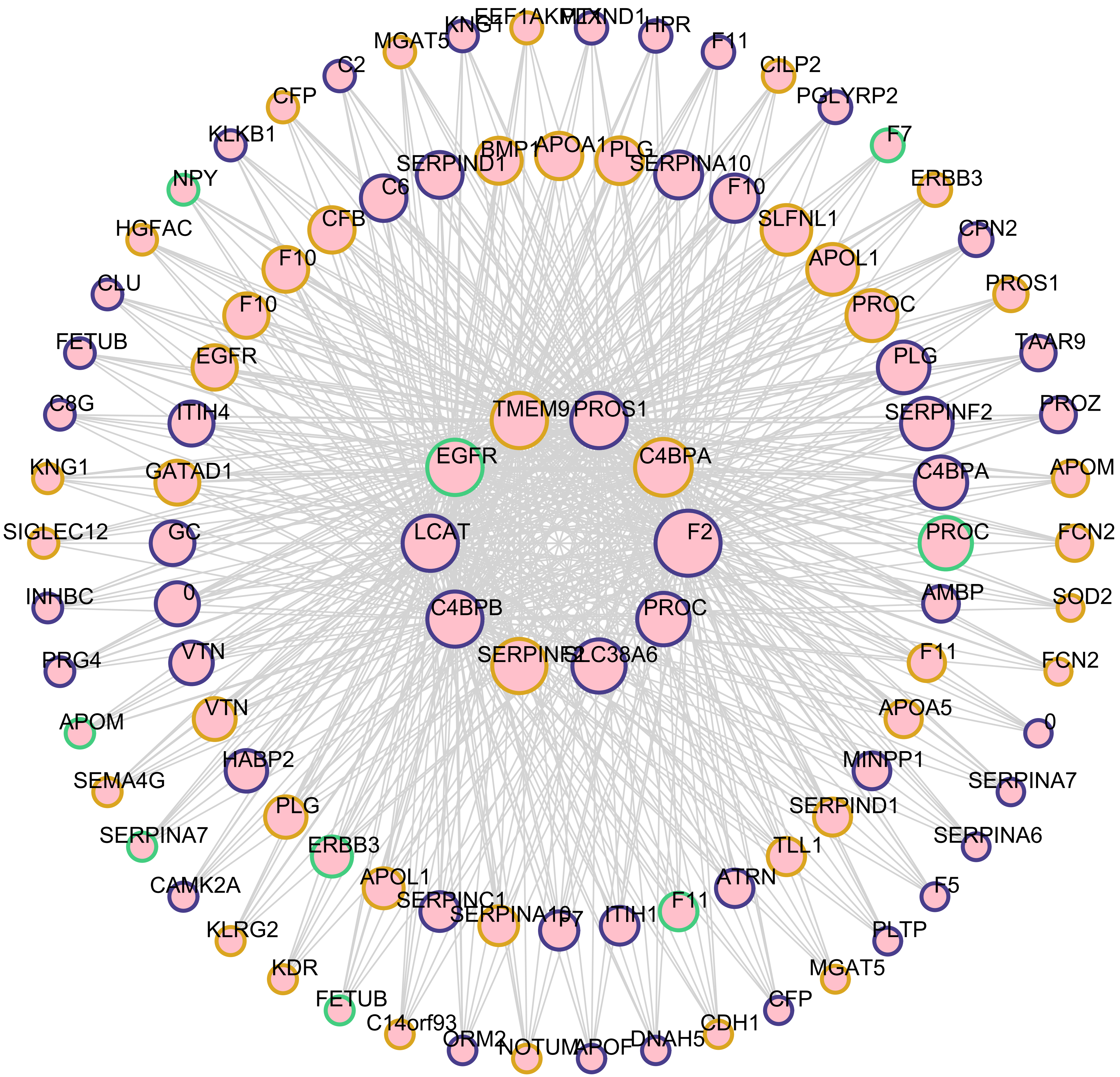

M17 grey60 module: Glycosylation

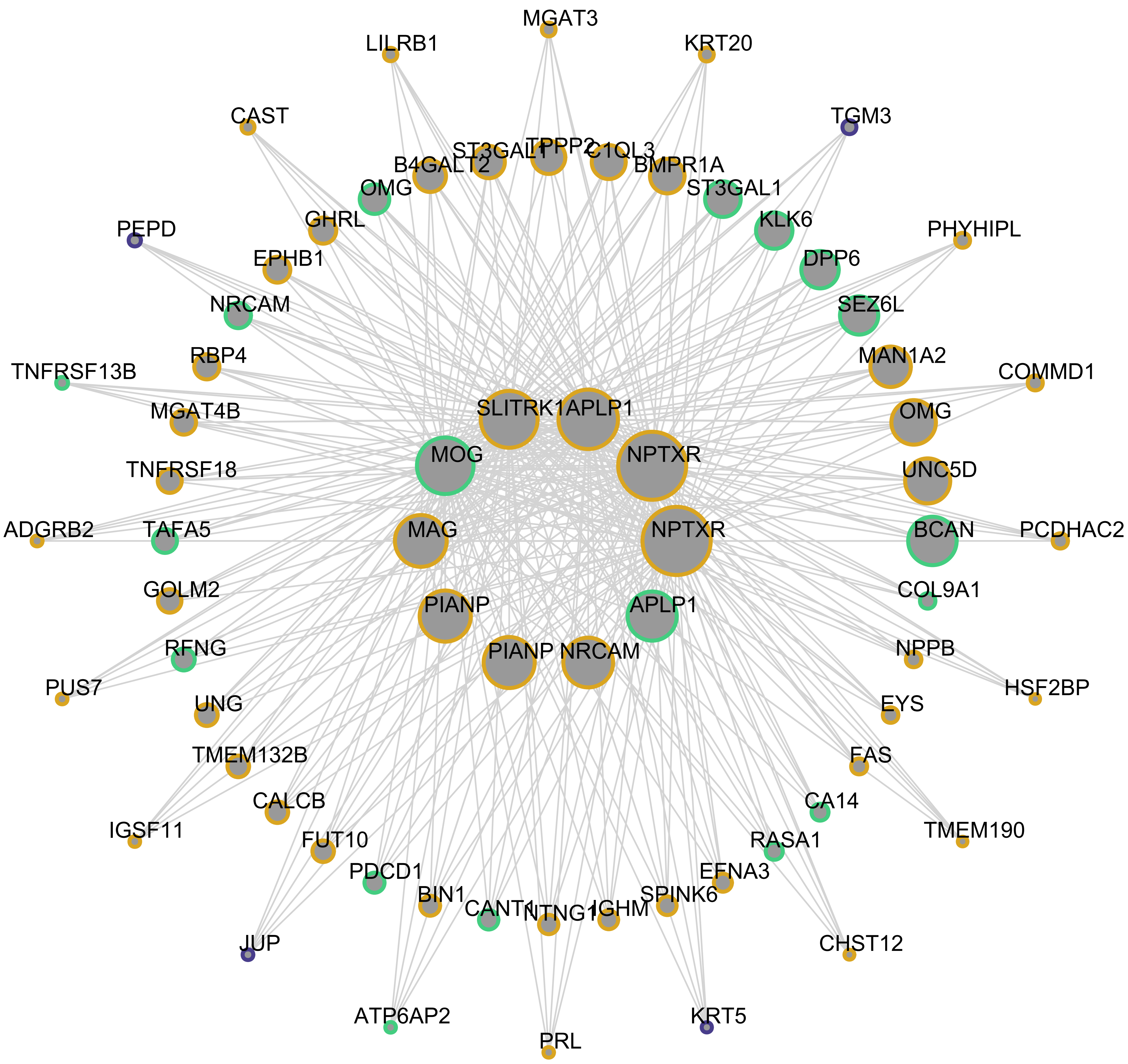

**M24 darkgrey module: Endocytosis**

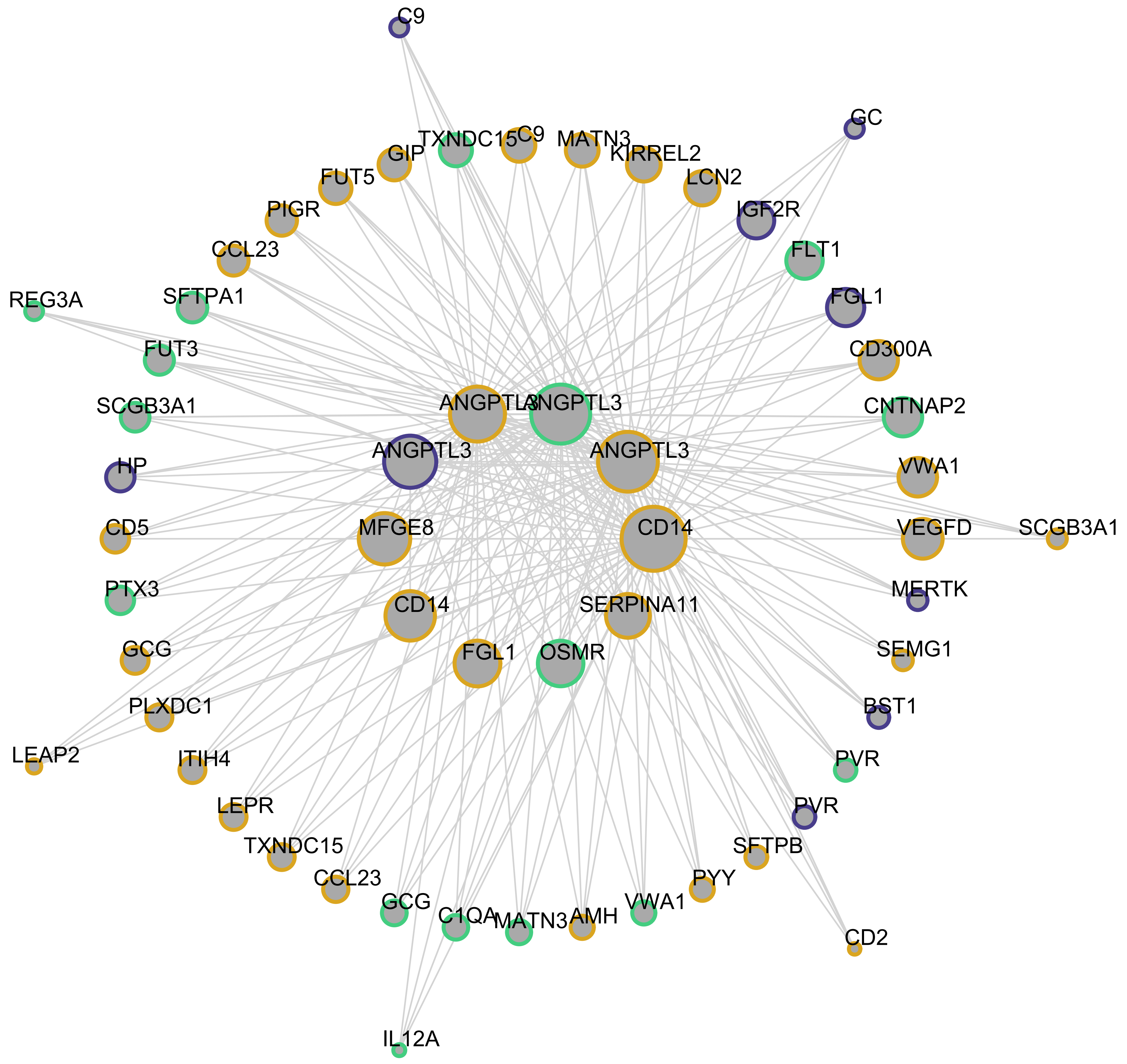

M33 darkolivegreen module: Adhesion/ECM/Wound Response

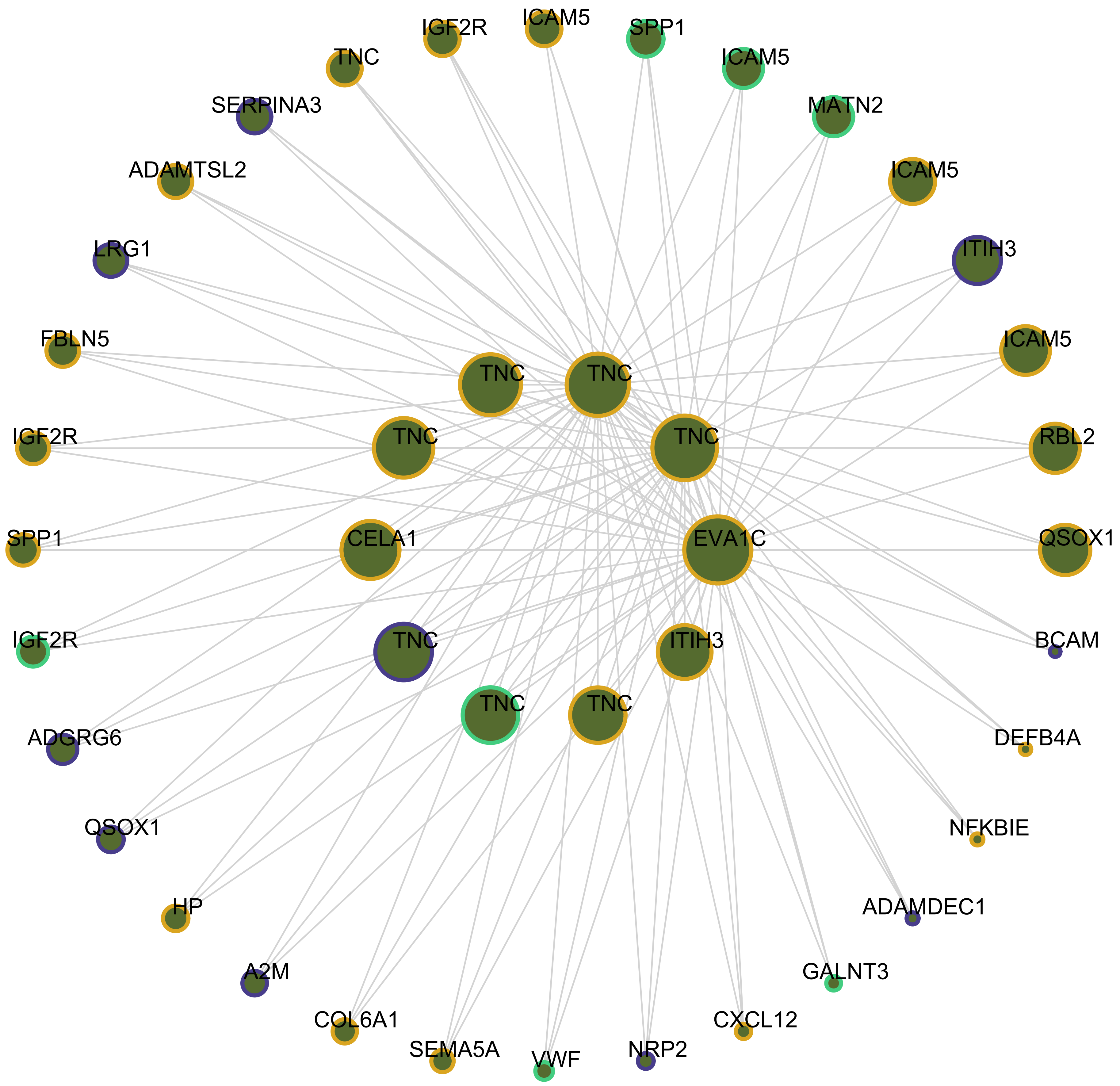

**M10 purple module: Ambiguous**

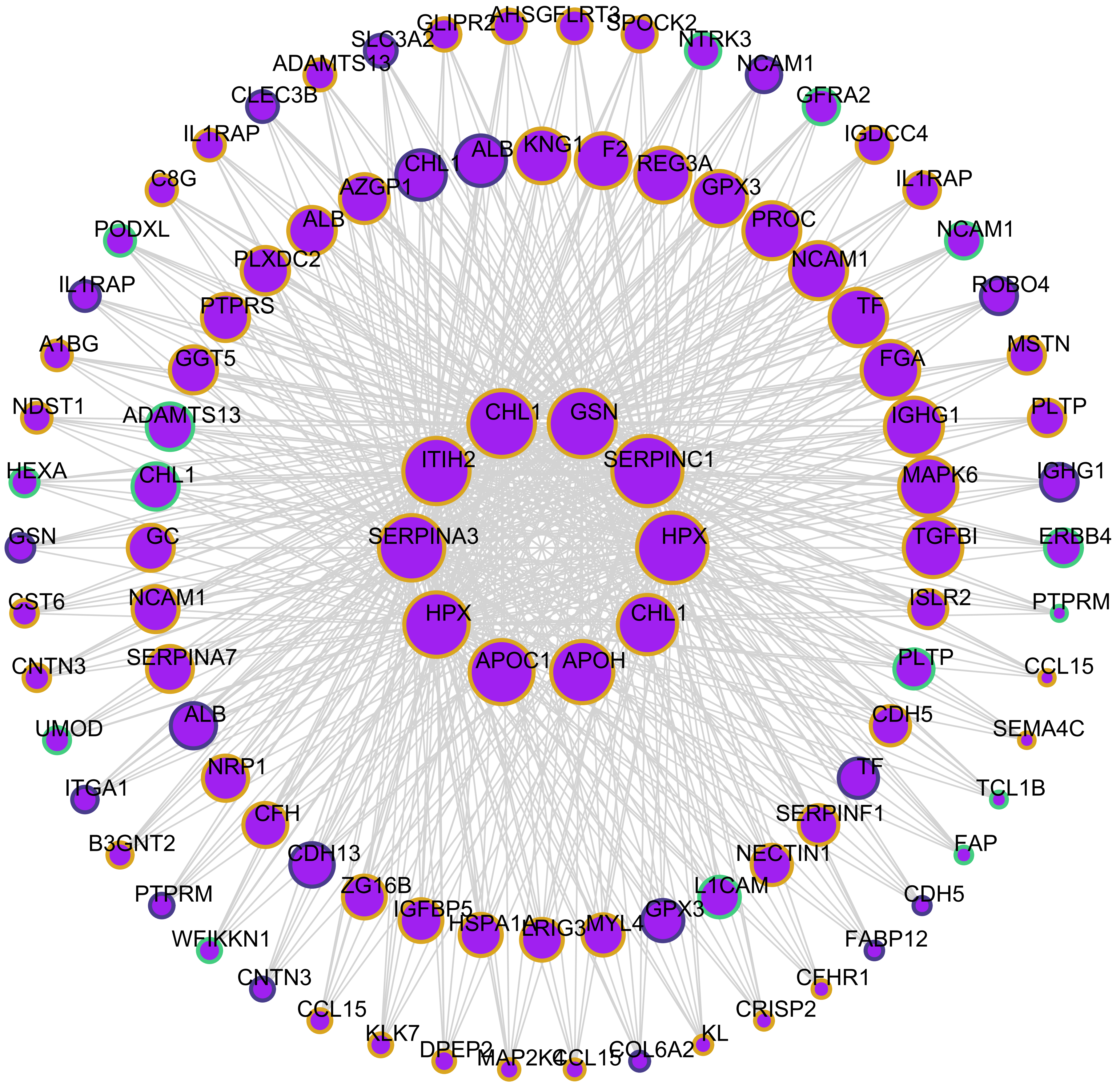

**M6 red module: Axon Guidance/Nervous System Dev**

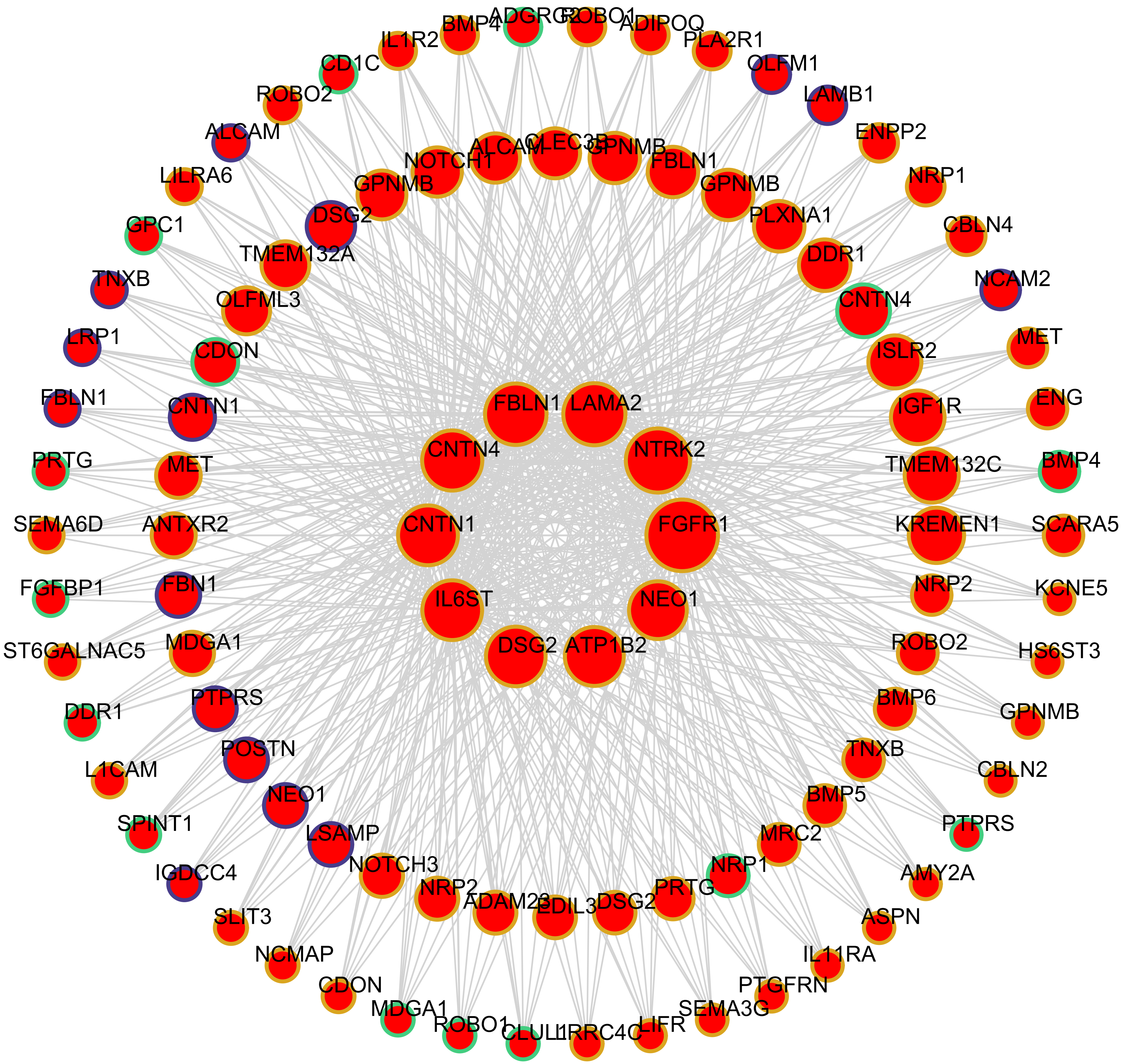

**M12 tan module: Matrisome**

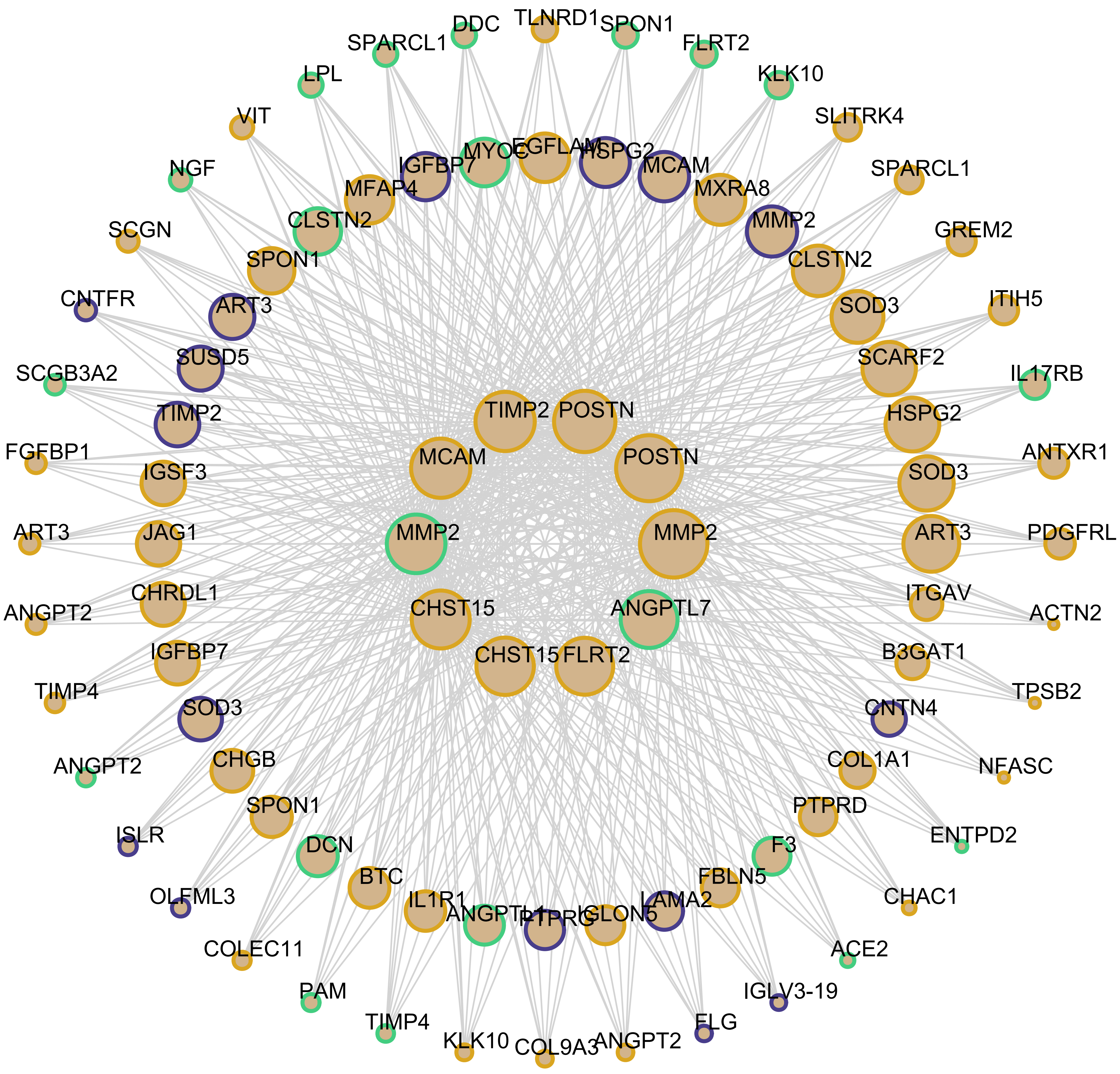

**M5 green module: ECM/IGF-PDGF Binding**

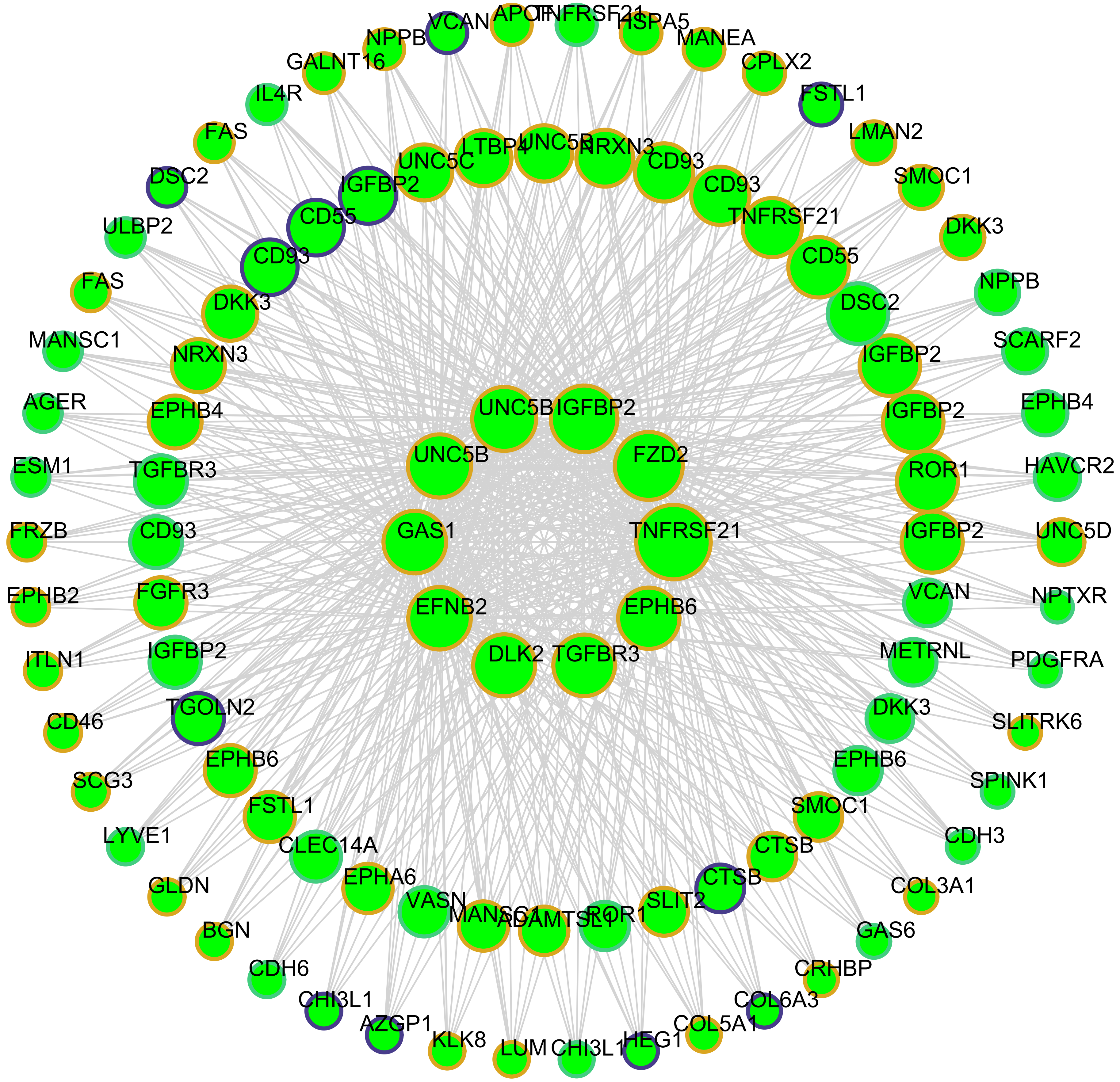

**M18 lightgreen module: Cellular Adhesion/ECM Organization**

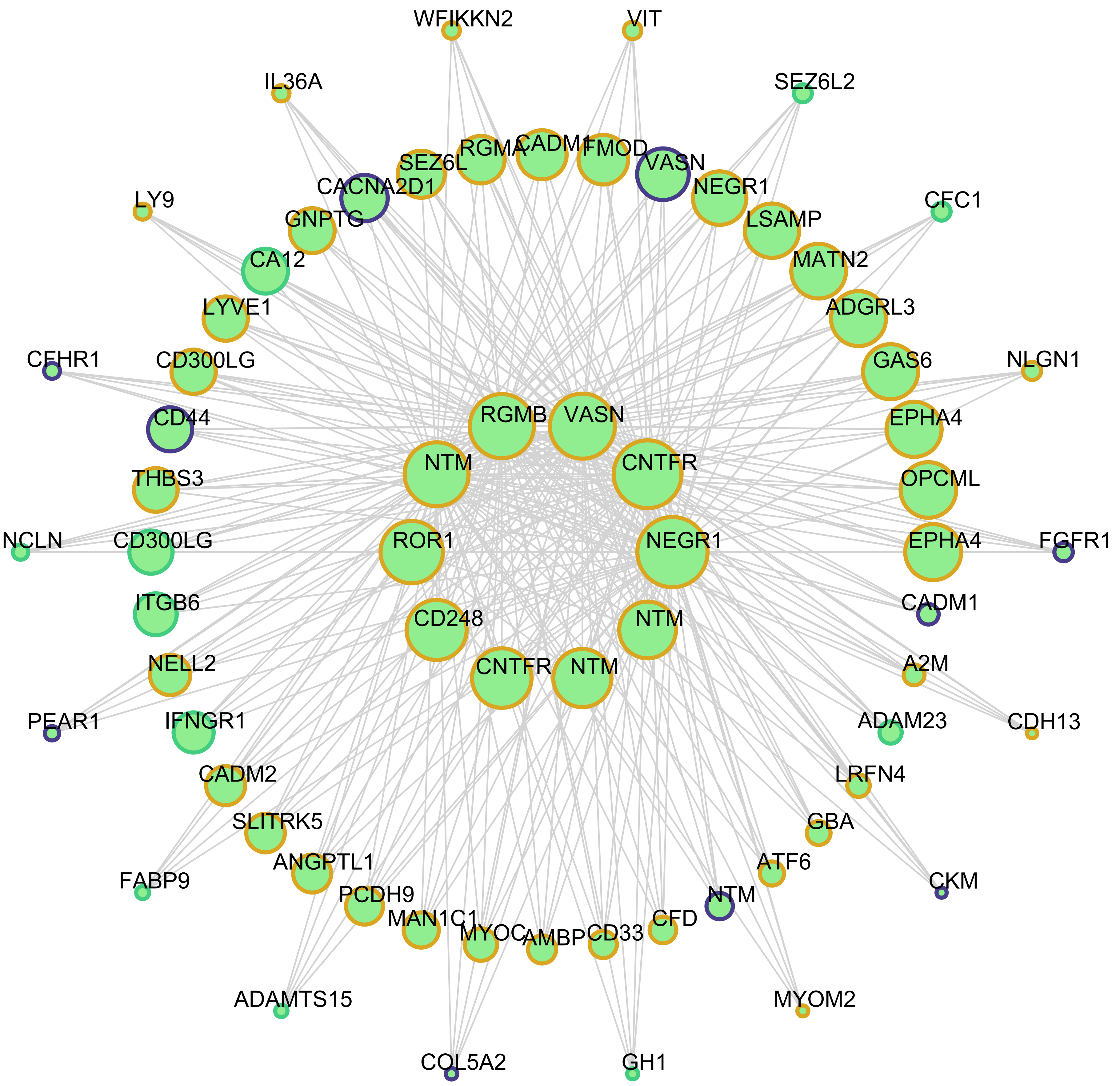

**M34 darkmagenta module: Carbohydrate Binding**

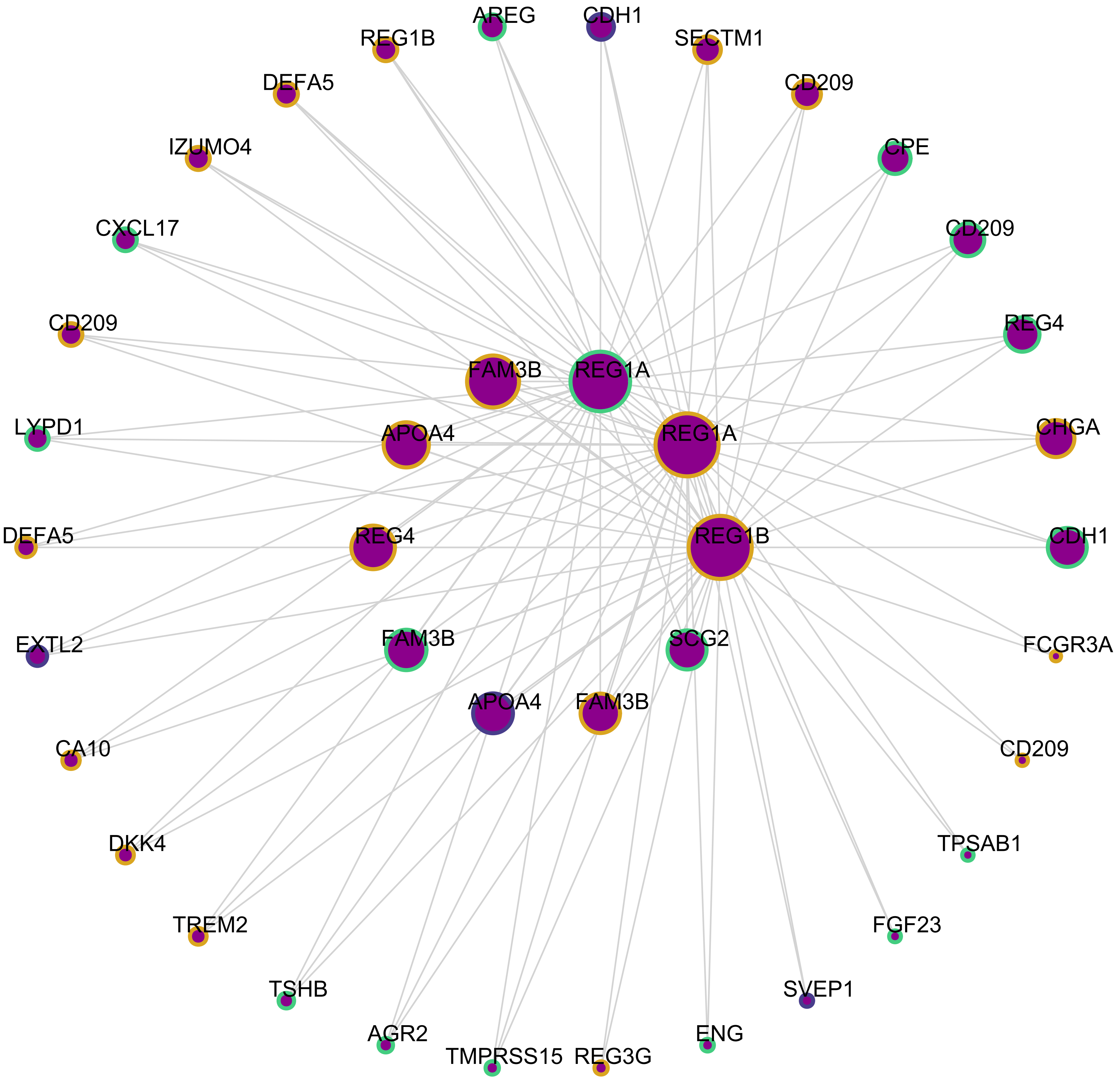

**M7 black module: TNF/Ephrin Signaling**

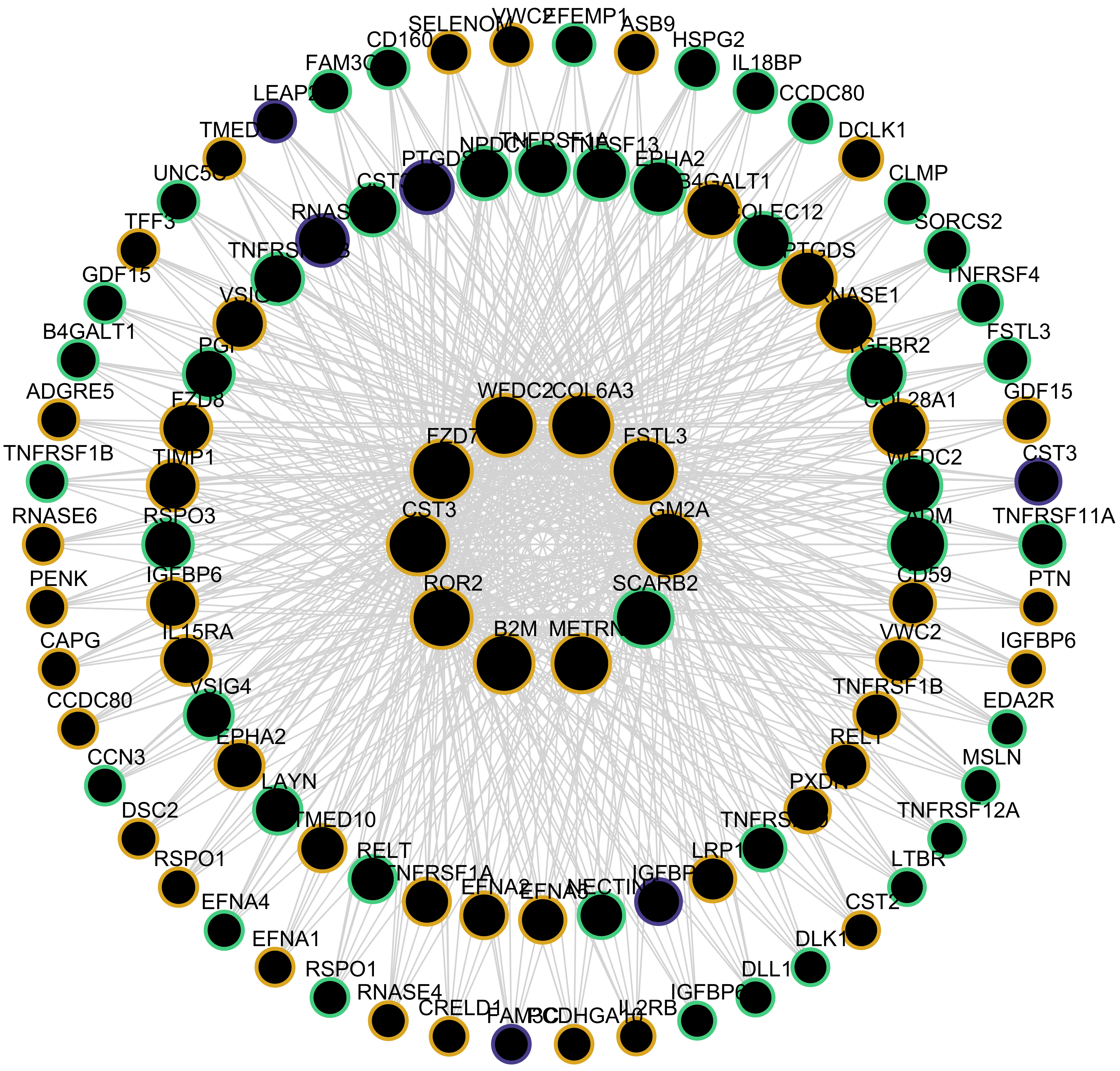

#### M23 darkturquoise module: Carbohydrate Binding/ECM

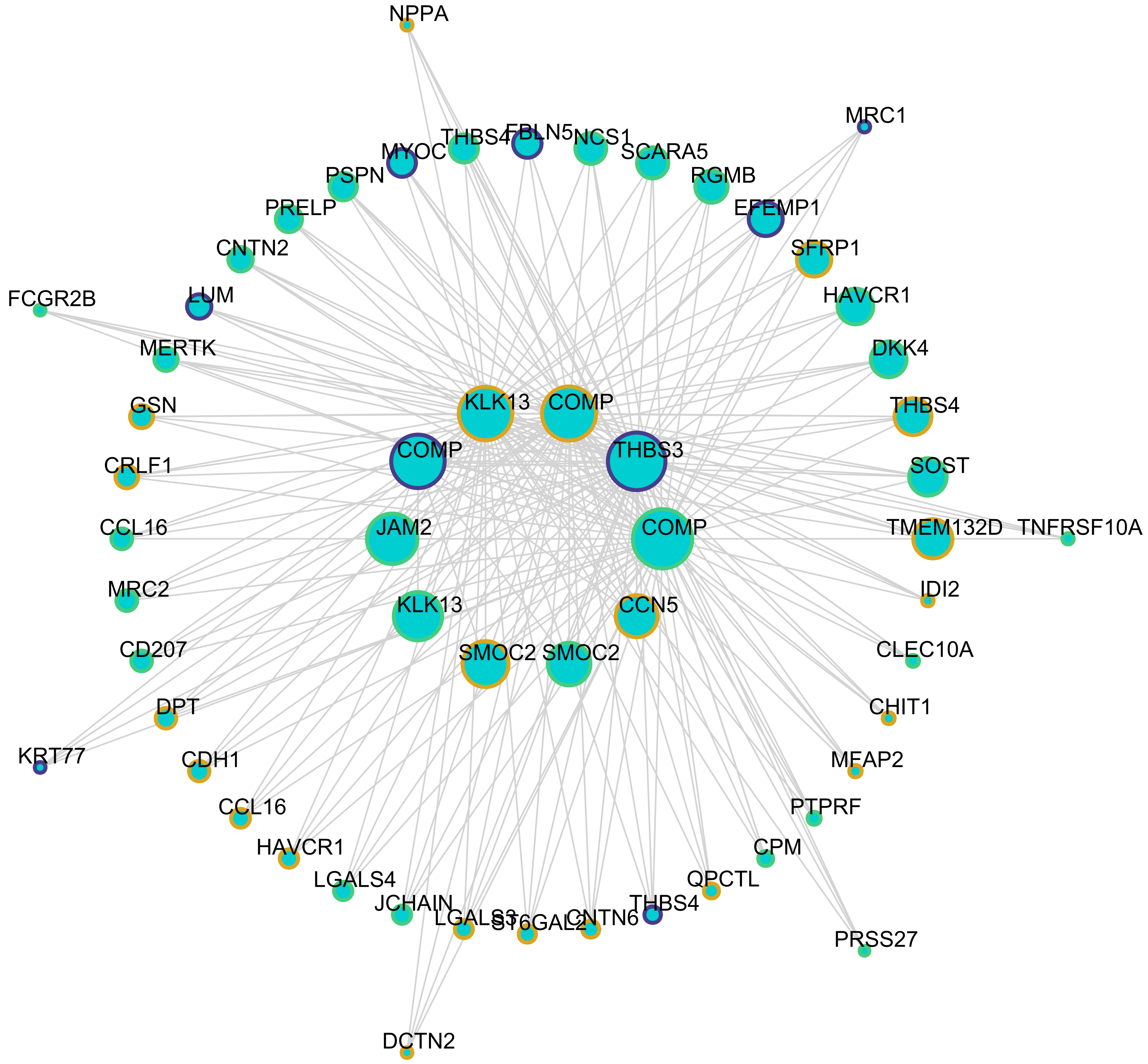

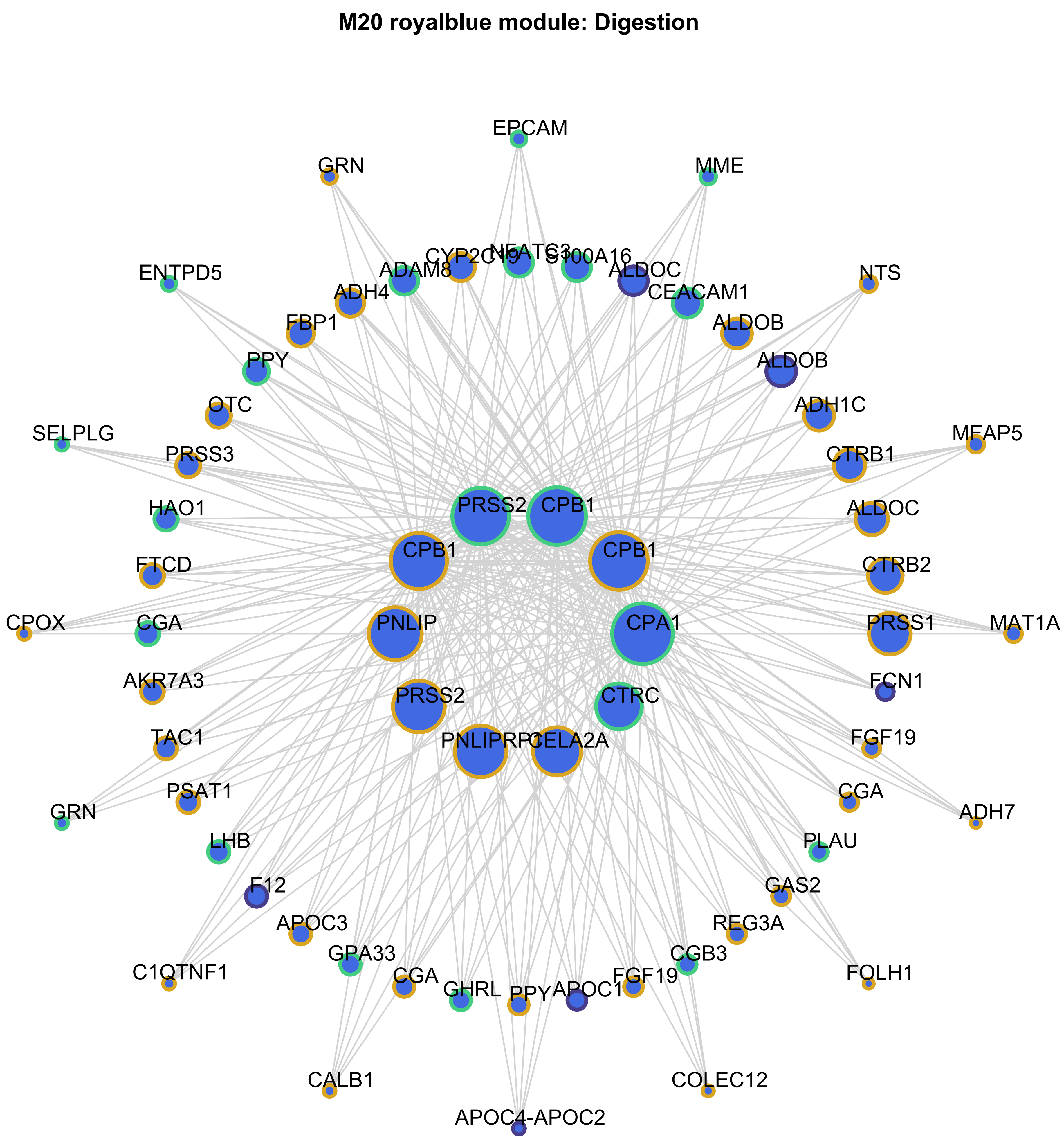
M20 royalblue module: Digestion
EPCAM
GRN
MME
ENTPD5
CYP2C19
NFATC3
S100A16
ADAM8
ALDOC
NTS
ADH4
CEACAM1
FBP1
ALDOB
PPY
ALDOB
OTC
ADH1C
SELPLG
MFAP5
PRSS3
CTRB1
PRSS2
CPB1
HAO1
ALDOC
CPB1
CPB1
FTCD
CTRB2
CPOX
CGA
PNLIP
CPA1
PRSS1
MAT1A
AKR7A3
FCN1
PRSS2
CTRC
TAC1
PNLIPRP1
CELA2A
FGF19
PSAT1
CGA
GRN
ADH7
LHB
PLAU
F12
GAS2
APOC3
REG3A
GPA33
CGB3
C1QTNF1
CGA
FOLH1
GHRL
PPY
APOC1
FGF19
CALB1
COLEC12
APOC4-APOC2

**M30 steelblue module: Ambiguous**

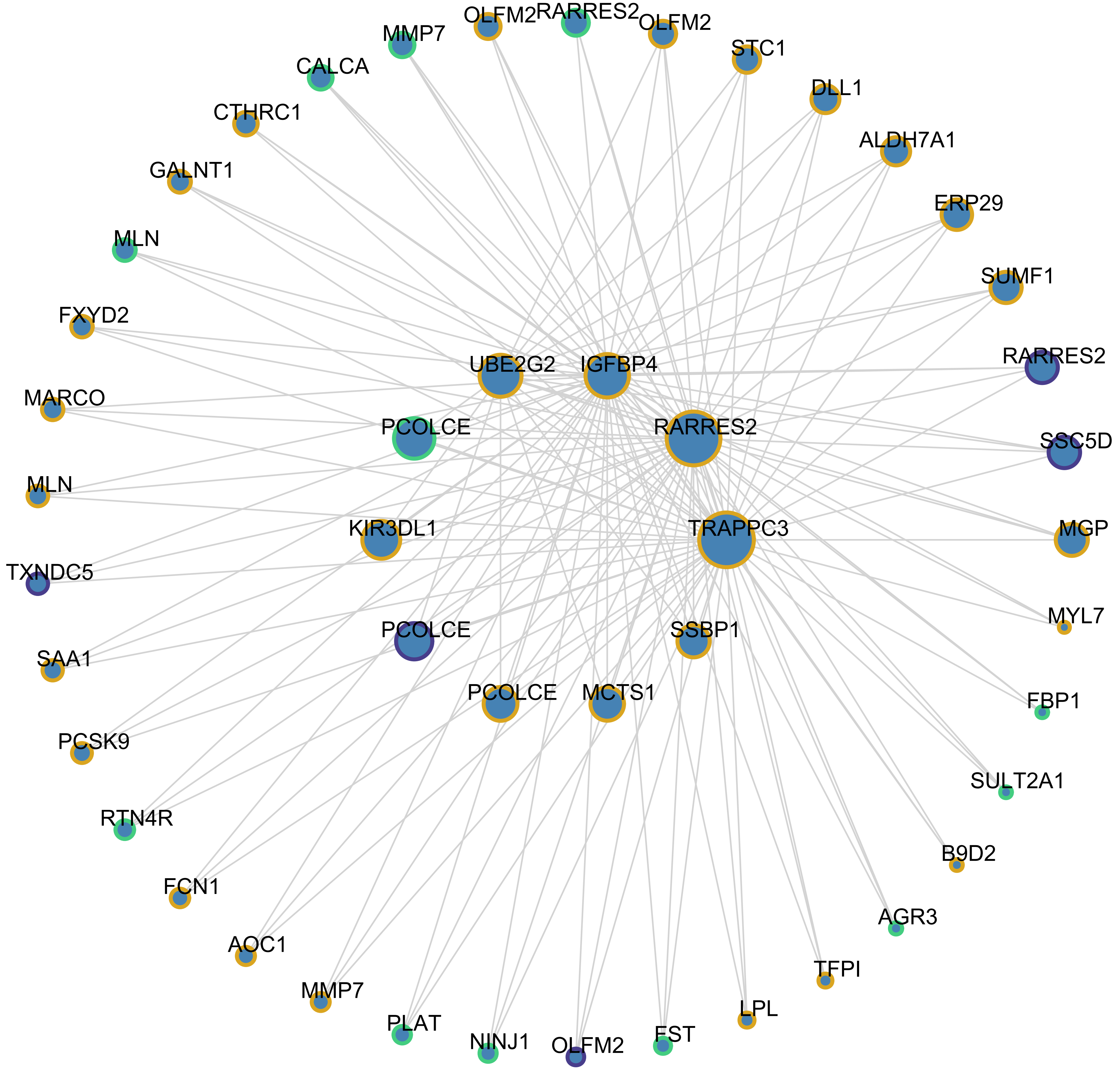

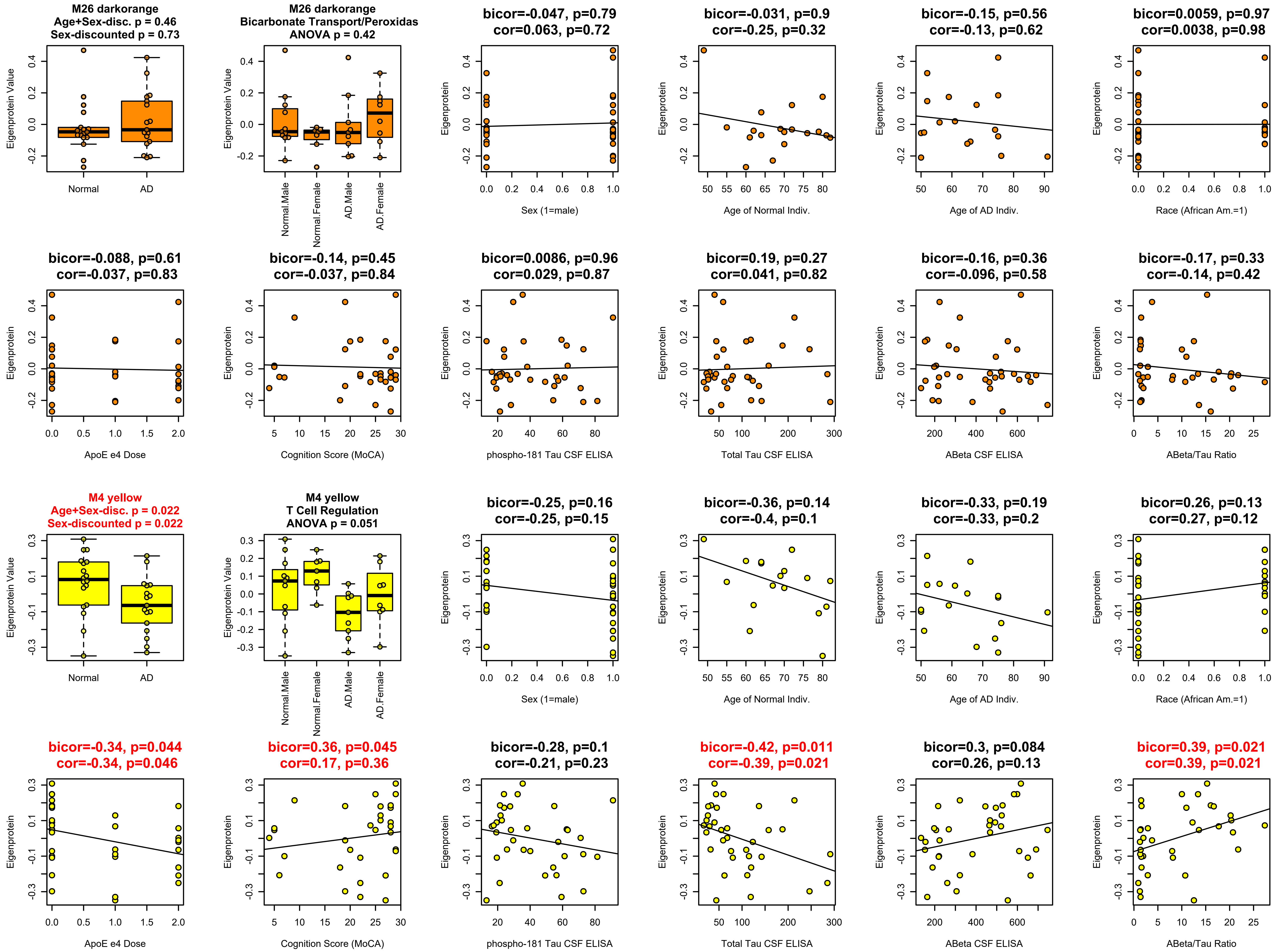
M26 darkorange
Age+Sex-disc. p = 0.46
Sex-discounted p = 0.73
Eigenprotein Value
Normal
AD
M26 darkorange
Bicarbonate Transport/Peroxidas
ANOVA p = 0.42
Eigenprotein Value
Normal.Male
Normal.Female
AD.Male
AD.Female
bicor=-0.047, p=0.79
cor=0.063, p=0.72
Eigenprotein
Sex (1=male)
bicor=-0.031, p=0.9
cor=-0.25, p=0.32
Eigenprotein
Age of Normal Indiv.
bicor=-0.15, p=0.56
cor=-0.13, p=0.62
Eigenprotein
Age of AD Indiv.
bicor=0.0059, p=0.97
cor=0.0038, p=0.98
Eigenprotein
Race (African Am.=1)
bicor=-0.088, p=0.61
cor=-0.037, p=0.83
Eigenprotein
ApoE e4 Dose
bicor=-0.14, p=0.45
cor=-0.037, p=0.84
Eigenprotein
Cognition Score (MoCA)
bicor=0.0086, p=0.96
cor=0.029, p=0.87
Eigenprotein
phospho-181 Tau CSF ELISA
bicor=0.19, p=0.27
cor=0.041, p=0.82
Eigenprotein
Total Tau CSF ELISA
bicor=-0.16, p=0.36
cor=-0.096, p=0.58
Eigenprotein
ABeta CSF ELISA
bicor=-0.17, p=0.33
cor=-0.14, p=0.42
Eigenprotein
ABeta/Tau Ratio
M4 yellow
Age+Sex-disc. p = 0.022
Sex-discounted p = 0.022
Eigenprotein Value
Normal
AD
M4 yellow
T Cell Regulation
ANOVA p = 0.051
Eigenprotein Value
Normal.Male
Normal.Female
AD.Male
AD.Female
bicor=-0.25, p=0.16
cor=-0.25, p=0.15
Eigenprotein
Sex (1=male)
bicor=-0.36, p=0.14
cor=-0.4, p=0.1
Eigenprotein
Age of Normal Indiv.
bicor=-0.33, p=0.19
cor=-0.33, p=0.2
Eigenprotein
Age of AD Indiv.
bicor=0.26, p=0.13
cor=0.27, p=0.12
Eigenprotein
Race (African Am.=1)
bicor=-0.34, p=0.044
cor=-0.34, p=0.046
Eigenprotein
ApoE e4 Dose
bicor=0.36, p=0.045
cor=0.17, p=0.36
Eigenprotein
Cognition Score (MoCA)
bicor=-0.28, p=0.1
cor=-0.21, p=0.23
Eigenprotein
phospho-181 Tau CSF ELISA
bicor=-0.42, p=0.011
cor=-0.39, p=0.021
Eigenprotein
Total Tau CSF ELISA
bicor=0.3, p=0.084
cor=0.26, p=0.13
Eigenprotein
ABeta CSF ELISA
bicor=0.39, p=0.021
cor=0.39, p=0.021
Eigenprotein
ABeta/Tau Ratio

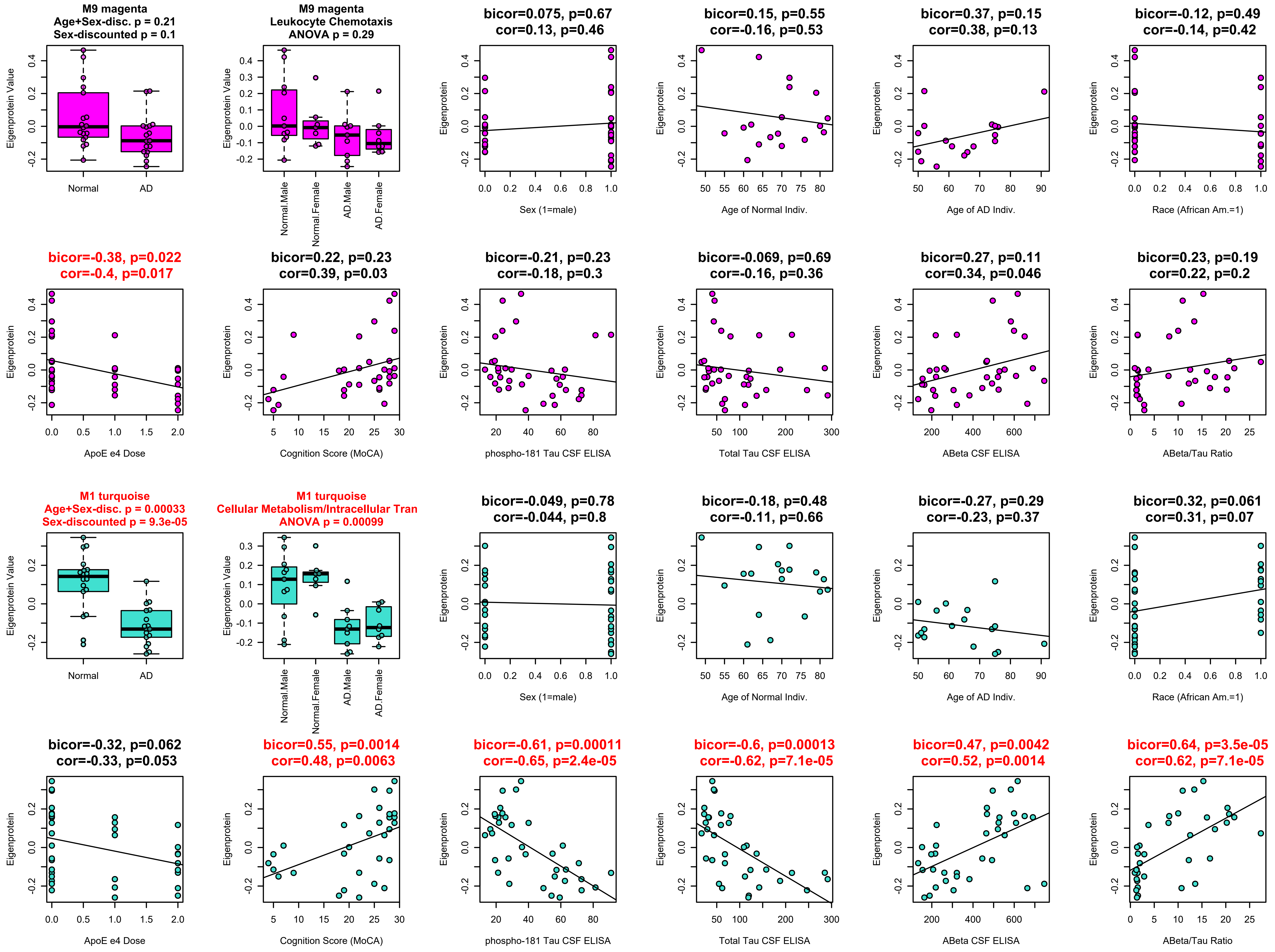

M9 magenta
Age+Sex-disc. p = 0.21
Sex-discounted p = 0.1
Eigenprotein Value
Normal
AD
M9 magenta
Leukocyte Chemotaxis
ANOVA p = 0.29
Normal.Male
Normal.Female
AD.Male
AD.Female
bicor=0.075, p=0.67
cor=0.13, p=0.46
Sex (1=male)
bicor=0.15, p=0.55
cor=-0.16, p=0.53
Age of Normal Indiv.
bicor=0.37, p=0.15
cor=0.38, p=0.13
Age of AD Indiv.
bicor=-0.12, p=0.49
cor=-0.14, p=0.42
Race (African Am.=1)
bicor=-0.38, p=0.022
cor=-0.4, p=0.017
ApoE e4 Dose
bicor=0.22, p=0.23
cor=0.39, p=0.03
Cognition Score (MoCA)
bicor=-0.21, p=0.23
cor=-0.18, p=0.3
phospho-181 Tau CSF ELISA
bicor=-0.069, p=0.69
cor=-0.16, p=0.36
Total Tau CSF ELISA
bicor=0.27, p=0.11
cor=0.34, p=0.046
ABeta CSF ELISA
bicor=0.23, p=0.19
cor=0.22, p=0.2
ABeta/Tau Ratio
M1 turquoise
Age+Sex-disc. p = 0.00033
Sex-discounted p = 9.3e-05
M1 turquoise
Cellular Metabolism/Intracellular Tran
ANOVA p = 0.00099
bicor=-0.049, p=0.78
cor=-0.044, p=0.8
bicor=-0.18, p=0.48
cor=-0.11, p=0.66
bicor=-0.27, p=0.29
cor=-0.23, p=0.37
bicor=0.32, p=0.061
cor=0.31, p=0.07
bicor=-0.32, p=0.062
cor=-0.33, p=0.053
bicor=0.55, p=0.0014
cor=0.48, p=0.0063
bicor=-0.61, p=0.00011
cor=-0.65, p=2.4e-05
bicor=-0.6, p=0.00013
cor=-0.62, p=7.1e-05
bicor=0.47, p=0.0042
cor=0.52, p=0.0014
bicor=0.64, p=3.5e-05
cor=0.62, p=7.1e-05
Eigenprotein

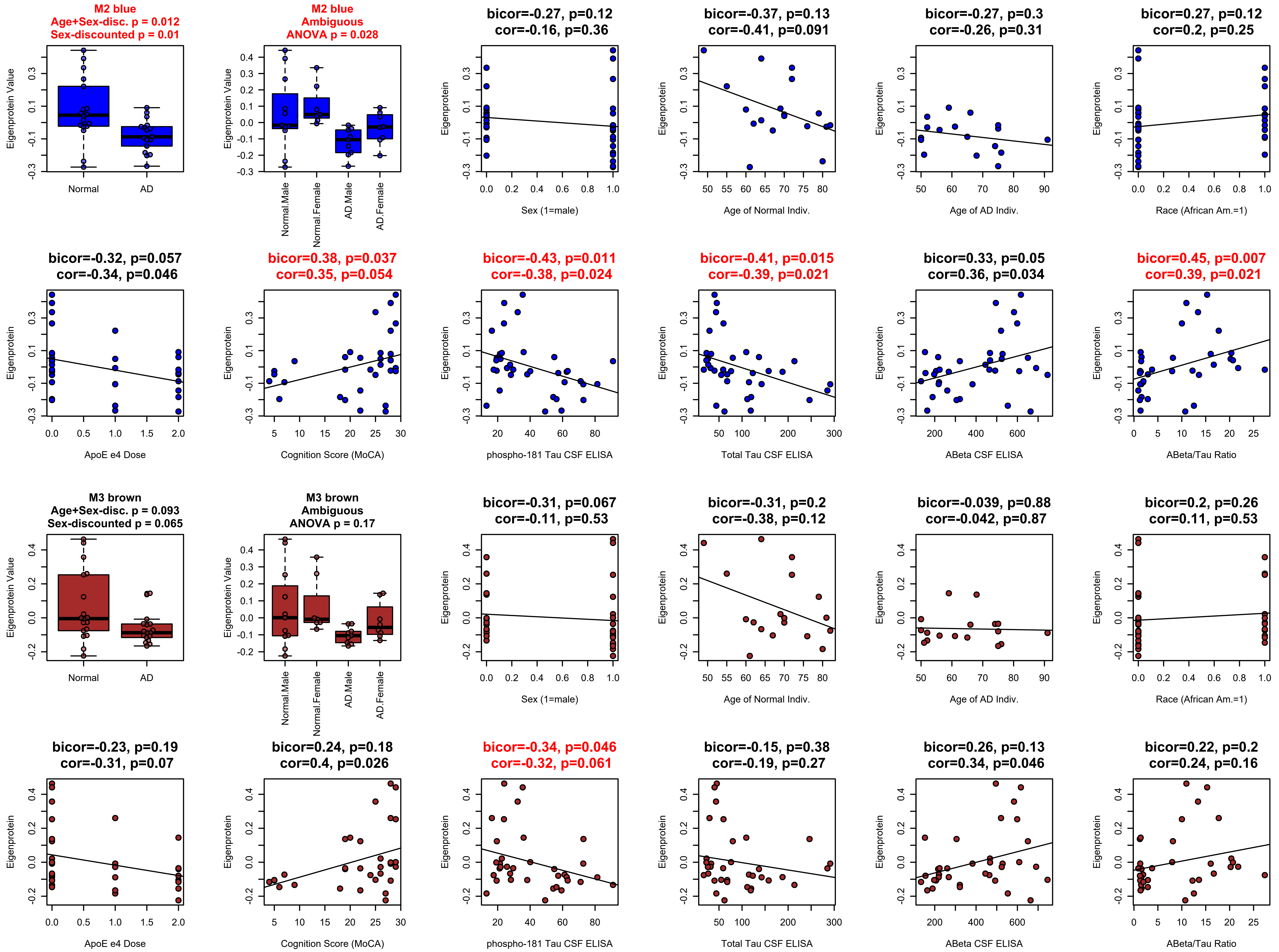
M2 blue
Age+Sex-disc. p = 0.012
Sex-discounted p = 0.01
Eigenprotein Value
Normal
AD
M2 blue
Ambiguous
ANOVA p = 0.028
Eigenprotein Value
Normal.Male
Normal.Female
AD.Male
AD.Female
bicor=-0.27, p=0.12
cor=-0.16, p=0.36
Eigenprotein
Sex (1=male)
bicor=-0.37, p=0.13
cor=-0.41, p=0.091
Eigenprotein
Age of Normal Indiv.
bicor=-0.27, p=0.3
cor=-0.26, p=0.31
Eigenprotein
Age of AD Indiv.
bicor=0.27, p=0.12
cor=0.2, p=0.25
Eigenprotein
Race (African Am.=1)
bicor=-0.32, p=0.057
cor=-0.34, p=0.046
Eigenprotein
ApoE e4 Dose
bicor=0.38, p=0.037
cor=0.35, p=0.054
Eigenprotein
Cognition Score (MoCA)
bicor=-0.43, p=0.011
cor=-0.38, p=0.024
Eigenprotein
phospho-181 Tau CSF ELISA
bicor=-0.41, p=0.015
cor=-0.39, p=0.021
Eigenprotein
Total Tau CSF ELISA
bicor=0.33, p=0.05
cor=0.36, p=0.034
Eigenprotein
ABeta CSF ELISA
bicor=0.45, p=0.007
cor=0.39, p=0.021
Eigenprotein
ABeta/Tau Ratio
M3 brown
Age+Sex-disc. p = 0.093
Sex-discounted p = 0.065
Eigenprotein Value
Normal
AD
M3 brown
Ambiguous
ANOVA p = 0.17
Eigenprotein Value
Normal.Male
Normal.Female
AD.Male
AD.Female
bicor=-0.31, p=0.067
cor=-0.11, p=0.53
Eigenprotein
Sex (1=male)
bicor=-0.31, p=0.2
cor=-0.38, p=0.12
Eigenprotein
Age of Normal Indiv.
bicor=-0.039, p=0.88
cor=-0.042, p=0.87
Eigenprotein
Age of AD Indiv.
bicor=0.2, p=0.26
cor=0.11, p=0.53
Eigenprotein
Race (African Am.=1)
bicor=-0.23, p=0.19
cor=-0.31, p=0.07
Eigenprotein
ApoE e4 Dose
bicor=0.24, p=0.18
cor=0.4, p=0.026
Eigenprotein
Cognition Score (MoCA)
bicor=-0.34, p=0.046
cor=-0.32, p=0.061
Eigenprotein
phospho-181 Tau CSF ELISA
bicor=-0.15, p=0.38
cor=-0.19, p=0.27
Eigenprotein
Total Tau CSF ELISA
bicor=0.26, p=0.13
cor=0.34, p=0.046
Eigenprotein
ABeta CSF ELISA
bicor=0.22, p=0.2
cor=0.24, p=0.16
Eigenprotein
ABeta/Tau Ratio

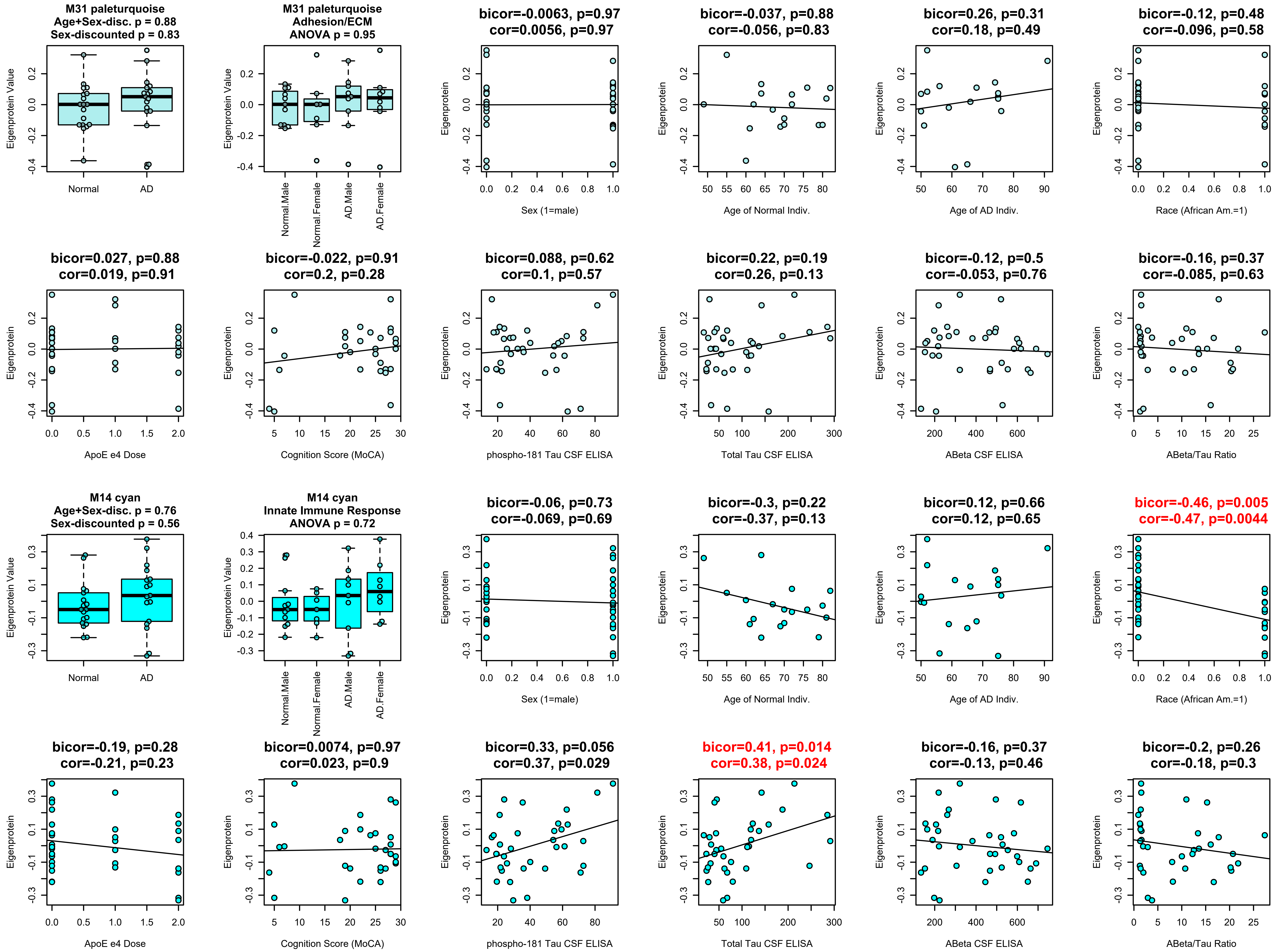
M31 paleturquoise
Age+Sex-disc. p = 0.88
Sex-discounted p = 0.83
Eigenprotein Value
Normal
AD
M31 paleturquoise
Adhesion/ECM
ANOVA p = 0.95
Normal.Male
Normal.Female
AD.Male
AD.Female
bicor=-0.0063, p=0.97
cor=0.0056, p=0.97
Eigenprotein
Sex (1=male)
bicor=-0.037, p=0.88
cor=-0.056, p=0.83
Age of Normal Indiv.
bicor=0.26, p=0.31
cor=0.18, p=0.49
Age of AD Indiv.
bicor=-0.12, p=0.48
cor=-0.096, p=0.58
Race (African Am.=1)
bicor=0.027, p=0.88
cor=0.019, p=0.91
ApoE e4 Dose
bicor=-0.022, p=0.91
cor=0.2, p=0.28
Cognition Score (MoCA)
bicor=0.088, p=0.62
cor=0.1, p=0.57
phospho-181 Tau CSF ELISA
bicor=0.22, p=0.19
cor=0.26, p=0.13
Total Tau CSF ELISA
bicor=-0.12, p=0.5
cor=-0.053, p=0.76
ABeta CSF ELISA
bicor=-0.16, p=0.37
cor=-0.085, p=0.63
ABeta/Tau Ratio
M14 cyan
Age+Sex-disc. p = 0.76
Sex-discounted p = 0.56
M14 cyan
Innate Immune Response
ANOVA p = 0.72
bicor=-0.06, p=0.73
cor=-0.069, p=0.69
bicor=-0.3, p=0.22
cor=-0.37, p=0.13
bicor=0.12, p=0.66
cor=0.12, p=0.65
bicor=-0.46, p=0.005
cor=-0.47, p=0.0044
bicor=-0.19, p=0.28
cor=-0.21, p=0.23
bicor=0.0074, p=0.97
cor=0.023, p=0.9
bicor=0.33, p=0.056
cor=0.37, p=0.029
bicor=0.41, p=0.014
cor=0.38, p=0.024
bicor=-0.16, p=0.37
cor=-0.13, p=0.46
bicor=-0.2, p=0.26
cor=-0.18, p=0.3

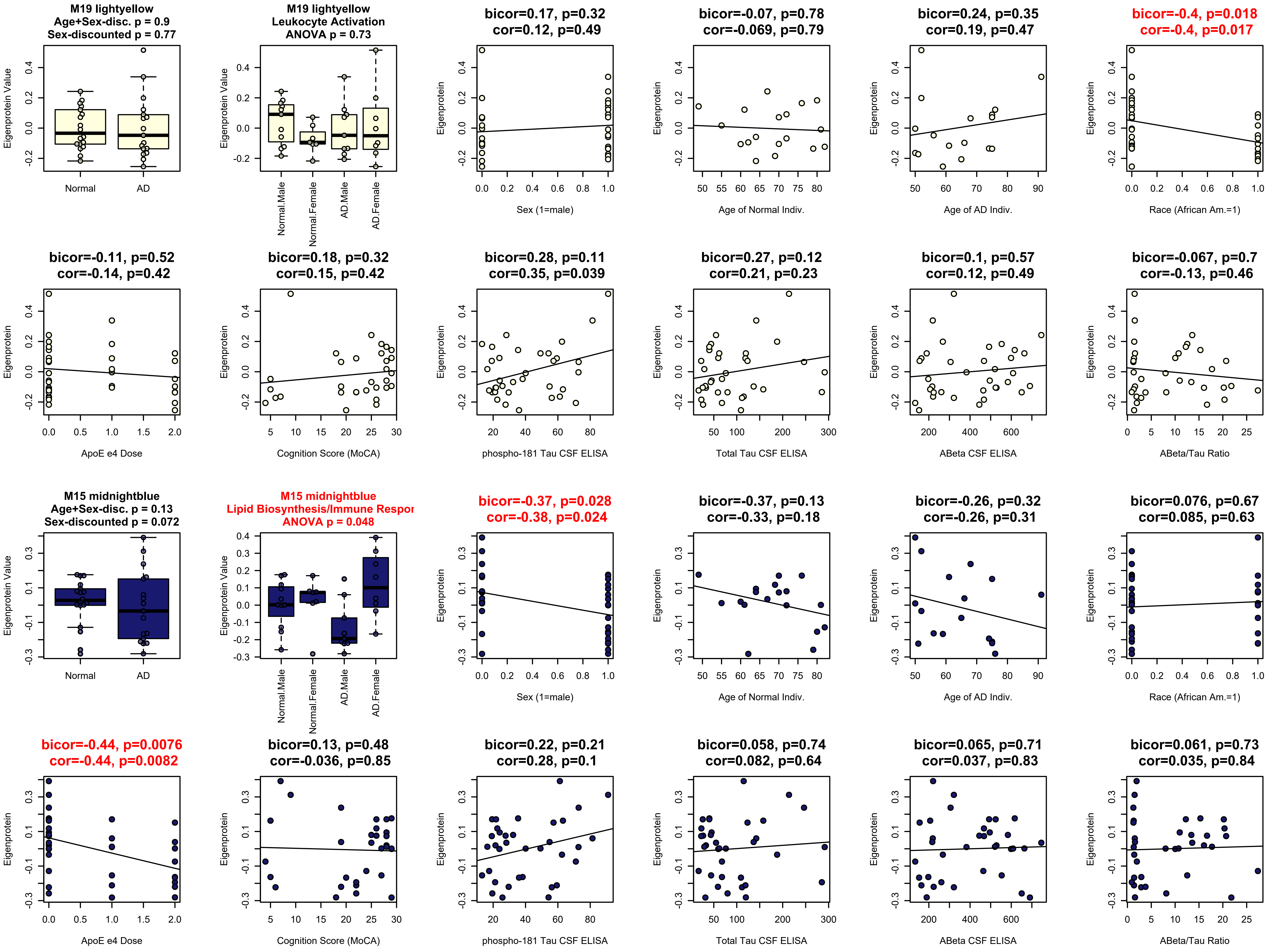

M19 lightyellow
Age+Sex-disc. p = 0.9
Sex-discounted p = 0.77
Eigenprotein Value
Normal
AD
M19 lightyellow
Leukocyte Activation
ANOVA p = 0.73
Eigenprotein Value
Normal.Male
Normal.Female
AD.Male
AD.Female
bicor=0.17, p=0.32
cor=0.12, p=0.49
Eigenprotein
Sex (1=male)
bicor=-0.07, p=0.78
cor=-0.069, p=0.79
Eigenprotein
Age of Normal Indiv.
bicor=0.24, p=0.35
cor=0.19, p=0.47
Eigenprotein
Age of AD Indiv.
bicor=-0.4, p=0.018
cor=-0.4, p=0.017
Eigenprotein
Race (African Am.=1)
bicor=-0.11, p=0.52
cor=-0.14, p=0.42
Eigenprotein
ApoE e4 Dose
bicor=0.18, p=0.32
cor=0.15, p=0.42
Eigenprotein
Cognition Score (MoCA)
bicor=0.28, p=0.11
cor=0.35, p=0.039
Eigenprotein
phospho-181 Tau CSF ELISA
bicor=0.27, p=0.12
cor=0.21, p=0.23
Eigenprotein
Total Tau CSF ELISA
bicor=0.1, p=0.57
cor=0.12, p=0.49
Eigenprotein
ABeta CSF ELISA
bicor=-0.067, p=0.7
cor=-0.13, p=0.46
Eigenprotein
ABeta/Tau Ratio
M15 midnightblue
Age+Sex-disc. p = 0.13
Sex-discounted p = 0.072
Eigenprotein Value
Normal
AD
M15 midnightblue
Lipid Biosynthesis/Immune Respor
ANOVA p = 0.048
Eigenprotein Value
Normal.Male
Normal.Female
AD.Male
AD.Female
bicor=-0.37, p=0.028
cor=-0.38, p=0.024
Eigenprotein
Sex (1=male)
bicor=-0.37, p=0.13
cor=-0.33, p=0.18
Eigenprotein
Age of Normal Indiv.
bicor=-0.26, p=0.32
cor=-0.26, p=0.31
Eigenprotein
Age of AD Indiv.
bicor=0.076, p=0.67
cor=0.085, p=0.63
Eigenprotein
Race (African Am.=1)
bicor=-0.44, p=0.0076
cor=-0.44, p=0.0082
Eigenprotein
ApoE e4 Dose
bicor=0.13, p=0.48
cor=-0.036, p=0.85
Eigenprotein
Cognition Score (MoCA)
bicor=0.22, p=0.21
cor=0.28, p=0.1
Eigenprotein
phospho-181 Tau CSF ELISA
bicor=0.058, p=0.74
cor=0.082, p=0.64
Eigenprotein
Total Tau CSF ELISA
bicor=0.065, p=0.71
cor=0.037, p=0.83
Eigenprotein
ABeta CSF ELISA
bicor=0.061, p=0.73
cor=0.035, p=0.84
Eigenprotein
ABeta/Tau Ratio

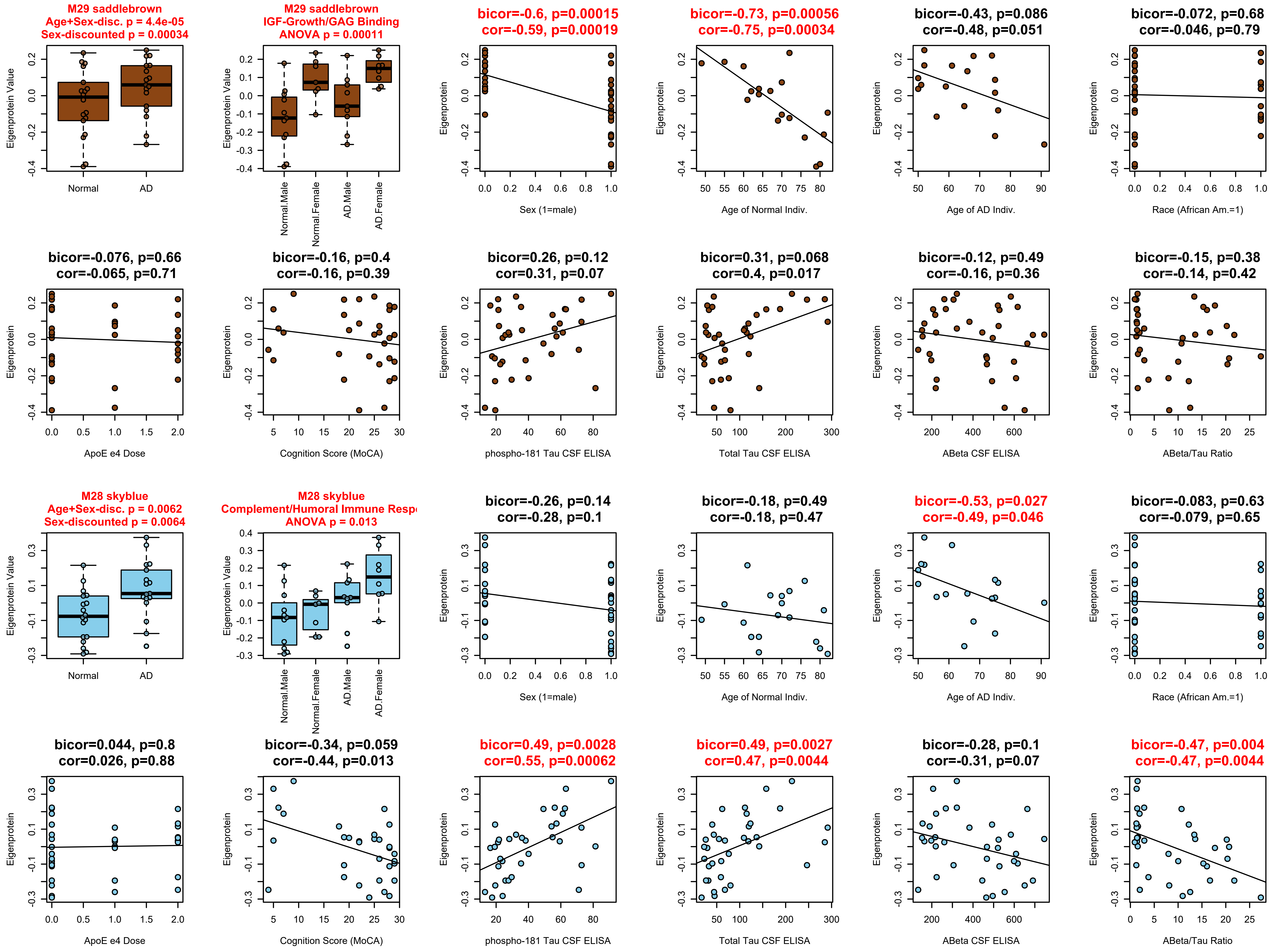
M29 saddlebrown
Age+Sex-disc. p = 4.4e-05
Sex-discounted p = 0.00034
M29 saddlebrown
IGF-Growth/GAG Binding
ANOVA p = 0.00011
bicor=-0.6, p=0.00015
cor=-0.59, p=0.00019
bicor=-0.73, p=0.00056
cor=-0.75, p=0.00034
bicor=-0.43, p=0.086
cor=-0.48, p=0.051
bicor=-0.072, p=0.68
cor=-0.046, p=0.79
bicor=-0.076, p=0.66
cor=-0.065, p=0.71
bicor=-0.16, p=0.4
cor=-0.16, p=0.39
bicor=0.26, p=0.12
cor=0.31, p=0.07
bicor=0.31, p=0.068
cor=0.4, p=0.017
bicor=-0.12, p=0.49
cor=-0.16, p=0.36
bicor=-0.15, p=0.38
cor=-0.14, p=0.42
M28 skyblue
Age+Sex-disc. p = 0.0062
Sex-discounted p = 0.0064
M28 skyblue
Complement/Humoral Immune Resp
ANOVA p = 0.013
bicor=-0.26, p=0.14
cor=-0.28, p=0.1
bicor=-0.18, p=0.49
cor=-0.18, p=0.47
bicor=-0.53, p=0.027
cor=-0.49, p=0.046
bicor=-0.083, p=0.63
cor=-0.079, p=0.65
bicor=0.044, p=0.8
cor=0.026, p=0.88
bicor=-0.34, p=0.059
cor=-0.44, p=0.013
bicor=0.49, p=0.0028
cor=0.55, p=0.00062
bicor=0.49, p=0.0027
cor=0.47, p=0.0044
bicor=-0.28, p=0.1
cor=-0.31, p=0.07
bicor=-0.47, p=0.004
cor=-0.47, p=0.0044
Eigenprotein Value
Eigenprotein
Normal
AD
Normal.Male
Normal.Female
AD.Male
AD.Female
Sex (1=male)
Age of Normal Indiv.
Age of AD Indiv.
Race (African Am.=1)
ApoE e4 Dose
Cognition Score (MoCA)
phospho-181 Tau CSF ELISA
Total Tau CSF ELISA
ABeta CSF ELISA
ABeta/Tau Ratio

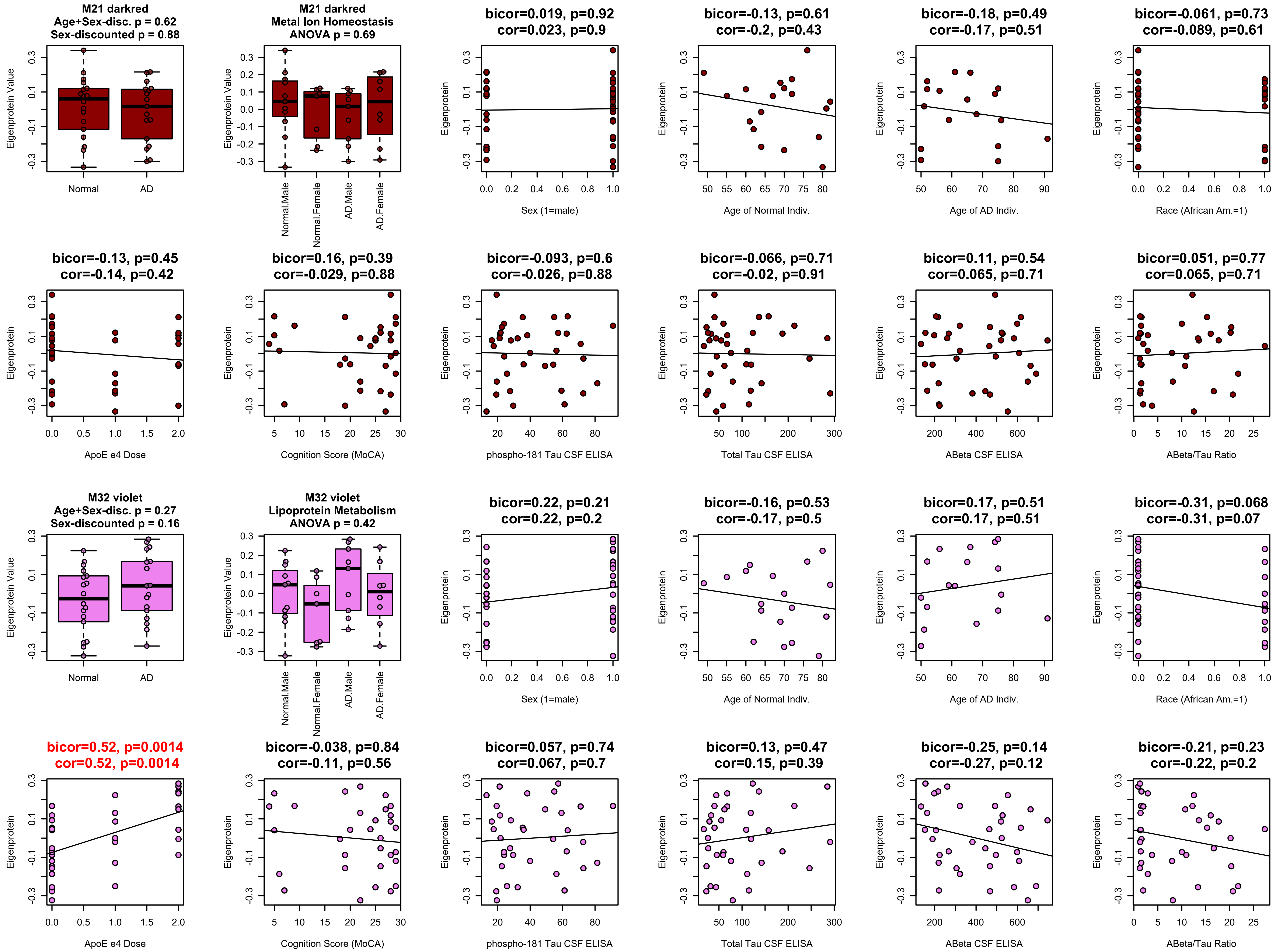
M21 darkred
Age+Sex-disc. p = 0.62
Sex-discounted p = 0.88
M21 darkred
Metal Ion Homeostasis
ANOVA p = 0.69
bicor=0.019, p=0.92
cor=0.023, p=0.9
bicor=-0.13, p=0.61
cor=-0.2, p=0.43
bicor=-0.18, p=0.49
cor=-0.17, p=0.51
bicor=-0.061, p=0.73
cor=-0.089, p=0.61
Eigenprotein Value
Eigenprotein
Normal
AD
Normal.Male
Normal.Female
AD.Male
AD.Female
Sex (1=male)
Age of Normal Indiv.
Age of AD Indiv.
Race (African Am.=1)
bicor=-0.13, p=0.45
cor=-0.14, p=0.42
bicor=0.16, p=0.39
cor=-0.029, p=0.88
bicor=-0.093, p=0.6
cor=-0.026, p=0.88
bicor=-0.066, p=0.71
cor=-0.02, p=0.91
bicor=0.11, p=0.54
cor=0.065, p=0.71
bicor=0.051, p=0.77
cor=0.065, p=0.71
ApoE e4 Dose
Cognition Score (MoCA)
phospho-181 Tau CSF ELISA
Total Tau CSF ELISA
ABeta CSF ELISA
ABeta/Tau Ratio
M32 violet
Age+Sex-disc. p = 0.27
Sex-discounted p = 0.16
M32 violet
Lipoprotein Metabolism
ANOVA p = 0.42
bicor=0.22, p=0.21
cor=0.22, p=0.2
bicor=-0.16, p=0.53
cor=-0.17, p=0.5
bicor=0.17, p=0.51
cor=0.17, p=0.51
bicor=-0.31, p=0.068
cor=-0.31, p=0.07
bicor=0.52, p=0.0014
cor=0.52, p=0.0014
bicor=-0.038, p=0.84
cor=-0.11, p=0.56
bicor=0.057, p=0.74
cor=0.067, p=0.7
bicor=0.13, p=0.47
cor=0.15, p=0.39
bicor=-0.25, p=0.14
cor=-0.27, p=0.12
bicor=-0.21, p=0.23
cor=-0.22, p=0.2

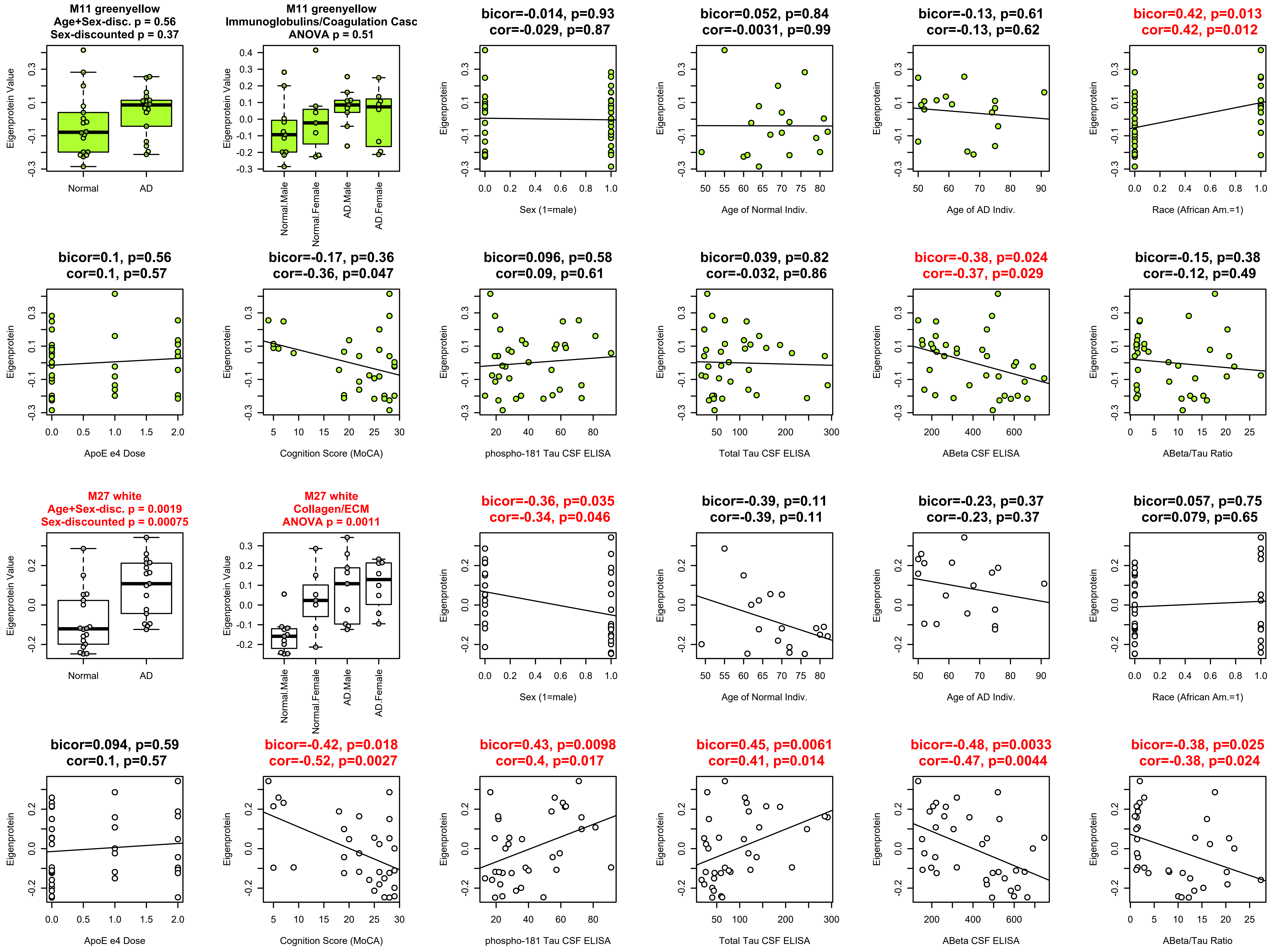

M11 greenyellow
Age+Sex-disc. p = 0.56
Sex-discounted p = 0.37
M11 greenyellow
Immunoglobulins/Coagulation Casc
ANOVA p = 0.51
bicor=-0.014, p=0.93
cor=-0.029, p=0.87
bicor=0.052, p=0.84
cor=-0.0031, p=0.99
bicor=-0.13, p=0.61
cor=-0.13, p=0.62
bicor=0.42, p=0.013
cor=0.42, p=0.012
Eigenprotein Value
Eigenprotein
Normal
AD
Normal.Male
Normal.Female
AD.Male
AD.Female
Sex (1=male)
Age of Normal Indiv.
Age of AD Indiv.
Race (African Am.=1)
bicor=0.1, p=0.56
cor=0.1, p=0.57
bicor=-0.17, p=0.36
cor=-0.36, p=0.047
bicor=0.096, p=0.58
cor=0.09, p=0.61
bicor=0.039, p=0.82
cor=-0.032, p=0.86
bicor=-0.38, p=0.024
cor=-0.37, p=0.029
bicor=-0.15, p=0.38
cor=-0.12, p=0.49
ApoE e4 Dose
Cognition Score (MoCA)
phospho-181 Tau CSF ELISA
Total Tau CSF ELISA
ABeta CSF ELISA
ABeta/Tau Ratio
M27 white
Age+Sex-disc. p = 0.0019
Sex-discounted p = 0.00075
M27 white
Collagen/ECM
ANOVA p = 0.0011
bicor=-0.36, p=0.035
cor=-0.34, p=0.046
bicor=-0.39, p=0.11
cor=-0.39, p=0.11
bicor=-0.23, p=0.37
cor=-0.23, p=0.37
bicor=0.057, p=0.75
cor=0.079, p=0.65
bicor=0.094, p=0.59
cor=0.1, p=0.57
bicor=-0.42, p=0.018
cor=-0.52, p=0.0027
bicor=0.43, p=0.0098
cor=0.4, p=0.017
bicor=0.45, p=0.0061
cor=0.41, p=0.014
bicor=-0.48, p=0.0033
cor=-0.47, p=0.0044
bicor=-0.38, p=0.025
cor=-0.38, p=0.024

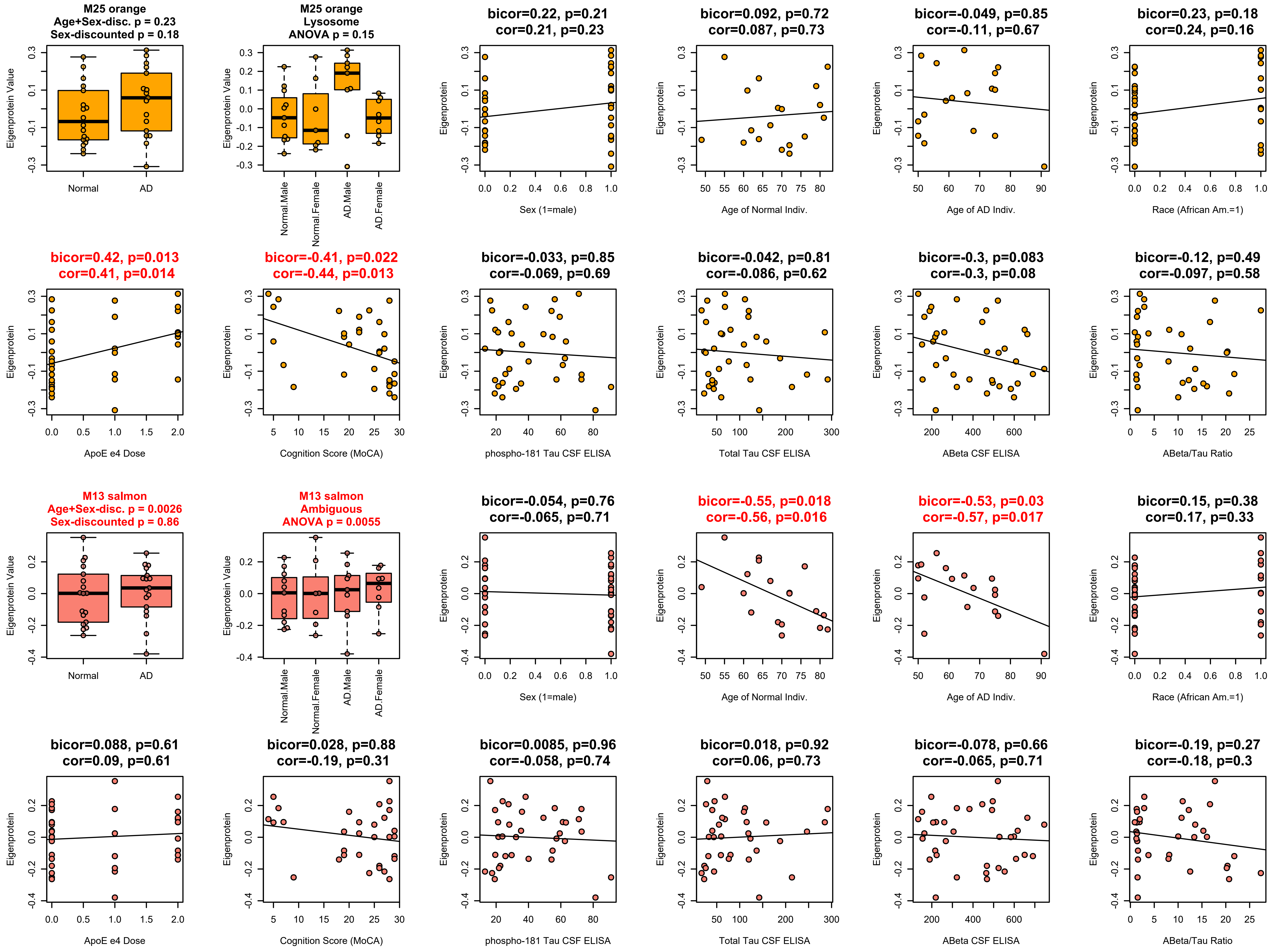
M25 orange
Age+Sex-disc. p = 0.23
Sex-discounted p = 0.18
Eigenprotein Value
Normal
AD
M25 orange
Lysosome
ANOVA p = 0.15
Eigenprotein Value
Normal.Male
Normal.Female
AD.Male
AD.Female
bicor=0.22, p=0.21
cor=0.21, p=0.23
Eigenprotein
Sex (1=male)
bicor=0.092, p=0.72
cor=0.087, p=0.73
Eigenprotein
Age of Normal Indiv.
bicor=-0.049, p=0.85
cor=-0.11, p=0.67
Eigenprotein
Age of AD Indiv.
bicor=0.23, p=0.18
cor=0.24, p=0.16
Eigenprotein
Race (African Am.=1)
bicor=0.42, p=0.013
cor=0.41, p=0.014
Eigenprotein
ApoE e4 Dose
bicor=-0.41, p=0.022
cor=-0.44, p=0.013
Eigenprotein
Cognition Score (MoCA)
bicor=-0.033, p=0.85
cor=-0.069, p=0.69
Eigenprotein
phospho-181 Tau CSF ELISA
bicor=-0.042, p=0.81
cor=-0.086, p=0.62
Eigenprotein
Total Tau CSF ELISA
bicor=-0.3, p=0.083
cor=-0.3, p=0.08
Eigenprotein
ABeta CSF ELISA
bicor=-0.12, p=0.49
cor=-0.097, p=0.58
Eigenprotein
ABeta/Tau Ratio
M13 salmon
Age+Sex-disc. p = 0.0026
Sex-discounted p = 0.86
Eigenprotein Value
Normal
AD
M13 salmon
Ambiguous
ANOVA p = 0.0055
Eigenprotein Value
Normal.Male
Normal.Female
AD.Male
AD.Female
bicor=-0.054, p=0.76
cor=-0.065, p=0.71
Eigenprotein
Sex (1=male)
bicor=-0.55, p=0.018
cor=-0.56, p=0.016
Eigenprotein
Age of Normal Indiv.
bicor=-0.53, p=0.03
cor=-0.57, p=0.017
Eigenprotein
Age of AD Indiv.
bicor=0.15, p=0.38
cor=0.17, p=0.33
Eigenprotein
Race (African Am.=1)
bicor=0.088, p=0.61
cor=0.09, p=0.61
Eigenprotein
ApoE e4 Dose
bicor=0.028, p=0.88
cor=-0.19, p=0.31
Eigenprotein
Cognition Score (MoCA)
bicor=0.0085, p=0.96
cor=-0.058, p=0.74
Eigenprotein
phospho-181 Tau CSF ELISA
bicor=0.018, p=0.92
cor=0.06, p=0.73
Eigenprotein
Total Tau CSF ELISA
bicor=-0.078, p=0.66
cor=-0.065, p=0.71
Eigenprotein
ABeta CSF ELISA
bicor=-0.19, p=0.27
cor=-0.18, p=0.3
Eigenprotein
ABeta/Tau Ratio

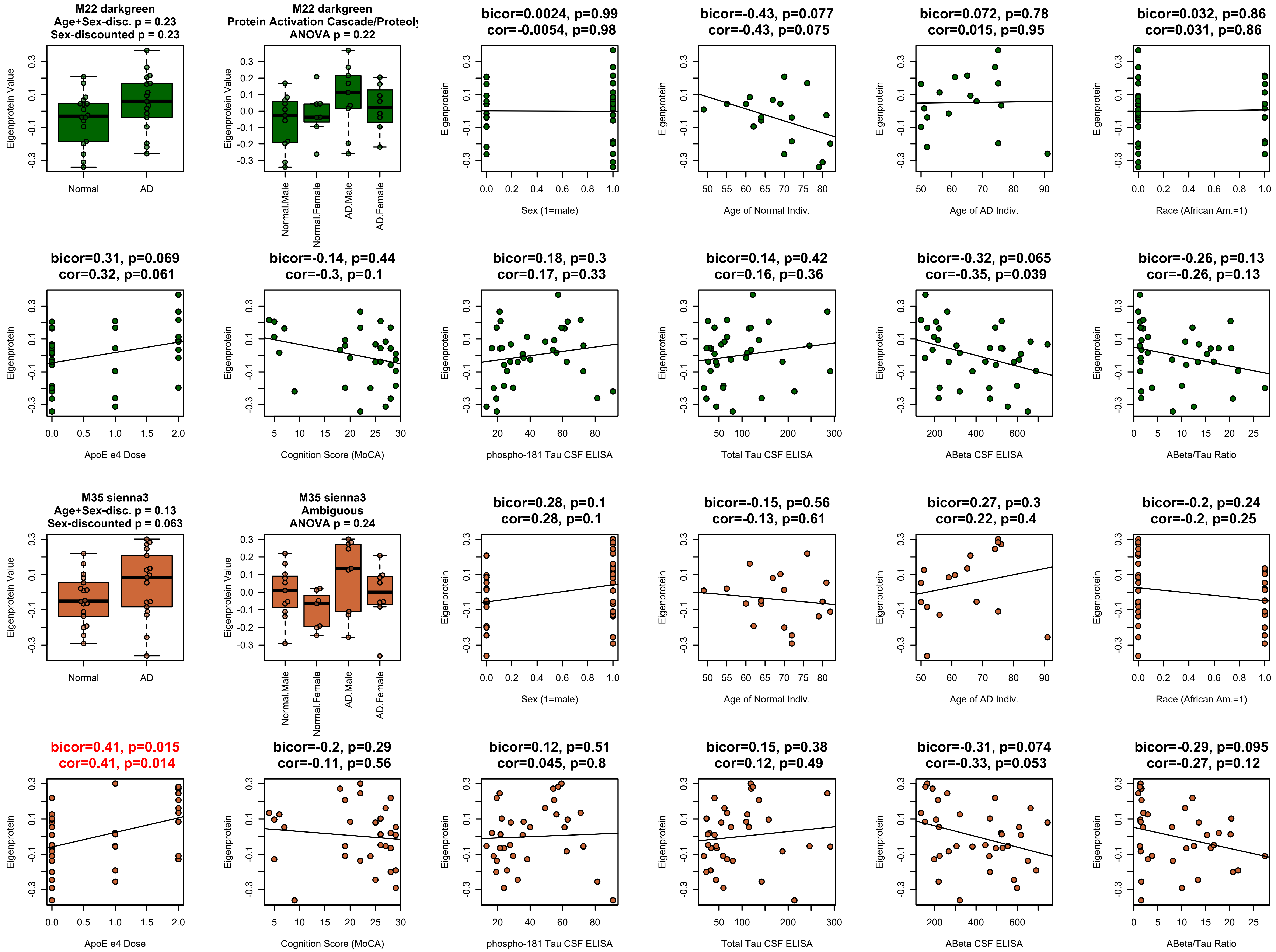

M22 darkgreen
Age+Sex-disc. p = 0.23
Sex-discounted p = 0.23
M22 darkgreen
Protein Activation Cascade/Proteoly
ANOVA p = 0.22
bicor=0.0024, p=0.99
cor=-0.0054, p=0.98
bicor=-0.43, p=0.077
cor=-0.43, p=0.075
bicor=0.072, p=0.78
cor=0.015, p=0.95
bicor=0.032, p=0.86
cor=0.031, p=0.86
Eigenprotein Value
Eigenprotein
Normal
AD
Normal.Male
Normal.Female
AD.Male
AD.Female
Sex (1=male)
Age of Normal Indiv.
Age of AD Indiv.
Race (African Am.=1)
bicor=0.31, p=0.069
cor=0.32, p=0.061
bicor=-0.14, p=0.44
cor=-0.3, p=0.1
bicor=0.18, p=0.3
cor=0.17, p=0.33
bicor=0.14, p=0.42
cor=0.16, p=0.36
bicor=-0.32, p=0.065
cor=-0.35, p=0.039
bicor=-0.26, p=0.13
cor=-0.26, p=0.13
ApoE e4 Dose
Cognition Score (MoCA)
phospho-181 Tau CSF ELISA
Total Tau CSF ELISA
ABeta CSF ELISA
ABeta/Tau Ratio
M35 sienna3
Age+Sex-disc. p = 0.13
Sex-discounted p = 0.063
M35 sienna3
Ambiguous
ANOVA p = 0.24
bicor=0.28, p=0.1
cor=0.28, p=0.1
bicor=-0.15, p=0.56
cor=-0.13, p=0.61
bicor=0.27, p=0.3
cor=0.22, p=0.4
bicor=-0.2, p=0.24
cor=-0.2, p=0.25
bicor=0.41, p=0.015
cor=0.41, p=0.014
bicor=-0.2, p=0.29
cor=-0.11, p=0.56
bicor=0.12, p=0.51
cor=0.045, p=0.8
bicor=0.15, p=0.38
cor=0.12, p=0.49
bicor=-0.31, p=0.074
cor=-0.33, p=0.053
bicor=-0.29, p=0.095
cor=-0.27, p=0.12

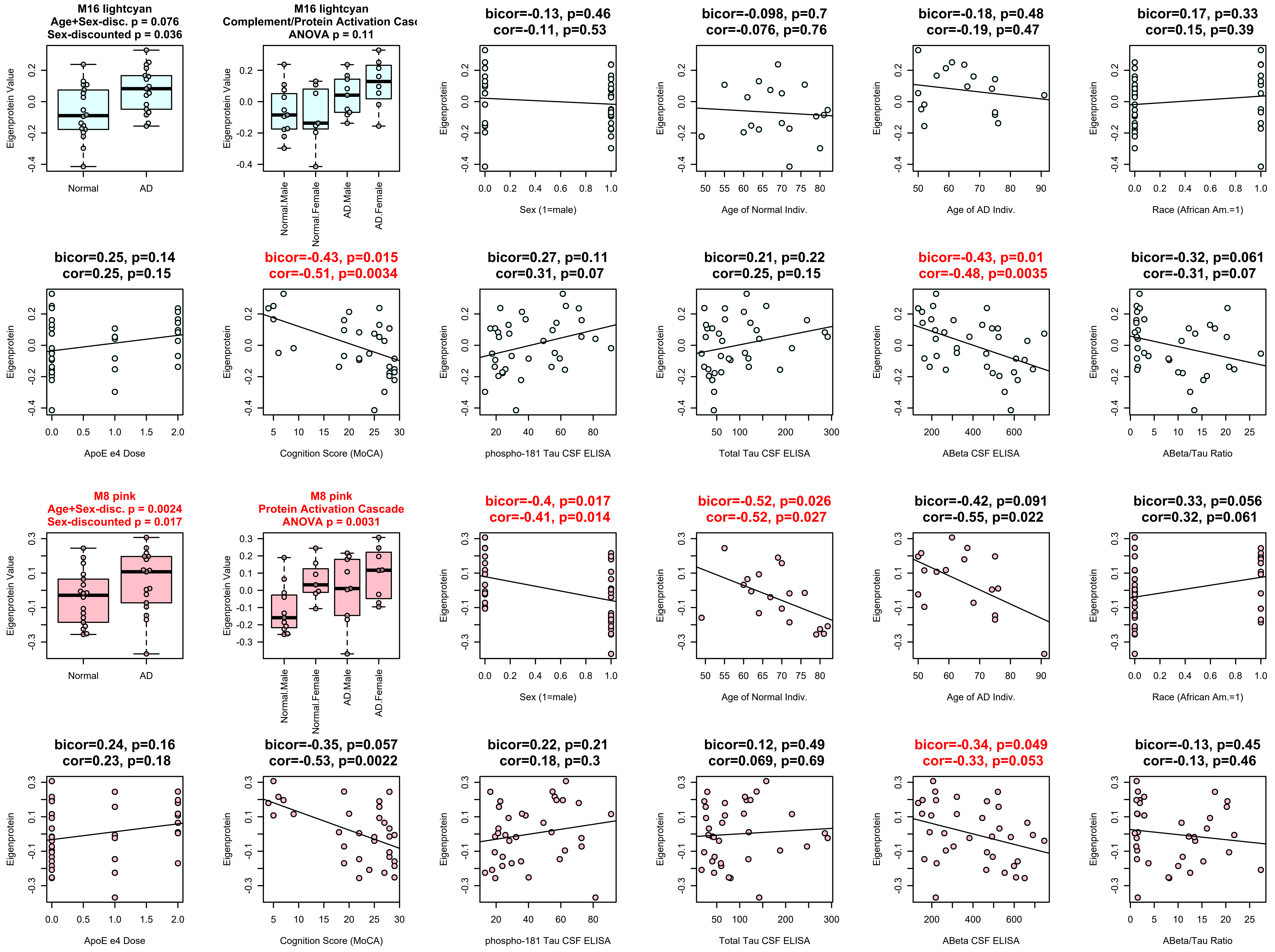
M16 lightcyan
Age+Sex-disc. p = 0.076
Sex-discounted p = 0.036
Eigenprotein Value
Normal
AD
M16 lightcyan
Complement/Protein Activation Casc
ANOVA p = 0.11
Normal.Male
Normal.Female
AD.Male
AD.Female
bicor=-0.13, p=0.46
cor=-0.11, p=0.53
Sex (1=male)
bicor=-0.098, p=0.7
cor=-0.076, p=0.76
Age of Normal Indiv.
bicor=-0.18, p=0.48
cor=-0.19, p=0.47
Age of AD Indiv.
bicor=0.17, p=0.33
cor=0.15, p=0.39
Race (African Am.=1)
Eigenprotein
bicor=0.25, p=0.14
cor=0.25, p=0.15
ApoE e4 Dose
bicor=-0.43, p=0.015
cor=-0.51, p=0.0034
Cognition Score (MoCA)
bicor=0.27, p=0.11
cor=0.31, p=0.07
phospho-181 Tau CSF ELISA
bicor=0.21, p=0.22
cor=0.25, p=0.15
Total Tau CSF ELISA
bicor=-0.43, p=0.01
cor=-0.48, p=0.0035
ABeta CSF ELISA
bicor=-0.32, p=0.061
cor=-0.31, p=0.07
ABeta/Tau Ratio
M8 pink
Age+Sex-disc. p = 0.0024
Sex-discounted p = 0.017
M8 pink
Protein Activation Cascade
ANOVA p = 0.0031
bicor=-0.4, p=0.017
cor=-0.41, p=0.014
bicor=-0.52, p=0.026
cor=-0.52, p=0.027
bicor=-0.42, p=0.091
cor=-0.55, p=0.022
bicor=0.33, p=0.056
cor=0.32, p=0.061
bicor=0.24, p=0.16
cor=0.23, p=0.18
bicor=-0.35, p=0.057
cor=-0.53, p=0.0022
bicor=0.22, p=0.21
cor=0.18, p=0.3
bicor=0.12, p=0.49
cor=0.069, p=0.69
bicor=-0.34, p=0.049
cor=-0.33, p=0.053
bicor=-0.13, p=0.45
cor=-0.13, p=0.46

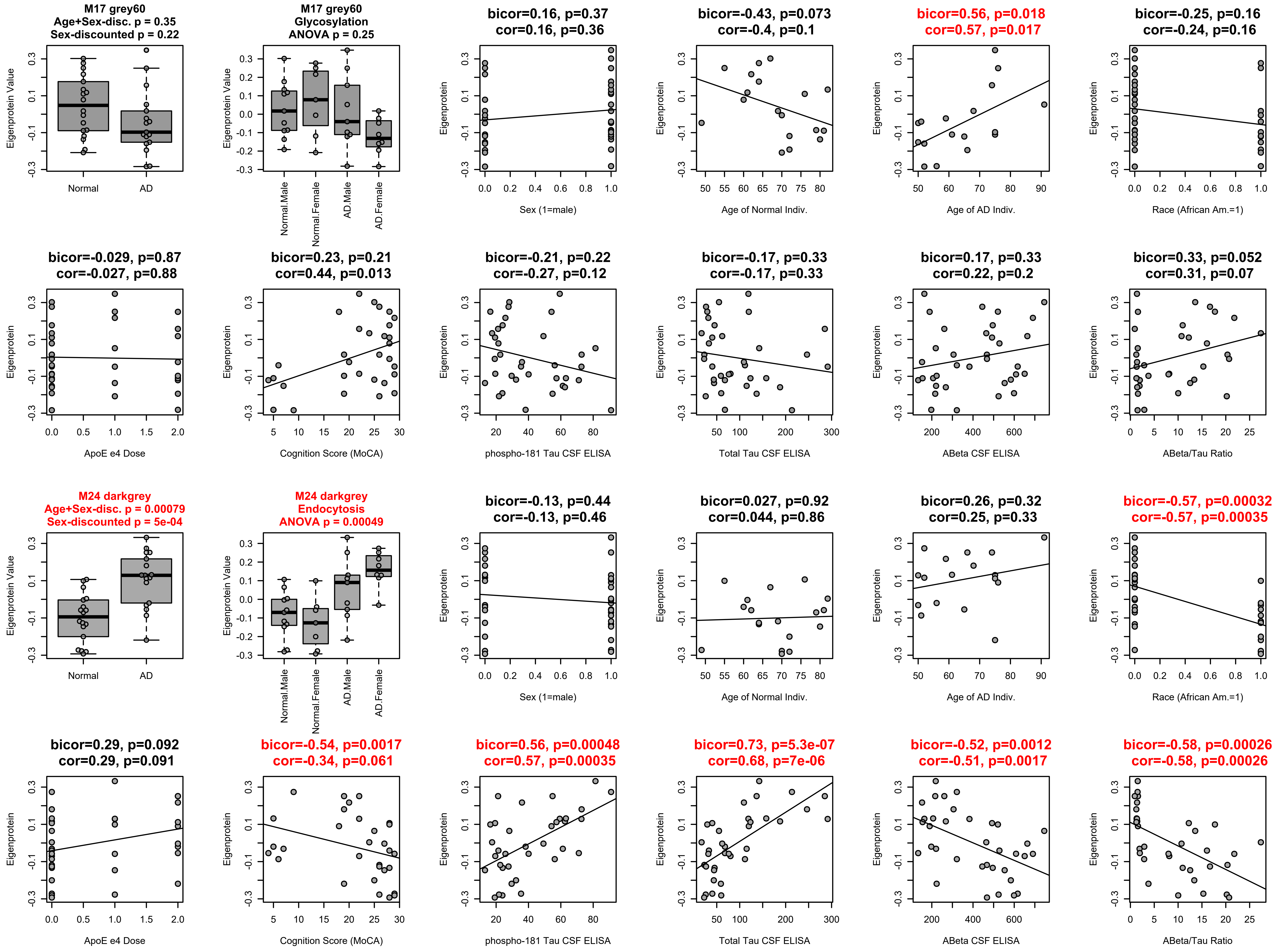
M17 grey60
Age+Sex-disc. p = 0.35
Sex-discounted p = 0.22
Eigenprotein Value
Normal
AD
M17 grey60
Glycosylation
ANOVA p = 0.25
Normal.Male
Normal.Female
AD.Male
AD.Female
bicor=0.16, p=0.37
cor=0.16, p=0.36
Sex (1=male)
bicor=-0.43, p=0.073
cor=-0.4, p=0.1
Age of Normal Indiv.
bicor=0.56, p=0.018
cor=0.57, p=0.017
Age of AD Indiv.
bicor=-0.25, p=0.16
cor=-0.24, p=0.16
Race (African Am.=1)
bicor=-0.029, p=0.87
cor=-0.027, p=0.88
ApoE e4 Dose
bicor=0.23, p=0.21
cor=0.44, p=0.013
Cognition Score (MoCA)
bicor=-0.21, p=0.22
cor=-0.27, p=0.12
phospho-181 Tau CSF ELISA
bicor=-0.17, p=0.33
cor=-0.17, p=0.33
Total Tau CSF ELISA
bicor=0.17, p=0.33
cor=0.22, p=0.2
ABeta CSF ELISA
bicor=0.33, p=0.052
cor=0.31, p=0.07
ABeta/Tau Ratio
M24 darkgrey
Age+Sex-disc. p = 0.00079
Sex-discounted p = 5e-04
M24 darkgrey
Endocytosis
ANOVA p = 0.00049
bicor=-0.13, p=0.44
cor=-0.13, p=0.46
bicor=0.027, p=0.92
cor=0.044, p=0.86
bicor=0.26, p=0.32
cor=0.25, p=0.33
bicor=-0.57, p=0.00032
cor=-0.57, p=0.00035
bicor=0.29, p=0.092
cor=0.29, p=0.091
bicor=-0.54, p=0.0017
cor=-0.34, p=0.061
bicor=0.56, p=0.00048
cor=0.57, p=0.00035
bicor=0.73, p=5.3e-07
cor=0.68, p=7e-06
bicor=-0.52, p=0.0012
cor=-0.51, p=0.0017
bicor=-0.58, p=0.00026
cor=-0.58, p=0.00026
Eigenprotein

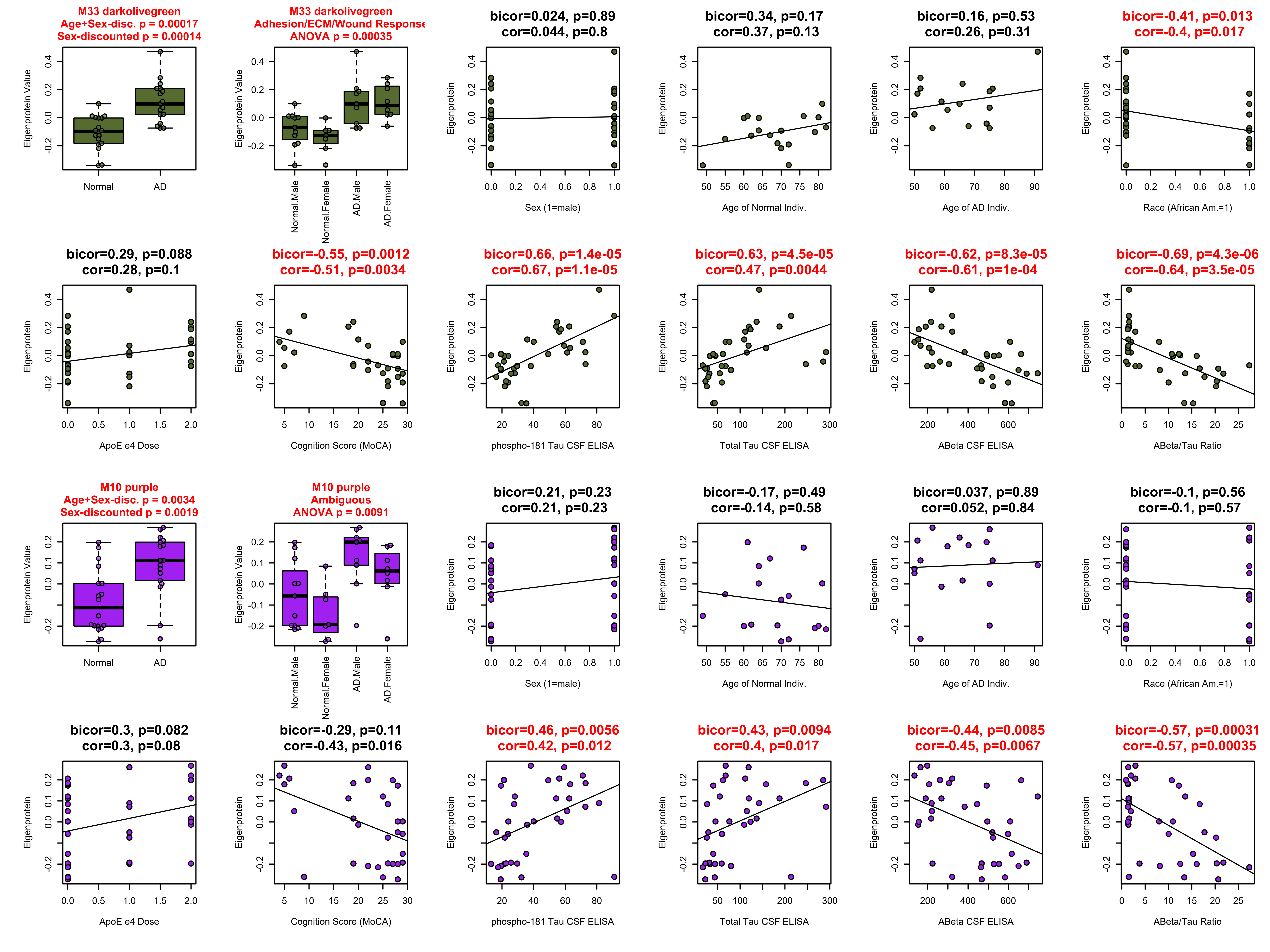
M33 darkolivegreen
Age+Sex-disc. p = 0.00017
Sex-discounted p = 0.00014
Eigenprotein Value
Normal
AD
M33 darkolivegreen
Adhesion/ECM/Wound Response
ANOVA p = 0.00035
Normal.Male
Normal.Female
AD.Male
AD.Female
bicor=0.024, p=0.89
cor=0.044, p=0.8
Sex (1=male)
bicor=0.34, p=0.17
cor=0.37, p=0.13
Age of Normal Indiv.
bicor=0.16, p=0.53
cor=0.26, p=0.31
Age of AD Indiv.
bicor=-0.41, p=0.013
cor=-0.4, p=0.017
Race (African Am.=1)
bicor=0.29, p=0.088
cor=0.28, p=0.1
ApoE e4 Dose
bicor=-0.55, p=0.0012
cor=-0.51, p=0.0034
Cognition Score (MoCA)
bicor=0.66, p=1.4e-05
cor=0.67, p=1.1e-05
phospho-181 Tau CSF ELISA
bicor=0.63, p=4.5e-05
cor=0.47, p=0.0044
Total Tau CSF ELISA
bicor=-0.62, p=8.3e-05
cor=-0.61, p=1e-04
ABeta CSF ELISA
bicor=-0.69, p=4.3e-06
cor=-0.64, p=3.5e-05
ABeta/Tau Ratio
Eigenprotein
M10 purple
Age+Sex-disc. p = 0.0034
Sex-discounted p = 0.0019
M10 purple
Ambiguous
ANOVA p = 0.0091
bicor=0.21, p=0.23
cor=0.21, p=0.23
bicor=-0.17, p=0.49
cor=-0.14, p=0.58
bicor=0.037, p=0.89
cor=0.052, p=0.84
bicor=-0.1, p=0.56
cor=-0.1, p=0.57
bicor=0.3, p=0.082
cor=0.3, p=0.08
bicor=-0.29, p=0.11
cor=-0.43, p=0.016
bicor=0.46, p=0.0056
cor=0.42, p=0.012
bicor=0.43, p=0.0094
cor=0.4, p=0.017
bicor=-0.44, p=0.0085
cor=-0.45, p=0.0067
bicor=-0.57, p=0.00031
cor=-0.57, p=0.00035

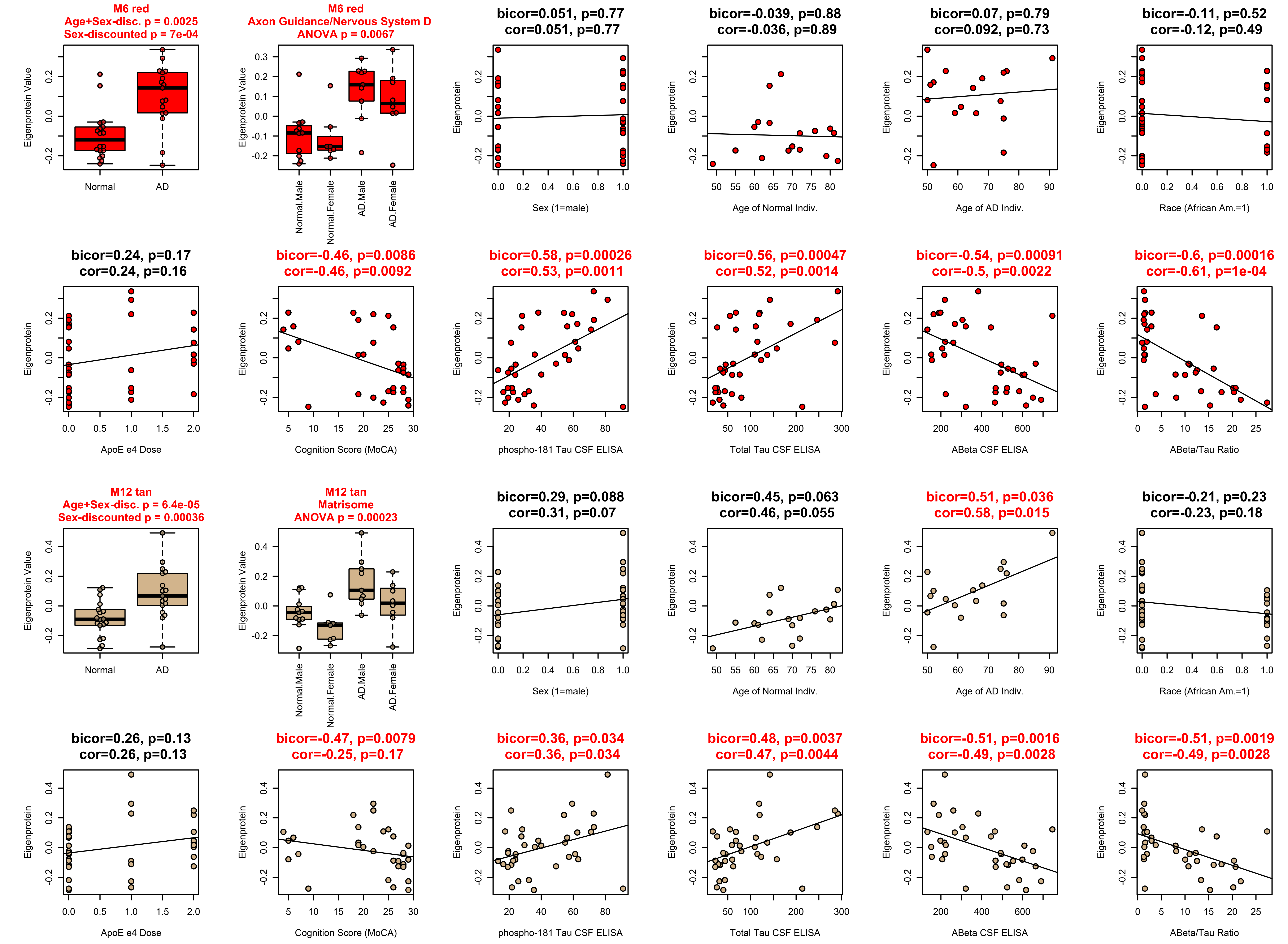
M6 red
Age+Sex-disc. p = 0.0025
Sex-discounted p = 7e-04
M6 red
Axon Guidance/Nervous System D
ANOVA p = 0.0067
bicor=0.051, p=0.77
cor=0.051, p=0.77
bicor=-0.039, p=0.88
cor=-0.036, p=0.89
bicor=0.07, p=0.79
cor=0.092, p=0.73
bicor=-0.11, p=0.52
cor=-0.12, p=0.49
bicor=0.24, p=0.17
cor=0.24, p=0.16
bicor=-0.46, p=0.0086
cor=-0.46, p=0.0092
bicor=0.58, p=0.00026
cor=0.53, p=0.0011
bicor=0.56, p=0.00047
cor=0.52, p=0.0014
bicor=-0.54, p=0.00091
cor=-0.5, p=0.0022
bicor=-0.6, p=0.00016
cor=-0.61, p=1e-04
M12 tan
Age+Sex-disc. p = 6.4e-05
Sex-discounted p = 0.00036
M12 tan
Matrisome
ANOVA p = 0.00023
bicor=0.29, p=0.088
cor=0.31, p=0.07
bicor=0.45, p=0.063
cor=0.46, p=0.055
bicor=0.51, p=0.036
cor=0.58, p=0.015
bicor=-0.21, p=0.23
cor=-0.23, p=0.18
bicor=0.26, p=0.13
cor=0.26, p=0.13
bicor=-0.47, p=0.0079
cor=-0.25, p=0.17
bicor=0.36, p=0.034
cor=0.36, p=0.034
bicor=0.48, p=0.0037
cor=0.47, p=0.0044
bicor=-0.51, p=0.0016
cor=-0.49, p=0.0028
bicor=-0.51, p=0.0019
cor=-0.49, p=0.0028
Eigenprotein Value
Eigenprotein
Normal
AD
Normal.Male
Normal.Female
AD.Male
AD.Female
Sex (1=male)
Age of Normal Indiv.
Age of AD Indiv.
Race (African Am.=1)
ApoE e4 Dose
Cognition Score (MoCA)
phospho-181 Tau CSF ELISA
Total Tau CSF ELISA
ABeta CSF ELISA
ABeta/Tau Ratio

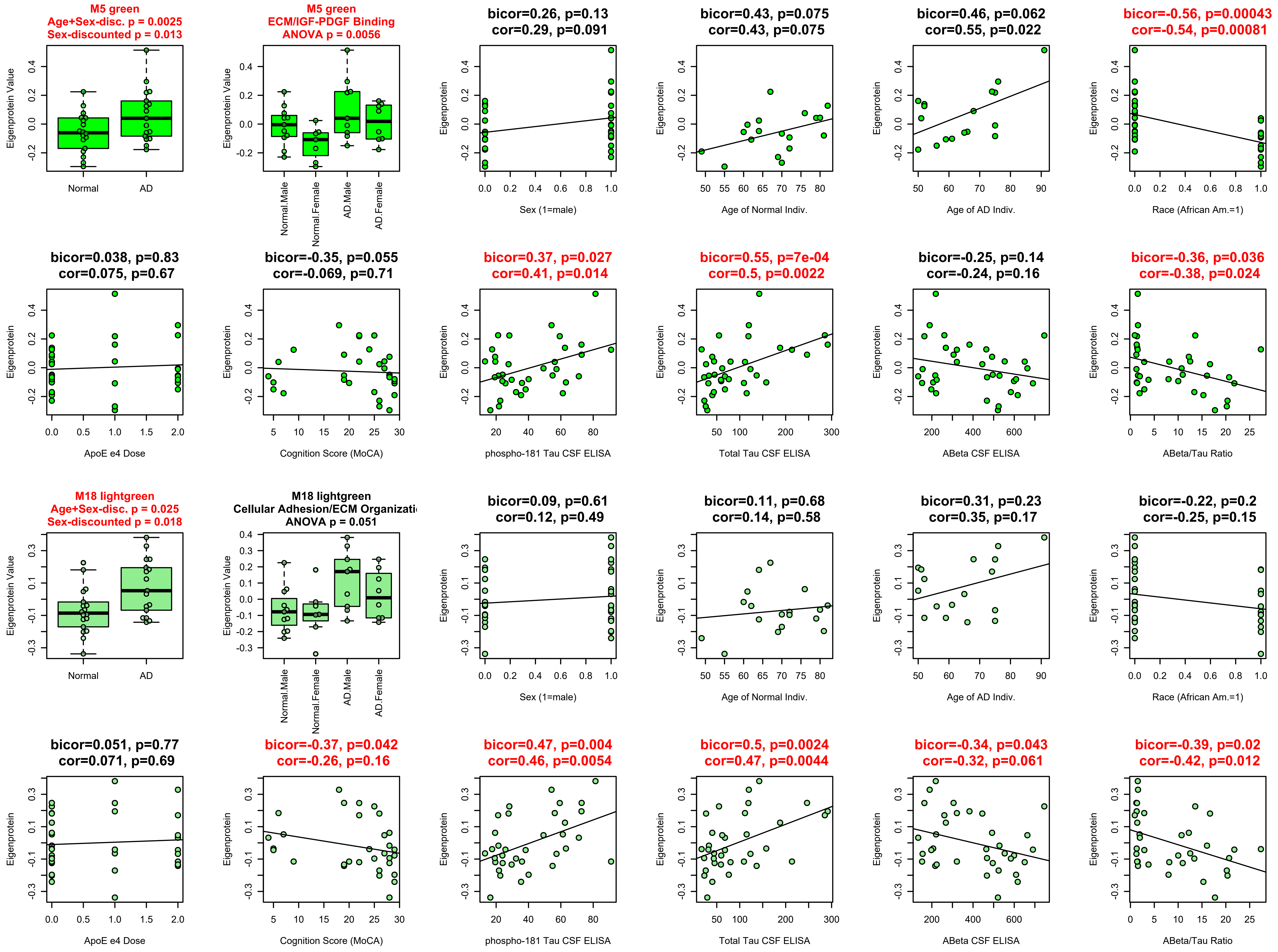
M5 green
Age+Sex-disc. p = 0.0025
Sex-discounted p = 0.013
Eigenprotein Value
Normal
AD
M5 green
ECM/IGF-PDGF Binding
ANOVA p = 0.0056
Normal.Male
Normal.Female
AD.Male
AD.Female
bicor=0.26, p=0.13
cor=0.29, p=0.091
Sex (1=male)
bicor=0.43, p=0.075
cor=0.43, p=0.075
Age of Normal Indiv.
bicor=0.46, p=0.062
cor=0.55, p=0.022
Age of AD Indiv.
bicor=-0.56, p=0.00043
cor=-0.54, p=0.00081
Race (African Am.=1)
bicor=0.038, p=0.83
cor=0.075, p=0.67
ApoE e4 Dose
bicor=-0.35, p=0.055
cor=-0.069, p=0.71
Cognition Score (MoCA)
bicor=0.37, p=0.027
cor=0.41, p=0.014
phospho-181 Tau CSF ELISA
bicor=0.55, p=7e-04
cor=0.5, p=0.0022
Total Tau CSF ELISA
bicor=-0.25, p=0.14
cor=-0.24, p=0.16
ABeta CSF ELISA
bicor=-0.36, p=0.036
cor=-0.38, p=0.024
ABeta/Tau Ratio
M18 lightgreen
Age+Sex-disc. p = 0.025
Sex-discounted p = 0.018
M18 lightgreen
Cellular Adhesion/ECM Organizati
ANOVA p = 0.051
bicor=0.09, p=0.61
cor=0.12, p=0.49
bicor=0.11, p=0.68
cor=0.14, p=0.58
bicor=0.31, p=0.23
cor=0.35, p=0.17
bicor=-0.22, p=0.2
cor=-0.25, p=0.15
bicor=0.051, p=0.77
cor=0.071, p=0.69
bicor=-0.37, p=0.042
cor=-0.26, p=0.16
bicor=0.47, p=0.004
cor=0.46, p=0.0054
bicor=0.5, p=0.0024
cor=0.47, p=0.0044
bicor=-0.34, p=0.043
cor=-0.32, p=0.061
bicor=-0.39, p=0.02
cor=-0.42, p=0.012
Eigenprotein

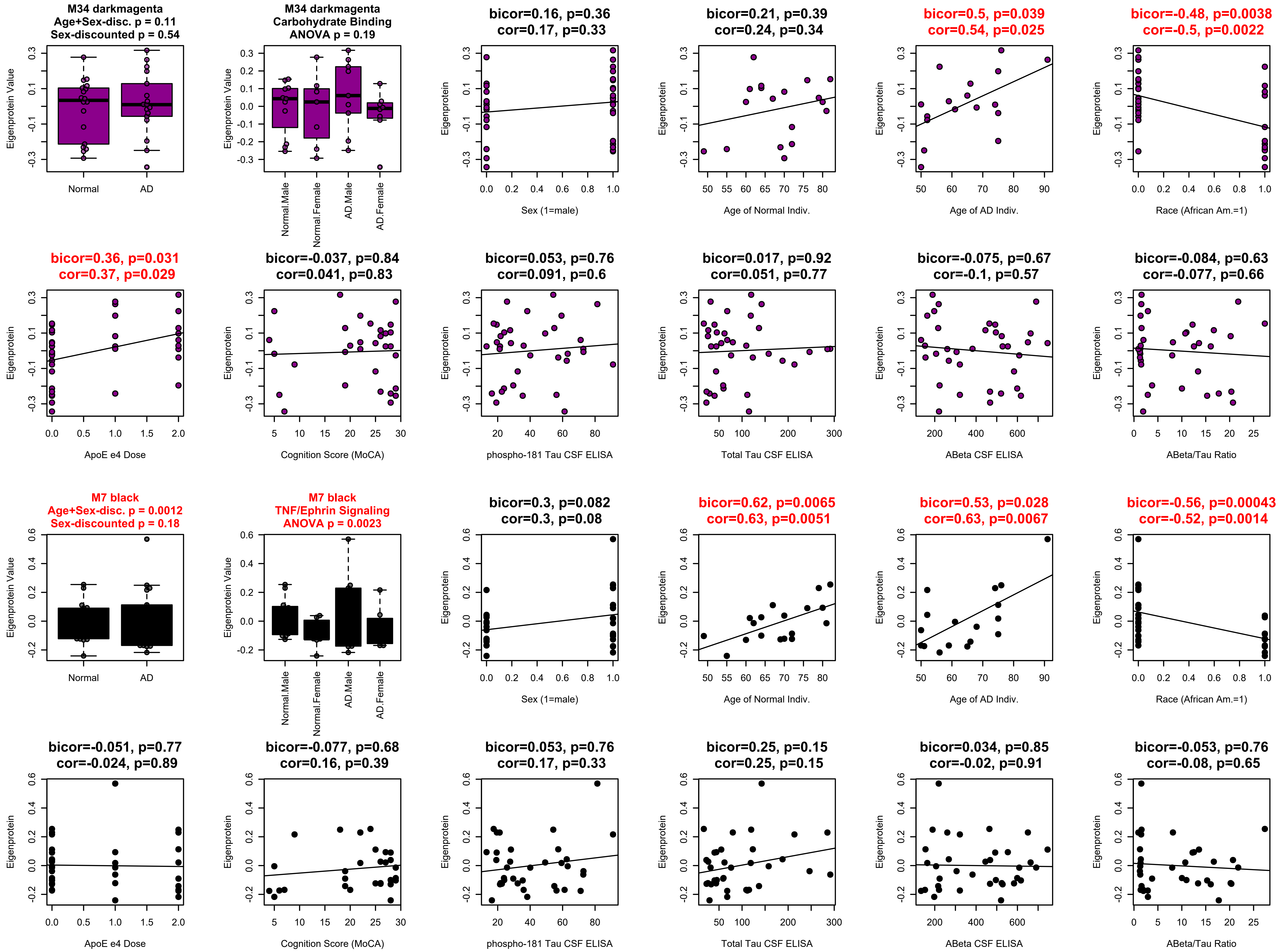
M34 darkmagenta
Age+Sex-disc. p = 0.11
Sex-discounted p = 0.54
Eigenprotein Value
Normal
AD
M34 darkmagenta
Carbohydrate Binding
ANOVA p = 0.19
Normal.Male
Normal.Female
AD.Male
AD.Female
bicor=0.16, p=0.36
cor=0.17, p=0.33
Sex (1=male)
bicor=0.21, p=0.39
cor=0.24, p=0.34
Age of Normal Indiv.
bicor=0.5, p=0.039
cor=0.54, p=0.025
Age of AD Indiv.
bicor=-0.48, p=0.0038
cor=-0.5, p=0.0022
Race (African Am.=1)
bicor=0.36, p=0.031
cor=0.37, p=0.029
ApoE e4 Dose
bicor=-0.037, p=0.84
cor=0.041, p=0.83
Cognition Score (MoCA)
bicor=0.053, p=0.76
cor=0.091, p=0.6
phospho-181 Tau CSF ELISA
bicor=0.017, p=0.92
cor=0.051, p=0.77
Total Tau CSF ELISA
bicor=-0.075, p=0.67
cor=-0.1, p=0.57
ABeta CSF ELISA
bicor=-0.084, p=0.63
cor=-0.077, p=0.66
ABeta/Tau Ratio
M7 black
Age+Sex-disc. p = 0.0012
Sex-discounted p = 0.18
M7 black
TNF/Ephrin Signaling
ANOVA p = 0.0023
bicor=0.3, p=0.082
cor=0.3, p=0.08
bicor=0.62, p=0.0065
cor=0.63, p=0.0051
bicor=0.53, p=0.028
cor=0.63, p=0.0067
bicor=-0.56, p=0.00043
cor=-0.52, p=0.0014
bicor=-0.051, p=0.77
cor=-0.024, p=0.89
bicor=-0.077, p=0.68
cor=0.16, p=0.39
bicor=0.053, p=0.76
cor=0.17, p=0.33
bicor=0.25, p=0.15
cor=0.25, p=0.15
bicor=0.034, p=0.85
cor=-0.02, p=0.91
bicor=-0.053, p=0.76
cor=-0.08, p=0.65
Eigenprotein

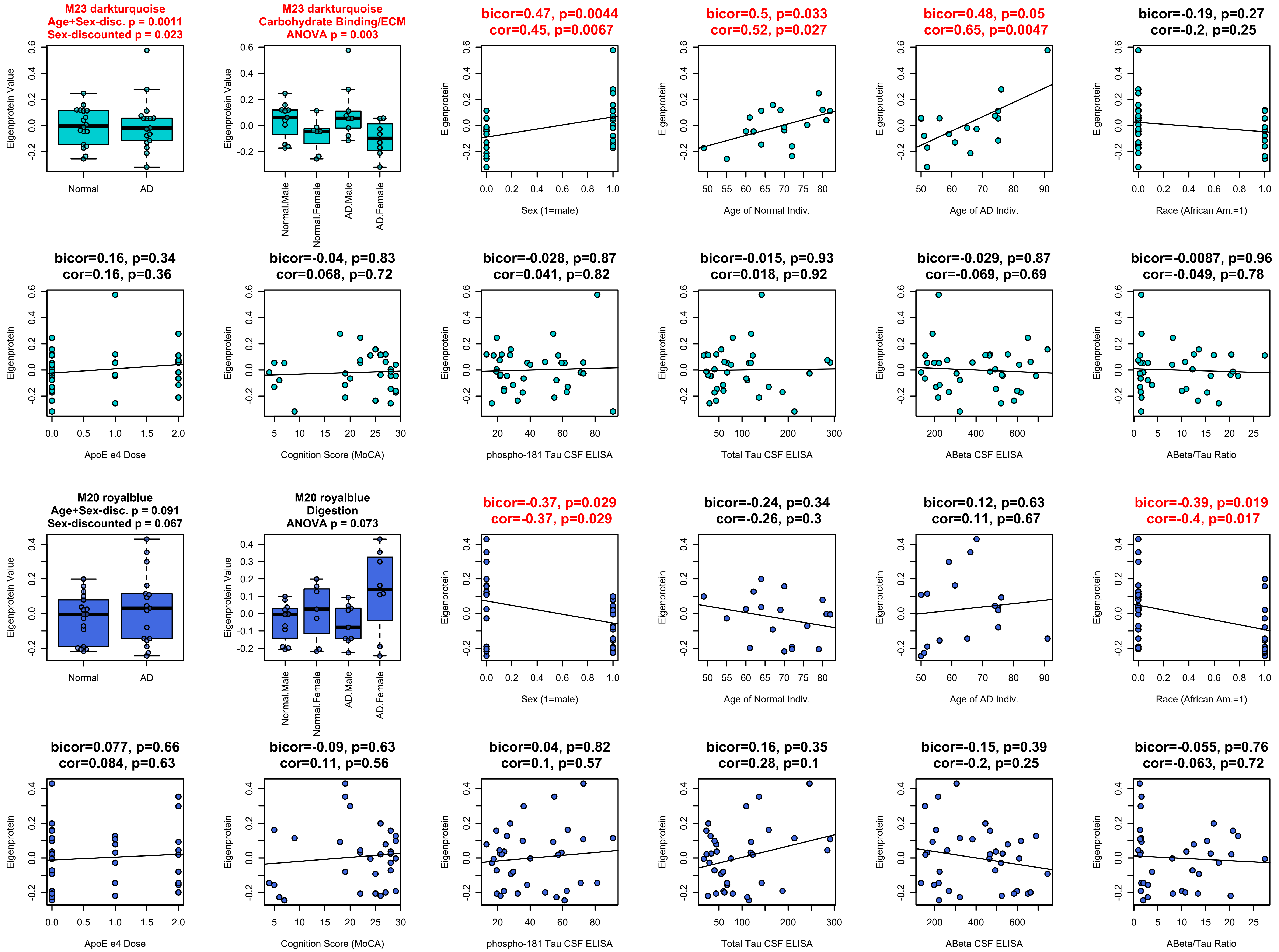
M23 darkturquoise
Age+Sex-disc. p = 0.0011
Sex-discounted p = 0.023
Eigenprotein Value
Normal
AD
M23 darkturquoise
Carbohydrate Binding/ECM
ANOVA p = 0.003
Normal.Male
Normal.Female
AD.Male
AD.Female
bicor=0.47, p=0.0044
cor=0.45, p=0.0067
Eigenprotein
Sex (1=male)
bicor=0.5, p=0.033
cor=0.52, p=0.027
Age of Normal Indiv.
bicor=0.48, p=0.05
cor=0.65, p=0.0047
Age of AD Indiv.
bicor=-0.19, p=0.27
cor=-0.2, p=0.25
Race (African Am.=1)
bicor=0.16, p=0.34
cor=0.16, p=0.36
ApoE e4 Dose
bicor=-0.04, p=0.83
cor=0.068, p=0.72
Cognition Score (MoCA)
bicor=-0.028, p=0.87
cor=0.041, p=0.82
phospho-181 Tau CSF ELISA
bicor=-0.015, p=0.93
cor=0.018, p=0.92
Total Tau CSF ELISA
bicor=-0.029, p=0.87
cor=-0.069, p=0.69
ABeta CSF ELISA
bicor=-0.0087, p=0.96
cor=-0.049, p=0.78
ABeta/Tau Ratio
M20 royalblue
Age+Sex-disc. p = 0.091
Sex-discounted p = 0.067
M20 royalblue
Digestion
ANOVA p = 0.073
bicor=-0.37, p=0.029
cor=-0.37, p=0.029
bicor=-0.24, p=0.34
cor=-0.26, p=0.3
bicor=0.12, p=0.63
cor=0.11, p=0.67
bicor=-0.39, p=0.019
cor=-0.4, p=0.017
bicor=0.077, p=0.66
cor=0.084, p=0.63
bicor=-0.09, p=0.63
cor=0.11, p=0.56
bicor=0.04, p=0.82
cor=0.1, p=0.57
bicor=0.16, p=0.35
cor=0.28, p=0.1
bicor=-0.15, p=0.39
cor=-0.2, p=0.25
bicor=-0.055, p=0.76
cor=-0.063, p=0.72

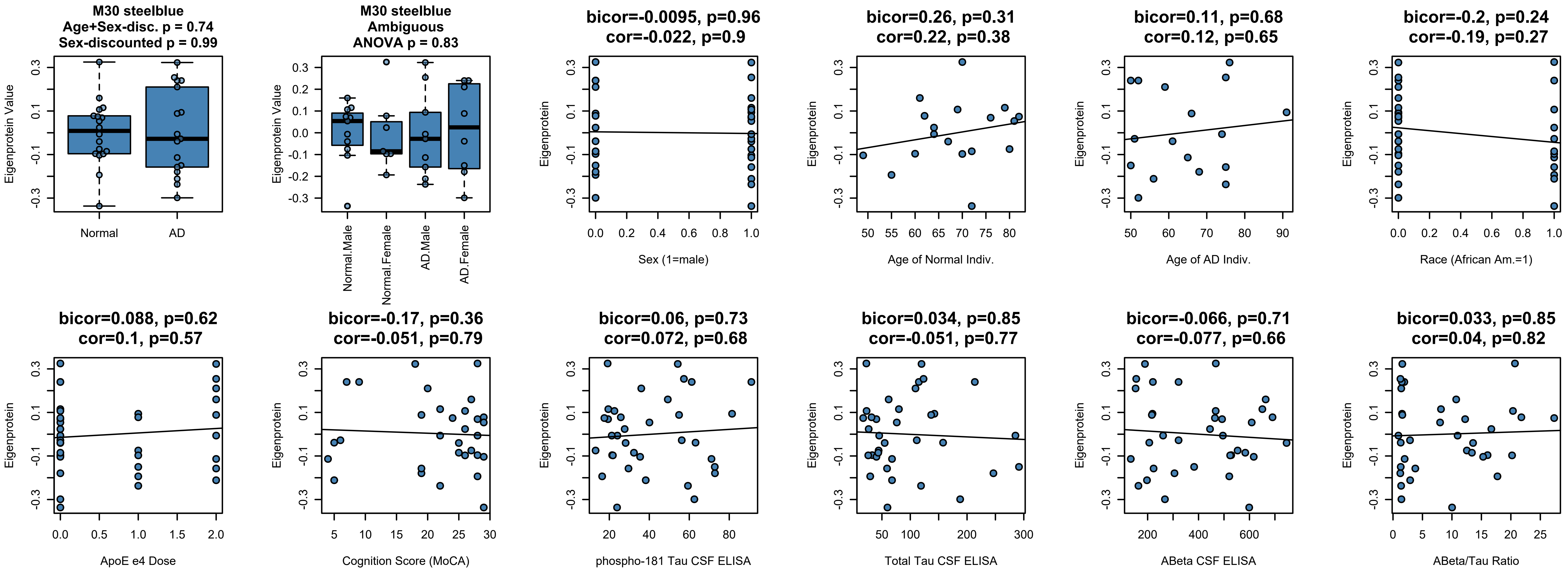

M30 steelblue
Age+Sex-disc. p = 0.74
Sex-discounted p = 0.99
Eigenprotein Value
Normal
AD
M30 steelblue
Ambiguous
ANOVA p = 0.83
Normal.Male
Normal.Female
AD.Male
AD.Female
bicor=-0.0095, p=0.96
cor=-0.022, p=0.9
Sex (1=male)
bicor=0.26, p=0.31
cor=0.22, p=0.38
Age of Normal Indiv.
bicor=0.11, p=0.68
cor=0.12, p=0.65
Age of AD Indiv.
bicor=-0.2, p=0.24
cor=-0.19, p=0.27
Race (African Am.=1)
bicor=0.088, p=0.62
cor=0.1, p=0.57
ApoE e4 Dose
bicor=-0.17, p=0.36
cor=-0.051, p=0.79
Cognition Score (MoCA)
bicor=0.06, p=0.73
cor=0.072, p=0.68
phospho-181 Tau CSF ELISA
bicor=0.034, p=0.85
cor=-0.051, p=0.77
Total Tau CSF ELISA
bicor=-0.066, p=0.71
cor=-0.077, p=0.66
ABeta CSF ELISA
bicor=0.033, p=0.85
cor=0.04, p=0.82
ABeta/Tau Ratio
Eigenprotein

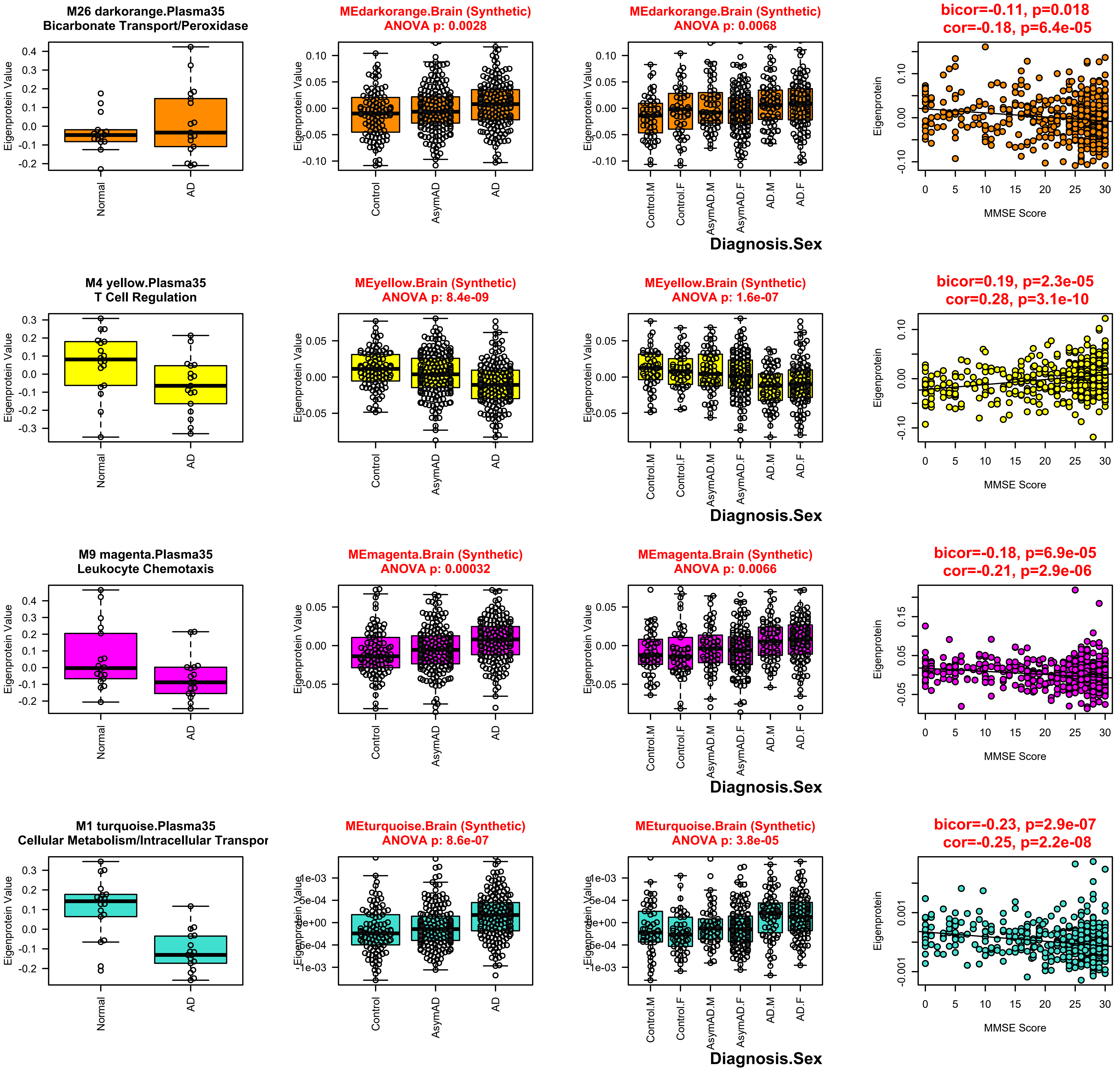
M26 darkorange.Plasma35
Bicarbonate Transport/Peroxidase
MEdarkorange.Brain (Synthetic)
ANOVA p: 0.0028
MEdarkorange.Brain (Synthetic)
ANOVA p: 0.0068
bicor=-0.11, p=0.018
cor=-0.18, p=6.4e-05
M4 yellow.Plasma35
T Cell Regulation
MEyellow.Brain (Synthetic)
ANOVA p: 8.4e-09
MEyellow.Brain (Synthetic)
ANOVA p: 1.6e-07
bicor=0.19, p=2.3e-05
cor=0.28, p=3.1e-10
M9 magenta.Plasma35
Leukocyte Chemotaxis
MEmagenta.Brain (Synthetic)
ANOVA p: 0.00032
MEmagenta.Brain (Synthetic)
ANOVA p: 0.0066
bicor=-0.18, p=6.9e-05
cor=-0.21, p=2.9e-06
M1 turquoise.Plasma35
Cellular Metabolism/Intracellular Transpor
MEturquoise.Brain (Synthetic)
ANOVA p: 8.6e-07
MEturquoise.Brain (Synthetic)
ANOVA p: 3.8e-05
bicor=-0.23, p=2.9e-07
cor=-0.25, p=2.2e-08
Eigenprotein Value
Eigenprotein
Normal
AD
Control
AsymAD
Control.M
Control.F
AsymAD.M
AsymAD.F
AD.M
AD.F
Diagnosis.Sex
MMSE Score

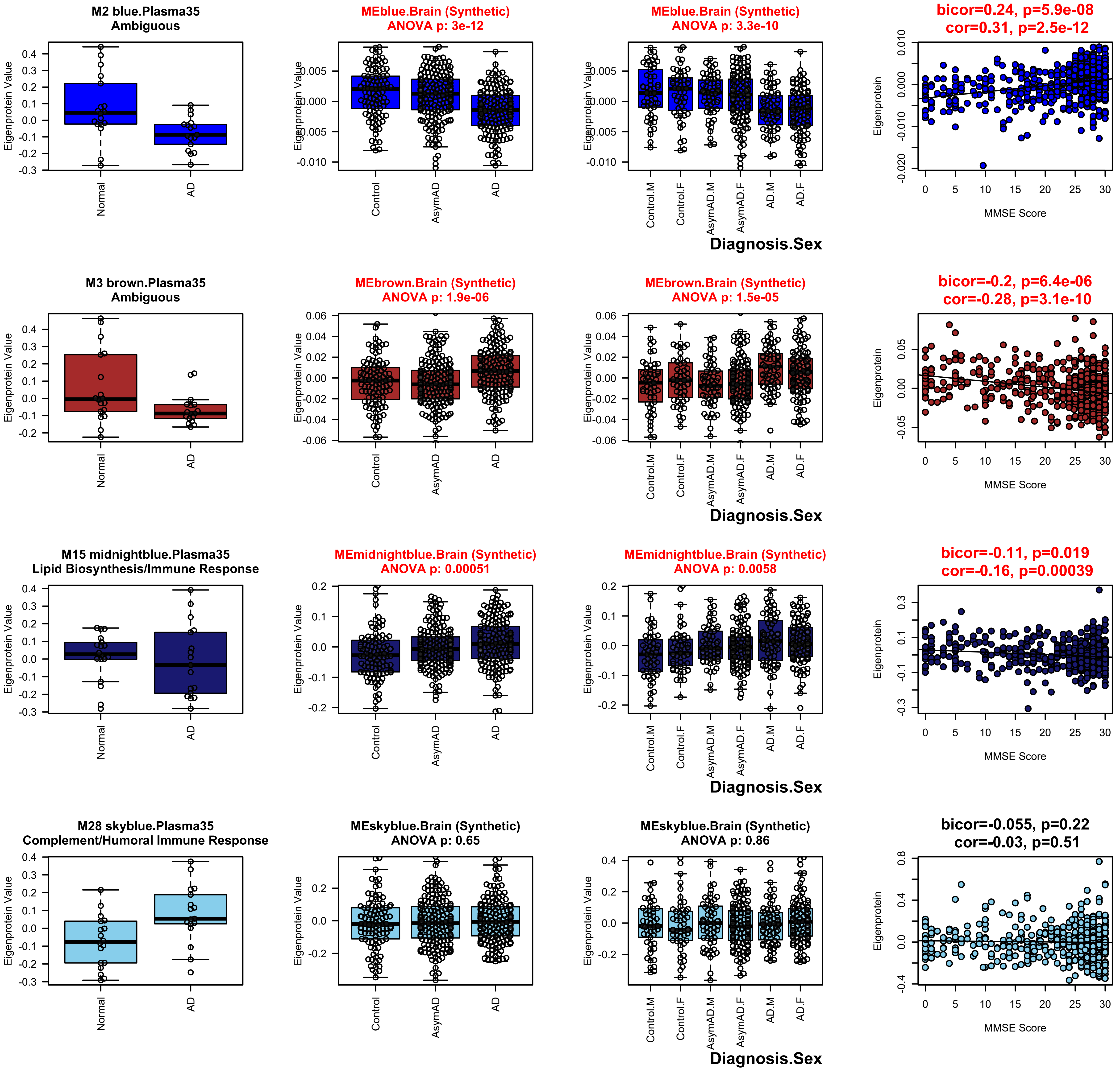

M2 blue.Plasma35
Ambiguous
MEblue.Brain (Synthetic)
ANOVA p: 3e-12
MEblue.Brain (Synthetic)
ANOVA p: 3.3e-10
bicor=0.24, p=5.9e-08
cor=0.31, p=2.5e-12
M3 brown.Plasma35
Ambiguous
MEbrown.Brain (Synthetic)
ANOVA p: 1.9e-06
MEbrown.Brain (Synthetic)
ANOVA p: 1.5e-05
bicor=-0.2, p=6.4e-06
cor=-0.28, p=3.1e-10
M15 midnightblue.Plasma35
Lipid Biosynthesis/Immune Response
MEmidnightblue.Brain (Synthetic)
ANOVA p: 0.00051
MEmidnightblue.Brain (Synthetic)
ANOVA p: 0.0058
bicor=-0.11, p=0.019
cor=-0.16, p=0.00039
M28 skyblue.Plasma35
Complement/Humoral Immune Response
MEskyblue.Brain (Synthetic)
ANOVA p: 0.65
MEskyblue.Brain (Synthetic)
ANOVA p: 0.86
bicor=-0.055, p=0.22
cor=-0.03, p=0.51
Eigenprotein Value
Eigenprotein
Normal
AD
Control
AsymAD
Control.M
Control.F
AsymAD.M
AsymAD.F
AD.M
AD.F
Diagnosis.Sex
MMSE Score

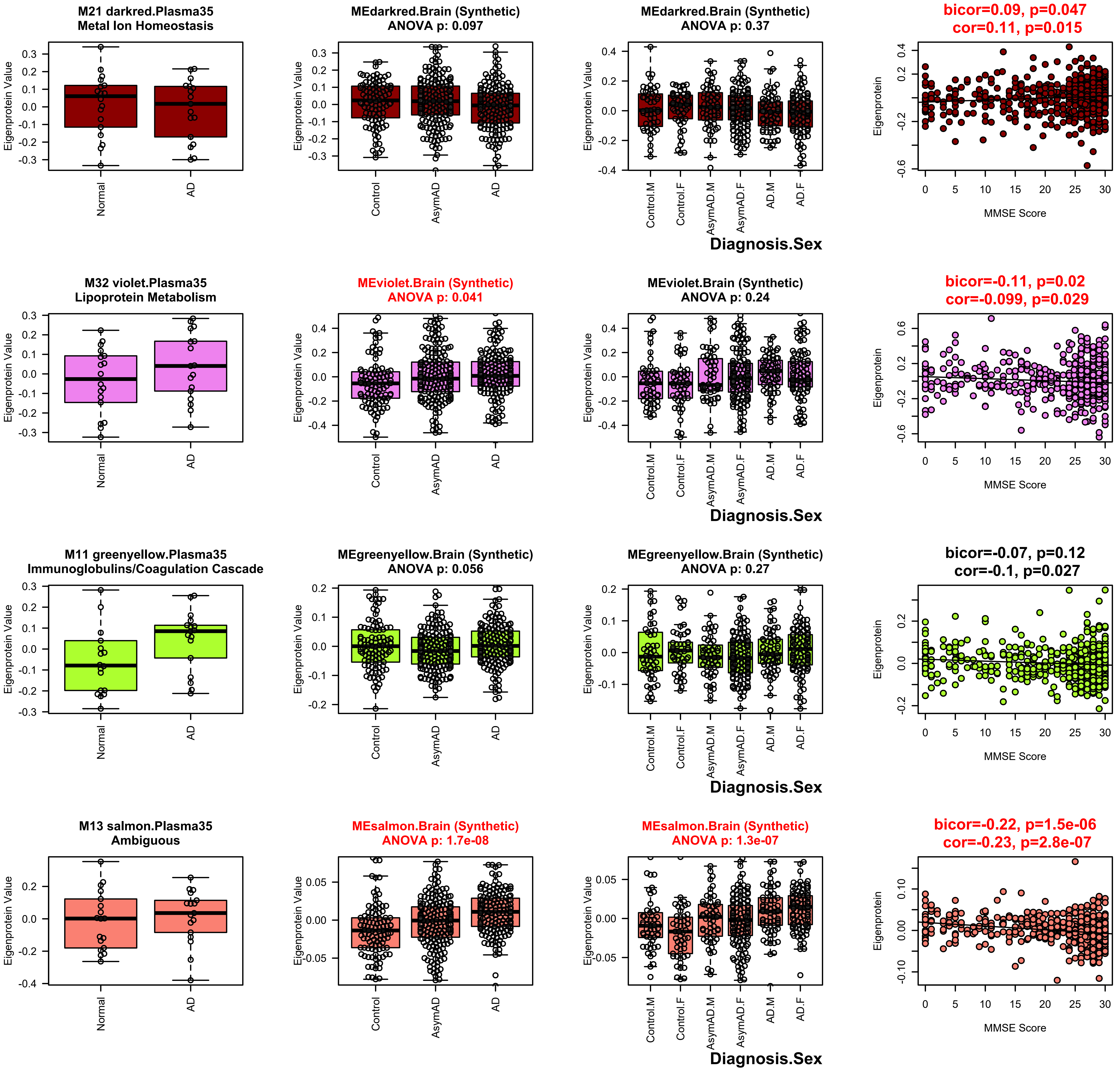
M21 darkred.Plasma35
Metal Ion Homeostasis
MEdarkred.Brain (Synthetic)
ANOVA p: 0.097
MEdarkred.Brain (Synthetic)
ANOVA p: 0.37
bicor=0.09, p=0.047
cor=0.11, p=0.015
M32 violet.Plasma35
Lipoprotein Metabolism
MEviolet.Brain (Synthetic)
ANOVA p: 0.041
MEviolet.Brain (Synthetic)
ANOVA p: 0.24
bicor=-0.11, p=0.02
cor=-0.099, p=0.029
M11 greenyellow.Plasma35
Immunoglobulins/Coagulation Cascade
MEgreenyellow.Brain (Synthetic)
ANOVA p: 0.056
MEgreenyellow.Brain (Synthetic)
ANOVA p: 0.27
bicor=-0.07, p=0.12
cor=-0.1, p=0.027
M13 salmon.Plasma35
Ambiguous
MEsalmon.Brain (Synthetic)
ANOVA p: 1.7e-08
MEsalmon.Brain (Synthetic)
ANOVA p: 1.3e-07
bicor=-0.22, p=1.5e-06
cor=-0.23, p=2.8e-07
Eigenprotein Value
Eigenprotein
Normal
AD
Control
AsymAD
Control.M
Control.F
AsymAD.M
AsymAD.F
AD.M
AD.F
Diagnosis.Sex
MMSE Score

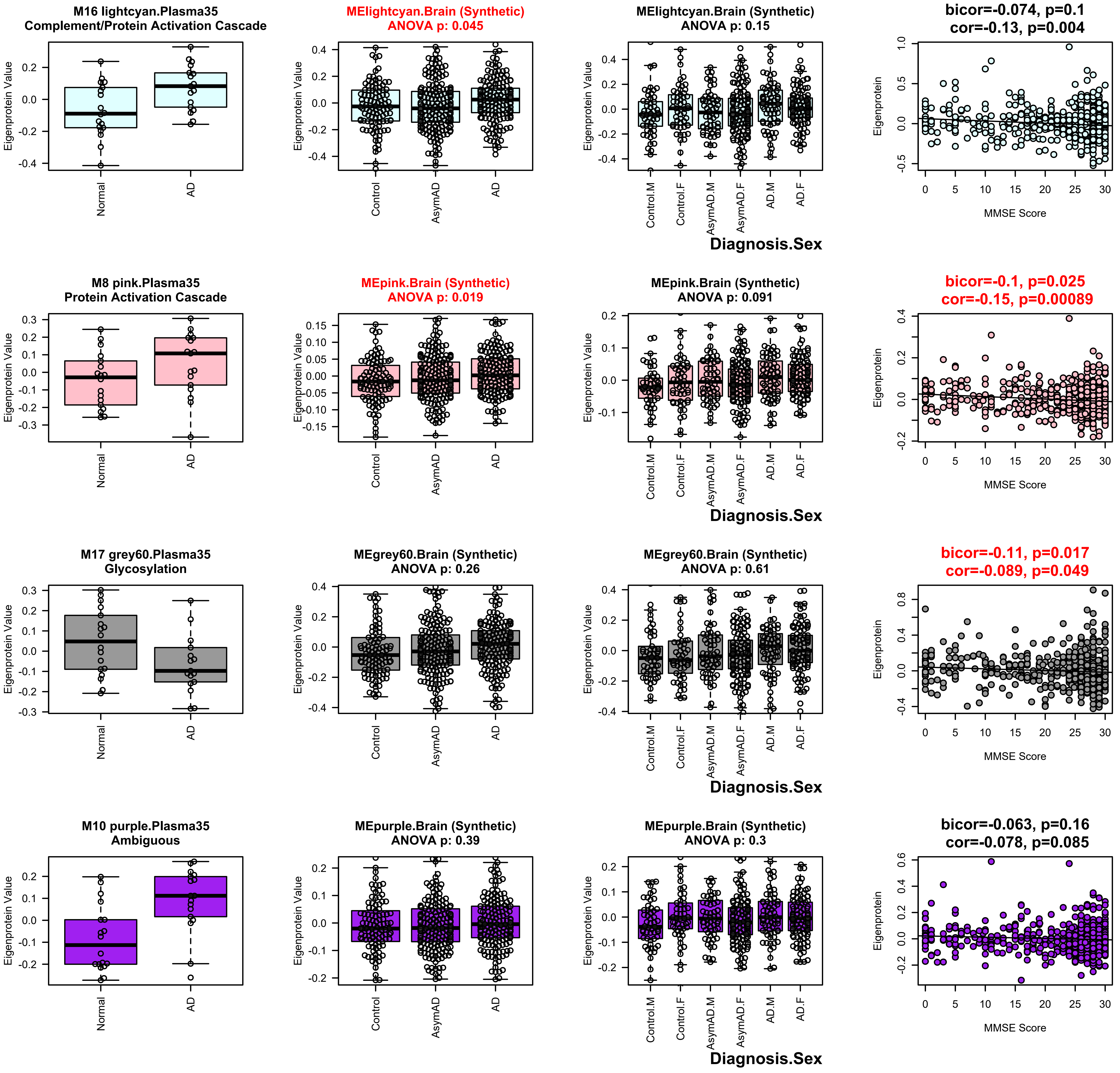
M16 lightcyan.Plasma35
Complement/Protein Activation Cascade
Eigenprotein Value
Normal
AD
MElightcyan.Brain (Synthetic)
ANOVA p: 0.045
Control
AsymAD
AD
MElightcyan.Brain (Synthetic)
ANOVA p: 0.15
Control.M
Control.F
AsymAD.M
AsymAD.F
AD.M
AD.F
Diagnosis.Sex
bicor=-0.074, p=0.1
cor=-0.13, p=0.004
Eigenprotein
MMSE Score
M8 pink.Plasma35
Protein Activation Cascade
MEpink.Brain (Synthetic)
ANOVA p: 0.019
MEpink.Brain (Synthetic)
ANOVA p: 0.091
bicor=-0.1, p=0.025
cor=-0.15, p=0.00089
M17 grey60.Plasma35
Glycosylation
MEgrey60.Brain (Synthetic)
ANOVA p: 0.26
MEgrey60.Brain (Synthetic)
ANOVA p: 0.61
bicor=-0.11, p=0.017
cor=-0.089, p=0.049
M10 purple.Plasma35
Ambiguous
MEpurple.Brain (Synthetic)
ANOVA p: 0.39
MEpurple.Brain (Synthetic)
ANOVA p: 0.3
bicor=-0.063, p=0.16
cor=-0.078, p=0.085

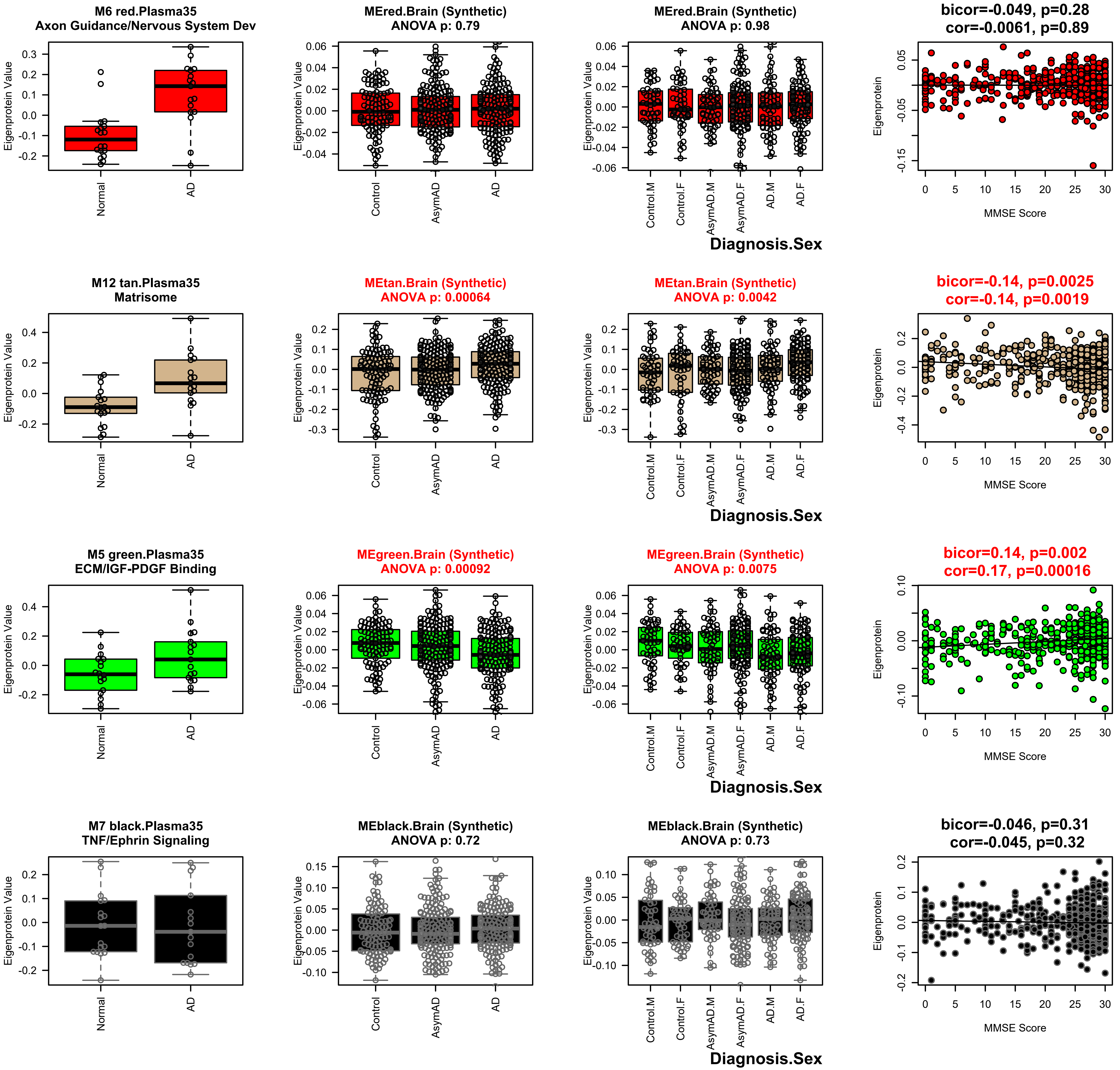

M6 red.Plasma35
Axon Guidance/Nervous System Dev
MEred.Brain (Synthetic)
ANOVA p: 0.79
MEred.Brain (Synthetic)
ANOVA p: 0.98
bicor=-0.049, p=0.28
cor=-0.0061, p=0.89
M12 tan.Plasma35
Matrisome
MEtan.Brain (Synthetic)
ANOVA p: 0.00064
MEtan.Brain (Synthetic)
ANOVA p: 0.0042
bicor=-0.14, p=0.0025
cor=-0.14, p=0.0019
M5 green.Plasma35
ECM/IGF-PDGF Binding
MEgreen.Brain (Synthetic)
ANOVA p: 0.00092
MEgreen.Brain (Synthetic)
ANOVA p: 0.0075
bicor=0.14, p=0.002
cor=0.17, p=0.00016
M7 black.Plasma35
TNF/Ephrin Signaling
MEblack.Brain (Synthetic)
ANOVA p: 0.72
MEblack.Brain (Synthetic)
ANOVA p: 0.73
bicor=-0.046, p=0.31
cor=-0.045, p=0.32
Eigenprotein Value
Eigenprotein
Normal
AD
Control
AsymAD
Control.M
Control.F
AsymAD.M
AsymAD.F
AD.M
AD.F
Diagnosis.Sex
MMSE Score

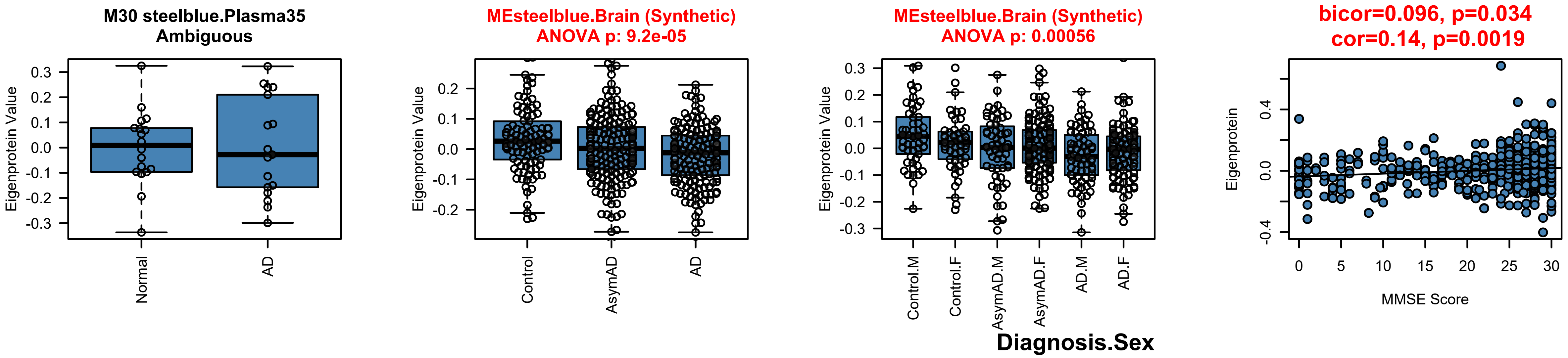
M30 steelblue.Plasma35
Ambiguous
Eigenprotein Value
Normal
AD
MEsteelblue.Brain (Synthetic)
ANOVA p: 9.2e-05
Eigenprotein Value
Control
AsymAD
AD
MEsteelblue.Brain (Synthetic)
ANOVA p: 0.00056
Eigenprotein Value
Control.M
Control.F
AsymAD.M
AsymAD.F
AD.M
AD.F
Diagnosis.Sex
bicor=0.096, p=0.034
cor=0.14, p=0.0019
Eigenprotein
MMSE Score

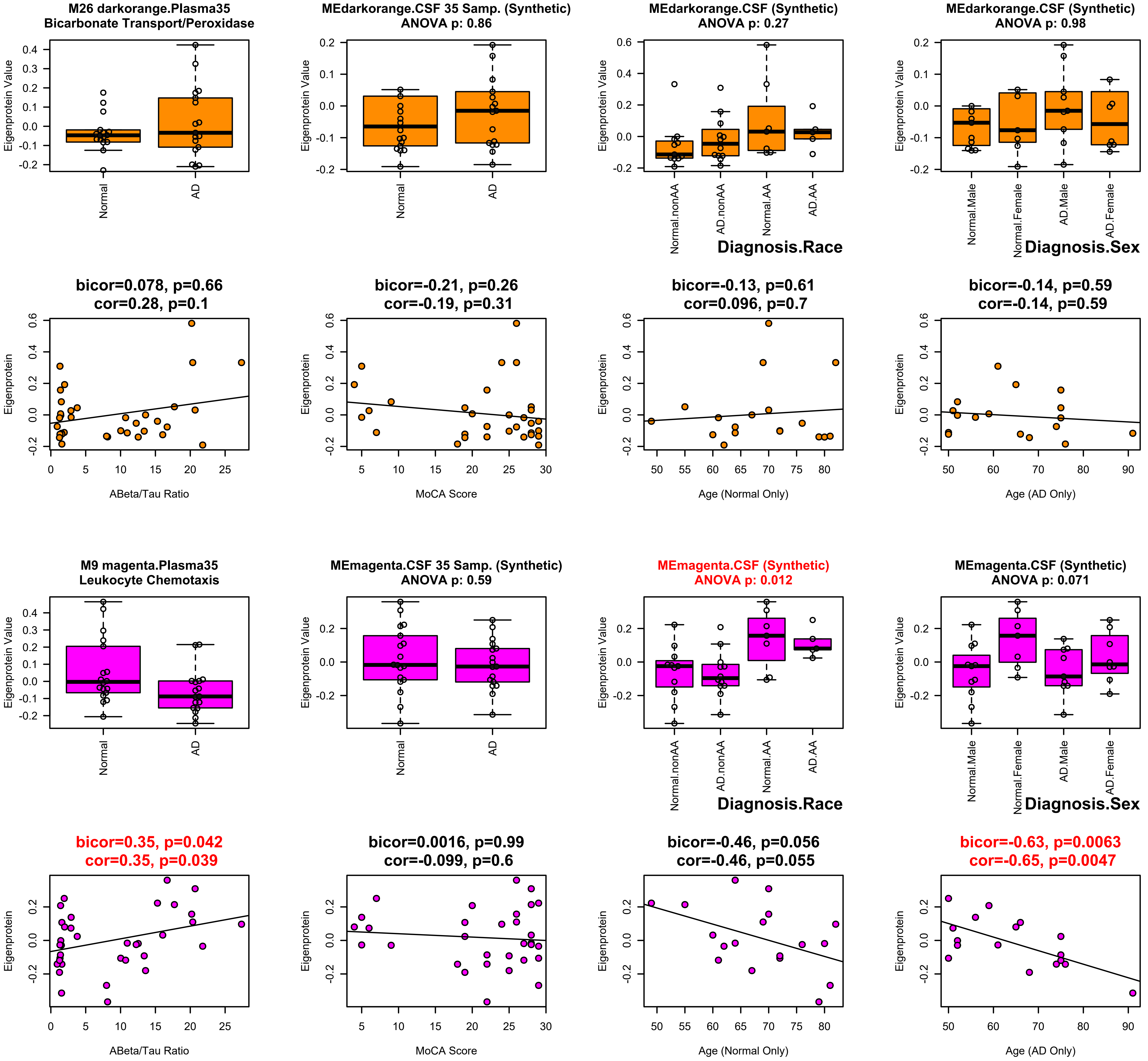
M26 darkorange.Plasma35
Bicarbonate Transport/Peroxidase
MEdarkorange.CSF 35 Samp. (Synthetic)
ANOVA p: 0.86
MEdarkorange.CSF (Synthetic)
ANOVA p: 0.27
MEdarkorange.CSF (Synthetic)
ANOVA p: 0.98
Eigenprotein Value
Normal
AD
Normal.nonAA
AD.nonAA
Normal.AA
AD.AA
Diagnosis.Race
Normal.Male
Normal.Female
AD.Male
AD.Female
Diagnosis.Sex
bicor=0.078, p=0.66
cor=0.28, p=0.1
bicor=-0.21, p=0.26
cor=-0.19, p=0.31
bicor=-0.13, p=0.61
cor=0.096, p=0.7
bicor=-0.14, p=0.59
cor=-0.14, p=0.59
Eigenprotein
ABeta/Tau Ratio
MoCA Score
Age (Normal Only)
Age (AD Only)
M9 magenta.Plasma35
Leukocyte Chemotaxis
MEmagenta.CSF 35 Samp. (Synthetic)
ANOVA p: 0.59
MEmagenta.CSF (Synthetic)
ANOVA p: 0.012
MEmagenta.CSF (Synthetic)
ANOVA p: 0.071
Eigenprotein Value
Normal
AD
Normal.nonAA
AD.nonAA
Normal.AA
AD.AA
Diagnosis.Race
Normal.Male
Normal.Female
AD.Male
AD.Female
Diagnosis.Sex
bicor=0.35, p=0.042
cor=0.35, p=0.039
bicor=0.0016, p=0.99
cor=-0.099, p=0.6
bicor=-0.46, p=0.056
cor=-0.46, p=0.055
bicor=-0.63, p=0.0063
cor=-0.65, p=0.0047
Eigenprotein
ABeta/Tau Ratio
MoCA Score
Age (Normal Only)
Age (AD Only)

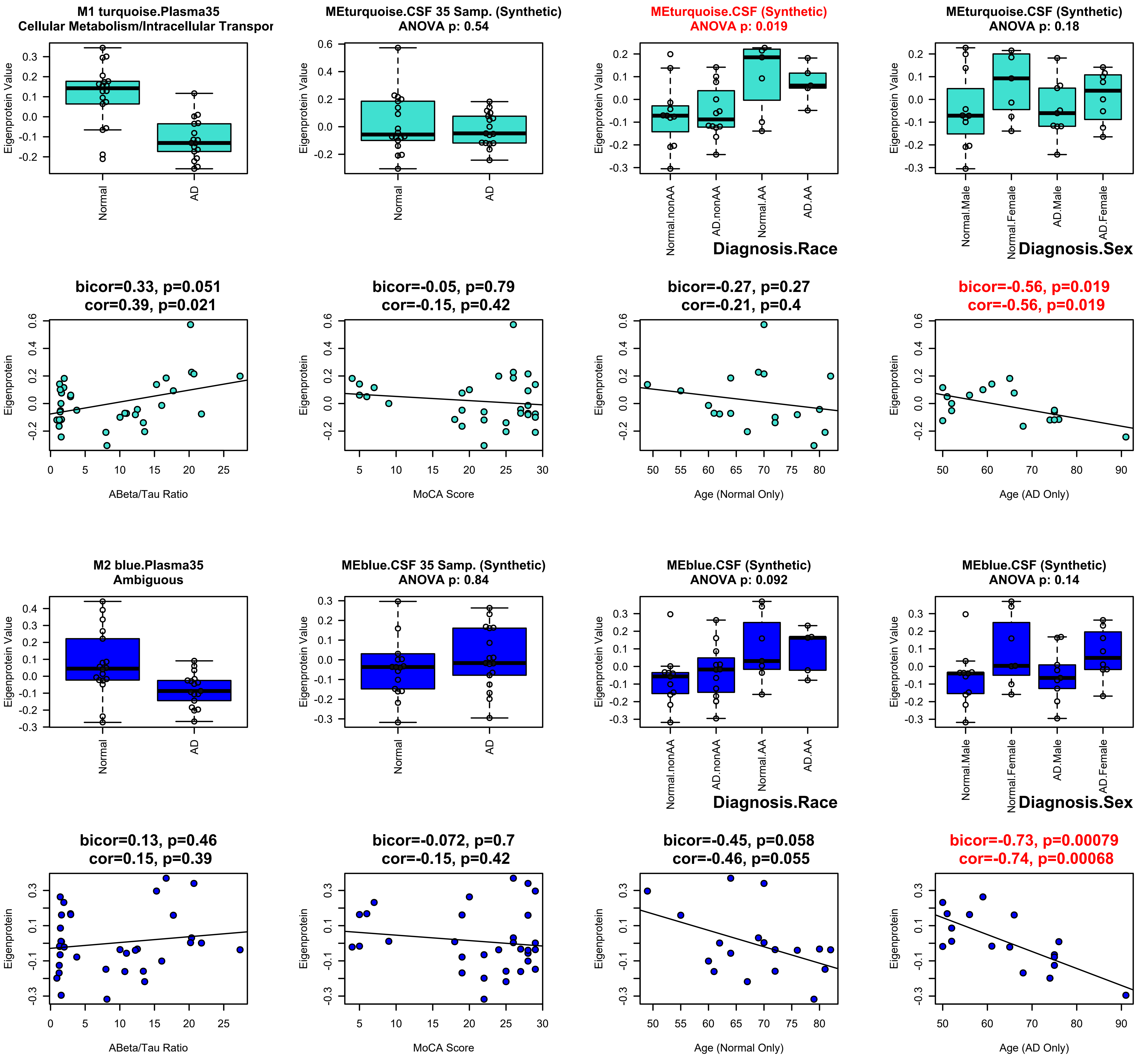

M1 turquoise.Plasma35
Cellular Metabolism/Intracellular Transpor
MEturquoise.CSF 35 Samp. (Synthetic)
ANOVA p: 0.54
MEturquoise.CSF (Synthetic)
ANOVA p: 0.019
MEturquoise.CSF (Synthetic)
ANOVA p: 0.18
Eigenprotein Value
Normal
AD
Normal.nonAA
AD.nonAA
Normal.AA
AD.AA
Diagnosis.Race
Normal.Male
Normal.Female
AD.Male
AD.Female
Diagnosis.Sex
bicor=0.33, p=0.051
cor=0.39, p=0.021
bicor=-0.05, p=0.79
cor=-0.15, p=0.42
bicor=-0.27, p=0.27
cor=-0.21, p=0.4
bicor=-0.56, p=0.019
cor=-0.56, p=0.019
Eigenprotein
ABeta/Tau Ratio
MoCA Score
Age (Normal Only)
Age (AD Only)
M2 blue.Plasma35
Ambiguous
MEblue.CSF 35 Samp. (Synthetic)
ANOVA p: 0.84
MEblue.CSF (Synthetic)
ANOVA p: 0.092
MEblue.CSF (Synthetic)
ANOVA p: 0.14
Eigenprotein Value
Normal
AD
Normal.nonAA
AD.nonAA
Normal.AA
AD.AA
Diagnosis.Race
Normal.Male
Normal.Female
AD.Male
AD.Female
Diagnosis.Sex
bicor=0.13, p=0.46
cor=0.15, p=0.39
bicor=-0.072, p=0.7
cor=-0.15, p=0.42
bicor=-0.45, p=0.058
cor=-0.46, p=0.055
bicor=-0.73, p=0.00079
cor=-0.74, p=0.00068
Eigenprotein
ABeta/Tau Ratio
MoCA Score
Age (Normal Only)
Age (AD Only)

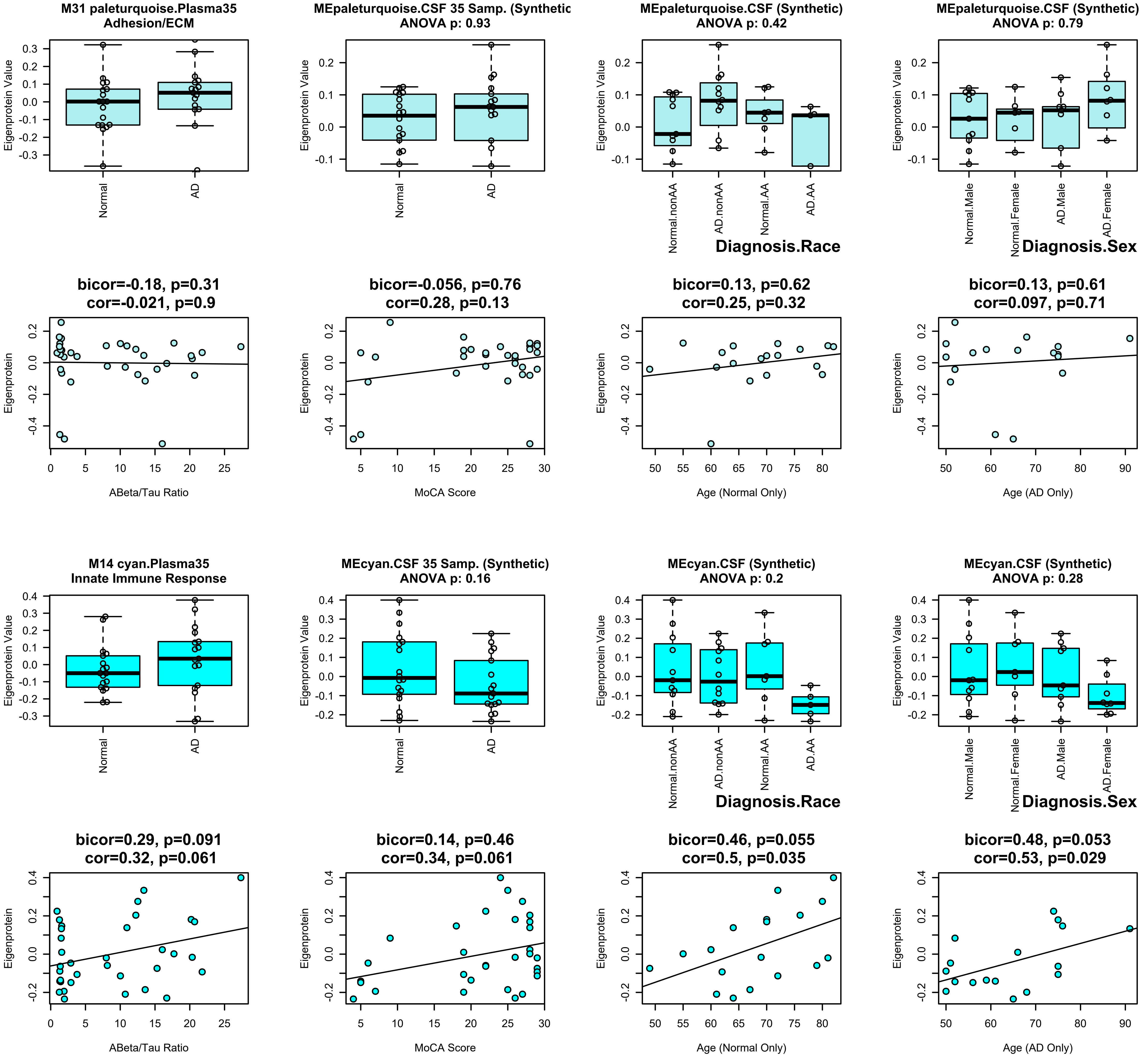
M31 paleturquoise.Plasma35
Adhesion/ECM
MEpaleturquoise.CSF 35 Samp. (Synthetic
ANOVA p: 0.93
MEpaleturquoise.CSF (Synthetic)
ANOVA p: 0.42
MEpaleturquoise.CSF (Synthetic)
ANOVA p: 0.79
Eigenprotein Value
Normal
AD
Normal.nonAA
AD.nonAA
Normal.AA
AD.AA
Diagnosis.Race
Normal.Male
Normal.Female
AD.Male
AD.Female
Diagnosis.Sex
bicor=-0.18, p=0.31
cor=-0.021, p=0.9
bicor=-0.056, p=0.76
cor=0.28, p=0.13
bicor=0.13, p=0.62
cor=0.25, p=0.32
bicor=0.13, p=0.61
cor=0.097, p=0.71
Eigenprotein
ABeta/Tau Ratio
MoCA Score
Age (Normal Only)
Age (AD Only)
M14 cyan.Plasma35
Innate Immune Response
MEcyan.CSF 35 Samp. (Synthetic)
ANOVA p: 0.16
MEcyan.CSF (Synthetic)
ANOVA p: 0.2
MEcyan.CSF (Synthetic)
ANOVA p: 0.28
bicor=0.29, p=0.091
cor=0.32, p=0.061
bicor=0.14, p=0.46
cor=0.34, p=0.061
bicor=0.46, p=0.055
cor=0.5, p=0.035
bicor=0.48, p=0.053
cor=0.53, p=0.029

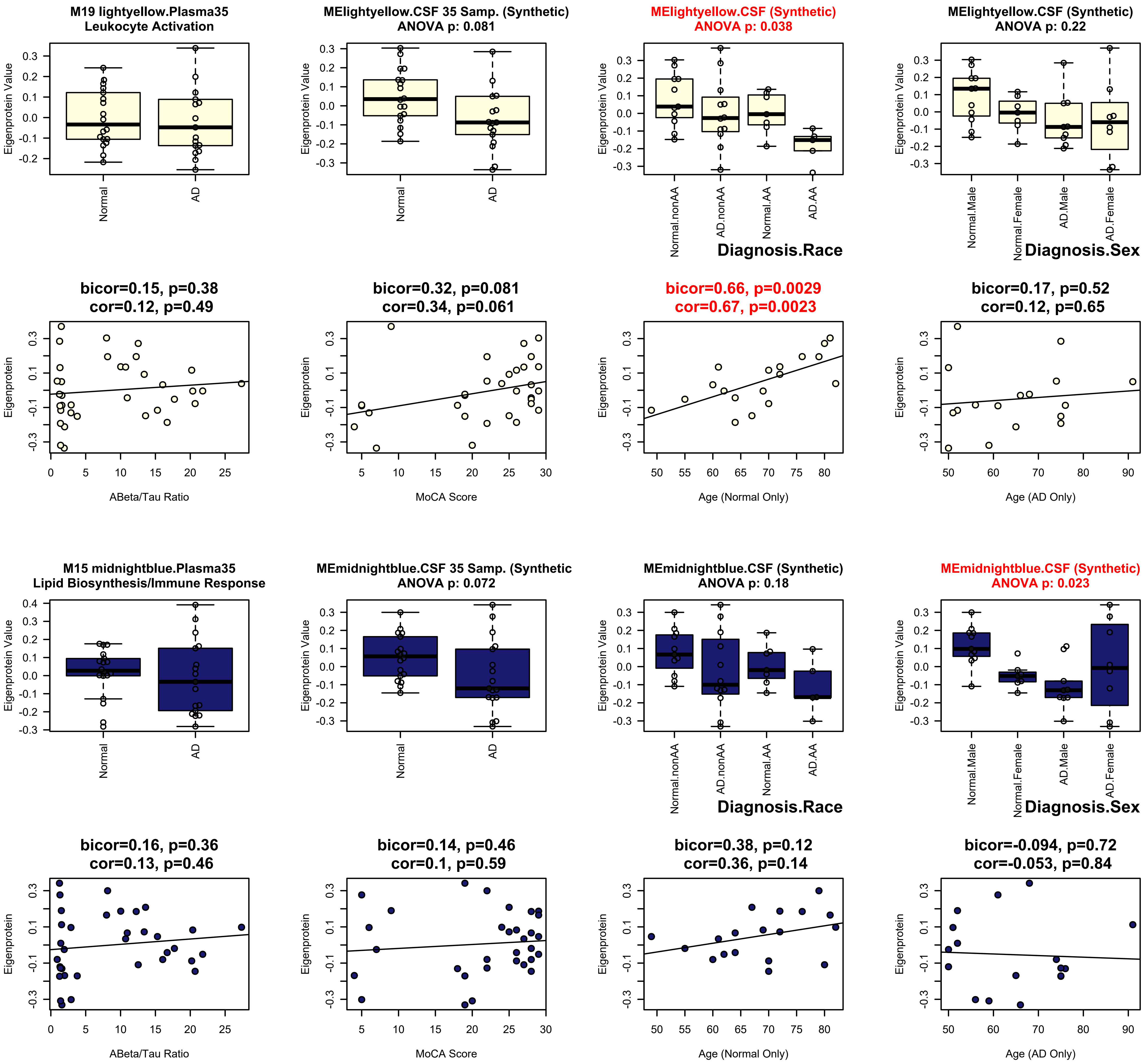
M19 lightyellow.Plasma35
Leukocyte Activation
MElightyellow.CSF 35 Samp. (Synthetic)
ANOVA p: 0.081
MElightyellow.CSF (Synthetic)
ANOVA p: 0.038
MElightyellow.CSF (Synthetic)
ANOVA p: 0.22
Eigenprotein Value
Normal
AD
Normal.nonAA
AD.nonAA
Normal.AA
AD.AA
Diagnosis.Race
Normal.Male
Normal.Female
AD.Male
AD.Female
Diagnosis.Sex
bicor=0.15, p=0.38
cor=0.12, p=0.49
bicor=0.32, p=0.081
cor=0.34, p=0.061
bicor=0.66, p=0.0029
cor=0.67, p=0.0023
bicor=0.17, p=0.52
cor=0.12, p=0.65
Eigenprotein
ABeta/Tau Ratio
MoCA Score
Age (Normal Only)
Age (AD Only)
M15 midnightblue.Plasma35
Lipid Biosynthesis/Immune Response
MEmidnightblue.CSF 35 Samp. (Synthetic
ANOVA p: 0.072
MEmidnightblue.CSF (Synthetic)
ANOVA p: 0.18
MEmidnightblue.CSF (Synthetic)
ANOVA p: 0.023
bicor=0.16, p=0.36
cor=0.13, p=0.46
bicor=0.14, p=0.46
cor=0.1, p=0.59
bicor=0.38, p=0.12
cor=0.36, p=0.14
bicor=-0.094, p=0.72
cor=-0.053, p=0.84

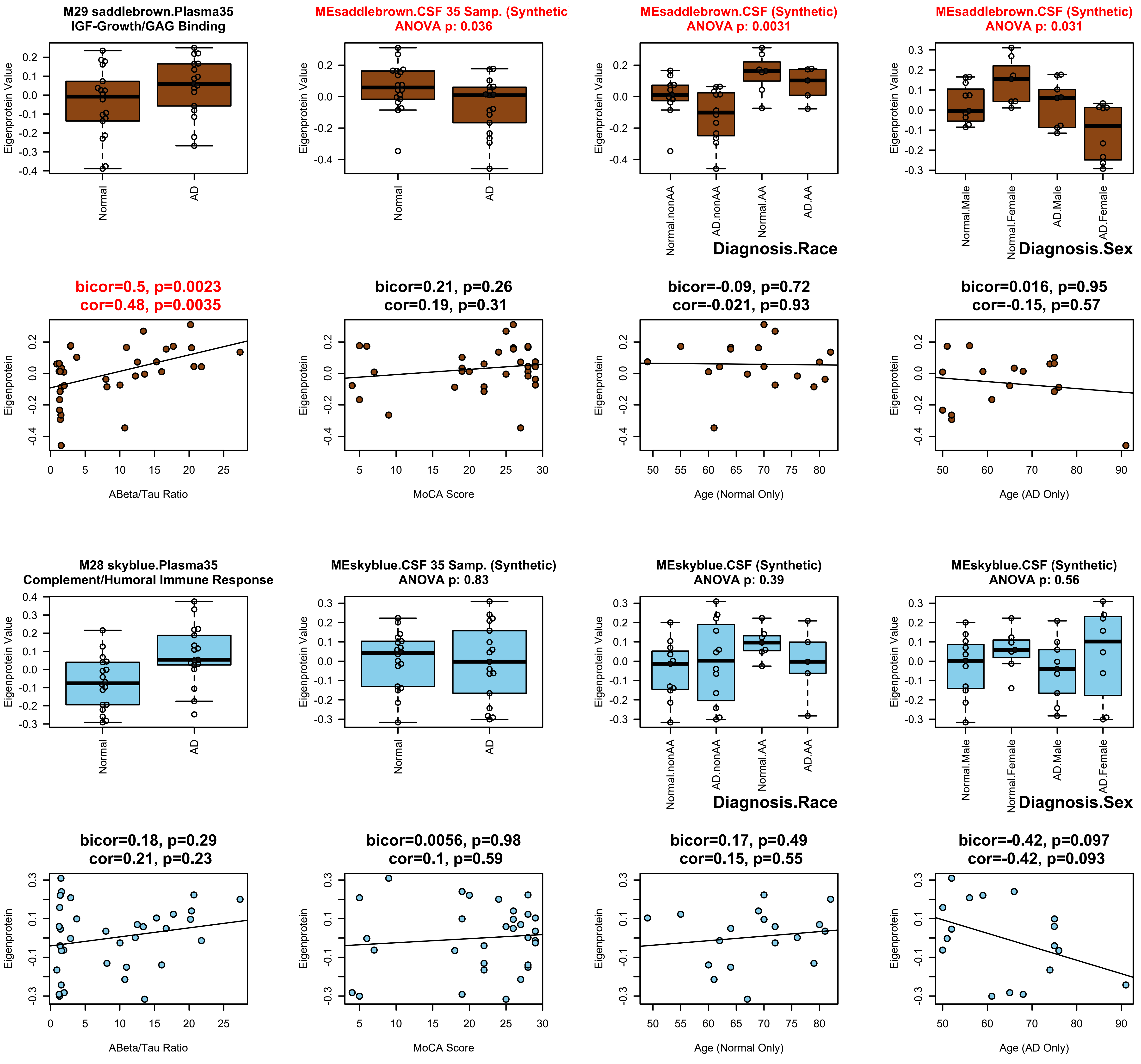
M29 saddlebrown.Plasma35
IGF-Growth/GAG Binding
MEsaddlebrown.CSF 35 Samp. (Synthetic
ANOVA p: 0.036
MEsaddlebrown.CSF (Synthetic)
ANOVA p: 0.0031
MEsaddlebrown.CSF (Synthetic)
ANOVA p: 0.031
Eigenprotein Value
Normal
AD
Normal.nonAA
AD.nonAA
Normal.AA
AD.AA
Diagnosis.Race
Normal.Male
Normal.Female
AD.Male
AD.Female
Diagnosis.Sex
bicor=0.5, p=0.0023
cor=0.48, p=0.0035
bicor=0.21, p=0.26
cor=0.19, p=0.31
bicor=-0.09, p=0.72
cor=-0.021, p=0.93
bicor=0.016, p=0.95
cor=-0.15, p=0.57
Eigenprotein
ABeta/Tau Ratio
MoCA Score
Age (Normal Only)
Age (AD Only)
M28 skyblue.Plasma35
Complement/Humoral Immune Response
MEskyblue.CSF 35 Samp. (Synthetic)
ANOVA p: 0.83
MEskyblue.CSF (Synthetic)
ANOVA p: 0.39
MEskyblue.CSF (Synthetic)
ANOVA p: 0.56
bicor=0.18, p=0.29
cor=0.21, p=0.23
bicor=0.0056, p=0.98
cor=0.1, p=0.59
bicor=0.17, p=0.49
cor=0.15, p=0.55
bicor=-0.42, p=0.097
cor=-0.42, p=0.093

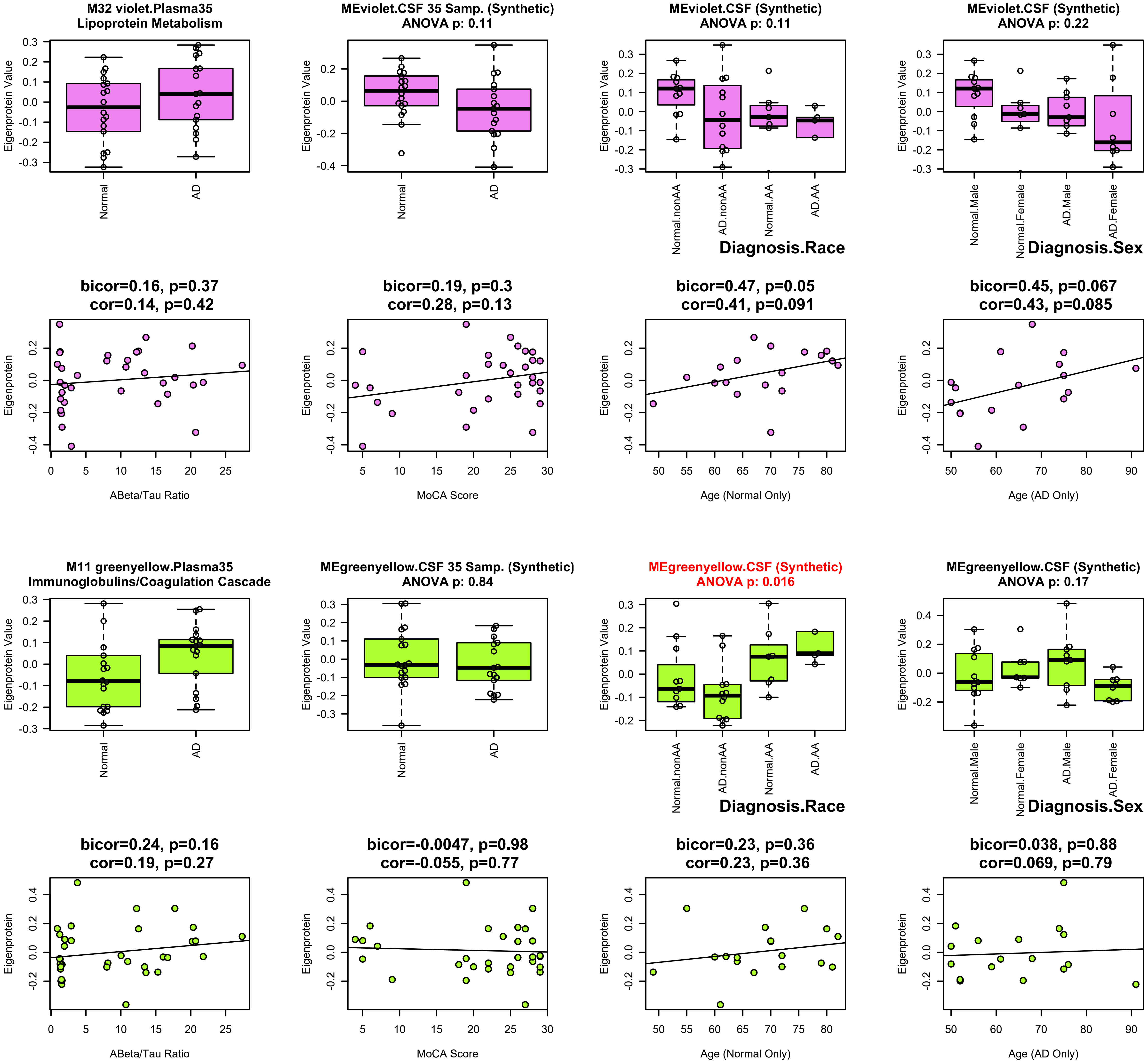

M32 violet.Plasma35
Lipoprotein Metabolism
MEviolet.CSF 35 Samp. (Synthetic)
ANOVA p: 0.11
MEviolet.CSF (Synthetic)
ANOVA p: 0.11
MEviolet.CSF (Synthetic)
ANOVA p: 0.22
Eigenprotein Value
Normal
AD
Normal.nonAA
AD.nonAA
Normal.AA
AD.AA
Diagnosis.Race
Normal.Male
Normal.Female
AD.Male
AD.Female
Diagnosis.Sex
bicor=0.16, p=0.37
cor=0.14, p=0.42
bicor=0.19, p=0.3
cor=0.28, p=0.13
bicor=0.47, p=0.05
cor=0.41, p=0.091
bicor=0.45, p=0.067
cor=0.43, p=0.085
Eigenprotein
ABeta/Tau Ratio
MoCA Score
Age (Normal Only)
Age (AD Only)
M11 greenyellow.Plasma35
Immunoglobulins/Coagulation Cascade
MEgreenyellow.CSF 35 Samp. (Synthetic)
ANOVA p: 0.84
MEgreenyellow.CSF (Synthetic)
ANOVA p: 0.016
MEgreenyellow.CSF (Synthetic)
ANOVA p: 0.17
Eigenprotein Value
Normal
AD
Normal.nonAA
AD.nonAA
Normal.AA
AD.AA
Diagnosis.Race
Normal.Male
Normal.Female
AD.Male
AD.Female
Diagnosis.Sex
bicor=0.24, p=0.16
cor=0.19, p=0.27
bicor=-0.0047, p=0.98
cor=-0.055, p=0.77
bicor=0.23, p=0.36
cor=0.23, p=0.36
bicor=0.038, p=0.88
cor=0.069, p=0.79
Eigenprotein
ABeta/Tau Ratio
MoCA Score
Age (Normal Only)
Age (AD Only)

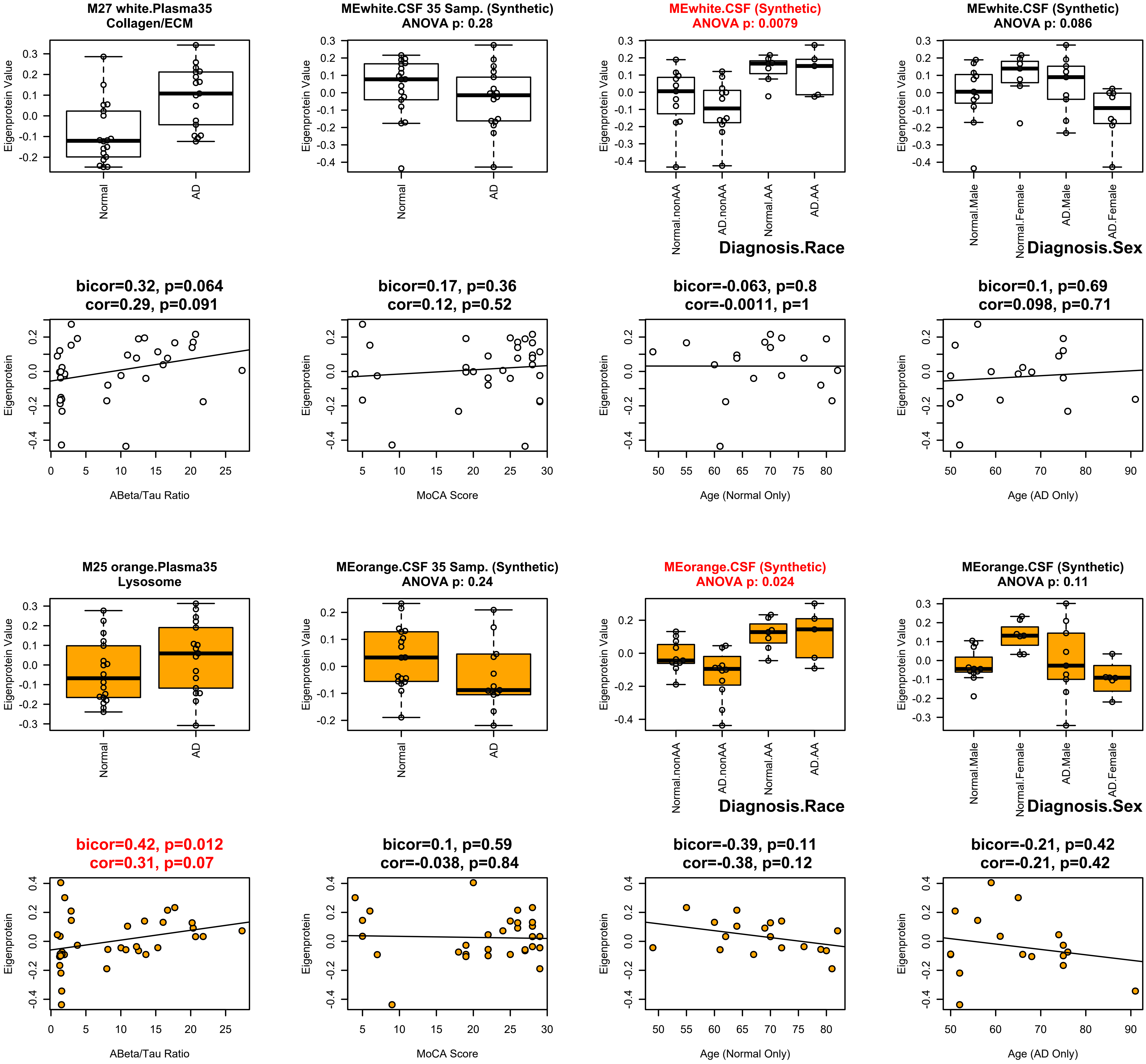
M27 white.Plasma35
Collagen/ECM
Eigenprotein Value
Normal
AD
MEwhite.CSF 35 Samp. (Synthetic)
ANOVA p: 0.28
Eigenprotein Value
Normal
AD
MEwhite.CSF (Synthetic)
ANOVA p: 0.0079
Eigenprotein Value
Normal.nonAA
AD.nonAA
Normal.AA
AD.AA
Diagnosis.Race
MEwhite.CSF (Synthetic)
ANOVA p: 0.086
Eigenprotein Value
Normal.Male
Normal.Female
AD.Male
AD.Female
Diagnosis.Sex
bicor=0.32, p=0.064
cor=0.29, p=0.091
Eigenprotein
ABeta/Tau Ratio
bicor=0.17, p=0.36
cor=0.12, p=0.52
Eigenprotein
MoCA Score
bicor=-0.063, p=0.8
cor=-0.0011, p=1
Eigenprotein
Age (Normal Only)
bicor=0.1, p=0.69
cor=0.098, p=0.71
Eigenprotein
Age (AD Only)
M25 orange.Plasma35
Lysosome
Eigenprotein Value
Normal
AD
MEorange.CSF 35 Samp. (Synthetic)
ANOVA p: 0.24
Eigenprotein Value
Normal
AD
MEorange.CSF (Synthetic)
ANOVA p: 0.024
Eigenprotein Value
Normal.nonAA
AD.nonAA
Normal.AA
AD.AA
Diagnosis.Race
MEorange.CSF (Synthetic)
ANOVA p: 0.11
Eigenprotein Value
Normal.Male
Normal.Female
AD.Male
AD.Female
Diagnosis.Sex
bicor=0.42, p=0.012
cor=0.31, p=0.07
Eigenprotein
ABeta/Tau Ratio
bicor=0.1, p=0.59
cor=-0.038, p=0.84
Eigenprotein
MoCA Score
bicor=-0.39, p=0.11
cor=-0.38, p=0.12
Eigenprotein
Age (Normal Only)
bicor=-0.21, p=0.42
cor=-0.21, p=0.42
Eigenprotein
Age (AD Only)

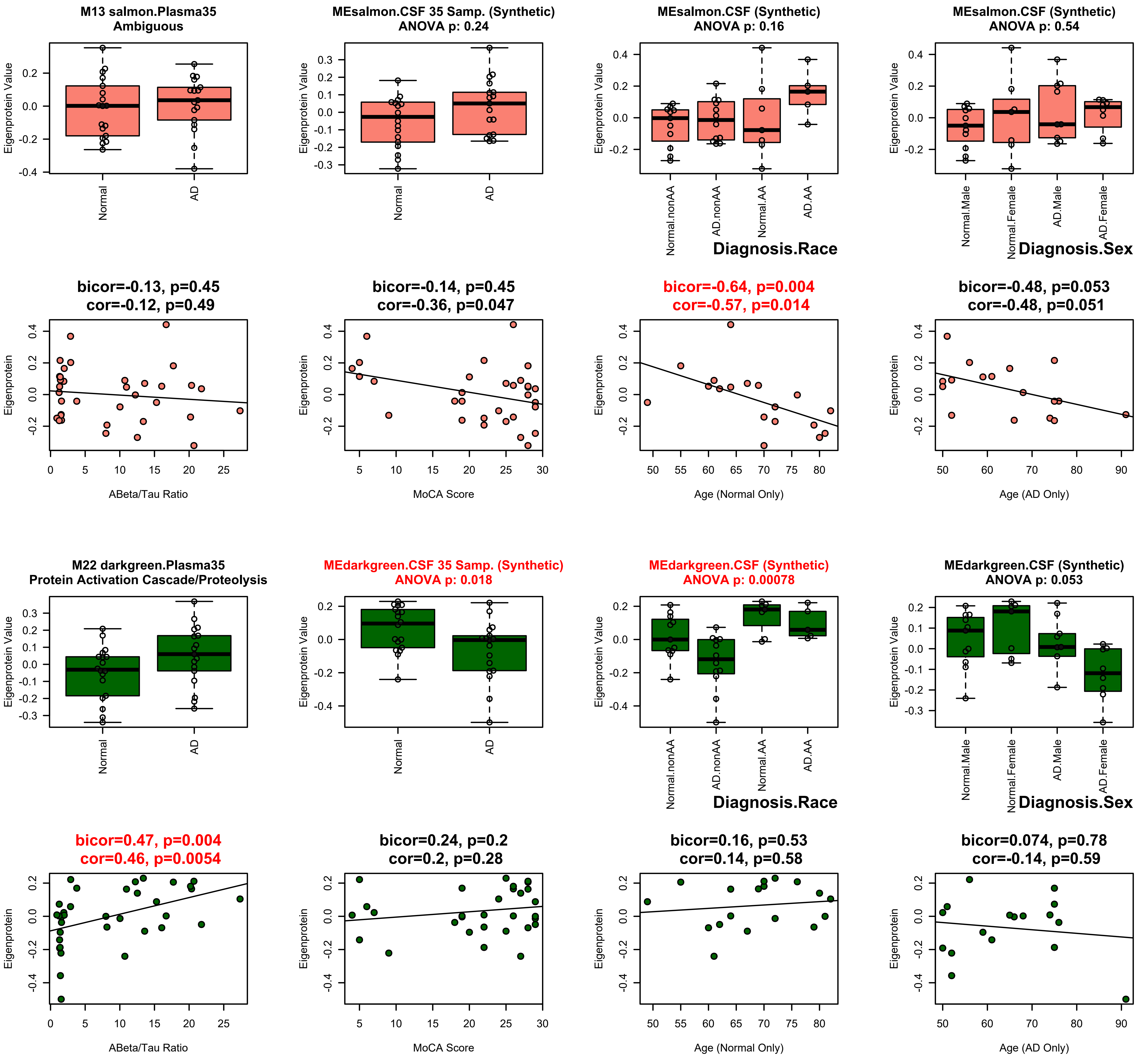

M13 salmon.Plasma35
Ambiguous
Eigenprotein Value
Normal
AD
MEsalmon.CSF 35 Samp. (Synthetic)
ANOVA p: 0.24
Eigenprotein Value
Normal
AD
MEsalmon.CSF (Synthetic)
ANOVA p: 0.16
Eigenprotein Value
Normal.nonAA
AD.nonAA
Normal.AA
AD.AA
Diagnosis.Race
MEsalmon.CSF (Synthetic)
ANOVA p: 0.54
Eigenprotein Value
Normal.Male
Normal.Female
AD.Male
AD.Female
Diagnosis.Sex
bicor=-0.13, p=0.45
cor=-0.12, p=0.49
Eigenprotein
ABeta/Tau Ratio
bicor=-0.14, p=0.45
cor=-0.36, p=0.047
Eigenprotein
MoCA Score
bicor=-0.64, p=0.004
cor=-0.57, p=0.014
Eigenprotein
Age (Normal Only)
bicor=-0.48, p=0.053
cor=-0.48, p=0.051
Eigenprotein
Age (AD Only)
M22 darkgreen.Plasma35
Protein Activation Cascade/Proteolysis
Eigenprotein Value
Normal
AD
MEdarkgreen.CSF 35 Samp. (Synthetic)
ANOVA p: 0.018
Eigenprotein Value
Normal
AD
MEdarkgreen.CSF (Synthetic)
ANOVA p: 0.00078
Eigenprotein Value
Normal.nonAA
AD.nonAA
Normal.AA
AD.AA
Diagnosis.Race
MEdarkgreen.CSF (Synthetic)
ANOVA p: 0.053
Eigenprotein Value
Normal.Male
Normal.Female
AD.Male
AD.Female
Diagnosis.Sex
bicor=0.47, p=0.004
cor=0.46, p=0.0054
Eigenprotein
ABeta/Tau Ratio
bicor=0.24, p=0.2
cor=0.2, p=0.28
Eigenprotein
MoCA Score
bicor=0.16, p=0.53
cor=0.14, p=0.58
Eigenprotein
Age (Normal Only)
bicor=0.074, p=0.78
cor=-0.14, p=0.59
Eigenprotein
Age (AD Only)

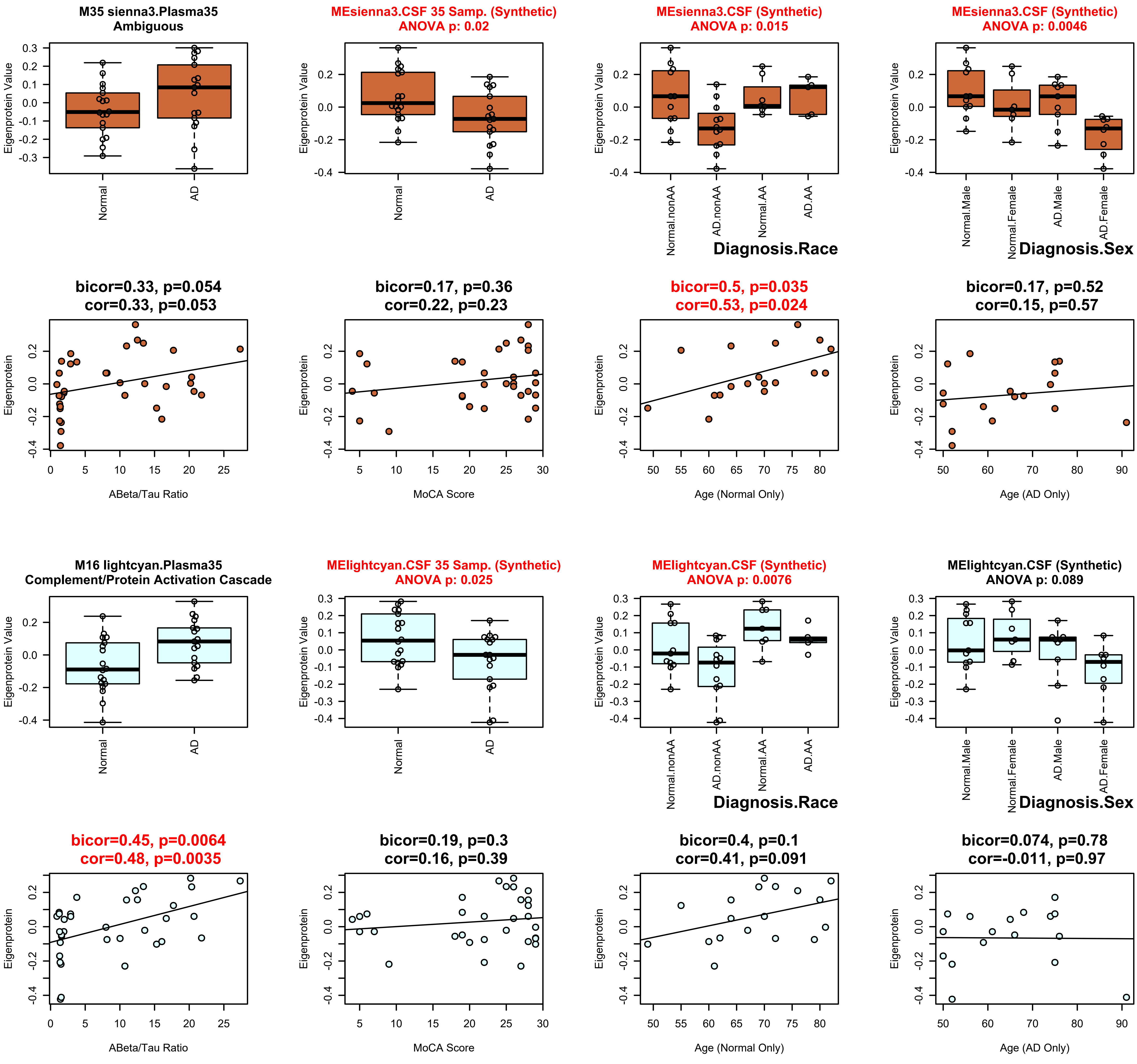

M35 sienna3.Plasma35
Ambiguous
MEsienna3.CSF 35 Samp. (Synthetic)
ANOVA p: 0.02
MEsienna3.CSF (Synthetic)
ANOVA p: 0.015
MEsienna3.CSF (Synthetic)
ANOVA p: 0.0046
Eigenprotein Value
Normal
AD
Normal.nonAA
AD.nonAA
Normal.AA
AD.AA
Diagnosis.Race
Normal.Male
Normal.Female
AD.Male
AD.Female
Diagnosis.Sex
bicor=0.33, p=0.054
cor=0.33, p=0.053
bicor=0.17, p=0.36
cor=0.22, p=0.23
bicor=0.5, p=0.035
cor=0.53, p=0.024
bicor=0.17, p=0.52
cor=0.15, p=0.57
Eigenprotein
ABeta/Tau Ratio
MoCA Score
Age (Normal Only)
Age (AD Only)
M16 lightcyan.Plasma35
Complement/Protein Activation Cascade
MElightcyan.CSF 35 Samp. (Synthetic)
ANOVA p: 0.025
MElightcyan.CSF (Synthetic)
ANOVA p: 0.0076
MElightcyan.CSF (Synthetic)
ANOVA p: 0.089
Eigenprotein Value
Normal
AD
Normal.nonAA
AD.nonAA
Normal.AA
AD.AA
Diagnosis.Race
Normal.Male
Normal.Female
AD.Male
AD.Female
Diagnosis.Sex
bicor=0.45, p=0.0064
cor=0.48, p=0.0035
bicor=0.19, p=0.3
cor=0.16, p=0.39
bicor=0.4, p=0.1
cor=0.41, p=0.091
bicor=0.074, p=0.78
cor=-0.011, p=0.97
Eigenprotein
ABeta/Tau Ratio
MoCA Score
Age (Normal Only)
Age (AD Only)

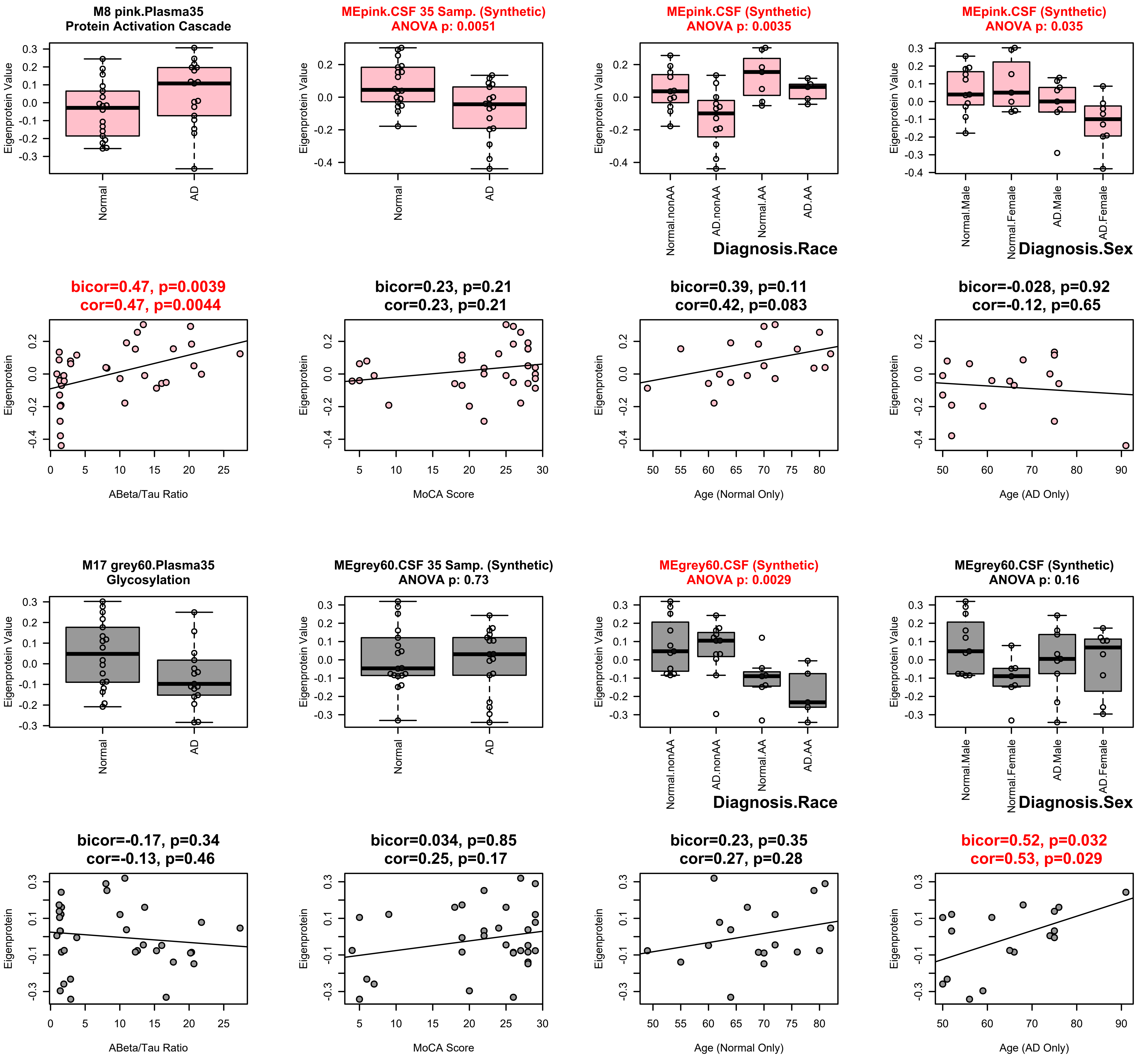

M8 pink.Plasma35
Protein Activation Cascade
Eigenprotein Value
Normal
AD
MEpink.CSF 35 Samp. (Synthetic)
ANOVA p: 0.0051
MEpink.CSF (Synthetic)
ANOVA p: 0.0035
Normal.nonAA
AD.nonAA
Normal.AA
AD.AA
Diagnosis.Race
MEpink.CSF (Synthetic)
ANOVA p: 0.035
Normal.Male
Normal.Female
AD.Male
AD.Female
Diagnosis.Sex
bicor=0.47, p=0.0039
cor=0.47, p=0.0044
Eigenprotein
ABeta/Tau Ratio
bicor=0.23, p=0.21
cor=0.23, p=0.21
MoCA Score
bicor=0.39, p=0.11
cor=0.42, p=0.083
Age (Normal Only)
bicor=-0.028, p=0.92
cor=-0.12, p=0.65
Age (AD Only)
M17 grey60.Plasma35
Glycosylation
MEgrey60.CSF 35 Samp. (Synthetic)
ANOVA p: 0.73
MEgrey60.CSF (Synthetic)
ANOVA p: 0.0029
MEgrey60.CSF (Synthetic)
ANOVA p: 0.16
bicor=-0.17, p=0.34
cor=-0.13, p=0.46
bicor=0.034, p=0.85
cor=0.25, p=0.17
bicor=0.23, p=0.35
cor=0.27, p=0.28
bicor=0.52, p=0.032
cor=0.53, p=0.029

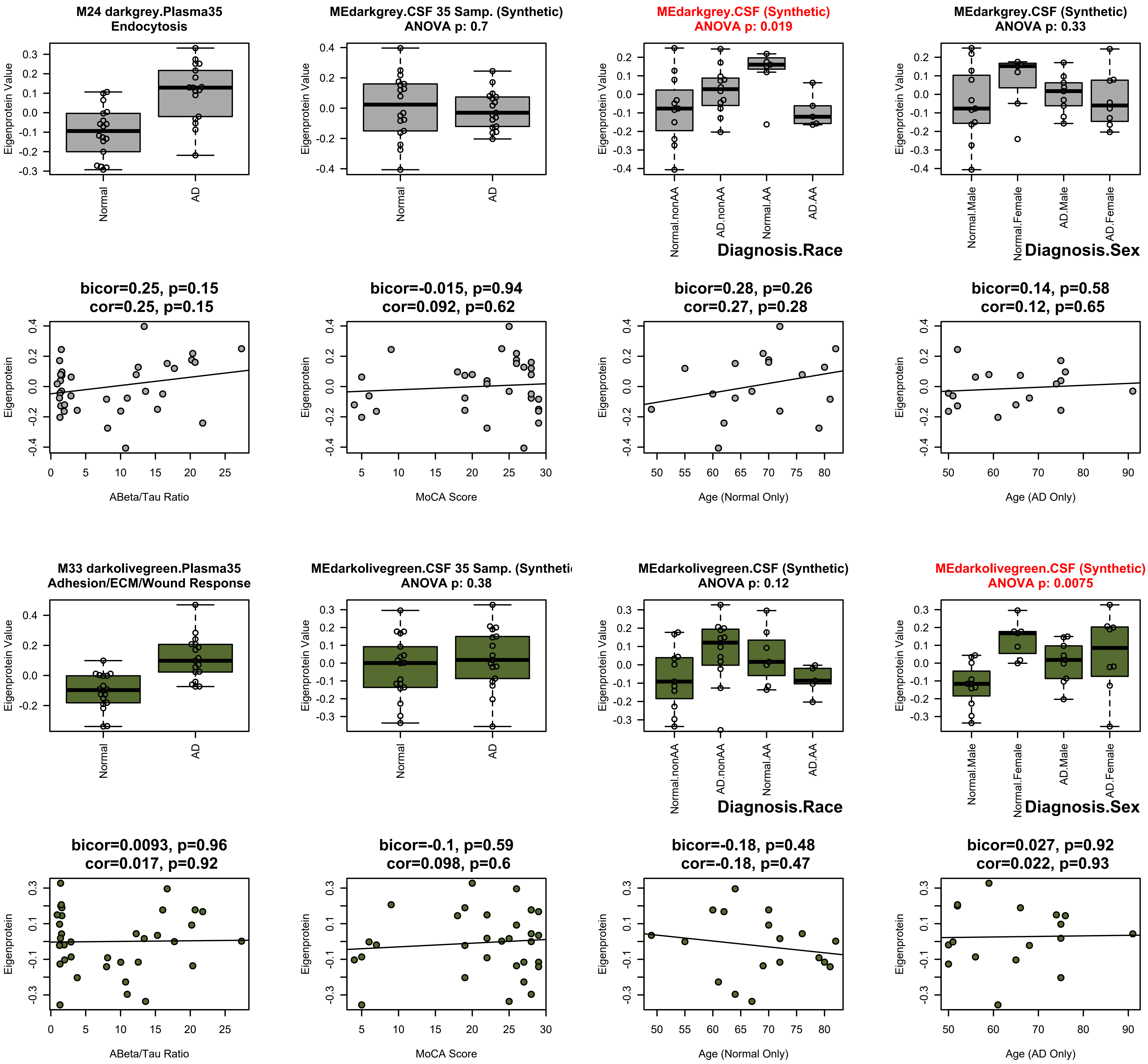
M24 darkgrey.Plasma35
Endocytosis
Eigenprotein Value
Normal
AD
MEdarkgrey.CSF 35 Samp. (Synthetic)
ANOVA p: 0.7
MEdarkgrey.CSF (Synthetic)
ANOVA p: 0.019
Normal.nonAA
AD.nonAA
Normal.AA
AD.AA
Diagnosis.Race
MEdarkgrey.CSF (Synthetic)
ANOVA p: 0.33
Normal.Male
Normal.Female
AD.Male
AD.Female
Diagnosis.Sex
bicor=0.25, p=0.15
cor=0.25, p=0.15
Eigenprotein
ABeta/Tau Ratio
bicor=-0.015, p=0.94
cor=0.092, p=0.62
MoCA Score
bicor=0.28, p=0.26
cor=0.27, p=0.28
Age (Normal Only)
bicor=0.14, p=0.58
cor=0.12, p=0.65
Age (AD Only)
M33 darkolivegreen.Plasma35
Adhesion/ECM/Wound Response
MEdarkolivegreen.CSF 35 Samp. (Syntheti
ANOVA p: 0.38
MEdarkolivegreen.CSF (Synthetic)
ANOVA p: 0.12
MEdarkolivegreen.CSF (Synthetic)
ANOVA p: 0.0075
bicor=0.0093, p=0.96
cor=0.017, p=0.92
bicor=-0.1, p=0.59
cor=0.098, p=0.6
bicor=-0.18, p=0.48
cor=-0.18, p=0.47
bicor=0.027, p=0.92
cor=0.022, p=0.93

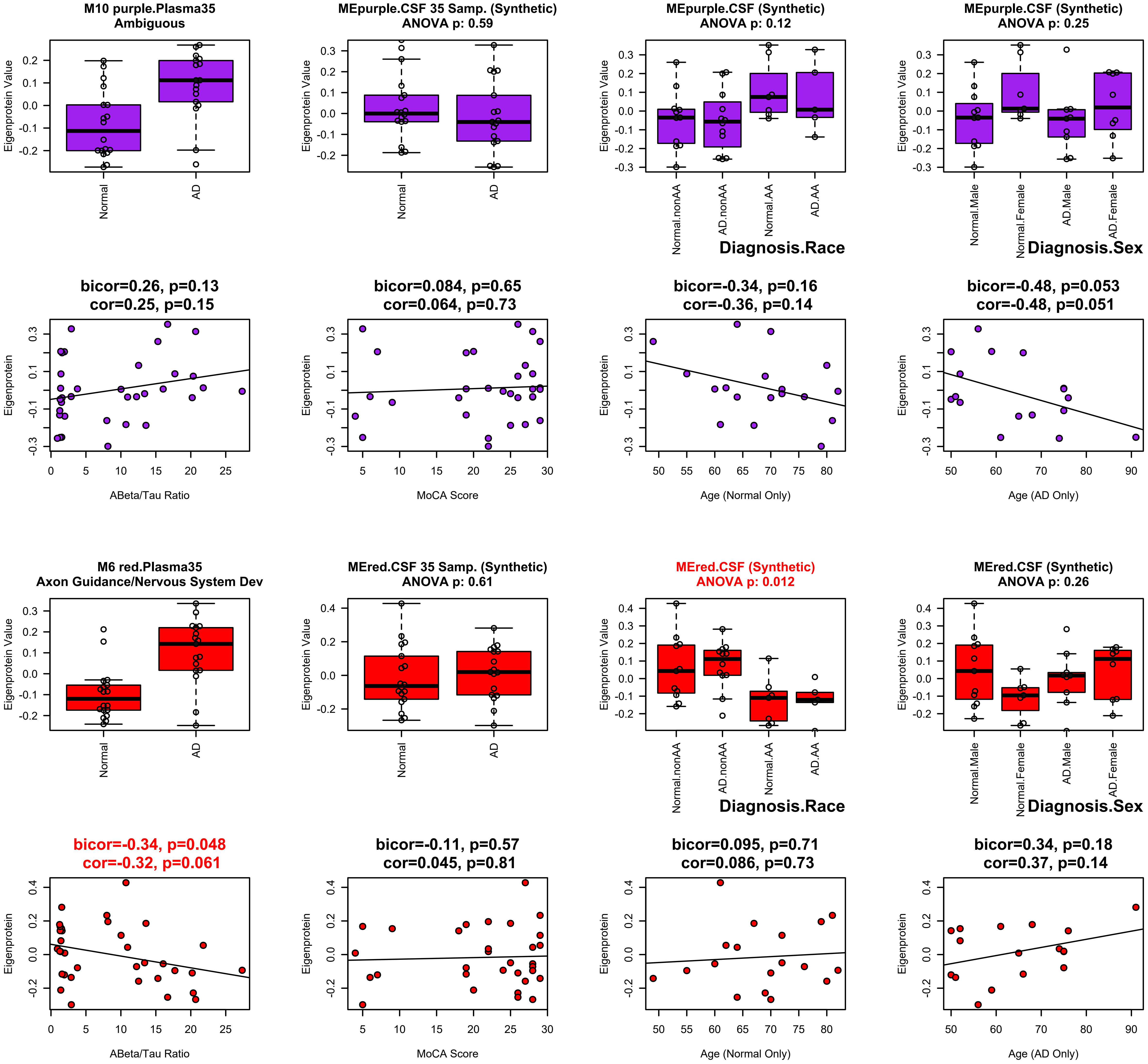
M10 purple.Plasma35
Ambiguous
Eigenprotein Value
Normal
AD
MEpurple.CSF 35 Samp. (Synthetic)
ANOVA p: 0.59
Eigenprotein Value
Normal
AD
MEpurple.CSF (Synthetic)
ANOVA p: 0.12
Eigenprotein Value
Normal.nonAA
AD.nonAA
Normal.AA
AD.AA
Diagnosis.Race
MEpurple.CSF (Synthetic)
ANOVA p: 0.25
Eigenprotein Value
Normal.Male
Normal.Female
AD.Male
AD.Female
Diagnosis.Sex
bicor=0.26, p=0.13
cor=0.25, p=0.15
Eigenprotein
ABeta/Tau Ratio
bicor=0.084, p=0.65
cor=0.064, p=0.73
Eigenprotein
MoCA Score
bicor=-0.34, p=0.16
cor=-0.36, p=0.14
Eigenprotein
Age (Normal Only)
bicor=-0.48, p=0.053
cor=-0.48, p=0.051
Eigenprotein
Age (AD Only)
M6 red.Plasma35
Axon Guidance/Nervous System Dev
Eigenprotein Value
Normal
AD
MEred.CSF 35 Samp. (Synthetic)
ANOVA p: 0.61
Eigenprotein Value
Normal
AD
MEred.CSF (Synthetic)
ANOVA p: 0.012
Eigenprotein Value
Normal.nonAA
AD.nonAA
Normal.AA
AD.AA
Diagnosis.Race
MEred.CSF (Synthetic)
ANOVA p: 0.26
Eigenprotein Value
Normal.Male
Normal.Female
AD.Male
AD.Female
Diagnosis.Sex
bicor=-0.34, p=0.048
cor=-0.32, p=0.061
Eigenprotein
ABeta/Tau Ratio
bicor=-0.11, p=0.57
cor=0.045, p=0.81
Eigenprotein
MoCA Score
bicor=0.095, p=0.71
cor=0.086, p=0.73
Eigenprotein
Age (Normal Only)
bicor=0.34, p=0.18
cor=0.37, p=0.14
Eigenprotein
Age (AD Only)

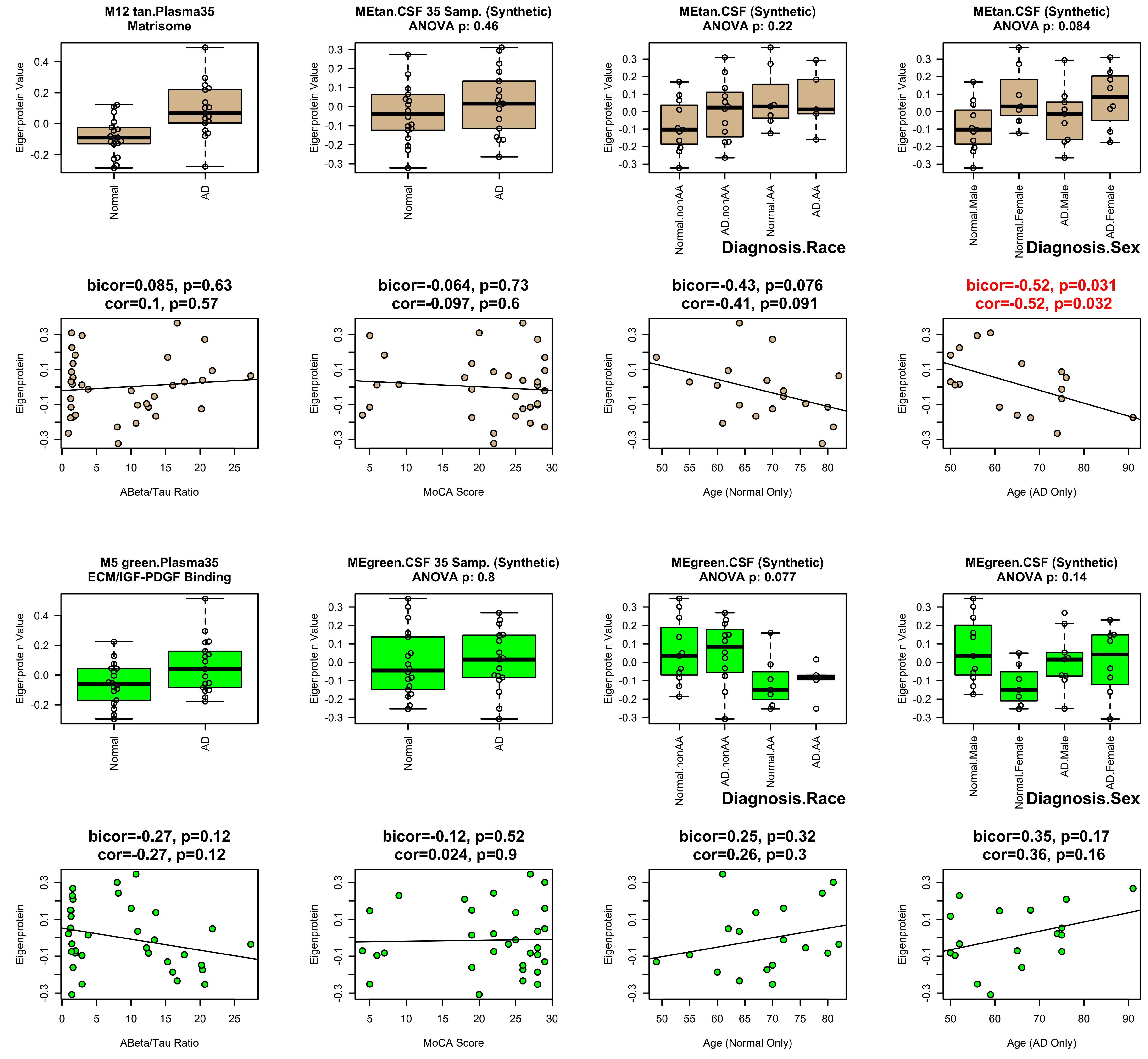
M12 tan.Plasma35
Matrisome
MEtan.CSF 35 Samp. (Synthetic)
ANOVA p: 0.46
MEtan.CSF (Synthetic)
ANOVA p: 0.22
MEtan.CSF (Synthetic)
ANOVA p: 0.084
Eigenprotein Value
Normal
AD
Normal.nonAA
AD.nonAA
Normal.AA
AD.AA
Diagnosis.Race
Normal.Male
Normal.Female
AD.Male
AD.Female
Diagnosis.Sex
bicor=0.085, p=0.63
cor=0.1, p=0.57
bicor=-0.064, p=0.73
cor=-0.097, p=0.6
bicor=-0.43, p=0.076
cor=-0.41, p=0.091
bicor=-0.52, p=0.031
cor=-0.52, p=0.032
Eigenprotein
ABeta/Tau Ratio
MoCA Score
Age (Normal Only)
Age (AD Only)
M5 green.Plasma35
ECM/IGF-PDGF Binding
MEgreen.CSF 35 Samp. (Synthetic)
ANOVA p: 0.8
MEgreen.CSF (Synthetic)
ANOVA p: 0.077
MEgreen.CSF (Synthetic)
ANOVA p: 0.14
bicor=-0.27, p=0.12
cor=-0.27, p=0.12
bicor=-0.12, p=0.52
cor=0.024, p=0.9
bicor=0.25, p=0.32
cor=0.26, p=0.3
bicor=0.35, p=0.17
cor=0.36, p=0.16

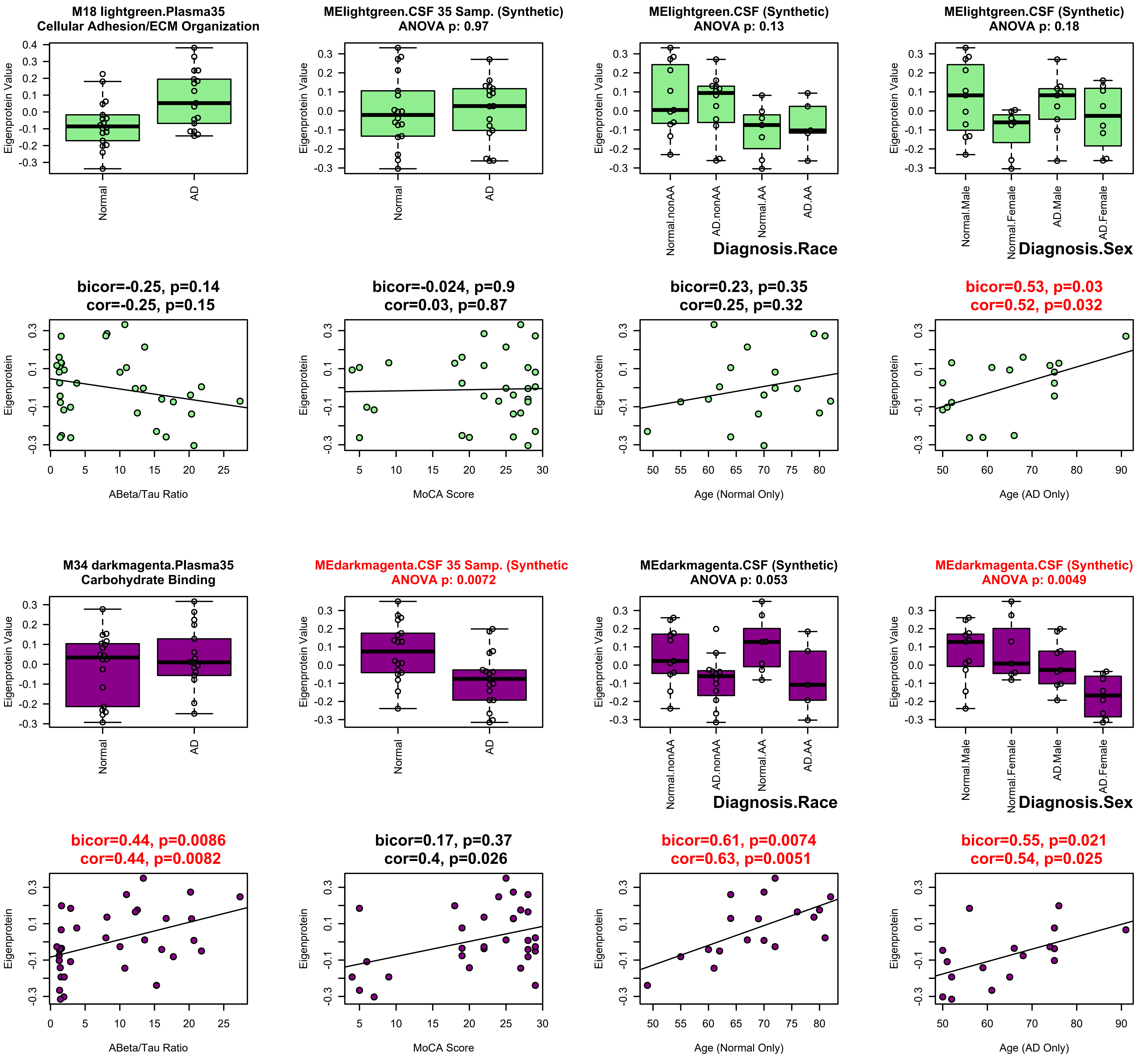
M18 lightgreen.Plasma35
Cellular Adhesion/ECM Organization
Eigenprotein Value
Normal
AD
MElightgreen.CSF 35 Samp. (Synthetic)
ANOVA p: 0.97
MElightgreen.CSF (Synthetic)
ANOVA p: 0.13
Normal.nonAA
AD.nonAA
Normal.AA
AD.AA
Diagnosis.Race
MElightgreen.CSF (Synthetic)
ANOVA p: 0.18
Normal.Male
Normal.Female
AD.Male
AD.Female
Diagnosis.Sex
bicor=-0.25, p=0.14
cor=-0.25, p=0.15
Eigenprotein
ABeta/Tau Ratio
bicor=-0.024, p=0.9
cor=0.03, p=0.87
MoCA Score
bicor=0.23, p=0.35
cor=0.25, p=0.32
Age (Normal Only)
bicor=0.53, p=0.03
cor=0.52, p=0.032
Age (AD Only)
M34 darkmagenta.Plasma35
Carbohydrate Binding
MEdarkmagenta.CSF 35 Samp. (Synthetic
ANOVA p: 0.0072
MEdarkmagenta.CSF (Synthetic)
ANOVA p: 0.053
MEdarkmagenta.CSF (Synthetic
ANOVA p: 0.0049
bicor=0.44, p=0.0086
cor=0.44, p=0.0082
bicor=0.17, p=0.37
cor=0.4, p=0.026
bicor=0.61, p=0.0074
cor=0.63, p=0.0051
bicor=0.55, p=0.021
cor=0.54, p=0.025

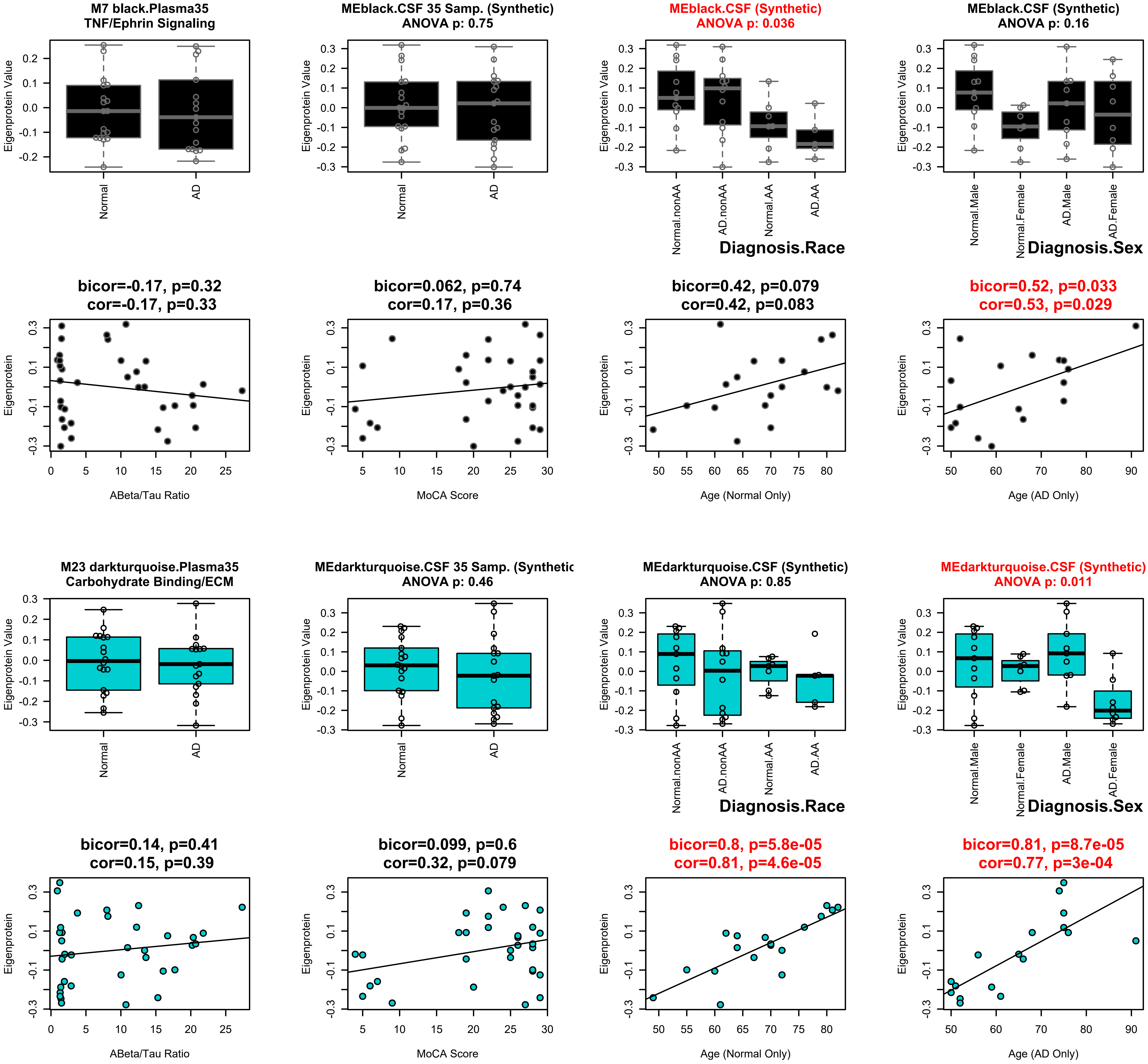
M7 black.Plasma35
TNF/Ephrin Signaling
Eigenprotein Value
Normal
AD
MEblack.CSF 35 Samp. (Synthetic)
ANOVA p: 0.75
MEblack.CSF (Synthetic)
ANOVA p: 0.036
Normal.nonAA
AD.nonAA
Normal.AA
AD.AA
Diagnosis.Race
MEblack.CSF (Synthetic)
ANOVA p: 0.16
Normal.Male
Normal.Female
AD.Male
AD.Female
Diagnosis.Sex
bicor=-0.17, p=0.32
cor=-0.17, p=0.33
ABeta/Tau Ratio
Eigenprotein
bicor=0.062, p=0.74
cor=0.17, p=0.36
MoCA Score
bicor=0.42, p=0.079
cor=0.42, p=0.083
Age (Normal Only)
bicor=0.52, p=0.033
cor=0.53, p=0.029
Age (AD Only)
M23 darkturquoise.Plasma35
Carbohydrate Binding/ECM
MEdarkturquoise.CSF 35 Samp. (Synthetic
ANOVA p: 0.46
MEdarkturquoise.CSF (Synthetic)
ANOVA p: 0.85
Diagnosis.Race
MEdarkturquoise.CSF (Synthetic)
ANOVA p: 0.011
Diagnosis.Sex
bicor=0.14, p=0.41
cor=0.15, p=0.39
ABeta/Tau Ratio
bicor=0.099, p=0.6
cor=0.32, p=0.079
MoCA Score
bicor=0.8, p=5.8e-05
cor=0.81, p=4.6e-05
Age (Normal Only)
bicor=0.81, p=8.7e-05
cor=0.77, p=3e-04
Age (AD Only)

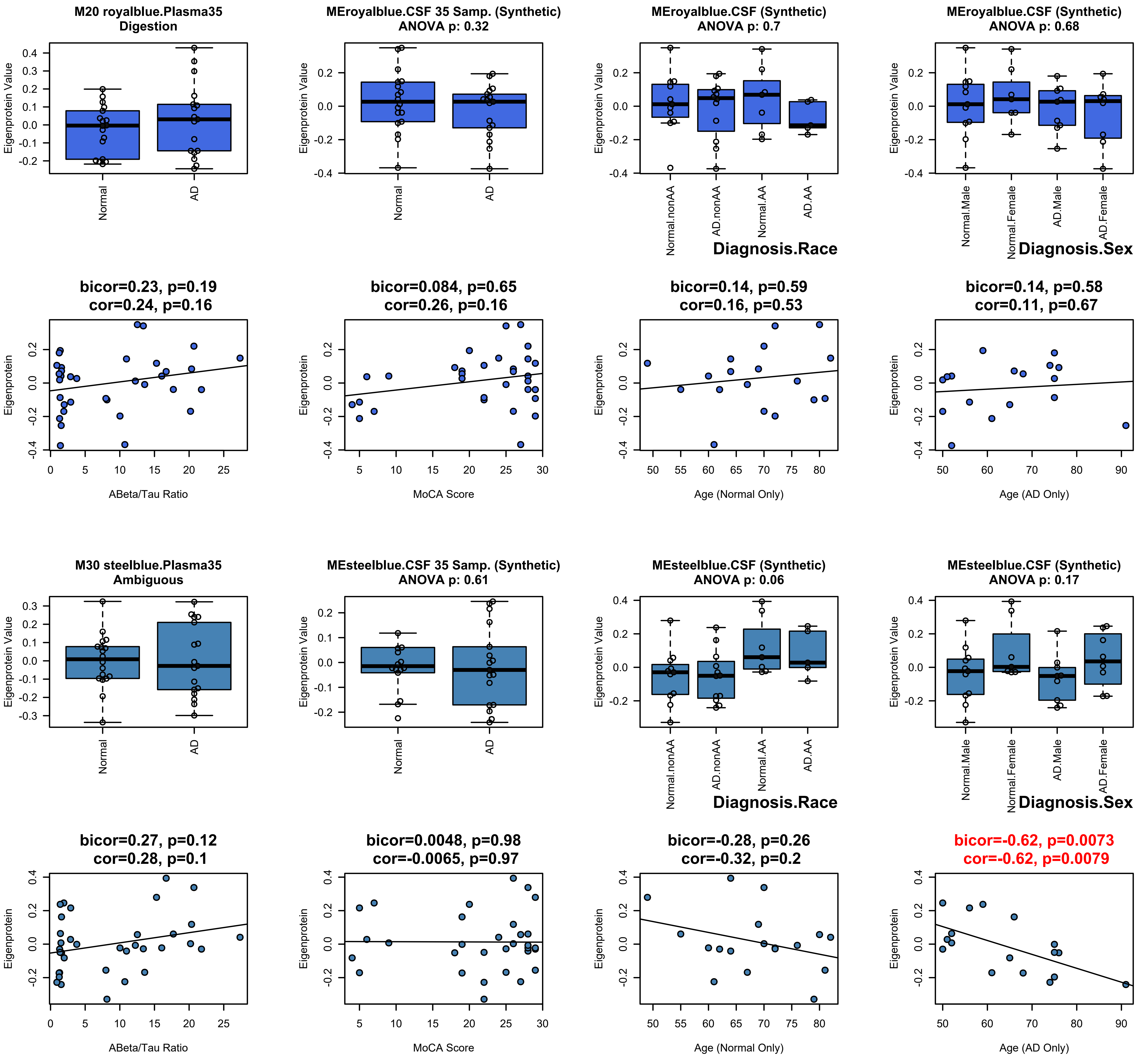
M20 royalblue.Plasma35
Digestion
Eigenprotein Value
Normal
AD
MEroyalblue.CSF 35 Samp. (Synthetic)
ANOVA p: 0.32
MEroyalblue.CSF (Synthetic)
ANOVA p: 0.7
Normal.nonAA
AD.nonAA
Normal.AA
AD.AA
Diagnosis.Race
MEroyalblue.CSF (Synthetic)
ANOVA p: 0.68
Normal.Male
Normal.Female
AD.Male
AD.Female
Diagnosis.Sex
bicor=0.23, p=0.19
cor=0.24, p=0.16
Eigenprotein
ABeta/Tau Ratio
bicor=0.084, p=0.65
cor=0.26, p=0.16
MoCA Score
bicor=0.14, p=0.59
cor=0.16, p=0.53
Age (Normal Only)
bicor=0.14, p=0.58
cor=0.11, p=0.67
Age (AD Only)
M30 steelblue.Plasma35
Ambiguous
MEsteelblue.CSF 35 Samp. (Synthetic)
ANOVA p: 0.61
MEsteelblue.CSF (Synthetic)
ANOVA p: 0.06
MEsteelblue.CSF (Synthetic)
ANOVA p: 0.17
bicor=0.27, p=0.12
cor=0.28, p=0.1
bicor=0.0048, p=0.98
cor=-0.0065, p=0.97
bicor=-0.28, p=0.26
cor=-0.32, p=0.2
bicor=-0.62, p=0.0073
cor=-0.62, p=0.0079

### Data Subset:

### Plasma 32 of 35 modules as template

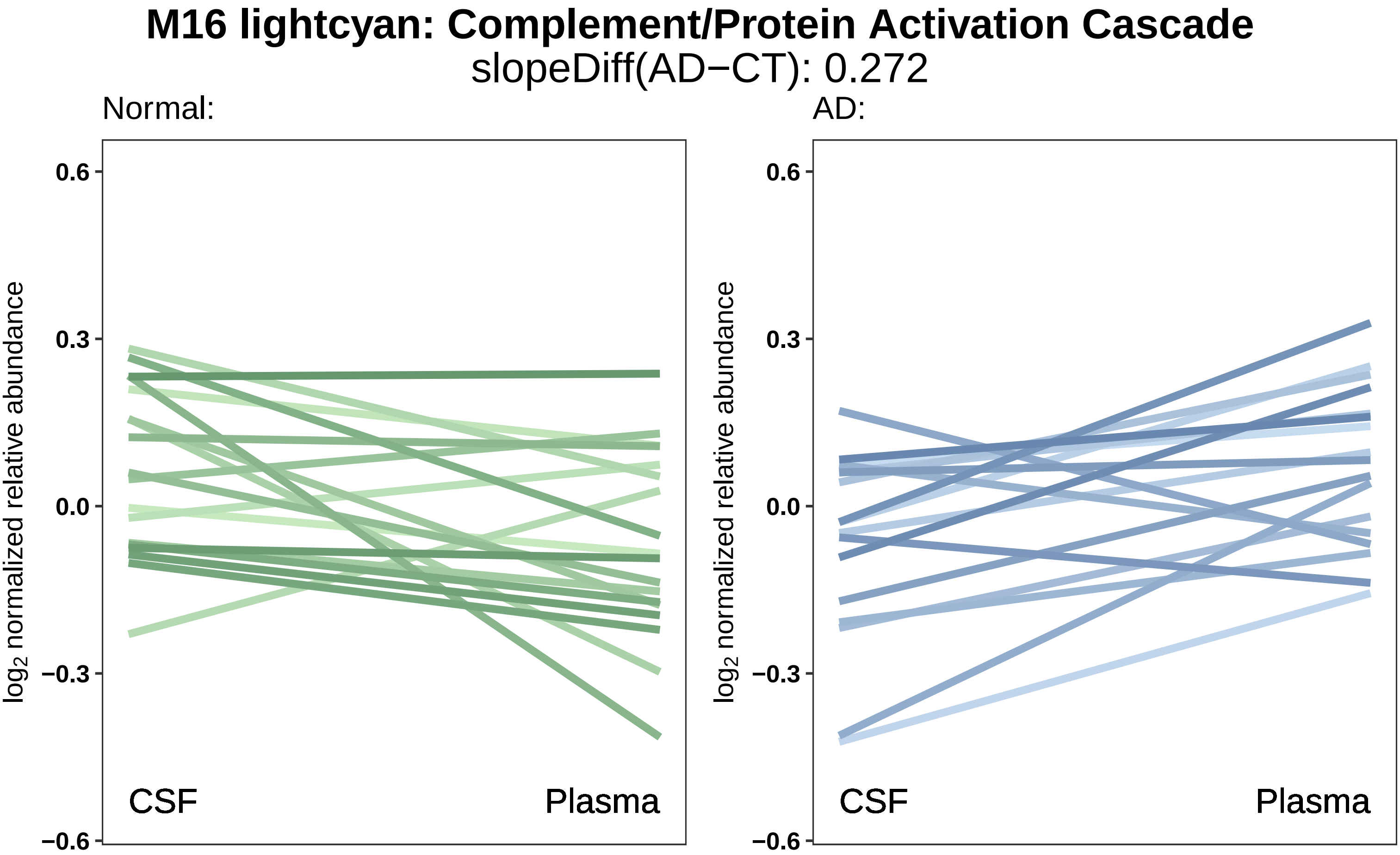

M16 lightcyan: Complement/Protein Activation Cascade
slopeDiff(AD−CT): 0.272
Normal:
AD:
0.6
0.3
0.0
−0.3
−0.6
log2 normalized relative abundance
CSF
Plasma

### M8 pink: Protein Activation Cascade

slopeDiff(AD−CT): 0.25

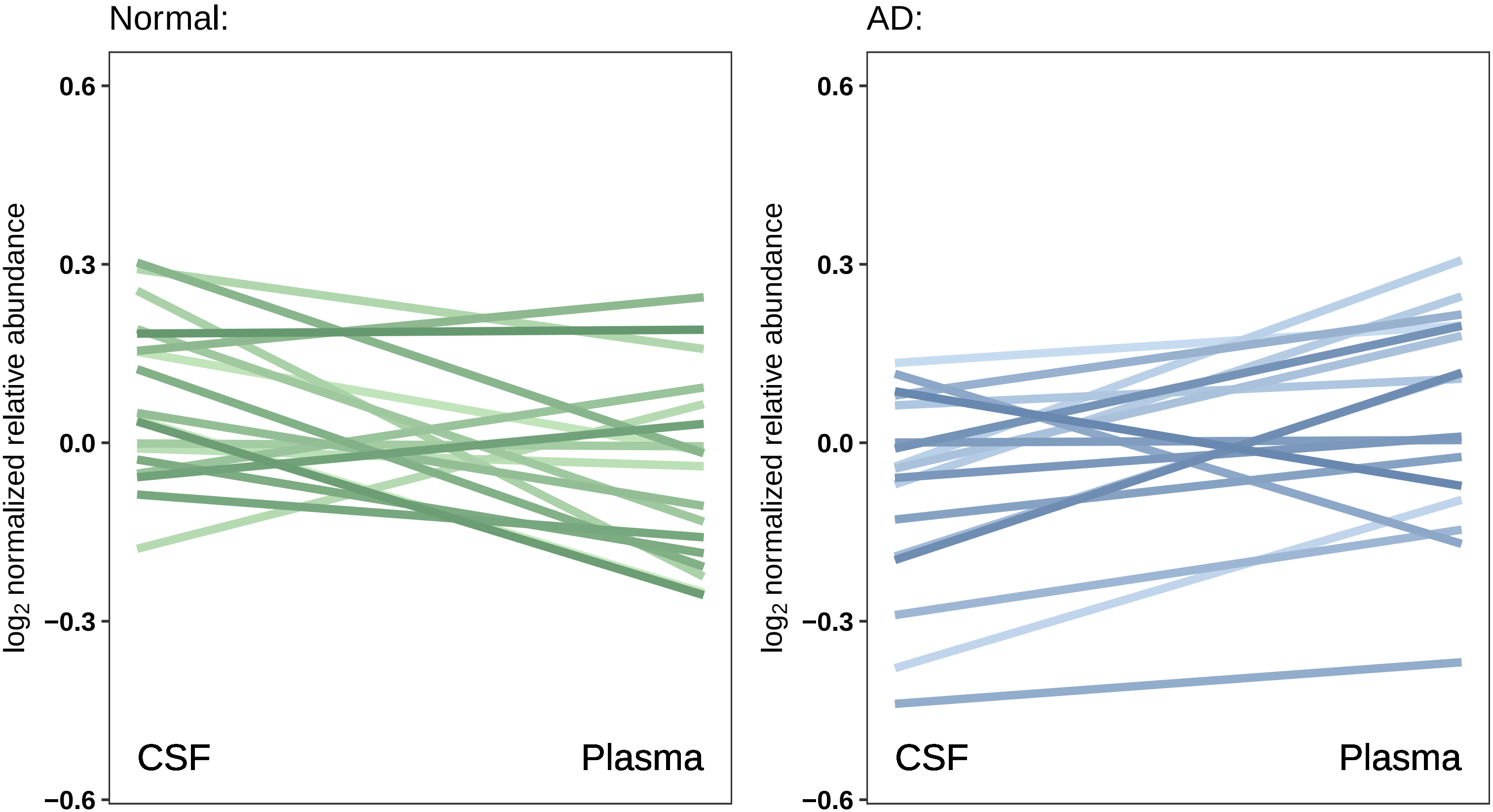

### M27 white: Collagen/ECM

slopeDiff(AD−CT): 0.24

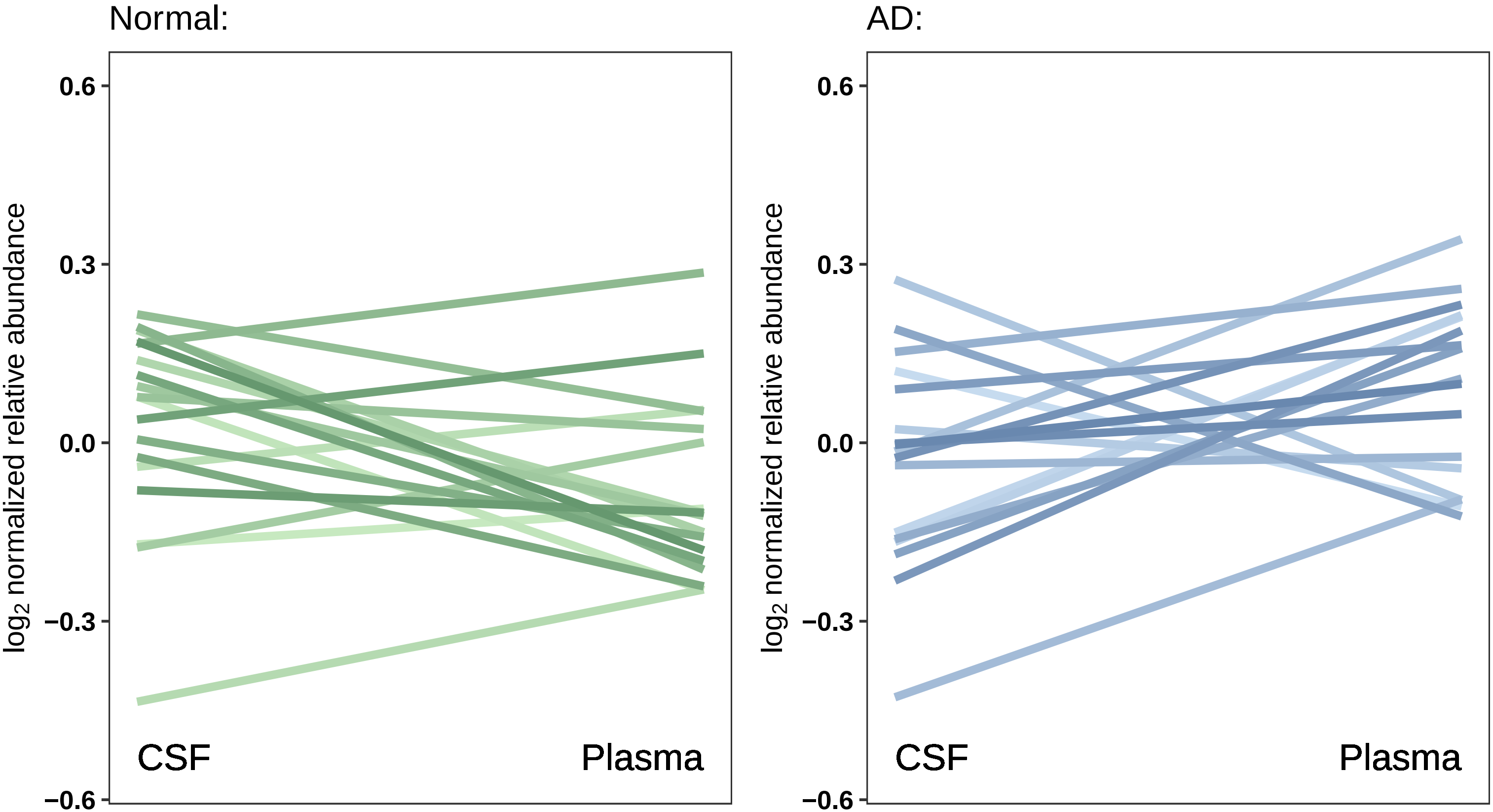

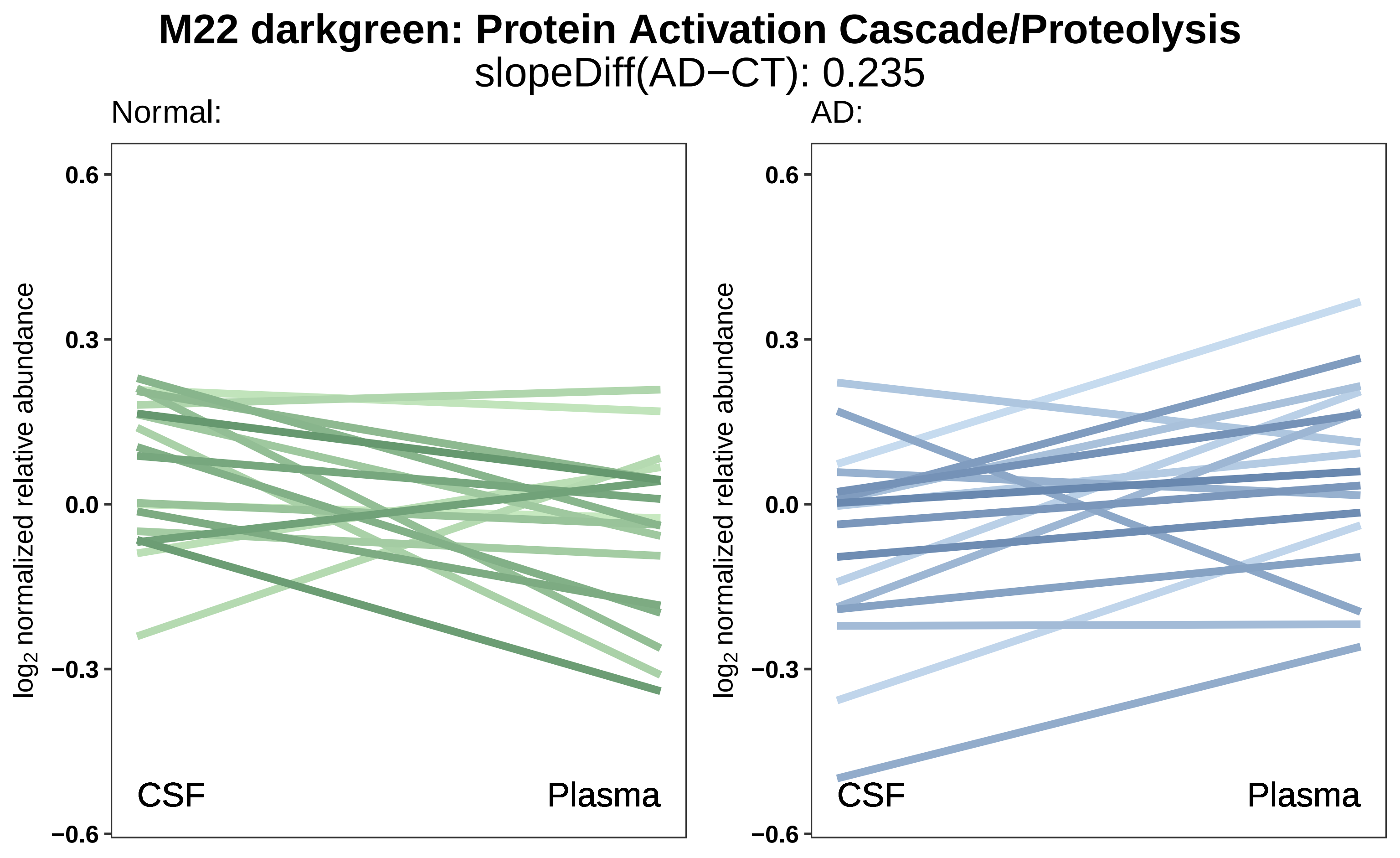
M22 darkgreen: Protein Activation Cascade/Proteolysis
slopeDiff(AD−CT): 0.235
Normal:
AD:
log2 normalized relative abundance
0.6
0.3
0.0
−0.3
−0.6
CSF
Plasma

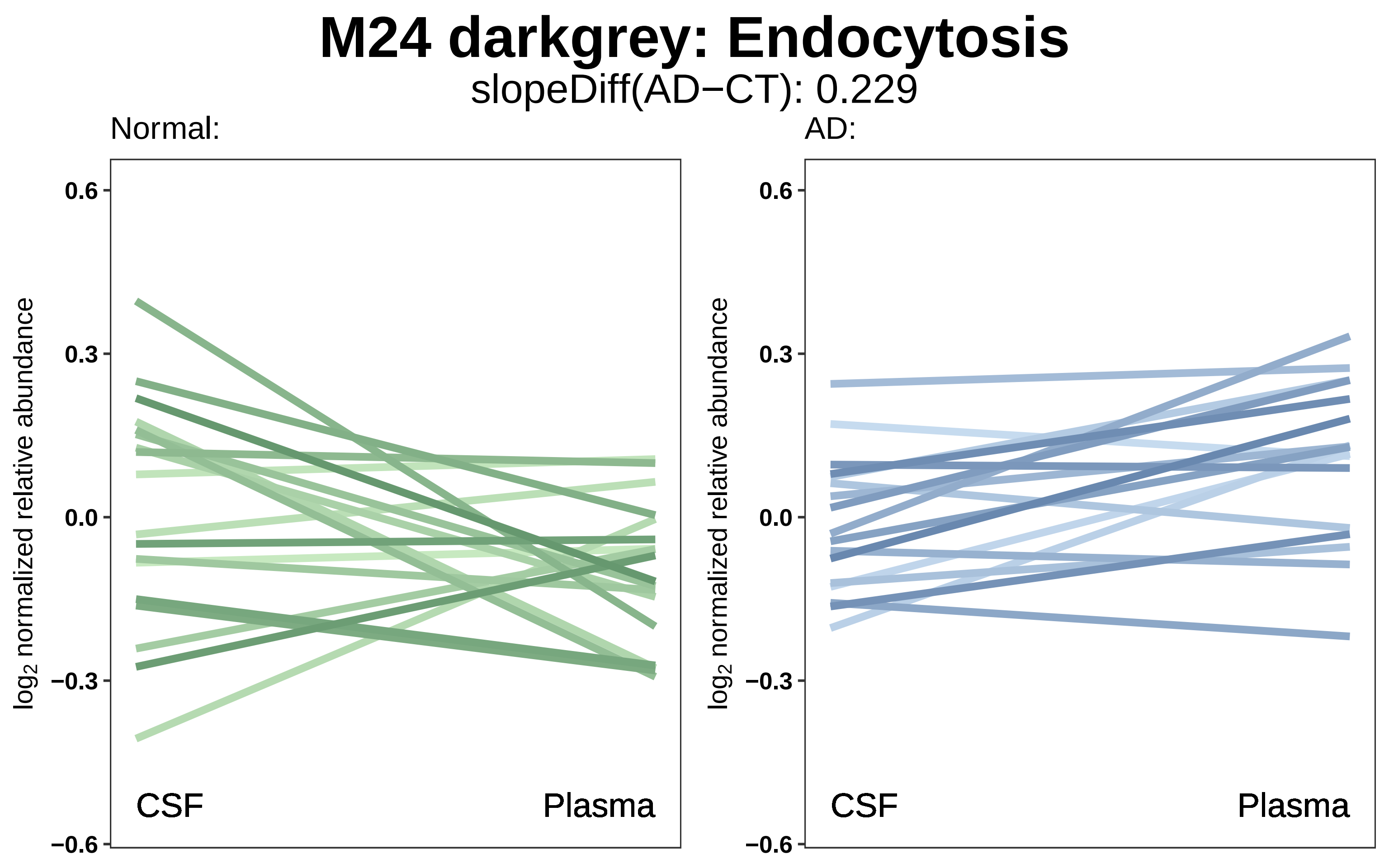

M24 darkgrey: Endocytosis
slopeDiff(AD−CT): 0.229
Normal:
AD:
0.6
0.3
0.0
−0.3
−0.6
log2 normalized relative abundance
CSF
Plasma

### M35 sienna3: Ambiguous

slopeDiff(AD−CT): 0.218

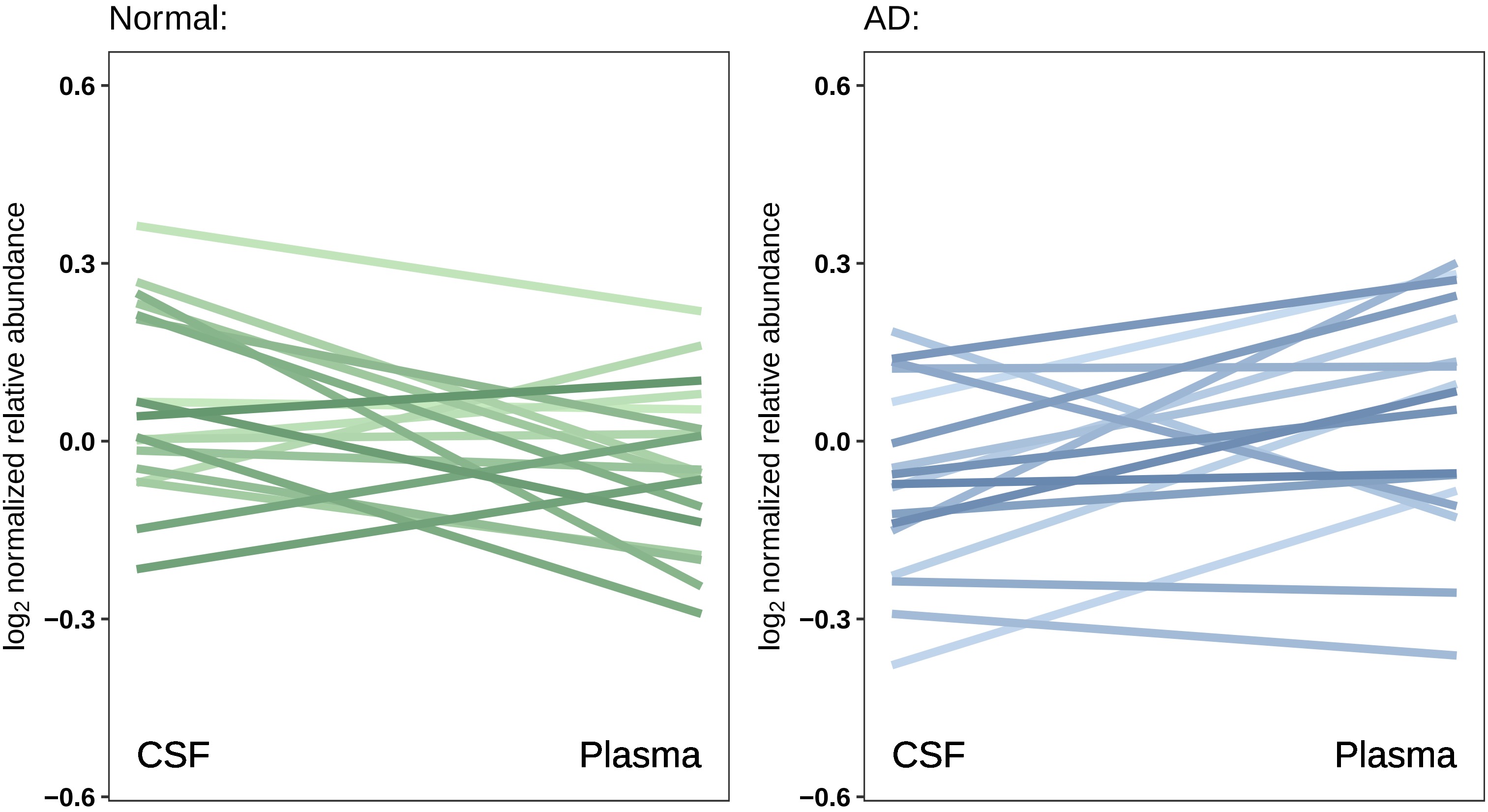

### M29 saddlebrown: IGF−Growth/GAG Binding

slopeDiff(AD−CT): 0.207

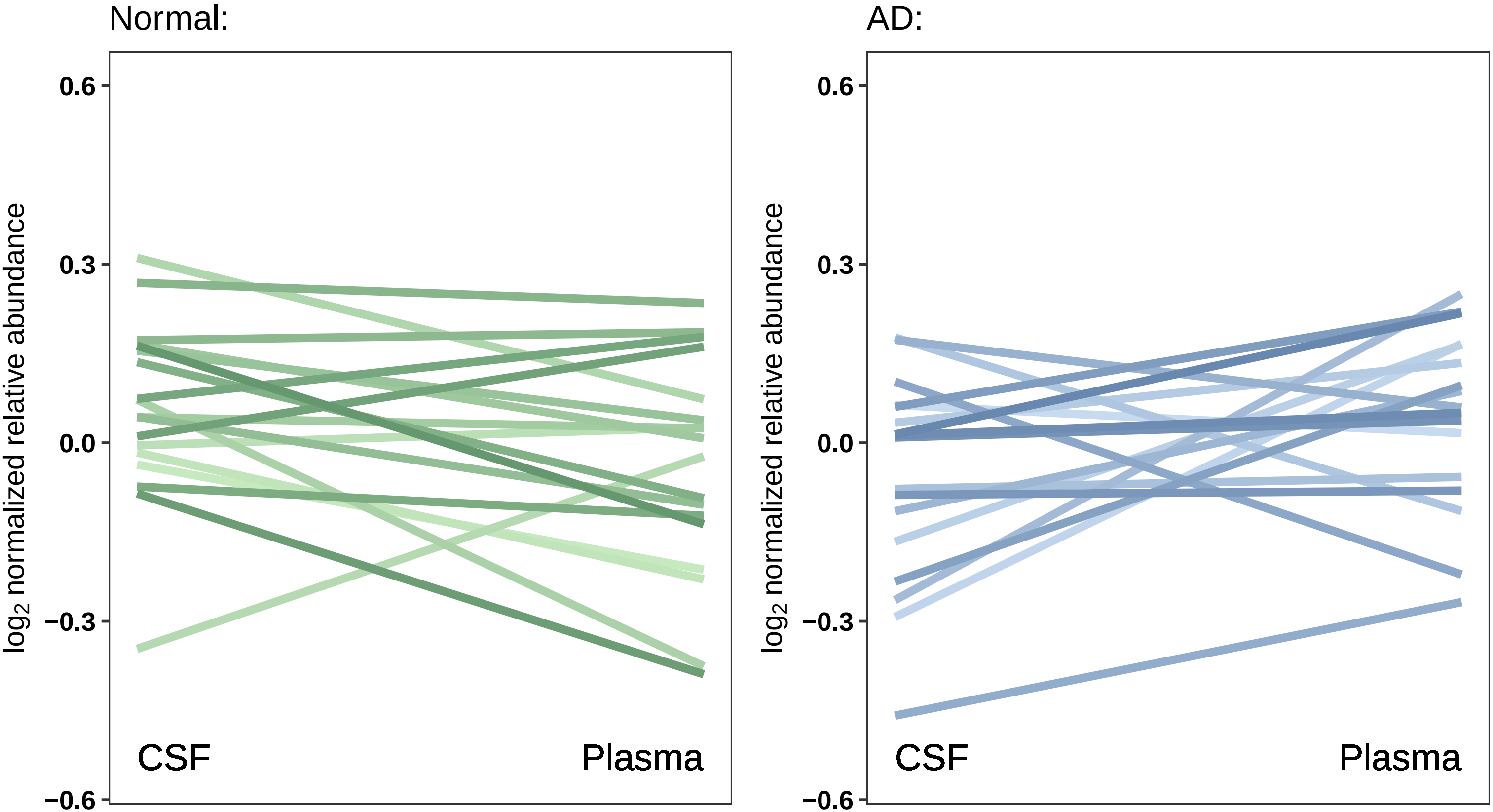

### M10 purple: Ambiguous

slopeDiff(AD−CT): 0.204

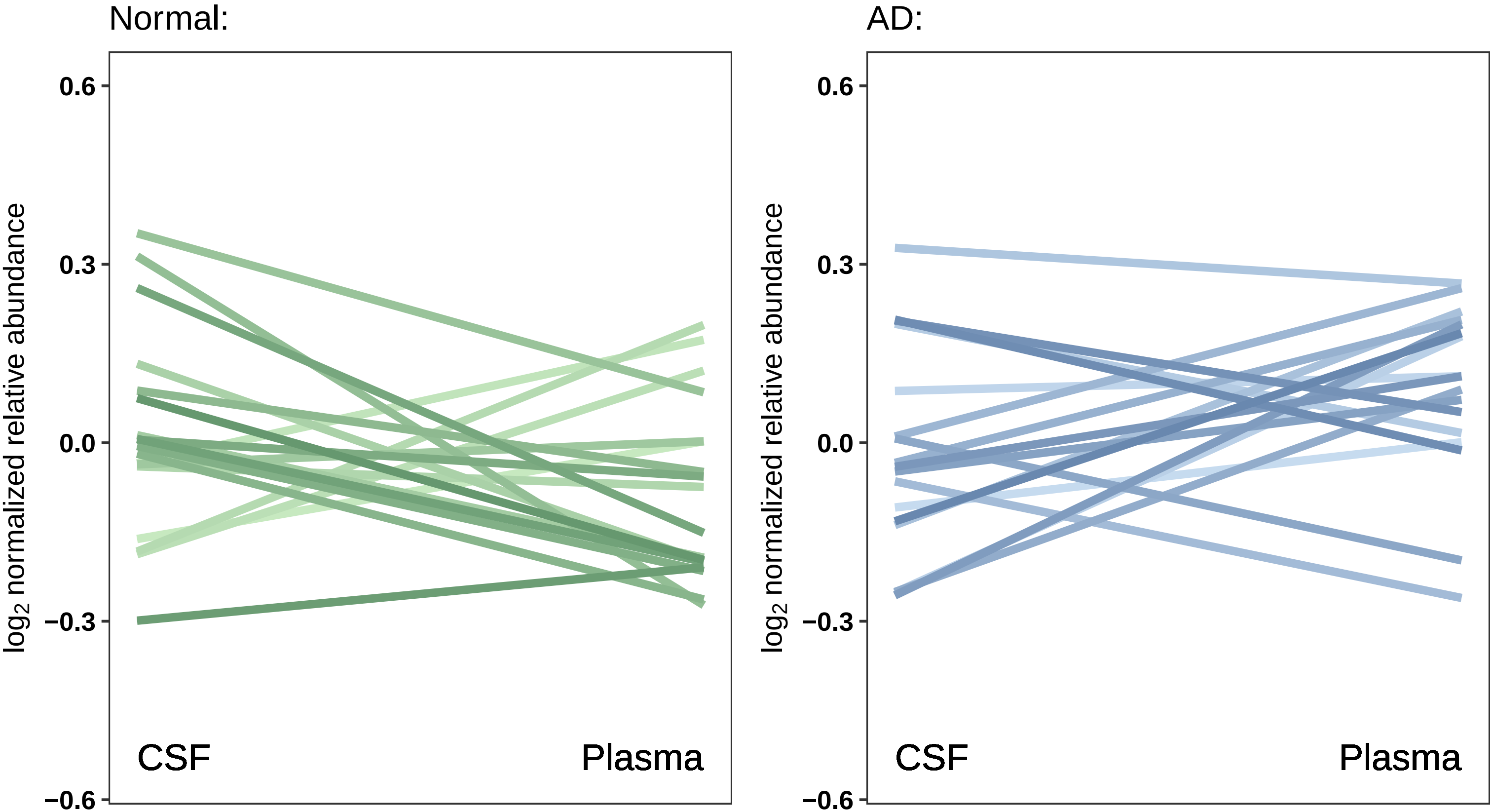

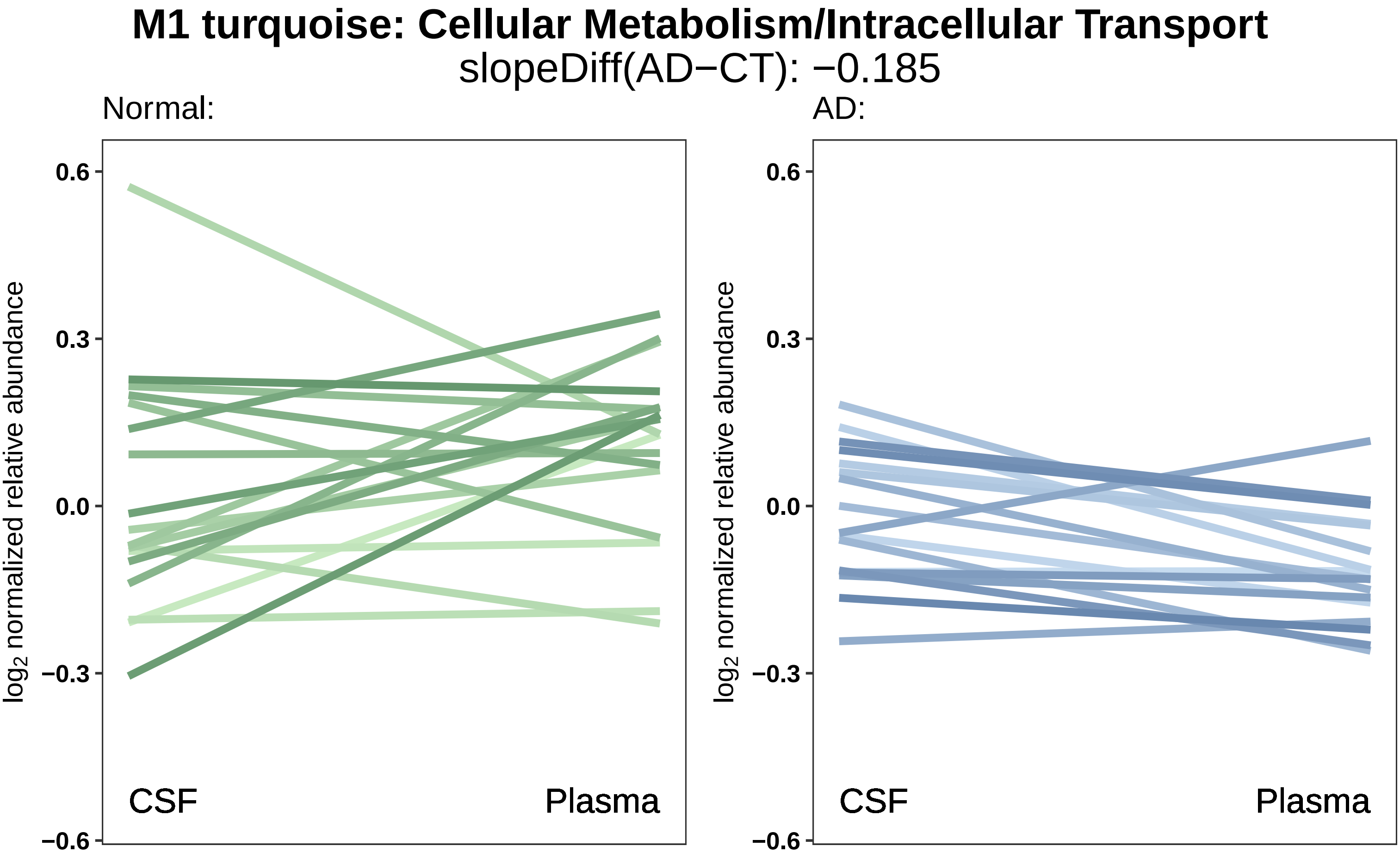
M1 turquoise: Cellular Metabolism/Intracellular Transport
slopeDiff(AD−CT): −0.185
Normal:
AD:
0.6
0.3
0.0
−0.3
−0.6
log2 normalized relative abundance
CSF
Plasma
0.6
0.3
0.0
−0.3
−0.6
log2 normalized relative abundance
CSF
Plasma

### M34 darkmagenta: Carbohydrate Binding

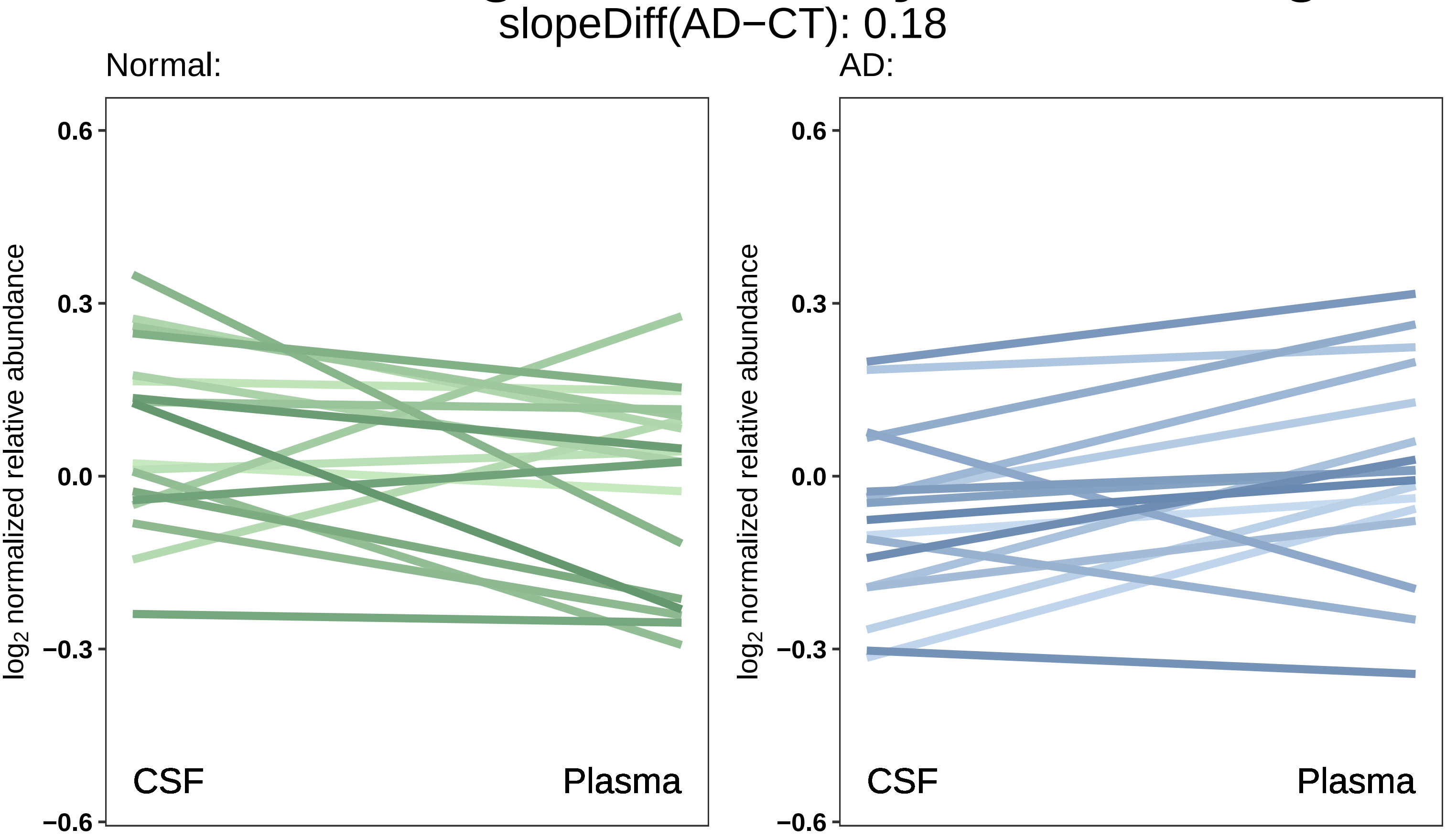

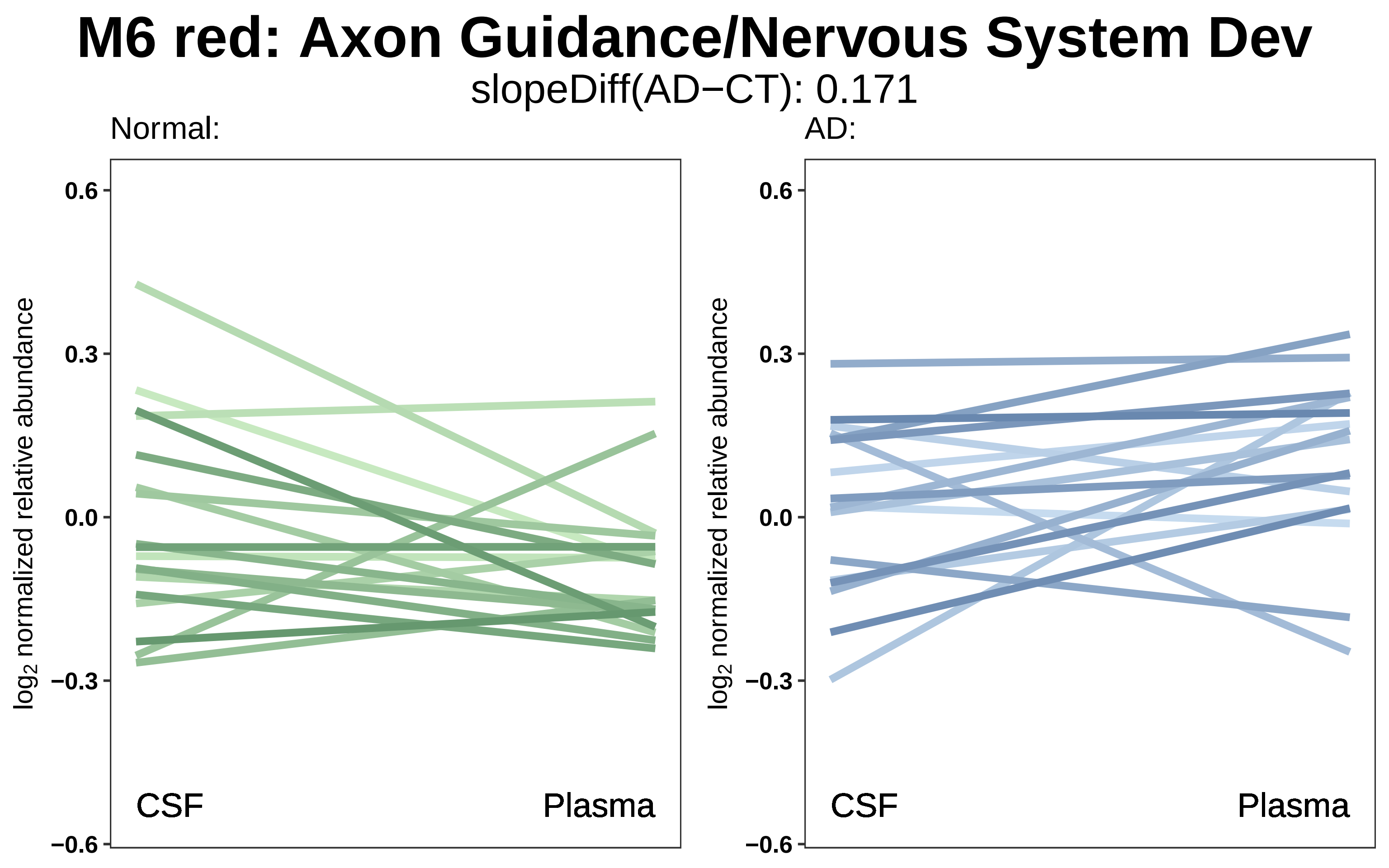
M6 red: Axon Guidance/Nervous System Dev
slopeDiff(AD−CT): 0.171
Normal:
AD:
log2 normalized relative abundance
0.6
0.3
0.0
−0.3
−0.6
CSF
Plasma

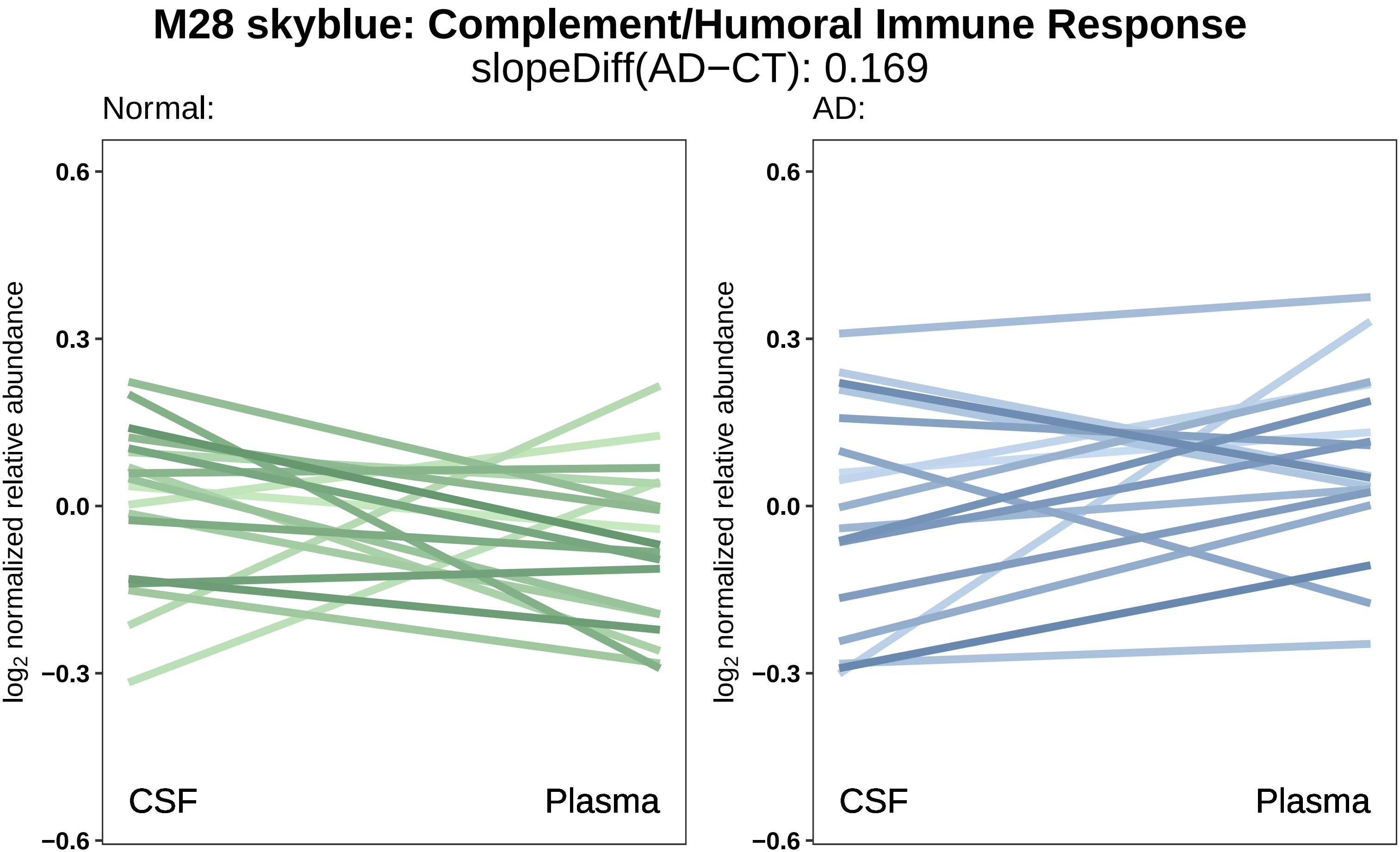

M28 skyblue: Complement/Humoral Immune Response
slopeDiff(AD−CT): 0.169
Normal:
AD:
log2 normalized relative abundance
0.6
0.3
0.0
−0.3
−0.6
CSF
Plasma

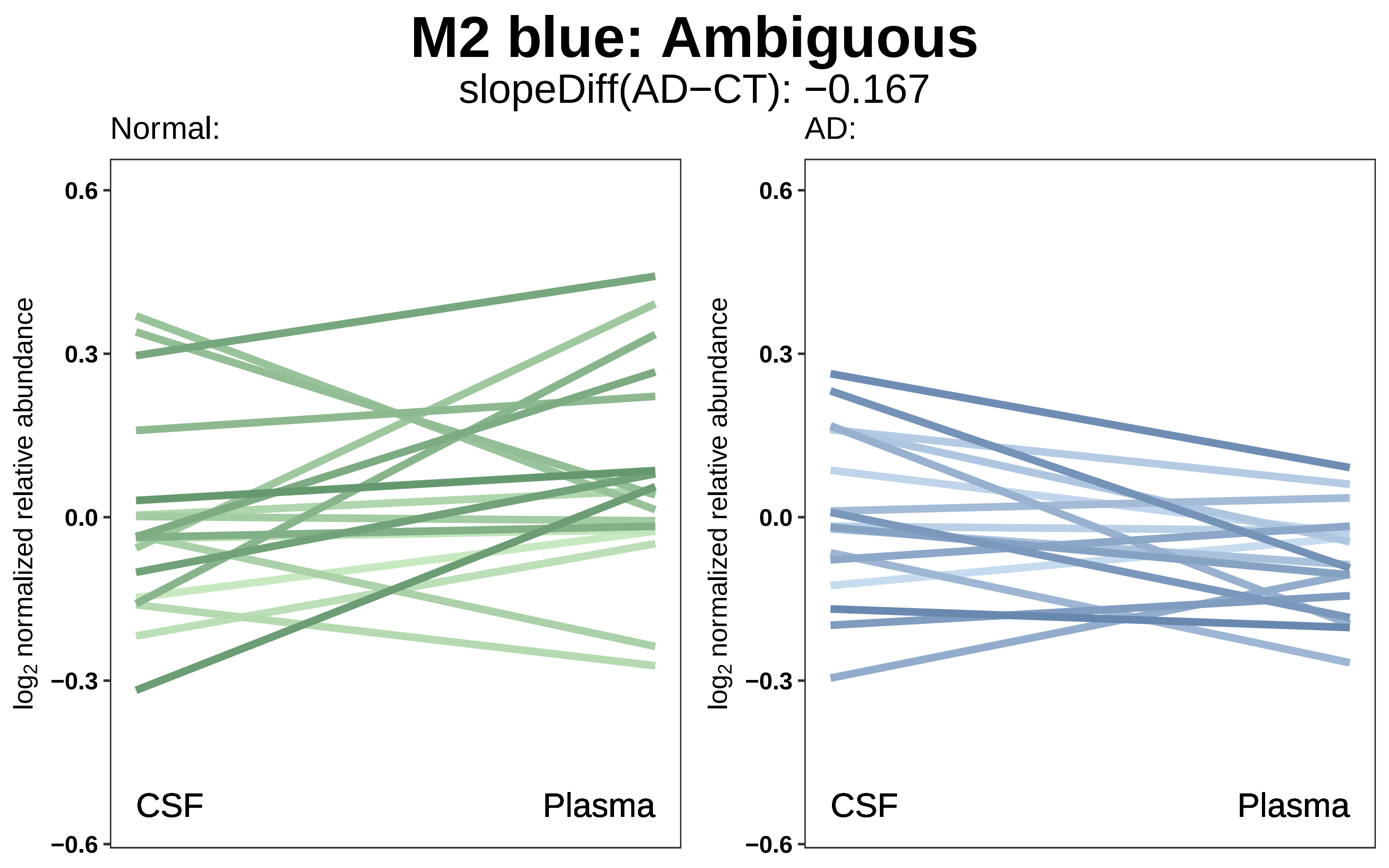
M2 blue: Ambiguous
slopeDiff(AD−CT): −0.167
Normal:
AD:
0.6
0.3
0.0
−0.3
−0.6
log2 normalized relative abundance
CSF
Plasma

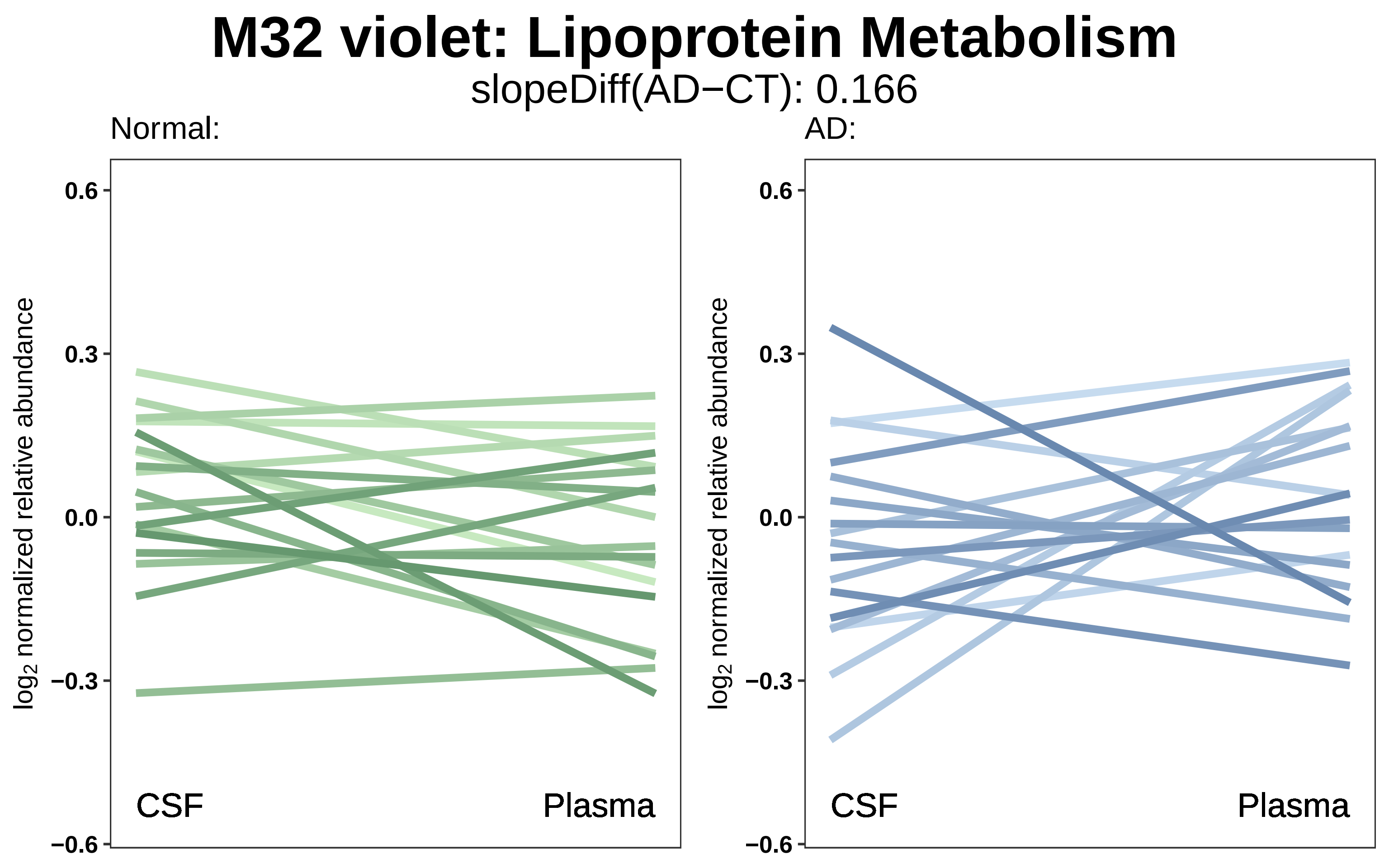

M32 violet: Lipoprotein Metabolism
slopeDiff(AD−CT): 0.166
Normal:
AD:
0.6
0.3
0.0
−0.3
−0.6
log2 normalized relative abundance
CSF
Plasma

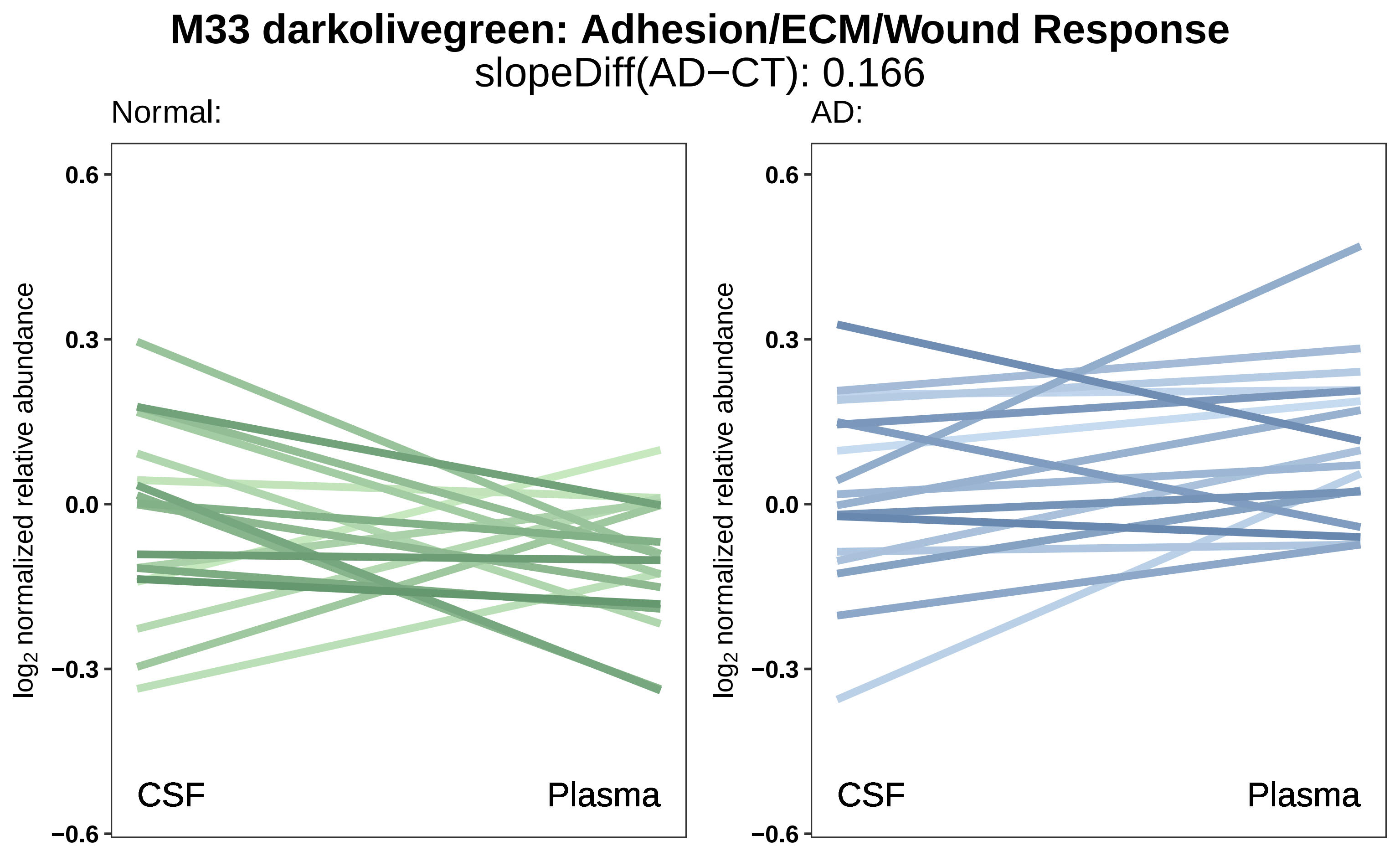
M33 darkolivegreen: Adhesion/ECM/Wound Response
slopeDiff(AD−CT): 0.166
Normal:
AD:
0.6
0.3
0.0
−0.3
−0.6
log2 normalized relative abundance
CSF
Plasma

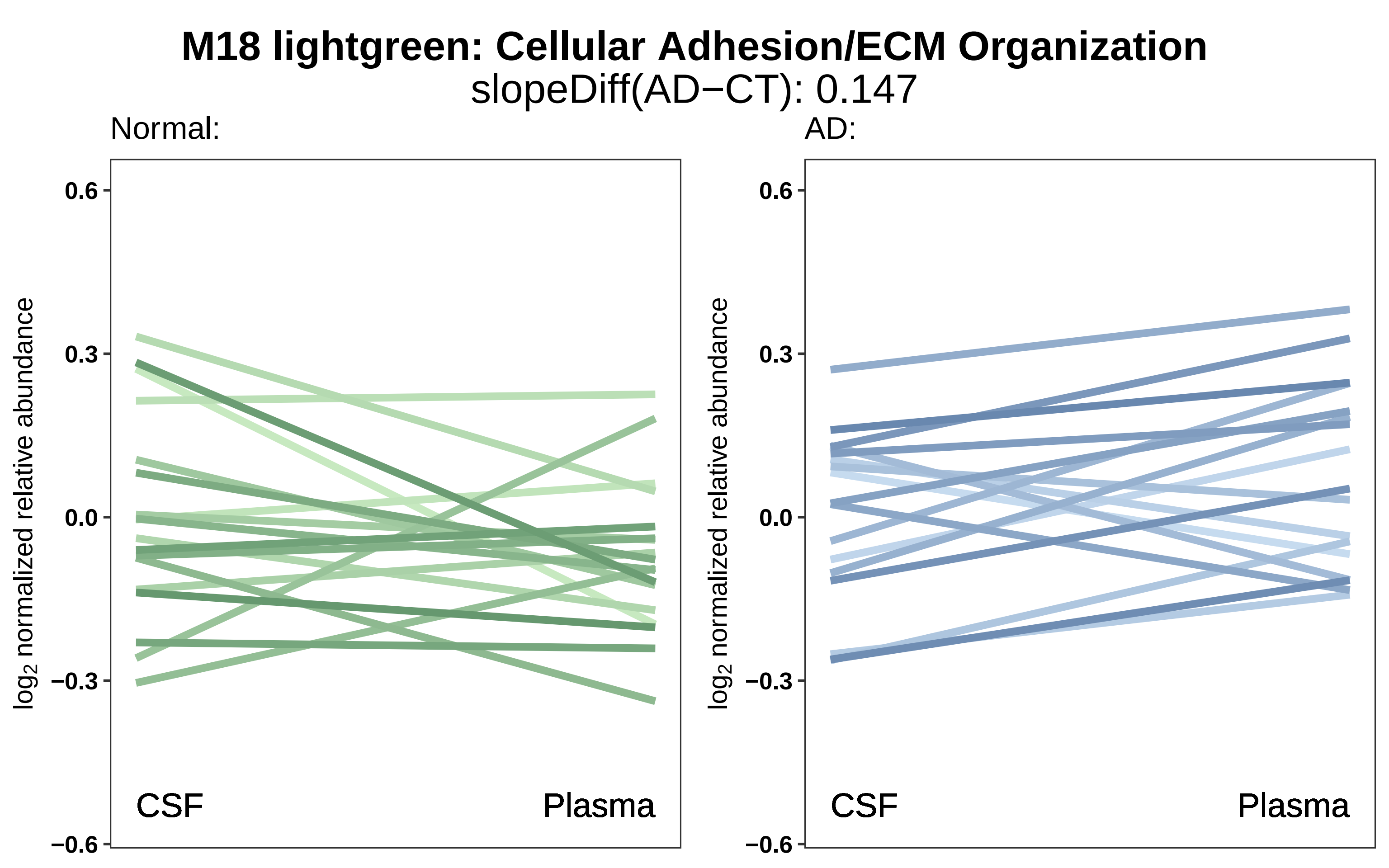
M18 lightgreen: Cellular Adhesion/ECM Organization
slopeDiff(AD−CT): 0.147
Normal:
AD:
log2 normalized relative abundance
0.6
0.3
0.0
−0.3
−0.6
CSF
Plasma

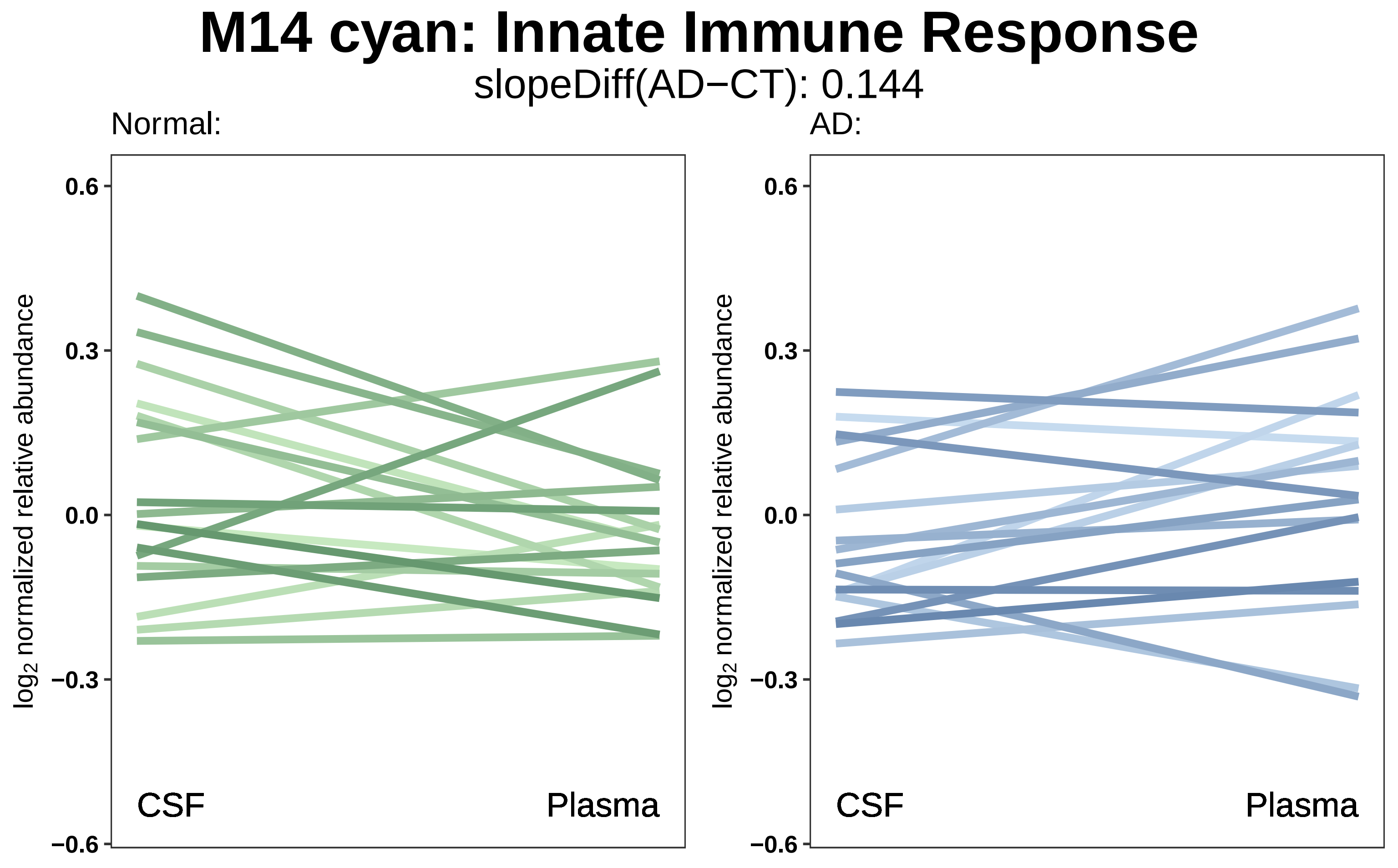
M14 cyan: Innate Immune Response
slopeDiff(AD−CT): 0.144
Normal:
AD:
0.6
0.3
0.0
−0.3
−0.6
log2 normalized relative abundance
CSF
Plasma
CSF
Plasma

### M25 orange: Lysosome

slopeDiff(AD−CT): 0.143

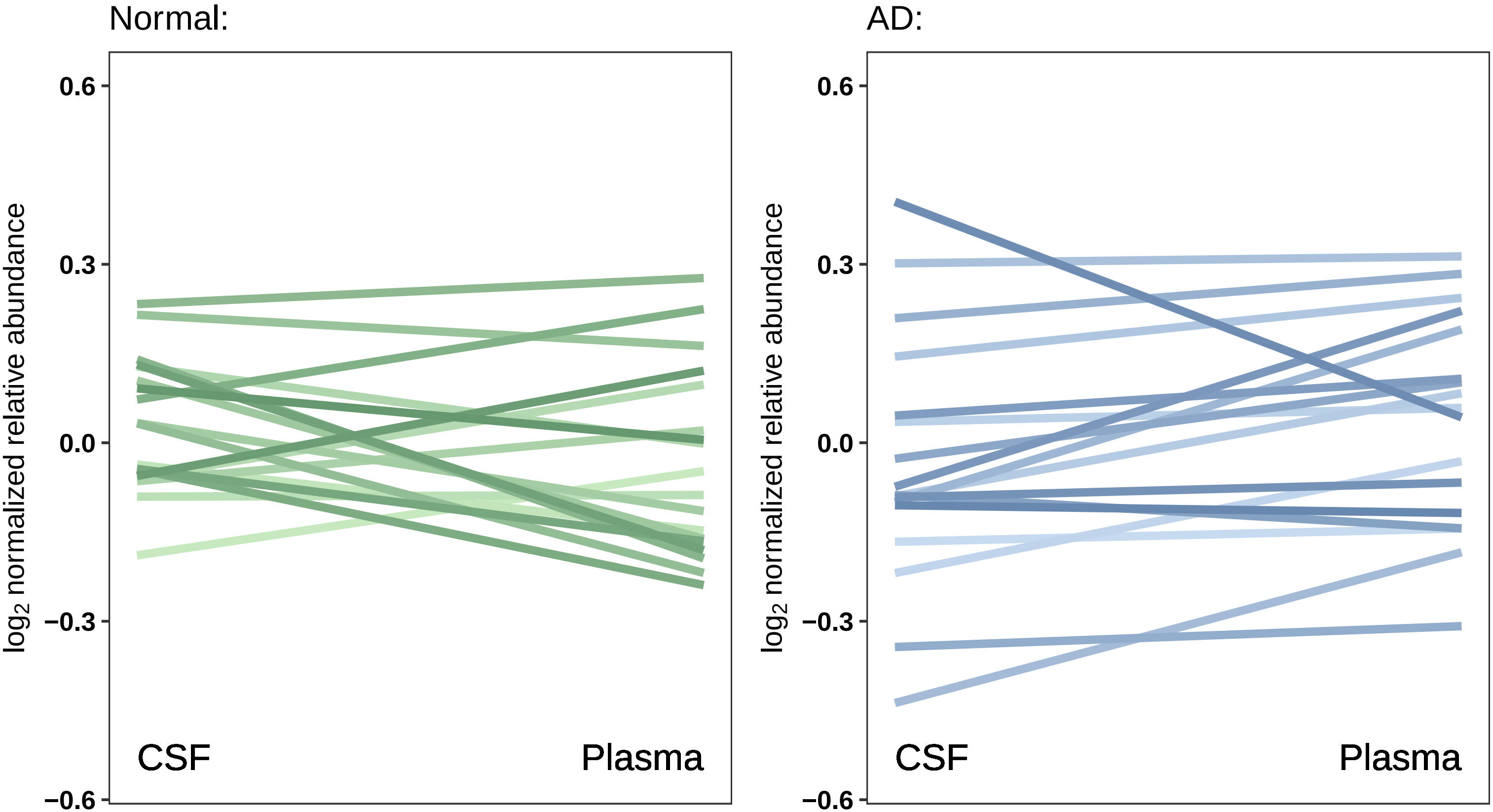

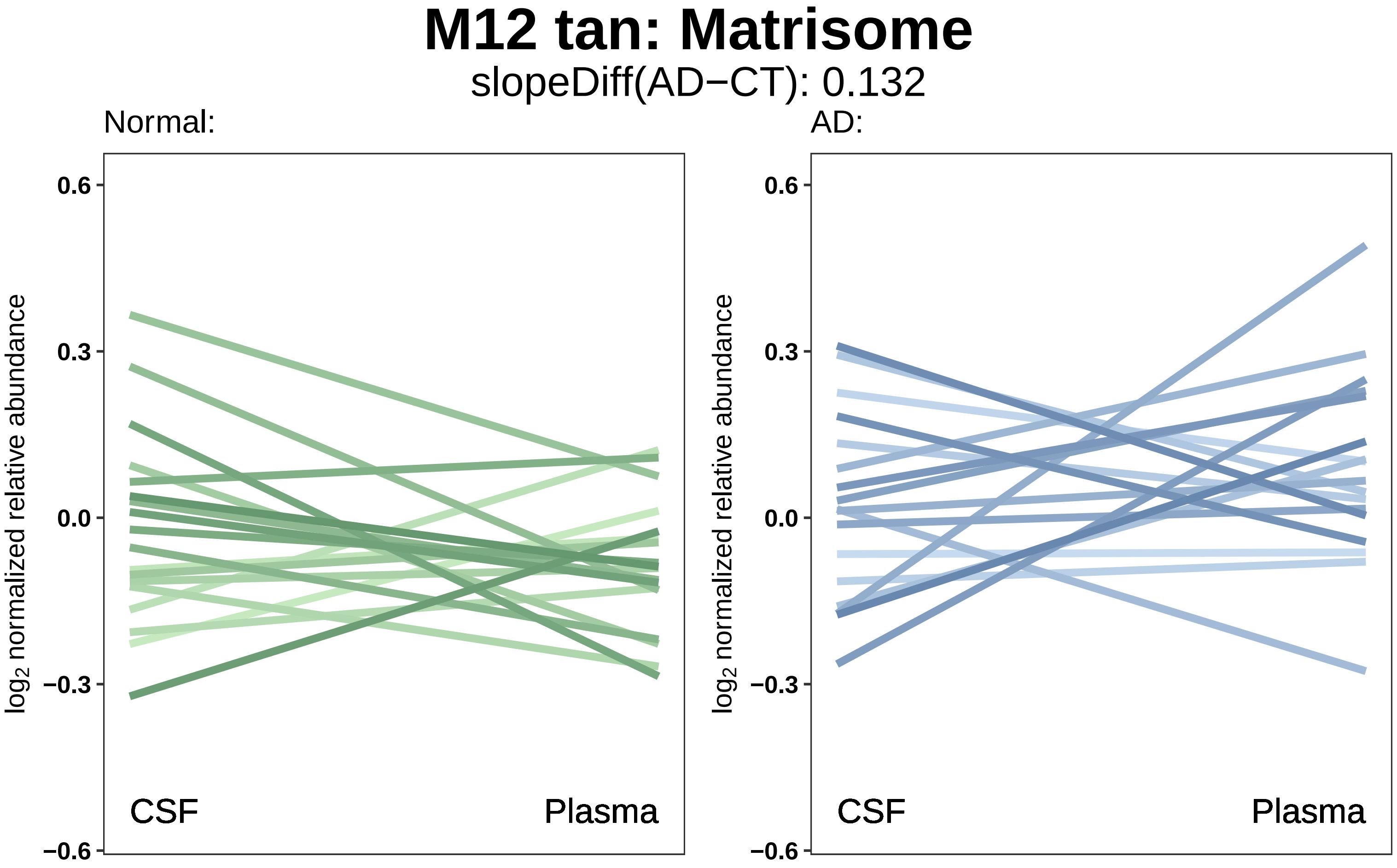
M12 tan: Matrisome
slopeDiff(AD−CT): 0.132
Normal:
AD:
log2 normalized relative abundance
0.6
0.3
0.0
−0.3
−0.6
CSF
Plasma

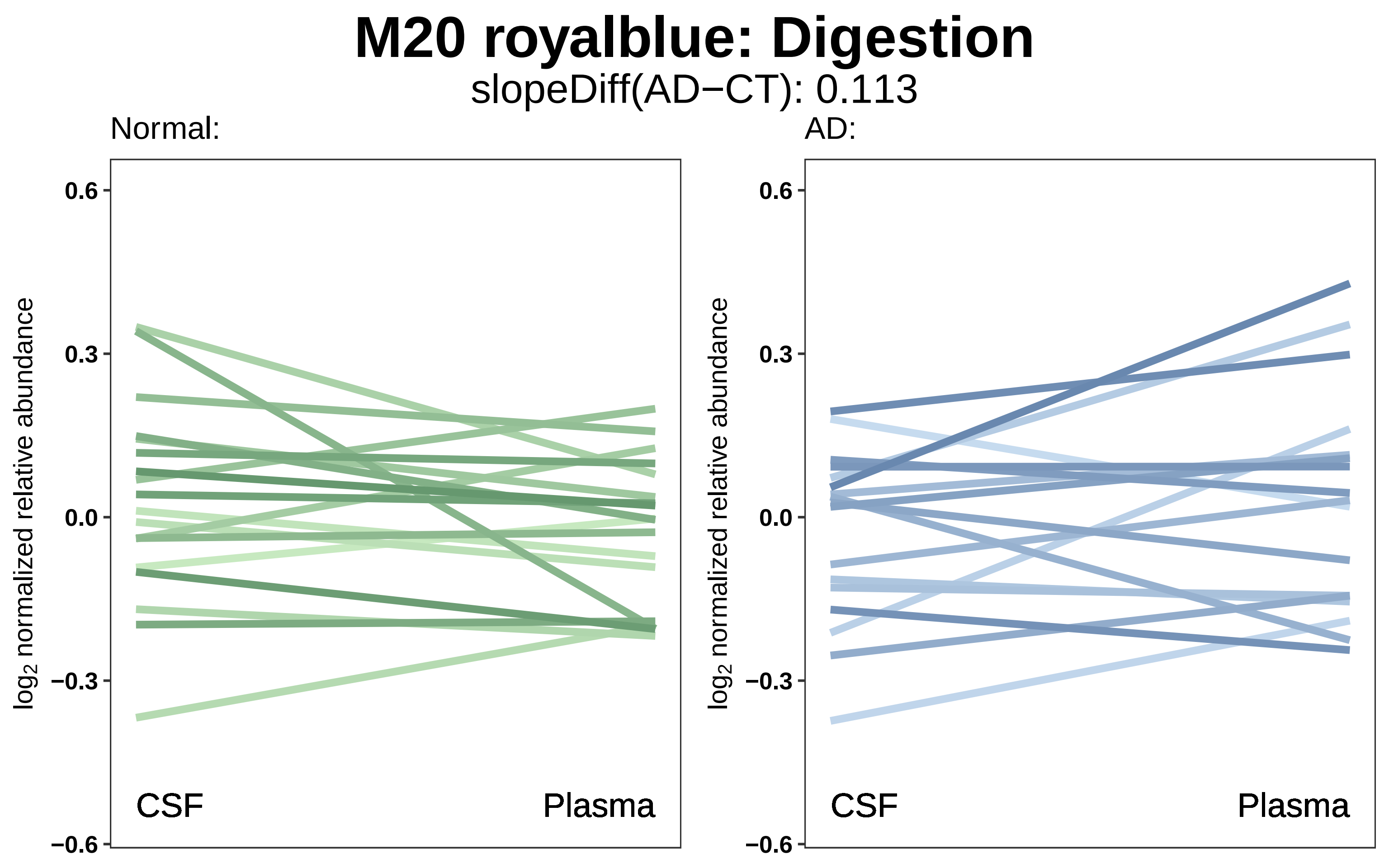
M20 royalblue: Digestion
slopeDiff(AD−CT): 0.113
Normal:
AD:
0.6
0.3
0.0
−0.3
−0.6
log2 normalized relative abundance
CSF
Plasma

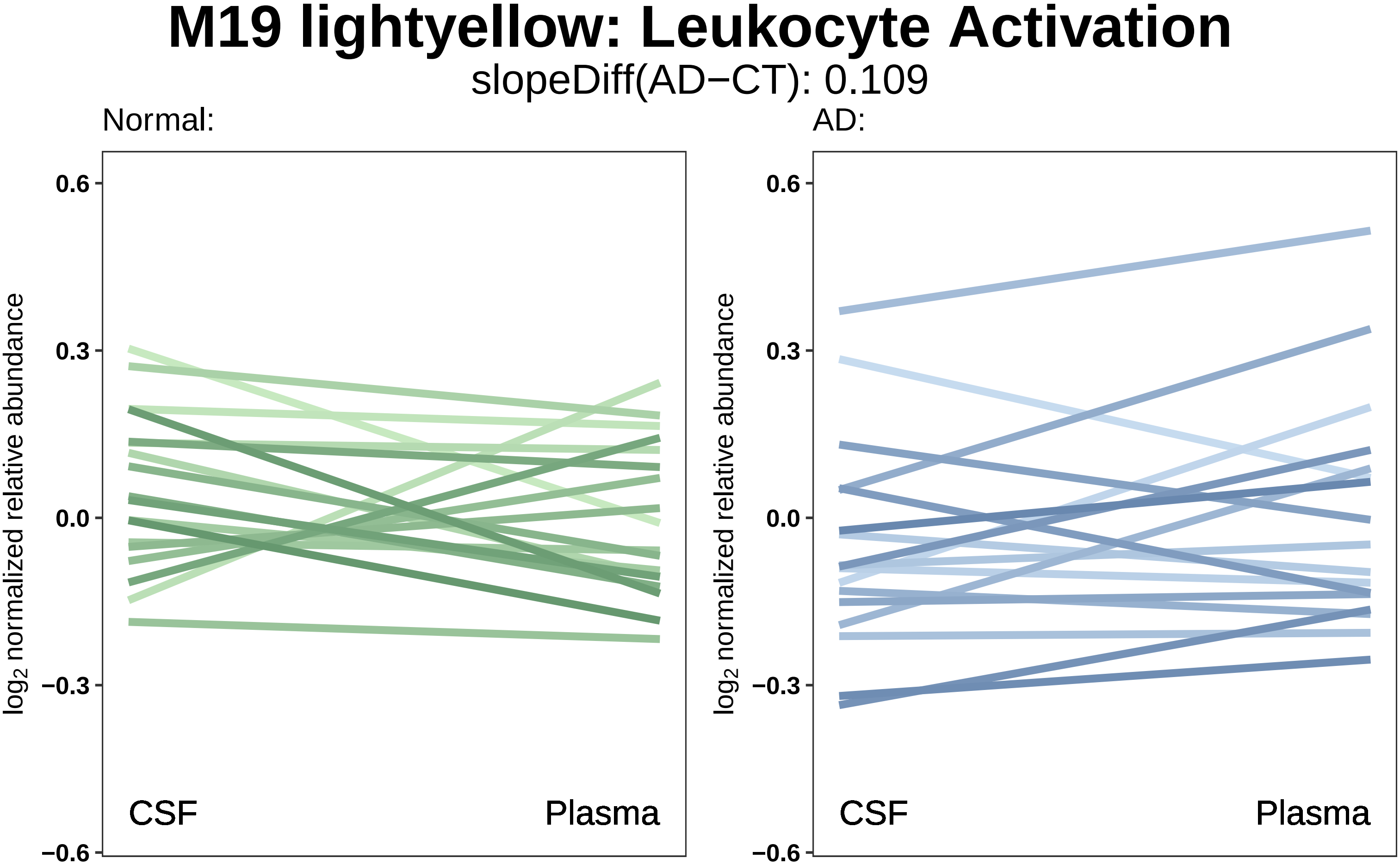
M19 lightyellow: Leukocyte Activation
slopeDiff(AD−CT): 0.109
Normal:
AD:
0.6
0.3
0.0
−0.3
−0.6
$log_2$ normalized relative abundance
CSF
Plasma

### M5 green: ECM/IGF−PDGF Binding

slopeDiff(AD−CT): 0.107

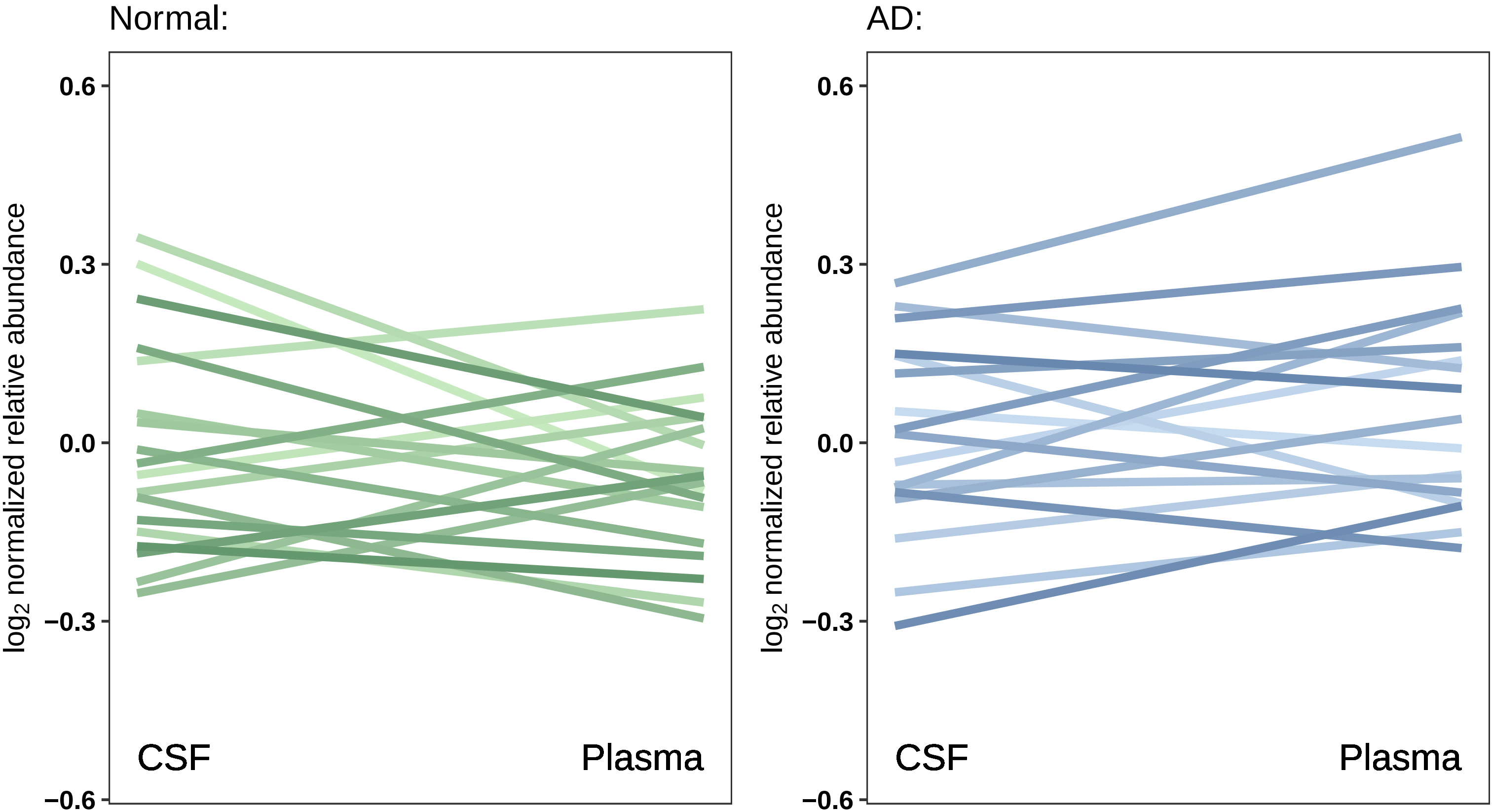

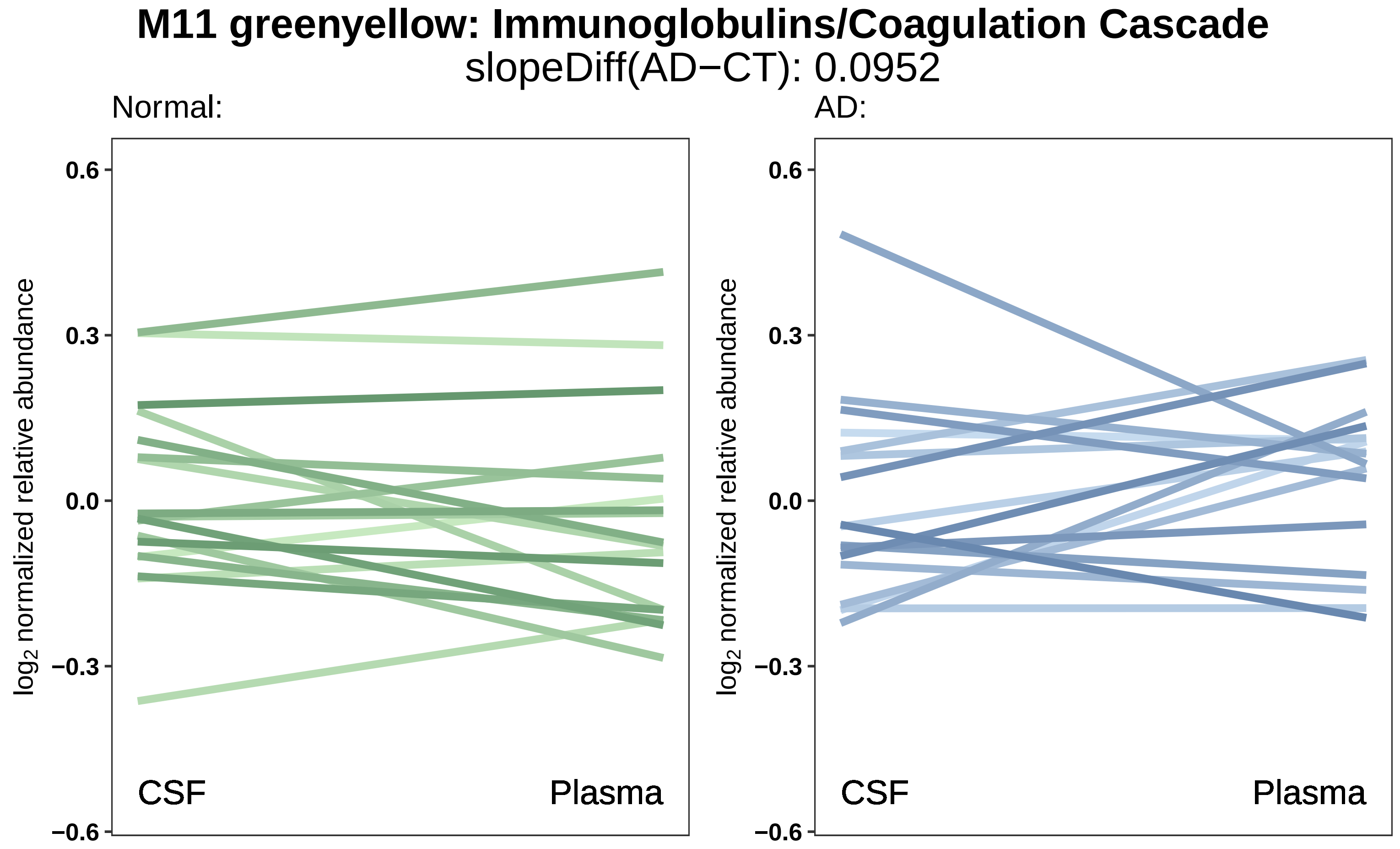

M11 greenyellow: Immunoglobulins/Coagulation Cascade
slopeDiff(AD−CT): 0.0952
Normal:
AD:
0.6
0.3
0.0
−0.3
−0.6
log2 normalized relative abundance
CSF
Plasma

### M9 magenta: Leukocyte Chemotaxis

slopeDiff(AD−CT): −0.0861

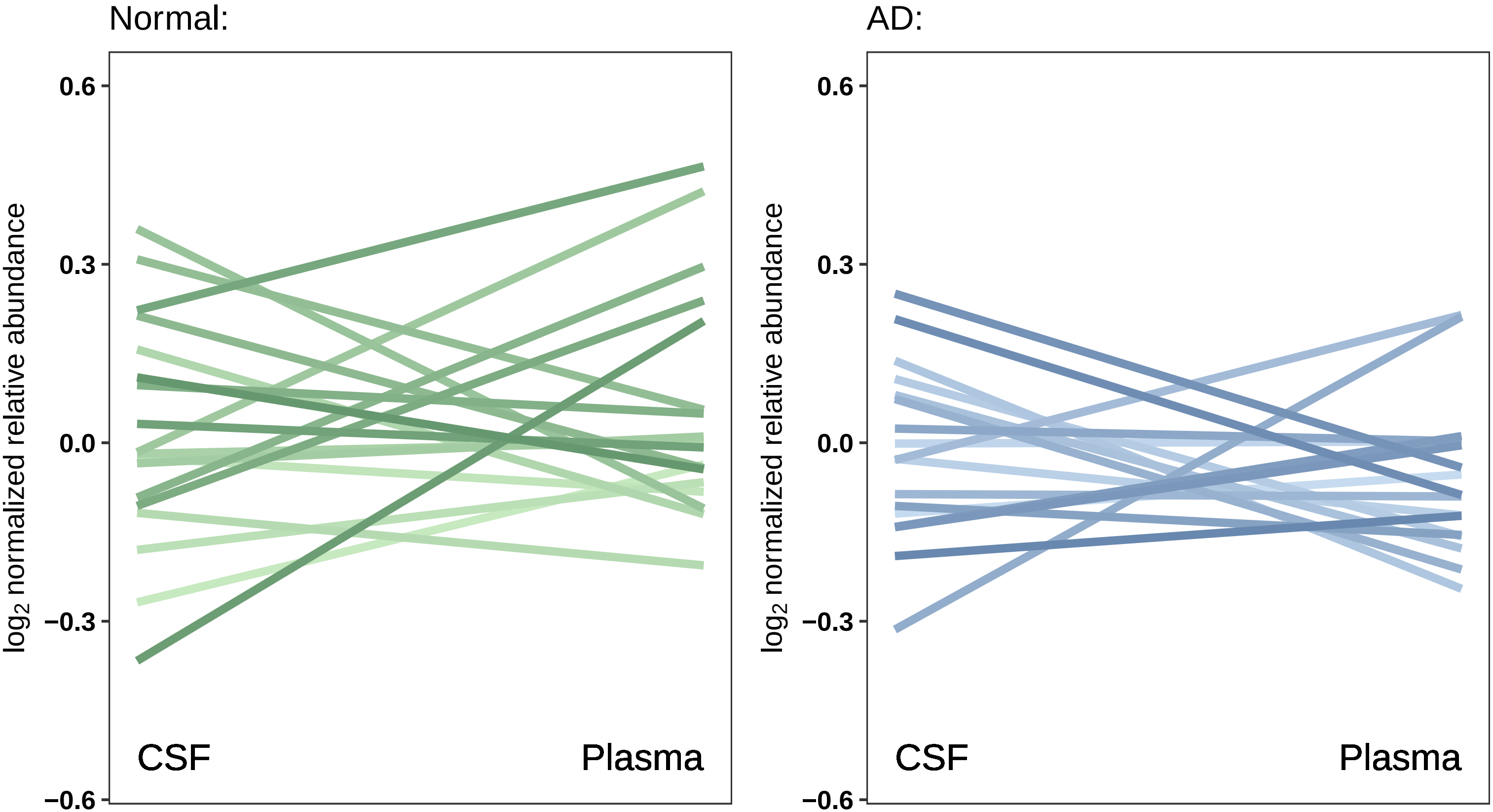

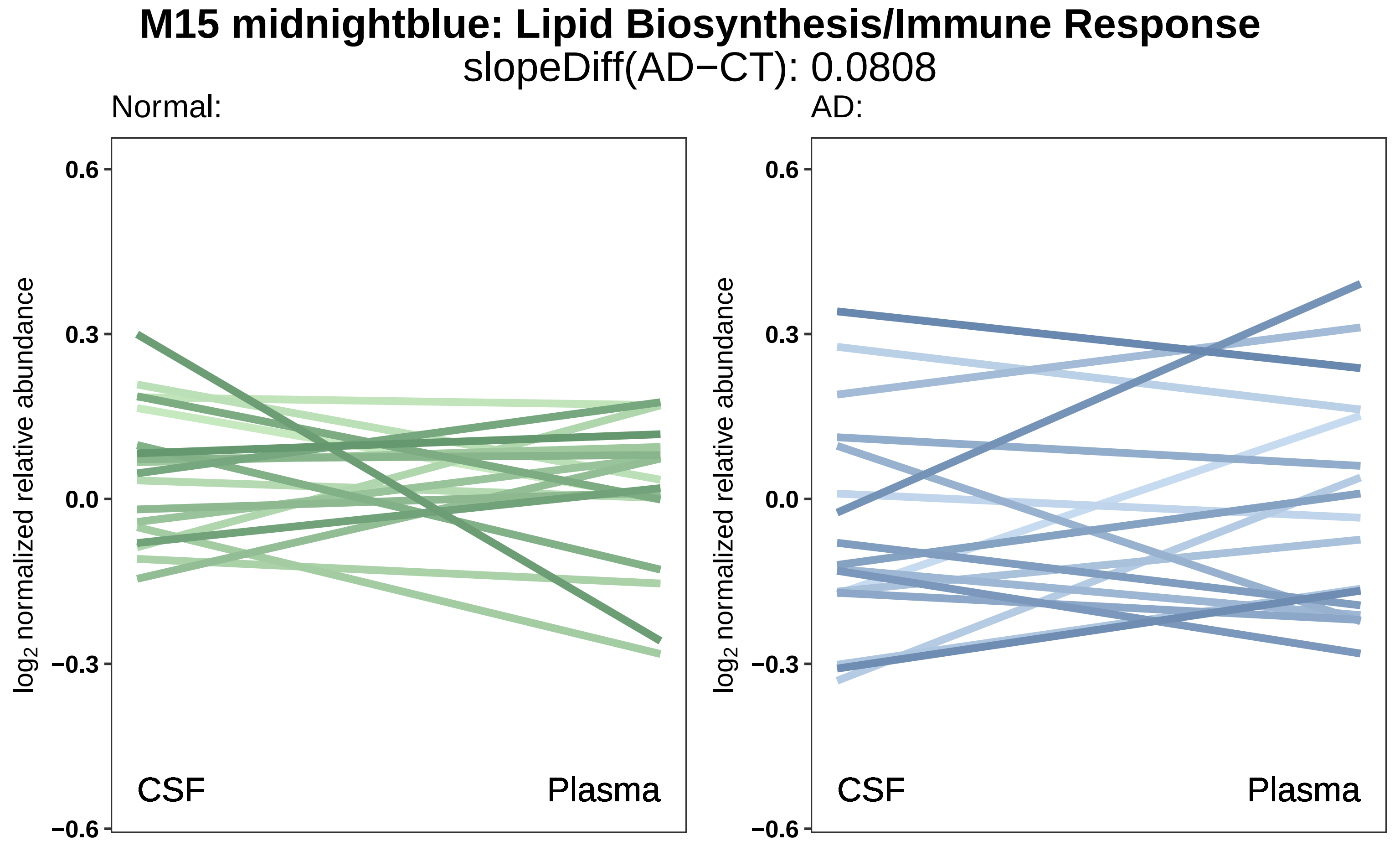
M15 midnightblue: Lipid Biosynthesis/Immune Response
slopeDiff(AD−CT): 0.0808
Normal:
AD:
0.6
0.3
0.0
−0.3
−0.6
log2 normalized relative abundance
CSF
Plasma

### M17 grey60: Glycosylation

slopeDiff(AD−CT): −0.0706

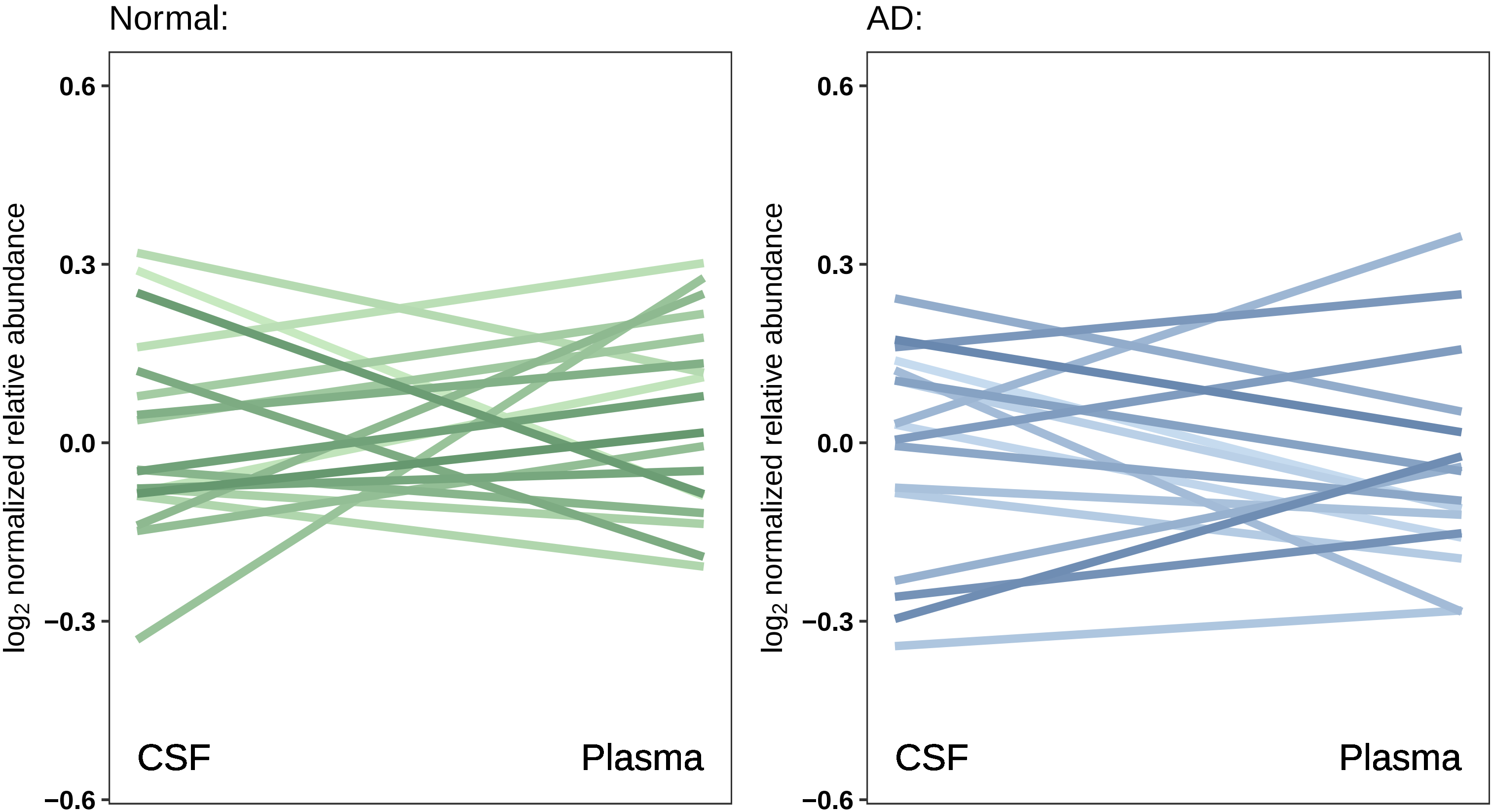

### M23 darkturquoise: Carbohydrate Binding/ECM

slopeDiff(AD−CT): 0.0583

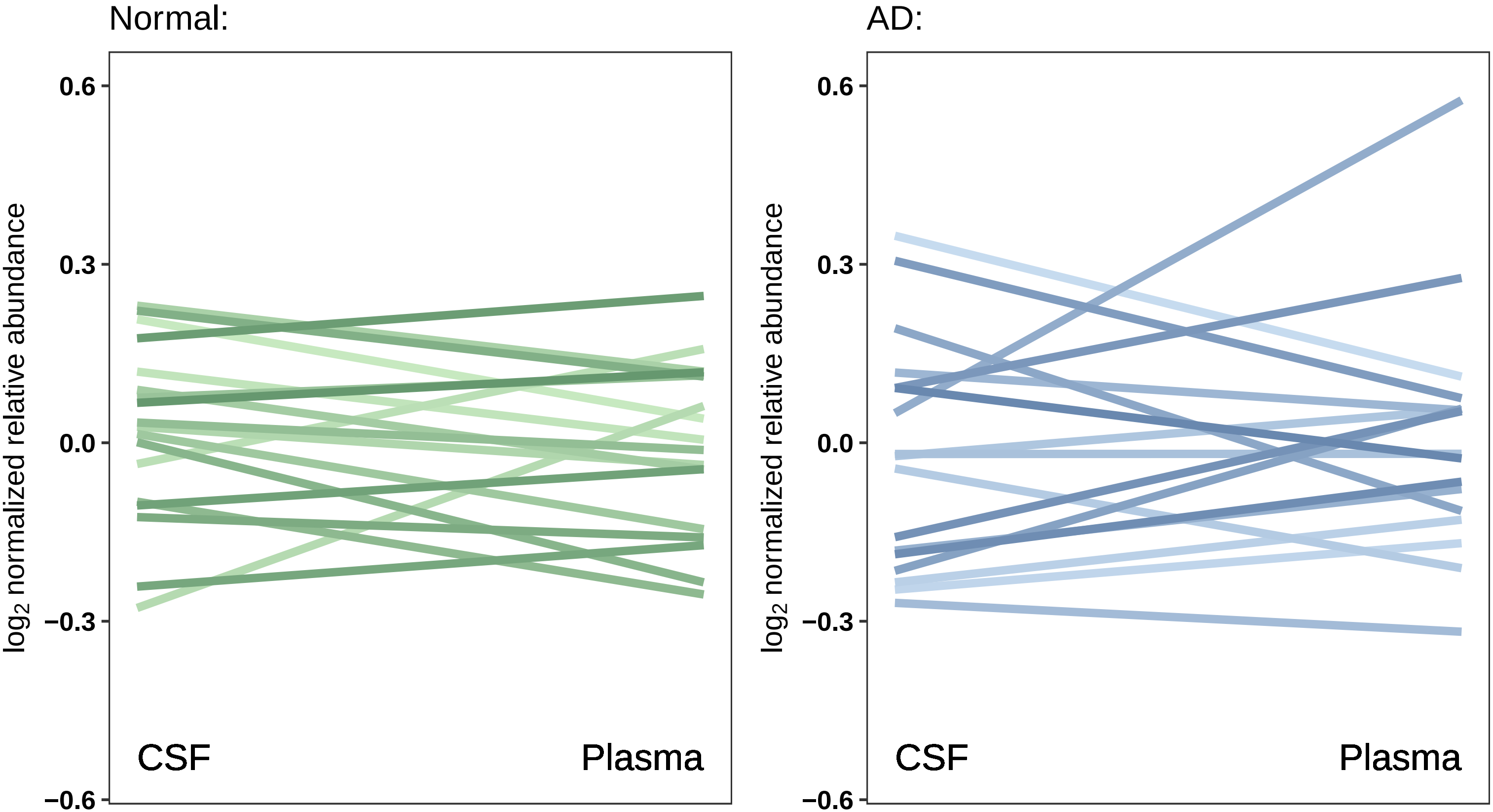

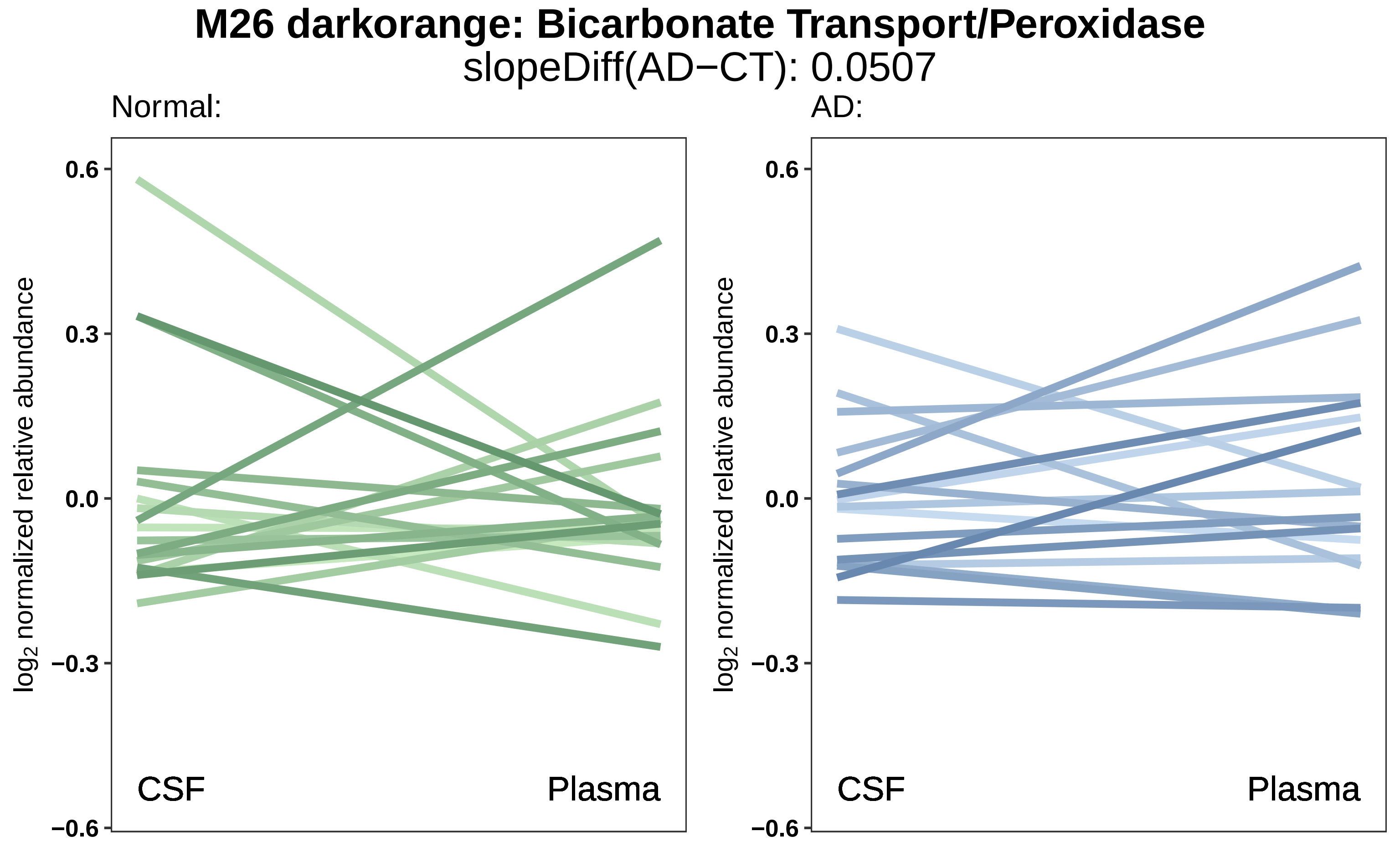
M26 darkorange: Bicarbonate Transport/Peroxidase
slopeDiff(AD−CT): 0.0507
Normal:
AD:
0.6
0.3
0.0
−0.3
−0.6
log2 normalized relative abundance
CSF
Plasma

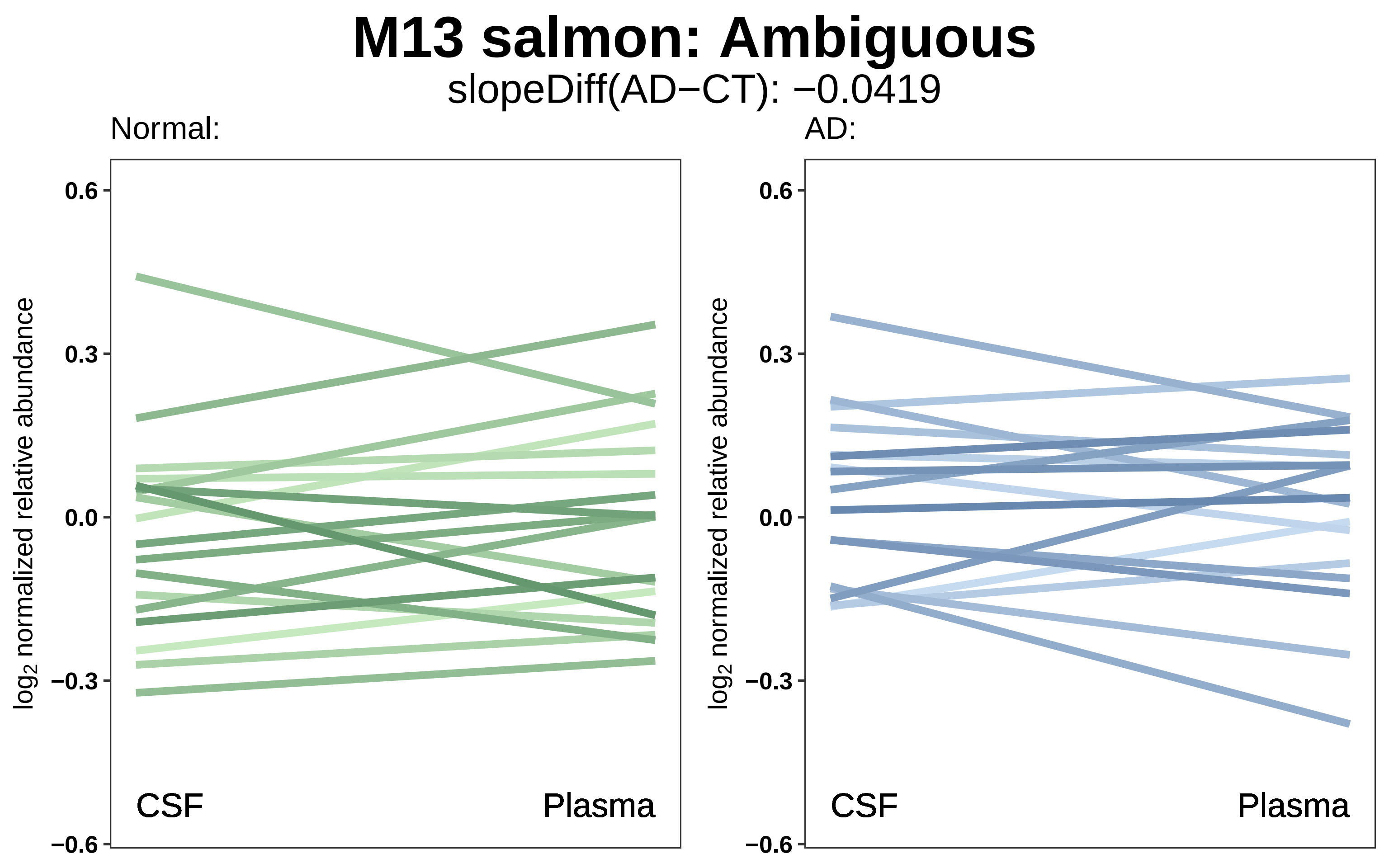
M13 salmon: Ambiguous
slopeDiff(AD−CT): −0.0419
Normal:
AD:
0.6
0.3
0.0
−0.3
−0.6
log2 normalized relative abundance
CSF
Plasma

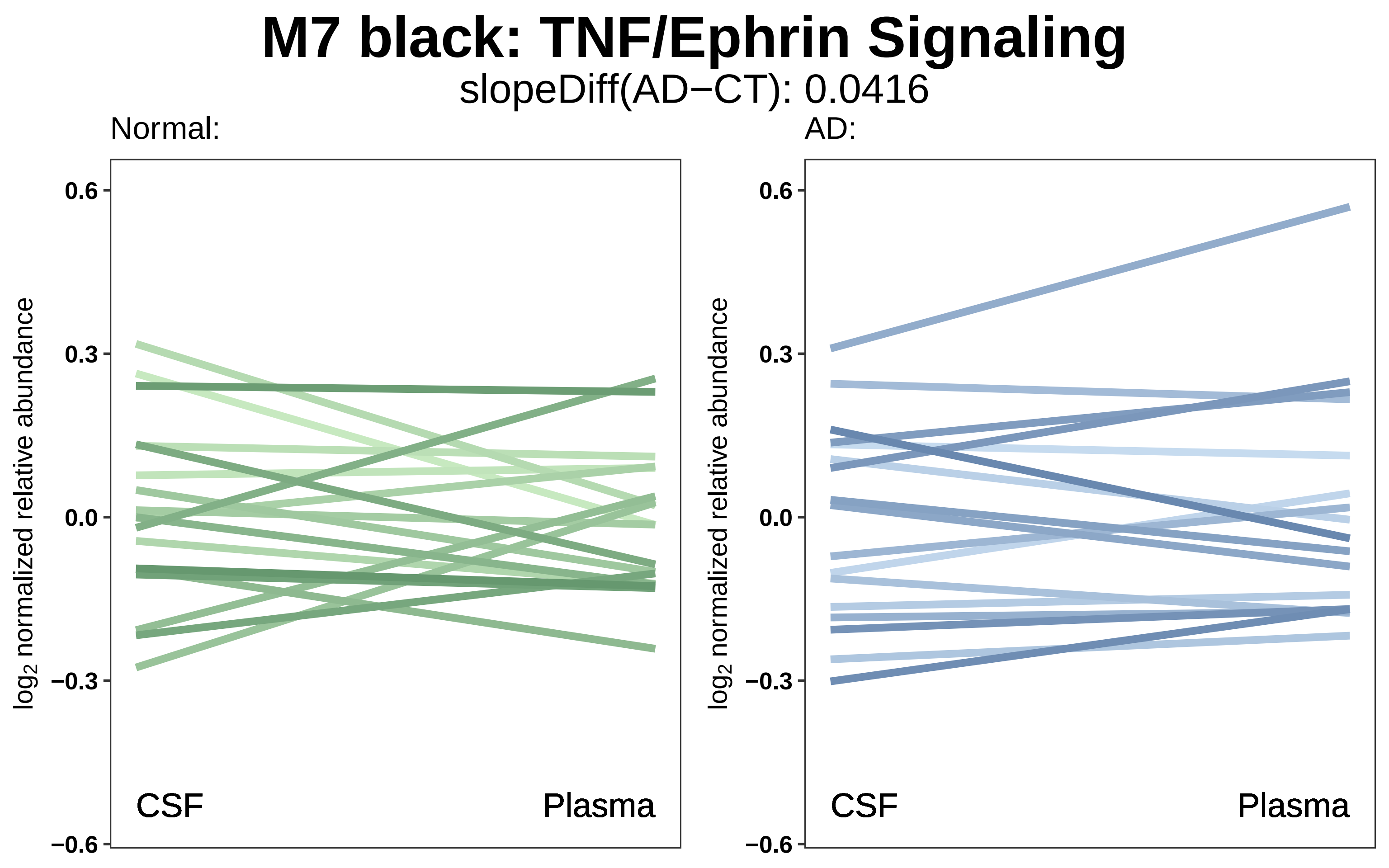
M7 black: TNF/Ephrin Signaling
slopeDiff(AD−CT): 0.0416
Normal:
AD:
log2 normalized relative abundance
0.6
0.3
0.0
−0.3
−0.6
CSF
Plasma

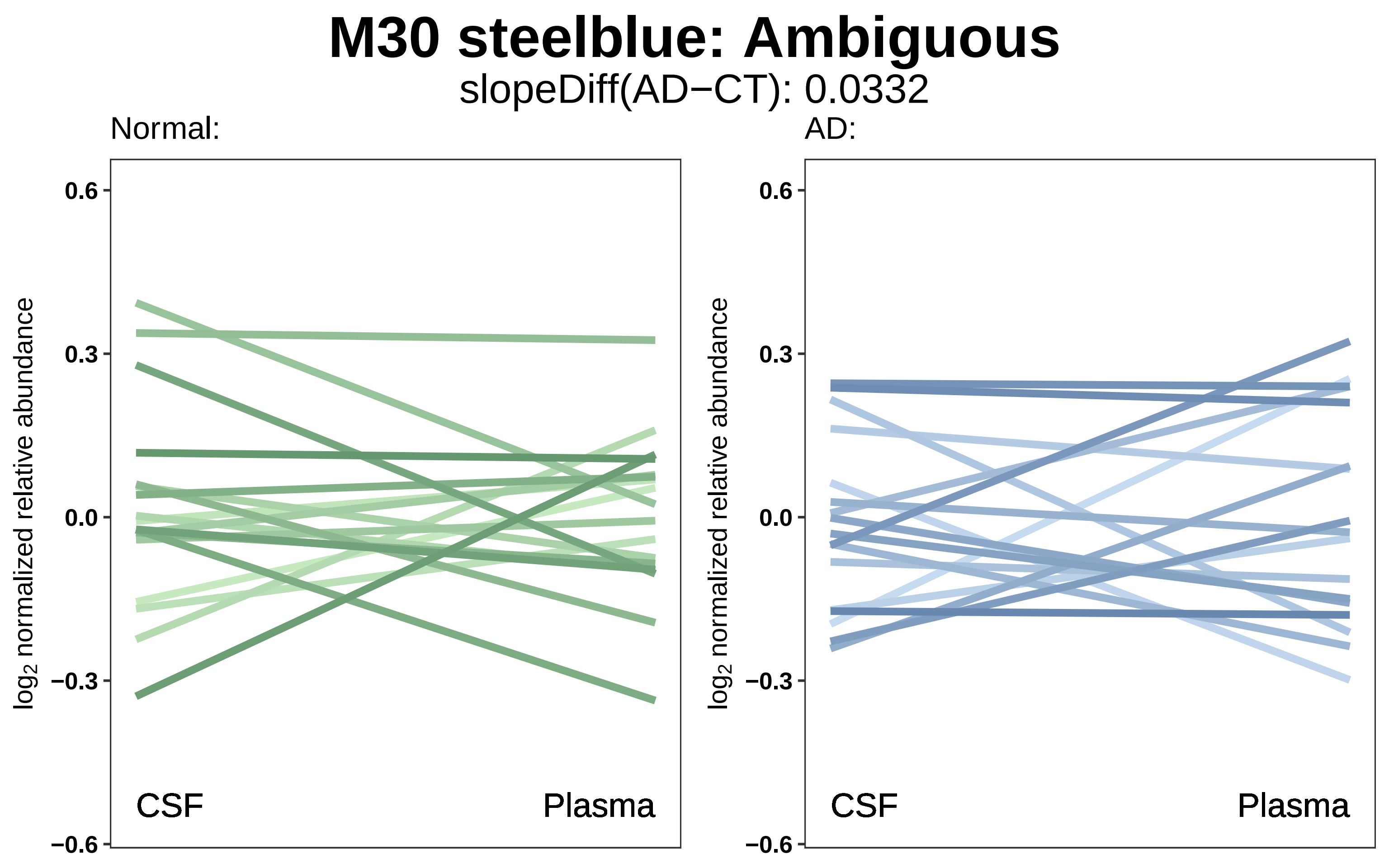

M30 steelblue: Ambiguous
slopeDiff(AD−CT): 0.0332
Normal:
AD:
0.6
0.3
0.0
−0.3
−0.6
log2 normalized relative abundance
CSF
Plasma

### M31 paleturquoise: Adhesion/ECM

slopeDiff(AD−CT): 0.0308

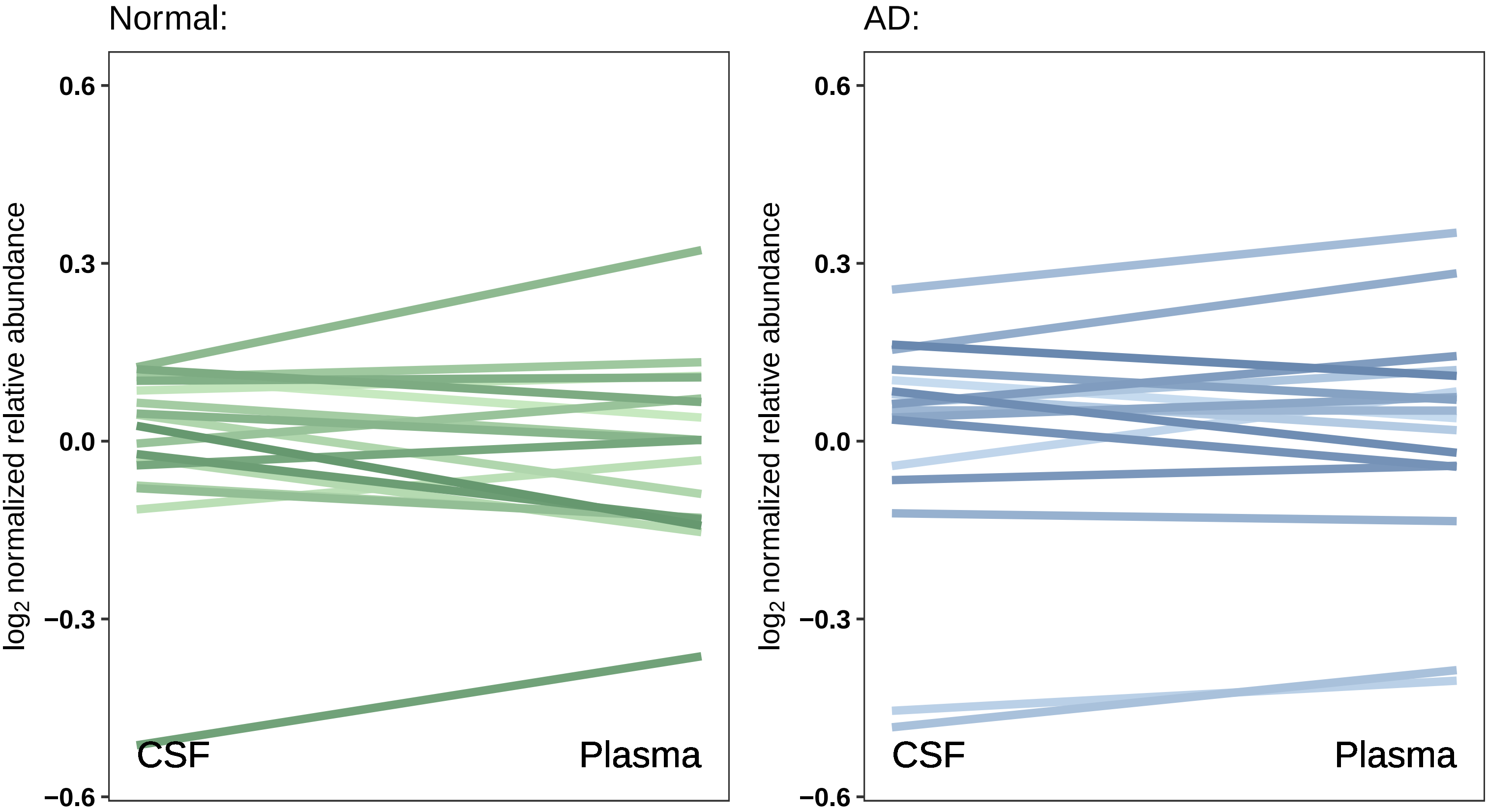

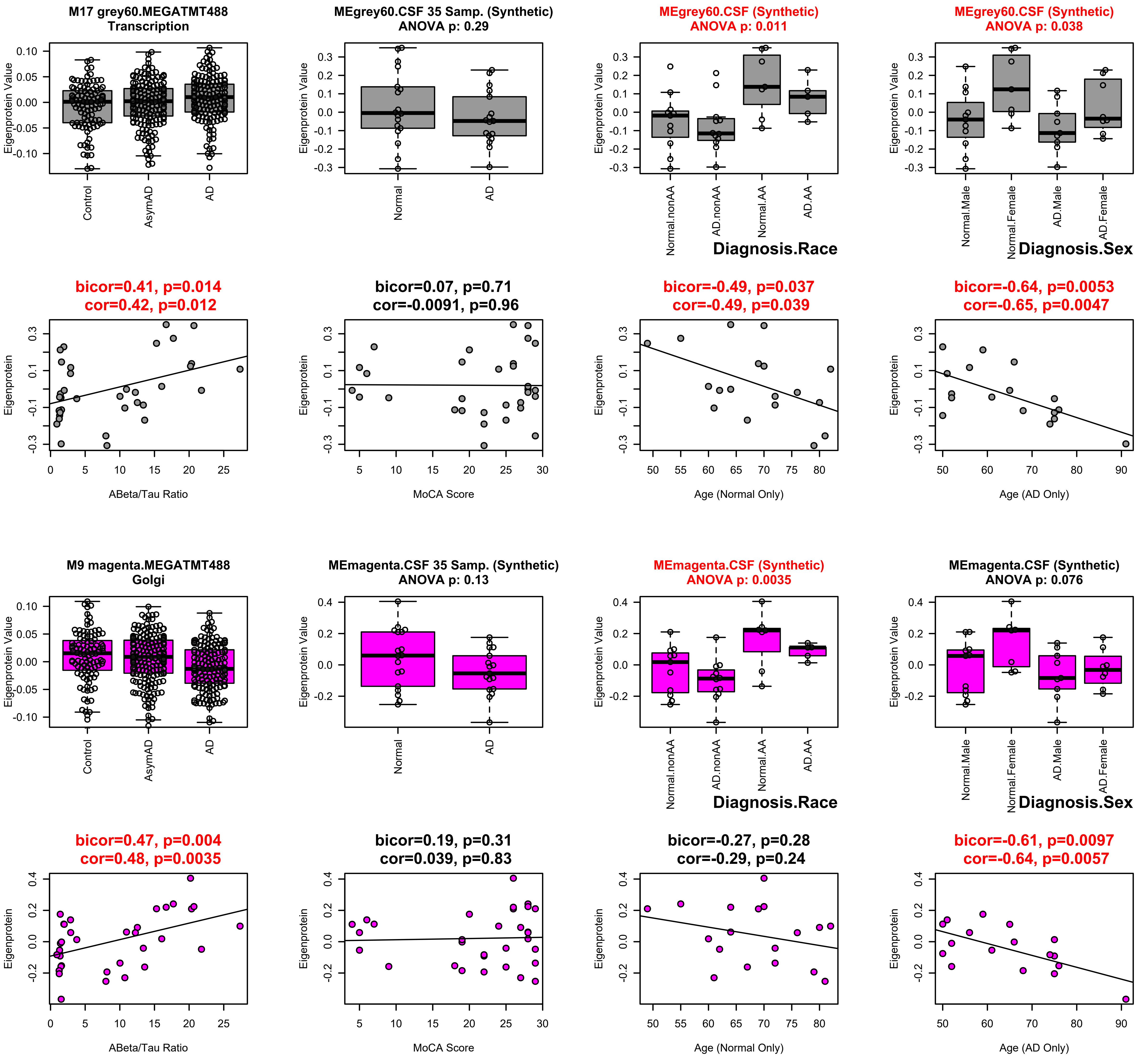
M17 grey60.MEGATMT488
Transcription
Eigenprotein Value
Control
AsymAD
AD
MEgrey60.CSF 35 Samp. (Synthetic)
ANOVA p: 0.29
Normal
AD
MEgrey60.CSF (Synthetic)
ANOVA p: 0.011
Normal.nonAA
AD.nonAA
Normal.AA
AD.AA
Diagnosis.Race
MEgrey60.CSF (Synthetic)
ANOVA p: 0.038
Normal.Male
Normal.Female
AD.Male
AD.Female
Diagnosis.Sex
bicor=0.41, p=0.014
cor=0.42, p=0.012
Eigenprotein
ABeta/Tau Ratio
bicor=0.07, p=0.71
cor=-0.0091, p=0.96
MoCA Score
bicor=-0.49, p=0.037
cor=-0.49, p=0.039
Age (Normal Only)
bicor=-0.64, p=0.0053
cor=-0.65, p=0.0047
Age (AD Only)
M9 magenta.MEGATMT488
Golgi
Eigenprotein Value
Control
AsymAD
AD
MEmagenta.CSF 35 Samp. (Synthetic)
ANOVA p: 0.13
Normal
AD
MEmagenta.CSF (Synthetic)
ANOVA p: 0.0035
Normal.nonAA
AD.nonAA
Normal.AA
AD.AA
Diagnosis.Race
MEmagenta.CSF (Synthetic)
ANOVA p: 0.076
Normal.Male
Normal.Female
AD.Male
AD.Female
Diagnosis.Sex
bicor=0.47, p=0.004
cor=0.48, p=0.0035
Eigenprotein
ABeta/Tau Ratio
bicor=0.19, p=0.31
cor=0.039, p=0.83
MoCA Score
bicor=-0.27, p=0.28
cor=-0.29, p=0.24
Age (Normal Only)
bicor=-0.61, p=0.0097
cor=-0.64, p=0.0057
Age (AD Only)

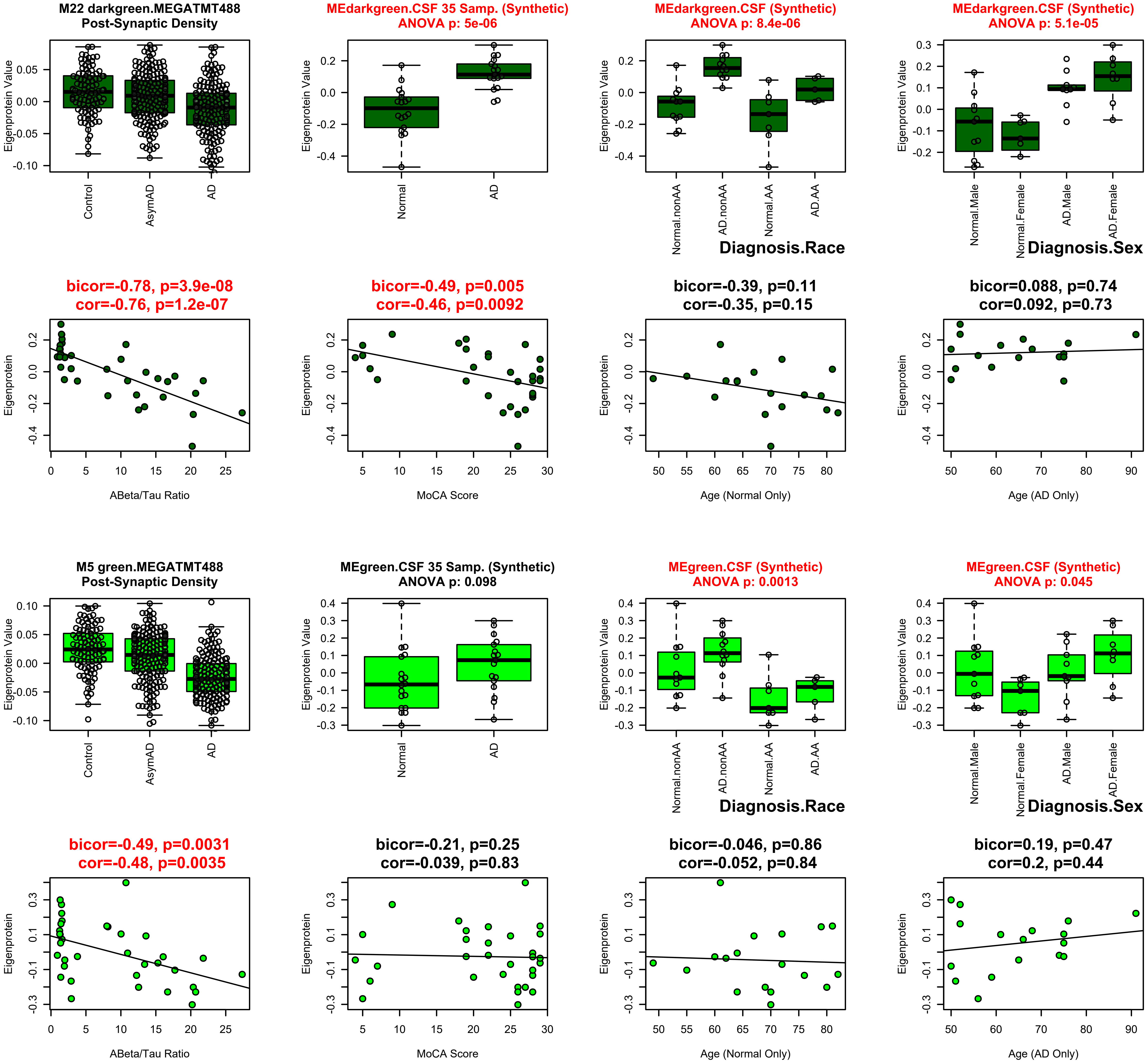

M22 darkgreen.MEGATMT488
Post-Synaptic Density
Eigenprotein Value
Control
AsymAD
AD
MEdarkgreen.CSF 35 Samp. (Synthetic)
ANOVA p: 5e-06
Normal
AD
MEdarkgreen.CSF (Synthetic)
ANOVA p: 8.4e-06
Normal.nonAA
AD.nonAA
Normal.AA
AD.AA
Diagnosis.Race
MEdarkgreen.CSF (Synthetic)
ANOVA p: 5.1e-05
Normal.Male
Normal.Female
AD.Male
AD.Female
Diagnosis.Sex
bicor=-0.78, p=3.9e-08
cor=-0.76, p=1.2e-07
Eigenprotein
ABeta/Tau Ratio
bicor=-0.49, p=0.005
cor=-0.46, p=0.0092
MoCA Score
bicor=-0.39, p=0.11
cor=-0.35, p=0.15
Age (Normal Only)
bicor=0.088, p=0.74
cor=0.092, p=0.73
Age (AD Only)
M5 green.MEGATMT488
Post-Synaptic Density
Eigenprotein Value
Control
AsymAD
AD
MEgreen.CSF 35 Samp. (Synthetic)
ANOVA p: 0.098
Normal
AD
MEgreen.CSF (Synthetic)
ANOVA p: 0.0013
Normal.nonAA
AD.nonAA
Normal.AA
AD.AA
Diagnosis.Race
MEgreen.CSF (Synthetic)
ANOVA p: 0.045
Normal.Male
Normal.Female
AD.Male
AD.Female
Diagnosis.Sex
bicor=-0.49, p=0.0031
cor=-0.48, p=0.0035
Eigenprotein
ABeta/Tau Ratio
bicor=-0.21, p=0.25
cor=-0.039, p=0.83
MoCA Score
bicor=-0.046, p=0.86
cor=-0.052, p=0.84
Age (Normal Only)
bicor=0.19, p=0.47
cor=0.2, p=0.44
Age (AD Only)

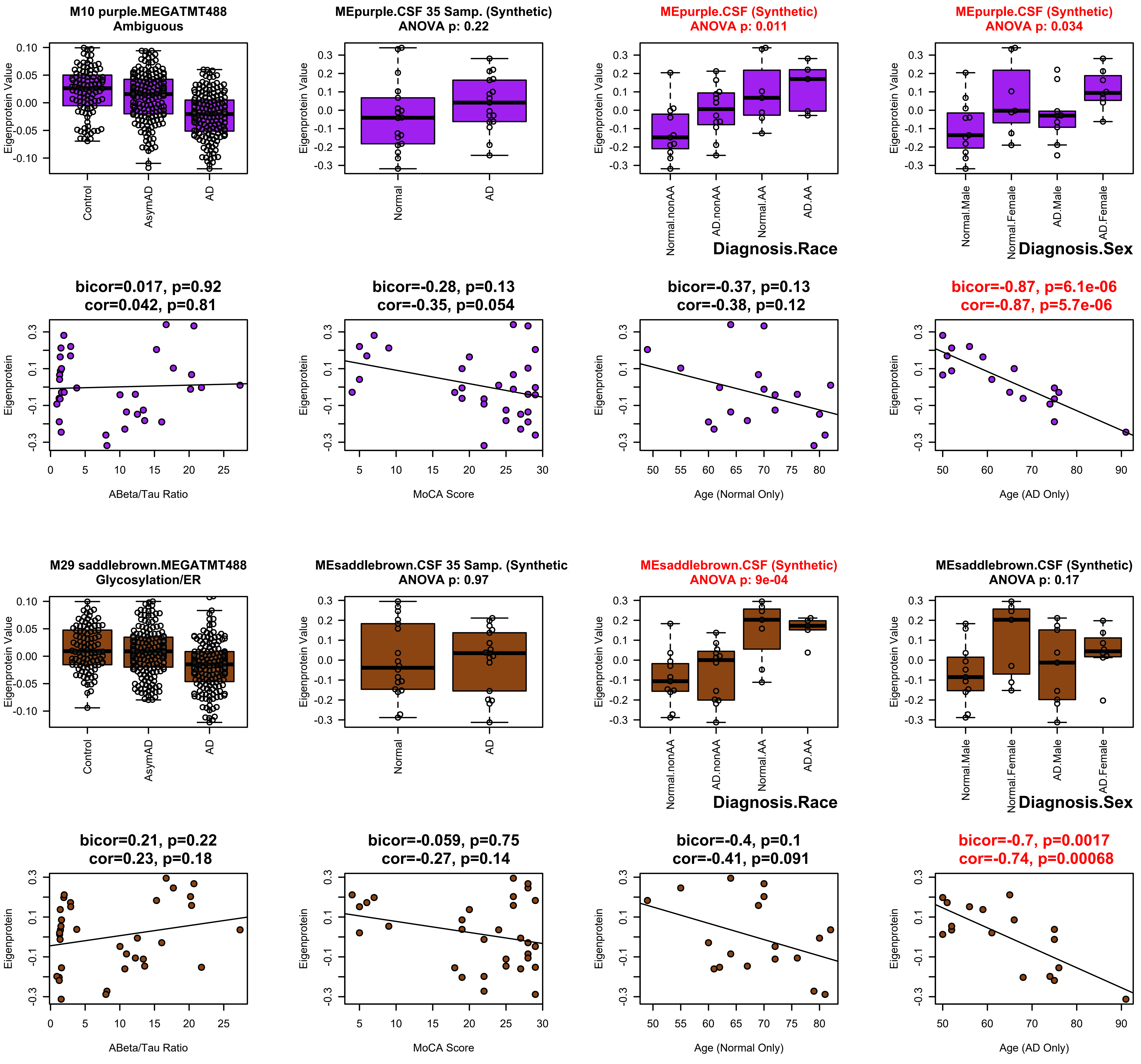
M10 purple.MEGATMT488
Ambiguous
Eigenprotein Value
Control
AsymAD
AD
MEpurple.CSF 35 Samp. (Synthetic)
ANOVA p: 0.22
Normal
AD
MEpurple.CSF (Synthetic)
ANOVA p: 0.011
Normal.nonAA
AD.nonAA
Normal.AA
AD.AA
Diagnosis.Race
MEpurple.CSF (Synthetic)
ANOVA p: 0.034
Normal.Male
Normal.Female
AD.Male
AD.Female
Diagnosis.Sex
bicor=0.017, p=0.92
cor=0.042, p=0.81
Eigenprotein
ABeta/Tau Ratio
bicor=-0.28, p=0.13
cor=-0.35, p=0.054
MoCA Score
bicor=-0.37, p=0.13
cor=-0.38, p=0.12
Age (Normal Only)
bicor=-0.87, p=6.1e-06
cor=-0.87, p=5.7e-06
Age (AD Only)
M29 saddlebrown.MEGATMT488
Glycosylation/ER
MEsaddlebrown.CSF 35 Samp. (Synthetic
ANOVA p: 0.97
MEsaddlebrown.CSF (Synthetic)
ANOVA p: 9e-04
MEsaddlebrown.CSF (Synthetic)
ANOVA p: 0.17
bicor=0.21, p=0.22
cor=0.23, p=0.18
bicor=-0.059, p=0.75
cor=-0.27, p=0.14
bicor=-0.4, p=0.1
cor=-0.41, p=0.091
bicor=-0.7, p=0.0017
cor=-0.74, p=0.00068

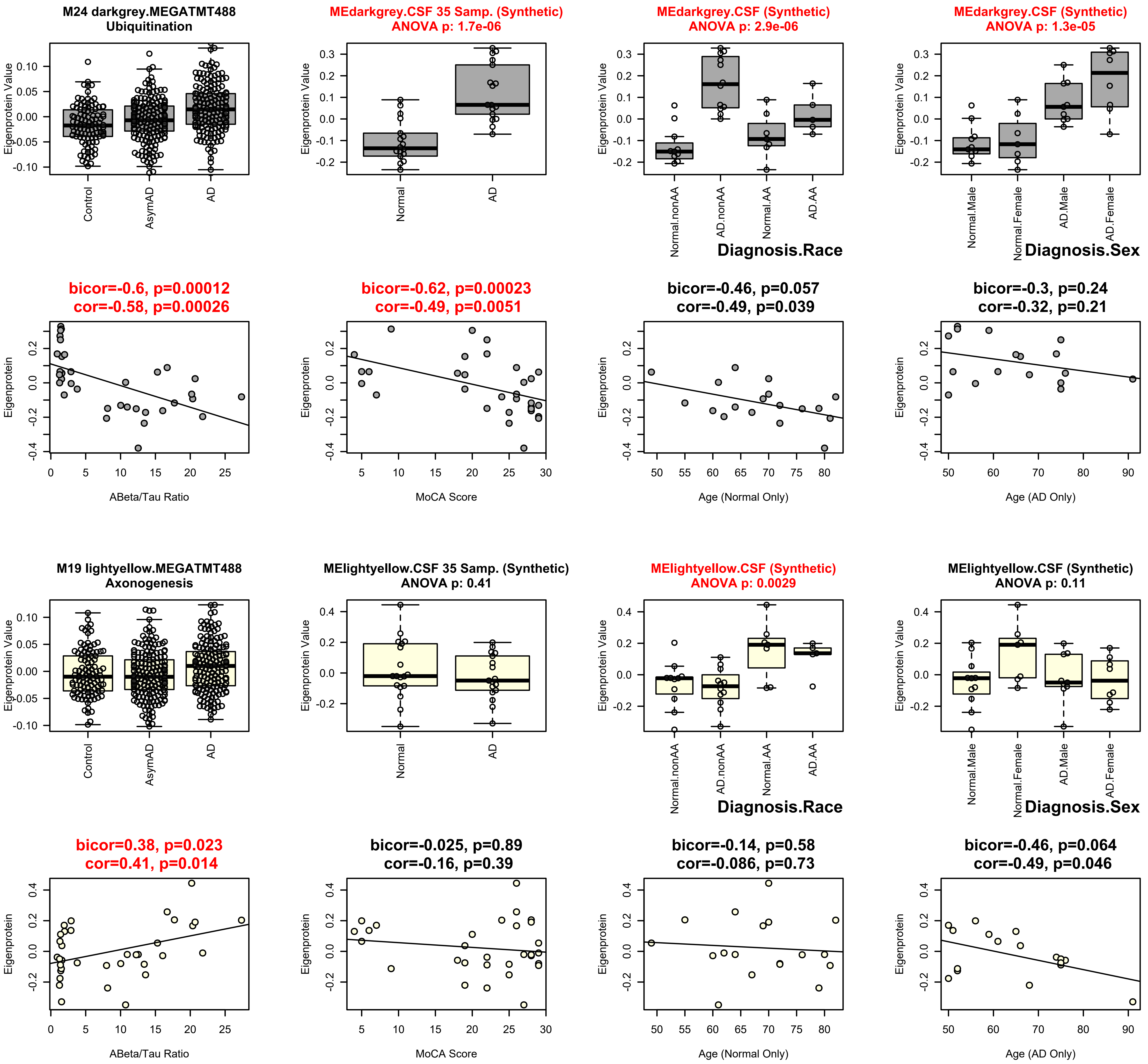

M24 darkgrey.MEGATMT488
Ubiquitination
MEdarkgrey.CSF 35 Samp. (Synthetic)
ANOVA p: 1.7e-06
MEdarkgrey.CSF (Synthetic)
ANOVA p: 2.9e-06
MEdarkgrey.CSF (Synthetic)
ANOVA p: 1.3e-05
Eigenprotein Value
Control
AsymAD
AD
Normal
Normal.nonAA
AD.nonAA
Normal.AA
AD.AA
Diagnosis.Race
Normal.Male
Normal.Female
AD.Male
AD.Female
Diagnosis.Sex
bicor=-0.6, p=0.00012
cor=-0.58, p=0.00026
bicor=-0.62, p=0.00023
cor=-0.49, p=0.0051
bicor=-0.46, p=0.057
cor=-0.49, p=0.039
bicor=-0.3, p=0.24
cor=-0.32, p=0.21
Eigenprotein
ABeta/Tau Ratio
MoCA Score
Age (Normal Only)
Age (AD Only)
M19 lightyellow.MEGATMT488
Axonogenesis
MElightyellow.CSF 35 Samp. (Synthetic)
ANOVA p: 0.41
MElightyellow.CSF (Synthetic)
ANOVA p: 0.0029
MElightyellow.CSF (Synthetic)
ANOVA p: 0.11
Eigenprotein Value
Control
AsymAD
AD
Normal
Normal.nonAA
AD.nonAA
Normal.AA
AD.AA
Diagnosis.Race
Normal.Male
Normal.Female
AD.Male
AD.Female
Diagnosis.Sex
bicor=0.38, p=0.023
cor=0.41, p=0.014
bicor=-0.025, p=0.89
cor=-0.16, p=0.39
bicor=-0.14, p=0.58
cor=-0.086, p=0.73
bicor=-0.46, p=0.064
cor=-0.49, p=0.046
Eigenprotein
ABeta/Tau Ratio
MoCA Score
Age (Normal Only)
Age (AD Only)

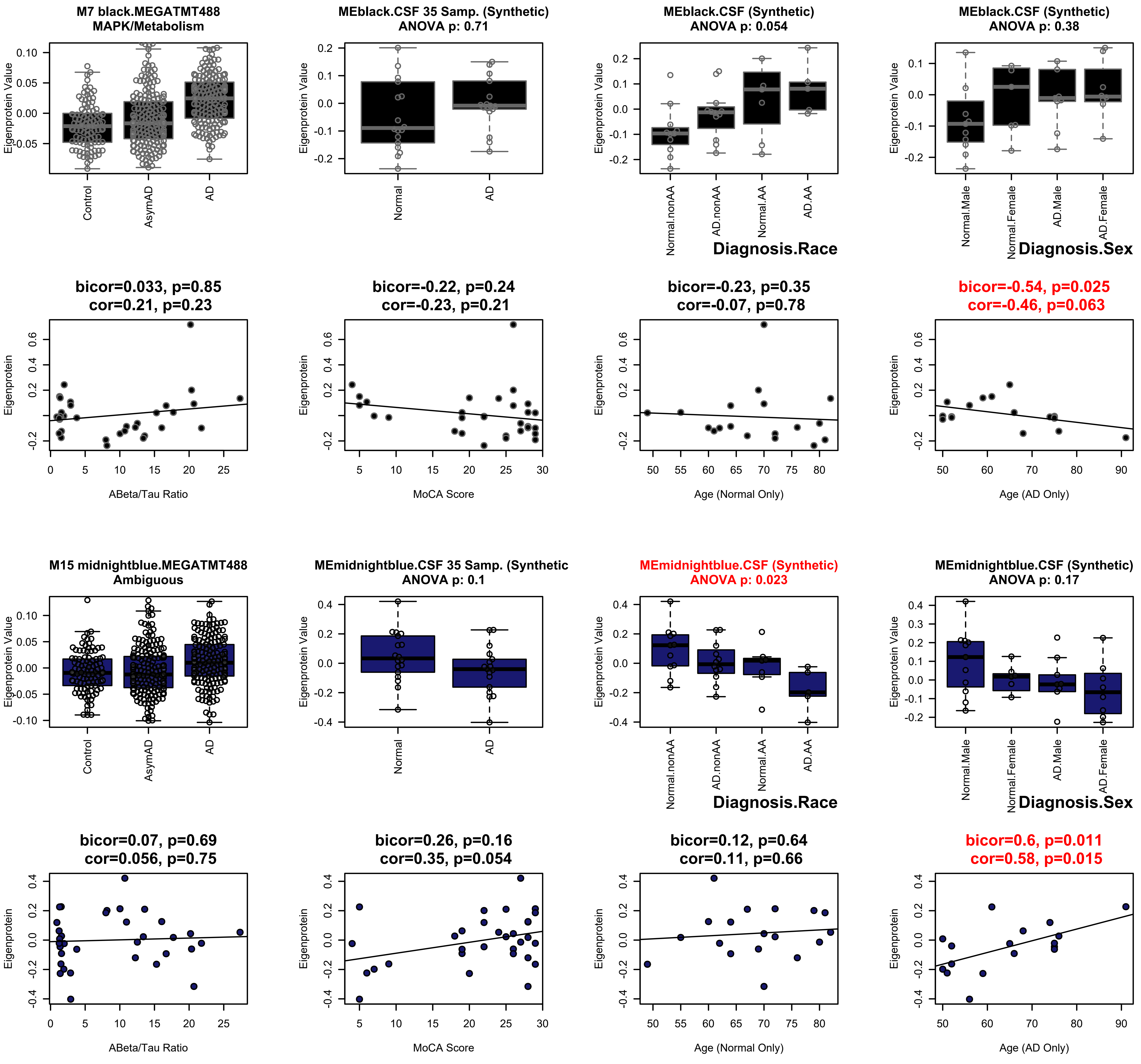

M7 black.MEGATMT488
MAPK/Metabolism
MEblack.CSF 35 Samp. (Synthetic)
ANOVA p: 0.71
MEblack.CSF (Synthetic)
ANOVA p: 0.054
MEblack.CSF (Synthetic)
ANOVA p: 0.38
Eigenprotein Value
Control
AsymAD
AD
Normal
Normal.nonAA
AD.nonAA
Normal.AA
AD.AA
Diagnosis.Race
Normal.Male
Normal.Female
AD.Male
AD.Female
Diagnosis.Sex
bicor=0.033, p=0.85
cor=0.21, p=0.23
bicor=-0.22, p=0.24
cor=-0.23, p=0.21
bicor=-0.23, p=0.35
cor=-0.07, p=0.78
bicor=-0.54, p=0.025
cor=-0.46, p=0.063
Eigenprotein
ABeta/Tau Ratio
MoCA Score
Age (Normal Only)
Age (AD Only)
M15 midnightblue.MEGATMT488
Ambiguous
MEmidnightblue.CSF 35 Samp. (Synthetic
ANOVA p: 0.1
MEmidnightblue.CSF (Synthetic)
ANOVA p: 0.023
MEmidnightblue.CSF (Synthetic)
ANOVA p: 0.17
bicor=0.07, p=0.69
cor=0.056, p=0.75
bicor=0.26, p=0.16
cor=0.35, p=0.054
bicor=0.12, p=0.64
cor=0.11, p=0.66
bicor=0.6, p=0.011
cor=0.58, p=0.015

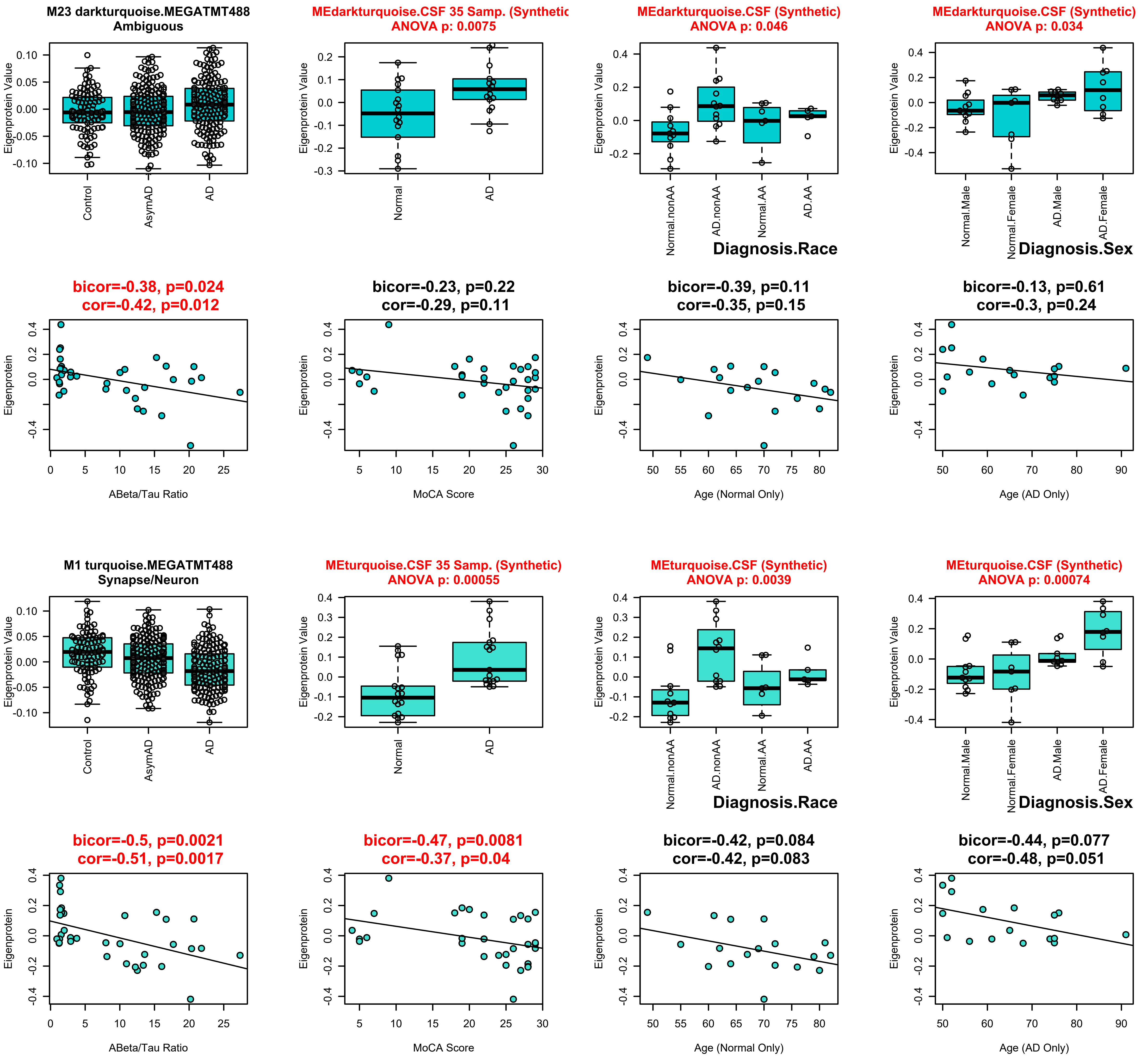
M23 darkturquoise.MEGATMT488
Ambiguous
Eigenprotein Value
Control
AsymAD
AD
MEdarkturquoise.CSF 35 Samp. (Synthetic
ANOVA p: 0.0075
Normal
AD
MEdarkturquoise.CSF (Synthetic)
ANOVA p: 0.046
Normal.nonAA
AD.nonAA
Normal.AA
AD.AA
Diagnosis.Race
MEdarkturquoise.CSF (Synthetic)
ANOVA p: 0.034
Normal.Male
Normal.Female
AD.Male
AD.Female
Diagnosis.Sex
bicor=-0.38, p=0.024
cor=-0.42, p=0.012
Eigenprotein
ABeta/Tau Ratio
bicor=-0.23, p=0.22
cor=-0.29, p=0.11
MoCA Score
bicor=-0.39, p=0.11
cor=-0.35, p=0.15
Age (Normal Only)
bicor=-0.13, p=0.61
cor=-0.3, p=0.24
Age (AD Only)
M1 turquoise.MEGATMT488
Synapse/Neuron
Eigenprotein Value
Control
AsymAD
AD
MEturquoise.CSF 35 Samp. (Synthetic)
ANOVA p: 0.00055
Normal
AD
MEturquoise.CSF (Synthetic)
ANOVA p: 0.0039
Normal.nonAA
AD.nonAA
Normal.AA
AD.AA
Diagnosis.Race
MEturquoise.CSF (Synthetic)
ANOVA p: 0.00074
Normal.Male
Normal.Female
AD.Male
AD.Female
Diagnosis.Sex
bicor=-0.5, p=0.0021
cor=-0.51, p=0.0017
Eigenprotein
ABeta/Tau Ratio
bicor=-0.47, p=0.0081
cor=-0.37, p=0.04
MoCA Score
bicor=-0.42, p=0.084
cor=-0.42, p=0.083
Age (Normal Only)
bicor=-0.44, p=0.077
cor=-0.48, p=0.051
Age (AD Only)

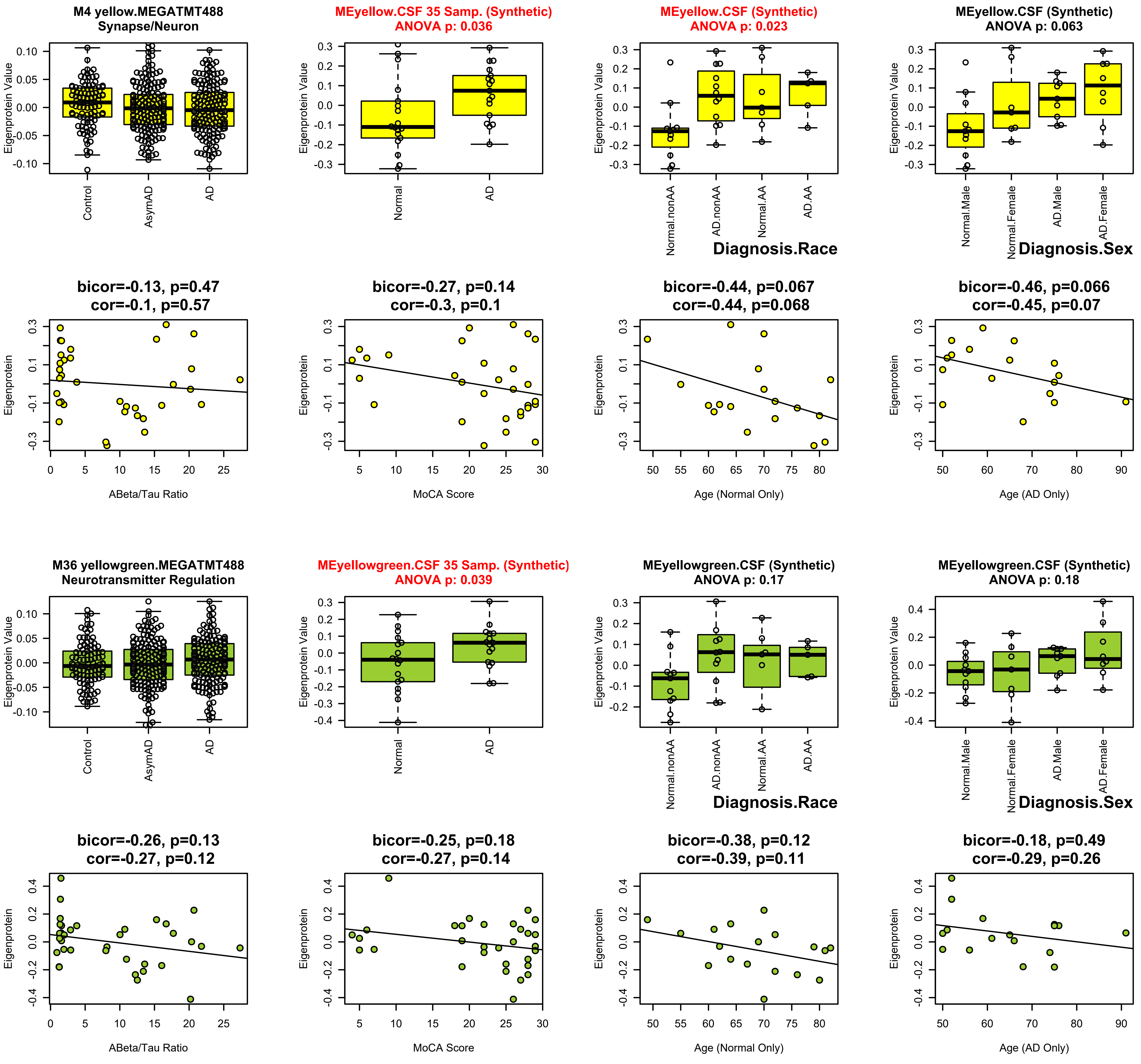
M4 yellow.MEGATMT488
Synapse/Neuron
Eigenprotein Value
Control
AsymAD
AD
MEyellow.CSF 35 Samp. (Synthetic)
ANOVA p: 0.036
Normal
AD
MEyellow.CSF (Synthetic)
ANOVA p: 0.023
Normal.nonAA
AD.nonAA
Normal.AA
AD.AA
Diagnosis.Race
MEyellow.CSF (Synthetic)
ANOVA p: 0.063
Normal.Male
Normal.Female
AD.Male
AD.Female
Diagnosis.Sex
bicor=-0.13, p=0.47
cor=-0.1, p=0.57
ABeta/Tau Ratio
Eigenprotein
bicor=-0.27, p=0.14
cor=-0.3, p=0.1
MoCA Score
bicor=-0.44, p=0.067
cor=-0.44, p=0.068
Age (Normal Only)
bicor=-0.46, p=0.066
cor=-0.45, p=0.07
Age (AD Only)
M36 yellowgreen.MEGATMT488
Neurotransmitter Regulation
MEyellowgreen.CSF 35 Samp. (Synthetic)
ANOVA p: 0.039
MEyellowgreen.CSF (Synthetic)
ANOVA p: 0.17
MEyellowgreen.CSF (Synthetic)
ANOVA p: 0.18
bicor=-0.26, p=0.13
cor=-0.27, p=0.12
bicor=-0.25, p=0.18
cor=-0.27, p=0.14
bicor=-0.38, p=0.12
cor=-0.39, p=0.11
bicor=-0.18, p=0.49
cor=-0.29, p=0.26

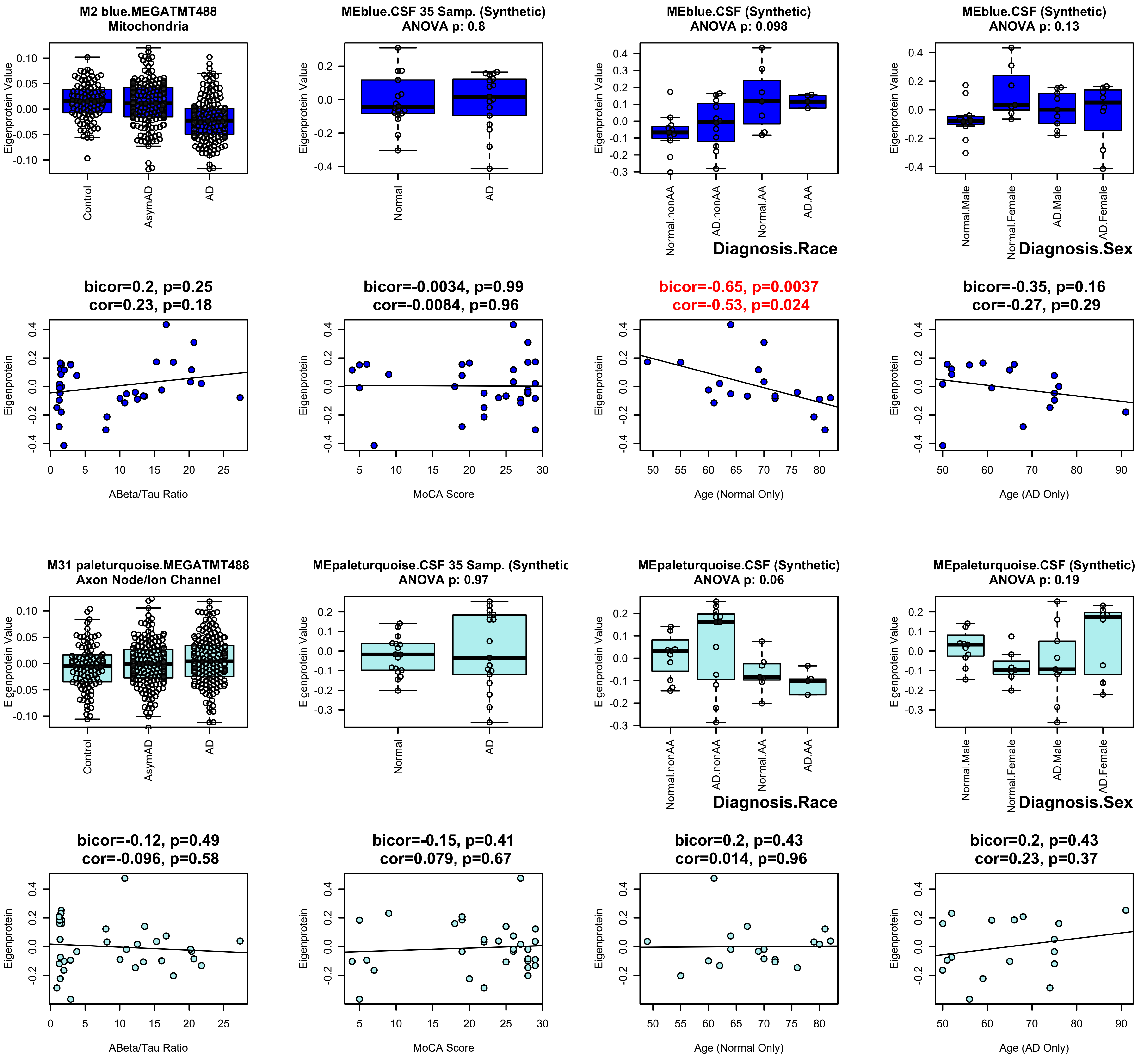
M2 blue.MEGATMT488
Mitochondria
Eigenprotein Value
Control
AsymAD
AD
MEblue.CSF 35 Samp. (Synthetic)
ANOVA p: 0.8
Normal
AD
MEblue.CSF (Synthetic)
ANOVA p: 0.098
Normal.nonAA
AD.nonAA
Normal.AA
AD.AA
Diagnosis.Race
MEblue.CSF (Synthetic)
ANOVA p: 0.13
Normal.Male
Normal.Female
AD.Male
AD.Female
Diagnosis.Sex
bicor=0.2, p=0.25
cor=0.23, p=0.18
Eigenprotein
ABeta/Tau Ratio
bicor=-0.0034, p=0.99
cor=-0.0084, p=0.96
MoCA Score
bicor=-0.65, p=0.0037
cor=-0.53, p=0.024
Age (Normal Only)
bicor=-0.35, p=0.16
cor=-0.27, p=0.29
Age (AD Only)
M31 paleturquoise.MEGATMT488
Axon Node/Ion Channel
MEpaleturquoise.CSF 35 Samp. (Synthetic
ANOVA p: 0.97
MEpaleturquoise.CSF (Synthetic)
ANOVA p: 0.06
Diagnosis.Race
MEpaleturquoise.CSF (Synthetic)
ANOVA p: 0.19
Diagnosis.Sex
bicor=-0.12, p=0.49
cor=-0.096, p=0.58
ABeta/Tau Ratio
bicor=-0.15, p=0.41
cor=0.079, p=0.67
MoCA Score
bicor=0.2, p=0.43
cor=0.014, p=0.96
Age (Normal Only)
bicor=0.2, p=0.43
cor=0.23, p=0.37
Age (AD Only)

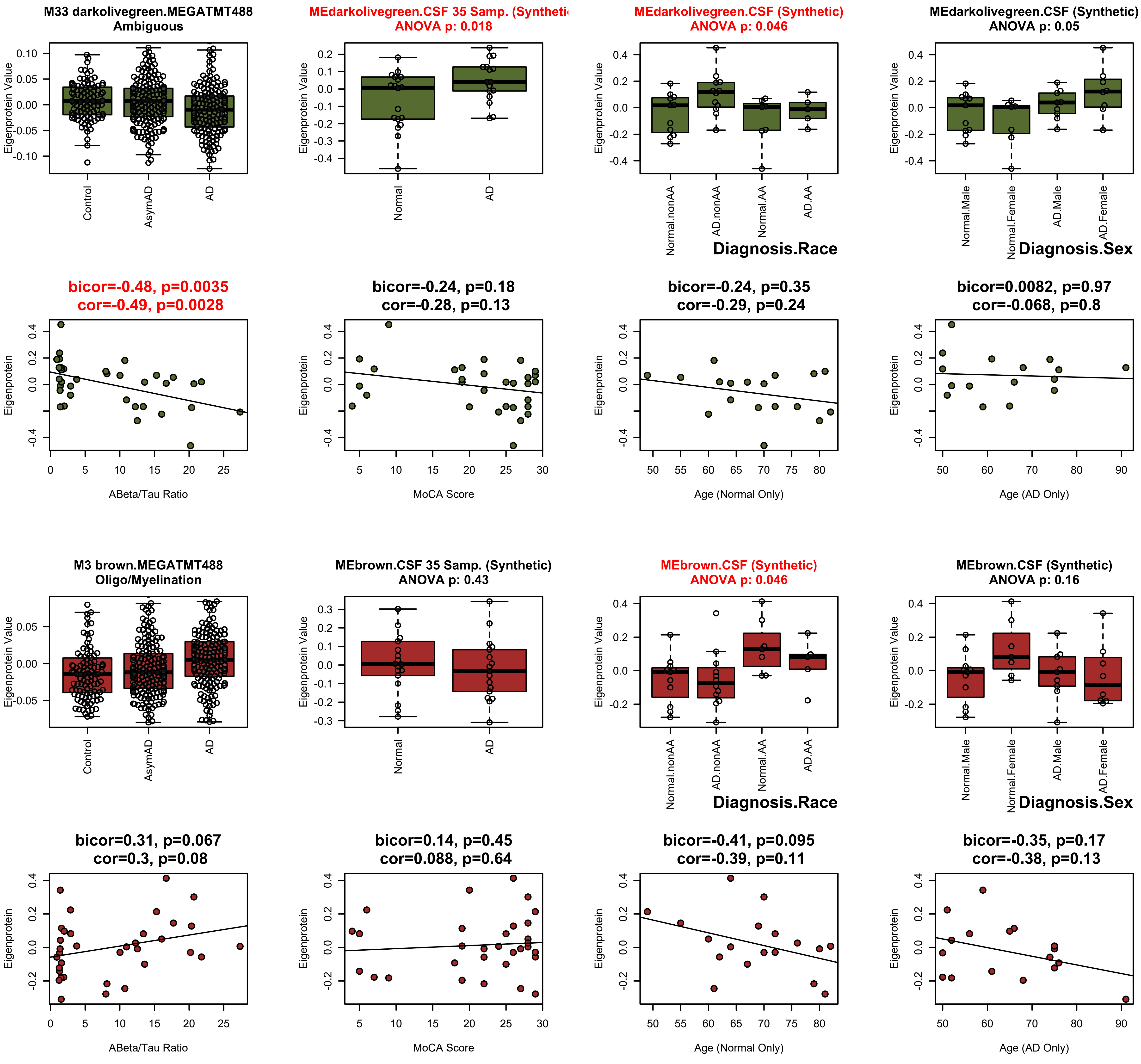

M33 darkolivegreen.MEGATMT488
Ambiguous
Eigenprotein Value
Control
AsymAD
AD
MEdarkolivegreen.CSF 35 Samp. (Syntheti
ANOVA p: 0.018
Eigenprotein Value
Normal
AD
MEdarkolivegreen.CSF (Synthetic)
ANOVA p: 0.046
Eigenprotein Value
Normal.nonAA
AD.nonAA
Normal.AA
AD.AA
Diagnosis.Race
MEdarkolivegreen.CSF (Synthetic)
ANOVA p: 0.05
Eigenprotein Value
Normal.Male
Normal.Female
AD.Male
AD.Female
Diagnosis.Sex
bicor=-0.48, p=0.0035
cor=-0.49, p=0.0028
Eigenprotein
ABeta/Tau Ratio
bicor=-0.24, p=0.18
cor=-0.28, p=0.13
Eigenprotein
MoCA Score
bicor=-0.24, p=0.35
cor=-0.29, p=0.24
Eigenprotein
Age (Normal Only)
bicor=0.0082, p=0.97
cor=-0.068, p=0.8
Eigenprotein
Age (AD Only)
M3 brown.MEGATMT488
Oligo/Myelination
Eigenprotein Value
Control
AsymAD
AD
MEbrown.CSF 35 Samp. (Synthetic)
ANOVA p: 0.43
Eigenprotein Value
Normal
AD
MEbrown.CSF (Synthetic)
ANOVA p: 0.046
Eigenprotein Value
Normal.nonAA
AD.nonAA
Normal.AA
AD.AA
Diagnosis.Race
MEbrown.CSF (Synthetic)
ANOVA p: 0.16
Eigenprotein Value
Normal.Male
Normal.Female
AD.Male
AD.Female
Diagnosis.Sex
bicor=0.31, p=0.067
cor=0.3, p=0.08
Eigenprotein
ABeta/Tau Ratio
bicor=0.14, p=0.45
cor=0.088, p=0.64
Eigenprotein
MoCA Score
bicor=-0.41, p=0.095
cor=-0.39, p=0.11
Eigenprotein
Age (Normal Only)
bicor=-0.35, p=0.17
cor=-0.38, p=0.13
Eigenprotein
Age (AD Only)

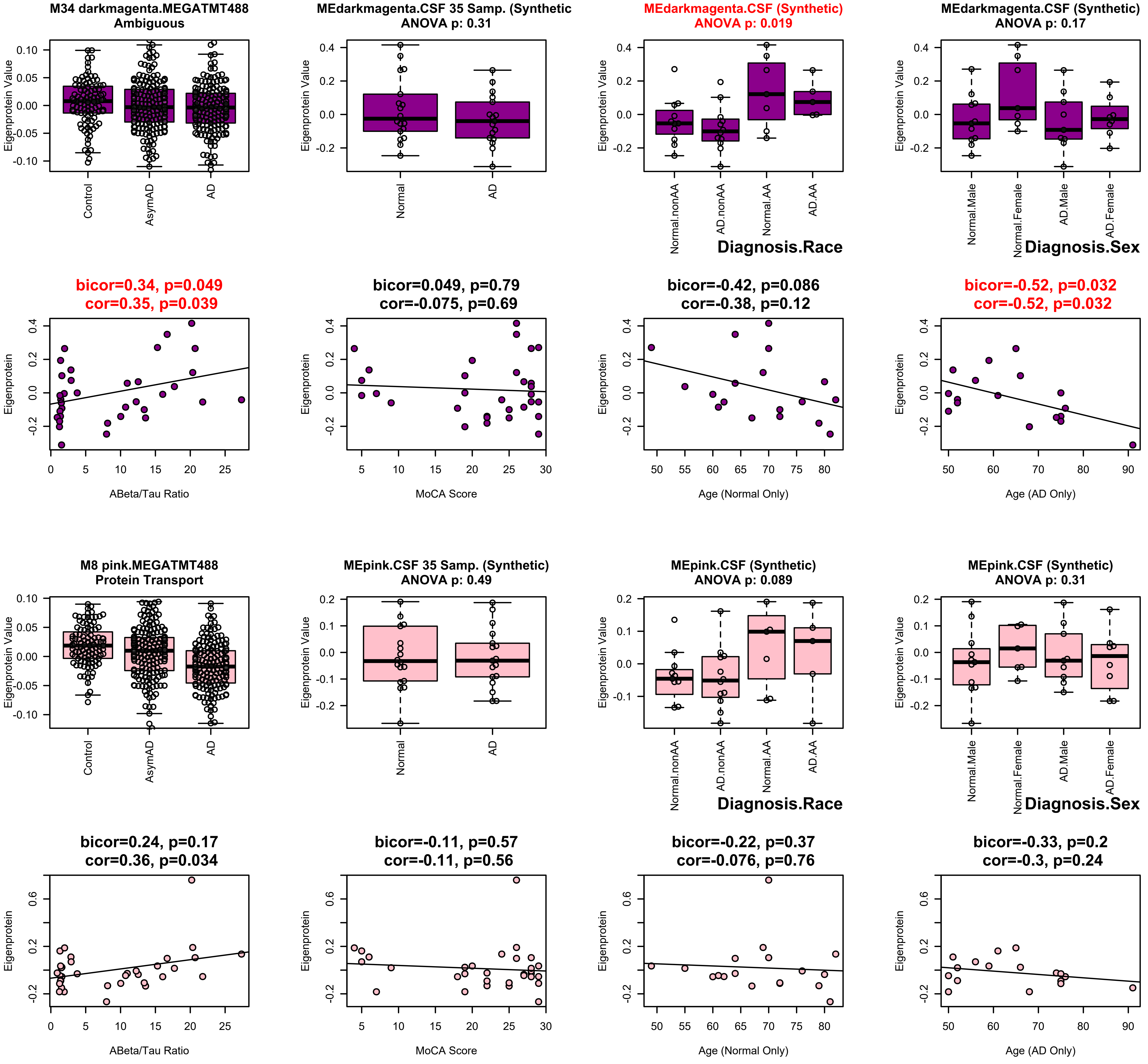

M34 darkmagenta.MEGATMT488
Ambiguous
Eigenprotein Value
Control
AsymAD
AD
MEdarkmagenta.CSF 35 Samp. (Synthetic
ANOVA p: 0.31
Eigenprotein Value
Normal
AD
MEdarkmagenta.CSF (Synthetic)
ANOVA p: 0.019
Eigenprotein Value
Normal.nonAA
AD.nonAA
Normal.AA
AD.AA
Diagnosis.Race
MEdarkmagenta.CSF (Synthetic)
ANOVA p: 0.17
Eigenprotein Value
Normal.Male
Normal.Female
AD.Male
AD.Female
Diagnosis.Sex
bicor=0.34, p=0.049
cor=0.35, p=0.039
Eigenprotein
ABeta/Tau Ratio
bicor=0.049, p=0.79
cor=-0.075, p=0.69
Eigenprotein
MoCA Score
bicor=-0.42, p=0.086
cor=-0.38, p=0.12
Eigenprotein
Age (Normal Only)
bicor=-0.52, p=0.032
cor=-0.52, p=0.032
Eigenprotein
Age (AD Only)
M8 pink.MEGATMT488
Protein Transport
Eigenprotein Value
Control
AsymAD
AD
MEpink.CSF 35 Samp. (Synthetic)
ANOVA p: 0.49
Eigenprotein Value
Normal
AD
MEpink.CSF (Synthetic)
ANOVA p: 0.089
Eigenprotein Value
Normal.nonAA
AD.nonAA
Normal.AA
AD.AA
Diagnosis.Race
MEpink.CSF (Synthetic)
ANOVA p: 0.31
Eigenprotein Value
Normal.Male
Normal.Female
AD.Male
AD.Female
Diagnosis.Sex
bicor=0.24, p=0.17
cor=0.36, p=0.034
Eigenprotein
ABeta/Tau Ratio
bicor=-0.11, p=0.57
cor=-0.11, p=0.56
Eigenprotein
MoCA Score
bicor=-0.22, p=0.37
cor=-0.076, p=0.76
Eigenprotein
Age (Normal Only)
bicor=-0.33, p=0.2
cor=-0.3, p=0.24
Eigenprotein
Age (AD Only)

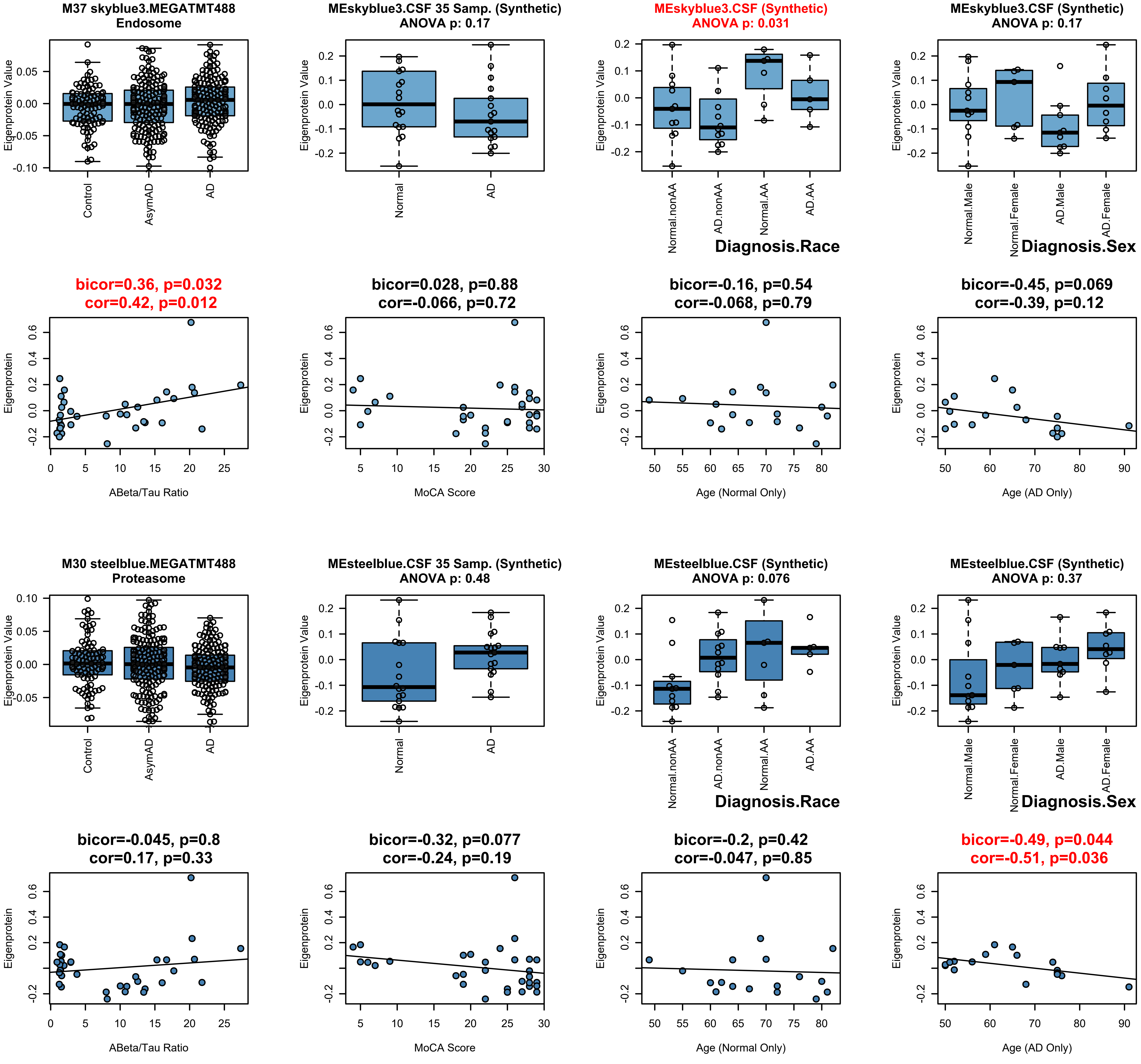
M37 skyblue3.MEGATMT488
Endosome
Eigenprotein Value
Control
AsymAD
AD
MEskyblue3.CSF 35 Samp. (Synthetic)
ANOVA p: 0.17
Normal
AD
MEskyblue3.CSF (Synthetic)
ANOVA p: 0.031
Normal.nonAA
AD.nonAA
Normal.AA
AD.AA
Diagnosis.Race
MEskyblue3.CSF (Synthetic)
ANOVA p: 0.17
Normal.Male
Normal.Female
AD.Male
AD.Female
Diagnosis.Sex
bicor=0.36, p=0.032
cor=0.42, p=0.012
Eigenprotein
ABeta/Tau Ratio
bicor=0.028, p=0.88
cor=-0.066, p=0.72
MoCA Score
bicor=-0.16, p=0.54
cor=-0.068, p=0.79
Age (Normal Only)
bicor=-0.45, p=0.069
cor=-0.39, p=0.12
Age (AD Only)
M30 steelblue.MEGATMT488
Proteasome
MEsteelblue.CSF 35 Samp. (Synthetic)
ANOVA p: 0.48
MEsteelblue.CSF (Synthetic)
ANOVA p: 0.076
MEsteelblue.CSF (Synthetic)
ANOVA p: 0.37
bicor=-0.045, p=0.8
cor=0.17, p=0.33
bicor=-0.32, p=0.077
cor=-0.24, p=0.19
bicor=-0.2, p=0.42
cor=-0.047, p=0.85
bicor=-0.49, p=0.044
cor=-0.51, p=0.036

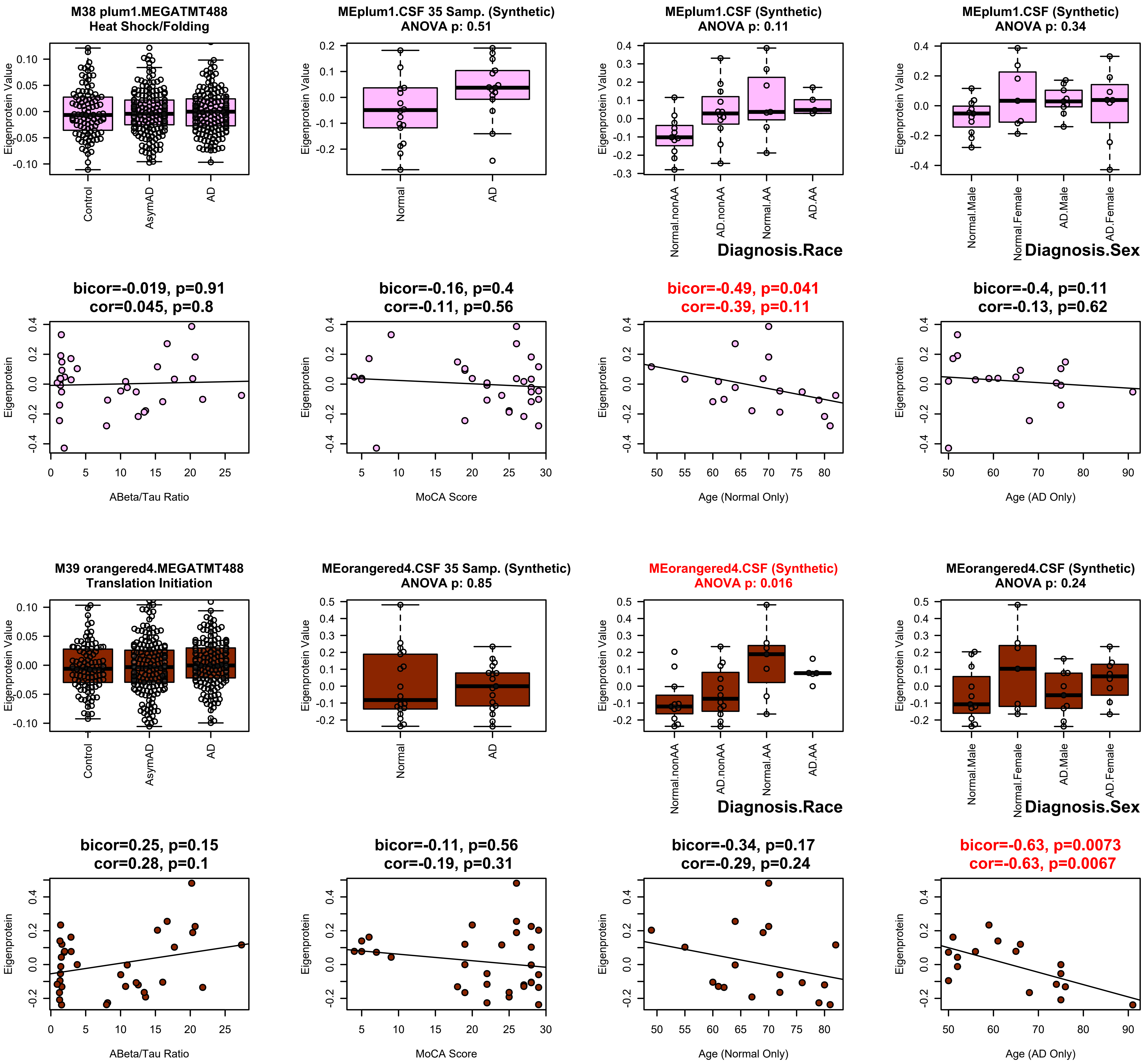
M38 plum1.MEGATMT488
Heat Shock/Folding
Eigenprotein Value
Control
AsymAD
AD
MEplum1.CSF 35 Samp. (Synthetic)
ANOVA p: 0.51
Normal
AD
MEplum1.CSF (Synthetic)
ANOVA p: 0.11
Normal.nonAA
AD.nonAA
Normal.AA
AD.AA
Diagnosis.Race
MEplum1.CSF (Synthetic)
ANOVA p: 0.34
Normal.Male
Normal.Female
AD.Male
AD.Female
Diagnosis.Sex
bicor=-0.019, p=0.91
cor=0.045, p=0.8
Eigenprotein
ABeta/Tau Ratio
bicor=-0.16, p=0.4
cor=-0.11, p=0.56
MoCA Score
bicor=-0.49, p=0.041
cor=-0.39, p=0.11
Age (Normal Only)
bicor=-0.4, p=0.11
cor=-0.13, p=0.62
Age (AD Only)
M39 orangered4.MEGATMT488
Translation Initiation
MEorangered4.CSF 35 Samp. (Synthetic)
ANOVA p: 0.85
MEorangered4.CSF (Synthetic)
ANOVA p: 0.016
MEorangered4.CSF (Synthetic)
ANOVA p: 0.24
bicor=0.25, p=0.15
cor=0.28, p=0.1
bicor=-0.11, p=0.56
cor=-0.19, p=0.31
bicor=-0.34, p=0.17
cor=-0.29, p=0.24
bicor=-0.63, p=0.0073
cor=-0.63, p=0.0067

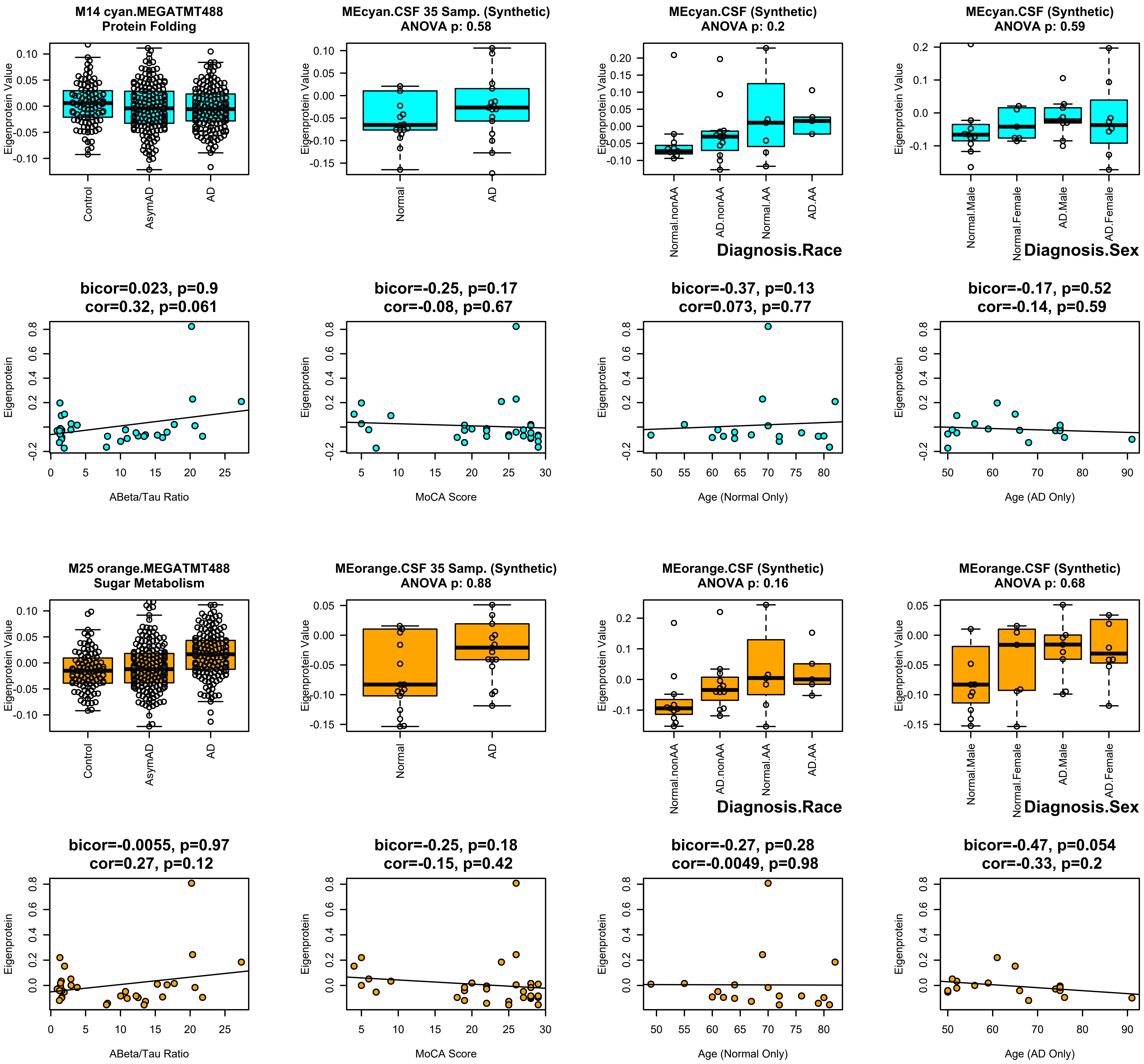
M14 cyan.MEGATMT488
Protein Folding
MEcyan.CSF 35 Samp. (Synthetic)
ANOVA p: 0.58
MEcyan.CSF (Synthetic)
ANOVA p: 0.2
MEcyan.CSF (Synthetic)
ANOVA p: 0.59
Eigenprotein Value
Control
AsymAD
AD
Normal
AD
Normal.nonAA
AD.nonAA
Normal.AA
AD.AA
Diagnosis.Race
Normal.Male
Normal.Female
AD.Male
AD.Female
Diagnosis.Sex
bicor=0.023, p=0.9
cor=0.32, p=0.061
bicor=-0.25, p=0.17
cor=-0.08, p=0.67
bicor=-0.37, p=0.13
cor=0.073, p=0.77
bicor=-0.17, p=0.52
cor=-0.14, p=0.59
Eigenprotein
ABeta/Tau Ratio
MoCA Score
Age (Normal Only)
Age (AD Only)
M25 orange.MEGATMT488
Sugar Metabolism
MEorange.CSF 35 Samp. (Synthetic)
ANOVA p: 0.88
MEorange.CSF (Synthetic)
ANOVA p: 0.16
MEorange.CSF (Synthetic)
ANOVA p: 0.68
Eigenprotein Value
Control
AsymAD
AD
Normal
AD
Normal.nonAA
AD.nonAA
Normal.AA
AD.AA
Diagnosis.Race
Normal.Male
Normal.Female
AD.Male
AD.Female
Diagnosis.Sex
bicor=-0.0055, p=0.97
cor=0.27, p=0.12
bicor=-0.25, p=0.18
cor=-0.15, p=0.42
bicor=-0.27, p=0.28
cor=-0.0049, p=0.98
bicor=-0.47, p=0.054
cor=-0.33, p=0.2
Eigenprotein
ABeta/Tau Ratio
MoCA Score
Age (Normal Only)
Age (AD Only)

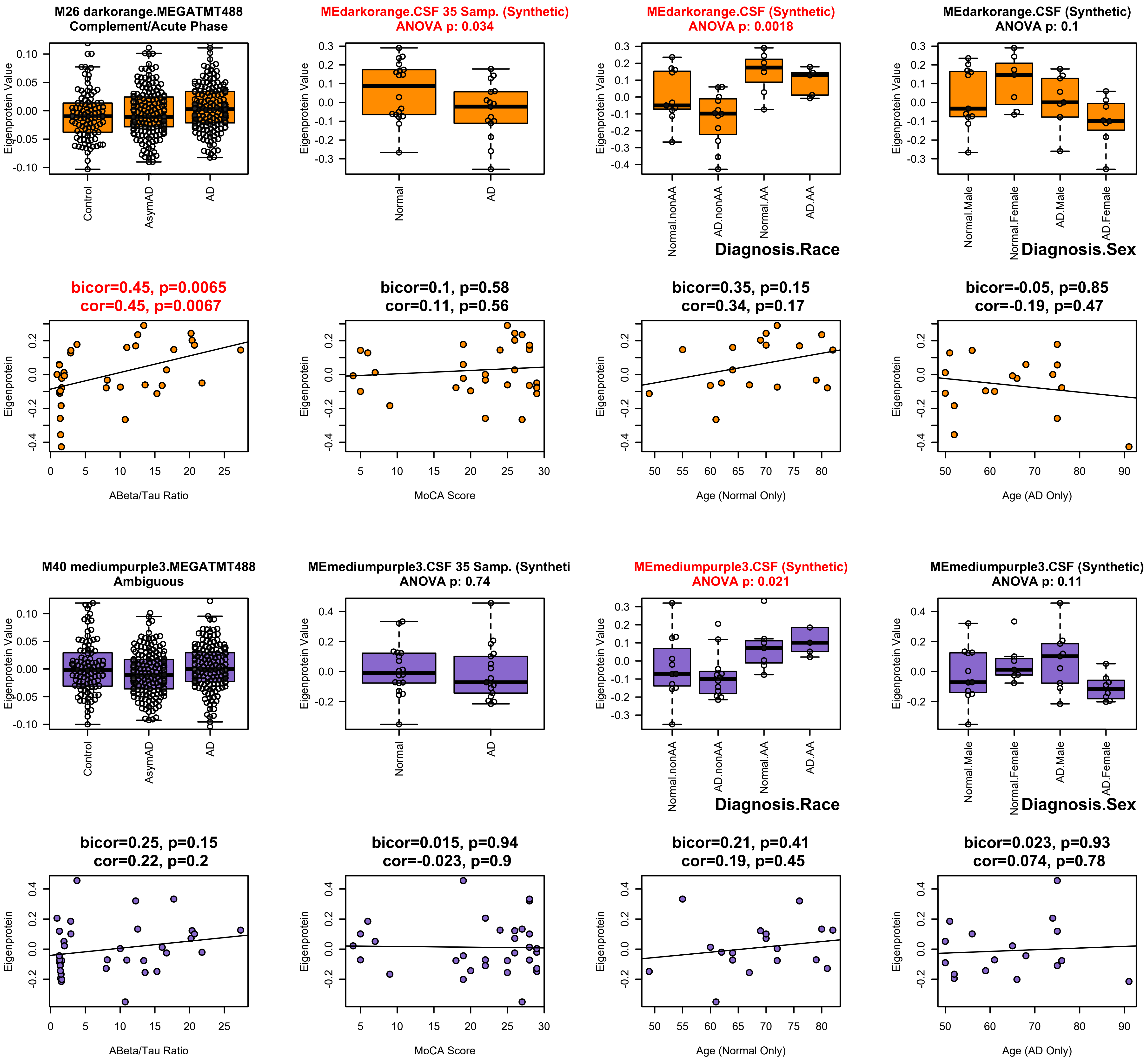
M26 darkorange.MEGATMT488
Complement/Acute Phase
Eigenprotein Value
Control
AsymAD
AD
MEdarkorange.CSF 35 Samp. (Synthetic)
ANOVA p: 0.034
Normal
AD
MEdarkorange.CSF (Synthetic)
ANOVA p: 0.0018
Normal.nonAA
AD.nonAA
Normal.AA
AD.AA
Diagnosis.Race
MEdarkorange.CSF (Synthetic)
ANOVA p: 0.1
Normal.Male
Normal.Female
AD.Male
AD.Female
Diagnosis.Sex
bicor=0.45, p=0.0065
cor=0.45, p=0.0067
Eigenprotein
ABeta/Tau Ratio
bicor=0.1, p=0.58
cor=0.11, p=0.56
MoCA Score
bicor=0.35, p=0.15
cor=0.34, p=0.17
Age (Normal Only)
bicor=-0.05, p=0.85
cor=-0.19, p=0.47
Age (AD Only)
M40 mediumpurple3.MEGATMT488
Ambiguous
MEmediumpurple3.CSF 35 Samp. (Syntheti
ANOVA p: 0.74
MEmediumpurple3.CSF (Synthetic)
ANOVA p: 0.021
MEmediumpurple3.CSF (Synthetic)
ANOVA p: 0.11
bicor=0.25, p=0.15
cor=0.22, p=0.2
bicor=0.015, p=0.94
cor=-0.023, p=0.9
bicor=0.21, p=0.41
cor=0.19, p=0.45
bicor=0.023, p=0.93
cor=0.074, p=0.78

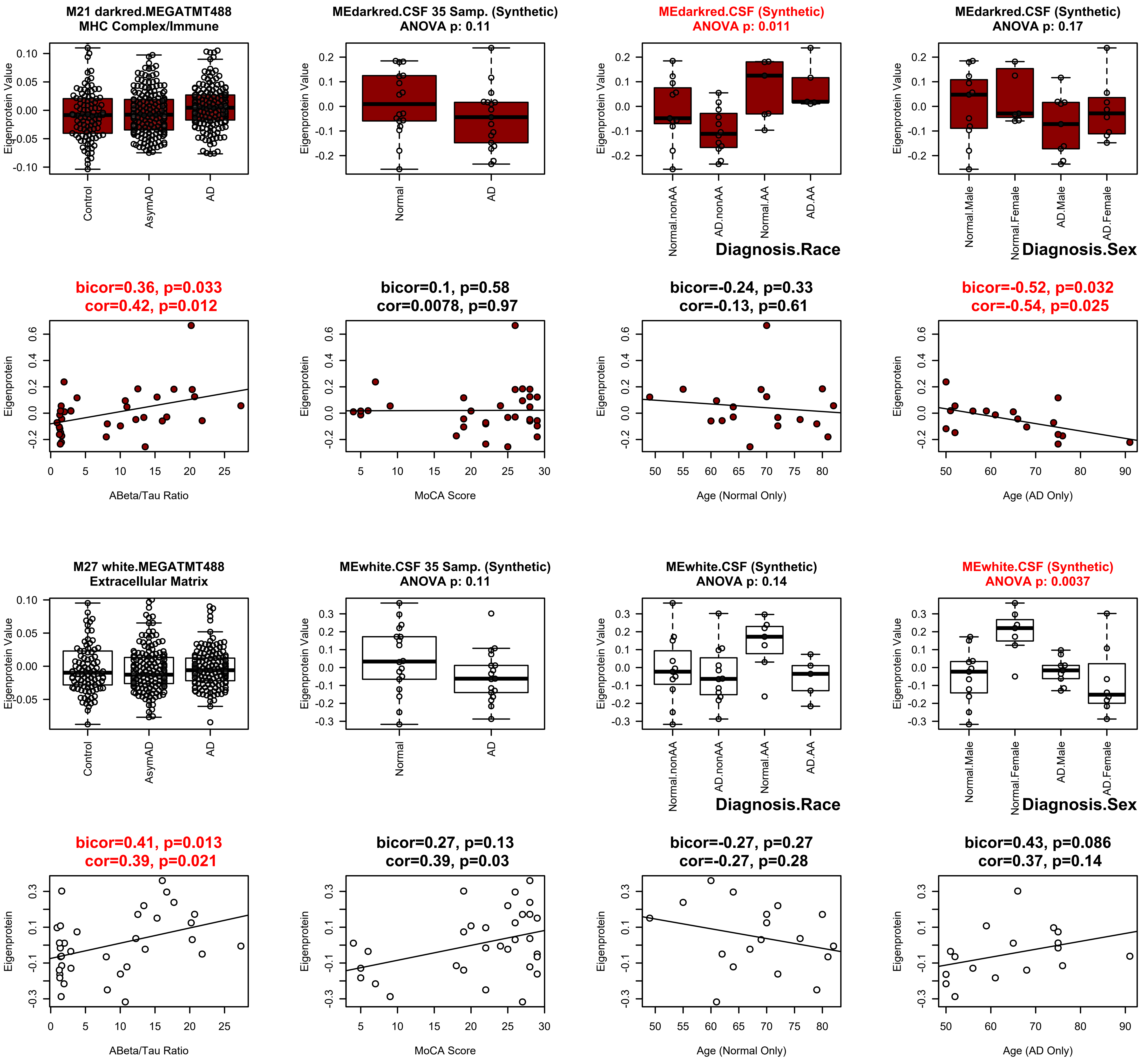

M21 darkred.MEGATMT488
MHC Complex/Immune
Eigenprotein Value
Control
AsymAD
AD
MEdarkred.CSF 35 Samp. (Synthetic)
ANOVA p: 0.11
Normal
AD
MEdarkred.CSF (Synthetic)
ANOVA p: 0.011
Normal.nonAA
AD.nonAA
Normal.AA
AD.AA
Diagnosis.Race
MEdarkred.CSF (Synthetic)
ANOVA p: 0.17
Normal.Male
Normal.Female
AD.Male
AD.Female
Diagnosis.Sex
bicor=0.36, p=0.033
cor=0.42, p=0.012
Eigenprotein
ABeta/Tau Ratio
bicor=0.1, p=0.58
cor=0.0078, p=0.97
MoCA Score
bicor=-0.24, p=0.33
cor=-0.13, p=0.61
Age (Normal Only)
bicor=-0.52, p=0.032
cor=-0.54, p=0.025
Age (AD Only)
M27 white.MEGATMT488
Extracellular Matrix
MEwhite.CSF 35 Samp. (Synthetic)
ANOVA p: 0.11
MEwhite.CSF (Synthetic)
ANOVA p: 0.14
MEwhite.CSF (Synthetic)
ANOVA p: 0.0037
bicor=0.41, p=0.013
cor=0.39, p=0.021
bicor=0.27, p=0.13
cor=0.39, p=0.03
bicor=-0.27, p=0.27
cor=-0.27, p=0.28
bicor=0.43, p=0.086
cor=0.37, p=0.14

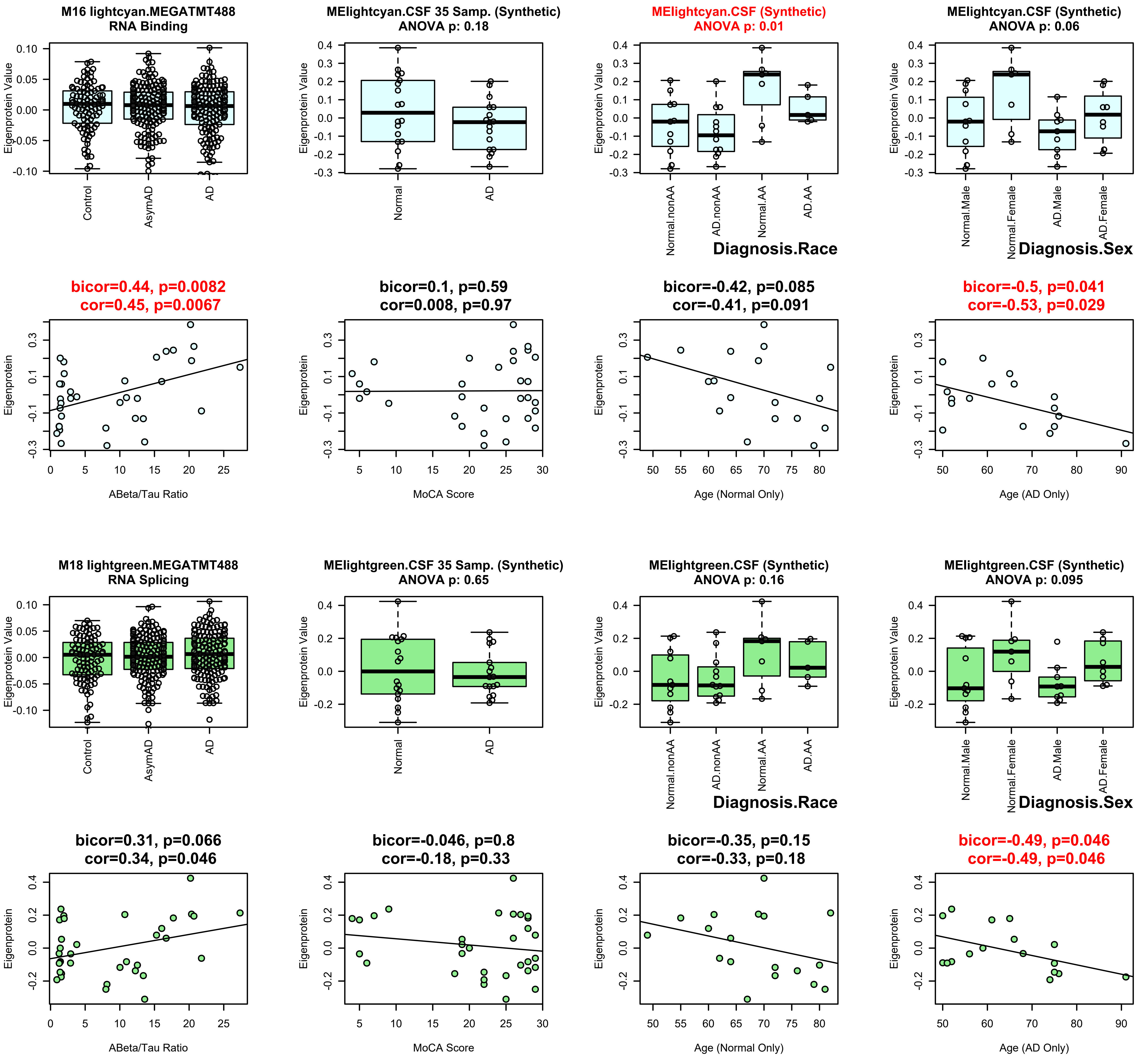
M16 lightcyan.MEGATMT488
RNA Binding
Eigenprotein Value
Control
AsymAD
AD
MElightcyan.CSF 35 Samp. (Synthetic)
ANOVA p: 0.18
Normal
AD
MElightcyan.CSF (Synthetic)
ANOVA p: 0.01
Normal.nonAA
AD.nonAA
Normal.AA
AD.AA
Diagnosis.Race
MElightcyan.CSF (Synthetic)
ANOVA p: 0.06
Normal.Male
Normal.Female
AD.Male
AD.Female
Diagnosis.Sex
bicor=0.44, p=0.0082
cor=0.45, p=0.0067
Eigenprotein
ABeta/Tau Ratio
bicor=0.1, p=0.59
cor=0.008, p=0.97
MoCA Score
bicor=-0.42, p=0.085
cor=-0.41, p=0.091
Age (Normal Only)
bicor=-0.5, p=0.041
cor=-0.53, p=0.029
Age (AD Only)
M18 lightgreen.MEGATMT488
RNA Splicing
MElightgreen.CSF 35 Samp. (Synthetic)
ANOVA p: 0.65
MElightgreen.CSF (Synthetic)
ANOVA p: 0.16
MElightgreen.CSF (Synthetic)
ANOVA p: 0.095
bicor=0.31, p=0.066
cor=0.34, p=0.046
bicor=-0.046, p=0.8
cor=-0.18, p=0.33
bicor=-0.35, p=0.15
cor=-0.33, p=0.18
bicor=-0.49, p=0.046
cor=-0.49, p=0.046

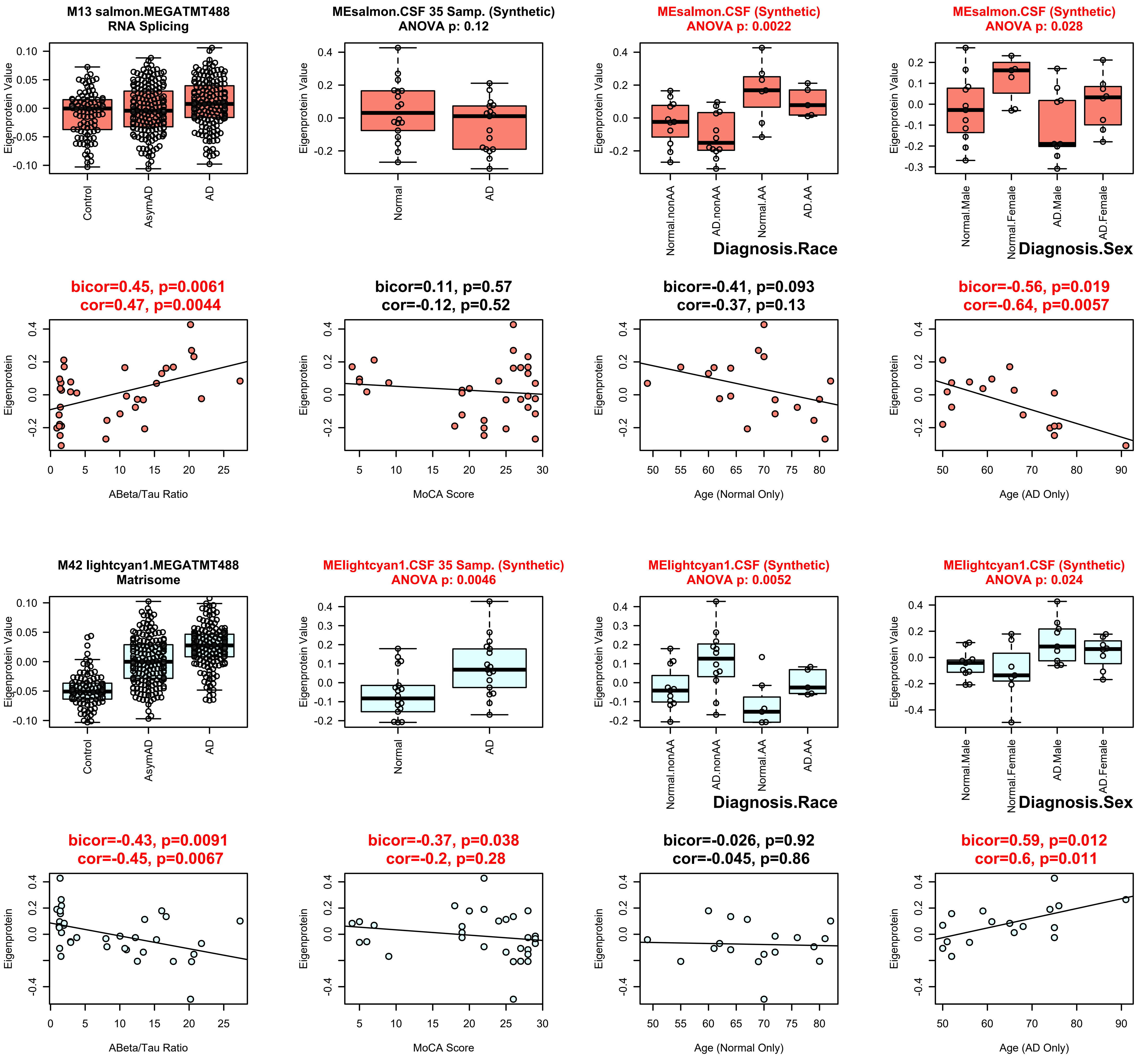

M13 salmon.MEGATMT488
RNA Splicing
MEsalmon.CSF 35 Samp. (Synthetic)
ANOVA p: 0.12
MEsalmon.CSF (Synthetic)
ANOVA p: 0.0022
MEsalmon.CSF (Synthetic)
ANOVA p: 0.028
Eigenprotein Value
Control
AsymAD
AD
Normal
Normal.nonAA
AD.nonAA
Normal.AA
AD.AA
Diagnosis.Race
Normal.Male
Normal.Female
AD.Male
AD.Female
Diagnosis.Sex
bicor=0.45, p=0.0061
cor=0.47, p=0.0044
bicor=0.11, p=0.57
cor=-0.12, p=0.52
bicor=-0.41, p=0.093
cor=-0.37, p=0.13
bicor=-0.56, p=0.019
cor=-0.64, p=0.0057
Eigenprotein
ABeta/Tau Ratio
MoCA Score
Age (Normal Only)
Age (AD Only)
M42 lightcyan1.MEGATMT488
Matrisome
MElightcyan1.CSF 35 Samp. (Synthetic)
ANOVA p: 0.0046
MElightcyan1.CSF (Synthetic)
ANOVA p: 0.0052
MElightcyan1.CSF (Synthetic)
ANOVA p: 0.024
bicor=-0.43, p=0.0091
cor=-0.45, p=0.0067
bicor=-0.37, p=0.038
cor=-0.2, p=0.28
bicor=-0.026, p=0.92
cor=-0.045, p=0.86
bicor=0.59, p=0.012
cor=0.6, p=0.011

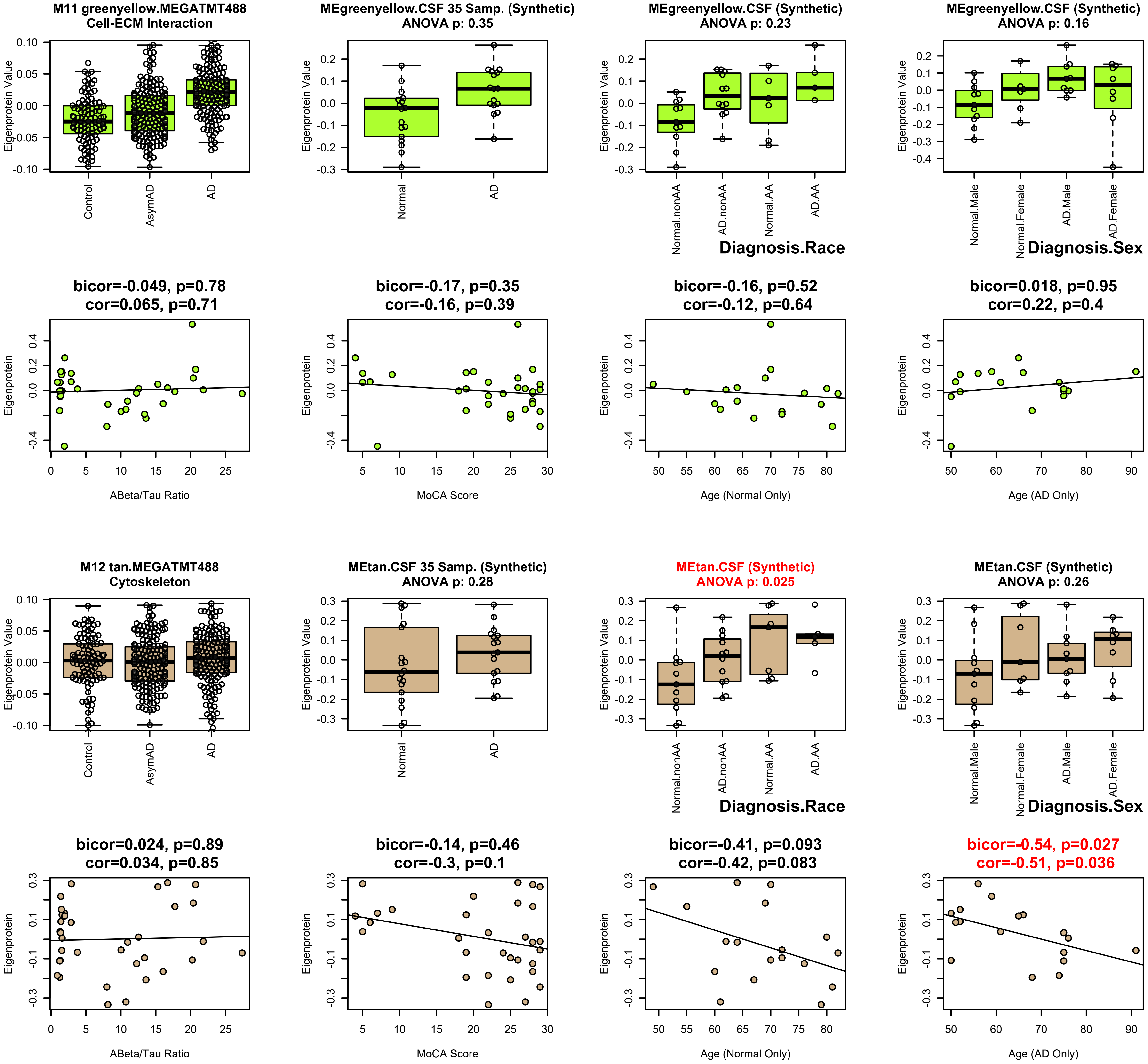
M11 greenyellow.MEGATMT488
Cell-ECM Interaction
Eigenprotein Value
Control
AsymAD
AD
MEgreenyellow.CSF 35 Samp. (Synthetic)
ANOVA p: 0.35
Normal
AD
MEgreenyellow.CSF (Synthetic)
ANOVA p: 0.23
Normal.nonAA
AD.nonAA
Normal.AA
AD.AA
Diagnosis.Race
MEgreenyellow.CSF (Synthetic)
ANOVA p: 0.16
Normal.Male
Normal.Female
AD.Male
AD.Female
Diagnosis.Sex
bicor=-0.049, p=0.78
cor=0.065, p=0.71
ABeta/Tau Ratio
bicor=-0.17, p=0.35
cor=-0.16, p=0.39
MoCA Score
bicor=-0.16, p=0.52
cor=-0.12, p=0.64
Age (Normal Only)
bicor=0.018, p=0.95
cor=0.22, p=0.4
Age (AD Only)
Eigenprotein
M12 tan.MEGATMT488
Cytoskeleton
METan.CSF 35 Samp. (Synthetic)
ANOVA p: 0.28
METan.CSF (Synthetic)
ANOVA p: 0.025
METan.CSF (Synthetic)
ANOVA p: 0.26
bicor=0.024, p=0.89
cor=0.034, p=0.85
bicor=-0.14, p=0.46
cor=-0.3, p=0.1
bicor=-0.41, p=0.093
cor=-0.42, p=0.083
bicor=-0.54, p=0.027
cor=-0.51, p=0.036

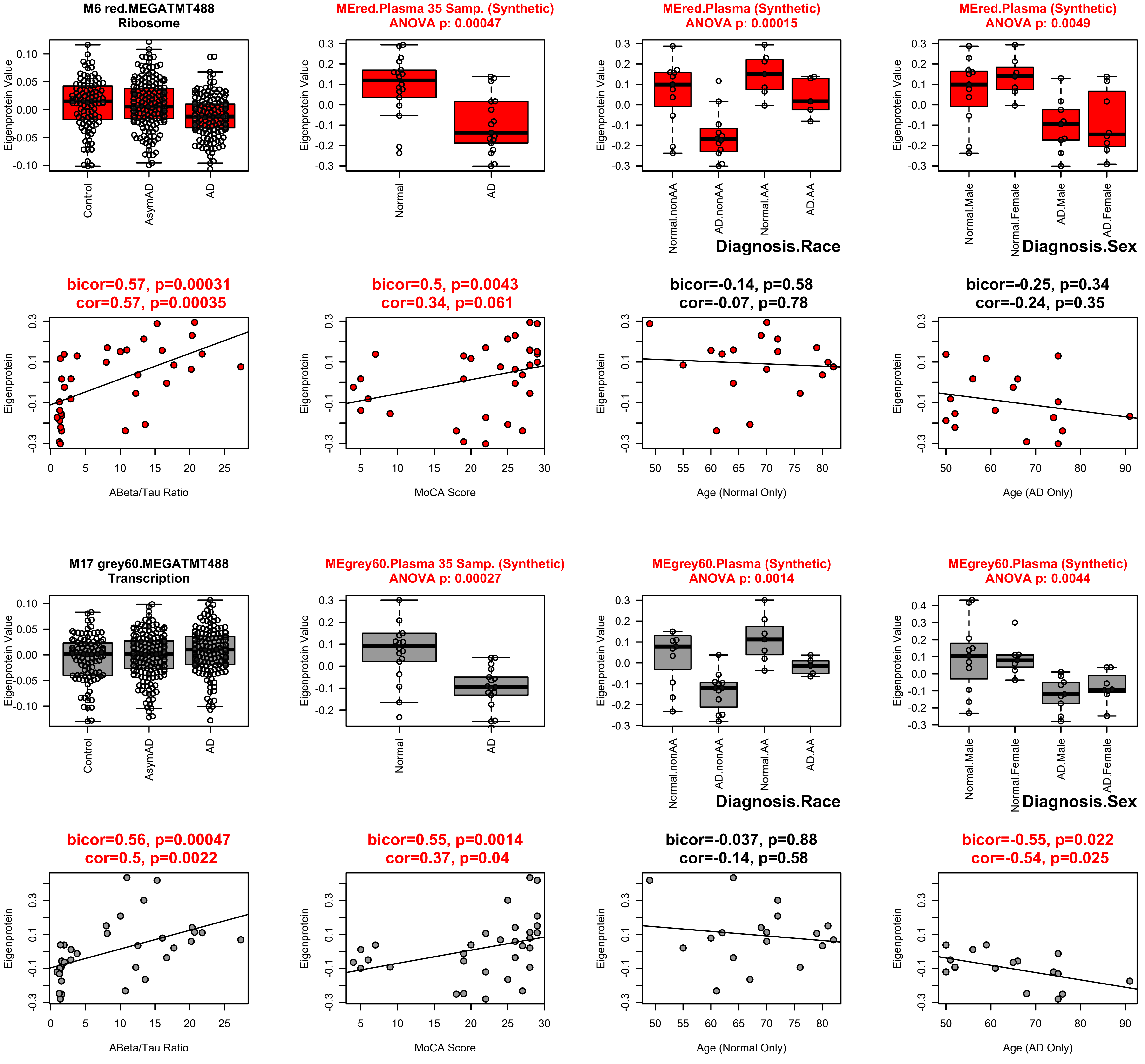
M6 red.MEGATMT488
Ribosome
MEred.Plasma 35 Samp. (Synthetic)
ANOVA p: 0.00047
MEred.Plasma (Synthetic)
ANOVA p: 0.00015
MEred.Plasma (Synthetic)
ANOVA p: 0.0049
Eigenprotein Value
Control
AsymAD
AD
Normal
Normal.nonAA
AD.nonAA
Normal.AA
AD.AA
Diagnosis.Race
Normal.Male
Normal.Female
AD.Male
AD.Female
Diagnosis.Sex
bicor=0.57, p=0.00031
cor=0.57, p=0.00035
bicor=0.5, p=0.0043
cor=0.34, p=0.061
bicor=-0.14, p=0.58
cor=-0.07, p=0.78
bicor=-0.25, p=0.34
cor=-0.24, p=0.35
Eigenprotein
ABeta/Tau Ratio
MoCA Score
Age (Normal Only)
Age (AD Only)
M17 grey60.MEGATMT488
Transcription
MEgrey60.Plasma 35 Samp. (Synthetic)
ANOVA p: 0.00027
MEgrey60.Plasma (Synthetic)
ANOVA p: 0.0014
MEgrey60.Plasma (Synthetic)
ANOVA p: 0.0044
Eigenprotein Value
Control
AsymAD
AD
Normal
Normal.nonAA
AD.nonAA
Normal.AA
AD.AA
Diagnosis.Race
Normal.Male
Normal.Female
AD.Male
AD.Female
Diagnosis.Sex
bicor=0.56, p=0.00047
cor=0.5, p=0.0022
bicor=0.55, p=0.0014
cor=0.37, p=0.04
bicor=-0.037, p=0.88
cor=-0.14, p=0.58
bicor=-0.55, p=0.022
cor=-0.54, p=0.025
Eigenprotein
ABeta/Tau Ratio
MoCA Score
Age (Normal Only)
Age (AD Only)

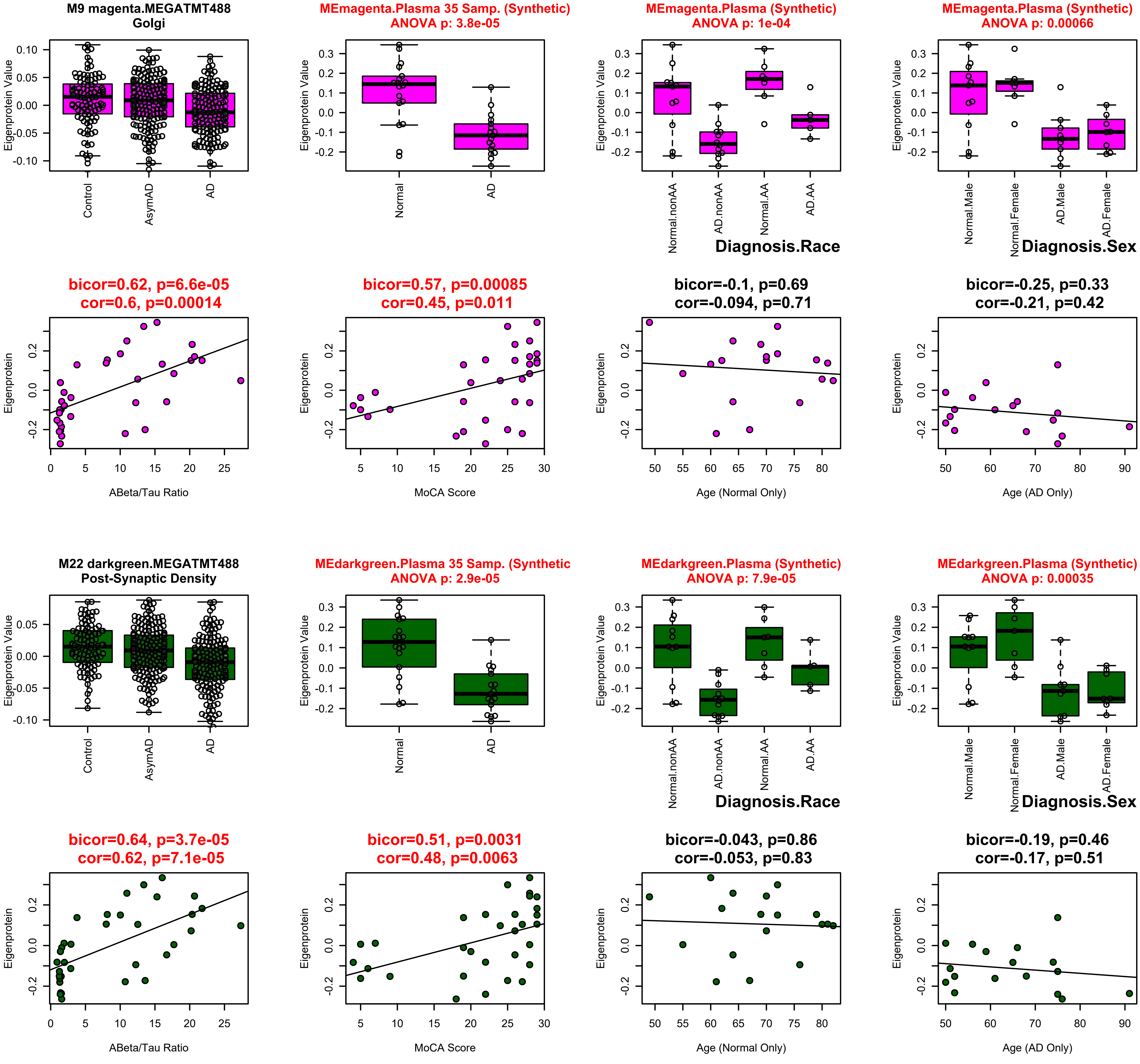

M9 magenta.MEGATMT488
Golgi
Eigenprotein Value
Control
AsymAD
AD
MEmagenta.Plasma 35 Samp. (Synthetic)
ANOVA p: 3.8e-05
Normal
AD
MEmagenta.Plasma (Synthetic)
ANOVA p: 1e-04
Normal.nonAA
AD.nonAA
Normal.AA
AD.AA
Diagnosis.Race
MEmagenta.Plasma (Synthetic)
ANOVA p: 0.00066
Normal.Male
Normal.Female
AD.Male
AD.Female
Diagnosis.Sex
bicor=0.62, p=6.6e-05
cor=0.6, p=0.00014
Eigenprotein
ABeta/Tau Ratio
bicor=0.57, p=0.00085
cor=0.45, p=0.011
MoCA Score
bicor=-0.1, p=0.69
cor=-0.094, p=0.71
Age (Normal Only)
bicor=-0.25, p=0.33
cor=-0.21, p=0.42
Age (AD Only)
M22 darkgreen.MEGATMT488
Post-Synaptic Density
MEdarkgreen.Plasma 35 Samp. (Synthetic
ANOVA p: 2.9e-05
MEdarkgreen.Plasma (Synthetic)
ANOVA p: 7.9e-05
MEdarkgreen.Plasma (Synthetic)
ANOVA p: 0.00035
bicor=0.64, p=3.7e-05
cor=0.62, p=7.1e-05
bicor=0.51, p=0.0031
cor=0.48, p=0.0063
bicor=-0.043, p=0.86
cor=-0.053, p=0.83
bicor=-0.19, p=0.46
cor=-0.17, p=0.51

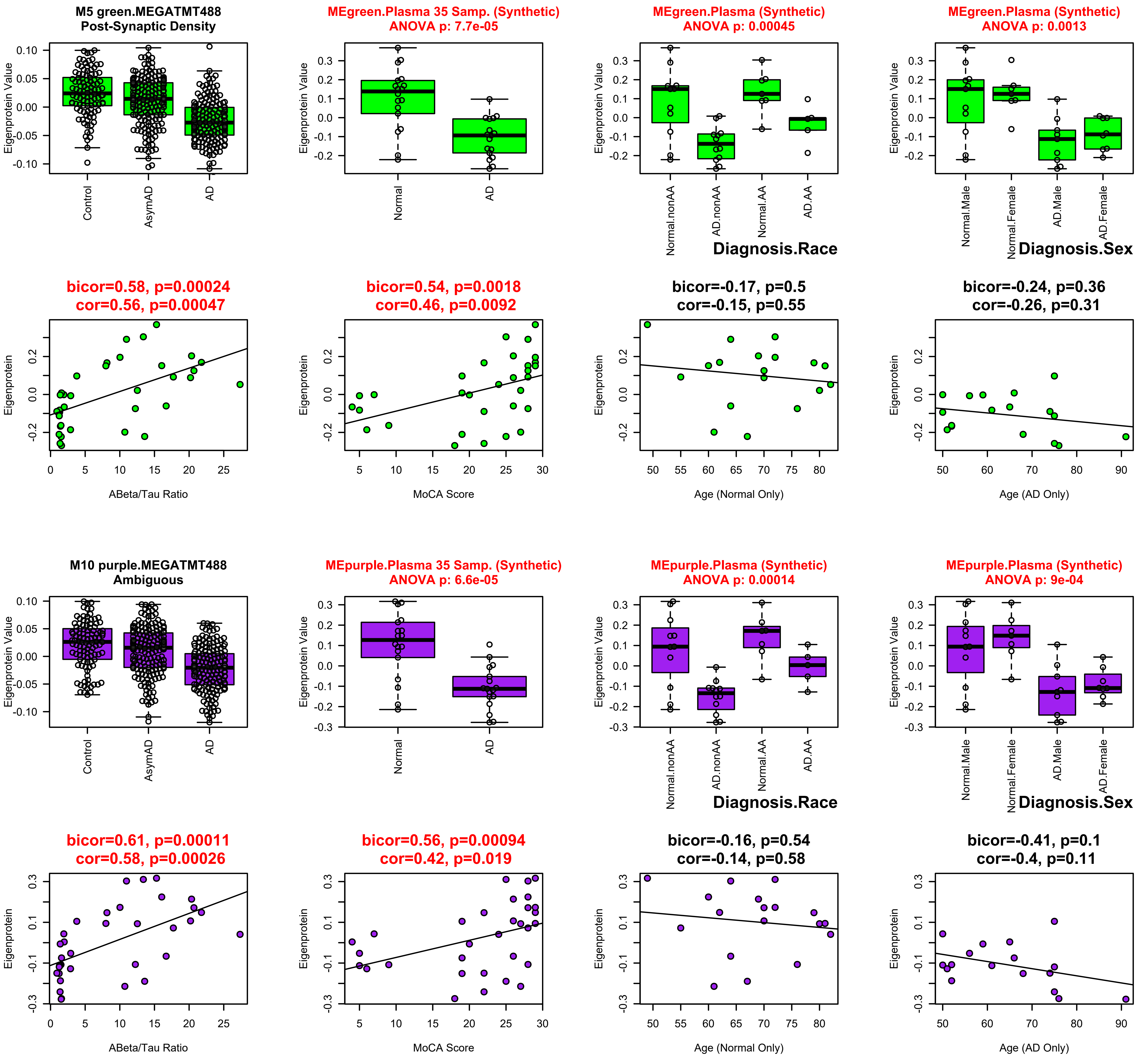
M5 green.MEGATMT488
Post-Synaptic Density
Eigenprotein Value
Control
AsymAD
AD
MEgreen.Plasma 35 Samp. (Synthetic)
ANOVA p: 7.7e-05
Normal
AD
MEgreen.Plasma (Synthetic)
ANOVA p: 0.00045
Normal.nonAA
AD.nonAA
Normal.AA
AD.AA
Diagnosis.Race
MEgreen.Plasma (Synthetic)
ANOVA p: 0.0013
Normal.Male
Normal.Female
AD.Male
AD.Female
Diagnosis.Sex
bicor=0.58, p=0.00024
cor=0.56, p=0.00047
Eigenprotein
ABeta/Tau Ratio
bicor=0.54, p=0.0018
cor=0.46, p=0.0092
MoCA Score
bicor=-0.17, p=0.5
cor=-0.15, p=0.55
Age (Normal Only)
bicor=-0.24, p=0.36
cor=-0.26, p=0.31
Age (AD Only)
M10 purple.MEGATMT488
Ambiguous
Eigenprotein Value
Control
AsymAD
AD
MEpurple.Plasma 35 Samp. (Synthetic)
ANOVA p: 6.6e-05
Normal
AD
MEpurple.Plasma (Synthetic)
ANOVA p: 0.00014
Normal.nonAA
AD.nonAA
Normal.AA
AD.AA
Diagnosis.Race
MEpurple.Plasma (Synthetic)
ANOVA p: 9e-04
Normal.Male
Normal.Female
AD.Male
AD.Female
Diagnosis.Sex
bicor=0.61, p=0.00011
cor=0.58, p=0.00026
Eigenprotein
ABeta/Tau Ratio
bicor=0.56, p=0.00094
cor=0.42, p=0.019
MoCA Score
bicor=-0.16, p=0.54
cor=-0.14, p=0.58
Age (Normal Only)
bicor=-0.41, p=0.1
cor=-0.4, p=0.11
Age (AD Only)

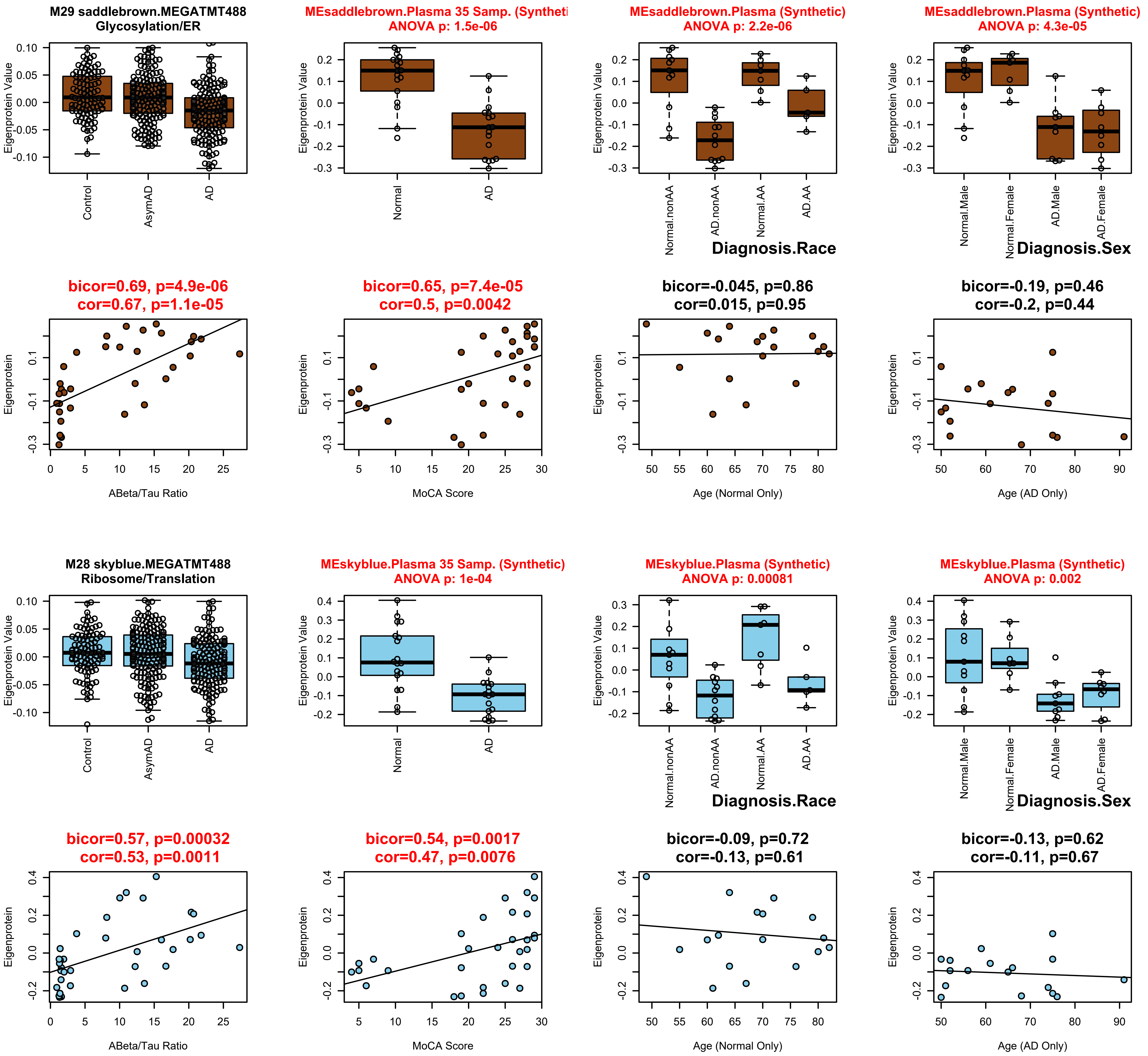
M29 saddlebrown.MEGATMT488
Glycosylation/ER
Eigenprotein Value
Control
AsymAD
AD
MEsaddlebrown.Plasma 35 Samp. (Syntheti
ANOVA p: 1.5e-06
Normal
AD
MEsaddlebrown.Plasma (Synthetic)
ANOVA p: 2.2e-06
Normal.nonAA
AD.nonAA
Normal.AA
AD.AA
Diagnosis.Race
MEsaddlebrown.Plasma (Synthetic)
ANOVA p: 4.3e-05
Normal.Male
Normal.Female
AD.Male
AD.Female
Diagnosis.Sex
bicor=0.69, p=4.9e-06
cor=0.67, p=1.1e-05
Eigenprotein
ABeta/Tau Ratio
bicor=0.65, p=7.4e-05
cor=0.5, p=0.0042
MoCA Score
bicor=-0.045, p=0.86
cor=0.015, p=0.95
Age (Normal Only)
bicor=-0.19, p=0.46
cor=-0.2, p=0.44
Age (AD Only)
M28 skyblue.MEGATMT488
Ribosome/Translation
Eigenprotein Value
Control
AsymAD
AD
MEskyblue.Plasma 35 Samp. (Synthetic)
ANOVA p: 1e-04
Normal
AD
MEskyblue.Plasma (Synthetic)
ANOVA p: 0.00081
Normal.nonAA
AD.nonAA
Normal.AA
AD.AA
Diagnosis.Race
MEskyblue.Plasma (Synthetic)
ANOVA p: 0.002
Normal.Male
Normal.Female
AD.Male
AD.Female
Diagnosis.Sex
bicor=0.57, p=0.00032
cor=0.53, p=0.0011
Eigenprotein
ABeta/Tau Ratio
bicor=0.54, p=0.0017
cor=0.47, p=0.0076
MoCA Score
bicor=-0.09, p=0.72
cor=-0.13, p=0.61
Age (Normal Only)
bicor=-0.13, p=0.62
cor=-0.11, p=0.67
Age (AD Only)

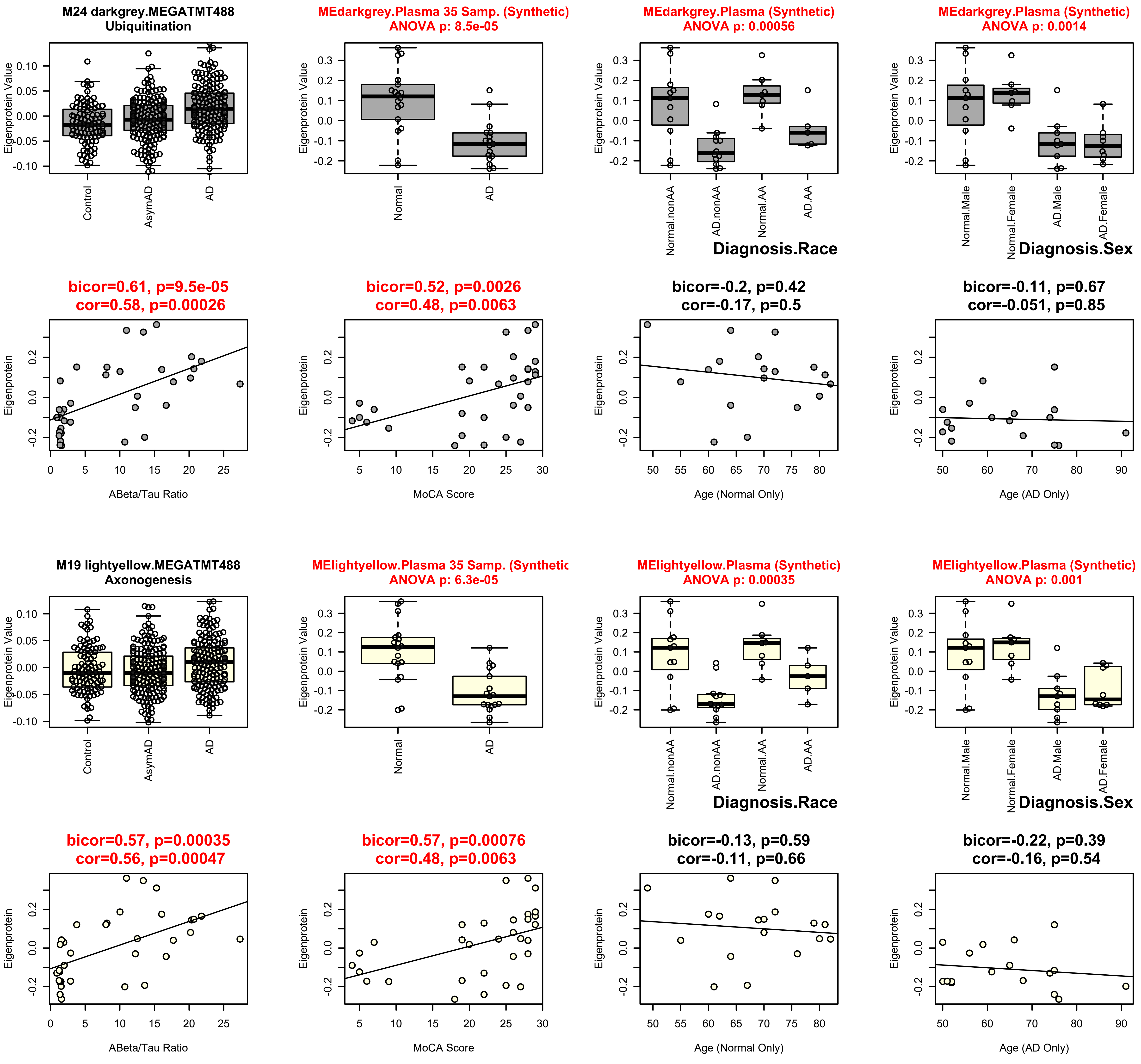
M24 darkgrey.MEGATMT488
Ubiquitination
MEdarkgrey.Plasma 35 Samp. (Synthetic)
ANOVA p: 8.5e-05
MEdarkgrey.Plasma (Synthetic)
ANOVA p: 0.00056
MEdarkgrey.Plasma (Synthetic)
ANOVA p: 0.0014
Eigenprotein Value
Control
AsymAD
AD
Normal
Normal.nonAA
AD.nonAA
Normal.AA
AD.AA
Diagnosis.Race
Normal.Male
Normal.Female
AD.Male
AD.Female
Diagnosis.Sex
bicor=0.61, p=9.5e-05
cor=0.58, p=0.00026
bicor=0.52, p=0.0026
cor=0.48, p=0.0063
bicor=-0.2, p=0.42
cor=-0.17, p=0.5
bicor=-0.11, p=0.67
cor=-0.051, p=0.85
Eigenprotein
ABeta/Tau Ratio
MoCA Score
Age (Normal Only)
Age (AD Only)
M19 lightyellow.MEGATMT488
Axonogenesis
MElightyellow.Plasma 35 Samp. (Synthetic
ANOVA p: 6.3e-05
MElightyellow.Plasma (Synthetic)
ANOVA p: 0.00035
MElightyellow.Plasma (Synthetic)
ANOVA p: 0.001
bicor=0.57, p=0.00035
cor=0.56, p=0.00047
bicor=0.57, p=0.00076
cor=0.48, p=0.0063
bicor=-0.13, p=0.59
cor=-0.11, p=0.66
bicor=-0.22, p=0.39
cor=-0.16, p=0.54

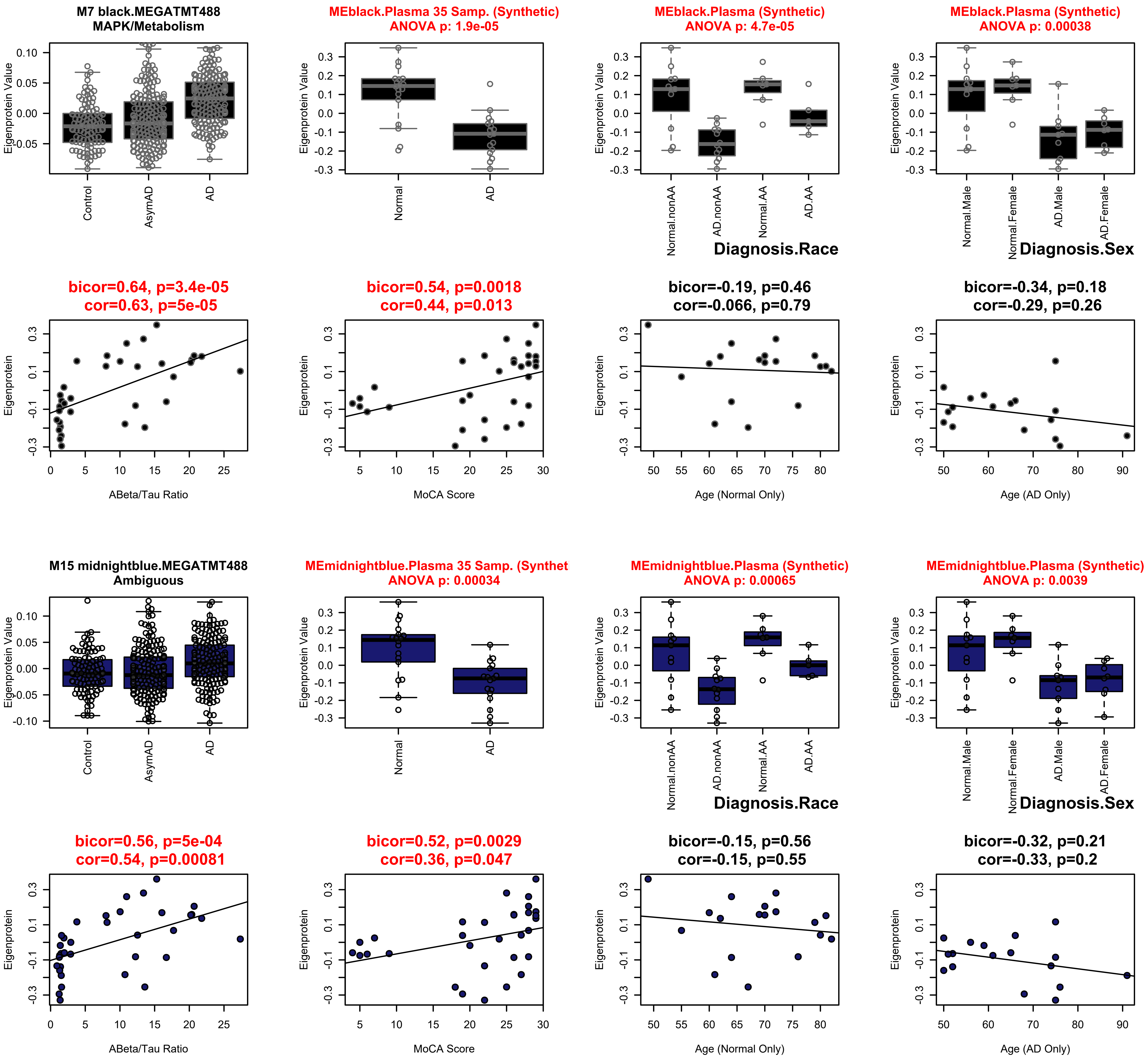

M7 black.MEGATMT488
MAPK/Metabolism
Eigenprotein Value
Control
AsymAD
AD
MEblack.Plasma 35 Samp. (Synthetic)
ANOVA p: 1.9e-05
Normal
AD
MEblack.Plasma (Synthetic)
ANOVA p: 4.7e-05
Normal.nonAA
AD.nonAA
Normal.AA
AD.AA
Diagnosis.Race
MEblack.Plasma (Synthetic)
ANOVA p: 0.00038
Normal.Male
Normal.Female
AD.Male
AD.Female
Diagnosis.Sex
bicor=0.64, p=3.4e-05
cor=0.63, p=5e-05
Eigenprotein
ABeta/Tau Ratio
bicor=0.54, p=0.0018
cor=0.44, p=0.013
MoCA Score
bicor=-0.19, p=0.46
cor=-0.066, p=0.79
Age (Normal Only)
bicor=-0.34, p=0.18
cor=-0.29, p=0.26
Age (AD Only)
M15 midnightblue.MEGATMT488
Ambiguous
MEmidnightblue.Plasma 35 Samp. (Synthet
ANOVA p: 0.00034
MEmidnightblue.Plasma (Synthetic)
ANOVA p: 0.00065
MEmidnightblue.Plasma (Synthetic)
ANOVA p: 0.0039
bicor=0.56, p=5e-04
cor=0.54, p=0.00081
bicor=0.52, p=0.0029
cor=0.36, p=0.047
bicor=-0.15, p=0.56
cor=-0.15, p=0.55
bicor=-0.32, p=0.21
cor=-0.33, p=0.2

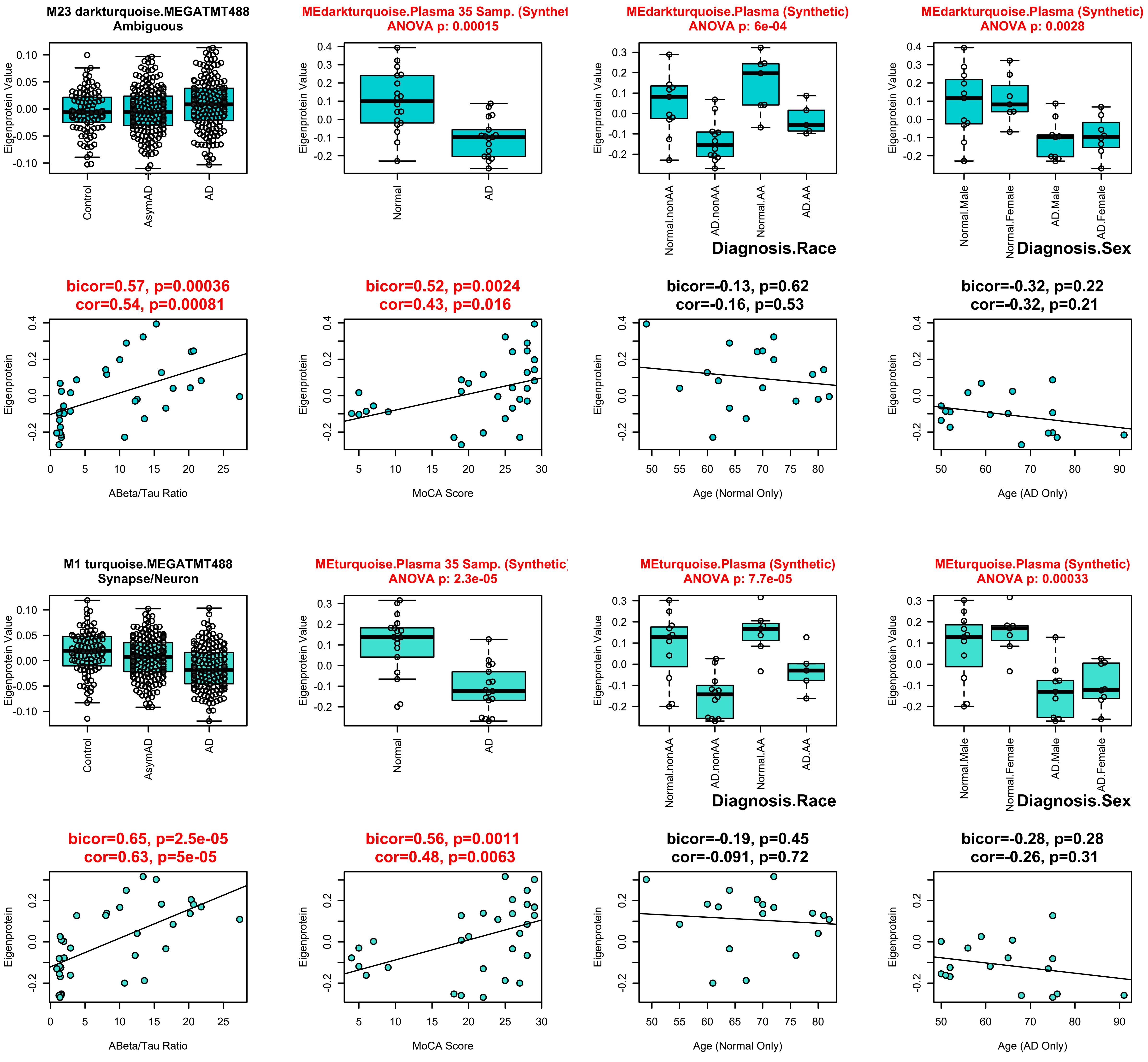
M23 darkturquoise.MEGATMT488
Ambiguous
Eigenprotein Value
Control
AsymAD
AD
MEdarkturquoise.Plasma 35 Samp. (Synthet
ANOVA p: 0.00015
Normal
AD
MEdarkturquoise.Plasma (Synthetic)
ANOVA p: 6e-04
Normal.nonAA
AD.nonAA
Normal.AA
AD.AA
Diagnosis.Race
MEdarkturquoise.Plasma (Synthetic)
ANOVA p: 0.0028
Normal.Male
Normal.Female
AD.Male
AD.Female
Diagnosis.Sex
bicor=0.57, p=0.00036
cor=0.54, p=0.00081
Eigenprotein
ABeta/Tau Ratio
bicor=0.52, p=0.0024
cor=0.43, p=0.016
MoCA Score
bicor=-0.13, p=0.62
cor=-0.16, p=0.53
Age (Normal Only)
bicor=-0.32, p=0.22
cor=-0.32, p=0.21
Age (AD Only)
M1 turquoise.MEGATMT488
Synapse/Neuron
Eigenprotein Value
Control
AsymAD
AD
MEturquoise.Plasma 35 Samp. (Synthetic)
ANOVA p: 2.3e-05
Normal
AD
MEturquoise.Plasma (Synthetic)
ANOVA p: 7.7e-05
Normal.nonAA
AD.nonAA
Normal.AA
AD.AA
Diagnosis.Race
MEturquoise.Plasma (Synthetic)
ANOVA p: 0.00033
Normal.Male
Normal.Female
AD.Male
AD.Female
Diagnosis.Sex
bicor=0.65, p=2.5e-05
cor=0.63, p=5e-05
Eigenprotein
ABeta/Tau Ratio
bicor=0.56, p=0.0011
cor=0.48, p=0.0063
MoCA Score
bicor=-0.19, p=0.45
cor=-0.091, p=0.72
Age (Normal Only)
bicor=-0.28, p=0.28
cor=-0.26, p=0.31
Age (AD Only)

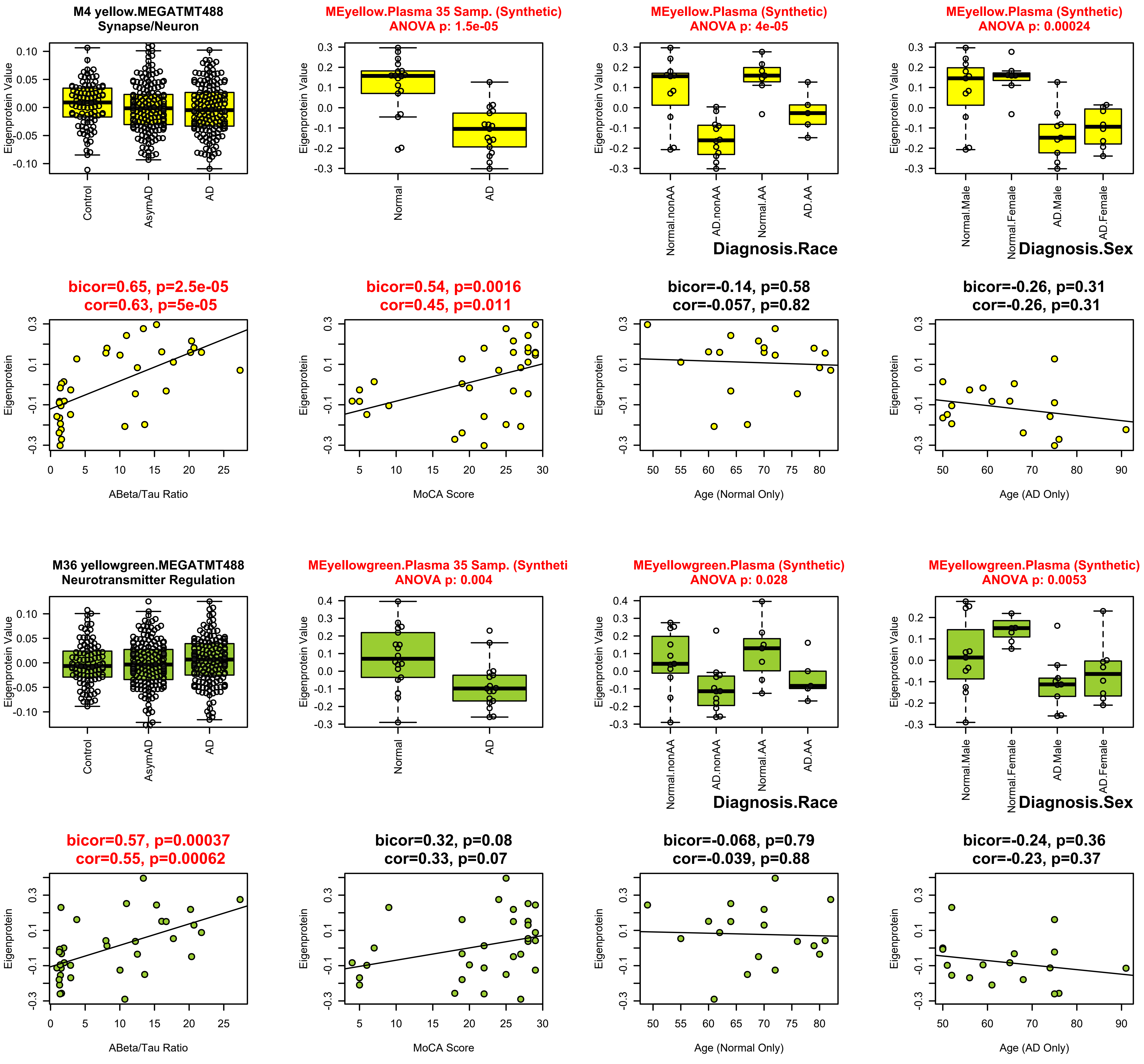
M4 yellow.MEGATMT488
Synapse/Neuron
Eigenprotein Value
Control
AsymAD
AD
MEyellow.Plasma 35 Samp. (Synthetic)
ANOVA p: 1.5e-05
Normal
AD
MEyellow.Plasma (Synthetic)
ANOVA p: 4e-05
Normal.nonAA
AD.nonAA
Normal.AA
AD.AA
Diagnosis.Race
MEyellow.Plasma (Synthetic)
ANOVA p: 0.00024
Normal.Male
Normal.Female
AD.Male
AD.Female
Diagnosis.Sex
bicor=0.65, p=2.5e-05
cor=0.63, p=5e-05
Eigenprotein
ABeta/Tau Ratio
bicor=0.54, p=0.0016
cor=0.45, p=0.011
MoCA Score
bicor=-0.14, p=0.58
cor=-0.057, p=0.82
Age (Normal Only)
bicor=-0.26, p=0.31
cor=-0.26, p=0.31
Age (AD Only)
M36 yellowgreen.MEGATMT488
Neurotransmitter Regulation
MEyellowgreen.Plasma 35 Samp. (Syntheti
ANOVA p: 0.004
MEyellowgreen.Plasma (Synthetic)
ANOVA p: 0.028
MEyellowgreen.Plasma (Synthetic)
ANOVA p: 0.0053
bicor=0.57, p=0.00037
cor=0.55, p=0.00062
bicor=0.32, p=0.08
cor=0.33, p=0.07
bicor=-0.068, p=0.79
cor=-0.039, p=0.88
bicor=-0.24, p=0.36
cor=-0.23, p=0.37

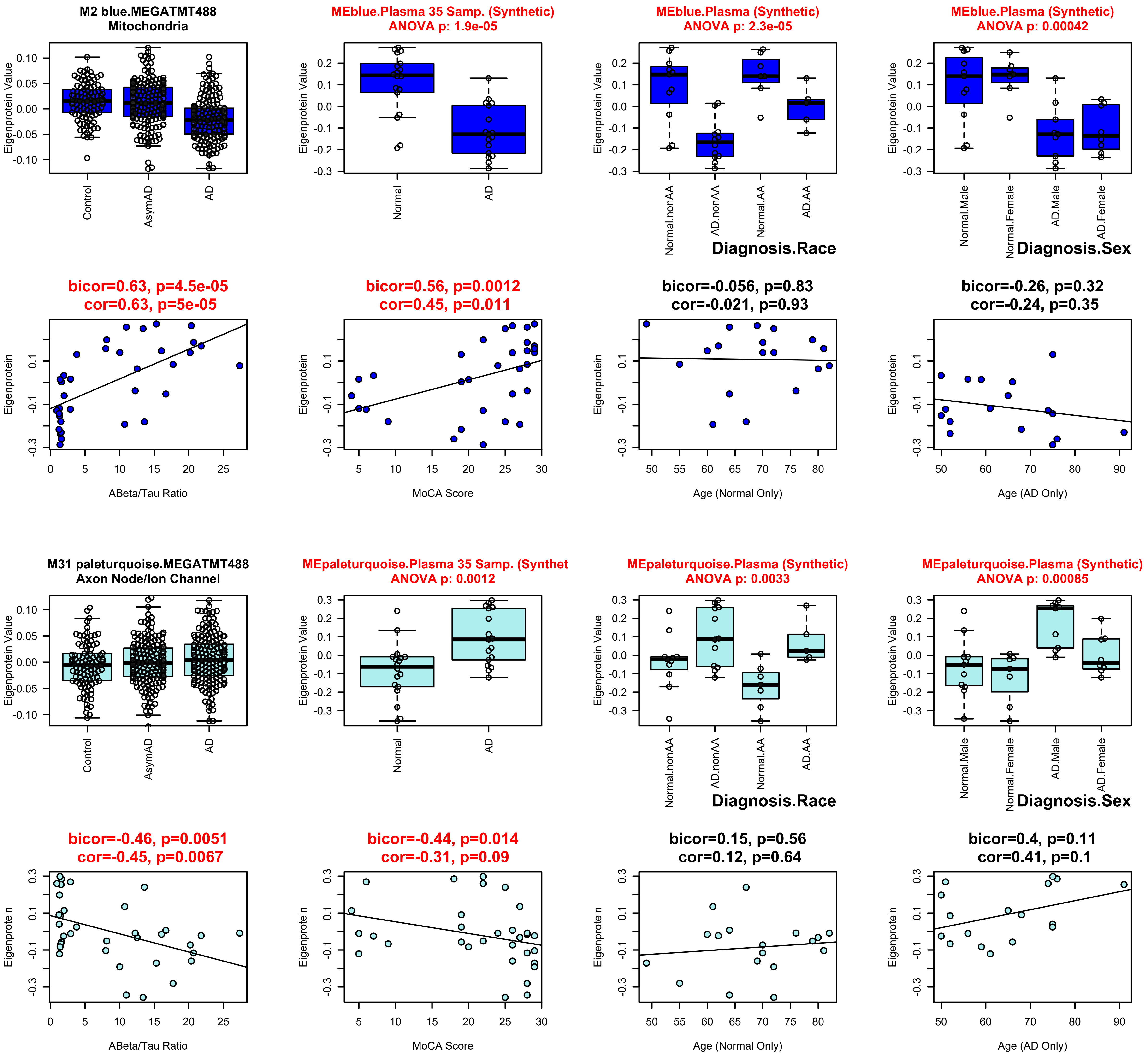
M2 blue.MEGATMT488
Mitochondria
Eigenprotein Value
Control
AsymAD
AD
MEblue.Plasma 35 Samp. (Synthetic)
ANOVA p: 1.9e-05
Normal
AD
MEblue.Plasma (Synthetic)
ANOVA p: 2.3e-05
Normal.nonAA
AD.nonAA
Normal.AA
AD.AA
Diagnosis.Race
MEblue.Plasma (Synthetic)
ANOVA p: 0.00042
Normal.Male
Normal.Female
AD.Male
AD.Female
Diagnosis.Sex
bicor=0.63, p=4.5e-05
cor=0.63, p=5e-05
ABeta/Tau Ratio
bicor=0.56, p=0.0012
cor=0.45, p=0.011
MoCA Score
bicor=-0.056, p=0.83
cor=-0.021, p=0.93
Age (Normal Only)
bicor=-0.26, p=0.32
cor=-0.24, p=0.35
Age (AD Only)
Eigenprotein
M31 paleturquoise.MEGATMT488
Axon Node/Ion Channel
MEpaleturquoise.Plasma 35 Samp. (Synthet
ANOVA p: 0.0012
MEpaleturquoise.Plasma (Synthetic)
ANOVA p: 0.0033
MEpaleturquoise.Plasma (Synthetic)
ANOVA p: 0.00085
bicor=-0.46, p=0.0051
cor=-0.45, p=0.0067
bicor=-0.44, p=0.014
cor=-0.31, p=0.09
bicor=0.15, p=0.56
cor=0.12, p=0.64
bicor=0.4, p=0.11
cor=0.41, p=0.1

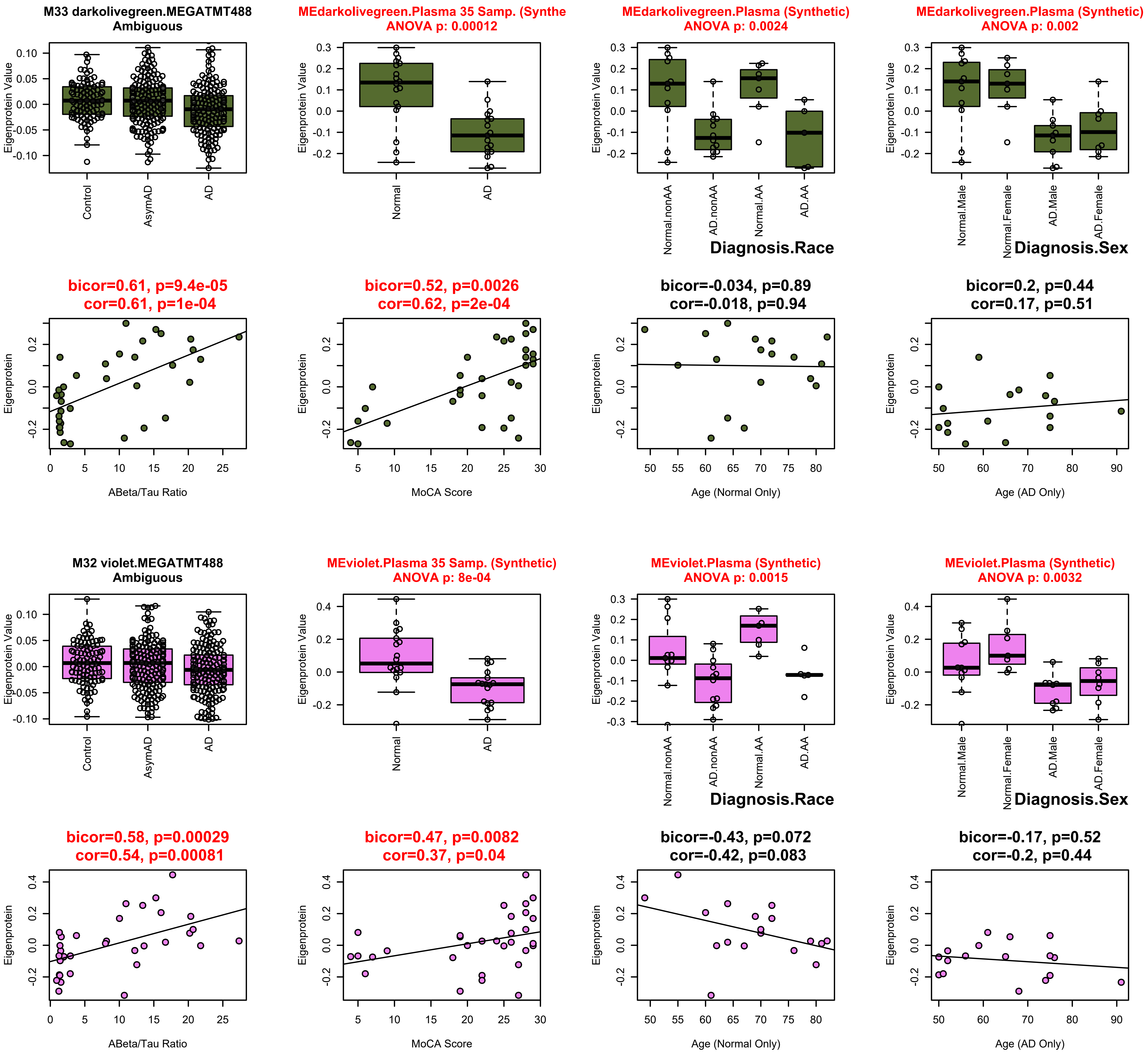

M33 darkolivegreen.MEGATMT488
Ambiguous
Eigenprotein Value
Control
AsymAD
AD
MEdarkolivegreen.Plasma 35 Samp. (Synthe
ANOVA p: 0.00012
Normal
AD
MEdarkolivegreen.Plasma (Synthetic)
ANOVA p: 0.0024
Normal.nonAA
AD.nonAA
Normal.AA
AD.AA
Diagnosis.Race
MEdarkolivegreen.Plasma (Synthetic)
ANOVA p: 0.002
Normal.Male
Normal.Female
AD.Male
AD.Female
Diagnosis.Sex
bicor=0.61, p=9.4e-05
cor=0.61, p=1e-04
Eigenprotein
ABeta/Tau Ratio
bicor=0.52, p=0.0026
cor=0.62, p=2e-04
MoCA Score
bicor=-0.034, p=0.89
cor=-0.018, p=0.94
Age (Normal Only)
bicor=0.2, p=0.44
cor=0.17, p=0.51
Age (AD Only)
M32 violet.MEGATMT488
Ambiguous
Eigenprotein Value
Control
AsymAD
AD
MEviolet.Plasma 35 Samp. (Synthetic)
ANOVA p: 8e-04
Normal
AD
MEviolet.Plasma (Synthetic)
ANOVA p: 0.0015
Normal.nonAA
AD.nonAA
Normal.AA
AD.AA
Diagnosis.Race
MEviolet.Plasma (Synthetic)
ANOVA p: 0.0032
Normal.Male
Normal.Female
AD.Male
AD.Female
Diagnosis.Sex
bicor=0.58, p=0.00029
cor=0.54, p=0.00081
Eigenprotein
ABeta/Tau Ratio
bicor=0.47, p=0.0082
cor=0.37, p=0.04
MoCA Score
bicor=-0.43, p=0.072
cor=-0.42, p=0.083
Age (Normal Only)
bicor=-0.17, p=0.52
cor=-0.2, p=0.44
Age (AD Only)

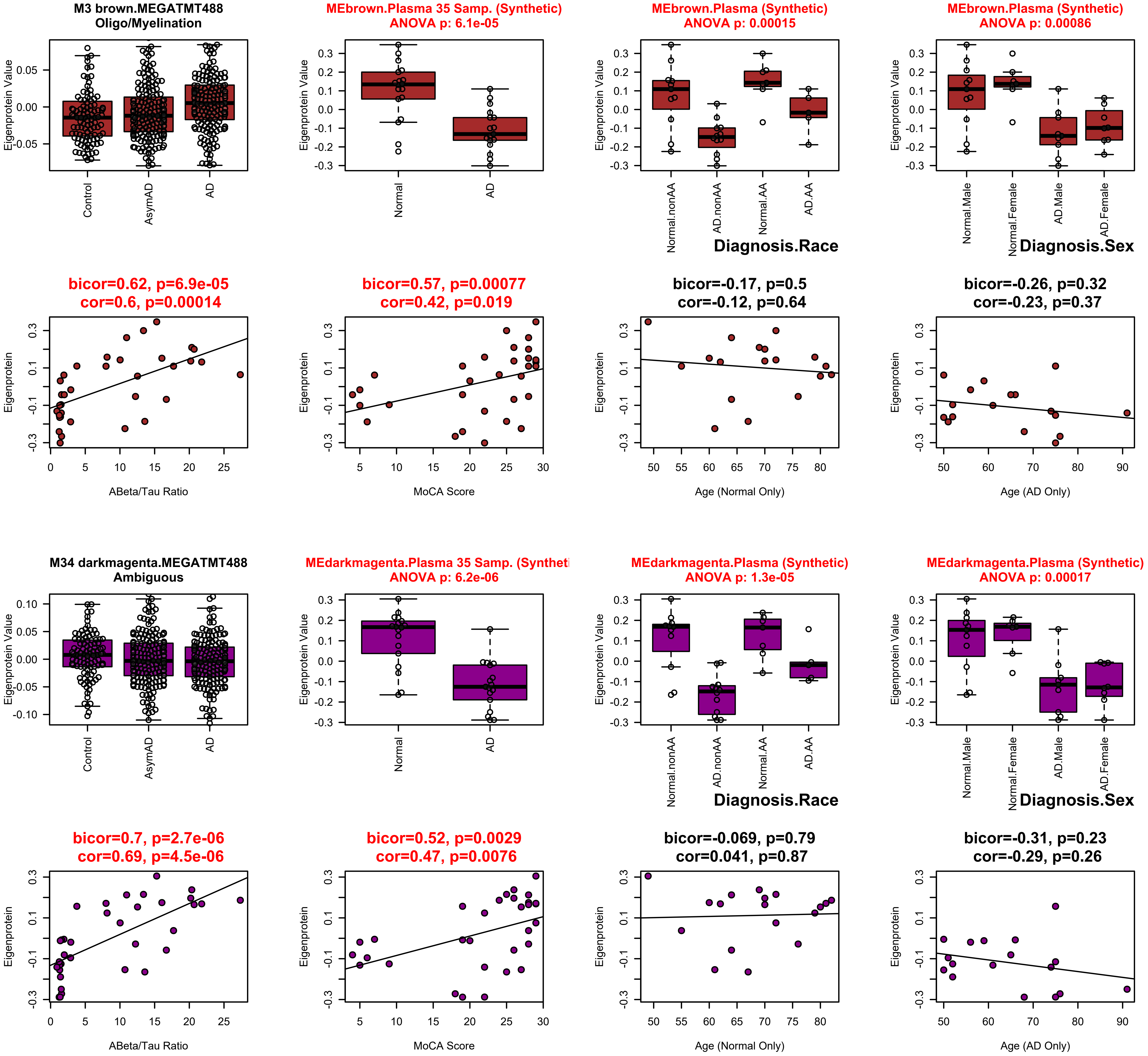

M3 brown.MEGATMT488
Oligo/Myelination
MEbrown.Plasma 35 Samp. (Synthetic)
ANOVA p: 6.1e-05
MEbrown.Plasma (Synthetic)
ANOVA p: 0.00015
MEbrown.Plasma (Synthetic)
ANOVA p: 0.00086
Eigenprotein Value
Control
AsymAD
AD
Normal
Normal.nonAA
AD.nonAA
Normal.AA
AD.AA
Diagnosis.Race
Normal.Male
Normal.Female
AD.Male
AD.Female
Diagnosis.Sex
bicor=0.62, p=6.9e-05
cor=0.6, p=0.00014
bicor=0.57, p=0.00077
cor=0.42, p=0.019
bicor=-0.17, p=0.5
cor=-0.12, p=0.64
bicor=-0.26, p=0.32
cor=-0.23, p=0.37
Eigenprotein
ABeta/Tau Ratio
MoCA Score
Age (Normal Only)
Age (AD Only)
M34 darkmagenta.MEGATMT488
Ambiguous
MEdarkmagenta.Plasma 35 Samp. (Syntheti
ANOVA p: 6.2e-06
MEdarkmagenta.Plasma (Synthetic)
ANOVA p: 1.3e-05
MEdarkmagenta.Plasma (Synthetic)
ANOVA p: 0.00017
bicor=0.7, p=2.7e-06
cor=0.69, p=4.5e-06
bicor=0.52, p=0.0029
cor=0.47, p=0.0076
bicor=-0.069, p=0.79
cor=0.041, p=0.87
bicor=-0.31, p=0.23
cor=-0.29, p=0.26

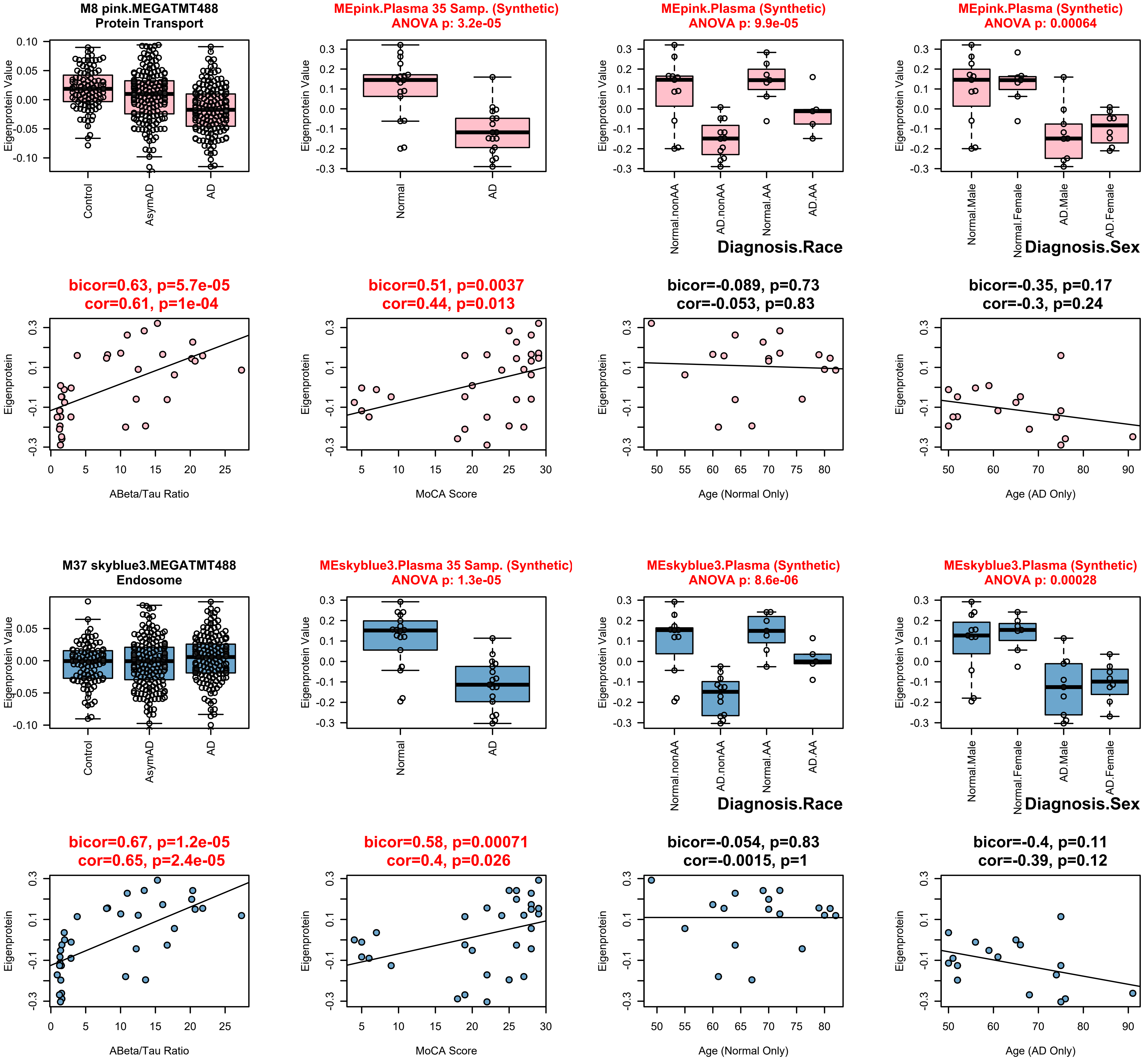
M8 pink.MEGATMT488
Protein Transport
Eigenprotein Value
Control
AsymAD
AD
MEpink.Plasma 35 Samp. (Synthetic)
ANOVA p: 3.2e-05
Normal
AD
MEpink.Plasma (Synthetic)
ANOVA p: 9.9e-05
Normal.nonAA
AD.nonAA
Normal.AA
AD.AA
Diagnosis.Race
MEpink.Plasma (Synthetic)
ANOVA p: 0.00064
Normal.Male
Normal.Female
AD.Male
AD.Female
Diagnosis.Sex
bicor=0.63, p=5.7e-05
cor=0.61, p=1e-04
Eigenprotein
ABeta/Tau Ratio
bicor=0.51, p=0.0037
cor=0.44, p=0.013
MoCA Score
bicor=-0.089, p=0.73
cor=-0.053, p=0.83
Age (Normal Only)
bicor=-0.35, p=0.17
cor=-0.3, p=0.24
Age (AD Only)
M37 skyblue3.MEGATMT488
Endosome
MEskyblue3.Plasma 35 Samp. (Synthetic)
ANOVA p: 1.3e-05
MEskyblue3.Plasma (Synthetic)
ANOVA p: 8.6e-06
Diagnosis.Race
MEskyblue3.Plasma (Synthetic)
ANOVA p: 0.00028
Diagnosis.Sex
bicor=0.67, p=1.2e-05
cor=0.65, p=2.4e-05
ABeta/Tau Ratio
bicor=0.58, p=0.00071
cor=0.4, p=0.026
MoCA Score
bicor=-0.054, p=0.83
cor=-0.0015, p=1
Age (Normal Only)
bicor=-0.4, p=0.11
cor=-0.39, p=0.12
Age (AD Only)

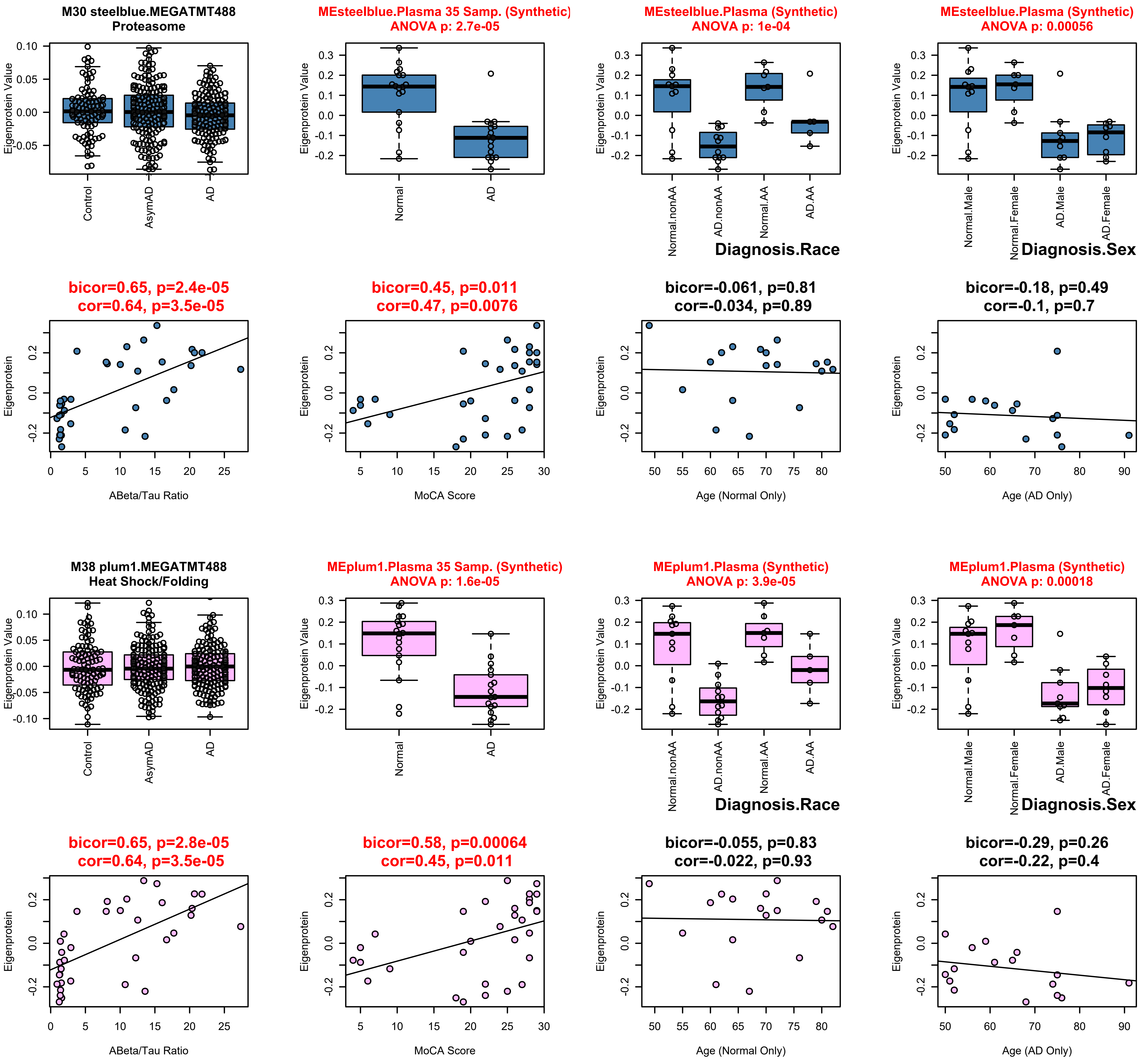
M30 steelblue.MEGATMT488
Proteasome
Eigenprotein Value
Control
AsymAD
AD
MEsteelblue.Plasma 35 Samp. (Synthetic)
ANOVA p: 2.7e-05
Normal
AD
MEsteelblue.Plasma (Synthetic)
ANOVA p: 1e-04
Normal.nonAA
AD.nonAA
Normal.AA
AD.AA
Diagnosis.Race
MEsteelblue.Plasma (Synthetic)
ANOVA p: 0.00056
Normal.Male
Normal.Female
AD.Male
AD.Female
Diagnosis.Sex
bicor=0.65, p=2.4e-05
cor=0.64, p=3.5e-05
Eigenprotein
ABeta/Tau Ratio
bicor=0.45, p=0.011
cor=0.47, p=0.0076
MoCA Score
bicor=-0.061, p=0.81
cor=-0.034, p=0.89
Age (Normal Only)
bicor=-0.18, p=0.49
cor=-0.1, p=0.7
Age (AD Only)
M38 plum1.MEGATMT488
Heat Shock/Folding
Eigenprotein Value
Control
AsymAD
AD
MEplum1.Plasma 35 Samp. (Synthetic)
ANOVA p: 1.6e-05
Normal
AD
MEplum1.Plasma (Synthetic)
ANOVA p: 3.9e-05
Normal.nonAA
AD.nonAA
Normal.AA
AD.AA
Diagnosis.Race
MEplum1.Plasma (Synthetic)
ANOVA p: 0.00018
Normal.Male
Normal.Female
AD.Male
AD.Female
Diagnosis.Sex
bicor=0.65, p=2.8e-05
cor=0.64, p=3.5e-05
Eigenprotein
ABeta/Tau Ratio
bicor=0.58, p=0.00064
cor=0.45, p=0.011
MoCA Score
bicor=-0.055, p=0.83
cor=-0.022, p=0.93
Age (Normal Only)
bicor=-0.29, p=0.26
cor=-0.22, p=0.4
Age (AD Only)

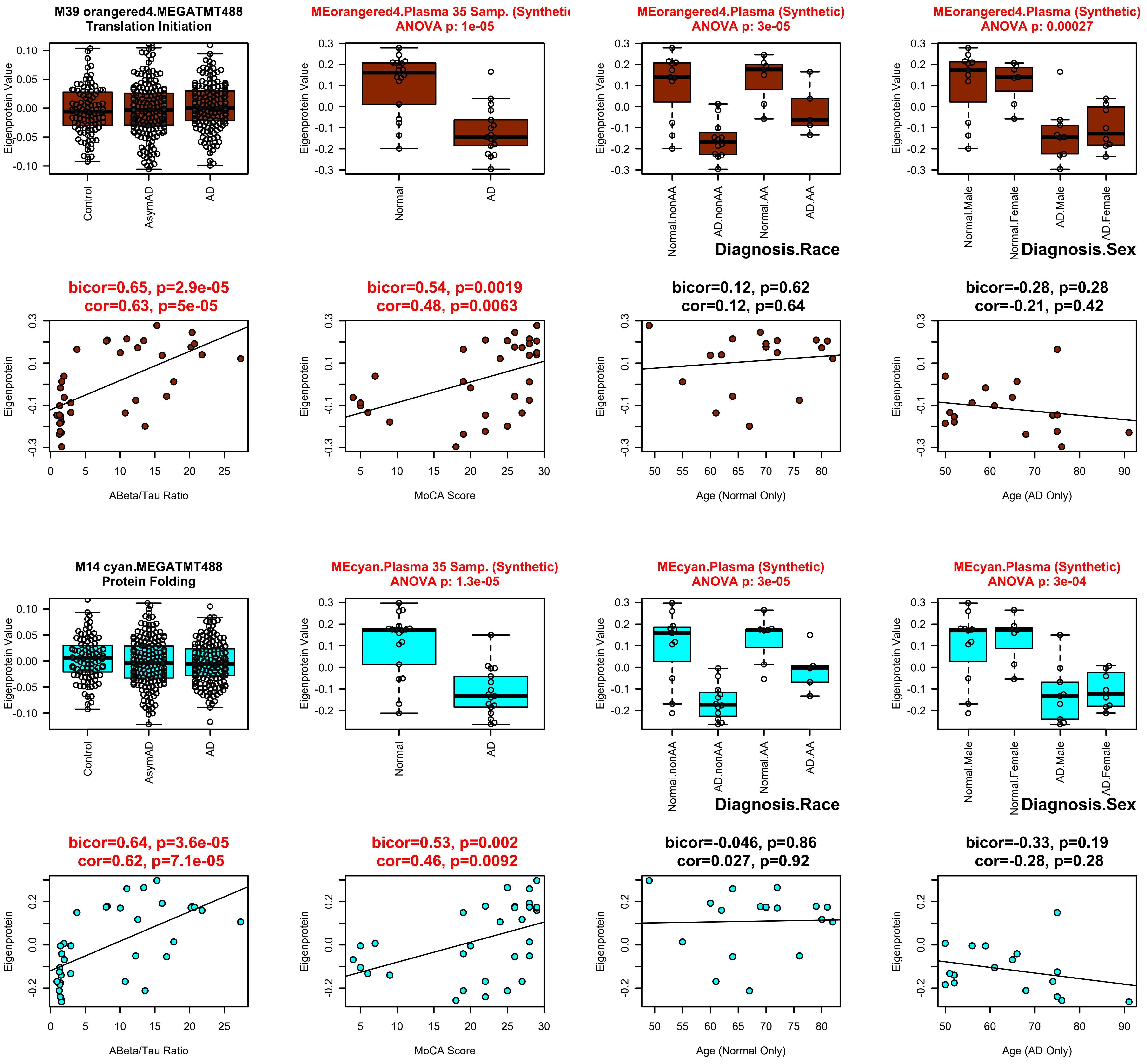
M39 orangered4.MEGATMT488
Translation Initiation
Eigenprotein Value
Control
AsymAD
AD
MEorangered4.Plasma 35 Samp. (Synthetic
ANOVA p: 1e-05
Normal
AD
MEorangered4.Plasma (Synthetic)
ANOVA p: 3e-05
Normal.nonAA
AD.nonAA
Normal.AA
AD.AA
Diagnosis.Race
MEorangered4.Plasma (Synthetic)
ANOVA p: 0.00027
Normal.Male
Normal.Female
AD.Male
AD.Female
Diagnosis.Sex
bicor=0.65, p=2.9e-05
cor=0.63, p=5e-05
Eigenprotein
ABeta/Tau Ratio
bicor=0.54, p=0.0019
cor=0.48, p=0.0063
MoCA Score
bicor=0.12, p=0.62
cor=0.12, p=0.64
Age (Normal Only)
bicor=-0.28, p=0.28
cor=-0.21, p=0.42
Age (AD Only)
M14 cyan.MEGATMT488
Protein Folding
Eigenprotein Value
Control
AsymAD
AD
MEcyan.Plasma 35 Samp. (Synthetic)
ANOVA p: 1.3e-05
Normal
AD
MEcyan.Plasma (Synthetic)
ANOVA p: 3e-05
Normal.nonAA
AD.nonAA
Normal.AA
AD.AA
Diagnosis.Race
MEcyan.Plasma (Synthetic)
ANOVA p: 3e-04
Normal.Male
Normal.Female
AD.Male
AD.Female
Diagnosis.Sex
bicor=0.64, p=3.6e-05
cor=0.62, p=7.1e-05
Eigenprotein
ABeta/Tau Ratio
bicor=0.53, p=0.002
cor=0.46, p=0.0092
MoCA Score
bicor=-0.046, p=0.86
cor=0.027, p=0.92
Age (Normal Only)
bicor=-0.33, p=0.19
cor=-0.28, p=0.28
Age (AD Only)

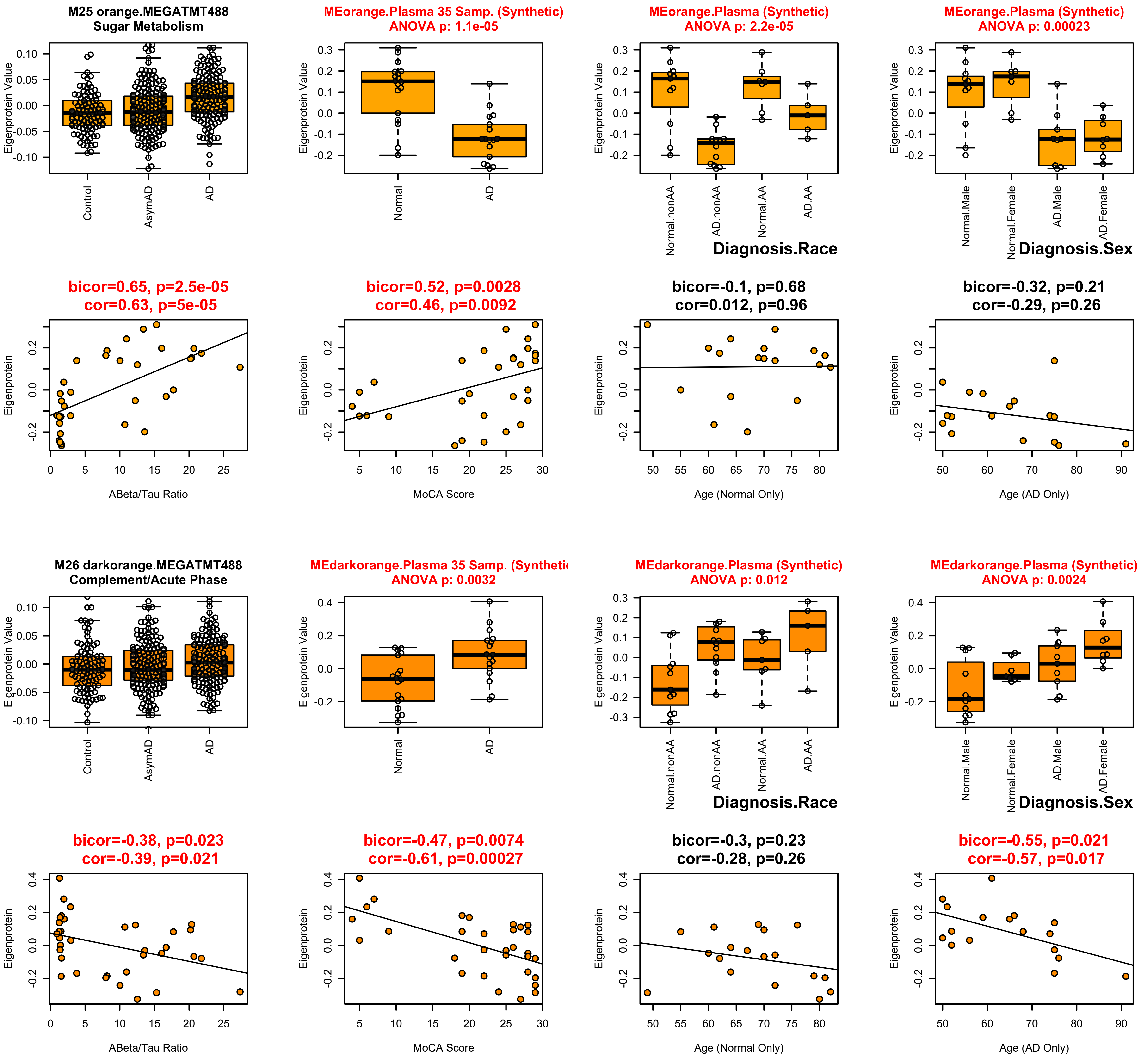
M25 orange.MEGATMT488
Sugar Metabolism
Eigenprotein Value
Control
AsymAD
AD
MEorange.Plasma 35 Samp. (Synthetic)
ANOVA p: 1.1e-05
Normal
AD
MEorange.Plasma (Synthetic)
ANOVA p: 2.2e-05
Normal.nonAA
AD.nonAA
Normal.AA
AD.AA
Diagnosis.Race
MEorange.Plasma (Synthetic)
ANOVA p: 0.00023
Normal.Male
Normal.Female
AD.Male
AD.Female
Diagnosis.Sex
bicor=0.65, p=2.5e-05
cor=0.63, p=5e-05
Eigenprotein
ABeta/Tau Ratio
bicor=0.52, p=0.0028
cor=0.46, p=0.0092
MoCA Score
bicor=-0.1, p=0.68
cor=0.012, p=0.96
Age (Normal Only)
bicor=-0.32, p=0.21
cor=-0.29, p=0.26
Age (AD Only)
M26 darkorange.MEGATMT488
Complement/Acute Phase
Eigenprotein Value
Control
AsymAD
AD
MEdarkorange.Plasma 35 Samp. (Syntheti
ANOVA p: 0.0032
Normal
AD
MEdarkorange.Plasma (Synthetic)
ANOVA p: 0.012
Normal.nonAA
AD.nonAA
Normal.AA
AD.AA
Diagnosis.Race
MEdarkorange.Plasma (Synthetic)
ANOVA p: 0.0024
Normal.Male
Normal.Female
AD.Male
AD.Female
Diagnosis.Sex
bicor=-0.38, p=0.023
cor=-0.39, p=0.021
Eigenprotein
ABeta/Tau Ratio
bicor=-0.47, p=0.0074
cor=-0.61, p=0.00027
MoCA Score
bicor=-0.3, p=0.23
cor=-0.28, p=0.26
Age (Normal Only)
bicor=-0.55, p=0.021
cor=-0.57, p=0.017
Age (AD Only)

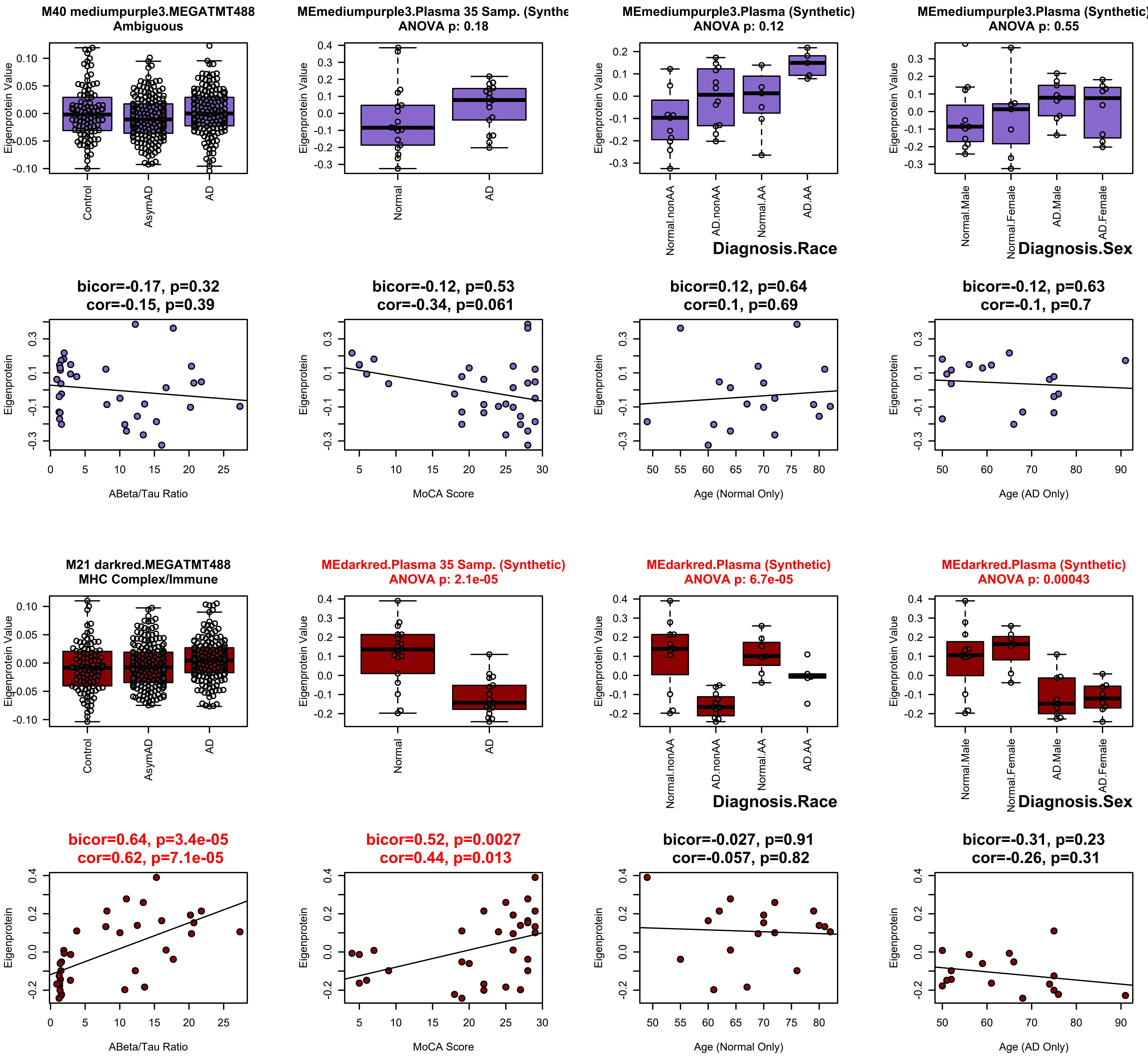

M40 mediumpurple3.MEGATMT488
Ambiguous
Eigenprotein Value
Control
AsymAD
AD
MEmediumpurple3.Plasma 35 Samp. (Synthe
ANOVA p: 0.18
Normal
AD
MEmediumpurple3.Plasma (Synthetic)
ANOVA p: 0.12
Normal.nonAA
AD.nonAA
Normal.AA
AD.AA
Diagnosis.Race
MEmediumpurple3.Plasma (Synthetic)
ANOVA p: 0.55
Normal.Male
Normal.Female
AD.Male
AD.Female
Diagnosis.Sex
bicor=-0.17, p=0.32
cor=-0.15, p=0.39
Eigenprotein
ABeta/Tau Ratio
bicor=-0.12, p=0.53
cor=-0.34, p=0.061
MoCA Score
bicor=0.12, p=0.64
cor=0.1, p=0.69
Age (Normal Only)
bicor=-0.12, p=0.63
cor=-0.1, p=0.7
Age (AD Only)
M21 darkred.MEGATMT488
MHC Complex/Immune
MEdarkred.Plasma 35 Samp. (Synthetic)
ANOVA p: 2.1e-05
MEdarkred.Plasma (Synthetic)
ANOVA p: 6.7e-05
Diagnosis.Race
MEdarkred.Plasma (Synthetic)
ANOVA p: 0.00043
Diagnosis.Sex
bicor=0.64, p=3.4e-05
cor=0.62, p=7.1e-05
ABeta/Tau Ratio
bicor=0.52, p=0.0027
cor=0.44, p=0.013
MoCA Score
bicor=-0.027, p=0.91
cor=-0.057, p=0.82
Age (Normal Only)
bicor=-0.31, p=0.23
cor=-0.26, p=0.31
Age (AD Only)

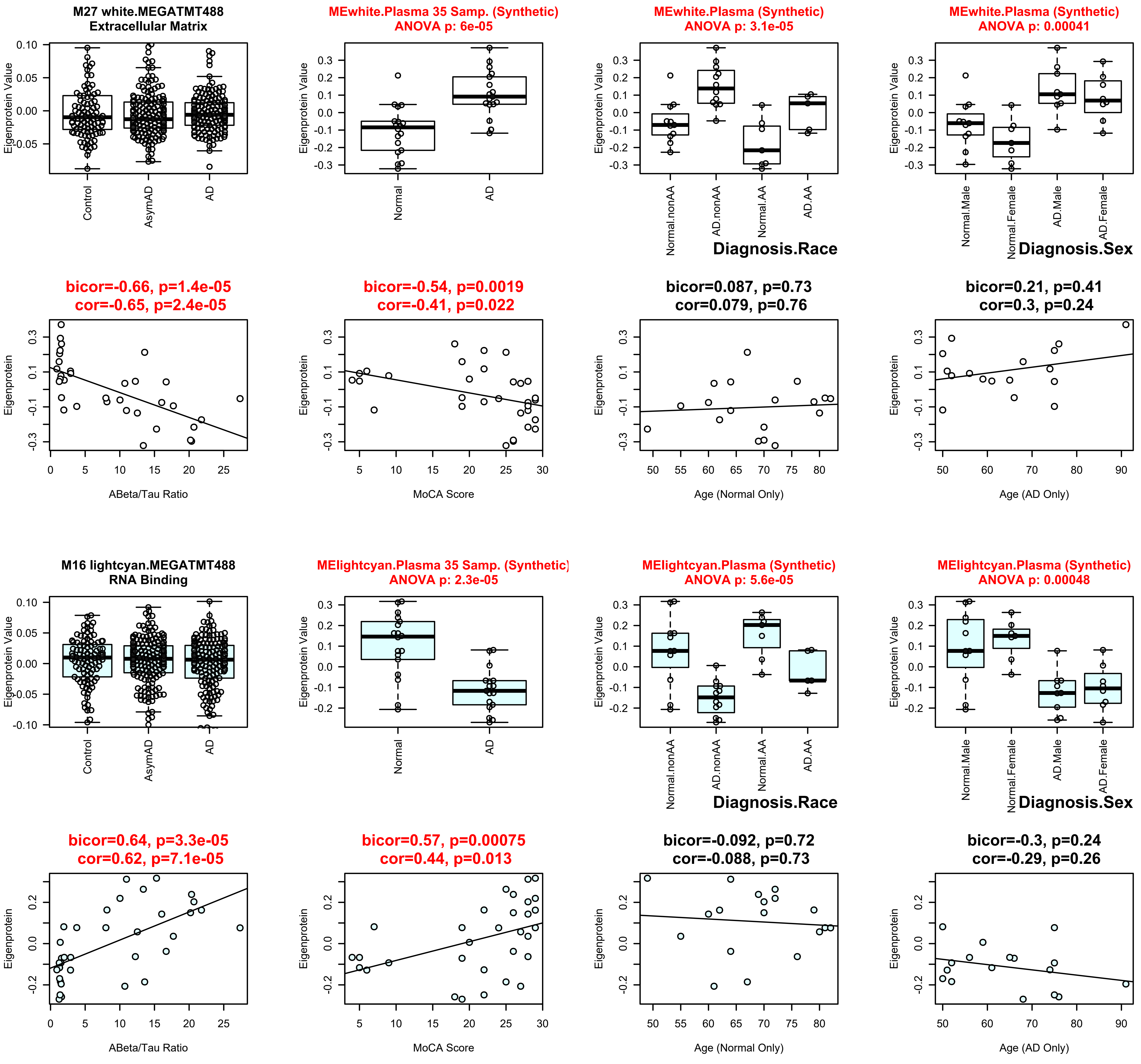
M27 white.MEGATMT488
Extracellular Matrix
Eigenprotein Value
Control
AsymAD
AD
MEwhite.Plasma 35 Samp. (Synthetic)
ANOVA p: 6e-05
Normal
AD
MEwhite.Plasma (Synthetic)
ANOVA p: 3.1e-05
Normal.nonAA
AD.nonAA
Normal.AA
AD.AA
Diagnosis.Race
MEwhite.Plasma (Synthetic)
ANOVA p: 0.00041
Normal.Male
Normal.Female
AD.Male
AD.Female
Diagnosis.Sex
bicor=-0.66, p=1.4e-05
cor=-0.65, p=2.4e-05
Eigenprotein
ABeta/Tau Ratio
bicor=-0.54, p=0.0019
cor=-0.41, p=0.022
MoCA Score
bicor=0.087, p=0.73
cor=0.079, p=0.76
Age (Normal Only)
bicor=0.21, p=0.41
cor=0.3, p=0.24
Age (AD Only)
M16 lightcyan.MEGATMT488
RNA Binding
MElightcyan.Plasma 35 Samp. (Synthetic)
ANOVA p: 2.3e-05
MElightcyan.Plasma (Synthetic)
ANOVA p: 5.6e-05
MElightcyan.Plasma (Synthetic)
ANOVA p: 0.00048
bicor=0.64, p=3.3e-05
cor=0.62, p=7.1e-05
bicor=0.57, p=0.00075
cor=0.44, p=0.013
bicor=-0.092, p=0.72
cor=-0.088, p=0.73
bicor=-0.3, p=0.24
cor=-0.29, p=0.26

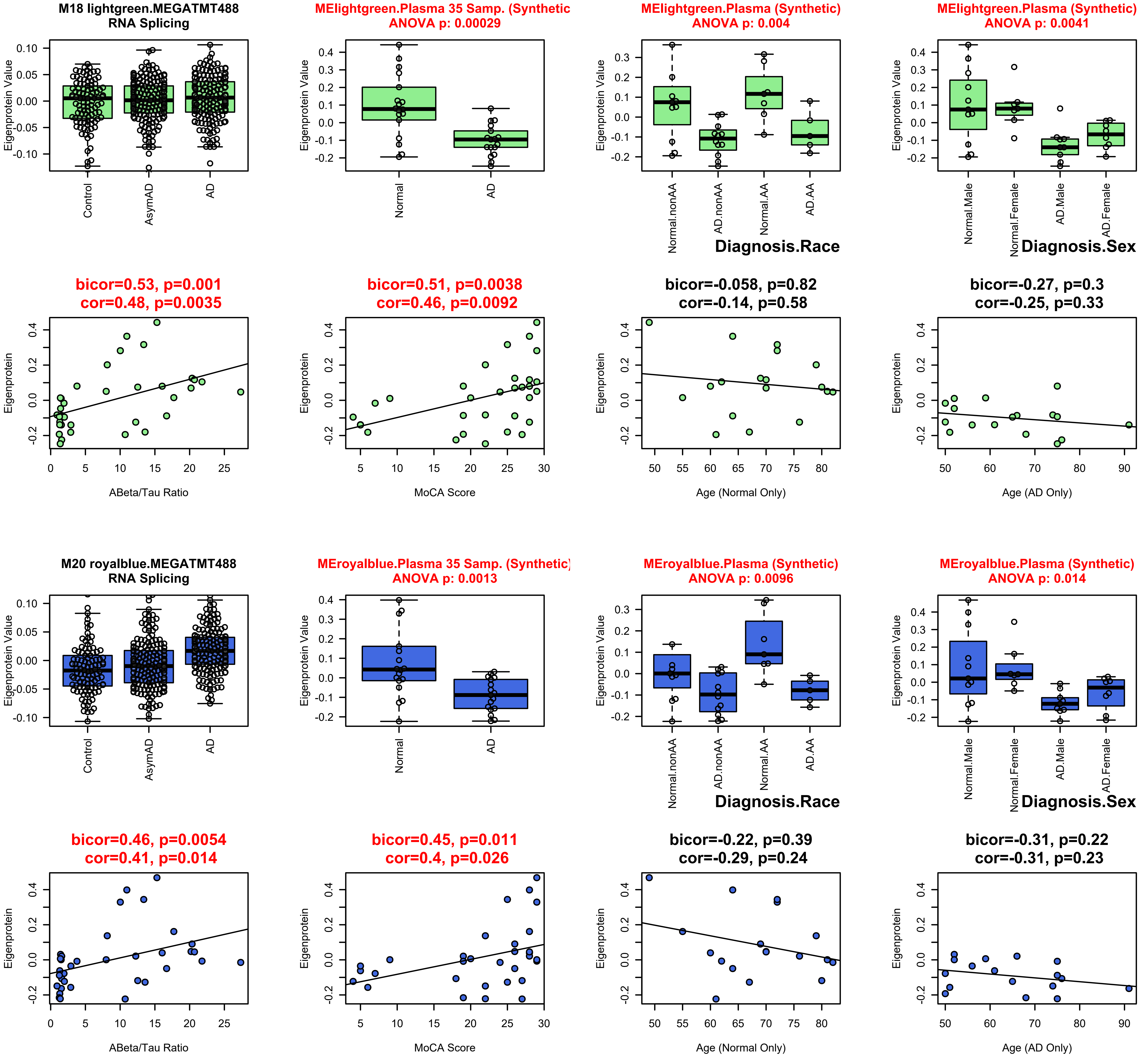
M18 lightgreen.MEGATMT488
RNA Splicing
Eigenprotein Value
Control
AsymAD
AD
MElightgreen.Plasma 35 Samp. (Synthetic
ANOVA p: 0.00029
Normal
AD
MElightgreen.Plasma (Synthetic)
ANOVA p: 0.004
Normal.nonAA
AD.nonAA
Normal.AA
AD.AA
Diagnosis.Race
MElightgreen.Plasma (Synthetic)
ANOVA p: 0.0041
Normal.Male
Normal.Female
AD.Male
AD.Female
Diagnosis.Sex
bicor=0.53, p=0.001
cor=0.48, p=0.0035
Eigenprotein
ABeta/Tau Ratio
bicor=0.51, p=0.0038
cor=0.46, p=0.0092
MoCA Score
bicor=-0.058, p=0.82
cor=-0.14, p=0.58
Age (Normal Only)
bicor=-0.27, p=0.3
cor=-0.25, p=0.33
Age (AD Only)
M20 royalblue.MEGATMT488
RNA Splicing
MEroyalblue.Plasma 35 Samp. (Synthetic)
ANOVA p: 0.0013
MEroyalblue.Plasma (Synthetic)
ANOVA p: 0.0096
MEroyalblue.Plasma (Synthetic)
ANOVA p: 0.014
bicor=0.46, p=0.0054
cor=0.41, p=0.014
bicor=0.45, p=0.011
cor=0.4, p=0.026
bicor=-0.22, p=0.39
cor=-0.29, p=0.24
bicor=-0.31, p=0.22
cor=-0.31, p=0.23

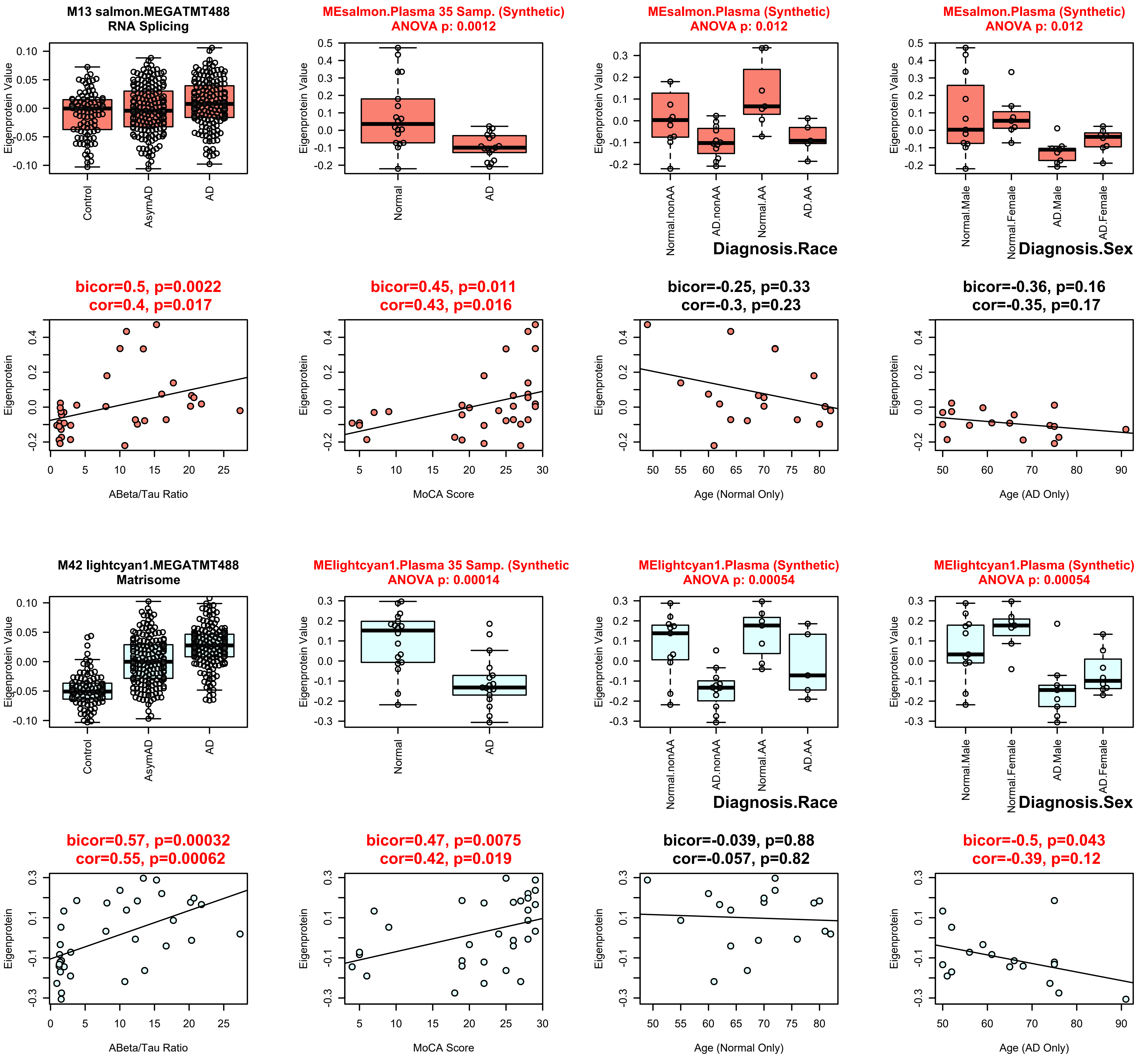

M13 salmon.MEGATMT488
RNA Splicing
Eigenprotein Value
Control
AsymAD
AD
MEsalmon.Plasma 35 Samp. (Synthetic)
ANOVA p: 0.0012
Normal
AD
MEsalmon.Plasma (Synthetic)
ANOVA p: 0.012
Normal.nonAA
AD.nonAA
Normal.AA
AD.AA
Diagnosis.Race
MEsalmon.Plasma (Synthetic)
ANOVA p: 0.012
Normal.Male
Normal.Female
AD.Male
AD.Female
Diagnosis.Sex
bicor=0.5, p=0.0022
cor=0.4, p=0.017
Eigenprotein
ABeta/Tau Ratio
bicor=0.45, p=0.011
cor=0.43, p=0.016
MoCA Score
bicor=-0.25, p=0.33
cor=-0.3, p=0.23
Age (Normal Only)
bicor=-0.36, p=0.16
cor=-0.35, p=0.17
Age (AD Only)
M42 lightcyan1.MEGATMT488
Matrisome
Eigenprotein Value
Control
AsymAD
AD
MElightcyan1.Plasma 35 Samp. (Synthetic
ANOVA p: 0.00014
Normal
AD
MElightcyan1.Plasma (Synthetic)
ANOVA p: 0.00054
Normal.nonAA
AD.nonAA
Normal.AA
AD.AA
Diagnosis.Race
MElightcyan1.Plasma (Synthetic)
ANOVA p: 0.00054
Normal.Male
Normal.Female
AD.Male
AD.Female
Diagnosis.Sex
bicor=0.57, p=0.00032
cor=0.55, p=0.00062
Eigenprotein
ABeta/Tau Ratio
bicor=0.47, p=0.0075
cor=0.42, p=0.019
MoCA Score
bicor=-0.039, p=0.88
cor=-0.057, p=0.82
Age (Normal Only)
bicor=-0.5, p=0.043
cor=-0.39, p=0.12
Age (AD Only)

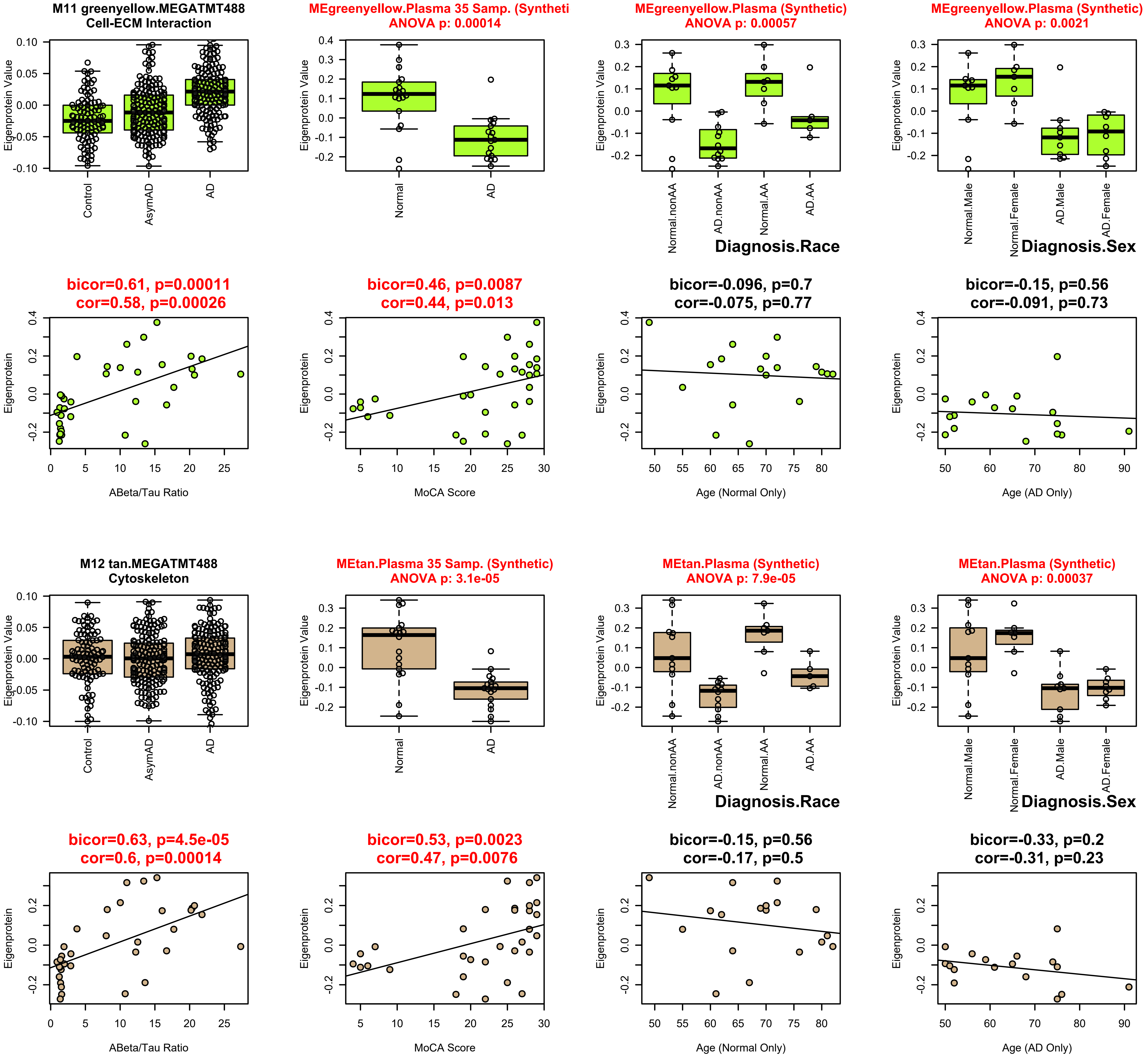
M11 greenyellow.MEGATMT488
Cell-ECM Interaction
Eigenprotein Value
Control
AsymAD
AD
MEgreenyellow.Plasma 35 Samp. (Syntheti
ANOVA p: 0.00014
Normal
AD
MEgreenyellow.Plasma (Synthetic)
ANOVA p: 0.00057
Normal.nonAA
AD.nonAA
Normal.AA
AD.AA
Diagnosis.Race
MEgreenyellow.Plasma (Synthetic)
ANOVA p: 0.0021
Normal.Male
Normal.Female
AD.Male
AD.Female
Diagnosis.Sex
bicor=0.61, p=0.00011
cor=0.58, p=0.00026
Eigenprotein
ABeta/Tau Ratio
bicor=0.46, p=0.0087
cor=0.44, p=0.013
MoCA Score
bicor=-0.096, p=0.7
cor=-0.075, p=0.77
Age (Normal Only)
bicor=-0.15, p=0.56
cor=-0.091, p=0.73
Age (AD Only)
M12 tan.MEGATMT488
Cytoskeleton
Eigenprotein Value
Control
AsymAD
AD
MEtan.Plasma 35 Samp. (Synthetic)
ANOVA p: 3.1e-05
Normal
AD
MEtan.Plasma (Synthetic)
ANOVA p: 7.9e-05
Normal.nonAA
AD.nonAA
Normal.AA
AD.AA
Diagnosis.Race
MEtan.Plasma (Synthetic)
ANOVA p: 0.00037
Normal.Male
Normal.Female
AD.Male
AD.Female
Diagnosis.Sex
bicor=0.63, p=4.5e-05
cor=0.6, p=0.00014
Eigenprotein
ABeta/Tau Ratio
bicor=0.53, p=0.0023
cor=0.47, p=0.0076
MoCA Score
bicor=-0.15, p=0.56
cor=-0.17, p=0.5
Age (Normal Only)
bicor=-0.33, p=0.2
cor=-0.31, p=0.23
Age (AD Only)
